# Hybrid CNN and Multi-Head Attention Model for Analyzing Epigenetic Mechanisms and Gene Expression Across Fungal Phylogenetic Distances

**DOI:** 10.1101/2024.12.12.628183

**Authors:** Laura Weinstock, Jenna Schambach, Anna Fisher, Cameron Kunstadt, Ethan Lee, Elizabeth Koning, William Morrell, Wittney Mays, Warren Davis, Raga Krishnakumar

**Affiliations:** Sandia National Laboratories, Livermore, CA 94550; Sandia National Laboratories, Albuquerque, NM 87123; The University of California, Berkeley, Berkeley, CA 94720; University of Illinois Urbana-Champaign, Urbana, IL 61801; Lawrence Berkeley National Laboratory, Berkeley, CA 94720

## Abstract

Understanding gene expression is crucial for optimizing biological processes in bioeconomic processes, human health, and environmental regulation. Epigenetic modifications significantly influence gene expression by altering chromatin structure and DNA accessibility. However, knowledge about the conservation of these mechanisms across species, especially in non-model organisms, is limited. This study predicts gene expression levels based on epigenetic modifications across fungal species, facilitating knowledge transfer from well-characterized to less understood species. We developed a deep learning model, MAPLE (Model predictions Across Phylogenetic distances by Learning Expression from Epigenetics), which integrates convolutional layers and multi-head attention to capture dependencies in epigenetic data. MAPLE shows strong cross-species performance in fungi, achieving up to 80% accuracy and 89% AUROC for intra-species validation, and 77% accuracy and 83% AUROC in cross-species tasks, outperforming benchmarks. SHAP analysis reveals key epigenetic features driving gene expression, providing insights for future experimental design. Our findings highlight MAPLE’s potential to generalize across fungal species, offering a versatile tool for optimizing gene expression.

## Introduction

Understanding the dynamic regulation of gene expression is a critical step towards robust regulation of biological processes. The need for precise, reversible, and dynamic control of gene expression in organisms is paramount for applications ranging from control of disease states to sustainable biomanufacturing^2–11^. Precise control ensures that the desired genes are expressed at the right levels, optimizing the production of target compounds or behavior while minimizing waste and unwanted byproducts. Reversible control allows for the temporary activation or suppression of genes, providing flexibility in the production process and enabling researchers to study gene function and metabolic pathways in a controlled manner. Dynamic control, which involves the ability to adjust gene expression in response to environmental or developmental cues, is crucial for adapting to changing conditions and real-time process optimization.

There are many regulatory mechanisms for gene expression, that, while broadly conserved in function, vary significantly between species in the details of how they impact genes. One significant regulatory mechanism is epigenetics, an umbrella term for factors, modifications and molecules that affect gene expression without altering genetic sequence, usually in a stable, reversible, and heritable manner^12, 13^. While there are broader definitions of the term, epigenetics is often used to refer to DNA and histone modifications that act to regulate gene expression. Epigenetics play a crucial role in regulating gene expression by altering chromatin structure and accessibility of DNA, and by recruiting and releasing factors from the genome^12, 13^. The location and combinations of epigenetic modifications on the genome significantly impacts gene expression outcomes.

Understanding the relationship between epigenetic modifications and gene expression can provide valuable insights into the regulation of cellular and population-level processes. Advances in genomic sequencing technologies have enabled the identification of epigenetic modifications at a genome-wide scale in more and broader organisms^14–18^. In mammals, there is a large body of literature on these epigenetic relationships. This rapidly-growing repository of epigenetic information has resulted in the ability to make increasingly accurate predictions both about how epigenetic modifications are distributed as well as their downstream functions. In recent years, machine learning models have shown great potential for use in predicting gene expression levels based on epigenetic modifications, aiding in identifying predictive features among the large and complex datasets^19–23^. This approach has been successful in predicting gene expression in various organisms, including humans, mice, and plants^1, 24–26^. Extending such approaches to broader phylogenetic distances can help shed light on regulatory mechanisms that are critical across a wide range of biological systems and processes, particularly of use for less well characterized taxa. One such example is fungi.

Fungi and engineered fungi are increasingly recognized as valuable chassis for biomanufacturing and bioproduction processes due to their versatility, efficiency, and sustainability. Fungi naturally produce a wide array of bioactive compounds, enzymes, and metabolites that are used for various industrial applications, including pharmaceuticals, biofuels, and food production. They are also capable of degrading and converting materials making them essential for bioremediation. Understanding the fungal stress response can facilitate the development of more effective bioremediation strategies. Finally, fungi are frequently used as materials themselves^27^. By leveraging genetic engineering, we can enhance these capabilities, tailoring fungi to produce specific compounds at higher yields or optimally exhibit specific desirable properties. Understanding how to broadly modify and control fungi to inform genetic engineering efforts would greatly advance several bioproduction applications. This could enable understanding of how to optimize the growth and flocculation of non-model organisms using model organism data, or how to gain and obtain a desired novel attribute obtained from non-model organisms in optimized model organism systems.

By predicting the effect of epigenetic modifications on gene expression, we can design strategies to optimize gene expression for a variety of applications. This can have significant implications for biomanufacturing, for instance, increasing the production of high-value compounds. However, despite the overall evolutionary conservation of epigenetic relationships, there is still a significant amount of variability in the impact of modifications across species, where the function of individual modifications or combinations of modifications may diverge^28^. This motivated the question of whether epigenetic regulatory patterns are identifiable across broad evolutionary distances in the fungal kingdom.

A challenge in predicting gene expression levels in fungi is the limited availability of epigenetic modification data, particularly in non-model organisms. In such cases, cross-species predictions can be used to overcome this limitation. Cross-species predictions involve training a machine learning model on one fungal species and using it to predict gene expression in another fungal species. However, cross-species predictions are limited by differences in the availability of epigenetic modification data and the variation in the epigenetic modification patterns across species. To address these limitations, it is essential to carefully choose the fungal species for cross-species predictions based on the availability and quality of the epigenetic modification data and develop models that can utilize this data appropriately.

In this study, we present MAPLE (Model predictions Across Phylogenetic distances by Learning Expression from Epigenetics) as a framework tool to predict gene expression levels based on epigenetic modification presence on fungal chromatin and based on underlying DNA sequences. Specifically, we are interested in identifying the relationship between modification type and the position and magnitude, or signal profile, of modifications profile near genes. By analyzing the relationship between epigenetic modifications and gene expression across different fungal species, we sought to develop predictive models that can be used to identify modifications that govern cross-species cellular activity, which can then be used as experimental tools to broadly engineer organisms like fungal chassis for optimal function^29–31^.

## Results

### Unique gene expression states form based on epigenetic modifications but differ between fungal species

To evaluate the use of conserved epigenetic modification patterns as the primary method by which predict changes in gene expression, we first curated a dataset of histone modification data matched with RNA sequencing data for four fungal species, called out in red boxes, that span a range taxonomic distances (Figure 1). We accessed publicly available datasets for four fungal species: *Neurospora crassa* (*N. crassa*), *Fusarium graminearum* (*F. graminearum*), *Leptosphaeria maculans* (*L. maculans*) and *Aspergillus nidulans* (*A. nidulans*). These species were chosen due to their range of biological functions, evolutionary relationships, and overlapping epigenomic and gene expression data availability. While they are all within the phylum Ascomycota, they are sufficiently distant from each other to evaluate the utility of cross-species models (Figure 1). Phylogenetic distance spread was confirmed using an in-house developed ancestral tree building tool specialized for fungal species called Poplar^32^. The organisms selected for this study reasonably span the phylogenetic tree when compared with other commonly studied species (Figure 1). Further information about the datasets used, the data matching and selection, and the pre-processing pipeline is described in Methods.

**Figure 1.**
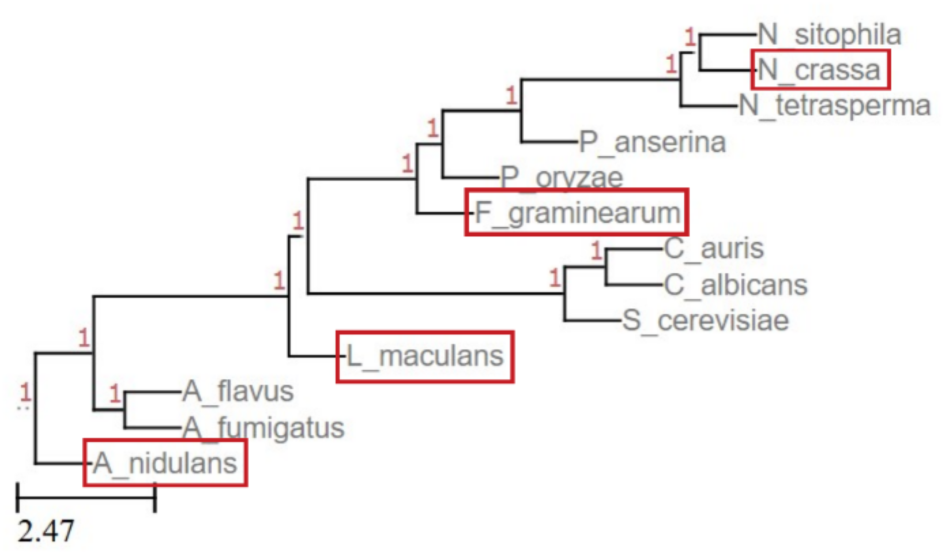
Phylogenetic tree inferred from genomes for candidate and common species. Species used in this study are called out in red boxes and show the relationship and span of evolutionary distances of the species.

We show that the curated epigenetic modification data can be used to identify unique chromatin states within fungal species using ChromHMM^33^, a pre-trained multivariate hidden Markov model that uses multiple histone modification datasets and patterns at promoters and enhancers to discover chromatin states. The hidden states are estimated based on chromatin accessibility, influenced by the epigenetic markers. The probability that each histone modification will be present in each state implicates the marker in the definition of that state, and connecting the gene expression levels for genes in each state links the states and markers to activity (Figure 2).

**Figure 2.**
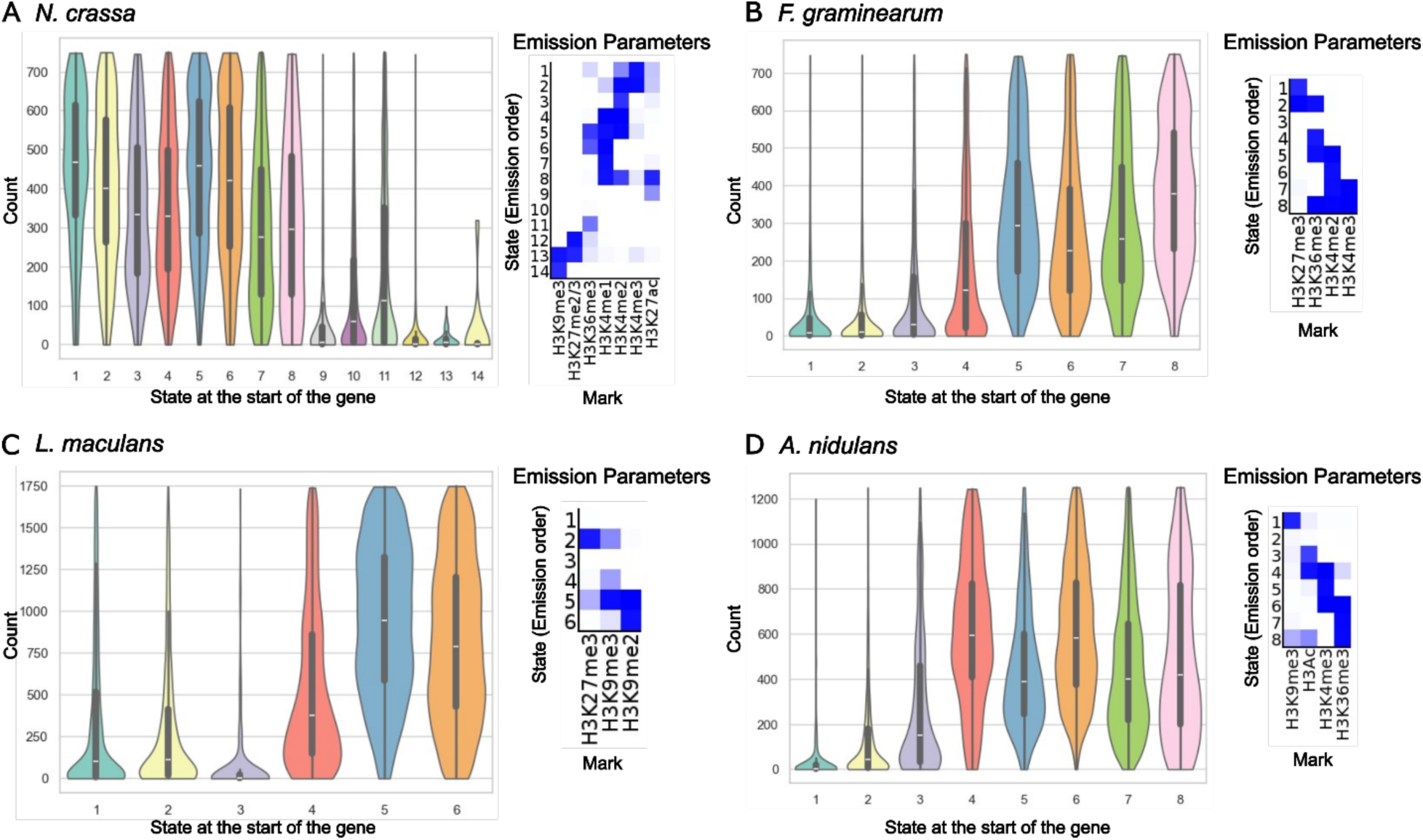
Midden Markov model ChromHMM identifies hidden states of chromatin accessibility around gene transcription start sites based on histone modification markers for (A) *N. crassa*, (B) *F. graminearum*, (C) *L. maculans*, and (D) *A. nidulans.* Violin plots show the distribution of RNAseq read counts for genes in each state. The number of states is determined by the model. to observe combinatorial correlation. Heatmaps show the probability that a histone mark will be present in that state.

Consistent with the literature, H3K4 methylation broadly associates with gene expression activity, and H3K27 methylation is expected in states with low levels of gene expression. When H3K36me3 is present alone in *N. crassa* and *F. graminearum*, it is associated with low RNA read counts, which increase when H3K36me3 is expected with H3K4 modifications (Figure 2A, B). However, in *A. nidulans*, H3K36me3 presence alone defines a hidden state, state 7, with active gene expression (200 read counts below the highest median state read count value (Figure 2D). In all species, the state with no histone markers (of the ones surveyed) is repressive. However, despite these noted similarities and differences, it is difficult to understand whether the gene states and associated marker patterns from one species would translate to another species, or whether they would correctly identify activating or repressive regimes for gene transcription.

### Shallow machine learning models perform well on intra-species but not inter-species prediction tasks

To evaluate whether complex patterns of epigenetic modifications are predictive of gene expression changes across fungal species, we tested a battery of models subject to several selection criteria and objectives, as outlined in the Methods.

Starting with established shallow learning models, we used a battery of six classifiers (Methods) to test intra- and inter-species prediction performance (see Methods and Supplemental Figure 1 & 8 for further details). We elected to use binary classification models to predict whether a gene is expected to show high or low/no expression given its epigenetic state. This is due to a number of considerations, including the read distributions seen in the ChromHMM analysis (Figure 2) and that RNA expression levels are not solely regulated epigenetics, possibly limiting sensitivity in output predictions. The binary prediction tasks were performed using each feature design strategy (Supplementary Figure 7) for peri-transcription start site (peri-TSS) epigenetic modifications in 200bp window around the TSS and averaged across reads; a 5kb window around the TSS and averaged reads, and a 5kb window with binned but not fully averaged reads, which retained more modification positional information (see Methods for further details). The tasks were also performed for each combination of the number of overlapping histone modifications (Table 3) and for each species pairing between for *N. crassa*, *F. graminearum*, *L. maculans* and *A. nidulans*.

Figure 3 shows average performance – evaluated using accuracy (top), precision (center), and AUROC (bottom) – for each of those conditions for the intra-(left) and inter-(right) species tasks. Individual classifier performance results for each test varying feature design, species specific performance, and available modification number are shown in Supplementary Figures 2 & 3.

**Figure 3.**
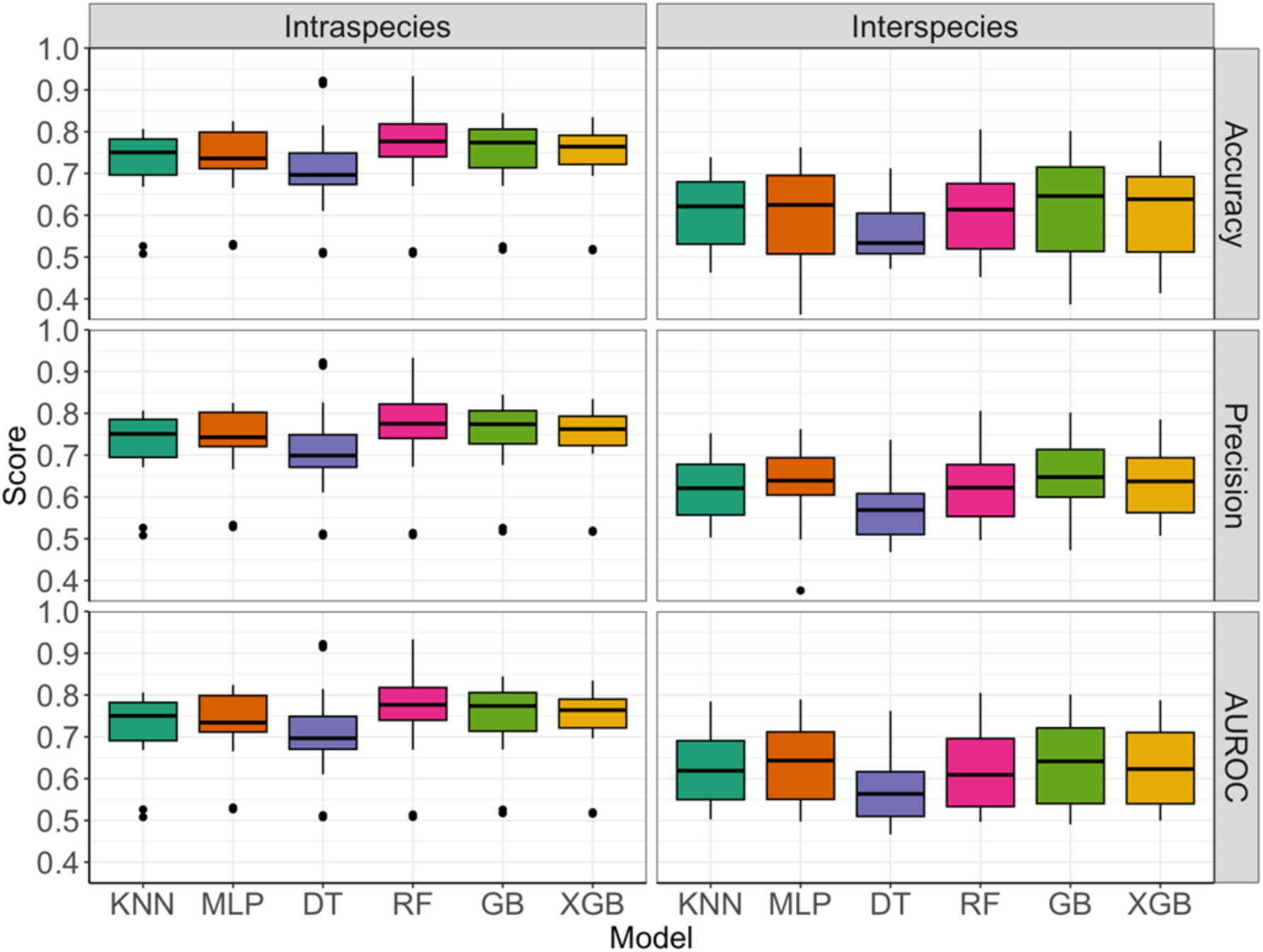
Intra-(left) and inter-(right) species prediction performance on accuracy (top), precision (center), and AUROC (bottom) metrics for the battery of shallow learning models. Results are averaged across species, TSS window, averaged signal and signal profile, and modification combinations. Boxes show median +/- quartile for n= 75 (intra) and 105 (inter) replicates. KNN: K-nearest neighbors; MLP: multilayer perceptron; DT: decision tree; RF: random forest; GB: gradient boosting; XGB: eXtreme gradient boosting.

For the intra-species prediction task, all six classification models achieved prediction accuracy, precision, and AUROC (Area Under the Receiver Operating Characteristic curve) scores between 70%-80% when the metrics were averaged across these conditions (Figure 3, left). Unsurprisingly, the intra-species classifier model performance surpassed inter-species performance across all conditions (Figure 3, right, Supplementary Figures 1, 2, 3). The Random Forest model yielded the highest intra-species performance of (77% averaged) across all three prediction performance metrics (Figure 4). Model performance decreased substantially on inter-species evaluations. Accuracy, precision, and AUROC scores ranged from 55-63% across the six classifiers (Figure 3, right). On the inter-species prediction tasks, the multilayer perceptron (MLP) and Gradient Boosting classifiers yield the best performance (Figure 3, right).

**Figure 4.**
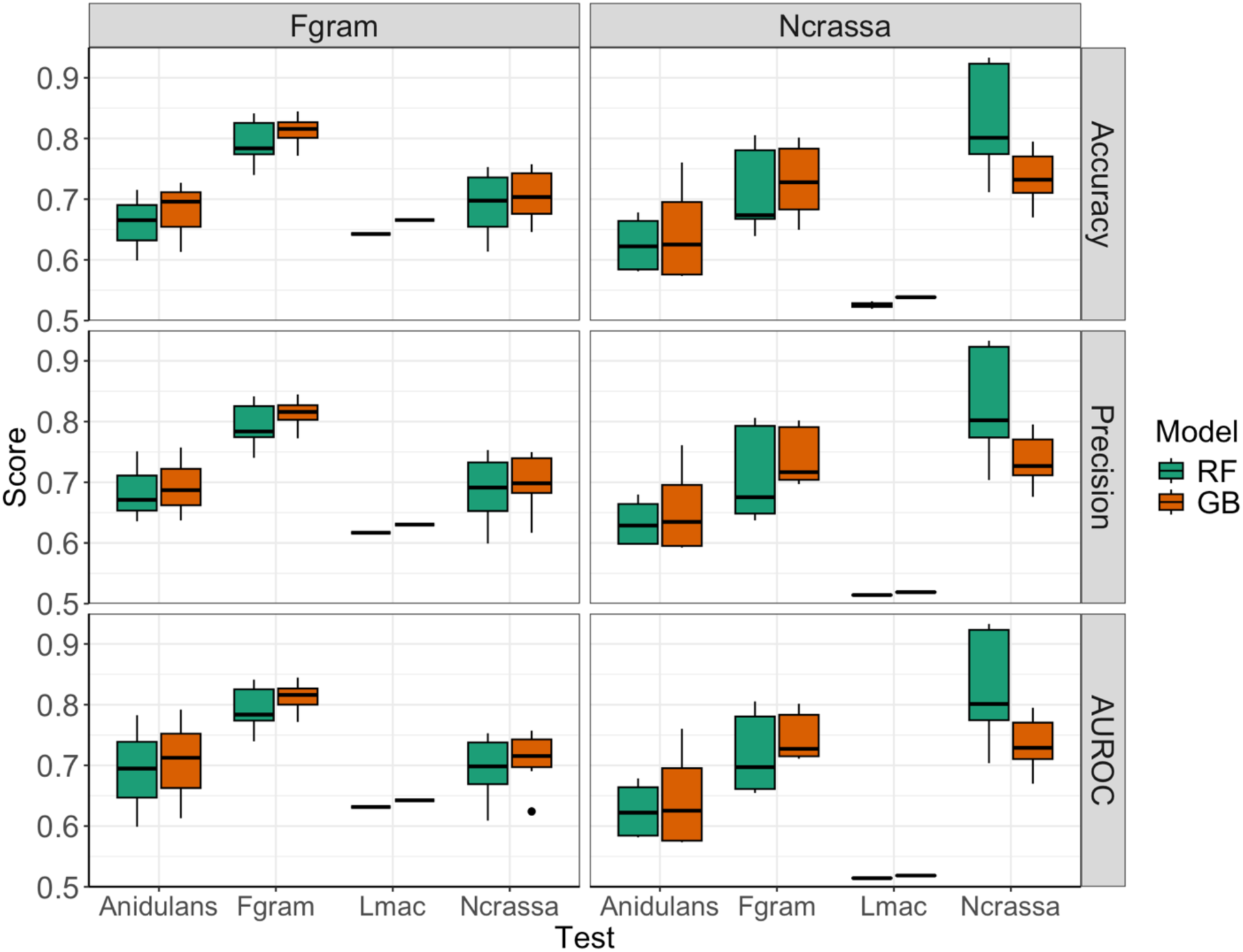
Intra-(left) and inter-(right) species prediction performance on accuracy (top), precision (center), and AUROC (bottom) metrics for the highest performing shallow learning models, random forest (RF) and gradient boosting (GB). Results are shown for each species comparison (boxes show median +/- quartile for n = 3 replicates for each TSS window, signal and signal profile, and modification combination). Species abbreviations: F. gram is *F. graminearum* and L. mac is *L. maculans*.

The number and type of overlapping modifications has some effect on performance, on average, for both the intra-, and interspecies evaluations with four modifications yielding the best performance across all the test conditions (Supplementary Figure 3 & 4).

We next sought to evaluate whether increasing the TSS window and/or using a binned signal profile rather than average signal for the ChIP-seq data would improve model performances. Increasing the TSS window (200 bp versus 5000 bp) and including more positional signal information (i.e., read signal averaged across window versus the signal profile where the reads are averaged in 100 bp bins) did not have a marked effect on model performance metrics across all models and task combinations (Supplementary Figure 3 & 5). The highest overall performance (93-95% for all metrics) is achieved with the 200 bp window average signal for the *N. crassa* intra-species evaluation using Random Forest and Decision Tree classifiers and either three or four histone modifications as features (Supplementary Figure 3). Extending the TSS window and including signal profile information does not improve intra-species predictive performance for other species nor recovers prediction power on inter-species tasks (Supplementary Figure 3 & 5).

Given the shallow machine learning models perform adequately on the intra-but not inter-species prediction tasks, the complexity of the combinatorial and positional epigenetic state may not be adequately captured by binned averages or flattened positional information of the epigenetic read signals. However, as shallow models require featurized data, we next sought to leverage deep learning methods that could use more of the raw epigenetic signal data as an input, and potentially capture more nuance in the spatial and combinatorial aspects of epigenomics.

### Developing a custom hybrid deep learning model for Model predictions Across Phylogenetic distances by Learning Expression from Epigenetics (MAPLE)

Motivated by achieving desired cross-species predictive power, and informed by the approaches taken in the DeepChrome, AttentiveChrome, and Chromoformer models^1, 24, 25^, we implemented a model with three stacks of 1D convolutional layers with batch normalization, 10-head multi-head attention layer, followed by two fully-connected layers with dropout (Figure 5).

**Figure 5.**
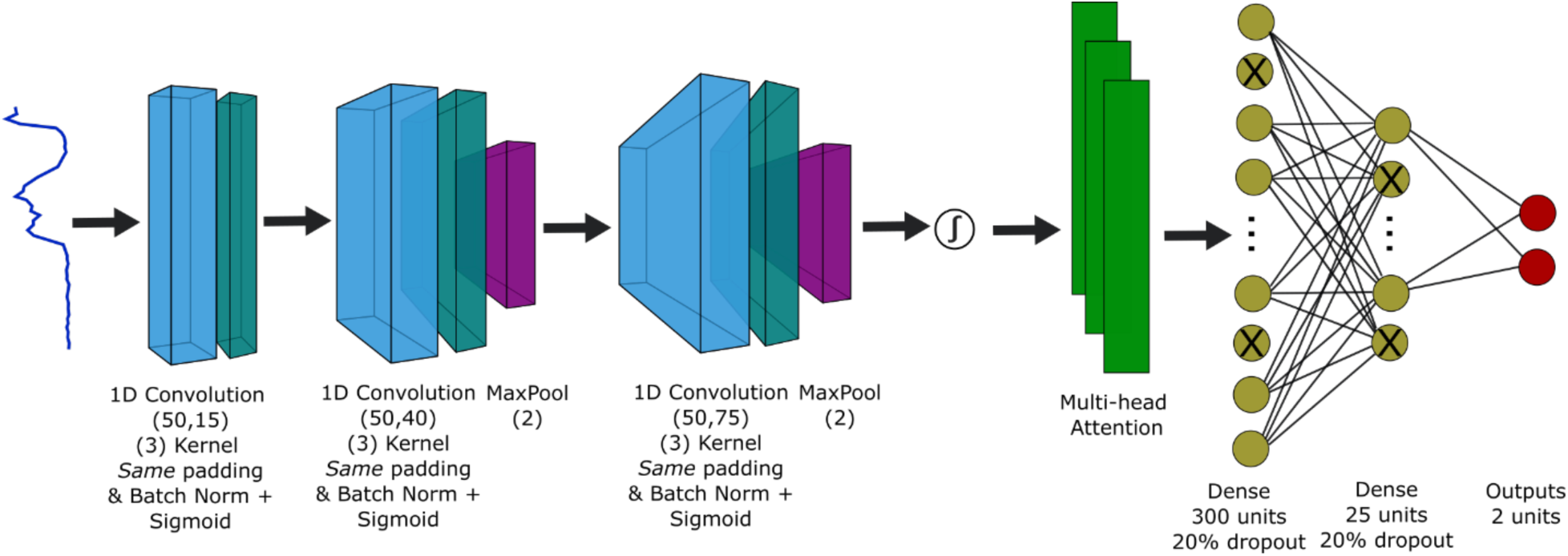
MAPLE model architecture. MAPLE accepts sequence data, here from ChIPseq epigenetic marker data, for any number of modifications. The signal profiles pass through 3 1D-convolutional layers each with batch normalization and sigmoid activation functions. Max pooling is performed on all but the first layer. Another sigmoid activation is applied before sending to a 10-head multi-head self-attention network, then through 2 fully connected dense neuron layers with 20% dropout. The model is shown with 2 output neurons as it was configured for this study, but it can be adapted to predict for any number of output classes.

This architecture combines the layers with strengths for each task we needed to implement. Convolution layers are a good way to process latent features epigenetic signal data that may comprise more useful information than a lexical tokenization approach, particularly in seeking to avoid over-fitting to any one species or modification. Multi-head attention layers help capture relationships in combinatorial spatial epigenomic profiles among multiple epigenetic markers. We did survey existing models and identified several with similar architectures for different goals, such as using genetic sequence to predict epigenetic sequence, or with similar goals but used data unavailable for fungi, such as distal cis-regulatory element interaction information. We also identified some with similar architectures to or components of MAPLE but were not readily available or functional for application in our study.

The output for this study is a binary classification prediction (Figure 5, far right). We have implemented MAPLE with two output units, representing p(0) and p(1), and perform a softmax on the output in an effort to make the model more readily extendible to greater than two output prediction categories, as the categorical cross-entropy loss function can be applied to models with two and more output neurons. The convolutional and pooling layers were selected to find representations of the sequences of epigenetic signal along the peri-TSS (+/- 5kb for the majority of work) region (Figure 5, far left). The attention heads then identify the sequences of those higher dimensional feature representations. By convention, the fully connected layers before the prediction are included to further ‘learn about and process’ the outputs of the upstream layers and taper to the result. Hyperparameters were selected using RandomizedSearchCV and are detailed in the accompanying supplemental information. Details about training strategy and output variable categorical embedding approaches are described in the Methods.

Initially, we did evaluate models with only fully connected layers and convolutional with fully connected layers, essentially only using the signal features in the epigenomic expression profile. The multilayer perceptron did not exhibit promising predictive performance across species. The model with CNN and dense layers was reasonably successful but less so than with the inclusion of attention. By then adding the attention mechanism to the model, this interspecies performance increased greatly, especially with regards to performance across evolutionary distances as inferred by Poplar (Figure 6)^32^.

**Figure 6.**
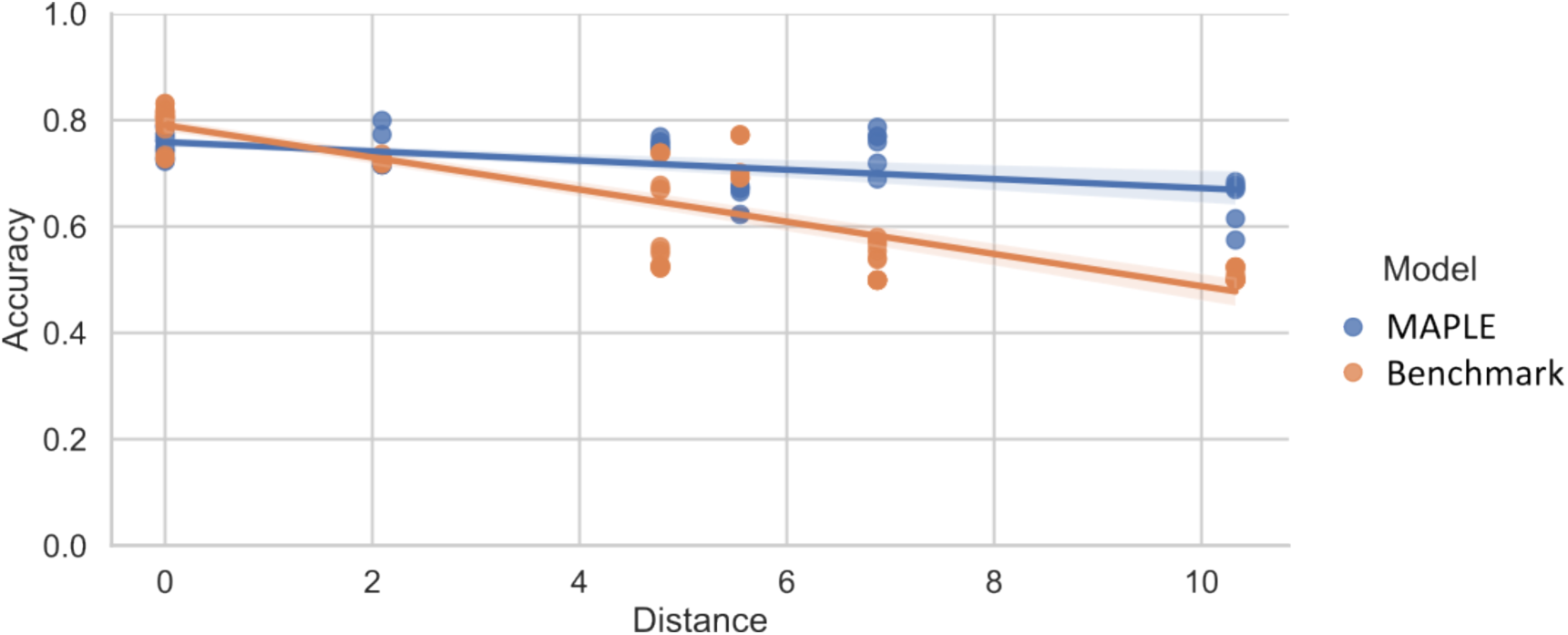
MAPLE preserves greater performance despite evolutionary distance. Scatterplot showing model prediction accuracy for MAPLE (blue) and the benchmark (orange) vs phylogenetic tree inferred distance. The interaction between distance and model is −0.022 (p-value = 2.165e-14 in a regression hypothesis test), or the benchmark model statistically significantly loses 2.2% more accuracy than MAPLE for every unit of evolutionary distance.

### MAPLE outperforms shallow and benchmarking transformer models on cross-species prediction tasks

We benchmarked our model and prediction tasks on a modified version of the Chromoformer model^1^, which in turn was benchmarked on DeepChrome^24^ and AttentiveChrome^25^. Chromoformer is a transformer-based model that incorporates three-level hierarchical histone cis-regulatory mechanisms involving core promoters and pCREs that is used for gene expression prediction tasks. Chromoformer was state-of-the-art for the task we were interested in and uses several foundational models in this domain as references. However, the model required modification for our specific purposes as the hierarchical and conformational data used by Lee *et al.* for the mammalian system studies were largely not available for fungi. In summary, we tested our model against a state-of-the-art transformer architecture.

Here again, to test the ability of the model to predict gene expression within and across fungal species using epigenetic marker data, the models were first trained on each candidate species, *N. crassa, F. graminearum, L. maculans, and A nidulans*. For intra-species prediction tasks, the models were then evaluated on withheld testing data from the same species. For cross-species predictions tasks, the model was evaluated on each of the other species in the set, and the epigenetic markers used as inputs were the maximum overlapping set. Results are averaged for different sets of markers included for intra- and relevant cross-species predictions tasks.

All prediction tasks were binary classification problems for gene expression levels, representing expression of gene or no expression. Each result was the average of three-fold cross validation where the model was run on re-split data. The prediction of gene expression by MAPLE on intra-species tasks was on par with the benchmark model. However, on cross-species tasks, MAPLE resulted in improved accuracy, AUROC, and precision over the benchmark (Figure 7). We also see that the benchmark appears to more frequently have a large imbalance between sensitivity and specificity, while MAPLE shows more even performance between these metrics (Supplementary Table 1). Additional epigenetic and species data would improve the predictive power of this modeling approach and architecture, which is extensible to the data and prediction task at hand. MAPLE’s performance began to drop for tasks where epigenetic marker overlap was low, where there were only one or two input features for the model to learn on (Figure 7, Supplementary Table 2). However, the performance broadly remained at or above that of the benchmark.

**Figure 7.**
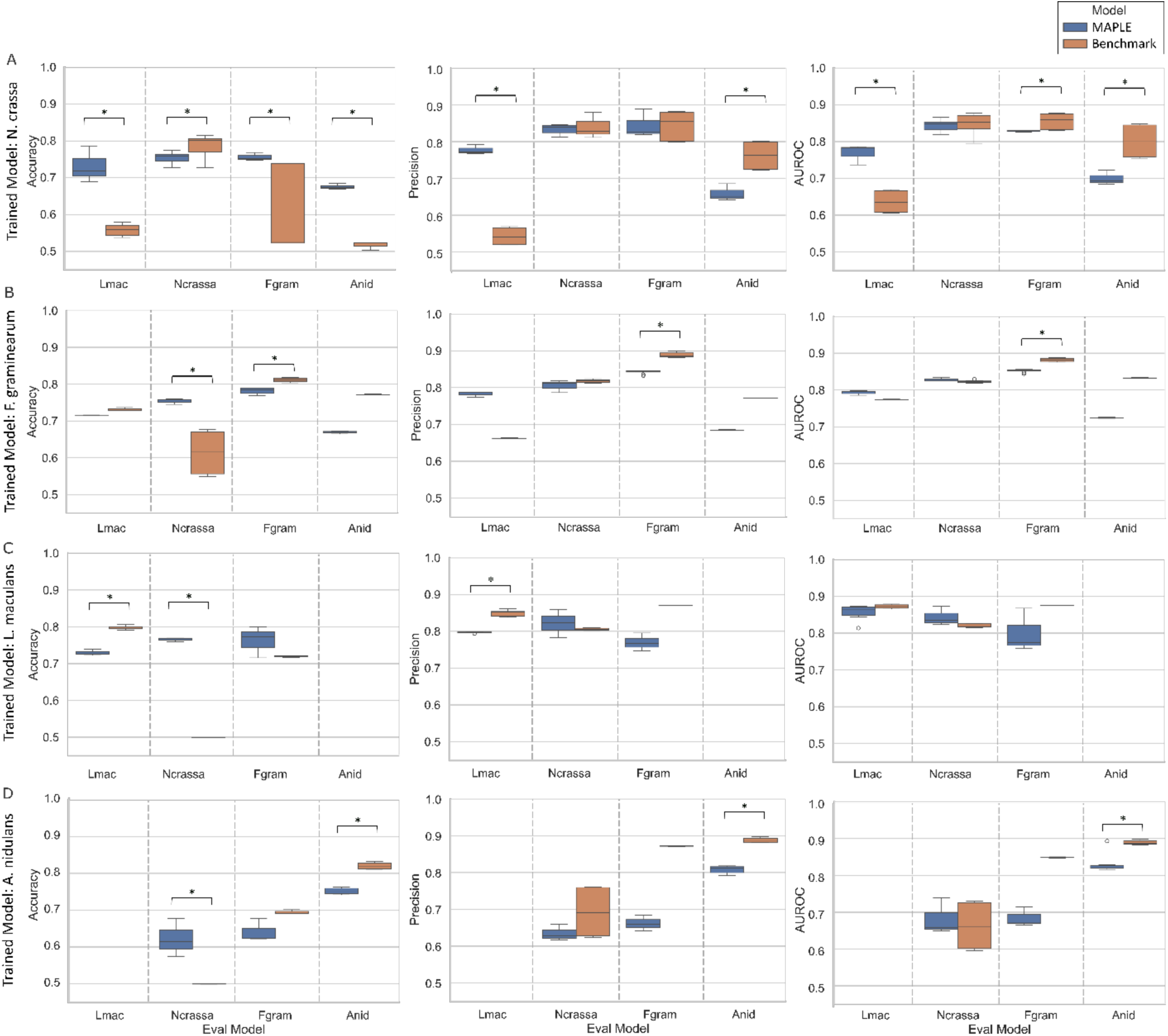
Intra- and cross-species model performance of (A) *N. crassa*, (B) *F. graminearum*, (C) *L. maculans,* and (D) *A. nidulans* trained models for each pairwise prediction task, using MAPLE (blue) or the benchmark transformer (orange) model (modified from Lee *et al*.^1^) on accuracy (left), precision (center) and AUROC (right) metrics. Repeats were 3 runs without seeding for replicates and reshuffling of test and train data. The results shown include only comparisons with at least 2 overlapping modifications. The * indicates a statistically significant difference (p < 0.05) on a pair-wise non-parametric Mann–Whitney U test. Fields without boxes are did not sufficient overlapping epigenetic modification data to train models, so performance was not evaluated. See Supplementary Table 1 for sensitivity and specificity performance metrics.

To address whether the modest performance degradation in the *A. nidulans* results was due to the number of features available or the window width of epigenetic marker signal around the TSS used by the model, we widened the window on data provided to MAPLE to 40kb, matching the benchmark. This did not markedly improve performance on the intra-species tasks and worsened the cross-species predictive performance (Table 1). This shows that while our model can improve upon the state-of-the-art for cross-species training, there are limitations to prediction power across evolutionary tree distance, and accuracy for *A. nidulans* would require more data from a species that is more closely related. One of the benefits of MAPLE is that a drop in cross-species predictive power can be used to determine where additional data is required, since we know that the architecture is inherently designed to maximally capture inter-species patterns given enough data, based on our analysis. Moreover, the majority of *A. nidulans* model performance looks to come from the inclusion of H3K36me3, as evaluated by comparing the use of all markers or only H3K36me3 on intra-species prediction (Supplementary Table 1), more so than widening the input signal window. MAPLE will accept any length of signal input but, given the significant increase of data preprocessing burden and limited performance increase, we continued with a 5kb signal window.

**Table 1.** Model performance on *N. crassa* and *A. nidulans* prediction tasks with MAPLE TSS window at original (5kb) and expanded (40kb) width. Models used features H3K36me3, H3K4me3, and H3K9me3.

|  |  | MAPLE |  |  |  | Benchmark |  |
| --- | --- | --- | --- | --- | --- | --- | --- |
| Train | Test | AUROC | Accuracy | AUROC | Accuracy | AUROC | Accuracy |
|  |  | 5kb | 5kb | 40 kb | 40 kb | 40 kb | 40 kb |
| *N. crassa* | *N. crassa* | 0.83 | 0.74 |  | 0.58 | 0.86 | 0.81 |
| *N. crassa* | *A. nidulans* | 0.7 | 0.68 | 0.75 | 0.68 | 0.76 | 0.52 |
| *A. nidulans* | *A. nidulans* | 0.82 | 0.75 |  | 0.72 | 0.89 | 0.82 |
| *A. nidulans* | *N. crassa* | 0.67 | 0.62 | 0.58 | 0.56 | 0.67 | 0.50 |

We were next interested in why MAPLE is performing above benchmark for the majority of the inter-species predictions tasks (Figure 7), particularly with *N. crassa* and *F. graminearum*, and whether it was tied to actual discriminatory power or some artifact. To inspect this, we extracted the latent states of the data after the final CNN and after the multi-head attention layer in MAPLE as well as after the multi-head attention layer in the benchmark model. We then reduced the latent space dimensionality using tSNE for visualization (Figure 8). By inspection, MAPLE better separates the data classes and has more defined latent space boundaries than after the CNN layers alone and when compared to the final benchmark model layer (Figure 8).

**Figure 8.**
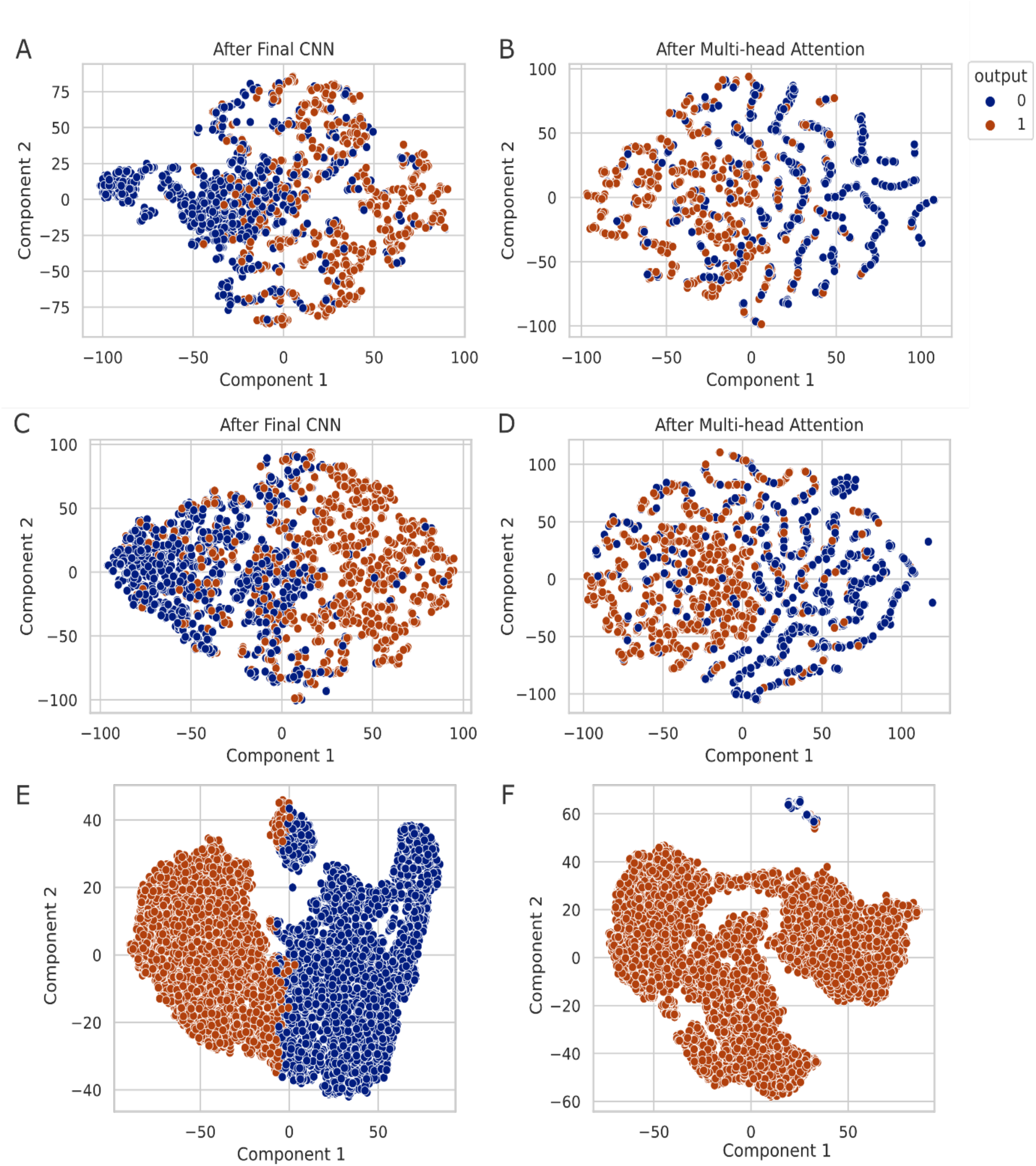
Intermediary model layer data separations of output predictions reduced to two component dimensions using t-SNE after key model layers in MAPLE and the benchmark model. t-SNE results for MAPLE intra-species prediction for (A) after the final CNN and (B) after the multi-head attention layer, and MAPLE inter-species prediction (C) after the final CNN and (D) after the multi-head attention layer. t-SNE results for the benchmark model (E) intra-species prediction and (F) inter-species prediction after the multi-head attention layer but before the dense layers. Benchmark tSNEs combine embeddings for 100, 500, and 2000 bp sequence windows. Inter-species predictions used *N. crassa* as the training species and *F. graminearum* as the testing species. Datapoint colors are model-predicted category at defined layer output; blue is a predicted value of zero, or low expression, and orange is a predicted value of one, or high expression.

### Explainability analysis identifies important and conserved features in epigenetic data

We performed SHAP (SHapley Additive exPlanations) analysis on *N. crassa* and *F. graminearum* models, which had the most overlap of epigenetic modifications, to investigate which epigenetic signal profiles were the most influential in MAPLE predictions, and to evaluate whether the signal profiles or position indices being considered across species were similar. The SHAP GradientExplainer was used to explain model predictions for a small set of genes, we inspected the SHAP importance profile (Figure 9A,B) and the corresponding signal profile (Figure 9A,B) for each epigenetic marker over the TSS window. We see that there are three genes for which H3K4 methylation markers (Figure 9A) in areas close to the TSS are highly influential in class prediction, here for class 1 or relatively high expression, while H3K27me3 and H3K36me3 are less influential in class prediction given the smaller SHAP value magnitudes for all window positions (Figure 9B). The same three genes exhibit high read signal peaks close to the TSS for the H3K4 markers (Figure 9C, pale traces), while the less influential marker signal profiles are also smaller in magnitude and noisier (Figure 9D).

**Figure 9.**
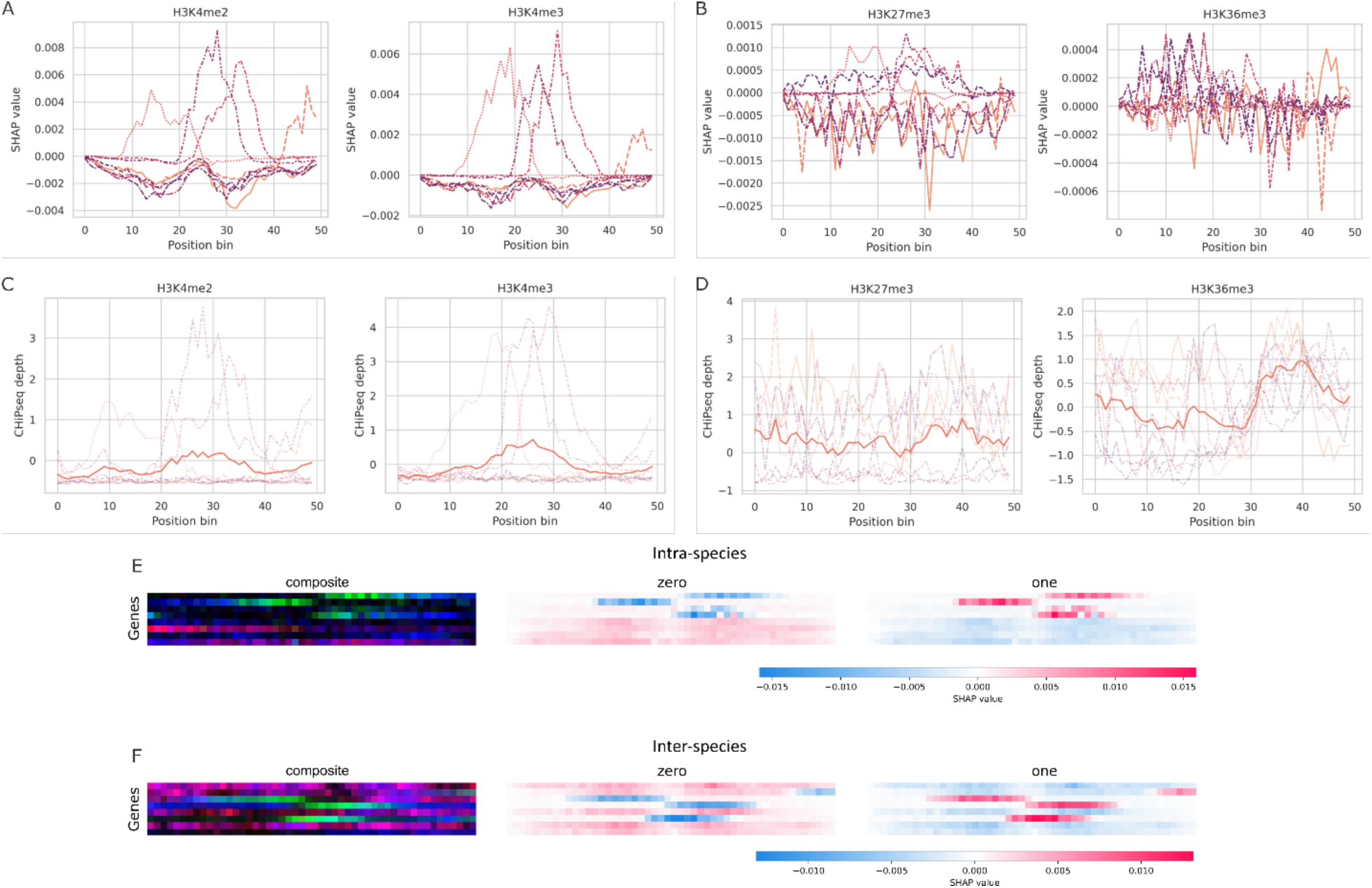
Explainability analysis using SHAP. SHAP importance values vs sequence position in TSS window for (A) comparatively influential modifications and (B) low-importance features. ChiP read depth vs sequence position in TSS window for (C) H3K4me2 and H3K4me3 and (D) H3K27me3 and H3K36me3, corresponding with categorizations in A and B, respectively. Aggregate SHAP signals for all features for each explained gene predicted for (E) intra-species and (F) inter-species tasks.

We then compared the aggregate epigenetic signal profiles’ relative importance for predicting gene class membership between intra-(Figure 9E) and inter-species (Figure 9F) evaluations. We find that the influential regions are similar between species near the TSS for a subset of the inter-species gene predictions (Figure 9E, F), which are evaluated to be in in the same class 1. These analyses can support the transfer of understanding of influential modifications patterns and corresponding signal profiles from established to less well-characterized species. The advantage of identifying specific signals that are used by the model to classify the genes that, in combination with synthetic biology tools, the information can be used to engineer the epigenome to modulate said expression^34^.

### MAPLE sees patterns in epigenomic signals that go beyond sequence

We next wanted to determine whether the common epigenomic signals recognized by MAPLE were as a result of common underlying DNA sequence. A number of studies have demonstrated that within given cell types, epigenomics can often be predicted with reasonable accuracy from underlying sequence information alone^26, 35, 36^. However, due to the dynamic nature of epigenomic regulation, cross cell-line and cross-species predictions using sequence are more challenging. Based on its cross-species predictive power, we wanted to determine whether the regions that SHAP determined as important for MAPLE are mainly driven by sequence, whether there are sequence-agnostic features of the signal that play an important role, or a hybrid of both. To this end, we used HOMER^37^, a software suite that uses established databases for motif identification, to analyze the sequences underlying regions that SHAP denoted as significant for MAPLE. We focused on the *N. crassa* and *F. graminearum* models which have the maximum number of overlapping epigenetic modifications. We performed motif finding and determined the fraction of motifs that overlap (in the top 50 and top 100 motifs) for each model when tested on the non-training species (i.e. *N. crassa* model tested on *F. graminearum* and vice versa).

We find that there are a number of overlapping motifs, which suggests that there are some sequence-specific factors that could have common binding motifs across species, which agrees with the literature highlighting examples of such conserved factors^38–40^. Importantly however, we see that anywhere between 40 – 70% of motifs identified (depending on the modification examined) are specific to a single model (Table 2). This shows that MAPLE is driven significantly by epigenetic signatures which are not necessarily linked directly to DNA sequence. On the flip side, the presence of common sequences also provides anchors for downstream epigenomic engineering across species, and using Shapley methods can point to these regions to narrow down experimental testing.

**Table 2.** Fraction of overlapping motifs discovered by HOMER for genomic regions identified by SHAP as important for MAPLE’s prediction-making.

|  | **K4me3** | **K4me2** | **K27me3** | **K36me3** |
| --- | --- | --- | --- | --- |
| **Top 50 motifs** | 0.38 | 0.54 | 0.46 | 0.36 |
| **Top 100 motifs** | 0.53 | 0.58 | 0.37 | 0.45 |
| **Right** |  |  |  |  |
| **Top 50 motifs** | 0.38 | 0.58 | 0.46 | 0.34 |
| **Top 100 motifs** | 0.45 | 0.62 | 0.5 | 0.41 |
| **Wrong** |  |  |  |  |
| **Top 50 motifs** | 0.46 | 0.52 | 0.5 | 0.36 |
| **Top 100 motifs** | 0.47 | 0.53 | 0.47 | 0.43 |

**Table 3.** ChIPseq histone modification data available for each fungal species. The X denotes that data are available for that particular species and modification.

|  | H3K27me3 | H3K36me3 | H3K4me2 | H3K4me3 | H3K9me3 |
| --- | --- | --- | --- | --- | --- |
| *A. nidulans* |  | X |  | X | X |
| *F. graminearum* | X | X | X | X |  |
| *L. maculans* | X |  | X |  | X |
| *N. crassa* | X | X | X | X | X |

The result was maintained when we examined results that the model got right versus wrong. We saw that while there were some specific differences between modifications when it came to overlapping motifs (e.g. overlapping motifs at H3K4me3 sites seem higher in wrong guesses), regardless of whether MAPLE mis-categorizes a gene, it is using features of the epigenome that are not solely correlated with underlying sequence, suggesting that there are conserved epigenome patterns across species that MAPLE can recognize (Table 2). To further investigate whether sequence alone can be equally predictive, we utilized the 7-billion parameter version of Evo2, a foundation model trained on genomic sequences, fine-tuned on each species we are investigating, and tested these fine-tuned models across species. We find that while models perform well within their respective species (intra-species task), this performance drops dramatically when tested on an outgroup (cross-species prediction task) for all pairwise comparisons (Supplementary Figure 9 & 10). Therefore, while sequence alone cannot predict gene expression outcomes, these results do highlight the potential for identification of novel factors for modulation of gene expression that can function across species, and even taxa. For instance, a few prominent motifs appear to be identified through Shapley analysis as being important in both *N. crassa* and *F. graminearum* (supplementary data). An example is the GATA-family motif, which suggests that GATA factors may be responsible for producing reproducible epigenomic patterns across the fungal kindgom, with insight from one species being directly applicable to another. Indeed, GATA family members have been shown to play critical roles across fungal species ranging from light response, development, virulence and metabolism^41–43^. Other such biological insights can be gleaned from the explainability analysis, and tested across species.

## Discussion

The primary goal of this work was to understand whether fungal epigenetic modification information could be used to predict gene expression within and across fungal species, to enable transferring information about well characterized species to poorly understood species. This could be leveraged to facilitate engineering of epigenomes across fungi by repurposing information of influential epigenomic patterns from related species. This insight would provide controls to optimize new fungal chassis that possess an attribute of interest, or to defend against novel fungal pathogens by targeting their epigenomes and engineer new synthetic biology tools to achieve these goals. To establish how well epigenetic data can be used to predict gene expression across species, we evaluated the predictive performance in a battery of machine learning models, ultimately finding a custom deep learning model that performed well on tasks that support these goals.

Here we demonstrate the mixed convolutional and multi-head attention model that performs well not only on the prediction of intra-species fungal gene expression from epigenetic modification signal profiles but also on the same task but across fungal species. Shallow learning models that use featurized epigenomic data are not sufficient to accurately predict gene expression from epigenomic data in cross-species prediction tasks. MAPLE outperforms a transformer model benchmark, which was based on the Chromoformer architecture but heavily modified to remove the distal cis-regulatory element components, as we did not have access to enough high-quality 3D interaction data across the multiple species chosen for this study. While there is more information about epigenomic regulation in fungi at cis-promoters and enhancers, we know that there are 3D interactions in fungal chromatin that can also influence gene expression^44, 45^. With increasing amount of 3D data becoming available, future work on MAPLE can incorporate this 3D information as another layer of cross-species predictive power, as was done with Chromoformer^1^. Additionally, MAPLE excels with regards to the computational and resource requirements needed to run it, as it can be trained and run on 2.0 or 2.4 GHz CPUs with an typical training time of 4-5 minutes and evaluation time of 12.3 seconds (varies with number of predictions), while the benchmark model requires GPU and has an average training time of 49 minutes and evaluation time of 29.3 seconds.

In MAPLE, we can interpret that the CNN layers extract and encode features in the epigenetic signal data that can then be operated on by the self-attention layers for contextual significance. We posit that the novel model architecture was able to perform well on the prediction tasks by capturing both the local patterns within the epigenomic data, through the convolutional layers, and the global dependencies between distant genomic regions, via the multi-head attention mechanism, without overfitting to the training data of a single species. This dual approach allowed our model to effectively learn from the complex, spatially distributed features characteristic of epigenetic data. Our dimensionality-reduction analysis of the embedding space within each model type shows significant difference in how the data points are being separated by the different deep learning architectures. We see that tokenization of epigenomic data (as seen by looking at the tSNE projection of the embeddings) after the attention layers in the benchmark transformer model separates the outputs less effectively than the CNN-embedded attention layer output in MAPLE (Figure 8). We therefore conclude that while tokenization can be very valuable for intra-species and even intra-cell type predictions, during inter-species prediction, the hybrid model is more successful at separating classes in its embedding space.

We demonstrate that by using the explainability tool SHAP, we can identify influential regions of epigenetic signal seen by MAPLE responsible for inter-species accuracy, across each epigenetic modification. There are multiple benefits to having explainability in a machine learning model. First, we can use biological precedent to verify that the model is extracting data from expected regions. For example, we show in Figure 9 that while H3K4me2/3 impact is found in a relatively narrow range of the region on average, H3K27me3 and H3K36me3 impacts are spread over a wider range. This is in line with the functions of H3K4me2/3 closer to promoter regions, and H3K27me3 and H3K36me3 across larger stretches of genome, both in fungi and in mammals^46–48^. In addition, identifying regions of genome that MAPLE considers important for epigenomic regulation of gene expression can allow for development of targeting strategies for engineering the epigenome. In a scenario where one is interested in using a new or non-model species for a given property, MAPLE can support using species data for which there is ample information to train the model and then evaluate the model on the new species and extract the epigenetic marker data in influential regions. This also provides potential for identifying DNA sequence motifs to target individual transcription factors that might be expressed in specific species, which could in turn affect a number of epigenomic marks simultaneously and at many genes^49, 50^. This strategy can be used together with or in addition to specifically targeting epigenetic enzymes to genomic regions^34, 51, 52^.

MAPLE represents, to our knowledge, the first demonstration of any model able to predict gene expression outcomes of one fungal species using a model built and trained for another. As such, we propose a use-case for MAPLE in a novel species where only the genome is known, and the goal is to use synthetic biology to impact gene expression. The use of a tree-building pipeline such as Poplar^32^ will demonstrate the closest MAPLE model or organism that has sufficient data for predictive power. Synthetic epigenomic data for the novel organism in question can be used to determine which combination of epigenomic modifications will result in the desired outcome, and based on the SHAP analysis outlined above, individual regions of interest for targeting can be identified and tested. Universal patterns of gene expression via epigenetic regulation provides a cross-species guidebook for fungal chassis performance and potentially for discovering broadly applicable and anti-treatment resistant antifungals. Computational hypothesis generation for novel target identification and ideal states reduces the number of experimental conditions that need to be tested, streamlining the path to biomanufacturing and bioproduction optimization and novel therapeutic development.

### Limitations of the study

We recognize that additional areas of study in the future can strengthen and improve on this current study. Future work will need to address a number of outstanding research directions that are beyond the scope of this initial study but address critical follow-up questions. In addition to finding novel fungal strains and testing the use case of MAPLE outlined above, the issue of limited data for epigenetic modifications shared among all the species we sought to use needs to be tackled to make MAPLE reach more broadly across species. For example, while MAPLE has superior performance overall where benchmark models are prone to overfitting, there are some exceptions (eg. precision of *A. nidulans* models predicting *N. crassa*). We postulate that the cause of this is related to the volume and quality of epigenetic data available. Specifically, there were very few overlapping modifications between *A. nidulans* and most of the other species, significantly reducing the amount of input data the model was trained with. Despite this, MAPLE performed either on par or better than benchmark models for most metrics. Additional work would focus on expanding the range of epigenetic modifications included in the model and integrating additional data types to further enhance predictive power and applicability. With more modifications included as channels in the model, perhaps yet better predictive performance could be achieved. Second, we did not include any positional information ahead of the attention layer, which could be a useful avenue to explore in the future. Third, exploring various resolutions of signal (including binning at various window sizes) could be impactful as sequencing technologies evolve, which we hope to address with additional data sets being created on novel platforms and iterations of next generation sequencing. Finally, we can incorporate a weighting system based on evolutionary distances from a program such as Poplar^32^ (as an alternative to allowing users to incorporate their own evolutionary distance analysis).

Another direction to build upon this work includes exploring other new frontier models, varying the peri-transcription start site window size, and using more data types such as CUT&RUN/CUT&Tag, methylation profiling and accessibility profiling such as ATAC-seq as model inputs^53–55^. This will not only add additional data, but also increase the depth of information provided to the model, likely contributing to further increases in accuracy and applicability across evolutionary distance. As technology continues to improve, research should focus on using single-cell technologies to significantly increase the amount of transcriptomic and epigenomic data available for training future models. MAPLE and its findings can inform experimental work seeking to identify epigenetic targets to dynamically and precisely control fungal gene expression in an array of species. Additionally, beyond fungi, MAPLE can be trained on data sets from other eukaryotic organisms such as plants, algae, protists and non-mammalian animals to help bridge epigenomic regulation data gaps across evolution. Further, while DNA sequence is not the only determining factor in gene expression prediction, there is clearly a strong sequence-specific component at play, which provides fertile ground for investigating the relationship between DNA sequence, epigenetic modification and gene expression across phylogenetic distances in fungi and other organisms. Future work will incorporate modules within MAPLE that can bring in DNA sequence, which will likely entail incorporating existing software in this domain such as Enformer^35^ and connecting physiological traits to epigenomic features as has been shown for genomic features in subsets of fungi^56^. In addition, we could use large language models to identify conserved motifs based on the SHAP analysis from MAPLE ^57^. Recent work has shown that these models can identify cross-species motifs, particularly in fungi, which would be an actionable module to implement downstream of MAPLE ^58^.

## Star Methods

### METHOD DETAILS

#### Dataset Collection

Gene expression and epigenetic modification datasets for each species were chosen based on the following criteria: (a) the peer-reviewed study had ChIPseq histone modification data and had corresponding gene expression data (RNAseq), (b) samples were sequenced on the Illumina platform, (c) data were available from the Sequence Read Archive (SRA) at NCBI, and (d) the project included samples with WT and untreated, unconditioned samples. We identified publicly available datasets that satisfied these criteria, as well as comprise a range of biological functions and span evolutionary distance, for four fungal species: *Neurospora crassa* (*N. crassa*), *Fusarium graminearum* (*F. graminearum*), *Leptosphaeria maculans* (*L. maculans*) and *Aspergillus nidulans* (*A. nidulans*). In the case of the *N. crassa* data, the RNAseq data are from JGI with matched treatment conditions, and validity was assessed using a clustering correlation analysis. Work conducted in this study used the data from the *L. maculans* lepidii group. All accession numbers provided in the Key Resources Table in STAR Methods.

#### Sequence Data Pre-processing

Data were downloaded using the Prefetch software^59^. Downloaded fastq files were quality checked using FastQC^60^. Illumina adapters were trimmed, and low-quality sequences were removed using Trimmomatic^61^ (paired end, LEADING: 3, TRAILING: 3 SLIDINGWINDOW: 4:15, MINLEN: 36). RNAseq data were aligned using STAR^62^ (paired end, ran with defaults except, --runThreadN 20), and CHiPseq data were aligned using Bowtie2^63^(paired end, ran with defaults except, -p 20). PCR duplicates were removed from aligned SAM files and the files were subsequently converted to BAM files using SAMtools^64^using the default parameters. Read counts for RNAseq data were assigned to genes using featureCounts^65^using default parameters. Read counts were then TMM-normalized (trimmed means of M-values) using the python package conorm. Bedtools genomecov^66^ (run with defaults except, - scale [number of reads per sample* 0.0000001]) was used to compute the per-base depth of ChIPseq feature coverage. This pipeline is outlined in Supplementary Figure 6. We used the publicly available fungal reference genomes that were used in the original publications that the data were acquired from for alignment^67–71^.

#### ChromHMM Usage

We re-aligned the histone modification files by running ChromHMM’s BinarizeBam command, followed by the LearnModel command^33^. We selected the number of emission states to be double the number of histone modifications per species. Then the ChIPseq output from ChromHMM containing the emission states was merged with normalized RNA-seq read counts of each coding sequence (CDS) to demonstrate a relationship between chromatin behavior upstream and gene expression downstream. We selected the nearest chromatin state at least 500 base pairs upstream from the start codon of the CDS. The output of ChromHMM shows how each chromatin state corresponds with levels of downstream gene expression.

#### Matching ChIPseq Modifications Data

The data used in this study were selected based on species relationship proximity (Figure 1), availability of both epigenetic and RNA expression data, and the greatest possible extent of shared histone modifications. However, the availability of fungal epigenetic sequencing data varies across species. As such, we identified three strategies to learn about the relationships between these features and gene expression when predicting across species: 1) use all features available for the species used to train the model and treat the unshared species as empty or null data for the cross-species predictions; 2) use only the shared features to train the model and evaluate cross-species predictions; and 3) to select only maximally important features determined through SHAP methods in the intra-species model training and evaluation. The third is limited by the fact that it is only useful when the maximally important feature is shared across species of interest. For the work in this study, we employed the second strategy unless otherwise noted. The modification data present for each species used to develop the models in this study are shown in 4.

#### Criteria for Model Development

Outlined below are the criteria that informed model development approach and selection: **Effective:** Sufficiently high performance on performance evaluation metrics (mean absolute error (MAE) and regression area under the receiver operating curve (AUROC) for regression models and accuracy, precision, and/or AUROC for classifier models). Sufficient performance will be subject and application specific. Here, we considered an accuracy score at or above 80% and an AUROC at or above 85% a good performer based on benchmarking model performance on related tasks and in considering the noise introduced with fungi rather than mammalian tissue as subjects^72^. Similar levels were considered good for inverse mean absolute error.

#### Explainability and Interpretability

Maintaining a high ability to find causal factors in predictions, to interpret and identify decision logic, and/or easily obtain parameters with physical relevance is highly desirable for applications concerned with trustworthy AI and AI security and to inform experimental tools design or treatment strategies^73^. This factor is related to model parsimony.

#### Economical and Efficient

Satisfying this criteria has many potential positive impacts. The model is able to train and run faster. Reduced energy consumption helps reduce the environmental burden. The option to run on CPU rather than GPU may mean that the model is more widely and more equitably available to interested researchers. A lighter model – running faster, on CPUs, or both – could make the model more relevant for real time and deployed applications in industry processes. This factor is related to model parsimony.

#### Parsimony

All else being equivalent, parsimonious models win out. In other words, models that use the fewest parameters to achieve desired accuracy in explaining outcomes are favored. Model selection methods like information criteria^74^ methods can guide model performance evaluation^75^. This has become less of an issue as datasets have become massive, but remains useful in principle.

### Feature Preparation, Data Splitting, and Output Variable Categorization

GFF3 files for each species were converted to bed files using gff2bed. From those files we extracted gene coordinates [start, stop], chromosome, and gene name, for 200 bp and 5000 bp windows around transcription start sites (TSS). From there, we used that information to extract the feature coverage for these windows from the ChIPseq data. For the shallow learning models we took the average depths across each of the windows (Supplementary Figure 7). In addition, for the 5000 bp window we also took the average read depth in 100 bp bins across the window, which was used for both shallow learning models and MAPLE. In the shallow learning models, this was included as additional independent feature without any explicit ties to the contiguous signal profile. This binned signal profile was used for MAPLE but maintained its context in the 3-dimensional tensor structure.

For the predicted variable, the RNAseq expression data, we used the quantile-based qcut function from the pandas module in python to perform dynamic binning of the RNAseq data, which assigns an equivalent number of samples in each class, which for the majority of findings presented here are binary. We explored between 3 and 10 class outputs as well, but this did not improve performance.

### Modeling Training and Evaluation Schema for Intra- and Inter-Species Prediction Tasks

We use intra-species prediction to describe training, validation and optimization, and testing or evaluation on the same species (Supplementary Figure 8). Inter-species or cross-species prediction describes when the model is trained and optimized on one species, then tested or evaluated (used to predict gene expression) in a second species (Supplementary Figure 8). For MAPLE, optimization epochs used 10-fold cross validation. Test data were withheld from training epochs. 20% of data were split into test data for intra-species tasks. For inter-species tasks, since the data were not needed for training, 75% of data were used for evaluation.

### Details of Shallow Learning Models

The following nine supervised-learning regression models from the scikit learn python module were evaluated: K-nearest neighbors, Multi-Layer Perceptron, Support Vector Machine, Decision Tree, Random Forest, Gradient Boost, eXtreme Gradient Boost, Dummy regressor, and Linear regressor (Supplementary Figure 1).

The following six classification models from the scikit learn python module were evaluated: K-nearest neighbors (KNN), Multi-Layer Perceptron (MLP), Decision Tree (DT), Random Forest (RF), Gradient Boost (BG), and eXtreme Gradient Boost (XGB). We first ran all models using the default parameters in triplicate (changing the train/test split) for each comparison.

Modification overlap dictated which intraspecies evaluations could be conducted, and *F. graminearum* and *N. crassa* were the only species to overlap on four modifications in our dataset. We performed Bayesian optimization on all models with *F. graminearum* data across four modifications with marginal improvement in performance metrics.

### Feature Preparation and Model Details for Benchmark Model

A multi-head attention transformer modeled from the previously published Chromoformer^1^ was trained with fungal epigenomic data. The Chromoformer architecture was modified for this purpose – specifically, Chromoformer has a ‘pairwise transformer’ that is used to model 3-dimensional interactions of DNA, which was removed as we are not using any 3D interaction information. Instead, the embedding transformer was directly connected to a regulation transformer and a 10-fold cross validation and hyperparameter optimization module was added using Optuna^76^. The parameters optimized were learning rate and weight decay in the AdamW optimizer, and gamma, the multiplicative factor applied to the learning rate after every epoch. We utilized the Gaussian process-based algorithm implemented in <u>GPSampler</u> option to search our parameter space, defined as lr=(1e-6, 1e-2), l2=(1e-6, 1e-1), and gamma=(1e-2, 1). Features were used according to the specifications of Chromoformer^1^. Hyperparameters were optimized using Optuna (details in Supplementary Information).

#### Fine-tuning Evo2

We used the pretrained EVO2-7B model as a frozen feature extractor, extracting embeddings from all transformer blocks. A trainable classification head (convolutional layer, EfficientNet-B0, dropout, and linear classifier) was added for binary classification. Stratified 5-fold cross-validation was performed on genomic sequences in a 200bp or 5kb window around the transcription start site from a single species (labeled as either expressed or not). AdamW optimization (lr=1×10⁻⁴, weight decay=1×10⁻⁴), cross-entropy loss, batch size 16, and early stopping (patience=15 epochs) were used. Dropout (p=0.3) and gradient clipping provided regularization. Each fold was saved as a separate model. Validation was done with a leave-out dataset from the training group, and testing was performed across species that were not seen during training (as with MAPLE and other models). Performance metrics are reported as mean ± SD across models.

## QUANTIFICATION AND STATISTICAL ANALYSIS

Figures were either plotted in Python or R. For statistical analysis, data were analyzed using the scipy stats package. For Figure 7, statistical significance was evaluated with a pair-wise non-parametric Mann-Whitney U test; p-values < 0.05 were considered significant.

## Summary of Supplemental Information

### Supplementary Information

- Supplementary Figure 1 - Intra-species shallow regressor model battery performance using N. crassa on mean absolute error and regression AUROC metrics (related to Figure 3)
- Supplementary Figure 2 - Confusion matrices and metrics output table show K-nearest neighbors classifier performance across (A) 2, (B) 3, (C) 4, (D) 5 output bins for intra-species predictions using N. crassa data (related to Figure 3)
- Supplementary Figure 3 - Detailed shallow classifier model prediction performance results for each combination of species, model, TSS window, signal vs averaged features, and class output number for both inter- and intra-species prediction (related to Figure 3)
- Supplementary Figure 4 - Average shallow learning classifier performance for each number of overlapping modifications available for use in test for intra-(left) and inter-(right) species predictions (related to Figure 3 and Figure 4)
- Supplementary Figure 5 - Feature design impact on prediction performance. Average shallow learning classifier performance for intra-(left) and inter-(right) species predictions using varying feature input approaches (related to Figure 3 and Figure 4).
- Supplementary Figure 6 – Data pre-processing pipelines (related to Figures 5, 6 and 7)
- Supplementary Figure 7 – Feature design strategy (related to Figures 5, 6 and 7)
- Supplementary Figure 8 – Machine learning training and evaluation strategy (related to Figures 5, 6 and 7)
- Supplementary Figure 9: Model performance of Evo2, a state-of-the-art *genomic sequence only* model for performance comparison (related to Figure 7), with 5kb TSS window inputs.
- Supplementary Figure 10: Model performance of Evo2, a state-of-the-art *genomic sequence only* model for performance comparison (related to Figure 7), with 200bp TSS window inputs.
- Supplementary Table 1 – MAPLE vs. Benchmark model intra- and inter-species prediction specificity and sensitivity (related to Figures 6 and 7)
- Supplementary Table 2 – MAPLE intra-species prediction performance for *A. nidulans* using modification subsets and varying TSS windows (related to Figures 6 and 7) Supplemental Data – outputs from HOMER for SHAP explainability analysis (related to Figures 8 and 9)

## Acknowledgements

We would like to thank Chris Ebsch, Emily Hollister, and Michael Darling for critical review of the manuscript. This work was supported by the Laboratory Directed Research and Development program at Sandia National Laboratories, a multi-mission laboratory managed and operated by National Technology and Engineering Solutions of Sandia, LLC, a wholly owned subsidiary of Honeywell International, Inc., for the US Department of Energy’s National Nuclear Security Administration under contract DE-NA0003525.

## Author information

### Contributions

L.W. and R.K. conceptualized and designed the study. L.W., J.S., A.F., C.K., W.M., and R.K acquired, processed, and performed preliminary analyses of data. L.W. and W.D. conceptualized model architecture. L.W., J.S., C.K., E.L, E.K., and R.K performed model development, model implementation and data analysis, and data interpretation. L.W. and W.M. productionized data analysis and model code. L.W, J.S., and R.K. wrote the manuscript. All authors reviewed and approved the manuscript.

Corresponding authors: Correspondence to Laura Weinstock or Raga Krishnakumar

### Ethics declarations

Competing interests: The authors declare no competing interests.

### Supplementary Information

Supplementary figures and tables provided in accompanying Supplementary Information file.

### Data Availability

All source data are available on NCBI at the project numbers provided in The Key Resources table in STAR Methods. ChIPseq histone modification data available for each fungal species (Table 3). Processed data and trained models can be found with the code on https://github.com/sandialabs/MAPLE.

### Code Availability

Source code available at https://github.com/sandialabs/MAPLE.

