## Supplemental dataset for "Hybrid CNN and Multi-Head Attention Model for Analyzing Epigenetic Mechanisms and Gene Expression Across Fungal Phylogenetic Distances": FgramModel_NcrassaTest_K4me2locs_homerResults.html

/projects/wg-feeds/SHAP/FgramModel\_NcrassaTest\_K4me2locs\_SHAP\_noDup\_HOMER// - Homer de novo Motif Results


### Homer *de novo* Motif Results (/projects/wg-feeds/SHAP/FgramModel\_NcrassaTest\_K4me2locs\_SHAP\_noDup\_HOMER//)

Non-redundant Motif File of Results  
Known Motif Enrichment Results  
Gene Ontology Enrichment Results  
If Homer is having trouble matching a motif to a known motif, try copy/pasting the matrix file into
STAMP  
More information on motif finding results: HOMER
| Description of Results
| Tips
  
Total target sequences = 32283  
Total background sequences = 145822  
\* - possible false positive  

|  |  |  |  |  |  |  |  |  |
| --- | --- | --- | --- | --- | --- | --- | --- | --- |
| Rank | Motif | P-value | log P-pvalue | % of Targets | % of Background | STD(Bg STD) | Best Match/Details | Motif File |
| 1 | T C A G T C G A T C A G T C A G C T G A T A C G T C A G C T G A T C A G T C A G C T G A T A C G | 1e-5562 | -1.281e+04 | 57.28% | 13.51% | 716.8bp (793.5bp) | TF3A(C2H2)/col-TF3A-DAP-Seq(GSE60143)/Homer(0.794) More Information \| Similar Motifs Found | motif file (matrix) |
| 2 | A G C T A C G T A T C G A C G T A G C T T A C G G C A T A G C T A T C G G C A T A G C T A C T G | 1e-5408 | -1.245e+04 | 47.30% | 8.23% | 695.0bp (765.3bp) | ZML2(C2C2gata)/col-ZML2-DAP-Seq(GSE60143)/Homer(0.774) More Information \| Similar Motifs Found | motif file (matrix) |
| 3 | A G C T A G C T A G C T A G C T G A C T A G C T A G C T A G C T A G C T A G C T A G C T A G C T | 1e-4050 | -9.327e+03 | 39.87% | 7.69% | 668.5bp (829.2bp) | VRN1(ABI3VP1)/col-VRN1-DAP-Seq(GSE60143)/Homer(0.920) More Information \| Similar Motifs Found | motif file (matrix) |
| 4 | A G T C G A C T C A T G A G T C A G C T C A T G A G T C A G C T C T A G A G T C A G T C C T A G | 1e-2774 | -6.388e+03 | 61.55% | 27.78% | 720.7bp (732.1bp) | ZNF93/MA1721.2/Jaspar(0.803) More Information \| Similar Motifs Found | motif file (matrix) |
| 5 | T A G C A G T C C G T A C A G T A G T C C G A T A G C T A T G C | 1e-2581 | -5.944e+03 | 75.89% | 43.00% | 722.8bp (763.8bp) | Yy1/MA0095.4/Jaspar(0.776) More Information \| Similar Motifs Found | motif file (matrix) |
| 6 | A G T C T A G C C G A T A G C T A T C G C G T A G A C T T A C G C T G A A G T C T A G C C G T A | 1e-2556 | -5.888e+03 | 58.41% | 26.27% | 725.2bp (742.7bp) | ARF7/MA1698.2/Jaspar(0.695) More Information \| Similar Motifs Found | motif file (matrix) |
| 7 | A T G C G A C T A G C T A T C G T C G A T A C G A T G C G C A T A G C T A T C G | 1e-2479 | -5.709e+03 | 67.31% | 34.83% | 717.6bp (752.8bp) | NR6A1/MA1541.2/Jaspar(0.810) More Information \| Similar Motifs Found | motif file (matrix) |
| 8 | T C A G T C G A G C A T C A T G C T A G C G T A C G A T C A T G C T A G C T G A | 1e-2222 | -5.117e+03 | 62.59% | 32.01% | 721.8bp (757.6bp) | HOXA1(Homeobox)/mES-Hoxa1-ChIP-Seq(SRP084292)/Homer(0.814) More Information \| Similar Motifs Found | motif file (matrix) |
| 9 | A G C T A G T C C T A G C T A G G A T C C T A G A T G C A G T C T C A G A T C G G A T C T C A G | 1e-2081 | -4.794e+03 | 63.38% | 33.68% | 719.1bp (720.6bp) | ERF8/MA0994.3/Jaspar(0.723) More Information \| Similar Motifs Found | motif file (matrix) |
| 10 | C T A G C T A G C T G A C T A G C T G A C A T G C T G A C A T G C T A G C A T G | 1e-1609 | -3.706e+03 | 44.14% | 20.24% | 707.2bp (769.0bp) | SeqBias: G/A bias(0.743) More Information \| Similar Motifs Found | motif file (matrix) |
| 11 | C T A G G T C A G T A C C G T A G C A T A C T G C A T G G A T C T A G C T C G A | 1e-1561 | -3.595e+03 | 69.23% | 43.41% | 710.1bp (744.4bp) | Tv\_0259(RRM)/Trichomonas\_vaginalis-RNCMPT00259-PBM/HughesRNA(0.715) More Information \| Similar Motifs Found | motif file (matrix) |
| 12 | A C T G A C G T A C G T A C T G A T C G G A C T A C G T A C T G | 1e-1450 | -3.339e+03 | 62.97% | 38.02% | 719.3bp (758.2bp) | NAC037/MA2047.2/Jaspar(0.842) More Information \| Similar Motifs Found | motif file (matrix) |
| 13 | T A C G C G T A A G C T T A C G A G C T A T G C A C T G G T C A | 1e-1391 | -3.205e+03 | 54.60% | 30.64% | 721.0bp (740.0bp) | XBP1/MA0414.2/Jaspar(0.844) More Information \| Similar Motifs Found | motif file (matrix) |
| 14 | A C T G T C G A C A G T C T G A A T C G T A C G G C A T C G T A A T C G T A C G A G C T G T C A | 1e-1207 | -2.780e+03 | 11.90% | 1.88% | 669.8bp (652.4bp) | PK06182.1/MA2354.1/Jaspar(0.732) More Information \| Similar Motifs Found | motif file (matrix) |
| 15 | T A G C A C G T C G T A A C T G C G T A A C T G A C T G A C G T C G T A A T G C | 1e-1134 | -2.613e+03 | 11.02% | 1.70% | 689.9bp (702.9bp) | MOT3/Literature(Harbison)/Yeast(0.651) More Information \| Similar Motifs Found | motif file (matrix) |
| 16 | G T A C T G C A C G T A G T C A T C A G A G T C G T A C C T G A | 1e-1060 | -2.442e+03 | 59.30% | 37.98% | 716.2bp (759.9bp) | Nr5a2(NR)/mES-Nr5a2-ChIP-Seq(GSE19019)/Homer(0.723) More Information \| Similar Motifs Found | motif file (matrix) |
| 17 | A C G T A C G T A G T C A G T C C G T A A G T C A C G T A C G T | 1e-785 | -1.810e+03 | 13.10% | 3.74% | 749.6bp (741.1bp) | MOD(RRM)/Drosophila\_melanogaster-RNCMPT00140-PBM/HughesRNA(0.888) More Information \| Similar Motifs Found | motif file (matrix) |
| 18 | C A G T C T A G C A G T C T A G C G A T C T A G C G A T C T A G C G A T C T A G C G A T C T A G | 1e-578 | -1.332e+03 | 9.23% | 2.49% | 684.1bp (737.9bp) | cg/MA2107.1/Jaspar(0.914) More Information \| Similar Motifs Found | motif file (matrix) |
| 19 | A C G T C G T A A G T C C G T A A G T C A C G T C G T A A G T C C G T A G A T C G C A T G T C A | 1e-396 | -9.132e+02 | 4.59% | 0.87% | 694.5bp (682.5bp) | SFPQ(RRM)/Homo\_sapiens-RNCMPT00177-PBM/HughesRNA(0.756) More Information \| Similar Motifs Found | motif file (matrix) |
| 20 | C T G A G T C A A G T C A G T C A G T C A G C T C T G A C G T A A G T C A G T C G T A C G A C T | 1e-300 | -6.930e+02 | 2.01% | 0.14% | 708.1bp (756.3bp) | TBF1/MA0403.3/Jaspar(0.816) More Information \| Similar Motifs Found | motif file (matrix) |
| 21 | C G A T C T G A A G C T T C G A A C G T C T G A A C G T C G T A A C G T C T G A | 1e-171 | -3.949e+02 | 2.70% | 0.70% | 684.9bp (641.6bp) | SeqBias: TA-repeat(0.992) More Information \| Similar Motifs Found | motif file (matrix) |
| 22 | T A C G T C G A A G T C A C G T C T A G C G T A A T G C G C A T C T A G T C G A A G T C A G C T | 1e-147 | -3.403e+02 | 1.44% | 0.21% | 619.7bp (726.7bp) | ASH1/Literature(Harbison)/Yeast(0.687) More Information \| Similar Motifs Found | motif file (matrix) |
