## Supplemental dataset for "Hybrid CNN and Multi-Head Attention Model for Analyzing Epigenetic Mechanisms and Gene Expression Across Fungal Phylogenetic Distances": FgramModel_NcrassaTest_K4me2locs_knownResults.html

/projects/wg-feeds/SHAP/FgramModel\_NcrassaTest\_K4me2locs\_SHAP\_noDup\_HOMER/ - Homer Known Motif Enrichment Results


### Homer Known Motif Enrichment Results (/projects/wg-feeds/SHAP/FgramModel\_NcrassaTest\_K4me2locs\_SHAP\_noDup\_HOMER/)

Homer *de novo* Motif Results  
Gene Ontology Enrichment Results  
Known Motif Enrichment Results (txt file)  
Total Target Sequences = 32277, Total Background Sequences = 145780

|  |  |  |  |  |  |  |  |  |  |  |  |
| --- | --- | --- | --- | --- | --- | --- | --- | --- | --- | --- | --- |
| Rank | Motif | Name | P-value | log P-pvalue | q-value (Benjamini) | # Target Sequences with Motif | % of Targets Sequences with Motif | # Background Sequences with Motif | % of Background Sequences with Motif | Motif File | SVG |
| 1 | T A G C G T C A G A C T T A G C G T C A G A C T A G T C G C T A G A C T G A T C | ZML2(C2C2gata)/col-ZML2-DAP-Seq(GSE60143)/Homer | 1e-3557 | -8.192e+03 | 0.0000 | 12175.0 | 37.72% | 11458.3 | 7.86% | motif file (matrix) | svg |
| 2 | C T A G T C A G C T G A C T G A C T A G C T G A C A T G C A T G C T G A C T A G C T A G C G T A C T A G C G T A G T C A | TF3A(C2H2)/col-TF3A-DAP-Seq(GSE60143)/Homer | 1e-3089 | -7.113e+03 | 0.0000 | 20259.0 | 62.76% | 39597.8 | 27.16% | motif file (matrix) | svg |
| 3 | G C T A G C A T A C G T A C G T A G C T G A T C G A C T G A C T A G C T A C G T A C G T A G C T | RLR1?/SacCer-Promoters/Homer | 1e-2848 | -6.559e+03 | 0.0000 | 8028.0 | 24.87% | 5217.3 | 3.58% | motif file (matrix) | svg |
| 4 | A T G C C A G T A C G T A G T C C A T G A C G T A G T C A C G T A G C T G A T C | Unknown4/Arabidopsis-Promoters/Homer | 1e-2729 | -6.286e+03 | 0.0000 | 22731.0 | 70.42% | 52973.7 | 36.34% | motif file (matrix) | svg |
| 5 | A C G T A C G T A C G T A C G T A C G T A C G T A C G T A C G T A C G T A C G T | VRN1(ABI3VP1)/col-VRN1-DAP-Seq(GSE60143)/Homer | 1e-2580 | -5.941e+03 | 0.0000 | 4059.0 | 12.57% | 373.8 | 0.26% | motif file (matrix) | svg |
| 6 | G T C A T G C A T G C A G C T A C G T A G C T A G C T A G C T A | REM19(REM)/colamp-REM19-DAP-Seq(GSE60143)/Homer | 1e-2023 | -4.660e+03 | 0.0000 | 9019.0 | 27.94% | 10600.0 | 7.27% | motif file (matrix) | svg |
| 7 | A C G T A T G C A C G T A C G T A G C T A G T C A G C T A G C T A G C T A G C T A G C T | hTCT(CPE) | 1e-1583 | -3.647e+03 | 0.0000 | 23595.0 | 73.09% | 68997.0 | 47.33% | motif file (matrix) | svg |
| 8 | G C A T A G C T A C G T A C G T A C T G A C G T G A T C A C G T A C G T A G C T C G A T G C A T A G T C G A C T C A G T | IDD5(C2H2)/colamp-IDD5-DAP-Seq(GSE60143)/Homer | 1e-1508 | -3.474e+03 | 0.0000 | 9719.0 | 30.11% | 15596.6 | 10.70% | motif file (matrix) | svg |
| 9 | T G A C C T A G T C A G G T C A C G T A T C A G C G A T T C A G T C G A T G C A C T G A T A G C | PU.1-IRF(ETS:IRF)/Bcell-PU.1-ChIP-Seq(GSE21512)/Homer | 1e-1409 | -3.245e+03 | 0.0000 | 16331.0 | 50.59% | 39302.1 | 26.96% | motif file (matrix) | svg |
| 10 | T G A C C G T A C T G A A C T G A C T G G A C T G A T C T G C A G T A C T A C G | SF1(NR)/H295R-Nr5a1-ChIP-Seq(GSE44220)/Homer | 1e-1201 | -2.767e+03 | 0.0000 | 10257.0 | 31.77% | 19698.4 | 13.51% | motif file (matrix) | svg |
| 11 | C A T G G A C T C T A G C A T G C A G T C G A T C T A G C A T G C G A T C G T A C T A G C A G T C G A T C T A G C A T G | AT1G24250(Orphan)/col-AT1G24250-DAP-Seq(GSE60143)/Homer | 1e-1144 | -2.636e+03 | 0.0000 | 9380.0 | 29.06% | 17404.4 | 11.94% | motif file (matrix) | svg |
| 12 | C T A G C T A G T C G A C T A G C G T A A T C G T C G A A C T G C T G A T C G A C T G A T A C G | FRS9(ND)/col-FRS9-DAP-Seq(GSE60143)/Homer | 1e-1065 | -2.454e+03 | 0.0000 | 4279.0 | 13.26% | 4149.2 | 2.85% | motif file (matrix) | svg |
| 13 | A C G T A C T G C G T A A C G T A C T G A C T G C G T A C G T A | HAP3(CCAATHAP3)/col-HAP3-DAP-Seq(GSE60143)/Homer | 1e-1044 | -2.405e+03 | 0.0000 | 10217.0 | 31.65% | 21043.5 | 14.44% | motif file (matrix) | svg |
| 14 | G A T C G C A T A G T C A G C T G A T C A G C T G A T C G A C T G A T C G A C T A G T C A C G T G A T C A G C T G A T C | GAGA-repeat/SacCer-Promoters/Homer | 1e-1036 | -2.388e+03 | 0.0000 | 26740.0 | 82.84% | 92992.2 | 63.79% | motif file (matrix) | svg |
| 15 | G A C T A G C T A G C T C T A G A C G T G A T C A C G T A C G T G A C T C G A T G C A T A G T C | IDD4(C2H2)/col-IDD4-DAP-Seq(GSE60143)/Homer | 1e-1019 | -2.347e+03 | 0.0000 | 11992.0 | 37.15% | 27452.8 | 18.83% | motif file (matrix) | svg |
| 16 | C A T G G C T A C T A G T A C G C G T A T C A G C G T A A C T G C G T A C A T G C T G A C G T A | BPC1(BBRBPC)/colamp-BPC1-DAP-Seq(GSE60143)/Homer | 1e-1015 | -2.339e+03 | 0.0000 | 8541.0 | 26.46% | 15903.0 | 10.91% | motif file (matrix) | svg |
| 17 | A C G T G A C T A T G C G C T A C T G A C T A G A C T G G A C T A G T C C G T A | Nr5a2(NR)/mES-Nr5a2-ChIP-Seq(GSE19019)/Homer | 1e-1007 | -2.320e+03 | 0.0000 | 11644.0 | 36.07% | 26342.3 | 18.07% | motif file (matrix) | svg |
| 18 | C T G A G C A T A C T G C T A G A G T C A C T G A C T G A G T C A C T G T C A G | AT4G18450(AP2EREBP)/col-AT4G18450-DAP-Seq(GSE60143)/Homer | 1e-1003 | -2.311e+03 | 0.0000 | 14744.0 | 45.67% | 37947.6 | 26.03% | motif file (matrix) | svg |
| 19 | A C G T G A C T T A G C C G T A C T G A C A T G C T A G G A C T G A T C C G T A | Nr5a2(NR)/Pancreas-LRH1-ChIP-Seq(GSE34295)/Homer | 1e-983 | -2.264e+03 | 0.0000 | 13958.0 | 43.24% | 35173.8 | 24.13% | motif file (matrix) | svg |
| 20 | G A T C C A G T T A G C A G T C A C T G A G T C A G T C C T A G G A C T G T A C | LEP(AP2EREBP)/col-LEP-DAP-Seq(GSE60143)/Homer | 1e-973 | -2.242e+03 | 0.0000 | 12188.0 | 37.76% | 28687.6 | 19.68% | motif file (matrix) | svg |
| 21 | G A C T G A T C G A T C G C T A G T A C A G T C G C T A C T G A G T A C G A T C G C T A G A C T | MYB13(MYB)/col-MYB13-DAP-Seq(GSE60143)/Homer | 1e-954 | -2.199e+03 | 0.0000 | 15742.0 | 48.77% | 42567.7 | 29.20% | motif file (matrix) | svg |
| 22 | C A T G A G T C G T A C A C T G A T G C A G T C C A T G G A T C G A T C C T G A | ERF5(AP2EREBP)/colamp-ERF5-DAP-Seq(GSE60143)/Homer | 1e-910 | -2.097e+03 | 0.0000 | 14916.0 | 46.21% | 39873.4 | 27.35% | motif file (matrix) | svg |
| 23 | A T G C A G T C A C T G A T G C A G T C A C T G A G T C G T A C | SHN3(AP2EREBP)/col-SHN3-DAP-Seq(GSE60143)/Homer | 1e-882 | -2.031e+03 | 0.0000 | 10643.0 | 32.97% | 24234.9 | 16.62% | motif file (matrix) | svg |
| 24 | C T G A C G A T C A T G A T C G G C A T C A T G G C T A A G T C | ASHR1(ND)/col-ASHR1-DAP-Seq(GSE60143)/Homer | 1e-877 | -2.021e+03 | 0.0000 | 20494.0 | 63.49% | 64197.3 | 44.04% | motif file (matrix) | svg |
| 25 | G C A T G A T C C T A G C G T A G C A T C G T A G C A T A G T C C T A G C G T A G C A T C G A T | AT5G22990(C2H2)/col-AT5G22990-DAP-Seq(GSE60143)/Homer | 1e-874 | -2.013e+03 | 0.0000 | 16176.0 | 50.11% | 45491.2 | 31.21% | motif file (matrix) | svg |
| 26 | C G T A C T A G C A T G G A C T C T G A A C T G A C G T A C G T C T A G C T A G C A T G T C G A | MYB94(MYB)/col-MYB94-DAP-Seq(GSE60143)/Homer | 1e-831 | -1.915e+03 | 0.0000 | 10856.0 | 33.63% | 25586.3 | 17.55% | motif file (matrix) | svg |
| 27 | A T G C G T A C A G T C A G T C A C G T A C G T C G A T A C G T | AT5G02460(C2C2dof)/col-AT5G02460-DAP-Seq(GSE60143)/Homer | 1e-826 | -1.902e+03 | 0.0000 | 22686.0 | 70.28% | 75388.0 | 51.72% | motif file (matrix) | svg |
| 28 | C G T A T A G C T A G C T G C A A C T G C T A G C G T A C G T A T C A G G A C T | EHF(ETS)/LoVo-EHF-ChIP-Seq(GSE49402)/Homer | 1e-813 | -1.872e+03 | 0.0000 | 17637.0 | 54.64% | 52580.5 | 36.07% | motif file (matrix) | svg |
| 29 | G C T A C G T A C G T A G A C T C A T G C T A G G A T C A C T G T C A G G A T C A C T G T A C G | ERF9(AP2EREBP)/colamp-ERF9-DAP-Seq(GSE60143)/Homer | 1e-810 | -1.867e+03 | 0.0000 | 11845.0 | 36.69% | 29445.9 | 20.20% | motif file (matrix) | svg |
| 30 | C T G A C G A T C T A G T C A G G A T C C T G A T C A G G A T C C T G A A C T G A G T C G C T A A C G T A G T C G C A T | PRDM9(Zf)/Testis-DMC1-ChIP-Seq(GSE35498)/Homer | 1e-786 | -1.810e+03 | 0.0000 | 7903.0 | 24.48% | 16040.9 | 11.00% | motif file (matrix) | svg |
| 31 | G T A C A C T G A G T C A G T C C T A G G A T C G T A C C T G A | CRF4(AP2EREBP)/colamp-CRF4-DAP-Seq(GSE60143)/Homer | 1e-769 | -1.772e+03 | 0.0000 | 17896.0 | 55.44% | 54409.5 | 37.32% | motif file (matrix) | svg |
| 32 | A G T C G A C T G A T C C G T A G T A C A G T C G C T A C G T A G T A C A G T C G T A C G T A C | MYB63(MYB)/col-MYB63-DAP-Seq(GSE60143)/Homer | 1e-767 | -1.768e+03 | 0.0000 | 13228.0 | 40.98% | 35276.4 | 24.20% | motif file (matrix) | svg |
| 33 | C G T A C T A G C A T G A G C T C T G A C A T G C A G T C G A T C T A G C T A G | MYB30(MYB)/colamp-MYB30-DAP-Seq(GSE60143)/Homer | 1e-761 | -1.753e+03 | 0.0000 | 19810.0 | 61.37% | 63033.0 | 43.24% | motif file (matrix) | svg |
| 34 | A T G C T C G A T A C G A C G T A T G C A G T C A C G T A G T C A G T C G A T C | Znf263(Zf)/K562-Znf263-ChIP-Seq(GSE31477)/Homer | 1e-752 | -1.732e+03 | 0.0000 | 21066.0 | 65.26% | 68962.8 | 47.31% | motif file (matrix) | svg |
| 35 | C G A T A C G T A C G T A G C T A G C T G A T C G A T C G C T A A G C T A C G T A T C G T A C G | NFATC2(RHD)/Islets-NFATC2-ChIP-Seq(GSE158496)/Homer | 1e-696 | -1.603e+03 | 0.0000 | 20345.0 | 63.02% | 66628.2 | 45.71% | motif file (matrix) | svg |
| 36 | C G A T C T A G A C T G A G C T C T G A A C T G A C G T A C G T C T A G C T A G | MYB96(MYB)/colamp-MYB96-DAP-Seq(GSE60143)/Homer | 1e-686 | -1.581e+03 | 0.0000 | 18047.0 | 55.91% | 56498.8 | 38.76% | motif file (matrix) | svg |
| 37 | A C T G A C T G A G T C A C T G A C T G A G T C A C G T C T A G | ERF1(AP2EREBP)/colamp-ERF1-DAP-Seq(GSE60143)/Homer | 1e-684 | -1.577e+03 | 0.0000 | 16235.0 | 50.29% | 48749.6 | 33.44% | motif file (matrix) | svg |
| 38 | C T A G G C A T A C T G C T A G A G T C A C T G A C T G A G T C A C T G T C A G | ERF10(AP2EREBP)/col-ERF10-DAP-Seq(GSE60143)/Homer | 1e-659 | -1.518e+03 | 0.0000 | 19564.0 | 60.61% | 63748.7 | 43.73% | motif file (matrix) | svg |
| 39 | A C T G A C T G A G T C A C T G A C T G A G T C A C G T T C A G | ERF2(AP2EREBP)/colamp-ERF2-DAP-Seq(GSE60143)/Homer | 1e-647 | -1.492e+03 | 0.0000 | 17129.0 | 53.06% | 53206.4 | 36.50% | motif file (matrix) | svg |
| 40 | A C T G C T A G A G T C A C T G A C T G A T G C A C T G T A C G | ESE1(AP2EREBP)/col-ESE1-DAP-Seq(GSE60143)/Homer | 1e-647 | -1.490e+03 | 0.0000 | 18744.0 | 58.07% | 60291.6 | 41.36% | motif file (matrix) | svg |
| 41 | C A T G A G C T T A C G G T C A G T A C T A G C A G C T G A C T A T C G T C G A | Esrrb(NR)/mES-Esrrb-ChIP-Seq(GSE11431)/Homer | 1e-646 | -1.489e+03 | 0.0000 | 12949.0 | 40.11% | 36024.5 | 24.71% | motif file (matrix) | svg |
| 42 | G T A C C T G A T A G C C G T A G C T A T C G A T G C A T G A C C T A G G T C A A G T C C G T A C T G A C T G A C G T A | At1g14580(C2H2)/colamp-At1g14580-DAP-Seq(GSE60143)/Homer | 1e-642 | -1.479e+03 | 0.0000 | 3775.0 | 11.69% | 5195.2 | 3.56% | motif file (matrix) | svg |
| 43 | C T G A T C A G A G T C C G T A A T C G A T G C C G A T A C T G A G T C G A C T A T C G A G T C | MyoD(bHLH)/Myotube-MyoD-ChIP-Seq(GSE21614)/Homer | 1e-627 | -1.444e+03 | 0.0000 | 10263.0 | 31.79% | 26113.4 | 17.91% | motif file (matrix) | svg |
| 44 | G A T C G C T A G T A C A G T C G C T A T G C A G T A C G A T C C G T A G A C T | MYB83(MYB)/colamp-MYB83-DAP-Seq(GSE60143)/Homer | 1e-623 | -1.437e+03 | 0.0000 | 24288.0 | 75.24% | 87005.0 | 59.68% | motif file (matrix) | svg |
| 45 | G A T C G A T C A G T C G T A C C G A T G T A C G T A C A G T C A G T C A G T C G C T A G A T C | ZNF148(Zf)/MDAMB231-ZNF148-ChIP-Seq(GSE147020)/Homer | 1e-623 | -1.436e+03 | 0.0000 | 7303.0 | 22.62% | 15817.4 | 10.85% | motif file (matrix) | svg |
| 46 | C G T A G A C T C A T G C T A G A G T C A C T G A C T G G T A C C A T G T A C G | ERF3(AP2EREBP)/colamp-ERF3-DAP-Seq(GSE60143)/Homer | 1e-615 | -1.417e+03 | 0.0000 | 19925.0 | 61.72% | 66228.3 | 45.43% | motif file (matrix) | svg |
| 47 | T C G A T G C A C A G T T C G A G A T C A G T C C G T A C G T A A C T G A G T C C G T A C G T A T C A G C G A T A G T C | AT5G25475(ABI3VP1)/col-AT5G25475-DAP-Seq(GSE60143)/Homer | 1e-614 | -1.415e+03 | 0.0000 | 19506.0 | 60.43% | 64338.8 | 44.14% | motif file (matrix) | svg |
| 48 | C G T A T G A C T A G C T G C A A C T G A C T G C G T A C G T A T C A G G A C T | ELF3(ETS)/PDAC-ELF3-ChIP-Seq(GSE64557)/Homer | 1e-611 | -1.407e+03 | 0.0000 | 10258.0 | 31.78% | 26320.9 | 18.06% | motif file (matrix) | svg |
| 49 | A T G C G T A C A C T G A G T C A G T C A C T G A G T C G T A C | ERF73(AP2EREBP)/col-ERF73-DAP-Seq(GSE60143)/Homer | 1e-598 | -1.378e+03 | 0.0000 | 16926.0 | 52.43% | 53246.2 | 36.53% | motif file (matrix) | svg |
| 50 | A G C T G A T C G A T C C G T A G T A C A G T C C G A T C T G A G T A C G A T C C G T A G A C T | ATY19(MYB)/col-ATY19-DAP-Seq(GSE60143)/Homer | 1e-594 | -1.369e+03 | 0.0000 | 16203.0 | 50.19% | 50221.4 | 34.45% | motif file (matrix) | svg |
| 51 | G A C T G C T A T G C A A G T C A C G T A C G T A C G T C G A T A C G T T A C G | At3g45610(C2C2dof)/col-At3g45610-DAP-Seq(GSE60143)/Homer | 1e-586 | -1.350e+03 | 0.0000 | 18131.0 | 56.17% | 58735.4 | 40.29% | motif file (matrix) | svg |
| 52 | T A C G G A C T T G A C C G T A A C G T G A T C G T C A C G T A A C G T A T G C C G T A G A C T | HOXA2(Homeobox)/mES-Hoxa2-ChIP-Seq(Donaldson\_et\_al.)/Homer | 1e-581 | -1.338e+03 | 0.0000 | 3918.0 | 12.14% | 5981.9 | 4.10% | motif file (matrix) | svg |
| 53 | A G T C A G T C C G A T A C G T A C G T A C T G A C G T A G C T A G T C A G T C | Sox4(HMG)/proB-Sox4-ChIP-Seq(GSE50066)/Homer | 1e-575 | -1.326e+03 | 0.0000 | 13204.0 | 40.90% | 38205.8 | 26.21% | motif file (matrix) | svg |
| 54 | C T A G C A T G G A C T C G T A C T A G A C T G A C G T C T A G C T A G T C A G | MYB17(MYB)/colamp-MYB17-DAP-Seq(GSE60143)/Homer | 1e-559 | -1.289e+03 | 0.0000 | 14165.0 | 43.88% | 42372.7 | 29.07% | motif file (matrix) | svg |
| 55 | C G A T C T A G C G T A G A C T C A G T C T A G C G T A A G C T C A T G C T A G | HOXA1(Homeobox)/mES-Hoxa1-ChIP-Seq(SRP084292)/Homer | 1e-550 | -1.268e+03 | 0.0000 | 7248.0 | 22.45% | 16489.9 | 11.31% | motif file (matrix) | svg |
| 56 | T C A G T C A G G C T A C G T A T A C G G A C T T C A G T C G A C T G A C G T A T A C G G A C T | IRF8(IRF)/BMDM-IRF8-ChIP-Seq(GSE77884)/Homer | 1e-546 | -1.257e+03 | 0.0000 | 5489.0 | 17.00% | 10839.2 | 7.44% | motif file (matrix) | svg |
| 57 | C A G T A C T G T C A G T G C A G C T A A T G C T C G A A T C G G T C A T G C A | ZNF189(Zf)/HEK293-ZNF189.GFP-ChIP-Seq(GSE58341)/Homer | 1e-544 | -1.253e+03 | 0.0000 | 12811.0 | 39.69% | 37185.2 | 25.51% | motif file (matrix) | svg |
| 58 | C A T G G A C T T A C G G T C A G T A C G A T C G A C T A G C T A T C G T C G A T A C G T A G C | ERRg(NR)/Kidney-ESRRG-ChIP-Seq(GSE104905)/Homer | 1e-532 | -1.226e+03 | 0.0000 | 15007.0 | 46.49% | 46361.6 | 31.80% | motif file (matrix) | svg |
| 59 | G A C T G A C T G A T C C G T A G T A C A G T C G C A T C G T A G T A C G A T C G C A T G C T A | MYB74(MYB)/colamp-MYB74-DAP-Seq(GSE60143)/Homer | 1e-527 | -1.214e+03 | 0.0000 | 13865.0 | 42.95% | 41742.4 | 28.63% | motif file (matrix) | svg |
| 60 | G A C T G A T C A G T C C G T A T G A C A G T C G C A T C T G A G T A C G A T C G C A T G A C T | MYB10(MYB)/col-MYB10-DAP-Seq(GSE60143)/Homer | 1e-518 | -1.193e+03 | 0.0000 | 11168.0 | 34.60% | 31199.7 | 21.40% | motif file (matrix) | svg |
| 61 | C G T A C T A G C A G T A C G T C G T A A C T G C A T G G C A T T C A G C T G A | MYB49(MYB)/col-MYB49-DAP-Seq(GSE60143)/Homer | 1e-487 | -1.122e+03 | 0.0000 | 18999.0 | 58.86% | 64645.3 | 44.35% | motif file (matrix) | svg |
| 62 | G A C T A C G T A C G T A C T G A C G T A G T C G C A T A G C T G C A T G C A T G A C T A G C T | SGR5(C2H2)/colamp-SGR5-DAP-Seq(GSE60143)/Homer | 1e-485 | -1.117e+03 | 0.0000 | 9807.0 | 30.38% | 26574.3 | 18.23% | motif file (matrix) | svg |
| 63 | G A T C G A C T G A C T A C G T A G T C A C G T A G T C A C G T A G T C A C G T A G T C A C G T G T A C C G A T G T C A | BPC6(BBRBPC)/col-BPC6-DAP-Seq(GSE60143)/Homer | 1e-483 | -1.114e+03 | 0.0000 | 1014.0 | 3.14% | 290.4 | 0.20% | motif file (matrix) | svg |
| 64 | A T G C A G T C G C A T A G C T A C G T T C A G C G A T A G C T G A T C A T C G | Sox10(HMG)/SciaticNerve-Sox3-ChIP-Seq(GSE35132)/Homer | 1e-483 | -1.113e+03 | 0.0000 | 20382.0 | 63.14% | 71057.0 | 48.74% | motif file (matrix) | svg |
| 65 | C T A G C T A G A T G C G T A C T C A G A T G C A G T C G C A T G A T C G A T C | ZNF91(Zf)/HEK-ZNF91.HA-ChIP-Seq(GSE162571)/Homer | 1e-483 | -1.113e+03 | 0.0000 | 14297.0 | 44.29% | 44363.8 | 30.43% | motif file (matrix) | svg |
| 66 | A G C T A G C T A G C T A C T G A C G T A G T C A C T G A C G T G A C T C G A T G C A T A T C G | IDD7(C2H2)/col-IDD7-DAP-Seq(GSE60143)/Homer | 1e-479 | -1.103e+03 | 0.0000 | 7704.0 | 23.87% | 19008.9 | 13.04% | motif file (matrix) | svg |
| 67 | G T A C A C G T A C G T A T C G C A G T C G A T A T C G G C T A T G C A T A G C C G T A G T C A C A T G A G C T G C T A | ANAC013(NAC)/col-ANAC013-DAP-Seq(GSE60143)/Homer | 1e-477 | -1.100e+03 | 0.0000 | 10056.0 | 31.15% | 27632.9 | 18.96% | motif file (matrix) | svg |
| 68 | C G A T C T A G A C G T A C G T A C G T C G T A A G C T C G A T A G C T C G T A C T A G T A G C | FoxD3(forkhead)/ZebrafishEmbryo-Foxd3.biotin-ChIP-seq(GSE106676)/Homer | 1e-476 | -1.097e+03 | 0.0000 | 9743.0 | 30.18% | 26471.5 | 18.16% | motif file (matrix) | svg |
| 69 | A T G C G A T C C G A T A C G T A C G T A C T G C A G T A G C T | Sox3(HMG)/NPC-Sox3-ChIP-Seq(GSE33059)/Homer | 1e-468 | -1.078e+03 | 0.0000 | 21053.0 | 65.22% | 74525.8 | 51.12% | motif file (matrix) | svg |
| 70 | G A T C C T G A A G T C A G C T A C G T A C G T A C G T A C G T | At1g64620(C2C2dof)/colamp-At1g64620-DAP-Seq(GSE60143)/Homer | 1e-464 | -1.070e+03 | 0.0000 | 18284.0 | 56.64% | 61925.5 | 42.48% | motif file (matrix) | svg |
| 71 | G T A C G T A C G T C A G C T A C G T A C G T A C G T A C T A G C T A G C T A G | SEP3(MADS)/Arabidoposis-Flower-Sep3-ChIP-Seq/Homer | 1e-463 | -1.068e+03 | 0.0000 | 15748.0 | 48.78% | 50871.6 | 34.90% | motif file (matrix) | svg |
| 72 | C T A G C T G A C G T A C G T A C G T A C G T A A C T G A C G T C T A G G T C A | COG1(C2C2dof)/col-COG1-DAP-Seq(GSE60143)/Homer | 1e-460 | -1.061e+03 | 0.0000 | 17343.0 | 53.73% | 57842.1 | 39.68% | motif file (matrix) | svg |
| 73 | C T A G A C T G A G T C A C T G A C T G A G C T A C T G T C A G | AT3G57600(AP2EREBP)/col-AT3G57600-DAP-Seq(GSE60143)/Homer | 1e-460 | -1.061e+03 | 0.0000 | 17300.0 | 53.59% | 57655.2 | 39.55% | motif file (matrix) | svg |
| 74 | G A C T A C G T A G C T G A C T A C T G C A G T A G T C A T C G A C G T G C A T G C A T G C A T | MGP(C2H2)/colamp-MGP-DAP-Seq(GSE60143)/Homer | 1e-460 | -1.060e+03 | 0.0000 | 7237.0 | 22.42% | 17637.1 | 12.10% | motif file (matrix) | svg |
| 75 | A G T C G A T C G C T A C G A T C A G T T A C G C G A T A G C T A G T C A T C G | SOX1(HMG)/NPC-SOX1-ChIP-Seq(GSE138215)/Homer | 1e-459 | -1.059e+03 | 0.0000 | 22636.0 | 70.12% | 82264.5 | 56.43% | motif file (matrix) | svg |
| 76 | G C A T A C T G C T A G A G T C A C T G A C T G A G T C A C G T | ERF105(AP2EREBP)/colamp-ERF105-DAP-Seq(GSE60143)/Homer | 1e-450 | -1.038e+03 | 0.0000 | 23908.0 | 74.06% | 88711.2 | 60.85% | motif file (matrix) | svg |
| 77 | C G A T C G A T G C A T G A C T A C G T C G T A C G T A A C T G T A G C C G T A C G T A C G T A | AT5G60130(ABI3VP1)/col-AT5G60130-DAP-Seq(GSE60143)/Homer | 1e-448 | -1.032e+03 | 0.0000 | 15197.0 | 47.08% | 48860.7 | 33.52% | motif file (matrix) | svg |
| 78 | A G C T T G A C C G A T C G A T C T A G A C G T C A G T C A G T G C T A A G T C | FOXK1(Forkhead)/HEK293-FOXK1-ChIP-Seq(GSE51673)/Homer | 1e-443 | -1.020e+03 | 0.0000 | 15287.0 | 47.36% | 49347.6 | 33.85% | motif file (matrix) | svg |
| 79 | G T A C A C T G A T G C A G T C C T A G G A T C G T A C C T G A G A C T G C A T C G A T G A C T | RAP212(AP2EREBP)/col-RAP212-DAP-Seq(GSE60143)/Homer | 1e-443 | -1.020e+03 | 0.0000 | 20851.0 | 64.59% | 74150.2 | 50.87% | motif file (matrix) | svg |
| 80 | A T G C G C A T T A G C C G A T T A G C G C A T T A G C G C A T A T G C G A C T | GAGA-repeat/Arabidopsis-Promoters/Homer | 1e-441 | -1.016e+03 | 0.0000 | 12217.0 | 37.85% | 36714.7 | 25.19% | motif file (matrix) | svg |
| 81 | G A C T G C A T A C G T A C G T A C T G C G T A A G T C A G C T C G A T A T C G G C A T A C T G C G A T C T A G C G T A | WRKY50(WRKY)/col-WRKY50-DAP-Seq(GSE60143)/Homer | 1e-436 | -1.005e+03 | 0.0000 | 17273.0 | 53.51% | 58071.4 | 39.84% | motif file (matrix) | svg |
| 82 | C G A T A G C T T G C A A C T G A G T C T G A C C T A G G T A C A G T C C G T A G C A T G C A T | ERF13(AP2EREBP)/colamp-ERF13-DAP-Seq(GSE60143)/Homer | 1e-434 | -1.001e+03 | 0.0000 | 22440.0 | 69.51% | 81887.7 | 56.17% | motif file (matrix) | svg |
| 83 | A T G C A G T C G A T C C G T A A C G T A C G T A C T G A C G T A G C T G A T C | Sox2(HMG)/mES-Sox2-ChIP-Seq(GSE11431)/Homer | 1e-433 | -9.977e+02 | 0.0000 | 13839.0 | 42.87% | 43482.9 | 29.83% | motif file (matrix) | svg |
| 84 | C G T A A C G T A C G T A C G T A C G T A G T C A G T C C T G A A G C T A G C T | NFAT(RHD)/Jurkat-NFATC1-ChIP-Seq(Jolma\_et\_al.)/Homer | 1e-431 | -9.935e+02 | 0.0000 | 12444.0 | 38.55% | 37818.8 | 25.94% | motif file (matrix) | svg |
| 85 | C T A G A C T G A C G T C G T A A C T G A C T G A G C T C T A G T C A G C T A G | MYB93(MYB)/colamp-MYB93-DAP-Seq(GSE60143)/Homer | 1e-425 | -9.809e+02 | 0.0000 | 20064.0 | 62.15% | 70891.0 | 48.63% | motif file (matrix) | svg |
| 86 | A T G C G T A C C T G A A G T C A G T C A C T G G T C A A G T C G T C A G C A T G C A T G A C T | At5g65130(AP2EREBP)/colamp-At5g65130-DAP-Seq(GSE60143)/Homer | 1e-425 | -9.788e+02 | 0.0000 | 10261.0 | 31.79% | 29330.4 | 20.12% | motif file (matrix) | svg |
| 87 | G C T A C G T A C G A T G A C T G C A T T G C A A G T C A G C T A C G T A C G T C G A T G A C T | DAG2(C2C2dof)/col-DAG2-DAP-Seq(GSE60143)/Homer | 1e-424 | -9.770e+02 | 0.0000 | 16855.0 | 52.21% | 56516.7 | 38.77% | motif file (matrix) | svg |
| 88 | A C G T A G T C A G T C C G A T A C G T A C G T A C T G A C G T A T G C G A C T A C T G T A C G | Sox21(HMG)/ESC-SOX21-ChIP-Seq(GSE110505)/Homer | 1e-420 | -9.672e+02 | 0.0000 | 20578.0 | 63.75% | 73413.8 | 50.36% | motif file (matrix) | svg |
| 89 | G A T C G A T C G A T C C G T A G T A C A G T C G C A T C G T A G T A C G A T C | MYB58(MYB)/colamp-MYB58-DAP-Seq(GSE60143)/Homer | 1e-417 | -9.611e+02 | 0.0000 | 19041.0 | 58.99% | 66411.5 | 45.56% | motif file (matrix) | svg |
| 90 | C A G T C G T A C G T A G C A T G A C T G C A T G T A C A G C T A C T G G A C T A C G T C A T G | RAV1(RAV)/colamp-RAV1-DAP-Seq(GSE60143)/Homer | 1e-415 | -9.561e+02 | 0.0000 | 9434.0 | 29.22% | 26348.9 | 18.08% | motif file (matrix) | svg |
| 91 | A C T G C T A G A G T C A C T G A C T G A G T C A C T G T A C G | ERF104(AP2EREBP)/col-ERF104-DAP-Seq(GSE60143)/Homer | 1e-414 | -9.551e+02 | 0.0000 | 21870.0 | 67.75% | 79628.6 | 54.62% | motif file (matrix) | svg |
| 92 | G C A T A G C T A G C T A G C T A C T G A C G T A G T C A C T G A C G T G A C T C G A T G C A T | JKD(C2H2)/col-JKD-DAP-Seq(GSE60143)/Homer | 1e-406 | -9.355e+02 | 0.0000 | 4497.0 | 13.93% | 9214.6 | 6.32% | motif file (matrix) | svg |
| 93 | C T A G T C A G C T G A T C A G T G C A A C T G T C G A T C A G | Trl(Zf)/S2-GAGAfactor-ChIP-Seq(GSE40646)/Homer | 1e-403 | -9.290e+02 | 0.0000 | 24801.0 | 76.83% | 94290.3 | 64.68% | motif file (matrix) | svg |
| 94 | T C G A A C T G C A T G A G C T A G T C C G T A C T G A C T A G A C T G C G A T A T G C C T G A | RAR:RXR(NR),DR0/ES-RAR-ChIP-Seq(GSE56893)/Homer | 1e-400 | -9.224e+02 | 0.0000 | 3162.0 | 9.80% | 5309.4 | 3.64% | motif file (matrix) | svg |
| 95 | C G A T C G A T G C A T G C A T G T C A A G T C A G C T A C G T A C G T C G A T G A C T A C G T | OBP4(C2C2dof)/col-OBP4-DAP-Seq(GSE60143)/Homer | 1e-397 | -9.148e+02 | 0.0000 | 16422.0 | 50.87% | 55249.0 | 37.90% | motif file (matrix) | svg |
| 96 | C T G A T A C G G C A T A G C T A G C T A G T C T C G A A C T G C A G T A G C T A G C T G A T C | IRF3(IRF)/BMDM-Irf3-ChIP-Seq(GSE67343)/Homer | 1e-396 | -9.122e+02 | 0.0000 | 4085.0 | 12.65% | 8049.1 | 5.52% | motif file (matrix) | svg |
| 97 | A G T C G A T C C T G A A G T C A G T C C A T G G T C A G A T C C G T A G A T C | DREB26(AP2EREBP)/col-DREB26-DAP-Seq(GSE60143)/Homer | 1e-392 | -9.034e+02 | 0.0000 | 9319.0 | 28.87% | 26320.0 | 18.06% | motif file (matrix) | svg |
| 98 | C T A G G C T A A G T C A C T G A C G T G A C T G A C T A T G C T C G A C A G T G A T C C G A T G A C T G A T C G A T C | RKD2(RWPRK)/colamp-RKD2-DAP-Seq(GSE60143)/Homer | 1e-388 | -8.956e+02 | 0.0000 | 13616.0 | 42.18% | 43506.7 | 29.85% | motif file (matrix) | svg |
| 99 | T A C G T A G C G C T A C G A T C T A G A C G T C A G T C A G T G C T A A G T C G T C A G C A T | FOXK2(Forkhead)/U2OS-FOXK2-ChIP-Seq(E-MTAB-2204)/Homer | 1e-388 | -8.939e+02 | 0.0000 | 11000.0 | 34.08% | 32915.1 | 22.58% | motif file (matrix) | svg |
| 100 | C G T A C G T A C T A G A C G T A C G T C G T A A C T G A C T G A C G T C T G A T C G A T C G A | MYB4(MYB)/col200-MYB4-DAP-Seq(GSE60143)/Homer | 1e-387 | -8.919e+02 | 0.0000 | 13278.0 | 41.13% | 42142.2 | 28.91% | motif file (matrix) | svg |
| 101 | G T C A G C T A G C T A T C G A A T C G A C G T A G T C T C G A T C G A T G A C | WRKY40(WRKY)/colamp-WRKY40-DAP-Seq(GSE60143)/Homer | 1e-385 | -8.887e+02 | 0.0000 | 12239.0 | 37.91% | 37922.0 | 26.01% | motif file (matrix) | svg |
| 102 | T C A G A C T G A C G T C G T A A C T G A C T G A C G T C T A G | MYB51(MYB)/col-MYB51-DAP-Seq(GSE60143)/Homer | 1e-385 | -8.878e+02 | 0.0000 | 18140.0 | 56.19% | 63111.3 | 43.29% | motif file (matrix) | svg |
| 103 | A T G C G T A C C G T A A G C T G C A T T A C G A G C T A G C T A G T C A G C T | Sox6(HMG)/Myotubes-Sox6-ChIP-Seq(GSE32627)/Homer | 1e-381 | -8.783e+02 | 0.0000 | 20567.0 | 63.71% | 74299.7 | 50.97% | motif file (matrix) | svg |
| 104 | A G C T G A C T A C G T A C T G A C G T A G T C A C T G A C G T G C A T C G A T | AtIDD11(C2H2)/colamp-AtIDD11-DAP-Seq(GSE60143)/Homer | 1e-381 | -8.774e+02 | 0.0000 | 7713.0 | 23.89% | 20533.9 | 14.09% | motif file (matrix) | svg |
| 105 | A C T G C T A G A G T C A C T G A C T G A G T C A C G T C T A G | AT5G23930(mTERF)/col-AT5G23930-DAP-Seq(GSE60143)/Homer | 1e-378 | -8.712e+02 | 0.0000 | 22988.0 | 71.21% | 85902.0 | 58.93% | motif file (matrix) | svg |
| 106 | C G A T C G T A G T A C A C G T A C G T T C A G G C A T C A G T T A C G G T C A C G T A A G T C C G T A T G C A C A T G | NAC2(NAC)/colamp-NAC2-DAP-Seq(GSE60143)/Homer | 1e-376 | -8.673e+02 | 0.0000 | 12898.0 | 39.96% | 40810.9 | 28.00% | motif file (matrix) | svg |
| 107 | C T A G C T A G C G T A C G T A T A C G C G A T C T A G C T G A C T G A C G T A T A C G G A C T | PU.1:IRF8(ETS:IRF)/pDC-Irf8-ChIP-Seq(GSE66899)/Homer | 1e-374 | -8.627e+02 | 0.0000 | 2984.0 | 9.24% | 5028.4 | 3.45% | motif file (matrix) | svg |
| 108 | C T G A T C G A C G T A A T G C C G T A C G T A C G A T C T A G T C A G G A T C | Sox15(HMG)/CPA-Sox15-ChIP-Seq(GSE62909)/Homer | 1e-374 | -8.623e+02 | 0.0000 | 14243.0 | 44.12% | 46452.1 | 31.87% | motif file (matrix) | svg |
| 109 | T A C G C G T A G A C T T C A G A G C T A G T C A C T G T C A G A G T C C T G A | DDF2(AP2EREBP)/col-DDF2-DAP-Seq(GSE60143)/Homer | 1e-373 | -8.612e+02 | 0.0000 | 4712.0 | 14.60% | 10264.3 | 7.04% | motif file (matrix) | svg |
| 110 | C T A G A G T C A G T C A C T G C G T A A G T C C T G A G A C T | DDF1(AP2EREBP)/col-DDF1-DAP-Seq(GSE60143)/Homer | 1e-373 | -8.607e+02 | 0.0000 | 19198.0 | 59.47% | 68180.7 | 46.77% | motif file (matrix) | svg |
| 111 | A G T C C G T A T G A C A T G C G C A T C T G A G T A C G A T C | MYB55(MYB)/colamp-MYB55-DAP-Seq(GSE60143)/Homer | 1e-373 | -8.592e+02 | 0.0000 | 20341.0 | 63.01% | 73455.5 | 50.39% | motif file (matrix) | svg |
| 112 | T A C G T C G A C G T A C G T A C G T A C T G A A C T G A C G T C G T A T C G A | AT2G28810(C2C2dof)/colamp-AT2G28810-DAP-Seq(GSE60143)/Homer | 1e-369 | -8.504e+02 | 0.0000 | 22765.0 | 70.52% | 85044.0 | 58.34% | motif file (matrix) | svg |
| 113 | A C T G A C G T C G A T C A G T C A T G C A T G C A G T G C A T C A G T C A T G | HuR(?)/HEK293-HuR-CLIP-Seq(GSE87887)/Homer | 1e-365 | -8.424e+02 | 0.0000 | 25621.0 | 79.37% | 99374.3 | 68.17% | motif file (matrix) | svg |
| 114 | A G T C G A T C A G C T C G T A G T A C A G T C G C A T C T G A G T A C G A T C | AT4G26030(C2H2)/col-AT4G26030-DAP-Seq(GSE60143)/Homer | 1e-365 | -8.412e+02 | 0.0000 | 18822.0 | 58.31% | 66684.6 | 45.74% | motif file (matrix) | svg |
| 115 | C T G A T C G A G T A C A C G T A C G T A T C G A C G T C G A T A T C G G C T A G T A C A T G C C G T A T G C A C A T G | ANAC103(NAC)/col-ANAC103-DAP-Seq(GSE60143)/Homer | 1e-364 | -8.400e+02 | 0.0000 | 8639.0 | 26.76% | 24247.6 | 16.63% | motif file (matrix) | svg |
| 116 | C G A T C T G A A G T C A C G T A C G T A T C G G A C T C T A G C G A T G A C T C G T A A T G C C G T A G T C A A C T G | ANAC011(NAC)/col-ANAC011-DAP-Seq(GSE60143)/Homer | 1e-362 | -8.349e+02 | 0.0000 | 6596.0 | 20.43% | 16812.6 | 11.53% | motif file (matrix) | svg |
| 117 | A G C T A G C T A G C T A C T G A C G T A G T C A C T G A C G T G C A T G C A T G C A T A C G T | At5g66730(C2H2)/colamp-At5g66730-DAP-Seq(GSE60143)/Homer | 1e-354 | -8.152e+02 | 0.0000 | 5725.0 | 17.73% | 13902.5 | 9.54% | motif file (matrix) | svg |
| 118 | G C A T C T A G A C T G A C G T C G T A A C T G A C T G C G A T C T A G T C G A T C G A G C T A | MYB40(MYB)/col-MYB40-DAP-Seq(GSE60143)/Homer | 1e-351 | -8.103e+02 | 0.0000 | 8776.0 | 27.19% | 25002.7 | 17.15% | motif file (matrix) | svg |
| 119 | A G T C A C G T A C T G A G C T A C G T A C G T G T C A A G T C | Foxo1(Forkhead)/RAW-Foxo1-ChIP-Seq(Fan\_et\_al.)/Homer | 1e-351 | -8.091e+02 | 0.0000 | 19873.0 | 61.56% | 71844.4 | 49.28% | motif file (matrix) | svg |
| 120 | C G T A G A T C C T A G A C G T G T A C C T G A A G C T G A T C G C T A G A C T | TGA2(bZIP)/colamp-TGA2-DAP-Seq(GSE60143)/Homer | 1e-347 | -7.997e+02 | 0.0000 | 19496.0 | 60.39% | 70216.1 | 48.17% | motif file (matrix) | svg |
| 121 | C G T A G A C T C A T G C T A G A G T C A C T G C T A G A G T C C A T G C T A G | ERF7(AP2EREBP)/col-ERF7-DAP-Seq(GSE60143)/Homer | 1e-346 | -7.987e+02 | 0.0000 | 25128.0 | 77.84% | 97322.0 | 66.76% | motif file (matrix) | svg |
| 122 | T A G C C G T A C T G A T A C G C G T A A C G T A C T G A C T G A G T C T A C G C T A G G T A C | YY1(Zf)/Promoter/Homer | 1e-345 | -7.966e+02 | 0.0000 | 2145.0 | 6.64% | 3013.4 | 2.07% | motif file (matrix) | svg |
| 123 | C T G A A T G C G C T A G C A T A T G C C G T A T C G A C T G A C T A G T C A G T A C G G T C A | Tcf4(HMG)/Hct116-Tcf4-ChIP-Seq(SRA012054)/Homer | 1e-345 | -7.959e+02 | 0.0000 | 7591.0 | 23.52% | 20684.7 | 14.19% | motif file (matrix) | svg |
| 124 | C G A T C G T A G T A C A C G T A C G T T C A G G C A T C A G T T A C G G T C A C G T A A G T C C G T A T G C A C A T G | ANAC053(NAC)/colamp-ANAC053-DAP-Seq(GSE60143)/Homer | 1e-344 | -7.934e+02 | 0.0000 | 11588.0 | 35.90% | 36177.0 | 24.82% | motif file (matrix) | svg |
| 125 | G A T C A T G C A G T C C G T A A G T C A G T C A C T G G C T A A G T C C G T A | AT1G44830(AP2EREBP)/col-AT1G44830-DAP-Seq(GSE60143)/Homer | 1e-344 | -7.926e+02 | 0.0000 | 11587.0 | 35.89% | 36180.8 | 24.82% | motif file (matrix) | svg |
| 126 | C T G A A G T C C G A T A G C T A T G C G T A C A C G T A T C G C A G T G C A T | Elf4(ETS)/BMDM-Elf4-ChIP-Seq(GSE88699)/Homer | 1e-336 | -7.745e+02 | 0.0000 | 15032.0 | 46.57% | 50711.6 | 34.79% | motif file (matrix) | svg |
| 127 | G T C A T C G A T C G A C G T A G C T A C G T A T C G A T G A C A C T G C G T A A G T C C G T A C G T A T C G A G C T A | IDD2(C2H2)/colamp-IDD2-DAP-Seq(GSE60143)/Homer | 1e-335 | -7.716e+02 | 0.0000 | 2806.0 | 8.69% | 4865.8 | 3.34% | motif file (matrix) | svg |
| 128 | G C T A A G T C T A C G T G C A A T C G T C A G G C T A T C G A T C A G A G C T | ELF5(ETS)/T47D-ELF5-ChIP-Seq(GSE30407)/Homer | 1e-331 | -7.643e+02 | 0.0000 | 11804.0 | 36.57% | 37334.1 | 25.61% | motif file (matrix) | svg |
| 129 | C T G A T C A G G T A C G C T A A C T G T G A C G C A T C A T G | SCL(bHLH)/HPC7-Scl-ChIP-Seq(GSE13511)/Homer | 1e-325 | -7.494e+02 | 0.0000 | 27221.0 | 84.33% | 108745.3 | 74.60% | motif file (matrix) | svg |
| 130 | A G T C G A C T A G C T C G A T A T C G G C T A C G A T A T C G C G A T A C T G T A C G A C G T | Tcf7(HMG)/GM12878-TCF7-ChIP-Seq(Encode)/Homer | 1e-325 | -7.494e+02 | 0.0000 | 5786.0 | 17.92% | 14534.8 | 9.97% | motif file (matrix) | svg |
| 131 | C T G A G A C T C A T G C T A G A G T C A C T G A C T G A G T C A C T G T C A G | ERF11(AP2EREBP)/col-ERF11-DAP-Seq(GSE60143)/Homer | 1e-324 | -7.475e+02 | 0.0000 | 22157.0 | 68.64% | 83276.1 | 57.13% | motif file (matrix) | svg |
| 132 | G C A T C G T A C T A G A G T C G T C A C G T A A T G C A C G T A C G T A C T G G A T C G C A T C G T A G C T A G C T A | bHLH122(bHLH)/col100-bHLH122-DAP-Seq(GSE60143)/Homer | 1e-322 | -7.423e+02 | 0.0000 | 15119.0 | 46.84% | 51436.5 | 35.28% | motif file (matrix) | svg |
| 133 | C G T A C G A T C T A G C G T A A G C T C A G T T A C G C G T A A C G T C A T G C T A G A T G C | HOXA3(Homeobox)/mEmbryo-Hoxa3-ChIP-Seq(E-MTAB-8607)/Homer | 1e-321 | -7.398e+02 | 0.0000 | 4278.0 | 13.25% | 9503.5 | 6.52% | motif file (matrix) | svg |
| 134 | C A T G A T G C T A G C C T G A A G T C A G T C A C T G G C T A A G T C G T A C G C T A G C A T | At4g28140(AP2EREBP)/colamp-At4g28140-DAP-Seq(GSE60143)/Homer | 1e-321 | -7.396e+02 | 0.0000 | 13539.0 | 41.94% | 44754.5 | 30.70% | motif file (matrix) | svg |
| 135 | A G T C C A T G A C G T A C G T A C T G C G T A A G T C G A C T G C A T G C T A | WRKY28(WRKY)/col-WRKY28-DAP-Seq(GSE60143)/Homer | 1e-318 | -7.334e+02 | 0.0000 | 19243.0 | 59.61% | 69821.2 | 47.90% | motif file (matrix) | svg |
| 136 | C G A T C T G A A G T C A C G T A C G T T A C G G C T A C A T G C T A G G C A T C G A T A G T C C G T A G T C A A C T G | ANAC096(NAC)/colamp-ANAC096-DAP-Seq(GSE60143)/Homer | 1e-314 | -7.251e+02 | 0.0000 | 12096.0 | 37.47% | 38919.4 | 26.70% | motif file (matrix) | svg |
| 137 | T C G A T G A C G C A T A G C T C A G T G A T C G C T A G A T C G A C T A C G T G C A T A G T C | PRDM1(Zf)/Hela-PRDM1-ChIP-Seq(GSE31477)/Homer | 1e-314 | -7.243e+02 | 0.0000 | 6518.0 | 20.19% | 17306.4 | 11.87% | motif file (matrix) | svg |
| 138 | A G T C G T A C C T G A A G T C G T A C C T A G G C T A T G A C T G C A G C T A C G T A C G T A | At1g22810(AP2EREBP)/colamp-At1g22810-DAP-Seq(GSE60143)/Homer | 1e-313 | -7.228e+02 | 0.0000 | 17087.0 | 52.93% | 60246.8 | 41.33% | motif file (matrix) | svg |
| 139 | G A C T A C T G C A G T A G T C A C T G A C T G A G C T A C T G C T A G G T C A | At1g77640(AP2EREBP)/col-At1g77640-DAP-Seq(GSE60143)/Homer | 1e-312 | -7.199e+02 | 0.0000 | 7990.0 | 24.75% | 22772.6 | 15.62% | motif file (matrix) | svg |
| 140 | C T A G A C T G A C G T C G T A A C T G C A T G G C A T T C A G | MYB92(MYB)/colamp-MYB92-DAP-Seq(GSE60143)/Homer | 1e-312 | -7.194e+02 | 0.0000 | 18313.0 | 56.73% | 65765.5 | 45.11% | motif file (matrix) | svg |
| 141 | G T A C T C G A T A G C C G T A C G T A C T G A T G C A T G A C A C T G C G T A A G T C C T G A C T G A T C G A C G T A | NUC(C2H2)/col-NUC-DAP-Seq(GSE60143)/Homer | 1e-310 | -7.145e+02 | 0.0000 | 2443.0 | 7.57% | 4063.7 | 2.79% | motif file (matrix) | svg |
| 142 | C T A G C T G A C T A G C T G A C T A G C T G A C T A G C T G A C T A G C T G A | SeqBias: GA-repeat | 1e-306 | -7.066e+02 | 0.0000 | 29774.0 | 92.23% | 123501.3 | 84.72% | motif file (matrix) | svg |
| 143 | A G C T G C T A T G C A A G T C A C G T A C G T A C G T C G A T A G C T T C A G | dof24(C2C2dof)/col-dof24-DAP-Seq(GSE60143)/Homer | 1e-304 | -7.006e+02 | 0.0000 | 21599.0 | 66.91% | 81175.9 | 55.69% | motif file (matrix) | svg |
| 144 | A G T C T G C A T C G A C T G A A C T G C A T G A C G T A T G C G T C A T A C G | Erra(NR)/HepG2-Erra-ChIP-Seq(GSE31477)/Homer | 1e-302 | -6.966e+02 | 0.0000 | 21014.0 | 65.10% | 78465.1 | 53.83% | motif file (matrix) | svg |
| 145 | G C A T C G A T G A C T T G C A A C T G A G T C T G A C A C T G G A T C A G T C C G T A G A C T | ERF15(AP2EREBP)/colamp-ERF15-DAP-Seq(GSE60143)/Homer | 1e-298 | -6.878e+02 | 0.0000 | 24682.0 | 76.46% | 96324.6 | 66.08% | motif file (matrix) | svg |
| 146 | G C T A C T G A T C G A A G T C A G T C C T G A A G T C G T C A C T G A T G C A | RUNX1(Runt)/Jurkat-RUNX1-ChIP-Seq(GSE29180)/Homer | 1e-296 | -6.835e+02 | 0.0000 | 15179.0 | 47.02% | 52358.4 | 35.92% | motif file (matrix) | svg |
| 147 | G C T A C G T A C G T A G C A T C A T G C T A G G A T C A C T G T C A G G A T C A C T G T C A G | ERF4(AP2EREBP)/colamp-ERF4-DAP-Seq(GSE60143)/Homer | 1e-295 | -6.811e+02 | 0.0000 | 23990.0 | 74.32% | 92966.4 | 63.77% | motif file (matrix) | svg |
| 148 | G A T C A G T C G A C T G C T A G T A C A G T C G C A T G C T A G T A C G A T C | MYB61(MYB)/colamp-MYB61-DAP-Seq(GSE60143)/Homer | 1e-291 | -6.714e+02 | 0.0000 | 22241.0 | 68.90% | 84587.2 | 58.03% | motif file (matrix) | svg |
| 149 | T C G A T A G C G T C A A C T G A C T G C G T A C G T A C T A G A G C T T C A G | ERG(ETS)/VCaP-ERG-ChIP-Seq(GSE14097)/Homer | 1e-291 | -6.704e+02 | 0.0000 | 16538.0 | 51.23% | 58439.8 | 40.09% | motif file (matrix) | svg |
| 150 | A G C T G T C A T G C A A G T C A C G T A C G T A C G T C G A T G A C T T A C G | AT3G12130(C3H)/colamp-AT3G12130-DAP-Seq(GSE60143)/Homer | 1e-289 | -6.670e+02 | 0.0000 | 23306.0 | 72.20% | 89783.1 | 61.59% | motif file (matrix) | svg |
| 151 | A T G C A G T C A G C T A G C T A C G T A T C G C G T A C G A T T A G C G A C T | LEF1(HMG)/H1-LEF1-ChIP-Seq(GSE64758)/Homer | 1e-287 | -6.609e+02 | 0.0000 | 9709.0 | 30.08% | 29946.3 | 20.54% | motif file (matrix) | svg |
| 152 | C A T G A G T C G T C A C G T A A T G C A C G T A C G T A C T G | bHLH130(bHLH)/col-bHLH130-DAP-Seq(GSE60143)/Homer | 1e-286 | -6.607e+02 | 0.0000 | 13573.0 | 42.05% | 45762.6 | 31.39% | motif file (matrix) | svg |
| 153 | G C T A C G T A C G A T C A G T A C T G C G A T G T A C A C T G A T C G G A C T C A T G C T A G G C A T C A G T C A T G | DEAR5(AP2EREBP)/col-DEAR5-DAP-Seq(GSE60143)/Homer | 1e-285 | -6.582e+02 | 0.0000 | 9071.0 | 28.10% | 27466.8 | 18.84% | motif file (matrix) | svg |
| 154 | G A C T A G T C C T G A A G T C A G T C A C T G C T G A A G T C G C T A G C A T G T A C C G A T G C A T G A C T C G A T | CBF2(AP2EREBP)/colamp-CBF2-DAP-Seq(GSE60143)/Homer | 1e-285 | -6.571e+02 | 0.0000 | 19494.0 | 60.39% | 71895.0 | 49.32% | motif file (matrix) | svg |
| 155 | T C G A C T G A C G T A C G T A C G T A C T G A A C T G A C G T C T G A C T G A | AT5G63260(C3H)/col-AT5G63260-DAP-Seq(GSE60143)/Homer | 1e-285 | -6.568e+02 | 0.0000 | 22050.0 | 68.31% | 83855.9 | 57.52% | motif file (matrix) | svg |
| 156 | C G A T T C G A G A T C C G A T G C A T T C A G G A C T C G A T G C A T G C T A C T G A A G T C C G T A G T C A C T A G | ANAC005(NAC)/col-ANAC005-DAP-Seq(GSE60143)/Homer | 1e-284 | -6.540e+02 | 0.0000 | 7405.0 | 22.94% | 21123.5 | 14.49% | motif file (matrix) | svg |
| 157 | C G A T T G C A A G T C A C G T A C G T T A C G C G A T C G A T T A C G G C T A G C T A A T G C C G T A G T C A C A T G | ANAC016(NAC)/col-ANAC016-DAP-Seq(GSE60143)/Homer | 1e-283 | -6.530e+02 | 0.0000 | 16505.0 | 51.13% | 58503.3 | 40.13% | motif file (matrix) | svg |
| 158 | G C A T G C T A C G T A A G C T G C T A T G C A A G T C A C G T A C G T A C G T G C A T G C A T | At4g38000(C2C2dof)/col-At4g38000-DAP-Seq(GSE60143)/Homer | 1e-282 | -6.494e+02 | 0.0000 | 12114.0 | 37.53% | 39793.3 | 27.30% | motif file (matrix) | svg |
| 159 | G A C T A C T G C G A T A G T C A C T G C T A G A G T C C T G A | AT1G12630(AP2EREBP)/colamp-AT1G12630-DAP-Seq(GSE60143)/Homer | 1e-279 | -6.442e+02 | 0.0000 | 18826.0 | 58.32% | 69002.5 | 47.33% | motif file (matrix) | svg |
| 160 | G A T C A G C T C T A G G A T C T G A C C T A G C G T A G T A C C G T A G C A T G T C A C T G A | CBF3(AP2EREBP)/colamp-CBF3-DAP-Seq(GSE60143)/Homer | 1e-279 | -6.429e+02 | 0.0000 | 19291.0 | 59.76% | 71141.5 | 48.80% | motif file (matrix) | svg |
| 161 | C A G T T C A G T C G A A G T C C G T A A C T G T G A C C G A T A C T G A C T G A C G T A T C G | Atoh7(bHLH)/Retina-Atoh7-CutnRun(GSE156756)/Homer | 1e-279 | -6.429e+02 | 0.0000 | 9765.0 | 30.25% | 30345.2 | 20.82% | motif file (matrix) | svg |
| 162 | C G A T T C G A G A T C A C G T A C G T T C A G G C A T C T G A C G T A G C T A C G T A A G T C C G T A T G C A C A T G | ANAC050(NAC)/colamp-ANAC050-DAP-Seq(GSE60143)/Homer | 1e-279 | -6.426e+02 | 0.0000 | 12068.0 | 37.38% | 39677.7 | 27.22% | motif file (matrix) | svg |
| 163 | G A C T A G T C G A T C C G T A G T A C A G T C G C A T C G T A G T C A G A T C | MYB67(MYB)/col-MYB67-DAP-Seq(GSE60143)/Homer | 1e-278 | -6.424e+02 | 0.0000 | 17977.0 | 55.69% | 65185.7 | 44.72% | motif file (matrix) | svg |
| 164 | A T G C C A G T A G C T A G C T T C A G G T C A T A G C G A C T C G T A C G A T | WRKY20(WRKY)/col-WRKY20-DAP-Seq(GSE60143)/Homer | 1e-278 | -6.413e+02 | 0.0000 | 14004.0 | 43.38% | 47812.4 | 32.80% | motif file (matrix) | svg |
| 165 | C G A T T C G A G A T C A C G T A C G T T C A G G C T A C G A T C G T A C G T A C G T A A T G C C G T A T G C A C T A G | ANAC028(NAC)/col-ANAC028-DAP-Seq(GSE60143)/Homer | 1e-278 | -6.403e+02 | 0.0000 | 13190.0 | 40.86% | 44380.1 | 30.44% | motif file (matrix) | svg |
| 166 | C G A T C G T A A G T C A C G T A C G T T C G A T G C A G C A T G C T A C G T A A C G T A G C T C G T A C G T A A C T G | ANAC062(NAC)/colamp-ANAC062-DAP-Seq(GSE60143)/Homer | 1e-277 | -6.384e+02 | 0.0000 | 6458.0 | 20.01% | 17740.1 | 12.17% | motif file (matrix) | svg |
| 167 | C G T A G A T C A G C T A C G T A C G T A C T G C G T A G T A C A G C T G C T A C G A T C G A T C G A T G C A T G C T A | WRKY18(WRKY)/col-WRKY18-DAP-Seq(GSE60143)/Homer | 1e-273 | -6.293e+02 | 0.0000 | 22551.0 | 69.86% | 86598.0 | 59.41% | motif file (matrix) | svg |
| 168 | C G T A C G T A C G T A C T G A C T A G A C G T C T A G G T C A | CDF3(C2C2dof)/colamp-CDF3-DAP-Seq(GSE60143)/Homer | 1e-273 | -6.292e+02 | 0.0000 | 19329.0 | 59.88% | 71488.9 | 49.04% | motif file (matrix) | svg |
| 169 | G A C T G T A C C T G A G A T C A G T C C T A G G C T A G T A C C T G A G C T A G C A T C G A T G C A T G A C T C G T A | AT3G16280(AP2EREBP)/colamp-AT3G16280-DAP-Seq(GSE60143)/Homer | 1e-272 | -6.285e+02 | 0.0000 | 17574.0 | 54.44% | 63551.6 | 43.60% | motif file (matrix) | svg |
| 170 | A G T C T G A C C T G A A G T C A G T C A C T G C G T A A G T C G T C A G C T A G C A T C G T A G C A T G C T A C G T A | DEAR3(AP2EREBP)/colamp-DEAR3-DAP-Seq(GSE60143)/Homer | 1e-272 | -6.281e+02 | 0.0000 | 15263.0 | 47.28% | 53377.6 | 36.62% | motif file (matrix) | svg |
| 171 | T C A G A G C T G T C A C G T A A C G T A T G C C G T A A C G T A C G T C T G A | PHV(HB)/col-PHV-DAP-Seq(GSE60143)/Homer | 1e-272 | -6.269e+02 | 0.0000 | 7487.0 | 23.19% | 21664.6 | 14.86% | motif file (matrix) | svg |
| 172 | A T C G A G T C A C T G A G T C A G T C A C T G G A C T G A C T | PUCHI(AP2EREBP)/colamp-PUCHI-DAP-Seq(GSE60143)/Homer | 1e-271 | -6.258e+02 | 0.0000 | 20388.0 | 63.16% | 76420.2 | 52.42% | motif file (matrix) | svg |
| 173 | C G T A C G A T C G T A C G A T C A T G A C T G C G A T A G T C A T C G T C A G G A C T A C T G | At1g36060(AP2EREBP)/colamp-At1g36060-DAP-Seq(GSE60143)/Homer | 1e-271 | -6.248e+02 | 0.0000 | 20511.0 | 63.54% | 77005.4 | 52.82% | motif file (matrix) | svg |
| 174 | T C G A T A G C T G C A A C T G A C T G C G T A C G T A C T A G G A C T T A C G | ETS1(ETS)/Jurkat-ETS1-ChIP-Seq(GSE17954)/Homer | 1e-270 | -6.230e+02 | 0.0000 | 14876.0 | 46.08% | 51765.1 | 35.51% | motif file (matrix) | svg |
| 175 | C A T G G T A C A C T G G T C A A G C T T A C G T G C A A T C G T G A C C A G T | TOD6?/SacCer-Promoters/Homer | 1e-267 | -6.162e+02 | 0.0000 | 5957.0 | 18.45% | 16090.8 | 11.04% | motif file (matrix) | svg |
| 176 | A G C T C T A G A G T C A G T C A C T G C G T A A G T C C T G A G C A T G C T A C T G A G C A T G C A T C G A T G C A T | CBF4(AP2EREBP)/colamp-CBF4-DAP-Seq(GSE60143)/Homer | 1e-265 | -6.124e+02 | 0.0000 | 23095.0 | 71.54% | 89436.1 | 61.35% | motif file (matrix) | svg |
| 177 | G C A T C G A T G C A T C G T A C T A G A G T C G T C A C G T A A T C G A C G T A C G T A C T G G T A C G C A T C G A T | bHLH80(bHLH)/col-bHLH80-DAP-Seq(GSE60143)/Homer | 1e-264 | -6.081e+02 | 0.0000 | 15557.0 | 48.19% | 54900.1 | 37.66% | motif file (matrix) | svg |
| 178 | G T A C A C T G A T G C T G A C C T A G G A C T G T A C C G T A G C A T G C A T | ERF8(AP2EREBP)/colamp-ERF8-DAP-Seq(GSE60143)/Homer | 1e-263 | -6.063e+02 | 0.0000 | 24607.0 | 76.23% | 96935.5 | 66.50% | motif file (matrix) | svg |
| 179 | G A T C C T G A A G T C A G T C A C T G C G T A A G T C C T G A | ERF38(AP2EREBP)/col-ERF38-DAP-Seq(GSE60143)/Homer | 1e-262 | -6.039e+02 | 0.0000 | 18461.0 | 57.19% | 67858.2 | 46.55% | motif file (matrix) | svg |
| 180 | G A C T C T A G C T A G C T A G A C T G T C G A C T G A C T A G C T A G C T A G G T A C G T C A | ZNF467(Zf)/HEK293-ZNF467.GFP-ChIP-Seq(GSE58341)/Homer | 1e-260 | -6.004e+02 | 0.0000 | 10377.0 | 32.15% | 33230.3 | 22.80% | motif file (matrix) | svg |
| 181 | A G T C G A T C A G T C C G T A A T C G C A G T A G T C G T A C C T G A A C T G T C A G A G C T A G C T A G C T A G C T | PRDM15(Zf)/ESC-Prdm15-ChIP-Seq(GSE73694)/Homer | 1e-260 | -6.001e+02 | 0.0000 | 14656.0 | 45.40% | 51096.3 | 35.05% | motif file (matrix) | svg |
| 182 | G A T C C T G A A G T C G T A C A C T G G C T A G A T C C T G A G C T A G C T A | At4g31060(AP2EREBP)/colamp-At4g31060-DAP-Seq(GSE60143)/Homer | 1e-259 | -5.964e+02 | 0.0000 | 17064.0 | 52.86% | 61683.3 | 42.31% | motif file (matrix) | svg |
| 183 | G A C T A C T G C G A T A G T C A C T G C T A G A G T C C G T A | Rap210(AP2EREBP)/col-Rap210-DAP-Seq(GSE60143)/Homer | 1e-258 | -5.949e+02 | 0.0000 | 21487.0 | 66.56% | 81971.4 | 56.23% | motif file (matrix) | svg |
| 184 | A G C T A G C T C A T G C T G A G T A C A G T C A G C T A G C T C A G T C T A G | RARa(NR)/K562-RARa-ChIP-Seq(Encode)/Homer | 1e-256 | -5.916e+02 | 0.0000 | 24836.0 | 76.94% | 98260.5 | 67.41% | motif file (matrix) | svg |
| 185 | T A G C G T A C A G T C G T A C C G A T A G T C A G T C A G T C A G T C A G T C C G T A G A T C | Zfp281(Zf)/ES-Zfp281-ChIP-Seq(GSE81042)/Homer | 1e-255 | -5.884e+02 | 0.0000 | 1775.0 | 5.50% | 2698.9 | 1.85% | motif file (matrix) | svg |
| 186 | A G C T G A T C C T G A A G T C A G T C A C T G C G T A A G T C C T G A G T C A G C A T C G A T G C T A G C A T C G T A | At2g44940(AP2EREBP)/colamp-At2g44940-DAP-Seq(GSE60143)/Homer | 1e-255 | -5.873e+02 | 0.0000 | 12137.0 | 37.60% | 40586.6 | 27.84% | motif file (matrix) | svg |
| 187 | A T G C C A T G A G C T C A G T C A T G T C G A A G T C G A C T C G A T C G A T C A G T C A G T | WRKY26(WRKY)/colamp-WRKY26-DAP-Seq(GSE60143)/Homer | 1e-253 | -5.843e+02 | 0.0000 | 11001.0 | 34.08% | 35938.3 | 24.65% | motif file (matrix) | svg |
| 188 | C T G A A T C G G T A C C T G A A G T C A G T C A C T G C G T A A G T C C T G A | TINY(AP2EREBP)/col-TINY-DAP-Seq(GSE60143)/Homer | 1e-250 | -5.777e+02 | 0.0000 | 12999.0 | 40.27% | 44308.0 | 30.39% | motif file (matrix) | svg |
| 189 | T C G A A G T C C G T A A T C G A T G C C G A T A C T G A G T C A G C T A C T G | Tcf12(bHLH)/GM12878-Tcf12-ChIP-Seq(GSE32465)/Homer | 1e-250 | -5.768e+02 | 0.0000 | 10393.0 | 32.20% | 33548.0 | 23.01% | motif file (matrix) | svg |
| 190 | G C T A C G T A C G T A G C A T C A T G C T A G A G T C A C T G A C T G A G T C A C T G T C A G | ABR1(AP2EREBP)/colamp-ABR1-DAP-Seq(GSE60143)/Homer | 1e-249 | -5.754e+02 | 0.0000 | 23248.0 | 72.02% | 90654.1 | 62.19% | motif file (matrix) | svg |
| 191 | C G T A T A C G T C G A A C T G A C T G C G T A C G T A T A C G A G C T T A C G | PU.1(ETS)/ThioMac-PU.1-ChIP-Seq(GSE21512)/Homer | 1e-246 | -5.682e+02 | 0.0000 | 6530.0 | 20.23% | 18582.0 | 12.75% | motif file (matrix) | svg |
| 192 | T G A C G C T A T C G A T G C A A G T C A G T C C G T A A G T C C G T A C T G A G C T A G T A C | RUNX2(Runt)/PCa-RUNX2-ChIP-Seq(GSE33889)/Homer | 1e-241 | -5.570e+02 | 0.0000 | 12527.0 | 38.81% | 42572.8 | 29.20% | motif file (matrix) | svg |
| 193 | C T A G G T A C C A T G G A C T C G A T C A T G G T C A G T A C G A C T C G A T C G A T C G A T | WRKY27(WRKY)/colamp-WRKY27-DAP-Seq(GSE60143)/Homer | 1e-241 | -5.553e+02 | 0.0000 | 14586.0 | 45.18% | 51355.1 | 35.23% | motif file (matrix) | svg |
| 194 | A G T C A G T C C T G A A G T C A G T C A C T G C G T A A G T C C T G A T C G A G C A T G A T C C G A T C G A T A C T G | AT3G60490(AP2EREBP)/colamp-AT3G60490-DAP-Seq(GSE60143)/Homer | 1e-240 | -5.545e+02 | 0.0000 | 14465.0 | 44.81% | 50843.6 | 34.88% | motif file (matrix) | svg |
| 195 | T G A C C G A T C T G A C T A G C T A G A C G T A T G C T G C A T C G A C T G A C T A G C A T G A C G T A G T C C G T A | PPARa(NR),DR1/Liver-Ppara-ChIP-Seq(GSE47954)/Homer | 1e-239 | -5.515e+02 | 0.0000 | 11884.0 | 36.81% | 39954.5 | 27.41% | motif file (matrix) | svg |
| 196 | T C A G C T G A C G T A C G T A T A C G G C A T C T A G C T G A C G T A C G T A T A C G G A C T | IRF1(IRF)/PBMC-IRF1-ChIP-Seq(GSE43036)/Homer | 1e-239 | -5.514e+02 | 0.0000 | 1902.0 | 5.89% | 3163.6 | 2.17% | motif file (matrix) | svg |
| 197 | T C A G A C G T T C G A T A G C A G T C C G T A A C T G G T A C A C G T A C T G A T C G A G T C | Atoh1(bHLH)/Cerebellum-Atoh1-ChIP-Seq(GSE22111)/Homer | 1e-238 | -5.497e+02 | 0.0000 | 13441.0 | 41.64% | 46522.9 | 31.91% | motif file (matrix) | svg |
| 198 | G C A T T G A C C A T G G A C T C A G T C A T G T C G A G T A C G A C T G C T A C G A T C G A T | WRKY6(WRKY)/colamp-WRKY6-DAP-Seq(GSE60143)/Homer | 1e-238 | -5.490e+02 | 0.0000 | 15158.0 | 46.96% | 53914.5 | 36.98% | motif file (matrix) | svg |
| 199 | C G T A C T G A C G T A C T A G T C G A C T A G A C T G C G T A C G T A T A C G A G C T A T C G | SpiB(ETS)/OCILY3-SPIB-ChIP-Seq(GSE56857)/Homer | 1e-237 | -5.461e+02 | 0.0000 | 3768.0 | 11.67% | 8940.3 | 6.13% | motif file (matrix) | svg |
| 200 | A T G C A G T C C T G A A G T C C G A T A C G T A G T C A G T C A C G T A T C G G A C T A C G T | Etv2(ETS)/ES-ER71-ChIP-Seq(GSE59402)/Homer | 1e-234 | -5.409e+02 | 0.0000 | 11277.0 | 34.93% | 37566.4 | 25.77% | motif file (matrix) | svg |
| 201 | C G T A G C A T C G T A C G T A G C A T A C T G C G A T A G T C A C T G A C T G G A C T C T A G | AT1G71450(AP2EREBP)/col-AT1G71450-DAP-Seq(GSE60143)/Homer | 1e-234 | -5.404e+02 | 0.0000 | 27337.0 | 84.68% | 111696.1 | 76.62% | motif file (matrix) | svg |
| 202 | C G T A G C T A C G A T C T A G A C G T G T C A C G T A C G T A A G T C C G T A T G C A T A C G | FoxL2(Forkhead)/Ovary-FoxL2-ChIP-Seq(GSE60858)/Homer | 1e-233 | -5.374e+02 | 0.0000 | 10349.0 | 32.06% | 33808.8 | 23.19% | motif file (matrix) | svg |
| 203 | G C A T A C G T A C G T A T C G C G T A C G T A C G T A C G T A | At2g41835(C2H2)/col-At2g41835-DAP-Seq(GSE60143)/Homer | 1e-233 | -5.371e+02 | 0.0000 | 6185.0 | 19.16% | 17567.8 | 12.05% | motif file (matrix) | svg |
| 204 | T C A G A T C G G A C T A C T G G A C T C A G T C T A G C G T A G T A C C G T A C T A G A T C G | Tbx20(T-box)/Heart-Tbx20-ChIP-Seq(GSE29636)/Homer | 1e-230 | -5.299e+02 | 0.0000 | 4838.0 | 14.99% | 12736.8 | 8.74% | motif file (matrix) | svg |
| 205 | C T A G A C T G A C G T C G T A A C T G A C T G A C G T T C A G C T G A T C G A | MYB107(MYB)/col-MYB107-DAP-Seq(GSE60143)/Homer | 1e-229 | -5.292e+02 | 0.0000 | 22775.0 | 70.55% | 88987.4 | 61.04% | motif file (matrix) | svg |
| 206 | A T G C C A T G G C A T G A C T C T A G T C G A G T A C A G C T C G T A G C T A | WRKY75(WRKY)/col-WRKY75-DAP-Seq(GSE60143)/Homer | 1e-229 | -5.289e+02 | 0.0000 | 16045.0 | 49.70% | 58062.8 | 39.83% | motif file (matrix) | svg |
| 207 | A T G C C G T A C G T A C G T A C G T A C G T A A C T G A C G T C G A T C T G A | dof43(C2C2dof)/colamp-dof43-DAP-Seq(GSE60143)/Homer | 1e-229 | -5.285e+02 | 0.0000 | 17032.0 | 52.76% | 62440.1 | 42.83% | motif file (matrix) | svg |
| 208 | C G T A G A T C C A T G G C A T G A C T C T A G T C G A T A G C A G C T G C A T | WRKY55(WRKY)/col-WRKY55-DAP-Seq(GSE60143)/Homer | 1e-228 | -5.255e+02 | 0.0000 | 17850.0 | 55.30% | 66145.8 | 45.38% | motif file (matrix) | svg |
| 209 | G C T A T C G A C G T A C T A G A G C T G T C A G T C A C G T A A G T C C G T A | FOXA1(Forkhead)/LNCAP-FOXA1-ChIP-Seq(GSE27824)/Homer | 1e-228 | -5.252e+02 | 0.0000 | 11670.0 | 36.15% | 39380.2 | 27.01% | motif file (matrix) | svg |
| 210 | G A T C C A T G A C G T A C G T A C T G C G T A A G T C A G C T C G A T G A C T | WRKY8(WRKY)/colamp-WRKY8-DAP-Seq(GSE60143)/Homer | 1e-226 | -5.211e+02 | 0.0000 | 2684.0 | 8.31% | 5584.4 | 3.83% | motif file (matrix) | svg |
| 211 | C G T A C G T A C G T A C G T A C T G A C A G T A C G T C G T A A C T G A C T G A C G T C T A G C T G A T C G A C T G A | MYB39(MYB)/col-MYB39-DAP-Seq(GSE60143)/Homer | 1e-224 | -5.165e+02 | 0.0000 | 3921.0 | 12.15% | 9644.9 | 6.62% | motif file (matrix) | svg |
| 212 | G C T A C G T A C G T A G C A T C A T G C T A G A G T C A C T G T A C G A G T C C A T G T A C G | RAP26(AP2EREBP)/colamp-RAP26-DAP-Seq(GSE60143)/Homer | 1e-224 | -5.163e+02 | 0.0000 | 24932.0 | 77.23% | 99710.8 | 68.40% | motif file (matrix) | svg |
| 213 | C T A G A C T G A G C T C G T A A C T G A C T G A C G T C T A G | MYB99(MYB)/colamp-MYB99-DAP-Seq(GSE60143)/Homer | 1e-223 | -5.144e+02 | 0.0000 | 18265.0 | 56.58% | 68173.1 | 46.77% | motif file (matrix) | svg |
| 214 | G A C T C T G A A G T C A G T C A C T G C G T A A G T C C T G A | bHLH10(bHLH)/colamp-bHLH10-DAP-Seq(GSE60143)/Homer | 1e-223 | -5.143e+02 | 0.0000 | 13925.0 | 43.14% | 49042.8 | 33.64% | motif file (matrix) | svg |
| 215 | T C G A C T G A C G T A C G T A C G T A C G T A A C T G A C G T C G A T C T G A | BBX31(Orphan)/col-BBX31-DAP-Seq(GSE60143)/Homer | 1e-222 | -5.120e+02 | 0.0000 | 17749.0 | 54.98% | 65878.8 | 45.19% | motif file (matrix) | svg |
| 216 | G A C T T C G A C G T A C G T A C G T A C G T A C G T A C T A G A G C T C G T A | dof45(C2C2dof)/col-dof45-DAP-Seq(GSE60143)/Homer | 1e-221 | -5.102e+02 | 0.0000 | 22488.0 | 69.66% | 87874.9 | 60.28% | motif file (matrix) | svg |
| 217 | G T A C C A T G A G C T A G C T T C A G T G C A T G A C A G C T C G T A C G T A | WRKY33(WRKY)/col-WRKY33-DAP-Seq(GSE60143)/Homer | 1e-221 | -5.099e+02 | 0.0000 | 15567.0 | 48.22% | 56220.7 | 38.57% | motif file (matrix) | svg |
| 218 | C A T G A C T G C T A G T C G A T C G A T C G A T C G A T C A G T C A G T C A G T G A C T G A C C G T A A C T G T G C A C G A T A C T G | RBPJ:Ebox(?,bHLH)/Panc1-Rbpj1-ChIP-Seq(GSE47459)/Homer | 1e-220 | -5.072e+02 | 0.0000 | 3648.0 | 11.30% | 8781.4 | 6.02% | motif file (matrix) | svg |
| 219 | A T G C G A C T A C G T C T A G A C G T A C G T A C G T C T G A G A T C G C T A A G C T C G T A | Foxa2(Forkhead)/Liver-Foxa2-ChIP-Seq(GSE25694)/Homer | 1e-219 | -5.053e+02 | 0.0000 | 10849.0 | 33.61% | 36226.8 | 24.85% | motif file (matrix) | svg |
| 220 | A T G C G A T C C G T A A G C T C A G T A T C G G C A T A G C T G A C T A C T G | Sox17(HMG)/Endoderm-Sox17-ChIP-Seq(GSE61475)/Homer | 1e-218 | -5.040e+02 | 0.0000 | 12083.0 | 37.43% | 41364.7 | 28.38% | motif file (matrix) | svg |
| 221 | T C G A G T A C C A T G A G C T A C G T C A T G G T C A G T A C A G C T G C T A C G A T C A G T | WRKY31(WRKY)/colamp-WRKY31-DAP-Seq(GSE60143)/Homer | 1e-218 | -5.026e+02 | 0.0000 | 13035.0 | 40.38% | 45397.3 | 31.14% | motif file (matrix) | svg |
| 222 | G A C T T C A G G C A T A G T C G C T A G A T C C T G A A C G T A G T C G T C A | Replumless(BLH)/Arabidopsis-RPL.GFP-ChIP-Seq(GSE78727)/Homer | 1e-218 | -5.022e+02 | 0.0000 | 19732.0 | 61.13% | 75048.9 | 51.48% | motif file (matrix) | svg |
| 223 | G T A C A C G T A C G T T C A G G A C T G C A T T C A G C G T A C T G A A G T C C G T A G T C A A C T G A C G T G C T A | NTM2(NAC)/col-NTM2-DAP-Seq(GSE60143)/Homer | 1e-216 | -4.991e+02 | 0.0000 | 10391.0 | 32.19% | 34423.6 | 23.61% | motif file (matrix) | svg |
| 224 | C T G A T G A C T G A C C G T A A C G T T G A C A G C T C T A G A C G T G A C T | Olig2(bHLH)/Neuron-Olig2-ChIP-Seq(GSE30882)/Homer | 1e-216 | -4.981e+02 | 0.0000 | 20382.0 | 63.14% | 78116.0 | 53.59% | motif file (matrix) | svg |
| 225 | A T C G T G C A G A T C C T A G A C G T A T C G C G T A A G T C T C A G A C T G T C A G G C T A | Knotted(Homeobox)/Corn-KN1-ChIP-Seq(GSE39161)/Homer | 1e-215 | -4.951e+02 | 0.0000 | 24055.0 | 74.52% | 95667.7 | 65.63% | motif file (matrix) | svg |
| 226 | C G T A C G T A C T A G C A G T G A C T C G T A C A T G C A T G C G A T C T G A C T G A C T G A | MS188(MYB)/colamp-MS188-DAP-Seq(GSE60143)/Homer | 1e-214 | -4.949e+02 | 0.0000 | 13456.0 | 41.68% | 47290.5 | 32.44% | motif file (matrix) | svg |
| 227 | C G A T T C G A A T G C C G A T G C A T T C G A A G C T G C A T G C A T C G A T T C G A A G C T T C G A G T C A C T A G | ANAC004(NAC)/colamp-ANAC004-DAP-Seq(GSE60143)/Homer | 1e-214 | -4.947e+02 | 0.0000 | 5815.0 | 18.01% | 16563.1 | 11.36% | motif file (matrix) | svg |
| 228 | A G T C G T A C C T G A A G T C A G T C C A T G G C T A A G T C T G C A G C T A G C A T G C A T | RAP21(AP2EREBP)/colamp-RAP21-DAP-Seq(GSE60143)/Homer | 1e-214 | -4.945e+02 | 0.0000 | 10096.0 | 31.28% | 33274.7 | 22.83% | motif file (matrix) | svg |
| 229 | G T A C C A T G A G C T A C G T A C T G C G T A A G T C G A C T G C A T C G A T | WRKY29(WRKY)/colamp-WRKY29-DAP-Seq(GSE60143)/Homer | 1e-214 | -4.939e+02 | 0.0000 | 16749.0 | 51.89% | 61660.7 | 42.30% | motif file (matrix) | svg |
| 230 | G A T C G T A C C T G A A G T C A G T C A C T G G C T A G T A C G T C A G C A T G C A T C G A T | DEAR2(AP2EREBP)/colamp-DEAR2-DAP-Seq(GSE60143)/Homer | 1e-212 | -4.889e+02 | 0.0000 | 24825.0 | 76.90% | 99547.8 | 68.29% | motif file (matrix) | svg |
| 231 | C G T A C G T A C G A T A C T G C A G T A G T C A C T G A C T G A G C T A C T G | DREB19(AP2EREBP)/colamp-DREB19-DAP-Seq(GSE60143)/Homer | 1e-211 | -4.876e+02 | 0.0000 | 18309.0 | 56.72% | 68753.4 | 47.16% | motif file (matrix) | svg |
| 232 | A G T C A C G T A C G T T A C G G C A T G C A T A T G C G C T A C G T A A T G C C G T A G T C A A C T G G A T C G C A T | ANAC075(NAC)/col-ANAC075-DAP-Seq(GSE60143)/Homer | 1e-210 | -4.850e+02 | 0.0000 | 8491.0 | 26.30% | 26936.6 | 18.48% | motif file (matrix) | svg |
| 233 | T C A G T G A C G T A C C G T A A C G T T G A C A C G T T C A G A G C T G A C T | NeuroD1(bHLH)/Islet-NeuroD1-ChIP-Seq(GSE30298)/Homer | 1e-208 | -4.805e+02 | 0.0000 | 10432.0 | 32.32% | 34814.3 | 23.88% | motif file (matrix) | svg |
| 234 | C G A T T C G A G T A C A C G T A C G T T C A G G C A T G C A T G C T A C G T A C G T A A G T C C G T A T G C A C A T G | ANAC020(NAC)/col-ANAC020-DAP-Seq(GSE60143)/Homer | 1e-207 | -4.776e+02 | 0.0000 | 12977.0 | 40.20% | 45479.7 | 31.20% | motif file (matrix) | svg |
| 235 | A G C T C T A G G A T C A G T C C T A G C T G A A G T C G C T A G C A T T G C A | CBF1(AP2EREBP)/colamp-CBF1-DAP-Seq(GSE60143)/Homer | 1e-206 | -4.750e+02 | 0.0000 | 21941.0 | 67.97% | 85770.2 | 58.84% | motif file (matrix) | svg |
| 236 | G A C T C A G T G C A T C G A T T G A C A C G T A T G C G T A C C T G A A C T G A C T G A G C T | WIP5(C2H2)/colamp-WIP5-DAP-Seq(GSE60143)/Homer | 1e-205 | -4.741e+02 | 0.0000 | 16391.0 | 50.78% | 60351.2 | 41.40% | motif file (matrix) | svg |
| 237 | G C A T T G A C C T G A A G T C A G T C A C T G G T C A A G T C G C T A G A C T G C T A C T G A | DREB2(AP2EREBP)/col-DREB2-DAP-Seq(GSE60143)/Homer | 1e-205 | -4.736e+02 | 0.0000 | 16432.0 | 50.90% | 60540.8 | 41.53% | motif file (matrix) | svg |
| 238 | A T G C C A T G A C G T A C G T A C T G C G T A A G T C G A C T G C A T C G A T | WRKY71(WRKY)/col-WRKY71-DAP-Seq(GSE60143)/Homer | 1e-204 | -4.717e+02 | 0.0000 | 13328.0 | 41.29% | 47054.6 | 32.28% | motif file (matrix) | svg |
| 239 | C G A T T G A C C A T G G A C T A C G T C A T G C G T A G A T C G A C T G C A T G C A T C G A T | WRKY14(WRKY)/colamp-WRKY14-DAP-Seq(GSE60143)/Homer | 1e-203 | -4.683e+02 | 0.0000 | 9630.0 | 29.83% | 31691.8 | 21.74% | motif file (matrix) | svg |
| 240 | C G A T G C T A G C T A G C A T G C T A C G T A A G T C A C G T A C G T A C G T C G A T A G C T | At5g62940(C2C2dof)/col-At5g62940-DAP-Seq(GSE60143)/Homer | 1e-201 | -4.647e+02 | 0.0000 | 25790.0 | 79.89% | 104702.5 | 71.82% | motif file (matrix) | svg |
| 241 | T C G A A G T C A C G T A C G T T C A G C A G T C T G A C T A G T C G A C G T A A T C G C G T A C G T A A C T G A G C T | NTM1(NAC)/col-NTM1-DAP-Seq(GSE60143)/Homer | 1e-201 | -4.637e+02 | 0.0000 | 7542.0 | 23.36% | 23435.8 | 16.08% | motif file (matrix) | svg |
| 242 | T G A C T A G C T C A G T C G A T C G A C G T A A G T C C G T A C G T A C G A T C T A G T A C G | Sox7(HMG)/ESC-Sox7-ChIP-Seq(GSE133899)/Homer | 1e-199 | -4.602e+02 | 0.0000 | 6552.0 | 20.30% | 19652.8 | 13.48% | motif file (matrix) | svg |
| 243 | C A G T A G C T G A C T T G C A A G T C A G C T A C G T A C G T C G A T G A C T | AT3G52440(C2C2dof)/colamp-AT3G52440-DAP-Seq(GSE60143)/Homer | 1e-198 | -4.573e+02 | 0.0000 | 21710.0 | 67.25% | 84936.4 | 58.27% | motif file (matrix) | svg |
| 244 | A G T C C T A G A C G T A C G T A C T G C G T A A G T C A G C T G C T A G C A T | WRKY24(WRKY)/colamp-WRKY24-DAP-Seq(GSE60143)/Homer | 1e-198 | -4.571e+02 | 0.0000 | 15250.0 | 47.24% | 55570.2 | 38.12% | motif file (matrix) | svg |
| 245 | A G C T C T G A C T A G C T A G A C T G T A G C T G C A T C G A C T G A C T A G C A T G A C G T A T G C T C G A | RXR(NR),DR1/3T3L1-RXR-ChIP-Seq(GSE13511)/Homer | 1e-197 | -4.543e+02 | 0.0000 | 11810.0 | 36.58% | 40861.7 | 28.03% | motif file (matrix) | svg |
| 246 | C G A T G A T C G A T C C T G A G A T C G A T C C A T G T G C A G T A C T C G A G T C A G C A T C G A T C G A T G C A T | At4g32800(AP2EREBP)/colamp-At4g32800-DAP-Seq(GSE60143)/Homer | 1e-197 | -4.537e+02 | 0.0000 | 7657.0 | 23.72% | 23991.8 | 16.46% | motif file (matrix) | svg |
| 247 | A G T C A C G T A C G T T C A G G C T A G C T A A T G C C G T A C G A T A G T C C G T A G T C A A C T G G A T C G C A T | SND3(NAC)/col-SND3-DAP-Seq(GSE60143)/Homer | 1e-196 | -4.531e+02 | 0.0000 | 13508.0 | 41.85% | 48079.4 | 32.98% | motif file (matrix) | svg |
| 248 | G A C T C G A T T C A G G A T C G A C T A G C T A G C T A G T C G A T C C G T A C T A G C T A G T C G A T C G A C T G A | Bcl6(Zf)/Liver-Bcl6-ChIP-Seq(GSE31578)/Homer | 1e-195 | -4.512e+02 | 0.0000 | 11883.0 | 36.81% | 41208.5 | 28.27% | motif file (matrix) | svg |
| 249 | G A C T G C A T G C A T A G T C A G C T T C G A T A C G G C T A C G T A A C T G G T A C G C A T C G A T A G T C G A C T | HSF3(HSF)/colamp-HSF3-DAP-Seq(GSE60143)/Homer | 1e-195 | -4.492e+02 | 0.0000 | 11289.0 | 34.97% | 38749.9 | 26.58% | motif file (matrix) | svg |
| 250 | C G T A C G A T C A G T C A T G C G A T G T A C C A T G A C T G G A C T C A T G | CEJ1(AP2EREBP)/col-CEJ1-DAP-Seq(GSE60143)/Homer | 1e-193 | -4.450e+02 | 0.0000 | 24371.0 | 75.50% | 97924.2 | 67.17% | motif file (matrix) | svg |
| 251 | C G A T T G C A T G C A G A T C C G T A A C T G T G A C G A C T C A T G A C T G | Tcf21(bHLH)/ArterySmoothMuscle-Tcf21-ChIP-Seq(GSE61369)/Homer | 1e-190 | -4.386e+02 | 0.0000 | 10415.0 | 32.26% | 35266.0 | 24.19% | motif file (matrix) | svg |
| 252 | C A T G C T A G A G T C A C T G A C T G G T A C C A T G T A C G | AT1G28160(AP2EREBP)/colamp-AT1G28160-DAP-Seq(GSE60143)/Homer | 1e-190 | -4.384e+02 | 0.0000 | 26046.0 | 80.69% | 106354.4 | 72.96% | motif file (matrix) | svg |
| 253 | G T C A T G C A G C T A A G T C C G T A A C T G T G A C G C A T T C A G C A G T | Ap4(bHLH)/AML-Tfap4-ChIP-Seq(GSE45738)/Homer | 1e-188 | -4.338e+02 | 0.0000 | 12427.0 | 38.50% | 43739.0 | 30.00% | motif file (matrix) | svg |
| 254 | C T A G T C G A T G A C A G T C C G T A A C T G G T A C A C G T A C T G A C T G | BHLHA15(bHLH)/NIH3T3-BHLHB8.HA-ChIP-Seq(GSE119782)/Homer | 1e-187 | -4.320e+02 | 0.0000 | 15255.0 | 47.26% | 55958.1 | 38.39% | motif file (matrix) | svg |
| 255 | G A C T G C A T C T A G C G A T G A T C T C G A C A T G G A T C | Tgif1(Homeobox)/mES-Tgif1-ChIP-Seq(GSE55404)/Homer | 1e-187 | -4.319e+02 | 0.0000 | 25607.0 | 79.33% | 104237.6 | 71.51% | motif file (matrix) | svg |
| 256 | G C A T C G A T C G T A G A T C C A T G A C G T A C G T A C T G C G T A A G T C A G C T G C A T G C A T C G T A G C T A | WRKY45(WRKY)/col-WRKY45-DAP-Seq(GSE60143)/Homer | 1e-186 | -4.292e+02 | 0.0000 | 6899.0 | 21.37% | 21300.7 | 14.61% | motif file (matrix) | svg |
| 257 | G C T A T C G A C G T A C T A G A G C T G T C A G T C A C G T A A G T C C G T A | FOXA1(Forkhead)/MCF7-FOXA1-ChIP-Seq(GSE26831)/Homer | 1e-184 | -4.257e+02 | 0.0000 | 9106.0 | 28.21% | 30082.2 | 20.64% | motif file (matrix) | svg |
| 258 | C T A G T A G C A T G C C T A G A G T C A G T C C T A G G A C T G A C T G C T A | CRF10(AP2EREBP)/col100-CRF10-DAP-Seq(GSE60143)/Homer | 1e-183 | -4.236e+02 | 0.0000 | 25345.0 | 78.51% | 103048.0 | 70.69% | motif file (matrix) | svg |
| 259 | A G C T A C G T A C T G A T G C A G T C C G T A C T G A T A C G | NF1-halfsite(CTF)/LNCaP-NF1-ChIP-Seq(Unpublished)/Homer | 1e-183 | -4.229e+02 | 0.0000 | 19809.0 | 61.36% | 76583.1 | 52.54% | motif file (matrix) | svg |
| 260 | G C T A G C T A C G T A C G T A C T G A C T A G A C G T A G T C C G T A C T G A G T A C A C T G | WRKY65(WRKY)/colamp-WRKY65-DAP-Seq(GSE60143)/Homer | 1e-183 | -4.224e+02 | 0.0000 | 9904.0 | 30.68% | 33372.5 | 22.89% | motif file (matrix) | svg |
| 261 | T C G A T G A C G T A C C G T A C A G T T G A C A C G T A C T G A G C T A G C T | NeuroG2(bHLH)/Fibroblast-NeuroG2-ChIP-Seq(GSE75910)/Homer | 1e-183 | -4.219e+02 | 0.0000 | 16989.0 | 52.63% | 63789.4 | 43.76% | motif file (matrix) | svg |
| 262 | C G A T C G T A G C T A G A C T T C G A A G C T A G T C A C T G T C G A A G C T C T G A C G A T | ZBTB38(Zf)/Hela-ZBTB38-ChIP-seq(GSE108618)/Homer | 1e-182 | -4.211e+02 | 0.0000 | 28862.0 | 89.41% | 121292.0 | 83.20% | motif file (matrix) | svg |
| 263 | T C A G T G A C C A T G G C A T C A G T A C T G C G T A T G A C G A C T C G A T C G A T C G T A | WRKY3(WRKY)/col-WRKY3-DAP-Seq(GSE60143)/Homer | 1e-182 | -4.204e+02 | 0.0000 | 11967.0 | 37.07% | 41980.2 | 28.80% | motif file (matrix) | svg |
| 264 | C G T A G C A T C A T G C T A G A G T C A C T G A T C G G T A C A C T G T C A G | At2g33710(AP2EREBP)/colamp-At2g33710-DAP-Seq(GSE60143)/Homer | 1e-182 | -4.191e+02 | 0.0000 | 26814.0 | 83.06% | 110528.9 | 75.82% | motif file (matrix) | svg |
| 265 | T A G C C A T G G A C T G A C T T C A G G T C A G A T C G A C T G C A T G C T A | WRKY15(WRKY)/col-WRKY15-DAP-Seq(GSE60143)/Homer | 1e-179 | -4.139e+02 | 0.0000 | 16318.0 | 50.55% | 60921.7 | 41.79% | motif file (matrix) | svg |
| 266 | T C G A A G C T A C G T A C G T A G T C A G T C A C G T A T C G G A C T A T C G | EWS:ERG-fusion(ETS)/CADO\_ES1-EWS:ERG-ChIP-Seq(SRA014231)/Homer | 1e-178 | -4.101e+02 | 0.0000 | 7801.0 | 24.17% | 25044.5 | 17.18% | motif file (matrix) | svg |
| 267 | G T A C G A T C C A G T A G T C A G T C A G T C T G C A G A T C C T G A A T G C G T C A A C G T | WT1(Zf)/Kidney-WT1-ChIP-Seq(GSE90016)/Homer | 1e-176 | -4.057e+02 | 0.0000 | 8585.0 | 26.59% | 28225.0 | 19.36% | motif file (matrix) | svg |
| 268 | C G A T C G T A G C T A G C A T G C A T C T G A A C T G A C G T A G T C C G T A C G T A G T A C T C A G G C T A C G A T | WRKY25(WRKY)/colamp-WRKY25-DAP-Seq(GSE60143)/Homer | 1e-175 | -4.047e+02 | 0.0000 | 17851.0 | 55.30% | 67930.1 | 46.60% | motif file (matrix) | svg |
| 269 | C T A G A C T G T G C A G T C A A T G C C G T A A T C G A T G C A G T C C T A G | ZNF341(Zf)/EBV-ZNF341-ChIP-Seq(GSE113194)/Homer | 1e-175 | -4.043e+02 | 0.0000 | 10875.0 | 33.69% | 37624.6 | 25.81% | motif file (matrix) | svg |
| 270 | G A T C C T G A G A T C G A T C C T A G G C T A A G T C C T G A G C T A C G T A | At4g16750(AP2EREBP)/col-At4g16750-DAP-Seq(GSE60143)/Homer | 1e-173 | -4.006e+02 | 0.0000 | 23033.0 | 71.35% | 92098.1 | 63.18% | motif file (matrix) | svg |
| 271 | G A T C C G T A G A C T C T A G G A T C C T G A G A C T C T G A G A C T C T A G G A T C C T G A G A C T C T G A G A C T | OCT:OCT(POU,Homeobox)/NPC-OCT6-ChIP-Seq(GSE43916)/Homer | 1e-173 | -4.005e+02 | 0.0000 | 1088.0 | 3.37% | 1522.9 | 1.04% | motif file (matrix) | svg |
| 272 | C G A T A G T C C A T G G A C T A C G T C T A G C G T A G A T C G A C T C G A T G C A T G A C T | WRKY43(WRKY)/colamp-WRKY43-DAP-Seq(GSE60143)/Homer | 1e-173 | -3.998e+02 | 0.0000 | 7233.0 | 22.41% | 22922.4 | 15.72% | motif file (matrix) | svg |
| 273 | C G T A C G A T C T A G G T C A G A C T C G A T C T A G C G T A A C G T C A T G | LIN-39(Homeobox)/cElegans.L3-LIN39-ChIP-Seq(modEncode)/Homer | 1e-171 | -3.946e+02 | 0.0000 | 13618.0 | 42.19% | 49397.6 | 33.89% | motif file (matrix) | svg |
| 274 | G A T C G C A T T C G A A G T C A C G T A C G T A C G T C G A T A C G T A T C G | AT1G47655(C2C2dof)/colamp-AT1G47655-DAP-Seq(GSE60143)/Homer | 1e-170 | -3.930e+02 | 0.0000 | 26110.0 | 80.88% | 107320.7 | 73.62% | motif file (matrix) | svg |
| 275 | G C T A C G T A G C T A C G T A C T G A A C T G A C G T G T A C C G T A C T G A G T A C A C T G | WRKY22(WRKY)/colamp-WRKY22-DAP-Seq(GSE60143)/Homer | 1e-169 | -3.906e+02 | 0.0000 | 10496.0 | 32.51% | 36235.9 | 24.86% | motif file (matrix) | svg |
| 276 | A C G T T C G A T C G A A G T C G T C A T A C G A T G C A C G T A C T G A G C T | Myf5(bHLH)/GM-Myf5-ChIP-Seq(GSE24852)/Homer | 1e-169 | -3.894e+02 | 0.0000 | 8040.0 | 24.91% | 26239.2 | 18.00% | motif file (matrix) | svg |
| 277 | C G A T C G T A A G T C A C G T A C G T T C A G G C A T C G T A G C T A G C T A C G T A A G T C C G T A G T C A A C T G | ANAC058(NAC)/col-ANAC058-DAP-Seq(GSE60143)/Homer | 1e-168 | -3.891e+02 | 0.0000 | 11595.0 | 35.92% | 40852.8 | 28.02% | motif file (matrix) | svg |
| 278 | G C T A T C G A C G T A C T G A A C T G A C G T A G T C C G T A C G T A A G T C C T A G T G C A | WRKY42(WRKY)/colamp-WRKY42-DAP-Seq(GSE60143)/Homer | 1e-168 | -3.883e+02 | 0.0000 | 9977.0 | 30.91% | 34118.2 | 23.40% | motif file (matrix) | svg |
| 279 | T C G A T C G A C T G A C G T A A C T G A T G C A C G T A G T C | Lola-I(Zf)/Embryo-LolaI-ChIP-Seq(GSE200870)/Homer | 1e-167 | -3.848e+02 | 0.0000 | 8284.0 | 25.66% | 27272.3 | 18.71% | motif file (matrix) | svg |
| 280 | G T A C A C G T A C G T T A C G A T G C C A T G T A C G G T A C T C A G A T G C C G T A G T C A A C T G A G C T G C T A | AT1G19040(NAC)/col-AT1G19040-DAP-Seq(GSE60143)/Homer | 1e-166 | -3.844e+02 | 0.0000 | 3390.0 | 10.50% | 8735.9 | 5.99% | motif file (matrix) | svg |
| 281 | A C G T C T A G A G C T A C G T A C G T C T G A A G T C G A C T A G C T C G T A | FOXM1(Forkhead)/MCF7-FOXM1-ChIP-Seq(GSE72977)/Homer | 1e-162 | -3.745e+02 | 0.0000 | 10529.0 | 32.62% | 36595.0 | 25.10% | motif file (matrix) | svg |
| 282 | G A C T G A T C C T G A A G T C A G T C A C T G C G T A A G T C G T A C G C T A G C A T C G A T | At1g19210(AP2EREBP)/colamp-At1g19210-DAP-Seq(GSE60143)/Homer | 1e-162 | -3.745e+02 | 0.0000 | 26361.0 | 81.66% | 108855.7 | 74.67% | motif file (matrix) | svg |
| 283 | A C T G A C G T C A T G A T C G A T C G T G A C A C T G A T C G A T C G T G C A C T G A C G T A | E2F3(E2F)/MEF-E2F3-ChIP-Seq(GSE71376)/Homer | 1e-161 | -3.718e+02 | 0.0000 | 14875.0 | 46.08% | 55217.8 | 37.88% | motif file (matrix) | svg |
| 284 | C A T G T G A C C A T G G A C T C A G T C T A G G C T A G T A C G A C T G C A T G C A T C G A T | WRKY21(WRKY)/colamp-WRKY21-DAP-Seq(GSE60143)/Homer | 1e-158 | -3.647e+02 | 0.0000 | 2508.0 | 7.77% | 5885.6 | 4.04% | motif file (matrix) | svg |
| 285 | C T A G C A T G A C G T C G T A C T A G C A T G C G A T C T A G T C A G T C A G | MYB3(MYB)/Arabidopsis-MYB3-ChIP-Seq(GSE80564)/Homer | 1e-155 | -3.573e+02 | 0.0000 | 23026.0 | 71.33% | 92756.0 | 63.63% | motif file (matrix) | svg |
| 286 | T C G A C T G A C G A T C G T A C G T A C G T A C T A G A G C T C T G A T C A G | Adof1(C2C2dof)/col-Adof1-DAP-Seq(GSE60143)/Homer | 1e-154 | -3.547e+02 | 0.0000 | 23445.0 | 72.63% | 94809.1 | 65.04% | motif file (matrix) | svg |
| 287 | C G T A C G T A C G T A C G T A C G T A A C T G A C T G A G T C | dof42(C2C2dof)/col-dof42-DAP-Seq(GSE60143)/Homer | 1e-151 | -3.491e+02 | 0.0000 | 10088.0 | 31.25% | 35121.9 | 24.09% | motif file (matrix) | svg |
| 288 | A C T G G A T C G A C T A C T G A C G T C A T G A C T G A C G T A G C T C G A T | RUNX-AML(Runt)/CD4+-PolII-ChIP-Seq(Barski\_et\_al.)/Homer | 1e-150 | -3.457e+02 | 0.0000 | 9870.0 | 30.58% | 34265.9 | 23.51% | motif file (matrix) | svg |
| 289 | C A T G T G C A G A C T C A T G C G T A A G T C T C A G G C A T T G A C C G T A | bZIP50(bZIP)/colamp-bZIP50-DAP-Seq(GSE60143)/Homer | 1e-150 | -3.457e+02 | 0.0000 | 21803.0 | 67.54% | 87140.8 | 59.78% | motif file (matrix) | svg |
| 290 | T C G A T C G A A G T C C G T A C T A G T A G C A C G T A C T G | MyoG(bHLH)/C2C12-MyoG-ChIP-Seq(GSE36024)/Homer | 1e-150 | -3.454e+02 | 0.0000 | 11501.0 | 35.63% | 41099.7 | 28.19% | motif file (matrix) | svg |
| 291 | C G T A A C T G C G T A A C G T A T C G C A G T T A G C C G T A T C G A G T A C C T G A T A G C C G T A A C T G C G T A A C G T C G T A C T G A A T C G G C T A | GATA3(Zf),DR8/iTreg-Gata3-ChIP-Seq(GSE20898)/Homer | 1e-149 | -3.446e+02 | 0.0000 | 1673.0 | 5.18% | 3356.8 | 2.30% | motif file (matrix) | svg |
| 292 | T G A C A G T C C G T A A C T G G T A C A C G T A C T G A C G T G A C T G A T C | Twist2(bHLH)/Myoblast-Twist2.Ty1-ChIP-Seq(GSE127998)/Homer | 1e-148 | -3.421e+02 | 0.0000 | 17896.0 | 55.44% | 69165.2 | 47.45% | motif file (matrix) | svg |
| 293 | G T A C G T C A G T A C G T C A G T A C G T C A G T A C G T C A G T A C G T C A | SeqBias: CA-repeat | 1e-147 | -3.393e+02 | 0.0000 | 28843.0 | 89.35% | 122220.9 | 83.84% | motif file (matrix) | svg |
| 294 | C A T G C T G A A G T C A C T G A C T G A G C T A C T G A T C G | ESE3(AP2EREBP)/col-ESE3-DAP-Seq(GSE60143)/Homer | 1e-146 | -3.381e+02 | 0.0000 | 24341.0 | 75.40% | 99417.7 | 68.20% | motif file (matrix) | svg |
| 295 | A T C G T C G A G A C T A T C G T G A C A C G T C T A G A C T G C G T A A C T G A G T C G T A C | ZNF415(Zf)/HEK293-ZNF415.GFP-ChIP-Seq(GSE58341)/Homer | 1e-145 | -3.362e+02 | 0.0000 | 9114.0 | 28.23% | 31283.1 | 21.46% | motif file (matrix) | svg |
| 296 | A C T G T C A G A G C T G A C T C A T G A G T C A G T C G C T A C G A T C T A G T C A G G T A C C T G A T C G A | Rfx1(HTH)/NPC-H3K4me1-ChIP-Seq(GSE16256)/Homer | 1e-144 | -3.328e+02 | 0.0000 | 3820.0 | 11.83% | 10671.0 | 7.32% | motif file (matrix) | svg |
| 297 | T C G A T G A C G T A C C G T A A C G T G A C T A C G T A C T G A C T G A G C T | Mesp1(bHLH)/ESC-Mesp1-ChIP-Seq(GSE165102)/Homer | 1e-140 | -3.241e+02 | 0.0000 | 8668.0 | 26.85% | 29625.5 | 20.32% | motif file (matrix) | svg |
| 298 | A C G T T G C A A G C T G A T C C T A G C T G A A G C T G T C A T C G A C G T A | CUX1(Homeobox)/K562-CUX1-ChIP-Seq(GSE92882)/Homer | 1e-140 | -3.237e+02 | 0.0000 | 16911.0 | 52.39% | 65049.0 | 44.62% | motif file (matrix) | svg |
| 299 | A G C T C A T G G C A T G A T C T G C A C T A G G A T C A C G T | Tgif2(Homeobox)/mES-Tgif2-ChIP-Seq(GSE55404)/Homer | 1e-140 | -3.237e+02 | 0.0000 | 26158.0 | 81.03% | 108608.6 | 74.50% | motif file (matrix) | svg |
| 300 | T A C G T G C A A G T C C G T A A C G T T G A C A C G T A C T G A C T G G C A T | TCF4(bHLH)/SHSY5Y-TCF4-ChIP-Seq(GSE96915)/Homer | 1e-139 | -3.221e+02 | 0.0000 | 16476.0 | 51.04% | 63130.2 | 43.31% | motif file (matrix) | svg |
| 301 | C A G T G A C T G C A T T C G A A G T C A C G T A C G T A C G T C G A T G A C T | OBP3(C2C2dof)/col-OBP3-DAP-Seq(GSE60143)/Homer | 1e-139 | -3.218e+02 | 0.0000 | 24123.0 | 74.73% | 98630.9 | 67.66% | motif file (matrix) | svg |
| 302 | G T A C G C T A C G A T C A G T A G T C G C T A C G A T C G A T A G T C G C T A | WUS1(Homeobox)/colamp-WUS1-DAP-Seq(GSE60143)/Homer | 1e-139 | -3.207e+02 | 0.0000 | 6452.0 | 19.99% | 20776.0 | 14.25% | motif file (matrix) | svg |
| 303 | C G T A G C A T C A G T C T A G A G T C A C T G A C T G G T A C A C T G A T C G | ERF115(AP2EREBP)/colamp-ERF115-DAP-Seq(GSE60143)/Homer | 1e-139 | -3.205e+02 | 0.0000 | 26081.0 | 80.79% | 108274.7 | 74.28% | motif file (matrix) | svg |
| 304 | G C A T C G T A G C T A G A C T C G A T G A C T A G T C C A G T A G T C A G T C A C T G C T A G G T A C C T A G C T G A | AT5G05550(Trihelix)/col-AT5G05550-DAP-Seq(GSE60143)/Homer | 1e-139 | -3.203e+02 | 0.0000 | 25461.0 | 78.87% | 105198.2 | 72.16% | motif file (matrix) | svg |
| 305 | C G A T T C G A A G T C A C G T A C G T T A C G C G T A G C T A G C T A C G A T G C A T A T G C C G T A G T C A A C T G | ANAC071(NAC)/col-ANAC071-DAP-Seq(GSE60143)/Homer | 1e-137 | -3.162e+02 | 0.0000 | 15853.0 | 49.11% | 60460.8 | 41.48% | motif file (matrix) | svg |
| 306 | C T G A A G T C G A T C C A T G G C T A G A T C C T G A G C T A G C T A C G A T | AT1G77200(AP2EREBP)/colamp-AT1G77200-DAP-Seq(GSE60143)/Homer | 1e-135 | -3.128e+02 | 0.0000 | 23221.0 | 71.93% | 94442.2 | 64.79% | motif file (matrix) | svg |
| 307 | G A C T C G A T C G A T C T G A G T A C A G T C C G A T C G T A G T C A G A T C G C A T G C A T | MYB121(MYB)/col-MYB121-DAP-Seq(GSE60143)/Homer | 1e-135 | -3.120e+02 | 0.0000 | 8553.0 | 26.50% | 29324.4 | 20.12% | motif file (matrix) | svg |
| 308 | C T G A C T A G A C T G G C A T A T G C C G T A C T G A C T A G A C T G A C G T A G T C C T G A | RARg(NR)/ES-RARg-ChIP-Seq(GSE30538)/Homer | 1e-134 | -3.096e+02 | 0.0000 | 782.0 | 2.42% | 1031.6 | 0.71% | motif file (matrix) | svg |
| 309 | A T C G T G A C A T G C C T G A T C A G G A C T A G T C C G A T T C A G T C G A C A T G C T A G C T A G C G T A C T A G C T A G C T G A C T A G C T A G A T G C | ZSCAN22(Zf)/HEK293-ZSCAN22.GFP-ChIP-Seq(GSE58341)/Homer | 1e-134 | -3.094e+02 | 0.0000 | 983.0 | 3.05% | 1538.9 | 1.06% | motif file (matrix) | svg |
| 310 | C T G A A T G C G C T A C G A T A T G C C G T A C G T A C G T A C T A G T A C G | Tcf3(HMG)/mES-Tcf3-ChIP-Seq(GSE11724)/Homer | 1e-131 | -3.027e+02 | 0.0000 | 4082.0 | 12.65% | 11905.2 | 8.17% | motif file (matrix) | svg |
| 311 | A G T C C G T A A C G T A G T C A C G T A C T G | Tal1 | 1e-131 | -3.025e+02 | 0.0000 | 16249.0 | 50.34% | 62466.1 | 42.85% | motif file (matrix) | svg |
| 312 | A G T C A G T C C T G A A G T C A G T C A C T G C G T A A G T C T C G A G A T C C G A T C G T A | AT1G01250(AP2EREBP)/col-AT1G01250-DAP-Seq(GSE60143)/Homer | 1e-130 | -2.994e+02 | 0.0000 | 4957.0 | 15.36% | 15228.1 | 10.45% | motif file (matrix) | svg |
| 313 | A G T C T A G C G A C T A C G T C T A G A C G T A C G T A C G T C T G A A G T C G C T A G A C T C G T A C T A G A C T G | Foxa3(Forkhead)/Liver-Foxa3-ChIP-Seq(GSE77670)/Homer | 1e-129 | -2.986e+02 | 0.0000 | 4499.0 | 13.94% | 13503.0 | 9.26% | motif file (matrix) | svg |
| 314 | C G A T C T A G A G T C A C G T A C G T T C A G G C T A C G T A G C A T G C A T C G A T A G T C C G T A G T C A A C T G | VND3(NAC)/colamp-VND3-DAP-Seq(GSE60143)/Homer | 1e-128 | -2.963e+02 | 0.0000 | 11967.0 | 37.07% | 43866.3 | 30.09% | motif file (matrix) | svg |
| 315 | T G A C C T G A C T A G T C G A C T G A A T G C C G T A A C T G G C A T G T A C G C A T A T C G G C A T A G C T G A T C | PR(NR)/T47D-PR-ChIP-Seq(GSE31130)/Homer | 1e-126 | -2.910e+02 | 0.0000 | 20265.0 | 62.78% | 80929.5 | 55.52% | motif file (matrix) | svg |
| 316 | C T G A C T G A C T G A A T G C G A T C C A T G A C T G G A C T G A C T G C A T C G T A C G T A A G T C G T A C C T G A A T C G G C A T G A C T G A C T A G C T | GRHL2(CP2)/HBE-GRHL2-ChIP-Seq(GSE46194)/Homer | 1e-126 | -2.904e+02 | 0.0000 | 6514.0 | 20.18% | 21400.7 | 14.68% | motif file (matrix) | svg |
| 317 | T G A C C T G A A G T C A G T C A C T G G A T C G A C T G C A T | At5g18450(AP2EREBP)/col-At5g18450-DAP-Seq(GSE60143)/Homer | 1e-125 | -2.896e+02 | 0.0000 | 24726.0 | 76.60% | 102112.5 | 70.05% | motif file (matrix) | svg |
| 318 | A G T C A C G T A C G T T C A G G C T A C G T A G C T A G C A T C G A T A G T C C G T A G T C A A C T G G A C T G C T A | SMB(NAC)/colamp-SMB-DAP-Seq(GSE60143)/Homer | 1e-125 | -2.892e+02 | 0.0000 | 16105.0 | 49.89% | 62068.6 | 42.58% | motif file (matrix) | svg |
| 319 | G C A T C G T A G C A T C G T A T C G A C G T A C T G A A C T G C G T A C G T A C G T A A C G T A C T G G T C A G C A T | AT2G31460(REMB3)/col-AT2G31460-DAP-Seq(GSE60143)/Homer | 1e-124 | -2.877e+02 | 0.0000 | 4211.0 | 13.04% | 12534.2 | 8.60% | motif file (matrix) | svg |
| 320 | G T A C G C T A T C A G C T G A C T A G C A T G A G C T G A T C T G C A T C G A C T G A A C T G C A G T A G T C G A T C G C T A | HNF4a(NR),DR1/HepG2-HNF4a-ChIP-Seq(GSE25021)/Homer | 1e-124 | -2.870e+02 | 0.0000 | 5215.0 | 16.16% | 16357.0 | 11.22% | motif file (matrix) | svg |
| 321 | C T A G T C G A C T G A C G T A T A C G G A C T T C A G T C G A G T C A T G C A T A C G A G C T | IRF2(IRF)/Erythroblas-IRF2-ChIP-Seq(GSE36985)/Homer | 1e-124 | -2.864e+02 | 0.0000 | 1759.0 | 5.45% | 3927.0 | 2.69% | motif file (matrix) | svg |
| 322 | G A C T G T A C T G C A A C G T G A T C G C T A T C G A A C G T A G T C C G T A | Pdx1(Homeobox)/Islet-Pdx1-ChIP-Seq(SRA008281)/Homer | 1e-123 | -2.833e+02 | 0.0000 | 13084.0 | 40.53% | 48901.0 | 33.55% | motif file (matrix) | svg |
| 323 | C G A T T C A G G T A C A C G T A C G T T C A G C G A T C G T A G T C A G C T A C G T A A G T C C G T A G T C A C A T G | ANAC057(NAC)/colamp-ANAC057-DAP-Seq(GSE60143)/Homer | 1e-122 | -2.813e+02 | 0.0000 | 14466.0 | 44.81% | 54964.7 | 37.71% | motif file (matrix) | svg |
| 324 | G A C T A C T G C G T A A G T C T C A G G C A T G T A C C G T A A C G T G A T C | TGA1(bZIP)/colamp-TGA1-DAP-Seq(GSE60143)/Homer | 1e-121 | -2.806e+02 | 0.0000 | 9815.0 | 30.40% | 35020.4 | 24.02% | motif file (matrix) | svg |
| 325 | G A C T C T A G A T G C A G T C G T C A T A C G A T G C A T C G | HIC1(Zf)/Treg-ZBTB29-ChIP-Seq(GSE99889)/Homer | 1e-121 | -2.788e+02 | 0.0000 | 22581.0 | 69.95% | 92006.9 | 63.12% | motif file (matrix) | svg |
| 326 | C A G T T C A G A G C T G A C T A C G T A G T C G A T C G A C T C T G A A C T G G A T C C G T A C T G A A G T C G T A C | Rfx6(HTH)/Min6b1-Rfx6.HA-ChIP-Seq(GSE62844)/Homer | 1e-120 | -2.773e+02 | 0.0000 | 14627.0 | 45.31% | 55745.2 | 38.24% | motif file (matrix) | svg |
| 327 | T A C G T A C G G T A C A T C G A C T G T A C G T C G A C T G A T C G A A T C G | E2F6(E2F)/Hela-E2F6-ChIP-Seq(GSE31477)/Homer | 1e-120 | -2.766e+02 | 0.0000 | 10740.0 | 33.27% | 38977.9 | 26.74% | motif file (matrix) | svg |
| 328 | T C G A C T G A T A G C T G A C T C A G T C A G C G T A C G T A T C A G A G C T | ETV1(ETS)/GIST48-ETV1-ChIP-Seq(GSE22441)/Homer | 1e-119 | -2.744e+02 | 0.0000 | 18068.0 | 55.97% | 71175.9 | 48.83% | motif file (matrix) | svg |
| 329 | C G A T G T C A A G T C A C G T A C G T A C T G G A C T C G A T A T C G G C T A G T C A A G T C C G T A G T C A A C T G | ANAC017(NAC)/colamp-ANAC017-DAP-Seq(GSE60143)/Homer | 1e-118 | -2.728e+02 | 0.0000 | 3666.0 | 11.36% | 10658.9 | 7.31% | motif file (matrix) | svg |
| 330 | G C A T A C G T A C T G A C G T A G T C A C T G A T C G G T C A C G A T C G T A | ARF2(ARF)/col-ARF2-DAP-Seq(GSE60143)/Homer | 1e-117 | -2.703e+02 | 0.0000 | 26620.0 | 82.46% | 111787.4 | 76.68% | motif file (matrix) | svg |
| 331 | C A G T A G C T G C A T T C G A A G T C A C G T A C G T A C G T C G A T G C A T | AT5G66940(C2C2dof)/col-AT5G66940-DAP-Seq(GSE60143)/Homer | 1e-117 | -2.703e+02 | 0.0000 | 18904.0 | 58.56% | 75062.0 | 51.49% | motif file (matrix) | svg |
| 332 | C T G A T C A G C A G T C T A G A C T G C T A G G A T C A T C G A C T G C T G A T C A G G A T C | Sp5(Zf)/mES-Sp5.Flag-ChIP-Seq(GSE72989)/Homer | 1e-115 | -2.653e+02 | 0.0000 | 12138.0 | 37.60% | 45138.5 | 30.96% | motif file (matrix) | svg |
| 333 | A T G C G T A C A C T G A G T C A G T C A C T G G A T C G T C A C G T A C G A T G C A T C G A T | RRTF1(AP2EREBP)/colamp-RRTF1-DAP-Seq(GSE60143)/Homer | 1e-113 | -2.625e+02 | 0.0000 | 11853.0 | 36.72% | 43964.6 | 30.16% | motif file (matrix) | svg |
| 334 | C G A T C T A G C T G A A T G C C T G A T C G A C G T A C T G A T C G A T A G C A G T C C G T A A C T G T C G A A T G C | Hand2(bHLH)/Mesoderm-Hand2-ChIP-Seq(GSE61475)/Homer | 1e-113 | -2.621e+02 | 0.0000 | 5276.0 | 16.34% | 16888.3 | 11.59% | motif file (matrix) | svg |
| 335 | C G A T T A C G T G C A G T A C G A T C G A C T A G C T A C G T A T C G G T A C G A T C G T A C G A T C G T C A | PPARE(NR),DR1/3T3L1-Pparg-ChIP-Seq(GSE13511)/Homer | 1e-113 | -2.603e+02 | 0.0000 | 9859.0 | 30.54% | 35535.5 | 24.38% | motif file (matrix) | svg |
| 336 | A G T C C T G A A T C G A G C T A G C T G A C T A G T C G C T A A C G T C G A T G C A T C G A T A T C G C G T A T A G C G C A T A T G C C G T A | bZIP:IRF(bZIP,IRF)/Th17-BatF-ChIP-Seq(GSE39756)/Homer | 1e-112 | -2.598e+02 | 0.0000 | 3741.0 | 11.59% | 11063.7 | 7.59% | motif file (matrix) | svg |
| 337 | C T A G C T A G T G A C G T A C C A T G A C T G G A T C G A T C C G T A C G T A | RAP211(AP2EREBP)/colamp-RAP211-DAP-Seq(GSE60143)/Homer | 1e-111 | -2.566e+02 | 0.0000 | 26234.0 | 81.27% | 110098.9 | 75.53% | motif file (matrix) | svg |
| 338 | A G T C C G A T A C T G A T C G T G A C G C T A C A T G A T C G T G A C C G A T A C T G T A G C G T A C G T C A | Tlx?(NR)/NPC-H3K4me1-ChIP-Seq(GSE16256)/Homer | 1e-111 | -2.565e+02 | 0.0000 | 4565.0 | 14.14% | 14209.7 | 9.75% | motif file (matrix) | svg |
| 339 | C T G A G A C T G A T C C T G A A G T C G C A T A C G T G A C T G C T A G C A T | OBP1(C2C2dof)/col-OBP1-DAP-Seq(GSE60143)/Homer | 1e-110 | -2.550e+02 | 0.0000 | 23067.0 | 71.46% | 94764.8 | 65.01% | motif file (matrix) | svg |
| 340 | C G A T C T A G A C G T G T C A C G T A C G T A A G T C C G T A | Foxo3(Forkhead)/U2OS-Foxo3-ChIP-Seq(E-MTAB-2701)/Homer | 1e-110 | -2.537e+02 | 0.0000 | 9732.0 | 30.15% | 35113.9 | 24.09% | motif file (matrix) | svg |
| 341 | C G A T C G T A A G T C A C G T A C G T T C A G C G T A C G T A G C A T G C A T G C A T A G T C C G T A G T C A A C T G | VND2(NAC)/col-VND2-DAP-Seq(GSE60143)/Homer | 1e-109 | -2.528e+02 | 0.0000 | 16147.0 | 50.02% | 62951.9 | 43.18% | motif file (matrix) | svg |
| 342 | C T A G A G T C T A C G T A C G T G A C C G T A A C T G T A G C G C A T C A T G A T G C A G C T | Ascl1(bHLH)/NeuralTubes-Ascl1-ChIP-Seq(GSE55840)/Homer | 1e-109 | -2.516e+02 | 0.0000 | 15943.0 | 49.39% | 62065.2 | 42.58% | motif file (matrix) | svg |
| 343 | T C G A C G T A A G T C C G T A C T A G A G T C C G A T A C T G G A C T A G C T A C T G G A C T | HLH-1(bHLH)/cElegans-Embryo-HLH1-ChIP-Seq(modEncode)/Homer | 1e-108 | -2.489e+02 | 0.0000 | 10284.0 | 31.86% | 37521.9 | 25.74% | motif file (matrix) | svg |
| 344 | G A C T C T A G C T A G G T A C A G T C G A T C G A C T G A C T T A G C T C A G | NLP7(RWPRK)/col-NLP7-DAP-Seq(GSE60143)/Homer | 1e-108 | -2.487e+02 | 0.0000 | 21754.0 | 67.39% | 88685.7 | 60.84% | motif file (matrix) | svg |
| 345 | T C A G G A C T G T C A C G T A A C G T A T C G C G T A A C G T A C G T C T G A | ATHB15(HB)/col-ATHB15-DAP-Seq(GSE60143)/Homer | 1e-106 | -2.457e+02 | 0.0000 | 6021.0 | 18.65% | 20033.7 | 13.74% | motif file (matrix) | svg |
| 346 | C G T A C T A G T C A G T C A G A G T C A T G C A G T C G C A T A G C T A C G T A T C G C G A T | Sox9(HMG)/Limb-SOX9-ChIP-Seq(GSE73225)/Homer | 1e-106 | -2.455e+02 | 0.0000 | 10933.0 | 33.87% | 40336.5 | 27.67% | motif file (matrix) | svg |
| 347 | C A G T C A T G T G C A G T A C C G T A T C A G G T A C G A C T T C A G C T G A | bZIP18(bZIP)/colamp-bZIP18-DAP-Seq(GSE60143)/Homer | 1e-105 | -2.433e+02 | 0.0000 | 28766.0 | 89.11% | 123172.6 | 84.49% | motif file (matrix) | svg |
| 348 | A G T C A C G T A C G T T A C G G C T A G C T A C G T A C G A T C G A T A T G C C G T A G T C A A C T G G A C T G C A T | SND2(NAC)/colamp-SND2-DAP-Seq(GSE60143)/Homer | 1e-105 | -2.422e+02 | 0.0000 | 11904.0 | 36.88% | 44554.4 | 30.56% | motif file (matrix) | svg |
| 349 | G C T A G C T A C T G A A C T G A C G T A G T C C G T A C G T A G T A C A C T G A T G C G C A T | WRKY47(WRKY)/colamp-WRKY47-DAP-Seq(GSE60143)/Homer | 1e-104 | -2.417e+02 | 0.0000 | 7057.0 | 21.86% | 24255.1 | 16.64% | motif file (matrix) | svg |
| 350 | C A G T G C T A G C A T T A C G C T G A C A G T T A G C C T G A | GATA15(C2C2gata)/col-GATA15-DAP-Seq(GSE60143)/Homer | 1e-104 | -2.415e+02 | 0.0000 | 22438.0 | 69.51% | 92049.3 | 63.14% | motif file (matrix) | svg |
| 351 | T A C G C T G A C A T G G A T C G T A C G C A T T C A G T A C G A G C T G T C A G A T C G C A T T A C G C G T A C T A G G A T C G A T C C G A T A C T G T C A G | ZNF322(Zf)/HEK293-ZNF322.GFP-ChIP-Seq(GSE58341)/Homer | 1e-104 | -2.412e+02 | 0.0000 | 2677.0 | 8.29% | 7359.4 | 5.05% | motif file (matrix) | svg |
| 352 | A T G C G A C T A G C T C T A G C G T A C T A G C G A T C T A G A T C G G A T C | Nkx2.2(Homeobox)/NPC-Nkx2.2-ChIP-Seq(GSE61673)/Homer | 1e-103 | -2.382e+02 | 0.0000 | 22360.0 | 69.27% | 91746.2 | 62.94% | motif file (matrix) | svg |
| 353 | C T G A T C A G C T G A C T A G C A T G A C G T A T G C C G T A A T G C G C A T T C A G C T G A A C T G A C G T C A G T A G T C C G T A C A G T C T A G C A T G | VDR(NR),DR3/GM10855-VDR+vitD-ChIP-Seq(GSE22484)/Homer | 1e-101 | -2.332e+02 | 0.0000 | 3520.0 | 10.90% | 10513.6 | 7.21% | motif file (matrix) | svg |
| 354 | C G A T G A T C T A C G C T G A G C T A C G T A G C A T A G T C C T A G C G T A G C A T C G A T | AT2G15740(C2H2)/col-AT2G15740-DAP-Seq(GSE60143)/Homer | 1e-100 | -2.322e+02 | 0.0000 | 24720.0 | 76.58% | 103135.4 | 70.75% | motif file (matrix) | svg |
| 355 | T C A G A G C T A C G T A C G T G T A C G A T C C G T A C T A G C A T G G T C A C G T A T C G A | STAT4(Stat)/CD4-Stat4-ChIP-Seq(GSE22104)/Homer | 1e-95 | -2.207e+02 | 0.0000 | 10009.0 | 31.01% | 36862.2 | 25.29% | motif file (matrix) | svg |
| 356 | C T G A A C T G C G T A A C G T G T C A A G C T A G C T G A C T G A C T C A G T | CCA(Myb)/Arabidopsis-CCA.GFP-ChIP-Seq(GSE70533)/Homer | 1e-94 | -2.181e+02 | 0.0000 | 12856.0 | 39.83% | 49143.5 | 33.71% | motif file (matrix) | svg |
| 357 | G T C A C G T A A C G T A T C G C G T A A C G T A C G T C T A G | ATHB7(Homeobox)/col-ATHB7-DAP-Seq(GSE60143)/Homer | 1e-92 | -2.137e+02 | 0.0000 | 11860.0 | 36.74% | 44914.4 | 30.81% | motif file (matrix) | svg |
| 358 | A C G T T G A C A G T C A G C T A G T C A G C T A C T G G A C T A G C T G A C T | REF6(Zf)/Arabidopsis-REF6-ChIP-Seq(GSE106942)/Homer | 1e-92 | -2.134e+02 | 0.0000 | 6354.0 | 19.68% | 21835.8 | 14.98% | motif file (matrix) | svg |
| 359 | T A C G A T G C G A C T A C T G A G C T A G T C G T C A T G C A A C G T A G T C G C T A T G C A | Pknox1(Homeobox)/ES-Prep1-ChIP-Seq(GSE63282)/Homer | 1e-92 | -2.133e+02 | 0.0000 | 4401.0 | 13.63% | 14093.6 | 9.67% | motif file (matrix) | svg |
| 360 | C A G T A G C T C G T A G C A T A G T C G A C T C T A G C T A G C A G T C T A G T C G A T G C A C T A G C A T G G A C T | STOP1(C2H2)/colamp-STOP1-DAP-Seq(GSE60143)/Homer | 1e-89 | -2.070e+02 | 0.0000 | 7445.0 | 23.06% | 26390.8 | 18.10% | motif file (matrix) | svg |
| 361 | T A C G T C G A G A C T A C T G C T G A A G T C T C A G G A C T T G A C C T G A | Atf1(bZIP)/K562-ATF1-ChIP-Seq(GSE31477)/Homer | 1e-89 | -2.058e+02 | 0.0000 | 13941.0 | 43.19% | 54148.0 | 37.14% | motif file (matrix) | svg |
| 362 | G C A T G C A T G C A T A T G C A G C T T C G A T A C G G C T A C G T A C A T G G T A C G C A T G C A T A G T C A G C T | HSFA6B(HSF)/colamp-HSFA6B-DAP-Seq(GSE60143)/Homer | 1e-88 | -2.047e+02 | 0.0000 | 6141.0 | 19.02% | 21107.4 | 14.48% | motif file (matrix) | svg |
| 363 | C T G A C G A T C T A G C G T A A G C T C G A T C A G T C T G A G A C T C T A G C T A G A T G C | PBX2(Homeobox)/K562-PBX2-ChIP-Seq(Encode)/Homer | 1e-88 | -2.037e+02 | 0.0000 | 13031.0 | 40.37% | 50204.9 | 34.44% | motif file (matrix) | svg |
| 364 | C T A G T A C G G A C T T G C A T G C A C G A T T A C G C T G A T C G A C T G A | Hoxa10(Homeobox)/ChickenMSG-Hoxa10.Flag-ChIP-Seq(GSE86088)/Homer | 1e-86 | -1.997e+02 | 0.0000 | 7870.0 | 24.38% | 28272.3 | 19.39% | motif file (matrix) | svg |
| 365 | A C T G C G T A A C T G A T G C T G A C G A T C A T C G T G C A A C T G A G T C | ZNF519(Zf)/HEK293-ZNF519.GFP-ChIP-Seq(GSE58341)/Homer | 1e-86 | -1.993e+02 | 0.0000 | 3608.0 | 11.18% | 11217.4 | 7.69% | motif file (matrix) | svg |
| 366 | T A G C T A G C G A C T C T A G A G C T A G T C G T C A T G C A A C G T A T G C G C T A T G C A | Pbx3(Homeobox)/GM12878-PBX3-ChIP-Seq(GSE32465)/Homer | 1e-85 | -1.972e+02 | 0.0000 | 3892.0 | 12.06% | 12328.4 | 8.46% | motif file (matrix) | svg |
| 367 | A T G C T C A G T C G A G C A T A C T G C G T A A G T C T C A G G A C T T G A C C G T A A G C T | Atf2(bZIP)/3T3L1-Atf2-ChIP-Seq(GSE56872)/Homer | 1e-83 | -1.930e+02 | 0.0000 | 5785.0 | 17.92% | 19851.9 | 13.62% | motif file (matrix) | svg |
| 368 | T A G C G T A C C T A G C A G T T C G A C G T A C G T A G C A T G A C T T G A C A G T C A C T G A T C G A G T C C T A G | AS2(LOBAS2)/col-AS2-DAP-Seq(GSE60143)/Homer | 1e-82 | -1.903e+02 | 0.0000 | 4472.0 | 13.85% | 14670.3 | 10.06% | motif file (matrix) | svg |
| 369 | C T A G T A C G G A T C G T A C G C T A A G C T A G C T G T C A T C G A T A G C | Nanog(Homeobox)/mES-Nanog-ChIP-Seq(GSE11724)/Homer | 1e-81 | -1.873e+02 | 0.0000 | 28677.0 | 88.84% | 123618.7 | 84.80% | motif file (matrix) | svg |
| 370 | A T G C T C G A A G T C A G C T A C G T G T A C A G T C G C T A C T A G C A T G G T C A C T G A T C A G A G T C | Stat3+il21(Stat)/CD4-Stat3-ChIP-Seq(GSE19198)/Homer | 1e-80 | -1.861e+02 | 0.0000 | 8422.0 | 26.09% | 30821.3 | 21.14% | motif file (matrix) | svg |
| 371 | T G C A C T G A A T G C G T C A A C G T A T G C A C G T A C T G A C T G T G C A | ZBTB18(Zf)/HEK293-ZBTB18.GFP-ChIP-Seq(GSE58341)/Homer | 1e-80 | -1.855e+02 | 0.0000 | 6003.0 | 18.60% | 20850.8 | 14.30% | motif file (matrix) | svg |
| 372 | G T A C A C G T A C G T T C A G G C T A C G T A C G A T G C A T G C A T A G T C C G T A G T C A C A T G G A C T G C T A | ANAC070(NAC)/colamp-ANAC070-DAP-Seq(GSE60143)/Homer | 1e-80 | -1.842e+02 | 0.0000 | 16965.0 | 52.55% | 68097.5 | 46.71% | motif file (matrix) | svg |
| 373 | T A G C G C T A T C G A C T G A A G T C A G T C C T G A A G T C C G T A C T A G | RUNX(Runt)/HPC7-Runx1-ChIP-Seq(GSE22178)/Homer | 1e-79 | -1.838e+02 | 0.0000 | 11682.0 | 36.19% | 44765.1 | 30.71% | motif file (matrix) | svg |
| 374 | A G C T G C A T G T C A C G A T T A G C C G T A A C G T G C T A | CRC(C2C2YABBY)/col-CRC-DAP-Seq(GSE60143)/Homer | 1e-79 | -1.826e+02 | 0.0000 | 17214.0 | 53.33% | 69258.5 | 47.51% | motif file (matrix) | svg |
| 375 | A G T C C T G A A G T C C G A T C A G T G A T C A T G C A C T G A T C G G A C T | Fli1(ETS)/CD8-FLI-ChIP-Seq(GSE20898)/Homer | 1e-78 | -1.796e+02 | 0.0000 | 18969.0 | 58.76% | 77301.5 | 53.03% | motif file (matrix) | svg |
| 376 | C G A T C T G A A G T C A C G T A C G T T C A G G C A T C G A T G C T A G C T A C G T A A G T C C G T A G T C A A C T G | CUC1(NAC)/col-CUC1-DAP-Seq(GSE60143)/Homer | 1e-77 | -1.784e+02 | 0.0000 | 9795.0 | 30.34% | 36776.5 | 25.23% | motif file (matrix) | svg |
| 377 | C T A G T C G A C G A T C T A G G C A T C A G T C T A G G A T C C G T A G T C A | CEBP:AP1(bZIP)/ThioMac-CEBPb-ChIP-Seq(GSE21512)/Homer | 1e-77 | -1.780e+02 | 0.0000 | 11301.0 | 35.01% | 43243.1 | 29.66% | motif file (matrix) | svg |
| 378 | G C A T G A T C T C A G G C T A G A C T A G T C C T A G C G T A C A T G G T C A | GATA20(C2C2gata)/colamp-GATA20-DAP-Seq(GSE60143)/Homer | 1e-76 | -1.772e+02 | 0.0000 | 26925.0 | 83.41% | 115000.0 | 78.89% | motif file (matrix) | svg |
| 379 | G A C T C T G A G T A C A G T C C G A T C G T A G T C A G A T C G C A T G C A T G C A T C G A T | AT3G10580(MYBrelated)/colamp-AT3G10580-DAP-Seq(GSE60143)/Homer | 1e-76 | -1.767e+02 | 0.0000 | 9294.0 | 28.79% | 34682.9 | 23.79% | motif file (matrix) | svg |
| 380 | T C G A G A C T A T C G C G T A A G T C C T A G G C A T G T A C C T G A A C G T G A T C G C T A | TGA4(bZIP)/colamp-TGA4-DAP-Seq(GSE60143)/Homer | 1e-76 | -1.757e+02 | 0.0000 | 7386.0 | 22.88% | 26695.4 | 18.31% | motif file (matrix) | svg |
| 381 | T C A G G C A T A C T G C G T A A G T C C T A G G C A T T G A C | TGA9(bZIP)/colamp-TGA9-DAP-Seq(GSE60143)/Homer | 1e-76 | -1.755e+02 | 0.0000 | 20960.0 | 64.93% | 86570.3 | 59.39% | motif file (matrix) | svg |
| 382 | C A G T A T C G C T G A A G T C T C A G C A G T T A G C C T G A A T G C T A C G | FEA4(bZIP)/Corn-FEA4-ChIP-Seq(GSE61954)/Homer | 1e-76 | -1.751e+02 | 0.0000 | 18751.0 | 58.09% | 76413.5 | 52.42% | motif file (matrix) | svg |
| 383 | G C A T T C A G C T G A A T C G A C T G C G A T G A T C C T G A | THRb(NR)/Liver-NR1A2-ChIP-Seq(GSE52613)/Homer | 1e-75 | -1.742e+02 | 0.0000 | 26413.0 | 81.82% | 112540.4 | 77.20% | motif file (matrix) | svg |
| 384 | C A T G C T A G A G C T G A C T C A T G A G T C G A T C G C T A C G A T C T A G T C A G G T A C C T G A T C G A | X-box(HTH)/NPC-H3K4me1-ChIP-Seq(GSE16256)/Homer | 1e-74 | -1.706e+02 | 0.0000 | 1591.0 | 4.93% | 4130.3 | 2.83% | motif file (matrix) | svg |
| 385 | T A C G T A G C C A T G C A G T A C G T C T A G C G T A A G T C G A C T G C A T G C A T C A G T | WRKY11(WRKY)/col-WRKY11-DAP-Seq(GSE60143)/Homer | 1e-73 | -1.698e+02 | 0.0000 | 3127.0 | 9.69% | 9735.6 | 6.68% | motif file (matrix) | svg |
| 386 | T A G C C A T G A G C T A C G T A C T G C G T A A G T C G A C T G C A T C T G A | AT3G42860(zfGRF)/col-AT3G42860-DAP-Seq(GSE60143)/Homer | 1e-73 | -1.693e+02 | 0.0000 | 6956.0 | 21.55% | 25030.8 | 17.17% | motif file (matrix) | svg |
| 387 | G T C A T C G A C T A G C T A G A G T C G T C A C G A T C T A G G A C T G A T C G A T C T C A G C T A G C T G A A G T C G C T A C A G T T C A G G A T C G A T C | p63(p53)/Keratinocyte-p63-ChIP-Seq(GSE17611)/Homer | 1e-73 | -1.689e+02 | 0.0000 | 6363.0 | 19.71% | 22596.8 | 15.50% | motif file (matrix) | svg |
| 388 | C A T G G A T C C T G A G T A C C T A G C T G A G C T A G C A T G A T C G A T C A G T C C T A G C G T A C A T G C T A G | AIL7(AP2EREBP)/colamp-AIL7-DAP-Seq(GSE60143)/Homer | 1e-73 | -1.688e+02 | 0.0000 | 11207.0 | 34.72% | 43039.3 | 29.52% | motif file (matrix) | svg |
| 389 | T G A C C A T G A C G T A C G T A C T G C G T A A G T C A G C T G C A T T C G A | WRKY30(WRKY)/colamp-WRKY30-DAP-Seq(GSE60143)/Homer | 1e-72 | -1.671e+02 | 0.0000 | 7725.0 | 23.93% | 28267.7 | 19.39% | motif file (matrix) | svg |
| 390 | G A C T C T A G C T A G A G T C T G C A A C T G A C G T A C G T C T A G T C A G | AMYB(HTH)/Testes-AMYB-ChIP-Seq(GSE44588)/Homer | 1e-72 | -1.671e+02 | 0.0000 | 23964.0 | 74.24% | 100880.8 | 69.20% | motif file (matrix) | svg |
| 391 | A G T C G A T C G A T C C G T A G T C A A G T C A G C T C T G A G A C T G A C T | ATY13(MYB)/col-ATY13-DAP-Seq(GSE60143)/Homer | 1e-71 | -1.641e+02 | 0.0000 | 27716.0 | 85.86% | 119186.1 | 81.76% | motif file (matrix) | svg |
| 392 | C A G T T A G C A G T C C A T G C A G T C A T G C G A T C G A T G A C T C G A T A T C G G T A C A C T G A T C G G T A C | LBD13(LOBAS2)/colamp-LBD13-DAP-Seq(GSE60143)/Homer | 1e-71 | -1.639e+02 | 0.0000 | 20355.0 | 63.06% | 84052.0 | 57.66% | motif file (matrix) | svg |
| 393 | G C A T C T A G G T A C A G T C C G A T A C T G C T A G C T A G G T A C G C T A | ZNF416(Zf)/HEK293-ZNF416.GFP-ChIP-Seq(GSE58341)/Homer | 1e-70 | -1.621e+02 | 0.0000 | 12069.0 | 37.39% | 46932.5 | 32.20% | motif file (matrix) | svg |
| 394 | T C G A T A G C G T C A A C T G C T A G C G T A C G A T A C T G A C G T A C T G A C T G A C G T | ETS:RUNX(ETS,Runt)/Jurkat-RUNX1-ChIP-Seq(GSE17954)/Homer | 1e-70 | -1.619e+02 | 0.0000 | 1553.0 | 4.81% | 4067.2 | 2.79% | motif file (matrix) | svg |
| 395 | C A G T T C G A A G T C A C G T A C G T T C A G C G A T G C T A G C T A C G T A G C A T A G T C C G T A T G C A A C T G | ANAC045(NAC)/col-ANAC045-DAP-Seq(GSE60143)/Homer | 1e-69 | -1.603e+02 | 0.0000 | 23799.0 | 73.72% | 100254.9 | 68.77% | motif file (matrix) | svg |
| 396 | G T A C C A T G T A G C A G T C C T A G C A T G C T G A C G T A G C A T G C A T A C G T G C A T G T A C A C T G A T C G | LOB(LOBAS2)/col-LOB-DAP-Seq(GSE60143)/Homer | 1e-69 | -1.595e+02 | 0.0000 | 10893.0 | 33.74% | 41893.6 | 28.74% | motif file (matrix) | svg |
| 397 | C T G A A T C G A G C T A G C T A C G T T A G C C T G A T A C G C G A T A C G T G A C T A G T C | ISRE(IRF)/ThioMac-LPS-Expression(GSE23622)/Homer | 1e-69 | -1.594e+02 | 0.0000 | 726.0 | 2.25% | 1405.5 | 0.96% | motif file (matrix) | svg |
| 398 | C G A T C T A G T C A G C A G T C G T A A G T C G C T A A C G T G A C T A T G C A G T C G C T A | PRDM10(Zf)/HEK293-PRDM10.eGFP-ChIP-Seq(Encode)/Homer | 1e-68 | -1.574e+02 | 0.0000 | 8362.0 | 25.90% | 31134.1 | 21.36% | motif file (matrix) | svg |
| 399 | G T A C A C G T A C G T T C A G C G T A C G T A C G A T G C A T G C A T A G T C C G T A G T C A C A T G G A C T G C T A | VND1(NAC)/col-VND1-DAP-Seq(GSE60143)/Homer | 1e-67 | -1.555e+02 | 0.0000 | 13348.0 | 41.35% | 52687.1 | 36.14% | motif file (matrix) | svg |
| 400 | C T A G T C A G C A G T T C A G A C T G A C T G G A T C C T A G A C T G C T A G T C A G A T G C | KLF14(Zf)/HEK293-KLF14.GFP-ChIP-Seq(GSE58341)/Homer | 1e-67 | -1.548e+02 | 0.0000 | 17827.0 | 55.22% | 72717.8 | 49.88% | motif file (matrix) | svg |
| 401 | G T A C C T G A A G T C A G T C A C T G G T C A G A T C G C A T | At1g75490(AP2EREBP)/colamp-At1g75490-DAP-Seq(GSE60143)/Homer | 1e-66 | -1.542e+02 | 0.0000 | 27465.0 | 85.08% | 118137.7 | 81.04% | motif file (matrix) | svg |
| 402 | A T G C C T G A A T C G T A C G A G T C C G A T T C A G C G A T C T A G A G C T G T C A G T C A C G T A A G T C C G T A T A C G C T G A | Fox:Ebox(Forkhead,bHLH)/Panc1-Foxa2-ChIP-Seq(GSE47459)/Homer | 1e-66 | -1.537e+02 | 0.0000 | 9863.0 | 30.55% | 37594.3 | 25.79% | motif file (matrix) | svg |
| 403 | G A C T C A G T A G C T C G A T A G T C G A T C A G T C C G T A A T G C T C A G | Rbpj1(?)/Panc1-Rbpj1-ChIP-Seq(GSE47459)/Homer | 1e-66 | -1.533e+02 | 0.0000 | 14236.0 | 44.10% | 56661.1 | 38.87% | motif file (matrix) | svg |
| 404 | T C G A G C A T A C G T C T A G G T A C T C G A G C A T T G A C T C G A A C G T | Chop(bZIP)/MEF-Chop-ChIP-Seq(GSE35681)/Homer | 1e-66 | -1.524e+02 | 0.0000 | 4285.0 | 13.27% | 14473.4 | 9.93% | motif file (matrix) | svg |
| 405 | G C T A T G A C G A T C C G A T G A C T A T G C C T G A A T C G G C A T A C G T | JGL(C2H2)/col-JGL-DAP-Seq(GSE60143)/Homer | 1e-65 | -1.514e+02 | 0.0000 | 17891.0 | 55.42% | 73095.1 | 50.14% | motif file (matrix) | svg |
| 406 | C G A T C A G T C T A G G C T A A G T C C G T A T C A G A G T C A C G T A C T G A C G T G T A C G C T A G C T A G C T A | bZIP52(bZIP)/colamp-bZIP52-DAP-Seq(GSE60143)/Homer | 1e-64 | -1.490e+02 | 0.0000 | 14823.0 | 45.92% | 59378.0 | 40.73% | motif file (matrix) | svg |
| 407 | T A G C G T A C C T A G A T C G C T G A C G T A G C T A G C A T A C G T T G A C G T A C A C T G T A C G G T C A C T A G | ASL18(LOBAS2)/colamp-ASL18-DAP-Seq(GSE60143)/Homer | 1e-63 | -1.471e+02 | 0.0000 | 21762.0 | 67.41% | 91003.5 | 62.43% | motif file (matrix) | svg |
| 408 | T A G C C T A G T C G A G A C T A C T G C T G A A G T C T C A G G C A T T G A C C T G A A G C T | Atf7(bZIP)/3T3L1-Atf7-ChIP-Seq(GSE56872)/Homer | 1e-63 | -1.461e+02 | 0.0000 | 8724.0 | 27.03% | 32905.2 | 22.57% | motif file (matrix) | svg |
| 409 | T A C G C T G A T C G A C G A T C T A G C T A G T C G A C T G A T C G A T C G A C G T A T C G A G C A T C A T G C G T A T A C G G C A T T G A C C G T A A G C T | NFAT:AP1(RHD,bZIP)/Jurkat-NFATC1-ChIP-Seq(Jolma\_et\_al.)/Homer | 1e-62 | -1.444e+02 | 0.0000 | 1689.0 | 5.23% | 4690.2 | 3.22% | motif file (matrix) | svg |
| 410 | C G T A A C T G C G T A A C G T C A G T A G T C A G C T G C A T G C T A C G A T | At2g01060(G2like)/colamp-At2g01060-DAP-Seq(GSE60143)/Homer | 1e-62 | -1.431e+02 | 0.0000 | 26347.0 | 81.62% | 112891.7 | 77.44% | motif file (matrix) | svg |
| 411 | T C G A C G T A C G T A T C G A A C T G G T A C C G T A A G C T G T C A G C A T | At3g24120(G2like)/col-At3g24120-DAP-Seq(GSE60143)/Homer | 1e-59 | -1.366e+02 | 0.0000 | 26612.0 | 82.44% | 114329.0 | 78.43% | motif file (matrix) | svg |
| 412 | C T A G A C T G C T A G T C A G T C A G T A C G C T A G A C T G | Maz(Zf)/HepG2-Maz-ChIP-Seq(GSE31477)/Homer | 1e-58 | -1.356e+02 | 0.0000 | 12994.0 | 40.25% | 51635.2 | 35.42% | motif file (matrix) | svg |
| 413 | C T A G G T A C A C G T A C G T A T C G G C A T A G C T A G C T A G C T G C A T G A C T C G T A G T C A A C T G G A C T | VND6(NAC)/col-VND6-DAP-Seq(GSE60143)/Homer | 1e-58 | -1.355e+02 | 0.0000 | 18380.0 | 56.94% | 75742.5 | 51.96% | motif file (matrix) | svg |
| 414 | T C G A T C G A T A G C G T A C T C A G T A C G C G T A C G T A T C A G A G C T | GABPA(ETS)/Jurkat-GABPa-ChIP-Seq(GSE17954)/Homer | 1e-58 | -1.348e+02 | 0.0000 | 13982.0 | 43.31% | 56017.3 | 38.43% | motif file (matrix) | svg |
| 415 | C G T A C T A G G A C T G T C A G T C A C G T A A G T C C G T A T C G A T C G A T C G A C G T A C T G A C T A G G C T A C G T A T A G C C G T A C G A T C G T A | FOXA1:AR(Forkhead,NR)/LNCAP-AR-ChIP-Seq(GSE27824)/Homer | 1e-57 | -1.319e+02 | 0.0000 | 485.0 | 1.50% | 835.0 | 0.57% | motif file (matrix) | svg |
| 416 | C G A T C T A G T C G A A G C T C G A T C T G A C G T A A G C T A C T G C T A G A T G C G A T C | Hoxb4(Homeobox)/ES-Hoxb4-ChIP-Seq(GSE34014)/Homer | 1e-55 | -1.288e+02 | 0.0000 | 3936.0 | 12.19% | 13450.3 | 9.23% | motif file (matrix) | svg |
| 417 | C T A G T A C G G A T C G T C A T G C A A C G T T G C A G C T A T C G A T G C A | Hoxa9(Homeobox)/ChickenMSG-Hoxa9.Flag-ChIP-Seq(GSE86088)/Homer | 1e-55 | -1.280e+02 | 0.0000 | 23718.0 | 73.47% | 100676.7 | 69.06% | motif file (matrix) | svg |
| 418 | C G A T C T G A A G T C A C G T A C G T T C A G C G T A C G T A C G T A G C A T C G A T A G T C C G T A G T C A A C T G | VND4(NAC)/colamp-VND4-DAP-Seq(GSE60143)/Homer | 1e-55 | -1.275e+02 | 0.0000 | 13343.0 | 41.33% | 53389.3 | 36.62% | motif file (matrix) | svg |
| 419 | T C A G T A C G T A G C A C G T A C T G C G A T A G T C C G T A T A C G A G T C | Meis1(Homeobox)/MastCells-Meis1-ChIP-Seq(GSE48085)/Homer | 1e-54 | -1.266e+02 | 0.0000 | 19781.0 | 61.28% | 82395.7 | 56.52% | motif file (matrix) | svg |
| 420 | T G C A C G T A A C T G T C A G C A G T C A T G T C A G G A T C T A C G A G T C T G C A A C T G A C T G T G A C G T C A | ZNF165(Zf)/WHIM12-ZNF165-ChIP-Seq(GSE65937)/Homer | 1e-54 | -1.261e+02 | 0.0000 | 1718.0 | 5.32% | 4967.9 | 3.41% | motif file (matrix) | svg |
| 421 | C G A T T C G A G T A C A C G T A C G T T C A G G C A T G C T A T G C A G C A T C G T A A G T C C G T A T G A C C A T G | ANAC092(NAC)/colamp-ANAC092-DAP-Seq(GSE60143)/Homer | 1e-54 | -1.261e+02 | 0.0000 | 9786.0 | 30.32% | 37913.6 | 26.01% | motif file (matrix) | svg |
| 422 | T A G C G T A C C G T A C T A G A C T G T G C A C G T A A T G C C G T A A T C G | AR-halfsite(NR)/LNCaP-AR-ChIP-Seq(GSE27824)/Homer | 1e-53 | -1.233e+02 | 0.0000 | 25503.0 | 79.00% | 109291.0 | 74.97% | motif file (matrix) | svg |
| 423 | A C T G A G C T A G T C G T C A A G C T T C A G A T G C G A T C G C A T A T C G T C G A T A G C C G A T C A T G T A G C | Pax8(Paired,Homeobox)/Thyroid-Pax8-ChIP-Seq(GSE26938)/Homer | 1e-53 | -1.224e+02 | 0.0000 | 4620.0 | 14.31% | 16310.0 | 11.19% | motif file (matrix) | svg |
| 424 | A G T C G A C T A C T G G A T C G T A C C G T A T G A C A G T C C G A T A G C T A C G T A C G T C T A G G A C T C T G A | ZNF7(Zf)/HepG2-ZNF7.Flag-ChIP-Seq(Encode)/Homer | 1e-52 | -1.207e+02 | 0.0000 | 7200.0 | 22.30% | 27025.8 | 18.54% | motif file (matrix) | svg |
| 425 | C G A T C T A G G A T C G C T A A G C T C T A G G A T C C G T A | RBFox2(?)/Heart-RBFox2-CLIP-Seq(GSE57926)/Homer | 1e-52 | -1.206e+02 | 0.0000 | 22629.0 | 70.10% | 95760.3 | 65.69% | motif file (matrix) | svg |
| 426 | G C T A A C T G T C G A C T G A C T G A A C G T T A G C C T G A C G T A C G A T | Cux2(Homeobox)/Liver-Cux2-ChIP-Seq(GSE35985)/Homer | 1e-52 | -1.201e+02 | 0.0000 | 12485.0 | 38.68% | 49813.5 | 34.17% | motif file (matrix) | svg |
| 427 | A G T C A T C G C T A G A G C T G A C T C T A G A G T C A G T C G C T A C A G T T C A G T C A G G A T C C T G A T C G A G A T C | RFX(HTH)/K562-RFX3-ChIP-Seq(SRA012198)/Homer | 1e-50 | -1.169e+02 | 0.0000 | 1419.0 | 4.40% | 3978.9 | 2.73% | motif file (matrix) | svg |
| 428 | T C A G T C A G A C G T G T A C G C T A T C A G C T G A A C T G A C T G A G C T A G T C C G T A | EAR2(NR)/K562-NR2F6-ChIP-Seq(Encode)/Homer | 1e-49 | -1.149e+02 | 0.0000 | 16078.0 | 49.81% | 65927.5 | 45.23% | motif file (matrix) | svg |
| 429 | C G A T C T G A G T A C A C G T A C G T T C A G C G T A C G T A G C T A G C A T G C A T A G T C C G T A G T C A C A T G | NST1(NAC)/colamp-NST1-DAP-Seq(GSE60143)/Homer | 1e-49 | -1.149e+02 | 0.0000 | 13112.0 | 40.62% | 52720.5 | 36.17% | motif file (matrix) | svg |
| 430 | A G C T A G T C A G T C A C G T C T A G A C G T A C G T A C G T C G T A A G T C G A T C C G T A | FOXP1(Forkhead)/H9-FOXP1-ChIP-Seq(GSE31006)/Homer | 1e-49 | -1.136e+02 | 0.0000 | 5752.0 | 17.82% | 21126.0 | 14.49% | motif file (matrix) | svg |
| 431 | T C G A G C A T A C T G C T G A A G T C T C A G G A C T G T A C C G T A A G C T A G T C G A T C | c-Jun-CRE(bZIP)/K562-cJun-ChIP-Seq(GSE31477)/Homer | 1e-48 | -1.124e+02 | 0.0000 | 4211.0 | 13.04% | 14832.6 | 10.17% | motif file (matrix) | svg |
| 432 | G A C T C A G T G A T C G A T C A C G T G A T C C T G A T A C G C G T A G T C A | STAT6(Stat)/Macrophage-Stat6-ChIP-Seq(GSE38377)/Homer | 1e-48 | -1.118e+02 | 0.0000 | 5713.0 | 17.70% | 21002.9 | 14.41% | motif file (matrix) | svg |
| 433 | C T G A A T G C C G T A A C G T A G T C A G T C A C G T A C T G A T C G G C A T | SPDEF(ETS)/VCaP-SPDEF-ChIP-Seq(SRA014231)/Homer | 1e-48 | -1.114e+02 | 0.0000 | 13752.0 | 42.60% | 55653.5 | 38.18% | motif file (matrix) | svg |
| 434 | T G C A C T G A A T G C G T C A A C T G A C T G C G T A C G T A C T A G A G C T | Ets1-distal(ETS)/CD4+-PolII-ChIP-Seq(Barski\_et\_al.)/Homer | 1e-47 | -1.102e+02 | 0.0000 | 3128.0 | 9.69% | 10538.4 | 7.23% | motif file (matrix) | svg |
| 435 | T G C A G C A T A G C T G C A T A G T C A G T C A G T C C T G A A C T G T C G A T C G A C A G T A T C G A G T C G A T C | ZNF143\|STAF(Zf)/CUTLL-ZNF143-ChIP-Seq(GSE29600)/Homer | 1e-47 | -1.086e+02 | 0.0000 | 3305.0 | 10.24% | 11264.6 | 7.73% | motif file (matrix) | svg |
| 436 | A T G C G A C T A C T G C A G T G A T C A C G T T A C G T A C G | Smad2(MAD)/ES-SMAD2-ChIP-Seq(GSE29422)/Homer | 1e-46 | -1.076e+02 | 0.0000 | 20552.0 | 63.67% | 86500.7 | 59.34% | motif file (matrix) | svg |
| 437 | A G T C C A G T T C A G A T G C A G T C C G A T C G T A G T C A G A T C G C A T | BOS1(MYB)/col-BOS1-DAP-Seq(GSE60143)/Homer | 1e-45 | -1.049e+02 | 0.0000 | 17059.0 | 52.85% | 70652.8 | 48.47% | motif file (matrix) | svg |
| 438 | C T A G C T A G A G T C T C A G A C T G A C G T A C G T C T G A | MYB(HTH)/ERMYB-Myb-ChIPSeq(GSE22095)/Homer | 1e-45 | -1.048e+02 | 0.0000 | 26021.0 | 80.61% | 112258.2 | 77.01% | motif file (matrix) | svg |
| 439 | T A C G C G T A T C A G G A C T C T A G A C T G C A G T T A G C T C G A A C G T G T A C C T A G A G T C A G T C G A T C | ZNF669(Zf)/HEK293-ZNF669.GFP-ChIP-Seq(GSE58341)/Homer | 1e-45 | -1.041e+02 | 0.0000 | 3104.0 | 9.62% | 10538.7 | 7.23% | motif file (matrix) | svg |
| 440 | C A T G A C G T C T A G A C G T C A G T A C G T C T A G A G T C | PHA-4(Forkhead)/cElegans-Embryos-PHA4-ChIP-Seq(modEncode)/Homer | 1e-43 | -9.999e+01 | 0.0000 | 23069.0 | 71.46% | 98418.3 | 67.51% | motif file (matrix) | svg |
| 441 | C G T A T C G A G A T C G C A T C G T A A C G T G T A C T C A G G T C A G A C T C G T A C T A G | DREF/Drosophila-Promoters/Homer | 1e-43 | -9.909e+01 | 0.0000 | 1609.0 | 4.98% | 4850.2 | 3.33% | motif file (matrix) | svg |
| 442 | C G A T G A C T C G A T T C A G G A C T A C G T C A G T C T G A G A C T G A C T A G C T C G A T A C T G A T C G G T A C G C T A | NF1:FOXA1(CTF,Forkhead)/LNCAP-FOXA1-ChIP-Seq(GSE27824)/Homer | 1e-42 | -9.875e+01 | 0.0000 | 672.0 | 2.08% | 1560.6 | 1.07% | motif file (matrix) | svg |
| 443 | T C G A A C T G A C T G C G T A C G T A T C G A A G T C C T G A A T C G G T A C G C A T C A T G | ETS:E-box(ETS,bHLH)/HPC7-Scl-ChIP-Seq(GSE22178)/Homer | 1e-42 | -9.691e+01 | 0.0000 | 941.0 | 2.92% | 2472.9 | 1.70% | motif file (matrix) | svg |
| 444 | C T G A A G C T A C G T A C G T A G T C G A C T G A C T C T G A C T G A C T A G C G T A C G T A | STAT6(Stat)/CD4-Stat6-ChIP-Seq(GSE22104)/Homer | 1e-41 | -9.589e+01 | 0.0000 | 5362.0 | 16.61% | 19886.1 | 13.64% | motif file (matrix) | svg |
| 445 | C G T A T G A C T C G A A G T C C G T A A T C G A T G C A C G T A C T G A G T C | E2A(bHLH)/proBcell-E2A-ChIP-Seq(GSE21978)/Homer | 1e-41 | -9.517e+01 | 0.0000 | 14016.0 | 43.42% | 57323.3 | 39.32% | motif file (matrix) | svg |
| 446 | G A C T C T A G G A T C C A G T A C T G C T G A A T G C G C A T A T G C C T G A | MafA(bZIP)/Islet-MafA-ChIP-Seq(GSE30298)/Homer | 1e-40 | -9.421e+01 | 0.0000 | 10862.0 | 33.65% | 43449.2 | 29.81% | motif file (matrix) | svg |
| 447 | T C A G A C T G C A G T A G T C A G T C G T C A C G T A C G T A A C T G C A G T A G T C A G T C C T G A T G C A A G C T | dHNF4(NR)/Fly-HNF4-ChIP-Seq(GSE73675)/Homer | 1e-40 | -9.287e+01 | 0.0000 | 922.0 | 2.86% | 2440.0 | 1.67% | motif file (matrix) | svg |
| 448 | C T A G C T A G T C A G G T C A C T A G T C A G G C T A A G T C A T C G A G C T C T A G | DPR(core promoter) | 1e-40 | -9.223e+01 | 0.0000 | 30646.0 | 94.94% | 135500.9 | 92.95% | motif file (matrix) | svg |
| 449 | G A C T A T C G C T G A A G T C T C A G G A C T G T A C C T G A A G C T G T A C | TGA6(bZIP)/colamp-TGA6-DAP-Seq(GSE60143)/Homer | 1e-39 | -9.141e+01 | 0.0000 | 14792.0 | 45.82% | 60904.1 | 41.78% | motif file (matrix) | svg |
| 450 | A G T C G A C T C A G T A C T G C T A G T G A C G C T A A T G C G C A T A T C G C G A T A C T G G A T C G T A C G T C A C T G A | NF1(CTF)/LNCAP-NF1-ChIP-Seq(Unpublished)/Homer | 1e-39 | -9.001e+01 | 0.0000 | 4559.0 | 14.12% | 16688.0 | 11.45% | motif file (matrix) | svg |
| 451 | G C A T G C A T G A T C G C A T T C G A A C T G C G T A C G T A A T C G T A G C C G A T A C G T A G T C A G C T C G T A | HSF6(HSF)/col-HSF6-DAP-Seq(GSE60143)/Homer | 1e-38 | -8.945e+01 | 0.0000 | 2639.0 | 8.18% | 8934.9 | 6.13% | motif file (matrix) | svg |
| 452 | T A G C G A T C A G C T T G A C G C T A A G C T C A T G A C T G A C G T T C A G A G T C G A T C G A C T A G C T G C T A A G T C A G C T A G T C G A T C A T G C A G C T G A C T C A T G A C G T A T C G | ZNF41(Zf)/HEK293-ZNF41.GFP-ChIP-Seq(GSE58341)/Homer | 1e-38 | -8.895e+01 | 0.0000 | 421.0 | 1.30% | 834.2 | 0.57% | motif file (matrix) | svg |
| 453 | T G C A G C A T C G A T C G T A C A G T A C T G G T A C C G T A C T G A A G C T G T C A A C T G C T A G G T C A C G A T A C T G G T A C T G C A C G T A A G C T | CEBP:CEBP(bZIP)/MEF-Chop-ChIP-Seq(GSE35681)/Homer | 1e-38 | -8.791e+01 | 0.0000 | 1831.0 | 5.67% | 5819.4 | 3.99% | motif file (matrix) | svg |
| 454 | T C G A G C T A T G A C G C T A C T A G G A T C C G A T A C T G C G A T A G C T G A C T C T A G | E-box/Drosophila-Promoters/Homer | 1e-37 | -8.632e+01 | 0.0000 | 2999.0 | 9.29% | 10416.9 | 7.15% | motif file (matrix) | svg |
| 455 | T A C G A T C G T A G C G A T C A C T G A C G T A G T C A C G T C T A G A T C G | Smad4(MAD)/ESC-SMAD4-ChIP-Seq(GSE29422)/Homer | 1e-37 | -8.544e+01 | 0.0000 | 20618.0 | 63.87% | 87520.3 | 60.04% | motif file (matrix) | svg |
| 456 | A T G C A G T C G T A C A G C T T C G A C T A G G A T C C T G A G T C A A G T C G C T A T C A G | Rfx5(HTH)/GM12878-Rfx5-ChIP-Seq(GSE31477)/Homer | 1e-36 | -8.304e+01 | 0.0000 | 5425.0 | 16.81% | 20446.7 | 14.03% | motif file (matrix) | svg |
| 457 | C T A G A G C T G A C T C A T G A G T C A G T C G T C A C A G T C T A G T C A G G T A C C T G A T C G A G A T C T G A C | Rfx2(HTH)/LoVo-RFX2-ChIP-Seq(GSE49402)/Homer | 1e-35 | -8.240e+01 | 0.0000 | 1457.0 | 4.51% | 4475.4 | 3.07% | motif file (matrix) | svg |
| 458 | C T A G C A G T C G T A A C G T A G T C A C T G C G T A A G C T A G T C G A T C | HNF6(Homeobox)/Liver-Hnf6-ChIP-Seq(ERP000394)/Homer | 1e-35 | -8.215e+01 | 0.0000 | 15398.0 | 47.70% | 63932.4 | 43.86% | motif file (matrix) | svg |
| 459 | C T A G T A C G A G T C C G T A A G T C A C G T A G T C T C G A C G T A T A C G | Nkx2.1(Homeobox)/LungAC-Nkx2.1-ChIP-Seq(GSE43252)/Homer | 1e-35 | -8.142e+01 | 0.0000 | 24869.0 | 77.04% | 107455.8 | 73.71% | motif file (matrix) | svg |
| 460 | G C A T C T A G C T A G C G T A A G C T C G T A C T G A C A T G C T A G G C A T | AT5G56840(MYBrelated)/colamp-AT5G56840-DAP-Seq(GSE60143)/Homer | 1e-35 | -8.068e+01 | 0.0000 | 17187.0 | 53.24% | 72040.4 | 49.42% | motif file (matrix) | svg |
| 461 | T G A C G C T A T G A C C G T A T C A G G A T C C G T A C A T G C A T G C T A G C T A G C T A G | Unknown-ESC-element(?)/mES-Nanog-ChIP-Seq(GSE11724)/Homer | 1e-34 | -7.993e+01 | 0.0000 | 5476.0 | 16.96% | 20736.8 | 14.23% | motif file (matrix) | svg |
| 462 | C G T A C G T A G C A T A C T G C G T A A G C T C T G A C G T A T A C G C T G A | ELT-3(Gata)/cElegans-L1-ELT3-ChIP-Seq(modEncode)/Homer | 1e-34 | -7.859e+01 | 0.0000 | 6546.0 | 20.28% | 25291.6 | 17.35% | motif file (matrix) | svg |
| 463 | T A C G T A C G C T A G T C A G A G T C C G T A A T C G A T G C A C G T A C T G A G T C G A C T | Ascl2(bHLH)/ESC-Ascl2-ChIP-Seq(GSE97712)/Homer | 1e-32 | -7.576e+01 | 0.0000 | 12036.0 | 37.29% | 49201.3 | 33.75% | motif file (matrix) | svg |
| 464 | C T G A C T A G A T C G A G C T A C T G G A C T A G T C C T G A | Tbx5(T-box)/HL1-Tbx5.biotin-ChIP-Seq(GSE21529)/Homer | 1e-32 | -7.564e+01 | 0.0000 | 25252.0 | 78.23% | 109450.6 | 75.08% | motif file (matrix) | svg |
| 465 | G C T A G C A T G C T A G C A T G C A T C G T A C G T A A G T C A G T C A C T G G C A T G C A T C G T A G C T A G C T A | MYB73(MYB)/col-MYB73-DAP-Seq(GSE60143)/Homer | 1e-32 | -7.503e+01 | 0.0000 | 22863.0 | 70.82% | 98269.0 | 67.41% | motif file (matrix) | svg |
| 466 | G A T C G T A C C G T A A G C T G A C T G C T A C T G A A C G T G A T C G C T A | Hoxc6(Homeobox)/EB-Hoxc6.iFlag-ChIP-Seq(GSE142377)/Homer | 1e-32 | -7.482e+01 | 0.0000 | 23581.0 | 73.05% | 101624.7 | 69.71% | motif file (matrix) | svg |
| 467 | C G T A C G T A C T G A A C T G A C G T A G T C C G T A C G T A A G T C A C T G A T G C G A T C | WRKY46(WRKY)/colamp-WRKY46-DAP-Seq(GSE60143)/Homer | 1e-32 | -7.442e+01 | 0.0000 | 2502.0 | 7.75% | 8639.7 | 5.93% | motif file (matrix) | svg |
| 468 | G C A T G C A T C T G A A C G T C T G A A C G T C G T A C G T A C G T A A G T C G T C A G T C A | Foxf1(Forkhead)/Lung-Foxf1-ChIP-Seq(GSE77951)/Homer | 1e-32 | -7.408e+01 | 0.0000 | 9069.0 | 28.09% | 36268.7 | 24.88% | motif file (matrix) | svg |
| 469 | C T G A T A G C T G A C T C A G C T A G G T C A C G T A T C A G A G C T T C A G | ETV4(ETS)/HepG2-ETV4-ChIP-Seq(ENCODE)/Homer | 1e-32 | -7.405e+01 | 0.0000 | 17729.0 | 54.92% | 74737.1 | 51.27% | motif file (matrix) | svg |
| 470 | T G C A C G T A G T C A A G C T A G T C G C T A T A G C C G A T C T A G G A T C | Gfi1b(Zf)/HPC7-Gfi1b-ChIP-Seq(GSE22178)/Homer | 1e-32 | -7.379e+01 | 0.0000 | 7792.0 | 24.14% | 30758.9 | 21.10% | motif file (matrix) | svg |
| 471 | T G C A A G C T A C G T C T A G G A T C C T A G G A T C G T C A C T G A A G T C | CEBP(bZIP)/ThioMac-CEBPb-ChIP-Seq(GSE21512)/Homer | 1e-31 | -7.333e+01 | 0.0000 | 12008.0 | 37.20% | 49163.9 | 33.73% | motif file (matrix) | svg |
| 472 | T C G A A C G T A C T G C G T A A G T C C T A G A G C T T G A C | TGA10(bZIP)/colamp-TGA10-DAP-Seq(GSE60143)/Homer | 1e-31 | -7.202e+01 | 0.0000 | 13820.0 | 42.81% | 57246.9 | 39.27% | motif file (matrix) | svg |
| 473 | G C T A G C A T G A C T G C A T T C A G G T A C G C T A G C A T C T G A G C T A T A G C G C T A C T G A C G A T C T A G | OCT4-SOX2-TCF-NANOG(POU,Homeobox,HMG)/mES-Oct4-ChIP-Seq(GSE11431)/Homer | 1e-31 | -7.147e+01 | 0.0000 | 1226.0 | 3.80% | 3741.2 | 2.57% | motif file (matrix) | svg |
| 474 | C T G A C T G A C T A G T C G A C G T A A T G C C G T A A C T G C G T A A C G T C T G A C G A T A G C T C G T A A C G T A G T C C G A T T A C G G T C A G C A T | GATA(Zf),IR3/iTreg-Gata3-ChIP-Seq(GSE20898)/Homer | 1e-30 | -7.071e+01 | 0.0000 | 2362.0 | 7.32% | 8146.1 | 5.59% | motif file (matrix) | svg |
| 475 | G C A T G C A T G A C T G C A T T A G C A T G C G A T C C A T G A G T C A G T C | DEL2(E2FDP)/col-DEL2-DAP-Seq(GSE60143)/Homer | 1e-30 | -7.023e+01 | 0.0000 | 11259.0 | 34.88% | 45972.5 | 31.54% | motif file (matrix) | svg |
| 476 | A T G C A G C T T C A G T G A C T C A G A T G C T G C A A C G T A T C G G A T C A C T G A G T C | NRF1(NRF)/MCF7-NRF1-ChIP-Seq(Unpublished)/Homer | 1e-30 | -6.978e+01 | 0.0000 | 2006.0 | 6.21% | 6755.8 | 4.63% | motif file (matrix) | svg |
| 477 | A G T C T A G C A C T G A C G T A C G T C G T A C G T A C A G T C G A T A G T C C T A G A C T G A C G T A C G T C T G A | MYB44(MYB)/colamp-MYB44-DAP-Seq(GSE60143)/Homer | 1e-30 | -6.933e+01 | 0.0000 | 1840.0 | 5.70% | 6113.9 | 4.19% | motif file (matrix) | svg |
| 478 | T A G C T C A G C A T G G C A T A G C T C G A T A T G C C G T A C G T A G T C A | CHR(?)/Hela-CellCycle-Expression/Homer | 1e-29 | -6.837e+01 | 0.0000 | 5961.0 | 18.47% | 23089.0 | 15.84% | motif file (matrix) | svg |
| 479 | C T A G C T G A A G T C G C T A C G A T A C T G G A C T G A T C G A T C C T G A C T A G C T G A T G A C G C T A C G A T T C A G G A C T G A T C G A T C T G A C | p53(p53)/Saos-p53-ChIP-Seq(GSE15780)/Homer | 1e-28 | -6.545e+01 | 0.0000 | 1732.0 | 5.37% | 5751.5 | 3.95% | motif file (matrix) | svg |
| 480 | C T A G C T G A A G T C G C T A C G A T A C T G G A C T G A T C G A T C C T G A C T A G C T G A T G A C G C T A C G A T T C A G G A C T G A T C G A T C T G A C | p53(p53)/Saos-p53-ChIP-Seq/Homer | 1e-28 | -6.545e+01 | 0.0000 | 1732.0 | 5.37% | 5751.5 | 3.95% | motif file (matrix) | svg |
| 481 | C G T A A G T C T G A C A G C T A C G T C G T A A C G T A G T C | At5g05790(MYBrelated)/col-At5g05790-DAP-Seq(GSE60143)/Homer | 1e-28 | -6.525e+01 | 0.0000 | 16169.0 | 50.09% | 68029.5 | 46.67% | motif file (matrix) | svg |
| 482 | T A C G A C T G A G C T G T A C C G T A T C G A C T G A A C T G C A T G A C G T A G T C C G T A | COUP-TFII(NR)/K562-NR2F1-ChIP-Seq(Encode)/Homer | 1e-27 | -6.385e+01 | 0.0000 | 16702.0 | 51.74% | 70488.9 | 48.35% | motif file (matrix) | svg |
| 483 | T G C A C T A G C T A G C T G A C A T G A C T G T G C A G A T C G T C A T G C A G T C A G T C A A G C T C T A G G C A T | ZNF675(Zf)/HEK293-ZNF675.GFP-ChIP-Seq(GSE58341)/Homer | 1e-27 | -6.218e+01 | 0.0000 | 2033.0 | 6.30% | 6986.8 | 4.79% | motif file (matrix) | svg |
| 484 | T C A G C T G A C T A G C A T G A C G T A T G C C T G A C T G A C T G A C T A G C A T G A C G T A T G C C T G A | TR4(NR),DR1/Hela-TR4-ChIP-Seq(GSE24685)/Homer | 1e-26 | -6.182e+01 | 0.0000 | 1110.0 | 3.44% | 3422.3 | 2.35% | motif file (matrix) | svg |
| 485 | A G C T G C A T A C T G A C G T A G T C A C G T C T A G T A C G | Smad3(MAD)/NPC-Smad3-ChIP-Seq(GSE36673)/Homer | 1e-26 | -6.165e+01 | 0.0000 | 23277.0 | 72.11% | 100685.3 | 69.07% | motif file (matrix) | svg |
| 486 | A C G T C T A G C G T A A G T C G T A C A C G T A C G T A C G T G T C A G T A C T G A C G A C T | Nur77(NR)/K562-NR4A1-ChIP-Seq(GSE31363)/Homer | 1e-26 | -6.160e+01 | 0.0000 | 2434.0 | 7.54% | 8598.0 | 5.90% | motif file (matrix) | svg |
| 487 | G A C T A G T C C G T A C G T A A G T C A G C T A C T G G A C T G T A C A T G C | MYB77(MYB)/col-MYB77-DAP-Seq(GSE60143)/Homer | 1e-26 | -6.072e+01 | 0.0000 | 23458.0 | 72.67% | 101563.8 | 69.67% | motif file (matrix) | svg |
| 488 | C T G A A C G T A C G T A C G T A G T C G A C T C G A T C T G A A C T G C G T A C G T A T C G A | STAT5(Stat)/mCD4+-Stat5-ChIP-Seq(GSE12346)/Homer | 1e-25 | -5.971e+01 | 0.0000 | 3027.0 | 9.38% | 11044.4 | 7.58% | motif file (matrix) | svg |
| 489 | A G C T G A C T A C T G A C G T G T C A A G T C A C G T C G A T | SPL9(SBP)/colamp-SPL9-DAP-Seq(GSE60143)/Homer | 1e-25 | -5.913e+01 | 0.0000 | 25937.0 | 80.35% | 113247.8 | 77.69% | motif file (matrix) | svg |
| 490 | C G T A C G T A C G T A C G T A C G T A A C T G A G C T C T A G G T A C G C T A | AT1G69570(C2C2dof)/col-AT1G69570-DAP-Seq(GSE60143)/Homer | 1e-25 | -5.882e+01 | 0.0000 | 13972.0 | 43.28% | 58439.4 | 40.09% | motif file (matrix) | svg |
| 491 | C G T A A T G C C G A T A C G T A G T C C G T A C G T A C G T A C T A G A T C G | TCFL2(HMG)/K562-TCF7L2-ChIP-Seq(GSE29196)/Homer | 1e-25 | -5.845e+01 | 0.0000 | 1154.0 | 3.57% | 3629.8 | 2.49% | motif file (matrix) | svg |
| 492 | A G T C G A T C G C T A C G A T A C G T T A C G G C A T C T G A G A C T A C T G A G T C G C T A C T G A T C G A C A G T | Oct4:Sox17(POU,Homeobox,HMG)/F9-Sox17-ChIP-Seq(GSE44553)/Homer | 1e-24 | -5.716e+01 | 0.0000 | 1506.0 | 4.67% | 4997.5 | 3.43% | motif file (matrix) | svg |
| 493 | C A T G G T C A A G T C C G T A C T A G G A T C C G A T A C T G A C G T G T A C C G T A C G T A | bZIP69(bZIP)/col-bZIP69-DAP-Seq(GSE60143)/Homer | 1e-24 | -5.627e+01 | 0.0000 | 1492.0 | 4.62% | 4956.5 | 3.40% | motif file (matrix) | svg |
| 494 | G C T A C T G A C G T A C G T A C T G A C T G A C G A T G T C A A C G T A G T C G C A T G C A T | At5g52660(MYBrelated)/colamp-At5g52660-DAP-Seq(GSE60143)/Homer | 1e-24 | -5.584e+01 | 0.0000 | 4253.0 | 13.17% | 16224.7 | 11.13% | motif file (matrix) | svg |
| 495 | A C T G G A C T A G T C C T G A G A T C T C A G A T G C G A C T A G T C A T G C T A G C A G C T A T C G T G C A | PAX5(Paired,Homeobox),condensed/GM12878-PAX5-ChIP-Seq(GSE32465)/Homer | 1e-23 | -5.465e+01 | 0.0000 | 2668.0 | 8.26% | 9686.5 | 6.64% | motif file (matrix) | svg |
| 496 | A G C T G A T C A G C T G A C T G A C T T C G A A G T C C G T A A C T G T C A G | SpliceAcceptor/Homer | 1e-23 | -5.368e+01 | 0.0000 | 27668.0 | 85.71% | 121669.5 | 83.46% | motif file (matrix) | svg |
| 497 | A T G C C T G A G A C T A C G T A C G T G T A C G A T C C G A T C T A G C A T G C G T A C G T A C T G A G A C T | STAT1(Stat)/HelaS3-STAT1-ChIP-Seq(GSE12782)/Homer | 1e-23 | -5.316e+01 | 0.0000 | 2848.0 | 8.82% | 10455.3 | 7.17% | motif file (matrix) | svg |
| 498 | G C T A C G T A A C G T A T C G C G T A A C G T A C G T C T A G | ATHB6(Homeobox)/col-ATHB6-DAP-Seq(GSE60143)/Homer | 1e-23 | -5.299e+01 | 0.0000 | 15364.0 | 47.59% | 64922.4 | 44.54% | motif file (matrix) | svg |
| 499 | T G A C T C G A C T G A C T G A A T G C G A T C C T A G T A C G G A C T G A C T G A T C T C G A C T G A C T G A A T G C G A T C C T A G A T C G G A C T G A C T | Tcfcp2l1(CP2)/mES-Tcfcp2l1-ChIP-Seq(GSE11431)/Homer | 1e-22 | -5.204e+01 | 0.0000 | 2976.0 | 9.22% | 11008.0 | 7.55% | motif file (matrix) | svg |
| 500 | C G T A C T G A C T A G C G T A C G T A A G T C C G T A C A G T G C A T G T C A C G A T A C T G A C G T G C A T G A T C | PGR(NR)/EndoStromal-PGR-ChIP-Seq(GSE69539)/Homer | 1e-22 | -5.108e+01 | 0.0000 | 2818.0 | 8.73% | 10380.0 | 7.12% | motif file (matrix) | svg |
| 501 | G A C T A G T C C G A T A C T G C T G A T G A C G T A C C G T A A T C G G C A T C T G A C T A G | Bcl11a(Zf)/HSPC-BCL11A-ChIP-Seq(GSE104676)/Homer | 1e-22 | -5.101e+01 | 0.0000 | 8144.0 | 25.23% | 33048.3 | 22.67% | motif file (matrix) | svg |
| 502 | A C T G C G T A A C G T C G T A C T G A A C T G T C A G G C A T | At3g11280(MYBrelated)/col-At3g11280-DAP-Seq(GSE60143)/Homer | 1e-22 | -5.086e+01 | 0.0000 | 15614.0 | 48.37% | 66141.8 | 45.37% | motif file (matrix) | svg |
| 503 | C T A G A T C G G T C A C A T G A G T C G A C T T A C G C A G T A G T C A G T C C T G A C G A T C T A G A T C G G A C T A T C G A G T C G A C T C T A G T C G A | REST-NRSF(Zf)/Jurkat-NRSF-ChIP-Seq/Homer | 1e-21 | -5.045e+01 | 0.0000 | 98.0 | 0.30% | 105.5 | 0.07% | motif file (matrix) | svg |
| 504 | G T A C C G T A C G T A T A C G G C A T G T A C C G T A C A T G A G T C C G T A C G T A C G A T G C A T G C A T G A C T | MafF(bZIP)/HepG2-MafF-ChIP-Seq(GSE31477)/Homer | 1e-21 | -4.950e+01 | 0.0000 | 3029.0 | 9.38% | 11286.2 | 7.74% | motif file (matrix) | svg |
| 505 | T C G A A G T C C G T A A T C G T A G C A C G T A C T G A G C T A C G T A G T C | Ptf1a(bHLH)/Panc1-Ptf1a-ChIP-Seq(GSE47459)/Homer | 1e-21 | -4.906e+01 | 0.0000 | 21129.0 | 65.45% | 91283.4 | 62.62% | motif file (matrix) | svg |
| 506 | G C A T C G A T A T G C A G C T T C G A T A C G G C T A C G T A C A T G T G A C G C A T C G A T A G T C A G C T C G T A | AT3G09735(S1Falike)/col-AT3G09735-DAP-Seq(GSE60143)/Homer | 1e-20 | -4.826e+01 | 0.0000 | 4625.0 | 14.33% | 18009.6 | 12.35% | motif file (matrix) | svg |
| 507 | T G C A A G C T C A T G C G T A A G C T A C T G G A T C G T C A C G T A A G C T | Atf4(bZIP)/MEF-Atf4-ChIP-Seq(GSE35681)/Homer | 1e-20 | -4.782e+01 | 0.0000 | 5317.0 | 16.47% | 20967.4 | 14.38% | motif file (matrix) | svg |
| 508 | C T G A T A C G G C A T C T A G A T G C G A T C C G A T A C T G C T A G G A T C C T G A A T G C | MYRF(MYRF)/CFPAC1-MYRF-ChIP-Seq(GSE145627)/Homer | 1e-20 | -4.632e+01 | 0.0000 | 5112.0 | 15.84% | 20141.3 | 13.82% | motif file (matrix) | svg |
| 509 | T G C A A T G C A C G T A C G T A C G T A T G C C T A G A C G T A C G T A G C T G A T C A G C T | T1ISRE(IRF)/ThioMac-Ifnb-Expression/Homer | 1e-20 | -4.612e+01 | 0.0000 | 206.0 | 0.64% | 401.6 | 0.28% | motif file (matrix) | svg |
| 510 | T G A C C G T A C T A G T A C G G A C T C T G A C T G A T C A G C A G T T C A G | SpliceDonor/U1snRNP/Homer | 1e-19 | -4.486e+01 | 0.0000 | 29287.0 | 90.73% | 129762.4 | 89.02% | motif file (matrix) | svg |
| 511 | T C G A T C A G T C G A A C T G C A T G A C G T A G T C C T G A | COUP-TFII(NR)/Artia-Nr2f2-ChIP-Seq(GSE46497)/Homer | 1e-19 | -4.474e+01 | 0.0000 | 19400.0 | 60.10% | 83573.1 | 57.33% | motif file (matrix) | svg |
| 512 | A T G C C G T A A C T G C G T A A C G T G C T A T C G A A G C T C G A T C G T A A C G T A G T C C G A T A C T G G A T C | GATA(Zf),IR4/iTreg-Gata3-ChIP-Seq(GSE20898)/Homer | 1e-19 | -4.459e+01 | 0.0000 | 1444.0 | 4.47% | 4956.8 | 3.40% | motif file (matrix) | svg |
| 513 | C T G A G T A C G A C T A G T C C A G T T G C A C T G A A C G T A G C T G A T C C T A G C G A T A C T G A T G C G A C T C T G A G A T C G A C T A G C T G A T C | Mouse\_Recombination\_Hotspot(Zf)/Testis-DMC1-ChIP-Seq(GSE24438)/Homer | 1e-18 | -4.344e+01 | 0.0000 | 866.0 | 2.68% | 2733.5 | 1.88% | motif file (matrix) | svg |
| 514 | C G A T C A G T C A G T C A T G G T C A G A T C C G T A T C A G A G T C A C G T C T A G A C G T G T A C G T C A G C T A | VIP1(bZIP)/col-VIP1-DAP-Seq(GSE60143)/Homer | 1e-18 | -4.277e+01 | 0.0000 | 2262.0 | 7.01% | 8288.1 | 5.69% | motif file (matrix) | svg |
| 515 | A C G T T A C G G A T C A C T G A C G T C T A G A C T G A C T G G A T C C T A G C A T G C T A G | Egr2(Zf)/Thymocytes-Egr2-ChIP-Seq(GSE34254)/Homer | 1e-18 | -4.243e+01 | 0.0000 | 3167.0 | 9.81% | 12040.0 | 8.26% | motif file (matrix) | svg |
| 516 | C G T A G A C T C G T A A C G T C A G T A G T C A G C T G A C T | KAN2(G2like)/colamp-KAN2-DAP-Seq(GSE60143)/Homer | 1e-18 | -4.205e+01 | 0.0000 | 14846.0 | 45.99% | 63108.3 | 43.29% | motif file (matrix) | svg |
| 517 | C G A T C T G A G T A C C A T G G C A T T C A G G C A T C G T A C G T A G C T A C G T A A G T C G C T A G T A C C A T G | CUC2(NAC)/colamp-CUC2-DAP-Seq(GSE60143)/Homer | 1e-18 | -4.186e+01 | 0.0000 | 7976.0 | 24.71% | 32678.7 | 22.42% | motif file (matrix) | svg |
| 518 | C G T A C G T A T C G A C G T A C G A T C G T A A C G T A G T C G C A T G C A T | At3g09600(MYBrelated)/colamp-At3g09600-DAP-Seq(GSE60143)/Homer | 1e-18 | -4.176e+01 | 0.0000 | 4701.0 | 14.56% | 18538.4 | 12.72% | motif file (matrix) | svg |
| 519 | G A C T G A T C G A C T A C G T C G T A A C G T A G T C A G T C C G T A G C A T G C T A G C A T | At1g74840(MYBrelated)/col100-At1g74840-DAP-Seq(GSE60143)/Homer | 1e-17 | -4.135e+01 | 0.0000 | 10598.0 | 32.83% | 44219.5 | 30.33% | motif file (matrix) | svg |
| 520 | T G C A C T G A C A T G C T A G C A G T A G T C C G T A A T G C A T G C T A C G G C A T T C A G G T C A G A T C G T A C | ERE(NR),IR3/MCF7-ERa-ChIP-Seq(Unpublished)/Homer | 1e-17 | -4.124e+01 | 0.0000 | 3604.0 | 11.16% | 13906.0 | 9.54% | motif file (matrix) | svg |
| 521 | G C A T T C G A C T G A G A T C A G T C G A T C G T C A G T C A A C G T A G T C C G T A C T G A | Duxbl(Homeobox)/NIH3T3-Duxbl.HA-ChIP-Seq(GSE119782)/Homer | 1e-17 | -4.069e+01 | 0.0000 | 1197.0 | 3.71% | 4049.2 | 2.78% | motif file (matrix) | svg |
| 522 | C G T A C T G A T C A G A C T G G T C A C G T A A C G T G T A C C G A T G C A T | AT5G45580(G2like)/colamp-AT5G45580-DAP-Seq(GSE60143)/Homer | 1e-17 | -3.982e+01 | 0.0000 | 18990.0 | 58.83% | 81950.5 | 56.22% | motif file (matrix) | svg |
| 523 | T G C A T A G C G A C T T G C A T G A C T G C A C G T A A G C T A G C T A G T C A G T C G T A C | GFY(?)/Promoter/Homer | 1e-17 | -3.976e+01 | 0.0000 | 1087.0 | 3.37% | 3634.0 | 2.49% | motif file (matrix) | svg |
| 524 | T G A C A T G C C G T A A T C G A T G C C A G T C A T G A C T G A G T C G T A C | HEB(bHLH)/mES-Heb-ChIP-Seq(GSE53233)/Homer | 1e-17 | -3.962e+01 | 0.0000 | 17493.0 | 54.19% | 75165.2 | 51.56% | motif file (matrix) | svg |
| 525 | C A T G G A T C C T G A A G T C C T A G C G T A G C T A G C A T G A T C G A T C A G T C C T A G C G T A C A T G C T A G | PLT1(AP2EREBP)/colamp-PLT1-DAP-Seq(GSE60143)/Homer | 1e-17 | -3.949e+01 | 0.0000 | 2799.0 | 8.67% | 10585.1 | 7.26% | motif file (matrix) | svg |
| 526 | A C G T A G T C A G C T A G T C C G T A G T A C A G T C C G A T C G T A G T C A | MYB41(MYB)/col-MYB41-DAP-Seq(GSE60143)/Homer | 1e-16 | -3.844e+01 | 0.0000 | 7612.0 | 23.58% | 31234.7 | 21.43% | motif file (matrix) | svg |
| 527 | G A T C G A T C G C T A G T C A G A C T A T G C T C G A C G A T C G A T C T A G | HAT2(Homeobox)/colamp-HAT2-DAP-Seq(GSE60143)/Homer | 1e-16 | -3.707e+01 | 0.0000 | 11730.0 | 36.34% | 49443.8 | 33.92% | motif file (matrix) | svg |
| 528 | T C G A C G T A A G T C A G C T C G T A A G T C T C G A G C T A G A C T C G A T A G T C A G T C A G T C C T G A T C A G T G C A T C G A C A G T A T C G A G T C | GFY-Staf(?,Zf)/Promoter/Homer | 1e-16 | -3.704e+01 | 0.0000 | 565.0 | 1.75% | 1687.4 | 1.16% | motif file (matrix) | svg |
| 529 | C G T A C T G A C T A G C T G A A G T C G C T A C G A T A T C G G A C T G A T C A G T C C T G A C T A G C T A G A G T C G C T A C G A T C T A G G A T C G A T C | p73(p53)/Trachea-p73-ChIP-Seq(PRJNA310161)/Homer | 1e-15 | -3.660e+01 | 0.0000 | 824.0 | 2.55% | 2665.2 | 1.83% | motif file (matrix) | svg |
| 530 | C G T A A C T G G T C A A C G T A T C G C A G T C T A G T C A G C G T A A C T G C G T A A C G T C G T A C T G A T A C G | GATA3(Zf),DR4/iTreg-Gata3-ChIP-Seq(GSE20898)/Homer | 1e-15 | -3.558e+01 | 0.0000 | 1254.0 | 3.88% | 4362.3 | 2.99% | motif file (matrix) | svg |
| 531 | A G T C G A C T C A G T G T A C A G T C A T C G T C A G A C T G G T C A C G T A | Stat3(Stat)/mES-Stat3-ChIP-Seq(GSE11431)/Homer | 1e-15 | -3.523e+01 | 0.0000 | 6657.0 | 20.62% | 27212.1 | 18.67% | motif file (matrix) | svg |
| 532 | T G C A C T G A A G T C G T C A A C T G A C T G C G T A C G T A C T G A A G C T | EWS:FLI1-fusion(ETS)/SK\_N\_MC-EWS:FLI1-ChIP-Seq(SRA014231)/Homer | 1e-15 | -3.510e+01 | 0.0000 | 7499.0 | 23.23% | 30887.9 | 21.19% | motif file (matrix) | svg |
| 533 | C T A G T C G A A C G T A C G T C A T G A G T C C T G A C G A T A G T C C G T A | AARE(HLH)/mES-cMyc-ChIP-Seq/Homer | 1e-15 | -3.504e+01 | 0.0000 | 1891.0 | 5.86% | 6946.4 | 4.77% | motif file (matrix) | svg |
| 534 | G T C A G C A T G C T A C A G T C T A G G A T C C G T A C T G A C G T A C G A T | Oct2(POU,Homeobox)/Bcell-Oct2-ChIP-Seq(GSE21512)/Homer | 1e-15 | -3.479e+01 | 0.0000 | 2746.0 | 8.51% | 10492.3 | 7.20% | motif file (matrix) | svg |
| 535 | T C A G C A T G C A T G A C T G A C T G A G C T A C T G A C G T A C T G C A G T A T G C A G T C | KLF10(Zf)/HEK293-KLF10.GFP-ChIP-Seq(GSE58341)/Homer | 1e-14 | -3.419e+01 | 0.0000 | 6376.0 | 19.75% | 26035.2 | 17.86% | motif file (matrix) | svg |
| 536 | C G A T C T G A G T A C A C T G A C G T T C A G G C A T C G T A C G T A G C A T C G T A A G T C C G T A G T A C C A T G | CUC3(NAC)/col-CUC3-DAP-Seq(GSE60143)/Homer | 1e-14 | -3.402e+01 | 0.0000 | 7834.0 | 24.27% | 32401.4 | 22.23% | motif file (matrix) | svg |
| 537 | G A T C G C A T G C A T A G T C A G C T T C G A T A C G G C T A C G T A C T A G T G A C G C A T C G A T G A T C A G C T | HSFC1(HSF)/col-HSFC1-DAP-Seq(GSE60143)/Homer | 1e-14 | -3.402e+01 | 0.0000 | 1873.0 | 5.80% | 6896.4 | 4.73% | motif file (matrix) | svg |
| 538 | T A G C C G A T T A C G A C T G A G T C A C T G A T C G A T C G C G T A C T G A | E2F1(E2F)/Hela-E2F1-ChIP-Seq(GSE22478)/Homer | 1e-14 | -3.353e+01 | 0.0000 | 6809.0 | 21.09% | 27947.8 | 19.17% | motif file (matrix) | svg |
| 539 | C T A G C T A G T C G A C G T A A T G C C G T A A T C G T C G A T A C G G C A T A C T G C A G T T A G C G A T C G A C T | MRE(NR)/Neuro2A-NR3C2-ChIPnexus(GSE115417)/Homer | 1e-14 | -3.322e+01 | 0.0000 | 14977.0 | 46.40% | 64162.7 | 44.01% | motif file (matrix) | svg |
| 540 | G T A C A C T G A C G T T C A G G C A T C G T A C G A T G C A T C G T A A G T C C G T A T G A C C A T G G A C T G C T A | ANAC083(NAC)/col-ANAC083-DAP-Seq(GSE60143)/Homer | 1e-14 | -3.304e+01 | 0.0000 | 13724.0 | 42.51% | 58553.0 | 40.17% | motif file (matrix) | svg |
| 541 | A C T G T G A C A C T G A C G T A C G T A C T G C G T A A G T C A G C T C G A T G C A T A C G T | WRKY17(WRKY)/colamp-WRKY17-DAP-Seq(GSE60143)/Homer | 1e-14 | -3.299e+01 | 0.0000 | 313.0 | 0.97% | 826.9 | 0.57% | motif file (matrix) | svg |
| 542 | T C G A A C G T A C T G C T G A A G T C T C A G A G C T G T A C C G T A A G C T G A T C T C G A | JunD(bZIP)/K562-JunD-ChIP-Seq/Homer | 1e-14 | -3.247e+01 | 0.0000 | 1156.0 | 3.58% | 4029.7 | 2.76% | motif file (matrix) | svg |
| 543 | T C A G C G T A A G T C A G C T C G T A A G T C C T G A C G T A A G T C G C A T A G T C A G T C A G T C C T G A A C T G T G C A T C G A C A T G A T C G G A T C | Ronin(THAP)/ES-Thap11-ChIP-Seq(GSE51522)/Homer | 1e-14 | -3.227e+01 | 0.0000 | 192.0 | 0.59% | 431.0 | 0.30% | motif file (matrix) | svg |
| 544 | C T A G C A T G C A T G T A C G A G T C G C A T A G C T C T A G A C G T A G T C G A C T A C T G A C T G A C T G T C G A | Zfp809(Zf)/ES-Zfp809-ChIP-Seq(GSE70799)/Homer | 1e-13 | -3.039e+01 | 0.0000 | 2361.0 | 7.31% | 9008.2 | 6.18% | motif file (matrix) | svg |
| 545 | C T A G A T G C A T G C C G A T A C T G G A C T A T G C G C T A T G A C A G C T T A G C G C T A | PBX1(Homeobox)/MCF7-PBX1-ChIP-Seq(GSE28007)/Homer | 1e-12 | -2.936e+01 | 0.0000 | 873.0 | 2.70% | 2965.3 | 2.03% | motif file (matrix) | svg |
| 546 | A T C G A G T C A G T C C G T A A C T G G C A T | hINR(CPE) | 1e-12 | -2.848e+01 | 0.0000 | 15100.0 | 46.78% | 64993.8 | 44.58% | motif file (matrix) | svg |
| 547 | A C T G A G T C G T C A C G T A A G T C C G T A C T A G C T A G G A C T C A T G | SCRT1(Zf)/HEK293-SCRT1.eGFP-ChIP-Seq(Encode)/Homer | 1e-12 | -2.846e+01 | 0.0000 | 5723.0 | 17.73% | 23449.2 | 16.09% | motif file (matrix) | svg |
| 548 | A T G C G C A T C G A T G A T C A G C T C T G A A C T G C G T A C G T A T C A G T G A C C G A T G C A T G A T C C G A T | HSF21(HSF)/col-HSF21-DAP-Seq(GSE60143)/Homer | 1e-12 | -2.803e+01 | 0.0000 | 940.0 | 2.91% | 3252.4 | 2.23% | motif file (matrix) | svg |
| 549 | G C T A T A G C A G C T A T C G G T C A C G T A G C T A A T G C G A T C C T G A | IRF4(IRF)/GM12878-IRF4-ChIP-Seq(GSE32465)/Homer | 1e-11 | -2.714e+01 | 0.0000 | 6841.0 | 21.19% | 28385.5 | 19.47% | motif file (matrix) | svg |
| 550 | T A G C A G T C T G A C A G T C C T A G A T C G A G T C C A T G T G A C A G T C G T A C A G T C A G T C G C A T C T A G A T C G G C A T A C T G A T C G G A T C | BORIS(Zf)/K562-CTCFL-ChIP-Seq(GSE32465)/Homer | 1e-11 | -2.672e+01 | 0.0000 | 1916.0 | 5.94% | 7257.1 | 4.98% | motif file (matrix) | svg |
| 551 | T A G C C G A T A C G T A G C T A G C T A G T C A T G C A G T C A C T G A T G C A T G C G C T A | E2F7(E2F)/Hela-E2F7-ChIP-Seq(GSE32673)/Homer | 1e-11 | -2.640e+01 | 0.0000 | 4045.0 | 12.53% | 16287.3 | 11.17% | motif file (matrix) | svg |
| 552 | C T A G G A C T G A T C A C G T A T C G A G C T C T G A A T C G C G A T C T A G G A T C G A C T C A T G A T C G G T A C G A C T A G T C G C A T A G C T C G A T | ZNF382(Zf)/HEK293-ZNF382.GFP-ChIP-Seq(GSE58341)/Homer | 1e-11 | -2.616e+01 | 0.0000 | 363.0 | 1.12% | 1066.5 | 0.73% | motif file (matrix) | svg |
| 553 | C T G A T A C G A C G T C T A G C G T A T C G A C T G A C G A T | At5g04390(C2H2)/col200-At5g04390-DAP-Seq(GSE60143)/Homer | 1e-10 | -2.516e+01 | 0.0000 | 28549.0 | 88.44% | 126946.8 | 87.08% | motif file (matrix) | svg |
| 554 | C T G A T G C A T A G C T G A C T A C G T C A G C T G A G C T A T C A G G A C T | ELF1(ETS)/Jurkat-ELF1-ChIP-Seq(SRA014231)/Homer | 1e-10 | -2.499e+01 | 0.0000 | 11825.0 | 36.63% | 50542.5 | 34.67% | motif file (matrix) | svg |
| 555 | A G T C G C A T C G T A C G T A G T A C A C G T A C T G G A T C G A T C T C G A | BMYB(HTH)/Hela-BMYB-ChIP-Seq(GSE27030)/Homer | 1e-10 | -2.353e+01 | 0.0000 | 23226.0 | 71.95% | 102258.1 | 70.15% | motif file (matrix) | svg |
| 556 | C G T A C G T A C T G A A C T G C G T A C G T A A C G T G T C A A C G T G C A T A G T C G A C T | At2g03500(G2like)/col-At2g03500-DAP-Seq(GSE60143)/Homer | 1e-10 | -2.330e+01 | 0.0000 | 4268.0 | 13.22% | 17377.3 | 11.92% | motif file (matrix) | svg |
| 557 | G A C T A C G T C G A T A G C T A G T C C G T A A C T G A C T G C G A T C T A G | NGA4(ABI3VP1)/col-NGA4-DAP-Seq(GSE60143)/Homer | 1e-9 | -2.299e+01 | 0.0000 | 21910.0 | 67.87% | 96254.7 | 66.03% | motif file (matrix) | svg |
| 558 | C G T A C T G A T A C G G A C T G A T C T C G A G A T C G A T C T G C A G A C T T C G A C G T A A T C G A G T C C G A T C G T A C G T A G T C A C G T A C T A G | PSE(SNAPc)/K562-mStart-Seq/Homer | 1e-9 | -2.193e+01 | 0.0000 | 7049.0 | 21.84% | 29573.4 | 20.29% | motif file (matrix) | svg |
| 559 | C G A T C G T A G C A T C T A G A C T G C G T A A C G T G T C A C G T A C T A G C T A G G C A T | At1g19000(MYBrelated)/colamp-At1g19000-DAP-Seq(GSE60143)/Homer | 1e-9 | -2.186e+01 | 0.0000 | 11484.0 | 35.58% | 49223.2 | 33.77% | motif file (matrix) | svg |
| 560 | G C A T G C A T G A T C G A C T T C G A T C A G G C T A C G T A A C T G G T A C G C A T G C A T A G T C A G C T C G T A | HSF7(HSF)/colamp-HSF7-DAP-Seq(GSE60143)/Homer | 1e-9 | -2.149e+01 | 0.0000 | 1720.0 | 5.33% | 6588.6 | 4.52% | motif file (matrix) | svg |
| 561 | T A C G T A C G G T A C A T C G T A C G T A C G G T C A C T G A C G T A G A C T | E2F4(E2F)/K562-E2F4-ChIP-Seq(GSE31477)/Homer | 1e-9 | -2.135e+01 | 0.0000 | 15165.0 | 46.98% | 65755.7 | 45.11% | motif file (matrix) | svg |
| 562 | C T A G T A C G G A T C G T C A T C G A A C G T T C A G C G T A C G T A C G T A | Hoxd10(Homeobox)/ChickenMSG-Hoxd10.Flag-ChIP-Seq(GSE86088)/Homer | 1e-9 | -2.129e+01 | 0.0000 | 12918.0 | 40.02% | 55671.5 | 38.19% | motif file (matrix) | svg |
| 563 | C A G T T C A G G A T C A C T G A C G T C T A G A C T G A C T G G A C T C T A G | Egr1(Zf)/K562-Egr1-ChIP-Seq(GSE32465)/Homer | 1e-9 | -2.113e+01 | 0.0000 | 9468.0 | 29.33% | 40304.4 | 27.65% | motif file (matrix) | svg |
| 564 | G C T A C G T A C G T A C G T A A C T G A C G T A G T C C G T A C G T A A G T C C A T G T A G C G T A C C G T A C G T A | WRKY7(WRKY)/colamp-WRKY7-DAP-Seq(GSE60143)/Homer | 1e-9 | -2.108e+01 | 0.0000 | 98.0 | 0.30% | 200.8 | 0.14% | motif file (matrix) | svg |
| 565 | A G T C C T A G A T C G C A G T C G A T A G C T G T A C A C T G C A T G C A T G | ZBED2(Zf)/SUIT2-ZBED2.HA-ChIP-Seq(GSE141606)/Homer | 1e-9 | -2.091e+01 | 0.0000 | 15695.0 | 48.62% | 68174.4 | 46.77% | motif file (matrix) | svg |
| 566 | G A T C G C T A A G C T G C A T A G T C C T G A T A G C G A C T | STZ(C2H2)/colamp-STZ-DAP-Seq(GSE60143)/Homer | 1e-9 | -2.074e+01 | 0.0000 | 28552.0 | 88.45% | 127166.5 | 87.23% | motif file (matrix) | svg |
| 567 | G A T C G T A C C G A T A C T G A C T G C G T A C G T A A C G T A C T G G A T C | TEAD(TEA)/Fibroblast-PU.1-ChIP-Seq(Unpublished)/Homer | 1e-8 | -1.953e+01 | 0.0000 | 6883.0 | 21.32% | 28982.7 | 19.88% | motif file (matrix) | svg |
| 568 | C T A G T G A C G A C T A T C G T C G A A G T C C T A G C A G T C T A G A T C G G T A C T C G A | O2(bZIP)/Corn-O2-ChIP-Seq(GSE63991)/Homer | 1e-8 | -1.939e+01 | 0.0000 | 3477.0 | 10.77% | 14138.0 | 9.70% | motif file (matrix) | svg |
| 569 | C G A T T C G A A C T G G T C A C G T A C G A T G T A C G A C T | At3g04030(G2like)/col-At3g04030-DAP-Seq(GSE60143)/Homer | 1e-8 | -1.902e+01 | 0.0000 | 12511.0 | 38.76% | 54012.0 | 37.05% | motif file (matrix) | svg |
| 570 | C T A G A G T C A G C T A C T G C G T A C A G T C G T A C T G A T A G C T G A C | Unknown5/Drosophila-Promoters/Homer | 1e-8 | -1.886e+01 | 0.0000 | 12804.0 | 39.66% | 55335.3 | 37.96% | motif file (matrix) | svg |
| 571 | C G T A C G A T C G T A T C G A T C G A A C G T C G T A A C G T A G T C G C A T | LHY(Myb)/Seedling-LHY-ChIP-Seq(GSE52175)/Homer | 1e-8 | -1.865e+01 | 0.0000 | 13218.0 | 40.95% | 57206.2 | 39.24% | motif file (matrix) | svg |
| 572 | G A T C G C A T A C G T C G T A A C G T A G T C A G T C C G T A | AT5G61620(MYBrelated)/colamp-AT5G61620-DAP-Seq(GSE60143)/Homer | 1e-7 | -1.722e+01 | 0.0000 | 19416.0 | 60.15% | 85296.4 | 58.51% | motif file (matrix) | svg |
| 573 | T G A C C T G A C T A G C T G A C G T A A G T C C T G A A C G T G C A T T A G C G C A T A T C G G A C T G A C T G A T C | GRE(NR),IR3/RAW264.7-GRE-ChIP-Seq(Unpublished)/Homer | 1e-7 | -1.648e+01 | 0.0000 | 3355.0 | 10.39% | 13744.7 | 9.43% | motif file (matrix) | svg |
| 574 | G C T A C G T A A C T G C G T A C G A T A C G T A G T C A G C T | At3g12730(G2like)/colamp-At3g12730-DAP-Seq(GSE60143)/Homer | 1e-7 | -1.641e+01 | 0.0000 | 18058.0 | 55.94% | 79200.0 | 54.33% | motif file (matrix) | svg |
| 575 | G C A T C G T A C G A T G A C T A C T G C T G A G A C T G A T C | Hnf6b(Homeobox)/LNCaP-Hnf6b-ChIP-Seq(GSE106305)/Homer | 1e-6 | -1.594e+01 | 0.0000 | 15600.0 | 48.33% | 68131.3 | 46.74% | motif file (matrix) | svg |
| 576 | A C T G G A T C C T G A A T C G A G T C T A G C C T G A C G T A T A C G A G T C C T A G C A G T T C A G T C G A T G A C G A T C | PAX5(Paired,Homeobox)/GM12878-PAX5-ChIP-Seq(GSE32465)/Homer | 1e-6 | -1.589e+01 | 0.0000 | 7409.0 | 22.95% | 31535.7 | 21.63% | motif file (matrix) | svg |
| 577 | A G T C A T C G G C A T C A T G A C T G A T C G C G A T C T A G A C T G A G C T T G A C G A C T | Gli2(Zf)/GM2-Gli2-ChIP-Chip(GSE112702)/Homer | 1e-6 | -1.586e+01 | 0.0000 | 3720.0 | 11.52% | 15356.3 | 10.53% | motif file (matrix) | svg |
| 578 | C T A G A G T C G A T C G T A C A G T C C T A G A G T C G T A C G A T C G T A C G A T C G C A T | KLF17(Zf)/2cell-Klf17-CutnTag(GSE211845)/Homer | 1e-6 | -1.581e+01 | 0.0000 | 7816.0 | 24.21% | 33342.0 | 22.87% | motif file (matrix) | svg |
| 579 | C T G A A G T C C G T A G T A C A C T G G A C T G C T A C G T A G A C T A G T C | ANAC038(NAC)/col-ANAC038-DAP-Seq(GSE60143)/Homer | 1e-6 | -1.566e+01 | 0.0000 | 25080.0 | 77.69% | 111321.6 | 76.37% | motif file (matrix) | svg |
| 580 | C A T G G A C T G C A T A C T G A G C T A C T G A C T G C G T A G C A T A G C T A T C G T A C G | Foxh1(Forkhead)/hESC-FOXH1-ChIP-Seq(GSE29422)/Homer | 1e-6 | -1.554e+01 | 0.0000 | 9822.0 | 30.43% | 42269.9 | 29.00% | motif file (matrix) | svg |
| 581 | G C A T T A G C G T A C C A T G C T G A G C A T G C A T G C A T G A C T G C A T G A C T G T A C A C T G A T C G C G T A | LBD2(LOBAS2)/colamp-LBD2-DAP-Seq(GSE60143)/Homer | 1e-6 | -1.545e+01 | 0.0000 | 13043.0 | 40.40% | 56673.0 | 38.88% | motif file (matrix) | svg |
| 582 | C G T A G A C T C G A T A T C G G T A C G C A T C A T G C G T A T A C G G C A T G T A C C G T A C A T G A T G C G C T A C T A G G C A T G C A T G C A T G A C T | MafB(bZIP)/BMM-Mafb-ChIP-Seq(GSE75722)/Homer | 1e-6 | -1.529e+01 | 0.0000 | 3687.0 | 11.42% | 15242.1 | 10.46% | motif file (matrix) | svg |
| 583 | C T G A T C A G G C T A A G C T A G T C G A C T C T G A C T A G T G C A C T G A A G T C G T A C G A T C A C T G T C G A | ZBTB12(Zf)/HEK293-ZBTB12.GFP-ChIP-Seq(GSE58341)/Homer | 1e-6 | -1.524e+01 | 0.0000 | 6163.0 | 19.09% | 26081.3 | 17.89% | motif file (matrix) | svg |
| 584 | G C A T G C A T G C T A G C A T G A T C C G A T A C G T C G T A A C G T A G T C G A C T G C A T G C A T G C A T G C A T | At5g58900(MYBrelated)/colamp-At5g58900-DAP-Seq(GSE60143)/Homer | 1e-6 | -1.515e+01 | 0.0000 | 14881.0 | 46.10% | 64959.9 | 44.56% | motif file (matrix) | svg |
| 585 | G T A C T G A C A T G C A G T C A C T G G A T C A T C G G A T C | SUT1?/SacCer-Promoters/Homer | 1e-6 | -1.495e+01 | 0.0000 | 30681.0 | 95.04% | 137540.1 | 94.35% | motif file (matrix) | svg |
| 586 | G A C T G A C T A T C G C G A T G T C A G T A C A G C T C G A T A C G T G T A C | SPL11(SBP)/col100-SPL11-DAP-Seq(GSE60143)/Homer | 1e-6 | -1.475e+01 | 0.0000 | 10913.0 | 33.81% | 47197.3 | 32.38% | motif file (matrix) | svg |
| 587 | A T G C G A C T A G C T A G C T A G T C G C T A C A G T C G A T G C T A A C G T A C T G G C T A T A G C G C A T T G A C | IRF:BATF(IRF:bZIP)/pDC-Irf8-ChIP-Seq(GSE66899)/Homer | 1e-6 | -1.456e+01 | 0.0000 | 883.0 | 2.74% | 3306.2 | 2.27% | motif file (matrix) | svg |
| 588 | C G A T T C G A G T A C A C T G A C G T T C A G G C A T T G C A G C T A G C A T C G T A A G T C C G T A G T A C C A T G | ANAC087(NAC)/col-ANAC087-DAP-Seq(GSE60143)/Homer | 1e-6 | -1.447e+01 | 0.0000 | 7792.0 | 24.14% | 33332.2 | 22.87% | motif file (matrix) | svg |
| 589 | C G T A C A G T G T A C A T G C C T A G C G T A A C G T A G T C T C G A T C A G | GATA19(C2C2gata)/colamp-GATA19-DAP-Seq(GSE60143)/Homer | 1e-6 | -1.437e+01 | 0.0000 | 3152.0 | 9.76% | 12969.9 | 8.90% | motif file (matrix) | svg |
| 590 | C T G A A C G T A T G C G C A T A G T C C G T A A T G C A C G T | AZF1(C2H2)/colamp-AZF1-DAP-Seq(GSE60143)/Homer | 1e-6 | -1.431e+01 | 0.0000 | 28144.0 | 87.18% | 125607.0 | 86.16% | motif file (matrix) | svg |
| 591 | A G C T C T A G T G A C C G T A A C G T C G A T A G T C A G T C C T G A C A T G | TEAD3(TEA)/HepG2-TEAD3-ChIP-Seq(Encode)/Homer | 1e-6 | -1.397e+01 | 0.0000 | 14514.0 | 44.96% | 63410.3 | 43.50% | motif file (matrix) | svg |
| 592 | G T A C G C A T A C G T C G T A A C G T A G T C A G T C C T G A | At5g47390(MYBrelated)/col-At5g47390-DAP-Seq(GSE60143)/Homer | 1e-6 | -1.387e+01 | 0.0000 | 17789.0 | 55.11% | 78201.4 | 53.65% | motif file (matrix) | svg |
| 593 | G C A T A C G T A G T C A T G C A G T C C T A G T A G C G T A C C T G A G C T A | DEL1(E2FDP)/colamp-DEL1-DAP-Seq(GSE60143)/Homer | 1e-5 | -1.361e+01 | 0.0000 | 152.0 | 0.47% | 431.6 | 0.30% | motif file (matrix) | svg |
| 594 | T G A C G T A C C G T A A C T G T G A C C G A T A C T G A T C G A G C T T A C G T C G A T A G C G T A C C G T A A T C G T G A C G C A T A C T G A C T G A T G C | Twist(bHLH)/HMLE-TWIST1-ChIP-Seq(Chang\_et\_al)/Homer | 1e-5 | -1.341e+01 | 0.0000 | 1082.0 | 3.35% | 4164.3 | 2.86% | motif file (matrix) | svg |
| 595 | G A C T G T A C G A C T A G T C T C A G C T G A A G T C A G T C C T A G C G A T A G C T A T G C C T G A C A G T A G C T | AT4G27900(C2C2COlike)/col-AT4G27900-DAP-Seq(GSE60143)/Homer | 1e-5 | -1.277e+01 | 0.0000 | 342.0 | 1.06% | 1159.9 | 0.80% | motif file (matrix) | svg |
| 596 | C T G A G A T C G C A T A C T G C G T A A C G T C G T A C G T A T A C G T C G A | PQM-1(?)/cElegans-L3-ChIP-Seq(modEncode)/Homer | 1e-5 | -1.233e+01 | 0.0000 | 6001.0 | 18.59% | 25568.8 | 17.54% | motif file (matrix) | svg |
| 597 | C T G A C T A G T C G A C G T A A T G C C G T A A T C G C G A T T A G C G C A T A T C G G C A T A G C T G A T C G A C T A G C T | ARE(NR)/LNCAP-AR-ChIP-Seq(GSE27824)/Homer | 1e-5 | -1.190e+01 | 0.0000 | 2539.0 | 7.87% | 10440.8 | 7.16% | motif file (matrix) | svg |
| 598 | G A T C G C T A C A G T A C G T T A C G A G T C A T G C C T A G A G T C T C G A | Zfp57(Zf)/H1-ZFP57.HA-ChIP-Seq(GSE115387)/Homer | 1e-5 | -1.171e+01 | 0.0000 | 15072.0 | 46.69% | 66134.0 | 45.37% | motif file (matrix) | svg |
| 599 | C G T A G C T A C G A T A C G T A G C T A G C T C T G A G T C A C G T A G C T A | Unknown6/Drosophila-Promoters/Homer | 1e-5 | -1.167e+01 | 0.0000 | 3695.0 | 11.45% | 15480.2 | 10.62% | motif file (matrix) | svg |
| 600 | C G A T C A G T G T C A G C T A C A G T A G C T A C G T A C T G A G T C C G T A A C G T A C T G A C G T C T G A T C G A | FUS3(ABI3VP1)/col-FUS3-DAP-Seq(GSE60143)/Homer | 1e-4 | -1.132e+01 | 0.0000 | 12815.0 | 39.70% | 56023.0 | 38.43% | motif file (matrix) | svg |
| 601 | T G C A T G C A A G T C A G T C G A C T C A G T A T G C G A T C C T G A A C G T C T A G C T A G A G T C A C G T A G T C A G T C A G T C G A C T C G T A A C G T A G C T C T A G G A T C G A T C G A T C | ZNF16(Zf)/HEK293-ZNF16.GFP-ChIP-Seq(GSE58341)/Homer | 1e-4 | -1.124e+01 | 0.0000 | 65.0 | 0.20% | 151.3 | 0.10% | motif file (matrix) | svg |
| 602 | C T G A C T G A T A G C G A T C G C T A G T A C A C G T G A T C T G C A C G T A | Nkx2.5(Homeobox)/HL1-Nkx2.5.biotin-ChIP-Seq(GSE21529)/Homer | 1e-4 | -1.105e+01 | 0.0000 | 22066.0 | 68.36% | 97898.9 | 67.16% | motif file (matrix) | svg |
| 603 | T C A G G T C A G C T A A G T C C G T A A T C G T C G A T C A G C G T A A C T G A C T G A C T G | ZNF768(Zf)/Rajj-ZNF768-ChIP-Seq(GSE111879)/Homer | 1e-4 | -1.104e+01 | 0.0000 | 276.0 | 0.85% | 929.4 | 0.64% | motif file (matrix) | svg |
| 604 | G C T A G A C T A G T C T C G A T C A G T C G A A C G T A G T C G A C T T C A G | GATA14(C2C2gata)/col-GATA14-DAP-Seq(GSE60143)/Homer | 1e-4 | -1.103e+01 | 0.0000 | 11287.0 | 34.96% | 49199.5 | 33.75% | motif file (matrix) | svg |
| 605 | T C A G T G A C G T A C T G C A G T A C C T A G G T A C A T G C A G T C G T C A A G T C G A C T | Klf9(Zf)/GBM-Klf9-ChIP-Seq(GSE62211)/Homer | 1e-4 | -1.101e+01 | 0.0000 | 3526.0 | 10.92% | 14782.7 | 10.14% | motif file (matrix) | svg |
| 606 | T G C A T C G A T A G C G T A C T C A G C T A G G T C A G C T A T C A G G A C T | ETS(ETS)/Promoter/Homer | 1e-4 | -1.078e+01 | 0.0000 | 7069.0 | 21.90% | 30416.2 | 20.87% | motif file (matrix) | svg |
| 607 | G A C T A G C T G T A C G A C T C T G A A C T G G T C A C T G A A T G C T A C G G A C T A C G T A G T C G A C T C T G A | HRE(HSF)/Striatum-HSF1-ChIP-Seq(GSE38000)/Homer | 1e-4 | -1.069e+01 | 0.0000 | 1648.0 | 5.11% | 6660.9 | 4.57% | motif file (matrix) | svg |
| 608 | G C A T A C G T A G C T A G C T A G C T C G T A A G T C A C G T | At3g60580(C2H2)/col-At3g60580-DAP-Seq(GSE60143)/Homer | 1e-4 | -1.045e+01 | 0.0000 | 28298.0 | 87.66% | 126579.0 | 86.83% | motif file (matrix) | svg |
| 609 | C T A G A C G T A G T C C G T A A C T G A G T C G C A T A C T G G C A T A G T C G A C T G A T C G C A T A G T C A G C T | ZNF317(Zf)/HEK293-ZNF317.GFP-ChIP-Seq(GSE58341)/Homer | 1e-4 | -1.035e+01 | 0.0001 | 1224.0 | 3.79% | 4865.6 | 3.34% | motif file (matrix) | svg |
| 610 | T C G A G A C T T C A G T G C A G T A C G T A C A G C T G T A C C A T G T C G A C A T G C A T G A C G T A G T C C T G A | FXR(NR),ER2/Liver-FXR-ChIP-Seq(GSE133700)/Homer | 1e-4 | -1.014e+01 | 0.0001 | 7373.0 | 22.84% | 31821.1 | 21.83% | motif file (matrix) | svg |
| 611 | A T G C G A T C C G A T C T A G A C T G G C T A C G T A A G C T A C T G A G C T | TEAD2(TEA)/Py2T-Tead2-ChIP-Seq(GSE55709)/Homer | 1e-4 | -9.863e+00 | 0.0001 | 6222.0 | 19.27% | 26738.5 | 18.34% | motif file (matrix) | svg |
| 612 | G C A T A C G T A T G C C T G A C T A G G A C T G A T C A C T G | Initiator/Drosophila-Promoters/Homer | 1e-4 | -9.739e+00 | 0.0001 | 21575.0 | 66.83% | 95792.2 | 65.71% | motif file (matrix) | svg |
| 613 | G T A C G A T C C T A G C G A T G T C A G T A C C T A G C A G T G C A T G A C T | SPL1(SBP)/colamp-SPL1-DAP-Seq(GSE60143)/Homer | 1e-4 | -9.686e+00 | 0.0001 | 24738.0 | 76.63% | 110240.3 | 75.62% | motif file (matrix) | svg |
| 614 | A G C T G A C T C T A G C G T A C A T G C G A T C T A G A T C G G A C T C A G T | Bapx1(Homeobox)/VertebralCol-Bapx1-ChIP-Seq(GSE36672)/Homer | 1e-4 | -9.646e+00 | 0.0001 | 21376.0 | 66.22% | 94896.5 | 65.10% | motif file (matrix) | svg |
| 615 | A G T C C G T A C G A T A G T C G T C A A G T C A C G T C T G A | Unknown2/Drosophila-Promoters/Homer | 1e-4 | -9.330e+00 | 0.0002 | 13269.0 | 41.10% | 58268.1 | 39.97% | motif file (matrix) | svg |
| 616 | G C A T G C A T G T A C G A C T T C G A A C T G G C T A C G T A A T C G T G A C G C A T G A C T A G T C A G C T C T G A | AGL95(ND)/col-AGL95-DAP-Seq(GSE60143)/Homer | 1e-3 | -8.846e+00 | 0.0002 | 889.0 | 2.75% | 3501.3 | 2.40% | motif file (matrix) | svg |
| 617 | G C T A G C T A T G C A A G T C C T A G C T G A G A T C C T A G G A C T G A T C C T A G A C G T C G A T C G A T G A C T | Unknown2/Arabidopsis-Promoters/Homer | 1e-3 | -8.678e+00 | 0.0003 | 502.0 | 1.56% | 1888.1 | 1.30% | motif file (matrix) | svg |
| 618 | G C A T C G T A C G A T A C T G A G T C G C T A C T G A C G T A C A G T A C T G C G T A T C A G | Oct6(POU,Homeobox)/NPC-Pou3f1-ChIP-Seq(GSE35496)/Homer | 1e-3 | -8.416e+00 | 0.0004 | 3809.0 | 11.80% | 16198.0 | 11.11% | motif file (matrix) | svg |
| 619 | G T C A C T G A T C A G G A T C T G C A T G C A A C G T T C A G C G T A C G T A C G T A G C T A | Hoxd12(Homeobox)/ChickenMSG-Hoxd12.Flag-ChIP-Seq(GSE86088)/Homer | 1e-3 | -8.258e+00 | 0.0004 | 16217.0 | 50.24% | 71674.4 | 49.17% | motif file (matrix) | svg |
| 620 | C T G A C T A G C A T G C T A G A C T G A C T G G A T C A C T G A C T G C T A G T C A G A G T C | Klf15(Zf)/Liver-Klf15-ChIP-Seq(GSE166083)/Homer | 1e-3 | -7.877e+00 | 0.0006 | 13292.0 | 41.18% | 58539.3 | 40.16% | motif file (matrix) | svg |
| 621 | G C A T G C T A G C T A C G T A G C A T G C T A C T A G C G T A C G T A A C T G C G T A C G A T A C G T A G T C G A C T | At1g68670(G2like)/colamp-At1g68670-DAP-Seq(GSE60143)/Homer | 1e-3 | -7.839e+00 | 0.0007 | 5024.0 | 15.56% | 21607.9 | 14.82% | motif file (matrix) | svg |
| 622 | G C T A G A C T G A C T T G C A C G T A A G T C C G T A T A G C G A T C G A C T | Eomes(T-box)/H9-Eomes-ChIP-Seq(GSE26097)/Homer | 1e-3 | -7.824e+00 | 0.0007 | 19617.0 | 60.77% | 87112.1 | 59.76% | motif file (matrix) | svg |
| 623 | T C G A G T A C T C G A T C G A C A T G A T G C A C G T A C T G A C T G A G T C C G T A C T A G A G T C A T C G A G T C | Unknown3/Drosophila-Promoters/Homer | 1e-3 | -7.715e+00 | 0.0008 | 1509.0 | 4.67% | 6200.7 | 4.25% | motif file (matrix) | svg |
| 624 | A G C T C G T A C G T A A G T C C T A G C T A G G T A C G A C T | MYB101(MYB)/colamp-MYB101-DAP-Seq(GSE60143)/Homer | 1e-3 | -7.632e+00 | 0.0008 | 24302.0 | 75.28% | 108455.1 | 74.40% | motif file (matrix) | svg |
| 625 | G C A T G A T C A G C T G T A C G A T C C T A G C T A G G A T C T A C G C T G A | AT3G58630(Trihelix)/col-AT3G58630-DAP-Seq(GSE60143)/Homer | 1e-3 | -7.350e+00 | 0.0011 | 6343.0 | 19.65% | 27505.3 | 18.87% | motif file (matrix) | svg |
| 626 | C G A T C G T A A C T G C G T A A C G T C G T A A C G T A C G T C G A T G C A T C G A T C G A T | AT2G28920(ND)/col-AT2G28920-DAP-Seq(GSE60143)/Homer | 1e-2 | -6.448e+00 | 0.0027 | 2345.0 | 7.26% | 9913.9 | 6.80% | motif file (matrix) | svg |
| 627 | C T A G T A C G A G T C G T A C T C G A A C G T G T C A G C T A G C T A C G A T A G T C G C T A | HOXA9(Homeobox)/HSC-Hoxa9-ChIP-Seq(GSE33509)/Homer | 1e-2 | -6.364e+00 | 0.0029 | 9405.0 | 29.13% | 41283.9 | 28.32% | motif file (matrix) | svg |
| 628 | T G C A A G C T C T G A A T C G G A C T C T A G G T A C G A T C G T C A A G T C G T A C G A C T C T A G A T C G G C A T C A T G C A T G G A T C G T A C C T G A | CTCF(Zf)/CD4+-CTCF-ChIP-Seq(Barski\_et\_al.)/Homer | 1e-2 | -6.280e+00 | 0.0031 | 1065.0 | 3.30% | 4356.4 | 2.99% | motif file (matrix) | svg |
| 629 | C T G A C A T G A C T G A C G T A T G C C G T A C A T G T A C G A T G C G C T A T A C G C T G A C T A G A C T G A C G T A T G C C G T A T A G C | RAR:RXR(NR),DR5/ES-RAR-ChIP-Seq(GSE56893)/Homer | 1e-2 | -6.145e+00 | 0.0036 | 267.0 | 0.83% | 985.3 | 0.68% | motif file (matrix) | svg |
| 630 | C G T A C G T A C T G A A C T G C G T A C G T A A C G T C T A G C G A T C G A T | AT2G38300(G2like)/col-AT2G38300-DAP-Seq(GSE60143)/Homer | 1e-2 | -6.124e+00 | 0.0036 | 12279.0 | 38.04% | 54211.0 | 37.19% | motif file (matrix) | svg |
| 631 | C G T A C G T A G C T A C G T A G A T C C T G A A C G T A C G T A G T C A G C T G C A T G C A T | AT2G40260(G2like)/colamp-AT2G40260-DAP-Seq(GSE60143)/Homer | 1e-2 | -6.034e+00 | 0.0040 | 14837.0 | 45.96% | 65739.6 | 45.10% | motif file (matrix) | svg |
| 632 | A G C T G A T C G A C T A G T C T C G A C T G A A G T C A G T C C T A G A G C T G A C T T A G C T C G A C G A T G A C T | AT5G59990(C2C2COlike)/colamp-AT5G59990-DAP-Seq(GSE60143)/Homer | 1e-2 | -5.700e+00 | 0.0056 | 1026.0 | 3.18% | 4216.2 | 2.89% | motif file (matrix) | svg |
| 633 | A T G C A T C G A T G C A T C G A T G C A T C G A T G C A T C G A T G C A T C G | SeqBias: CG-repeat | 1e-2 | -5.639e+00 | 0.0059 | 22903.0 | 70.95% | 102322.4 | 70.19% | motif file (matrix) | svg |
| 634 | C G T A C G T A C G T A C G T A C G T A C G T A C G T A C G T A C G T A C G T A | SeqBias: polyA-repeat | 1e-2 | -5.620e+00 | 0.0060 | 32260.0 | 99.93% | 145602.9 | 99.88% | motif file (matrix) | svg |
| 635 | G C A T C A G T C G A T A G T C G A T C G C T A C G A T C G A T C G A T G C T A C G A T C T A G A C T G G C T A G C T A | AGL25(MADS)/colamp-AGL25-DAP-Seq(GSE60143)/Homer | 1e-2 | -5.475e+00 | 0.0069 | 326.0 | 1.01% | 1245.0 | 0.85% | motif file (matrix) | svg |
| 636 | G C T A A G C T G T A C G C A T A G C T T C G A C T G A A G T C A G T C T A C G A C G T G A C T T A C G C T A G C G T A | ZML1(C2C2gata)/colamp-ZML1-DAP-Seq(GSE60143)/Homer | 1e-2 | -5.415e+00 | 0.0073 | 943.0 | 2.92% | 3872.9 | 2.66% | motif file (matrix) | svg |
| 637 | A G T C C G T A A C G T G T A C A G T C A G T C C G T A A C G T C G T A C G T A C G A T T G C A G T A C A G C T A T G C | ZNF410(Zf)/CD34-ZNF410-ChIP-Seq(GSE154960)/Homer | 1e-2 | -5.371e+00 | 0.0077 | 34.0 | 0.11% | 87.7 | 0.06% | motif file (matrix) | svg |
| 638 | G A C T C G A T C T G A G T C A G A C T C G A T T C G A C G T A G C T A G C T A T G A C G T A C C G T A A C T G T G C A C G A T A C T G A C G T | Pitx1:Ebox(Homeobox,bHLH)/Hindlimb-Pitx1-ChIP-Seq(GSE41591)/Homer | 1e-2 | -4.911e+00 | 0.0121 | 1343.0 | 4.16% | 5635.5 | 3.87% | motif file (matrix) | svg |
| 639 | A G C T T A C G C G A T T C A G A G C T A C G T G A T C C T G A T A G C A G C T A T G C C T G A C G T A A T C G A G T C C T A G A C T G C T G A G T C A T C G A | PAX6(Paired,Homeobox)/Forebrain-Pax6-ChIP-Seq(GSE66961)/Homer | 1e-2 | -4.804e+00 | 0.0135 | 1800.0 | 5.58% | 7641.4 | 5.24% | motif file (matrix) | svg |
| 640 | A C T G G T C A A C G T G C A T C G A T T C A G G T A C G T C A A C G T C T G A | Oct11(POU,Homeobox)/NCIH1048-POU2F3-ChIP-seq(GSE115123)/Homer | 1e-2 | -4.729e+00 | 0.0145 | 3195.0 | 9.90% | 13798.4 | 9.47% | motif file (matrix) | svg |
