## Supplemental dataset for "Hybrid CNN and Multi-Head Attention Model for Analyzing Epigenetic Mechanisms and Gene Expression Across Fungal Phylogenetic Distances": FgramModel_NcrassaTest_K4me3locs_homerResults.html

/projects/wg-feeds/SHAP/FgramModel\_NcrassaTest\_K4me3locs\_SHAP\_noDup\_HOMER// - Homer de novo Motif Results


### Homer *de novo* Motif Results (/projects/wg-feeds/SHAP/FgramModel\_NcrassaTest\_K4me3locs\_SHAP\_noDup\_HOMER//)

Non-redundant Motif File of Results  
Known Motif Enrichment Results  
Gene Ontology Enrichment Results  
If Homer is having trouble matching a motif to a known motif, try copy/pasting the matrix file into
STAMP  
More information on motif finding results: HOMER
| Description of Results
| Tips
  
Total target sequences = 39596  
Total background sequences = 181025  
\* - possible false positive  

|  |  |  |  |  |  |  |  |  |
| --- | --- | --- | --- | --- | --- | --- | --- | --- |
| Rank | Motif | P-value | log P-pvalue | % of Targets | % of Background | STD(Bg STD) | Best Match/Details | Motif File |
| 1 | G C A T A G C T A T C G G C T A A G C T A C T G G C T A A G C T A T C G G T C A A G C T A C T G | 1e-2660 | -6.126e+03 | 41.28% | 15.07% | 224.7bp (232.3bp) | ZML2(C2C2gata)/col-ZML2-DAP-Seq(GSE60143)/Homer(0.795) More Information \| Similar Motifs Found | motif file (matrix) |
| 2 | T A G C T C G A C G T A A T C G T C A G G C A T A T G C T C G A C G T A T A C G | 1e-2224 | -5.122e+03 | 51.33% | 24.78% | 227.6bp (236.4bp) | NR6A1/MA1541.2/Jaspar(0.785) More Information \| Similar Motifs Found | motif file (matrix) |
| 3 | C G T A C A G T A T G C G C T A C G A T A T G C G C T A C G A T A T G C G C T A C G A T A T C G | 1e-1999 | -4.605e+03 | 64.15% | 37.74% | 226.2bp (235.4bp) | ZML2(C2C2gata)/col-ZML2-DAP-Seq(GSE60143)/Homer(0.600) More Information \| Similar Motifs Found | motif file (matrix) |
| 4 | A G C T A G T C A G C T A G C T A G T C A G C T G A C T A T G C | 1e-1922 | -4.426e+03 | 45.31% | 21.47% | 237.3bp (234.2bp) | Unknown4/Arabidopsis-Promoters/Homer(0.842) More Information \| Similar Motifs Found | motif file (matrix) |
| 5 | G A C T A G T C T G C A C G T A T A C G T C G A C G T A T A C G | 1e-1643 | -3.784e+03 | 57.62% | 33.86% | 227.9bp (236.8bp) | MATR3(RRM)/Homo\_sapiens-RNCMPT00037-PBM/HughesRNA(0.782) More Information \| Similar Motifs Found | motif file (matrix) |
| 6 | A G C T A G C T A G C T A G C T A G C T A G C T A G C T A G C T A G C T A G C T A G C T A G C T | 1e-1612 | -3.713e+03 | 32.44% | 13.33% | 234.4bp (212.8bp) | SeqBias: G/A bias(0.849) More Information \| Similar Motifs Found | motif file (matrix) |
| 7 | T A G C C G T A T C A G T C A G C G T A T C A G G T A C C G A T A T C G T A C G T G C A T C A G | 1e-1241 | -2.858e+03 | 52.51% | 32.06% | 226.4bp (232.3bp) | ZFP14/MA1972.1/Jaspar(0.676) More Information \| Similar Motifs Found | motif file (matrix) |
| 8 | T A G C G T A C C G A T A C G T A C T G G C T A G A C T T C A G T C G A G A T C | 1e-1152 | -2.653e+03 | 52.17% | 32.46% | 225.3bp (239.4bp) | Aef1/dmmpmm(Bergman)/fly(0.721) More Information \| Similar Motifs Found | motif file (matrix) |
| 9 | G C A T A C G T A C T G A T G C A G T C C T A G C A G T A G T C G A C T A C G T A C T G T A C G | 1e-1108 | -2.553e+03 | 59.58% | 39.88% | 235.4bp (234.2bp) | pan/MA0237.2/Jaspar(0.636) More Information \| Similar Motifs Found | motif file (matrix) |
| 10 | C T A G T A C G G T A C G T C A C T G A T A C G T G C A G T C A C T G A C T G A | 1e-870 | -2.005e+03 | 47.30% | 30.41% | 230.6bp (232.9bp) | NAC101/MA2050.2/Jaspar(0.818) More Information \| Similar Motifs Found | motif file (matrix) |
| 11 | A G T C G T C A C G T A C G A T A C T G C A G T A G T C C T A G | 1e-776 | -1.788e+03 | 59.12% | 42.60% | 236.2bp (236.5bp) | Hr39/MA2244.1/Jaspar(0.762) More Information \| Similar Motifs Found | motif file (matrix) |
| 12 | T A G C A C G T C G T A A G T C G T A C C G A T A T G C A C G T C G T A A T G C | 1e-619 | -1.427e+03 | 4.30% | 0.51% | 217.7bp (200.5bp) | vfl/MA1462.2/Jaspar(0.677) More Information \| Similar Motifs Found | motif file (matrix) |
| 13 | G C A T T C G A A T G C T A G C A C G T C G T A A T G C T A G C G A C T G C T A | 1e-583 | -1.342e+03 | 4.24% | 0.55% | 210.5bp (198.8bp) | PK06182.1/MA2354.1/Jaspar(0.795) More Information \| Similar Motifs Found | motif file (matrix) |
| 14 | A G T C C G T A G C A T A G T C T A C G C T G A G C A T G C T A A G C T A G T C | 1e-469 | -1.082e+03 | 28.95% | 18.25% | 234.7bp (229.7bp) | OsCCA1/MA2349.1/Jaspar(0.748) More Information \| Similar Motifs Found | motif file (matrix) |
| 15 | A C T G A C G T A G T C G T C A G C T A C T A G C A G T A G T C | 1e-445 | -1.026e+03 | 37.14% | 25.66% | 230.6bp (240.3bp) | SNRNP70(RRM)/Homo\_sapiens-RNCMPT00070-PBM/HughesRNA(0.790) More Information \| Similar Motifs Found | motif file (matrix) |
| 16 | C T A G A G C T C T A G A C G T C T A G A C G T C T A G A C G T T C A G A G C T C T A G A G C T | 1e-209 | -4.825e+02 | 1.80% | 0.30% | 247.4bp (232.9bp) | cg/MA2107.1/Jaspar(0.895) More Information \| Similar Motifs Found | motif file (matrix) |
| 17 | A C G T C G T A A T G C G C A T T A C G A C G T C T G A A T C G C G A T T C A G A C G T C G T A | 1e-184 | -4.238e+02 | 2.49% | 0.68% | 229.0bp (214.3bp) | SFPQ(RRM)/Homo\_sapiens-RNCMPT00177-PBM/HughesRNA(0.806) More Information \| Similar Motifs Found | motif file (matrix) |
| 18 | C T A G A C T G A C T G A C T G A C T G A C T G A C T G A C T G A C T G A C T G A C T G C T A G | 1e-138 | -3.187e+02 | 0.70% | 0.04% | 228.9bp (154.7bp) | SeqBias: polyC-repeat(0.923) More Information \| Similar Motifs Found | motif file (matrix) |
| 19 | A C T G C G T A A C T G C G T A A C T G C G T A A C T G C G T A A C T G C G T A | 1e-121 | -2.796e+02 | 0.94% | 0.13% | 275.4bp (211.0bp) | SeqBias: GA-repeat(0.994) More Information \| Similar Motifs Found | motif file (matrix) |
| 20 | C T G A T G C A A T G C A G T C G T A C G A C T C G T A C G T A A G T C A G T C A T G C A G C T | 1e-105 | -2.429e+02 | 0.64% | 0.06% | 284.2bp (206.4bp) | TBF1/MA0403.3/Jaspar(0.811) More Information \| Similar Motifs Found | motif file (matrix) |
| 21 | A C T G G C A T G A T C C G A T C A T G G C A T G A T C C G T A A C T G A C G T | 1e-67 | -1.546e+02 | 2.03% | 0.93% | 245.6bp (232.2bp) | HOW(KH)/Drosophila\_melanogaster-RNCMPT00118-PBM/HughesRNA(0.723) More Information \| Similar Motifs Found | motif file (matrix) |
| 22 | C G A T G A C T A G T C G C T A C A T G A C G T A C G T A G T C G C T A T C A G C A G T G A C T | 1e-34 | -7.997e+01 | 0.25% | 0.03% | 251.1bp (277.1bp) | IRF9/MA0653.1/Jaspar(0.703) More Information \| Similar Motifs Found | motif file (matrix) |
| 23 | A G T C A G T C C T A G C G T A A G T C A G T C A C G T C G T A A G T C A G T C A C T G C G T A | 1e-31 | -7.200e+01 | 0.15% | 0.01% | 222.7bp (262.0bp) | MYB55/MA1041.1/Jaspar(0.789) More Information \| Similar Motifs Found | motif file (matrix) |
| 24 | C G A T C G A T C G T A G C T A A C G T C G T A G C T A C G A T C G T A C G T A C G A T C G T A | 1e-30 | -7.102e+01 | 0.20% | 0.02% | 270.8bp (170.4bp) | Ubx/dmmpmm(Down)/fly(0.799) More Information \| Similar Motifs Found | motif file (matrix) |
| 25 | A C G T A C T G A G T C A G T C A G T C A C G T A C T G A G T C A T G C A G T C A C G T A C T G | 1e-29 | -6.875e+01 | 0.25% | 0.04% | 212.1bp (208.8bp) | THAP1/MA0597.3/Jaspar(0.760) More Information \| Similar Motifs Found | motif file (matrix) |
| 26 | A G T C A G T C A C G T A C T G A G T C C G T A A C T G A G C T C G T A A G T C C G A T A C G T | 1e-14 | -3.452e+01 | 0.10% | 0.01% | 548.2bp (196.5bp) | WIP5/MA2367.1/Jaspar(0.720) More Information \| Similar Motifs Found | motif file (matrix) |
