## Supplemental dataset for "Hybrid CNN and Multi-Head Attention Model for Analyzing Epigenetic Mechanisms and Gene Expression Across Fungal Phylogenetic Distances": FgramModel_NcrassaTest_K4me3locs_knownResults.html

/projects/wg-feeds/SHAP/FgramModel\_NcrassaTest\_K4me3locs\_SHAP\_noDup\_HOMER/ - Homer Known Motif Enrichment Results


### Homer Known Motif Enrichment Results (/projects/wg-feeds/SHAP/FgramModel\_NcrassaTest\_K4me3locs\_SHAP\_noDup\_HOMER/)

Homer *de novo* Motif Results  
Gene Ontology Enrichment Results  
Known Motif Enrichment Results (txt file)  
Total Target Sequences = 39599, Total Background Sequences = 180966

|  |  |  |  |  |  |  |  |  |  |  |  |
| --- | --- | --- | --- | --- | --- | --- | --- | --- | --- | --- | --- |
| Rank | Motif | Name | P-value | log P-pvalue | q-value (Benjamini) | # Target Sequences with Motif | % of Targets Sequences with Motif | # Background Sequences with Motif | % of Background Sequences with Motif | Motif File | SVG |
| 1 | A T G C C A G T A C G T A G T C C A T G A C G T A G T C A C G T A G C T G A T C | Unknown4/Arabidopsis-Promoters/Homer | 1e-1323 | -3.047e+03 | 0.0000 | 12735.0 | 32.16% | 26421.1 | 14.60% | motif file (matrix) | svg |
| 2 | T A G C G T C A G A C T T A G C G T C A G A C T A G T C G C T A G A C T G A T C | ZML2(C2C2gata)/col-ZML2-DAP-Seq(GSE60143)/Homer | 1e-1216 | -2.800e+03 | 0.0000 | 4840.0 | 12.22% | 4693.1 | 2.59% | motif file (matrix) | svg |
| 3 | C T A G T C A G C T G A C T G A C T A G C T G A C A T G C A T G C T G A C T A G C T A G C G T A C T A G C G T A G T C A | TF3A(C2H2)/col-TF3A-DAP-Seq(GSE60143)/Homer | 1e-1017 | -2.343e+03 | 0.0000 | 10359.0 | 26.16% | 21571.7 | 11.92% | motif file (matrix) | svg |
| 4 | A C G T A C G T A C G T A C G T A C G T A C G T A C G T A C G T A C G T A C G T | VRN1(ABI3VP1)/col-VRN1-DAP-Seq(GSE60143)/Homer | 1e-847 | -1.953e+03 | 0.0000 | 1593.0 | 4.02% | 327.9 | 0.18% | motif file (matrix) | svg |
| 5 | G C T A G C A T A C G T A C G T A G C T G A T C G A C T G A C T A G C T A C G T A C G T A G C T | RLR1?/SacCer-Promoters/Homer | 1e-811 | -1.868e+03 | 0.0000 | 3422.0 | 8.64% | 3470.5 | 1.92% | motif file (matrix) | svg |
| 6 | A C G T A T G C A C G T A C G T A G C T A G T C A G C T A G C T A G C T A G C T A G C T | hTCT(CPE) | 1e-702 | -1.619e+03 | 0.0000 | 14275.0 | 36.05% | 39878.4 | 22.03% | motif file (matrix) | svg |
| 7 | G A T C G C A T A G T C A G C T G A T C A G C T G A T C G A C T G A T C G A C T A G T C A C G T G A T C A G C T G A T C | GAGA-repeat/SacCer-Promoters/Homer | 1e-674 | -1.554e+03 | 0.0000 | 20238.0 | 51.11% | 64997.9 | 35.91% | motif file (matrix) | svg |
| 8 | C T A G C T G A C T A G C T G A C T A G C T G A C T A G C T G A C T A G C T G A | SeqBias: GA-repeat | 1e-562 | -1.295e+03 | 0.0000 | 29633.0 | 74.84% | 111492.3 | 61.60% | motif file (matrix) | svg |
| 9 | G T C A T G C A T G C A G C T A C G T A G C T A G C T A G C T A | REM19(REM)/colamp-REM19-DAP-Seq(GSE60143)/Homer | 1e-508 | -1.170e+03 | 0.0000 | 3715.0 | 9.38% | 6008.0 | 3.32% | motif file (matrix) | svg |
| 10 | T G A C C T A G T C A G G T C A C G T A T C A G C G A T T C A G T C G A T G C A C T G A T A G C | PU.1-IRF(ETS:IRF)/Bcell-PU.1-ChIP-Seq(GSE21512)/Homer | 1e-433 | -9.974e+02 | 0.0000 | 7595.0 | 19.18% | 19296.8 | 10.66% | motif file (matrix) | svg |
| 11 | C T G A C G A T C A T G A T C G G C A T C A T G G C T A A G T C | ASHR1(ND)/col-ASHR1-DAP-Seq(GSE60143)/Homer | 1e-420 | -9.673e+02 | 0.0000 | 10596.0 | 26.76% | 30625.4 | 16.92% | motif file (matrix) | svg |
| 12 | A C G T G A C T T A G C C G T A C T G A C A T G C T A G G A C T G A T C C G T A | Nr5a2(NR)/Pancreas-LRH1-ChIP-Seq(GSE34295)/Homer | 1e-404 | -9.304e+02 | 0.0000 | 5740.0 | 14.50% | 13311.5 | 7.35% | motif file (matrix) | svg |
| 13 | G A T C G C T A G T A C A G T C G C T A T G C A G T A C G A T C C G T A G A C T | MYB83(MYB)/colamp-MYB83-DAP-Seq(GSE60143)/Homer | 1e-398 | -9.182e+02 | 0.0000 | 15239.0 | 38.49% | 49706.5 | 27.46% | motif file (matrix) | svg |
| 14 | G T A C A C T G A G T C A G T C C T A G G A T C G T A C C T G A | CRF4(AP2EREBP)/colamp-CRF4-DAP-Seq(GSE60143)/Homer | 1e-388 | -8.945e+02 | 0.0000 | 8111.0 | 20.48% | 21875.5 | 12.09% | motif file (matrix) | svg |
| 15 | C G T A G A C T C A T G C T A G A G T C A C T G A C T G G T A C C A T G T A C G | ERF3(AP2EREBP)/colamp-ERF3-DAP-Seq(GSE60143)/Homer | 1e-385 | -8.877e+02 | 0.0000 | 9780.0 | 24.70% | 28165.5 | 15.56% | motif file (matrix) | svg |
| 16 | C T G A G C A T A C T G C T A G A G T C A C T G A C T G A G T C A C T G T C A G | AT4G18450(AP2EREBP)/col-AT4G18450-DAP-Seq(GSE60143)/Homer | 1e-377 | -8.700e+02 | 0.0000 | 5760.0 | 14.55% | 13733.0 | 7.59% | motif file (matrix) | svg |
| 17 | C T A G G C A T A C T G C T A G A G T C A C T G A C T G A G T C A C T G T C A G | ERF10(AP2EREBP)/col-ERF10-DAP-Seq(GSE60143)/Homer | 1e-376 | -8.678e+02 | 0.0000 | 9383.0 | 23.70% | 26822.7 | 14.82% | motif file (matrix) | svg |
| 18 | A C G T G A C T A T G C G C T A C T G A C T A G A C T G G A C T A G T C C G T A | Nr5a2(NR)/mES-Nr5a2-ChIP-Seq(GSE19019)/Homer | 1e-365 | -8.419e+02 | 0.0000 | 4453.0 | 11.25% | 9580.0 | 5.29% | motif file (matrix) | svg |
| 19 | C G T A G A C T C A T G C T A G A G T C A C T G C T A G A G T C C A T G C T A G | ERF7(AP2EREBP)/col-ERF7-DAP-Seq(GSE60143)/Homer | 1e-354 | -8.165e+02 | 0.0000 | 16319.0 | 41.21% | 55351.4 | 30.58% | motif file (matrix) | svg |
| 20 | G C A T G A T C C T A G C G T A G C A T C G T A G C A T A G T C C T A G C G T A G C A T C G A T | AT5G22990(C2H2)/col-AT5G22990-DAP-Seq(GSE60143)/Homer | 1e-352 | -8.114e+02 | 0.0000 | 6645.0 | 16.78% | 17187.4 | 9.50% | motif file (matrix) | svg |
| 21 | C A T G A G T C G T A C A C T G A T G C A G T C C A T G G A T C G A T C C T G A | ERF5(AP2EREBP)/colamp-ERF5-DAP-Seq(GSE60143)/Homer | 1e-350 | -8.065e+02 | 0.0000 | 5903.0 | 14.91% | 14619.9 | 8.08% | motif file (matrix) | svg |
| 22 | G A T C C A G T T A G C A G T C A C T G A G T C A G T C C T A G G A C T G T A C | LEP(AP2EREBP)/col-LEP-DAP-Seq(GSE60143)/Homer | 1e-348 | -8.032e+02 | 0.0000 | 4499.0 | 11.36% | 9930.0 | 5.49% | motif file (matrix) | svg |
| 23 | G C A T A C T G C T A G A G T C A C T G A C T G A G T C A C G T | ERF105(AP2EREBP)/colamp-ERF105-DAP-Seq(GSE60143)/Homer | 1e-348 | -8.025e+02 | 0.0000 | 14260.0 | 36.01% | 46855.7 | 25.89% | motif file (matrix) | svg |
| 24 | T G A C C G T A C T G A A C T G A C T G G A C T G A T C T G C A G T A C T A C G | SF1(NR)/H295R-Nr5a1-ChIP-Seq(GSE44220)/Homer | 1e-346 | -7.968e+02 | 0.0000 | 3650.0 | 9.22% | 7288.2 | 4.03% | motif file (matrix) | svg |
| 25 | C T G A T C A G G T A C G C T A A C T G T G A C G C A T C A T G | SCL(bHLH)/HPC7-Scl-ChIP-Seq(GSE13511)/Homer | 1e-345 | -7.957e+02 | 0.0000 | 20567.0 | 51.94% | 74147.3 | 40.97% | motif file (matrix) | svg |
| 26 | A C G T A C T G C G T A A C G T A C T G A C T G C G T A C G T A | HAP3(CCAATHAP3)/col-HAP3-DAP-Seq(GSE60143)/Homer | 1e-331 | -7.630e+02 | 0.0000 | 3961.0 | 10.00% | 8418.9 | 4.65% | motif file (matrix) | svg |
| 27 | A T G C G T A C A G T C A G T C A C G T A C G T C G A T A C G T | AT5G02460(C2C2dof)/col-AT5G02460-DAP-Seq(GSE60143)/Homer | 1e-322 | -7.421e+02 | 0.0000 | 13363.0 | 33.75% | 43805.5 | 24.20% | motif file (matrix) | svg |
| 28 | C G A T A C G T A C G T A G C T A G C T G A T C G A T C G C T A A G C T A C G T A T C G T A C G | NFATC2(RHD)/Islets-NFATC2-ChIP-Seq(GSE158496)/Homer | 1e-310 | -7.158e+02 | 0.0000 | 10902.0 | 27.53% | 34117.7 | 18.85% | motif file (matrix) | svg |
| 29 | A T G C T C G A T A C G A C G T A T G C A G T C A C G T A G T C A G T C G A T C | Znf263(Zf)/K562-Znf263-ChIP-Seq(GSE31477)/Homer | 1e-309 | -7.133e+02 | 0.0000 | 10987.0 | 27.75% | 34481.8 | 19.05% | motif file (matrix) | svg |
| 30 | C G T A G C A T C G T A C G T A G C A T A C T G C G A T A G T C A C T G A C T G G A C T C T A G | AT1G71450(AP2EREBP)/col-AT1G71450-DAP-Seq(GSE60143)/Homer | 1e-308 | -7.111e+02 | 0.0000 | 21259.0 | 53.69% | 78339.8 | 43.28% | motif file (matrix) | svg |
| 31 | A C T G C T A G A G T C A C T G A C T G A T G C A C T G T A C G | ESE1(AP2EREBP)/col-ESE1-DAP-Seq(GSE60143)/Homer | 1e-299 | -6.900e+02 | 0.0000 | 8645.0 | 21.83% | 25552.9 | 14.12% | motif file (matrix) | svg |
| 32 | G T A C A C T G A T G C A G T C C T A G G A T C G T A C C T G A G A C T G C A T C G A T G A C T | RAP212(AP2EREBP)/col-RAP212-DAP-Seq(GSE60143)/Homer | 1e-297 | -6.857e+02 | 0.0000 | 10608.0 | 26.79% | 33254.3 | 18.37% | motif file (matrix) | svg |
| 33 | C G A T A G C T T G C A A C T G A G T C T G A C C T A G G T A C A G T C C G T A G C A T G C A T | ERF13(AP2EREBP)/colamp-ERF13-DAP-Seq(GSE60143)/Homer | 1e-288 | -6.643e+02 | 0.0000 | 12144.0 | 30.67% | 39669.5 | 21.92% | motif file (matrix) | svg |
| 34 | C T G A C G A T C T A G T C A G G A T C C T G A T C A G G A T C C T G A A C T G A G T C G C T A A C G T A G T C G C A T | PRDM9(Zf)/Testis-DMC1-ChIP-Seq(GSE35498)/Homer | 1e-286 | -6.605e+02 | 0.0000 | 2722.0 | 6.87% | 5105.2 | 2.82% | motif file (matrix) | svg |
| 35 | G C T A C G T A C G T A G A C T C A T G C T A G G A T C A C T G T C A G G A T C A C T G T A C G | ERF9(AP2EREBP)/colamp-ERF9-DAP-Seq(GSE60143)/Homer | 1e-286 | -6.594e+02 | 0.0000 | 4274.0 | 10.79% | 10002.8 | 5.53% | motif file (matrix) | svg |
| 36 | G A C T G A T C G A T C G C T A G T A C A G T C G C T A C T G A G T A C G A T C G C T A G A C T | MYB13(MYB)/col-MYB13-DAP-Seq(GSE60143)/Homer | 1e-284 | -6.562e+02 | 0.0000 | 7058.0 | 17.82% | 19867.6 | 10.98% | motif file (matrix) | svg |
| 37 | G C A T C G A T G A C T T G C A A C T G A G T C T G A C A C T G G A T C A G T C C G T A G A C T | ERF15(AP2EREBP)/colamp-ERF15-DAP-Seq(GSE60143)/Homer | 1e-279 | -6.437e+02 | 0.0000 | 15554.0 | 39.28% | 54156.9 | 29.92% | motif file (matrix) | svg |
| 38 | C G T A T A G C T A G C T G C A A C T G C T A G C G T A C G T A T C A G G A C T | EHF(ETS)/LoVo-EHF-ChIP-Seq(GSE49402)/Homer | 1e-274 | -6.320e+02 | 0.0000 | 8060.0 | 20.36% | 23848.1 | 13.18% | motif file (matrix) | svg |
| 39 | C G T A C T A G C A T G A G C T C T G A C A T G C A G T C G A T C T A G C T A G | MYB30(MYB)/colamp-MYB30-DAP-Seq(GSE60143)/Homer | 1e-266 | -6.140e+02 | 0.0000 | 9968.0 | 25.17% | 31458.2 | 17.38% | motif file (matrix) | svg |
| 40 | A C T G A C T G A G T C A C T G A C T G A G T C A C G T C T A G | ERF1(AP2EREBP)/colamp-ERF1-DAP-Seq(GSE60143)/Homer | 1e-265 | -6.107e+02 | 0.0000 | 6766.0 | 17.09% | 19167.9 | 10.59% | motif file (matrix) | svg |
| 41 | A T G C A G T C A C T G A T G C A G T C A C T G A G T C G T A C | SHN3(AP2EREBP)/col-SHN3-DAP-Seq(GSE60143)/Homer | 1e-263 | -6.072e+02 | 0.0000 | 3671.0 | 9.27% | 8319.5 | 4.60% | motif file (matrix) | svg |
| 42 | A C T G A C T G A G T C A C T G A C T G A G T C A C G T T C A G | ERF2(AP2EREBP)/colamp-ERF2-DAP-Seq(GSE60143)/Homer | 1e-260 | -6.000e+02 | 0.0000 | 7305.0 | 18.45% | 21277.2 | 11.76% | motif file (matrix) | svg |
| 43 | A C T G C T A G A G T C A C T G A C T G A G T C A C T G T A C G | ERF104(AP2EREBP)/col-ERF104-DAP-Seq(GSE60143)/Homer | 1e-260 | -5.996e+02 | 0.0000 | 11594.0 | 29.28% | 38162.4 | 21.09% | motif file (matrix) | svg |
| 44 | T C G A T G C A C A G T T C G A G A T C A G T C C G T A C G T A A C T G A G T C C G T A C G T A T C A G C G A T A G T C | AT5G25475(ABI3VP1)/col-AT5G25475-DAP-Seq(GSE60143)/Homer | 1e-259 | -5.987e+02 | 0.0000 | 9542.0 | 24.10% | 29933.0 | 16.54% | motif file (matrix) | svg |
| 45 | G C T A C G T A C G T A G C A T C A T G C T A G G A T C A C T G T C A G G A T C A C T G T C A G | ERF4(AP2EREBP)/colamp-ERF4-DAP-Seq(GSE60143)/Homer | 1e-259 | -5.972e+02 | 0.0000 | 14421.0 | 36.42% | 49936.1 | 27.59% | motif file (matrix) | svg |
| 46 | A C T G C T A G A G T C A C T G A C T G A G T C A C G T C T A G | AT5G23930(mTERF)/col-AT5G23930-DAP-Seq(GSE60143)/Homer | 1e-257 | -5.926e+02 | 0.0000 | 13176.0 | 33.28% | 44768.6 | 24.74% | motif file (matrix) | svg |
| 47 | A T G C G T A C A C T G A G T C A G T C A C T G A G T C G T A C | ERF73(AP2EREBP)/col-ERF73-DAP-Seq(GSE60143)/Homer | 1e-244 | -5.637e+02 | 0.0000 | 7237.0 | 18.28% | 21352.3 | 11.80% | motif file (matrix) | svg |
| 48 | C G A T C G T A G C T A G A C T T C G A A G C T A G T C A C T G T C G A A G C T C T G A C G A T | ZBTB38(Zf)/Hela-ZBTB38-ChIP-seq(GSE108618)/Homer | 1e-241 | -5.550e+02 | 0.0000 | 25846.0 | 65.27% | 101811.9 | 56.25% | motif file (matrix) | svg |
| 49 | G T A C A C T G A T G C T G A C C T A G G A C T G T A C C G T A G C A T G C A T | ERF8(AP2EREBP)/colamp-ERF8-DAP-Seq(GSE60143)/Homer | 1e-240 | -5.528e+02 | 0.0000 | 15347.0 | 38.76% | 54468.4 | 30.09% | motif file (matrix) | svg |
| 50 | C T G A G A C T C A T G C T A G A G T C A C T G A C T G A G T C A C T G T C A G | ERF11(AP2EREBP)/col-ERF11-DAP-Seq(GSE60143)/Homer | 1e-239 | -5.513e+02 | 0.0000 | 12030.0 | 30.38% | 40529.0 | 22.39% | motif file (matrix) | svg |
| 51 | C G A T C T A G A C T G A G C T C T G A A C T G A C G T A C G T C T A G C T A G | MYB96(MYB)/colamp-MYB96-DAP-Seq(GSE60143)/Homer | 1e-238 | -5.491e+02 | 0.0000 | 8704.0 | 21.98% | 27156.5 | 15.00% | motif file (matrix) | svg |
| 52 | C A T G G A C T C T A G C A T G C A G T C G A T C T A G C A T G C G A T C G T A C T A G C A G T C G A T C T A G C A T G | AT1G24250(Orphan)/col-AT1G24250-DAP-Seq(GSE60143)/Homer | 1e-235 | -5.412e+02 | 0.0000 | 3633.0 | 9.17% | 8582.5 | 4.74% | motif file (matrix) | svg |
| 53 | C A T G A G C T T A C G G T C A G T A C T A G C A G C T G A C T A T C G T C G A | Esrrb(NR)/mES-Esrrb-ChIP-Seq(GSE11431)/Homer | 1e-232 | -5.346e+02 | 0.0000 | 5139.0 | 12.98% | 13849.2 | 7.65% | motif file (matrix) | svg |
| 54 | G C A T A G C T A C G T A C G T A C T G A C G T G A T C A C G T A C G T A G C T C G A T G C A T A G T C G A C T C A G T | IDD5(C2H2)/colamp-IDD5-DAP-Seq(GSE60143)/Homer | 1e-228 | -5.251e+02 | 0.0000 | 3668.0 | 9.26% | 8799.3 | 4.86% | motif file (matrix) | svg |
| 55 | A G T C G A C T G A T C C G T A G T A C A G T C G C T A C G T A G T A C A G T C G T A C G T A C | MYB63(MYB)/col-MYB63-DAP-Seq(GSE60143)/Homer | 1e-222 | -5.117e+02 | 0.0000 | 5560.0 | 14.04% | 15561.9 | 8.60% | motif file (matrix) | svg |
| 56 | A G C T G A T C G A T C C G T A G T A C A G T C C G A T C T G A G T A C G A T C C G T A G A C T | ATY19(MYB)/col-ATY19-DAP-Seq(GSE60143)/Homer | 1e-219 | -5.062e+02 | 0.0000 | 7442.0 | 18.79% | 22694.2 | 12.54% | motif file (matrix) | svg |
| 57 | C G T A C T A G C A G T A C G T C G T A A C T G C A T G G C A T T C A G C T G A | MYB49(MYB)/col-MYB49-DAP-Seq(GSE60143)/Homer | 1e-218 | -5.041e+02 | 0.0000 | 9515.0 | 24.03% | 30868.7 | 17.06% | motif file (matrix) | svg |
| 58 | C T A G T A G C A T G C C T A G A G T C A G T C C T A G G A C T G A C T G C T A | CRF10(AP2EREBP)/col100-CRF10-DAP-Seq(GSE60143)/Homer | 1e-216 | -4.985e+02 | 0.0000 | 16939.0 | 42.78% | 62140.7 | 34.33% | motif file (matrix) | svg |
| 59 | G C T A C G T A C G T A G C A T C A T G C T A G A G T C A C T G T A C G A G T C C A T G T A C G | RAP26(AP2EREBP)/colamp-RAP26-DAP-Seq(GSE60143)/Homer | 1e-216 | -4.981e+02 | 0.0000 | 15946.0 | 40.27% | 57825.5 | 31.95% | motif file (matrix) | svg |
| 60 | C T G A T C A G A G T C C G T A A T C G A T G C C G A T A C T G A G T C G A C T A T C G A G T C | MyoD(bHLH)/Myotube-MyoD-ChIP-Seq(GSE21614)/Homer | 1e-214 | -4.934e+02 | 0.0000 | 3647.0 | 9.21% | 8931.1 | 4.93% | motif file (matrix) | svg |
| 61 | C A T G G A C T T A C G G T C A G T A C G A T C G A C T A G C T A T C G T C G A T A C G T A G C | ERRg(NR)/Kidney-ESRRG-ChIP-Seq(GSE104905)/Homer | 1e-207 | -4.775e+02 | 0.0000 | 6323.0 | 15.97% | 18707.7 | 10.34% | motif file (matrix) | svg |
| 62 | G C T A C G T A C G T A G C A T C A T G C T A G A G T C A C T G A C T G A G T C A C T G T C A G | ABR1(AP2EREBP)/colamp-ABR1-DAP-Seq(GSE60143)/Homer | 1e-204 | -4.707e+02 | 0.0000 | 13388.0 | 33.81% | 47273.1 | 26.12% | motif file (matrix) | svg |
| 63 | G A C T G A C T G A T C C G T A G T A C A G T C G C A T C G T A G T A C G A T C G C A T G C T A | MYB74(MYB)/colamp-MYB74-DAP-Seq(GSE60143)/Homer | 1e-199 | -4.603e+02 | 0.0000 | 6104.0 | 15.42% | 18045.1 | 9.97% | motif file (matrix) | svg |
| 64 | A G T C C G T A T G A C A T G C G C A T C T G A G T A C G A T C | MYB55(MYB)/colamp-MYB55-DAP-Seq(GSE60143)/Homer | 1e-197 | -4.541e+02 | 0.0000 | 10734.0 | 27.11% | 36432.8 | 20.13% | motif file (matrix) | svg |
| 65 | G A T C G A T C G A T C C G T A G T A C A G T C G C A T C G T A G T A C G A T C | MYB58(MYB)/colamp-MYB58-DAP-Seq(GSE60143)/Homer | 1e-194 | -4.489e+02 | 0.0000 | 9553.0 | 24.13% | 31685.6 | 17.51% | motif file (matrix) | svg |
| 66 | C T A G A G T C A G T C A C T G C G T A A G T C C T G A G A C T | DDF1(AP2EREBP)/col-DDF1-DAP-Seq(GSE60143)/Homer | 1e-194 | -4.470e+02 | 0.0000 | 9096.0 | 22.97% | 29867.3 | 16.50% | motif file (matrix) | svg |
| 67 | T A C G G A C T T G A C C G T A A C G T G A T C G T C A C G T A A C G T A T G C C G T A G A C T | HOXA2(Homeobox)/mES-Hoxa2-ChIP-Seq(Donaldson\_et\_al.)/Homer | 1e-187 | -4.312e+02 | 0.0000 | 1470.0 | 3.71% | 2443.2 | 1.35% | motif file (matrix) | svg |
| 68 | C T A G C T A G T C G A C T A G C G T A A T C G T C G A A C T G C T G A T C G A C T G A T A C G | FRS9(ND)/col-FRS9-DAP-Seq(GSE60143)/Homer | 1e-185 | -4.280e+02 | 0.0000 | 1461.0 | 3.69% | 2430.9 | 1.34% | motif file (matrix) | svg |
| 69 | C T A G A C T G A C G T C G T A A C T G A C T G A G C T C T A G T C A G C T A G | MYB93(MYB)/colamp-MYB93-DAP-Seq(GSE60143)/Homer | 1e-184 | -4.237e+02 | 0.0000 | 10360.0 | 26.16% | 35300.5 | 19.50% | motif file (matrix) | svg |
| 70 | G A C T G A T C A G T C C G T A T G A C A G T C G C A T C T G A G T A C G A T C G C A T G A C T | MYB10(MYB)/col-MYB10-DAP-Seq(GSE60143)/Homer | 1e-180 | -4.167e+02 | 0.0000 | 4613.0 | 11.65% | 12911.4 | 7.13% | motif file (matrix) | svg |
| 71 | C G T A C T A G C A T G G A C T C T G A A C T G A C G T A C G T C T A G C T A G C A T G T C G A | MYB94(MYB)/col-MYB94-DAP-Seq(GSE60143)/Homer | 1e-179 | -4.129e+02 | 0.0000 | 4126.0 | 10.42% | 11185.8 | 6.18% | motif file (matrix) | svg |
| 72 | C A T G C T A G A G T C A C T G A C T G G T A C C A T G T A C G | AT1G28160(AP2EREBP)/colamp-AT1G28160-DAP-Seq(GSE60143)/Homer | 1e-179 | -4.123e+02 | 0.0000 | 18432.0 | 46.55% | 70106.7 | 38.74% | motif file (matrix) | svg |
| 73 | C G T A G C A T C A T G C T A G A G T C A C T G A T C G G T A C A C T G T C A G | At2g33710(AP2EREBP)/colamp-At2g33710-DAP-Seq(GSE60143)/Homer | 1e-178 | -4.117e+02 | 0.0000 | 19753.0 | 49.89% | 76030.0 | 42.01% | motif file (matrix) | svg |
| 74 | G A C T G C A T A C G T A C G T A C T G C G T A A G T C A G C T C G A T A T C G G C A T A C T G C G A T C T A G C G T A | WRKY50(WRKY)/col-WRKY50-DAP-Seq(GSE60143)/Homer | 1e-176 | -4.070e+02 | 0.0000 | 7816.0 | 19.74% | 25242.8 | 13.95% | motif file (matrix) | svg |
| 75 | T C A G A C T G A C G T C G T A A C T G A C T G A C G T C T A G | MYB51(MYB)/col-MYB51-DAP-Seq(GSE60143)/Homer | 1e-176 | -4.070e+02 | 0.0000 | 8879.0 | 22.42% | 29492.8 | 16.30% | motif file (matrix) | svg |
| 76 | A G C T C T A G A G T C A G T C A C T G C G T A A G T C C T G A G C A T G C T A C T G A G C A T G C A T C G A T G C A T | CBF4(AP2EREBP)/colamp-CBF4-DAP-Seq(GSE60143)/Homer | 1e-172 | -3.973e+02 | 0.0000 | 12929.0 | 32.65% | 46412.0 | 25.64% | motif file (matrix) | svg |
| 77 | C T A G C A T G G A C T C G T A C T A G A C T G A C G T C T A G C T A G T C A G | MYB17(MYB)/colamp-MYB17-DAP-Seq(GSE60143)/Homer | 1e-169 | -3.902e+02 | 0.0000 | 6033.0 | 15.24% | 18478.2 | 10.21% | motif file (matrix) | svg |
| 78 | C T A G T C A G C T G A T C A G T G C A A C T G T C G A T C A G | Trl(Zf)/S2-GAGAfactor-ChIP-Seq(GSE40646)/Homer | 1e-169 | -3.893e+02 | 0.0000 | 16083.0 | 40.62% | 60133.3 | 33.22% | motif file (matrix) | svg |
| 79 | A C T G A C G T C G A T C A G T C A T G C A T G C A G T G C A T C A G T C A T G | HuR(?)/HEK293-HuR-CLIP-Seq(GSE87887)/Homer | 1e-167 | -3.861e+02 | 0.0000 | 17966.0 | 45.37% | 68487.0 | 37.84% | motif file (matrix) | svg |
| 80 | A G T C G A T C A G C T C G T A G T A C A G T C G C A T C T G A G T A C G A T C | AT4G26030(C2H2)/col-AT4G26030-DAP-Seq(GSE60143)/Homer | 1e-166 | -3.840e+02 | 0.0000 | 9370.0 | 23.66% | 31784.7 | 17.56% | motif file (matrix) | svg |
| 81 | C G A T C T A G C G T A G A C T C A G T C T A G C G T A A G C T C A T G C T A G | HOXA1(Homeobox)/mES-Hoxa1-ChIP-Seq(SRP084292)/Homer | 1e-166 | -3.826e+02 | 0.0000 | 2749.0 | 6.94% | 6628.0 | 3.66% | motif file (matrix) | svg |
| 82 | G A C T A C T G C G A T A G T C A C T G C T A G A G T C C G T A | Rap210(AP2EREBP)/col-Rap210-DAP-Seq(GSE60143)/Homer | 1e-163 | -3.771e+02 | 0.0000 | 11296.0 | 28.53% | 39839.2 | 22.01% | motif file (matrix) | svg |
| 83 | C A T G G C T A C T A G T A C G C G T A T C A G C G T A A C T G C G T A C A T G C T G A C G T A | BPC1(BBRBPC)/colamp-BPC1-DAP-Seq(GSE60143)/Homer | 1e-163 | -3.769e+02 | 0.0000 | 3227.0 | 8.15% | 8294.8 | 4.58% | motif file (matrix) | svg |
| 84 | G A T C A G T C G A C T G C T A G T A C A G T C G C A T G C T A G T A C G A T C | MYB61(MYB)/colamp-MYB61-DAP-Seq(GSE60143)/Homer | 1e-156 | -3.614e+02 | 0.0000 | 12686.0 | 32.04% | 45944.6 | 25.39% | motif file (matrix) | svg |
| 85 | G T A C A C G T A C G T A T C G C A G T C G A T A T C G G C T A T G C A T A G C C G T A G T C A C A T G A G C T G C T A | ANAC013(NAC)/col-ANAC013-DAP-Seq(GSE60143)/Homer | 1e-147 | -3.408e+02 | 0.0000 | 3657.0 | 9.24% | 10097.6 | 5.58% | motif file (matrix) | svg |
| 86 | G A T C G T A C C T G A A G T C A G T C A C T G G C T A G T A C G T C A G C A T G C A T C G A T | DEAR2(AP2EREBP)/colamp-DEAR2-DAP-Seq(GSE60143)/Homer | 1e-147 | -3.407e+02 | 0.0000 | 15749.0 | 39.77% | 59517.1 | 32.88% | motif file (matrix) | svg |
| 87 | A T C G A G T C A C T G A G T C A G T C A C T G G A C T G A C T | PUCHI(AP2EREBP)/colamp-PUCHI-DAP-Seq(GSE60143)/Homer | 1e-147 | -3.391e+02 | 0.0000 | 9981.0 | 25.21% | 34934.7 | 19.30% | motif file (matrix) | svg |
| 88 | G C A T C G T A C T A G A G T C G T C A C G T A A T G C A C G T A C G T A C T G G A T C G C A T C G T A G C T A G C T A | bHLH122(bHLH)/col100-bHLH122-DAP-Seq(GSE60143)/Homer | 1e-147 | -3.386e+02 | 0.0000 | 6471.0 | 16.34% | 20756.7 | 11.47% | motif file (matrix) | svg |
| 89 | A G C T A G C T C A T G C T G A G T A C A G T C A G C T A G C T C A G T C T A G | RARa(NR)/K562-RARa-ChIP-Seq(Encode)/Homer | 1e-146 | -3.382e+02 | 0.0000 | 15632.0 | 39.48% | 59051.3 | 32.63% | motif file (matrix) | svg |
| 90 | A G T C T G C A T C G A C T G A A C T G C A T G A C G T A T G C G T C A T A C G | Erra(NR)/HepG2-Erra-ChIP-Seq(GSE31477)/Homer | 1e-146 | -3.367e+02 | 0.0000 | 10870.0 | 27.45% | 38673.6 | 21.37% | motif file (matrix) | svg |
| 91 | C G T A G A T C A G C T A C G T A C G T A C T G C G T A G T A C A G C T G C T A C G A T C G A T C G A T G C A T G C T A | WRKY18(WRKY)/col-WRKY18-DAP-Seq(GSE60143)/Homer | 1e-146 | -3.362e+02 | 0.0000 | 12612.0 | 31.85% | 46042.4 | 25.44% | motif file (matrix) | svg |
| 92 | G C A T C T A G A C T G A C G T C G T A A C T G A C T G C G A T C T A G T C G A T C G A G C T A | MYB40(MYB)/col-MYB40-DAP-Seq(GSE60143)/Homer | 1e-145 | -3.341e+02 | 0.0000 | 3523.0 | 8.90% | 9672.6 | 5.34% | motif file (matrix) | svg |
| 93 | A T G C G A T C C G A T A C G T A C G T A C T G C A G T A G C T | Sox3(HMG)/NPC-Sox3-ChIP-Seq(GSE33059)/Homer | 1e-143 | -3.307e+02 | 0.0000 | 11081.0 | 27.98% | 39654.5 | 21.91% | motif file (matrix) | svg |
| 94 | A G T C G A T C G C T A C G A T C A G T T A C G C G A T A G C T A G T C A T C G | SOX1(HMG)/NPC-SOX1-ChIP-Seq(GSE138215)/Homer | 1e-142 | -3.292e+02 | 0.0000 | 12766.0 | 32.24% | 46816.1 | 25.87% | motif file (matrix) | svg |
| 95 | G A C T G C T A T G C A A G T C A C G T A C G T A C G T C G A T A C G T T A C G | At3g45610(C2C2dof)/col-At3g45610-DAP-Seq(GSE60143)/Homer | 1e-142 | -3.280e+02 | 0.0000 | 8920.0 | 22.53% | 30728.5 | 16.98% | motif file (matrix) | svg |
| 96 | C T A G A C T G A C G T C G T A A C T G C A T G G C A T T C A G | MYB92(MYB)/colamp-MYB92-DAP-Seq(GSE60143)/Homer | 1e-141 | -3.268e+02 | 0.0000 | 9063.0 | 22.89% | 31331.4 | 17.31% | motif file (matrix) | svg |
| 97 | G A T C A G C T C T A G G A T C T G A C C T A G C G T A G T A C C G T A G C A T G T C A C T G A | CBF3(AP2EREBP)/colamp-CBF3-DAP-Seq(GSE60143)/Homer | 1e-141 | -3.259e+02 | 0.0000 | 9113.0 | 23.01% | 31550.7 | 17.43% | motif file (matrix) | svg |
| 98 | C G T A T G A C T A G C T G C A A C T G A C T G C G T A C G T A T C A G G A C T | ELF3(ETS)/PDAC-ELF3-ChIP-Seq(GSE64557)/Homer | 1e-141 | -3.259e+02 | 0.0000 | 3759.0 | 9.49% | 10595.4 | 5.85% | motif file (matrix) | svg |
| 99 | G A C T A C T G C G A T A G T C A C T G C T A G A G T C C T G A | AT1G12630(AP2EREBP)/colamp-AT1G12630-DAP-Seq(GSE60143)/Homer | 1e-140 | -3.245e+02 | 0.0000 | 8801.0 | 22.23% | 30291.6 | 16.74% | motif file (matrix) | svg |
| 100 | C G A T T G C A A G T C A C G T A C G T T A C G C G A T C G A T T A C G G C T A G C T A A T G C C G T A G T C A C A T G | ANAC016(NAC)/col-ANAC016-DAP-Seq(GSE60143)/Homer | 1e-140 | -3.237e+02 | 0.0000 | 7239.0 | 18.28% | 23984.5 | 13.25% | motif file (matrix) | svg |
| 101 | A G T C C A T G A C G T A C G T A C T G C G T A A G T C G A C T G C A T G C T A | WRKY28(WRKY)/col-WRKY28-DAP-Seq(GSE60143)/Homer | 1e-140 | -3.235e+02 | 0.0000 | 9457.0 | 23.88% | 33001.5 | 18.23% | motif file (matrix) | svg |
| 102 | C G T A C G T A C T A G A C G T A C G T C G T A A C T G A C T G A C G T C T G A T C G A T C G A | MYB4(MYB)/col200-MYB4-DAP-Seq(GSE60143)/Homer | 1e-140 | -3.235e+02 | 0.0000 | 5799.0 | 14.65% | 18312.6 | 10.12% | motif file (matrix) | svg |
| 103 | G C A T C G A T G C A T C G T A C T A G A G T C G T C A C G T A A T C G A C G T A C G T A C T G G T A C G C A T C G A T | bHLH80(bHLH)/col-bHLH80-DAP-Seq(GSE60143)/Homer | 1e-139 | -3.215e+02 | 0.0000 | 6849.0 | 17.30% | 22459.1 | 12.41% | motif file (matrix) | svg |
| 104 | G A C T A G C T A G C T C T A G A C G T G A T C A C G T A C G T G A C T C G A T G C A T A G T C | IDD4(C2H2)/col-IDD4-DAP-Seq(GSE60143)/Homer | 1e-139 | -3.203e+02 | 0.0000 | 4651.0 | 11.75% | 13953.5 | 7.71% | motif file (matrix) | svg |
| 105 | A T G C A G T C G C A T A G C T A C G T T C A G C G A T A G C T G A T C A T C G | Sox10(HMG)/SciaticNerve-Sox3-ChIP-Seq(GSE35132)/Homer | 1e-137 | -3.175e+02 | 0.0000 | 10308.0 | 26.03% | 36626.9 | 20.24% | motif file (matrix) | svg |
| 106 | G A C T A G T C C T G A A G T C A G T C A C T G C T G A A G T C G C T A G C A T G T A C C G A T G C A T G A C T C G A T | CBF2(AP2EREBP)/colamp-CBF2-DAP-Seq(GSE60143)/Homer | 1e-137 | -3.171e+02 | 0.0000 | 9176.0 | 23.17% | 31939.9 | 17.65% | motif file (matrix) | svg |
| 107 | C G A T C G T A G T A C A C G T A C G T T C A G G C A T C A G T T A C G G T C A C G T A A G T C C G T A T G C A C A T G | NAC2(NAC)/colamp-NAC2-DAP-Seq(GSE60143)/Homer | 1e-137 | -3.170e+02 | 0.0000 | 5118.0 | 12.93% | 15761.7 | 8.71% | motif file (matrix) | svg |
| 108 | C T A G C T A G A T G C G T A C T C A G A T G C A G T C G C A T G A T C G A T C | ZNF91(Zf)/HEK-ZNF91.HA-ChIP-Seq(GSE162571)/Homer | 1e-137 | -3.166e+02 | 0.0000 | 5654.0 | 14.28% | 17829.0 | 9.85% | motif file (matrix) | svg |
| 109 | C A G T A C T G T C A G T G C A G C T A A T G C T C G A A T C G G T C A T G C A | ZNF189(Zf)/HEK293-ZNF189.GFP-ChIP-Seq(GSE58341)/Homer | 1e-136 | -3.145e+02 | 0.0000 | 5162.0 | 13.04% | 15956.1 | 8.82% | motif file (matrix) | svg |
| 110 | G T C A G C T A G C T A T C G A A T C G A C G T A G T C T C G A T C G A T G A C | WRKY40(WRKY)/colamp-WRKY40-DAP-Seq(GSE60143)/Homer | 1e-136 | -3.143e+02 | 0.0000 | 4884.0 | 12.33% | 14898.7 | 8.23% | motif file (matrix) | svg |
| 111 | G T A C G T C A G T A C G T C A G T A C G T C A G T A C G T C A G T A C G T C A | SeqBias: CA-repeat | 1e-131 | -3.038e+02 | 0.0000 | 26068.0 | 65.83% | 107212.8 | 59.24% | motif file (matrix) | svg |
| 112 | C G T A G A T C C T A G A C G T G T A C C T G A A G C T G A T C G C T A G A C T | TGA2(bZIP)/colamp-TGA2-DAP-Seq(GSE60143)/Homer | 1e-130 | -3.007e+02 | 0.0000 | 9069.0 | 22.90% | 31747.3 | 17.54% | motif file (matrix) | svg |
| 113 | G A C T A G T C G A T C C G T A G T A C A G T C G C A T C G T A G T C A G A T C | MYB67(MYB)/col-MYB67-DAP-Seq(GSE60143)/Homer | 1e-126 | -2.909e+02 | 0.0000 | 8755.0 | 22.11% | 30603.8 | 16.91% | motif file (matrix) | svg |
| 114 | C G A T C G T A G T A C A C G T A C G T T C A G G C A T C A G T T A C G G T C A C G T A A G T C C G T A T G C A C A T G | ANAC053(NAC)/colamp-ANAC053-DAP-Seq(GSE60143)/Homer | 1e-123 | -2.839e+02 | 0.0000 | 4402.0 | 11.12% | 13393.5 | 7.40% | motif file (matrix) | svg |
| 115 | C G T A G C A T C A G T C T A G A G T C A C T G A C T G G T A C A C T G A T C G | ERF115(AP2EREBP)/colamp-ERF115-DAP-Seq(GSE60143)/Homer | 1e-122 | -2.827e+02 | 0.0000 | 18123.0 | 45.77% | 71146.8 | 39.31% | motif file (matrix) | svg |
| 116 | G A C T G C A T C T A G C G A T G A T C T C G A C A T G G A T C | Tgif1(Homeobox)/mES-Tgif1-ChIP-Seq(GSE55404)/Homer | 1e-120 | -2.774e+02 | 0.0000 | 17348.0 | 43.81% | 67797.5 | 37.46% | motif file (matrix) | svg |
| 117 | G A T C G A C T G A C T A C G T A G T C A C G T A G T C A C G T A G T C A C G T A G T C A C G T G T A C C G A T G T C A | BPC6(BBRBPC)/col-BPC6-DAP-Seq(GSE60143)/Homer | 1e-120 | -2.766e+02 | 0.0000 | 319.0 | 0.81% | 162.1 | 0.09% | motif file (matrix) | svg |
| 118 | T A C G T C G A C G T A C G T A C G T A C T G A A C T G A C G T C G T A T C G A | AT2G28810(C2C2dof)/colamp-AT2G28810-DAP-Seq(GSE60143)/Homer | 1e-119 | -2.760e+02 | 0.0000 | 13378.0 | 33.79% | 50402.9 | 27.85% | motif file (matrix) | svg |
| 119 | G A C T G T A C C T G A G A T C A G T C C T A G G C T A G T A C C T G A G C T A G C A T C G A T G C A T G A C T C G T A | AT3G16280(AP2EREBP)/colamp-AT3G16280-DAP-Seq(GSE60143)/Homer | 1e-117 | -2.709e+02 | 0.0000 | 7734.0 | 19.53% | 26723.6 | 14.77% | motif file (matrix) | svg |
| 120 | C A T G A G T C G T C A C G T A A T G C A C G T A C G T A C T G | bHLH130(bHLH)/col-bHLH130-DAP-Seq(GSE60143)/Homer | 1e-116 | -2.690e+02 | 0.0000 | 5584.0 | 14.10% | 18131.9 | 10.02% | motif file (matrix) | svg |
| 121 | C G T A A C G T A C G T A C G T A C G T A G T C A G T C C T G A A G C T A G C T | NFAT(RHD)/Jurkat-NFATC1-ChIP-Seq(Jolma\_et\_al.)/Homer | 1e-115 | -2.669e+02 | 0.0000 | 5236.0 | 13.22% | 16795.1 | 9.28% | motif file (matrix) | svg |
| 122 | A G T C A G T C C G A T A C G T A C G T A C T G A C G T A G C T A G T C A G T C | Sox4(HMG)/proB-Sox4-ChIP-Seq(GSE50066)/Homer | 1e-115 | -2.650e+02 | 0.0000 | 5381.0 | 13.59% | 17386.7 | 9.61% | motif file (matrix) | svg |
| 123 | C T A G A C T G A G T C A C T G A C T G A G C T A C T G T C A G | AT3G57600(AP2EREBP)/col-AT3G57600-DAP-Seq(GSE60143)/Homer | 1e-114 | -2.646e+02 | 0.0000 | 7562.0 | 19.10% | 26116.5 | 14.43% | motif file (matrix) | svg |
| 124 | C G T A C G A T C A G T C A T G C G A T G T A C C A T G A C T G G A C T C A T G | CEJ1(AP2EREBP)/col-CEJ1-DAP-Seq(GSE60143)/Homer | 1e-114 | -2.644e+02 | 0.0000 | 14931.0 | 37.71% | 57390.4 | 31.71% | motif file (matrix) | svg |
| 125 | C T G A T G A C T G A C C G T A A C G T T G A C A G C T C T A G A C G T G A C T | Olig2(bHLH)/Neuron-Olig2-ChIP-Seq(GSE30882)/Homer | 1e-113 | -2.624e+02 | 0.0000 | 10392.0 | 26.24% | 37897.1 | 20.94% | motif file (matrix) | svg |
| 126 | C T A G A C T G A G C T C G T A A C T G A C T G A C G T C T A G | MYB99(MYB)/colamp-MYB99-DAP-Seq(GSE60143)/Homer | 1e-113 | -2.618e+02 | 0.0000 | 8954.0 | 22.61% | 31888.4 | 17.62% | motif file (matrix) | svg |
| 127 | A G C T T G A C C G A T C G A T C T A G A C G T C A G T C A G T G C T A A G T C | FOXK1(Forkhead)/HEK293-FOXK1-ChIP-Seq(GSE51673)/Homer | 1e-112 | -2.596e+02 | 0.0000 | 6423.0 | 16.22% | 21586.3 | 11.93% | motif file (matrix) | svg |
| 128 | C T A G A C T G A C G T C G T A A C T G A C T G A C G T T C A G C T G A T C G A | MYB107(MYB)/col-MYB107-DAP-Seq(GSE60143)/Homer | 1e-112 | -2.591e+02 | 0.0000 | 13151.0 | 33.21% | 49748.5 | 27.49% | motif file (matrix) | svg |
| 129 | C G A T C T G A A G T C A C G T A C G T A T C G G A C T C T A G C G A T G A C T C G T A A T G C C G T A G T C A A C T G | ANAC011(NAC)/col-ANAC011-DAP-Seq(GSE60143)/Homer | 1e-111 | -2.570e+02 | 0.0000 | 2325.0 | 5.87% | 6057.3 | 3.35% | motif file (matrix) | svg |
| 130 | A G C T C T A G G A T C A G T C C T A G C T G A A G T C G C T A G C A T T G C A | CBF1(AP2EREBP)/colamp-CBF1-DAP-Seq(GSE60143)/Homer | 1e-110 | -2.547e+02 | 0.0000 | 11594.0 | 29.28% | 43139.3 | 23.84% | motif file (matrix) | svg |
| 131 | A G T C G T A C C T G A A G T C G T A C C T A G G C T A T G A C T G C A G C T A C G T A C G T A | At1g22810(AP2EREBP)/colamp-At1g22810-DAP-Seq(GSE60143)/Homer | 1e-110 | -2.534e+02 | 0.0000 | 7472.0 | 18.87% | 25919.5 | 14.32% | motif file (matrix) | svg |
| 132 | C G A T T C G A G A T C A C G T A C G T T C A G G C A T C T G A C G T A G C T A C G T A A G T C C G T A T G C A C A T G | ANAC050(NAC)/colamp-ANAC050-DAP-Seq(GSE60143)/Homer | 1e-109 | -2.530e+02 | 0.0000 | 4648.0 | 11.74% | 14681.9 | 8.11% | motif file (matrix) | svg |
| 133 | C G A T C G A T G C A T G A C T A C G T C G T A C G T A A C T G T A G C C G T A C G T A C G T A | AT5G60130(ABI3VP1)/col-AT5G60130-DAP-Seq(GSE60143)/Homer | 1e-109 | -2.516e+02 | 0.0000 | 6997.0 | 17.67% | 24014.3 | 13.27% | motif file (matrix) | svg |
| 134 | C G T A C G A T C G T A C G A T C A T G A C T G C G A T A G T C A T C G T C A G G A C T A C T G | At1g36060(AP2EREBP)/colamp-At1g36060-DAP-Seq(GSE60143)/Homer | 1e-108 | -2.509e+02 | 0.0000 | 10227.0 | 25.83% | 37406.9 | 20.67% | motif file (matrix) | svg |
| 135 | G A T C C T G A A G T C A G C T A C G T A C G T A C G T A C G T | At1g64620(C2C2dof)/colamp-At1g64620-DAP-Seq(GSE60143)/Homer | 1e-106 | -2.456e+02 | 0.0000 | 8926.0 | 22.54% | 32044.4 | 17.71% | motif file (matrix) | svg |
| 136 | C G A T C T G A A G T C A C G T A C G T T A C G G C T A C A T G C T A G G C A T C G A T A G T C C G T A G T C A A C T G | ANAC096(NAC)/colamp-ANAC096-DAP-Seq(GSE60143)/Homer | 1e-106 | -2.450e+02 | 0.0000 | 4624.0 | 11.68% | 14684.7 | 8.11% | motif file (matrix) | svg |
| 137 | A T G C G T A C C G T A A G C T G C A T T A C G A G C T A G C T A G T C A G C T | Sox6(HMG)/Myotubes-Sox6-ChIP-Seq(GSE32627)/Homer | 1e-105 | -2.438e+02 | 0.0000 | 10665.0 | 26.93% | 39389.8 | 21.76% | motif file (matrix) | svg |
| 138 | G A T C C T G A A G T C A G T C A C T G C G T A A G T C C T G A | ERF38(AP2EREBP)/col-ERF38-DAP-Seq(GSE60143)/Homer | 1e-105 | -2.431e+02 | 0.0000 | 8504.0 | 21.48% | 30333.0 | 16.76% | motif file (matrix) | svg |
| 139 | G A C T C G A T C G A T C T G A G T A C A G T C C G A T C G T A G T C A G A T C G C A T G C A T | MYB121(MYB)/col-MYB121-DAP-Seq(GSE60143)/Homer | 1e-105 | -2.425e+02 | 0.0000 | 3607.0 | 9.11% | 10834.5 | 5.99% | motif file (matrix) | svg |
| 140 | A G C T A C G T A C T G A T G C A G T C C G T A C T G A T A C G | NF1-halfsite(CTF)/LNCaP-NF1-ChIP-Seq(Unpublished)/Homer | 1e-104 | -2.412e+02 | 0.0000 | 9950.0 | 25.13% | 36413.8 | 20.12% | motif file (matrix) | svg |
| 141 | C A G T T C A G T C G A A G T C C G T A A C T G T G A C C G A T A C T G A C T G A C G T A T C G | Atoh7(bHLH)/Retina-Atoh7-CutnRun(GSE156756)/Homer | 1e-104 | -2.409e+02 | 0.0000 | 3614.0 | 9.13% | 10876.5 | 6.01% | motif file (matrix) | svg |
| 142 | C G A T T C G A G A T C A C G T A C G T T C A G G C T A C G A T C G T A C G T A C G T A A T G C C G T A T G C A C T A G | ANAC028(NAC)/col-ANAC028-DAP-Seq(GSE60143)/Homer | 1e-102 | -2.359e+02 | 0.0000 | 5250.0 | 13.26% | 17245.2 | 9.53% | motif file (matrix) | svg |
| 143 | G A C T G A T C C T G A A G T C A G T C A C T G C G T A A G T C G T A C G C T A G C A T C G A T | At1g19210(AP2EREBP)/colamp-At1g19210-DAP-Seq(GSE60143)/Homer | 1e-102 | -2.351e+02 | 0.0000 | 18497.0 | 46.71% | 73855.3 | 40.81% | motif file (matrix) | svg |
| 144 | A T C G T G C A G A T C C T A G A C G T A T C G C G T A A G T C T C A G A C T G T C A G G C T A | Knotted(Homeobox)/Corn-KN1-ChIP-Seq(GSE39161)/Homer | 1e-101 | -2.336e+02 | 0.0000 | 14525.0 | 36.68% | 56252.1 | 31.08% | motif file (matrix) | svg |
| 145 | C G T A C G T A C G T A C G T A C T G A C A G T A C G T C G T A A C T G A C T G A C G T C T A G C T G A T C G A C T G A | MYB39(MYB)/col-MYB39-DAP-Seq(GSE60143)/Homer | 1e-101 | -2.334e+02 | 0.0000 | 1556.0 | 3.93% | 3624.2 | 2.00% | motif file (matrix) | svg |
| 146 | A T G C C A G T A G C T A G C T T C A G G T C A T A G C G A C T C G T A C G A T | WRKY20(WRKY)/col-WRKY20-DAP-Seq(GSE60143)/Homer | 1e-101 | -2.330e+02 | 0.0000 | 5879.0 | 14.85% | 19783.4 | 10.93% | motif file (matrix) | svg |
| 147 | T A C G T A G C G C T A C G A T C T A G A C G T C A G T C A G T G C T A A G T C G T C A G C A T | FOXK2(Forkhead)/U2OS-FOXK2-ChIP-Seq(E-MTAB-2204)/Homer | 1e-100 | -2.322e+02 | 0.0000 | 4141.0 | 10.46% | 12974.3 | 7.17% | motif file (matrix) | svg |
| 148 | A T G C G T A C C T G A A G T C A G T C A C T G G T C A A G T C G T C A G C A T G C A T G A C T | At5g65130(AP2EREBP)/colamp-At5g65130-DAP-Seq(GSE60143)/Homer | 1e-100 | -2.318e+02 | 0.0000 | 3637.0 | 9.19% | 11060.3 | 6.11% | motif file (matrix) | svg |
| 149 | C T G A T C G A G T A C A C G T A C G T A T C G A C G T C G A T A T C G G C T A G T A C A T G C C G T A T G C A C A T G | ANAC103(NAC)/col-ANAC103-DAP-Seq(GSE60143)/Homer | 1e-100 | -2.305e+02 | 0.0000 | 3025.0 | 7.64% | 8792.0 | 4.86% | motif file (matrix) | svg |
| 150 | C T G A A G T C C G A T A G C T A T G C G T A C A C G T A T C G C A G T G C A T | Elf4(ETS)/BMDM-Elf4-ChIP-Seq(GSE88699)/Homer | 1e-100 | -2.304e+02 | 0.0000 | 6376.0 | 16.10% | 21819.4 | 12.06% | motif file (matrix) | svg |
| 151 | C G A T C T A G A C G T A C G T A C G T C G T A A G C T C G A T A G C T C G T A C T A G T A G C | FoxD3(forkhead)/ZebrafishEmbryo-Foxd3.biotin-ChIP-seq(GSE106676)/Homer | 1e-99 | -2.292e+02 | 0.0000 | 3786.0 | 9.56% | 11653.4 | 6.44% | motif file (matrix) | svg |
| 152 | G T A C G T A C G T C A G C T A C G T A C G T A C G T A C T A G C T A G C T A G | SEP3(MADS)/Arabidoposis-Flower-Sep3-ChIP-Seq/Homer | 1e-99 | -2.288e+02 | 0.0000 | 7075.0 | 17.87% | 24683.2 | 13.64% | motif file (matrix) | svg |
| 153 | T C A G A C G T T C G A T A G C A G T C C G T A A C T G G T A C A C G T A C T G A T C G A G T C | Atoh1(bHLH)/Cerebellum-Atoh1-ChIP-Seq(GSE22111)/Homer | 1e-99 | -2.284e+02 | 0.0000 | 5404.0 | 13.65% | 17955.4 | 9.92% | motif file (matrix) | svg |
| 154 | C G T A C G T A C T A G C A G T G A C T C G T A C A T G C A T G C G A T C T G A C T G A C T G A | MS188(MYB)/colamp-MS188-DAP-Seq(GSE60143)/Homer | 1e-98 | -2.261e+02 | 0.0000 | 6027.0 | 15.22% | 20476.2 | 11.31% | motif file (matrix) | svg |
| 155 | A T G C C A T G A G C T C A G T C A T G T C G A A G T C G A C T C G A T C G A T C A G T C A G T | WRKY26(WRKY)/colamp-WRKY26-DAP-Seq(GSE60143)/Homer | 1e-97 | -2.252e+02 | 0.0000 | 4355.0 | 11.00% | 13883.6 | 7.67% | motif file (matrix) | svg |
| 156 | T A G C C G T A C T G A T A C G C G T A A C G T A C T G A C T G A G T C T A C G C T A G G T A C | YY1(Zf)/Promoter/Homer | 1e-96 | -2.229e+02 | 0.0000 | 680.0 | 1.72% | 1047.4 | 0.58% | motif file (matrix) | svg |
| 157 | T C G A A C T G C A T G A G C T A G T C C G T A C T G A C T A G A C T G C G A T A T G C C T G A | RAR:RXR(NR),DR0/ES-RAR-ChIP-Seq(GSE56893)/Homer | 1e-96 | -2.216e+02 | 0.0000 | 961.0 | 2.43% | 1834.1 | 1.01% | motif file (matrix) | svg |
| 158 | C T A G C T G A C G T A C G T A C G T A C G T A A C T G A C G T C T A G G T C A | COG1(C2C2dof)/col-COG1-DAP-Seq(GSE60143)/Homer | 1e-95 | -2.190e+02 | 0.0000 | 8238.0 | 20.80% | 29640.6 | 16.38% | motif file (matrix) | svg |
| 159 | A G T C G A T C C T G A A G T C A G T C C A T G G T C A G A T C C G T A G A T C | DREB26(AP2EREBP)/col-DREB26-DAP-Seq(GSE60143)/Homer | 1e-95 | -2.188e+02 | 0.0000 | 3263.0 | 8.24% | 9794.0 | 5.41% | motif file (matrix) | svg |
| 160 | C T A G C T A G C G T A C G T A T A C G C G A T C T A G C T G A C T G A C G T A T A C G G A C T | PU.1:IRF8(ETS:IRF)/pDC-Irf8-ChIP-Seq(GSE66899)/Homer | 1e-93 | -2.158e+02 | 0.0000 | 1083.0 | 2.74% | 2226.7 | 1.23% | motif file (matrix) | svg |
| 161 | A C G T A G T C A G T C C G A T A C G T A C G T A C T G A C G T A T G C G A C T A C T G T A C G | Sox21(HMG)/ESC-SOX21-ChIP-Seq(GSE110505)/Homer | 1e-93 | -2.158e+02 | 0.0000 | 10613.0 | 26.80% | 39706.4 | 21.94% | motif file (matrix) | svg |
| 162 | G A C T C A G T G C A T C G A T T G A C A C G T A T G C G T A C C T G A A C T G A C T G A G C T | WIP5(C2H2)/colamp-WIP5-DAP-Seq(GSE60143)/Homer | 1e-93 | -2.153e+02 | 0.0000 | 7176.0 | 18.12% | 25314.8 | 13.99% | motif file (matrix) | svg |
| 163 | C T G A A T G C G C T A G C A T A T G C C G T A T C G A C T G A C T A G T C A G T A C G G T C A | Tcf4(HMG)/Hct116-Tcf4-ChIP-Seq(SRA012054)/Homer | 1e-93 | -2.142e+02 | 0.0000 | 2928.0 | 7.39% | 8594.2 | 4.75% | motif file (matrix) | svg |
| 164 | C T A G G C T A A G T C A C T G A C G T G A C T G A C T A T G C T C G A C A G T G A T C C G A T G A C T G A T C G A T C | RKD2(RWPRK)/colamp-RKD2-DAP-Seq(GSE60143)/Homer | 1e-91 | -2.113e+02 | 0.0000 | 5663.0 | 14.30% | 19233.8 | 10.63% | motif file (matrix) | svg |
| 165 | A G C T C A T G G C A T G A T C T G C A C T A G G A T C A C G T | Tgif2(Homeobox)/mES-Tgif2-ChIP-Seq(GSE55404)/Homer | 1e-91 | -2.100e+02 | 0.0000 | 18453.0 | 46.60% | 74246.3 | 41.02% | motif file (matrix) | svg |
| 166 | G C A T C G T A G C T A G A C T C G A T G A C T A G T C C A G T A G T C A G T C A C T G C T A G G T A C C T A G C T G A | AT5G05550(Trihelix)/col-AT5G05550-DAP-Seq(GSE60143)/Homer | 1e-90 | -2.090e+02 | 0.0000 | 16676.0 | 42.11% | 66313.6 | 36.64% | motif file (matrix) | svg |
| 167 | C T A G G T A C C A T G G A C T C G A T C A T G G T C A G T A C G A C T C G A T C G A T C G A T | WRKY27(WRKY)/colamp-WRKY27-DAP-Seq(GSE60143)/Homer | 1e-90 | -2.083e+02 | 0.0000 | 6279.0 | 15.86% | 21765.7 | 12.03% | motif file (matrix) | svg |
| 168 | A G T C A C G T A C T G A G C T A C G T A C G T G T C A A G T C | Foxo1(Forkhead)/RAW-Foxo1-ChIP-Seq(Fan\_et\_al.)/Homer | 1e-90 | -2.082e+02 | 0.0000 | 9701.0 | 24.50% | 35981.5 | 19.88% | motif file (matrix) | svg |
| 169 | C G T A C G T A C G A T A C T G C A G T A G T C A C T G A C T G A G C T A C T G | DREB19(AP2EREBP)/colamp-DREB19-DAP-Seq(GSE60143)/Homer | 1e-89 | -2.057e+02 | 0.0000 | 8548.0 | 21.59% | 31173.0 | 17.22% | motif file (matrix) | svg |
| 170 | G A T C A T G C A G T C C G T A A G T C A G T C A C T G G C T A A G T C C G T A | AT1G44830(AP2EREBP)/col-AT1G44830-DAP-Seq(GSE60143)/Homer | 1e-88 | -2.045e+02 | 0.0000 | 4277.0 | 10.80% | 13839.2 | 7.65% | motif file (matrix) | svg |
| 171 | A T G C A G T C G A T C C G T A A C G T A C G T A C T G A C G T A G C T G A T C | Sox2(HMG)/mES-Sox2-ChIP-Seq(GSE11431)/Homer | 1e-88 | -2.035e+02 | 0.0000 | 5724.0 | 14.46% | 19594.7 | 10.83% | motif file (matrix) | svg |
| 172 | G A C T A C T G C A G T A G T C A C T G A C T G A G C T A C T G C T A G G T C A | At1g77640(AP2EREBP)/col-At1g77640-DAP-Seq(GSE60143)/Homer | 1e-88 | -2.030e+02 | 0.0000 | 2790.0 | 7.05% | 8196.6 | 4.53% | motif file (matrix) | svg |
| 173 | T A C G C G T A G A C T T C A G A G C T A G T C A C T G T C A G A G T C C T G A | DDF2(AP2EREBP)/col-DDF2-DAP-Seq(GSE60143)/Homer | 1e-88 | -2.028e+02 | 0.0000 | 1441.0 | 3.64% | 3439.4 | 1.90% | motif file (matrix) | svg |
| 174 | A G T C T G A C C T G A A G T C A G T C A C T G C G T A A G T C G T C A G C T A G C A T C G T A G C A T G C T A C G T A | DEAR3(AP2EREBP)/colamp-DEAR3-DAP-Seq(GSE60143)/Homer | 1e-87 | -2.020e+02 | 0.0000 | 6296.0 | 15.90% | 21934.8 | 12.12% | motif file (matrix) | svg |
| 175 | T C A G T C A G G C T A C G T A T A C G G A C T T C A G T C G A C T G A C G T A T A C G G A C T | IRF8(IRF)/BMDM-IRF8-ChIP-Seq(GSE77884)/Homer | 1e-87 | -2.017e+02 | 0.0000 | 1873.0 | 4.73% | 4912.8 | 2.71% | motif file (matrix) | svg |
| 176 | C G T A C G A T C T A G C G T A A G C T C A G T T A C G C G T A A C G T C A T G C T A G A T G C | HOXA3(Homeobox)/mEmbryo-Hoxa3-ChIP-Seq(E-MTAB-8607)/Homer | 1e-87 | -2.017e+02 | 0.0000 | 1521.0 | 3.84% | 3712.9 | 2.05% | motif file (matrix) | svg |
| 177 | C A T G C T G A A G T C A C T G A C T G A G C T A C T G A T C G | ESE3(AP2EREBP)/col-ESE3-DAP-Seq(GSE60143)/Homer | 1e-87 | -2.012e+02 | 0.0000 | 14986.0 | 37.85% | 59007.5 | 32.60% | motif file (matrix) | svg |
| 178 | G T A C A C G T A C G T T C A G G A C T G C A T T C A G C G T A C T G A A G T C C G T A G T C A A C T G A C G T G C T A | NTM2(NAC)/col-NTM2-DAP-Seq(GSE60143)/Homer | 1e-87 | -2.010e+02 | 0.0000 | 3938.0 | 9.95% | 12567.4 | 6.94% | motif file (matrix) | svg |
| 179 | G A T C C T G A A G T C G T A C A C T G G C T A G A T C C T G A G C T A G C T A | At4g31060(AP2EREBP)/colamp-At4g31060-DAP-Seq(GSE60143)/Homer | 1e-87 | -2.007e+02 | 0.0000 | 7417.0 | 18.73% | 26556.0 | 14.67% | motif file (matrix) | svg |
| 180 | T C G A T G A C G T A C C G T A C A G T T G A C A C G T A C T G A G C T A G C T | NeuroG2(bHLH)/Fibroblast-NeuroG2-ChIP-Seq(GSE75910)/Homer | 1e-86 | -2.001e+02 | 0.0000 | 7764.0 | 19.61% | 28004.9 | 15.47% | motif file (matrix) | svg |
| 181 | A T G C C A T G G C A T G A C T C T A G T C G A G T A C A G C T C G T A G C T A | WRKY75(WRKY)/col-WRKY75-DAP-Seq(GSE60143)/Homer | 1e-86 | -1.997e+02 | 0.0000 | 7133.0 | 18.01% | 25400.7 | 14.03% | motif file (matrix) | svg |
| 182 | G C A T T G A C C A T G G A C T C A G T C A T G T C G A G T A C G A C T G C T A C G A T C G A T | WRKY6(WRKY)/colamp-WRKY6-DAP-Seq(GSE60143)/Homer | 1e-86 | -1.994e+02 | 0.0000 | 6520.0 | 16.47% | 22889.3 | 12.65% | motif file (matrix) | svg |
| 183 | A T G C A G T C A G C T A G C T A C G T A T C G C G T A C G A T T A G C G A C T | LEF1(HMG)/H1-LEF1-ChIP-Seq(GSE64758)/Homer | 1e-86 | -1.992e+02 | 0.0000 | 3989.0 | 10.07% | 12787.7 | 7.07% | motif file (matrix) | svg |
| 184 | C T A G T C G A T G A C A G T C C G T A A C T G G T A C A C G T A C T G A C T G | BHLHA15(bHLH)/NIH3T3-BHLHB8.HA-ChIP-Seq(GSE119782)/Homer | 1e-86 | -1.982e+02 | 0.0000 | 6522.0 | 16.47% | 22917.5 | 12.66% | motif file (matrix) | svg |
| 185 | G C T A C G T A C G A T G A C T G C A T T G C A A G T C A G C T A C G T A C G T C G A T G A C T | DAG2(C2C2dof)/col-DAG2-DAP-Seq(GSE60143)/Homer | 1e-85 | -1.969e+02 | 0.0000 | 8047.0 | 20.32% | 29242.2 | 16.16% | motif file (matrix) | svg |
| 186 | G A C T C T G A G T A C A G T C C G A T C G T A G T C A G A T C G C A T G C A T G C A T C G A T | AT3G10580(MYBrelated)/colamp-AT3G10580-DAP-Seq(GSE60143)/Homer | 1e-85 | -1.967e+02 | 0.0000 | 3964.0 | 10.01% | 12721.5 | 7.03% | motif file (matrix) | svg |
| 187 | C T G A T C G A C G T A A T G C C G T A C G T A C G A T C T A G T C A G G A T C | Sox15(HMG)/CPA-Sox15-ChIP-Seq(GSE62909)/Homer | 1e-85 | -1.960e+02 | 0.0000 | 6186.0 | 15.62% | 21583.5 | 11.93% | motif file (matrix) | svg |
| 188 | G C T A A G T C T A C G T G C A A T C G T C A G G C T A T C G A T C A G A G C T | ELF5(ETS)/T47D-ELF5-ChIP-Seq(GSE30407)/Homer | 1e-84 | -1.954e+02 | 0.0000 | 4566.0 | 11.53% | 15094.8 | 8.34% | motif file (matrix) | svg |
| 189 | C G A T G C T A G C T A G C A T G C T A C G T A A G T C A C G T A C G T A C G T C G A T A G C T | At5g62940(C2C2dof)/col-At5g62940-DAP-Seq(GSE60143)/Homer | 1e-84 | -1.952e+02 | 0.0000 | 18113.0 | 45.74% | 73081.4 | 40.38% | motif file (matrix) | svg |
| 190 | A G C T G T C A T G C A A G T C A C G T A C G T A C G T C G A T G A C T T A C G | AT3G12130(C3H)/colamp-AT3G12130-DAP-Seq(GSE60143)/Homer | 1e-84 | -1.949e+02 | 0.0000 | 13879.0 | 35.05% | 54285.1 | 29.99% | motif file (matrix) | svg |
| 191 | T C G A A G T C C G T A A T C G A T G C C G A T A C T G A G T C A G C T A C T G | Tcf12(bHLH)/GM12878-Tcf12-ChIP-Seq(GSE32465)/Homer | 1e-84 | -1.942e+02 | 0.0000 | 3712.0 | 9.37% | 11778.2 | 6.51% | motif file (matrix) | svg |
| 192 | C G A T C G T A A G T C A C G T A C G T T C G A T G C A G C A T G C T A C G T A A C G T A G C T C G T A C G T A A C T G | ANAC062(NAC)/colamp-ANAC062-DAP-Seq(GSE60143)/Homer | 1e-84 | -1.939e+02 | 0.0000 | 2200.0 | 5.56% | 6133.0 | 3.39% | motif file (matrix) | svg |
| 193 | C G A T T C G A G A T C C G A T G C A T T C A G G A C T C G A T G C A T G C T A C T G A A G T C C G T A G T C A C T A G | ANAC005(NAC)/col-ANAC005-DAP-Seq(GSE60143)/Homer | 1e-84 | -1.937e+02 | 0.0000 | 2617.0 | 6.61% | 7650.4 | 4.23% | motif file (matrix) | svg |
| 194 | G T A C C A T G A G C T A G C T T C A G T G C A T G A C A G C T C G T A C G T A | WRKY33(WRKY)/col-WRKY33-DAP-Seq(GSE60143)/Homer | 1e-83 | -1.919e+02 | 0.0000 | 6738.0 | 17.02% | 23908.2 | 13.21% | motif file (matrix) | svg |
| 195 | G T A C C T G A T A G C C G T A G C T A T C G A T G C A T G A C C T A G G T C A A G T C C G T A C T G A C T G A C G T A | At1g14580(C2H2)/colamp-At1g14580-DAP-Seq(GSE60143)/Homer | 1e-81 | -1.865e+02 | 0.0000 | 1182.0 | 2.99% | 2696.0 | 1.49% | motif file (matrix) | svg |
| 196 | A G C T G A T C C T G A A G T C A G T C A C T G C G T A A G T C C T G A G T C A G C A T C G A T G C T A G C A T C G T A | At2g44940(AP2EREBP)/colamp-At2g44940-DAP-Seq(GSE60143)/Homer | 1e-80 | -1.864e+02 | 0.0000 | 4558.0 | 11.51% | 15187.0 | 8.39% | motif file (matrix) | svg |
| 197 | C G A T C G A T G C A T G C A T G T C A A G T C A G C T A C G T A C G T C G A T G A C T A C G T | OBP4(C2C2dof)/col-OBP4-DAP-Seq(GSE60143)/Homer | 1e-80 | -1.862e+02 | 0.0000 | 7634.0 | 19.28% | 27718.4 | 15.31% | motif file (matrix) | svg |
| 198 | C G T A G A T C C A T G G C A T G A C T C T A G T C G A T A G C A G C T G C A T | WRKY55(WRKY)/col-WRKY55-DAP-Seq(GSE60143)/Homer | 1e-80 | -1.857e+02 | 0.0000 | 8273.0 | 20.89% | 30397.6 | 16.80% | motif file (matrix) | svg |
| 199 | T G A C A G T C C G T A A C T G G T A C A C G T A C T G A C G T G A C T G A T C | Twist2(bHLH)/Myoblast-Twist2.Ty1-ChIP-Seq(GSE127998)/Homer | 1e-80 | -1.856e+02 | 0.0000 | 8378.0 | 21.16% | 30841.3 | 17.04% | motif file (matrix) | svg |
| 200 | C A T G T G C A G A C T C A T G C G T A A G T C T C A G G C A T T G A C C G T A | bZIP50(bZIP)/colamp-bZIP50-DAP-Seq(GSE60143)/Homer | 1e-80 | -1.850e+02 | 0.0000 | 11593.0 | 29.28% | 44561.9 | 24.62% | motif file (matrix) | svg |
| 201 | T C A G T G A C G T A C C G T A A C G T T G A C A C G T T C A G A G C T G A C T | NeuroD1(bHLH)/Islet-NeuroD1-ChIP-Seq(GSE30298)/Homer | 1e-80 | -1.850e+02 | 0.0000 | 3973.0 | 10.03% | 12907.3 | 7.13% | motif file (matrix) | svg |
| 202 | G T A C C A T G A G C T A C G T A C T G C G T A A G T C G A C T G C A T C G A T | WRKY29(WRKY)/colamp-WRKY29-DAP-Seq(GSE60143)/Homer | 1e-80 | -1.846e+02 | 0.0000 | 7596.0 | 19.18% | 27588.5 | 15.24% | motif file (matrix) | svg |
| 203 | C A T G A T G C T A G C C T G A A G T C A G T C A C T G G C T A A G T C G T A C G C T A G C A T | At4g28140(AP2EREBP)/colamp-At4g28140-DAP-Seq(GSE60143)/Homer | 1e-78 | -1.802e+02 | 0.0000 | 5294.0 | 13.37% | 18220.2 | 10.07% | motif file (matrix) | svg |
| 204 | G A T C C T G A G A T C G A T C C T A G G C T A A G T C C T G A G C T A C G T A | At4g16750(AP2EREBP)/col-At4g16750-DAP-Seq(GSE60143)/Homer | 1e-78 | -1.801e+02 | 0.0000 | 12793.0 | 32.31% | 49883.9 | 27.56% | motif file (matrix) | svg |
| 205 | A G T C A G T C C T G A A G T C A G T C A C T G C G T A A G T C C T G A T C G A G C A T G A T C C G A T C G A T A C T G | AT3G60490(AP2EREBP)/colamp-AT3G60490-DAP-Seq(GSE60143)/Homer | 1e-78 | -1.796e+02 | 0.0000 | 5705.0 | 14.41% | 19893.2 | 10.99% | motif file (matrix) | svg |
| 206 | T C G A G T A C C A T G A G C T A C G T C A T G G T C A G T A C A G C T G C T A C G A T C A G T | WRKY31(WRKY)/colamp-WRKY31-DAP-Seq(GSE60143)/Homer | 1e-77 | -1.796e+02 | 0.0000 | 5309.0 | 13.41% | 18291.2 | 10.11% | motif file (matrix) | svg |
| 207 | T C A G A G C T G T C A C G T A A C G T A T G C C G T A A C G T A C G T C T G A | PHV(HB)/col-PHV-DAP-Seq(GSE60143)/Homer | 1e-77 | -1.795e+02 | 0.0000 | 2722.0 | 6.87% | 8187.0 | 4.52% | motif file (matrix) | svg |
| 208 | C G A T T G A C C A T G G A C T A C G T C A T G C G T A G A T C G A C T G C A T G C A T C G A T | WRKY14(WRKY)/colamp-WRKY14-DAP-Seq(GSE60143)/Homer | 1e-77 | -1.788e+02 | 0.0000 | 3735.0 | 9.43% | 12060.6 | 6.66% | motif file (matrix) | svg |
| 209 | A G T C A C G T A C G T T A C G G C A T G C A T A T G C G C T A C G T A A T G C C G T A G T C A A C T G G A T C G C A T | ANAC075(NAC)/col-ANAC075-DAP-Seq(GSE60143)/Homer | 1e-77 | -1.783e+02 | 0.0000 | 3051.0 | 7.71% | 9438.4 | 5.21% | motif file (matrix) | svg |
| 210 | C G A T T C G A G T A C A C G T A C G T T C A G G C A T G C A T G C T A C G T A C G T A A G T C C G T A T G C A C A T G | ANAC020(NAC)/col-ANAC020-DAP-Seq(GSE60143)/Homer | 1e-77 | -1.783e+02 | 0.0000 | 5084.0 | 12.84% | 17405.4 | 9.62% | motif file (matrix) | svg |
| 211 | C G A T T G C A T G C A G A T C C G T A A C T G T G A C G A C T C A T G A C T G | Tcf21(bHLH)/ArterySmoothMuscle-Tcf21-ChIP-Seq(GSE61369)/Homer | 1e-77 | -1.776e+02 | 0.0000 | 3854.0 | 9.73% | 12540.0 | 6.93% | motif file (matrix) | svg |
| 212 | T C G A T A G C G T C A A C T G A C T G C G T A C G T A C T A G A G C T T C A G | ERG(ETS)/VCaP-ERG-ChIP-Seq(GSE14097)/Homer | 1e-76 | -1.767e+02 | 0.0000 | 7340.0 | 18.54% | 26670.1 | 14.74% | motif file (matrix) | svg |
| 213 | C T A G C T A G T G A C G T A C C A T G A C T G G A T C G A T C C G T A C G T A | RAP211(AP2EREBP)/colamp-RAP211-DAP-Seq(GSE60143)/Homer | 1e-75 | -1.744e+02 | 0.0000 | 18497.0 | 46.71% | 75348.9 | 41.63% | motif file (matrix) | svg |
| 214 | G A C T T C A G G C A T A G T C G C T A G A T C C T G A A C G T A G T C G T C A | Replumless(BLH)/Arabidopsis-RPL.GFP-ChIP-Seq(GSE78727)/Homer | 1e-75 | -1.741e+02 | 0.0000 | 9780.0 | 24.70% | 37015.8 | 20.45% | motif file (matrix) | svg |
| 215 | A G T C C T A G A C G T A C G T A C T G C G T A A G T C A G C T G C T A G C A T | WRKY24(WRKY)/colamp-WRKY24-DAP-Seq(GSE60143)/Homer | 1e-74 | -1.726e+02 | 0.0000 | 6618.0 | 16.71% | 23753.6 | 13.12% | motif file (matrix) | svg |
| 216 | T C G A C T G A C G T A C G T A C G T A C T G A A C T G A C G T C T G A C T G A | AT5G63260(C3H)/col-AT5G63260-DAP-Seq(GSE60143)/Homer | 1e-74 | -1.719e+02 | 0.0000 | 12650.0 | 31.95% | 49455.1 | 27.32% | motif file (matrix) | svg |
| 217 | G A C T C T G A A G T C A G T C A C T G C G T A A G T C C T G A | bHLH10(bHLH)/colamp-bHLH10-DAP-Seq(GSE60143)/Homer | 1e-74 | -1.705e+02 | 0.0000 | 5514.0 | 13.93% | 19266.0 | 10.64% | motif file (matrix) | svg |
| 218 | G C A T T G A C C T G A A G T C A G T C A C T G G T C A A G T C G C T A G A C T G C T A C T G A | DREB2(AP2EREBP)/col-DREB2-DAP-Seq(GSE60143)/Homer | 1e-73 | -1.686e+02 | 0.0000 | 7176.0 | 18.12% | 26140.0 | 14.44% | motif file (matrix) | svg |
| 219 | T A G C C A T G G A C T G A C T T C A G G T C A G A T C G A C T G C A T G C T A | WRKY15(WRKY)/col-WRKY15-DAP-Seq(GSE60143)/Homer | 1e-72 | -1.672e+02 | 0.0000 | 7359.0 | 18.58% | 26930.8 | 14.88% | motif file (matrix) | svg |
| 220 | A G T C A C G T A C G T T C A G G C T A G C T A A T G C C G T A C G A T A G T C C G T A G T C A A C T G G A T C G C A T | SND3(NAC)/col-SND3-DAP-Seq(GSE60143)/Homer | 1e-72 | -1.669e+02 | 0.0000 | 5374.0 | 13.57% | 18757.4 | 10.36% | motif file (matrix) | svg |
| 221 | T C G A T A G C T G C A A C T G A C T G C G T A C G T A C T A G G A C T T A C G | ETS1(ETS)/Jurkat-ETS1-ChIP-Seq(GSE17954)/Homer | 1e-72 | -1.669e+02 | 0.0000 | 6233.0 | 15.74% | 22268.4 | 12.30% | motif file (matrix) | svg |
| 222 | T C A G A T C G G A C T A C T G G A C T C A G T C T A G C G T A G T A C C G T A C T A G A T C G | Tbx20(T-box)/Heart-Tbx20-ChIP-Seq(GSE29636)/Homer | 1e-72 | -1.665e+02 | 0.0000 | 1707.0 | 4.31% | 4613.7 | 2.55% | motif file (matrix) | svg |
| 223 | C G A T T C G A A T G C C G A T G C A T T C G A A G C T G C A T G C A T C G A T T C G A A G C T T C G A G T C A C T A G | ANAC004(NAC)/colamp-ANAC004-DAP-Seq(GSE60143)/Homer | 1e-72 | -1.659e+02 | 0.0000 | 1999.0 | 5.05% | 5661.9 | 3.13% | motif file (matrix) | svg |
| 224 | G C T A C G T A G C T A C G T A C T G A A C T G A C G T G T A C C G T A C T G A G T A C A C T G | WRKY22(WRKY)/colamp-WRKY22-DAP-Seq(GSE60143)/Homer | 1e-71 | -1.646e+02 | 0.0000 | 4204.0 | 10.62% | 14097.5 | 7.79% | motif file (matrix) | svg |
| 225 | C A G T C G T A C G T A G C A T G A C T G C A T G T A C A G C T A C T G G A C T A C G T C A T G | RAV1(RAV)/colamp-RAV1-DAP-Seq(GSE60143)/Homer | 1e-71 | -1.636e+02 | 0.0000 | 3631.0 | 9.17% | 11855.3 | 6.55% | motif file (matrix) | svg |
| 226 | A T G C C A T G A C G T A C G T A C T G C G T A A G T C G A C T G C A T C G A T | WRKY71(WRKY)/col-WRKY71-DAP-Seq(GSE60143)/Homer | 1e-70 | -1.618e+02 | 0.0000 | 5537.0 | 13.98% | 19506.5 | 10.78% | motif file (matrix) | svg |
| 227 | T C A G T G A C C A T G G C A T C A G T A C T G C G T A T G A C G A C T C G A T C G A T C G T A | WRKY3(WRKY)/col-WRKY3-DAP-Seq(GSE60143)/Homer | 1e-70 | -1.616e+02 | 0.0000 | 4819.0 | 12.17% | 16600.1 | 9.17% | motif file (matrix) | svg |
| 228 | G C T A C G T A C G A T C A G T A C T G C G A T G T A C A C T G A T C G G A C T C A T G C T A G G C A T C A G T C A T G | DEAR5(AP2EREBP)/col-DEAR5-DAP-Seq(GSE60143)/Homer | 1e-70 | -1.613e+02 | 0.0000 | 3246.0 | 8.20% | 10389.9 | 5.74% | motif file (matrix) | svg |
| 229 | C G A T C G T A A G T C A C G T A C G T T C A G G C A T C G T A G C T A G C T A C G T A A G T C C G T A G T C A A C T G | ANAC058(NAC)/col-ANAC058-DAP-Seq(GSE60143)/Homer | 1e-69 | -1.612e+02 | 0.0000 | 4444.0 | 11.22% | 15103.1 | 8.34% | motif file (matrix) | svg |
| 230 | C T G A T A C G G C A T A G C T A G C T A G T C T C G A A C T G C A G T A G C T A G C T G A T C | IRF3(IRF)/BMDM-Irf3-ChIP-Seq(GSE67343)/Homer | 1e-69 | -1.610e+02 | 0.0000 | 1422.0 | 3.59% | 3665.2 | 2.03% | motif file (matrix) | svg |
| 231 | A G T C G A C T A G C T C G A T A T C G G C T A C G A T A T C G C G A T A C T G T A C G A C G T | Tcf7(HMG)/GM12878-TCF7-ChIP-Seq(Encode)/Homer | 1e-69 | -1.594e+02 | 0.0000 | 2118.0 | 5.35% | 6156.8 | 3.40% | motif file (matrix) | svg |
| 232 | C T G A A T C G G T A C C T G A A G T C A G T C A C T G C G T A A G T C C T G A | TINY(AP2EREBP)/col-TINY-DAP-Seq(GSE60143)/Homer | 1e-69 | -1.594e+02 | 0.0000 | 4949.0 | 12.50% | 17160.9 | 9.48% | motif file (matrix) | svg |
| 233 | T G A C C T G A A G T C A G T C A C T G G A T C G A C T G C A T | At5g18450(AP2EREBP)/col-At5g18450-DAP-Seq(GSE60143)/Homer | 1e-68 | -1.576e+02 | 0.0000 | 15746.0 | 39.77% | 63471.5 | 35.07% | motif file (matrix) | svg |
| 234 | A G T C C G T A A C G T A G T C A C G T A C T G | Tal1 | 1e-67 | -1.564e+02 | 0.0000 | 7584.0 | 19.15% | 28086.1 | 15.52% | motif file (matrix) | svg |
| 235 | A T G C G C A T T A G C C G A T T A G C G C A T T A G C G C A T A T G C G A C T | GAGA-repeat/Arabidopsis-Promoters/Homer | 1e-67 | -1.556e+02 | 0.0000 | 4935.0 | 12.46% | 17164.3 | 9.48% | motif file (matrix) | svg |
| 236 | C A G T C A T G T G C A G T A C C G T A T C A G G T A C G A C T T C A G C T G A | bZIP18(bZIP)/colamp-bZIP18-DAP-Seq(GSE60143)/Homer | 1e-67 | -1.549e+02 | 0.0000 | 25500.0 | 64.40% | 108060.3 | 59.71% | motif file (matrix) | svg |
| 237 | A G C T G C T A T G C A A G T C A C G T A C G T A C G T C G A T A G C T T C A G | dof24(C2C2dof)/col-dof24-DAP-Seq(GSE60143)/Homer | 1e-66 | -1.539e+02 | 0.0000 | 11802.0 | 29.81% | 46202.2 | 25.53% | motif file (matrix) | svg |
| 238 | G A T C G C A T T C G A A G T C A C G T A C G T A C G T C G A T A C G T A T C G | AT1G47655(C2C2dof)/colamp-AT1G47655-DAP-Seq(GSE60143)/Homer | 1e-66 | -1.520e+02 | 0.0000 | 18548.0 | 46.84% | 76195.3 | 42.10% | motif file (matrix) | svg |
| 239 | T C G A T G A C G C A T A G C T C A G T G A T C G C T A G A T C G A C T A C G T G C A T A G T C | PRDM1(Zf)/Hela-PRDM1-ChIP-Seq(GSE31477)/Homer | 1e-65 | -1.517e+02 | 0.0000 | 2538.0 | 6.41% | 7800.5 | 4.31% | motif file (matrix) | svg |
| 240 | G T C A T G C A G C T A A G T C C G T A A C T G T G A C G C A T T C A G C A G T | Ap4(bHLH)/AML-Tfap4-ChIP-Seq(GSE45738)/Homer | 1e-65 | -1.508e+02 | 0.0000 | 4703.0 | 11.88% | 16306.4 | 9.01% | motif file (matrix) | svg |
| 241 | T A C G T G C A A G T C C G T A A C G T T G A C A C G T A C T G A C T G G C A T | TCF4(bHLH)/SHSY5Y-TCF4-ChIP-Seq(GSE96915)/Homer | 1e-65 | -1.499e+02 | 0.0000 | 7350.0 | 18.56% | 27237.8 | 15.05% | motif file (matrix) | svg |
| 242 | C T G A A G T C G A T C C A T G G C T A G A T C C T G A G C T A G C T A C G A T | AT1G77200(AP2EREBP)/colamp-AT1G77200-DAP-Seq(GSE60143)/Homer | 1e-63 | -1.464e+02 | 0.0000 | 13102.0 | 33.09% | 52079.7 | 28.78% | motif file (matrix) | svg |
| 243 | G C T A G C T A C G T A C G T A C T G A C T A G A C G T A G T C C G T A C T G A G T A C A C T G | WRKY65(WRKY)/colamp-WRKY65-DAP-Seq(GSE60143)/Homer | 1e-62 | -1.444e+02 | 0.0000 | 3792.0 | 9.58% | 12760.0 | 7.05% | motif file (matrix) | svg |
| 244 | G A T C G A T C A G T C G T A C C G A T G T A C G T A C A G T C A G T C A G T C G C T A G A T C | ZNF148(Zf)/MDAMB231-ZNF148-ChIP-Seq(GSE147020)/Homer | 1e-62 | -1.430e+02 | 0.0000 | 2652.0 | 6.70% | 8335.8 | 4.61% | motif file (matrix) | svg |
| 245 | G C T A T C G A C G T A C T G A A C T G A C G T A G T C C G T A C G T A A G T C C T A G T G C A | WRKY42(WRKY)/colamp-WRKY42-DAP-Seq(GSE60143)/Homer | 1e-60 | -1.400e+02 | 0.0000 | 3743.0 | 9.45% | 12630.4 | 6.98% | motif file (matrix) | svg |
| 246 | C G A T G A T C G A T C C T G A G A T C G A T C C A T G T G C A G T A C T C G A G T C A G C A T C G A T C G A T G C A T | At4g32800(AP2EREBP)/colamp-At4g32800-DAP-Seq(GSE60143)/Homer | 1e-58 | -1.345e+02 | 0.0000 | 2627.0 | 6.63% | 8344.4 | 4.61% | motif file (matrix) | svg |
| 247 | C G A T C G T A G C T A G C A T G C A T C T G A A C T G A C G T A G T C C G T A C G T A G T A C T C A G G C T A C G A T | WRKY25(WRKY)/colamp-WRKY25-DAP-Seq(GSE60143)/Homer | 1e-58 | -1.343e+02 | 0.0000 | 8236.0 | 20.80% | 31321.4 | 17.31% | motif file (matrix) | svg |
| 248 | G A C T C G A T T C A G G A T C G A C T A G C T A G C T A G T C G A T C C G T A C T A G C T A G T C G A T C G A C T G A | Bcl6(Zf)/Liver-Bcl6-ChIP-Seq(GSE31578)/Homer | 1e-57 | -1.333e+02 | 0.0000 | 4534.0 | 11.45% | 15914.1 | 8.79% | motif file (matrix) | svg |
| 249 | G A C T C T A G A T G C A G T C G T C A T A C G A T G C A T C G | HIC1(Zf)/Treg-ZBTB29-ChIP-Seq(GSE99889)/Homer | 1e-56 | -1.305e+02 | 0.0000 | 12633.0 | 31.90% | 50452.1 | 27.88% | motif file (matrix) | svg |
| 250 | T C G A A G T C A C G T A C G T T C A G C A G T C T G A C T A G T C G A C G T A A T C G C G T A C G T A A C T G A G C T | NTM1(NAC)/col-NTM1-DAP-Seq(GSE60143)/Homer | 1e-55 | -1.288e+02 | 0.0000 | 2631.0 | 6.64% | 8432.1 | 4.66% | motif file (matrix) | svg |
| 251 | G T A C C A T G T A G C A G T C C T A G C A T G C T G A C G T A G C A T G C A T A C G T G C A T G T A C A C T G A T C G | LOB(LOBAS2)/col-LOB-DAP-Seq(GSE60143)/Homer | 1e-55 | -1.287e+02 | 0.0000 | 4113.0 | 10.39% | 14290.6 | 7.90% | motif file (matrix) | svg |
| 252 | A T G C A G T C C T G A A G T C C G A T A C G T A G T C A G T C A C G T A T C G G A C T A C G T | Etv2(ETS)/ES-ER71-ChIP-Seq(GSE59402)/Homer | 1e-55 | -1.280e+02 | 0.0000 | 4387.0 | 11.08% | 15410.7 | 8.51% | motif file (matrix) | svg |
| 253 | C G T A C G T A C G T A C T G A C T A G A C G T C T A G G T C A | CDF3(C2C2dof)/colamp-CDF3-DAP-Seq(GSE60143)/Homer | 1e-54 | -1.258e+02 | 0.0000 | 9753.0 | 24.63% | 38032.9 | 21.01% | motif file (matrix) | svg |
| 254 | C G A T T C G A A G T C A C G T A C G T T A C G C G T A G C T A G C T A C G A T G C A T A T G C C G T A G T C A A C T G | ANAC071(NAC)/col-ANAC071-DAP-Seq(GSE60143)/Homer | 1e-54 | -1.252e+02 | 0.0000 | 6772.0 | 17.10% | 25332.2 | 14.00% | motif file (matrix) | svg |
| 255 | A G C T A G C T A G C T A C T G A C G T A G T C A C T G A C G T G A C T C G A T G C A T A T C G | IDD7(C2H2)/col-IDD7-DAP-Seq(GSE60143)/Homer | 1e-54 | -1.249e+02 | 0.0000 | 2678.0 | 6.76% | 8664.8 | 4.79% | motif file (matrix) | svg |
| 256 | G C A T A G C T A G C T A G C T A C T G A C G T A G T C A C T G A C G T G A C T C G A T G C A T | JKD(C2H2)/col-JKD-DAP-Seq(GSE60143)/Homer | 1e-53 | -1.238e+02 | 0.0000 | 1498.0 | 3.78% | 4244.0 | 2.34% | motif file (matrix) | svg |
| 257 | C T A G A C T G T G C A G T C A A T G C C G T A A T C G A T G C A G T C C T A G | ZNF341(Zf)/EBV-ZNF341-ChIP-Seq(GSE113194)/Homer | 1e-53 | -1.224e+02 | 0.0000 | 4213.0 | 10.64% | 14801.5 | 8.18% | motif file (matrix) | svg |
| 258 | C G A T A G T C C A T G G A C T A C G T C T A G C G T A G A T C G A C T C G A T G C A T G A C T | WRKY43(WRKY)/colamp-WRKY43-DAP-Seq(GSE60143)/Homer | 1e-53 | -1.223e+02 | 0.0000 | 2618.0 | 6.61% | 8466.6 | 4.68% | motif file (matrix) | svg |
| 259 | A G T C G T A C C T G A A G T C A G T C C A T G G C T A A G T C T G C A G C T A G C A T G C A T | RAP21(AP2EREBP)/colamp-RAP21-DAP-Seq(GSE60143)/Homer | 1e-52 | -1.212e+02 | 0.0000 | 3674.0 | 9.28% | 12648.0 | 6.99% | motif file (matrix) | svg |
| 260 | A T G C G A C T A C G T C T A G A C G T A C G T A C G T C T G A G A T C G C T A A G C T C G T A | Foxa2(Forkhead)/Liver-Foxa2-ChIP-Seq(GSE25694)/Homer | 1e-52 | -1.211e+02 | 0.0000 | 4141.0 | 10.46% | 14532.4 | 8.03% | motif file (matrix) | svg |
| 261 | G A C T A C T G C G T A A G T C T C A G G C A T G T A C C G T A A C G T G A T C | TGA1(bZIP)/colamp-TGA1-DAP-Seq(GSE60143)/Homer | 1e-51 | -1.196e+02 | 0.0000 | 3707.0 | 9.36% | 12806.2 | 7.08% | motif file (matrix) | svg |
| 262 | G A C T T C G A C G T A C G T A C G T A C G T A C G T A C T A G A G C T C G T A | dof45(C2C2dof)/col-dof45-DAP-Seq(GSE60143)/Homer | 1e-51 | -1.195e+02 | 0.0000 | 13185.0 | 33.30% | 53199.5 | 29.39% | motif file (matrix) | svg |
| 263 | A G T C A C G T A C G T T A C G G C T A G C T A C G T A C G A T C G A T A T G C C G T A G T C A A C T G G A C T G C A T | SND2(NAC)/colamp-SND2-DAP-Seq(GSE60143)/Homer | 1e-51 | -1.191e+02 | 0.0000 | 4617.0 | 11.66% | 16506.7 | 9.12% | motif file (matrix) | svg |
| 264 | G A C T A C G T A C G T A C T G A C G T A G T C G C A T A G C T G C A T G C A T G A C T A G C T | SGR5(C2H2)/colamp-SGR5-DAP-Seq(GSE60143)/Homer | 1e-50 | -1.171e+02 | 0.0000 | 3662.0 | 9.25% | 12667.0 | 7.00% | motif file (matrix) | svg |
| 265 | G C T A G C T A C T G A A C T G A C G T A G T C C G T A C G T A G T A C A C T G A T G C G C A T | WRKY47(WRKY)/colamp-WRKY47-DAP-Seq(GSE60143)/Homer | 1e-50 | -1.165e+02 | 0.0000 | 2596.0 | 6.56% | 8459.6 | 4.67% | motif file (matrix) | svg |
| 266 | C G A T C T A G A G T C A C G T A C G T T C A G G C T A C G T A G C A T G C A T C G A T A G T C C G T A G T C A A C T G | VND3(NAC)/colamp-VND3-DAP-Seq(GSE60143)/Homer | 1e-50 | -1.165e+02 | 0.0000 | 4652.0 | 11.75% | 16698.5 | 9.23% | motif file (matrix) | svg |
| 267 | C G A T T C A G G T A C A C G T A C G T T C A G C G A T C G T A G T C A G C T A C G T A A G T C C G T A G T C A C A T G | ANAC057(NAC)/colamp-ANAC057-DAP-Seq(GSE60143)/Homer | 1e-50 | -1.159e+02 | 0.0000 | 5913.0 | 14.93% | 21932.7 | 12.12% | motif file (matrix) | svg |
| 268 | G A C T G C A T G C A T A G T C A G C T T C G A T A C G G C T A C G T A A C T G G T A C G C A T C G A T A G T C G A C T | HSF3(HSF)/colamp-HSF3-DAP-Seq(GSE60143)/Homer | 1e-49 | -1.135e+02 | 0.0000 | 4463.0 | 11.27% | 15980.6 | 8.83% | motif file (matrix) | svg |
| 269 | A T G C G T A C A C T G A G T C A G T C A C T G G A T C G T C A C G T A C G A T G C A T C G A T | RRTF1(AP2EREBP)/colamp-RRTF1-DAP-Seq(GSE60143)/Homer | 1e-48 | -1.126e+02 | 0.0000 | 4398.0 | 11.11% | 15730.3 | 8.69% | motif file (matrix) | svg |
| 270 | A C G T T C G A T C G A A G T C G T C A T A C G A T G C A C G T A C T G A G C T | Myf5(bHLH)/GM-Myf5-ChIP-Seq(GSE24852)/Homer | 1e-48 | -1.122e+02 | 0.0000 | 2709.0 | 6.84% | 8959.0 | 4.95% | motif file (matrix) | svg |
| 271 | T C G A T C G A C T G A C G T A A C T G A T G C A C G T A G T C | Lola-I(Zf)/Embryo-LolaI-ChIP-Seq(GSE200870)/Homer | 1e-48 | -1.119e+02 | 0.0000 | 3085.0 | 7.79% | 10446.3 | 5.77% | motif file (matrix) | svg |
| 272 | G A T C C A T G A C G T A C G T A C T G C G T A A G T C A G C T C G A T G A C T | WRKY8(WRKY)/colamp-WRKY8-DAP-Seq(GSE60143)/Homer | 1e-48 | -1.115e+02 | 0.0000 | 878.0 | 2.22% | 2176.6 | 1.20% | motif file (matrix) | svg |
| 273 | G C A T G C T A C G T A A G C T G C T A T G C A A G T C A C G T A C G T A C G T G C A T G C A T | At4g38000(C2C2dof)/col-At4g38000-DAP-Seq(GSE60143)/Homer | 1e-48 | -1.109e+02 | 0.0000 | 5047.0 | 12.75% | 18434.9 | 10.19% | motif file (matrix) | svg |
| 274 | C G T A T A C G T C G A A C T G A C T G C G T A C G T A T A C G A G C T T A C G | PU.1(ETS)/ThioMac-PU.1-ChIP-Seq(GSE21512)/Homer | 1e-47 | -1.099e+02 | 0.0000 | 2360.0 | 5.96% | 7632.3 | 4.22% | motif file (matrix) | svg |
| 275 | A G T C G A T C A G T C C G T A A T C G C A G T A G T C G T A C C T G A A C T G T C A G A G C T A G C T A G C T A G C T | PRDM15(Zf)/ESC-Prdm15-ChIP-Seq(GSE73694)/Homer | 1e-47 | -1.099e+02 | 0.0000 | 6059.0 | 15.30% | 22675.1 | 12.53% | motif file (matrix) | svg |
| 276 | G T A C G C T A C G A T C A G T A G T C G C T A C G A T C G A T A G T C G C T A | WUS1(Homeobox)/colamp-WUS1-DAP-Seq(GSE60143)/Homer | 1e-47 | -1.098e+02 | 0.0000 | 2573.0 | 6.50% | 8461.3 | 4.68% | motif file (matrix) | svg |
| 277 | C T A G C A T G A C G T C G T A C T A G C A T G C G A T C T A G T C A G T C A G | MYB3(MYB)/Arabidopsis-MYB3-ChIP-Seq(GSE80564)/Homer | 1e-47 | -1.096e+02 | 0.0000 | 13314.0 | 33.62% | 54072.9 | 29.88% | motif file (matrix) | svg |
| 278 | G C T A C T G A T C G A A G T C A G T C C T G A A G T C G T C A C T G A T G C A | RUNX1(Runt)/Jurkat-RUNX1-ChIP-Seq(GSE29180)/Homer | 1e-47 | -1.093e+02 | 0.0000 | 6401.0 | 16.17% | 24127.5 | 13.33% | motif file (matrix) | svg |
| 279 | G A C T A C G T A G C T G A C T A C T G C A G T A G T C A T C G A C G T G C A T G C A T G C A T | MGP(C2H2)/colamp-MGP-DAP-Seq(GSE60143)/Homer | 1e-46 | -1.078e+02 | 0.0000 | 2337.0 | 5.90% | 7571.1 | 4.18% | motif file (matrix) | svg |
| 280 | C T A G A G T C T A C G T A C G T G A C C G T A A C T G T A G C G C A T C A T G A T G C A G C T | Ascl1(bHLH)/NeuralTubes-Ascl1-ChIP-Seq(GSE55840)/Homer | 1e-46 | -1.074e+02 | 0.0000 | 6891.0 | 17.40% | 26242.7 | 14.50% | motif file (matrix) | svg |
| 281 | G A T C C G T A G A C T C T A G G A T C C T G A G A C T C T G A G A C T C T A G G A T C C T G A G A C T C T G A G A C T | OCT:OCT(POU,Homeobox)/NPC-OCT6-ChIP-Seq(GSE43916)/Homer | 1e-46 | -1.062e+02 | 0.0000 | 319.0 | 0.81% | 489.1 | 0.27% | motif file (matrix) | svg |
| 282 | G C A T A C G T A C G T A T C G C G T A C G T A C G T A C G T A | At2g41835(C2H2)/col-At2g41835-DAP-Seq(GSE60143)/Homer | 1e-46 | -1.061e+02 | 0.0000 | 2231.0 | 5.63% | 7184.2 | 3.97% | motif file (matrix) | svg |
| 283 | A G T C A C G T A C G T T C A G G C T A C G T A G C T A G C A T C G A T A G T C C G T A G T C A A C T G G A C T G C T A | SMB(NAC)/colamp-SMB-DAP-Seq(GSE60143)/Homer | 1e-45 | -1.058e+02 | 0.0000 | 6868.0 | 17.34% | 26184.8 | 14.47% | motif file (matrix) | svg |
| 284 | T G A C C A T G A C G T A C G T A C T G C G T A A G T C A G C T G C A T T C G A | WRKY30(WRKY)/colamp-WRKY30-DAP-Seq(GSE60143)/Homer | 1e-45 | -1.052e+02 | 0.0000 | 2951.0 | 7.45% | 10018.4 | 5.54% | motif file (matrix) | svg |
| 285 | C A G T T C A G A G C T G A C T A C G T A G T C G A T C G A C T C T G A A C T G G A T C C G T A C T G A A G T C G T A C | Rfx6(HTH)/Min6b1-Rfx6.HA-ChIP-Seq(GSE62844)/Homer | 1e-45 | -1.045e+02 | 0.0000 | 5940.0 | 15.00% | 22297.8 | 12.32% | motif file (matrix) | svg |
| 286 | C T G A C T A G A C T G G C A T A T G C C G T A C T G A C T A G A C T G A C G T A G T C C T G A | RARg(NR)/ES-RARg-ChIP-Seq(GSE30538)/Homer | 1e-44 | -1.035e+02 | 0.0000 | 242.0 | 0.61% | 307.0 | 0.17% | motif file (matrix) | svg |
| 287 | A T G C G A T C C G T A A G C T C A G T A T C G G C A T A G C T G A C T A C T G | Sox17(HMG)/Endoderm-Sox17-ChIP-Seq(GSE61475)/Homer | 1e-44 | -1.031e+02 | 0.0000 | 4830.0 | 12.20% | 17695.2 | 9.78% | motif file (matrix) | svg |
| 288 | C T A G T A C G G A T C G T A C G C T A A G C T A G C T G T C A T C G A T A G C | Nanog(Homeobox)/mES-Nanog-ChIP-Seq(GSE11724)/Homer | 1e-44 | -1.024e+02 | 0.0000 | 25025.0 | 63.20% | 107492.9 | 59.39% | motif file (matrix) | svg |
| 289 | T C G A T C G A A G T C C G T A C T A G T A G C A C G T A C T G | MyoG(bHLH)/C2C12-MyoG-ChIP-Seq(GSE36024)/Homer | 1e-43 | -1.009e+02 | 0.0000 | 4282.0 | 10.81% | 15478.2 | 8.55% | motif file (matrix) | svg |
| 290 | C A G T A G C T G A C T T G C A A G T C A G C T A C G T A C G T C G A T G A C T | AT3G52440(C2C2dof)/colamp-AT3G52440-DAP-Seq(GSE60143)/Homer | 1e-43 | -1.005e+02 | 0.0000 | 12168.0 | 30.73% | 49298.2 | 27.24% | motif file (matrix) | svg |
| 291 | T G A C G C T A T C G A T G C A A G T C A G T C C G T A A G T C C G T A C T G A G C T A G T A C | RUNX2(Runt)/PCa-RUNX2-ChIP-Seq(GSE33889)/Homer | 1e-43 | -1.005e+02 | 0.0000 | 5083.0 | 12.84% | 18801.5 | 10.39% | motif file (matrix) | svg |
| 292 | A C T G T C A G A G C T G A C T C A T G A G T C A G T C G C T A C G A T C T A G T C A G G T A C C T G A T C G A | Rfx1(HTH)/NPC-H3K4me1-ChIP-Seq(GSE16256)/Homer | 1e-43 | -1.004e+02 | 0.0000 | 1201.0 | 3.03% | 3391.3 | 1.87% | motif file (matrix) | svg |
| 293 | G T A C A C G T A C G T T A C G A T G C C A T G T A C G G T A C T C A G A T G C C G T A G T C A A C T G A G C T G C T A | AT1G19040(NAC)/col-AT1G19040-DAP-Seq(GSE60143)/Homer | 1e-43 | -1.002e+02 | 0.0000 | 1039.0 | 2.62% | 2817.3 | 1.56% | motif file (matrix) | svg |
| 294 | C G T A G C T A C G A T C T A G A C G T G T C A C G T A C G T A A G T C C G T A T G C A T A C G | FoxL2(Forkhead)/Ovary-FoxL2-ChIP-Seq(GSE60858)/Homer | 1e-43 | -9.954e+01 | 0.0000 | 3907.0 | 9.87% | 13968.7 | 7.72% | motif file (matrix) | svg |
| 295 | G C A T C G A T C G T A G A T C C A T G A C G T A C G T A C T G C G T A A G T C A G C T G C A T G C A T C G T A G C T A | WRKY45(WRKY)/col-WRKY45-DAP-Seq(GSE60143)/Homer | 1e-43 | -9.942e+01 | 0.0000 | 2452.0 | 6.19% | 8136.3 | 4.50% | motif file (matrix) | svg |
| 296 | C G A T C G T A A G T C A C G T A C G T T C A G C G T A C G T A G C A T G C A T G C A T A G T C C G T A G T C A A C T G | VND2(NAC)/col-VND2-DAP-Seq(GSE60143)/Homer | 1e-42 | -9.801e+01 | 0.0000 | 6921.0 | 17.48% | 26602.9 | 14.70% | motif file (matrix) | svg |
| 297 | C G T A C T G A C G T A C T A G T C G A C T A G A C T G C G T A C G T A T A C G A G C T A T C G | SpiB(ETS)/OCILY3-SPIB-ChIP-Seq(GSE56857)/Homer | 1e-42 | -9.791e+01 | 0.0000 | 1293.0 | 3.27% | 3748.3 | 2.07% | motif file (matrix) | svg |
| 298 | C A T G A C T G C T A G T C G A T C G A T C G A T C G A T C A G T C A G T C A G T G A C T G A C C G T A A C T G T G C A C G A T A C T G | RBPJ:Ebox(?,bHLH)/Panc1-Rbpj1-ChIP-Seq(GSE47459)/Homer | 1e-42 | -9.765e+01 | 0.0000 | 1275.0 | 3.22% | 3685.6 | 2.04% | motif file (matrix) | svg |
| 299 | T A G C C A T G A G C T A C G T A C T G C G T A A G T C G A C T G C A T C T G A | AT3G42860(zfGRF)/col-AT3G42860-DAP-Seq(GSE60143)/Homer | 1e-42 | -9.732e+01 | 0.0000 | 2626.0 | 6.63% | 8852.5 | 4.89% | motif file (matrix) | svg |
| 300 | A C G T T G C A A G C T G A T C C T A G C T G A A G C T G T C A T C G A C G T A | CUX1(Homeobox)/K562-CUX1-ChIP-Seq(GSE92882)/Homer | 1e-42 | -9.708e+01 | 0.0000 | 7712.0 | 19.48% | 30006.3 | 16.58% | motif file (matrix) | svg |
| 301 | A T C G T C G A G A C T A T C G T G A C A C G T C T A G A C T G C G T A A C T G A G T C G T A C | ZNF415(Zf)/HEK293-ZNF415.GFP-ChIP-Seq(GSE58341)/Homer | 1e-42 | -9.684e+01 | 0.0000 | 3166.0 | 8.00% | 11013.3 | 6.09% | motif file (matrix) | svg |
| 302 | C T G A A T G C G C T A C G A T A T G C C G T A C G T A C G T A C T A G T A C G | Tcf3(HMG)/mES-Tcf3-ChIP-Seq(GSE11724)/Homer | 1e-41 | -9.497e+01 | 0.0000 | 1531.0 | 3.87% | 4658.8 | 2.57% | motif file (matrix) | svg |
| 303 | C A T G G T A C A C T G G T C A A G C T T A C G T G C A A T C G T G A C C A G T | TOD6?/SacCer-Promoters/Homer | 1e-41 | -9.476e+01 | 0.0000 | 1886.0 | 4.76% | 6005.5 | 3.32% | motif file (matrix) | svg |
| 304 | G C A T A C G T A C T G A C G T A G T C A C T G A T C G G T C A C G A T C G T A | ARF2(ARF)/col-ARF2-DAP-Seq(GSE60143)/Homer | 1e-40 | -9.272e+01 | 0.0000 | 19109.0 | 48.26% | 80656.6 | 44.56% | motif file (matrix) | svg |
| 305 | A G C T A G C T A G C T A C T G A C G T A G T C A C T G A C G T G C A T G C A T G C A T A C G T | At5g66730(C2H2)/colamp-At5g66730-DAP-Seq(GSE60143)/Homer | 1e-40 | -9.250e+01 | 0.0000 | 1959.0 | 4.95% | 6315.3 | 3.49% | motif file (matrix) | svg |
| 306 | G C T A T C G A C G T A C T A G A G C T G T C A G T C A C G T A A G T C C G T A | FOXA1(Forkhead)/LNCAP-FOXA1-ChIP-Seq(GSE27824)/Homer | 1e-40 | -9.224e+01 | 0.0000 | 4580.0 | 11.57% | 16886.5 | 9.33% | motif file (matrix) | svg |
| 307 | C G T A C G T A C G T A C G T A C G T A A C T G A C T G A G T C | dof42(C2C2dof)/col-dof42-DAP-Seq(GSE60143)/Homer | 1e-39 | -9.098e+01 | 0.0000 | 3989.0 | 10.07% | 14470.9 | 8.00% | motif file (matrix) | svg |
| 308 | A T G C C G T A C G T A C G T A C G T A C G T A A C T G A C G T C G A T C T G A | dof43(C2C2dof)/colamp-dof43-DAP-Seq(GSE60143)/Homer | 1e-39 | -9.071e+01 | 0.0000 | 8051.0 | 20.33% | 31639.2 | 17.48% | motif file (matrix) | svg |
| 309 | C G T A C G A T C T A G G T C A G A C T C G A T C T A G C G T A A C G T C A T G | LIN-39(Homeobox)/cElegans.L3-LIN39-ChIP-Seq(modEncode)/Homer | 1e-39 | -9.055e+01 | 0.0000 | 5896.0 | 14.89% | 22440.6 | 12.40% | motif file (matrix) | svg |
| 310 | T C G A C G T A A G T C C G T A C T A G A G T C C G A T A C T G G A C T A G C T A C T G G A C T | HLH-1(bHLH)/cElegans-Embryo-HLH1-ChIP-Seq(modEncode)/Homer | 1e-38 | -8.840e+01 | 0.0000 | 3928.0 | 9.92% | 14271.5 | 7.89% | motif file (matrix) | svg |
| 311 | T C G A C T G A C G A T C G T A C G T A C G T A C T A G A G C T C T G A T C A G | Adof1(C2C2dof)/col-Adof1-DAP-Seq(GSE60143)/Homer | 1e-37 | -8.721e+01 | 0.0000 | 14425.0 | 36.43% | 59761.5 | 33.02% | motif file (matrix) | svg |
| 312 | T G A C T A G C T C A G T C G A T C G A C G T A A G T C C G T A C G T A C G A T C T A G T A C G | Sox7(HMG)/ESC-Sox7-ChIP-Seq(GSE133899)/Homer | 1e-37 | -8.708e+01 | 0.0000 | 2275.0 | 5.75% | 7624.0 | 4.21% | motif file (matrix) | svg |
| 313 | T G A C C T G A C T A G T C G A C T G A A T G C C G T A A C T G G C A T G T A C G C A T A T C G G C A T A G C T G A T C | PR(NR)/T47D-PR-ChIP-Seq(GSE31130)/Homer | 1e-37 | -8.583e+01 | 0.0000 | 9925.0 | 25.07% | 39921.1 | 22.06% | motif file (matrix) | svg |
| 314 | T A C G C T G A C A T G G A T C G T A C G C A T T C A G T A C G A G C T G T C A G A T C G C A T T A C G C G T A C T A G G A T C G A T C C G A T A C T G T C A G | ZNF322(Zf)/HEK293-ZNF322.GFP-ChIP-Seq(GSE58341)/Homer | 1e-37 | -8.570e+01 | 0.0000 | 810.0 | 2.05% | 2136.5 | 1.18% | motif file (matrix) | svg |
| 315 | G A C T G T A C T G C A A C G T G A T C G C T A T C G A A C G T A G T C C G T A | Pdx1(Homeobox)/Islet-Pdx1-ChIP-Seq(SRA008281)/Homer | 1e-36 | -8.502e+01 | 0.0000 | 5686.0 | 14.36% | 21690.0 | 11.98% | motif file (matrix) | svg |
| 316 | G A C T C T A G C T A G A G T C T G C A A C T G A C G T A C G T C T A G T C A G | AMYB(HTH)/Testes-AMYB-ChIP-Seq(GSE44588)/Homer | 1e-36 | -8.310e+01 | 0.0000 | 14510.0 | 36.64% | 60291.1 | 33.31% | motif file (matrix) | svg |
| 317 | C G A T G A T C T A C G C T G A G C T A C G T A G C A T A G T C C T A G C G T A G C A T C G A T | AT2G15740(C2H2)/col-AT2G15740-DAP-Seq(GSE60143)/Homer | 1e-36 | -8.295e+01 | 0.0000 | 15672.0 | 39.58% | 65501.9 | 36.19% | motif file (matrix) | svg |
| 318 | T G C A C T G A A T G C G T C A A C G T A T G C A C G T A C T G A C T G T G C A | ZBTB18(Zf)/HEK293-ZBTB18.GFP-ChIP-Seq(GSE58341)/Homer | 1e-35 | -8.237e+01 | 0.0000 | 2123.0 | 5.36% | 7096.1 | 3.92% | motif file (matrix) | svg |
| 319 | T C G A A G C T A C G T A C G T A G T C A G T C A C G T A T C G G A C T A T C G | EWS:ERG-fusion(ETS)/CADO\_ES1-EWS:ERG-ChIP-Seq(SRA014231)/Homer | 1e-34 | -8.025e+01 | 0.0000 | 2837.0 | 7.16% | 9977.8 | 5.51% | motif file (matrix) | svg |
| 320 | C T G A C T G A C T G A A T G C G A T C C A T G A C T G G A C T G A C T G C A T C G T A C G T A A G T C G T A C C T G A A T C G G C A T G A C T G A C T A G C T | GRHL2(CP2)/HBE-GRHL2-ChIP-Seq(GSE46194)/Homer | 1e-34 | -7.998e+01 | 0.0000 | 2193.0 | 5.54% | 7408.6 | 4.09% | motif file (matrix) | svg |
| 321 | C T G A A C T G C G T A A C G T G T C A A G C T A G C T G A C T G A C T C A G T | CCA(Myb)/Arabidopsis-CCA.GFP-ChIP-Seq(GSE70533)/Homer | 1e-34 | -7.902e+01 | 0.0000 | 5313.0 | 13.42% | 20263.8 | 11.20% | motif file (matrix) | svg |
| 322 | G T A C T C G A T A G C C G T A C G T A C T G A T G C A T G A C A C T G C G T A A G T C C T G A C T G A T C G A C G T A | NUC(C2H2)/col-NUC-DAP-Seq(GSE60143)/Homer | 1e-34 | -7.878e+01 | 0.0000 | 743.0 | 1.88% | 1959.6 | 1.08% | motif file (matrix) | svg |
| 323 | T C G A C T G A C G T A C G T A C G T A C G T A A C T G A C G T C G A T C T G A | BBX31(Orphan)/col-BBX31-DAP-Seq(GSE60143)/Homer | 1e-34 | -7.866e+01 | 0.0000 | 8588.0 | 21.69% | 34323.5 | 18.96% | motif file (matrix) | svg |
| 324 | C A G T T C G A A G T C A C G T A C G T T C A G C G A T G C T A G C T A C G T A G C A T A G T C C G T A T G C A A C T G | ANAC045(NAC)/col-ANAC045-DAP-Seq(GSE60143)/Homer | 1e-34 | -7.837e+01 | 0.0000 | 14171.0 | 35.79% | 58955.0 | 32.57% | motif file (matrix) | svg |
| 325 | T C G A C T G A T A G C T G A C T C A G T C A G C G T A C G T A T C A G A G C T | ETV1(ETS)/GIST48-ETV1-ChIP-Seq(GSE22441)/Homer | 1e-33 | -7.820e+01 | 0.0000 | 8467.0 | 21.38% | 33812.7 | 18.68% | motif file (matrix) | svg |
| 326 | A T G C T C G A A G T C A G C T A C G T G T A C A G T C G C T A C T A G C A T G G T C A C T G A T C A G A G T C | Stat3+il21(Stat)/CD4-Stat3-ChIP-Seq(GSE19198)/Homer | 1e-33 | -7.763e+01 | 0.0000 | 3039.0 | 7.67% | 10844.8 | 5.99% | motif file (matrix) | svg |
| 327 | T C G A T G A C G T A C C G T A A C G T G A C T A C G T A C T G A C T G A G C T | Mesp1(bHLH)/ESC-Mesp1-ChIP-Seq(GSE165102)/Homer | 1e-33 | -7.669e+01 | 0.0000 | 3120.0 | 7.88% | 11192.4 | 6.18% | motif file (matrix) | svg |
| 328 | G T C A T C G A C T A G C T A G A G T C G T C A C G A T C T A G G A C T G A T C G A T C T C A G C T A G C T G A A G T C G C T A C A G T T C A G G A T C G A T C | p63(p53)/Keratinocyte-p63-ChIP-Seq(GSE17611)/Homer | 1e-33 | -7.602e+01 | 0.0000 | 2236.0 | 5.65% | 7641.1 | 4.22% | motif file (matrix) | svg |
| 329 | C A G T G C T A G C A T T A C G C T G A C A G T T A G C C T G A | GATA15(C2C2gata)/col-GATA15-DAP-Seq(GSE60143)/Homer | 1e-32 | -7.567e+01 | 0.0000 | 12788.0 | 32.30% | 52895.3 | 29.23% | motif file (matrix) | svg |
| 330 | T A G C G T A C C T A G C A G T T C G A C G T A C G T A G C A T G A C T T G A C A G T C A C T G A T C G A G T C C T A G | AS2(LOBAS2)/col-AS2-DAP-Seq(GSE60143)/Homer | 1e-32 | -7.417e+01 | 0.0000 | 1387.0 | 3.50% | 4360.6 | 2.41% | motif file (matrix) | svg |
| 331 | G T A C G C T A T C A G C T G A C T A G C A T G A G C T G A T C T G C A T C G A C T G A A C T G C A G T A G T C G A T C G C T A | HNF4a(NR),DR1/HepG2-HNF4a-ChIP-Seq(GSE25021)/Homer | 1e-32 | -7.403e+01 | 0.0000 | 1798.0 | 4.54% | 5947.0 | 3.29% | motif file (matrix) | svg |
| 332 | A G T C C G A T A C T G A T C G T G A C G C T A C A T G A T C G T G A C C G A T A C T G T A G C G T A C G T C A | Tlx?(NR)/NPC-H3K4me1-ChIP-Seq(GSE16256)/Homer | 1e-31 | -7.364e+01 | 0.0000 | 1519.0 | 3.84% | 4871.9 | 2.69% | motif file (matrix) | svg |
| 333 | C G A T C T G A A G T C A C G T A C G T T C A G G C A T C G A T G C T A G C T A C G T A A G T C C G T A G T C A A C T G | CUC1(NAC)/col-CUC1-DAP-Seq(GSE60143)/Homer | 1e-31 | -7.339e+01 | 0.0000 | 3556.0 | 8.98% | 13049.2 | 7.21% | motif file (matrix) | svg |
| 334 | T A G C T A G C G A C T C T A G A G C T A G T C G T C A T G C A A C G T A T G C G C T A T G C A | Pbx3(Homeobox)/GM12878-PBX3-ChIP-Seq(GSE32465)/Homer | 1e-31 | -7.334e+01 | 0.0000 | 1321.0 | 3.34% | 4120.0 | 2.28% | motif file (matrix) | svg |
| 335 | T C A G A G C T A C G T A C G T G T A C G A T C C G T A C T A G C A T G G T C A C G T A T C G A | STAT4(Stat)/CD4-Stat4-ChIP-Seq(GSE22104)/Homer | 1e-31 | -7.326e+01 | 0.0000 | 3773.0 | 9.53% | 13950.4 | 7.71% | motif file (matrix) | svg |
| 336 | C G A T G T C A A G T C A C G T A C G T A C T G G A C T C G A T A T C G G C T A G T C A A G T C C G T A G T C A A C T G | ANAC017(NAC)/colamp-ANAC017-DAP-Seq(GSE60143)/Homer | 1e-31 | -7.273e+01 | 0.0000 | 1150.0 | 2.90% | 3485.8 | 1.93% | motif file (matrix) | svg |
| 337 | A C G T C T A G A G C T A C G T A C G T C T G A A G T C G A C T A G C T C G T A | FOXM1(Forkhead)/MCF7-FOXM1-ChIP-Seq(GSE72977)/Homer | 1e-31 | -7.241e+01 | 0.0000 | 4104.0 | 10.36% | 15346.9 | 8.48% | motif file (matrix) | svg |
| 338 | C A T G T G A C C A T G G A C T C A G T C T A G G C T A G T A C G A C T G C A T G C A T C G A T | WRKY21(WRKY)/colamp-WRKY21-DAP-Seq(GSE60143)/Homer | 1e-31 | -7.154e+01 | 0.0000 | 789.0 | 1.99% | 2181.2 | 1.21% | motif file (matrix) | svg |
| 339 | T C A G C T G A C G T A C G T A T A C G G C A T C T A G C T G A C G T A C G T A T A C G G A C T | IRF1(IRF)/PBMC-IRF1-ChIP-Seq(GSE43036)/Homer | 1e-30 | -7.011e+01 | 0.0000 | 578.0 | 1.46% | 1461.0 | 0.81% | motif file (matrix) | svg |
| 340 | C A G T T A G C A G T C C A T G C A G T C A T G C G A T C G A T G A C T C G A T A T C G G T A C A C T G A T C G G T A C | LBD13(LOBAS2)/colamp-LBD13-DAP-Seq(GSE60143)/Homer | 1e-30 | -6.987e+01 | 0.0000 | 10069.0 | 25.43% | 41092.3 | 22.70% | motif file (matrix) | svg |
| 341 | C A T G G A T C C T G A G T A C C T A G C T G A G C T A G C A T G A T C G A T C A G T C C T A G C G T A C A T G C T A G | AIL7(AP2EREBP)/colamp-AIL7-DAP-Seq(GSE60143)/Homer | 1e-29 | -6.760e+01 | 0.0000 | 4246.0 | 10.72% | 16055.9 | 8.87% | motif file (matrix) | svg |
| 342 | G T A C A C G T A C G T T C A G G C T A C G T A C G A T G C A T G C A T A G T C C G T A G T C A C A T G G A C T G C T A | ANAC070(NAC)/colamp-ANAC070-DAP-Seq(GSE60143)/Homer | 1e-29 | -6.753e+01 | 0.0000 | 7520.0 | 18.99% | 30045.8 | 16.60% | motif file (matrix) | svg |
| 343 | C A G T G A C T G C A T T C G A A G T C A C G T A C G T A C G T C G A T G A C T | OBP3(C2C2dof)/col-OBP3-DAP-Seq(GSE60143)/Homer | 1e-29 | -6.733e+01 | 0.0000 | 15215.0 | 38.42% | 64071.4 | 35.40% | motif file (matrix) | svg |
| 344 | C A T G C T A G A G C T G A C T C A T G A G T C G A T C G C T A C G A T C T A G T C A G G T A C C T G A T C G A | X-box(HTH)/NPC-H3K4me1-ChIP-Seq(GSE16256)/Homer | 1e-29 | -6.728e+01 | 0.0000 | 517.0 | 1.31% | 1275.6 | 0.70% | motif file (matrix) | svg |
| 345 | T G A C C G A T C T G A C T A G C T A G A C G T A T G C T G C A T C G A C T G A C T A G C A T G A C G T A G T C C G T A | PPARa(NR),DR1/Liver-Ppara-ChIP-Seq(GSE47954)/Homer | 1e-28 | -6.654e+01 | 0.0000 | 4590.0 | 11.59% | 17529.7 | 9.69% | motif file (matrix) | svg |
| 346 | T A C G T C G A G A C T A C T G C T G A A G T C T C A G G A C T T G A C C T G A | Atf1(bZIP)/K562-ATF1-ChIP-Seq(GSE31477)/Homer | 1e-28 | -6.622e+01 | 0.0000 | 5677.0 | 14.34% | 22153.2 | 12.24% | motif file (matrix) | svg |
| 347 | A G T C A G T C C T G A A G T C A G T C A C T G C G T A A G T C T C G A G A T C C G A T C G T A | AT1G01250(AP2EREBP)/col-AT1G01250-DAP-Seq(GSE60143)/Homer | 1e-28 | -6.594e+01 | 0.0000 | 1587.0 | 4.01% | 5241.0 | 2.90% | motif file (matrix) | svg |
| 348 | G C A T C T A G G T A C A G T C C G A T A C T G C T A G C T A G G T A C G C T A | ZNF416(Zf)/HEK293-ZNF416.GFP-ChIP-Seq(GSE58341)/Homer | 1e-28 | -6.586e+01 | 0.0000 | 4687.0 | 11.84% | 17956.2 | 9.92% | motif file (matrix) | svg |
| 349 | G C A T G C A T G C A T A T G C A G C T T C G A T A C G G C T A C G T A C A T G G T A C G C A T G C A T A G T C A G C T | HSFA6B(HSF)/colamp-HSFA6B-DAP-Seq(GSE60143)/Homer | 1e-28 | -6.572e+01 | 0.0000 | 2243.0 | 5.66% | 7843.6 | 4.33% | motif file (matrix) | svg |
| 350 | G C T A T C G A C G T A C T A G A G C T G T C A G T C A C G T A A G T C C G T A | FOXA1(Forkhead)/MCF7-FOXA1-ChIP-Seq(GSE26831)/Homer | 1e-28 | -6.542e+01 | 0.0000 | 3383.0 | 8.54% | 12507.4 | 6.91% | motif file (matrix) | svg |
| 351 | G T A C A C G T A C G T T C A G C G T A C G T A C G A T G C A T G C A T A G T C C G T A G T C A C A T G G A C T G C T A | VND1(NAC)/col-VND1-DAP-Seq(GSE60143)/Homer | 1e-28 | -6.518e+01 | 0.0000 | 5389.0 | 13.61% | 20954.9 | 11.58% | motif file (matrix) | svg |
| 352 | C G A T C T A G T C A G C A G T C G T A A G T C G C T A A C G T G A C T A T G C A G T C G C T A | PRDM10(Zf)/HEK293-PRDM10.eGFP-ChIP-Seq(Encode)/Homer | 1e-27 | -6.407e+01 | 0.0000 | 3091.0 | 7.81% | 11330.6 | 6.26% | motif file (matrix) | svg |
| 353 | T C A G G A C T G T C A C G T A A C G T A T C G C G T A A C G T A C G T C T G A | ATHB15(HB)/col-ATHB15-DAP-Seq(GSE60143)/Homer | 1e-27 | -6.351e+01 | 0.0000 | 2090.0 | 5.28% | 7268.0 | 4.02% | motif file (matrix) | svg |
| 354 | G C T A T G A C G A T C C G A T G A C T A T G C C T G A A T C G G C A T A C G T | JGL(C2H2)/col-JGL-DAP-Seq(GSE60143)/Homer | 1e-27 | -6.343e+01 | 0.0000 | 8594.0 | 21.70% | 34858.4 | 19.26% | motif file (matrix) | svg |
| 355 | T C A G G C A T A C T G C G T A A G T C C T A G G C A T T G A C | TGA9(bZIP)/colamp-TGA9-DAP-Seq(GSE60143)/Homer | 1e-27 | -6.336e+01 | 0.0000 | 10860.0 | 27.43% | 44822.7 | 24.77% | motif file (matrix) | svg |
| 356 | G T C A T C G A T C G A C G T A G C T A C G T A T C G A T G A C A C T G C G T A A G T C C G T A C G T A T C G A G C T A | IDD2(C2H2)/colamp-IDD2-DAP-Seq(GSE60143)/Homer | 1e-27 | -6.313e+01 | 0.0000 | 845.0 | 2.13% | 2462.6 | 1.36% | motif file (matrix) | svg |
| 357 | C G A T C T G A A G T C A C G T A C G T T C A G C G T A C G T A C G T A G C A T C G A T A G T C C G T A G T C A A C T G | VND4(NAC)/colamp-VND4-DAP-Seq(GSE60143)/Homer | 1e-26 | -6.204e+01 | 0.0000 | 5371.0 | 13.56% | 20967.1 | 11.58% | motif file (matrix) | svg |
| 358 | A G T C C A G T T C A G A T G C A G T C C G A T C G T A G T C A G A T C G C A T | BOS1(MYB)/col-BOS1-DAP-Seq(GSE60143)/Homer | 1e-26 | -6.171e+01 | 0.0000 | 7960.0 | 20.10% | 32152.6 | 17.76% | motif file (matrix) | svg |
| 359 | G A C T A T C G C T G A A G T C T C A G G A C T G T A C C T G A A G C T G T A C | TGA6(bZIP)/colamp-TGA6-DAP-Seq(GSE60143)/Homer | 1e-26 | -6.148e+01 | 0.0000 | 6356.0 | 16.05% | 25209.3 | 13.93% | motif file (matrix) | svg |
| 360 | A T G C G A C T A G C T C T A G C G T A C T A G C G A T C T A G A T C G G A T C | Nkx2.2(Homeobox)/NPC-Nkx2.2-ChIP-Seq(GSE61673)/Homer | 1e-26 | -6.077e+01 | 0.0000 | 12442.0 | 31.42% | 51945.3 | 28.70% | motif file (matrix) | svg |
| 361 | T C G A G A C T A T C G C G T A A G T C C T A G G C A T G T A C C T G A A C G T G A T C G C T A | TGA4(bZIP)/colamp-TGA4-DAP-Seq(GSE60143)/Homer | 1e-25 | -5.966e+01 | 0.0000 | 2624.0 | 6.63% | 9504.1 | 5.25% | motif file (matrix) | svg |
| 362 | A G C T G A C T A C G T A C T G A C G T A G T C A C T G A C G T G C A T C G A T | AtIDD11(C2H2)/colamp-AtIDD11-DAP-Seq(GSE60143)/Homer | 1e-25 | -5.944e+01 | 0.0000 | 2538.0 | 6.41% | 9157.4 | 5.06% | motif file (matrix) | svg |
| 363 | A C T G A C G T C A T G A T C G A T C G T G A C A C T G A T C G A T C G T G C A C T G A C G T A | E2F3(E2F)/MEF-E2F3-ChIP-Seq(GSE71376)/Homer | 1e-25 | -5.938e+01 | 0.0000 | 5975.0 | 15.09% | 23635.7 | 13.06% | motif file (matrix) | svg |
| 364 | G C A T C G T A G C A T C G T A T C G A C G T A C T G A A C T G C G T A C G T A C G T A A C G T A C T G G T C A G C A T | AT2G31460(REMB3)/col-AT2G31460-DAP-Seq(GSE60143)/Homer | 1e-25 | -5.922e+01 | 0.0000 | 1550.0 | 3.91% | 5195.1 | 2.87% | motif file (matrix) | svg |
| 365 | C G T A C T A G T C A G T C A G A G T C A T G C A G T C G C A T A G C T A C G T A T C G C G A T | Sox9(HMG)/Limb-SOX9-ChIP-Seq(GSE73225)/Homer | 1e-25 | -5.802e+01 | 0.0000 | 4276.0 | 10.80% | 16428.0 | 9.08% | motif file (matrix) | svg |
| 366 | C A G T A T C G C T G A A G T C T C A G C A G T T A G C C T G A A T G C T A C G | FEA4(bZIP)/Corn-FEA4-ChIP-Seq(GSE61954)/Homer | 1e-24 | -5.756e+01 | 0.0000 | 8915.0 | 22.51% | 36479.0 | 20.16% | motif file (matrix) | svg |
| 367 | C G T A A C T G C G T A A C G T A T C G C A G T T A G C C G T A T C G A G T A C C T G A T A G C C G T A A C T G C G T A A C G T C G T A C T G A A T C G G C T A | GATA3(Zf),DR8/iTreg-Gata3-ChIP-Seq(GSE20898)/Homer | 1e-24 | -5.742e+01 | 0.0000 | 493.0 | 1.25% | 1265.2 | 0.70% | motif file (matrix) | svg |
| 368 | A G C T C T G A C T A G C T A G A C T G T A G C T G C A T C G A C T G A C T A G C A T G A C G T A T G C T C G A | RXR(NR),DR1/3T3L1-RXR-ChIP-Seq(GSE13511)/Homer | 1e-24 | -5.607e+01 | 0.0000 | 4619.0 | 11.67% | 17935.8 | 9.91% | motif file (matrix) | svg |
| 369 | A G T C A T C G C T A G A G C T G A C T C T A G A G T C A G T C G C T A C A G T T C A G T C A G G A T C C T G A T C G A G A T C | RFX(HTH)/K562-RFX3-ChIP-Seq(SRA012198)/Homer | 1e-24 | -5.582e+01 | 0.0000 | 458.0 | 1.16% | 1158.8 | 0.64% | motif file (matrix) | svg |
| 370 | T A C G T A G C C A T G C A G T A C G T C T A G C G T A A G T C G A C T G C A T G C A T C A G T | WRKY11(WRKY)/col-WRKY11-DAP-Seq(GSE60143)/Homer | 1e-24 | -5.539e+01 | 0.0000 | 1043.0 | 2.63% | 3287.9 | 1.82% | motif file (matrix) | svg |
| 371 | A G T C T A G C G A C T A C G T C T A G A C G T A C G T A C G T C T G A A G T C G C T A G A C T C G T A C T A G A C T G | Foxa3(Forkhead)/Liver-Foxa3-ChIP-Seq(GSE77670)/Homer | 1e-23 | -5.511e+01 | 0.0000 | 1451.0 | 3.66% | 4869.2 | 2.69% | motif file (matrix) | svg |
| 372 | A T G C T C A G T C G A G C A T A C T G C G T A A G T C T C A G G A C T T G A C C G T A A G C T | Atf2(bZIP)/3T3L1-Atf2-ChIP-Seq(GSE56872)/Homer | 1e-23 | -5.446e+01 | 0.0000 | 1931.0 | 4.88% | 6790.9 | 3.75% | motif file (matrix) | svg |
| 373 | T A G C G T A C A G T C G T A C C G A T A G T C A G T C A G T C A G T C A G T C C G T A G A T C | Zfp281(Zf)/ES-Zfp281-ChIP-Seq(GSE81042)/Homer | 1e-23 | -5.349e+01 | 0.0000 | 560.0 | 1.41% | 1529.7 | 0.85% | motif file (matrix) | svg |
| 374 | C T A G G T A C A C G T A C G T A T C G G C A T A G C T A G C T A G C T G C A T G A C T C G T A G T C A A C T G G A C T | VND6(NAC)/col-VND6-DAP-Seq(GSE60143)/Homer | 1e-23 | -5.321e+01 | 0.0000 | 8555.0 | 21.61% | 35067.4 | 19.38% | motif file (matrix) | svg |
| 375 | C G A T C T A G C T G A A T G C C T G A T C G A C G T A C T G A T C G A T A G C A G T C C G T A A C T G T C G A A T G C | Hand2(bHLH)/Mesoderm-Hand2-ChIP-Seq(GSE61475)/Homer | 1e-22 | -5.145e+01 | 0.0000 | 1815.0 | 4.58% | 6378.4 | 3.52% | motif file (matrix) | svg |
| 376 | C G A T T C G A G T A C A C G T A C G T T C A G G C A T G C T A T G C A G C A T C G T A A G T C C G T A T G A C C A T G | ANAC092(NAC)/colamp-ANAC092-DAP-Seq(GSE60143)/Homer | 1e-22 | -5.107e+01 | 0.0000 | 3543.0 | 8.95% | 13521.8 | 7.47% | motif file (matrix) | svg |
| 377 | C A G T A G C T C G T A G C A T A G T C G A C T C T A G C T A G C A G T C T A G T C G A T G C A C T A G C A T G G A C T | STOP1(C2H2)/colamp-STOP1-DAP-Seq(GSE60143)/Homer | 1e-22 | -5.101e+01 | 0.0000 | 2706.0 | 6.83% | 10028.5 | 5.54% | motif file (matrix) | svg |
| 378 | A T G C A G T C G T A C A G C T T C G A C T A G G A T C C T G A G T C A A G T C G C T A T C A G | Rfx5(HTH)/GM12878-Rfx5-ChIP-Seq(GSE31477)/Homer | 1e-21 | -5.058e+01 | 0.0000 | 1927.0 | 4.87% | 6846.1 | 3.78% | motif file (matrix) | svg |
| 379 | C T A G C T A G T C A G G T C A C T A G T C A G G C T A A G T C A T C G A G C T C T A G | DPR(core promoter) | 1e-21 | -5.053e+01 | 0.0000 | 33420.0 | 84.40% | 149099.0 | 82.38% | motif file (matrix) | svg |
| 380 | G T C A C G T A A C G T A T C G C G T A A C G T A C G T C T A G | ATHB7(Homeobox)/col-ATHB7-DAP-Seq(GSE60143)/Homer | 1e-21 | -5.049e+01 | 0.0000 | 4857.0 | 12.27% | 19116.9 | 10.56% | motif file (matrix) | svg |
| 381 | G A C T C A G T G A T C G A T C A C G T G A T C C T G A T A C G C G T A G T C A | STAT6(Stat)/Macrophage-Stat6-ChIP-Seq(GSE38377)/Homer | 1e-21 | -5.035e+01 | 0.0000 | 2116.0 | 5.34% | 7618.4 | 4.21% | motif file (matrix) | svg |
| 382 | C T A G T C G A C G A T C T A G G C A T C A G T C T A G G A T C C G T A G T C A | CEBP:AP1(bZIP)/ThioMac-CEBPb-ChIP-Seq(GSE21512)/Homer | 1e-21 | -5.017e+01 | 0.0000 | 4512.0 | 11.39% | 17652.5 | 9.75% | motif file (matrix) | svg |
| 383 | T A C G A T G C G A C T A C T G A G C T A G T C G T C A T G C A A C G T A G T C G C T A T G C A | Pknox1(Homeobox)/ES-Prep1-ChIP-Seq(GSE63282)/Homer | 1e-21 | -4.994e+01 | 0.0000 | 1417.0 | 3.58% | 4816.4 | 2.66% | motif file (matrix) | svg |
| 384 | A T C G T G A C A T G C C T G A T C A G G A C T A G T C C G A T T C A G T C G A C A T G C T A G C T A G C G T A C T A G C T A G C T G A C T A G C T A G A T G C | ZSCAN22(Zf)/HEK293-ZSCAN22.GFP-ChIP-Seq(GSE58341)/Homer | 1e-21 | -4.887e+01 | 0.0000 | 297.0 | 0.75% | 673.5 | 0.37% | motif file (matrix) | svg |
| 385 | A T C G A G T C A G T C C G T A A C T G G C A T | hINR(CPE) | 1e-21 | -4.847e+01 | 0.0000 | 6719.0 | 16.97% | 27224.5 | 15.04% | motif file (matrix) | svg |
| 386 | C G A T C T A G T C G A A G C T C G A T C T G A C G T A A G C T A C T G C T A G A T G C G A T C | Hoxb4(Homeobox)/ES-Hoxb4-ChIP-Seq(GSE34014)/Homer | 1e-20 | -4.755e+01 | 0.0000 | 1390.0 | 3.51% | 4748.4 | 2.62% | motif file (matrix) | svg |
| 387 | T A G C C T A G T C G A G A C T A C T G C T G A A G T C T C A G G C A T T G A C C T G A A G C T | Atf7(bZIP)/3T3L1-Atf7-ChIP-Seq(GSE56872)/Homer | 1e-20 | -4.694e+01 | 0.0000 | 3136.0 | 7.92% | 11919.8 | 6.59% | motif file (matrix) | svg |
| 388 | C G A T C A G T C T A G G C T A A G T C C G T A T C A G A G T C A C G T A C T G A C G T G T A C G C T A G C T A G C T A | bZIP52(bZIP)/colamp-bZIP52-DAP-Seq(GSE60143)/Homer | 1e-20 | -4.669e+01 | 0.0000 | 6034.0 | 15.24% | 24314.5 | 13.43% | motif file (matrix) | svg |
| 389 | C T A G A G C T G A C T C A T G A G T C A G T C G T C A C A G T C T A G T C A G G T A C C T G A T C G A G A T C T G A C | Rfx2(HTH)/LoVo-RFX2-ChIP-Seq(GSE49402)/Homer | 1e-20 | -4.643e+01 | 0.0000 | 472.0 | 1.19% | 1280.2 | 0.71% | motif file (matrix) | svg |
| 390 | T C A G T A C G T A G C A C G T A C T G C G A T A G T C C G T A T A C G A G T C | Meis1(Homeobox)/MastCells-Meis1-ChIP-Seq(GSE48085)/Homer | 1e-19 | -4.602e+01 | 0.0000 | 9897.0 | 24.99% | 41283.4 | 22.81% | motif file (matrix) | svg |
| 391 | C T G A A G C T A C G T A C G T A G T C G A C T G A C T C T G A C T G A C T A G C G T A C G T A | STAT6(Stat)/CD4-Stat6-ChIP-Seq(GSE22104)/Homer | 1e-19 | -4.542e+01 | 0.0000 | 2000.0 | 5.05% | 7245.9 | 4.00% | motif file (matrix) | svg |
| 392 | T A G C G T A C C T A G A T C G C T G A C G T A G C T A G C A T A C G T T G A C G T A C A C T G T A C G G T C A C T A G | ASL18(LOBAS2)/colamp-ASL18-DAP-Seq(GSE60143)/Homer | 1e-19 | -4.529e+01 | 0.0000 | 11548.0 | 29.16% | 48652.3 | 26.88% | motif file (matrix) | svg |
| 393 | C T G A G A C T G A T C C T G A A G T C G C A T A C G T G A C T G C T A G C A T | OBP1(C2C2dof)/col-OBP1-DAP-Seq(GSE60143)/Homer | 1e-19 | -4.522e+01 | 0.0000 | 13975.0 | 35.29% | 59516.2 | 32.88% | motif file (matrix) | svg |
| 394 | C G T A A C T G C G T A A C G T C A G T A G T C A G C T G C A T G C T A C G A T | At2g01060(G2like)/colamp-At2g01060-DAP-Seq(GSE60143)/Homer | 1e-19 | -4.490e+01 | 0.0000 | 19063.0 | 48.14% | 82552.3 | 45.61% | motif file (matrix) | svg |
| 395 | A C T G C G T A A C T G A T G C T G A C G A T C A T C G T G C A A C T G A G T C | ZNF519(Zf)/HEK293-ZNF519.GFP-ChIP-Seq(GSE58341)/Homer | 1e-19 | -4.398e+01 | 0.0000 | 1085.0 | 2.74% | 3604.5 | 1.99% | motif file (matrix) | svg |
| 396 | C T G A C G A T C T A G C G T A A G C T C G A T C A G T C T G A G A C T C T A G C T A G A T G C | PBX2(Homeobox)/K562-PBX2-ChIP-Seq(Encode)/Homer | 1e-19 | -4.384e+01 | 0.0000 | 5362.0 | 13.54% | 21507.1 | 11.88% | motif file (matrix) | svg |
| 397 | G A C T C T A G C T A G G T A C A G T C G A T C G A C T G A C T T A G C T C A G | NLP7(RWPRK)/col-NLP7-DAP-Seq(GSE60143)/Homer | 1e-18 | -4.359e+01 | 0.0000 | 11616.0 | 29.34% | 49037.9 | 27.09% | motif file (matrix) | svg |
| 398 | C T G A A T G C C G T A A C G T A G T C A G T C A C G T A C T G A T C G G C A T | SPDEF(ETS)/VCaP-SPDEF-ChIP-Seq(SRA014231)/Homer | 1e-18 | -4.344e+01 | 0.0000 | 5622.0 | 14.20% | 22646.9 | 12.51% | motif file (matrix) | svg |
| 399 | G A C T C A G T A G C T C G A T A G T C G A T C A G T C C G T A A T G C T C A G | Rbpj1(?)/Panc1-Rbpj1-ChIP-Seq(GSE47459)/Homer | 1e-18 | -4.291e+01 | 0.0000 | 6135.0 | 15.49% | 24892.8 | 13.75% | motif file (matrix) | svg |
| 400 | A C T G G A T C G A C T A C T G A C G T C A T G A C T G A C G T A G C T C G A T | RUNX-AML(Runt)/CD4+-PolII-ChIP-Seq(Barski\_et\_al.)/Homer | 1e-18 | -4.274e+01 | 0.0000 | 3720.0 | 9.39% | 14502.3 | 8.01% | motif file (matrix) | svg |
| 401 | T A C G C T G A T C G A C G A T C T A G C T A G T C G A C T G A T C G A T C G A C G T A T C G A G C A T C A T G C G T A T A C G G C A T T G A C C G T A A G C T | NFAT:AP1(RHD,bZIP)/Jurkat-NFATC1-ChIP-Seq(Jolma\_et\_al.)/Homer | 1e-18 | -4.271e+01 | 0.0000 | 650.0 | 1.64% | 1960.5 | 1.08% | motif file (matrix) | svg |
| 402 | A G T C C T G A A T C G A G C T A G C T G A C T A G T C G C T A A C G T C G A T G C A T C G A T A T C G C G T A T A G C G C A T A T G C C G T A | bZIP:IRF(bZIP,IRF)/Th17-BatF-ChIP-Seq(GSE39756)/Homer | 1e-18 | -4.162e+01 | 0.0000 | 1339.0 | 3.38% | 4645.4 | 2.57% | motif file (matrix) | svg |
| 403 | C T A G T A C G G A C T T G C A T G C A C G A T T A C G C T G A T C G A C T G A | Hoxa10(Homeobox)/ChickenMSG-Hoxa10.Flag-ChIP-Seq(GSE86088)/Homer | 1e-17 | -4.144e+01 | 0.0000 | 3037.0 | 7.67% | 11649.1 | 6.44% | motif file (matrix) | svg |
| 404 | C G A T C T A G A C G T G T C A C G T A C G T A A G T C C G T A | Foxo3(Forkhead)/U2OS-Foxo3-ChIP-Seq(E-MTAB-2701)/Homer | 1e-17 | -4.006e+01 | 0.0000 | 3622.0 | 9.15% | 14166.0 | 7.83% | motif file (matrix) | svg |
| 405 | T C G A G C T A T G A C G C T A C T A G G A T C C G A T A C T G C G A T A G C T G A C T C T A G | E-box/Drosophila-Promoters/Homer | 1e-17 | -3.968e+01 | 0.0000 | 982.0 | 2.48% | 3266.0 | 1.80% | motif file (matrix) | svg |
| 406 | C T A G T C G A C T G A C G T A T A C G G A C T T C A G T C G A G T C A T G C A T A C G A G C T | IRF2(IRF)/Erythroblas-IRF2-ChIP-Seq(GSE36985)/Homer | 1e-17 | -3.962e+01 | 0.0000 | 551.0 | 1.39% | 1628.8 | 0.90% | motif file (matrix) | svg |
| 407 | T A C G T A C G G T A C A T C G A C T G T A C G T C G A C T G A T C G A A T C G | E2F6(E2F)/Hela-E2F6-ChIP-Seq(GSE31477)/Homer | 1e-16 | -3.851e+01 | 0.0000 | 4025.0 | 10.16% | 15936.7 | 8.81% | motif file (matrix) | svg |
| 408 | C G T A T G A C T C G A A G T C C G T A A T C G A T G C A C G T A C T G A G T C | E2A(bHLH)/proBcell-E2A-ChIP-Seq(GSE21978)/Homer | 1e-16 | -3.799e+01 | 0.0000 | 5787.0 | 14.61% | 23568.1 | 13.02% | motif file (matrix) | svg |
| 409 | C G A T T A C G T G C A G T A C G A T C G A C T A G C T A C G T A T C G G T A C G A T C G T A C G A T C G T C A | PPARE(NR),DR1/3T3L1-Pparg-ChIP-Seq(GSE13511)/Homer | 1e-15 | -3.676e+01 | 0.0000 | 3835.0 | 9.69% | 15181.0 | 8.39% | motif file (matrix) | svg |
| 410 | A C T G A G C T A G T C G T C A A G C T T C A G A T G C G A T C G C A T A T C G T C G A T A G C C G A T C A T G T A G C | Pax8(Paired,Homeobox)/Thyroid-Pax8-ChIP-Seq(GSE26938)/Homer | 1e-15 | -3.662e+01 | 0.0000 | 1547.0 | 3.91% | 5578.2 | 3.08% | motif file (matrix) | svg |
| 411 | C G T A C G T A C T G A A C T G A C G T A G T C C G T A C G T A A G T C A C T G A T G C G A T C | WRKY46(WRKY)/colamp-WRKY46-DAP-Seq(GSE60143)/Homer | 1e-15 | -3.647e+01 | 0.0000 | 828.0 | 2.09% | 2714.9 | 1.50% | motif file (matrix) | svg |
| 412 | C G A T C T G A G T A C A C G T A C G T T C A G C G T A C G T A G C T A G C A T G C A T A G T C C G T A G T C A C A T G | NST1(NAC)/colamp-NST1-DAP-Seq(GSE60143)/Homer | 1e-15 | -3.638e+01 | 0.0000 | 5266.0 | 13.30% | 21367.0 | 11.81% | motif file (matrix) | svg |
| 413 | T C G A T C G A T A G C G T A C T C A G T A C G C G T A C G T A T C A G A G C T | GABPA(ETS)/Jurkat-GABPa-ChIP-Seq(GSE17954)/Homer | 1e-15 | -3.621e+01 | 0.0000 | 5797.0 | 14.64% | 23683.1 | 13.09% | motif file (matrix) | svg |
| 414 | C A G T A G C T G C A T T C G A A G T C A C G T A C G T A C G T C G A T G C A T | AT5G66940(C2C2dof)/col-AT5G66940-DAP-Seq(GSE60143)/Homer | 1e-15 | -3.467e+01 | 0.0000 | 9630.0 | 24.32% | 40642.9 | 22.46% | motif file (matrix) | svg |
| 415 | T C G A T A G C G T C A A C T G C T A G C G T A C G A T A C T G A C G T A C T G A C T G A C G T | ETS:RUNX(ETS,Runt)/Jurkat-RUNX1-ChIP-Seq(GSE17954)/Homer | 1e-14 | -3.416e+01 | 0.0000 | 498.0 | 1.26% | 1491.9 | 0.82% | motif file (matrix) | svg |
| 416 | T C G A A C T G A C T G C G T A C G T A T C G A A G T C C T G A A T C G G T A C G C A T C A T G | ETS:E-box(ETS,bHLH)/HPC7-Scl-ChIP-Seq(GSE22178)/Homer | 1e-14 | -3.375e+01 | 0.0000 | 343.0 | 0.87% | 935.8 | 0.52% | motif file (matrix) | svg |
| 417 | A G T C C T G A A G T C C G A T C A G T G A T C A T G C A C T G A T C G G A C T | Fli1(ETS)/CD8-FLI-ChIP-Seq(GSE20898)/Homer | 1e-14 | -3.357e+01 | 0.0000 | 9046.0 | 22.85% | 38105.4 | 21.05% | motif file (matrix) | svg |
| 418 | T G C A C T G A A T G C G T C A A C T G A C T G C G T A C G T A C T A G A G C T | Ets1-distal(ETS)/CD4+-PolII-ChIP-Seq(Barski\_et\_al.)/Homer | 1e-14 | -3.327e+01 | 0.0000 | 1130.0 | 2.85% | 3957.5 | 2.19% | motif file (matrix) | svg |
| 419 | T C G A G C A T A C T G C T G A A G T C T C A G G A C T G T A C C G T A A G C T A G T C G A T C | c-Jun-CRE(bZIP)/K562-cJun-ChIP-Seq(GSE31477)/Homer | 1e-14 | -3.319e+01 | 0.0000 | 1393.0 | 3.52% | 5021.2 | 2.77% | motif file (matrix) | svg |
| 420 | T A C G C G T A T C A G G A C T C T A G A C T G C A G T T A G C T C G A A C G T G T A C C T A G A G T C A G T C G A T C | ZNF669(Zf)/HEK293-ZNF669.GFP-ChIP-Seq(GSE58341)/Homer | 1e-14 | -3.307e+01 | 0.0000 | 992.0 | 2.51% | 3410.2 | 1.88% | motif file (matrix) | svg |
| 421 | C G A T C A G T C A G T C A T G G T C A G A T C C G T A T C A G A G T C A C G T C T A G A C G T G T A C G T C A G C T A | VIP1(bZIP)/col-VIP1-DAP-Seq(GSE60143)/Homer | 1e-13 | -3.200e+01 | 0.0000 | 747.0 | 1.89% | 2464.2 | 1.36% | motif file (matrix) | svg |
| 422 | A T G C C T G A A T C G T A C G A G T C C G A T T C A G C G A T C T A G A G C T G T C A G T C A C G T A A G T C C G T A T A C G C T G A | Fox:Ebox(Forkhead,bHLH)/Panc1-Foxa2-ChIP-Seq(GSE47459)/Homer | 1e-13 | -3.161e+01 | 0.0000 | 3716.0 | 9.38% | 14845.3 | 8.20% | motif file (matrix) | svg |
| 423 | A G T C G A C T C A G T A C T G C T A G T G A C G C T A A T G C G C A T A T C G C G A T A C T G G A T C G T A C G T C A C T G A | NF1(CTF)/LNCAP-NF1-ChIP-Seq(Unpublished)/Homer | 1e-13 | -3.137e+01 | 0.0000 | 1529.0 | 3.86% | 5616.0 | 3.10% | motif file (matrix) | svg |
| 424 | G A C T C T A G G A T C C A G T A C T G C T G A A T G C G C A T A T G C C T G A | MafA(bZIP)/Islet-MafA-ChIP-Seq(GSE30298)/Homer | 1e-13 | -3.089e+01 | 0.0000 | 4203.0 | 10.61% | 16971.9 | 9.38% | motif file (matrix) | svg |
| 425 | G C A T T C A G C T G A A T C G A C T G C G A T G A T C C T G A | THRb(NR)/Liver-NR1A2-ChIP-Seq(GSE52613)/Homer | 1e-13 | -3.079e+01 | 0.0000 | 18678.0 | 47.17% | 81636.7 | 45.11% | motif file (matrix) | svg |
| 426 | T C G A G C A T A C G T C T A G G T A C T C G A G C A T T G A C T C G A A C G T | Chop(bZIP)/MEF-Chop-ChIP-Seq(GSE35681)/Homer | 1e-13 | -3.073e+01 | 0.0000 | 1461.0 | 3.69% | 5351.7 | 2.96% | motif file (matrix) | svg |
| 427 | C T G A C T G A C T A G T C G A C G T A A T G C C G T A A C T G C G T A A C G T C T G A C G A T A G C T C G T A A C G T A G T C C G A T T A C G G T C A G C A T | GATA(Zf),IR3/iTreg-Gata3-ChIP-Seq(GSE20898)/Homer | 1e-13 | -3.058e+01 | 0.0000 | 835.0 | 2.11% | 2830.8 | 1.56% | motif file (matrix) | svg |
| 428 | T A C G T A C G C T A G T C A G A G T C C G T A A T C G A T G C A C G T A C T G A G T C G A C T | Ascl2(bHLH)/ESC-Ascl2-ChIP-Seq(GSE97712)/Homer | 1e-13 | -3.011e+01 | 0.0000 | 4620.0 | 11.67% | 18809.7 | 10.39% | motif file (matrix) | svg |
| 429 | A G T C G A C T C A G T G T A C A G T C A T C G T C A G A C T G G T C A C G T A | Stat3(Stat)/mES-Stat3-ChIP-Seq(GSE11431)/Homer | 1e-13 | -3.009e+01 | 0.0000 | 2390.0 | 6.04% | 9240.0 | 5.11% | motif file (matrix) | svg |
| 430 | G C A T G A T C T C A G G C T A G A C T A G T C C T A G C G T A C A T G G T C A | GATA20(C2C2gata)/colamp-GATA20-DAP-Seq(GSE60143)/Homer | 1e-12 | -2.983e+01 | 0.0000 | 19876.0 | 50.20% | 87164.8 | 48.16% | motif file (matrix) | svg |
| 431 | T C G A C G T A C G T A T C G A A C T G G T A C C G T A A G C T G T C A G C A T | At3g24120(G2like)/col-At3g24120-DAP-Seq(GSE60143)/Homer | 1e-12 | -2.865e+01 | 0.0000 | 19433.0 | 49.08% | 85222.8 | 47.09% | motif file (matrix) | svg |
| 432 | T G A C T C G A C T G A C T G A A T G C G A T C C T A G T A C G G A C T G A C T G A T C T C G A C T G A C T G A A T G C G A T C C T A G A T C G G A C T G A C T | Tcfcp2l1(CP2)/mES-Tcfcp2l1-ChIP-Seq(GSE11431)/Homer | 1e-11 | -2.758e+01 | 0.0000 | 965.0 | 2.44% | 3399.9 | 1.88% | motif file (matrix) | svg |
| 433 | C A T G G T C A A G T C C G T A C T A G G A T C C G A T A C T G A C G T G T A C C G T A C G T A | bZIP69(bZIP)/col-bZIP69-DAP-Seq(GSE60143)/Homer | 1e-11 | -2.696e+01 | 0.0000 | 471.0 | 1.19% | 1473.8 | 0.81% | motif file (matrix) | svg |
| 434 | C G A T C T G A G T A C C A T G G C A T T C A G G C A T C G T A C G T A G C T A C G T A A G T C G C T A G T A C C A T G | CUC2(NAC)/colamp-CUC2-DAP-Seq(GSE60143)/Homer | 1e-11 | -2.688e+01 | 0.0000 | 2819.0 | 7.12% | 11163.4 | 6.17% | motif file (matrix) | svg |
| 435 | A C G T C T A G C G T A A G T C G T A C A C G T A C G T A C G T G T C A G T A C T G A C G A C T | Nur77(NR)/K562-NR4A1-ChIP-Seq(GSE31363)/Homer | 1e-11 | -2.655e+01 | 0.0000 | 871.0 | 2.20% | 3042.0 | 1.68% | motif file (matrix) | svg |
| 436 | C T G A T C A G C T G A C T A G C A T G A C G T A T G C C G T A A T G C G C A T T C A G C T G A A C T G A C G T C A G T A G T C C G T A C A G T C T A G C A T G | VDR(NR),DR3/GM10855-VDR+vitD-ChIP-Seq(GSE22484)/Homer | 1e-11 | -2.552e+01 | 0.0000 | 1070.0 | 2.70% | 3864.9 | 2.14% | motif file (matrix) | svg |
| 437 | G A C T A G T C C G A T A C T G C T G A T G A C G T A C C G T A A T C G G C A T C T G A C T A G | Bcl11a(Zf)/HSPC-BCL11A-ChIP-Seq(GSE104676)/Homer | 1e-10 | -2.508e+01 | 0.0000 | 2950.0 | 7.45% | 11787.7 | 6.51% | motif file (matrix) | svg |
| 438 | T G C A G C A T A G C T G C A T A G T C A G T C A G T C C T G A A C T G T C G A T C G A C A G T A T C G A G T C G A T C | ZNF143\|STAF(Zf)/CUTLL-ZNF143-ChIP-Seq(GSE29600)/Homer | 1e-10 | -2.503e+01 | 0.0000 | 1120.0 | 2.83% | 4078.8 | 2.25% | motif file (matrix) | svg |
| 439 | A G T C G A T C G A T C C G T A G T C A A G T C A G C T C T G A G A C T G A C T | ATY13(MYB)/col-ATY13-DAP-Seq(GSE60143)/Homer | 1e-10 | -2.442e+01 | 0.0000 | 21786.0 | 55.02% | 96286.2 | 53.20% | motif file (matrix) | svg |
| 440 | G A C T G A C T A T C G C G A T G T C A G T A C A G C T C G A T A C G T G T A C | SPL11(SBP)/col100-SPL11-DAP-Seq(GSE60143)/Homer | 1e-10 | -2.427e+01 | 0.0000 | 4456.0 | 11.25% | 18343.6 | 10.14% | motif file (matrix) | svg |
| 441 | C T A G T G A C G A C T A T C G T C G A A G T C C T A G C A G T C T A G A T C G G T A C T C G A | O2(bZIP)/Corn-O2-ChIP-Seq(GSE63991)/Homer | 1e-10 | -2.414e+01 | 0.0000 | 1236.0 | 3.12% | 4574.5 | 2.53% | motif file (matrix) | svg |
| 442 | C G A T C T G A G T A C A C T G A C G T T C A G G C A T C G T A C G T A G C A T C G T A A G T C C G T A G T A C C A T G | CUC3(NAC)/col-CUC3-DAP-Seq(GSE60143)/Homer | 1e-10 | -2.408e+01 | 0.0000 | 2804.0 | 7.08% | 11196.1 | 6.19% | motif file (matrix) | svg |
| 443 | G C A T C G A T A T G C A G C T T C G A T A C G G C T A C G T A C A T G T G A C G C A T C G A T A G T C A G C T C G T A | AT3G09735(S1Falike)/col-AT3G09735-DAP-Seq(GSE60143)/Homer | 1e-10 | -2.405e+01 | 0.0000 | 1670.0 | 4.22% | 6381.4 | 3.53% | motif file (matrix) | svg |
| 444 | G T A C G A T C C A G T A G T C A G T C A G T C T G C A G A T C C T G A A T G C G T C A A C G T | WT1(Zf)/Kidney-WT1-ChIP-Seq(GSE90016)/Homer | 1e-10 | -2.399e+01 | 0.0000 | 3158.0 | 7.98% | 12722.6 | 7.03% | motif file (matrix) | svg |
| 445 | A T G C G A C T A C T G C A G T G A T C A C G T T A C G T A C G | Smad2(MAD)/ES-SMAD2-ChIP-Seq(GSE29422)/Homer | 1e-10 | -2.384e+01 | 0.0000 | 10492.0 | 26.50% | 45114.9 | 24.93% | motif file (matrix) | svg |
| 446 | C G T A A T G C C G A T A C G T A G T C C G T A C G T A C G T A C T A G A T C G | TCFL2(HMG)/K562-TCF7L2-ChIP-Seq(GSE29196)/Homer | 1e-9 | -2.277e+01 | 0.0000 | 448.0 | 1.13% | 1439.2 | 0.80% | motif file (matrix) | svg |
| 447 | T G C A C G T A G T C A A G C T A G T C G C T A T A G C C G A T C T A G G A T C | Gfi1b(Zf)/HPC7-Gfi1b-ChIP-Seq(GSE22178)/Homer | 1e-9 | -2.239e+01 | 0.0000 | 2854.0 | 7.21% | 11473.3 | 6.34% | motif file (matrix) | svg |
| 448 | C T G A T A C G G C A T C T A G A T G C G A T C C G A T A C T G C T A G G A T C C T G A A T G C | MYRF(MYRF)/CFPAC1-MYRF-ChIP-Seq(GSE145627)/Homer | 1e-9 | -2.222e+01 | 0.0000 | 1716.0 | 4.33% | 6626.7 | 3.66% | motif file (matrix) | svg |
| 449 | A T G C C T G A G A C T A C G T A C G T G T A C G A T C C G A T C T A G C A T G C G T A C G T A C T G A G A C T | STAT1(Stat)/HelaS3-STAT1-ChIP-Seq(GSE12782)/Homer | 1e-9 | -2.222e+01 | 0.0000 | 965.0 | 2.44% | 3505.7 | 1.94% | motif file (matrix) | svg |
| 450 | T G C A C T G A C A T G C T A G C A G T A G T C C G T A A T G C A T G C T A C G G C A T T C A G G T C A G A T C G T A C | ERE(NR),IR3/MCF7-ERa-ChIP-Seq(Unpublished)/Homer | 1e-9 | -2.151e+01 | 0.0000 | 1188.0 | 3.00% | 4438.9 | 2.45% | motif file (matrix) | svg |
| 451 | A C G T A G T C A G C T A G T C C G T A G T A C A G T C C G A T C G T A G T C A | MYB41(MYB)/col-MYB41-DAP-Seq(GSE60143)/Homer | 1e-9 | -2.147e+01 | 0.0000 | 2802.0 | 7.08% | 11284.4 | 6.23% | motif file (matrix) | svg |
| 452 | G C A T G C A T G A T C G C A T T C G A A C T G C G T A C G T A A T C G T A G C C G A T A C G T A G T C A G C T C G T A | HSF6(HSF)/col-HSF6-DAP-Seq(GSE60143)/Homer | 1e-9 | -2.111e+01 | 0.0000 | 884.0 | 2.23% | 3198.4 | 1.77% | motif file (matrix) | svg |
| 453 | G C T A A C T G T C G A C T G A C T G A A C G T T A G C C T G A C G T A C G A T | Cux2(Homeobox)/Liver-Cux2-ChIP-Seq(GSE35985)/Homer | 1e-8 | -2.044e+01 | 0.0000 | 5011.0 | 12.65% | 20960.1 | 11.58% | motif file (matrix) | svg |
| 454 | A C T G G A C T A G T C C T G A G A T C T C A G A T G C G A C T A G T C A T G C T A G C A G C T A T C G T G C A | PAX5(Paired,Homeobox),condensed/GM12878-PAX5-ChIP-Seq(GSE32465)/Homer | 1e-8 | -2.018e+01 | 0.0000 | 821.0 | 2.07% | 2961.5 | 1.64% | motif file (matrix) | svg |
| 455 | T G C A A G C T A C G T C T A G G A T C C T A G G A T C G T C A C T G A A G T C | CEBP(bZIP)/ThioMac-CEBPb-ChIP-Seq(GSE21512)/Homer | 1e-8 | -1.987e+01 | 0.0000 | 4816.0 | 12.16% | 20132.6 | 11.12% | motif file (matrix) | svg |
| 456 | T C G A A C G T A C T G C G T A A G T C C T A G A G C T T G A C | TGA10(bZIP)/colamp-TGA10-DAP-Seq(GSE60143)/Homer | 1e-8 | -1.955e+01 | 0.0000 | 5592.0 | 14.12% | 23569.4 | 13.02% | motif file (matrix) | svg |
| 457 | G T A C A C T G A C G T T C A G G C A T C G T A C G A T G C A T C G T A A G T C C G T A T G A C C A T G G A C T G C T A | ANAC083(NAC)/col-ANAC083-DAP-Seq(GSE60143)/Homer | 1e-8 | -1.891e+01 | 0.0000 | 5559.0 | 14.04% | 23459.7 | 12.96% | motif file (matrix) | svg |
| 458 | T C A G A C T G C A G T A G T C A G T C G T C A C G T A C G T A A C T G C A G T A G T C A G T C C T G A T G C A A G C T | dHNF4(NR)/Fly-HNF4-ChIP-Seq(GSE73675)/Homer | 1e-8 | -1.854e+01 | 0.0000 | 251.0 | 0.63% | 748.1 | 0.41% | motif file (matrix) | svg |
| 459 | C G A T C T A G G A T C G C T A A G C T C T A G G A T C C G T A | RBFox2(?)/Heart-RBFox2-CLIP-Seq(GSE57926)/Homer | 1e-8 | -1.851e+01 | 0.0000 | 12670.0 | 32.00% | 55293.0 | 30.55% | motif file (matrix) | svg |
| 460 | C T G A A C G T A C G T A C G T A G T C G A C T C G A T C T G A A C T G C G T A C G T A T C G A | STAT5(Stat)/mCD4+-Stat5-ChIP-Seq(GSE12346)/Homer | 1e-7 | -1.836e+01 | 0.0000 | 1028.0 | 2.60% | 3852.8 | 2.13% | motif file (matrix) | svg |
| 461 | G A C T C T A G C T A G C T A G A C T G T C G A C T G A C T A G C T A G C T A G G T A C G T C A | ZNF467(Zf)/HEK293-ZNF467.GFP-ChIP-Seq(GSE58341)/Homer | 1e-7 | -1.828e+01 | 0.0000 | 3817.0 | 9.64% | 15831.5 | 8.75% | motif file (matrix) | svg |
| 462 | A G T C T A G C A C T G A C G T A C G T C G T A C G T A C A G T C G A T A G T C C T A G A C T G A C G T A C G T C T G A | MYB44(MYB)/colamp-MYB44-DAP-Seq(GSE60143)/Homer | 1e-7 | -1.800e+01 | 0.0000 | 629.0 | 1.59% | 2227.7 | 1.23% | motif file (matrix) | svg |
| 463 | T G C A C G T A A C T G T C A G C A G T C A T G T C A G G A T C T A C G A G T C T G C A A C T G A C T G T G A C G T C A | ZNF165(Zf)/WHIM12-ZNF165-ChIP-Seq(GSE65937)/Homer | 1e-7 | -1.770e+01 | 0.0000 | 535.0 | 1.35% | 1856.4 | 1.03% | motif file (matrix) | svg |
| 464 | C T A G C T G A A G T C G C T A C G A T A C T G G A C T G A T C G A T C C T G A C T A G C T G A T G A C G C T A C G A T T C A G G A C T G A T C G A T C T G A C | p53(p53)/Saos-p53-ChIP-Seq(GSE15780)/Homer | 1e-7 | -1.748e+01 | 0.0000 | 531.0 | 1.34% | 1844.1 | 1.02% | motif file (matrix) | svg |
| 465 | C T A G C T G A A G T C G C T A C G A T A C T G G A C T G A T C G A T C C T G A C T A G C T G A T G A C G C T A C G A T T C A G G A C T G A T C G A T C T G A C | p53(p53)/Saos-p53-ChIP-Seq/Homer | 1e-7 | -1.748e+01 | 0.0000 | 531.0 | 1.34% | 1844.1 | 1.02% | motif file (matrix) | svg |
| 466 | A T G C A G C T T C A G T G A C T C A G A T G C T G C A A C G T A T C G G A T C A C T G A G T C | NRF1(NRF)/MCF7-NRF1-ChIP-Seq(Unpublished)/Homer | 1e-7 | -1.736e+01 | 0.0000 | 573.0 | 1.45% | 2014.4 | 1.11% | motif file (matrix) | svg |
| 467 | C T A G T A C G G A T C G T C A T G C A A C G T T G C A G C T A T C G A T G C A | Hoxa9(Homeobox)/ChickenMSG-Hoxa9.Flag-ChIP-Seq(GSE86088)/Homer | 1e-7 | -1.714e+01 | 0.0000 | 14382.0 | 36.32% | 63147.7 | 34.89% | motif file (matrix) | svg |
| 468 | T G C A G C A T C G A T C G T A C A G T A C T G G T A C C G T A C T G A A G C T G T C A A C T G C T A G G T C A C G A T A C T G G T A C T G C A C G T A A G C T | CEBP:CEBP(bZIP)/MEF-Chop-ChIP-Seq(GSE35681)/Homer | 1e-7 | -1.691e+01 | 0.0000 | 629.0 | 1.59% | 2248.8 | 1.24% | motif file (matrix) | svg |
| 469 | C G T A T C G A G A T C G C A T C G T A A C G T G T A C T C A G G T C A G A C T C G T A C T A G | DREF/Drosophila-Promoters/Homer | 1e-7 | -1.679e+01 | 0.0000 | 457.0 | 1.15% | 1562.2 | 0.86% | motif file (matrix) | svg |
| 470 | C G A T G A C T C G A T T C A G G A C T A C G T C A G T C T G A G A C T G A C T A G C T C G A T A C T G A T C G G T A C G C T A | NF1:FOXA1(CTF,Forkhead)/LNCAP-FOXA1-ChIP-Seq(GSE27824)/Homer | 1e-7 | -1.639e+01 | 0.0000 | 232.0 | 0.59% | 701.6 | 0.39% | motif file (matrix) | svg |
| 471 | T C A G C T G A C T A G C A T G A C G T A T G C C T G A C T G A C T G A C T A G C A T G A C G T A T G C C T G A | TR4(NR),DR1/Hela-TR4-ChIP-Seq(GSE24685)/Homer | 1e-7 | -1.627e+01 | 0.0000 | 376.0 | 0.95% | 1252.5 | 0.69% | motif file (matrix) | svg |
| 472 | T G A C G T A C C G T A A C T G T G A C C G A T A C T G A T C G A G C T T A C G T C G A T A G C G T A C C G T A A T C G T G A C G C A T A C T G A C T G A T G C | Twist(bHLH)/HMLE-TWIST1-ChIP-Seq(Chang\_et\_al)/Homer | 1e-7 | -1.614e+01 | 0.0000 | 375.0 | 0.95% | 1250.8 | 0.69% | motif file (matrix) | svg |
| 473 | T C G A C A T G C A T G A C G T A T G C T C G A C T G A A G C T T A C G T G C A G T A C G A T C A G C T A G T C | FXR(NR),IR1/Liver-FXR-ChIP-Seq(Chong\_et\_al.)/Homer | 1e-6 | -1.603e+01 | 0.0000 | 2088.0 | 5.27% | 8419.0 | 4.65% | motif file (matrix) | svg |
| 474 | C T A G A T C G G T C A C A T G A G T C G A C T T A C G C A G T A G T C A G T C C T G A C G A T C T A G A T C G G A C T A T C G A G T C G A C T C T A G T C G A | REST-NRSF(Zf)/Jurkat-NRSF-ChIP-Seq/Homer | 1e-6 | -1.568e+01 | 0.0000 | 23.0 | 0.06% | 20.9 | 0.01% | motif file (matrix) | svg |
| 475 | C G T A A C T G G T C A A C G T A T C G C A G T C T A G T C A G C G T A A C T G C G T A A C G T C G T A C T G A T A C G | GATA3(Zf),DR4/iTreg-Gata3-ChIP-Seq(GSE20898)/Homer | 1e-6 | -1.566e+01 | 0.0000 | 450.0 | 1.14% | 1553.1 | 0.86% | motif file (matrix) | svg |
| 476 | T G A C G C T A T G A C C G T A T C A G G A T C C G T A C A T G C A T G C T A G C T A G C T A G | Unknown-ESC-element(?)/mES-Nanog-ChIP-Seq(GSE11724)/Homer | 1e-6 | -1.552e+01 | 0.0000 | 1755.0 | 4.43% | 7008.0 | 3.87% | motif file (matrix) | svg |
| 477 | A T G C G A T C C G A T C T A G A C T G G C T A C G T A A G C T A C T G A G C T | TEAD2(TEA)/Py2T-Tead2-ChIP-Seq(GSE55709)/Homer | 1e-6 | -1.538e+01 | 0.0000 | 2336.0 | 5.90% | 9517.0 | 5.26% | motif file (matrix) | svg |
| 478 | C G A T C G A T G T A C G A T C G A T C C G T A G C T A C G A T C G A T C T G A C T A G C A T G G C T A G C T A C G T A | AGL16(MADS)/col-AGL16-DAP-Seq(GSE60143)/Homer | 1e-6 | -1.503e+01 | 0.0000 | 363.0 | 0.92% | 1220.1 | 0.67% | motif file (matrix) | svg |
| 479 | A G C T G C A T G T C A C G A T T A G C C G T A A C G T G C T A | CRC(C2C2YABBY)/col-CRC-DAP-Seq(GSE60143)/Homer | 1e-6 | -1.499e+01 | 0.0000 | 7844.0 | 19.81% | 33881.9 | 18.72% | motif file (matrix) | svg |
| 480 | T G C A A G C T C A T G C G T A A G C T A C T G G A T C G T C A C G T A A G C T | Atf4(bZIP)/MEF-Atf4-ChIP-Seq(GSE35681)/Homer | 1e-6 | -1.422e+01 | 0.0000 | 1883.0 | 4.76% | 7608.0 | 4.20% | motif file (matrix) | svg |
| 481 | C A T G G A T C C T G A A G T C C T A G C G T A G C T A G C A T G A T C G A T C A G T C C T A G C G T A C A T G C T A G | PLT1(AP2EREBP)/colamp-PLT1-DAP-Seq(GSE60143)/Homer | 1e-6 | -1.396e+01 | 0.0000 | 870.0 | 2.20% | 3307.1 | 1.83% | motif file (matrix) | svg |
| 482 | C T G A T A G C T G A C T C A G C T A G G T C A C G T A T C A G A G C T T C A G | ETV4(ETS)/HepG2-ETV4-ChIP-Seq(ENCODE)/Homer | 1e-6 | -1.393e+01 | 0.0000 | 8184.0 | 20.67% | 35485.4 | 19.61% | motif file (matrix) | svg |
| 483 | G C A T A C G T C G A T A G T C A G T C G C A T C G T A C G T A C G A T C G A T C G A T C T A G A C T G G C T A G C T A | AGL15(MADS)/col-AGL15-DAP-Seq(GSE60143)/Homer | 1e-5 | -1.335e+01 | 0.0000 | 493.0 | 1.25% | 1768.1 | 0.98% | motif file (matrix) | svg |
| 484 | C G T A C G A T C G T A T C G A T C G A A C G T C G T A A C G T A G T C G C A T | LHY(Myb)/Seedling-LHY-ChIP-Seq(GSE52175)/Homer | 1e-5 | -1.333e+01 | 0.0000 | 5723.0 | 14.45% | 24539.7 | 13.56% | motif file (matrix) | svg |
| 485 | C T G A A T C G A G C T A G C T A C G T T A G C C T G A T A C G C G A T A C G T G A C T A G T C | ISRE(IRF)/ThioMac-LPS-Expression(GSE23622)/Homer | 1e-5 | -1.320e+01 | 0.0000 | 203.0 | 0.51% | 629.2 | 0.35% | motif file (matrix) | svg |
| 486 | T C A G T C A G A C G T G T A C G C T A T C A G C T G A A C T G A C T G A G C T A G T C C G T A | EAR2(NR)/K562-NR2F6-ChIP-Seq(Encode)/Homer | 1e-5 | -1.311e+01 | 0.0000 | 6817.0 | 17.22% | 29433.3 | 16.26% | motif file (matrix) | svg |
| 487 | G T A C C T G A A G T C A G T C A C T G G T C A G A T C G C A T | At1g75490(AP2EREBP)/colamp-At1g75490-DAP-Seq(GSE60143)/Homer | 1e-5 | -1.294e+01 | 0.0000 | 21133.0 | 53.37% | 94298.3 | 52.10% | motif file (matrix) | svg |
| 488 | C T A G C A G T C G T A A C G T A G T C A C T G C G T A A G C T A G T C G A T C | HNF6(Homeobox)/Liver-Hnf6-ChIP-Seq(ERP000394)/Homer | 1e-5 | -1.206e+01 | 0.0000 | 6657.0 | 16.81% | 28800.6 | 15.91% | motif file (matrix) | svg |
| 489 | C G T A G A C T C G A T A T C G G T A C G C A T C A T G C G T A T A C G G C A T G T A C C G T A C A T G A T G C G C T A C T A G G C A T G C A T G C A T G A C T | MafB(bZIP)/BMM-Mafb-ChIP-Seq(GSE75722)/Homer | 1e-5 | -1.204e+01 | 0.0000 | 1278.0 | 3.23% | 5092.1 | 2.81% | motif file (matrix) | svg |
| 490 | T C G A A G T C C G T A A T C G T A G C A C G T A C T G A G C T A C G T A G T C | Ptf1a(bHLH)/Panc1-Ptf1a-ChIP-Seq(GSE47459)/Homer | 1e-5 | -1.193e+01 | 0.0000 | 11199.0 | 28.28% | 49229.5 | 27.20% | motif file (matrix) | svg |
| 491 | G A T C G C A T G C A T A G T C A G C T T C G A T A C G G C T A C G T A C T A G T G A C G C A T C G A T G A T C A G C T | HSFC1(HSF)/col-HSFC1-DAP-Seq(GSE60143)/Homer | 1e-5 | -1.184e+01 | 0.0000 | 637.0 | 1.61% | 2392.5 | 1.32% | motif file (matrix) | svg |
| 492 | A G C T A G T C A G T C A C G T C T A G A C G T A C G T A C G T C G T A A G T C G A T C C G T A | FOXP1(Forkhead)/H9-FOXP1-ChIP-Seq(GSE31006)/Homer | 1e-4 | -1.149e+01 | 0.0000 | 1892.0 | 4.78% | 7761.9 | 4.29% | motif file (matrix) | svg |
| 493 | A T G C C G T A A C T G C G T A A C G T G C T A T C G A A G C T C G A T C G T A A C G T A G T C C G A T A C T G G A T C | GATA(Zf),IR4/iTreg-Gata3-ChIP-Seq(GSE20898)/Homer | 1e-4 | -1.148e+01 | 0.0000 | 461.0 | 1.16% | 1676.8 | 0.93% | motif file (matrix) | svg |
| 494 | A G T C A G T C A G T C A G T C A G T C A G T C A G T C A G T C A G T C A G T C | SeqBias: polyC-repeat | 1e-4 | -1.148e+01 | 0.0000 | 39580.0 | 99.96% | 180784.0 | 99.89% | motif file (matrix) | svg |
| 495 | T C G A A C G T A C T G C T G A A G T C T C A G A G C T G T A C C G T A A G C T G A T C T C G A | JunD(bZIP)/K562-JunD-ChIP-Seq/Homer | 1e-4 | -1.145e+01 | 0.0000 | 372.0 | 0.94% | 1316.1 | 0.73% | motif file (matrix) | svg |
| 496 | C T G A C T A G A T C G G C A T A C T G G T A C A T G C C G T A A C T G G C T A A G T C C G T A | Tbox:Smad(T-box,MAD)/ESCd5-Smad2\_3-ChIP-Seq(GSE29422)/Homer | 1e-4 | -1.123e+01 | 0.0000 | 868.0 | 2.19% | 3376.5 | 1.87% | motif file (matrix) | svg |
| 497 | T A G C G C T A T C G A C T G A A G T C A G T C C T G A A G T C C G T A C T A G | RUNX(Runt)/HPC7-Runx1-ChIP-Seq(GSE22178)/Homer | 1e-4 | -1.113e+01 | 0.0000 | 4458.0 | 11.26% | 19071.1 | 10.54% | motif file (matrix) | svg |
| 498 | C T A G C T A G T C G A C G T A A T G C C G T A A T C G T C G A T A C G G C A T A C T G C A G T T A G C G A T C G A C T | MRE(NR)/Neuro2A-NR3C2-ChIPnexus(GSE115417)/Homer | 1e-4 | -1.104e+01 | 0.0000 | 6233.0 | 15.74% | 26987.1 | 14.91% | motif file (matrix) | svg |
| 499 | T C G A G T A C T C G A T C G A C A T G A T G C A C G T A C T G A C T G A G T C C G T A C T A G A G T C A T C G A G T C | Unknown3/Drosophila-Promoters/Homer | 1e-4 | -1.099e+01 | 0.0000 | 505.0 | 1.28% | 1867.7 | 1.03% | motif file (matrix) | svg |
| 500 | A C T G G A T C C T G A A T C G A G T C T A G C C T G A C G T A T A C G A G T C C T A G C A G T T C A G T C G A T G A C G A T C | PAX5(Paired,Homeobox)/GM12878-PAX5-ChIP-Seq(GSE32465)/Homer | 1e-4 | -1.098e+01 | 0.0000 | 2664.0 | 6.73% | 11157.9 | 6.16% | motif file (matrix) | svg |
| 501 | C T A G A T G C A T G C C G A T A C T G G A C T A T G C G C T A T G A C A G C T T A G C G C T A | PBX1(Homeobox)/MCF7-PBX1-ChIP-Seq(GSE28007)/Homer | 1e-4 | -1.094e+01 | 0.0000 | 278.0 | 0.70% | 950.2 | 0.53% | motif file (matrix) | svg |
| 502 | G A C T A G T C C G T A C G T A A G T C A G C T A C T G G A C T G T A C A T G C | MYB77(MYB)/col-MYB77-DAP-Seq(GSE60143)/Homer | 1e-4 | -1.085e+01 | 0.0000 | 13760.0 | 34.75% | 60933.7 | 33.67% | motif file (matrix) | svg |
| 503 | C G A T T C G A G T A C A C T G A C G T T C A G G C A T T G C A G C T A G C A T C G T A A G T C C G T A G T A C C A T G | ANAC087(NAC)/col-ANAC087-DAP-Seq(GSE60143)/Homer | 1e-4 | -1.050e+01 | 0.0001 | 2686.0 | 6.78% | 11280.6 | 6.23% | motif file (matrix) | svg |
| 504 | T G C A A T G C A C G T A C G T A C G T A T G C C T A G A C G T A C G T A G C T G A T C A G C T | T1ISRE(IRF)/ThioMac-Ifnb-Expression/Homer | 1e-4 | -1.027e+01 | 0.0001 | 70.0 | 0.18% | 176.0 | 0.10% | motif file (matrix) | svg |
| 505 | C G T A C T G A C T A G C T G A A G T C G C T A C G A T A T C G G A C T G A T C A G T C C T G A C T A G C T A G A G T C G C T A C G A T C T A G G A T C G A T C | p73(p53)/Trachea-p73-ChIP-Seq(PRJNA310161)/Homer | 1e-4 | -1.004e+01 | 0.0001 | 248.0 | 0.63% | 846.4 | 0.47% | motif file (matrix) | svg |
| 506 | C G T A C G T A C T G A A C T G C G T A C G T A A C G T G T C A A C G T G C A T A G T C G A C T | At2g03500(G2like)/col-At2g03500-DAP-Seq(GSE60143)/Homer | 1e-4 | -9.988e+00 | 0.0001 | 1451.0 | 3.66% | 5916.7 | 3.27% | motif file (matrix) | svg |
| 507 | C G T A C A G T G T A C A T G C C T A G C G T A A C G T A G T C T C G A T C A G | GATA19(C2C2gata)/colamp-GATA19-DAP-Seq(GSE60143)/Homer | 1e-4 | -9.907e+00 | 0.0001 | 1091.0 | 2.76% | 4369.3 | 2.41% | motif file (matrix) | svg |
| 508 | C G T A C G T A G C A T A C T G C G T A A G C T C T G A C G T A T A C G C T G A | ELT-3(Gata)/cElegans-L1-ELT3-ChIP-Seq(modEncode)/Homer | 1e-4 | -9.907e+00 | 0.0001 | 2349.0 | 5.93% | 9834.1 | 5.43% | motif file (matrix) | svg |
| 509 | T A C G A T C G T A G C G A T C A C T G A C G T A G T C A C G T C T A G A T C G | Smad4(MAD)/ESC-SMAD4-ChIP-Seq(GSE29422)/Homer | 1e-4 | -9.709e+00 | 0.0001 | 10452.0 | 26.40% | 46082.2 | 25.46% | motif file (matrix) | svg |
| 510 | T C G A G A C T T C A G T G C A G T A C G T A C A G C T G T A C C A T G T C G A C A T G C A T G A C G T A G T C C T G A | FXR(NR),ER2/Liver-FXR-ChIP-Seq(GSE133700)/Homer | 1e-4 | -9.603e+00 | 0.0001 | 2500.0 | 6.31% | 10514.8 | 5.81% | motif file (matrix) | svg |
| 511 | C T G A C T A G T G C A C T G A T C G A A G C T C A T G T C G A A G T C G A C T A C G T A G T C G A T C G A T C G A C T | ZNF528(Zf)/HEK293-ZNF528.GFP-ChIP-Seq(GSE58341)/Homer | 1e-4 | -9.446e+00 | 0.0002 | 29.0 | 0.07% | 51.6 | 0.03% | motif file (matrix) | svg |
| 512 | T G A C A T G C C G T A A T C G A T G C C A G T C A T G A C T G A G T C G T A C | HEB(bHLH)/mES-Heb-ChIP-Seq(GSE53233)/Homer | 1e-4 | -9.404e+00 | 0.0002 | 7982.0 | 20.16% | 34978.9 | 19.33% | motif file (matrix) | svg |
| 513 | G C T A C T G A C G T A C G T A C T G A C T G A C G A T G T C A A C G T A G T C G C A T G C A T | At5g52660(MYBrelated)/colamp-At5g52660-DAP-Seq(GSE60143)/Homer | 1e-4 | -9.345e+00 | 0.0002 | 1525.0 | 3.85% | 6265.1 | 3.46% | motif file (matrix) | svg |
| 514 | C G T A C T G A C T A G C G T A C G T A A G T C C G T A C A G T G C A T G T C A C G A T A C T G A C G T G C A T G A T C | PGR(NR)/EndoStromal-PGR-ChIP-Seq(GSE69539)/Homer | 1e-3 | -9.135e+00 | 0.0002 | 867.0 | 2.19% | 3439.2 | 1.90% | motif file (matrix) | svg |
| 515 | G C T A G C A T G C T A G C A T G C A T C G T A C G T A A G T C A G T C A C T G G C A T G C A T C G T A G C T A G C T A | MYB73(MYB)/col-MYB73-DAP-Seq(GSE60143)/Homer | 1e-3 | -9.093e+00 | 0.0002 | 13217.0 | 33.38% | 58670.2 | 32.42% | motif file (matrix) | svg |
| 516 | A C T G A G T C G T C A C G T A A G T C C G T A C T A G C T A G G A C T C A T G | SCRT1(Zf)/HEK293-SCRT1.eGFP-ChIP-Seq(Encode)/Homer | 1e-3 | -8.989e+00 | 0.0003 | 1949.0 | 4.92% | 8131.1 | 4.49% | motif file (matrix) | svg |
| 517 | C T G A G T A C G A C T A G T C C A G T T G C A C T G A A C G T A G C T G A T C C T A G C G A T A C T G A T G C G A C T C T G A G A T C G A C T A G C T G A T C | Mouse\_Recombination\_Hotspot(Zf)/Testis-DMC1-ChIP-Seq(GSE24438)/Homer | 1e-3 | -8.467e+00 | 0.0004 | 291.0 | 0.73% | 1046.9 | 0.58% | motif file (matrix) | svg |
| 518 | C T A G T C G A A C G T A C G T C A T G A G T C C T G A C G A T A G T C C G T A | AARE(HLH)/mES-cMyc-ChIP-Seq/Homer | 1e-3 | -8.420e+00 | 0.0004 | 639.0 | 1.61% | 2494.8 | 1.38% | motif file (matrix) | svg |
| 519 | G T A C C G T A C G T A T A C G G C A T G T A C C G T A C A T G A G T C C G T A C G T A C G A T G C A T G C A T G A C T | MafF(bZIP)/HepG2-MafF-ChIP-Seq(GSE31477)/Homer | 1e-3 | -8.367e+00 | 0.0005 | 1024.0 | 2.59% | 4140.5 | 2.29% | motif file (matrix) | svg |
| 520 | G C A T C T A G C T A G C G T A A G C T C G T A C T G A C A T G C T A G G C A T | AT5G56840(MYBrelated)/colamp-AT5G56840-DAP-Seq(GSE60143)/Homer | 1e-3 | -8.277e+00 | 0.0005 | 8029.0 | 20.28% | 35305.8 | 19.51% | motif file (matrix) | svg |
| 521 | T G C A T A G C G A C T T G C A T G A C T G C A C G T A A G C T A G C T A G T C A G T C G T A C | GFY(?)/Promoter/Homer | 1e-3 | -7.994e+00 | 0.0007 | 365.0 | 0.92% | 1359.3 | 0.75% | motif file (matrix) | svg |
| 522 | C T A G C A T G C A T G T A C G A G T C G C A T A G C T C T A G A C G T A G T C G A C T A C T G A C T G A C T G T C G A | Zfp809(Zf)/ES-Zfp809-ChIP-Seq(GSE70799)/Homer | 1e-3 | -7.587e+00 | 0.0010 | 748.0 | 1.89% | 2985.4 | 1.65% | motif file (matrix) | svg |
| 523 | G C A T G C A T G A T C G A C T T C G A T C A G G C T A C G T A A C T G G T A C G C A T G C A T A G T C A G C T C G T A | HSF7(HSF)/colamp-HSF7-DAP-Seq(GSE60143)/Homer | 1e-3 | -7.501e+00 | 0.0011 | 598.0 | 1.51% | 2349.0 | 1.30% | motif file (matrix) | svg |
| 524 | T G C A C T G A A G T C G T C A A C T G A C T G C G T A C G T A C T G A A G C T | EWS:FLI1-fusion(ETS)/SK\_N\_MC-EWS:FLI1-ChIP-Seq(SRA014231)/Homer | 1e-3 | -7.380e+00 | 0.0012 | 2637.0 | 6.66% | 11258.6 | 6.22% | motif file (matrix) | svg |
| 525 | T A G C T C A G C A T G G C A T A G C T C G A T A T G C C G T A C G T A G T C A | CHR(?)/Hela-CellCycle-Expression/Homer | 1e-3 | -7.350e+00 | 0.0013 | 2189.0 | 5.53% | 9280.7 | 5.13% | motif file (matrix) | svg |
| 526 | G C A T G A T C A G C T G T A C G A T C C T A G C T A G G A T C T A C G C T G A | AT3G58630(Trihelix)/col-AT3G58630-DAP-Seq(GSE60143)/Homer | 1e-3 | -7.336e+00 | 0.0013 | 2143.0 | 5.41% | 9078.2 | 5.02% | motif file (matrix) | svg |
| 527 | G T C A G C A T G C T A C A G T C T A G G A T C C G T A C T G A C G T A C G A T | Oct2(POU,Homeobox)/Bcell-Oct2-ChIP-Seq(GSE21512)/Homer | 1e-3 | -7.191e+00 | 0.0015 | 945.0 | 2.39% | 3847.7 | 2.13% | motif file (matrix) | svg |
| 528 | A T G C G C A T C G A T G A T C A G C T C T G A A C T G C G T A C G T A T C A G T G A C C G A T G C A T G A T C C G A T | HSF21(HSF)/col-HSF21-DAP-Seq(GSE60143)/Homer | 1e-3 | -7.140e+00 | 0.0016 | 295.0 | 0.75% | 1090.8 | 0.60% | motif file (matrix) | svg |
| 529 | A C G T T G A C A G T C A G C T A G T C A G C T A C T G G A C T A G C T G A C T | REF6(Zf)/Arabidopsis-REF6-ChIP-Seq(GSE106942)/Homer | 1e-3 | -6.962e+00 | 0.0019 | 2146.0 | 5.42% | 9115.6 | 5.04% | motif file (matrix) | svg |
| 530 | C T G A T G C A T A G C T G A C T A C G T C A G C T G A G C T A T C A G G A C T | ELF1(ETS)/Jurkat-ELF1-ChIP-Seq(SRA014231)/Homer | 1e-3 | -6.939e+00 | 0.0019 | 4658.0 | 11.76% | 20299.0 | 11.22% | motif file (matrix) | svg |
| 531 | C T A G A C G T A G T C C G T A A C T G A G T C G C A T A C T G G C A T A G T C G A C T G A T C G C A T A G T C A G C T | ZNF317(Zf)/HEK293-ZNF317.GFP-ChIP-Seq(GSE58341)/Homer | 1e-2 | -6.424e+00 | 0.0032 | 415.0 | 1.05% | 1609.0 | 0.89% | motif file (matrix) | svg |
| 532 | A G T C G A C T A C T G G A T C G T A C C G T A T G A C A G T C C G A T A G C T A C G T A C G T C T A G G A C T C T G A | ZNF7(Zf)/HepG2-ZNF7.Flag-ChIP-Seq(Encode)/Homer | 1e-2 | -6.400e+00 | 0.0033 | 2490.0 | 6.29% | 10676.5 | 5.90% | motif file (matrix) | svg |
| 533 | A G C T C T A G T G A C C G T A A C G T C G A T A G T C A G T C C T G A C A T G | TEAD3(TEA)/HepG2-TEAD3-ChIP-Seq(Encode)/Homer | 1e-2 | -6.081e+00 | 0.0045 | 6401.0 | 16.17% | 28216.7 | 15.59% | motif file (matrix) | svg |
| 534 | A T G C A C T G C G A T T C A G A T G C C G T A C T G A T G C A C T G A G A C T A C T G G T C A | ABF1/SacCer-Promoters/Homer | 1e-2 | -5.890e+00 | 0.0054 | 4537.0 | 11.46% | 19858.4 | 10.97% | motif file (matrix) | svg |
| 535 | G A T C G A T C G C T A G T C A G A C T A T G C T C G A C G A T C G A T C T A G | HAT2(Homeobox)/colamp-HAT2-DAP-Seq(GSE60143)/Homer | 1e-2 | -5.802e+00 | 0.0059 | 4771.0 | 12.05% | 20917.5 | 11.56% | motif file (matrix) | svg |
| 536 | G A T C G T A C C G A T A C T G A C T G C G T A C G T A A C G T A C T G G A T C | TEAD(TEA)/Fibroblast-PU.1-ChIP-Seq(Unpublished)/Homer | 1e-2 | -5.751e+00 | 0.0062 | 2562.0 | 6.47% | 11045.8 | 6.10% | motif file (matrix) | svg |
| 537 | C T A G C T A G A G T C T C A G A C T G A C G T A C G T C T G A | MYB(HTH)/ERMYB-Myb-ChIPSeq(GSE22095)/Homer | 1e-2 | -5.699e+00 | 0.0065 | 17742.0 | 44.81% | 79739.4 | 44.06% | motif file (matrix) | svg |
| 538 | G C T A G C A T G A C T G C A T T C A G G T A C G C T A G C A T C T G A G C T A T A G C G C T A C T G A C G A T C T A G | OCT4-SOX2-TCF-NANOG(POU,Homeobox,HMG)/mES-Oct4-ChIP-Seq(GSE11431)/Homer | 1e-2 | -5.673e+00 | 0.0067 | 379.0 | 0.96% | 1478.1 | 0.82% | motif file (matrix) | svg |
| 539 | A G T C G A T C G C T A C G A T A C G T T A C G G C A T C T G A G A C T A C T G A G T C G C T A C T G A T C G A C A G T | Oct4:Sox17(POU,Homeobox,HMG)/F9-Sox17-ChIP-Seq(GSE44553)/Homer | 1e-2 | -5.428e+00 | 0.0085 | 472.0 | 1.19% | 1881.2 | 1.04% | motif file (matrix) | svg |
| 540 | C G T A C T A G G A C T G T C A G T C A C G T A A G T C C G T A T C G A T C G A T C G A C G T A C T G A C T A G G C T A C G T A T A G C C G T A C G A T C G T A | FOXA1:AR(Forkhead,NR)/LNCAP-AR-ChIP-Seq(GSE27824)/Homer | 1e-2 | -5.377e+00 | 0.0090 | 121.0 | 0.31% | 418.5 | 0.23% | motif file (matrix) | svg |
| 541 | G C T A A G C T G T A C G C A T A G C T T C G A C T G A A G T C A G T C T A C G A C G T G A C T T A C G C T A G C G T A | ZML1(C2C2gata)/colamp-ZML1-DAP-Seq(GSE60143)/Homer | 1e-2 | -5.299e+00 | 0.0097 | 296.0 | 0.75% | 1139.8 | 0.63% | motif file (matrix) | svg |
| 542 | T C A G C G T A A G T C A G C T C G T A A G T C C T G A C G T A A G T C G C A T A G T C A G T C A G T C C T G A A C T G T G C A T C G A C A T G A T C G G A T C | Ronin(THAP)/ES-Thap11-ChIP-Seq(GSE51522)/Homer | 1e-2 | -5.234e+00 | 0.0103 | 48.0 | 0.12% | 139.2 | 0.08% | motif file (matrix) | svg |
| 543 | T G C A A G C T C T G A A T C G G A C T C T A G G T A C G A T C G T C A A G T C G T A C G A C T C T A G A T C G G C A T C A T G C A T G G A T C G T A C C T G A | CTCF(Zf)/CD4+-CTCF-ChIP-Seq(Barski\_et\_al.)/Homer | 1e-2 | -5.227e+00 | 0.0104 | 334.0 | 0.84% | 1301.0 | 0.72% | motif file (matrix) | svg |
| 544 | G C A T G C A T G T A C G A C T T C G A A C T G G C T A C G T A A T C G T G A C G C A T G A C T A G T C A G C T C T G A | AGL95(ND)/col-AGL95-DAP-Seq(GSE60143)/Homer | 1e-2 | -5.001e+00 | 0.0130 | 309.0 | 0.78% | 1202.3 | 0.66% | motif file (matrix) | svg |
| 545 | T G C A C T A G C T A G C T G A C A T G A C T G T G C A G A T C G T C A T G C A G T C A G T C A A G C T C T A G G C A T | ZNF675(Zf)/HEK293-ZNF675.GFP-ChIP-Seq(GSE58341)/Homer | 1e-2 | -4.866e+00 | 0.0148 | 699.0 | 1.77% | 2882.2 | 1.59% | motif file (matrix) | svg |
