## Supplemental dataset for "Hybrid CNN and Multi-Head Attention Model for Analyzing Epigenetic Mechanisms and Gene Expression Across Fungal Phylogenetic Distances": FgramModel_NcrassaTest_K27me3locs_homerResults.html

/projects/wg-feeds/SHAP/FgramModel\_NcrassaTest\_K27me3locs\_SHAP\_noDup\_HOMER// - Homer de novo Motif Results


### Homer *de novo* Motif Results (/projects/wg-feeds/SHAP/FgramModel\_NcrassaTest\_K27me3locs\_SHAP\_noDup\_HOMER//)

Non-redundant Motif File of Results  
Known Motif Enrichment Results  
Gene Ontology Enrichment Results  
If Homer is having trouble matching a motif to a known motif, try copy/pasting the matrix file into
STAMP  
More information on motif finding results: HOMER
| Description of Results
| Tips
  
Total target sequences = 39597  
Total background sequences = 179793  
\* - possible false positive  

|  |  |  |  |  |  |  |  |  |
| --- | --- | --- | --- | --- | --- | --- | --- | --- |
| Rank | Motif | P-value | log P-pvalue | % of Targets | % of Background | STD(Bg STD) | Best Match/Details | Motif File |
| 1 | T A G C T C G A C G A T T A G C T C G A G C A T T A G C T C G A G C A T T A G C T C G A G C T A | 1e-3136 | -7.221e+03 | 43.39% | 14.76% | 510.9bp (480.9bp) | ZML2(C2C2gata)/col-ZML2-DAP-Seq(GSE60143)/Homer(0.818) More Information \| Similar Motifs Found | motif file (matrix) |
| 2 | A G T C T A G C C G A T A C G T A T G C G C A T A C G T A T G C G C A T A G C T A T G C C G T A | 1e-2486 | -5.725e+03 | 46.71% | 19.78% | 518.0bp (483.3bp) | ZML2(C2C2gata)/col-ZML2-DAP-Seq(GSE60143)/Homer(0.659) More Information \| Similar Motifs Found | motif file (matrix) |
| 3 | G A C T A G C T A G C T A G C T A G C T A G C T A G C T A G C T A G C T A G C T A G C T A G C T | 1e-2049 | -4.719e+03 | 28.59% | 9.01% | 514.1bp (391.5bp) | SeqBias: polyA-repeat(0.850) More Information \| Similar Motifs Found | motif file (matrix) |
| 4 | A T C G C A G T A C G T A T C G C G T A C G A T A C T G C T A G G C T A C A T G C A T G A T G C | 1e-1459 | -3.361e+03 | 50.41% | 28.57% | 499.3bp (471.0bp) | SRSF9(RRM)/Homo\_sapiens-RNCMPT00067-PBM/HughesRNA(0.648) More Information \| Similar Motifs Found | motif file (matrix) |
| 5 | G A C T A T G C T C G A C G T A A T C G T C G A G C A T A T G C | 1e-1349 | -3.107e+03 | 58.66% | 36.98% | 517.7bp (486.3bp) | ftz-f1/MA2311.1/Jaspar(0.889) More Information \| Similar Motifs Found | motif file (matrix) |
| 6 | A T C G G C A T G A C T A T C G G C A T A G T C T C A G C G T A | 1e-1322 | -3.044e+03 | 49.09% | 28.37% | 511.9bp (490.1bp) | ZBTB38(Zf)/Hela-ZBTB38-ChIP-seq(GSE108618)/Homer(0.673) More Information \| Similar Motifs Found | motif file (matrix) |
| 7 | C A T G A T C G C T A G C T G A C G T A C T A G C T A G C T G A C T G A C T A G | 1e-1008 | -2.323e+03 | 37.50% | 20.76% | 477.5bp (438.0bp) | PCBP2(KH)/Homo\_sapiens-RNCMPT00044-PBM/HughesRNA(0.838) More Information \| Similar Motifs Found | motif file (matrix) |
| 8 | A T G C T G A C T G C A C T G A T A C G T G A C G T C A T C G A T C A G T G A C T G C A C T G A | 1e-975 | -2.247e+03 | 40.51% | 23.53% | 516.9bp (476.0bp) | NAC037/MA2047.2/Jaspar(0.682) More Information \| Similar Motifs Found | motif file (matrix) |
| 9 | T C A G A C G T T C G A T A C G T A C G A G C T C G T A A T C G T A C G G A C T C G T A A T C G | 1e-784 | -1.807e+03 | 7.84% | 1.61% | 498.4bp (459.5bp) | PK06182.1/MA2354.1/Jaspar(0.842) More Information \| Similar Motifs Found | motif file (matrix) |
| 10 | A G T C C T A G A G T C A G T C C T A G G A T C A G T C C T A G | 1e-752 | -1.732e+03 | 19.78% | 8.89% | 536.8bp (469.5bp) | Zm00001d005892/MA1819.2/Jaspar(0.997) More Information \| Similar Motifs Found | motif file (matrix) |
| 11 | T A G C C T G A C G T A G C A T A C T G T A C G G T A C T C A G | 1e-750 | -1.728e+03 | 45.37% | 29.79% | 509.8bp (490.3bp) | Hr39/MA2244.1/Jaspar(0.745) More Information \| Similar Motifs Found | motif file (matrix) |
| 12 | A C G T C G T A A G T C A G T C A C G T A G T C A C G T C G T A | 1e-680 | -1.566e+03 | 6.09% | 1.08% | 492.3bp (472.5bp) | MOT3/Literature(Harbison)/Yeast(0.671) More Information \| Similar Motifs Found | motif file (matrix) |
| 13 | A G T C G A C T A T G C A G C T A T G C A G C T T A G C A G C T T A G C A G C T G A T C A G C T | 1e-378 | -8.707e+02 | 22.52% | 13.83% | 482.7bp (454.2bp) | RAMOSA1/MA1416.1/Jaspar(0.947) More Information \| Similar Motifs Found | motif file (matrix) |
| 14 | C G T A C G T A A C T G A C G T A C T G A C T G C G T A C G T A | 1e-345 | -7.952e+02 | 5.03% | 1.47% | 493.7bp (470.6bp) | MOD(RRM)/Drosophila\_melanogaster-RNCMPT00140-PBM/HughesRNA(0.888) More Information \| Similar Motifs Found | motif file (matrix) |
| 15 | A G T C C G T A A C G T A G T C C G T A A C G T C T G A A G C T | 1e-314 | -7.231e+02 | 13.25% | 7.12% | 521.0bp (475.4bp) | RIN(RRM)/Drosophila\_melanogaster-RNCMPT00138-PBM/HughesRNA(0.813) More Information \| Similar Motifs Found | motif file (matrix) |
| 16 | G A C T C T A G A G C T T C A G C G A T T C A G G A C T T C A G A G C T T C A G G A C T T C A G | 1e-252 | -5.809e+02 | 1.99% | 0.29% | 451.9bp (440.0bp) | cg/MA2107.1/Jaspar(0.902) More Information \| Similar Motifs Found | motif file (matrix) |
| 17 | A C T G A C G T C G T A A C T G A C G T A C T G A C G T C G T A | 1e-208 | -4.797e+02 | 3.33% | 1.04% | 442.2bp (472.2bp) | SFPQ(RRM)/Homo\_sapiens-RNCMPT00177-PBM/HughesRNA(0.858) More Information \| Similar Motifs Found | motif file (matrix) |
| 18 | C T G A A T G C A G C T C T G A A G T C G A C T C T G A A G T C A G C T C T G A A G T C A G C T | 1e-195 | -4.502e+02 | 2.48% | 0.64% | 517.3bp (451.5bp) | SPL4D/MA2445.1/Jaspar(0.735) More Information \| Similar Motifs Found | motif file (matrix) |
| 19 | C T A G A C T G C G T A A C G T A C T G A C T G C G T A A C G T A C T G A C T G | 1e-190 | -4.392e+02 | 1.59% | 0.25% | 403.9bp (461.7bp) | HOXA1(Homeobox)/mES-Hoxa1-ChIP-Seq(SRP084292)/Homer(0.779) More Information \| Similar Motifs Found | motif file (matrix) |
| 20 | C T A G A C T G A C T G A C T G A C T G A C T G A C T G A C T G A C T G A C T G A C T G C A T G | 1e-187 | -4.318e+02 | 0.87% | 0.03% | 597.9bp (184.3bp) | SeqBias: polyC-repeat(0.922) More Information \| Similar Motifs Found | motif file (matrix) |
| 21 | C T A G G C T A G T A C C A G T A T C G C T A G T C G A A G T C C A G T A C T G T C A G T C G A | 1e-121 | -2.804e+02 | 1.53% | 0.39% | 439.4bp (502.5bp) | Knotted(Homeobox)/Corn-KN1-ChIP-Seq(GSE39161)/Homer(0.733) More Information \| Similar Motifs Found | motif file (matrix) |
| 22 | C T G A C T A G A C G T A G T C C G T A A C T G A G C T A G T C C G T A C T A G A C G T A G T C | 1e-54 | -1.256e+02 | 0.30% | 0.02% | 748.8bp (476.3bp) | ASH1/Literature(Harbison)/Yeast(0.668) More Information \| Similar Motifs Found | motif file (matrix) |
| 23 | A C G T C G T A A G C T C G T A A C G T C G T A A G C T C G T A | 1e-51 | -1.197e+02 | 1.03% | 0.37% | 458.6bp (440.9bp) | Cf2/MA0015.2/Jaspar(0.933) More Information \| Similar Motifs Found | motif file (matrix) |
| 24 | C G T A A G T C A G T C A C T G C G T A A G T C A G T C A C T G C G T A A G T C A G T C A C T G | 1e-28 | -6.507e+01 | 0.12% | 0.00% | 353.7bp (295.3bp) | MYB55/MA1041.1/Jaspar(0.689) More Information \| Similar Motifs Found | motif file (matrix) |
| 25 | C G A T C G T A G C A T C G A T C G T A C G T A C G A T C G T A C G A T C G A T C G T A C G A T | 1e-17 | -4.141e+01 | 0.26% | 0.08% | 347.0bp (301.9bp) | SeqBias: A/T bias(0.824) More Information \| Similar Motifs Found | motif file (matrix) |
