## Supplemental dataset for "Hybrid CNN and Multi-Head Attention Model for Analyzing Epigenetic Mechanisms and Gene Expression Across Fungal Phylogenetic Distances": FgramModel_NcrassaTest_K27me3locs_knownResults.html

/projects/wg-feeds/SHAP/FgramModel\_NcrassaTest\_K27me3locs\_SHAP\_noDup\_HOMER/ - Homer Known Motif Enrichment Results


### Homer Known Motif Enrichment Results (/projects/wg-feeds/SHAP/FgramModel\_NcrassaTest\_K27me3locs\_SHAP\_noDup\_HOMER/)

Homer *de novo* Motif Results  
Gene Ontology Enrichment Results  
Known Motif Enrichment Results (txt file)  
Total Target Sequences = 39586, Total Background Sequences = 179517

|  |  |  |  |  |  |  |  |  |  |  |  |
| --- | --- | --- | --- | --- | --- | --- | --- | --- | --- | --- | --- |
| Rank | Motif | Name | P-value | log P-pvalue | q-value (Benjamini) | # Target Sequences with Motif | % of Targets Sequences with Motif | # Background Sequences with Motif | % of Background Sequences with Motif | Motif File | SVG |
| 1 | T A G C G T C A G A C T T A G C G T C A G A C T A G T C G C T A G A C T G A T C | ZML2(C2C2gata)/col-ZML2-DAP-Seq(GSE60143)/Homer | 1e-1663 | -3.829e+03 | 0.0000 | 6211.0 | 15.69% | 5654.8 | 3.15% | motif file (matrix) | svg |
| 2 | A T G C C A G T A C G T A G T C C A T G A C G T A G T C A C G T A G C T G A T C | Unknown4/Arabidopsis-Promoters/Homer | 1e-1624 | -3.740e+03 | 0.0000 | 14790.0 | 37.35% | 30285.9 | 16.85% | motif file (matrix) | svg |
| 3 | C T A G T C A G C T G A C T G A C T A G C T G A C A T G C A T G C T G A C T A G C T A G C G T A C T A G C G T A G T C A | TF3A(C2H2)/col-TF3A-DAP-Seq(GSE60143)/Homer | 1e-1394 | -3.211e+03 | 0.0000 | 12276.0 | 31.00% | 24029.5 | 13.37% | motif file (matrix) | svg |
| 4 | G C T A G C A T A C G T A C G T A G C T G A T C G A C T G A C T A G C T A C G T A C G T A G C T | RLR1?/SacCer-Promoters/Homer | 1e-1103 | -2.541e+03 | 0.0000 | 4082.0 | 10.31% | 3559.4 | 1.98% | motif file (matrix) | svg |
| 5 | A C G T A C G T A C G T A C G T A C G T A C G T A C G T A C G T A C G T A C G T | VRN1(ABI3VP1)/col-VRN1-DAP-Seq(GSE60143)/Homer | 1e-1071 | -2.468e+03 | 0.0000 | 1921.0 | 4.85% | 323.1 | 0.18% | motif file (matrix) | svg |
| 6 | A C G T A T G C A C G T A C G T A G C T A G T C A G C T A G C T A G C T A G C T A G C T | hTCT(CPE) | 1e-829 | -1.909e+03 | 0.0000 | 16154.0 | 40.80% | 44921.8 | 24.99% | motif file (matrix) | svg |
| 7 | G A T C G C A T A G T C A G C T G A T C A G C T G A T C G A C T G A T C G A C T A G T C A C G T G A T C A G C T G A T C | GAGA-repeat/SacCer-Promoters/Homer | 1e-755 | -1.740e+03 | 0.0000 | 21854.0 | 55.19% | 70011.6 | 38.95% | motif file (matrix) | svg |
| 8 | G T C A T G C A T G C A G C T A C G T A G C T A G C T A G C T A | REM19(REM)/colamp-REM19-DAP-Seq(GSE60143)/Homer | 1e-737 | -1.698e+03 | 0.0000 | 4509.0 | 11.39% | 6426.6 | 3.58% | motif file (matrix) | svg |
| 9 | T G A C C T A G T C A G G T C A C G T A T C A G C G A T T C A G T C G A T G C A C T G A T A G C | PU.1-IRF(ETS:IRF)/Bcell-PU.1-ChIP-Seq(GSE21512)/Homer | 1e-622 | -1.433e+03 | 0.0000 | 9238.0 | 23.33% | 22217.1 | 12.36% | motif file (matrix) | svg |
| 10 | A C G T G A C T T A G C C G T A C T G A C A T G C T A G G A C T G A T C C G T A | Nr5a2(NR)/Pancreas-LRH1-ChIP-Seq(GSE34295)/Homer | 1e-547 | -1.260e+03 | 0.0000 | 7292.0 | 18.42% | 16548.7 | 9.21% | motif file (matrix) | svg |
| 11 | G A T C C A G T T A G C A G T C A C T G A G T C A G T C C T A G G A C T G T A C | LEP(AP2EREBP)/col-LEP-DAP-Seq(GSE60143)/Homer | 1e-547 | -1.260e+03 | 0.0000 | 6023.0 | 15.21% | 12429.0 | 6.92% | motif file (matrix) | svg |
| 12 | C T G A G C A T A C T G C T A G A G T C A C T G A C T G A G T C A C T G T C A G | AT4G18450(AP2EREBP)/col-AT4G18450-DAP-Seq(GSE60143)/Homer | 1e-541 | -1.246e+03 | 0.0000 | 7467.0 | 18.86% | 17214.3 | 9.58% | motif file (matrix) | svg |
| 13 | C T G A C G A T C A T G A T C G G C A T C A T G G C T A A G T C | ASHR1(ND)/col-ASHR1-DAP-Seq(GSE60143)/Homer | 1e-540 | -1.245e+03 | 0.0000 | 12508.0 | 31.59% | 35518.6 | 19.76% | motif file (matrix) | svg |
| 14 | G C A T G A T C C T A G C G T A G C A T C G T A G C A T A G T C C T A G C G T A G C A T C G A T | AT5G22990(C2H2)/col-AT5G22990-DAP-Seq(GSE60143)/Homer | 1e-534 | -1.231e+03 | 0.0000 | 8464.0 | 21.38% | 20724.2 | 11.53% | motif file (matrix) | svg |
| 15 | G T A C A C T G A G T C A G T C C T A G G A T C G T A C C T G A | CRF4(AP2EREBP)/colamp-CRF4-DAP-Seq(GSE60143)/Homer | 1e-527 | -1.215e+03 | 0.0000 | 10056.0 | 25.40% | 26521.2 | 14.76% | motif file (matrix) | svg |
| 16 | C A T G A G T C G T A C A C T G A T G C A G T C C A T G G A T C G A T C C T G A | ERF5(AP2EREBP)/colamp-ERF5-DAP-Seq(GSE60143)/Homer | 1e-521 | -1.202e+03 | 0.0000 | 7663.0 | 19.35% | 18131.1 | 10.09% | motif file (matrix) | svg |
| 17 | T G A C C G T A C T G A A C T G A C T G G A C T G A T C T G C A G T A C T A C G | SF1(NR)/H295R-Nr5a1-ChIP-Seq(GSE44220)/Homer | 1e-510 | -1.175e+03 | 0.0000 | 4925.0 | 12.44% | 9420.4 | 5.24% | motif file (matrix) | svg |
| 18 | C T A G C T G A C T A G C T G A C T A G C T G A C T A G C T G A C T A G C T G A | SeqBias: GA-repeat | 1e-509 | -1.173e+03 | 0.0000 | 30272.0 | 76.45% | 115162.4 | 64.07% | motif file (matrix) | svg |
| 19 | A C G T G A C T A T G C G C T A C T G A C T A G A C T G G A C T A G T C C G T A | Nr5a2(NR)/mES-Nr5a2-ChIP-Seq(GSE19019)/Homer | 1e-504 | -1.163e+03 | 0.0000 | 5713.0 | 14.43% | 11911.8 | 6.63% | motif file (matrix) | svg |
| 20 | C T A G G C A T A C T G C T A G A G T C A C T G A C T G A G T C A C T G T C A G | ERF10(AP2EREBP)/col-ERF10-DAP-Seq(GSE60143)/Homer | 1e-499 | -1.149e+03 | 0.0000 | 11422.0 | 28.85% | 32099.7 | 17.86% | motif file (matrix) | svg |
| 21 | C G T A G A C T C A T G C T A G A G T C A C T G A C T G G T A C C A T G T A C G | ERF3(AP2EREBP)/colamp-ERF3-DAP-Seq(GSE60143)/Homer | 1e-480 | -1.106e+03 | 0.0000 | 11718.0 | 29.59% | 33575.7 | 18.68% | motif file (matrix) | svg |
| 22 | G A T C G C T A G T A C A G T C G C T A T G C A G T A C G A T C C G T A G A C T | MYB83(MYB)/colamp-MYB83-DAP-Seq(GSE60143)/Homer | 1e-472 | -1.087e+03 | 0.0000 | 17058.0 | 43.08% | 55250.0 | 30.74% | motif file (matrix) | svg |
| 23 | A C G T A C T G C G T A A C G T A C T G A C T G C G T A C G T A | HAP3(CCAATHAP3)/col-HAP3-DAP-Seq(GSE60143)/Homer | 1e-450 | -1.037e+03 | 0.0000 | 4972.0 | 12.56% | 10182.2 | 5.67% | motif file (matrix) | svg |
| 24 | G C A T A C T G C T A G A G T C A C T G A C T G A G T C A C G T | ERF105(AP2EREBP)/colamp-ERF105-DAP-Seq(GSE60143)/Homer | 1e-449 | -1.035e+03 | 0.0000 | 16274.0 | 41.10% | 52480.7 | 29.20% | motif file (matrix) | svg |
| 25 | A T G C A G T C A C T G A T G C A G T C A C T G A G T C G T A C | SHN3(AP2EREBP)/col-SHN3-DAP-Seq(GSE60143)/Homer | 1e-443 | -1.020e+03 | 0.0000 | 5033.0 | 12.71% | 10455.7 | 5.82% | motif file (matrix) | svg |
| 26 | C G T A C T A G C A T G A G C T C T G A C A T G C A G T C G A T C T A G C T A G | MYB30(MYB)/colamp-MYB30-DAP-Seq(GSE60143)/Homer | 1e-433 | -9.980e+02 | 0.0000 | 12087.0 | 30.53% | 35902.4 | 19.98% | motif file (matrix) | svg |
| 27 | G A C T G A T C G A T C G C T A G T A C A G T C G C T A C T G A G T A C G A T C G C T A G A C T | MYB13(MYB)/col-MYB13-DAP-Seq(GSE60143)/Homer | 1e-431 | -9.944e+02 | 0.0000 | 8608.0 | 21.74% | 22795.8 | 12.68% | motif file (matrix) | svg |
| 28 | G C T A C G T A C G T A G A C T C A T G C T A G G A T C A C T G T C A G G A T C A C T G T A C G | ERF9(AP2EREBP)/colamp-ERF9-DAP-Seq(GSE60143)/Homer | 1e-425 | -9.787e+02 | 0.0000 | 5732.0 | 14.48% | 12916.7 | 7.19% | motif file (matrix) | svg |
| 29 | G C A T A G C T A C G T A C G T A C T G A C G T G A T C A C G T A C G T A G C T C G A T G C A T A G T C G A C T C A G T | IDD5(C2H2)/colamp-IDD5-DAP-Seq(GSE60143)/Homer | 1e-422 | -9.721e+02 | 0.0000 | 4667.0 | 11.79% | 9541.2 | 5.31% | motif file (matrix) | svg |
| 30 | C G T A T A G C T A G C T G C A A C T G C T A G C G T A C G T A T C A G G A C T | EHF(ETS)/LoVo-EHF-ChIP-Seq(GSE49402)/Homer | 1e-406 | -9.358e+02 | 0.0000 | 9874.0 | 24.94% | 27923.7 | 15.54% | motif file (matrix) | svg |
| 31 | C T G A C G A T C T A G T C A G G A T C C T G A T C A G G A T C C T G A A C T G A G T C G C T A A C G T A G T C G C A T | PRDM9(Zf)/Testis-DMC1-ChIP-Seq(GSE35498)/Homer | 1e-404 | -9.324e+02 | 0.0000 | 3677.0 | 9.29% | 6731.4 | 3.75% | motif file (matrix) | svg |
| 32 | A C T G A C T G A G T C A C T G A C T G A G T C A C G T C T A G | ERF1(AP2EREBP)/colamp-ERF1-DAP-Seq(GSE60143)/Homer | 1e-402 | -9.271e+02 | 0.0000 | 8602.0 | 21.72% | 23264.7 | 12.94% | motif file (matrix) | svg |
| 33 | A C T G C T A G A G T C A C T G A C T G A T G C A C T G T A C G | ESE1(AP2EREBP)/col-ESE1-DAP-Seq(GSE60143)/Homer | 1e-401 | -9.244e+02 | 0.0000 | 10592.0 | 26.75% | 30746.7 | 17.11% | motif file (matrix) | svg |
| 34 | C A T G G A C T C T A G C A T G C A G T C G A T C T A G C A T G C G A T C G T A C T A G C A G T C G A T C T A G C A T G | AT1G24250(Orphan)/col-AT1G24250-DAP-Seq(GSE60143)/Homer | 1e-396 | -9.128e+02 | 0.0000 | 4632.0 | 11.70% | 9718.3 | 5.41% | motif file (matrix) | svg |
| 35 | A T G C T C G A T A C G A C G T A T G C A G T C A C G T A G T C A G T C G A T C | Znf263(Zf)/K562-Znf263-ChIP-Seq(GSE31477)/Homer | 1e-395 | -9.114e+02 | 0.0000 | 12889.0 | 32.55% | 39859.2 | 22.18% | motif file (matrix) | svg |
| 36 | C T G A T C A G G T A C G C T A A C T G T G A C G C A T C A T G | SCL(bHLH)/HPC7-Scl-ChIP-Seq(GSE13511)/Homer | 1e-395 | -9.112e+02 | 0.0000 | 22202.0 | 56.07% | 79570.6 | 44.27% | motif file (matrix) | svg |
| 37 | A C T G A C T G A G T C A C T G A C T G A G T C A C G T T C A G | ERF2(AP2EREBP)/colamp-ERF2-DAP-Seq(GSE60143)/Homer | 1e-393 | -9.070e+02 | 0.0000 | 9261.0 | 23.39% | 25855.9 | 14.39% | motif file (matrix) | svg |
| 38 | C G T A G A C T C A T G C T A G A G T C A C T G C T A G A G T C C A T G C T A G | ERF7(AP2EREBP)/col-ERF7-DAP-Seq(GSE60143)/Homer | 1e-389 | -8.973e+02 | 0.0000 | 17987.0 | 45.43% | 61162.4 | 34.03% | motif file (matrix) | svg |
| 39 | A T G C G T A C A G T C A G T C A C G T A C G T C G A T A C G T | AT5G02460(C2C2dof)/col-AT5G02460-DAP-Seq(GSE60143)/Homer | 1e-384 | -8.862e+02 | 0.0000 | 15035.0 | 37.97% | 48838.6 | 27.17% | motif file (matrix) | svg |
| 40 | G T A C A C T G A T G C A G T C C T A G G A T C G T A C C T G A G A C T G C A T C G A T G A C T | RAP212(AP2EREBP)/col-RAP212-DAP-Seq(GSE60143)/Homer | 1e-381 | -8.792e+02 | 0.0000 | 12536.0 | 31.66% | 38756.6 | 21.56% | motif file (matrix) | svg |
| 41 | A T G C G T A C A C T G A G T C A G T C A C T G A G T C G T A C | ERF73(AP2EREBP)/col-ERF73-DAP-Seq(GSE60143)/Homer | 1e-377 | -8.695e+02 | 0.0000 | 9154.0 | 23.12% | 25755.3 | 14.33% | motif file (matrix) | svg |
| 42 | C G T A G C A T C G T A C G T A G C A T A C T G C G A T A G T C A C T G A C T G G A C T C T A G | AT1G71450(AP2EREBP)/col-AT1G71450-DAP-Seq(GSE60143)/Homer | 1e-376 | -8.667e+02 | 0.0000 | 22796.0 | 57.57% | 82789.0 | 46.06% | motif file (matrix) | svg |
| 43 | C G A T A C G T A C G T A G C T A G C T G A T C G A T C G C T A A G C T A C G T A T C G T A C G | NFATC2(RHD)/Islets-NFATC2-ChIP-Seq(GSE158496)/Homer | 1e-371 | -8.552e+02 | 0.0000 | 12544.0 | 31.68% | 39020.0 | 21.71% | motif file (matrix) | svg |
| 44 | C G A T A G C T T G C A A C T G A G T C T G A C C T A G G T A C A G T C C G T A G C A T G C A T | ERF13(AP2EREBP)/colamp-ERF13-DAP-Seq(GSE60143)/Homer | 1e-368 | -8.476e+02 | 0.0000 | 14182.0 | 35.82% | 45726.2 | 25.44% | motif file (matrix) | svg |
| 45 | C T G A T C A G A G T C C G T A A T C G A T G C C G A T A C T G A G T C G A C T A T C G A G T C | MyoD(bHLH)/Myotube-MyoD-ChIP-Seq(GSE21614)/Homer | 1e-357 | -8.229e+02 | 0.0000 | 5036.0 | 12.72% | 11494.4 | 6.40% | motif file (matrix) | svg |
| 46 | A C T G C T A G A G T C A C T G A C T G A G T C A C T G T A C G | ERF104(AP2EREBP)/col-ERF104-DAP-Seq(GSE60143)/Homer | 1e-346 | -7.975e+02 | 0.0000 | 13655.0 | 34.49% | 44094.1 | 24.53% | motif file (matrix) | svg |
| 47 | A G T C G A C T G A T C C G T A G T A C A G T C G C T A C G T A G T A C A G T C G T A C G T A C | MYB63(MYB)/col-MYB63-DAP-Seq(GSE60143)/Homer | 1e-343 | -7.909e+02 | 0.0000 | 6937.0 | 17.52% | 18243.7 | 10.15% | motif file (matrix) | svg |
| 48 | C T A G C T A G T C G A C T A G C G T A A T C G T C G A A C T G C T G A T C G A C T G A T A C G | FRS9(ND)/col-FRS9-DAP-Seq(GSE60143)/Homer | 1e-339 | -7.818e+02 | 0.0000 | 2016.0 | 5.09% | 2737.7 | 1.52% | motif file (matrix) | svg |
| 49 | T C G A T G C A C A G T T C G A G A T C A G T C C G T A C G T A A C T G A G T C C G T A C G T A T C A G C G A T A G T C | AT5G25475(ABI3VP1)/col-AT5G25475-DAP-Seq(GSE60143)/Homer | 1e-338 | -7.787e+02 | 0.0000 | 11420.0 | 28.84% | 35297.5 | 19.64% | motif file (matrix) | svg |
| 50 | C A T G G C T A C T A G T A C G C G T A T C A G C G T A A C T G C G T A C A T G C T G A C G T A | BPC1(BBRBPC)/colamp-BPC1-DAP-Seq(GSE60143)/Homer | 1e-333 | -7.674e+02 | 0.0000 | 4278.0 | 10.80% | 9337.7 | 5.20% | motif file (matrix) | svg |
| 51 | A G C T G A T C G A T C C G T A G T A C A G T C C G A T C T G A G T A C G A T C C G T A G A C T | ATY19(MYB)/col-ATY19-DAP-Seq(GSE60143)/Homer | 1e-332 | -7.646e+02 | 0.0000 | 9148.0 | 23.10% | 26614.7 | 14.81% | motif file (matrix) | svg |
| 52 | C G A T C T A G A C T G A G C T C T G A A C T G A C G T A C G T C T A G C T A G | MYB96(MYB)/colamp-MYB96-DAP-Seq(GSE60143)/Homer | 1e-327 | -7.546e+02 | 0.0000 | 10383.0 | 26.22% | 31458.7 | 17.50% | motif file (matrix) | svg |
| 53 | C G T A C T A G C A T G G A C T C T G A A C T G A C G T A C G T C T A G C T A G C A T G T C G A | MYB94(MYB)/col-MYB94-DAP-Seq(GSE60143)/Homer | 1e-321 | -7.404e+02 | 0.0000 | 5416.0 | 13.68% | 13267.3 | 7.38% | motif file (matrix) | svg |
| 54 | C T G A G A C T C A T G C T A G A G T C A C T G A C T G A G T C A C T G T C A G | ERF11(AP2EREBP)/col-ERF11-DAP-Seq(GSE60143)/Homer | 1e-321 | -7.393e+02 | 0.0000 | 14029.0 | 35.43% | 46260.2 | 25.74% | motif file (matrix) | svg |
| 55 | A C T G C T A G A G T C A C T G A C T G A G T C A C G T C T A G | AT5G23930(mTERF)/col-AT5G23930-DAP-Seq(GSE60143)/Homer | 1e-316 | -7.291e+02 | 0.0000 | 15150.0 | 38.26% | 51039.1 | 28.40% | motif file (matrix) | svg |
| 56 | G C A T C G A T G A C T T G C A A C T G A G T C T G A C A C T G G A T C A G T C C G T A G A C T | ERF15(AP2EREBP)/colamp-ERF15-DAP-Seq(GSE60143)/Homer | 1e-308 | -7.115e+02 | 0.0000 | 17331.0 | 43.77% | 60514.8 | 33.67% | motif file (matrix) | svg |
| 57 | C G T A C T A G C A G T A C G T C G T A A C T G C A T G G C A T T C A G C T G A | MYB49(MYB)/col-MYB49-DAP-Seq(GSE60143)/Homer | 1e-305 | -7.035e+02 | 0.0000 | 11291.0 | 28.52% | 35541.2 | 19.77% | motif file (matrix) | svg |
| 58 | G C T A C G T A C G T A G C A T C A T G C T A G G A T C A C T G T C A G G A T C A C T G T C A G | ERF4(AP2EREBP)/colamp-ERF4-DAP-Seq(GSE60143)/Homer | 1e-301 | -6.933e+02 | 0.0000 | 16137.0 | 40.76% | 55632.7 | 30.95% | motif file (matrix) | svg |
| 59 | C A T G A G C T T A C G G T C A G T A C T A G C A G C T G A C T A T C G T C G A | Esrrb(NR)/mES-Esrrb-ChIP-Seq(GSE11431)/Homer | 1e-297 | -6.860e+02 | 0.0000 | 6417.0 | 16.21% | 17157.3 | 9.55% | motif file (matrix) | svg |
| 60 | C T A G C A T G G A C T C G T A C T A G A C T G A C G T C T A G C T A G T C A G | MYB17(MYB)/colamp-MYB17-DAP-Seq(GSE60143)/Homer | 1e-292 | -6.740e+02 | 0.0000 | 7723.0 | 19.50% | 22037.4 | 12.26% | motif file (matrix) | svg |
| 61 | G A T C G A T C G A T C C G T A G T A C A G T C G C A T C G T A G T A C G A T C | MYB58(MYB)/colamp-MYB58-DAP-Seq(GSE60143)/Homer | 1e-287 | -6.612e+02 | 0.0000 | 11435.0 | 28.88% | 36563.0 | 20.34% | motif file (matrix) | svg |
| 62 | G A C T G A C T G A T C C G T A G T A C A G T C G C A T C G T A G T A C G A T C G C A T G C T A | MYB74(MYB)/colamp-MYB74-DAP-Seq(GSE60143)/Homer | 1e-286 | -6.594e+02 | 0.0000 | 7482.0 | 18.90% | 21264.8 | 11.83% | motif file (matrix) | svg |
| 63 | G T A C A C T G A T G C T G A C C T A G G A C T G T A C C G T A G C A T G C A T | ERF8(AP2EREBP)/colamp-ERF8-DAP-Seq(GSE60143)/Homer | 1e-285 | -6.578e+02 | 0.0000 | 17065.0 | 43.10% | 60048.0 | 33.41% | motif file (matrix) | svg |
| 64 | G A C T A G C T A G C T C T A G A C G T G A T C A C G T A C G T G A C T C G A T G C A T A G T C | IDD4(C2H2)/col-IDD4-DAP-Seq(GSE60143)/Homer | 1e-280 | -6.467e+02 | 0.0000 | 5916.0 | 14.94% | 15662.8 | 8.71% | motif file (matrix) | svg |
| 65 | C T A G A C T G A C G T C G T A A C T G A C T G A G C T C T A G T C A G C T A G | MYB93(MYB)/colamp-MYB93-DAP-Seq(GSE60143)/Homer | 1e-270 | -6.217e+02 | 0.0000 | 12244.0 | 30.92% | 40276.4 | 22.41% | motif file (matrix) | svg |
| 66 | C A T G G A C T T A C G G T C A G T A C G A T C G A C T A G C T A T C G T C G A T A C G T A G C | ERRg(NR)/Kidney-ESRRG-ChIP-Seq(GSE104905)/Homer | 1e-268 | -6.190e+02 | 0.0000 | 7759.0 | 19.60% | 22658.5 | 12.61% | motif file (matrix) | svg |
| 67 | G C T A C G T A C G T A G C A T C A T G C T A G A G T C A C T G T A C G A G T C C A T G T A C G | RAP26(AP2EREBP)/colamp-RAP26-DAP-Seq(GSE60143)/Homer | 1e-266 | -6.136e+02 | 0.0000 | 17691.0 | 44.68% | 63337.1 | 35.24% | motif file (matrix) | svg |
| 68 | C G T A T G A C T A G C T G C A A C T G A C T G C G T A C G T A T C A G G A C T | ELF3(ETS)/PDAC-ELF3-ChIP-Seq(GSE64557)/Homer | 1e-261 | -6.023e+02 | 0.0000 | 5047.0 | 12.75% | 12923.7 | 7.19% | motif file (matrix) | svg |
| 69 | A G T C C G T A T G A C A T G C G C A T C T G A G T A C G A T C | MYB55(MYB)/colamp-MYB55-DAP-Seq(GSE60143)/Homer | 1e-260 | -6.002e+02 | 0.0000 | 12540.0 | 31.67% | 41739.2 | 23.22% | motif file (matrix) | svg |
| 70 | G A C T G A T C A G T C C G T A T G A C A G T C G C A T C T G A G T A C G A T C G C A T G A C T | MYB10(MYB)/col-MYB10-DAP-Seq(GSE60143)/Homer | 1e-256 | -5.905e+02 | 0.0000 | 5719.0 | 14.44% | 15386.1 | 8.56% | motif file (matrix) | svg |
| 71 | A G T C G A T C A G C T C G T A G T A C A G T C G C A T C T G A G T A C G A T C | AT4G26030(C2H2)/col-AT4G26030-DAP-Seq(GSE60143)/Homer | 1e-256 | -5.897e+02 | 0.0000 | 11276.0 | 28.48% | 36719.2 | 20.43% | motif file (matrix) | svg |
| 72 | G A C T G C A T A C G T A C G T A C T G C G T A A G T C A G C T C G A T A T C G G C A T A C T G C G A T C T A G C G T A | WRKY50(WRKY)/col-WRKY50-DAP-Seq(GSE60143)/Homer | 1e-253 | -5.845e+02 | 0.0000 | 9645.0 | 24.36% | 30273.7 | 16.84% | motif file (matrix) | svg |
| 73 | T C A G A C T G A C G T C G T A A C T G A C T G A C G T C T A G | MYB51(MYB)/col-MYB51-DAP-Seq(GSE60143)/Homer | 1e-253 | -5.840e+02 | 0.0000 | 10651.0 | 26.90% | 34272.8 | 19.07% | motif file (matrix) | svg |
| 74 | G C T A C G T A C G T A G C A T C A T G C T A G A G T C A C T G A C T G A G T C A C T G T C A G | ABR1(AP2EREBP)/colamp-ABR1-DAP-Seq(GSE60143)/Homer | 1e-249 | -5.739e+02 | 0.0000 | 15174.0 | 38.32% | 53065.9 | 29.52% | motif file (matrix) | svg |
| 75 | C T A G A G T C A G T C A C T G C G T A A G T C C T G A G A C T | DDF1(AP2EREBP)/col-DDF1-DAP-Seq(GSE60143)/Homer | 1e-246 | -5.678e+02 | 0.0000 | 10912.0 | 27.56% | 35504.5 | 19.75% | motif file (matrix) | svg |
| 76 | C T A G A C T G A G T C A C T G A C T G A G C T A C T G T C A G | AT3G57600(AP2EREBP)/col-AT3G57600-DAP-Seq(GSE60143)/Homer | 1e-246 | -5.670e+02 | 0.0000 | 9614.0 | 24.28% | 30338.3 | 16.88% | motif file (matrix) | svg |
| 77 | C T A G T A G C A T G C C T A G A G T C A G T C C T A G G A C T G A C T G C T A | CRF10(AP2EREBP)/col100-CRF10-DAP-Seq(GSE60143)/Homer | 1e-246 | -5.667e+02 | 0.0000 | 18563.0 | 46.88% | 67800.1 | 37.72% | motif file (matrix) | svg |
| 78 | C G T A C G T A C T A G A C G T A C G T C G T A A C T G A C T G A C G T C T G A T C G A T C G A | MYB4(MYB)/col200-MYB4-DAP-Seq(GSE60143)/Homer | 1e-240 | -5.542e+02 | 0.0000 | 7282.0 | 18.39% | 21456.7 | 11.94% | motif file (matrix) | svg |
| 79 | A T C G A G T C A C T G A G T C A G T C A C T G G A C T G A C T | PUCHI(AP2EREBP)/colamp-PUCHI-DAP-Seq(GSE60143)/Homer | 1e-237 | -5.461e+02 | 0.0000 | 12198.0 | 30.81% | 40995.1 | 22.81% | motif file (matrix) | svg |
| 80 | C T A G C T A G A T G C G T A C T C A G A T G C A G T C G C A T G A T C G A T C | ZNF91(Zf)/HEK-ZNF91.HA-ChIP-Seq(GSE162571)/Homer | 1e-236 | -5.453e+02 | 0.0000 | 7260.0 | 18.34% | 21457.3 | 11.94% | motif file (matrix) | svg |
| 81 | G A C T G C T A T G C A A G T C A C G T A C G T A C G T C G A T A C G T T A C G | At3g45610(C2C2dof)/col-At3g45610-DAP-Seq(GSE60143)/Homer | 1e-229 | -5.289e+02 | 0.0000 | 10624.0 | 26.83% | 34799.9 | 19.36% | motif file (matrix) | svg |
| 82 | G A T C G T A C C T G A A G T C A G T C A C T G G C T A G T A C G T C A G C A T G C A T C G A T | DEAR2(AP2EREBP)/colamp-DEAR2-DAP-Seq(GSE60143)/Homer | 1e-228 | -5.253e+02 | 0.0000 | 17784.0 | 44.91% | 64987.2 | 36.16% | motif file (matrix) | svg |
| 83 | G C A T C T A G A C T G A C G T C G T A A C T G A C T G C G A T C T A G T C G A T C G A G C T A | MYB40(MYB)/col-MYB40-DAP-Seq(GSE60143)/Homer | 1e-227 | -5.230e+02 | 0.0000 | 4520.0 | 11.42% | 11655.0 | 6.48% | motif file (matrix) | svg |
| 84 | A G C T C T A G A G T C A G T C A C T G C G T A A G T C C T G A G C A T G C T A C T G A G C A T G C A T C G A T G C A T | CBF4(AP2EREBP)/colamp-CBF4-DAP-Seq(GSE60143)/Homer | 1e-226 | -5.223e+02 | 0.0000 | 14894.0 | 37.62% | 52577.1 | 29.25% | motif file (matrix) | svg |
| 85 | G C A T C G T A C T A G A G T C G T C A C G T A A T G C A C G T A C G T A C T G G A T C G C A T C G T A G C T A G C T A | bHLH122(bHLH)/col100-bHLH122-DAP-Seq(GSE60143)/Homer | 1e-223 | -5.149e+02 | 0.0000 | 8241.0 | 20.81% | 25521.8 | 14.20% | motif file (matrix) | svg |
| 86 | A T G C G T A C C T G A A G T C A G T C A C T G G T C A A G T C G T C A G C A T G C A T G A C T | At5g65130(AP2EREBP)/colamp-At5g65130-DAP-Seq(GSE60143)/Homer | 1e-222 | -5.115e+02 | 0.0000 | 4992.0 | 12.61% | 13404.1 | 7.46% | motif file (matrix) | svg |
| 87 | A G T C G A T C C T G A A G T C A G T C C A T G G T C A G A T C C G T A G A T C | DREB26(AP2EREBP)/col-DREB26-DAP-Seq(GSE60143)/Homer | 1e-220 | -5.081e+02 | 0.0000 | 4578.0 | 11.56% | 11966.5 | 6.66% | motif file (matrix) | svg |
| 88 | G T A C A C G T A C G T A T C G C A G T C G A T A T C G G C T A T G C A T A G C C G T A G T C A C A T G A G C T G C T A | ANAC013(NAC)/col-ANAC013-DAP-Seq(GSE60143)/Homer | 1e-219 | -5.046e+02 | 0.0000 | 4692.0 | 11.85% | 12393.4 | 6.90% | motif file (matrix) | svg |
| 89 | A G T C G T A C C T G A A G T C G T A C C T A G G C T A T G A C T G C A G C T A C G T A C G T A | At1g22810(AP2EREBP)/colamp-At1g22810-DAP-Seq(GSE60143)/Homer | 1e-219 | -5.044e+02 | 0.0000 | 9506.0 | 24.01% | 30609.5 | 17.03% | motif file (matrix) | svg |
| 90 | C G T A G A T C C T A G A C G T G T A C C T G A A G C T G A T C G C T A G A C T | TGA2(bZIP)/colamp-TGA2-DAP-Seq(GSE60143)/Homer | 1e-218 | -5.042e+02 | 0.0000 | 11204.0 | 28.30% | 37451.9 | 20.84% | motif file (matrix) | svg |
| 91 | G A C T A G T C C T G A A G T C A G T C A C T G C T G A A G T C G C T A G C A T G T A C C G A T G C A T G A C T C G A T | CBF2(AP2EREBP)/colamp-CBF2-DAP-Seq(GSE60143)/Homer | 1e-218 | -5.036e+02 | 0.0000 | 11272.0 | 28.47% | 37735.8 | 21.00% | motif file (matrix) | svg |
| 92 | G A T C A G T C G A C T G C T A G T A C A G T C G C A T G C T A G T A C G A T C | MYB61(MYB)/colamp-MYB61-DAP-Seq(GSE60143)/Homer | 1e-217 | -5.019e+02 | 0.0000 | 14608.0 | 36.89% | 51649.8 | 28.74% | motif file (matrix) | svg |
| 93 | C G T A G C A T C A T G C T A G A G T C A C T G A T C G G T A C A C T G T C A G | At2g33710(AP2EREBP)/colamp-At2g33710-DAP-Seq(GSE60143)/Homer | 1e-216 | -4.975e+02 | 0.0000 | 21079.0 | 53.24% | 80038.6 | 44.53% | motif file (matrix) | svg |
| 94 | G A C T A C T G C G A T A G T C A C T G C T A G A G T C C T G A | AT1G12630(AP2EREBP)/colamp-AT1G12630-DAP-Seq(GSE60143)/Homer | 1e-214 | -4.929e+02 | 0.0000 | 10748.0 | 27.14% | 35738.5 | 19.88% | motif file (matrix) | svg |
| 95 | C G A T C G T A G C T A G A C T T C G A A G C T A G T C A C T G T C G A A G C T C T G A C G A T | ZBTB38(Zf)/Hela-ZBTB38-ChIP-seq(GSE108618)/Homer | 1e-212 | -4.902e+02 | 0.0000 | 26415.0 | 66.71% | 104791.6 | 58.30% | motif file (matrix) | svg |
| 96 | A T G C A G T C G C A T A G C T A C G T T C A G C G A T A G C T G A T C A T C G | Sox10(HMG)/SciaticNerve-Sox3-ChIP-Seq(GSE35132)/Homer | 1e-212 | -4.901e+02 | 0.0000 | 12167.0 | 30.73% | 41585.6 | 23.14% | motif file (matrix) | svg |
| 97 | A C T G A C G T C G A T C A G T C A T G C A T G C A G T G C A T C A G T C A T G | HuR(?)/HEK293-HuR-CLIP-Seq(GSE87887)/Homer | 1e-212 | -4.898e+02 | 0.0000 | 19416.0 | 49.04% | 72718.3 | 40.46% | motif file (matrix) | svg |
| 98 | A G T C A G T C C G A T A C G T A C G T A C T G A C G T A G C T A G T C A G T C | Sox4(HMG)/proB-Sox4-ChIP-Seq(GSE50066)/Homer | 1e-211 | -4.863e+02 | 0.0000 | 6809.0 | 17.20% | 20319.9 | 11.31% | motif file (matrix) | svg |
| 99 | C G T A G A T C A G C T A C G T A C G T A C T G C G T A G T A C A G C T G C T A C G A T C G A T C G A T G C A T G C T A | WRKY18(WRKY)/col-WRKY18-DAP-Seq(GSE60143)/Homer | 1e-210 | -4.854e+02 | 0.0000 | 14626.0 | 36.94% | 51960.4 | 28.91% | motif file (matrix) | svg |
| 100 | A T G C G A T C C G A T A C G T A C G T A C T G C A G T A G C T | Sox3(HMG)/NPC-Sox3-ChIP-Seq(GSE33059)/Homer | 1e-208 | -4.793e+02 | 0.0000 | 12906.0 | 32.60% | 44801.6 | 24.93% | motif file (matrix) | svg |
| 101 | A G T C C A T G A C G T A C G T A C T G C G T A A G T C G A C T G C A T G C T A | WRKY28(WRKY)/col-WRKY28-DAP-Seq(GSE60143)/Homer | 1e-207 | -4.784e+02 | 0.0000 | 11310.0 | 28.56% | 38213.8 | 21.26% | motif file (matrix) | svg |
| 102 | C T A G A C T G A C G T C G T A A C T G C A T G G C A T T C A G | MYB92(MYB)/colamp-MYB92-DAP-Seq(GSE60143)/Homer | 1e-207 | -4.780e+02 | 0.0000 | 10823.0 | 27.33% | 36230.8 | 20.16% | motif file (matrix) | svg |
| 103 | A G C T A G C T C A T G C T G A G T A C A G T C A G C T A G C T C A G T C T A G | RARa(NR)/K562-RARa-ChIP-Seq(Encode)/Homer | 1e-207 | -4.774e+02 | 0.0000 | 17614.0 | 44.49% | 64967.1 | 36.15% | motif file (matrix) | svg |
| 104 | G A C T A C T G C G A T A G T C A C T G C T A G A G T C C G T A | Rap210(AP2EREBP)/col-Rap210-DAP-Seq(GSE60143)/Homer | 1e-206 | -4.756e+02 | 0.0000 | 13155.0 | 33.22% | 45893.7 | 25.53% | motif file (matrix) | svg |
| 105 | T A C G G A C T T G A C C G T A A C G T G A T C G T C A C G T A A C G T A T G C C G T A G A C T | HOXA2(Homeobox)/mES-Hoxa2-ChIP-Seq(Donaldson\_et\_al.)/Homer | 1e-206 | -4.754e+02 | 0.0000 | 1710.0 | 4.32% | 2920.3 | 1.62% | motif file (matrix) | svg |
| 106 | A G C T T G A C C G A T C G A T C T A G A C G T C A G T C A G T G C T A A G T C | FOXK1(Forkhead)/HEK293-FOXK1-ChIP-Seq(GSE51673)/Homer | 1e-203 | -4.693e+02 | 0.0000 | 8054.0 | 20.34% | 25285.4 | 14.07% | motif file (matrix) | svg |
| 107 | G C A T C G A T G C A T C G T A C T A G A G T C G T C A C G T A A T C G A C G T A C G T A C T G G T A C G C A T C G A T | bHLH80(bHLH)/col-bHLH80-DAP-Seq(GSE60143)/Homer | 1e-203 | -4.689e+02 | 0.0000 | 8598.0 | 21.71% | 27422.7 | 15.26% | motif file (matrix) | svg |
| 108 | G A T C A G C T C T A G G A T C T G A C C T A G C G T A G T A C C G T A G C A T G T C A C T G A | CBF3(AP2EREBP)/colamp-CBF3-DAP-Seq(GSE60143)/Homer | 1e-201 | -4.628e+02 | 0.0000 | 11049.0 | 27.91% | 37347.4 | 20.78% | motif file (matrix) | svg |
| 109 | C A T G A G T C G T C A C G T A A T G C A C G T A C G T A C T G | bHLH130(bHLH)/col-bHLH130-DAP-Seq(GSE60143)/Homer | 1e-200 | -4.619e+02 | 0.0000 | 7238.0 | 18.28% | 22203.7 | 12.35% | motif file (matrix) | svg |
| 110 | C G T A C G A T C G T A C G A T C A T G A C T G C G A T A G T C A T C G T C A G G A C T A C T G | At1g36060(AP2EREBP)/colamp-At1g36060-DAP-Seq(GSE60143)/Homer | 1e-200 | -4.611e+02 | 0.0000 | 12394.0 | 31.30% | 42919.7 | 23.88% | motif file (matrix) | svg |
| 111 | C A G T A C T G T C A G T G C A G C T A A T G C T C G A A T C G G T C A T G C A | ZNF189(Zf)/HEK293-ZNF189.GFP-ChIP-Seq(GSE58341)/Homer | 1e-197 | -4.553e+02 | 0.0000 | 6416.0 | 16.20% | 19137.5 | 10.65% | motif file (matrix) | svg |
| 112 | G A T C G A C T G A C T A C G T A G T C A C G T A G T C A C G T A G T C A C G T A G T C A C G T G T A C C G A T G T C A | BPC6(BBRBPC)/col-BPC6-DAP-Seq(GSE60143)/Homer | 1e-197 | -4.542e+02 | 0.0000 | 451.0 | 1.14% | 163.7 | 0.09% | motif file (matrix) | svg |
| 113 | T C A G T C A G G C T A C G T A T A C G G A C T T C A G T C G A C T G A C G T A T A C G G A C T | IRF8(IRF)/BMDM-IRF8-ChIP-Seq(GSE77884)/Homer | 1e-197 | -4.538e+02 | 0.0000 | 2596.0 | 6.56% | 5663.7 | 3.15% | motif file (matrix) | svg |
| 114 | C G T A C G A T C A G T C A T G C G A T G T A C C A T G A C T G G A C T C A T G | CEJ1(AP2EREBP)/col-CEJ1-DAP-Seq(GSE60143)/Homer | 1e-196 | -4.534e+02 | 0.0000 | 17024.0 | 43.00% | 62765.0 | 34.92% | motif file (matrix) | svg |
| 115 | C T A G T C A G C T G A T C A G T G C A A C T G T C G A T C A G | Trl(Zf)/S2-GAGAfactor-ChIP-Seq(GSE40646)/Homer | 1e-196 | -4.524e+02 | 0.0000 | 17706.0 | 44.72% | 65762.0 | 36.59% | motif file (matrix) | svg |
| 116 | G A C T A C T G C A G T A G T C A C T G A C T G A G C T A C T G C T A G G T C A | At1g77640(AP2EREBP)/col-At1g77640-DAP-Seq(GSE60143)/Homer | 1e-195 | -4.510e+02 | 0.0000 | 3928.0 | 9.92% | 10118.1 | 5.63% | motif file (matrix) | svg |
| 117 | C G A T C G T A G T A C A C G T A C G T T C A G G C A T C A G T T A C G G T C A C G T A A G T C C G T A T G C A C A T G | NAC2(NAC)/colamp-NAC2-DAP-Seq(GSE60143)/Homer | 1e-193 | -4.464e+02 | 0.0000 | 6395.0 | 16.15% | 19146.1 | 10.65% | motif file (matrix) | svg |
| 118 | C A T G C T A G A G T C A C T G A C T G G T A C C A T G T A C G | AT1G28160(AP2EREBP)/colamp-AT1G28160-DAP-Seq(GSE60143)/Homer | 1e-193 | -4.453e+02 | 0.0000 | 19747.0 | 49.87% | 74907.3 | 41.68% | motif file (matrix) | svg |
| 119 | A G T C G A T C G C T A C G A T C A G T T A C G C G A T A G C T A G T C A T C G | SOX1(HMG)/NPC-SOX1-ChIP-Seq(GSE138215)/Homer | 1e-191 | -4.400e+02 | 0.0000 | 14618.0 | 36.92% | 52596.0 | 29.26% | motif file (matrix) | svg |
| 120 | C G A T C T A G C G T A G A C T C A G T C T A G C G T A A G C T C A T G C T A G | HOXA1(Homeobox)/mES-Hoxa1-ChIP-Seq(SRP084292)/Homer | 1e-189 | -4.372e+02 | 0.0000 | 3303.0 | 8.34% | 8077.0 | 4.49% | motif file (matrix) | svg |
| 121 | G T A C C T G A T A G C C G T A G C T A T C G A T G C A T G A C C T A G G T C A A G T C C G T A C T G A C T G A C G T A | At1g14580(C2H2)/colamp-At1g14580-DAP-Seq(GSE60143)/Homer | 1e-186 | -4.289e+02 | 0.0000 | 1625.0 | 4.10% | 2861.3 | 1.59% | motif file (matrix) | svg |
| 122 | G A T C C T G A A G T C A G T C A C T G C G T A A G T C C T G A | ERF38(AP2EREBP)/col-ERF38-DAP-Seq(GSE60143)/Homer | 1e-186 | -4.288e+02 | 0.0000 | 10494.0 | 26.50% | 35528.1 | 19.77% | motif file (matrix) | svg |
| 123 | C G A T C G A T G C A T G A C T A C G T C G T A C G T A A C T G T A G C C G T A C G T A C G T A | AT5G60130(ABI3VP1)/col-AT5G60130-DAP-Seq(GSE60143)/Homer | 1e-186 | -4.284e+02 | 0.0000 | 8516.0 | 21.51% | 27572.4 | 15.34% | motif file (matrix) | svg |
| 124 | C G A T T G C A A G T C A C G T A C G T T A C G C G A T C G A T T A C G G C T A G C T A A T G C C G T A G T C A C A T G | ANAC016(NAC)/col-ANAC016-DAP-Seq(GSE60143)/Homer | 1e-185 | -4.283e+02 | 0.0000 | 8839.0 | 22.32% | 28858.1 | 16.06% | motif file (matrix) | svg |
| 125 | G T C A G C T A G C T A T C G A A T C G A C G T A G T C T C G A T C G A T G A C | WRKY40(WRKY)/colamp-WRKY40-DAP-Seq(GSE60143)/Homer | 1e-185 | -4.261e+02 | 0.0000 | 6131.0 | 15.48% | 18350.8 | 10.21% | motif file (matrix) | svg |
| 126 | G A C T A G T C G A T C C G T A G T A C A G T C G C A T C G T A G T C A G A T C | MYB67(MYB)/col-MYB67-DAP-Seq(GSE60143)/Homer | 1e-183 | -4.229e+02 | 0.0000 | 10465.0 | 26.43% | 35489.7 | 19.75% | motif file (matrix) | svg |
| 127 | G A T C A T G C A G T C C G T A A G T C A G T C A C T G G C T A A G T C C G T A | AT1G44830(AP2EREBP)/col-AT1G44830-DAP-Seq(GSE60143)/Homer | 1e-183 | -4.224e+02 | 0.0000 | 5722.0 | 14.45% | 16849.3 | 9.37% | motif file (matrix) | svg |
| 128 | A G T C T G C A T C G A C T G A A C T G C A T G A C G T A T G C G T C A T A C G | Erra(NR)/HepG2-Erra-ChIP-Seq(GSE31477)/Homer | 1e-183 | -4.217e+02 | 0.0000 | 12652.0 | 31.95% | 44556.7 | 24.79% | motif file (matrix) | svg |
| 129 | T A C G T A G C G C T A C G A T C T A G A C G T C A G T C A G T G C T A A G T C G T C A G C A T | FOXK2(Forkhead)/U2OS-FOXK2-ChIP-Seq(E-MTAB-2204)/Homer | 1e-181 | -4.171e+02 | 0.0000 | 5368.0 | 13.56% | 15580.4 | 8.67% | motif file (matrix) | svg |
| 130 | C T A G C T G A C G T A C G T A C G T A C G T A A C T G A C G T C T A G G T C A | COG1(C2C2dof)/col-COG1-DAP-Seq(GSE60143)/Homer | 1e-180 | -4.152e+02 | 0.0000 | 9955.0 | 25.14% | 33515.8 | 18.65% | motif file (matrix) | svg |
| 131 | A G C T C T A G G A T C A G T C C T A G C T G A A G T C G C T A G C A T T G C A | CBF1(AP2EREBP)/colamp-CBF1-DAP-Seq(GSE60143)/Homer | 1e-179 | -4.137e+02 | 0.0000 | 13731.0 | 34.68% | 49225.8 | 27.39% | motif file (matrix) | svg |
| 132 | A G T C T G A C C T G A A G T C A G T C A C T G C G T A A G T C G T C A G C T A G C A T C G T A G C A T G C T A C G T A | DEAR3(AP2EREBP)/colamp-DEAR3-DAP-Seq(GSE60143)/Homer | 1e-179 | -4.136e+02 | 0.0000 | 8146.0 | 20.57% | 26285.3 | 14.62% | motif file (matrix) | svg |
| 133 | C G A T C T A G A C G T A C G T A C G T C G T A A G C T C G A T A G C T C G T A C T A G T A G C | FoxD3(forkhead)/ZebrafishEmbryo-Foxd3.biotin-ChIP-seq(GSE106676)/Homer | 1e-179 | -4.134e+02 | 0.0000 | 4773.0 | 12.05% | 13427.4 | 7.47% | motif file (matrix) | svg |
| 134 | T A C G T C G A C G T A C G T A C G T A C T G A A C T G A C G T C G T A T C G A | AT2G28810(C2C2dof)/colamp-AT2G28810-DAP-Seq(GSE60143)/Homer | 1e-179 | -4.132e+02 | 0.0000 | 15267.0 | 38.56% | 55794.2 | 31.04% | motif file (matrix) | svg |
| 135 | C A T G A T G C T A G C C T G A A G T C A G T C A C T G G C T A A G T C G T A C G C T A G C A T | At4g28140(AP2EREBP)/colamp-At4g28140-DAP-Seq(GSE60143)/Homer | 1e-178 | -4.114e+02 | 0.0000 | 6912.0 | 17.46% | 21488.6 | 11.96% | motif file (matrix) | svg |
| 136 | C G T A A C G T A C G T A C G T A C G T A G T C A G T C C T G A A G C T A G C T | NFAT(RHD)/Jurkat-NFATC1-ChIP-Seq(Jolma\_et\_al.)/Homer | 1e-176 | -4.053e+02 | 0.0000 | 6389.0 | 16.14% | 19545.8 | 10.87% | motif file (matrix) | svg |
| 137 | G A C T G T A C C T G A G A T C A G T C C T A G G C T A G T A C C T G A G C T A G C A T C G A T G C A T G A C T C G T A | AT3G16280(AP2EREBP)/colamp-AT3G16280-DAP-Seq(GSE60143)/Homer | 1e-175 | -4.052e+02 | 0.0000 | 9542.0 | 24.10% | 31973.8 | 17.79% | motif file (matrix) | svg |
| 138 | C T G A T C G A G T A C A C G T A C G T A T C G A C G T C G A T A T C G G C T A G T A C A T G C C G T A T G C A C A T G | ANAC103(NAC)/col-ANAC103-DAP-Seq(GSE60143)/Homer | 1e-174 | -4.009e+02 | 0.0000 | 4018.0 | 10.15% | 10816.4 | 6.02% | motif file (matrix) | svg |
| 139 | C T G A A G T C C G A T A G C T A T G C G T A C A C G T A T C G C A G T G C A T | Elf4(ETS)/BMDM-Elf4-ChIP-Seq(GSE88699)/Homer | 1e-172 | -3.968e+02 | 0.0000 | 8141.0 | 20.56% | 26469.3 | 14.73% | motif file (matrix) | svg |
| 140 | C G A T C G T A G T A C A C G T A C G T T C A G G C A T C A G T T A C G G T C A C G T A A G T C C G T A T G C A C A T G | ANAC053(NAC)/colamp-ANAC053-DAP-Seq(GSE60143)/Homer | 1e-171 | -3.950e+02 | 0.0000 | 5573.0 | 14.08% | 16558.9 | 9.21% | motif file (matrix) | svg |
| 141 | G A T C C T G A A G T C G T A C A C T G G C T A G A T C C T G A G C T A G C T A | At4g31060(AP2EREBP)/colamp-At4g31060-DAP-Seq(GSE60143)/Homer | 1e-170 | -3.933e+02 | 0.0000 | 9317.0 | 23.53% | 31221.9 | 17.37% | motif file (matrix) | svg |
| 142 | G A C T G C A T C T A G C G A T G A T C T C G A C A T G G A T C | Tgif1(Homeobox)/mES-Tgif1-ChIP-Seq(GSE55404)/Homer | 1e-169 | -3.901e+02 | 0.0000 | 19232.0 | 48.57% | 73549.8 | 40.92% | motif file (matrix) | svg |
| 143 | C G A T C T G A A G T C A C G T A C G T A T C G G A C T C T A G C G A T G A C T C G T A A T G C C G T A G T C A A C T G | ANAC011(NAC)/col-ANAC011-DAP-Seq(GSE60143)/Homer | 1e-168 | -3.876e+02 | 0.0000 | 2993.0 | 7.56% | 7362.8 | 4.10% | motif file (matrix) | svg |
| 144 | G A T C C T G A A G T C A G C T A C G T A C G T A C G T A C G T | At1g64620(C2C2dof)/colamp-At1g64620-DAP-Seq(GSE60143)/Homer | 1e-166 | -3.837e+02 | 0.0000 | 10564.0 | 26.68% | 36436.5 | 20.27% | motif file (matrix) | svg |
| 145 | C T A G A C T G A G C T C G T A A C T G A C T G A C G T C T A G | MYB99(MYB)/colamp-MYB99-DAP-Seq(GSE60143)/Homer | 1e-161 | -3.725e+02 | 0.0000 | 10696.0 | 27.01% | 37142.2 | 20.66% | motif file (matrix) | svg |
| 146 | T C G A A C T G C A T G A G C T A G T C C G T A C T G A C T A G A C T G C G A T A T G C C T G A | RAR:RXR(NR),DR0/ES-RAR-ChIP-Seq(GSE56893)/Homer | 1e-161 | -3.716e+02 | 0.0000 | 1373.0 | 3.47% | 2378.6 | 1.32% | motif file (matrix) | svg |
| 147 | C G T A G C A T C A G T C T A G A G T C A C T G A C T G G T A C A C T G A T C G | ERF115(AP2EREBP)/colamp-ERF115-DAP-Seq(GSE60143)/Homer | 1e-161 | -3.713e+02 | 0.0000 | 19636.0 | 49.59% | 75687.4 | 42.11% | motif file (matrix) | svg |
| 148 | C G A T C T G A A G T C A C G T A C G T T A C G G C T A C A T G C T A G G C A T C G A T A G T C C G T A G T C A A C T G | ANAC096(NAC)/colamp-ANAC096-DAP-Seq(GSE60143)/Homer | 1e-160 | -3.698e+02 | 0.0000 | 5875.0 | 14.84% | 17960.2 | 9.99% | motif file (matrix) | svg |
| 149 | A G C T A C G T A C T G A T G C A G T C C G T A C T G A T A C G | NF1-halfsite(CTF)/LNCaP-NF1-ChIP-Seq(Unpublished)/Homer | 1e-159 | -3.671e+02 | 0.0000 | 11951.0 | 30.18% | 42441.5 | 23.61% | motif file (matrix) | svg |
| 150 | A C G T A G T C A G T C C G A T A C G T A C G T A C T G A C G T A T G C G A C T A C T G T A C G | Sox21(HMG)/ESC-SOX21-ChIP-Seq(GSE110505)/Homer | 1e-158 | -3.660e+02 | 0.0000 | 12487.0 | 31.54% | 44708.2 | 24.87% | motif file (matrix) | svg |
| 151 | C T A G G C T A A G T C A C T G A C G T G A C T G A C T A T G C T C G A C A G T G A T C C G A T G A C T G A T C G A T C | RKD2(RWPRK)/colamp-RKD2-DAP-Seq(GSE60143)/Homer | 1e-156 | -3.613e+02 | 0.0000 | 6981.0 | 17.63% | 22329.0 | 12.42% | motif file (matrix) | svg |
| 152 | G C T A C G T A C G A T G A C T G C A T T G C A A G T C A G C T A C G T A C G T C G A T G A C T | DAG2(C2C2dof)/col-DAG2-DAP-Seq(GSE60143)/Homer | 1e-156 | -3.612e+02 | 0.0000 | 9701.0 | 24.50% | 33220.1 | 18.48% | motif file (matrix) | svg |
| 153 | T A C G C G T A G A C T T C A G A G C T A G T C A C T G T C A G A G T C C T G A | DDF2(AP2EREBP)/col-DDF2-DAP-Seq(GSE60143)/Homer | 1e-155 | -3.592e+02 | 0.0000 | 2021.0 | 5.10% | 4362.5 | 2.43% | motif file (matrix) | svg |
| 154 | G A C T G A T C C T G A A G T C A G T C A C T G C G T A A G T C G T A C G C T A G C A T C G A T | At1g19210(AP2EREBP)/colamp-At1g19210-DAP-Seq(GSE60143)/Homer | 1e-155 | -3.587e+02 | 0.0000 | 20232.0 | 51.10% | 78591.0 | 43.73% | motif file (matrix) | svg |
| 155 | G A C T C A G T G C A T C G A T T G A C A C G T A T G C G T A C C T G A A C T G A C T G A G C T | WIP5(C2H2)/colamp-WIP5-DAP-Seq(GSE60143)/Homer | 1e-154 | -3.547e+02 | 0.0000 | 8946.0 | 22.59% | 30241.8 | 16.83% | motif file (matrix) | svg |
| 156 | C T A G A C T G A C G T C G T A A C T G A C T G A C G T T C A G C T G A T C G A | MYB107(MYB)/col-MYB107-DAP-Seq(GSE60143)/Homer | 1e-153 | -3.537e+02 | 0.0000 | 15099.0 | 38.13% | 56056.4 | 31.19% | motif file (matrix) | svg |
| 157 | G C T A A G T C T A C G T G C A A T C G T C A G G C T A T C G A T C A G A G C T | ELF5(ETS)/T47D-ELF5-ChIP-Seq(GSE30407)/Homer | 1e-153 | -3.528e+02 | 0.0000 | 5937.0 | 14.99% | 18381.7 | 10.23% | motif file (matrix) | svg |
| 158 | C G A T T C G A G A T C A C G T A C G T T C A G G C T A C G A T C G T A C G T A C G T A A T G C C G T A T G C A C T A G | ANAC028(NAC)/col-ANAC028-DAP-Seq(GSE60143)/Homer | 1e-153 | -3.526e+02 | 0.0000 | 6588.0 | 16.64% | 20900.0 | 11.63% | motif file (matrix) | svg |
| 159 | C G A T T C G A G A T C A C G T A C G T T C A G G C A T C T G A C G T A G C T A C G T A A G T C C G T A T G C A C A T G | ANAC050(NAC)/colamp-ANAC050-DAP-Seq(GSE60143)/Homer | 1e-152 | -3.523e+02 | 0.0000 | 5891.0 | 14.88% | 18211.2 | 10.13% | motif file (matrix) | svg |
| 160 | G T A C G T C A G T A C G T C A G T A C G T C A G T A C G T C A G T A C G T C A | SeqBias: CA-repeat | 1e-152 | -3.500e+02 | 0.0000 | 26793.0 | 67.67% | 108987.8 | 60.64% | motif file (matrix) | svg |
| 161 | A T G C C A G T A G C T A G C T T C A G G T C A T A G C G A C T C G T A C G A T | WRKY20(WRKY)/col-WRKY20-DAP-Seq(GSE60143)/Homer | 1e-151 | -3.485e+02 | 0.0000 | 7377.0 | 18.63% | 24043.4 | 13.38% | motif file (matrix) | svg |
| 162 | A G T C A C G T A C T G A G C T A C G T A C G T G T C A A G T C | Foxo1(Forkhead)/RAW-Foxo1-ChIP-Seq(Fan\_et\_al.)/Homer | 1e-151 | -3.483e+02 | 0.0000 | 11596.0 | 29.29% | 41247.1 | 22.95% | motif file (matrix) | svg |
| 163 | G C T A C G T A C G A T C A G T A C T G C G A T G T A C A C T G A T C G G A C T C A T G C T A G G C A T C A G T C A T G | DEAR5(AP2EREBP)/col-DEAR5-DAP-Seq(GSE60143)/Homer | 1e-149 | -3.447e+02 | 0.0000 | 4356.0 | 11.00% | 12522.6 | 6.97% | motif file (matrix) | svg |
| 164 | C T G A A T G C G C T A G C A T A T G C C G T A T C G A C T G A C T A G T C A G T A C G G T C A | Tcf4(HMG)/Hct116-Tcf4-ChIP-Seq(SRA012054)/Homer | 1e-149 | -3.439e+02 | 0.0000 | 3715.0 | 9.38% | 10205.9 | 5.68% | motif file (matrix) | svg |
| 165 | T C G A T A G C G T C A A C T G A C T G C G T A C G T A C T A G A G C T T C A G | ERG(ETS)/VCaP-ERG-ChIP-Seq(GSE14097)/Homer | 1e-147 | -3.394e+02 | 0.0000 | 9188.0 | 23.20% | 31438.0 | 17.49% | motif file (matrix) | svg |
| 166 | A T C G T G C A G A T C C T A G A C G T A T C G C G T A A G T C T C A G A C T G T C A G G C T A | Knotted(Homeobox)/Corn-KN1-ChIP-Seq(GSE39161)/Homer | 1e-146 | -3.382e+02 | 0.0000 | 16537.0 | 41.77% | 62583.2 | 34.82% | motif file (matrix) | svg |
| 167 | C A G T C G T A C G T A G C A T G A C T G C A T G T A C A G C T A C T G G A C T A C G T C A T G | RAV1(RAV)/colamp-RAV1-DAP-Seq(GSE60143)/Homer | 1e-146 | -3.376e+02 | 0.0000 | 4685.0 | 11.83% | 13807.2 | 7.68% | motif file (matrix) | svg |
| 168 | C A T G C T G A A G T C A C T G A C T G A G C T A C T G A T C G | ESE3(AP2EREBP)/col-ESE3-DAP-Seq(GSE60143)/Homer | 1e-145 | -3.356e+02 | 0.0000 | 16959.0 | 42.83% | 64480.6 | 35.88% | motif file (matrix) | svg |
| 169 | C G T A C G T A C T A G C A G T G A C T C G T A C A T G C A T G C G A T C T G A C T G A C T G A | MS188(MYB)/colamp-MS188-DAP-Seq(GSE60143)/Homer | 1e-145 | -3.356e+02 | 0.0000 | 7320.0 | 18.49% | 23982.3 | 13.34% | motif file (matrix) | svg |
| 170 | G A T C C T G A G A T C G A T C C T A G G C T A A G T C C T G A G C T A C G T A | At4g16750(AP2EREBP)/col-At4g16750-DAP-Seq(GSE60143)/Homer | 1e-145 | -3.354e+02 | 0.0000 | 14955.0 | 37.77% | 55754.9 | 31.02% | motif file (matrix) | svg |
| 171 | G T A C G T A C G T C A G C T A C G T A C G T A C G T A C T A G C T A G C T A G | SEP3(MADS)/Arabidoposis-Flower-Sep3-ChIP-Seq/Homer | 1e-145 | -3.350e+02 | 0.0000 | 8411.0 | 21.24% | 28351.7 | 15.77% | motif file (matrix) | svg |
| 172 | C G T A C G T A C G A T A C T G C A G T A G T C A C T G A C T G A G C T A C T G | DREB19(AP2EREBP)/colamp-DREB19-DAP-Seq(GSE60143)/Homer | 1e-145 | -3.344e+02 | 0.0000 | 10474.0 | 26.45% | 36793.4 | 20.47% | motif file (matrix) | svg |
| 173 | A T G C G C A T T A G C C G A T T A G C G C A T T A G C G C A T A T G C G A C T | GAGA-repeat/Arabidopsis-Promoters/Homer | 1e-144 | -3.338e+02 | 0.0000 | 6333.0 | 15.99% | 20128.3 | 11.20% | motif file (matrix) | svg |
| 174 | T C G A T A G C T G C A A C T G A C T G C G T A C G T A C T A G G A C T T A C G | ETS1(ETS)/Jurkat-ETS1-ChIP-Seq(GSE17954)/Homer | 1e-142 | -3.286e+02 | 0.0000 | 8031.0 | 20.28% | 26909.2 | 14.97% | motif file (matrix) | svg |
| 175 | C G A T C G A T G C A T G C A T G T C A A G T C A G C T A C G T A C G T C G A T G A C T A C G T | OBP4(C2C2dof)/col-OBP4-DAP-Seq(GSE60143)/Homer | 1e-142 | -3.285e+02 | 0.0000 | 9229.0 | 23.31% | 31762.6 | 17.67% | motif file (matrix) | svg |
| 176 | T C G A A G T C C G T A A T C G A T G C C G A T A C T G A G T C A G C T A C T G | Tcf12(bHLH)/GM12878-Tcf12-ChIP-Seq(GSE32465)/Homer | 1e-141 | -3.254e+02 | 0.0000 | 5000.0 | 12.63% | 15111.9 | 8.41% | motif file (matrix) | svg |
| 177 | A G T C G A T C A G T C C G T A A T C G C A G T A G T C G T A C C T G A A C T G T C A G A G C T A G C T A G C T A G C T | PRDM15(Zf)/ESC-Prdm15-ChIP-Seq(GSE73694)/Homer | 1e-141 | -3.251e+02 | 0.0000 | 7931.0 | 20.03% | 26556.4 | 14.78% | motif file (matrix) | svg |
| 178 | G C A T T G A C C A T G G A C T C A G T C A T G T C G A G T A C G A C T G C T A C G A T C G A T | WRKY6(WRKY)/colamp-WRKY6-DAP-Seq(GSE60143)/Homer | 1e-140 | -3.245e+02 | 0.0000 | 8139.0 | 20.56% | 27400.4 | 15.24% | motif file (matrix) | svg |
| 179 | A G C T A G C T A G C T A C T G A C G T A G T C A C T G A C G T G A C T C G A T G C A T A T C G | IDD7(C2H2)/col-IDD7-DAP-Seq(GSE60143)/Homer | 1e-140 | -3.243e+02 | 0.0000 | 3634.0 | 9.18% | 10081.8 | 5.61% | motif file (matrix) | svg |
| 180 | A T G C G T A C C G T A A G C T G C A T T A C G A G C T A G C T A G T C A G C T | Sox6(HMG)/Myotubes-Sox6-ChIP-Seq(GSE32627)/Homer | 1e-139 | -3.201e+02 | 0.0000 | 12335.0 | 31.15% | 44816.1 | 24.93% | motif file (matrix) | svg |
| 181 | C T A G C T A G C G T A C G T A T A C G C G A T C T A G C T G A C T G A C G T A T A C G G A C T | PU.1:IRF8(ETS:IRF)/pDC-Irf8-ChIP-Seq(GSE66899)/Homer | 1e-138 | -3.191e+02 | 0.0000 | 1390.0 | 3.51% | 2642.5 | 1.47% | motif file (matrix) | svg |
| 182 | A G T C A G T C C T G A A G T C A G T C A C T G C G T A A G T C C T G A T C G A G C A T G A T C C G A T C G A T A C T G | AT3G60490(AP2EREBP)/colamp-AT3G60490-DAP-Seq(GSE60143)/Homer | 1e-137 | -3.169e+02 | 0.0000 | 7326.0 | 18.50% | 24249.6 | 13.49% | motif file (matrix) | svg |
| 183 | G T A C C A T G A G C T A G C T T C A G T G C A T G A C A G C T C G T A C G T A | WRKY33(WRKY)/col-WRKY33-DAP-Seq(GSE60143)/Homer | 1e-137 | -3.157e+02 | 0.0000 | 8399.0 | 21.21% | 28573.9 | 15.90% | motif file (matrix) | svg |
| 184 | T C G A G T A C C A T G A G C T A C G T C A T G G T C A G T A C A G C T G C T A C G A T C A G T | WRKY31(WRKY)/colamp-WRKY31-DAP-Seq(GSE60143)/Homer | 1e-135 | -3.131e+02 | 0.0000 | 6795.0 | 17.16% | 22194.6 | 12.35% | motif file (matrix) | svg |
| 185 | A G C T G A T C C T G A A G T C A G T C A C T G C G T A A G T C C T G A G T C A G C A T C G A T G C T A G C A T C G T A | At2g44940(AP2EREBP)/colamp-At2g44940-DAP-Seq(GSE60143)/Homer | 1e-135 | -3.129e+02 | 0.0000 | 5902.0 | 14.91% | 18703.1 | 10.41% | motif file (matrix) | svg |
| 186 | C G T A G A T C C A T G G C A T G A C T C T A G T C G A T A G C A G C T G C A T | WRKY55(WRKY)/col-WRKY55-DAP-Seq(GSE60143)/Homer | 1e-135 | -3.127e+02 | 0.0000 | 10152.0 | 25.64% | 35797.8 | 19.92% | motif file (matrix) | svg |
| 187 | C T G A A T C G G T A C C T G A A G T C A G T C A C T G C G T A A G T C C T G A | TINY(AP2EREBP)/col-TINY-DAP-Seq(GSE60143)/Homer | 1e-134 | -3.106e+02 | 0.0000 | 6472.0 | 16.35% | 20954.4 | 11.66% | motif file (matrix) | svg |
| 188 | A G T C A C G T A C G T T A C G G C A T G C A T A T G C G C T A C G T A A T G C C G T A G T C A A C T G G A T C G C A T | ANAC075(NAC)/col-ANAC075-DAP-Seq(GSE60143)/Homer | 1e-133 | -3.074e+02 | 0.0000 | 4021.0 | 10.16% | 11642.8 | 6.48% | motif file (matrix) | svg |
| 189 | C T G A T G A C T G A C C G T A A C G T T G A C A G C T C T A G A C G T G A C T | Olig2(bHLH)/Neuron-Olig2-ChIP-Seq(GSE30882)/Homer | 1e-133 | -3.065e+02 | 0.0000 | 12251.0 | 30.94% | 44693.8 | 24.87% | motif file (matrix) | svg |
| 190 | G C A T T G A C C T G A A G T C A G T C A C T G G T C A A G T C G C T A G A C T G C T A C T G A | DREB2(AP2EREBP)/col-DREB2-DAP-Seq(GSE60143)/Homer | 1e-132 | -3.055e+02 | 0.0000 | 8979.0 | 22.68% | 31083.0 | 17.29% | motif file (matrix) | svg |
| 191 | A G C T G T C A T G C A A G T C A C G T A C G T A C G T C G A T G A C T T A C G | AT3G12130(C3H)/colamp-AT3G12130-DAP-Seq(GSE60143)/Homer | 1e-132 | -3.051e+02 | 0.0000 | 15744.0 | 39.76% | 59734.1 | 33.23% | motif file (matrix) | svg |
| 192 | A T G C C A T G G C A T G A C T C T A G T C G A G T A C A G C T C G T A G C T A | WRKY75(WRKY)/col-WRKY75-DAP-Seq(GSE60143)/Homer | 1e-132 | -3.040e+02 | 0.0000 | 8752.0 | 22.10% | 30177.4 | 16.79% | motif file (matrix) | svg |
| 193 | C G A T T C G A G A T C C G A T G C A T T C A G G A C T C G A T G C A T G C T A C T G A A G T C C G T A G T C A C T A G | ANAC005(NAC)/col-ANAC005-DAP-Seq(GSE60143)/Homer | 1e-131 | -3.038e+02 | 0.0000 | 3408.0 | 8.61% | 9447.2 | 5.26% | motif file (matrix) | svg |
| 194 | A T G C C A T G A G C T C A G T C A T G T C G A A G T C G A C T C G A T C G A T C A G T C A G T | WRKY26(WRKY)/colamp-WRKY26-DAP-Seq(GSE60143)/Homer | 1e-131 | -3.033e+02 | 0.0000 | 5465.0 | 13.80% | 17128.2 | 9.53% | motif file (matrix) | svg |
| 195 | C T G A T C G A C G T A A T G C C G T A C G T A C G A T C T A G T C A G G A T C | Sox15(HMG)/CPA-Sox15-ChIP-Seq(GSE62909)/Homer | 1e-131 | -3.020e+02 | 0.0000 | 7473.0 | 18.87% | 25037.4 | 13.93% | motif file (matrix) | svg |
| 196 | C T A G G T A C C A T G G A C T C G A T C A T G G T C A G T A C G A C T C G A T C G A T C G A T | WRKY27(WRKY)/colamp-WRKY27-DAP-Seq(GSE60143)/Homer | 1e-130 | -3.016e+02 | 0.0000 | 7729.0 | 19.52% | 26069.9 | 14.50% | motif file (matrix) | svg |
| 197 | G A T C G A T C A G T C G T A C C G A T G T A C G T A C A G T C A G T C A G T C G C T A G A T C | ZNF148(Zf)/MDAMB231-ZNF148-ChIP-Seq(GSE147020)/Homer | 1e-130 | -3.014e+02 | 0.0000 | 3415.0 | 8.62% | 9493.5 | 5.28% | motif file (matrix) | svg |
| 198 | C A T G T G C A G A C T C A T G C G T A A G T C T C A G G C A T T G A C C G T A | bZIP50(bZIP)/colamp-bZIP50-DAP-Seq(GSE60143)/Homer | 1e-130 | -3.010e+02 | 0.0000 | 13736.0 | 34.69% | 51122.3 | 28.44% | motif file (matrix) | svg |
| 199 | G A C T C T G A A G T C A G T C A C T G C G T A A G T C C T G A | bHLH10(bHLH)/colamp-bHLH10-DAP-Seq(GSE60143)/Homer | 1e-130 | -3.002e+02 | 0.0000 | 7089.0 | 17.90% | 23528.9 | 13.09% | motif file (matrix) | svg |
| 200 | G A C T A C G T A C G T A C T G A C G T A G T C G C A T A G C T G C A T G C A T G A C T A G C T | SGR5(C2H2)/colamp-SGR5-DAP-Seq(GSE60143)/Homer | 1e-130 | -2.996e+02 | 0.0000 | 4792.0 | 12.10% | 14600.7 | 8.12% | motif file (matrix) | svg |
| 201 | C A G T T C A G T C G A A G T C C G T A A C T G T G A C C G A T A C T G A C T G A C G T A T C G | Atoh7(bHLH)/Retina-Atoh7-CutnRun(GSE156756)/Homer | 1e-130 | -2.996e+02 | 0.0000 | 4612.0 | 11.65% | 13921.1 | 7.75% | motif file (matrix) | svg |
| 202 | A T G C A G T C G A T C C G T A A C G T A C G T A C T G A C G T A G C T G A T C | Sox2(HMG)/mES-Sox2-ChIP-Seq(GSE11431)/Homer | 1e-129 | -2.983e+02 | 0.0000 | 7018.0 | 17.72% | 23271.8 | 12.95% | motif file (matrix) | svg |
| 203 | T A G C C G T A C T G A T A C G C G T A A C G T A C T G A C T G A G T C T A C G C T A G G T A C | YY1(Zf)/Promoter/Homer | 1e-128 | -2.965e+02 | 0.0000 | 885.0 | 2.24% | 1332.0 | 0.74% | motif file (matrix) | svg |
| 204 | G T A C C A T G A G C T A C G T A C T G C G T A A G T C G A C T G C A T C G A T | WRKY29(WRKY)/colamp-WRKY29-DAP-Seq(GSE60143)/Homer | 1e-128 | -2.952e+02 | 0.0000 | 9261.0 | 23.39% | 32394.1 | 18.02% | motif file (matrix) | svg |
| 205 | C G A T T C G A G T A C A C G T A C G T T C A G G C A T G C A T G C T A C G T A C G T A A G T C C G T A T G C A C A T G | ANAC020(NAC)/col-ANAC020-DAP-Seq(GSE60143)/Homer | 1e-128 | -2.949e+02 | 0.0000 | 6487.0 | 16.38% | 21213.5 | 11.80% | motif file (matrix) | svg |
| 206 | G C A T C G T A G C T A G A C T C G A T G A C T A G T C C A G T A G T C A G T C A C T G C T A G G T A C C T A G C T G A | AT5G05550(Trihelix)/col-AT5G05550-DAP-Seq(GSE60143)/Homer | 1e-127 | -2.945e+02 | 0.0000 | 18573.0 | 46.91% | 72411.7 | 40.29% | motif file (matrix) | svg |
| 207 | G C T A C T G A T C G A A G T C A G T C C T G A A G T C G T C A C T G A T G C A | RUNX1(Runt)/Jurkat-RUNX1-ChIP-Seq(GSE29180)/Homer | 1e-127 | -2.925e+02 | 0.0000 | 8098.0 | 20.45% | 27689.7 | 15.41% | motif file (matrix) | svg |
| 208 | A G T C C T A G A C G T A C G T A C T G C G T A A G T C A G C T G C T A G C A T | WRKY24(WRKY)/colamp-WRKY24-DAP-Seq(GSE60143)/Homer | 1e-124 | -2.877e+02 | 0.0000 | 8237.0 | 20.80% | 28323.7 | 15.76% | motif file (matrix) | svg |
| 209 | G A C T T C A G G C A T A G T C G C T A G A T C C T G A A C G T A G T C G T C A | Replumless(BLH)/Arabidopsis-RPL.GFP-ChIP-Seq(GSE78727)/Homer | 1e-124 | -2.863e+02 | 0.0000 | 11674.0 | 29.48% | 42604.4 | 23.70% | motif file (matrix) | svg |
| 210 | G A C T C G A T T C A G G A T C G A C T A G C T A G C T A G T C G A T C C G T A C T A G C T A G T C G A T C G A C T G A | Bcl6(Zf)/Liver-Bcl6-ChIP-Seq(GSE31578)/Homer | 1e-123 | -2.849e+02 | 0.0000 | 6073.0 | 15.34% | 19713.3 | 10.97% | motif file (matrix) | svg |
| 211 | C T G A T A C G G C A T A G C T A G C T A G T C T C G A A C T G C A G T A G C T A G C T G A T C | IRF3(IRF)/BMDM-Irf3-ChIP-Seq(GSE67343)/Homer | 1e-123 | -2.837e+02 | 0.0000 | 1886.0 | 4.76% | 4361.2 | 2.43% | motif file (matrix) | svg |
| 212 | G A C T C G A T C G A T C T G A G T A C A G T C C G A T C G T A G T C A G A T C G C A T G C A T | MYB121(MYB)/col-MYB121-DAP-Seq(GSE60143)/Homer | 1e-122 | -2.824e+02 | 0.0000 | 4380.0 | 11.06% | 13230.0 | 7.36% | motif file (matrix) | svg |
| 213 | A G C T G C T A T G C A A G T C A C G T A C G T A C G T C G A T A G C T T C A G | dof24(C2C2dof)/col-dof24-DAP-Seq(GSE60143)/Homer | 1e-122 | -2.819e+02 | 0.0000 | 13623.0 | 34.41% | 50992.8 | 28.37% | motif file (matrix) | svg |
| 214 | G A C T A C G T A G C T G A C T A C T G C A G T A G T C A T C G A C G T G C A T G C A T G C A T | MGP(C2H2)/colamp-MGP-DAP-Seq(GSE60143)/Homer | 1e-121 | -2.794e+02 | 0.0000 | 3157.0 | 7.97% | 8759.3 | 4.87% | motif file (matrix) | svg |
| 215 | C T G A A G T C G A T C C A T G G C T A G A T C C T G A G C T A G C T A C G A T | AT1G77200(AP2EREBP)/colamp-AT1G77200-DAP-Seq(GSE60143)/Homer | 1e-118 | -2.733e+02 | 0.0000 | 15211.0 | 38.42% | 58029.0 | 32.29% | motif file (matrix) | svg |
| 216 | G T C A T G C A G C T A A G T C C G T A A C T G T G A C G C A T T C A G C A G T | Ap4(bHLH)/AML-Tfap4-ChIP-Seq(GSE45738)/Homer | 1e-118 | -2.724e+02 | 0.0000 | 6301.0 | 15.91% | 20775.8 | 11.56% | motif file (matrix) | svg |
| 217 | A T G C A G T C A G C T A G C T A C G T A T C G C G T A C G A T T A G C G A C T | LEF1(HMG)/H1-LEF1-ChIP-Seq(GSE64758)/Homer | 1e-118 | -2.718e+02 | 0.0000 | 4841.0 | 12.23% | 15097.9 | 8.40% | motif file (matrix) | svg |
| 218 | A G T C A C G T A C G T T C A G G C T A G C T A A T G C C G T A C G A T A G T C C G T A G T C A A C T G G A T C G C A T | SND3(NAC)/col-SND3-DAP-Seq(GSE60143)/Homer | 1e-117 | -2.704e+02 | 0.0000 | 6816.0 | 17.21% | 22848.7 | 12.71% | motif file (matrix) | svg |
| 219 | C G A T G A T C G A T C C T G A G A T C G A T C C A T G T G C A G T A C T C G A G T C A G C A T C G A T C G A T G C A T | At4g32800(AP2EREBP)/colamp-At4g32800-DAP-Seq(GSE60143)/Homer | 1e-115 | -2.667e+02 | 0.0000 | 3613.0 | 9.12% | 10534.3 | 5.86% | motif file (matrix) | svg |
| 220 | T C A G A C G T T C G A T A G C A G T C C G T A A C T G G T A C A C G T A C T G A T C G A G T C | Atoh1(bHLH)/Cerebellum-Atoh1-ChIP-Seq(GSE22111)/Homer | 1e-115 | -2.653e+02 | 0.0000 | 6727.0 | 16.99% | 22565.9 | 12.56% | motif file (matrix) | svg |
| 221 | A G T C G T A C C T G A A G T C A G T C C A T G G C T A A G T C T G C A G C T A G C A T G C A T | RAP21(AP2EREBP)/colamp-RAP21-DAP-Seq(GSE60143)/Homer | 1e-114 | -2.632e+02 | 0.0000 | 4899.0 | 12.37% | 15419.7 | 8.58% | motif file (matrix) | svg |
| 222 | T C A G A G C T G T C A C G T A A C G T A T G C C G T A A C G T A C G T C T G A | PHV(HB)/col-PHV-DAP-Seq(GSE60143)/Homer | 1e-113 | -2.614e+02 | 0.0000 | 3485.0 | 8.80% | 10115.7 | 5.63% | motif file (matrix) | svg |
| 223 | C G T A C G T A C G T A C G T A C T G A C A G T A C G T C G T A A C T G A C T G A C G T C T A G C T G A T C G A C T G A | MYB39(MYB)/col-MYB39-DAP-Seq(GSE60143)/Homer | 1e-113 | -2.614e+02 | 0.0000 | 1867.0 | 4.72% | 4438.5 | 2.47% | motif file (matrix) | svg |
| 224 | A T G C C A T G A C G T A C G T A C T G C G T A A G T C G A C T G C A T C G A T | WRKY71(WRKY)/col-WRKY71-DAP-Seq(GSE60143)/Homer | 1e-113 | -2.613e+02 | 0.0000 | 6892.0 | 17.41% | 23282.6 | 12.95% | motif file (matrix) | svg |
| 225 | A G C T C A T G G C A T G A T C T G C A C T A G G A T C A C G T | Tgif2(Homeobox)/mES-Tgif2-ChIP-Seq(GSE55404)/Homer | 1e-113 | -2.611e+02 | 0.0000 | 20080.0 | 50.71% | 79862.8 | 44.43% | motif file (matrix) | svg |
| 226 | C G A T C G T A A G T C A C G T A C G T T C G A T G C A G C A T G C T A C G T A A C G T A G C T C G T A C G T A A C T G | ANAC062(NAC)/colamp-ANAC062-DAP-Seq(GSE60143)/Homer | 1e-113 | -2.606e+02 | 0.0000 | 2833.0 | 7.15% | 7764.8 | 4.32% | motif file (matrix) | svg |
| 227 | C G A T T G A C C A T G G A C T A C G T C A T G C G T A G A T C G A C T G C A T G C A T C G A T | WRKY14(WRKY)/colamp-WRKY14-DAP-Seq(GSE60143)/Homer | 1e-112 | -2.592e+02 | 0.0000 | 4761.0 | 12.02% | 14937.7 | 8.31% | motif file (matrix) | svg |
| 228 | T G A C C T G A A G T C A G T C A C T G G A T C G A C T G C A T | At5g18450(AP2EREBP)/col-At5g18450-DAP-Seq(GSE60143)/Homer | 1e-112 | -2.591e+02 | 0.0000 | 17640.0 | 44.55% | 69000.3 | 38.39% | motif file (matrix) | svg |
| 229 | T A G C C A T G G A C T G A C T T C A G G T C A G A T C G A C T G C A T G C T A | WRKY15(WRKY)/col-WRKY15-DAP-Seq(GSE60143)/Homer | 1e-109 | -2.531e+02 | 0.0000 | 8922.0 | 22.53% | 31675.8 | 17.62% | motif file (matrix) | svg |
| 230 | G T A C A C G T A C G T T C A G G A C T G C A T T C A G C G T A C T G A A G T C C G T A G T C A A C T G A C G T G C T A | NTM2(NAC)/col-NTM2-DAP-Seq(GSE60143)/Homer | 1e-109 | -2.522e+02 | 0.0000 | 4913.0 | 12.41% | 15606.0 | 8.68% | motif file (matrix) | svg |
| 231 | T C G A T G A C G T A C C G T A C A G T T G A C A C G T A C T G A G C T A G C T | NeuroG2(bHLH)/Fibroblast-NeuroG2-ChIP-Seq(GSE75910)/Homer | 1e-109 | -2.512e+02 | 0.0000 | 9335.0 | 23.58% | 33418.8 | 18.59% | motif file (matrix) | svg |
| 232 | T G A C A G T C C G T A A C T G G T A C A C G T A C T G A C G T G A C T G A T C | Twist2(bHLH)/Myoblast-Twist2.Ty1-ChIP-Seq(GSE127998)/Homer | 1e-107 | -2.483e+02 | 0.0000 | 10182.0 | 25.72% | 37001.4 | 20.59% | motif file (matrix) | svg |
| 233 | G A C T G C A T G C A T A G T C A G C T T C G A T A C G G C T A C G T A A C T G G T A C G C A T C G A T A G T C G A C T | HSF3(HSF)/colamp-HSF3-DAP-Seq(GSE60143)/Homer | 1e-107 | -2.482e+02 | 0.0000 | 5767.0 | 14.56% | 18991.5 | 10.57% | motif file (matrix) | svg |
| 234 | T C G A T G A C G C A T A G C T C A G T G A T C G C T A G A T C G A C T A C G T G C A T A G T C | PRDM1(Zf)/Hela-PRDM1-ChIP-Seq(GSE31477)/Homer | 1e-107 | -2.477e+02 | 0.0000 | 3193.0 | 8.06% | 9177.0 | 5.11% | motif file (matrix) | svg |
| 235 | C G A T G C T A G C T A G C A T G C T A C G T A A G T C A C G T A C G T A C G T C G A T A G C T | At5g62940(C2C2dof)/col-At5g62940-DAP-Seq(GSE60143)/Homer | 1e-107 | -2.476e+02 | 0.0000 | 19693.0 | 49.74% | 78417.2 | 43.63% | motif file (matrix) | svg |
| 236 | C G A T T G C A T G C A G A T C C G T A A C T G T G A C G A C T C A T G A C T G | Tcf21(bHLH)/ArterySmoothMuscle-Tcf21-ChIP-Seq(GSE61369)/Homer | 1e-107 | -2.476e+02 | 0.0000 | 5043.0 | 12.74% | 16166.9 | 8.99% | motif file (matrix) | svg |
| 237 | C G T A C G A T C T A G C G T A A G C T C A G T T A C G C G T A A C G T C A T G C T A G A T G C | HOXA3(Homeobox)/mEmbryo-Hoxa3-ChIP-Seq(E-MTAB-8607)/Homer | 1e-106 | -2.452e+02 | 0.0000 | 1874.0 | 4.73% | 4567.5 | 2.54% | motif file (matrix) | svg |
| 238 | C T A G T C G A T G A C A G T C C G T A A C T G G T A C A C G T A C T G A C T G | BHLHA15(bHLH)/NIH3T3-BHLHB8.HA-ChIP-Seq(GSE119782)/Homer | 1e-105 | -2.424e+02 | 0.0000 | 7971.0 | 20.13% | 27942.0 | 15.55% | motif file (matrix) | svg |
| 239 | G C T A G C T A C G T A C G T A C T G A C T A G A C G T A G T C C G T A C T G A G T A C A C T G | WRKY65(WRKY)/colamp-WRKY65-DAP-Seq(GSE60143)/Homer | 1e-105 | -2.420e+02 | 0.0000 | 4902.0 | 12.38% | 15688.9 | 8.73% | motif file (matrix) | svg |
| 240 | T C A G T G A C C A T G G C A T C A G T A C T G C G T A T G A C G A C T C G A T C G A T C G T A | WRKY3(WRKY)/col-WRKY3-DAP-Seq(GSE60143)/Homer | 1e-104 | -2.395e+02 | 0.0000 | 6084.0 | 15.37% | 20366.4 | 11.33% | motif file (matrix) | svg |
| 241 | C G A T C G T A G C T A G C A T G C A T C T G A A C T G A C G T A G T C C G T A C G T A G T A C T C A G G C T A C G A T | WRKY25(WRKY)/colamp-WRKY25-DAP-Seq(GSE60143)/Homer | 1e-103 | -2.383e+02 | 0.0000 | 10091.0 | 25.49% | 36801.1 | 20.48% | motif file (matrix) | svg |
| 242 | T C A G T G A C G T A C C G T A A C G T T G A C A C G T T C A G A G C T G A C T | NeuroD1(bHLH)/Islet-NeuroD1-ChIP-Seq(GSE30298)/Homer | 1e-103 | -2.377e+02 | 0.0000 | 5056.0 | 12.77% | 16341.2 | 9.09% | motif file (matrix) | svg |
| 243 | G C T A T C G A C G T A C T G A A C T G A C G T A G T C C G T A C G T A A G T C C T A G T G C A | WRKY42(WRKY)/colamp-WRKY42-DAP-Seq(GSE60143)/Homer | 1e-102 | -2.369e+02 | 0.0000 | 4898.0 | 12.37% | 15737.1 | 8.76% | motif file (matrix) | svg |
| 244 | G C A T A G C T A G C T A G C T A C T G A C G T A G T C A C T G A C G T G A C T C G A T G C A T | JKD(C2H2)/col-JKD-DAP-Seq(GSE60143)/Homer | 1e-102 | -2.353e+02 | 0.0000 | 1942.0 | 4.90% | 4864.9 | 2.71% | motif file (matrix) | svg |
| 245 | G A C T C T G A G T A C A G T C C G A T C G T A G T C A G A T C G C A T G C A T G C A T C G A T | AT3G10580(MYBrelated)/colamp-AT3G10580-DAP-Seq(GSE60143)/Homer | 1e-100 | -2.314e+02 | 0.0000 | 4865.0 | 12.29% | 15679.0 | 8.72% | motif file (matrix) | svg |
| 246 | C A G T T C A G A G C T G A C T A C G T A G T C G A T C G A C T C T G A A C T G G A T C C G T A C T G A A G T C G T A C | Rfx6(HTH)/Min6b1-Rfx6.HA-ChIP-Seq(GSE62844)/Homer | 1e-100 | -2.311e+02 | 0.0000 | 7732.0 | 19.53% | 27149.0 | 15.10% | motif file (matrix) | svg |
| 247 | G C A T G C T A C G T A A G C T G C T A T G C A A G T C A C G T A C G T A C G T G C A T G C A T | At4g38000(C2C2dof)/col-At4g38000-DAP-Seq(GSE60143)/Homer | 1e-99 | -2.298e+02 | 0.0000 | 6342.0 | 16.02% | 21537.7 | 11.98% | motif file (matrix) | svg |
| 248 | C G A T C G T A A G T C A C G T A C G T T C A G G C A T C G T A G C T A G C T A C G T A A G T C C G T A G T C A A C T G | ANAC058(NAC)/col-ANAC058-DAP-Seq(GSE60143)/Homer | 1e-99 | -2.294e+02 | 0.0000 | 5649.0 | 14.27% | 18782.0 | 10.45% | motif file (matrix) | svg |
| 249 | C T A G C T A G T G A C G T A C C A T G A C T G G A T C G A T C C G T A C G T A | RAP211(AP2EREBP)/colamp-RAP211-DAP-Seq(GSE60143)/Homer | 1e-99 | -2.292e+02 | 0.0000 | 19957.0 | 50.40% | 80024.4 | 44.52% | motif file (matrix) | svg |
| 250 | A T G C A G T C C T G A A G T C C G A T A C G T A G T C A G T C A C G T A T C G G A C T A C G T | Etv2(ETS)/ES-ER71-ChIP-Seq(GSE59402)/Homer | 1e-99 | -2.289e+02 | 0.0000 | 5655.0 | 14.28% | 18813.3 | 10.47% | motif file (matrix) | svg |
| 251 | A G C T A G C T A G C T A C T G A C G T A G T C A C T G A C G T G C A T G C A T G C A T A C G T | At5g66730(C2H2)/colamp-At5g66730-DAP-Seq(GSE60143)/Homer | 1e-98 | -2.272e+02 | 0.0000 | 2590.0 | 6.54% | 7186.4 | 4.00% | motif file (matrix) | svg |
| 252 | C A T G G T A C A C T G G T C A A G C T T A C G T G C A A T C G T G A C C A G T | TOD6?/SacCer-Promoters/Homer | 1e-98 | -2.267e+02 | 0.0000 | 2647.0 | 6.69% | 7395.5 | 4.11% | motif file (matrix) | svg |
| 253 | C G A T A G T C C A T G G A C T A C G T C T A G C G T A G A T C G A C T C G A T G C A T G A C T | WRKY43(WRKY)/colamp-WRKY43-DAP-Seq(GSE60143)/Homer | 1e-97 | -2.251e+02 | 0.0000 | 3492.0 | 8.82% | 10510.8 | 5.85% | motif file (matrix) | svg |
| 254 | C G A T T C A G G T A C A C G T A C G T T C A G C G A T C G T A G T C A G C T A C G T A A G T C C G T A G T C A C A T G | ANAC057(NAC)/colamp-ANAC057-DAP-Seq(GSE60143)/Homer | 1e-97 | -2.250e+02 | 0.0000 | 7540.0 | 19.04% | 26462.3 | 14.72% | motif file (matrix) | svg |
| 255 | T G A C G C T A T C G A T G C A A G T C A G T C C G T A A G T C C G T A C T G A G C T A G T A C | RUNX2(Runt)/PCa-RUNX2-ChIP-Seq(GSE33889)/Homer | 1e-96 | -2.221e+02 | 0.0000 | 6358.0 | 16.06% | 21717.9 | 12.08% | motif file (matrix) | svg |
| 256 | T C A G A T C G G A C T A C T G G A C T C A G T C T A G C G T A G T A C C G T A C T A G A T C G | Tbx20(T-box)/Heart-Tbx20-ChIP-Seq(GSE29636)/Homer | 1e-95 | -2.197e+02 | 0.0000 | 2137.0 | 5.40% | 5650.4 | 3.14% | motif file (matrix) | svg |
| 257 | T C G A C T G A C G T A C G T A C G T A C T G A A C T G A C G T C T G A C T G A | AT5G63260(C3H)/col-AT5G63260-DAP-Seq(GSE60143)/Homer | 1e-95 | -2.197e+02 | 0.0000 | 14252.0 | 35.99% | 54973.4 | 30.59% | motif file (matrix) | svg |
| 258 | A G C T G A C T A C G T A C T G A C G T A G T C A C T G A C G T G C A T C G A T | AtIDD11(C2H2)/colamp-AtIDD11-DAP-Seq(GSE60143)/Homer | 1e-95 | -2.194e+02 | 0.0000 | 3532.0 | 8.92% | 10721.3 | 5.97% | motif file (matrix) | svg |
| 259 | T C G A A G T C A C G T A C G T T C A G C A G T C T G A C T A G T C G A C G T A A T C G C G T A C G T A A C T G A G C T | NTM1(NAC)/col-NTM1-DAP-Seq(GSE60143)/Homer | 1e-95 | -2.191e+02 | 0.0000 | 3455.0 | 8.73% | 10436.5 | 5.81% | motif file (matrix) | svg |
| 260 | C A T G A C T G C T A G T C G A T C G A T C G A T C G A T C A G T C A G T C A G T G A C T G A C C G T A A C T G T G C A C G A T A C T G | RBPJ:Ebox(?,bHLH)/Panc1-Rbpj1-ChIP-Seq(GSE47459)/Homer | 1e-95 | -2.190e+02 | 0.0000 | 1701.0 | 4.30% | 4167.0 | 2.32% | motif file (matrix) | svg |
| 261 | A G T C G A C T A G C T C G A T A T C G G C T A C G A T A T C G C G A T A C T G T A C G A C G T | Tcf7(HMG)/GM12878-TCF7-ChIP-Seq(Encode)/Homer | 1e-94 | -2.184e+02 | 0.0000 | 2614.0 | 6.60% | 7352.0 | 4.09% | motif file (matrix) | svg |
| 262 | C G T A T A C G T C G A A C T G A C T G C G T A C G T A T A C G A G C T T A C G | PU.1(ETS)/ThioMac-PU.1-ChIP-Seq(GSE21512)/Homer | 1e-94 | -2.167e+02 | 0.0000 | 3156.0 | 7.97% | 9347.2 | 5.20% | motif file (matrix) | svg |
| 263 | G C T A C G T A G C T A C G T A C T G A A C T G A C G T G T A C C G T A C T G A G T A C A C T G | WRKY22(WRKY)/colamp-WRKY22-DAP-Seq(GSE60143)/Homer | 1e-93 | -2.164e+02 | 0.0000 | 5225.0 | 13.20% | 17288.2 | 9.62% | motif file (matrix) | svg |
| 264 | A T G C G A C T A C G T C T A G A C G T A C G T A C G T C T G A G A T C G C T A A G C T C G T A | Foxa2(Forkhead)/Liver-Foxa2-ChIP-Seq(GSE25694)/Homer | 1e-93 | -2.161e+02 | 0.0000 | 5289.0 | 13.36% | 17544.9 | 9.76% | motif file (matrix) | svg |
| 265 | T G A C C G A T C T G A C T A G C T A G A C G T A T G C T G C A T C G A C T G A C T A G C A T G A C G T A G T C C G T A | PPARa(NR),DR1/Liver-Ppara-ChIP-Seq(GSE47954)/Homer | 1e-93 | -2.158e+02 | 0.0000 | 6048.0 | 15.27% | 20568.3 | 11.44% | motif file (matrix) | svg |
| 266 | C G A T T C G A A T G C C G A T G C A T T C G A A G C T G C A T G C A T C G A T T C G A A G C T T C G A G T C A C T A G | ANAC004(NAC)/colamp-ANAC004-DAP-Seq(GSE60143)/Homer | 1e-92 | -2.126e+02 | 0.0000 | 2550.0 | 6.44% | 7173.9 | 3.99% | motif file (matrix) | svg |
| 267 | G A C T C T A G A T G C A G T C G T C A T A C G A T G C A T C G | HIC1(Zf)/Treg-ZBTB29-ChIP-Seq(GSE99889)/Homer | 1e-91 | -2.112e+02 | 0.0000 | 14681.0 | 37.08% | 57031.5 | 31.73% | motif file (matrix) | svg |
| 268 | G C T A T C G A C G T A C T A G A G C T G T C A G T C A C G T A A G T C C G T A | FOXA1(Forkhead)/LNCAP-FOXA1-ChIP-Seq(GSE27824)/Homer | 1e-91 | -2.097e+02 | 0.0000 | 5856.0 | 14.79% | 19891.7 | 11.07% | motif file (matrix) | svg |
| 269 | A C G T T C G A T C G A A G T C G T C A T A C G A T G C A C G T A C T G A G C T | Myf5(bHLH)/GM-Myf5-ChIP-Seq(GSE24852)/Homer | 1e-90 | -2.087e+02 | 0.0000 | 3755.0 | 9.48% | 11685.9 | 6.50% | motif file (matrix) | svg |
| 270 | C G T A C G T A C G T A C T G A C T A G A C G T C T A G G T C A | CDF3(C2C2dof)/colamp-CDF3-DAP-Seq(GSE60143)/Homer | 1e-90 | -2.079e+02 | 0.0000 | 11407.0 | 28.81% | 42962.0 | 23.90% | motif file (matrix) | svg |
| 271 | A T G C G T A C A C T G A G T C A G T C A C T G G A T C G T C A C G T A C G A T G C A T C G A T | RRTF1(AP2EREBP)/colamp-RRTF1-DAP-Seq(GSE60143)/Homer | 1e-90 | -2.077e+02 | 0.0000 | 5764.0 | 14.56% | 19553.8 | 10.88% | motif file (matrix) | svg |
| 272 | C T A G C A T G A C G T C G T A C T A G C A T G C G A T C T A G T C A G T C A G | MYB3(MYB)/Arabidopsis-MYB3-ChIP-Seq(GSE80564)/Homer | 1e-88 | -2.047e+02 | 0.0000 | 15439.0 | 38.99% | 60501.3 | 33.66% | motif file (matrix) | svg |
| 273 | C G A T G A T C T A C G C T G A G C T A C G T A G C A T A G T C C T A G C G T A G C A T C G A T | AT2G15740(C2H2)/col-AT2G15740-DAP-Seq(GSE60143)/Homer | 1e-87 | -2.016e+02 | 0.0000 | 17780.0 | 44.90% | 70931.1 | 39.46% | motif file (matrix) | svg |
| 274 | T C A G C T G A C G T A C G T A T A C G G C A T C T A G C T G A C G T A C G T A T A C G G A C T | IRF1(IRF)/PBMC-IRF1-ChIP-Seq(GSE43036)/Homer | 1e-87 | -2.014e+02 | 0.0000 | 882.0 | 2.23% | 1679.9 | 0.93% | motif file (matrix) | svg |
| 275 | C G A T T C G A A G T C A C G T A C G T T A C G C G T A G C T A G C T A C G A T G C A T A T G C C G T A G T C A A C T G | ANAC071(NAC)/col-ANAC071-DAP-Seq(GSE60143)/Homer | 1e-87 | -2.012e+02 | 0.0000 | 8399.0 | 21.21% | 30418.3 | 16.92% | motif file (matrix) | svg |
| 276 | C G T A G C T A C G A T C T A G A C G T G T C A C G T A C G T A A G T C C G T A T G C A T A C G | FoxL2(Forkhead)/Ovary-FoxL2-ChIP-Seq(GSE60858)/Homer | 1e-87 | -2.011e+02 | 0.0000 | 4948.0 | 12.50% | 16409.0 | 9.13% | motif file (matrix) | svg |
| 277 | T C G A T C G A A G T C C G T A C T A G T A G C A C G T A C T G | MyoG(bHLH)/C2C12-MyoG-ChIP-Seq(GSE36024)/Homer | 1e-86 | -1.991e+02 | 0.0000 | 5697.0 | 14.39% | 19415.6 | 10.80% | motif file (matrix) | svg |
| 278 | C G A T C T A G A G T C A C G T A C G T T C A G G C T A C G T A G C A T G C A T C G A T A G T C C G T A G T C A A C T G | VND3(NAC)/colamp-VND3-DAP-Seq(GSE60143)/Homer | 1e-85 | -1.968e+02 | 0.0000 | 5944.0 | 15.01% | 20442.4 | 11.37% | motif file (matrix) | svg |
| 279 | C T A G A C T G T G C A G T C A A T G C C G T A A T C G A T G C A G T C C T A G | ZNF341(Zf)/EBV-ZNF341-ChIP-Seq(GSE113194)/Homer | 1e-85 | -1.965e+02 | 0.0000 | 5330.0 | 13.46% | 17988.1 | 10.01% | motif file (matrix) | svg |
| 280 | T A C G T G C A A G T C C G T A A C G T T G A C A C G T A C T G A C T G G C A T | TCF4(bHLH)/SHSY5Y-TCF4-ChIP-Seq(GSE96915)/Homer | 1e-84 | -1.947e+02 | 0.0000 | 8957.0 | 22.62% | 32868.8 | 18.29% | motif file (matrix) | svg |
| 281 | G A T C C A T G A C G T A C G T A C T G C G T A A G T C A G C T C G A T G A C T | WRKY8(WRKY)/colamp-WRKY8-DAP-Seq(GSE60143)/Homer | 1e-81 | -1.885e+02 | 0.0000 | 1177.0 | 2.97% | 2647.8 | 1.47% | motif file (matrix) | svg |
| 282 | C T G A C T G A C T G A A T G C G A T C C A T G A C T G G A C T G A C T G C A T C G T A C G T A A G T C G T A C C T G A A T C G G C A T G A C T G A C T A G C T | GRHL2(CP2)/HBE-GRHL2-ChIP-Seq(GSE46194)/Homer | 1e-81 | -1.882e+02 | 0.0000 | 3054.0 | 7.71% | 9271.1 | 5.16% | motif file (matrix) | svg |
| 283 | A G T C A C G T A C G T T C A G G C T A C G T A G C T A G C A T C G A T A G T C C G T A G T C A A C T G G A C T G C T A | SMB(NAC)/colamp-SMB-DAP-Seq(GSE60143)/Homer | 1e-81 | -1.873e+02 | 0.0000 | 8563.0 | 21.63% | 31363.8 | 17.45% | motif file (matrix) | svg |
| 284 | G C A T C G A T C G T A G A T C C A T G A C G T A C G T A C T G C G T A A G T C A G C T G C A T G C A T C G T A G C T A | WRKY45(WRKY)/col-WRKY45-DAP-Seq(GSE60143)/Homer | 1e-80 | -1.862e+02 | 0.0000 | 3228.0 | 8.15% | 9948.2 | 5.53% | motif file (matrix) | svg |
| 285 | A C T G A C G T C A T G A T C G A T C G T G A C A C T G A T C G A T C G T G C A C T G A C G T A | E2F3(E2F)/MEF-E2F3-ChIP-Seq(GSE71376)/Homer | 1e-80 | -1.861e+02 | 0.0000 | 7785.0 | 19.66% | 28149.2 | 15.66% | motif file (matrix) | svg |
| 286 | A C G T T G C A A G C T G A T C C T A G C T G A A G C T G T C A T C G A C G T A | CUX1(Homeobox)/K562-CUX1-ChIP-Seq(GSE92882)/Homer | 1e-80 | -1.857e+02 | 0.0000 | 9481.0 | 23.94% | 35248.8 | 19.61% | motif file (matrix) | svg |
| 287 | A T G C C G T A C G T A C G T A C G T A C G T A A C T G A C G T C G A T C T G A | dof43(C2C2dof)/colamp-dof43-DAP-Seq(GSE60143)/Homer | 1e-80 | -1.855e+02 | 0.0000 | 9683.0 | 24.46% | 36105.9 | 20.09% | motif file (matrix) | svg |
| 288 | G A T C G C A T T C G A A G T C A C G T A C G T A C G T C G A T A C G T A T C G | AT1G47655(C2C2dof)/colamp-AT1G47655-DAP-Seq(GSE60143)/Homer | 1e-80 | -1.849e+02 | 0.0000 | 20047.0 | 50.63% | 81520.2 | 45.36% | motif file (matrix) | svg |
| 289 | A T G C G A T C C G T A A G C T C A G T A T C G G C A T A G C T G A C T A C T G | Sox17(HMG)/Endoderm-Sox17-ChIP-Seq(GSE61475)/Homer | 1e-79 | -1.837e+02 | 0.0000 | 6017.0 | 15.20% | 20947.2 | 11.65% | motif file (matrix) | svg |
| 290 | G A C T T C G A C G T A C G T A C G T A C G T A C G T A C T A G A G C T C G T A | dof45(C2C2dof)/col-dof45-DAP-Seq(GSE60143)/Homer | 1e-79 | -1.833e+02 | 0.0000 | 14887.0 | 37.60% | 58588.5 | 32.60% | motif file (matrix) | svg |
| 291 | T C G A C T G A C G T A C G T A C G T A C G T A A C T G A C G T C G A T C T G A | BBX31(Orphan)/col-BBX31-DAP-Seq(GSE60143)/Homer | 1e-79 | -1.822e+02 | 0.0000 | 10351.0 | 26.14% | 39006.1 | 21.70% | motif file (matrix) | svg |
| 292 | C A G T A G C T G A C T T G C A A G T C A G C T A C G T A C G T C G A T G A C T | AT3G52440(C2C2dof)/colamp-AT3G52440-DAP-Seq(GSE60143)/Homer | 1e-78 | -1.817e+02 | 0.0000 | 13984.0 | 35.32% | 54675.6 | 30.42% | motif file (matrix) | svg |
| 293 | C A G T C A T G T G C A G T A C C G T A T C A G G T A C G A C T T C A G C T G A | bZIP18(bZIP)/colamp-bZIP18-DAP-Seq(GSE60143)/Homer | 1e-78 | -1.802e+02 | 0.0000 | 26434.0 | 66.76% | 110984.2 | 61.75% | motif file (matrix) | svg |
| 294 | C G T A C G T A C G T A C G T A C G T A A C T G A C T G A G T C | dof42(C2C2dof)/col-dof42-DAP-Seq(GSE60143)/Homer | 1e-78 | -1.802e+02 | 0.0000 | 5106.0 | 12.90% | 17341.9 | 9.65% | motif file (matrix) | svg |
| 295 | G A T C C G T A G A C T C T A G G A T C C T G A G A C T C T G A G A C T C T A G G A T C C T G A G A C T C T G A G A C T | OCT:OCT(POU,Homeobox)/NPC-OCT6-ChIP-Seq(GSE43916)/Homer | 1e-77 | -1.786e+02 | 0.0000 | 467.0 | 1.18% | 637.7 | 0.35% | motif file (matrix) | svg |
| 296 | T G A C C T G A C T A G T C G A C T G A A T G C C G T A A C T G G C A T G T A C G C A T A T C G G C A T A G C T G A T C | PR(NR)/T47D-PR-ChIP-Seq(GSE31130)/Homer | 1e-76 | -1.756e+02 | 0.0000 | 12060.0 | 30.46% | 46474.3 | 25.86% | motif file (matrix) | svg |
| 297 | A C G T C T A G A G C T A C G T A C G T C T G A A G T C G A C T A G C T C G T A | FOXM1(Forkhead)/MCF7-FOXM1-ChIP-Seq(GSE72977)/Homer | 1e-76 | -1.755e+02 | 0.0000 | 5302.0 | 13.39% | 18198.6 | 10.13% | motif file (matrix) | svg |
| 298 | A G C T C T G A C T A G C T A G A C T G T A G C T G C A T C G A C T G A C T A G C A T G A C G T A T G C T C G A | RXR(NR),DR1/3T3L1-RXR-ChIP-Seq(GSE13511)/Homer | 1e-75 | -1.746e+02 | 0.0000 | 5988.0 | 15.12% | 20980.7 | 11.67% | motif file (matrix) | svg |
| 299 | A G T C C G T A A C G T A G T C A C G T A C T G | Tal1 | 1e-75 | -1.740e+02 | 0.0000 | 8960.0 | 22.63% | 33294.3 | 18.52% | motif file (matrix) | svg |
| 300 | T G A C T A G C T C A G T C G A T C G A C G T A A G T C C G T A C G T A C G A T C T A G T A C G | Sox7(HMG)/ESC-Sox7-ChIP-Seq(GSE133899)/Homer | 1e-74 | -1.724e+02 | 0.0000 | 2992.0 | 7.56% | 9215.4 | 5.13% | motif file (matrix) | svg |
| 301 | G C A T A C G T A C G T A T C G C G T A C G T A C G T A C G T A | At2g41835(C2H2)/col-At2g41835-DAP-Seq(GSE60143)/Homer | 1e-73 | -1.703e+02 | 0.0000 | 2920.0 | 7.37% | 8968.6 | 4.99% | motif file (matrix) | svg |
| 302 | A G T C A C G T A C G T T A C G G C T A G C T A C G T A C G A T C G A T A T G C C G T A G T C A A C T G G A C T G C A T | SND2(NAC)/colamp-SND2-DAP-Seq(GSE60143)/Homer | 1e-73 | -1.691e+02 | 0.0000 | 5820.0 | 14.70% | 20391.2 | 11.35% | motif file (matrix) | svg |
| 303 | T C G A C G T A A G T C C G T A C T A G A G T C C G A T A C T G G A C T A G C T A C T G G A C T | HLH-1(bHLH)/cElegans-Embryo-HLH1-ChIP-Seq(modEncode)/Homer | 1e-73 | -1.688e+02 | 0.0000 | 5114.0 | 12.92% | 17549.9 | 9.76% | motif file (matrix) | svg |
| 304 | T A G C G T A C A G T C G T A C C G A T A G T C A G T C A G T C A G T C A G T C C G T A G A T C | Zfp281(Zf)/ES-Zfp281-ChIP-Seq(GSE81042)/Homer | 1e-72 | -1.678e+02 | 0.0000 | 821.0 | 2.07% | 1654.8 | 0.92% | motif file (matrix) | svg |
| 305 | A C T G T C A G A G C T G A C T C A T G A G T C A G T C G C T A C G A T C T A G T C A G G T A C C T G A T C G A | Rfx1(HTH)/NPC-H3K4me1-ChIP-Seq(GSE16256)/Homer | 1e-71 | -1.654e+02 | 0.0000 | 1647.0 | 4.16% | 4379.9 | 2.44% | motif file (matrix) | svg |
| 306 | G C A T A C G T A C T G A C G T A G T C A C T G A T C G G T C A C G A T C G T A | ARF2(ARF)/col-ARF2-DAP-Seq(GSE60143)/Homer | 1e-71 | -1.647e+02 | 0.0000 | 20636.0 | 52.12% | 84726.8 | 47.14% | motif file (matrix) | svg |
| 307 | G T A C G C T A T C A G C T G A C T A G C A T G A G C T G A T C T G C A T C G A C T G A A C T G C A G T A G T C G A T C G C T A | HNF4a(NR),DR1/HepG2-HNF4a-ChIP-Seq(GSE25021)/Homer | 1e-70 | -1.623e+02 | 0.0000 | 2465.0 | 6.23% | 7356.8 | 4.09% | motif file (matrix) | svg |
| 308 | T C G A A G C T A C G T A C G T A G T C A G T C A C G T A T C G G A C T A T C G | EWS:ERG-fusion(ETS)/CADO\_ES1-EWS:ERG-ChIP-Seq(SRA014231)/Homer | 1e-70 | -1.620e+02 | 0.0000 | 3679.0 | 9.29% | 11979.3 | 6.66% | motif file (matrix) | svg |
| 309 | C T A G A G T C T A C G T A C G T G A C C G T A A C T G T A G C G C A T C A T G A T G C A G C T | Ascl1(bHLH)/NeuralTubes-Ascl1-ChIP-Seq(GSE55840)/Homer | 1e-69 | -1.605e+02 | 0.0000 | 8571.0 | 21.65% | 31939.4 | 17.77% | motif file (matrix) | svg |
| 310 | G T A C A C G T A C G T T A C G A T G C C A T G T A C G G T A C T C A G A T G C C G T A G T C A A C T G A G C T G C T A | AT1G19040(NAC)/col-AT1G19040-DAP-Seq(GSE60143)/Homer | 1e-69 | -1.602e+02 | 0.0000 | 1420.0 | 3.59% | 3637.0 | 2.02% | motif file (matrix) | svg |
| 311 | C A G T G C T A G C A T T A C G C T G A C A G T T A G C C T G A | GATA15(C2C2gata)/col-GATA15-DAP-Seq(GSE60143)/Homer | 1e-69 | -1.599e+02 | 0.0000 | 14672.0 | 37.06% | 58237.5 | 32.40% | motif file (matrix) | svg |
| 312 | G C T A G C T A C T G A A C T G A C G T A G T C C G T A C G T A G T A C A C T G A T G C G C A T | WRKY47(WRKY)/colamp-WRKY47-DAP-Seq(GSE60143)/Homer | 1e-68 | -1.575e+02 | 0.0000 | 3348.0 | 8.46% | 10756.7 | 5.98% | motif file (matrix) | svg |
| 313 | T C G A C T G A T A G C T G A C T C A G T C A G C G T A C G T A T C A G A G C T | ETV1(ETS)/GIST48-ETV1-ChIP-Seq(GSE22441)/Homer | 1e-67 | -1.562e+02 | 0.0000 | 10423.0 | 26.32% | 39895.6 | 22.20% | motif file (matrix) | svg |
| 314 | T C G A C T G A C G A T C G T A C G T A C G T A C T A G A G C T C T G A T C A G | Adof1(C2C2dof)/col-Adof1-DAP-Seq(GSE60143)/Homer | 1e-67 | -1.558e+02 | 0.0000 | 16174.0 | 40.85% | 64974.2 | 36.15% | motif file (matrix) | svg |
| 315 | T C A G A G C T A C G T A C G T G T A C G A T C C G T A C T A G C A T G G T C A C G T A T C G A | STAT4(Stat)/CD4-Stat4-ChIP-Seq(GSE22104)/Homer | 1e-66 | -1.535e+02 | 0.0000 | 4997.0 | 12.62% | 17327.8 | 9.64% | motif file (matrix) | svg |
| 316 | A T C G T C G A G A C T A T C G T G A C A C G T C T A G A C T G C G T A A C T G A G T C G T A C | ZNF415(Zf)/HEK293-ZNF415.GFP-ChIP-Seq(GSE58341)/Homer | 1e-66 | -1.535e+02 | 0.0000 | 4198.0 | 10.60% | 14137.2 | 7.87% | motif file (matrix) | svg |
| 317 | C A G T T A G C A G T C C A T G C A G T C A T G C G A T C G A T G A C T C G A T A T C G G T A C A C T G A T C G G T A C | LBD13(LOBAS2)/colamp-LBD13-DAP-Seq(GSE60143)/Homer | 1e-66 | -1.532e+02 | 0.0000 | 12163.0 | 30.72% | 47459.1 | 26.40% | motif file (matrix) | svg |
| 318 | G T A C T C G A T A G C C G T A C G T A C T G A T G C A T G A C A C T G C G T A A G T C C T G A C T G A T C G A C G T A | NUC(C2H2)/col-NUC-DAP-Seq(GSE60143)/Homer | 1e-66 | -1.529e+02 | 0.0000 | 991.0 | 2.50% | 2263.3 | 1.26% | motif file (matrix) | svg |
| 319 | G A C T A C T G C G T A A G T C T C A G G C A T G T A C C G T A A C G T G A T C | TGA1(bZIP)/colamp-TGA1-DAP-Seq(GSE60143)/Homer | 1e-66 | -1.526e+02 | 0.0000 | 4674.0 | 11.80% | 16046.1 | 8.93% | motif file (matrix) | svg |
| 320 | A G T C A G T C C T G A A G T C A G T C A C T G C G T A A G T C T C G A G A T C C G A T C G T A | AT1G01250(AP2EREBP)/col-AT1G01250-DAP-Seq(GSE60143)/Homer | 1e-65 | -1.507e+02 | 0.0000 | 2210.0 | 5.58% | 6537.0 | 3.64% | motif file (matrix) | svg |
| 321 | G C T A T C G A C G T A C T A G A G C T G T C A G T C A C G T A A G T C C G T A | FOXA1(Forkhead)/MCF7-FOXA1-ChIP-Seq(GSE26831)/Homer | 1e-65 | -1.506e+02 | 0.0000 | 4373.0 | 11.04% | 14876.4 | 8.28% | motif file (matrix) | svg |
| 322 | C T A G T A C G G A T C G T A C G C T A A G C T A G C T G T C A T C G A T A G C | Nanog(Homeobox)/mES-Nanog-ChIP-Seq(GSE11724)/Homer | 1e-63 | -1.458e+02 | 0.0000 | 26190.0 | 66.14% | 110782.3 | 61.64% | motif file (matrix) | svg |
| 323 | G T A C G C T A C G A T C A G T A G T C G C T A C G A T C G A T A G T C G C T A | WUS1(Homeobox)/colamp-WUS1-DAP-Seq(GSE60143)/Homer | 1e-62 | -1.448e+02 | 0.0000 | 3132.0 | 7.91% | 10090.6 | 5.61% | motif file (matrix) | svg |
| 324 | T A G C G T A C C T A G C A G T T C G A C G T A C G T A G C A T G A C T T G A C A G T C A C T G A T C G A G T C C T A G | AS2(LOBAS2)/col-AS2-DAP-Seq(GSE60143)/Homer | 1e-62 | -1.440e+02 | 0.0000 | 1964.0 | 4.96% | 5702.6 | 3.17% | motif file (matrix) | svg |
| 325 | C T G A A C T G C G T A A C G T G T C A A G C T A G C T G A C T G A C T C A G T | CCA(Myb)/Arabidopsis-CCA.GFP-ChIP-Seq(GSE70533)/Homer | 1e-62 | -1.434e+02 | 0.0000 | 6611.0 | 16.70% | 24107.7 | 13.41% | motif file (matrix) | svg |
| 326 | C A T G T G A C C A T G G A C T C A G T C T A G G C T A G T A C G A C T G C A T G C A T C G A T | WRKY21(WRKY)/colamp-WRKY21-DAP-Seq(GSE60143)/Homer | 1e-62 | -1.429e+02 | 0.0000 | 1091.0 | 2.76% | 2653.3 | 1.48% | motif file (matrix) | svg |
| 327 | G T C A T C G A T C G A C G T A G C T A C G T A T C G A T G A C A C T G C G T A A G T C C G T A C G T A T C G A G C T A | IDD2(C2H2)/colamp-IDD2-DAP-Seq(GSE60143)/Homer | 1e-61 | -1.427e+02 | 0.0000 | 1114.0 | 2.81% | 2731.2 | 1.52% | motif file (matrix) | svg |
| 328 | T C G A T C G A C T G A C G T A A C T G A T G C A C G T A G T C | Lola-I(Zf)/Embryo-LolaI-ChIP-Seq(GSE200870)/Homer | 1e-60 | -1.401e+02 | 0.0000 | 3912.0 | 9.88% | 13208.5 | 7.35% | motif file (matrix) | svg |
| 329 | C G A T C G T A A G T C A C G T A C G T T C A G C G T A C G T A G C A T G C A T G C A T A G T C C G T A G T C A A C T G | VND2(NAC)/col-VND2-DAP-Seq(GSE60143)/Homer | 1e-59 | -1.378e+02 | 0.0000 | 8453.0 | 21.35% | 31941.4 | 17.77% | motif file (matrix) | svg |
| 330 | C G T A C T G A C G T A C T A G T C G A C T A G A C T G C G T A C G T A T A C G A G C T A T C G | SpiB(ETS)/OCILY3-SPIB-ChIP-Seq(GSE56857)/Homer | 1e-59 | -1.364e+02 | 0.0000 | 1620.0 | 4.09% | 4533.8 | 2.52% | motif file (matrix) | svg |
| 331 | G T C A T C G A C T A G C T A G A G T C G T C A C G A T C T A G G A C T G A T C G A T C T C A G C T A G C T G A A G T C G C T A C A G T T C A G G A T C G A T C | p63(p53)/Keratinocyte-p63-ChIP-Seq(GSE17611)/Homer | 1e-59 | -1.362e+02 | 0.0000 | 2981.0 | 7.53% | 9620.3 | 5.35% | motif file (matrix) | svg |
| 332 | A G T C C G A T A C T G A T C G T G A C G C T A C A T G A T C G T G A C C G A T A C T G T A G C G T A C G T C A | Tlx?(NR)/NPC-H3K4me1-ChIP-Seq(GSE16256)/Homer | 1e-58 | -1.358e+02 | 0.0000 | 2040.0 | 5.15% | 6065.1 | 3.37% | motif file (matrix) | svg |
| 333 | C G T A C G A T C T A G G T C A G A C T C G A T C T A G C G T A A C G T C A T G | LIN-39(Homeobox)/cElegans.L3-LIN39-ChIP-Seq(modEncode)/Homer | 1e-58 | -1.357e+02 | 0.0000 | 7031.0 | 17.76% | 26011.5 | 14.47% | motif file (matrix) | svg |
| 334 | A G T C C A G T T C A G A T G C A G T C C G A T C G T A G T C A G A T C G C A T | BOS1(MYB)/col-BOS1-DAP-Seq(GSE60143)/Homer | 1e-58 | -1.354e+02 | 0.0000 | 9838.0 | 24.85% | 37891.9 | 21.08% | motif file (matrix) | svg |
| 335 | G T A C C A T G T A G C A G T C C T A G C A T G C T G A C G T A G C A T G C A T A C G T G C A T G T A C A C T G A T C G | LOB(LOBAS2)/col-LOB-DAP-Seq(GSE60143)/Homer | 1e-58 | -1.348e+02 | 0.0000 | 5180.0 | 13.08% | 18393.6 | 10.23% | motif file (matrix) | svg |
| 336 | C A G T T C G A A G T C A C G T A C G T T C A G C G A T G C T A G C T A C G T A G C A T A G T C C G T A T G C A A C T G | ANAC045(NAC)/col-ANAC045-DAP-Seq(GSE60143)/Homer | 1e-58 | -1.338e+02 | 0.0000 | 16123.0 | 40.72% | 65371.3 | 36.37% | motif file (matrix) | svg |
| 337 | T A C G C T G A C A T G G A T C G T A C G C A T T C A G T A C G A G C T G T C A G A T C G C A T T A C G C G T A C T A G G A T C G A T C C G A T A C T G T C A G | ZNF322(Zf)/HEK293-ZNF322.GFP-ChIP-Seq(GSE58341)/Homer | 1e-58 | -1.336e+02 | 0.0000 | 1146.0 | 2.89% | 2903.9 | 1.62% | motif file (matrix) | svg |
| 338 | T C A G G C A T A C T G C G T A A G T C C T A G G C A T T G A C | TGA9(bZIP)/colamp-TGA9-DAP-Seq(GSE60143)/Homer | 1e-57 | -1.332e+02 | 0.0000 | 12799.0 | 32.32% | 50752.6 | 28.24% | motif file (matrix) | svg |
| 339 | A T G C T C G A A G T C A G C T A C G T G T A C A G T C G C T A C T A G C A T G G T C A C T G A T C A G A G T C | Stat3+il21(Stat)/CD4-Stat3-ChIP-Seq(GSE19198)/Homer | 1e-57 | -1.318e+02 | 0.0000 | 4016.0 | 10.14% | 13750.2 | 7.65% | motif file (matrix) | svg |
| 340 | C G T A A C T G C G T A A C G T A T C G C A G T T A G C C G T A T C G A G T A C C T G A T A G C C G T A A C T G C G T A A C G T C G T A C T G A A T C G G C T A | GATA3(Zf),DR8/iTreg-Gata3-ChIP-Seq(GSE20898)/Homer | 1e-56 | -1.310e+02 | 0.0000 | 714.0 | 1.80% | 1515.6 | 0.84% | motif file (matrix) | svg |
| 341 | T G A C C A T G A C G T A C G T A C T G C G T A A G T C A G C T G C A T T C G A | WRKY30(WRKY)/colamp-WRKY30-DAP-Seq(GSE60143)/Homer | 1e-56 | -1.296e+02 | 0.0000 | 3753.0 | 9.48% | 12739.4 | 7.09% | motif file (matrix) | svg |
| 342 | G C A T C T A G G T A C A G T C C G A T A C T G C T A G C T A G G T A C G C T A | ZNF416(Zf)/HEK293-ZNF416.GFP-ChIP-Seq(GSE58341)/Homer | 1e-55 | -1.280e+02 | 0.0000 | 6046.0 | 15.27% | 22078.9 | 12.28% | motif file (matrix) | svg |
| 343 | C G A T G T C A A G T C A C G T A C G T A C T G G A C T C G A T A T C G G C T A G T C A A G T C C G T A G T C A A C T G | ANAC017(NAC)/colamp-ANAC017-DAP-Seq(GSE60143)/Homer | 1e-55 | -1.275e+02 | 0.0000 | 1585.0 | 4.00% | 4490.3 | 2.50% | motif file (matrix) | svg |
| 344 | T C G A T G A C G T A C C G T A A C G T G A C T A C G T A C T G A C T G A G C T | Mesp1(bHLH)/ESC-Mesp1-ChIP-Seq(GSE165102)/Homer | 1e-54 | -1.259e+02 | 0.0000 | 4001.0 | 10.10% | 13787.6 | 7.67% | motif file (matrix) | svg |
| 345 | G T A C A C G T A C G T T C A G G C T A C G T A C G A T G C A T G C A T A G T C C G T A G T C A C A T G G A C T G C T A | ANAC070(NAC)/colamp-ANAC070-DAP-Seq(GSE60143)/Homer | 1e-54 | -1.254e+02 | 0.0000 | 9250.0 | 23.36% | 35626.3 | 19.82% | motif file (matrix) | svg |
| 346 | C T G A C T A G A C T G G C A T A T G C C G T A C T G A C T A G A C T G A C G T A G T C C T G A | RARg(NR)/ES-RARg-ChIP-Seq(GSE30538)/Homer | 1e-53 | -1.237e+02 | 0.0000 | 332.0 | 0.84% | 464.6 | 0.26% | motif file (matrix) | svg |
| 347 | C A G T A G C T C G T A G C A T A G T C G A C T C T A G C T A G C A G T C T A G T C G A T G C A C T A G C A T G G A C T | STOP1(C2H2)/colamp-STOP1-DAP-Seq(GSE60143)/Homer | 1e-53 | -1.231e+02 | 0.0000 | 3649.0 | 9.22% | 12429.6 | 6.92% | motif file (matrix) | svg |
| 348 | G A C T C T A G C T A G A G T C T G C A A C T G A C G T A C G T C T A G T C A G | AMYB(HTH)/Testes-AMYB-ChIP-Seq(GSE44588)/Homer | 1e-52 | -1.202e+02 | 0.0000 | 16251.0 | 41.04% | 66354.6 | 36.92% | motif file (matrix) | svg |
| 349 | G A C T C T A G C T A G C T A G A C T G T C G A C T G A C T A G C T A G C T A G G T A C G T C A | ZNF467(Zf)/HEK293-ZNF467.GFP-ChIP-Seq(GSE58341)/Homer | 1e-52 | -1.201e+02 | 0.0000 | 5046.0 | 12.74% | 18119.6 | 10.08% | motif file (matrix) | svg |
| 350 | G T A C A C G T A C G T T C A G C G T A C G T A C G A T G C A T G C A T A G T C C G T A G T C A C A T G G A C T G C T A | VND1(NAC)/col-VND1-DAP-Seq(GSE60143)/Homer | 1e-51 | -1.197e+02 | 0.0000 | 6812.0 | 17.20% | 25443.4 | 14.16% | motif file (matrix) | svg |
| 351 | A C T G G A T C G A C T A C T G A C G T C A T G A C T G A C G T A G C T C G A T | RUNX-AML(Runt)/CD4+-PolII-ChIP-Seq(Barski\_et\_al.)/Homer | 1e-51 | -1.184e+02 | 0.0000 | 4756.0 | 12.01% | 16967.7 | 9.44% | motif file (matrix) | svg |
| 352 | T G C A C T G A A T G C G T C A A C G T A T G C A C G T A C T G A C T G T G C A | ZBTB18(Zf)/HEK293-ZBTB18.GFP-ChIP-Seq(GSE58341)/Homer | 1e-51 | -1.177e+02 | 0.0000 | 2812.0 | 7.10% | 9219.4 | 5.13% | motif file (matrix) | svg |
| 353 | C G A T T C G A G T A C A C G T A C G T T C A G G C A T G C T A T G C A G C A T C G T A A G T C C G T A T G A C C A T G | ANAC092(NAC)/colamp-ANAC092-DAP-Seq(GSE60143)/Homer | 1e-49 | -1.146e+02 | 0.0000 | 4700.0 | 11.87% | 16811.4 | 9.35% | motif file (matrix) | svg |
| 354 | T A G C C A T G A G C T A C G T A C T G C G T A A G T C G A C T G C A T C T G A | AT3G42860(zfGRF)/col-AT3G42860-DAP-Seq(GSE60143)/Homer | 1e-49 | -1.143e+02 | 0.0000 | 3304.0 | 8.34% | 11199.1 | 6.23% | motif file (matrix) | svg |
| 355 | T A G C G T A C C T A G A T C G C T G A C G T A G C T A G C A T A C G T T G A C G T A C A C T G T A C G G T C A C T A G | ASL18(LOBAS2)/colamp-ASL18-DAP-Seq(GSE60143)/Homer | 1e-48 | -1.112e+02 | 0.0000 | 13572.0 | 34.28% | 54782.7 | 30.48% | motif file (matrix) | svg |
| 356 | A T G C G A C T A G C T C T A G C G T A C T A G C G A T C T A G A T C G G A T C | Nkx2.2(Homeobox)/NPC-Nkx2.2-ChIP-Seq(GSE61673)/Homer | 1e-48 | -1.109e+02 | 0.0000 | 14394.0 | 36.35% | 58412.1 | 32.50% | motif file (matrix) | svg |
| 357 | C G A T C T G A A G T C A C G T A C G T T C A G G C A T C G A T G C T A G C T A C G T A A G T C C G T A G T C A A C T G | CUC1(NAC)/col-CUC1-DAP-Seq(GSE60143)/Homer | 1e-48 | -1.109e+02 | 0.0000 | 4560.0 | 11.52% | 16310.7 | 9.07% | motif file (matrix) | svg |
| 358 | C T G A A T G C G C T A C G A T A T G C C G T A C G T A C G T A C T A G T A C G | Tcf3(HMG)/mES-Tcf3-ChIP-Seq(GSE11724)/Homer | 1e-47 | -1.098e+02 | 0.0000 | 1816.0 | 4.59% | 5519.1 | 3.07% | motif file (matrix) | svg |
| 359 | C G T A A C T G C G T A A C G T C A G T A G T C A G C T G C A T G C T A C G A T | At2g01060(G2like)/colamp-At2g01060-DAP-Seq(GSE60143)/Homer | 1e-47 | -1.085e+02 | 0.0000 | 20789.0 | 52.50% | 87139.2 | 48.48% | motif file (matrix) | svg |
| 360 | A G T C T A G C G A C T A C G T C T A G A C G T A C G T A C G T C T G A A G T C G C T A G A C T C G T A C T A G A C T G | Foxa3(Forkhead)/Liver-Foxa3-ChIP-Seq(GSE77670)/Homer | 1e-46 | -1.078e+02 | 0.0000 | 1979.0 | 5.00% | 6153.2 | 3.42% | motif file (matrix) | svg |
| 361 | C G T A C T A G T C A G T C A G A G T C A T G C A G T C G C A T A G C T A C G T A T C G C G A T | Sox9(HMG)/Limb-SOX9-ChIP-Seq(GSE73225)/Homer | 1e-46 | -1.078e+02 | 0.0000 | 5381.0 | 13.59% | 19745.8 | 10.99% | motif file (matrix) | svg |
| 362 | G C T A T G A C G A T C C G A T G A C T A T G C C T G A A T C G G C A T A C G T | JGL(C2H2)/col-JGL-DAP-Seq(GSE60143)/Homer | 1e-46 | -1.078e+02 | 0.0000 | 10308.0 | 26.03% | 40644.8 | 22.61% | motif file (matrix) | svg |
| 363 | T C A G G A C T G T C A C G T A A C G T A T C G C G T A A C G T A C G T C T G A | ATHB15(HB)/col-ATHB15-DAP-Seq(GSE60143)/Homer | 1e-46 | -1.076e+02 | 0.0000 | 2745.0 | 6.93% | 9102.9 | 5.06% | motif file (matrix) | svg |
| 364 | G T A C G A T C C A G T A G T C A G T C A G T C T G C A G A T C C T G A A T G C G T C A A C G T | WT1(Zf)/Kidney-WT1-ChIP-Seq(GSE90016)/Homer | 1e-46 | -1.074e+02 | 0.0000 | 4157.0 | 10.50% | 14737.2 | 8.20% | motif file (matrix) | svg |
| 365 | C A G T A T C G C T G A A G T C T C A G C A G T T A G C C T G A A T G C T A C G | FEA4(bZIP)/Corn-FEA4-ChIP-Seq(GSE61954)/Homer | 1e-46 | -1.073e+02 | 0.0000 | 10690.0 | 27.00% | 42310.0 | 23.54% | motif file (matrix) | svg |
| 366 | G A C T G T A C T G C A A C G T G A T C G C T A T C G A A C G T A G T C C G T A | Pdx1(Homeobox)/Islet-Pdx1-ChIP-Seq(SRA008281)/Homer | 1e-46 | -1.071e+02 | 0.0000 | 6744.0 | 17.03% | 25447.6 | 14.16% | motif file (matrix) | svg |
| 367 | T A C G T C G A G A C T A C T G C T G A A G T C T C A G G A C T T G A C C T G A | Atf1(bZIP)/K562-ATF1-ChIP-Seq(GSE31477)/Homer | 1e-46 | -1.061e+02 | 0.0000 | 7133.0 | 18.01% | 27112.3 | 15.08% | motif file (matrix) | svg |
| 368 | G C A T G C A T G C A T A T G C A G C T T C G A T A C G G C T A C G T A C A T G G T A C G C A T G C A T A G T C A G C T | HSFA6B(HSF)/colamp-HSFA6B-DAP-Seq(GSE60143)/Homer | 1e-45 | -1.048e+02 | 0.0000 | 2880.0 | 7.27% | 9674.4 | 5.38% | motif file (matrix) | svg |
| 369 | C A G T G A C T G C A T T C G A A G T C A C G T A C G T A C G T C G A T G A C T | OBP3(C2C2dof)/col-OBP3-DAP-Seq(GSE60143)/Homer | 1e-45 | -1.042e+02 | 0.0000 | 16918.0 | 42.73% | 69850.6 | 38.86% | motif file (matrix) | svg |
| 370 | A G C T G C A T G T C A C G A T T A G C C G T A A C G T G C T A | CRC(C2C2YABBY)/col-CRC-DAP-Seq(GSE60143)/Homer | 1e-45 | -1.041e+02 | 0.0000 | 9845.0 | 24.86% | 38748.0 | 21.56% | motif file (matrix) | svg |
| 371 | A G T C C T G A A T C G A G C T A G C T G A C T A G T C G C T A A C G T C G A T G C A T C G A T A T C G C G T A T A G C G C A T A T G C C G T A | bZIP:IRF(bZIP,IRF)/Th17-BatF-ChIP-Seq(GSE39756)/Homer | 1e-45 | -1.038e+02 | 0.0000 | 1745.0 | 4.41% | 5322.8 | 2.96% | motif file (matrix) | svg |
| 372 | C G A T C T A G C T G A A T G C C T G A T C G A C G T A C T G A T C G A T A G C A G T C C G T A A C T G T C G A A T G C | Hand2(bHLH)/Mesoderm-Hand2-ChIP-Seq(GSE61475)/Homer | 1e-44 | -1.035e+02 | 0.0000 | 2409.0 | 6.08% | 7854.9 | 4.37% | motif file (matrix) | svg |
| 373 | A T G C T C A G T C G A G C A T A C T G C G T A A G T C T C A G G A C T T G A C C G T A A G C T | Atf2(bZIP)/3T3L1-Atf2-ChIP-Seq(GSE56872)/Homer | 1e-44 | -1.026e+02 | 0.0000 | 2574.0 | 6.50% | 8507.4 | 4.73% | motif file (matrix) | svg |
| 374 | C T G A A T G C C G T A A C G T A G T C A G T C A C G T A C T G A T C G G C A T | SPDEF(ETS)/VCaP-SPDEF-ChIP-Seq(SRA014231)/Homer | 1e-44 | -1.020e+02 | 0.0000 | 7351.0 | 18.57% | 28135.6 | 15.65% | motif file (matrix) | svg |
| 375 | G T C A C G T A A C G T A T C G C G T A A C G T A C G T C T A G | ATHB7(Homeobox)/col-ATHB7-DAP-Seq(GSE60143)/Homer | 1e-43 | -1.007e+02 | 0.0000 | 6072.0 | 15.34% | 22780.5 | 12.67% | motif file (matrix) | svg |
| 376 | C G A T C T G A A G T C A C G T A C G T T C A G C G T A C G T A C G T A G C A T C G A T A G T C C G T A G T C A A C T G | VND4(NAC)/colamp-VND4-DAP-Seq(GSE60143)/Homer | 1e-43 | -9.987e+01 | 0.0000 | 6747.0 | 17.04% | 25635.4 | 14.26% | motif file (matrix) | svg |
| 377 | C G A T C T A G T C A G C A G T C G T A A G T C G C T A A C G T G A C T A T G C A G T C G C T A | PRDM10(Zf)/HEK293-PRDM10.eGFP-ChIP-Seq(Encode)/Homer | 1e-43 | -9.986e+01 | 0.0000 | 4017.0 | 10.15% | 14310.6 | 7.96% | motif file (matrix) | svg |
| 378 | C T A G G T A C A C G T A C G T A T C G G C A T A G C T A G C T A G C T G C A T G A C T C G T A G T C A A C T G G A C T | VND6(NAC)/col-VND6-DAP-Seq(GSE60143)/Homer | 1e-41 | -9.668e+01 | 0.0000 | 10284.0 | 25.97% | 40865.1 | 22.74% | motif file (matrix) | svg |
| 379 | C T A G T A C G G A C T T G C A T G C A C G A T T A C G C T G A T C G A C T G A | Hoxa10(Homeobox)/ChickenMSG-Hoxa10.Flag-ChIP-Seq(GSE86088)/Homer | 1e-41 | -9.661e+01 | 0.0000 | 3842.0 | 9.70% | 13661.7 | 7.60% | motif file (matrix) | svg |
| 380 | C G A T T A C G T G C A G T A C G A T C G A C T A G C T A C G T A T C G G T A C G A T C G T A C G A T C G T C A | PPARE(NR),DR1/3T3L1-Pparg-ChIP-Seq(GSE13511)/Homer | 1e-41 | -9.590e+01 | 0.0000 | 4854.0 | 12.26% | 17820.6 | 9.91% | motif file (matrix) | svg |
| 381 | T A C G T A C G G T A C A T C G A C T G T A C G T C G A C T G A T C G A A T C G | E2F6(E2F)/Hela-E2F6-ChIP-Seq(GSE31477)/Homer | 1e-41 | -9.446e+01 | 0.0000 | 5214.0 | 13.17% | 19343.5 | 10.76% | motif file (matrix) | svg |
| 382 | C A T G G A T C C T G A G T A C C T A G C T G A G C T A G C A T G A T C G A T C A G T C C T A G C G T A C A T G C T A G | AIL7(AP2EREBP)/colamp-AIL7-DAP-Seq(GSE60143)/Homer | 1e-41 | -9.444e+01 | 0.0000 | 5411.0 | 13.67% | 20163.6 | 11.22% | motif file (matrix) | svg |
| 383 | G A C T C A G T A G C T C G A T A G T C G A T C A G T C C G T A A T G C T C A G | Rbpj1(?)/Panc1-Rbpj1-ChIP-Seq(GSE47459)/Homer | 1e-40 | -9.385e+01 | 0.0000 | 7657.0 | 19.34% | 29649.8 | 16.50% | motif file (matrix) | svg |
| 384 | C G A T C A G T C T A G G C T A A G T C C G T A T C A G A G T C A C G T A C T G A C G T G T A C G C T A G C T A G C T A | bZIP52(bZIP)/colamp-bZIP52-DAP-Seq(GSE60143)/Homer | 1e-40 | -9.240e+01 | 0.0000 | 7738.0 | 19.54% | 30034.9 | 16.71% | motif file (matrix) | svg |
| 385 | C G A T C T G A G T A C A C G T A C G T T C A G C G T A C G T A G C T A G C A T G C A T A G T C C G T A G T C A C A T G | NST1(NAC)/colamp-NST1-DAP-Seq(GSE60143)/Homer | 1e-39 | -9.115e+01 | 0.0000 | 6711.0 | 16.95% | 25703.6 | 14.30% | motif file (matrix) | svg |
| 386 | C T A G T C G A C T G A C G T A T A C G G A C T T C A G T C G A G T C A T G C A T A C G A G C T | IRF2(IRF)/Erythroblas-IRF2-ChIP-Seq(GSE36985)/Homer | 1e-39 | -9.086e+01 | 0.0000 | 765.0 | 1.93% | 1928.4 | 1.07% | motif file (matrix) | svg |
| 387 | C T A G T C G A C G A T C T A G G C A T C A G T C T A G G A T C C G T A G T C A | CEBP:AP1(bZIP)/ThioMac-CEBPb-ChIP-Seq(GSE21512)/Homer | 1e-39 | -9.034e+01 | 0.0000 | 5725.0 | 14.46% | 21571.8 | 12.00% | motif file (matrix) | svg |
| 388 | T C G A C G T A C G T A T C G A A C T G G T A C C G T A A G C T G T C A G C A T | At3g24120(G2like)/col-At3g24120-DAP-Seq(GSE60143)/Homer | 1e-39 | -9.022e+01 | 0.0000 | 21140.0 | 53.39% | 89391.6 | 49.73% | motif file (matrix) | svg |
| 389 | C T G A A G C T A C G T A C G T A G T C G A C T G A C T C T G A C T G A C T A G C G T A C G T A | STAT6(Stat)/CD4-Stat6-ChIP-Seq(GSE22104)/Homer | 1e-39 | -8.993e+01 | 0.0000 | 2628.0 | 6.64% | 8912.5 | 4.96% | motif file (matrix) | svg |
| 390 | C G A T C T A G A C G T G T C A C G T A C G T A A G T C C G T A | Foxo3(Forkhead)/U2OS-Foxo3-ChIP-Seq(E-MTAB-2701)/Homer | 1e-38 | -8.971e+01 | 0.0000 | 4590.0 | 11.59% | 16863.5 | 9.38% | motif file (matrix) | svg |
| 391 | G A C T C T A G C T A G G T A C A G T C G A T C G A C T G A C T T A G C T C A G | NLP7(RWPRK)/col-NLP7-DAP-Seq(GSE60143)/Homer | 1e-38 | -8.821e+01 | 0.0000 | 13484.0 | 34.05% | 55148.6 | 30.68% | motif file (matrix) | svg |
| 392 | T C A G T A C G T A G C A C G T A C T G C G A T A G T C C G T A T A C G A G T C | Meis1(Homeobox)/MastCells-Meis1-ChIP-Seq(GSE48085)/Homer | 1e-38 | -8.815e+01 | 0.0000 | 11719.0 | 29.60% | 47389.8 | 26.37% | motif file (matrix) | svg |
| 393 | G A C T A G T C C G A T A C T G C T G A T G A C G T A C C G T A A T C G G C A T C T G A C T A G | Bcl11a(Zf)/HSPC-BCL11A-ChIP-Seq(GSE104676)/Homer | 1e-38 | -8.804e+01 | 0.0000 | 4050.0 | 10.23% | 14677.4 | 8.17% | motif file (matrix) | svg |
| 394 | G C A T C G T A G C A T C G T A T C G A C G T A C T G A A C T G C G T A C G T A C G T A A C G T A C T G G T C A G C A T | AT2G31460(REMB3)/col-AT2G31460-DAP-Seq(GSE60143)/Homer | 1e-37 | -8.741e+01 | 0.0000 | 1947.0 | 4.92% | 6292.0 | 3.50% | motif file (matrix) | svg |
| 395 | T C G A T C G A T A G C G T A C T C A G T A C G C G T A C G T A T C A G A G C T | GABPA(ETS)/Jurkat-GABPa-ChIP-Seq(GSE17954)/Homer | 1e-37 | -8.633e+01 | 0.0000 | 7446.0 | 18.81% | 28956.8 | 16.11% | motif file (matrix) | svg |
| 396 | C T G A T C A G C T G A C T A G C A T G A C G T A T G C C G T A A T G C G C A T T C A G C T G A A C T G A C G T C A G T A G T C C G T A C A G T C T A G C A T G | VDR(NR),DR3/GM10855-VDR+vitD-ChIP-Seq(GSE22484)/Homer | 1e-37 | -8.613e+01 | 0.0000 | 1591.0 | 4.02% | 4952.4 | 2.76% | motif file (matrix) | svg |
| 397 | T A C G A T G C G A C T A C T G A G C T A G T C G T C A T G C A A C G T A G T C G C T A T G C A | Pknox1(Homeobox)/ES-Prep1-ChIP-Seq(GSE63282)/Homer | 1e-36 | -8.514e+01 | 0.0000 | 1888.0 | 4.77% | 6096.1 | 3.39% | motif file (matrix) | svg |
| 398 | A T G C C T G A A T C G T A C G A G T C C G A T T C A G C G A T C T A G A G C T G T C A G T C A C G T A A G T C C G T A T A C G C T G A | Fox:Ebox(Forkhead,bHLH)/Panc1-Foxa2-ChIP-Seq(GSE47459)/Homer | 1e-36 | -8.438e+01 | 0.0000 | 4792.0 | 12.10% | 17818.4 | 9.91% | motif file (matrix) | svg |
| 399 | A G T C C T G A A G T C C G A T C A G T G A T C A T G C A C T G A T C G G A C T | Fli1(ETS)/CD8-FLI-ChIP-Seq(GSE20898)/Homer | 1e-36 | -8.339e+01 | 0.0000 | 11036.0 | 27.87% | 44561.3 | 24.79% | motif file (matrix) | svg |
| 400 | A G T C A T C G C T A G A G C T G A C T C T A G A G T C A G T C G C T A C A G T T C A G T C A G G A T C C T G A T C G A G A T C | RFX(HTH)/K562-RFX3-ChIP-Seq(SRA012198)/Homer | 1e-35 | -8.281e+01 | 0.0000 | 618.0 | 1.56% | 1494.8 | 0.83% | motif file (matrix) | svg |
| 401 | G A C T C A G T G A T C G A T C A C G T G A T C C T G A T A C G C G T A G T C A | STAT6(Stat)/Macrophage-Stat6-ChIP-Seq(GSE38377)/Homer | 1e-35 | -8.266e+01 | 0.0000 | 2712.0 | 6.85% | 9365.1 | 5.21% | motif file (matrix) | svg |
| 402 | G C A T G A T C T C A G G C T A G A C T A G T C C T A G C G T A C A T G G T C A | GATA20(C2C2gata)/colamp-GATA20-DAP-Seq(GSE60143)/Homer | 1e-35 | -8.199e+01 | 0.0000 | 21549.0 | 54.42% | 91570.3 | 50.95% | motif file (matrix) | svg |
| 403 | T C G A G C A T A C G T C T A G G T A C T C G A G C A T T G A C T C G A A C G T | Chop(bZIP)/MEF-Chop-ChIP-Seq(GSE35681)/Homer | 1e-35 | -8.144e+01 | 0.0000 | 2039.0 | 5.15% | 6732.8 | 3.75% | motif file (matrix) | svg |
| 404 | T C G A G A C T A T C G C G T A A G T C C T A G G C A T G T A C C T G A A C G T G A T C G C T A | TGA4(bZIP)/colamp-TGA4-DAP-Seq(GSE60143)/Homer | 1e-34 | -8.004e+01 | 0.0000 | 3293.0 | 8.32% | 11750.5 | 6.54% | motif file (matrix) | svg |
| 405 | C T G A C G A T C T A G C G T A A G C T C G A T C A G T C T G A G A C T C T A G C T A G A T G C | PBX2(Homeobox)/K562-PBX2-ChIP-Seq(Encode)/Homer | 1e-34 | -7.977e+01 | 0.0000 | 6563.0 | 16.58% | 25379.1 | 14.12% | motif file (matrix) | svg |
| 406 | C A T G C T A G A G C T G A C T C A T G A G T C G A T C G C T A C G A T C T A G T C A G G T A C C T G A T C G A | X-box(HTH)/NPC-H3K4me1-ChIP-Seq(GSE16256)/Homer | 1e-34 | -7.893e+01 | 0.0000 | 657.0 | 1.66% | 1651.9 | 0.92% | motif file (matrix) | svg |
| 407 | C A G T A G C T G C A T T C G A A G T C A C G T A C G T A C G T C G A T G C A T | AT5G66940(C2C2dof)/col-AT5G66940-DAP-Seq(GSE60143)/Homer | 1e-34 | -7.884e+01 | 0.0000 | 11261.0 | 28.44% | 45701.6 | 25.43% | motif file (matrix) | svg |
| 408 | A G T C G A T C G A T C C G T A G T C A A G T C A G C T C T G A G A C T G A C T | ATY13(MYB)/col-ATY13-DAP-Seq(GSE60143)/Homer | 1e-33 | -7.784e+01 | 0.0000 | 23381.0 | 59.05% | 100106.6 | 55.70% | motif file (matrix) | svg |
| 409 | G T A C C T G A A G T C A G T C A C T G G T C A G A T C G C A T | At1g75490(AP2EREBP)/colamp-At1g75490-DAP-Seq(GSE60143)/Homer | 1e-33 | -7.690e+01 | 0.0000 | 22711.0 | 57.36% | 97077.7 | 54.01% | motif file (matrix) | svg |
| 410 | A T C G T G A C A T G C C T G A T C A G G A C T A G T C C G A T T C A G T C G A C A T G C T A G C T A G C G T A C T A G C T A G C T G A C T A G C T A G A T G C | ZSCAN22(Zf)/HEK293-ZSCAN22.GFP-ChIP-Seq(GSE58341)/Homer | 1e-33 | -7.666e+01 | 0.0000 | 417.0 | 1.05% | 888.5 | 0.49% | motif file (matrix) | svg |
| 411 | T A G C C T A G T C G A G A C T A C T G C T G A A G T C T C A G G C A T T G A C C T G A A G C T | Atf7(bZIP)/3T3L1-Atf7-ChIP-Seq(GSE56872)/Homer | 1e-33 | -7.618e+01 | 0.0000 | 3996.0 | 10.09% | 14707.2 | 8.18% | motif file (matrix) | svg |
| 412 | C G A T C T A G G A T C G C T A A G C T C T A G G A T C C G T A | RBFox2(?)/Heart-RBFox2-CLIP-Seq(GSE57926)/Homer | 1e-32 | -7.546e+01 | 0.0000 | 14805.0 | 37.39% | 61473.2 | 34.20% | motif file (matrix) | svg |
| 413 | G A C T C T A G G A T C C A G T A C T G C T G A A T G C G C A T A T G C C T G A | MafA(bZIP)/Islet-MafA-ChIP-Seq(GSE30298)/Homer | 1e-32 | -7.536e+01 | 0.0000 | 5415.0 | 13.68% | 20643.3 | 11.49% | motif file (matrix) | svg |
| 414 | T C G A A C G T A C T G C G T A A G T C C T A G A G C T T G A C | TGA10(bZIP)/colamp-TGA10-DAP-Seq(GSE60143)/Homer | 1e-32 | -7.517e+01 | 0.0000 | 7216.0 | 18.22% | 28295.3 | 15.74% | motif file (matrix) | svg |
| 415 | A G T C G A C T C A G T A C T G C T A G T G A C G C T A A T G C G C A T A T C G C G A T A C T G G A T C G T A C G T C A C T G A | NF1(CTF)/LNCAP-NF1-ChIP-Seq(Unpublished)/Homer | 1e-32 | -7.407e+01 | 0.0000 | 2076.0 | 5.24% | 6988.9 | 3.89% | motif file (matrix) | svg |
| 416 | G A C T A T C G C T G A A G T C T C A G G A C T G T A C C T G A A G C T G T A C | TGA6(bZIP)/colamp-TGA6-DAP-Seq(GSE60143)/Homer | 1e-31 | -7.277e+01 | 0.0000 | 7744.0 | 19.56% | 30633.4 | 17.04% | motif file (matrix) | svg |
| 417 | T A G C T A G C G A C T C T A G A G C T A G T C G T C A T G C A A C G T A T G C G C T A T G C A | Pbx3(Homeobox)/GM12878-PBX3-ChIP-Seq(GSE32465)/Homer | 1e-31 | -7.261e+01 | 0.0000 | 1637.0 | 4.13% | 5302.2 | 2.95% | motif file (matrix) | svg |
| 418 | G C A T T C A G C T G A A T C G A C T G C G A T G A T C C T G A | THRb(NR)/Liver-NR1A2-ChIP-Seq(GSE52613)/Homer | 1e-31 | -7.219e+01 | 0.0000 | 20379.0 | 51.47% | 86653.3 | 48.21% | motif file (matrix) | svg |
| 419 | C T A G A G C T G A C T C A T G A G T C A G T C G T C A C A G T C T A G T C A G G T A C C T G A T C G A G A T C T G A C | Rfx2(HTH)/LoVo-RFX2-ChIP-Seq(GSE49402)/Homer | 1e-31 | -7.157e+01 | 0.0000 | 654.0 | 1.65% | 1696.5 | 0.94% | motif file (matrix) | svg |
| 420 | C T G A A T C G A G C T A G C T A C G T T A G C C T G A T A C G C G A T A C G T G A C T A G T C | ISRE(IRF)/ThioMac-LPS-Expression(GSE23622)/Homer | 1e-30 | -7.078e+01 | 0.0000 | 348.0 | 0.88% | 708.9 | 0.39% | motif file (matrix) | svg |
| 421 | C T G A T A C G G C A T C T A G A T G C G A T C C G A T A C T G C T A G G A T C C T G A A T G C | MYRF(MYRF)/CFPAC1-MYRF-ChIP-Seq(GSE145627)/Homer | 1e-30 | -7.075e+01 | 0.0000 | 2404.0 | 6.07% | 8342.1 | 4.64% | motif file (matrix) | svg |
| 422 | A T G C A G T C G T A C A G C T T C G A C T A G G A T C C T G A G T C A A G T C G C T A T C A G | Rfx5(HTH)/GM12878-Rfx5-ChIP-Seq(GSE31477)/Homer | 1e-30 | -7.059e+01 | 0.0000 | 2538.0 | 6.41% | 8881.2 | 4.94% | motif file (matrix) | svg |
| 423 | C G T A T G A C T C G A A G T C C G T A A T C G A T G C A C G T A C T G A G T C | E2A(bHLH)/proBcell-E2A-ChIP-Seq(GSE21978)/Homer | 1e-30 | -6.928e+01 | 0.0000 | 7253.0 | 18.32% | 28633.4 | 15.93% | motif file (matrix) | svg |
| 424 | A C T G A G C T A G T C G T C A A G C T T C A G A T G C G A T C G C A T A T C G T C G A T A G C C G A T C A T G T A G C | Pax8(Paired,Homeobox)/Thyroid-Pax8-ChIP-Seq(GSE26938)/Homer | 1e-30 | -6.912e+01 | 0.0000 | 2096.0 | 5.29% | 7146.1 | 3.98% | motif file (matrix) | svg |
| 425 | T A C G T A C G C T A G T C A G A G T C C G T A A T C G A T G C A C G T A C T G A G T C G A C T | Ascl2(bHLH)/ESC-Ascl2-ChIP-Seq(GSE97712)/Homer | 1e-30 | -6.910e+01 | 0.0000 | 6027.0 | 15.22% | 23402.4 | 13.02% | motif file (matrix) | svg |
| 426 | T C G A G C A T A C T G C T G A A G T C T C A G G A C T G T A C C G T A A G C T A G T C G A T C | c-Jun-CRE(bZIP)/K562-cJun-ChIP-Seq(GSE31477)/Homer | 1e-29 | -6.770e+01 | 0.0000 | 1846.0 | 4.66% | 6185.3 | 3.44% | motif file (matrix) | svg |
| 427 | C T A G C T A G T C A G G T C A C T A G T C A G G C T A A G T C A T C G A G C T C T A G | DPR(core promoter) | 1e-29 | -6.753e+01 | 0.0000 | 33484.0 | 84.57% | 147761.9 | 82.21% | motif file (matrix) | svg |
| 428 | C T G A G A C T G A T C C T G A A G T C G C A T A C G T G A C T G C T A G C A T | OBP1(C2C2dof)/col-OBP1-DAP-Seq(GSE60143)/Homer | 1e-29 | -6.724e+01 | 0.0000 | 15577.0 | 39.34% | 65247.9 | 36.30% | motif file (matrix) | svg |
| 429 | T A G C G C T A T C G A C T G A A G T C A G T C C T G A A G T C C G T A C T A G | RUNX(Runt)/HPC7-Runx1-ChIP-Seq(GSE22178)/Homer | 1e-28 | -6.452e+01 | 0.0000 | 5817.0 | 14.69% | 22643.0 | 12.60% | motif file (matrix) | svg |
| 430 | T A C G T A G C C A T G C A G T A C G T C T A G C G T A A G T C G A C T G C A T G C A T C A G T | WRKY11(WRKY)/col-WRKY11-DAP-Seq(GSE60143)/Homer | 1e-27 | -6.418e+01 | 0.0000 | 1328.0 | 3.35% | 4232.1 | 2.35% | motif file (matrix) | svg |
| 431 | A T G C G A C T A C T G C A G T G A T C A C G T T A C G T A C G | Smad2(MAD)/ES-SMAD2-ChIP-Seq(GSE29422)/Homer | 1e-27 | -6.388e+01 | 0.0000 | 12378.0 | 31.26% | 51175.4 | 28.47% | motif file (matrix) | svg |
| 432 | A C G T A G T C A G C T A G T C C G T A G T A C A G T C C G A T C G T A G T C A | MYB41(MYB)/col-MYB41-DAP-Seq(GSE60143)/Homer | 1e-27 | -6.343e+01 | 0.0000 | 3771.0 | 9.52% | 14068.1 | 7.83% | motif file (matrix) | svg |
| 433 | A T C G A G T C A G T C C G T A A C T G G C A T | hINR(CPE) | 1e-27 | -6.285e+01 | 0.0000 | 8218.0 | 20.76% | 33006.5 | 18.36% | motif file (matrix) | svg |
| 434 | T C G A G C T A T G A C G C T A C T A G G A T C C G A T A C T G C G A T A G C T G A C T C T A G | E-box/Drosophila-Promoters/Homer | 1e-27 | -6.268e+01 | 0.0000 | 1379.0 | 3.48% | 4447.9 | 2.47% | motif file (matrix) | svg |
| 435 | T G C A C T G A A T G C G T C A A C T G A C T G C G T A C G T A C T A G A G C T | Ets1-distal(ETS)/CD4+-PolII-ChIP-Seq(Barski\_et\_al.)/Homer | 1e-27 | -6.264e+01 | 0.0000 | 1504.0 | 3.80% | 4929.9 | 2.74% | motif file (matrix) | svg |
| 436 | T G C A G C A T A G C T G C A T A G T C A G T C A G T C C T G A A C T G T C G A T C G A C A G T A T C G A G T C G A T C | ZNF143\|STAF(Zf)/CUTLL-ZNF143-ChIP-Seq(GSE29600)/Homer | 1e-27 | -6.226e+01 | 0.0000 | 1557.0 | 3.93% | 5140.4 | 2.86% | motif file (matrix) | svg |
| 437 | T C G A T A G C G T C A A C T G C T A G C G T A C G A T A C T G A C G T A C T G A C T G A C G T | ETS:RUNX(ETS,Runt)/Jurkat-RUNX1-ChIP-Seq(GSE17954)/Homer | 1e-26 | -6.165e+01 | 0.0000 | 689.0 | 1.74% | 1900.8 | 1.06% | motif file (matrix) | svg |
| 438 | A C T G C G T A A C T G A T G C T G A C G A T C A T C G T G C A A C T G A G T C | ZNF519(Zf)/HEK293-ZNF519.GFP-ChIP-Seq(GSE58341)/Homer | 1e-26 | -6.134e+01 | 0.0000 | 1472.0 | 3.72% | 4824.4 | 2.68% | motif file (matrix) | svg |
| 439 | C G A T C T A G T C G A A G C T C G A T C T G A C G T A A G C T A C T G C T A G A T G C G A T C | Hoxb4(Homeobox)/ES-Hoxb4-ChIP-Seq(GSE34014)/Homer | 1e-25 | -5.956e+01 | 0.0000 | 1794.0 | 4.53% | 6110.5 | 3.40% | motif file (matrix) | svg |
| 440 | G C T A A C T G T C G A C T G A C T G A A C G T T A G C C T G A C G T A C G A T | Cux2(Homeobox)/Liver-Cux2-ChIP-Seq(GSE35985)/Homer | 1e-25 | -5.799e+01 | 0.0000 | 6427.0 | 16.23% | 25449.2 | 14.16% | motif file (matrix) | svg |
| 441 | T C A G T C A G A C G T G T A C G C T A T C A G C T G A A C T G A C T G A G C T A G T C C G T A | EAR2(NR)/K562-NR2F6-ChIP-Seq(Encode)/Homer | 1e-24 | -5.721e+01 | 0.0000 | 8556.0 | 21.61% | 34676.6 | 19.29% | motif file (matrix) | svg |
| 442 | C T A G C T G A A G T C G C T A C G A T A C T G G A C T G A T C G A T C C T G A C T A G C T G A T G A C G C T A C G A T T C A G G A C T G A T C G A T C T G A C | p53(p53)/Saos-p53-ChIP-Seq(GSE15780)/Homer | 1e-24 | -5.656e+01 | 0.0000 | 799.0 | 2.02% | 2344.5 | 1.30% | motif file (matrix) | svg |
| 443 | C T A G C T G A A G T C G C T A C G A T A C T G G A C T G A T C G A T C C T G A C T A G C T G A T G A C G C T A C G A T T C A G G A C T G A T C G A T C T G A C | p53(p53)/Saos-p53-ChIP-Seq/Homer | 1e-24 | -5.656e+01 | 0.0000 | 799.0 | 2.02% | 2344.5 | 1.30% | motif file (matrix) | svg |
| 444 | T A C G C G T A T C A G G A C T C T A G A C T G C A G T T A G C T C G A A C G T G T A C C T A G A G T C A G T C G A T C | ZNF669(Zf)/HEK293-ZNF669.GFP-ChIP-Seq(GSE58341)/Homer | 1e-24 | -5.640e+01 | 0.0000 | 1374.0 | 3.47% | 4516.8 | 2.51% | motif file (matrix) | svg |
| 445 | T G C A C G T A G T C A A G C T A G T C G C T A T A G C C G A T C T A G G A T C | Gfi1b(Zf)/HPC7-Gfi1b-ChIP-Seq(GSE22178)/Homer | 1e-24 | -5.626e+01 | 0.0000 | 3774.0 | 9.53% | 14257.2 | 7.93% | motif file (matrix) | svg |
| 446 | T C A G A C T G C A G T A G T C A G T C G T C A C G T A C G T A A C T G C A G T A G T C A G T C C T G A T G C A A G C T | dHNF4(NR)/Fly-HNF4-ChIP-Seq(GSE73675)/Homer | 1e-24 | -5.565e+01 | 0.0000 | 396.0 | 1.00% | 945.8 | 0.53% | motif file (matrix) | svg |
| 447 | C T G A T C A G C A G T C T A G A C T G C T A G G A T C A T C G A C T G C T G A T C A G G A T C | Sp5(Zf)/mES-Sp5.Flag-ChIP-Seq(GSE72989)/Homer | 1e-24 | -5.541e+01 | 0.0000 | 6152.0 | 15.54% | 24354.9 | 13.55% | motif file (matrix) | svg |
| 448 | G C A T G C A T G A C T G C A T T A G C A T G C G A T C C A T G A G T C A G T C | DEL2(E2FDP)/col-DEL2-DAP-Seq(GSE60143)/Homer | 1e-23 | -5.376e+01 | 0.0000 | 5612.0 | 14.17% | 22097.2 | 12.29% | motif file (matrix) | svg |
| 449 | C T A G T A C G G A T C G T C A T G C A A C G T T G C A G C T A T C G A T G C A | Hoxa9(Homeobox)/ChickenMSG-Hoxa9.Flag-ChIP-Seq(GSE86088)/Homer | 1e-22 | -5.262e+01 | 0.0000 | 16335.0 | 41.26% | 69297.5 | 38.56% | motif file (matrix) | svg |
| 450 | C T G A C T G A C T A G T C G A C G T A A T G C C G T A A C T G C G T A A C G T C T G A C G A T A G C T C G T A A C G T A G T C C G A T T A C G G T C A G C A T | GATA(Zf),IR3/iTreg-Gata3-ChIP-Seq(GSE20898)/Homer | 1e-22 | -5.119e+01 | 0.0000 | 1097.0 | 2.77% | 3523.2 | 1.96% | motif file (matrix) | svg |
| 451 | T C A G C T G A C T A G C A T G A C G T A T G C C T G A C T G A C T G A C T A G C A T G A C G T A T G C C T G A | TR4(NR),DR1/Hela-TR4-ChIP-Seq(GSE24685)/Homer | 1e-22 | -5.099e+01 | 0.0000 | 555.0 | 1.40% | 1522.0 | 0.85% | motif file (matrix) | svg |
| 452 | T A C G C T G A T C G A C G A T C T A G C T A G T C G A C T G A T C G A T C G A C G T A T C G A G C A T C A T G C G T A T A C G G C A T T G A C C G T A A G C T | NFAT:AP1(RHD,bZIP)/Jurkat-NFATC1-ChIP-Seq(Jolma\_et\_al.)/Homer | 1e-21 | -5.055e+01 | 0.0000 | 754.0 | 1.90% | 2243.7 | 1.25% | motif file (matrix) | svg |
| 453 | A C G T T G A C A G T C A G C T A G T C A G C T A C T G G A C T A G C T G A C T | REF6(Zf)/Arabidopsis-REF6-ChIP-Seq(GSE106942)/Homer | 1e-21 | -4.972e+01 | 0.0000 | 2904.0 | 7.33% | 10804.1 | 6.01% | motif file (matrix) | svg |
| 454 | G C A T C G A T A T G C A G C T T C G A T A C G G C T A C G T A C A T G T G A C G C A T C G A T A G T C A G C T C G T A | AT3G09735(S1Falike)/col-AT3G09735-DAP-Seq(GSE60143)/Homer | 1e-20 | -4.709e+01 | 0.0000 | 2193.0 | 5.54% | 7944.7 | 4.42% | motif file (matrix) | svg |
| 455 | C G T A C G T A C T G A A C T G A C G T A G T C C G T A C G T A A G T C A C T G A T G C G A T C | WRKY46(WRKY)/colamp-WRKY46-DAP-Seq(GSE60143)/Homer | 1e-20 | -4.654e+01 | 0.0000 | 1116.0 | 2.82% | 3661.4 | 2.04% | motif file (matrix) | svg |
| 456 | T G C A C T G A C A T G C T A G C A G T A G T C C G T A A T G C A T G C T A C G G C A T T C A G G T C A G A T C G T A C | ERE(NR),IR3/MCF7-ERa-ChIP-Seq(Unpublished)/Homer | 1e-20 | -4.636e+01 | 0.0000 | 1703.0 | 4.30% | 5978.3 | 3.33% | motif file (matrix) | svg |
| 457 | C T G A A C G T A C G T A C G T A G T C G A C T C G A T C T G A A C T G C G T A C G T A T C G A | STAT5(Stat)/mCD4+-Stat5-ChIP-Seq(GSE12346)/Homer | 1e-20 | -4.614e+01 | 0.0000 | 1418.0 | 3.58% | 4848.7 | 2.70% | motif file (matrix) | svg |
| 458 | T G A C G C T A T G A C C G T A T C A G G A T C C G T A C A T G C A T G C T A G C T A G C T A G | Unknown-ESC-element(?)/mES-Nanog-ChIP-Seq(GSE11724)/Homer | 1e-19 | -4.563e+01 | 0.0000 | 2452.0 | 6.19% | 9036.3 | 5.03% | motif file (matrix) | svg |
| 459 | C A T G A C G T C T A G A C G T C A G T A C G T C T A G A G T C | PHA-4(Forkhead)/cElegans-Embryos-PHA4-ChIP-Seq(modEncode)/Homer | 1e-19 | -4.412e+01 | 0.0000 | 15560.0 | 39.30% | 66249.6 | 36.86% | motif file (matrix) | svg |
| 460 | C T G A T C A G G C T A A G C T A G T C G A C T C T G A C T A G T G C A C T G A A G T C G T A C G A T C A C T G T C G A | ZBTB12(Zf)/HEK293-ZBTB12.GFP-ChIP-Seq(GSE58341)/Homer | 1e-18 | -4.362e+01 | 0.0000 | 2953.0 | 7.46% | 11159.5 | 6.21% | motif file (matrix) | svg |
| 461 | G T A C A C T G A C G T T C A G G C A T C G T A C G A T G C A T C G T A A G T C C G T A T G A C C A T G G A C T G C T A | ANAC083(NAC)/col-ANAC083-DAP-Seq(GSE60143)/Homer | 1e-18 | -4.324e+01 | 0.0000 | 6977.0 | 17.62% | 28352.8 | 15.77% | motif file (matrix) | svg |
| 462 | G A C T A G T C C G T A C G T A A G T C A G C T A C T G G A C T G T A C A T G C | MYB77(MYB)/col-MYB77-DAP-Seq(GSE60143)/Homer | 1e-18 | -4.323e+01 | 0.0000 | 15699.0 | 39.65% | 66919.5 | 37.23% | motif file (matrix) | svg |
| 463 | A T G C C T G A G A C T A C G T A C G T G T A C G A T C C G A T C T A G C A T G C G T A C G T A C T G A G A C T | STAT1(Stat)/HelaS3-STAT1-ChIP-Seq(GSE12782)/Homer | 1e-18 | -4.299e+01 | 0.0000 | 1335.0 | 3.37% | 4573.7 | 2.54% | motif file (matrix) | svg |
| 464 | A G T C G A C T C A G T G T A C A G T C A T C G T C A G A C T G G T C A C G T A | Stat3(Stat)/mES-Stat3-ChIP-Seq(GSE11431)/Homer | 1e-18 | -4.293e+01 | 0.0000 | 3162.0 | 7.99% | 12050.0 | 6.70% | motif file (matrix) | svg |
| 465 | G A T C G C T A C A G T A C G T T A C G A G T C A T G C C T A G A G T C T C G A | Zfp57(Zf)/H1-ZFP57.HA-ChIP-Seq(GSE115387)/Homer | 1e-18 | -4.273e+01 | 0.0000 | 8079.0 | 20.40% | 33172.2 | 18.46% | motif file (matrix) | svg |
| 466 | G T C A G C A T G C T A C A G T C T A G G A T C C G T A C T G A C G T A C G A T | Oct2(POU,Homeobox)/Bcell-Oct2-ChIP-Seq(GSE21512)/Homer | 1e-18 | -4.257e+01 | 0.0000 | 1316.0 | 3.32% | 4505.5 | 2.51% | motif file (matrix) | svg |
| 467 | T A G C G T A C C G T A C T A G A C T G T G C A C G T A A T G C C G T A A T C G | AR-halfsite(NR)/LNCaP-AR-ChIP-Seq(GSE27824)/Homer | 1e-18 | -4.246e+01 | 0.0000 | 18662.0 | 47.13% | 80302.7 | 44.68% | motif file (matrix) | svg |
| 468 | T G C A A G C T C A T G C G T A A G C T A C T G G A T C G T C A C G T A A G C T | Atf4(bZIP)/MEF-Atf4-ChIP-Seq(GSE35681)/Homer | 1e-18 | -4.226e+01 | 0.0000 | 2557.0 | 6.46% | 9549.6 | 5.31% | motif file (matrix) | svg |
| 469 | A C G T C T A G C G T A A G T C G T A C A C G T A C G T A C G T G T C A G T A C T G A C G A C T | Nur77(NR)/K562-NR4A1-ChIP-Seq(GSE31363)/Homer | 1e-18 | -4.224e+01 | 0.0000 | 1127.0 | 2.85% | 3768.1 | 2.10% | motif file (matrix) | svg |
| 470 | T A C G A T C G T A G C G A T C A C T G A C G T A G T C A C G T C T A G A T C G | Smad4(MAD)/ESC-SMAD4-ChIP-Seq(GSE29422)/Homer | 1e-18 | -4.209e+01 | 0.0000 | 12318.0 | 31.11% | 51886.1 | 28.87% | motif file (matrix) | svg |
| 471 | T C G A A G T C C G T A A T C G T A G C A C G T A C T G A G C T A C G T A G T C | Ptf1a(bHLH)/Panc1-Ptf1a-ChIP-Seq(GSE47459)/Homer | 1e-18 | -4.183e+01 | 0.0000 | 13204.0 | 33.35% | 55840.4 | 31.07% | motif file (matrix) | svg |
| 472 | C G A T C T G A G T A C C A T G G C A T T C A G G C A T C G T A C G T A G C T A C G T A A G T C G C T A G T A C C A T G | CUC2(NAC)/colamp-CUC2-DAP-Seq(GSE60143)/Homer | 1e-17 | -4.137e+01 | 0.0000 | 3649.0 | 9.22% | 14142.3 | 7.87% | motif file (matrix) | svg |
| 473 | G C A T G C A T G A T C G C A T T C G A A C T G C G T A C G T A A T C G T A G C C G A T A C G T A G T C A G C T C G T A | HSF6(HSF)/col-HSF6-DAP-Seq(GSE60143)/Homer | 1e-17 | -4.105e+01 | 0.0000 | 1159.0 | 2.93% | 3912.1 | 2.18% | motif file (matrix) | svg |
| 474 | C T G A T A G C T G A C T C A G C T A G G T C A C G T A T C A G A G C T T C A G | ETV4(ETS)/HepG2-ETV4-ChIP-Seq(ENCODE)/Homer | 1e-17 | -4.069e+01 | 0.0000 | 10106.0 | 25.52% | 42163.5 | 23.46% | motif file (matrix) | svg |
| 475 | A G T C G A C T A C T G G A T C G T A C C G T A T G A C A G T C C G A T A G C T A C G T A C G T C T A G G A C T C T G A | ZNF7(Zf)/HepG2-ZNF7.Flag-ChIP-Seq(Encode)/Homer | 1e-17 | -4.052e+01 | 0.0000 | 3314.0 | 8.37% | 12755.9 | 7.10% | motif file (matrix) | svg |
| 476 | C G A T C T G A G T A C A C T G A C G T T C A G G C A T C G T A C G T A G C A T C G T A A G T C C G T A G T A C C A T G | CUC3(NAC)/col-CUC3-DAP-Seq(GSE60143)/Homer | 1e-17 | -4.028e+01 | 0.0000 | 3602.0 | 9.10% | 13976.7 | 7.78% | motif file (matrix) | svg |
| 477 | C G T A T C G A G A T C G C A T C G T A A C G T G T A C T C A G G T C A G A C T C G T A C T A G | DREF/Drosophila-Promoters/Homer | 1e-17 | -4.025e+01 | 0.0000 | 680.0 | 1.72% | 2085.3 | 1.16% | motif file (matrix) | svg |
| 478 | A G C T A G T C A G T C A C G T C T A G A C G T A C G T A C G T C G T A A G T C G A T C C G T A | FOXP1(Forkhead)/H9-FOXP1-ChIP-Seq(GSE31006)/Homer | 1e-16 | -3.907e+01 | 0.0000 | 2537.0 | 6.41% | 9546.5 | 5.31% | motif file (matrix) | svg |
| 479 | C G T A C G T A G C A T A C T G C G T A A G C T C T G A C G T A T A C G C T G A | ELT-3(Gata)/cElegans-L1-ELT3-ChIP-Seq(modEncode)/Homer | 1e-16 | -3.898e+01 | 0.0000 | 3131.0 | 7.91% | 12030.3 | 6.69% | motif file (matrix) | svg |
| 480 | G A T C G C A T G C A T A G T C A G C T T C G A T A C G G C T A C G T A C T A G T G A C G C A T C G A T G A T C A G C T | HSFC1(HSF)/col-HSFC1-DAP-Seq(GSE60143)/Homer | 1e-16 | -3.895e+01 | 0.0000 | 897.0 | 2.27% | 2925.1 | 1.63% | motif file (matrix) | svg |
| 481 | T G C A G C A T C G A T C G T A C A G T A C T G G T A C C G T A C T G A A G C T G T C A A C T G C T A G G T C A C G A T A C T G G T A C T G C A C G T A A G C T | CEBP:CEBP(bZIP)/MEF-Chop-ChIP-Seq(GSE35681)/Homer | 1e-16 | -3.889e+01 | 0.0000 | 813.0 | 2.05% | 2604.3 | 1.45% | motif file (matrix) | svg |
| 482 | C A T G G A T C C T G A A G T C C T A G C G T A G C T A G C A T G A T C G A T C A G T C C T A G C G T A C A T G C T A G | PLT1(AP2EREBP)/colamp-PLT1-DAP-Seq(GSE60143)/Homer | 1e-16 | -3.882e+01 | 0.0000 | 1246.0 | 3.15% | 4292.2 | 2.39% | motif file (matrix) | svg |
| 483 | C T A G C A G T C G T A A C G T A G T C A C T G C G T A A G C T A G T C G A T C | HNF6(Homeobox)/Liver-Hnf6-ChIP-Seq(ERP000394)/Homer | 1e-16 | -3.851e+01 | 0.0000 | 8264.0 | 20.87% | 34167.4 | 19.01% | motif file (matrix) | svg |
| 484 | C T A G C T A G A G T C T C A G A C T G A C G T A C G T C T G A | MYB(HTH)/ERMYB-Myb-ChIPSeq(GSE22095)/Homer | 1e-16 | -3.800e+01 | 0.0000 | 19544.0 | 49.36% | 84547.7 | 47.04% | motif file (matrix) | svg |
| 485 | A G T C T A G C A C T G A C G T A C G T C G T A C G T A C A G T C G A T A G T C C T A G A C T G A C G T A C G T C T G A | MYB44(MYB)/colamp-MYB44-DAP-Seq(GSE60143)/Homer | 1e-15 | -3.608e+01 | 0.0000 | 843.0 | 2.13% | 2758.7 | 1.53% | motif file (matrix) | svg |
| 486 | A T G C C G T A A C T G C G T A A C G T G C T A T C G A A G C T C G A T C G T A A C G T A G T C C G A T A C T G G A T C | GATA(Zf),IR4/iTreg-Gata3-ChIP-Seq(GSE20898)/Homer | 1e-15 | -3.594e+01 | 0.0000 | 665.0 | 1.68% | 2081.2 | 1.16% | motif file (matrix) | svg |
| 487 | A G C T G C A T A C T G A C G T A G T C A C G T C T A G T A C G | Smad3(MAD)/NPC-Smad3-ChIP-Seq(GSE36673)/Homer | 1e-15 | -3.580e+01 | 0.0000 | 15685.0 | 39.61% | 67270.9 | 37.43% | motif file (matrix) | svg |
| 488 | T A G C T C A G C A T G G C A T A G C T C G A T A T G C C G T A C G T A G T C A | CHR(?)/Hela-CellCycle-Expression/Homer | 1e-15 | -3.487e+01 | 0.0000 | 2871.0 | 7.25% | 11056.4 | 6.15% | motif file (matrix) | svg |
| 489 | T G C A C G T A A C T G T C A G C A G T C A T G T C A G G A T C T A C G A G T C T G C A A C T G A C T G T G A C G T C A | ZNF165(Zf)/WHIM12-ZNF165-ChIP-Seq(GSE65937)/Homer | 1e-14 | -3.453e+01 | 0.0000 | 713.0 | 1.80% | 2281.5 | 1.27% | motif file (matrix) | svg |
| 490 | T A C G A C T G A G C T G T A C C G T A T C G A C T G A A C T G C A T G A C G T A G T C C G T A | COUP-TFII(NR)/K562-NR2F1-ChIP-Seq(Encode)/Homer | 1e-14 | -3.384e+01 | 0.0000 | 9106.0 | 23.00% | 38092.6 | 21.19% | motif file (matrix) | svg |
| 491 | G C T A G C A T G C T A G C A T G C A T C G T A C G T A A G T C A G T C A C T G G C A T G C A T C G T A G C T A G C T A | MYB73(MYB)/col-MYB73-DAP-Seq(GSE60143)/Homer | 1e-14 | -3.225e+01 | 0.0000 | 15105.0 | 38.15% | 64881.0 | 36.10% | motif file (matrix) | svg |
| 492 | T G C A A G C T A C G T C T A G G A T C C T A G G A T C G T C A C T G A A G T C | CEBP(bZIP)/ThioMac-CEBPb-ChIP-Seq(GSE21512)/Homer | 1e-13 | -3.171e+01 | 0.0000 | 5961.0 | 15.05% | 24419.2 | 13.59% | motif file (matrix) | svg |
| 493 | C G A T G A C T C G A T T C A G G A C T A C G T C A G T C T G A G A C T G A C T A G C T C G A T A C T G A T C G G T A C G C T A | NF1:FOXA1(CTF,Forkhead)/LNCAP-FOXA1-ChIP-Seq(GSE27824)/Homer | 1e-13 | -3.147e+01 | 0.0000 | 273.0 | 0.69% | 705.8 | 0.39% | motif file (matrix) | svg |
| 494 | C T A G T C G A A C G T A C G T C A T G A G T C C T G A C G A T A G T C C G T A | AARE(HLH)/mES-cMyc-ChIP-Seq/Homer | 1e-13 | -3.127e+01 | 0.0000 | 894.0 | 2.26% | 3030.6 | 1.69% | motif file (matrix) | svg |
| 495 | T C G A G A C T T C A G T G C A G T A C G T A C A G C T G T A C C A T G T C G A C A T G C A T G A C G T A G T C C T G A | FXR(NR),ER2/Liver-FXR-ChIP-Seq(GSE133700)/Homer | 1e-13 | -3.109e+01 | 0.0000 | 3459.0 | 8.74% | 13664.9 | 7.60% | motif file (matrix) | svg |
| 496 | T G C A C T A G C T A G C T G A C A T G A C T G T G C A G A T C G T C A T G C A G T C A G T C A A G C T C T A G G C A T | ZNF675(Zf)/HEK293-ZNF675.GFP-ChIP-Seq(GSE58341)/Homer | 1e-12 | -2.992e+01 | 0.0000 | 942.0 | 2.38% | 3242.4 | 1.80% | motif file (matrix) | svg |
| 497 | C G T A C T A G G A C T G T C A G T C A C G T A A G T C C G T A T C G A T C G A T C G A C G T A C T G A C T A G G C T A C G T A T A G C C G T A C G A T C G T A | FOXA1:AR(Forkhead,NR)/LNCAP-AR-ChIP-Seq(GSE27824)/Homer | 1e-12 | -2.959e+01 | 0.0000 | 198.0 | 0.50% | 468.9 | 0.26% | motif file (matrix) | svg |
| 498 | C A T G G T C A A G T C C G T A C T A G G A T C C G A T A C T G A C G T G T A C C G T A C G T A | bZIP69(bZIP)/col-bZIP69-DAP-Seq(GSE60143)/Homer | 1e-12 | -2.934e+01 | 0.0000 | 651.0 | 1.64% | 2115.1 | 1.18% | motif file (matrix) | svg |
| 499 | G T A C C G T A C G T A T A C G G C A T G T A C C G T A C A T G A G T C C G T A C G T A C G A T G C A T G C A T G A C T | MafF(bZIP)/HepG2-MafF-ChIP-Seq(GSE31477)/Homer | 1e-12 | -2.906e+01 | 0.0000 | 1355.0 | 3.42% | 4918.3 | 2.74% | motif file (matrix) | svg |
| 500 | C G T A C G T A T C G A C G T A C G A T C G T A A C G T A G T C G C A T G C A T | At3g09600(MYBrelated)/colamp-At3g09600-DAP-Seq(GSE60143)/Homer | 1e-12 | -2.855e+01 | 0.0000 | 2276.0 | 5.75% | 8742.4 | 4.86% | motif file (matrix) | svg |
| 501 | A C T G G A C T A G T C C T G A G A T C T C A G A T G C G A C T A G T C A T G C T A G C A G C T A T C G T G C A | PAX5(Paired,Homeobox),condensed/GM12878-PAX5-ChIP-Seq(GSE32465)/Homer | 1e-12 | -2.845e+01 | 0.0000 | 1136.0 | 2.87% | 4044.4 | 2.25% | motif file (matrix) | svg |
| 502 | C T A G C A T G C A T G T A C G A G T C G C A T A G C T C T A G A C G T A G T C G A C T A C T G A C T G A C T G T C G A | Zfp809(Zf)/ES-Zfp809-ChIP-Seq(GSE70799)/Homer | 1e-12 | -2.822e+01 | 0.0000 | 1063.0 | 2.68% | 3755.1 | 2.09% | motif file (matrix) | svg |
| 503 | C G A T C A G T C A G T C A T G G T C A G A T C C G T A T C A G A G T C A C G T C T A G A C G T G T A C G T C A G C T A | VIP1(bZIP)/col-VIP1-DAP-Seq(GSE60143)/Homer | 1e-12 | -2.784e+01 | 0.0000 | 998.0 | 2.52% | 3502.9 | 1.95% | motif file (matrix) | svg |
| 504 | C G T A C T G A C T A G C G T A C G T A A G T C C G T A C A G T G C A T G T C A C G A T A C T G A C G T G C A T G A T C | PGR(NR)/EndoStromal-PGR-ChIP-Seq(GSE69539)/Homer | 1e-11 | -2.753e+01 | 0.0000 | 1217.0 | 3.07% | 4391.0 | 2.44% | motif file (matrix) | svg |
| 505 | C T A G C T A G T C G A C G T A A T G C C G T A A T C G T C G A T A C G G C A T A C T G C A G T T A G C G A T C G A C T | MRE(NR)/Neuro2A-NR3C2-ChIPnexus(GSE115417)/Homer | 1e-11 | -2.703e+01 | 0.0000 | 7859.0 | 19.85% | 32956.4 | 18.34% | motif file (matrix) | svg |
| 506 | G C T A G C A T G A C T G C A T T C A G G T A C G C T A G C A T C T G A G C T A T A G C G C T A C T G A C G A T C T A G | OCT4-SOX2-TCF-NANOG(POU,Homeobox,HMG)/mES-Oct4-ChIP-Seq(GSE11431)/Homer | 1e-11 | -2.691e+01 | 0.0000 | 549.0 | 1.39% | 1758.3 | 0.98% | motif file (matrix) | svg |
| 507 | G C T A C T G A C G T A C G T A C T G A C T G A C G A T G T C A A C G T A G T C G C A T G C A T | At5g52660(MYBrelated)/colamp-At5g52660-DAP-Seq(GSE60143)/Homer | 1e-11 | -2.681e+01 | 0.0000 | 1985.0 | 5.01% | 7574.4 | 4.21% | motif file (matrix) | svg |
| 508 | A T G C A G C T T C A G T G A C T C A G A T G C T G C A A C G T A T C G G A T C A C T G A G T C | NRF1(NRF)/MCF7-NRF1-ChIP-Seq(Unpublished)/Homer | 1e-11 | -2.663e+01 | 0.0000 | 801.0 | 2.02% | 2741.8 | 1.53% | motif file (matrix) | svg |
| 509 | C T A G T A C G A G T C C G T A A G T C A C G T A G T C T C G A C G T A T A C G | Nkx2.1(Homeobox)/LungAC-Nkx2.1-ChIP-Seq(GSE43252)/Homer | 1e-11 | -2.642e+01 | 0.0000 | 17864.0 | 45.12% | 77689.3 | 43.22% | motif file (matrix) | svg |
| 510 | G A C T A C G T C G A T A G C T A G T C C G T A A C T G A C T G C G A T C T A G | NGA4(ABI3VP1)/col-NGA4-DAP-Seq(GSE60143)/Homer | 1e-11 | -2.613e+01 | 0.0000 | 14084.0 | 35.57% | 60694.2 | 33.77% | motif file (matrix) | svg |
| 511 | T A G C G A T C A G C T T G A C G C T A A G C T C A T G A C T G A C G T T C A G A G T C G A T C G A C T A G C T G C T A A G T C A G C T A G T C G A T C A T G C A G C T G A C T C A T G A C G T A T C G | ZNF41(Zf)/HEK293-ZNF41.GFP-ChIP-Seq(GSE58341)/Homer | 1e-11 | -2.550e+01 | 0.0000 | 177.0 | 0.45% | 426.7 | 0.24% | motif file (matrix) | svg |
| 512 | A G T C C T A G A T C G C A G T C G A T A G C T G T A C A C T G C A T G C A T G | ZBED2(Zf)/SUIT2-ZBED2.HA-ChIP-Seq(GSE141606)/Homer | 1e-11 | -2.536e+01 | 0.0000 | 8520.0 | 21.52% | 35965.2 | 20.01% | motif file (matrix) | svg |
| 513 | G A C T G A C T A T C G C G A T G T C A G T A C A G C T C G A T A C G T G T A C | SPL11(SBP)/col100-SPL11-DAP-Seq(GSE60143)/Homer | 1e-10 | -2.533e+01 | 0.0000 | 5478.0 | 13.84% | 22609.4 | 12.58% | motif file (matrix) | svg |
| 514 | G C T A C G T A A C G T A T C G C G T A A C G T A C G T C T A G | ATHB6(Homeobox)/col-ATHB6-DAP-Seq(GSE60143)/Homer | 1e-10 | -2.511e+01 | 0.0000 | 8426.0 | 21.28% | 35564.7 | 19.79% | motif file (matrix) | svg |
| 515 | G A T C G C A T A C T G C T A G C T G A A G C T G C T A G C T A C T G A T C A G G C A T T G C A A C G T A C G T G A T C G A C T G C A T C T A G T A C G G A C T C T A G C A T G C T A G G T A C T C G A | ZNF136(Zf)/HEK293-ZNF136.GFP-ChIP-Seq(GSE58341)/Homer | 1e-10 | -2.501e+01 | 0.0000 | 461.0 | 1.16% | 1449.2 | 0.81% | motif file (matrix) | svg |
| 516 | C T A G A T C G G T C A C A T G A G T C G A C T T A C G C A G T A G T C A G T C C T G A C G A T C T A G A T C G G A C T A T C G A G T C G A C T C T A G T C G A | REST-NRSF(Zf)/Jurkat-NRSF-ChIP-Seq/Homer | 1e-10 | -2.484e+01 | 0.0000 | 44.0 | 0.11% | 45.6 | 0.03% | motif file (matrix) | svg |
| 517 | A C T G A G T C G T C A C G T A A G T C C G T A C T A G C T A G G A C T C A T G | SCRT1(Zf)/HEK293-SCRT1.eGFP-ChIP-Seq(Encode)/Homer | 1e-10 | -2.440e+01 | 0.0000 | 2602.0 | 6.57% | 10248.3 | 5.70% | motif file (matrix) | svg |
| 518 | G C A T G C A T C T G A A C G T C T G A A C G T C G T A C G T A C G T A A G T C G T C A G T C A | Foxf1(Forkhead)/Lung-Foxf1-ChIP-Seq(GSE77951)/Homer | 1e-10 | -2.389e+01 | 0.0000 | 4273.0 | 10.79% | 17440.8 | 9.70% | motif file (matrix) | svg |
| 519 | T C G A A C T G A C T G C G T A C G T A T C G A A G T C C T G A A T C G G T A C G C A T C A T G | ETS:E-box(ETS,bHLH)/HPC7-Scl-ChIP-Seq(GSE22178)/Homer | 1e-10 | -2.327e+01 | 0.0000 | 374.0 | 0.94% | 1144.6 | 0.64% | motif file (matrix) | svg |
| 520 | C T G A T G C A T A G C T G A C T A C G T C A G C T G A G C T A T C A G G A C T | ELF1(ETS)/Jurkat-ELF1-ChIP-Seq(SRA014231)/Homer | 1e-10 | -2.327e+01 | 0.0000 | 6040.0 | 15.25% | 25168.3 | 14.00% | motif file (matrix) | svg |
| 521 | C T A G A G T C A G C T A C T G C G T A C A G T C G T A C T G A T A G C T G A C | Unknown5/Drosophila-Promoters/Homer | 1e-10 | -2.313e+01 | 0.0000 | 6398.0 | 16.16% | 26745.2 | 14.88% | motif file (matrix) | svg |
| 522 | T G A C T C G A C T G A C T G A A T G C G A T C C T A G T A C G G A C T G A C T G A T C T C G A C T G A C T G A A T G C G A T C C T A G A T C G G A C T G A C T | Tcfcp2l1(CP2)/mES-Tcfcp2l1-ChIP-Seq(GSE11431)/Homer | 1e-9 | -2.244e+01 | 0.0000 | 1219.0 | 3.08% | 4512.5 | 2.51% | motif file (matrix) | svg |
| 523 | C G T A C G T A C T G A A C T G C G T A C G T A A C G T G T C A A C G T G C A T A G T C G A C T | At2g03500(G2like)/col-At2g03500-DAP-Seq(GSE60143)/Homer | 1e-9 | -2.232e+01 | 0.0000 | 1947.0 | 4.92% | 7546.9 | 4.20% | motif file (matrix) | svg |
| 524 | C G T A C G T A C G T A C G T A C G T A A C T G A G C T C T A G G T A C G C T A | AT1G69570(C2C2dof)/col-AT1G69570-DAP-Seq(GSE60143)/Homer | 1e-9 | -2.207e+01 | 0.0000 | 7489.0 | 18.91% | 31606.9 | 17.59% | motif file (matrix) | svg |
| 525 | C G T A A C T G G T C A A C G T A T C G C A G T C T A G T C A G C G T A A C T G C G T A A C G T C G T A C T G A T A C G | GATA3(Zf),DR4/iTreg-Gata3-ChIP-Seq(GSE20898)/Homer | 1e-9 | -2.206e+01 | 0.0000 | 603.0 | 1.52% | 2038.2 | 1.13% | motif file (matrix) | svg |
| 526 | C G T A C T G A T C A G A C T G G T C A C G T A A C G T G T A C C G A T G C A T | AT5G45580(G2like)/colamp-AT5G45580-DAP-Seq(GSE60143)/Homer | 1e-9 | -2.179e+01 | 0.0000 | 11234.0 | 28.37% | 48246.1 | 26.84% | motif file (matrix) | svg |
| 527 | C G A T T C G A A C T G G T C A C G T A C G A T G T A C G A C T | At3g04030(G2like)/col-At3g04030-DAP-Seq(GSE60143)/Homer | 1e-9 | -2.165e+01 | 0.0000 | 6473.0 | 16.35% | 27156.5 | 15.11% | motif file (matrix) | svg |
| 528 | T G A C A T G C C G T A A T C G A T G C C A G T C A T G A C T G A G T C G T A C | HEB(bHLH)/mES-Heb-ChIP-Seq(GSE53233)/Homer | 1e-9 | -2.119e+01 | 0.0000 | 9612.0 | 24.28% | 41064.1 | 22.85% | motif file (matrix) | svg |
| 529 | G C A T C G T A C G A T A C T G A G T C G C T A C T G A C G T A C A G T A C T G C G T A T C A G | Oct6(POU,Homeobox)/NPC-Pou3f1-ChIP-Seq(GSE35496)/Homer | 1e-9 | -2.103e+01 | 0.0000 | 1880.0 | 4.75% | 7304.4 | 4.06% | motif file (matrix) | svg |
| 530 | C G T A C G A T C G T A T C G A T C G A A C G T C G T A A C G T A G T C G C A T | LHY(Myb)/Seedling-LHY-ChIP-Seq(GSE52175)/Homer | 1e-9 | -2.082e+01 | 0.0000 | 6893.0 | 17.41% | 29052.0 | 16.16% | motif file (matrix) | svg |
| 531 | C G A T T C G A G T A C A C T G A C G T T C A G G C A T T G C A G C T A G C A T C G T A A G T C C G T A G T A C C A T G | ANAC087(NAC)/col-ANAC087-DAP-Seq(GSE60143)/Homer | 1e-9 | -2.075e+01 | 0.0000 | 3540.0 | 8.94% | 14408.8 | 8.02% | motif file (matrix) | svg |
| 532 | A G C T G A C T A C T G A C G T G T C A A G T C A C G T C G A T | SPL9(SBP)/colamp-SPL9-DAP-Seq(GSE60143)/Homer | 1e-8 | -2.065e+01 | 0.0000 | 19827.0 | 50.07% | 87013.7 | 48.41% | motif file (matrix) | svg |
| 533 | T A G C C G A T T A C G A C T G A G T C A C T G A T C G A T C G C G T A C T G A | E2F1(E2F)/Hela-E2F1-ChIP-Seq(GSE22478)/Homer | 1e-8 | -2.056e+01 | 0.0000 | 3216.0 | 8.12% | 13019.0 | 7.24% | motif file (matrix) | svg |
| 534 | T G A C C T G A C T A G C T G A C G T A A G T C C T G A A C G T G C A T T A G C G C A T A T C G G A C T G A C T G A T C | GRE(NR),IR3/RAW264.7-GRE-ChIP-Seq(Unpublished)/Homer | 1e-8 | -2.049e+01 | 0.0000 | 1532.0 | 3.87% | 5860.5 | 3.26% | motif file (matrix) | svg |
| 535 | G A C T A G C T G T A C G A C T C T G A A C T G G T C A C T G A A T G C T A C G G A C T A C G T A G T C G A C T C T G A | HRE(HSF)/Striatum-HSF1-ChIP-Seq(GSE38000)/Homer | 1e-8 | -2.021e+01 | 0.0000 | 766.0 | 1.93% | 2718.0 | 1.51% | motif file (matrix) | svg |
| 536 | A T G C G A T C C G A T C T A G A C T G G C T A C G T A A G C T A C T G A G C T | TEAD2(TEA)/Py2T-Tead2-ChIP-Seq(GSE55709)/Homer | 1e-8 | -2.015e+01 | 0.0000 | 2978.0 | 7.52% | 12012.1 | 6.68% | motif file (matrix) | svg |
| 537 | C T A G T G A C G A C T A T C G T C G A A G T C C T A G C A G T C T A G A T C G G T A C T C G A | O2(bZIP)/Corn-O2-ChIP-Seq(GSE63991)/Homer | 1e-8 | -2.000e+01 | 0.0000 | 1564.0 | 3.95% | 6008.1 | 3.34% | motif file (matrix) | svg |
| 538 | G C A T G C A T G A T C G A C T T C G A T C A G G C T A C G T A A C T G G T A C G C A T G C A T A G T C A G C T C G T A | HSF7(HSF)/colamp-HSF7-DAP-Seq(GSE60143)/Homer | 1e-8 | -1.998e+01 | 0.0000 | 786.0 | 1.99% | 2804.0 | 1.56% | motif file (matrix) | svg |
| 539 | C G T A G A C T C G T A A C G T C A G T A G T C A G C T G A C T | KAN2(G2like)/colamp-KAN2-DAP-Seq(GSE60143)/Homer | 1e-8 | -1.986e+01 | 0.0000 | 8152.0 | 20.59% | 34674.7 | 19.29% | motif file (matrix) | svg |
| 540 | C G T A C T G A C T A G C T G A A G T C G C T A C G A T A T C G G A C T G A T C A G T C C T G A C T A G C T A G A G T C G C T A C G A T C T A G G A T C G A T C | p73(p53)/Trachea-p73-ChIP-Seq(PRJNA310161)/Homer | 1e-8 | -1.963e+01 | 0.0000 | 346.0 | 0.87% | 1083.7 | 0.60% | motif file (matrix) | svg |
| 541 | A G T C G C A T C G T A C G T A G T A C A C G T A C T G G A T C G A T C T C G A | BMYB(HTH)/Hela-BMYB-ChIP-Seq(GSE27030)/Homer | 1e-8 | -1.946e+01 | 0.0000 | 15501.0 | 39.15% | 67555.9 | 37.59% | motif file (matrix) | svg |
| 542 | C G T A A T G C C G A T A C G T A G T C C G T A C G T A C G T A C T A G A T C G | TCFL2(HMG)/K562-TCF7L2-ChIP-Seq(GSE29196)/Homer | 1e-8 | -1.941e+01 | 0.0000 | 504.0 | 1.27% | 1693.4 | 0.94% | motif file (matrix) | svg |
| 543 | G A T C G T A C C G T A A G C T G A C T G C T A C T G A A C G T G A T C G C T A | Hoxc6(Homeobox)/EB-Hoxc6.iFlag-ChIP-Seq(GSE142377)/Homer | 1e-8 | -1.937e+01 | 0.0000 | 16299.0 | 41.16% | 71159.1 | 39.59% | motif file (matrix) | svg |
| 544 | C G T A A G T C T G A C A G C T A C G T C G T A A C G T A G T C | At5g05790(MYBrelated)/col-At5g05790-DAP-Seq(GSE60143)/Homer | 1e-8 | -1.889e+01 | 0.0000 | 8684.0 | 21.93% | 37098.0 | 20.64% | motif file (matrix) | svg |
| 545 | G A T C G A T C G C T A G T C A G A C T A T G C T C G A C G A T C G A T C T A G | HAT2(Homeobox)/colamp-HAT2-DAP-Seq(GSE60143)/Homer | 1e-8 | -1.878e+01 | 0.0000 | 5941.0 | 15.00% | 24982.3 | 13.90% | motif file (matrix) | svg |
| 546 | T A C G T A C G G T A C A T C G T A C G T A C G G T C A C T G A C G T A G A C T | E2F4(E2F)/K562-E2F4-ChIP-Seq(GSE31477)/Homer | 1e-8 | -1.878e+01 | 0.0000 | 7998.0 | 20.20% | 34063.9 | 18.95% | motif file (matrix) | svg |
| 547 | G C A T G C A T G T A C G A C T T C G A A C T G G C T A C G T A A T C G T G A C G C A T G A C T A G T C A G C T C T G A | AGL95(ND)/col-AGL95-DAP-Seq(GSE60143)/Homer | 1e-8 | -1.853e+01 | 0.0000 | 435.0 | 1.10% | 1438.0 | 0.80% | motif file (matrix) | svg |
| 548 | G C A T A G T C G A C T T C G A T A C G G T C A T C G A A C T G T A G C G C A T G C A T A T G C | AT2G01818(PLATZ)/col-AT2G01818-DAP-Seq(GSE60143)/Homer | 1e-7 | -1.752e+01 | 0.0000 | 864.0 | 2.18% | 3172.5 | 1.77% | motif file (matrix) | svg |
| 549 | C G T A G A C T C G A T A T C G G T A C G C A T C A T G C G T A T A C G G C A T G T A C C G T A C A T G A T G C G C T A C T A G G C A T G C A T G C A T G A C T | MafB(bZIP)/BMM-Mafb-ChIP-Seq(GSE75722)/Homer | 1e-7 | -1.734e+01 | 0.0000 | 1660.0 | 4.19% | 6494.4 | 3.61% | motif file (matrix) | svg |
| 550 | T G C A C T G A A G T C G T C A A C T G A C T G C G T A C G T A C T G A A G C T | EWS:FLI1-fusion(ETS)/SK\_N\_MC-EWS:FLI1-ChIP-Seq(SRA014231)/Homer | 1e-7 | -1.729e+01 | 0.0000 | 3513.0 | 8.87% | 14449.1 | 8.04% | motif file (matrix) | svg |
| 551 | T C G A C A T G C A T G A C G T A T G C T C G A C T G A A G C T T A C G T G C A G T A C G A T C A G C T A G T C | FXR(NR),IR1/Liver-FXR-ChIP-Seq(Chong\_et\_al.)/Homer | 1e-7 | -1.668e+01 | 0.0000 | 2690.0 | 6.79% | 10918.2 | 6.07% | motif file (matrix) | svg |
| 552 | G C A T C T A G C T A G C G T A A G C T C G T A C T G A C A T G C T A G G C A T | AT5G56840(MYBrelated)/colamp-AT5G56840-DAP-Seq(GSE60143)/Homer | 1e-7 | -1.666e+01 | 0.0000 | 9469.0 | 23.91% | 40750.8 | 22.67% | motif file (matrix) | svg |
| 553 | C T A G A T G C A T G C C G A T A C T G G A C T A T G C G C T A T G A C A G C T T A G C G C T A | PBX1(Homeobox)/MCF7-PBX1-ChIP-Seq(GSE28007)/Homer | 1e-7 | -1.618e+01 | 0.0000 | 357.0 | 0.90% | 1171.8 | 0.65% | motif file (matrix) | svg |
| 554 | A T G C T A G C A G C T A G C T T G A C G A C T T C A G T A C G G T C A C T G A A T C G T A G C G A C T C A G T A G T C A G C T T C G A A T C G T G C A T G C A | HRE(HSF)/HepG2-HSF1-ChIP-Seq(GSE31477)/Homer | 1e-7 | -1.617e+01 | 0.0000 | 663.0 | 1.67% | 2384.5 | 1.33% | motif file (matrix) | svg |
| 555 | G C T A C G T A A C T G C G T A C G A T A C G T A G T C A G C T | At3g12730(G2like)/colamp-At3g12730-DAP-Seq(GSE60143)/Homer | 1e-6 | -1.594e+01 | 0.0000 | 10591.0 | 26.75% | 45813.1 | 25.49% | motif file (matrix) | svg |
| 556 | C T A G A C G T A G T C C G T A A C T G A G T C G C A T A C T G G C A T A G T C G A C T G A T C G C A T A G T C A G C T | ZNF317(Zf)/HEK293-ZNF317.GFP-ChIP-Seq(GSE58341)/Homer | 1e-6 | -1.536e+01 | 0.0000 | 589.0 | 1.49% | 2102.3 | 1.17% | motif file (matrix) | svg |
| 557 | T C G A T C A G T C G A A C T G C A T G A C G T A G T C C T G A | COUP-TFII(NR)/Artia-Nr2f2-ChIP-Seq(GSE46497)/Homer | 1e-6 | -1.517e+01 | 0.0000 | 11211.0 | 28.31% | 48650.3 | 27.07% | motif file (matrix) | svg |
| 558 | T C G A A C G T A C T G C T G A A G T C T C A G A G C T G T A C C G T A A G C T G A T C T C G A | JunD(bZIP)/K562-JunD-ChIP-Seq/Homer | 1e-6 | -1.516e+01 | 0.0000 | 465.0 | 1.17% | 1610.7 | 0.90% | motif file (matrix) | svg |
| 559 | A C T G C G T A A C G T C G T A C T G A A C T G T C A G G C A T | At3g11280(MYBrelated)/col-At3g11280-DAP-Seq(GSE60143)/Homer | 1e-6 | -1.512e+01 | 0.0000 | 8303.0 | 20.97% | 35677.7 | 19.85% | motif file (matrix) | svg |
| 560 | C T G A C T A G A T C G A G C T A C T G G A C T A G T C C T G A | Tbx5(T-box)/HL1-Tbx5.biotin-ChIP-Seq(GSE21529)/Homer | 1e-6 | -1.510e+01 | 0.0000 | 18345.0 | 46.33% | 80784.9 | 44.95% | motif file (matrix) | svg |
| 561 | C T G A G T A C G A C T A G T C C A G T T G C A C T G A A C G T A G C T G A T C C T A G C G A T A C T G A T G C G A C T C T G A G A T C G A C T A G C T G A T C | Mouse\_Recombination\_Hotspot(Zf)/Testis-DMC1-ChIP-Seq(GSE24438)/Homer | 1e-6 | -1.491e+01 | 0.0000 | 374.0 | 0.94% | 1256.5 | 0.70% | motif file (matrix) | svg |
| 562 | A C T G G A T C C T G A A T C G A G T C T A G C C T G A C G T A T A C G A G T C C T A G C A G T T C A G T C G A T G A C G A T C | PAX5(Paired,Homeobox)/GM12878-PAX5-ChIP-Seq(GSE32465)/Homer | 1e-6 | -1.490e+01 | 0.0000 | 3406.0 | 8.60% | 14103.9 | 7.85% | motif file (matrix) | svg |
| 563 | A G T C C G T A C G A T A G T C G T C A A G T C A C G T C T G A | Unknown2/Drosophila-Promoters/Homer | 1e-6 | -1.486e+01 | 0.0000 | 6979.0 | 17.63% | 29819.2 | 16.59% | motif file (matrix) | svg |
| 564 | G C A T G A T C A G C T G T A C G A T C C T A G C T A G G A T C T A C G C T G A | AT3G58630(Trihelix)/col-AT3G58630-DAP-Seq(GSE60143)/Homer | 1e-6 | -1.478e+01 | 0.0000 | 2865.0 | 7.24% | 11760.9 | 6.54% | motif file (matrix) | svg |
| 565 | T C G A C G T A A G T C A G C T C G T A A G T C T C G A G C T A G A C T C G A T A G T C A G T C A G T C C T G A T C A G T G C A T C G A C A G T A T C G A G T C | GFY-Staf(?,Zf)/Promoter/Homer | 1e-6 | -1.462e+01 | 0.0000 | 225.0 | 0.57% | 690.8 | 0.38% | motif file (matrix) | svg |
| 566 | C G A T C A G T G T C A G C T A C A G T A G C T A C G T A C T G A G T C C G T A A C G T A C T G A C G T C T G A T C G A | FUS3(ABI3VP1)/col-FUS3-DAP-Seq(GSE60143)/Homer | 1e-6 | -1.448e+01 | 0.0000 | 6688.0 | 16.89% | 28557.3 | 15.89% | motif file (matrix) | svg |
| 567 | T A G C C G A T A C G T A G C T A G C T A G T C A T G C A G T C A C T G A T G C A T G C G C T A | E2F7(E2F)/Hela-E2F7-ChIP-Seq(GSE32673)/Homer | 1e-6 | -1.440e+01 | 0.0000 | 1832.0 | 4.63% | 7330.0 | 4.08% | motif file (matrix) | svg |
| 568 | A T G C G A C T A G C T A G C T A G T C G C T A C A G T C G A T G C T A A C G T A C T G G C T A T A G C G C A T T G A C | IRF:BATF(IRF:bZIP)/pDC-Irf8-ChIP-Seq(GSE66899)/Homer | 1e-6 | -1.392e+01 | 0.0000 | 429.0 | 1.08% | 1490.0 | 0.83% | motif file (matrix) | svg |
| 569 | G A T C G T A C C G A T A C T G A C T G C G T A C G T A A C G T A C T G G A T C | TEAD(TEA)/Fibroblast-PU.1-ChIP-Seq(Unpublished)/Homer | 1e-5 | -1.368e+01 | 0.0000 | 3262.0 | 8.24% | 13540.2 | 7.53% | motif file (matrix) | svg |
| 570 | G C A T T A G C G T A C C A T G C T G A G C A T G C A T G C A T G A C T G C A T G A C T G T A C A C T G A T C G C G T A | LBD2(LOBAS2)/colamp-LBD2-DAP-Seq(GSE60143)/Homer | 1e-5 | -1.354e+01 | 0.0000 | 6439.0 | 16.26% | 27522.1 | 15.31% | motif file (matrix) | svg |
| 571 | C G T A C G T A C G T A A C G T C G T A A C G T A G T C G C A T | EPR1(MYBrelated)/colamp-EPR1-DAP-Seq(GSE60143)/Homer | 1e-5 | -1.330e+01 | 0.0000 | 2157.0 | 5.45% | 8769.1 | 4.88% | motif file (matrix) | svg |
| 572 | C G T A C G T A C G T A A C G T C G T A A C G T A G T C G C A T | LHY1(MYBrelated)/col-LHY1-DAP-Seq(GSE60143)/Homer | 1e-5 | -1.330e+01 | 0.0000 | 2157.0 | 5.45% | 8769.1 | 4.88% | motif file (matrix) | svg |
| 573 | A G T C G A T C G C T A C G A T A C G T T A C G G C A T C T G A G A C T A C T G A G T C G C T A C T G A T C G A C A G T | Oct4:Sox17(POU,Homeobox,HMG)/F9-Sox17-ChIP-Seq(GSE44553)/Homer | 1e-5 | -1.313e+01 | 0.0000 | 630.0 | 1.59% | 2316.2 | 1.29% | motif file (matrix) | svg |
| 574 | A T G C G C A T C G A T G A T C A G C T C T G A A C T G C G T A C G T A T C A G T G A C C G A T G C A T G A T C C G A T | HSF21(HSF)/col-HSF21-DAP-Seq(GSE60143)/Homer | 1e-5 | -1.259e+01 | 0.0000 | 384.0 | 0.97% | 1334.7 | 0.74% | motif file (matrix) | svg |
| 575 | T G C A A T G C A C G T A C G T A C G T A T G C C T A G A C G T A C G T A G C T G A T C A G C T | T1ISRE(IRF)/ThioMac-Ifnb-Expression/Homer | 1e-5 | -1.227e+01 | 0.0000 | 82.0 | 0.21% | 201.6 | 0.11% | motif file (matrix) | svg |
| 576 | G C T A G A C T A G T C T C G A T C A G T C G A A C G T A G T C G A C T T C A G | GATA14(C2C2gata)/col-GATA14-DAP-Seq(GSE60143)/Homer | 1e-5 | -1.224e+01 | 0.0000 | 5633.0 | 14.23% | 24050.0 | 13.38% | motif file (matrix) | svg |
| 577 | C G T A C A T G C A T G A C T G C T A G T C G A G C A T C G A T A G C T A G T C G A T C G T A C | NFkB-p65(RHD)/GM12787-p65-ChIP-Seq(GSE19485)/Homer | 1e-5 | -1.216e+01 | 0.0000 | 2721.0 | 6.87% | 11265.4 | 6.27% | motif file (matrix) | svg |
| 578 | G C A T T C G A C T G A G A T C A G T C G A T C G T C A G T C A A C G T A G T C C G T A C T G A | Duxbl(Homeobox)/NIH3T3-Duxbl.HA-ChIP-Seq(GSE119782)/Homer | 1e-5 | -1.214e+01 | 0.0000 | 525.0 | 1.33% | 1910.2 | 1.06% | motif file (matrix) | svg |
| 579 | T G C A T A G C G A C T T G C A T G A C T G C A C G T A A G C T A G C T A G T C A G T C G T A C | GFY(?)/Promoter/Homer | 1e-5 | -1.182e+01 | 0.0000 | 478.0 | 1.21% | 1726.3 | 0.96% | motif file (matrix) | svg |
| 580 | G T A C A G C T T C G A G T A C A G T C C A G T C G T A G T C A G A T C G C A T | MYB62(MYB)/colamp-MYB62-DAP-Seq(GSE60143)/Homer | 1e-5 | -1.171e+01 | 0.0000 | 12550.0 | 31.70% | 54977.3 | 30.59% | motif file (matrix) | svg |
| 581 | T C A G C G T A A G T C A G C T C G T A A G T C C T G A C G T A A G T C G C A T A G T C A G T C A G T C C T G A A C T G T G C A T C G A C A T G A T C G G A T C | Ronin(THAP)/ES-Thap11-ChIP-Seq(GSE51522)/Homer | 1e-4 | -1.148e+01 | 0.0000 | 86.0 | 0.22% | 220.6 | 0.12% | motif file (matrix) | svg |
| 582 | T C G A G T A C T C G A T C G A C A T G A T G C A C G T A C T G A C T G A G T C C G T A C T A G A G T C A T C G A G T C | Unknown3/Drosophila-Promoters/Homer | 1e-4 | -1.126e+01 | 0.0000 | 668.0 | 1.69% | 2518.6 | 1.40% | motif file (matrix) | svg |
| 583 | G C T A T A G C A G C T A T C G G T C A C G T A G C T A A T G C G A T C C T G A | IRF4(IRF)/GM12878-IRF4-ChIP-Seq(GSE32465)/Homer | 1e-4 | -1.114e+01 | 0.0000 | 3154.0 | 7.97% | 13210.8 | 7.35% | motif file (matrix) | svg |
| 584 | T G C A A G C T C T G A A T C G G A C T C T A G G T A C G A T C G T C A A G T C G T A C G A C T C T A G A T C G G C A T C A T G C A T G G A T C G T A C C T G A | CTCF(Zf)/CD4+-CTCF-ChIP-Seq(Barski\_et\_al.)/Homer | 1e-4 | -1.086e+01 | 0.0000 | 471.0 | 1.19% | 1718.3 | 0.96% | motif file (matrix) | svg |
| 585 | G C T A A G C T G T A C G C A T A G C T T C G A C T G A A G T C A G T C T A C G A C G T G A C T T A C G C T A G C G T A | ZML1(C2C2gata)/colamp-ZML1-DAP-Seq(GSE60143)/Homer | 1e-4 | -1.082e+01 | 0.0000 | 421.0 | 1.06% | 1516.2 | 0.84% | motif file (matrix) | svg |
| 586 | A T G C A C T G C G A T T C A G A T G C C G T A C T G A T G C A C T G A G A C T A C T G G T C A | ABF1/SacCer-Promoters/Homer | 1e-4 | -1.057e+01 | 0.0000 | 5761.0 | 14.55% | 24744.3 | 13.77% | motif file (matrix) | svg |
| 587 | C G T A C A G T G T A C A T G C C T A G C G T A A C G T A G T C T C G A T C A G | GATA19(C2C2gata)/colamp-GATA19-DAP-Seq(GSE60143)/Homer | 1e-4 | -1.039e+01 | 0.0001 | 1416.0 | 3.58% | 5709.0 | 3.18% | motif file (matrix) | svg |
| 588 | C G A T C G T A A C T G C G T A A C G T C G T A A C G T A C G T C G A T G C A T C G A T C G A T | AT2G28920(ND)/col-AT2G28920-DAP-Seq(GSE60143)/Homer | 1e-4 | -1.035e+01 | 0.0001 | 1121.0 | 2.83% | 4450.2 | 2.48% | motif file (matrix) | svg |
| 589 | C A T G A G T C C T G A A T G C C T A G T C G A G C T A G C A T G A T C G A C T A G T C C T A G C G T A C A T G C T A G | PLT3(AP2EREBP)/col-PLT3-DAP-Seq(GSE60143)/Homer | 1e-4 | -1.015e+01 | 0.0001 | 942.0 | 2.38% | 3698.3 | 2.06% | motif file (matrix) | svg |
| 590 | C T G A C T A G T C G A C G T A A T G C C G T A A T C G C G A T T A G C G C A T A T C G G C A T A G C T G A T C G A C T A G C T | ARE(NR)/LNCAP-AR-ChIP-Seq(GSE27824)/Homer | 1e-4 | -1.011e+01 | 0.0001 | 1114.0 | 2.81% | 4429.9 | 2.46% | motif file (matrix) | svg |
| 591 | C T G A C T G A G C A T G A C T A G T C T C G A C T A G C G T A A C G T G A T C A G C T T C A G | GATA4(C2C2gata)/col-GATA4-DAP-Seq(GSE60143)/Homer | 1e-4 | -1.008e+01 | 0.0001 | 6868.0 | 17.35% | 29706.0 | 16.53% | motif file (matrix) | svg |
| 592 | T A G C A G T C T G A C A G T C C T A G A T C G A G T C C A T G T G A C A G T C G T A C A G T C A G T C G C A T C T A G A T C G G C A T A C T G A T C G G A T C | BORIS(Zf)/K562-CTCFL-ChIP-Seq(GSE32465)/Homer | 1e-4 | -9.889e+00 | 0.0001 | 771.0 | 1.95% | 2986.7 | 1.66% | motif file (matrix) | svg |
| 593 | C T A G T A C G G A T C G T C A T C G A A C G T T C A G C G T A C G T A C G T A | Hoxd10(Homeobox)/ChickenMSG-Hoxd10.Flag-ChIP-Seq(GSE86088)/Homer | 1e-4 | -9.767e+00 | 0.0001 | 6758.0 | 17.07% | 29243.3 | 16.27% | motif file (matrix) | svg |
| 594 | C G T A C T G A T A C G G A C T G A T C T C G A G A T C G A T C T G C A G A C T T C G A C G T A A T C G A G T C C G A T C G T A C G T A G T C A C G T A C T A G | PSE(SNAPc)/K562-mStart-Seq/Homer | 1e-4 | -9.684e+00 | 0.0001 | 3295.0 | 8.32% | 13918.7 | 7.74% | motif file (matrix) | svg |
| 595 | C T G A G A T C G C A T A C T G C G T A A C G T C G T A C G T A T A C G T C G A | PQM-1(?)/cElegans-L3-ChIP-Seq(modEncode)/Homer | 1e-4 | -9.385e+00 | 0.0001 | 2782.0 | 7.03% | 11687.9 | 6.50% | motif file (matrix) | svg |
| 596 | T G C A T C G A T A G C G T A C T C A G C T A G G T C A G C T A T C A G G A C T | ETS(ETS)/Promoter/Homer | 1e-4 | -9.299e+00 | 0.0002 | 3337.0 | 8.43% | 14128.4 | 7.86% | motif file (matrix) | svg |
| 597 | C G T A C G A T C G A T G C A T C G A T T C A G A G T C A C T G C T A G A G T C A C G T C T G A | At5g08750(C3H)/col-At5g08750-DAP-Seq(GSE60143)/Homer | 1e-3 | -9.047e+00 | 0.0002 | 9769.0 | 24.67% | 42772.9 | 23.80% | motif file (matrix) | svg |
| 598 | G C A T A C G T C G A T A G T C A G T C G C A T C G T A C G T A C G A T C G A T C G A T C T A G A C T G G C T A G C T A | AGL15(MADS)/col-AGL15-DAP-Seq(GSE60143)/Homer | 1e-3 | -8.709e+00 | 0.0003 | 579.0 | 1.46% | 2217.2 | 1.23% | motif file (matrix) | svg |
| 599 | A G C T C T A G T G A C C G T A A C G T C G A T A G T C A G T C C T G A C A T G | TEAD3(TEA)/HepG2-TEAD3-ChIP-Seq(Encode)/Homer | 1e-3 | -8.683e+00 | 0.0003 | 7686.0 | 19.41% | 33486.7 | 18.63% | motif file (matrix) | svg |
| 600 | G C T A A G C T T A C G G T C A C T G A C G A T C T G A G C A T C A G T A G T C | Brn2(POU,Homeobox)/NPC-Brn2-ChIP-Seq(GSE35496)/Homer | 1e-3 | -8.617e+00 | 0.0003 | 494.0 | 1.25% | 1866.2 | 1.04% | motif file (matrix) | svg |
| 601 | A G C T G A T C G A C T A G T C T C G A C T G A A G T C A G T C C T A G A G C T G A C T T A G C T C G A C G A T G A C T | AT5G59990(C2C2COlike)/colamp-AT5G59990-DAP-Seq(GSE60143)/Homer | 1e-3 | -8.465e+00 | 0.0004 | 466.0 | 1.18% | 1754.8 | 0.98% | motif file (matrix) | svg |
| 602 | C G A T C G A T G T A C G A T C G A T C C G T A G C T A C G A T C G A T C T G A C T A G C A T G G C T A G C T A C G T A | AGL16(MADS)/col-AGL16-DAP-Seq(GSE60143)/Homer | 1e-3 | -8.331e+00 | 0.0004 | 416.0 | 1.05% | 1552.0 | 0.86% | motif file (matrix) | svg |
| 603 | A C T G T G A C A C T G A C G T A C G T A C T G C G T A A G T C A G C T C G A T G C A T A C G T | WRKY17(WRKY)/colamp-WRKY17-DAP-Seq(GSE60143)/Homer | 1e-3 | -8.313e+00 | 0.0004 | 114.0 | 0.29% | 349.1 | 0.19% | motif file (matrix) | svg |
| 604 | C G T A C G T A C G T A A C G T C G T A A C G T A G T C G C A T | RVE1(MYBrelated)/col-RVE1-DAP-Seq(GSE60143)/Homer | 1e-3 | -8.215e+00 | 0.0005 | 3815.0 | 9.64% | 16313.1 | 9.08% | motif file (matrix) | svg |
| 605 | A G C T A G C T T A G C A T C G A G T C A C T G A T G C A T C G T C G A C T G A T C G A C T G A | E2F(E2F)/Hela-CellCycle-Expression/Homer | 1e-3 | -8.061e+00 | 0.0005 | 1204.0 | 3.04% | 4896.9 | 2.72% | motif file (matrix) | svg |
| 606 | A C T G G T C A A C G T G C A T C G A T T C A G G T A C G T C A A C G T C T G A | Oct11(POU,Homeobox)/NCIH1048-POU2F3-ChIP-seq(GSE115123)/Homer | 1e-3 | -7.899e+00 | 0.0006 | 1446.0 | 3.65% | 5948.4 | 3.31% | motif file (matrix) | svg |
| 607 | C T G A C T A G A T C G G C A T A C T G G T A C A T G C C G T A A C T G G C T A A G T C C G T A | Tbox:Smad(T-box,MAD)/ESCd5-Smad2\_3-ChIP-Seq(GSE29422)/Homer | 1e-3 | -7.630e+00 | 0.0008 | 1082.0 | 2.73% | 4390.9 | 2.44% | motif file (matrix) | svg |
| 608 | C G T A A C G T A G C T C G A T C T A G G T A C C G T A A G C T C G T A G C T A | Oct4(POU,Homeobox)/mES-Oct4-ChIP-Seq(GSE11431)/Homer | 1e-3 | -7.317e+00 | 0.0011 | 1993.0 | 5.03% | 8360.1 | 4.65% | motif file (matrix) | svg |
| 609 | G C T A C G T A C G T A C G T A A C T G A C G T A G T C C G T A C G T A A G T C C A T G T A G C G T A C C G T A C G T A | WRKY7(WRKY)/colamp-WRKY7-DAP-Seq(GSE60143)/Homer | 1e-3 | -7.229e+00 | 0.0012 | 35.0 | 0.09% | 79.5 | 0.04% | motif file (matrix) | svg |
| 610 | C G T A C G T A C G T A C G A T C G T A A C G T A G T C G C A T | At4g01280(MYBrelated)/colamp-At4g01280-DAP-Seq(GSE60143)/Homer | 1e-3 | -7.057e+00 | 0.0015 | 2924.0 | 7.38% | 12467.3 | 6.94% | motif file (matrix) | svg |
| 611 | C T G A C T A G T G C A C T G A T C G A A G C T C A T G T C G A A G T C G A C T A C G T A G T C G A T C G A T C G A C T | ZNF528(Zf)/HEK293-ZNF528.GFP-ChIP-Seq(GSE58341)/Homer | 1e-3 | -7.038e+00 | 0.0015 | 26.0 | 0.07% | 52.5 | 0.03% | motif file (matrix) | svg |
| 612 | T G A C G T A C C G T A A C T G T G A C C G A T A C T G A T C G A G C T T A C G T C G A T A G C G T A C C G T A A T C G T G A C G C A T A C T G A C T G A T G C | Twist(bHLH)/HMLE-TWIST1-ChIP-Seq(Chang\_et\_al)/Homer | 1e-3 | -6.938e+00 | 0.0017 | 453.0 | 1.14% | 1741.7 | 0.97% | motif file (matrix) | svg |
| 613 | G C A T C G T A C G A T G A C T A C T G C T G A G A C T G A T C | Hnf6b(Homeobox)/LNCaP-Hnf6b-ChIP-Seq(GSE106305)/Homer | 1e-3 | -6.930e+00 | 0.0017 | 8559.0 | 21.62% | 37587.6 | 20.91% | motif file (matrix) | svg |
| 614 | G C T A G C T A T G C A A G T C C T A G C T G A G A T C C T A G G A C T G A T C C T A G A C G T C G A T C G A T G A C T | Unknown2/Arabidopsis-Promoters/Homer | 1e-2 | -6.739e+00 | 0.0020 | 205.0 | 0.52% | 726.6 | 0.40% | motif file (matrix) | svg |
| 615 | C T A G C G A T C G A T A C G T A C G T A C G T A T C G A T G C A T G C A T C G T A C G T G A C C G T A C G T A C G T A | REM16(ABI3VP1)/col-REM16-DAP-Seq(GSE60143)/Homer | 1e-2 | -6.700e+00 | 0.0021 | 19.0 | 0.05% | 34.0 | 0.02% | motif file (matrix) | svg |
| 616 | A C G T T A C G G A T C A C T G A C G T C T A G A C T G A C T G G A T C C T A G C A T G C T A G | Egr2(Zf)/Thymocytes-Egr2-ChIP-Seq(GSE34254)/Homer | 1e-2 | -6.688e+00 | 0.0021 | 1389.0 | 3.51% | 5762.8 | 3.21% | motif file (matrix) | svg |
| 617 | C G T A C G T A T C G A A C T G C G T A C G T A A C G T C G T A A C G T C G A T A G T C A G C T G C A T G C A T C G A T | AT2G20400(G2like)/colamp-AT2G20400-DAP-Seq(GSE60143)/Homer | 1e-2 | -6.424e+00 | 0.0028 | 1244.0 | 3.14% | 5146.7 | 2.86% | motif file (matrix) | svg |
| 618 | T G A C A G T C C T G A T G A C C G T A A C G T A C G T A G T C A G T C C G T A | TEAD1(TEAD)/HepG2-TEAD1-ChIP-Seq(Encode)/Homer | 1e-2 | -6.142e+00 | 0.0037 | 6105.0 | 15.42% | 26691.4 | 14.85% | motif file (matrix) | svg |
| 619 | C T A G T C A G C A G T T C A G A C T G A C T G G A T C C T A G A C T G C T A G T C A G A T G C | KLF14(Zf)/HEK293-KLF14.GFP-ChIP-Seq(GSE58341)/Homer | 1e-2 | -5.898e+00 | 0.0047 | 10069.0 | 25.43% | 44504.3 | 24.76% | motif file (matrix) | svg |
| 620 | C T G A A C T G C G T A C G T A A C G T G T A C G A C T G C A T G C A T C G A T | AT4G37180(G2like)/col-AT4G37180-DAP-Seq(GSE60143)/Homer | 1e-2 | -5.655e+00 | 0.0059 | 7523.0 | 19.00% | 33099.0 | 18.42% | motif file (matrix) | svg |
| 621 | T G C A G T A C C G T A A T C G A C T G A C G T C T A G C G A T T C G A A G T C | ZEB1(Zf)/PDAC-ZEB1-ChIP-Seq(GSE64557)/Homer | 1e-2 | -5.651e+00 | 0.0059 | 9779.0 | 24.70% | 43234.1 | 24.05% | motif file (matrix) | svg |
| 622 | T G C A T G C A A G T C A G T C G A C T C A G T A T G C G A T C C T G A A C G T C T A G C T A G A G T C A C G T A G T C A G T C A G T C G A C T C G T A A C G T A G C T C T A G G A T C G A T C G A T C | ZNF16(Zf)/HEK293-ZNF16.GFP-ChIP-Seq(GSE58341)/Homer | 1e-2 | -5.470e+00 | 0.0071 | 32.0 | 0.08% | 80.3 | 0.04% | motif file (matrix) | svg |
| 623 | G A C T G A T C G A C T A C G T C G T A A C G T A G T C A G T C C G T A G C A T G C T A G C A T | At1g74840(MYBrelated)/col100-At1g74840-DAP-Seq(GSE60143)/Homer | 1e-2 | -5.275e+00 | 0.0086 | 5187.0 | 13.10% | 22688.0 | 12.62% | motif file (matrix) | svg |
| 624 | G A C T G T A C G A C T A G T C T C A G C T G A A G T C A G T C C T A G C G A T A G C T A T G C C T G A C A G T A G C T | AT4G27900(C2C2COlike)/col-AT4G27900-DAP-Seq(GSE60143)/Homer | 1e-2 | -5.082e+00 | 0.0104 | 153.0 | 0.39% | 548.2 | 0.30% | motif file (matrix) | svg |
| 625 | G C A T G A C T A T G C A G C T T C G A C T A G G C T A C G T A C A T G T G A C G C A T G A C T A G T C A G C T C T G A | HSFB4(HSF)/col-HSFB4-DAP-Seq(GSE60143)/Homer | 1e-2 | -4.936e+00 | 0.0121 | 103.0 | 0.26% | 351.4 | 0.20% | motif file (matrix) | svg |
| 626 | T C G A C T A G G C T A C G T A A C G T G T C A A C G T C G A T G A T C A G C T G C A T G C T A | At5g29000(G2like)/col-At5g29000-DAP-Seq(GSE60143)/Homer | 1e-2 | -4.817e+00 | 0.0136 | 2068.0 | 5.22% | 8860.6 | 4.93% | motif file (matrix) | svg |
| 627 | T A G C T A G C G C A T C A T G A C T G G C T A C G T A A C G T A C T G G A T C | TEAD4(TEA)/Tropoblast-Tead4-ChIP-Seq(GSE37350)/Homer | 1e-2 | -4.810e+00 | 0.0136 | 5361.0 | 13.54% | 23520.9 | 13.09% | motif file (matrix) | svg |
| 628 | C T G A A G T C C G T A G T A C A C T G G A C T G C T A C G T A G A C T A G T C | ANAC038(NAC)/col-ANAC038-DAP-Seq(GSE60143)/Homer | 1e-2 | -4.809e+00 | 0.0136 | 18500.0 | 46.72% | 82780.7 | 46.06% | motif file (matrix) | svg |
