## Supplemental dataset for "Hybrid CNN and Multi-Head Attention Model for Analyzing Epigenetic Mechanisms and Gene Expression Across Fungal Phylogenetic Distances": FgramModel_NcrassaTest_K36me3locs_homerResults.html

/projects/wg-feeds/SHAP/FgramModel\_NcrassaTest\_K36me3locs\_SHAP\_noDup\_HOMER// - Homer de novo Motif Results


### Homer *de novo* Motif Results (/projects/wg-feeds/SHAP/FgramModel\_NcrassaTest\_K36me3locs\_SHAP\_noDup\_HOMER//)

Non-redundant Motif File of Results  
Known Motif Enrichment Results  
Gene Ontology Enrichment Results  
If Homer is having trouble matching a motif to a known motif, try copy/pasting the matrix file into
STAMP  
More information on motif finding results: HOMER
| Description of Results
| Tips
  
Total target sequences = 39591  
Total background sequences = 182447  
\* - possible false positive  

|  |  |  |  |  |  |  |  |  |
| --- | --- | --- | --- | --- | --- | --- | --- | --- |
| Rank | Motif | P-value | log P-pvalue | % of Targets | % of Background | STD(Bg STD) | Best Match/Details | Motif File |
| 1 | C G A T A G T C A T C G C G T A A G C T A T C G C G T A G A C T A T C G C G T A G A C T A C T G | 1e-2189 | -5.041e+03 | 41.29% | 17.07% | 133.8bp (138.5bp) | ZML2(C2C2gata)/col-ZML2-DAP-Seq(GSE60143)/Homer(0.780) More Information \| Similar Motifs Found | motif file (matrix) |
| 2 | A T G C T C G A C G T A A T C G T C G A G C A T A T G C T C G A G C T A A T C G | 1e-1923 | -4.430e+03 | 53.60% | 28.44% | 133.7bp (143.4bp) | NR6A1/MA1541.2/Jaspar(0.737) More Information \| Similar Motifs Found | motif file (matrix) |
| 3 | T A C G C T G A T C G A T A C G C T G A T C G A T C A G T C G A | 1e-1334 | -3.073e+03 | 32.56% | 14.84% | 132.2bp (139.5bp) | Unknown4/Arabidopsis-Promoters/Homer(0.844) More Information \| Similar Motifs Found | motif file (matrix) |
| 4 | T G C A C G A T A C T G A T C G G C A T A G C T C A T G A C G T A T G C C G A T A C G T A T C G | 1e-1026 | -2.364e+03 | 44.34% | 26.46% | 135.1bp (142.0bp) | NAC004/MA2043.2/Jaspar(0.598) More Information \| Similar Motifs Found | motif file (matrix) |
| 5 | A T G C G A C T A C G T A T C G C A G T G A T C T C A G C T G A T C A G C T A G | 1e-984 | -2.267e+03 | 57.19% | 38.67% | 130.5bp (142.6bp) | NF1-halfsite(CTF)/LNCaP-NF1-ChIP-Seq(Unpublished)/Homer(0.695) More Information \| Similar Motifs Found | motif file (matrix) |
| 6 | A T G C C G A T A C G T A T G C C G T A A G T C G T C A C A G T A G T C G T A C | 1e-885 | -2.038e+03 | 55.39% | 37.87% | 132.5bp (140.9bp) | ZNF135/MA1587.1/Jaspar(0.695) More Information \| Similar Motifs Found | motif file (matrix) |
| 7 | G A C T A G C T A G C T A G C T A G C T A G C T A G C T A G C T A G C T A G C T A G C T A G C T | 1e-847 | -1.951e+03 | 26.39% | 13.13% | 126.9bp (135.5bp) | VRN1(ABI3VP1)/col-VRN1-DAP-Seq(GSE60143)/Homer(0.914) More Information \| Similar Motifs Found | motif file (matrix) |
| 8 | A G T C C T G A C G A T A T C G C G T A G A C T A C T G C T A G | 1e-827 | -1.906e+03 | 52.47% | 35.63% | 133.0bp (142.8bp) | Yy1/MA0095.4/Jaspar(0.755) More Information \| Similar Motifs Found | motif file (matrix) |
| 9 | G A C T C A T G A C T G A G T C G T C A C T G A A G C T A G T C C G T A C A G T A T G C A T G C | 1e-608 | -1.400e+03 | 39.20% | 25.70% | 133.3bp (142.5bp) | ZNF91(Zf)/HEK-ZNF91.HA-ChIP-Seq(GSE162571)/Homer(0.732) More Information \| Similar Motifs Found | motif file (matrix) |
| 10 | A T G C C T A G C G T A A T C G C T G A G T A C T A C G C T A G | 1e-599 | -1.381e+03 | 35.05% | 22.15% | 134.7bp (138.9bp) | TF3A(C2H2)/col-TF3A-DAP-Seq(GSE60143)/Homer(0.683) More Information \| Similar Motifs Found | motif file (matrix) |
| 11 | A G T C C G A T A C G T A T C G G A C T A G C T A C T G T A C G | 1e-577 | -1.330e+03 | 56.22% | 41.99% | 135.5bp (143.5bp) | grh/dmmpmm(Bigfoot)/fly(0.795) More Information \| Similar Motifs Found | motif file (matrix) |
| 12 | G A C T C A T G C A T G T A G C G T A C G C T A G C A T A C T G A C T G G T A C | 1e-518 | -1.193e+03 | 28.09% | 17.06% | 133.7bp (143.3bp) | Rfx1(HTH)/NPC-H3K4me1-ChIP-Seq(GSE16256)/Homer(0.761) More Information \| Similar Motifs Found | motif file (matrix) |
| 13 | C A G T C T G A A T C G T A C G A C G T C G T A A T C G T A C G G A C T C T G A | 1e-504 | -1.161e+03 | 4.26% | 0.70% | 111.1bp (117.4bp) | PK06182.1/MA2354.1/Jaspar(0.799) More Information \| Similar Motifs Found | motif file (matrix) |
| 14 | G A C T C T G A A G T C G T A C G C A T A G T C C G A T T G C A A T G C T A G C C G A T G T C A | 1e-480 | -1.107e+03 | 3.36% | 0.41% | 130.1bp (114.4bp) | vfl/MA1462.2/Jaspar(0.681) More Information \| Similar Motifs Found | motif file (matrix) |
| 15 | C T A G C A T G G C A T A G T C A C G T A G C T A C T G T A C G | 1e-432 | -9.966e+02 | 34.17% | 23.17% | 137.1bp (139.5bp) | HAP2/MA0313.1/Jaspar(0.839) More Information \| Similar Motifs Found | motif file (matrix) |
| 16 | C G A T G A C T A T C G T A C G A G T C G C A T A C G T G A T C T A G C C G T A | 1e-410 | -9.456e+02 | 29.60% | 19.44% | 135.3bp (140.5bp) | GCR1/MA0304.1/Jaspar(0.819) More Information \| Similar Motifs Found | motif file (matrix) |
| 17 | G A T C G A C T T C G A G A T C A G C T T C G A A T G C A G C T T C G A A T G C G A C T T C G A | 1e-372 | -8.572e+02 | 6.04% | 1.93% | 124.5bp (123.3bp) | SPL4D/MA2445.1/Jaspar(0.696) More Information \| Similar Motifs Found | motif file (matrix) |
| 18 | T C A G C T G A G A C T G C T A A G C T A T G C T C A G G C T A | 1e-332 | -7.661e+02 | 28.54% | 19.45% | 136.0bp (142.0bp) | PB0126.1\_Gata5\_2/Jaspar(0.821) More Information \| Similar Motifs Found | motif file (matrix) |
| 19 | A G C T A T C G C G T A C G T A A C T G A G T C G T A C C T G A | 1e-263 | -6.059e+02 | 17.27% | 10.76% | 127.8bp (143.9bp) | NAC007/MA2044.2/Jaspar(0.853) More Information \| Similar Motifs Found | motif file (matrix) |
| 20 | A C T G A C T G C G T A A C G T A C T G A C T G C G T A A C G T A C T G C T A G | 1e-214 | -4.946e+02 | 12.09% | 7.13% | 127.3bp (136.8bp) | HOXA1(Homeobox)/mES-Hoxa1-ChIP-Seq(SRP084292)/Homer(0.799) More Information \| Similar Motifs Found | motif file (matrix) |
| 21 | T A C G C G A T G C T A G A T C G C T A A T G C C G A T C T G A A T G C G C T A | 1e-173 | -4.006e+02 | 2.47% | 0.70% | 119.6bp (134.9bp) | SFPQ(RRM)/Homo\_sapiens-RNCMPT00177-PBM/HughesRNA(0.802) More Information \| Similar Motifs Found | motif file (matrix) |
| 22 | A C G T A C G T A G T C G T A C G T A C A C T G C G A T G T A C | 1e-166 | -3.835e+02 | 10.79% | 6.61% | 130.0bp (143.8bp) | Knotted(Homeobox)/Corn-KN1-ChIP-Seq(GSE39161)/Homer(0.822) More Information \| Similar Motifs Found | motif file (matrix) |
| 23 | A C G T C T A G G C A T C T G A C G A T C T A G G A C T T C G A A G C T C T A G G A C T T C A G | 1e-117 | -2.709e+02 | 0.96% | 0.15% | 126.3bp (117.4bp) | cg/MA2107.1/Jaspar(0.856) More Information \| Similar Motifs Found | motif file (matrix) |
| 24 | A C T G A C T G A C T G A C T G A C T G A C T G A C T G A C T G A C T G A C T G C T A G A C T G | 1e-85 | -1.976e+02 | 0.49% | 0.04% | 204.3bp (117.1bp) | SeqBias: polyC-repeat(0.918) More Information \| Similar Motifs Found | motif file (matrix) |
| 25 | C T G A T G C A A G T C A G T C A G T C A C G T C T G A C G T A A G T C A G T C A G T C A G C T | 1e-76 | -1.773e+02 | 0.38% | 0.02% | 175.6bp (130.1bp) | TBF1/MA0403.3/Jaspar(0.818) More Information \| Similar Motifs Found | motif file (matrix) |
| 26 | C T G A C T A G T C G A A G T C C T G A T C A G G T C A A G T C C G A T T C A G T C G A G T A C | 1e-49 | -1.149e+02 | 0.90% | 0.31% | 123.2bp (169.6bp) | CNOT4(RRM)/Homo\_sapiens-RNCMPT00156-PBM/HughesRNA(0.687) More Information \| Similar Motifs Found | motif file (matrix) |
| 27 | C G A T C G T A C G A T C G A T C G T A G C A T A C G T C G T A G C A T C G A T C G T A C G A T | 1e-36 | -8.390e+01 | 0.32% | 0.06% | 251.4bp (102.4bp) | Ubx/dmmpmm(Down)/fly(0.778) More Information \| Similar Motifs Found | motif file (matrix) |
| 28 | C T G A C T A G A G C T T C A G C T G A C A G T A G C T A T C G C T G A C G A T C G A T T C A G | 1e-26 | -6.100e+01 | 0.33% | 0.09% | 88.1bp (142.4bp) | WUS1(Homeobox)/colamp-WUS1-DAP-Seq(GSE60143)/Homer(0.745) More Information \| Similar Motifs Found | motif file (matrix) |
| 29 \* | A C T G A C G T C T A G G T C A A C T G A C G T C T A G G T C A T A C G G C A T | 1e-11 | -2.641e+01 | 0.19% | 0.06% | 101.5bp (132.2bp) | MSANTD3/MA1523.2/Jaspar(0.772) More Information \| Similar Motifs Found | motif file (matrix) |
