## Supplemental dataset for "Hybrid CNN and Multi-Head Attention Model for Analyzing Epigenetic Mechanisms and Gene Expression Across Fungal Phylogenetic Distances": FgramModel_NcrassaTest_K36me3locs_knownResults.html

/projects/wg-feeds/SHAP/FgramModel\_NcrassaTest\_K36me3locs\_SHAP\_noDup\_HOMER/ - Homer Known Motif Enrichment Results


### Homer Known Motif Enrichment Results (/projects/wg-feeds/SHAP/FgramModel\_NcrassaTest\_K36me3locs\_SHAP\_noDup\_HOMER/)

Homer *de novo* Motif Results  
Gene Ontology Enrichment Results  
Known Motif Enrichment Results (txt file)  
Total Target Sequences = 39614, Total Background Sequences = 182095

|  |  |  |  |  |  |  |  |  |  |  |  |
| --- | --- | --- | --- | --- | --- | --- | --- | --- | --- | --- | --- |
| Rank | Motif | Name | P-value | log P-pvalue | q-value (Benjamini) | # Target Sequences with Motif | % of Targets Sequences with Motif | # Background Sequences with Motif | % of Background Sequences with Motif | Motif File | SVG |
| 1 | T A G C G T C A G A C T T A G C G T C A G A C T A G T C G C T A G A C T G A T C | ZML2(C2C2gata)/col-ZML2-DAP-Seq(GSE60143)/Homer | 1e-810 | -1.866e+03 | 0.0000 | 3490.0 | 8.81% | 3660.1 | 2.01% | motif file (matrix) | svg |
| 2 | A T G C C A G T A C G T A G T C C A T G A C G T A G T C A C G T A G C T G A T C | Unknown4/Arabidopsis-Promoters/Homer | 1e-775 | -1.787e+03 | 0.0000 | 9400.0 | 23.74% | 21203.4 | 11.62% | motif file (matrix) | svg |
| 3 | C T A G T C A G C T G A C T G A C T A G C T G A C A T G C A T G C T G A C T A G C T A G C G T A C T A G C G T A G T C A | TF3A(C2H2)/col-TF3A-DAP-Seq(GSE60143)/Homer | 1e-571 | -1.316e+03 | 0.0000 | 7553.0 | 19.08% | 17403.4 | 9.54% | motif file (matrix) | svg |
| 4 | A C G T A C G T A C G T A C G T A C G T A C G T A C G T A C G T A C G T A C G T | VRN1(ABI3VP1)/col-VRN1-DAP-Seq(GSE60143)/Homer | 1e-498 | -1.149e+03 | 0.0000 | 1022.0 | 2.58% | 283.6 | 0.16% | motif file (matrix) | svg |
| 5 | C T A G C T G A C T A G C T G A C T A G C T G A C T A G C T G A C T A G C T G A | SeqBias: GA-repeat | 1e-426 | -9.829e+02 | 0.0000 | 26107.0 | 65.94% | 98310.5 | 53.88% | motif file (matrix) | svg |
| 6 | G C T A G C A T A C G T A C G T A G C T G A T C G A C T G A C T A G C T A C G T A C G T A G C T | RLR1?/SacCer-Promoters/Homer | 1e-421 | -9.717e+02 | 0.0000 | 2295.0 | 5.80% | 2960.3 | 1.62% | motif file (matrix) | svg |
| 7 | A C G T A T G C A C G T A C G T A G C T A G T C A G C T A G C T A G C T A G C T A G C T | hTCT(CPE) | 1e-388 | -8.943e+02 | 0.0000 | 10951.0 | 27.66% | 32906.9 | 18.04% | motif file (matrix) | svg |
| 8 | G A T C G C A T A G T C A G C T G A T C A G C T G A T C G A C T G A T C G A C T A G T C A C G T G A T C A G C T G A T C | GAGA-repeat/SacCer-Promoters/Homer | 1e-372 | -8.570e+02 | 0.0000 | 16301.0 | 41.17% | 55290.5 | 30.30% | motif file (matrix) | svg |
| 9 | G A T C G C T A G T A C A G T C G C T A T G C A G T A C G A T C C G T A G A C T | MYB83(MYB)/colamp-MYB83-DAP-Seq(GSE60143)/Homer | 1e-276 | -6.359e+02 | 0.0000 | 11763.0 | 29.71% | 38765.6 | 21.25% | motif file (matrix) | svg |
| 10 | A C G T G A C T T A G C C G T A C T G A C A T G C T A G G A C T G A T C C G T A | Nr5a2(NR)/Pancreas-LRH1-ChIP-Seq(GSE34295)/Homer | 1e-265 | -6.112e+02 | 0.0000 | 4106.0 | 10.37% | 9819.9 | 5.38% | motif file (matrix) | svg |
| 11 | G T A C A C T G A G T C A G T C C T A G G A T C G T A C C T G A | CRF4(AP2EREBP)/colamp-CRF4-DAP-Seq(GSE60143)/Homer | 1e-256 | -5.913e+02 | 0.0000 | 5769.0 | 14.57% | 15811.4 | 8.67% | motif file (matrix) | svg |
| 12 | C T G A C G A T C A T G A T C G G C A T C A T G G C T A A G T C | ASHR1(ND)/col-ASHR1-DAP-Seq(GSE60143)/Homer | 1e-256 | -5.912e+02 | 0.0000 | 7869.0 | 19.88% | 23704.8 | 12.99% | motif file (matrix) | svg |
| 13 | G C A T G A T C C T A G C G T A G C A T C G T A G C A T A G T C C T A G C G T A G C A T C G A T | AT5G22990(C2H2)/col-AT5G22990-DAP-Seq(GSE60143)/Homer | 1e-253 | -5.847e+02 | 0.0000 | 4776.0 | 12.06% | 12303.2 | 6.74% | motif file (matrix) | svg |
| 14 | C G T A G A C T C A T G C T A G A G T C A C T G A C T G G T A C C A T G T A C G | ERF3(AP2EREBP)/colamp-ERF3-DAP-Seq(GSE60143)/Homer | 1e-252 | -5.812e+02 | 0.0000 | 6962.0 | 17.58% | 20330.2 | 11.14% | motif file (matrix) | svg |
| 15 | G C A T A C T G C T A G A G T C A C T G A C T G A G T C A C G T | ERF105(AP2EREBP)/colamp-ERF105-DAP-Seq(GSE60143)/Homer | 1e-252 | -5.810e+02 | 0.0000 | 10716.0 | 27.07% | 35112.4 | 19.24% | motif file (matrix) | svg |
| 16 | C T G A G C A T A C T G C T A G A G T C A C T G A C T G A G T C A C T G T C A G | AT4G18450(AP2EREBP)/col-AT4G18450-DAP-Seq(GSE60143)/Homer | 1e-252 | -5.803e+02 | 0.0000 | 4057.0 | 10.25% | 9846.7 | 5.40% | motif file (matrix) | svg |
| 17 | C T A G G C A T A C T G C T A G A G T C A C T G A C T G A G T C A C T G T C A G | ERF10(AP2EREBP)/col-ERF10-DAP-Seq(GSE60143)/Homer | 1e-250 | -5.757e+02 | 0.0000 | 6774.0 | 17.11% | 19670.9 | 10.78% | motif file (matrix) | svg |
| 18 | C A T G A G T C G T A C A C T G A T G C A G T C C A T G G A T C G A T C C T G A | ERF5(AP2EREBP)/colamp-ERF5-DAP-Seq(GSE60143)/Homer | 1e-249 | -5.754e+02 | 0.0000 | 4224.0 | 10.67% | 10448.2 | 5.73% | motif file (matrix) | svg |
| 19 | C G T A G A C T C A T G C T A G A G T C A C T G C T A G A G T C C A T G C T A G | ERF7(AP2EREBP)/col-ERF7-DAP-Seq(GSE60143)/Homer | 1e-249 | -5.753e+02 | 0.0000 | 12413.0 | 31.35% | 42172.1 | 23.11% | motif file (matrix) | svg |
| 20 | A C G T G A C T A T G C G C T A C T G A C T A G A C T G G A C T A G T C C G T A | Nr5a2(NR)/mES-Nr5a2-ChIP-Seq(GSE19019)/Homer | 1e-246 | -5.685e+02 | 0.0000 | 3186.0 | 8.05% | 7031.9 | 3.85% | motif file (matrix) | svg |
| 21 | C T G A T C A G G T A C G C T A A C T G T G A C G C A T C A T G | SCL(bHLH)/HPC7-Scl-ChIP-Seq(GSE13511)/Homer | 1e-244 | -5.636e+02 | 0.0000 | 16624.0 | 41.99% | 60333.6 | 33.07% | motif file (matrix) | svg |
| 22 | G T C A T G C A T G C A G C T A C G T A G C T A G C T A G C T A | REM19(REM)/colamp-REM19-DAP-Seq(GSE60143)/Homer | 1e-235 | -5.411e+02 | 0.0000 | 2486.0 | 6.28% | 4973.5 | 2.73% | motif file (matrix) | svg |
| 23 | T G A C C T A G T C A G G T C A C G T A T C A G C G A T T C A G T C G A T G C A C T G A T A G C | PU.1-IRF(ETS:IRF)/Bcell-PU.1-ChIP-Seq(GSE21512)/Homer | 1e-219 | -5.059e+02 | 0.0000 | 5449.0 | 13.76% | 15336.8 | 8.41% | motif file (matrix) | svg |
| 24 | G A T C C A G T T A G C A G T C A C T G A G T C A G T C C T A G G A C T G T A C | LEP(AP2EREBP)/col-LEP-DAP-Seq(GSE60143)/Homer | 1e-207 | -4.781e+02 | 0.0000 | 3048.0 | 7.70% | 7092.1 | 3.89% | motif file (matrix) | svg |
| 25 | C G A T A G C T T G C A A C T G A G T C T G A C C T A G G T A C A G T C C G T A G C A T G C A T | ERF13(AP2EREBP)/colamp-ERF13-DAP-Seq(GSE60143)/Homer | 1e-204 | -4.718e+02 | 0.0000 | 9005.0 | 22.74% | 29456.5 | 16.14% | motif file (matrix) | svg |
| 26 | C G T A G C A T C G T A C G T A G C A T A C T G C G A T A G T C A C T G A C T G G A C T C T A G | AT1G71450(AP2EREBP)/col-AT1G71450-DAP-Seq(GSE60143)/Homer | 1e-204 | -4.714e+02 | 0.0000 | 16933.0 | 42.77% | 63055.4 | 34.56% | motif file (matrix) | svg |
| 27 | T G A C C G T A C T G A A C T G A C T G G A C T G A T C T G C A G T A C T A C G | SF1(NR)/H295R-Nr5a1-ChIP-Seq(GSE44220)/Homer | 1e-204 | -4.700e+02 | 0.0000 | 2546.0 | 6.43% | 5511.7 | 3.02% | motif file (matrix) | svg |
| 28 | G C A T C G A T G A C T T G C A A C T G A G T C T G A C A C T G G A T C A G T C C G T A G A C T | ERF15(AP2EREBP)/colamp-ERF15-DAP-Seq(GSE60143)/Homer | 1e-202 | -4.674e+02 | 0.0000 | 11891.0 | 30.03% | 41387.3 | 22.68% | motif file (matrix) | svg |
| 29 | A C G T A C T G C G T A A C G T A C T G A C T G C G T A C G T A | HAP3(CCAATHAP3)/col-HAP3-DAP-Seq(GSE60143)/Homer | 1e-199 | -4.587e+02 | 0.0000 | 2754.0 | 6.96% | 6241.9 | 3.42% | motif file (matrix) | svg |
| 30 | G T A C A C T G A T G C A G T C C T A G G A T C G T A C C T G A G A C T G C A T C G A T G A C T | RAP212(AP2EREBP)/col-RAP212-DAP-Seq(GSE60143)/Homer | 1e-196 | -4.515e+02 | 0.0000 | 7633.0 | 19.28% | 24219.7 | 13.27% | motif file (matrix) | svg |
| 31 | A C T G C T A G A G T C A C T G A C T G A T G C A C T G T A C G | ESE1(AP2EREBP)/col-ESE1-DAP-Seq(GSE60143)/Homer | 1e-186 | -4.285e+02 | 0.0000 | 6142.0 | 15.51% | 18661.3 | 10.23% | motif file (matrix) | svg |
| 32 | G C T A C G T A C G T A G C A T C A T G C T A G G A T C A C T G T C A G G A T C A C T G T C A G | ERF4(AP2EREBP)/colamp-ERF4-DAP-Seq(GSE60143)/Homer | 1e-184 | -4.246e+02 | 0.0000 | 10767.0 | 27.19% | 37278.3 | 20.43% | motif file (matrix) | svg |
| 33 | A C T G C T A G A G T C A C T G A C T G A G T C A C G T C T A G | AT5G23930(mTERF)/col-AT5G23930-DAP-Seq(GSE60143)/Homer | 1e-183 | -4.225e+02 | 0.0000 | 9851.0 | 24.88% | 33516.1 | 18.37% | motif file (matrix) | svg |
| 34 | A C T G C T A G A G T C A C T G A C T G A G T C A C T G T A C G | ERF104(AP2EREBP)/col-ERF104-DAP-Seq(GSE60143)/Homer | 1e-181 | -4.181e+02 | 0.0000 | 8530.0 | 21.54% | 28190.7 | 15.45% | motif file (matrix) | svg |
| 35 | C T G A C G A T C T A G T C A G G A T C C T G A T C A G G A T C C T G A A C T G A G T C G C T A A C G T A G T C G C A T | PRDM9(Zf)/Testis-DMC1-ChIP-Seq(GSE35498)/Homer | 1e-177 | -4.088e+02 | 0.0000 | 1849.0 | 4.67% | 3654.3 | 2.00% | motif file (matrix) | svg |
| 36 | C T G A T C A G A G T C C G T A A T C G A T G C C G A T A C T G A G T C G A C T A T C G A G T C | MyoD(bHLH)/Myotube-MyoD-ChIP-Seq(GSE21614)/Homer | 1e-176 | -4.064e+02 | 0.0000 | 2720.0 | 6.87% | 6439.8 | 3.53% | motif file (matrix) | svg |
| 37 | G A C T G A T C G A T C G C T A G T A C A G T C G C T A C T G A G T A C G A T C G C T A G A C T | MYB13(MYB)/col-MYB13-DAP-Seq(GSE60143)/Homer | 1e-174 | -4.010e+02 | 0.0000 | 5082.0 | 12.84% | 14907.2 | 8.17% | motif file (matrix) | svg |
| 38 | A C T G A C T G A G T C A C T G A C T G A G T C A C G T T C A G | ERF2(AP2EREBP)/colamp-ERF2-DAP-Seq(GSE60143)/Homer | 1e-173 | -3.997e+02 | 0.0000 | 5227.0 | 13.20% | 15465.8 | 8.48% | motif file (matrix) | svg |
| 39 | G C T A C G T A C G T A G A C T C A T G C T A G G A T C A C T G T C A G G A T C A C T G T A C G | ERF9(AP2EREBP)/colamp-ERF9-DAP-Seq(GSE60143)/Homer | 1e-172 | -3.977e+02 | 0.0000 | 2876.0 | 7.26% | 7017.7 | 3.85% | motif file (matrix) | svg |
| 40 | C G A T C G T A G C T A G A C T T C G A A G C T A G T C A C T G T C G A A G C T C T G A C G A T | ZBTB38(Zf)/Hela-ZBTB38-ChIP-seq(GSE108618)/Homer | 1e-171 | -3.951e+02 | 0.0000 | 21591.0 | 54.53% | 85366.8 | 46.79% | motif file (matrix) | svg |
| 41 | A T G C T C G A T A C G A C G T A T G C A G T C A C G T A G T C A G T C G A T C | Znf263(Zf)/K562-Znf263-ChIP-Seq(GSE31477)/Homer | 1e-167 | -3.850e+02 | 0.0000 | 7978.0 | 20.15% | 26379.0 | 14.46% | motif file (matrix) | svg |
| 42 | C G T A C T A G C A G T A C G T C G T A A C T G C A T G G C A T T C A G C T G A | MYB49(MYB)/col-MYB49-DAP-Seq(GSE60143)/Homer | 1e-162 | -3.733e+02 | 0.0000 | 7088.0 | 17.90% | 22964.0 | 12.59% | motif file (matrix) | svg |
| 43 | A C T G A C T G A G T C A C T G A C T G A G T C A C G T C T A G | ERF1(AP2EREBP)/colamp-ERF1-DAP-Seq(GSE60143)/Homer | 1e-160 | -3.689e+02 | 0.0000 | 4710.0 | 11.90% | 13816.2 | 7.57% | motif file (matrix) | svg |
| 44 | A T G C A G T C A C T G A T G C A G T C A C T G A G T C G T A C | SHN3(AP2EREBP)/col-SHN3-DAP-Seq(GSE60143)/Homer | 1e-159 | -3.669e+02 | 0.0000 | 2497.0 | 6.31% | 5942.1 | 3.26% | motif file (matrix) | svg |
| 45 | G T A C A C T G A T G C T G A C C T A G G A C T G T A C C G T A G C A T G C A T | ERF8(AP2EREBP)/colamp-ERF8-DAP-Seq(GSE60143)/Homer | 1e-158 | -3.649e+02 | 0.0000 | 11461.0 | 28.95% | 41059.6 | 22.50% | motif file (matrix) | svg |
| 46 | C T G A G A C T C A T G C T A G A G T C A C T G A C T G A G T C A C T G T C A G | ERF11(AP2EREBP)/col-ERF11-DAP-Seq(GSE60143)/Homer | 1e-156 | -3.604e+02 | 0.0000 | 8802.0 | 22.23% | 30046.2 | 16.47% | motif file (matrix) | svg |
| 47 | A G C T G A T C G A T C C G T A G T A C A G T C C G A T C T G A G T A C G A T C C G T A G A C T | ATY19(MYB)/col-ATY19-DAP-Seq(GSE60143)/Homer | 1e-155 | -3.585e+02 | 0.0000 | 5471.0 | 13.82% | 16808.4 | 9.21% | motif file (matrix) | svg |
| 48 | A G T C G A C T G A T C C G T A G T A C A G T C G C T A C G T A G T A C A G T C G T A C G T A C | MYB63(MYB)/col-MYB63-DAP-Seq(GSE60143)/Homer | 1e-154 | -3.567e+02 | 0.0000 | 4001.0 | 10.11% | 11302.7 | 6.19% | motif file (matrix) | svg |
| 49 | G C T A C G T A C G T A G C A T C A T G C T A G A G T C A C T G T A C G A G T C C A T G T A C G | RAP26(AP2EREBP)/colamp-RAP26-DAP-Seq(GSE60143)/Homer | 1e-153 | -3.525e+02 | 0.0000 | 12060.0 | 30.46% | 43800.1 | 24.01% | motif file (matrix) | svg |
| 50 | A T G C G T A C A G T C A G T C A C G T A C G T C G A T A C G T | AT5G02460(C2C2dof)/col-AT5G02460-DAP-Seq(GSE60143)/Homer | 1e-149 | -3.449e+02 | 0.0000 | 10158.0 | 25.66% | 35883.2 | 19.67% | motif file (matrix) | svg |
| 51 | A T G C G T A C A C T G A G T C A G T C A C T G A G T C G T A C | ERF73(AP2EREBP)/col-ERF73-DAP-Seq(GSE60143)/Homer | 1e-148 | -3.427e+02 | 0.0000 | 5119.0 | 12.93% | 15625.1 | 8.56% | motif file (matrix) | svg |
| 52 | C G T A C T A G C A T G A G C T C T G A C A T G C A G T C G A T C T A G C T A G | MYB30(MYB)/colamp-MYB30-DAP-Seq(GSE60143)/Homer | 1e-145 | -3.361e+02 | 0.0000 | 7265.0 | 18.35% | 24132.9 | 13.23% | motif file (matrix) | svg |
| 53 | C G T A T A G C T A G C T G C A A C T G C T A G C G T A C G T A T C A G G A C T | EHF(ETS)/LoVo-EHF-ChIP-Seq(GSE49402)/Homer | 1e-144 | -3.327e+02 | 0.0000 | 5855.0 | 14.79% | 18581.2 | 10.18% | motif file (matrix) | svg |
| 54 | C G A T A C G T A C G T A G C T A G C T G A T C G A T C G C T A A G C T A C G T A T C G T A C G | NFATC2(RHD)/Islets-NFATC2-ChIP-Seq(GSE158496)/Homer | 1e-143 | -3.300e+02 | 0.0000 | 7997.0 | 20.20% | 27175.1 | 14.89% | motif file (matrix) | svg |
| 55 | T C G A T G C A C A G T T C G A G A T C A G T C C G T A C G T A A C T G A G T C C G T A C G T A T C A G C G A T A G T C | AT5G25475(ABI3VP1)/col-AT5G25475-DAP-Seq(GSE60143)/Homer | 1e-142 | -3.270e+02 | 0.0000 | 6923.0 | 17.49% | 22877.0 | 12.54% | motif file (matrix) | svg |
| 56 | A G T C C G T A T G A C A T G C G C A T C T G A G T A C G A T C | MYB55(MYB)/colamp-MYB55-DAP-Seq(GSE60143)/Homer | 1e-140 | -3.230e+02 | 0.0000 | 8006.0 | 20.22% | 27309.1 | 14.97% | motif file (matrix) | svg |
| 57 | G A T C G A T C G A T C C G T A G T A C A G T C G C A T C G T A G T A C G A T C | MYB58(MYB)/colamp-MYB58-DAP-Seq(GSE60143)/Homer | 1e-140 | -3.225e+02 | 0.0000 | 7144.0 | 18.04% | 23823.5 | 13.06% | motif file (matrix) | svg |
| 58 | C A T G G A C T C T A G C A T G C A G T C G A T C T A G C A T G C G A T C G T A C T A G C A G T C G A T C T A G C A T G | AT1G24250(Orphan)/col-AT1G24250-DAP-Seq(GSE60143)/Homer | 1e-135 | -3.129e+02 | 0.0000 | 2560.0 | 6.47% | 6515.4 | 3.57% | motif file (matrix) | svg |
| 59 | C T A G T A G C A T G C C T A G A G T C A G T C C T A G G A C T G A C T G C T A | CRF10(AP2EREBP)/col100-CRF10-DAP-Seq(GSE60143)/Homer | 1e-131 | -3.039e+02 | 0.0000 | 12837.0 | 32.42% | 47963.0 | 26.29% | motif file (matrix) | svg |
| 60 | G C T A C G T A C G T A G C A T C A T G C T A G A G T C A C T G A C T G A G T C A C T G T C A G | ABR1(AP2EREBP)/colamp-ABR1-DAP-Seq(GSE60143)/Homer | 1e-129 | -2.990e+02 | 0.0000 | 9855.0 | 24.89% | 35329.9 | 19.36% | motif file (matrix) | svg |
| 61 | G A C T G A C T G A T C C G T A G T A C A G T C G C A T C G T A G T A C G A T C G C A T G C T A | MYB74(MYB)/colamp-MYB74-DAP-Seq(GSE60143)/Homer | 1e-128 | -2.950e+02 | 0.0000 | 4354.0 | 11.00% | 13209.0 | 7.24% | motif file (matrix) | svg |
| 62 | T C A G A C T G A C G T C G T A A C T G A C T G A C G T C T A G | MYB51(MYB)/col-MYB51-DAP-Seq(GSE60143)/Homer | 1e-125 | -2.895e+02 | 0.0000 | 6568.0 | 16.59% | 21944.0 | 12.03% | motif file (matrix) | svg |
| 63 | C G T A G C A T C A T G C T A G A G T C A C T G A T C G G T A C A C T G T C A G | At2g33710(AP2EREBP)/colamp-At2g33710-DAP-Seq(GSE60143)/Homer | 1e-125 | -2.880e+02 | 0.0000 | 15568.0 | 39.32% | 60216.4 | 33.00% | motif file (matrix) | svg |
| 64 | G A C T G A T C A G T C C G T A T G A C A G T C G C A T C T G A G T A C G A T C G C A T G A C T | MYB10(MYB)/col-MYB10-DAP-Seq(GSE60143)/Homer | 1e-124 | -2.869e+02 | 0.0000 | 3289.0 | 8.31% | 9315.2 | 5.11% | motif file (matrix) | svg |
| 65 | A G T C G A T C A G C T C G T A G T A C A G T C G C A T C T G A G T A C G A T C | AT4G26030(C2H2)/col-AT4G26030-DAP-Seq(GSE60143)/Homer | 1e-124 | -2.863e+02 | 0.0000 | 7029.0 | 17.75% | 23848.7 | 13.07% | motif file (matrix) | svg |
| 66 | C A T G A G C T T A C G G T C A G T A C T A G C A G C T G A C T A T C G T C G A | Esrrb(NR)/mES-Esrrb-ChIP-Seq(GSE11431)/Homer | 1e-122 | -2.822e+02 | 0.0000 | 3529.0 | 8.91% | 10240.9 | 5.61% | motif file (matrix) | svg |
| 67 | C T A G A C T G A C G T C G T A A C T G A C T G A G C T C T A G T C A G C T A G | MYB93(MYB)/colamp-MYB93-DAP-Seq(GSE60143)/Homer | 1e-121 | -2.788e+02 | 0.0000 | 7753.0 | 19.58% | 26909.3 | 14.75% | motif file (matrix) | svg |
| 68 | C T A G C T A G A T G C G T A C T C A G A T G C A G T C G C A T G A T C G A T C | ZNF91(Zf)/HEK-ZNF91.HA-ChIP-Seq(GSE162571)/Homer | 1e-120 | -2.785e+02 | 0.0000 | 4206.0 | 10.62% | 12819.2 | 7.03% | motif file (matrix) | svg |
| 69 | C G A T C T A G A C T G A G C T C T G A A C T G A C G T A C G T C T A G C T A G | MYB96(MYB)/colamp-MYB96-DAP-Seq(GSE60143)/Homer | 1e-120 | -2.771e+02 | 0.0000 | 6281.0 | 15.86% | 20958.7 | 11.49% | motif file (matrix) | svg |
| 70 | C T A G C A T G G A C T C G T A C T A G A C T G A C G T C T A G C T A G T C A G | MYB17(MYB)/colamp-MYB17-DAP-Seq(GSE60143)/Homer | 1e-118 | -2.731e+02 | 0.0000 | 4410.0 | 11.14% | 13657.0 | 7.48% | motif file (matrix) | svg |
| 71 | G C A T A G C T A C G T A C G T A C T G A C G T G A T C A C G T A C G T A G C T C G A T G C A T A G T C G A C T C A G T | IDD5(C2H2)/colamp-IDD5-DAP-Seq(GSE60143)/Homer | 1e-109 | -2.528e+02 | 0.0000 | 2606.0 | 6.58% | 7139.6 | 3.91% | motif file (matrix) | svg |
| 72 | C A T G C T A G A G T C A C T G A C T G G T A C C A T G T A C G | AT1G28160(AP2EREBP)/colamp-AT1G28160-DAP-Seq(GSE60143)/Homer | 1e-108 | -2.495e+02 | 0.0000 | 14339.0 | 36.22% | 55568.2 | 30.45% | motif file (matrix) | svg |
| 73 | C G T A C G T A C T A G A C G T A C G T C G T A A C T G A C T G A C G T C T G A T C G A T C G A | MYB4(MYB)/col200-MYB4-DAP-Seq(GSE60143)/Homer | 1e-108 | -2.493e+02 | 0.0000 | 4249.0 | 10.73% | 13304.3 | 7.29% | motif file (matrix) | svg |
| 74 | G A C T G C A T A C G T A C G T A C T G C G T A A G T C A G C T C G A T A T C G G C A T A C T G C G A T C T A G C G T A | WRKY50(WRKY)/col-WRKY50-DAP-Seq(GSE60143)/Homer | 1e-108 | -2.488e+02 | 0.0000 | 5768.0 | 14.57% | 19287.0 | 10.57% | motif file (matrix) | svg |
| 75 | C A T G G A C T T A C G G T C A G T A C G A T C G A C T A G C T A T C G T C G A T A C G T A G C | ERRg(NR)/Kidney-ESRRG-ChIP-Seq(GSE104905)/Homer | 1e-105 | -2.424e+02 | 0.0000 | 4381.0 | 11.07% | 13895.9 | 7.62% | motif file (matrix) | svg |
| 76 | G A T C G T A C C T G A A G T C A G T C A C T G G C T A G T A C G T C A G C A T G C A T C G A T | DEAR2(AP2EREBP)/colamp-DEAR2-DAP-Seq(GSE60143)/Homer | 1e-103 | -2.373e+02 | 0.0000 | 12099.0 | 30.56% | 46025.1 | 25.22% | motif file (matrix) | svg |
| 77 | G C A T C G T A C T A G A G T C G T C A C G T A A T G C A C G T A C G T A C T G G A T C G C A T C G T A G C T A G C T A | bHLH122(bHLH)/col100-bHLH122-DAP-Seq(GSE60143)/Homer | 1e-103 | -2.373e+02 | 0.0000 | 4804.0 | 12.13% | 15611.4 | 8.56% | motif file (matrix) | svg |
| 78 | G A T C A G T C G A C T G C T A G T A C A G T C G C A T G C T A G T A C G A T C | MYB61(MYB)/colamp-MYB61-DAP-Seq(GSE60143)/Homer | 1e-102 | -2.359e+02 | 0.0000 | 9693.0 | 24.48% | 35725.2 | 19.58% | motif file (matrix) | svg |
| 79 | G C A T C T A G A C T G A C G T C G T A A C T G A C T G C G A T C T A G T C G A T C G A G C T A | MYB40(MYB)/col-MYB40-DAP-Seq(GSE60143)/Homer | 1e-102 | -2.352e+02 | 0.0000 | 2538.0 | 6.41% | 7044.1 | 3.86% | motif file (matrix) | svg |
| 80 | G C A T C G A T G C A T C G T A C T A G A G T C G T C A C G T A A T C G A C G T A C G T A C T G G T A C G C A T C G A T | bHLH80(bHLH)/col-bHLH80-DAP-Seq(GSE60143)/Homer | 1e-100 | -2.306e+02 | 0.0000 | 5097.0 | 12.87% | 16855.3 | 9.24% | motif file (matrix) | svg |
| 81 | C G T A C T A G C A T G G A C T C T G A A C T G A C G T A C G T C T A G C T A G C A T G T C G A | MYB94(MYB)/col-MYB94-DAP-Seq(GSE60143)/Homer | 1e-99 | -2.291e+02 | 0.0000 | 2937.0 | 7.42% | 8556.4 | 4.69% | motif file (matrix) | svg |
| 82 | T A C G G A C T T G A C C G T A A C G T G A T C G T C A C G T A A C G T A T G C C G T A G A C T | HOXA2(Homeobox)/mES-Hoxa2-ChIP-Seq(Donaldson\_et\_al.)/Homer | 1e-97 | -2.250e+02 | 0.0000 | 962.0 | 2.43% | 1839.9 | 1.01% | motif file (matrix) | svg |
| 83 | A G T C C A T G A C G T A C G T A C T G C G T A A G T C G A C T G C A T G C T A | WRKY28(WRKY)/col-WRKY28-DAP-Seq(GSE60143)/Homer | 1e-97 | -2.239e+02 | 0.0000 | 7077.0 | 17.87% | 24978.7 | 13.69% | motif file (matrix) | svg |
| 84 | C T A G A C T G A C G T C G T A A C T G C A T G G C A T T C A G | MYB92(MYB)/colamp-MYB92-DAP-Seq(GSE60143)/Homer | 1e-96 | -2.223e+02 | 0.0000 | 6730.0 | 17.00% | 23577.0 | 12.92% | motif file (matrix) | svg |
| 85 | A G C T C T A G A G T C A G T C A C T G C G T A A G T C C T G A G C A T G C T A C T G A G C A T G C A T C G A T G C A T | CBF4(AP2EREBP)/colamp-CBF4-DAP-Seq(GSE60143)/Homer | 1e-96 | -2.219e+02 | 0.0000 | 9463.0 | 23.90% | 35008.4 | 19.19% | motif file (matrix) | svg |
| 86 | G A C T C G A T C G A T C T G A G T A C A G T C C G A T C G T A G T C A G A T C G C A T G C A T | MYB121(MYB)/col-MYB121-DAP-Seq(GSE60143)/Homer | 1e-95 | -2.201e+02 | 0.0000 | 2616.0 | 6.61% | 7461.9 | 4.09% | motif file (matrix) | svg |
| 87 | C G T A G A T C C T A G A C G T G T A C C T G A A G C T G A T C G C T A G A C T | TGA2(bZIP)/colamp-TGA2-DAP-Seq(GSE60143)/Homer | 1e-94 | -2.182e+02 | 0.0000 | 6803.0 | 17.18% | 23942.9 | 13.12% | motif file (matrix) | svg |
| 88 | G A C T G C A T C T A G C G A T G A T C T C G A C A T G G A T C | Tgif1(Homeobox)/mES-Tgif1-ChIP-Seq(GSE55404)/Homer | 1e-94 | -2.181e+02 | 0.0000 | 14036.0 | 35.45% | 54907.0 | 30.09% | motif file (matrix) | svg |
| 89 | C G A T C T A G C G T A G A C T C A G T C T A G C G T A A G C T C A T G C T A G | HOXA1(Homeobox)/mES-Hoxa1-ChIP-Seq(SRP084292)/Homer | 1e-94 | -2.171e+02 | 0.0000 | 1903.0 | 4.81% | 4942.5 | 2.71% | motif file (matrix) | svg |
| 90 | C T A G A G T C A G T C A C T G C G T A A G T C C T G A G A C T | DDF1(AP2EREBP)/col-DDF1-DAP-Seq(GSE60143)/Homer | 1e-93 | -2.163e+02 | 0.0000 | 6422.0 | 16.22% | 22409.8 | 12.28% | motif file (matrix) | svg |
| 91 | C T A G C T A G T C G A C T A G C G T A A T C G T C G A A C T G C T G A T C G A C T G A T A C G | FRS9(ND)/col-FRS9-DAP-Seq(GSE60143)/Homer | 1e-93 | -2.158e+02 | 0.0000 | 1018.0 | 2.57% | 2048.3 | 1.12% | motif file (matrix) | svg |
| 92 | C A T G A G T C G T C A C G T A A T G C A C G T A C G T A C T G | bHLH130(bHLH)/col-bHLH130-DAP-Seq(GSE60143)/Homer | 1e-92 | -2.123e+02 | 0.0000 | 4191.0 | 10.59% | 13521.3 | 7.41% | motif file (matrix) | svg |
| 93 | A T C G A G T C A C T G A G T C A G T C A C T G G A C T G A C T | PUCHI(AP2EREBP)/colamp-PUCHI-DAP-Seq(GSE60143)/Homer | 1e-89 | -2.054e+02 | 0.0000 | 7172.0 | 18.11% | 25680.8 | 14.07% | motif file (matrix) | svg |
| 94 | G T A C A C G T A C G T A T C G C A G T C G A T A T C G G C T A T G C A T A G C C G T A G T C A C A T G A G C T G C T A | ANAC013(NAC)/col-ANAC013-DAP-Seq(GSE60143)/Homer | 1e-88 | -2.033e+02 | 0.0000 | 2472.0 | 6.24% | 7089.8 | 3.89% | motif file (matrix) | svg |
| 95 | A G T C T G C A T C G A C T G A A C T G C A T G A C G T A T G C G T C A T A C G | Erra(NR)/HepG2-Erra-ChIP-Seq(GSE31477)/Homer | 1e-86 | -2.002e+02 | 0.0000 | 8088.0 | 20.43% | 29600.2 | 16.22% | motif file (matrix) | svg |
| 96 | C G T A G A T C A G C T A C G T A C G T A C T G C G T A G T A C A G C T G C T A C G A T C G A T C G A T G C A T G C T A | WRKY18(WRKY)/col-WRKY18-DAP-Seq(GSE60143)/Homer | 1e-86 | -1.994e+02 | 0.0000 | 9542.0 | 24.10% | 35778.7 | 19.61% | motif file (matrix) | svg |
| 97 | T A C G T A G C G C T A C G A T C T A G A C G T C A G T C A G T G C T A A G T C G T C A G C A T | FOXK2(Forkhead)/U2OS-FOXK2-ChIP-Seq(E-MTAB-2204)/Homer | 1e-86 | -1.993e+02 | 0.0000 | 3085.0 | 7.79% | 9414.0 | 5.16% | motif file (matrix) | svg |
| 98 | C G A T C G T A G T A C A C G T A C G T T C A G G C A T C A G T T A C G G T C A C G T A A G T C C G T A T G C A C A T G | NAC2(NAC)/colamp-NAC2-DAP-Seq(GSE60143)/Homer | 1e-85 | -1.976e+02 | 0.0000 | 3572.0 | 9.02% | 11295.9 | 6.19% | motif file (matrix) | svg |
| 99 | G A C T A G T C G A T C C G T A G T A C A G T C G C A T C G T A G T C A G A T C | MYB67(MYB)/col-MYB67-DAP-Seq(GSE60143)/Homer | 1e-84 | -1.952e+02 | 0.0000 | 6486.0 | 16.38% | 23014.9 | 12.61% | motif file (matrix) | svg |
| 100 | C T A G A C T G A G C T C G T A A C T G A C T G A C G T C T A G | MYB99(MYB)/colamp-MYB99-DAP-Seq(GSE60143)/Homer | 1e-83 | -1.916e+02 | 0.0000 | 6675.0 | 16.86% | 23856.7 | 13.08% | motif file (matrix) | svg |
| 101 | C G A T T G C A A G T C A C G T A C G T T A C G C G A T C G A T T A C G G C T A G C T A A T G C C G T A G T C A C A T G | ANAC016(NAC)/col-ANAC016-DAP-Seq(GSE60143)/Homer | 1e-81 | -1.879e+02 | 0.0000 | 5109.0 | 12.90% | 17508.8 | 9.60% | motif file (matrix) | svg |
| 102 | C T A G T C A G C T G A T C A G T G C A A C T G T C G A T C A G | Trl(Zf)/S2-GAGAfactor-ChIP-Seq(GSE40646)/Homer | 1e-80 | -1.864e+02 | 0.0000 | 12558.0 | 31.72% | 49123.3 | 26.92% | motif file (matrix) | svg |
| 103 | G A C T A G C T A G C T C T A G A C G T G A T C A C G T A C G T G A C T C G A T G C A T A G T C | IDD4(C2H2)/col-IDD4-DAP-Seq(GSE60143)/Homer | 1e-80 | -1.862e+02 | 0.0000 | 3388.0 | 8.56% | 10722.6 | 5.88% | motif file (matrix) | svg |
| 104 | G A C T C A G T G C A T C G A T T G A C A C G T A T G C G T A C C T G A A C T G A C T G A G C T | WIP5(C2H2)/colamp-WIP5-DAP-Seq(GSE60143)/Homer | 1e-80 | -1.861e+02 | 0.0000 | 5365.0 | 13.55% | 18572.2 | 10.18% | motif file (matrix) | svg |
| 105 | C G T A G C A T C A G T C T A G A G T C A C T G A C T G G T A C A C T G A T C G | ERF115(AP2EREBP)/colamp-ERF115-DAP-Seq(GSE60143)/Homer | 1e-80 | -1.848e+02 | 0.0000 | 13991.0 | 35.34% | 55477.8 | 30.41% | motif file (matrix) | svg |
| 106 | A G C T A C G T A C T G A T G C A G T C C G T A C T G A T A C G | NF1-halfsite(CTF)/LNCaP-NF1-ChIP-Seq(Unpublished)/Homer | 1e-79 | -1.833e+02 | 0.0000 | 7565.0 | 19.11% | 27717.9 | 15.19% | motif file (matrix) | svg |
| 107 | G A C T A C T G C G A T A G T C A C T G C T A G A G T C C G T A | Rap210(AP2EREBP)/col-Rap210-DAP-Seq(GSE60143)/Homer | 1e-79 | -1.826e+02 | 0.0000 | 8093.0 | 20.44% | 29954.8 | 16.42% | motif file (matrix) | svg |
| 108 | C G T A T G A C T A G C T G C A A C T G A C T G C G T A C G T A T C A G G A C T | ELF3(ETS)/PDAC-ELF3-ChIP-Seq(GSE64557)/Homer | 1e-79 | -1.822e+02 | 0.0000 | 2765.0 | 6.98% | 8389.0 | 4.60% | motif file (matrix) | svg |
| 109 | A G C T A G C T C A T G C T G A G T A C A G T C A G C T A G C T C A G T C T A G | RARa(NR)/K562-RARa-ChIP-Seq(Encode)/Homer | 1e-78 | -1.797e+02 | 0.0000 | 12061.0 | 30.46% | 47100.4 | 25.81% | motif file (matrix) | svg |
| 110 | C G A T C G T A G T A C A C G T A C G T T C A G G C A T C A G T T A C G G T C A C G T A A G T C C G T A T G C A C A T G | ANAC053(NAC)/colamp-ANAC053-DAP-Seq(GSE60143)/Homer | 1e-77 | -1.775e+02 | 0.0000 | 3077.0 | 7.77% | 9628.4 | 5.28% | motif file (matrix) | svg |
| 111 | G T C A G C T A G C T A T C G A A T C G A C G T A G T C T C G A T C G A T G A C | WRKY40(WRKY)/colamp-WRKY40-DAP-Seq(GSE60143)/Homer | 1e-76 | -1.750e+02 | 0.0000 | 3504.0 | 8.85% | 11309.3 | 6.20% | motif file (matrix) | svg |
| 112 | G A T C G A C T G A C T A C G T A G T C A C G T A G T C A C G T A G T C A C G T A G T C A C G T G T A C C G A T G T C A | BPC6(BBRBPC)/col-BPC6-DAP-Seq(GSE60143)/Homer | 1e-75 | -1.742e+02 | 0.0000 | 215.0 | 0.54% | 127.0 | 0.07% | motif file (matrix) | svg |
| 113 | T A C G C G T A G A C T T C A G A G C T A G T C A C T G T C A G A G T C C T G A | DDF2(AP2EREBP)/col-DDF2-DAP-Seq(GSE60143)/Homer | 1e-75 | -1.732e+02 | 0.0000 | 1062.0 | 2.68% | 2410.4 | 1.32% | motif file (matrix) | svg |
| 114 | C G A T T C G A G A T C A C G T A C G T T C A G G C T A C G A T C G T A C G T A C G T A A T G C C G T A T G C A C T A G | ANAC028(NAC)/col-ANAC028-DAP-Seq(GSE60143)/Homer | 1e-72 | -1.672e+02 | 0.0000 | 3774.0 | 9.53% | 12472.8 | 6.84% | motif file (matrix) | svg |
| 115 | T C G A A G T C C G T A A T C G A T G C C G A T A C T G A G T C A G C T A C T G | Tcf12(bHLH)/GM12878-Tcf12-ChIP-Seq(GSE32465)/Homer | 1e-71 | -1.657e+02 | 0.0000 | 2804.0 | 7.08% | 8721.8 | 4.78% | motif file (matrix) | svg |
| 116 | C G A T C T G A A G T C A C G T A C G T T A C G G C T A C A T G C T A G G C A T C G A T A G T C C G T A G T C A A C T G | ANAC096(NAC)/colamp-ANAC096-DAP-Seq(GSE60143)/Homer | 1e-71 | -1.647e+02 | 0.0000 | 3314.0 | 8.37% | 10700.1 | 5.86% | motif file (matrix) | svg |
| 117 | G A C T C T G A G T A C A G T C C G A T C G T A G T C A G A T C G C A T G C A T G C A T C G A T | AT3G10580(MYBrelated)/colamp-AT3G10580-DAP-Seq(GSE60143)/Homer | 1e-71 | -1.639e+02 | 0.0000 | 2885.0 | 7.29% | 9051.6 | 4.96% | motif file (matrix) | svg |
| 118 | C A T G G C T A C T A G T A C G C G T A T C A G C G T A A C T G C G T A C A T G C T G A C G T A | BPC1(BBRBPC)/colamp-BPC1-DAP-Seq(GSE60143)/Homer | 1e-71 | -1.638e+02 | 0.0000 | 2275.0 | 5.75% | 6747.1 | 3.70% | motif file (matrix) | svg |
| 119 | A G T C G A T C C T G A A G T C A G T C C A T G G T C A G A T C C G T A G A T C | DREB26(AP2EREBP)/col-DREB26-DAP-Seq(GSE60143)/Homer | 1e-71 | -1.635e+02 | 0.0000 | 2371.0 | 5.99% | 7108.9 | 3.90% | motif file (matrix) | svg |
| 120 | C T A G A C T G A C G T C G T A A C T G A C T G A C G T T C A G C T G A T C G A | MYB107(MYB)/col-MYB107-DAP-Seq(GSE60143)/Homer | 1e-70 | -1.635e+02 | 0.0000 | 10072.0 | 25.44% | 38822.3 | 21.28% | motif file (matrix) | svg |
| 121 | A C T G A C G T C G A T C A G T C A T G C A T G C A G T G C A T C A G T C A T G | HuR(?)/HEK293-HuR-CLIP-Seq(GSE87887)/Homer | 1e-69 | -1.600e+02 | 0.0000 | 14196.0 | 35.86% | 57012.7 | 31.25% | motif file (matrix) | svg |
| 122 | G A C T A G T C C T G A A G T C A G T C A C T G C T G A A G T C G C T A G C A T G T A C C G A T G C A T G A C T C G A T | CBF2(AP2EREBP)/colamp-CBF2-DAP-Seq(GSE60143)/Homer | 1e-69 | -1.592e+02 | 0.0000 | 6488.0 | 16.39% | 23660.8 | 12.97% | motif file (matrix) | svg |
| 123 | G T A C G T C A G T A C G T C A G T A C G T C A G T A C G T C A G T A C G T C A | SeqBias: CA-repeat | 1e-69 | -1.591e+02 | 0.0000 | 22120.0 | 55.87% | 93039.7 | 50.99% | motif file (matrix) | svg |
| 124 | C T G A T C G A G T A C A C G T A C G T A T C G A C G T C G A T A T C G G C T A G T A C A T G C C G T A T G C A C A T G | ANAC103(NAC)/col-ANAC103-DAP-Seq(GSE60143)/Homer | 1e-68 | -1.574e+02 | 0.0000 | 2081.0 | 5.26% | 6092.4 | 3.34% | motif file (matrix) | svg |
| 125 | A G C T T G A C C G A T C G A T C T A G A C G T C A G T C A G T G C T A A G T C | FOXK1(Forkhead)/HEK293-FOXK1-ChIP-Seq(GSE51673)/Homer | 1e-66 | -1.541e+02 | 0.0000 | 4628.0 | 11.69% | 16086.5 | 8.82% | motif file (matrix) | svg |
| 126 | C G T A C G T A C T A G C A G T G A C T C G T A C A T G C A T G C G A T C T G A C T G A C T G A | MS188(MYB)/colamp-MS188-DAP-Seq(GSE60143)/Homer | 1e-66 | -1.526e+02 | 0.0000 | 4318.0 | 10.91% | 14856.2 | 8.14% | motif file (matrix) | svg |
| 127 | C A G T A C T G T C A G T G C A G C T A A T G C T C G A A T C G G T C A T G C A | ZNF189(Zf)/HEK293-ZNF189.GFP-ChIP-Seq(GSE58341)/Homer | 1e-66 | -1.522e+02 | 0.0000 | 3680.0 | 9.29% | 12309.5 | 6.75% | motif file (matrix) | svg |
| 128 | C T A G A C T G A G T C A C T G A C T G A G C T A C T G T C A G | AT3G57600(AP2EREBP)/col-AT3G57600-DAP-Seq(GSE60143)/Homer | 1e-65 | -1.506e+02 | 0.0000 | 5373.0 | 13.57% | 19191.6 | 10.52% | motif file (matrix) | svg |
| 129 | A T G C A G T C G C A T A G C T A C G T T C A G C G A T A G C T G A T C A T C G | Sox10(HMG)/SciaticNerve-Sox3-ChIP-Seq(GSE35132)/Homer | 1e-65 | -1.500e+02 | 0.0000 | 7550.0 | 19.07% | 28312.7 | 15.52% | motif file (matrix) | svg |
| 130 | C G A T T C G A G A T C A C G T A C G T T C A G G C A T C T G A C G T A G C T A C G T A A G T C C G T A T G C A C A T G | ANAC050(NAC)/colamp-ANAC050-DAP-Seq(GSE60143)/Homer | 1e-64 | -1.496e+02 | 0.0000 | 3244.0 | 8.19% | 10623.2 | 5.82% | motif file (matrix) | svg |
| 131 | C G T A A C G T A C G T A C G T A C G T A G T C A G T C C T G A A G C T A G C T | NFAT(RHD)/Jurkat-NFATC1-ChIP-Seq(Jolma\_et\_al.)/Homer | 1e-64 | -1.478e+02 | 0.0000 | 3741.0 | 9.45% | 12614.9 | 6.91% | motif file (matrix) | svg |
| 132 | G A C T G A T C C T G A A G T C A G T C A C T G C G T A A G T C G T A C G C T A G C A T C G A T | At1g19210(AP2EREBP)/colamp-At1g19210-DAP-Seq(GSE60143)/Homer | 1e-63 | -1.473e+02 | 0.0000 | 14367.0 | 36.29% | 58115.3 | 31.85% | motif file (matrix) | svg |
| 133 | C G A T C T G A A G T C A C G T A C G T A T C G G A C T C T A G C G A T G A C T C G T A A T G C C G T A G T C A A C T G | ANAC011(NAC)/col-ANAC011-DAP-Seq(GSE60143)/Homer | 1e-63 | -1.467e+02 | 0.0000 | 1581.0 | 3.99% | 4373.3 | 2.40% | motif file (matrix) | svg |
| 134 | G A C T A C T G C A G T A G T C A C T G A C T G A G C T A C T G C T A G G T C A | At1g77640(AP2EREBP)/col-At1g77640-DAP-Seq(GSE60143)/Homer | 1e-63 | -1.456e+02 | 0.0000 | 2017.0 | 5.09% | 5974.6 | 3.27% | motif file (matrix) | svg |
| 135 | A G C T C T A G G A T C A G T C C T A G C T G A A G T C G C T A G C A T T G C A | CBF1(AP2EREBP)/colamp-CBF1-DAP-Seq(GSE60143)/Homer | 1e-62 | -1.440e+02 | 0.0000 | 8504.0 | 21.48% | 32512.6 | 17.82% | motif file (matrix) | svg |
| 136 | G C A T C G T A G C T A G A C T C G A T G A C T A G T C C A G T A G T C A G T C A C T G C T A G G T A C C T A G C T G A | AT5G05550(Trihelix)/col-AT5G05550-DAP-Seq(GSE60143)/Homer | 1e-62 | -1.438e+02 | 0.0000 | 12854.0 | 32.47% | 51488.9 | 28.22% | motif file (matrix) | svg |
| 137 | A G C T C A T G G C A T G A T C T G C A C T A G G A T C A C G T | Tgif2(Homeobox)/mES-Tgif2-ChIP-Seq(GSE55404)/Homer | 1e-62 | -1.434e+02 | 0.0000 | 14882.0 | 37.59% | 60525.1 | 33.17% | motif file (matrix) | svg |
| 138 | G A C T A C T G C G A T A G T C A C T G C T A G A G T C C T G A | AT1G12630(AP2EREBP)/colamp-AT1G12630-DAP-Seq(GSE60143)/Homer | 1e-61 | -1.416e+02 | 0.0000 | 6147.0 | 15.53% | 22573.9 | 12.37% | motif file (matrix) | svg |
| 139 | C G T A C G A T C T A G C G T A A G C T C A G T T A C G C G T A A C G T C A T G C T A G A T G C | HOXA3(Homeobox)/mEmbryo-Hoxa3-ChIP-Seq(E-MTAB-8607)/Homer | 1e-61 | -1.412e+02 | 0.0000 | 1084.0 | 2.74% | 2685.0 | 1.47% | motif file (matrix) | svg |
| 140 | C G T A C G A T C A G T C A T G C G A T G T A C C A T G A C T G G A C T C A T G | CEJ1(AP2EREBP)/col-CEJ1-DAP-Seq(GSE60143)/Homer | 1e-60 | -1.392e+02 | 0.0000 | 11164.0 | 28.20% | 44167.6 | 24.21% | motif file (matrix) | svg |
| 141 | C G A T T C G A G T A C A C G T A C G T T C A G G C A T G C A T G C T A C G T A C G T A A G T C C G T A T G C A C A T G | ANAC020(NAC)/col-ANAC020-DAP-Seq(GSE60143)/Homer | 1e-60 | -1.384e+02 | 0.0000 | 3683.0 | 9.30% | 12523.4 | 6.86% | motif file (matrix) | svg |
| 142 | A G T C G A T C G C T A C G A T C A G T T A C G C G A T A G C T A G T C A T C G | SOX1(HMG)/NPC-SOX1-ChIP-Seq(GSE138215)/Homer | 1e-59 | -1.374e+02 | 0.0000 | 9563.0 | 24.15% | 37233.7 | 20.41% | motif file (matrix) | svg |
| 143 | C T G A T G A C T G A C C G T A A C G T T G A C A G C T C T A G A C G T G A C T | Olig2(bHLH)/Neuron-Olig2-ChIP-Seq(GSE30882)/Homer | 1e-59 | -1.371e+02 | 0.0000 | 7707.0 | 19.47% | 29259.4 | 16.04% | motif file (matrix) | svg |
| 144 | G A T C A G C T C T A G G A T C T G A C C T A G C G T A G T A C C G T A G C A T G T C A C T G A | CBF3(AP2EREBP)/colamp-CBF3-DAP-Seq(GSE60143)/Homer | 1e-59 | -1.363e+02 | 0.0000 | 6295.0 | 15.90% | 23299.0 | 12.77% | motif file (matrix) | svg |
| 145 | A T G C G A T C C G A T A C G T A C G T A C T G C A G T A G C T | Sox3(HMG)/NPC-Sox3-ChIP-Seq(GSE33059)/Homer | 1e-57 | -1.335e+02 | 0.0000 | 8129.0 | 20.53% | 31146.2 | 17.07% | motif file (matrix) | svg |
| 146 | A G T C A C G T A C G T T C A G G C T A G C T A A T G C C G T A C G A T A G T C C G T A G T C A A C T G G A T C G C A T | SND3(NAC)/col-SND3-DAP-Seq(GSE60143)/Homer | 1e-57 | -1.329e+02 | 0.0000 | 3933.0 | 9.93% | 13612.7 | 7.46% | motif file (matrix) | svg |
| 147 | G T A C C A T G A G C T A C G T A C T G C G T A A G T C G A C T G C A T C G A T | WRKY29(WRKY)/colamp-WRKY29-DAP-Seq(GSE60143)/Homer | 1e-57 | -1.325e+02 | 0.0000 | 5655.0 | 14.28% | 20695.0 | 11.34% | motif file (matrix) | svg |
| 148 | T C A G A C G T T C G A T A G C A G T C C G T A A C T G G T A C A C G T A C T G A T C G A G T C | Atoh1(bHLH)/Cerebellum-Atoh1-ChIP-Seq(GSE22111)/Homer | 1e-56 | -1.297e+02 | 0.0000 | 3904.0 | 9.86% | 13547.3 | 7.42% | motif file (matrix) | svg |
| 149 | A G T C A C G T A C G T T A C G G C A T G C A T A T G C G C T A C G T A A T G C C G T A G T C A A C T G G A T C G C A T | ANAC075(NAC)/col-ANAC075-DAP-Seq(GSE60143)/Homer | 1e-56 | -1.294e+02 | 0.0000 | 2203.0 | 5.56% | 6849.9 | 3.75% | motif file (matrix) | svg |
| 150 | G A T C A T G C A G T C C G T A A G T C A G T C A C T G G C T A A G T C C G T A | AT1G44830(AP2EREBP)/col-AT1G44830-DAP-Seq(GSE60143)/Homer | 1e-56 | -1.291e+02 | 0.0000 | 3032.0 | 7.66% | 10071.4 | 5.52% | motif file (matrix) | svg |
| 151 | C G A T C G T A A G T C A C G T A C G T T C G A T G C A G C A T G C T A C G T A A C G T A G C T C G T A C G T A A C T G | ANAC062(NAC)/colamp-ANAC062-DAP-Seq(GSE60143)/Homer | 1e-55 | -1.286e+02 | 0.0000 | 1530.0 | 3.86% | 4352.0 | 2.39% | motif file (matrix) | svg |
| 152 | C G A T T C G A G A T C C G A T G C A T T C A G G A C T C G A T G C A T G C T A C T G A A G T C C G T A G T C A C T A G | ANAC005(NAC)/col-ANAC005-DAP-Seq(GSE60143)/Homer | 1e-55 | -1.284e+02 | 0.0000 | 1813.0 | 4.58% | 5393.8 | 2.96% | motif file (matrix) | svg |
| 153 | G T A C A C G T A C G T T C A G G A C T G C A T T C A G C G T A C T G A A G T C C G T A G T C A A C T G A C G T G C T A | NTM2(NAC)/col-NTM2-DAP-Seq(GSE60143)/Homer | 1e-55 | -1.277e+02 | 0.0000 | 2727.0 | 6.89% | 8893.4 | 4.87% | motif file (matrix) | svg |
| 154 | C A G T C A T G T G C A G T A C C G T A T C A G G T A C G A C T T C A G C T G A | bZIP18(bZIP)/colamp-bZIP18-DAP-Seq(GSE60143)/Homer | 1e-54 | -1.256e+02 | 0.0000 | 21895.0 | 55.30% | 93015.0 | 50.98% | motif file (matrix) | svg |
| 155 | C A T G C T G A A G T C A C T G A C T G A G C T A C T G A T C G | ESE3(AP2EREBP)/col-ESE3-DAP-Seq(GSE60143)/Homer | 1e-54 | -1.248e+02 | 0.0000 | 11333.0 | 28.62% | 45294.2 | 24.82% | motif file (matrix) | svg |
| 156 | A T G C C A G T A G C T A G C T T C A G G T C A T A G C G A C T C G T A C G A T | WRKY20(WRKY)/col-WRKY20-DAP-Seq(GSE60143)/Homer | 1e-54 | -1.245e+02 | 0.0000 | 4271.0 | 10.79% | 15127.6 | 8.29% | motif file (matrix) | svg |
| 157 | C T G A A G T C C G A T A G C T A T G C G T A C A C G T A T C G C A G T G C A T | Elf4(ETS)/BMDM-Elf4-ChIP-Seq(GSE88699)/Homer | 1e-54 | -1.244e+02 | 0.0000 | 4623.0 | 11.68% | 16573.7 | 9.08% | motif file (matrix) | svg |
| 158 | G A T C C T G A A G T C A G T C A C T G C G T A A G T C C T G A | ERF38(AP2EREBP)/col-ERF38-DAP-Seq(GSE60143)/Homer | 1e-53 | -1.241e+02 | 0.0000 | 6085.0 | 15.37% | 22668.7 | 12.42% | motif file (matrix) | svg |
| 159 | A G T C G T A C C T G A A G T C G T A C C T A G G C T A T G A C T G C A G C T A C G T A C G T A | At1g22810(AP2EREBP)/colamp-At1g22810-DAP-Seq(GSE60143)/Homer | 1e-53 | -1.237e+02 | 0.0000 | 5235.0 | 13.22% | 19119.5 | 10.48% | motif file (matrix) | svg |
| 160 | G C A T T G A C C A T G G A C T C A G T C A T G T C G A G T A C G A C T G C T A C G A T C G A T | WRKY6(WRKY)/colamp-WRKY6-DAP-Seq(GSE60143)/Homer | 1e-53 | -1.221e+02 | 0.0000 | 4816.0 | 12.16% | 17411.4 | 9.54% | motif file (matrix) | svg |
| 161 | C G T A G A T C C A T G G C A T G A C T C T A G T C G A T A G C A G C T G C A T | WRKY55(WRKY)/col-WRKY55-DAP-Seq(GSE60143)/Homer | 1e-52 | -1.214e+02 | 0.0000 | 6197.0 | 15.65% | 23197.2 | 12.71% | motif file (matrix) | svg |
| 162 | G A C T G C T A T G C A A G T C A C G T A C G T A C G T C G A T A C G T T A C G | At3g45610(C2C2dof)/col-At3g45610-DAP-Seq(GSE60143)/Homer | 1e-52 | -1.212e+02 | 0.0000 | 6466.0 | 16.33% | 24337.7 | 13.34% | motif file (matrix) | svg |
| 163 | A G T C A G T C C G A T A C G T A C G T A C T G A C G T A G C T A G T C A G T C | Sox4(HMG)/proB-Sox4-ChIP-Seq(GSE50066)/Homer | 1e-52 | -1.207e+02 | 0.0000 | 3829.0 | 9.67% | 13391.4 | 7.34% | motif file (matrix) | svg |
| 164 | A G T C A C G T A C T G A G C T A C G T A C G T G T C A A G T C | Foxo1(Forkhead)/RAW-Foxo1-ChIP-Seq(Fan\_et\_al.)/Homer | 1e-52 | -1.201e+02 | 0.0000 | 7166.0 | 18.10% | 27338.5 | 14.98% | motif file (matrix) | svg |
| 165 | A T G C G T A C C T G A A G T C A G T C A C T G G T C A A G T C G T C A G C A T G C A T G A C T | At5g65130(AP2EREBP)/colamp-At5g65130-DAP-Seq(GSE60143)/Homer | 1e-51 | -1.188e+02 | 0.0000 | 2436.0 | 6.15% | 7876.1 | 4.32% | motif file (matrix) | svg |
| 166 | T C A G T C A G G C T A C G T A T A C G G A C T T C A G T C G A C T G A C G T A T A C G G A C T | IRF8(IRF)/BMDM-IRF8-ChIP-Seq(GSE77884)/Homer | 1e-50 | -1.159e+02 | 0.0000 | 1299.0 | 3.28% | 3633.8 | 1.99% | motif file (matrix) | svg |
| 167 | G A C T G T A C C T G A G A T C A G T C C T A G G C T A G T A C C T G A G C T A G C A T C G A T G C A T G A C T C G T A | AT3G16280(AP2EREBP)/colamp-AT3G16280-DAP-Seq(GSE60143)/Homer | 1e-50 | -1.159e+02 | 0.0000 | 5358.0 | 13.53% | 19788.4 | 10.85% | motif file (matrix) | svg |
| 168 | C G T A C G T A C G T A C G T A C T G A C A G T A C G T C G T A A C T G A C T G A C G T C T A G C T G A T C G A C T G A | MYB39(MYB)/col-MYB39-DAP-Seq(GSE60143)/Homer | 1e-50 | -1.155e+02 | 0.0000 | 1021.0 | 2.58% | 2655.8 | 1.46% | motif file (matrix) | svg |
| 169 | C G T A C G A T C G T A C G A T C A T G A C T G C G A T A G T C A T C G T C A G G A C T A C T G | At1g36060(AP2EREBP)/colamp-At1g36060-DAP-Seq(GSE60143)/Homer | 1e-49 | -1.150e+02 | 0.0000 | 7352.0 | 18.57% | 28253.5 | 15.48% | motif file (matrix) | svg |
| 170 | G A C T C T A G A T G C A G T C G T C A T A C G A T G C A T C G | HIC1(Zf)/Treg-ZBTB29-ChIP-Seq(GSE99889)/Homer | 1e-49 | -1.143e+02 | 0.0000 | 9788.0 | 24.72% | 38810.2 | 21.27% | motif file (matrix) | svg |
| 171 | A T G C C A T G G C A T G A C T C T A G T C G A G T A C A G C T C G T A G C T A | WRKY75(WRKY)/col-WRKY75-DAP-Seq(GSE60143)/Homer | 1e-49 | -1.137e+02 | 0.0000 | 5198.0 | 13.13% | 19163.1 | 10.50% | motif file (matrix) | svg |
| 172 | G T A C C A T G A G C T A G C T T C A G T G C A T G A C A G C T C G T A C G T A | WRKY33(WRKY)/col-WRKY33-DAP-Seq(GSE60143)/Homer | 1e-48 | -1.122e+02 | 0.0000 | 4912.0 | 12.41% | 18001.6 | 9.87% | motif file (matrix) | svg |
| 173 | T A G C C G T A C T G A T A C G C G T A A C G T A C T G A C T G A G T C T A C G C T A G G T A C | YY1(Zf)/Promoter/Homer | 1e-47 | -1.093e+02 | 0.0000 | 457.0 | 1.15% | 866.8 | 0.48% | motif file (matrix) | svg |
| 174 | C A G T T C A G T C G A A G T C C G T A A C T G T G A C C G A T A C T G A C T G A C G T A T C G | Atoh7(bHLH)/Retina-Atoh7-CutnRun(GSE156756)/Homer | 1e-47 | -1.092e+02 | 0.0000 | 2511.0 | 6.34% | 8298.3 | 4.55% | motif file (matrix) | svg |
| 175 | C T A G T C G A T G A C A G T C C G T A A C T G G T A C A C G T A C T G A C T G | BHLHA15(bHLH)/NIH3T3-BHLHB8.HA-ChIP-Seq(GSE119782)/Homer | 1e-46 | -1.065e+02 | 0.0000 | 4682.0 | 11.83% | 17159.3 | 9.40% | motif file (matrix) | svg |
| 176 | C G A T C G A T G C A T G A C T A C G T C G T A C G T A A C T G T A G C C G T A C G T A C G T A | AT5G60130(ABI3VP1)/col-AT5G60130-DAP-Seq(GSE60143)/Homer | 1e-46 | -1.060e+02 | 0.0000 | 5128.0 | 12.95% | 19029.8 | 10.43% | motif file (matrix) | svg |
| 177 | A T G C G T A C C G T A A G C T G C A T T A C G A G C T A G C T A G T C A G C T | Sox6(HMG)/Myotubes-Sox6-ChIP-Seq(GSE32627)/Homer | 1e-46 | -1.060e+02 | 0.0000 | 7805.0 | 19.71% | 30423.8 | 16.67% | motif file (matrix) | svg |
| 178 | C G A T C G T A A G T C A C G T A C G T T C A G G C A T C G T A G C T A G C T A C G T A A G T C C G T A G T C A A C T G | ANAC058(NAC)/col-ANAC058-DAP-Seq(GSE60143)/Homer | 1e-45 | -1.059e+02 | 0.0000 | 3180.0 | 8.03% | 11012.6 | 6.04% | motif file (matrix) | svg |
| 179 | C G A T T G C A T G C A G A T C C G T A A C T G T G A C G A C T C A T G A C T G | Tcf21(bHLH)/ArterySmoothMuscle-Tcf21-ChIP-Seq(GSE61369)/Homer | 1e-45 | -1.059e+02 | 0.0000 | 2809.0 | 7.09% | 9526.8 | 5.22% | motif file (matrix) | svg |
| 180 | T C A G A G C T G T C A C G T A A C G T A T G C C G T A A C G T A C G T C T G A | PHV(HB)/col-PHV-DAP-Seq(GSE60143)/Homer | 1e-45 | -1.049e+02 | 0.0000 | 1907.0 | 4.82% | 6010.6 | 3.29% | motif file (matrix) | svg |
| 181 | G T A C C T G A T A G C C G T A G C T A T C G A T G C A T G A C C T A G G T C A A G T C C G T A C T G A C T G A C G T A | At1g14580(C2H2)/colamp-At1g14580-DAP-Seq(GSE60143)/Homer | 1e-45 | -1.047e+02 | 0.0000 | 856.0 | 2.16% | 2169.3 | 1.19% | motif file (matrix) | svg |
| 182 | G C T A A G T C T A C G T G C A A T C G T C A G G C T A T C G A T C A G A G C T | ELF5(ETS)/T47D-ELF5-ChIP-Seq(GSE30407)/Homer | 1e-44 | -1.026e+02 | 0.0000 | 3327.0 | 8.40% | 11661.2 | 6.39% | motif file (matrix) | svg |
| 183 | A T C G T G C A G A T C C T A G A C G T A T C G C G T A A G T C T C A G A C T G T C A G G C T A | Knotted(Homeobox)/Corn-KN1-ChIP-Seq(GSE39161)/Homer | 1e-44 | -1.026e+02 | 0.0000 | 10990.0 | 27.76% | 44428.9 | 24.35% | motif file (matrix) | svg |
| 184 | C T A G G T A C C A T G G A C T C G A T C A T G G T C A G T A C G A C T C G A T C G A T C G A T | WRKY27(WRKY)/colamp-WRKY27-DAP-Seq(GSE60143)/Homer | 1e-44 | -1.023e+02 | 0.0000 | 4505.0 | 11.38% | 16507.8 | 9.05% | motif file (matrix) | svg |
| 185 | T C A G T G A C G T A C C G T A A C G T T G A C A C G T T C A G A G C T G A C T | NeuroD1(bHLH)/Islet-NeuroD1-ChIP-Seq(GSE30298)/Homer | 1e-44 | -1.020e+02 | 0.0000 | 2836.0 | 7.16% | 9694.4 | 5.31% | motif file (matrix) | svg |
| 186 | C G A T T C G A A T G C C G A T G C A T T C G A A G C T G C A T G C A T C G A T T C G A A G C T T C G A G T C A C T A G | ANAC004(NAC)/colamp-ANAC004-DAP-Seq(GSE60143)/Homer | 1e-43 | -1.012e+02 | 0.0000 | 1384.0 | 3.50% | 4083.8 | 2.24% | motif file (matrix) | svg |
| 187 | T G A C A G T C C G T A A C T G G T A C A C G T A C T G A C G T G A C T G A T C | Twist2(bHLH)/Myoblast-Twist2.Ty1-ChIP-Seq(GSE127998)/Homer | 1e-43 | -9.990e+01 | 0.0000 | 6153.0 | 15.54% | 23496.3 | 12.88% | motif file (matrix) | svg |
| 188 | C A T G A T G C T A G C C T G A A G T C A G T C A C T G G C T A A G T C G T A C G C T A G C A T | At4g28140(AP2EREBP)/colamp-At4g28140-DAP-Seq(GSE60143)/Homer | 1e-42 | -9.822e+01 | 0.0000 | 3607.0 | 9.11% | 12879.8 | 7.06% | motif file (matrix) | svg |
| 189 | C G A T C T A G A C G T A C G T A C G T C G T A A G C T C G A T A G C T C G T A C T A G T A G C | FoxD3(forkhead)/ZebrafishEmbryo-Foxd3.biotin-ChIP-seq(GSE106676)/Homer | 1e-42 | -9.728e+01 | 0.0000 | 2578.0 | 6.51% | 8737.7 | 4.79% | motif file (matrix) | svg |
| 190 | T C G A G T A C C A T G A G C T A C G T C A T G G T C A G T A C A G C T G C T A C G A T C A G T | WRKY31(WRKY)/colamp-WRKY31-DAP-Seq(GSE60143)/Homer | 1e-42 | -9.721e+01 | 0.0000 | 3843.0 | 9.71% | 13867.9 | 7.60% | motif file (matrix) | svg |
| 191 | C A T G G T A C A C T G G T C A A G C T T A C G T G C A A T C G T G A C C A G T | TOD6?/SacCer-Promoters/Homer | 1e-42 | -9.672e+01 | 0.0000 | 1438.0 | 3.63% | 4330.6 | 2.37% | motif file (matrix) | svg |
| 192 | G T C A T G C A G C T A A G T C C G T A A C T G T G A C G C A T T C A G C A G T | Ap4(bHLH)/AML-Tfap4-ChIP-Seq(GSE45738)/Homer | 1e-41 | -9.575e+01 | 0.0000 | 3464.0 | 8.75% | 12339.5 | 6.76% | motif file (matrix) | svg |
| 193 | A T G C C A T G A G C T C A G T C A T G T C G A A G T C G A C T C G A T C G A T C A G T C A G T | WRKY26(WRKY)/colamp-WRKY26-DAP-Seq(GSE60143)/Homer | 1e-40 | -9.417e+01 | 0.0000 | 3001.0 | 7.58% | 10484.2 | 5.75% | motif file (matrix) | svg |
| 194 | A G T C A C G T A C G T T A C G G C T A G C T A C G T A C G A T C G A T A T G C C G T A G T C A A C T G G A C T G C A T | SND2(NAC)/colamp-SND2-DAP-Seq(GSE60143)/Homer | 1e-40 | -9.286e+01 | 0.0000 | 3342.0 | 8.44% | 11893.6 | 6.52% | motif file (matrix) | svg |
| 195 | T C G A T C G A A G T C C G T A C T A G T A G C A C G T A C T G | MyoG(bHLH)/C2C12-MyoG-ChIP-Seq(GSE36024)/Homer | 1e-40 | -9.254e+01 | 0.0000 | 3247.0 | 8.20% | 11511.3 | 6.31% | motif file (matrix) | svg |
| 196 | T C G A A C T G C A T G A G C T A G T C C G T A C T G A C T A G A C T G C G A T A T G C C T G A | RAR:RXR(NR),DR0/ES-RAR-ChIP-Seq(GSE56893)/Homer | 1e-40 | -9.230e+01 | 0.0000 | 597.0 | 1.51% | 1388.4 | 0.76% | motif file (matrix) | svg |
| 197 | C G A T T C G A A G T C A C G T A C G T T A C G C G T A G C T A G C T A C G A T G C A T A T G C C G T A G T C A A C T G | ANAC071(NAC)/col-ANAC071-DAP-Seq(GSE60143)/Homer | 1e-40 | -9.224e+01 | 0.0000 | 4943.0 | 12.48% | 18552.6 | 10.17% | motif file (matrix) | svg |
| 198 | A G C T A G C T A G C T A C T G A C G T A G T C A C T G A C G T G A C T C G A T G C A T A T C G | IDD7(C2H2)/col-IDD7-DAP-Seq(GSE60143)/Homer | 1e-40 | -9.213e+01 | 0.0000 | 2016.0 | 5.09% | 6596.9 | 3.62% | motif file (matrix) | svg |
| 199 | T C G A T A G C T G C A A C T G A C T G C G T A C G T A C T A G G A C T T A C G | ETS1(ETS)/Jurkat-ETS1-ChIP-Seq(GSE17954)/Homer | 1e-39 | -9.107e+01 | 0.0000 | 4555.0 | 11.50% | 16949.6 | 9.29% | motif file (matrix) | svg |
| 200 | C A T G T G C A G A C T C A T G C G T A A G T C T C A G G C A T T G A C C G T A | bZIP50(bZIP)/colamp-bZIP50-DAP-Seq(GSE60143)/Homer | 1e-39 | -9.085e+01 | 0.0000 | 8700.0 | 21.97% | 34727.6 | 19.03% | motif file (matrix) | svg |
| 201 | A T G C C A T G A C G T A C G T A C T G C G T A A G T C G A C T G C A T C G A T | WRKY71(WRKY)/col-WRKY71-DAP-Seq(GSE60143)/Homer | 1e-39 | -9.083e+01 | 0.0000 | 4020.0 | 10.15% | 14724.3 | 8.07% | motif file (matrix) | svg |
| 202 | G A C T C G A T T C A G G A T C G A C T A G C T A G C T A G T C G A T C C G T A C T A G C T A G T C G A T C G A C T G A | Bcl6(Zf)/Liver-Bcl6-ChIP-Seq(GSE31578)/Homer | 1e-39 | -9.029e+01 | 0.0000 | 3299.0 | 8.33% | 11764.9 | 6.45% | motif file (matrix) | svg |
| 203 | T C A G T G A C C A T G G C A T C A G T A C T G C G T A T G A C G A C T C G A T C G A T C G T A | WRKY3(WRKY)/col-WRKY3-DAP-Seq(GSE60143)/Homer | 1e-38 | -8.887e+01 | 0.0000 | 3504.0 | 8.85% | 12631.4 | 6.92% | motif file (matrix) | svg |
| 204 | G A T C C T G A G A T C G A T C C T A G G C T A A G T C C T G A G C T A C G T A | At4g16750(AP2EREBP)/col-At4g16750-DAP-Seq(GSE60143)/Homer | 1e-38 | -8.883e+01 | 0.0000 | 9462.0 | 23.90% | 38124.5 | 20.89% | motif file (matrix) | svg |
| 205 | T C G A T G A C G T A C C G T A C A G T T G A C A C G T A C T G A G C T A G C T | NeuroG2(bHLH)/Fibroblast-NeuroG2-ChIP-Seq(GSE75910)/Homer | 1e-38 | -8.882e+01 | 0.0000 | 5531.0 | 13.97% | 21118.5 | 11.57% | motif file (matrix) | svg |
| 206 | C T A G C T G A C G T A C G T A C G T A C G T A A C T G A C G T C T A G G T C A | COG1(C2C2dof)/col-COG1-DAP-Seq(GSE60143)/Homer | 1e-38 | -8.831e+01 | 0.0000 | 6075.0 | 15.34% | 23449.5 | 12.85% | motif file (matrix) | svg |
| 207 | T G A C C T G A A G T C A G T C A C T G G A T C G A C T G C A T | At5g18450(AP2EREBP)/col-At5g18450-DAP-Seq(GSE60143)/Homer | 1e-38 | -8.800e+01 | 0.0000 | 12006.0 | 30.32% | 49401.4 | 27.08% | motif file (matrix) | svg |
| 208 | C T G A A T G C G C T A G C A T A T G C C G T A T C G A C T G A C T A G T C A G T A C G G T C A | Tcf4(HMG)/Hct116-Tcf4-ChIP-Seq(SRA012054)/Homer | 1e-38 | -8.784e+01 | 0.0000 | 2034.0 | 5.14% | 6726.6 | 3.69% | motif file (matrix) | svg |
| 209 | A G T C C T A G A C G T A C G T A C T G C G T A A G T C A G C T G C T A G C A T | WRKY24(WRKY)/colamp-WRKY24-DAP-Seq(GSE60143)/Homer | 1e-38 | -8.772e+01 | 0.0000 | 4762.0 | 12.03% | 17892.5 | 9.81% | motif file (matrix) | svg |
| 210 | C T A G C T A G T G A C G T A C C A T G A C T G G A T C G A T C C G T A C G T A | RAP211(AP2EREBP)/colamp-RAP211-DAP-Seq(GSE60143)/Homer | 1e-37 | -8.683e+01 | 0.0000 | 14367.0 | 36.29% | 60012.2 | 32.89% | motif file (matrix) | svg |
| 211 | T C G A A G T C A C G T A C G T T C A G C A G T C T G A C T A G T C G A C G T A A T C G C G T A C G T A A C T G A G C T | NTM1(NAC)/col-NTM1-DAP-Seq(GSE60143)/Homer | 1e-37 | -8.666e+01 | 0.0000 | 1836.0 | 4.64% | 5969.4 | 3.27% | motif file (matrix) | svg |
| 212 | A T G C A G T C G A T C C G T A A C G T A C G T A C T G A C G T A G C T G A T C | Sox2(HMG)/mES-Sox2-ChIP-Seq(GSE11431)/Homer | 1e-37 | -8.655e+01 | 0.0000 | 4081.0 | 10.31% | 15066.8 | 8.26% | motif file (matrix) | svg |
| 213 | C T G A T A C G G C A T A G C T A G C T A G T C T C G A A C T G C A G T A G C T A G C T G A T C | IRF3(IRF)/BMDM-Irf3-ChIP-Seq(GSE67343)/Homer | 1e-37 | -8.625e+01 | 0.0000 | 995.0 | 2.51% | 2808.0 | 1.54% | motif file (matrix) | svg |
| 214 | T A G C C A T G G A C T G A C T T C A G G T C A G A T C G A C T G C A T G C T A | WRKY15(WRKY)/col-WRKY15-DAP-Seq(GSE60143)/Homer | 1e-37 | -8.591e+01 | 0.0000 | 5350.0 | 13.51% | 20420.9 | 11.19% | motif file (matrix) | svg |
| 215 | T A C G T C G A C G T A C G T A C G T A C T G A A C T G A C G T C G T A T C G A | AT2G28810(C2C2dof)/colamp-AT2G28810-DAP-Seq(GSE60143)/Homer | 1e-37 | -8.572e+01 | 0.0000 | 10090.0 | 25.48% | 40986.8 | 22.46% | motif file (matrix) | svg |
| 216 | G A T C C T G A A G T C G T A C A C T G G C T A G A T C C T G A G C T A G C T A | At4g31060(AP2EREBP)/colamp-At4g31060-DAP-Seq(GSE60143)/Homer | 1e-37 | -8.546e+01 | 0.0000 | 5151.0 | 13.01% | 19587.4 | 10.74% | motif file (matrix) | svg |
| 217 | C T A G A G T C T A C G T A C G T G A C C G T A A C T G T A G C G C A T C A T G A T G C A G C T | Ascl1(bHLH)/NeuralTubes-Ascl1-ChIP-Seq(GSE55840)/Homer | 1e-36 | -8.350e+01 | 0.0000 | 5096.0 | 12.87% | 19401.4 | 10.63% | motif file (matrix) | svg |
| 218 | A G T C T G A C C T G A A G T C A G T C A C T G C G T A A G T C G T C A G C T A G C A T C G T A G C A T G C T A C G T A | DEAR3(AP2EREBP)/colamp-DEAR3-DAP-Seq(GSE60143)/Homer | 1e-35 | -8.286e+01 | 0.0000 | 4349.0 | 10.98% | 16266.8 | 8.92% | motif file (matrix) | svg |
| 219 | G A T C C T G A A G T C A G C T A C G T A C G T A C G T A C G T | At1g64620(C2C2dof)/colamp-At1g64620-DAP-Seq(GSE60143)/Homer | 1e-35 | -8.262e+01 | 0.0000 | 6474.0 | 16.35% | 25310.1 | 13.87% | motif file (matrix) | svg |
| 220 | C G A T T C A G G T A C A C G T A C G T T C A G C G A T C G T A G T C A G C T A C G T A A G T C C G T A G T C A C A T G | ANAC057(NAC)/colamp-ANAC057-DAP-Seq(GSE60143)/Homer | 1e-35 | -8.216e+01 | 0.0000 | 4252.0 | 10.74% | 15875.2 | 8.70% | motif file (matrix) | svg |
| 221 | A C G T T C G A T C G A A G T C G T C A T A C G A T G C A C G T A C T G A G C T | Myf5(bHLH)/GM-Myf5-ChIP-Seq(GSE24852)/Homer | 1e-35 | -8.216e+01 | 0.0000 | 1995.0 | 5.04% | 6654.5 | 3.65% | motif file (matrix) | svg |
| 222 | G A C T T C A G G C A T A G T C G C T A G A T C C T G A A C G T A G T C G T C A | Replumless(BLH)/Arabidopsis-RPL.GFP-ChIP-Seq(GSE78727)/Homer | 1e-35 | -8.136e+01 | 0.0000 | 7283.0 | 18.40% | 28841.0 | 15.81% | motif file (matrix) | svg |
| 223 | A G T C A G T C C T G A A G T C A G T C A C T G C G T A A G T C C T G A T C G A G C A T G A T C C G A T C G A T A C T G | AT3G60490(AP2EREBP)/colamp-AT3G60490-DAP-Seq(GSE60143)/Homer | 1e-35 | -8.079e+01 | 0.0000 | 3901.0 | 9.85% | 14438.4 | 7.91% | motif file (matrix) | svg |
| 224 | C G T A C G T A C G A T A C T G C A G T A G T C A C T G A C T G A G C T A C T G | DREB19(AP2EREBP)/colamp-DREB19-DAP-Seq(GSE60143)/Homer | 1e-34 | -7.930e+01 | 0.0000 | 6024.0 | 15.22% | 23466.3 | 12.86% | motif file (matrix) | svg |
| 225 | C T A G C T A G C G T A C G T A T A C G C G A T C T A G C T G A C T G A C G T A T A C G G A C T | PU.1:IRF8(ETS:IRF)/pDC-Irf8-ChIP-Seq(GSE66899)/Homer | 1e-33 | -7.826e+01 | 0.0000 | 669.0 | 1.69% | 1724.9 | 0.95% | motif file (matrix) | svg |
| 226 | A C T G T C A G A G C T G A C T C A T G A G T C A G T C G C T A C G A T C T A G T C A G G T A C C T G A T C G A | Rfx1(HTH)/NPC-H3K4me1-ChIP-Seq(GSE16256)/Homer | 1e-33 | -7.785e+01 | 0.0000 | 867.0 | 2.19% | 2423.6 | 1.33% | motif file (matrix) | svg |
| 227 | A G T C A C G T A C G T T C A G G C T A C G T A G C T A G C A T C G A T A G T C C G T A G T C A A C T G G A C T G C T A | SMB(NAC)/colamp-SMB-DAP-Seq(GSE60143)/Homer | 1e-33 | -7.759e+01 | 0.0000 | 5011.0 | 12.66% | 19185.9 | 10.52% | motif file (matrix) | svg |
| 228 | C G A T T G A C C A T G G A C T A C G T C A T G C G T A G A T C G A C T G C A T G C A T C G A T | WRKY14(WRKY)/colamp-WRKY14-DAP-Seq(GSE60143)/Homer | 1e-33 | -7.750e+01 | 0.0000 | 2620.0 | 6.62% | 9229.2 | 5.06% | motif file (matrix) | svg |
| 229 | G C T A G C T A C G T A C G T A C T G A C T A G A C G T A G T C C G T A C T G A G T A C A C T G | WRKY65(WRKY)/colamp-WRKY65-DAP-Seq(GSE60143)/Homer | 1e-33 | -7.742e+01 | 0.0000 | 2696.0 | 6.81% | 9539.8 | 5.23% | motif file (matrix) | svg |
| 230 | C G A T C G T A G C T A G C A T G C A T C T G A A C T G A C G T A G T C C G T A C G T A G T A C T C A G G C T A C G A T | WRKY25(WRKY)/colamp-WRKY25-DAP-Seq(GSE60143)/Homer | 1e-33 | -7.730e+01 | 0.0000 | 6094.0 | 15.39% | 23821.3 | 13.06% | motif file (matrix) | svg |
| 231 | T C A G A T C G G A C T A C T G G A C T C A G T C T A G C G T A G T A C C G T A C T A G A T C G | Tbx20(T-box)/Heart-Tbx20-ChIP-Seq(GSE29636)/Homer | 1e-33 | -7.640e+01 | 0.0000 | 1087.0 | 2.75% | 3240.3 | 1.78% | motif file (matrix) | svg |
| 232 | G T A C C A T G T A G C A G T C C T A G C A T G C T G A C G T A G C A T G C A T A C G T G C A T G T A C A C T G A T C G | LOB(LOBAS2)/col-LOB-DAP-Seq(GSE60143)/Homer | 1e-32 | -7.571e+01 | 0.0000 | 2766.0 | 6.99% | 9855.9 | 5.40% | motif file (matrix) | svg |
| 233 | G C T A C G T A C G A T C A G T A C T G C G A T G T A C A C T G A T C G G A C T C A T G C T A G G C A T C A G T C A T G | DEAR5(AP2EREBP)/col-DEAR5-DAP-Seq(GSE60143)/Homer | 1e-32 | -7.534e+01 | 0.0000 | 2169.0 | 5.48% | 7449.9 | 4.08% | motif file (matrix) | svg |
| 234 | C G T A G C T A C G A T C T A G A C G T G T C A C G T A C G T A A G T C C G T A T G C A T A C G | FoxL2(Forkhead)/Ovary-FoxL2-ChIP-Seq(GSE60858)/Homer | 1e-32 | -7.482e+01 | 0.0000 | 2851.0 | 7.20% | 10219.7 | 5.60% | motif file (matrix) | svg |
| 235 | T C A G C T G A C G T A C G T A T A C G G C A T C T A G C T G A C G T A C G T A T A C G G A C T | IRF1(IRF)/PBMC-IRF1-ChIP-Seq(GSE43036)/Homer | 1e-32 | -7.443e+01 | 0.0000 | 472.0 | 1.19% | 1091.6 | 0.60% | motif file (matrix) | svg |
| 236 | C G A T C T A G A G T C A C G T A C G T T C A G G C T A C G T A G C A T G C A T C G A T A G T C C G T A G T C A A C T G | VND3(NAC)/colamp-VND3-DAP-Seq(GSE60143)/Homer | 1e-32 | -7.420e+01 | 0.0000 | 3312.0 | 8.37% | 12129.1 | 6.65% | motif file (matrix) | svg |
| 237 | C G A T G C T A G C T A G C A T G C T A C G T A A G T C A C G T A C G T A C G T C G A T A G C T | At5g62940(C2C2dof)/col-At5g62940-DAP-Seq(GSE60143)/Homer | 1e-32 | -7.408e+01 | 0.0000 | 14356.0 | 36.26% | 60446.3 | 33.13% | motif file (matrix) | svg |
| 238 | G A C T A C G T A G C T G A C T A C T G C A G T A G T C A T C G A C G T G C A T G C A T G C A T | MGP(C2H2)/colamp-MGP-DAP-Seq(GSE60143)/Homer | 1e-32 | -7.382e+01 | 0.0000 | 1725.0 | 4.36% | 5713.0 | 3.13% | motif file (matrix) | svg |
| 239 | C T A G G C T A A G T C A C T G A C G T G A C T G A C T A T G C T C G A C A G T G A T C C G A T G A C T G A T C G A T C | RKD2(RWPRK)/colamp-RKD2-DAP-Seq(GSE60143)/Homer | 1e-31 | -7.323e+01 | 0.0000 | 3916.0 | 9.89% | 14667.5 | 8.04% | motif file (matrix) | svg |
| 240 | T A G C G T A C C T A G C A G T T C G A C G T A C G T A G C A T G A C T T G A C A G T C A C T G A T C G A G T C C T A G | AS2(LOBAS2)/col-AS2-DAP-Seq(GSE60143)/Homer | 1e-31 | -7.287e+01 | 0.0000 | 965.0 | 2.44% | 2826.7 | 1.55% | motif file (matrix) | svg |
| 241 | G A T C C G T A G A C T C T A G G A T C C T G A G A C T C T G A G A C T C T A G G A T C C T G A G A C T C T G A G A C T | OCT:OCT(POU,Homeobox)/NPC-OCT6-ChIP-Seq(GSE43916)/Homer | 1e-31 | -7.272e+01 | 0.0000 | 211.0 | 0.53% | 321.6 | 0.18% | motif file (matrix) | svg |
| 242 | A T G C A G T C A G C T A G C T A C G T A T C G C G T A C G A T T A G C G A C T | LEF1(HMG)/H1-LEF1-ChIP-Seq(GSE64758)/Homer | 1e-31 | -7.184e+01 | 0.0000 | 2750.0 | 6.95% | 9862.9 | 5.41% | motif file (matrix) | svg |
| 243 | T A C G T G C A A G T C C G T A A C G T T G A C A C G T A C T G A C T G G C A T | TCF4(bHLH)/SHSY5Y-TCF4-ChIP-Seq(GSE96915)/Homer | 1e-31 | -7.163e+01 | 0.0000 | 5310.0 | 13.41% | 20613.0 | 11.30% | motif file (matrix) | svg |
| 244 | G C A T T G A C C T G A A G T C A G T C A C T G G T C A A G T C G C T A G A C T G C T A C T G A | DREB2(AP2EREBP)/col-DREB2-DAP-Seq(GSE60143)/Homer | 1e-30 | -7.134e+01 | 0.0000 | 4980.0 | 12.58% | 19212.5 | 10.53% | motif file (matrix) | svg |
| 245 | G C T A T C G A C G T A C T G A A C T G A C G T A G T C C G T A C G T A A G T C C T A G T G C A | WRKY42(WRKY)/colamp-WRKY42-DAP-Seq(GSE60143)/Homer | 1e-29 | -6.884e+01 | 0.0000 | 2596.0 | 6.56% | 9289.3 | 5.09% | motif file (matrix) | svg |
| 246 | C T G A T C G A C G T A A T G C C G T A C G T A C G A T C T A G T C A G G A T C | Sox15(HMG)/CPA-Sox15-ChIP-Seq(GSE62909)/Homer | 1e-29 | -6.863e+01 | 0.0000 | 4360.0 | 11.01% | 16647.7 | 9.12% | motif file (matrix) | svg |
| 247 | G A C T A C T G C G T A A G T C T C A G G C A T G T A C C G T A A C G T G A T C | TGA1(bZIP)/colamp-TGA1-DAP-Seq(GSE60143)/Homer | 1e-29 | -6.819e+01 | 0.0000 | 2617.0 | 6.61% | 9387.4 | 5.14% | motif file (matrix) | svg |
| 248 | A C G T A G T C A G T C C G A T A C G T A C G T A C T G A C G T A T G C G A C T A C T G T A C G | Sox21(HMG)/ESC-SOX21-ChIP-Seq(GSE110505)/Homer | 1e-29 | -6.746e+01 | 0.0000 | 7776.0 | 19.64% | 31418.7 | 17.22% | motif file (matrix) | svg |
| 249 | A T G C G C A T T A G C C G A T T A G C G C A T T A G C G C A T A T G C G A C T | GAGA-repeat/Arabidopsis-Promoters/Homer | 1e-29 | -6.740e+01 | 0.0000 | 3582.0 | 9.05% | 13400.6 | 7.34% | motif file (matrix) | svg |
| 250 | C G A T C G T A A G T C A C G T A C G T T C A G C G T A C G T A G C A T G C A T G C A T A G T C C G T A G T C A A C T G | VND2(NAC)/col-VND2-DAP-Seq(GSE60143)/Homer | 1e-28 | -6.651e+01 | 0.0000 | 5023.0 | 12.69% | 19524.7 | 10.70% | motif file (matrix) | svg |
| 251 | C A G T T C A G A G C T G A C T A C G T A G T C G A T C G A C T C T G A A C T G G A T C C G T A C T G A A G T C G T A C | Rfx6(HTH)/Min6b1-Rfx6.HA-ChIP-Seq(GSE62844)/Homer | 1e-28 | -6.535e+01 | 0.0000 | 4254.0 | 10.74% | 16280.5 | 8.92% | motif file (matrix) | svg |
| 252 | A T G C G A C T A C G T C T A G A C G T A C G T A C G T C T G A G A T C G C T A A G C T C G T A | Foxa2(Forkhead)/Liver-Foxa2-ChIP-Seq(GSE25694)/Homer | 1e-28 | -6.507e+01 | 0.0000 | 2953.0 | 7.46% | 10832.7 | 5.94% | motif file (matrix) | svg |
| 253 | G C T A C G T A G C T A C G T A C T G A A C T G A C G T G T A C C G T A C T G A G T A C A C T G | WRKY22(WRKY)/colamp-WRKY22-DAP-Seq(GSE60143)/Homer | 1e-28 | -6.494e+01 | 0.0000 | 2897.0 | 7.32% | 10603.7 | 5.81% | motif file (matrix) | svg |
| 254 | G T A C G T A C G T C A G C T A C G T A C G T A C G T A C T A G C T A G C T A G | SEP3(MADS)/Arabidoposis-Flower-Sep3-ChIP-Seq/Homer | 1e-27 | -6.392e+01 | 0.0000 | 4980.0 | 12.58% | 19411.7 | 10.64% | motif file (matrix) | svg |
| 255 | C G A T C G A T G C A T G C A T G T C A A G T C A G C T A C G T A C G T C G A T G A C T A C G T | OBP4(C2C2dof)/col-OBP4-DAP-Seq(GSE60143)/Homer | 1e-27 | -6.380e+01 | 0.0000 | 5620.0 | 14.19% | 22164.4 | 12.15% | motif file (matrix) | svg |
| 256 | A G C T G T C A T G C A A G T C A C G T A C G T A C G T C G A T G A C T T A C G | AT3G12130(C3H)/colamp-AT3G12130-DAP-Seq(GSE60143)/Homer | 1e-27 | -6.379e+01 | 0.0000 | 10587.0 | 26.74% | 43963.1 | 24.09% | motif file (matrix) | svg |
| 257 | T C G A T A G C G T C A A C T G A C T G C G T A C G T A C T A G A G C T T C A G | ERG(ETS)/VCaP-ERG-ChIP-Seq(GSE14097)/Homer | 1e-27 | -6.374e+01 | 0.0000 | 5248.0 | 13.26% | 20565.1 | 11.27% | motif file (matrix) | svg |
| 258 | C T G A A G T C G A T C C A T G G C T A G A T C C T G A G C T A G C T A C G A T | AT1G77200(AP2EREBP)/colamp-AT1G77200-DAP-Seq(GSE60143)/Homer | 1e-27 | -6.287e+01 | 0.0000 | 9683.0 | 24.46% | 39983.6 | 21.91% | motif file (matrix) | svg |
| 259 | G A C T A C G T A C G T A C T G A C G T A G T C G C A T A G C T G C A T G C A T G A C T A G C T | SGR5(C2H2)/colamp-SGR5-DAP-Seq(GSE60143)/Homer | 1e-27 | -6.247e+01 | 0.0000 | 2721.0 | 6.87% | 9928.0 | 5.44% | motif file (matrix) | svg |
| 260 | C G T A C T G A C G T A C T A G T C G A C T A G A C T G C G T A C G T A T A C G A G C T A T C G | SpiB(ETS)/OCILY3-SPIB-ChIP-Seq(GSE56857)/Homer | 1e-26 | -6.203e+01 | 0.0000 | 964.0 | 2.43% | 2937.7 | 1.61% | motif file (matrix) | svg |
| 261 | A G T C C G A T A C T G A T C G T G A C G C T A C A T G A T C G T G A C C G A T A C T G T A G C G T A C G T C A | Tlx?(NR)/NPC-H3K4me1-ChIP-Seq(GSE16256)/Homer | 1e-26 | -6.159e+01 | 0.0000 | 1065.0 | 2.69% | 3323.8 | 1.82% | motif file (matrix) | svg |
| 262 | A G T C G A C T A G C T C G A T A T C G G C T A C G A T A T C G C G A T A C T G T A C G A C G T | Tcf7(HMG)/GM12878-TCF7-ChIP-Seq(Encode)/Homer | 1e-26 | -6.128e+01 | 0.0000 | 1432.0 | 3.62% | 4744.6 | 2.60% | motif file (matrix) | svg |
| 263 | G A T C G A T C A G T C G T A C C G A T G T A C G T A C A G T C A G T C A G T C G C T A G A T C | ZNF148(Zf)/MDAMB231-ZNF148-ChIP-Seq(GSE147020)/Homer | 1e-25 | -5.976e+01 | 0.0000 | 1888.0 | 4.77% | 6580.3 | 3.61% | motif file (matrix) | svg |
| 264 | A T G C A G T C C T G A A G T C C G A T A C G T A G T C A G T C A C G T A T C G G A C T A C G T | Etv2(ETS)/ES-ER71-ChIP-Seq(GSE59402)/Homer | 1e-25 | -5.885e+01 | 0.0000 | 3130.0 | 7.91% | 11704.7 | 6.41% | motif file (matrix) | svg |
| 265 | C T G A C T G A C T G A A T G C G A T C C A T G A C T G G A C T G A C T G C A T C G T A C G T A A G T C G T A C C T G A A T C G G C A T G A C T G A C T A G C T | GRHL2(CP2)/HBE-GRHL2-ChIP-Seq(GSE46194)/Homer | 1e-25 | -5.878e+01 | 0.0000 | 1551.0 | 3.92% | 5250.0 | 2.88% | motif file (matrix) | svg |
| 266 | C T A G T A C G G A T C G T A C G C T A A G C T A G C T G T C A T C G A T A G C | Nanog(Homeobox)/mES-Nanog-ChIP-Seq(GSE11724)/Homer | 1e-25 | -5.826e+01 | 0.0000 | 21159.0 | 53.44% | 92208.4 | 50.54% | motif file (matrix) | svg |
| 267 | C T G A A C T G C G T A A C G T G T C A A G C T A G C T G A C T G A C T C A G T | CCA(Myb)/Arabidopsis-CCA.GFP-ChIP-Seq(GSE70533)/Homer | 1e-25 | -5.810e+01 | 0.0000 | 3918.0 | 9.90% | 15041.2 | 8.24% | motif file (matrix) | svg |
| 268 | A C G T T G C A A G C T G A T C C T A G C T G A A G C T G T C A T C G A C G T A | CUX1(Homeobox)/K562-CUX1-ChIP-Seq(GSE92882)/Homer | 1e-24 | -5.750e+01 | 0.0000 | 5686.0 | 14.36% | 22640.7 | 12.41% | motif file (matrix) | svg |
| 269 | G C T A T C G A C G T A C T A G A G C T G T C A G T C A C G T A A G T C C G T A | FOXA1(Forkhead)/LNCAP-FOXA1-ChIP-Seq(GSE27824)/Homer | 1e-24 | -5.603e+01 | 0.0000 | 3391.0 | 8.56% | 12867.0 | 7.05% | motif file (matrix) | svg |
| 270 | G A C T C T G A A G T C A G T C A C T G C G T A A G T C C T G A | bHLH10(bHLH)/colamp-bHLH10-DAP-Seq(GSE60143)/Homer | 1e-24 | -5.571e+01 | 0.0000 | 3709.0 | 9.37% | 14217.9 | 7.79% | motif file (matrix) | svg |
| 271 | T A C G A T G C G A C T A C T G A G C T A G T C G T C A T G C A A C G T A G T C G C T A T G C A | Pknox1(Homeobox)/ES-Prep1-ChIP-Seq(GSE63282)/Homer | 1e-24 | -5.546e+01 | 0.0000 | 1095.0 | 2.77% | 3515.0 | 1.93% | motif file (matrix) | svg |
| 272 | C T A G C A T G A C G T C G T A C T A G C A T G C G A T C T A G T C A G T C A G | MYB3(MYB)/Arabidopsis-MYB3-ChIP-Seq(GSE80564)/Homer | 1e-23 | -5.517e+01 | 0.0000 | 10208.0 | 25.78% | 42620.8 | 23.36% | motif file (matrix) | svg |
| 273 | C A T G A C T G C T A G T C G A T C G A T C G A T C G A T C A G T C A G T C A G T G A C T G A C C G T A A C T G T G C A C G A T A C T G | RBPJ:Ebox(?,bHLH)/Panc1-Rbpj1-ChIP-Seq(GSE47459)/Homer | 1e-23 | -5.503e+01 | 0.0000 | 870.0 | 2.20% | 2663.3 | 1.46% | motif file (matrix) | svg |
| 274 | A G C T A G C T A G C T A C T G A C G T A G T C A C T G A C G T G C A T G C A T G C A T A C G T | At5g66730(C2H2)/colamp-At5g66730-DAP-Seq(GSE60143)/Homer | 1e-23 | -5.463e+01 | 0.0000 | 1450.0 | 3.66% | 4914.5 | 2.69% | motif file (matrix) | svg |
| 275 | G C A T C G A T C G T A G A T C C A T G A C G T A C G T A C T G C G T A A G T C A G C T G C A T G C A T C G T A G C T A | WRKY45(WRKY)/col-WRKY45-DAP-Seq(GSE60143)/Homer | 1e-23 | -5.460e+01 | 0.0000 | 1774.0 | 4.48% | 6210.3 | 3.40% | motif file (matrix) | svg |
| 276 | C T G A A T C G G T A C C T G A A G T C A G T C A C T G C G T A A G T C C T G A | TINY(AP2EREBP)/col-TINY-DAP-Seq(GSE60143)/Homer | 1e-23 | -5.460e+01 | 0.0000 | 3329.0 | 8.41% | 12640.8 | 6.93% | motif file (matrix) | svg |
| 277 | G A C T G C A T G C A T A G T C A G C T T C G A T A C G G C T A C G T A A C T G G T A C G C A T C G A T A G T C G A C T | HSF3(HSF)/colamp-HSF3-DAP-Seq(GSE60143)/Homer | 1e-23 | -5.393e+01 | 0.0000 | 3139.0 | 7.93% | 11857.1 | 6.50% | motif file (matrix) | svg |
| 278 | T C G A C G T A A G T C C G T A C T A G A G T C C G A T A C T G G A C T A G C T A C T G G A C T | HLH-1(bHLH)/cElegans-Embryo-HLH1-ChIP-Seq(modEncode)/Homer | 1e-23 | -5.388e+01 | 0.0000 | 2786.0 | 7.04% | 10381.9 | 5.69% | motif file (matrix) | svg |
| 279 | T C G A T C G A C T G A C G T A A C T G A T G C A C G T A G T C | Lola-I(Zf)/Embryo-LolaI-ChIP-Seq(GSE200870)/Homer | 1e-23 | -5.313e+01 | 0.0000 | 2186.0 | 5.52% | 7915.0 | 4.34% | motif file (matrix) | svg |
| 280 | G C A T A G C T A G C T A G C T A C T G A C G T A G T C A C T G A C G T G A C T C G A T G C A T | JKD(C2H2)/col-JKD-DAP-Seq(GSE60143)/Homer | 1e-23 | -5.303e+01 | 0.0000 | 1059.0 | 2.67% | 3407.8 | 1.87% | motif file (matrix) | svg |
| 281 | C A G T T C G A A G T C A C G T A C G T T C A G C G A T G C T A G C T A C G T A G C A T A G T C C G T A T G C A A C T G | ANAC045(NAC)/col-ANAC045-DAP-Seq(GSE60143)/Homer | 1e-22 | -5.277e+01 | 0.0000 | 10818.0 | 27.32% | 45444.6 | 24.91% | motif file (matrix) | svg |
| 282 | A G T C C G T A A C G T A G T C A C G T A C T G | Tal1 | 1e-22 | -5.251e+01 | 0.0000 | 5373.0 | 13.57% | 21443.2 | 11.75% | motif file (matrix) | svg |
| 283 | G T A C A C G T A C G T T A C G A T G C C A T G T A C G G T A C T C A G A T G C C G T A G T C A A C T G A G C T G C T A | AT1G19040(NAC)/col-AT1G19040-DAP-Seq(GSE60143)/Homer | 1e-22 | -5.219e+01 | 0.0000 | 695.0 | 1.76% | 2043.7 | 1.12% | motif file (matrix) | svg |
| 284 | C G A T A G T C C A T G G A C T A C G T C T A G C G T A G A T C G A C T C G A T G C A T G A C T | WRKY43(WRKY)/colamp-WRKY43-DAP-Seq(GSE60143)/Homer | 1e-22 | -5.216e+01 | 0.0000 | 1849.0 | 4.67% | 6557.5 | 3.59% | motif file (matrix) | svg |
| 285 | C G A T G A T C G A T C C T G A G A T C G A T C C A T G T G C A G T A C T C G A G T C A G C A T C G A T C G A T G C A T | At4g32800(AP2EREBP)/colamp-At4g32800-DAP-Seq(GSE60143)/Homer | 1e-22 | -5.212e+01 | 0.0000 | 1728.0 | 4.36% | 6068.3 | 3.33% | motif file (matrix) | svg |
| 286 | G T A C G C T A T C A G C T G A C T A G C A T G A G C T G A T C T G C A T C G A C T G A A C T G C A G T A G T C G A T C G C T A | HNF4a(NR),DR1/HepG2-HNF4a-ChIP-Seq(GSE25021)/Homer | 1e-22 | -5.207e+01 | 0.0000 | 1321.0 | 3.34% | 4443.1 | 2.44% | motif file (matrix) | svg |
| 287 | A T G C G T A C A C T G A G T C A G T C A C T G G A T C G T C A C G T A C G A T G C A T C G A T | RRTF1(AP2EREBP)/colamp-RRTF1-DAP-Seq(GSE60143)/Homer | 1e-22 | -5.185e+01 | 0.0000 | 2962.0 | 7.48% | 11164.2 | 6.12% | motif file (matrix) | svg |
| 288 | G C T A C T G A T C G A A G T C A G T C C T G A A G T C G T C A C T G A T G C A | RUNX1(Runt)/Jurkat-RUNX1-ChIP-Seq(GSE29180)/Homer | 1e-22 | -5.144e+01 | 0.0000 | 4643.0 | 11.73% | 18325.9 | 10.04% | motif file (matrix) | svg |
| 289 | A G C T G A T C C T G A A G T C A G T C A C T G C G T A A G T C C T G A G T C A G C A T C G A T G C T A G C A T C G T A | At2g44940(AP2EREBP)/colamp-At2g44940-DAP-Seq(GSE60143)/Homer | 1e-22 | -5.068e+01 | 0.0000 | 2956.0 | 7.47% | 11167.6 | 6.12% | motif file (matrix) | svg |
| 290 | T G A C C A T G A C G T A C G T A C T G C G T A A G T C A G C T G C A T T C G A | WRKY30(WRKY)/colamp-WRKY30-DAP-Seq(GSE60143)/Homer | 1e-21 | -5.043e+01 | 0.0000 | 2080.0 | 5.25% | 7533.2 | 4.13% | motif file (matrix) | svg |
| 291 | A G T C G T A C C T G A A G T C A G T C C A T G G C T A A G T C T G C A G C T A G C A T G C A T | RAP21(AP2EREBP)/colamp-RAP21-DAP-Seq(GSE60143)/Homer | 1e-21 | -4.977e+01 | 0.0000 | 2458.0 | 6.21% | 9108.5 | 4.99% | motif file (matrix) | svg |
| 292 | T C G A T G A C G C A T A G C T C A G T G A T C G C T A G A T C G A C T A C G T G C A T A G T C | PRDM1(Zf)/Hela-PRDM1-ChIP-Seq(GSE31477)/Homer | 1e-21 | -4.951e+01 | 0.0000 | 1733.0 | 4.38% | 6135.5 | 3.36% | motif file (matrix) | svg |
| 293 | G C T A C G T A C G A T G A C T G C A T T G C A A G T C A G C T A C G T A C G T C G A T G A C T | DAG2(C2C2dof)/col-DAG2-DAP-Seq(GSE60143)/Homer | 1e-21 | -4.854e+01 | 0.0000 | 5827.0 | 14.72% | 23548.0 | 12.91% | motif file (matrix) | svg |
| 294 | A G C T G C T A T G C A A G T C A C G T A C G T A C G T C G A T A G C T T C A G | dof24(C2C2dof)/col-dof24-DAP-Seq(GSE60143)/Homer | 1e-21 | -4.836e+01 | 0.0000 | 8812.0 | 22.26% | 36695.0 | 20.11% | motif file (matrix) | svg |
| 295 | G T A C A C G T A C G T T C A G C G T A C G T A C G A T G C A T G C A T A G T C C G T A G T C A C A T G G A C T G C T A | VND1(NAC)/col-VND1-DAP-Seq(GSE60143)/Homer | 1e-20 | -4.832e+01 | 0.0000 | 3898.0 | 9.85% | 15222.8 | 8.34% | motif file (matrix) | svg |
| 296 | T C A G G A C T G T C A C G T A A C G T A T C G C G T A A C G T A C G T C T G A | ATHB15(HB)/col-ATHB15-DAP-Seq(GSE60143)/Homer | 1e-20 | -4.821e+01 | 0.0000 | 1506.0 | 3.80% | 5242.6 | 2.87% | motif file (matrix) | svg |
| 297 | G C T A T G A C G A T C C G A T G A C T A T G C C T G A A T C G G C A T A C G T | JGL(C2H2)/col-JGL-DAP-Seq(GSE60143)/Homer | 1e-20 | -4.811e+01 | 0.0000 | 6477.0 | 16.36% | 26404.6 | 14.47% | motif file (matrix) | svg |
| 298 | C G T A A C T G C G T A A C G T A T C G C A G T T A G C C G T A T C G A G T A C C T G A T A G C C G T A A C T G C G T A A C G T C G T A C T G A A T C G G C T A | GATA3(Zf),DR8/iTreg-Gata3-ChIP-Seq(GSE20898)/Homer | 1e-20 | -4.800e+01 | 0.0000 | 386.0 | 0.97% | 980.7 | 0.54% | motif file (matrix) | svg |
| 299 | T G A C C T G A C T A G T C G A C T G A A T G C C G T A A C T G G C A T G T A C G C A T A T C G G C A T A G C T G A T C | PR(NR)/T47D-PR-ChIP-Seq(GSE31130)/Homer | 1e-20 | -4.772e+01 | 0.0000 | 7332.0 | 18.52% | 30175.6 | 16.54% | motif file (matrix) | svg |
| 300 | A G C T G A C T A C G T A C T G A C G T A G T C A C T G A C G T G C A T C G A T | AtIDD11(C2H2)/colamp-AtIDD11-DAP-Seq(GSE60143)/Homer | 1e-20 | -4.770e+01 | 0.0000 | 1957.0 | 4.94% | 7083.9 | 3.88% | motif file (matrix) | svg |
| 301 | T G A C T A G C T C A G T C G A T C G A C G T A A G T C C G T A C G T A C G A T C T A G T A C G | Sox7(HMG)/ESC-Sox7-ChIP-Seq(GSE133899)/Homer | 1e-20 | -4.725e+01 | 0.0000 | 1557.0 | 3.93% | 5464.9 | 3.00% | motif file (matrix) | svg |
| 302 | G T A C A C G T A C G T T C A G G C T A C G T A C G A T G C A T G C A T A G T C C G T A G T C A C A T G G A C T G C T A | ANAC070(NAC)/colamp-ANAC070-DAP-Seq(GSE60143)/Homer | 1e-20 | -4.705e+01 | 0.0000 | 5544.0 | 14.00% | 22367.6 | 12.26% | motif file (matrix) | svg |
| 303 | A T G C T C G A A G T C A G C T A C G T G T A C A G T C G C T A C T A G C A T G G T C A C T G A T C A G A G T C | Stat3+il21(Stat)/CD4-Stat3-ChIP-Seq(GSE19198)/Homer | 1e-20 | -4.683e+01 | 0.0000 | 2139.0 | 5.40% | 7850.5 | 4.30% | motif file (matrix) | svg |
| 304 | C T A G A C T G T G C A G T C A A T G C C G T A A T C G A T G C A G T C C T A G | ZNF341(Zf)/EBV-ZNF341-ChIP-Seq(GSE113194)/Homer | 1e-20 | -4.674e+01 | 0.0000 | 2917.0 | 7.37% | 11100.8 | 6.08% | motif file (matrix) | svg |
| 305 | T C G A A G C T A C G T A C G T A G T C A G T C A C G T A T C G G A C T A T C G | EWS:ERG-fusion(ETS)/CADO\_ES1-EWS:ERG-ChIP-Seq(SRA014231)/Homer | 1e-20 | -4.644e+01 | 0.0000 | 2033.0 | 5.13% | 7421.6 | 4.07% | motif file (matrix) | svg |
| 306 | C A G T T A G C A G T C C A T G C A G T C A T G C G A T C G A T G A C T C G A T A T C G G T A C A C T G A T C G G T A C | LBD13(LOBAS2)/colamp-LBD13-DAP-Seq(GSE60143)/Homer | 1e-20 | -4.627e+01 | 0.0000 | 7405.0 | 18.70% | 30554.4 | 16.75% | motif file (matrix) | svg |
| 307 | C G A T G T C A A G T C A C G T A C G T A C T G G A C T C G A T A T C G G C T A G T C A A G T C C G T A G T C A A C T G | ANAC017(NAC)/colamp-ANAC017-DAP-Seq(GSE60143)/Homer | 1e-20 | -4.614e+01 | 0.0000 | 793.0 | 2.00% | 2476.4 | 1.36% | motif file (matrix) | svg |
| 308 | C A G T C G T A C G T A G C A T G A C T G C A T G T A C A G C T A C T G G A C T A C G T C A T G | RAV1(RAV)/colamp-RAV1-DAP-Seq(GSE60143)/Homer | 1e-19 | -4.568e+01 | 0.0000 | 2477.0 | 6.26% | 9280.3 | 5.09% | motif file (matrix) | svg |
| 309 | G C A T A C G T A C G T A T C G C G T A C G T A C G T A C G T A | At2g41835(C2H2)/col-At2g41835-DAP-Seq(GSE60143)/Homer | 1e-19 | -4.519e+01 | 0.0000 | 1582.0 | 4.00% | 5602.8 | 3.07% | motif file (matrix) | svg |
| 310 | C G T A C G T A C G T A C T G A C T A G A C G T C T A G G T C A | CDF3(C2C2dof)/colamp-CDF3-DAP-Seq(GSE60143)/Homer | 1e-19 | -4.517e+01 | 0.0000 | 7290.0 | 18.41% | 30091.4 | 16.49% | motif file (matrix) | svg |
| 311 | C G T A C G A T C T A G G T C A G A C T C G A T C T A G C G T A A C G T C A T G | LIN-39(Homeobox)/cElegans.L3-LIN39-ChIP-Seq(modEncode)/Homer | 1e-19 | -4.506e+01 | 0.0000 | 4392.0 | 11.09% | 17441.5 | 9.56% | motif file (matrix) | svg |
| 312 | C A T G G A T C C T G A G T A C C T A G C T G A G C T A G C A T G A T C G A T C A G T C C T A G C G T A C A T G C T A G | AIL7(AP2EREBP)/colamp-AIL7-DAP-Seq(GSE60143)/Homer | 1e-19 | -4.501e+01 | 0.0000 | 3024.0 | 7.64% | 11596.5 | 6.36% | motif file (matrix) | svg |
| 313 | C G A T C T G A A G T C A C G T A C G T T C A G G C A T C G A T G C T A G C T A C G T A A G T C C G T A G T C A A C T G | CUC1(NAC)/col-CUC1-DAP-Seq(GSE60143)/Homer | 1e-19 | -4.422e+01 | 0.0000 | 2508.0 | 6.33% | 9444.9 | 5.18% | motif file (matrix) | svg |
| 314 | G T A C T C G A T A G C C G T A C G T A C T G A T G C A T G A C A C T G C G T A A G T C C T G A C T G A T C G A C G T A | NUC(C2H2)/col-NUC-DAP-Seq(GSE60143)/Homer | 1e-19 | -4.397e+01 | 0.0000 | 542.0 | 1.37% | 1565.1 | 0.86% | motif file (matrix) | svg |
| 315 | A G T C T A G C G A C T A C G T C T A G A C G T A C G T A C G T C T G A A G T C G C T A G A C T C G T A C T A G A C T G | Foxa3(Forkhead)/Liver-Foxa3-ChIP-Seq(GSE77670)/Homer | 1e-19 | -4.397e+01 | 0.0000 | 1043.0 | 2.63% | 3470.8 | 1.90% | motif file (matrix) | svg |
| 316 | G C T A G C T A C T G A A C T G A C G T A G T C C G T A C G T A G T A C A C T G A T G C G C A T | WRKY47(WRKY)/colamp-WRKY47-DAP-Seq(GSE60143)/Homer | 1e-18 | -4.373e+01 | 0.0000 | 1771.0 | 4.47% | 6400.5 | 3.51% | motif file (matrix) | svg |
| 317 | C G A T G A T C T A C G C T G A G C T A C G T A G C A T A G T C C T A G C G T A G C A T C G A T | AT2G15740(C2H2)/col-AT2G15740-DAP-Seq(GSE60143)/Homer | 1e-18 | -4.369e+01 | 0.0000 | 12278.0 | 31.01% | 52421.5 | 28.73% | motif file (matrix) | svg |
| 318 | C A T G C T A G A G C T G A C T C A T G A G T C G A T C G C T A C G A T C T A G T C A G G T A C C T G A T C G A | X-box(HTH)/NPC-H3K4me1-ChIP-Seq(GSE16256)/Homer | 1e-18 | -4.367e+01 | 0.0000 | 362.0 | 0.91% | 930.9 | 0.51% | motif file (matrix) | svg |
| 319 | T G C A C T G A A T G C G T C A A C G T A T G C A C G T A C T G A C T G T G C A | ZBTB18(Zf)/HEK293-ZBTB18.GFP-ChIP-Seq(GSE58341)/Homer | 1e-18 | -4.335e+01 | 0.0000 | 1467.0 | 3.71% | 5170.3 | 2.83% | motif file (matrix) | svg |
| 320 | G A T C G C A T T C G A A G T C A C G T A C G T A C G T C G A T A C G T A T C G | AT1G47655(C2C2dof)/colamp-AT1G47655-DAP-Seq(GSE60143)/Homer | 1e-18 | -4.325e+01 | 0.0000 | 14716.0 | 37.17% | 63474.9 | 34.79% | motif file (matrix) | svg |
| 321 | A T G C T C A G T C G A G C A T A C T G C G T A A G T C T C A G G A C T T G A C C G T A A G C T | Atf2(bZIP)/3T3L1-Atf2-ChIP-Seq(GSE56872)/Homer | 1e-18 | -4.320e+01 | 0.0000 | 1418.0 | 3.58% | 4975.5 | 2.73% | motif file (matrix) | svg |
| 322 | T C G A C T G A C G T A C G T A C G T A C T G A A C T G A C G T C T G A C T G A | AT5G63260(C3H)/col-AT5G63260-DAP-Seq(GSE60143)/Homer | 1e-18 | -4.311e+01 | 0.0000 | 9497.0 | 23.99% | 39974.3 | 21.91% | motif file (matrix) | svg |
| 323 | T A G C C A T G A G C T A C G T A C T G C G T A A G T C G A C T G C A T C T G A | AT3G42860(zfGRF)/col-AT3G42860-DAP-Seq(GSE60143)/Homer | 1e-18 | -4.284e+01 | 0.0000 | 1830.0 | 4.62% | 6660.2 | 3.65% | motif file (matrix) | svg |
| 324 | A G T C A T C G C T A G A G C T G A C T C T A G A G T C A G T C G C T A C A G T T C A G T C A G G A T C C T G A T C G A G A T C | RFX(HTH)/K562-RFX3-ChIP-Seq(SRA012198)/Homer | 1e-18 | -4.254e+01 | 0.0000 | 324.0 | 0.82% | 809.1 | 0.44% | motif file (matrix) | svg |
| 325 | C G T A T A C G T C G A A C T G A C T G C G T A C G T A T A C G A G C T T A C G | PU.1(ETS)/ThioMac-PU.1-ChIP-Seq(GSE21512)/Homer | 1e-18 | -4.200e+01 | 0.0000 | 1660.0 | 4.19% | 5980.9 | 3.28% | motif file (matrix) | svg |
| 326 | T C A G A G C T A C G T A C G T G T A C G A T C C G T A C T A G C A T G G T C A C G T A T C G A | STAT4(Stat)/CD4-Stat4-ChIP-Seq(GSE22104)/Homer | 1e-18 | -4.189e+01 | 0.0000 | 2697.0 | 6.81% | 10296.8 | 5.64% | motif file (matrix) | svg |
| 327 | G C A T C T A G G T A C A G T C C G A T A C T G C T A G C T A G G T A C G C T A | ZNF416(Zf)/HEK293-ZNF416.GFP-ChIP-Seq(GSE58341)/Homer | 1e-18 | -4.156e+01 | 0.0000 | 3380.0 | 8.54% | 13204.8 | 7.24% | motif file (matrix) | svg |
| 328 | C G A T C T A G T C A G C A G T C G T A A G T C G C T A A C G T G A C T A T G C A G T C G C T A | PRDM10(Zf)/HEK293-PRDM10.eGFP-ChIP-Seq(Encode)/Homer | 1e-17 | -4.052e+01 | 0.0000 | 2211.0 | 5.58% | 8290.8 | 4.54% | motif file (matrix) | svg |
| 329 | A G T C A G T C C T G A A G T C A G T C A C T G C G T A A G T C T C G A G A T C C G A T C G T A | AT1G01250(AP2EREBP)/col-AT1G01250-DAP-Seq(GSE60143)/Homer | 1e-17 | -4.046e+01 | 0.0000 | 1112.0 | 2.81% | 3796.4 | 2.08% | motif file (matrix) | svg |
| 330 | C T A G T C G A C T G A C G T A T A C G G A C T T C A G T C G A G T C A T G C A T A C G A G C T | IRF2(IRF)/Erythroblas-IRF2-ChIP-Seq(GSE36985)/Homer | 1e-17 | -3.996e+01 | 0.0000 | 417.0 | 1.05% | 1152.5 | 0.63% | motif file (matrix) | svg |
| 331 | T C A G T A C G T A G C A C G T A C T G C G A T A G T C C G T A T A C G A G T C | Meis1(Homeobox)/MastCells-Meis1-ChIP-Seq(GSE48085)/Homer | 1e-17 | -3.980e+01 | 0.0000 | 7565.0 | 19.11% | 31531.8 | 17.28% | motif file (matrix) | svg |
| 332 | T A C G C T G A C A T G G A T C G T A C G C A T T C A G T A C G A G C T G T C A G A T C G C A T T A C G C G T A C T A G G A T C G A T C C G A T A C T G T C A G | ZNF322(Zf)/HEK293-ZNF322.GFP-ChIP-Seq(GSE58341)/Homer | 1e-17 | -3.964e+01 | 0.0000 | 528.0 | 1.33% | 1557.0 | 0.85% | motif file (matrix) | svg |
| 333 | A T C G T C G A G A C T A T C G T G A C A C G T C T A G A C T G C G T A A C T G A G T C G T A C | ZNF415(Zf)/HEK293-ZNF415.GFP-ChIP-Seq(GSE58341)/Homer | 1e-16 | -3.890e+01 | 0.0000 | 2163.0 | 5.46% | 8128.9 | 4.46% | motif file (matrix) | svg |
| 334 | C T A G G T A C A C G T A C G T A T C G G C A T A G C T A G C T A G C T G C A T G A C T C G T A G T C A A C T G G A C T | VND6(NAC)/col-VND6-DAP-Seq(GSE60143)/Homer | 1e-16 | -3.886e+01 | 0.0000 | 6276.0 | 15.85% | 25879.4 | 14.18% | motif file (matrix) | svg |
| 335 | T A G C T A G C G A C T C T A G A G C T A G T C G T C A T G C A A C G T A T G C G C T A T G C A | Pbx3(Homeobox)/GM12878-PBX3-ChIP-Seq(GSE32465)/Homer | 1e-16 | -3.812e+01 | 0.0000 | 926.0 | 2.34% | 3097.0 | 1.70% | motif file (matrix) | svg |
| 336 | C T G A A T G C G C T A C G A T A T G C C G T A C G T A C G T A C T A G T A C G | Tcf3(HMG)/mES-Tcf3-ChIP-Seq(GSE11724)/Homer | 1e-16 | -3.812e+01 | 0.0000 | 1023.0 | 2.58% | 3480.3 | 1.91% | motif file (matrix) | svg |
| 337 | G C T A T C G A C G T A C T A G A G C T G T C A G T C A C G T A A G T C C G T A | FOXA1(Forkhead)/MCF7-FOXA1-ChIP-Seq(GSE26831)/Homer | 1e-16 | -3.743e+01 | 0.0000 | 2479.0 | 6.26% | 9492.8 | 5.20% | motif file (matrix) | svg |
| 338 | G T C A T C G A T C G A C G T A G C T A C G T A T C G A T G A C A C T G C G T A A G T C C G T A C G T A T C G A G C T A | IDD2(C2H2)/colamp-IDD2-DAP-Seq(GSE60143)/Homer | 1e-16 | -3.729e+01 | 0.0000 | 650.0 | 1.64% | 2040.4 | 1.12% | motif file (matrix) | svg |
| 339 | A C T G A C G T C A T G A T C G A T C G T G A C A C T G A T C G A T C G T G C A C T G A C G T A | E2F3(E2F)/MEF-E2F3-ChIP-Seq(GSE71376)/Homer | 1e-16 | -3.725e+01 | 0.0000 | 4289.0 | 10.83% | 17255.7 | 9.46% | motif file (matrix) | svg |
| 340 | G T A C G C T A C G A T C A G T A G T C G C T A C G A T C G A T A G T C G C T A | WUS1(Homeobox)/colamp-WUS1-DAP-Seq(GSE60143)/Homer | 1e-16 | -3.717e+01 | 0.0000 | 1735.0 | 4.38% | 6388.7 | 3.50% | motif file (matrix) | svg |
| 341 | C G A T C T G A A G T C A C G T A C G T T C A G C G T A C G T A C G T A G C A T C G A T A G T C C G T A G T C A A C T G | VND4(NAC)/colamp-VND4-DAP-Seq(GSE60143)/Homer | 1e-15 | -3.681e+01 | 0.0000 | 3827.0 | 9.67% | 15271.8 | 8.37% | motif file (matrix) | svg |
| 342 | G A T C C A T G A C G T A C G T A C T G C G T A A G T C A G C T C G A T G A C T | WRKY8(WRKY)/colamp-WRKY8-DAP-Seq(GSE60143)/Homer | 1e-15 | -3.673e+01 | 0.0000 | 560.0 | 1.41% | 1707.5 | 0.94% | motif file (matrix) | svg |
| 343 | C T G A C T G A C T A G T C G A C G T A A T G C C G T A A C T G C G T A A C G T C T G A C G A T A G C T C G T A A C G T A G T C C G A T T A C G G T C A G C A T | GATA(Zf),IR3/iTreg-Gata3-ChIP-Seq(GSE20898)/Homer | 1e-15 | -3.647e+01 | 0.0000 | 645.0 | 1.63% | 2031.8 | 1.11% | motif file (matrix) | svg |
| 344 | C T A G A G C T G A C T C A T G A G T C A G T C G T C A C A G T C T A G T C A G G T A C C T G A T C G A G A T C T G A C | Rfx2(HTH)/LoVo-RFX2-ChIP-Seq(GSE49402)/Homer | 1e-15 | -3.620e+01 | 0.0000 | 344.0 | 0.87% | 926.5 | 0.51% | motif file (matrix) | svg |
| 345 | A G T C G A T C A G T C C G T A A T C G C A G T A G T C G T A C C T G A A C T G T C A G A G C T A G C T A G C T A G C T | PRDM15(Zf)/ESC-Prdm15-ChIP-Seq(GSE73694)/Homer | 1e-15 | -3.568e+01 | 0.0000 | 4288.0 | 10.83% | 17306.3 | 9.48% | motif file (matrix) | svg |
| 346 | G T C A T C G A C T A G C T A G A G T C G T C A C G A T C T A G G A C T G A T C G A T C T C A G C T A G C T G A A G T C G C T A C A G T T C A G G A T C G A T C | p63(p53)/Keratinocyte-p63-ChIP-Seq(GSE17611)/Homer | 1e-15 | -3.535e+01 | 0.0000 | 1542.0 | 3.89% | 5631.1 | 3.09% | motif file (matrix) | svg |
| 347 | C G A T T C G A G T A C A C G T A C G T T C A G G C A T G C T A T G C A G C A T C G T A A G T C C G T A T G A C C A T G | ANAC092(NAC)/colamp-ANAC092-DAP-Seq(GSE60143)/Homer | 1e-15 | -3.515e+01 | 0.0000 | 2538.0 | 6.41% | 9803.9 | 5.37% | motif file (matrix) | svg |
| 348 | A C G T C T A G A G C T A C G T A C G T C T G A A G T C G A C T A G C T C G T A | FOXM1(Forkhead)/MCF7-FOXM1-ChIP-Seq(GSE72977)/Homer | 1e-15 | -3.513e+01 | 0.0000 | 2928.0 | 7.40% | 11463.4 | 6.28% | motif file (matrix) | svg |
| 349 | A T G C G A T C C G T A A G C T C A G T A T C G G C A T A G C T G A C T A C T G | Sox17(HMG)/Endoderm-Sox17-ChIP-Seq(GSE61475)/Homer | 1e-15 | -3.477e+01 | 0.0000 | 3302.0 | 8.34% | 13076.1 | 7.17% | motif file (matrix) | svg |
| 350 | T A C G T C G A G A C T A C T G C T G A A G T C T C A G G A C T T G A C C T G A | Atf1(bZIP)/K562-ATF1-ChIP-Seq(GSE31477)/Homer | 1e-14 | -3.448e+01 | 0.0000 | 4125.0 | 10.42% | 16641.0 | 9.12% | motif file (matrix) | svg |
| 351 | A C G T A G T C A G C T A G T C C G T A G T A C A G T C C G A T C G T A G T C A | MYB41(MYB)/col-MYB41-DAP-Seq(GSE60143)/Homer | 1e-14 | -3.397e+01 | 0.0000 | 2039.0 | 5.15% | 7729.8 | 4.24% | motif file (matrix) | svg |
| 352 | T C G A T G A C G T A C C G T A A C G T G A C T A C G T A C T G A C T G A G C T | Mesp1(bHLH)/ESC-Mesp1-ChIP-Seq(GSE165102)/Homer | 1e-14 | -3.319e+01 | 0.0000 | 2185.0 | 5.52% | 8363.3 | 4.58% | motif file (matrix) | svg |
| 353 | C G A T C T G A G T A C A C G T A C G T T C A G C G T A C G T A G C T A G C A T G C A T A G T C C G T A G T C A C A T G | NST1(NAC)/colamp-NST1-DAP-Seq(GSE60143)/Homer | 1e-14 | -3.287e+01 | 0.0000 | 3867.0 | 9.77% | 15579.5 | 8.54% | motif file (matrix) | svg |
| 354 | G A C T C T A G C T A G A G T C T G C A A C T G A C G T A C G T C T A G T C A G | AMYB(HTH)/Testes-AMYB-ChIP-Seq(GSE44588)/Homer | 1e-14 | -3.260e+01 | 0.0000 | 10936.0 | 27.62% | 46961.4 | 25.74% | motif file (matrix) | svg |
| 355 | C T G A A T G C C G T A A C G T A G T C A G T C A C G T A C T G A T C G G C A T | SPDEF(ETS)/VCaP-SPDEF-ChIP-Seq(SRA014231)/Homer | 1e-13 | -3.145e+01 | 0.0000 | 4201.0 | 10.61% | 17083.2 | 9.36% | motif file (matrix) | svg |
| 356 | C G T A A C T G C G T A A C G T C A G T A G T C A G C T G C A T G C T A C G A T | At2g01060(G2like)/colamp-At2g01060-DAP-Seq(GSE60143)/Homer | 1e-13 | -3.134e+01 | 0.0000 | 15416.0 | 38.94% | 67348.3 | 36.91% | motif file (matrix) | svg |
| 357 | T C G A G A C T A T C G C G T A A G T C C T A G G C A T G T A C C T G A A C G T G A T C G C T A | TGA4(bZIP)/colamp-TGA4-DAP-Seq(GSE60143)/Homer | 1e-13 | -3.100e+01 | 0.0000 | 1795.0 | 4.53% | 6780.7 | 3.72% | motif file (matrix) | svg |
| 358 | G A C T G T A C T G C A A C G T G A T C G C T A T C G A A C G T A G T C C G T A | Pdx1(Homeobox)/Islet-Pdx1-ChIP-Seq(SRA008281)/Homer | 1e-13 | -3.081e+01 | 0.0000 | 4105.0 | 10.37% | 16689.1 | 9.15% | motif file (matrix) | svg |
| 359 | G C A T G C T A C G T A A G C T G C T A T G C A A G T C A C G T A C G T A C G T G C A T G C A T | At4g38000(C2C2dof)/col-At4g38000-DAP-Seq(GSE60143)/Homer | 1e-13 | -3.076e+01 | 0.0000 | 3592.0 | 9.07% | 14462.2 | 7.93% | motif file (matrix) | svg |
| 360 | G A C T T C G A C G T A C G T A C G T A C G T A C G T A C T A G A G C T C G T A | dof45(C2C2dof)/col-dof45-DAP-Seq(GSE60143)/Homer | 1e-13 | -3.062e+01 | 0.0000 | 10038.0 | 25.35% | 43031.8 | 23.58% | motif file (matrix) | svg |
| 361 | A G T C C A G T T C A G A T G C A G T C C G A T C G T A G T C A G A T C G C A T | BOS1(MYB)/col-BOS1-DAP-Seq(GSE60143)/Homer | 1e-13 | -2.998e+01 | 0.0000 | 5786.0 | 14.61% | 24100.9 | 13.21% | motif file (matrix) | svg |
| 362 | A C T G C G T A A C T G A T G C T G A C G A T C A T C G T G C A A C T G A G T C | ZNF519(Zf)/HEK293-ZNF519.GFP-ChIP-Seq(GSE58341)/Homer | 1e-12 | -2.992e+01 | 0.0000 | 751.0 | 1.90% | 2532.6 | 1.39% | motif file (matrix) | svg |
| 363 | C G T A T G A C T C G A A G T C C G T A A T C G A T G C A C G T A C T G A G T C | E2A(bHLH)/proBcell-E2A-ChIP-Seq(GSE21978)/Homer | 1e-12 | -2.926e+01 | 0.0000 | 4234.0 | 10.69% | 17312.8 | 9.49% | motif file (matrix) | svg |
| 364 | T G A C G C T A T C G A T G C A A G T C A G T C C G T A A G T C C G T A C T G A G C T A G T A C | RUNX2(Runt)/PCa-RUNX2-ChIP-Seq(GSE33889)/Homer | 1e-12 | -2.918e+01 | 0.0000 | 3500.0 | 8.84% | 14120.4 | 7.74% | motif file (matrix) | svg |
| 365 | T A C G T A C G C T A G T C A G A G T C C G T A A T C G A T G C A C G T A C T G A G T C G A C T | Ascl2(bHLH)/ESC-Ascl2-ChIP-Seq(GSE97712)/Homer | 1e-12 | -2.904e+01 | 0.0000 | 3389.0 | 8.56% | 13644.4 | 7.48% | motif file (matrix) | svg |
| 366 | C G A T C T A G A C G T G T C A C G T A C G T A A G T C C G T A | Foxo3(Forkhead)/U2OS-Foxo3-ChIP-Seq(E-MTAB-2701)/Homer | 1e-12 | -2.815e+01 | 0.0000 | 2608.0 | 6.59% | 10308.9 | 5.65% | motif file (matrix) | svg |
| 367 | T C G A G C T A T G A C G C T A C T A G G A T C C G A T A C T G C G A T A G C T G A C T C T A G | E-box/Drosophila-Promoters/Homer | 1e-12 | -2.790e+01 | 0.0000 | 716.0 | 1.81% | 2425.4 | 1.33% | motif file (matrix) | svg |
| 368 | T C G A C T G A T A G C T G A C T C A G T C A G C G T A C G T A T C A G A G C T | ETV1(ETS)/GIST48-ETV1-ChIP-Seq(GSE22441)/Homer | 1e-12 | -2.772e+01 | 0.0000 | 6142.0 | 15.51% | 25781.2 | 14.13% | motif file (matrix) | svg |
| 369 | G T C A C G T A A C G T A T C G C G T A A C G T A C G T C T A G | ATHB7(Homeobox)/col-ATHB7-DAP-Seq(GSE60143)/Homer | 1e-12 | -2.764e+01 | 0.0000 | 3582.0 | 9.05% | 14533.0 | 7.97% | motif file (matrix) | svg |
| 370 | C G A T C T A G T C G A A G C T C G A T C T G A C G T A A G C T A C T G C T A G A T G C G A T C | Hoxb4(Homeobox)/ES-Hoxb4-ChIP-Seq(GSE34014)/Homer | 1e-11 | -2.747e+01 | 0.0000 | 1002.0 | 2.53% | 3580.4 | 1.96% | motif file (matrix) | svg |
| 371 | T A C G C T G A T C G A C G A T C T A G C T A G T C G A C T G A T C G A T C G A C G T A T C G A G C A T C A T G C G T A T A C G G C A T T G A C C G T A A G C T | NFAT:AP1(RHD,bZIP)/Jurkat-NFATC1-ChIP-Seq(Jolma\_et\_al.)/Homer | 1e-11 | -2.732e+01 | 0.0000 | 472.0 | 1.19% | 1485.5 | 0.81% | motif file (matrix) | svg |
| 372 | C G T A C G T A C G T A C G T A C G T A A C T G A C T G A G T C | dof42(C2C2dof)/col-dof42-DAP-Seq(GSE60143)/Homer | 1e-11 | -2.665e+01 | 0.0000 | 2804.0 | 7.08% | 11200.3 | 6.14% | motif file (matrix) | svg |
| 373 | C T G A T A C G G C A T C T A G A T G C G A T C C G A T A C T G C T A G G A T C C T G A A T G C | MYRF(MYRF)/CFPAC1-MYRF-ChIP-Seq(GSE145627)/Homer | 1e-11 | -2.553e+01 | 0.0000 | 1245.0 | 3.14% | 4618.1 | 2.53% | motif file (matrix) | svg |
| 374 | C A G T A G C T G A C T T G C A A G T C A G C T A C G T A C G T C G A T G A C T | AT3G52440(C2C2dof)/colamp-AT3G52440-DAP-Seq(GSE60143)/Homer | 1e-10 | -2.519e+01 | 0.0000 | 9275.0 | 23.43% | 39918.1 | 21.88% | motif file (matrix) | svg |
| 375 | C A G T G C T A G C A T T A C G C T G A C A G T T A G C C T G A | GATA15(C2C2gata)/col-GATA15-DAP-Seq(GSE60143)/Homer | 1e-10 | -2.508e+01 | 0.0000 | 9639.0 | 24.35% | 41563.4 | 22.78% | motif file (matrix) | svg |
| 376 | T G C A C T G A A T G C G T C A A C T G A C T G C G T A C G T A C T A G A G C T | Ets1-distal(ETS)/CD4+-PolII-ChIP-Seq(Barski\_et\_al.)/Homer | 1e-10 | -2.413e+01 | 0.0000 | 821.0 | 2.07% | 2909.0 | 1.59% | motif file (matrix) | svg |
| 377 | G C A T G C A T G C A T A T G C A G C T T C G A T A C G G C T A C G T A C A T G G T A C G C A T G C A T A G T C A G C T | HSFA6B(HSF)/colamp-HSFA6B-DAP-Seq(GSE60143)/Homer | 1e-10 | -2.411e+01 | 0.0000 | 1530.0 | 3.86% | 5843.2 | 3.20% | motif file (matrix) | svg |
| 378 | C T G A C T A G A C T G G C A T A T G C C G T A C T G A C T A G A C T G A C G T A G T C C T G A | RARg(NR)/ES-RARg-ChIP-Seq(GSE30538)/Homer | 1e-10 | -2.388e+01 | 0.0000 | 119.0 | 0.30% | 255.8 | 0.14% | motif file (matrix) | svg |
| 379 | C A G T A G C T C G T A G C A T A G T C G A C T C T A G C T A G C A G T C T A G T C G A T G C A C T A G C A T G G A C T | STOP1(C2H2)/colamp-STOP1-DAP-Seq(GSE60143)/Homer | 1e-10 | -2.365e+01 | 0.0000 | 1830.0 | 4.62% | 7124.7 | 3.90% | motif file (matrix) | svg |
| 380 | G A C T A G T C C G A T A C T G C T G A T G A C G T A C C G T A A T C G G C A T C T G A C T A G | Bcl11a(Zf)/HSPC-BCL11A-ChIP-Seq(GSE104676)/Homer | 1e-10 | -2.347e+01 | 0.0000 | 2221.0 | 5.61% | 8799.1 | 4.82% | motif file (matrix) | svg |
| 381 | T A C G C G T A T C A G G A C T C T A G A C T G C A G T T A G C T C G A A C G T G T A C C T A G A G T C A G T C G A T C | ZNF669(Zf)/HEK293-ZNF669.GFP-ChIP-Seq(GSE58341)/Homer | 1e-10 | -2.321e+01 | 0.0000 | 678.0 | 1.71% | 2350.2 | 1.29% | motif file (matrix) | svg |
| 382 | T A G C G T A C A G T C G T A C C G A T A G T C A G T C A G T C A G T C A G T C C G T A G A T C | Zfp281(Zf)/ES-Zfp281-ChIP-Seq(GSE81042)/Homer | 1e-10 | -2.311e+01 | 0.0000 | 399.0 | 1.01% | 1259.3 | 0.69% | motif file (matrix) | svg |
| 383 | G C A T A C G T A C T G A C G T A G T C A C T G A T C G G T C A C G A T C G T A | ARF2(ARF)/col-ARF2-DAP-Seq(GSE60143)/Homer | 1e-10 | -2.307e+01 | 0.0000 | 14901.0 | 37.64% | 65566.9 | 35.93% | motif file (matrix) | svg |
| 384 | C T G A A T C G A G C T A G C T A C G T T A G C C T G A T A C G C G A T A C G T G A C T A G T C | ISRE(IRF)/ThioMac-LPS-Expression(GSE23622)/Homer | 1e-9 | -2.287e+01 | 0.0000 | 176.0 | 0.44% | 448.9 | 0.25% | motif file (matrix) | svg |
| 385 | C G A T C G A T G T A C G A T C G A T C C G T A G C T A C G A T C G A T C T G A C T A G C A T G G C T A G C T A C G T A | AGL16(MADS)/col-AGL16-DAP-Seq(GSE60143)/Homer | 1e-9 | -2.215e+01 | 0.0000 | 283.0 | 0.71% | 835.4 | 0.46% | motif file (matrix) | svg |
| 386 | C A T G T G A C C A T G G A C T C A G T C T A G G C T A G T A C G A C T G C A T G C A T C G A T | WRKY21(WRKY)/colamp-WRKY21-DAP-Seq(GSE60143)/Homer | 1e-9 | -2.209e+01 | 0.0000 | 525.0 | 1.33% | 1761.7 | 0.97% | motif file (matrix) | svg |
| 387 | C G A T C A G T C T A G G C T A A G T C C G T A T C A G A G T C A C G T A C T G A C G T G T A C G C T A G C T A G C T A | bZIP52(bZIP)/colamp-bZIP52-DAP-Seq(GSE60143)/Homer | 1e-9 | -2.199e+01 | 0.0000 | 4432.0 | 11.19% | 18496.4 | 10.14% | motif file (matrix) | svg |
| 388 | A T G C G A C T A G C T C T A G C G T A C T A G C G A T C T A G A T C G G A T C | Nkx2.2(Homeobox)/NPC-Nkx2.2-ChIP-Seq(GSE61673)/Homer | 1e-9 | -2.195e+01 | 0.0000 | 9393.0 | 23.72% | 40655.2 | 22.28% | motif file (matrix) | svg |
| 389 | G A C T C T A G C T A G G T A C A G T C G A T C G A C T G A C T T A G C T C A G | NLP7(RWPRK)/col-NLP7-DAP-Seq(GSE60143)/Homer | 1e-9 | -2.184e+01 | 0.0000 | 8861.0 | 22.38% | 38266.6 | 20.97% | motif file (matrix) | svg |
| 390 | T G A C C G A T C T G A C T A G C T A G A C G T A T G C T G C A T C G A C T G A C T A G C A T G A C G T A G T C C G T A | PPARa(NR),DR1/Liver-Ppara-ChIP-Seq(GSE47954)/Homer | 1e-9 | -2.180e+01 | 0.0000 | 3285.0 | 8.30% | 13469.5 | 7.38% | motif file (matrix) | svg |
| 391 | G A C T C A G T A G C T C G A T A G T C G A T C A G T C C G T A A T G C T C A G | Rbpj1(?)/Panc1-Rbpj1-ChIP-Seq(GSE47459)/Homer | 1e-9 | -2.164e+01 | 0.0000 | 4522.0 | 11.42% | 18910.5 | 10.36% | motif file (matrix) | svg |
| 392 | A G T C G A C T C A G T A C T G C T A G T G A C G C T A A T G C G C A T A T C G C G A T A C T G G A T C G T A C G T C A C T G A | NF1(CTF)/LNCAP-NF1-ChIP-Seq(Unpublished)/Homer | 1e-9 | -2.141e+01 | 0.0000 | 1060.0 | 2.68% | 3945.5 | 2.16% | motif file (matrix) | svg |
| 393 | A C T G A G C T A G T C G T C A A G C T T C A G A T G C G A T C G C A T A T C G T C G A T A G C C G A T C A T G T A G C | Pax8(Paired,Homeobox)/Thyroid-Pax8-ChIP-Seq(GSE26938)/Homer | 1e-9 | -2.124e+01 | 0.0000 | 1087.0 | 2.75% | 4061.6 | 2.23% | motif file (matrix) | svg |
| 394 | C G T A C T A G T C A G T C A G A G T C A T G C A G T C G C A T A G C T A C G T A T C G C G A T | Sox9(HMG)/Limb-SOX9-ChIP-Seq(GSE73225)/Homer | 1e-9 | -2.088e+01 | 0.0000 | 2918.0 | 7.37% | 11906.5 | 6.53% | motif file (matrix) | svg |
| 395 | T C G A C T G A C G A T C G T A C G T A C G T A C T A G A G C T C T G A T C A G | Adof1(C2C2dof)/col-Adof1-DAP-Seq(GSE60143)/Homer | 1e-8 | -2.029e+01 | 0.0000 | 11022.0 | 27.84% | 48132.2 | 26.38% | motif file (matrix) | svg |
| 396 | T A G C G T A C C T A G A T C G C T G A C G T A G C T A G C A T A C G T T G A C G T A C A C T G T A C G G T C A C T A G | ASL18(LOBAS2)/colamp-ASL18-DAP-Seq(GSE60143)/Homer | 1e-8 | -2.025e+01 | 0.0000 | 8599.0 | 21.72% | 37191.9 | 20.38% | motif file (matrix) | svg |
| 397 | T C A G G C A T A C T G C G T A A G T C C T A G G C A T T G A C | TGA9(bZIP)/colamp-TGA9-DAP-Seq(GSE60143)/Homer | 1e-8 | -2.023e+01 | 0.0000 | 7914.0 | 19.99% | 34112.4 | 18.70% | motif file (matrix) | svg |
| 398 | A T C G A G T C A G T C C G T A A C T G G C A T | hINR(CPE) | 1e-8 | -2.005e+01 | 0.0000 | 4734.0 | 11.96% | 19927.2 | 10.92% | motif file (matrix) | svg |
| 399 | T C G A C G T A C G T A T C G A A C T G G T A C C G T A A G C T G T C A G C A T | At3g24120(G2like)/col-At3g24120-DAP-Seq(GSE60143)/Homer | 1e-8 | -1.978e+01 | 0.0000 | 15766.0 | 39.82% | 69774.0 | 38.24% | motif file (matrix) | svg |
| 400 | T C G A A C T G A C T G C G T A C G T A T C G A A G T C C T G A A T C G G T A C G C A T C A T G | ETS:E-box(ETS,bHLH)/HPC7-Scl-ChIP-Seq(GSE22178)/Homer | 1e-8 | -1.973e+01 | 0.0000 | 233.0 | 0.59% | 676.8 | 0.37% | motif file (matrix) | svg |
| 401 | G C A T C G T A G C A T C G T A T C G A C G T A C T G A A C T G C G T A C G T A C G T A A C G T A C T G G T C A G C A T | AT2G31460(REMB3)/col-AT2G31460-DAP-Seq(GSE60143)/Homer | 1e-8 | -1.949e+01 | 0.0000 | 1092.0 | 2.76% | 4125.0 | 2.26% | motif file (matrix) | svg |
| 402 | G A C T A T C G C T G A A G T C T C A G G A C T G T A C C T G A A G C T G T A C | TGA6(bZIP)/colamp-TGA6-DAP-Seq(GSE60143)/Homer | 1e-8 | -1.939e+01 | 0.0000 | 4502.0 | 11.37% | 18934.4 | 10.38% | motif file (matrix) | svg |
| 403 | A T G C A G T C G T A C A G C T T C G A C T A G G A T C C T G A G T C A A G T C G C T A T C A G | Rfx5(HTH)/GM12878-Rfx5-ChIP-Seq(GSE31477)/Homer | 1e-8 | -1.930e+01 | 0.0000 | 1306.0 | 3.30% | 5028.8 | 2.76% | motif file (matrix) | svg |
| 404 | T C G A T C G A T A G C G T A C T C A G T A C G C G T A C G T A T C A G A G C T | GABPA(ETS)/Jurkat-GABPa-ChIP-Seq(GSE17954)/Homer | 1e-8 | -1.929e+01 | 0.0000 | 4191.0 | 10.59% | 17564.4 | 9.63% | motif file (matrix) | svg |
| 405 | T A G C C T A G T C G A G A C T A C T G C T G A A G T C T C A G G C A T T G A C C T G A A G C T | Atf7(bZIP)/3T3L1-Atf7-ChIP-Seq(GSE56872)/Homer | 1e-8 | -1.902e+01 | 0.0000 | 2202.0 | 5.56% | 8870.5 | 4.86% | motif file (matrix) | svg |
| 406 | C A G T A T C G C T G A A G T C T C A G C A G T T A G C C T G A A T G C T A C G | FEA4(bZIP)/Corn-FEA4-ChIP-Seq(GSE61954)/Homer | 1e-8 | -1.899e+01 | 0.0000 | 6572.0 | 16.60% | 28173.8 | 15.44% | motif file (matrix) | svg |
| 407 | C T G A A G C T A C G T A C G T A G T C G A C T G A C T C T G A C T G A C T A G C G T A C G T A | STAT6(Stat)/CD4-Stat6-ChIP-Seq(GSE22104)/Homer | 1e-8 | -1.860e+01 | 0.0000 | 1406.0 | 3.55% | 5471.6 | 3.00% | motif file (matrix) | svg |
| 408 | C G A T C T A G C T G A A T G C C T G A T C G A C G T A C T G A T C G A T A G C A G T C C G T A A C T G T C G A A T G C | Hand2(bHLH)/Mesoderm-Hand2-ChIP-Seq(GSE61475)/Homer | 1e-8 | -1.849e+01 | 0.0000 | 1265.0 | 3.20% | 4877.1 | 2.67% | motif file (matrix) | svg |
| 409 | C T A G A T G C A T G C C G A T A C T G G A C T A T G C G C T A T G A C A G C T T A G C G C T A | PBX1(Homeobox)/MCF7-PBX1-ChIP-Seq(GSE28007)/Homer | 1e-7 | -1.798e+01 | 0.0000 | 232.0 | 0.59% | 690.5 | 0.38% | motif file (matrix) | svg |
| 410 | G T A C A C T G A C G T T C A G G C A T C G T A C G A T G C A T C G T A A G T C C G T A T G A C C A T G G A C T G C T A | ANAC083(NAC)/col-ANAC083-DAP-Seq(GSE60143)/Homer | 1e-7 | -1.793e+01 | 0.0000 | 4046.0 | 10.22% | 16992.4 | 9.31% | motif file (matrix) | svg |
| 411 | C G A T C T G A G T A C A C T G A C G T T C A G G C A T C G T A C G T A G C A T C G T A A G T C C G T A G T A C C A T G | CUC3(NAC)/col-CUC3-DAP-Seq(GSE60143)/Homer | 1e-7 | -1.789e+01 | 0.0000 | 2008.0 | 5.07% | 8073.5 | 4.42% | motif file (matrix) | svg |
| 412 | C G T A T C G A G A T C G C A T C G T A A C G T G T A C T C A G G T C A G A C T C G T A C T A G | DREF/Drosophila-Promoters/Homer | 1e-7 | -1.767e+01 | 0.0000 | 344.0 | 0.87% | 1118.8 | 0.61% | motif file (matrix) | svg |
| 413 | A G C T A G T C A G T C A C G T C T A G A C G T A C G T A C G T C G T A A G T C G A T C C G T A | FOXP1(Forkhead)/H9-FOXP1-ChIP-Seq(GSE31006)/Homer | 1e-7 | -1.734e+01 | 0.0000 | 1439.0 | 3.63% | 5649.8 | 3.10% | motif file (matrix) | svg |
| 414 | A C T G G A T C G A C T A C T G A C G T C A T G A C T G A C G T A G C T C G A T | RUNX-AML(Runt)/CD4+-PolII-ChIP-Seq(Barski\_et\_al.)/Homer | 1e-7 | -1.718e+01 | 0.0000 | 2634.0 | 6.65% | 10820.3 | 5.93% | motif file (matrix) | svg |
| 415 | C A T G G T C A A G T C C G T A C T A G G A T C C G A T A C T G A C G T G T A C C G T A C G T A | bZIP69(bZIP)/col-bZIP69-DAP-Seq(GSE60143)/Homer | 1e-7 | -1.694e+01 | 0.0000 | 352.0 | 0.89% | 1159.3 | 0.64% | motif file (matrix) | svg |
| 416 | C A T G G A T C C T G A A G T C C T A G C G T A G C T A G C A T G A T C G A T C A G T C C T A G C G T A C A T G C T A G | PLT1(AP2EREBP)/colamp-PLT1-DAP-Seq(GSE60143)/Homer | 1e-7 | -1.683e+01 | 0.0000 | 620.0 | 1.57% | 2232.4 | 1.22% | motif file (matrix) | svg |
| 417 | A T G C C G T A C G T A C G T A C G T A C G T A A C T G A C G T C G A T C T G A | dof43(C2C2dof)/colamp-dof43-DAP-Seq(GSE60143)/Homer | 1e-7 | -1.681e+01 | 0.0000 | 5810.0 | 14.67% | 24901.2 | 13.65% | motif file (matrix) | svg |
| 418 | A G T C C T G A A T C G A G C T A G C T G A C T A G T C G C T A A C G T C G A T G C A T C G A T A T C G C G T A T A G C G C A T A T G C C G T A | bZIP:IRF(bZIP,IRF)/Th17-BatF-ChIP-Seq(GSE39756)/Homer | 1e-7 | -1.653e+01 | 0.0000 | 939.0 | 2.37% | 3558.3 | 1.95% | motif file (matrix) | svg |
| 419 | C T G A T C A G C T G A C T A G C A T G A C G T A T G C C G T A A T G C G C A T T C A G C T G A A C T G A C G T C A G T A G T C C G T A C A G T C T A G C A T G | VDR(NR),DR3/GM10855-VDR+vitD-ChIP-Seq(GSE22484)/Homer | 1e-7 | -1.649e+01 | 0.0000 | 788.0 | 1.99% | 2930.8 | 1.61% | motif file (matrix) | svg |
| 420 | T G C A C G T A A C T G T C A G C A G T C A T G T C A G G A T C T A C G A G T C T G C A A C T G A C T G T G A C G T C A | ZNF165(Zf)/WHIM12-ZNF165-ChIP-Seq(GSE65937)/Homer | 1e-7 | -1.635e+01 | 0.0000 | 384.0 | 0.97% | 1294.0 | 0.71% | motif file (matrix) | svg |
| 421 | C G A T C T G A G T A C C A T G G C A T T C A G G C A T C G T A C G T A G C T A C G T A A G T C G C T A G T A C C A T G | CUC2(NAC)/colamp-CUC2-DAP-Seq(GSE60143)/Homer | 1e-6 | -1.596e+01 | 0.0000 | 2008.0 | 5.07% | 8145.5 | 4.46% | motif file (matrix) | svg |
| 422 | A G C T C T G A C T A G C T A G A C T G T A G C T G C A T C G A C T G A C T A G C A T G A C G T A T G C T C G A | RXR(NR),DR1/3T3L1-RXR-ChIP-Seq(GSE13511)/Homer | 1e-6 | -1.579e+01 | 0.0000 | 3308.0 | 8.36% | 13843.0 | 7.59% | motif file (matrix) | svg |
| 423 | C G A T C A G T C A G T C A T G G T C A G A T C C G T A T C A G A G T C A C G T C T A G A C G T G T A C G T C A G C T A | VIP1(bZIP)/col-VIP1-DAP-Seq(GSE60143)/Homer | 1e-6 | -1.526e+01 | 0.0000 | 541.0 | 1.37% | 1941.1 | 1.06% | motif file (matrix) | svg |
| 424 | C T G A C G A T C T A G C G T A A G C T C G A T C A G T C T G A G A C T C T A G C T A G A T G C | PBX2(Homeobox)/K562-PBX2-ChIP-Seq(Encode)/Homer | 1e-6 | -1.524e+01 | 0.0000 | 3818.0 | 9.64% | 16125.3 | 8.84% | motif file (matrix) | svg |
| 425 | A C T G G A C T A G T C C T G A G A T C T C A G A T G C G A C T A G T C A T G C T A G C A G C T A T C G T G C A | PAX5(Paired,Homeobox),condensed/GM12878-PAX5-ChIP-Seq(GSE32465)/Homer | 1e-6 | -1.473e+01 | 0.0000 | 571.0 | 1.44% | 2074.0 | 1.14% | motif file (matrix) | svg |
| 426 | T C G A G C A T A C T G C T G A A G T C T C A G G A C T G T A C C G T A A G C T A G T C G A T C | c-Jun-CRE(bZIP)/K562-cJun-ChIP-Seq(GSE31477)/Homer | 1e-6 | -1.447e+01 | 0.0000 | 980.0 | 2.48% | 3785.2 | 2.07% | motif file (matrix) | svg |
| 427 | G A C T C A G T G A T C G A T C A C G T G A T C C T G A T A C G C G T A G T C A | STAT6(Stat)/Macrophage-Stat6-ChIP-Seq(GSE38377)/Homer | 1e-6 | -1.401e+01 | 0.0000 | 1442.0 | 3.64% | 5771.9 | 3.16% | motif file (matrix) | svg |
| 428 | G C A T A C G T C G A T A G T C A G T C G C A T C G T A C G T A C G A T C G A T C G A T C T A G A C T G G C T A G C T A | AGL15(MADS)/col-AGL15-DAP-Seq(GSE60143)/Homer | 1e-6 | -1.398e+01 | 0.0000 | 365.0 | 0.92% | 1255.8 | 0.69% | motif file (matrix) | svg |
| 429 | A T C G T G A C A T G C C T G A T C A G G A C T A G T C C G A T T C A G T C G A C A T G C T A G C T A G C G T A C T A G C T A G C T G A C T A G C T A G A T G C | ZSCAN22(Zf)/HEK293-ZSCAN22.GFP-ChIP-Seq(GSE58341)/Homer | 1e-6 | -1.391e+01 | 0.0000 | 188.0 | 0.47% | 570.2 | 0.31% | motif file (matrix) | svg |
| 430 | C G T A A T G C C G A T A C G T A G T C C G T A C G T A C G T A C T A G A T C G | TCFL2(HMG)/K562-TCF7L2-ChIP-Seq(GSE29196)/Homer | 1e-5 | -1.381e+01 | 0.0000 | 310.0 | 0.78% | 1041.0 | 0.57% | motif file (matrix) | svg |
| 431 | G C A T G C A T G A T C G C A T T C G A A C T G C G T A C G T A A T C G T A G C C G A T A C G T A G T C A G C T C G T A | HSF6(HSF)/col-HSF6-DAP-Seq(GSE60143)/Homer | 1e-5 | -1.296e+01 | 0.0000 | 604.0 | 1.53% | 2248.2 | 1.23% | motif file (matrix) | svg |
| 432 | A G T C C T G A A G T C C G A T C A G T G A T C A T G C A C T G A T C G G A C T | Fli1(ETS)/CD8-FLI-ChIP-Seq(GSE20898)/Homer | 1e-5 | -1.280e+01 | 0.0000 | 6663.0 | 16.83% | 29008.2 | 15.90% | motif file (matrix) | svg |
| 433 | T C G A C T G A C G T A C G T A C G T A C G T A A C T G A C G T C G A T C T G A | BBX31(Orphan)/col-BBX31-DAP-Seq(GSE60143)/Homer | 1e-5 | -1.237e+01 | 0.0000 | 6227.0 | 15.73% | 27078.6 | 14.84% | motif file (matrix) | svg |
| 434 | T C G A T A G C G T C A A C T G C T A G C G T A C G A T A C T G A C G T A C T G A C T G A C G T | ETS:RUNX(ETS,Runt)/Jurkat-RUNX1-ChIP-Seq(GSE17954)/Homer | 1e-5 | -1.219e+01 | 0.0000 | 339.0 | 0.86% | 1181.5 | 0.65% | motif file (matrix) | svg |
| 435 | C T A G T C G A C G A T C T A G G C A T C A G T C T A G G A T C C G T A G T C A | CEBP:AP1(bZIP)/ThioMac-CEBPb-ChIP-Seq(GSE21512)/Homer | 1e-5 | -1.201e+01 | 0.0000 | 3149.0 | 7.95% | 13340.0 | 7.31% | motif file (matrix) | svg |
| 436 | C T A G C T A G T C A G G T C A C T A G T C A G G C T A A G T C A T C G A G C T C T A G | DPR(core promoter) | 1e-5 | -1.190e+01 | 0.0000 | 30494.0 | 77.02% | 138660.1 | 75.99% | motif file (matrix) | svg |
| 437 | T A C G T A G C C A T G C A G T A C G T C T A G C G T A A G T C G A C T G C A T G C A T C A G T | WRKY11(WRKY)/col-WRKY11-DAP-Seq(GSE60143)/Homer | 1e-5 | -1.182e+01 | 0.0000 | 686.0 | 1.73% | 2618.3 | 1.43% | motif file (matrix) | svg |
| 438 | C G T A C G T A C T G A A C T G A C G T A G T C C G T A C G T A A G T C A C T G A T G C G A T C | WRKY46(WRKY)/colamp-WRKY46-DAP-Seq(GSE60143)/Homer | 1e-4 | -1.142e+01 | 0.0000 | 553.0 | 1.40% | 2072.5 | 1.14% | motif file (matrix) | svg |
| 439 | A T G C C T G A A T C G T A C G A G T C C G A T T C A G C G A T C T A G A G C T G T C A G T C A C G T A A G T C C G T A T A C G C T G A | Fox:Ebox(Forkhead,bHLH)/Panc1-Foxa2-ChIP-Seq(GSE47459)/Homer | 1e-4 | -1.127e+01 | 0.0000 | 2640.0 | 6.67% | 11128.9 | 6.10% | motif file (matrix) | svg |
| 440 | C G A T C G A T C G A T A G T C A G T C G C T A C G T A C G T A C G T A G C T A C G A T C T A G A C T G G C T A C G T A | AGL6(MADS)/col-AGL6-DAP-Seq(GSE60143)/Homer | 1e-4 | -1.107e+01 | 0.0000 | 317.0 | 0.80% | 1112.4 | 0.61% | motif file (matrix) | svg |
| 441 | T G C A A G C T A C G T C T A G G A T C C T A G G A T C G T C A C T G A A G T C | CEBP(bZIP)/ThioMac-CEBPb-ChIP-Seq(GSE21512)/Homer | 1e-4 | -1.093e+01 | 0.0000 | 3492.0 | 8.82% | 14929.3 | 8.18% | motif file (matrix) | svg |
| 442 | C G A T G A C T C G A T T C A G G A C T A C G T C A G T C T G A G A C T G A C T A G C T C G A T A C T G A T C G G T A C G C T A | NF1:FOXA1(CTF,Forkhead)/LNCAP-FOXA1-ChIP-Seq(GSE27824)/Homer | 1e-4 | -1.083e+01 | 0.0000 | 153.0 | 0.39% | 473.9 | 0.26% | motif file (matrix) | svg |
| 443 | T C G A G C A T A C G T C T A G G T A C T C G A G C A T T G A C T C G A A C G T | Chop(bZIP)/MEF-Chop-ChIP-Seq(GSE35681)/Homer | 1e-4 | -1.081e+01 | 0.0000 | 990.0 | 2.50% | 3939.7 | 2.16% | motif file (matrix) | svg |
| 444 | T C A G A C T G C A G T A G T C A G T C G T C A C G T A C G T A A C T G C A G T A G T C A G T C C T G A T G C A A G C T | dHNF4(NR)/Fly-HNF4-ChIP-Seq(GSE73675)/Homer | 1e-4 | -1.060e+01 | 0.0001 | 192.0 | 0.48% | 625.9 | 0.34% | motif file (matrix) | svg |
| 445 | T G A C T C G A C T G A C T G A A T G C G A T C C T A G T A C G G A C T G A C T G A T C T C G A C T G A C T G A A T G C G A T C C T A G A T C G G A C T G A C T | Tcfcp2l1(CP2)/mES-Tcfcp2l1-ChIP-Seq(GSE11431)/Homer | 1e-4 | -1.052e+01 | 0.0001 | 617.0 | 1.56% | 2362.8 | 1.29% | motif file (matrix) | svg |
| 446 | G A C T C T A G G A T C C A G T A C T G C T G A A T G C G C A T A T G C C T G A | MafA(bZIP)/Islet-MafA-ChIP-Seq(GSE30298)/Homer | 1e-4 | -1.043e+01 | 0.0001 | 2893.0 | 7.31% | 12297.0 | 6.74% | motif file (matrix) | svg |
| 447 | A G T C G A C T C A G T G T A C A G T C A T C G T C A G A C T G G T C A C G T A | Stat3(Stat)/mES-Stat3-ChIP-Seq(GSE11431)/Homer | 1e-4 | -1.032e+01 | 0.0001 | 1650.0 | 4.17% | 6820.4 | 3.74% | motif file (matrix) | svg |
| 448 | C G T A C G T A C G T A A C G T C G T A A C G T A G T C G C A T | EPR1(MYBrelated)/colamp-EPR1-DAP-Seq(GSE60143)/Homer | 1e-4 | -1.031e+01 | 0.0001 | 1264.0 | 3.19% | 5139.4 | 2.82% | motif file (matrix) | svg |
| 449 | C G T A C G T A C G T A A C G T C G T A A C G T A G T C G C A T | LHY1(MYBrelated)/col-LHY1-DAP-Seq(GSE60143)/Homer | 1e-4 | -1.031e+01 | 0.0001 | 1264.0 | 3.19% | 5139.4 | 2.82% | motif file (matrix) | svg |
| 450 | A T G C G A C T A C T G C A G T G A T C A C G T T A C G T A C G | Smad2(MAD)/ES-SMAD2-ChIP-Seq(GSE29422)/Homer | 1e-4 | -1.022e+01 | 0.0001 | 7790.0 | 19.68% | 34319.1 | 18.81% | motif file (matrix) | svg |
| 451 | G C A T C G A T A T G C A G C T T C G A T A C G G C T A C G T A C A T G T G A C G C A T C G A T A G T C A G C T C G T A | AT3G09735(S1Falike)/col-AT3G09735-DAP-Seq(GSE60143)/Homer | 1e-4 | -1.005e+01 | 0.0001 | 1134.0 | 2.86% | 4586.6 | 2.51% | motif file (matrix) | svg |
| 452 | G C A T G A T C T C A G G C T A G A C T A G T C C T A G C G T A C A T G G T C A | GATA20(C2C2gata)/colamp-GATA20-DAP-Seq(GSE60143)/Homer | 1e-4 | -9.974e+00 | 0.0001 | 15922.0 | 40.22% | 71441.3 | 39.15% | motif file (matrix) | svg |
| 453 | T A C G T A C G G T A C A T C G A C T G T A C G T C G A C T G A T C G A A T C G | E2F6(E2F)/Hela-E2F6-ChIP-Seq(GSE31477)/Homer | 1e-4 | -9.924e+00 | 0.0001 | 2792.0 | 7.05% | 11878.4 | 6.51% | motif file (matrix) | svg |
| 454 | G A T C G C A T G C A T A G T C A G C T T C G A T A C G G C T A C G T A C T A G T G A C G C A T C G A T G A T C A G C T | HSFC1(HSF)/col-HSFC1-DAP-Seq(GSE60143)/Homer | 1e-4 | -9.879e+00 | 0.0001 | 450.0 | 1.14% | 1680.1 | 0.92% | motif file (matrix) | svg |
| 455 | T G A C A T G C C G T A A T C G A T G C C A G T C A T G A C T G A G T C G T A C | HEB(bHLH)/mES-Heb-ChIP-Seq(GSE53233)/Homer | 1e-4 | -9.799e+00 | 0.0001 | 5956.0 | 15.04% | 26066.4 | 14.29% | motif file (matrix) | svg |
| 456 | C G T A C G A T C G T A T C G A T C G A A C G T C G T A A C G T A G T C G C A T | LHY(Myb)/Seedling-LHY-ChIP-Seq(GSE52175)/Homer | 1e-4 | -9.565e+00 | 0.0002 | 4188.0 | 10.58% | 18133.8 | 9.94% | motif file (matrix) | svg |
| 457 | T C G A A C G T A C T G C T G A A G T C T C A G A G C T G T A C C G T A A G C T G A T C T C G A | JunD(bZIP)/K562-JunD-ChIP-Seq/Homer | 1e-4 | -9.359e+00 | 0.0002 | 274.0 | 0.69% | 969.8 | 0.53% | motif file (matrix) | svg |
| 458 | A T G C G C A T C G A T G A T C A G C T C T G A A C T G C G T A C G T A T C A G T G A C C G A T G C A T G A T C C G A T | HSF21(HSF)/col-HSF21-DAP-Seq(GSE60143)/Homer | 1e-4 | -9.212e+00 | 0.0002 | 222.0 | 0.56% | 763.1 | 0.42% | motif file (matrix) | svg |
| 459 | G C T A G C A T G A C T G C A T T C A G G T A C G C T A G C A T C T G A G C T A T A G C G C T A C T G A C G A T C T A G | OCT4-SOX2-TCF-NANOG(POU,Homeobox,HMG)/mES-Oct4-ChIP-Seq(GSE11431)/Homer | 1e-3 | -9.041e+00 | 0.0003 | 295.0 | 0.75% | 1060.2 | 0.58% | motif file (matrix) | svg |
| 460 | A C T G A G T C G T C A C G T A A G T C C G T A C T A G C T A G G A C T C A T G | SCRT1(Zf)/HEK293-SCRT1.eGFP-ChIP-Seq(Encode)/Homer | 1e-3 | -9.038e+00 | 0.0003 | 1399.0 | 3.53% | 5780.1 | 3.17% | motif file (matrix) | svg |
| 461 | C A G T G A C T G C A T T C G A A G T C A C G T A C G T A C G T C G A T G A C T | OBP3(C2C2dof)/col-OBP3-DAP-Seq(GSE60143)/Homer | 1e-3 | -8.971e+00 | 0.0003 | 11677.0 | 29.49% | 52133.8 | 28.57% | motif file (matrix) | svg |
| 462 | C T G A G T A C G A C T A G T C C A G T T G C A C T G A A C G T A G C T G A T C C T A G C G A T A C T G A T G C G A C T C T G A G A T C G A C T A G C T G A T C | Mouse\_Recombination\_Hotspot(Zf)/Testis-DMC1-ChIP-Seq(GSE24438)/Homer | 1e-3 | -8.904e+00 | 0.0003 | 220.0 | 0.56% | 760.6 | 0.42% | motif file (matrix) | svg |
| 463 | A G C T G C A T G T C A C G A T T A G C C G T A A C G T G C T A | CRC(C2C2YABBY)/col-CRC-DAP-Seq(GSE60143)/Homer | 1e-3 | -8.641e+00 | 0.0004 | 5921.0 | 14.96% | 26011.2 | 14.26% | motif file (matrix) | svg |
| 464 | T C G A A G T C C G T A A T C G T A G C A C G T A C T G A G C T A C G T A G T C | Ptf1a(bHLH)/Panc1-Ptf1a-ChIP-Seq(GSE47459)/Homer | 1e-3 | -8.531e+00 | 0.0004 | 8483.0 | 21.43% | 37633.7 | 20.63% | motif file (matrix) | svg |
| 465 | G C A T C A G T C G A T A G T C G A T C G C T A C G A T C G A T C G A T G C T A C G A T C T A G A C T G G C T A G C T A | AGL25(MADS)/colamp-AGL25-DAP-Seq(GSE60143)/Homer | 1e-3 | -8.518e+00 | 0.0005 | 103.0 | 0.26% | 311.3 | 0.17% | motif file (matrix) | svg |
| 466 | T A G C G C T A T C G A C T G A A G T C A G T C C T G A A G T C C G T A C T A G | RUNX(Runt)/HPC7-Runx1-ChIP-Seq(GSE22178)/Homer | 1e-3 | -8.510e+00 | 0.0005 | 3257.0 | 8.23% | 14042.8 | 7.70% | motif file (matrix) | svg |
| 467 | G C T A A G C T G T A C G C A T A G C T T C G A C T G A A G T C A G T C T A C G A C G T G A C T T A C G C T A G C G T A | ZML1(C2C2gata)/colamp-ZML1-DAP-Seq(GSE60143)/Homer | 1e-3 | -8.490e+00 | 0.0005 | 230.0 | 0.58% | 807.3 | 0.44% | motif file (matrix) | svg |
| 468 | A T G C C T G A G A C T A C G T A C G T G T A C G A T C C G A T C T A G C A T G C G T A C G T A C T G A G A C T | STAT1(Stat)/HelaS3-STAT1-ChIP-Seq(GSE12782)/Homer | 1e-3 | -8.434e+00 | 0.0005 | 641.0 | 1.62% | 2523.6 | 1.38% | motif file (matrix) | svg |
| 469 | T G A C G T A C C G T A A C T G T G A C C G A T A C T G A T C G A G C T T A C G T C G A T A G C G T A C C G T A A T C G T G A C G C A T A C T G A C T G A T G C | Twist(bHLH)/HMLE-TWIST1-ChIP-Seq(Chang\_et\_al)/Homer | 1e-3 | -8.348e+00 | 0.0005 | 263.0 | 0.66% | 943.6 | 0.52% | motif file (matrix) | svg |
| 470 | G C T A A C T G T C G A C T G A C T G A A C G T T A G C C T G A C G T A C G A T | Cux2(Homeobox)/Liver-Cux2-ChIP-Seq(GSE35985)/Homer | 1e-3 | -7.974e+00 | 0.0008 | 3620.0 | 9.14% | 15707.9 | 8.61% | motif file (matrix) | svg |
| 471 | C T G A A C G T A C G T A C G T A G T C G A C T C G A T C T G A A C T G C G T A C G T A T C G A | STAT5(Stat)/mCD4+-Stat5-ChIP-Seq(GSE12346)/Homer | 1e-3 | -7.765e+00 | 0.0009 | 672.0 | 1.70% | 2677.9 | 1.47% | motif file (matrix) | svg |
| 472 | C G T A C G T A C T G A A C T G C G T A C G T A A C G T G T C A A C G T G C A T A G T C G A C T | At2g03500(G2like)/col-At2g03500-DAP-Seq(GSE60143)/Homer | 1e-3 | -7.659e+00 | 0.0010 | 1014.0 | 2.56% | 4160.1 | 2.28% | motif file (matrix) | svg |
| 473 | T A C G A T C G T A G C G A T C A C T G A C G T A G T C A C G T C T A G A T C G | Smad4(MAD)/ESC-SMAD4-ChIP-Seq(GSE29422)/Homer | 1e-3 | -7.629e+00 | 0.0011 | 7890.0 | 19.93% | 35037.8 | 19.20% | motif file (matrix) | svg |
| 474 | G C A T T C G A C T G A G A T C A G T C G A T C G T C A G T C A A C G T A G T C C G T A C T G A | Duxbl(Homeobox)/NIH3T3-Duxbl.HA-ChIP-Seq(GSE119782)/Homer | 1e-3 | -7.549e+00 | 0.0012 | 322.0 | 0.81% | 1202.5 | 0.66% | motif file (matrix) | svg |
| 475 | C T A G T A C G G A C T T G C A T G C A C G A T T A C G C T G A T C G A C T G A | Hoxa10(Homeobox)/ChickenMSG-Hoxa10.Flag-ChIP-Seq(GSE86088)/Homer | 1e-3 | -7.544e+00 | 0.0012 | 2085.0 | 5.27% | 8883.1 | 4.87% | motif file (matrix) | svg |
| 476 | T G C A G C A T C G A T C G T A C A G T A C T G G T A C C G T A C T G A A G C T G T C A A C T G C T A G G T C A C G A T A C T G G T A C T G C A C G T A A G C T | CEBP:CEBP(bZIP)/MEF-Chop-ChIP-Seq(GSE35681)/Homer | 1e-3 | -7.421e+00 | 0.0013 | 415.0 | 1.05% | 1594.4 | 0.87% | motif file (matrix) | svg |
| 477 | A T G C C G T A A C T G C G T A A C G T G C T A T C G A A G C T C G A T C G T A A C G T A G T C C G A T A C T G G A T C | GATA(Zf),IR4/iTreg-Gata3-ChIP-Seq(GSE20898)/Homer | 1e-3 | -7.357e+00 | 0.0014 | 326.0 | 0.82% | 1223.9 | 0.67% | motif file (matrix) | svg |
| 478 | C G T A C T G A C T A G C G T A C G T A A G T C C G T A C A G T G C A T G T C A C G A T A C T G A C G T G C A T G A T C | PGR(NR)/EndoStromal-PGR-ChIP-Seq(GSE69539)/Homer | 1e-3 | -7.350e+00 | 0.0014 | 596.0 | 1.51% | 2365.8 | 1.30% | motif file (matrix) | svg |
| 479 | A T G C A G C T T C A G T G A C T C A G A T G C T G C A A C G T A T C G G A T C A C T G A G T C | NRF1(NRF)/MCF7-NRF1-ChIP-Seq(Unpublished)/Homer | 1e-3 | -7.241e+00 | 0.0016 | 367.0 | 0.93% | 1397.9 | 0.77% | motif file (matrix) | svg |
| 480 | C G A T C G A T A G T C A G T C G C T A C G A T C G T A C G A T G C A T C G A T C T A G A C T G G C A T G C T A C G T A | AGL13(MADS)/col-AGL13-DAP-Seq(GSE60143)/Homer | 1e-3 | -7.112e+00 | 0.0018 | 102.0 | 0.26% | 323.3 | 0.18% | motif file (matrix) | svg |
| 481 | C G T A C G T A C G T A A C G T C G T A A C G T A G T C G C A T | RVE1(MYBrelated)/col-RVE1-DAP-Seq(GSE60143)/Homer | 1e-3 | -7.105e+00 | 0.0018 | 2240.0 | 5.66% | 9600.2 | 5.26% | motif file (matrix) | svg |
| 482 | C T A G T G A C G A C T A T C G T C G A A G T C C T A G C A G T C T A G A T C G G T A C T C G A | O2(bZIP)/Corn-O2-ChIP-Seq(GSE63991)/Homer | 1e-2 | -6.836e+00 | 0.0023 | 820.0 | 2.07% | 3350.4 | 1.84% | motif file (matrix) | svg |
| 483 | T G A C C T G A C T A G C T G A C G T A A G T C C T G A A C G T G C A T T A G C G C A T A T C G G A C T G A C T G A T C | GRE(NR),IR3/RAW264.7-GRE-ChIP-Seq(Unpublished)/Homer | 1e-2 | -6.621e+00 | 0.0029 | 748.0 | 1.89% | 3046.7 | 1.67% | motif file (matrix) | svg |
| 484 | G C T A C G T A C G T A A G T C G A C T C G T A A C G T C T G A A C G T C G T A C G A T A C G T C G T A C G A T C T G A | AT1G04880(ARID)/colamp-AT1G04880-DAP-Seq(GSE60143)/Homer | 1e-2 | -6.497e+00 | 0.0033 | 250.0 | 0.63% | 927.9 | 0.51% | motif file (matrix) | svg |
| 485 | G A C T G A C T A T C G C G A T G T C A G T A C A G C T C G A T A C G T G T A C | SPL11(SBP)/col100-SPL11-DAP-Seq(GSE60143)/Homer | 1e-2 | -6.445e+00 | 0.0034 | 3051.0 | 7.71% | 13275.2 | 7.28% | motif file (matrix) | svg |
| 486 | C G T A C G T A C G T A C G A T C G T A A C G T A G T C G C A T | At4g01280(MYBrelated)/colamp-At4g01280-DAP-Seq(GSE60143)/Homer | 1e-2 | -6.426e+00 | 0.0035 | 1716.0 | 4.33% | 7313.7 | 4.01% | motif file (matrix) | svg |
| 487 | G A C T A G T C C G T A C G T A A G T C A G C T A C T G G A C T G T A C A T G C | MYB77(MYB)/col-MYB77-DAP-Seq(GSE60143)/Homer | 1e-2 | -6.409e+00 | 0.0035 | 10531.0 | 26.60% | 47223.0 | 25.88% | motif file (matrix) | svg |
| 488 | G T A C C T G A A G T C A G T C A C T G G T C A G A T C G C A T | At1g75490(AP2EREBP)/colamp-At1g75490-DAP-Seq(GSE60143)/Homer | 1e-2 | -6.409e+00 | 0.0035 | 17118.0 | 43.24% | 77415.1 | 42.43% | motif file (matrix) | svg |
| 489 | G A C T C T A G C T A G C T A G A C T G T C G A C T G A C T A G C T A G C T A G G T A C G T C A | ZNF467(Zf)/HEK293-ZNF467.GFP-ChIP-Seq(GSE58341)/Homer | 1e-2 | -6.362e+00 | 0.0037 | 2811.0 | 7.10% | 12205.7 | 6.69% | motif file (matrix) | svg |
| 490 | G C T A C G T A C G T A C G T A C G T A A C G T C G T A A C G T A G T C A G C T G C A T G C T A G C T A G C T A G C T A | LCL1(MYBrelated)/colamp-LCL1-DAP-Seq(GSE60143)/Homer | 1e-2 | -6.360e+00 | 0.0037 | 330.0 | 0.83% | 1264.2 | 0.69% | motif file (matrix) | svg |
| 491 | T G A C G C T A T G A C C G T A T C A G G A T C C G T A C A T G C A T G C T A G C T A G C T A G | Unknown-ESC-element(?)/mES-Nanog-ChIP-Seq(GSE11724)/Homer | 1e-2 | -6.191e+00 | 0.0044 | 1231.0 | 3.11% | 5180.1 | 2.84% | motif file (matrix) | svg |
| 492 | C G A T T C G A G T A C A C T G A C G T T C A G G C A T T G C A G C T A G C A T C G T A A G T C C G T A G T A C C A T G | ANAC087(NAC)/col-ANAC087-DAP-Seq(GSE60143)/Homer | 1e-2 | -6.125e+00 | 0.0047 | 1906.0 | 4.81% | 8177.3 | 4.48% | motif file (matrix) | svg |
| 493 | T A G C A G T C T G A C A G T C C T A G A T C G A G T C C A T G T G A C A G T C G T A C A G T C A G T C G C A T C T A G A T C G G C A T A C T G A T C G G A T C | BORIS(Zf)/K562-CTCFL-ChIP-Seq(GSE32465)/Homer | 1e-2 | -6.124e+00 | 0.0047 | 386.0 | 0.97% | 1507.7 | 0.83% | motif file (matrix) | svg |
| 494 | T C A G C G T A A G T C A G C T C G T A A G T C C T G A C G T A A G T C G C A T A G T C A G T C A G T C C T G A A C T G T G C A T C G A C A T G A T C G G A T C | Ronin(THAP)/ES-Thap11-ChIP-Seq(GSE51522)/Homer | 1e-2 | -6.097e+00 | 0.0048 | 39.0 | 0.10% | 101.9 | 0.06% | motif file (matrix) | svg |
| 495 | C G T A C T G A C G T A C G T A C G T A A C G T C G T A A C G T A G T C G C A T C G A T C G A T | AT3G10113(MYBrelated)/col-AT3G10113-DAP-Seq(GSE60143)/Homer | 1e-2 | -5.962e+00 | 0.0055 | 1572.0 | 3.97% | 6702.4 | 3.67% | motif file (matrix) | svg |
| 496 | G A C T G T A C G A C T A G T C T C A G C T G A A G T C A G T C C T A G C G A T A G C T A T G C C T G A C A G T A G C T | AT4G27900(C2C2COlike)/col-AT4G27900-DAP-Seq(GSE60143)/Homer | 1e-2 | -5.956e+00 | 0.0055 | 84.0 | 0.21% | 268.9 | 0.15% | motif file (matrix) | svg |
| 497 | C G A T C G T A A C T G C G T A A C G T C G T A A C G T A C G T C G A T G C A T C G A T C G A T | AT2G28920(ND)/col-AT2G28920-DAP-Seq(GSE60143)/Homer | 1e-2 | -5.916e+00 | 0.0057 | 634.0 | 1.60% | 2580.0 | 1.41% | motif file (matrix) | svg |
| 498 | T C G A C A T G C A T G A C G T A T G C T C G A C T G A A G C T T A C G T G C A G T A C G A T C A G C T A G T C | FXR(NR),IR1/Liver-FXR-ChIP-Seq(Chong\_et\_al.)/Homer | 1e-2 | -5.737e+00 | 0.0068 | 1438.0 | 3.63% | 6121.2 | 3.35% | motif file (matrix) | svg |
| 499 | C G T A C A G T G T A C A T G C C T A G C G T A A C G T A G T C T C G A T C A G | GATA19(C2C2gata)/colamp-GATA19-DAP-Seq(GSE60143)/Homer | 1e-2 | -5.699e+00 | 0.0070 | 727.0 | 1.84% | 2992.9 | 1.64% | motif file (matrix) | svg |
| 500 | T G C A A G C T C T G A A T C G G A C T C T A G G T A C G A T C G T C A A G T C G T A C G A C T C T A G A T C G G C A T C A T G C A T G G A T C G T A C C T G A | CTCF(Zf)/CD4+-CTCF-ChIP-Seq(Barski\_et\_al.)/Homer | 1e-2 | -5.560e+00 | 0.0081 | 237.0 | 0.60% | 894.1 | 0.49% | motif file (matrix) | svg |
| 501 | T G C A A T G C A C G T A C G T A C G T A T G C C T A G A C G T A C G T A G C T G A T C A G C T | T1ISRE(IRF)/ThioMac-Ifnb-Expression/Homer | 1e-2 | -5.560e+00 | 0.0081 | 46.0 | 0.12% | 130.2 | 0.07% | motif file (matrix) | svg |
| 502 | G C T A C T G A C G T A C G T A C T G A C T G A C G A T G T C A A C G T A G T C G C A T G C A T | At5g52660(MYBrelated)/colamp-At5g52660-DAP-Seq(GSE60143)/Homer | 1e-2 | -5.467e+00 | 0.0088 | 1082.0 | 2.73% | 4561.2 | 2.50% | motif file (matrix) | svg |
| 503 | C T G A C T A G T C G A C G T A A T G C C G T A A T C G C G A T T A G C G C A T A T C G G C A T A G C T G A T C G A C T A G C T | ARE(NR)/LNCAP-AR-ChIP-Seq(GSE27824)/Homer | 1e-2 | -5.444e+00 | 0.0090 | 550.0 | 1.39% | 2233.9 | 1.22% | motif file (matrix) | svg |
| 504 | G A C T A G C T G T A C G A C T C T G A A C T G G T C A C T G A A T G C T A C G G A C T A C G T A G T C G A C T C T G A | HRE(HSF)/Striatum-HSF1-ChIP-Seq(GSE38000)/Homer | 1e-2 | -5.421e+00 | 0.0092 | 390.0 | 0.99% | 1545.8 | 0.85% | motif file (matrix) | svg |
| 505 | C T A G T A C G G A T C G T C A T G C A A C G T T G C A G C T A T C G A T G C A | Hoxa9(Homeobox)/ChickenMSG-Hoxa9.Flag-ChIP-Seq(GSE86088)/Homer | 1e-2 | -5.384e+00 | 0.0095 | 11268.0 | 28.46% | 50742.7 | 27.81% | motif file (matrix) | svg |
| 506 | C T A G C A T G C A T G T A C G A G T C G C A T A G C T C T A G A C G T A G T C G A C T A C T G A C T G A C T G T C G A | Zfp809(Zf)/ES-Zfp809-ChIP-Seq(GSE70799)/Homer | 1e-2 | -5.250e+00 | 0.0109 | 528.0 | 1.33% | 2145.7 | 1.18% | motif file (matrix) | svg |
| 507 | C T A G C T A G T C G A C G T A A T G C C G T A A T C G T C G A T A C G G C A T A C T G C A G T T A G C G A T C G A C T | MRE(NR)/Neuro2A-NR3C2-ChIPnexus(GSE115417)/Homer | 1e-2 | -5.129e+00 | 0.0123 | 4473.0 | 11.30% | 19814.9 | 10.86% | motif file (matrix) | svg |
| 508 | G C A T G C A T C T G A A C G T C T G A A C G T C G T A C G T A C G T A A G T C G T C A G T C A | Foxf1(Forkhead)/Lung-Foxf1-ChIP-Seq(GSE77951)/Homer | 1e-2 | -4.981e+00 | 0.0142 | 2456.0 | 6.20% | 10724.6 | 5.88% | motif file (matrix) | svg |
| 509 | G T A C G A T C C A G T A G T C A G T C A G T C T G C A G A T C C T G A A T G C G T C A A C G T | WT1(Zf)/Kidney-WT1-ChIP-Seq(GSE90016)/Homer | 1e-2 | -4.896e+00 | 0.0154 | 2199.0 | 5.55% | 9577.6 | 5.25% | motif file (matrix) | svg |
| 510 | C T A G C T G A A G T C G C T A C G A T A C T G G A C T G A T C G A T C C T G A C T A G C T G A T G A C G C T A C G A T T C A G G A C T G A T C G A T C T G A C | p53(p53)/Saos-p53-ChIP-Seq(GSE15780)/Homer | 1e-2 | -4.892e+00 | 0.0154 | 345.0 | 0.87% | 1369.1 | 0.75% | motif file (matrix) | svg |
| 511 | C T A G C T G A A G T C G C T A C G A T A C T G G A C T G A T C G A T C C T G A C T A G C T G A T G A C G C T A C G A T T C A G G A C T G A T C G A T C T G A C | p53(p53)/Saos-p53-ChIP-Seq/Homer | 1e-2 | -4.892e+00 | 0.0154 | 345.0 | 0.87% | 1369.1 | 0.75% | motif file (matrix) | svg |
| 512 | T G C A G C A T A G C T G C A T A G T C A G T C A G T C C T G A A C T G T C G A T C G A C A G T A T C G A G T C G A T C | ZNF143\|STAF(Zf)/CUTLL-ZNF143-ChIP-Seq(GSE29600)/Homer | 1e-2 | -4.841e+00 | 0.0162 | 758.0 | 1.91% | 3166.6 | 1.74% | motif file (matrix) | svg |
| 513 | G C A T T C A G C T G A A T C G A C T G C G A T G A T C C T G A | THRb(NR)/Liver-NR1A2-ChIP-Seq(GSE52613)/Homer | 1e-2 | -4.777e+00 | 0.0172 | 14773.0 | 37.31% | 66912.6 | 36.67% | motif file (matrix) | svg |
| 514 | T C G A A C G T A C T G C G T A A G T C C T A G A G C T T G A C | TGA10(bZIP)/colamp-TGA10-DAP-Seq(GSE60143)/Homer | 1e-2 | -4.769e+00 | 0.0173 | 3915.0 | 9.89% | 17327.9 | 9.50% | motif file (matrix) | svg |
