## Supplemental dataset for "Hybrid CNN and Multi-Head Attention Model for Analyzing Epigenetic Mechanisms and Gene Expression Across Fungal Phylogenetic Distances": NcrassaModel_FgramTest_K4me2locs_homerResults.html

/projects/wg-feeds/SHAP/NcrassaModel\_FgramTest\_K4me2locs\_SHAP\_noDup\_HOMER// - Homer de novo Motif Results


### Homer *de novo* Motif Results (/projects/wg-feeds/SHAP/NcrassaModel\_FgramTest\_K4me2locs\_SHAP\_noDup\_HOMER//)

Non-redundant Motif File of Results  
Known Motif Enrichment Results  
Gene Ontology Enrichment Results  
If Homer is having trouble matching a motif to a known motif, try copy/pasting the matrix file into
STAMP  
More information on motif finding results: HOMER
| Description of Results
| Tips
  
Total target sequences = 15607  
Total background sequences = 73243  
\* - possible false positive  

|  |  |  |  |  |  |  |  |  |
| --- | --- | --- | --- | --- | --- | --- | --- | --- |
| Rank | Motif | P-value | log P-pvalue | % of Targets | % of Background | STD(Bg STD) | Best Match/Details | Motif File |
| 1 | T G A C T C G A C G T A A T C G T C A G G A C T T G A C T C G A C G T A T A C G T C G A G C T A | 1e-2328 | -5.361e+03 | 70.49% | 26.18% | 647.1bp (703.3bp) | SF1(NR)/H295R-Nr5a1-ChIP-Seq(GSE44220)/Homer(0.736) More Information \| Similar Motifs Found | motif file (matrix) |
| 2 | A G T C G C A T A G C T A T G C G C A T A G C T A T G C G C A T A G C T A T G C G C A T A G C T | 1e-2226 | -5.126e+03 | 51.35% | 12.73% | 668.8bp (720.8bp) | Unknown4/Arabidopsis-Promoters/Homer(0.845) More Information \| Similar Motifs Found | motif file (matrix) |
| 3 | T A G C T C G A G C T A T A C G T G C A G C T A A T C G T G A C G A C T A T G C T C G A C G T A | 1e-1981 | -4.562e+03 | 76.74% | 35.42% | 652.8bp (719.7bp) | MATR3(RRM)/Homo\_sapiens-RNCMPT00037-PBM/HughesRNA(0.673) More Information \| Similar Motifs Found | motif file (matrix) |
| 4 | A T G C C G A T A G C T A T C G C G T A A T G C T C G A C T G A A T C G T G A C C G A T A G T C | 1e-1842 | -4.243e+03 | 59.52% | 21.27% | 643.5bp (705.4bp) | RIM101/MA0368.1/Jaspar(0.629) More Information \| Similar Motifs Found | motif file (matrix) |
| 5 | A G C T A G C T A C G T A G C T A G C T A C G T A G C T A G C T A C G T A G C T | 1e-1700 | -3.915e+03 | 69.67% | 31.36% | 672.4bp (710.9bp) | SeqBias: polyA-repeat(0.931) More Information \| Similar Motifs Found | motif file (matrix) |
| 6 | A T G C C T G A C G T A T A G C C T G A C T G A A T G C A G T C C G A T A G C T A T C G C T G A | 1e-1485 | -3.422e+03 | 80.77% | 45.54% | 642.8bp (710.5bp) | ftz-f1/MA2311.1/Jaspar(0.593) More Information \| Similar Motifs Found | motif file (matrix) |
| 7 | A G T C C G T A G C A T A T G C C G T A C G A T A T C G C T A G G T A C A G T C C T A G G T A C | 1e-1398 | -3.220e+03 | 76.38% | 41.68% | 643.5bp (707.3bp) | ZML2(C2C2gata)/col-ZML2-DAP-Seq(GSE60143)/Homer(0.701) More Information \| Similar Motifs Found | motif file (matrix) |
| 8 | C T A G A C G T A G T C T G A C C G T A C A G T T A C G A C G T A G T C G A C T A C G T A T G C | 1e-1245 | -2.867e+03 | 76.54% | 43.89% | 643.5bp (709.6bp) | Prdm15/MA1616.2/Jaspar(0.686) More Information \| Similar Motifs Found | motif file (matrix) |
| 9 | A G C T A G T C A C G T A G T C C A G T A T G C A G T C A G T C A C G T A G T C | 1e-1149 | -2.646e+03 | 73.17% | 41.56% | 666.5bp (713.8bp) | SeqBias: GA-repeat(0.883) More Information \| Similar Motifs Found | motif file (matrix) |
| 10 | C A G T A G T C C T A G T G C A A T G C G T C A C G A T A T G C | 1e-1085 | -2.499e+03 | 81.34% | 51.72% | 649.8bp (709.9bp) | ZNF135/MA1587.1/Jaspar(0.685) More Information \| Similar Motifs Found | motif file (matrix) |
| 11 | A T G C T G A C G C A T A T C G C G T A C T G A T A G C G A T C C G A T A T C G | 1e-965 | -2.222e+03 | 74.99% | 46.24% | 656.8bp (709.9bp) | ZNF708/MA1730.2/Jaspar(0.643) More Information \| Similar Motifs Found | motif file (matrix) |
| 12 | C T A G C T A G G T A C C T A G C T A G G T A C C T A G C T A G G T C A C T A G C T A G G T A C | 1e-939 | -2.164e+03 | 61.06% | 32.56% | 677.4bp (694.1bp) | DREB2F/MA1242.1/Jaspar(0.827) More Information \| Similar Motifs Found | motif file (matrix) |
| 13 | G T A C A G C T A G T C A G C T A C G T A T G C T G C A C G T A | 1e-746 | -1.719e+03 | 79.29% | 54.80% | 650.5bp (709.3bp) | At2g41835(C2H2)/col-At2g41835-DAP-Seq(GSE60143)/Homer(0.709) More Information \| Similar Motifs Found | motif file (matrix) |
| 14 | C T A G A G T C T G A C C G T A A C G T A C T G A C T G A G T C | 1e-705 | -1.624e+03 | 63.91% | 38.98% | 643.9bp (708.4bp) | Rfx1(HTH)/NPC-H3K4me1-ChIP-Seq(GSE16256)/Homer(0.748) More Information \| Similar Motifs Found | motif file (matrix) |
| 15 | A G C T A C G T A C G T A C G T A C T G G A C T A C G T A C T G | 1e-510 | -1.175e+03 | 62.16% | 40.93% | 663.5bp (715.7bp) | HuR(RRM)/Homo\_sapiens-RNCMPT00117-PBM/HughesRNA(0.850) More Information \| Similar Motifs Found | motif file (matrix) |
| 16 | G A C T G C A T A C G T A C T G A G T C A C T G G T C A G T A C T C G A C T G A | 1e-410 | -9.462e+02 | 21.90% | 8.86% | 671.3bp (695.7bp) | CG7903(RRM)/Drosophila\_melanogaster-RNCMPT00144-PBM/HughesRNA(0.766) More Information \| Similar Motifs Found | motif file (matrix) |
| 17 | C G A T C T A G A C G T G A C T C G T A A C T G G A T C G T C A C G T A G A C T A C G T C A T G | 1e-369 | -8.513e+02 | 22.70% | 9.95% | 653.3bp (686.2bp) | HOW(KH)/Drosophila\_melanogaster-RNCMPT00118-PBM/HughesRNA(0.766) More Information \| Similar Motifs Found | motif file (matrix) |
| 18 | A G T C C A G T G T C A G A T C A G T C A C G T C T G A A G T C G A T C G C A T T C G A A T G C | 1e-109 | -2.510e+02 | 2.98% | 0.66% | 663.3bp (713.1bp) | PK06182.1/MA2354.1/Jaspar(0.874) More Information \| Similar Motifs Found | motif file (matrix) |
| 19 | C T G A A C G T C T A G A C G T C T G A A C G T C T G A A C G T C T G A A C G T C T A G A C G T | 1e-95 | -2.194e+02 | 3.55% | 1.05% | 534.3bp (668.5bp) | cg/MA2107.1/Jaspar(0.799) More Information \| Similar Motifs Found | motif file (matrix) |
| 20 | C G T A G T C A G C A T C T A G C G T A A C G T C G A T A C G T A G T C T C A G C G A T A G T C | 1e-80 | -1.856e+02 | 1.40% | 0.17% | 678.9bp (772.5bp) | PH0037.1\_Hdx/Jaspar(0.729) More Information \| Similar Motifs Found | motif file (matrix) |
| 21 | C T A G A C T G A C T G C A G T C T G A C A G T A G T C C G T A A C T G C G T A C T G A C G T A | 1e-72 | -1.674e+02 | 1.04% | 0.08% | 573.2bp (513.0bp) | PB0059.1\_Six6\_1/Jaspar(0.744) More Information \| Similar Motifs Found | motif file (matrix) |
