## Supplemental dataset for "Hybrid CNN and Multi-Head Attention Model for Analyzing Epigenetic Mechanisms and Gene Expression Across Fungal Phylogenetic Distances": NcrassaModel_FgramTest_K4me2locs_knownResults.html

/projects/wg-feeds/SHAP/NcrassaModel\_FgramTest\_K4me2locs\_SHAP\_noDup\_HOMER/ - Homer Known Motif Enrichment Results


### Homer Known Motif Enrichment Results (/projects/wg-feeds/SHAP/NcrassaModel\_FgramTest\_K4me2locs\_SHAP\_noDup\_HOMER/)

Homer *de novo* Motif Results  
Gene Ontology Enrichment Results  
Known Motif Enrichment Results (txt file)  
Total Target Sequences = 15609, Total Background Sequences = 73240

|  |  |  |  |  |  |  |  |  |  |  |  |
| --- | --- | --- | --- | --- | --- | --- | --- | --- | --- | --- | --- |
| Rank | Motif | Name | P-value | log P-pvalue | q-value (Benjamini) | # Target Sequences with Motif | % of Targets Sequences with Motif | # Background Sequences with Motif | % of Background Sequences with Motif | Motif File | SVG |
| 1 | T A G C G T C A G A C T T A G C G T C A G A C T A G T C G C T A G A C T G A T C | ZML2(C2C2gata)/col-ZML2-DAP-Seq(GSE60143)/Homer | 1e-1354 | -3.118e+03 | 0.0000 | 5552.0 | 35.57% | 6613.6 | 9.03% | motif file (matrix) | svg |
| 2 | A T G C C A G T A C G T A G T C C A T G A C G T A G T C A C G T A G C T G A T C | Unknown4/Arabidopsis-Promoters/Homer | 1e-1180 | -2.719e+03 | 0.0000 | 11257.0 | 72.13% | 29317.4 | 40.03% | motif file (matrix) | svg |
| 3 | C T A G T C A G C T G A C T G A C T A G C T G A C A T G C A T G C T G A C T A G C T A G C G T A C T A G C G T A G T C A | TF3A(C2H2)/col-TF3A-DAP-Seq(GSE60143)/Homer | 1e-1154 | -2.658e+03 | 0.0000 | 9361.0 | 59.98% | 21046.9 | 28.74% | motif file (matrix) | svg |
| 4 | T G A C C G T A C T G A A C T G A C T G G A C T G A T C T G C A G T A C T A C G | SF1(NR)/H295R-Nr5a1-ChIP-Seq(GSE44220)/Homer | 1e-740 | -1.705e+03 | 0.0000 | 5669.0 | 36.32% | 10924.6 | 14.92% | motif file (matrix) | svg |
| 5 | A C G T G A C T A T G C G C T A C T G A C T A G A C T G G A C T A G T C C G T A | Nr5a2(NR)/mES-Nr5a2-ChIP-Seq(GSE19019)/Homer | 1e-679 | -1.565e+03 | 0.0000 | 6448.0 | 41.31% | 14236.0 | 19.44% | motif file (matrix) | svg |
| 6 | A C G T G A C T T A G C C G T A C T G A C A T G C T A G G A C T G A T C C G T A | Nr5a2(NR)/Pancreas-LRH1-ChIP-Seq(GSE34295)/Homer | 1e-639 | -1.473e+03 | 0.0000 | 7732.0 | 49.54% | 19637.3 | 26.81% | motif file (matrix) | svg |
| 7 | G C A T G A T C C T A G C G T A G C A T C G T A G C A T A G T C C T A G C G T A G C A T C G A T | AT5G22990(C2H2)/col-AT5G22990-DAP-Seq(GSE60143)/Homer | 1e-522 | -1.203e+03 | 0.0000 | 9027.0 | 57.84% | 26707.7 | 36.47% | motif file (matrix) | svg |
| 8 | A C G T A T G C A C G T A C G T A G C T A G T C A G C T A G C T A G C T A G C T A G C T | hTCT(CPE) | 1e-460 | -1.061e+03 | 0.0000 | 11463.0 | 73.45% | 39425.2 | 53.83% | motif file (matrix) | svg |
| 9 | G C A T A G C T A C G T A C G T A C T G A C G T G A T C A C G T A C G T A G C T C G A T G C A T A G T C G A C T C A G T | IDD5(C2H2)/colamp-IDD5-DAP-Seq(GSE60143)/Homer | 1e-429 | -9.883e+02 | 0.0000 | 4764.0 | 30.52% | 10821.3 | 14.78% | motif file (matrix) | svg |
| 10 | C A T G A G C T T A C G G T C A G T A C T A G C A G C T G A C T A T C G T C G A | Esrrb(NR)/mES-Esrrb-ChIP-Seq(GSE11431)/Homer | 1e-384 | -8.853e+02 | 0.0000 | 7146.0 | 45.79% | 20655.2 | 28.20% | motif file (matrix) | svg |
| 11 | G C T A G C A T A C G T A C G T A G C T G A T C G A C T G A C T A G C T A C G T A C G T A G C T | RLR1?/SacCer-Promoters/Homer | 1e-360 | -8.309e+02 | 0.0000 | 2279.0 | 14.60% | 3516.6 | 4.80% | motif file (matrix) | svg |
| 12 | G A T C G C A T A G T C A G C T G A T C A G C T G A T C G A C T G A T C G A C T A G T C A C G T G A T C A G C T G A T C | GAGA-repeat/SacCer-Promoters/Homer | 1e-354 | -8.155e+02 | 0.0000 | 13710.0 | 87.85% | 53879.5 | 73.57% | motif file (matrix) | svg |
| 13 | G A C T A G C T A G C T C T A G A C G T G A T C A C G T A C G T G A C T C G A T G C A T A G T C | IDD4(C2H2)/col-IDD4-DAP-Seq(GSE60143)/Homer | 1e-350 | -8.076e+02 | 0.0000 | 6466.0 | 41.43% | 18377.5 | 25.09% | motif file (matrix) | svg |
| 14 | A C G T A C T G C G T A A C G T A C T G A C T G C G T A C G T A | HAP3(CCAATHAP3)/col-HAP3-DAP-Seq(GSE60143)/Homer | 1e-350 | -8.066e+02 | 0.0000 | 4961.0 | 31.79% | 12504.3 | 17.07% | motif file (matrix) | svg |
| 15 | T C G A T G C A C A G T T C G A G A T C A G T C C G T A C G T A A C T G A G T C C G T A C G T A T C A G C G A T A G T C | AT5G25475(ABI3VP1)/col-AT5G25475-DAP-Seq(GSE60143)/Homer | 1e-348 | -8.032e+02 | 0.0000 | 10502.0 | 67.29% | 36535.7 | 49.89% | motif file (matrix) | svg |
| 16 | T G A C C T A G T C A G G T C A C G T A T C A G C G A T T C A G T C G A T G C A C T G A T A G C | PU.1-IRF(ETS:IRF)/Bcell-PU.1-ChIP-Seq(GSE21512)/Homer | 1e-329 | -7.592e+02 | 0.0000 | 7377.0 | 47.27% | 22531.3 | 30.76% | motif file (matrix) | svg |
| 17 | C A T G G A C T T A C G G T C A G T A C G A T C G A C T A G C T A T C G T C G A T A C G T A G C | ERRg(NR)/Kidney-ESRRG-ChIP-Seq(GSE104905)/Homer | 1e-315 | -7.271e+02 | 0.0000 | 8036.0 | 51.49% | 25650.4 | 35.02% | motif file (matrix) | svg |
| 18 | A G T C A G T C C G A T A C G T A C G T A C T G A C G T A G C T A G T C A G T C | Sox4(HMG)/proB-Sox4-ChIP-Seq(GSE50066)/Homer | 1e-302 | -6.973e+02 | 0.0000 | 7253.0 | 46.47% | 22475.2 | 30.69% | motif file (matrix) | svg |
| 19 | C T G A C G A T C T A G T C A G G A T C C T G A T C A G G A T C C T G A A C T G A G T C G C T A A C G T A G T C G C A T | PRDM9(Zf)/Testis-DMC1-ChIP-Seq(GSE35498)/Homer | 1e-297 | -6.839e+02 | 0.0000 | 3891.0 | 24.93% | 9299.9 | 12.70% | motif file (matrix) | svg |
| 20 | C T G A C G A T C A T G A T C G G C A T C A T G G C T A A G T C | ASHR1(ND)/col-ASHR1-DAP-Seq(GSE60143)/Homer | 1e-274 | -6.313e+02 | 0.0000 | 10252.0 | 65.69% | 36795.2 | 50.24% | motif file (matrix) | svg |
| 21 | G A C T G C A T A C G T A C G T A C T G C G T A A G T C A G C T C G A T A T C G G C A T A C T G C G A T C T A G C G T A | WRKY50(WRKY)/col-WRKY50-DAP-Seq(GSE60143)/Homer | 1e-265 | -6.115e+02 | 0.0000 | 10000.0 | 64.07% | 35753.8 | 48.82% | motif file (matrix) | svg |
| 22 | T C G A A C T G C A T G A G C T A G T C C G T A C T G A C T A G A C T G C G A T A T G C C T G A | RAR:RXR(NR),DR0/ES-RAR-ChIP-Seq(GSE56893)/Homer | 1e-263 | -6.072e+02 | 0.0000 | 1785.0 | 11.44% | 2857.0 | 3.90% | motif file (matrix) | svg |
| 23 | A T G C A G T C G C A T A G C T A C G T T C A G C G A T A G C T G A T C A T C G | Sox10(HMG)/SciaticNerve-Sox3-ChIP-Seq(GSE35132)/Homer | 1e-259 | -5.970e+02 | 0.0000 | 11205.0 | 71.79% | 41872.0 | 57.17% | motif file (matrix) | svg |
| 24 | A G C T A C G T A C T G A T G C A G T C C G T A C T G A T A C G | NF1-halfsite(CTF)/LNCaP-NF1-ChIP-Seq(Unpublished)/Homer | 1e-256 | -5.916e+02 | 0.0000 | 11113.0 | 71.21% | 41455.4 | 56.60% | motif file (matrix) | svg |
| 25 | C A T G G C T A C T A G T A C G C G T A T C A G C G T A A C T G C G T A C A T G C T G A C G T A | BPC1(BBRBPC)/colamp-BPC1-DAP-Seq(GSE60143)/Homer | 1e-251 | -5.792e+02 | 0.0000 | 3807.0 | 24.39% | 9604.9 | 13.11% | motif file (matrix) | svg |
| 26 | A T G C G A T C C G A T A C G T A C G T A C T G C A G T A G C T | Sox3(HMG)/NPC-Sox3-ChIP-Seq(GSE33059)/Homer | 1e-246 | -5.679e+02 | 0.0000 | 11658.0 | 74.70% | 44484.8 | 60.74% | motif file (matrix) | svg |
| 27 | G T C A G C T A G C T A T C G A A T C G A C G T A G T C T C G A T C G A T G A C | WRKY40(WRKY)/colamp-WRKY40-DAP-Seq(GSE60143)/Homer | 1e-228 | -5.268e+02 | 0.0000 | 7113.0 | 45.58% | 23312.6 | 31.83% | motif file (matrix) | svg |
| 28 | G A C T A C G T A C G T A C T G A C G T A G T C G C A T A G C T G C A T G C A T G A C T A G C T | SGR5(C2H2)/colamp-SGR5-DAP-Seq(GSE60143)/Homer | 1e-219 | -5.058e+02 | 0.0000 | 5674.0 | 36.36% | 17405.7 | 23.77% | motif file (matrix) | svg |
| 29 | A C G T A C G T A C G T A C G T A C G T A C G T A C G T A C G T A C G T A C G T | VRN1(ABI3VP1)/col-VRN1-DAP-Seq(GSE60143)/Homer | 1e-211 | -4.872e+02 | 0.0000 | 506.0 | 3.24% | 219.3 | 0.30% | motif file (matrix) | svg |
| 30 | G T A C A C G T A C G T A T C G C A G T C G A T A T C G G C T A T G C A T A G C C G T A G T C A C A T G A G C T G C T A | ANAC013(NAC)/col-ANAC013-DAP-Seq(GSE60143)/Homer | 1e-201 | -4.631e+02 | 0.0000 | 5234.0 | 33.54% | 15968.3 | 21.80% | motif file (matrix) | svg |
| 31 | C T A G A G T C A G T C A C T G C G T A A G T C C T G A G A C T | DDF1(AP2EREBP)/col-DDF1-DAP-Seq(GSE60143)/Homer | 1e-200 | -4.614e+02 | 0.0000 | 9986.0 | 63.98% | 37184.6 | 50.77% | motif file (matrix) | svg |
| 32 | C A T G G A C T C T A G C A T G C A G T C G A T C T A G C A T G C G A T C G T A C T A G C A G T C G A T C T A G C A T G | AT1G24250(Orphan)/col-AT1G24250-DAP-Seq(GSE60143)/Homer | 1e-199 | -4.597e+02 | 0.0000 | 3793.0 | 24.30% | 10338.1 | 14.12% | motif file (matrix) | svg |
| 33 | C G A T C T A G C G T A G A C T C A G T C T A G C G T A A G C T C A T G C T A G | HOXA1(Homeobox)/mES-Hoxa1-ChIP-Seq(SRP084292)/Homer | 1e-194 | -4.488e+02 | 0.0000 | 3982.0 | 25.51% | 11135.7 | 15.20% | motif file (matrix) | svg |
| 34 | A T G C T C G A T A C G A C G T A T G C A G T C A C G T A G T C A G T C G A T C | Znf263(Zf)/K562-Znf263-ChIP-Seq(GSE31477)/Homer | 1e-189 | -4.365e+02 | 0.0000 | 9619.0 | 61.63% | 35687.6 | 48.73% | motif file (matrix) | svg |
| 35 | A G T C T G C A T C G A C T G A A C T G C A T G A C G T A T G C G T C A T A C G | Erra(NR)/HepG2-Erra-ChIP-Seq(GSE31477)/Homer | 1e-187 | -4.329e+02 | 0.0000 | 11182.0 | 71.65% | 43424.6 | 59.29% | motif file (matrix) | svg |
| 36 | C T G A T C A G A G T C C G T A A T C G A T G C C G A T A C T G A G T C G A C T A T C G A G T C | MyoD(bHLH)/Myotube-MyoD-ChIP-Seq(GSE21614)/Homer | 1e-187 | -4.315e+02 | 0.0000 | 4538.0 | 29.08% | 13438.4 | 18.35% | motif file (matrix) | svg |
| 37 | C G A T C T G A A G T C A C G T A C G T A T C G G A C T C T A G C G A T G A C T C G T A A T G C C G T A G T C A A C T G | ANAC011(NAC)/col-ANAC011-DAP-Seq(GSE60143)/Homer | 1e-186 | -4.301e+02 | 0.0000 | 3732.0 | 23.91% | 10323.0 | 14.09% | motif file (matrix) | svg |
| 38 | A G T C C A T G A C G T A C G T A C T G C G T A A G T C G A C T G C A T G C T A | WRKY28(WRKY)/col-WRKY28-DAP-Seq(GSE60143)/Homer | 1e-183 | -4.235e+02 | 0.0000 | 10905.0 | 69.87% | 42131.1 | 57.53% | motif file (matrix) | svg |
| 39 | C G T A T G A C T A G C T G C A A C T G A C T G C G T A C G T A T C A G G A C T | ELF3(ETS)/PDAC-ELF3-ChIP-Seq(GSE64557)/Homer | 1e-181 | -4.188e+02 | 0.0000 | 4983.0 | 31.93% | 15332.1 | 20.93% | motif file (matrix) | svg |
| 40 | A G T C G A T C G C T A C G A T C A G T T A C G C G A T A G C T A G T C A T C G | SOX1(HMG)/NPC-SOX1-ChIP-Seq(GSE138215)/Homer | 1e-178 | -4.112e+02 | 0.0000 | 12182.0 | 78.05% | 48841.8 | 66.69% | motif file (matrix) | svg |
| 41 | C G T A T A G C T A G C T G C A A C T G C T A G C G T A C G T A T C A G G A C T | EHF(ETS)/LoVo-EHF-ChIP-Seq(GSE49402)/Homer | 1e-178 | -4.112e+02 | 0.0000 | 8321.0 | 53.32% | 29883.9 | 40.80% | motif file (matrix) | svg |
| 42 | A G C T A G C T A G C T A C T G A C G T A G T C A C T G A C G T G A C T C G A T G C A T A T C G | IDD7(C2H2)/col-IDD7-DAP-Seq(GSE60143)/Homer | 1e-176 | -4.060e+02 | 0.0000 | 4286.0 | 27.46% | 12653.5 | 17.28% | motif file (matrix) | svg |
| 43 | C T G A T C G A G T A C A C G T A C G T A T C G A C G T C G A T A T C G G C T A G T A C A T G C C G T A T G C A C A T G | ANAC103(NAC)/col-ANAC103-DAP-Seq(GSE60143)/Homer | 1e-173 | -3.991e+02 | 0.0000 | 4718.0 | 30.23% | 14434.3 | 19.71% | motif file (matrix) | svg |
| 44 | T A G C C G T A C T G A T A C G C G T A A C G T A C T G A C T G A G T C T A C G C T A G G T A C | YY1(Zf)/Promoter/Homer | 1e-173 | -3.987e+02 | 0.0000 | 1069.0 | 6.85% | 1573.9 | 2.15% | motif file (matrix) | svg |
| 45 | C A T G G T A C A C T G G T C A A G C T T A C G T G C A A T C G T G A C C A G T | TOD6?/SacCer-Promoters/Homer | 1e-170 | -3.915e+02 | 0.0000 | 3324.0 | 21.30% | 9070.0 | 12.38% | motif file (matrix) | svg |
| 46 | C T G A T C G A C G T A A T G C C G T A C G T A C G A T C T A G T C A G G A T C | Sox15(HMG)/CPA-Sox15-ChIP-Seq(GSE62909)/Homer | 1e-169 | -3.899e+02 | 0.0000 | 7975.0 | 51.10% | 28540.0 | 38.97% | motif file (matrix) | svg |
| 47 | T A C G G A C T T G A C C G T A A C G T G A T C G T C A C G T A A C G T A T G C C G T A G A C T | HOXA2(Homeobox)/mES-Hoxa2-ChIP-Seq(Donaldson\_et\_al.)/Homer | 1e-166 | -3.832e+02 | 0.0000 | 1907.0 | 12.22% | 4130.6 | 5.64% | motif file (matrix) | svg |
| 48 | C T A G G T A C C A T G G A C T C G A T C A T G G T C A G T A C G A C T C G A T C G A T C G A T | WRKY27(WRKY)/colamp-WRKY27-DAP-Seq(GSE60143)/Homer | 1e-166 | -3.825e+02 | 0.0000 | 8718.0 | 55.86% | 32044.5 | 43.75% | motif file (matrix) | svg |
| 49 | C T G A A T G C G C T A G C A T A T G C C G T A T C G A C T G A C T A G T C A G T A C G G T C A | Tcf4(HMG)/Hct116-Tcf4-ChIP-Seq(SRA012054)/Homer | 1e-163 | -3.773e+02 | 0.0000 | 4346.0 | 27.85% | 13135.4 | 17.93% | motif file (matrix) | svg |
| 50 | G A C T G C A T G C A T A G T C A G C T T C G A T A C G G C T A C G T A A C T G G T A C G C A T C G A T A G T C G A C T | HSF3(HSF)/colamp-HSF3-DAP-Seq(GSE60143)/Homer | 1e-161 | -3.715e+02 | 0.0000 | 7148.0 | 45.80% | 25029.4 | 34.17% | motif file (matrix) | svg |
| 51 | G T A C C T G A T A G C C G T A G C T A T C G A T G C A T G A C C T A G G T C A A G T C C G T A C T G A C T G A C G T A | At1g14580(C2H2)/colamp-At1g14580-DAP-Seq(GSE60143)/Homer | 1e-161 | -3.714e+02 | 0.0000 | 1683.0 | 10.78% | 3469.4 | 4.74% | motif file (matrix) | svg |
| 52 | A C G T A G T C A G T C C G A T A C G T A C G T A C T G A C G T A T G C G A C T A C T G T A C G | Sox21(HMG)/ESC-SOX21-ChIP-Seq(GSE110505)/Homer | 1e-160 | -3.690e+02 | 0.0000 | 10988.0 | 70.40% | 43176.6 | 58.95% | motif file (matrix) | svg |
| 53 | A G T C A C G T A C G T T A C G G C A T G C A T A T G C G C T A C G T A A T G C C G T A G T C A A C T G G A T C G C A T | ANAC075(NAC)/col-ANAC075-DAP-Seq(GSE60143)/Homer | 1e-158 | -3.648e+02 | 0.0000 | 5007.0 | 32.08% | 15929.0 | 21.75% | motif file (matrix) | svg |
| 54 | C G A T C G T A G T A C A C G T A C G T T C A G G C A T C A G T T A C G G T C A C G T A A G T C C G T A T G C A C A T G | NAC2(NAC)/colamp-NAC2-DAP-Seq(GSE60143)/Homer | 1e-156 | -3.603e+02 | 0.0000 | 7008.0 | 44.90% | 24535.3 | 33.50% | motif file (matrix) | svg |
| 55 | G T A C C A T G A G C T A C G T A C T G C G T A A G T C G A C T G C A T C G A T | WRKY29(WRKY)/colamp-WRKY29-DAP-Seq(GSE60143)/Homer | 1e-155 | -3.583e+02 | 0.0000 | 9789.0 | 62.72% | 37409.2 | 51.08% | motif file (matrix) | svg |
| 56 | A T G C A G T C G A T C C G T A A C G T A C G T A C T G A C G T A G C T G A T C | Sox2(HMG)/mES-Sox2-ChIP-Seq(GSE11431)/Homer | 1e-154 | -3.558e+02 | 0.0000 | 7355.0 | 47.13% | 26130.2 | 35.68% | motif file (matrix) | svg |
| 57 | A T G C G T A C C G T A A G C T G C A T T A C G A G C T A G C T A G T C A G C T | Sox6(HMG)/Myotubes-Sox6-ChIP-Seq(GSE32627)/Homer | 1e-153 | -3.537e+02 | 0.0000 | 11267.0 | 72.19% | 44768.6 | 61.13% | motif file (matrix) | svg |
| 58 | C T A G T C A G C T G A T C A G T G C A A C T G T C G A T C A G | Trl(Zf)/S2-GAGAfactor-ChIP-Seq(GSE40646)/Homer | 1e-152 | -3.514e+02 | 0.0000 | 12528.0 | 80.27% | 51355.2 | 70.12% | motif file (matrix) | svg |
| 59 | C G A T C G T A G T A C A C G T A C G T T C A G G C A T C A G T T A C G G T C A C G T A A G T C C G T A T G C A C A T G | ANAC053(NAC)/colamp-ANAC053-DAP-Seq(GSE60143)/Homer | 1e-150 | -3.462e+02 | 0.0000 | 6240.0 | 39.98% | 21332.0 | 29.13% | motif file (matrix) | svg |
| 60 | A G T C G A T C C T G A A G T C A G T C C A T G G T C A G A T C C G T A G A T C | DREB26(AP2EREBP)/col-DREB26-DAP-Seq(GSE60143)/Homer | 1e-149 | -3.452e+02 | 0.0000 | 4169.0 | 26.71% | 12715.1 | 17.36% | motif file (matrix) | svg |
| 61 | C G T A G A T C C A T G G C A T G A C T C T A G T C G A T A G C A G C T G C A T | WRKY55(WRKY)/col-WRKY55-DAP-Seq(GSE60143)/Homer | 1e-148 | -3.431e+02 | 0.0000 | 10210.0 | 65.42% | 39640.8 | 54.12% | motif file (matrix) | svg |
| 62 | T C G A T C G A C T G A C G T A A C T G A T G C A C G T A G T C | Lola-I(Zf)/Embryo-LolaI-ChIP-Seq(GSE200870)/Homer | 1e-148 | -3.411e+02 | 0.0000 | 5145.0 | 32.97% | 16735.5 | 22.85% | motif file (matrix) | svg |
| 63 | C G T A G A T C A G C T A C G T A C G T A C T G C G T A G T A C A G C T G C T A C G A T C G A T C G A T G C A T G C T A | WRKY18(WRKY)/col-WRKY18-DAP-Seq(GSE60143)/Homer | 1e-146 | -3.373e+02 | 0.0000 | 12486.0 | 80.00% | 51295.6 | 70.04% | motif file (matrix) | svg |
| 64 | G A C T G A T C G A T C G C T A G T A C A G T C G C T A C T G A G T A C G A T C G C T A G A C T | MYB13(MYB)/col-MYB13-DAP-Seq(GSE60143)/Homer | 1e-146 | -3.363e+02 | 0.0000 | 6874.0 | 44.04% | 24216.1 | 33.06% | motif file (matrix) | svg |
| 65 | G C T A C G T A G C T A C G T A C T G A A C T G A C G T G T A C C G T A C T G A G T A C A C T G | WRKY22(WRKY)/colamp-WRKY22-DAP-Seq(GSE60143)/Homer | 1e-143 | -3.302e+02 | 0.0000 | 6559.0 | 42.03% | 22900.3 | 31.27% | motif file (matrix) | svg |
| 66 | C G T A C T A G C A T G A G C T C T G A C A T G C A G T C G A T C T A G C T A G | MYB30(MYB)/colamp-MYB30-DAP-Seq(GSE60143)/Homer | 1e-143 | -3.300e+02 | 0.0000 | 9505.0 | 60.90% | 36397.6 | 49.70% | motif file (matrix) | svg |
| 67 | C G T A C T A G C A T G G A C T C T G A A C T G A C G T A C G T C T A G C T A G C A T G T C G A | MYB94(MYB)/col-MYB94-DAP-Seq(GSE60143)/Homer | 1e-142 | -3.279e+02 | 0.0000 | 4558.0 | 29.20% | 14447.6 | 19.73% | motif file (matrix) | svg |
| 68 | G T A C A C T G A G T C A G T C C T A G G A T C G T A C C T G A | CRF4(AP2EREBP)/colamp-CRF4-DAP-Seq(GSE60143)/Homer | 1e-142 | -3.274e+02 | 0.0000 | 7056.0 | 45.21% | 25129.8 | 34.31% | motif file (matrix) | svg |
| 69 | G A C T C A G T G C A T C G A T T G A C A C G T A T G C G T A C C T G A A C T G A C T G A G C T | WIP5(C2H2)/colamp-WIP5-DAP-Seq(GSE60143)/Homer | 1e-142 | -3.272e+02 | 0.0000 | 9323.0 | 59.74% | 35565.9 | 48.56% | motif file (matrix) | svg |
| 70 | C G A T C T G A A G T C A C G T A C G T T A C G G C T A C A T G C T A G G C A T C G A T A G T C C G T A G T C A A C T G | ANAC096(NAC)/colamp-ANAC096-DAP-Seq(GSE60143)/Homer | 1e-141 | -3.260e+02 | 0.0000 | 6789.0 | 43.50% | 23961.1 | 32.72% | motif file (matrix) | svg |
| 71 | G A C T A C G T A G C T G A C T A C T G C A G T A G T C A T C G A C G T G C A T G C A T G C A T | MGP(C2H2)/colamp-MGP-DAP-Seq(GSE60143)/Homer | 1e-141 | -3.252e+02 | 0.0000 | 3747.0 | 24.01% | 11224.4 | 15.33% | motif file (matrix) | svg |
| 72 | A G C T C T A G A G T C A G T C A C T G C G T A A G T C C T G A G C A T G C T A C T G A G C A T G C A T C G A T G C A T | CBF4(AP2EREBP)/colamp-CBF4-DAP-Seq(GSE60143)/Homer | 1e-140 | -3.228e+02 | 0.0000 | 12063.0 | 77.29% | 49238.5 | 67.23% | motif file (matrix) | svg |
| 73 | A G T C G A C T A G C T C G A T A T C G G C T A C G A T A T C G C G A T A C T G T A C G A C G T | Tcf7(HMG)/GM12878-TCF7-ChIP-Seq(Encode)/Homer | 1e-138 | -3.182e+02 | 0.0000 | 3280.0 | 21.02% | 9471.3 | 12.93% | motif file (matrix) | svg |
| 74 | G A T C C A T G A C G T A C G T A C T G C G T A A G T C A G C T C G A T G A C T | WRKY8(WRKY)/colamp-WRKY8-DAP-Seq(GSE60143)/Homer | 1e-137 | -3.159e+02 | 0.0000 | 1662.0 | 10.65% | 3676.8 | 5.02% | motif file (matrix) | svg |
| 75 | C G A T T G C A A G T C A C G T A C G T T A C G C G A T C G A T T A C G G C T A G C T A A T G C C G T A G T C A C A T G | ANAC016(NAC)/col-ANAC016-DAP-Seq(GSE60143)/Homer | 1e-136 | -3.139e+02 | 0.0000 | 8835.0 | 56.61% | 33431.6 | 45.65% | motif file (matrix) | svg |
| 76 | C T G A G C A T A C T G C T A G A G T C A C T G A C T G A G T C A C T G T C A G | AT4G18450(AP2EREBP)/col-AT4G18450-DAP-Seq(GSE60143)/Homer | 1e-136 | -3.134e+02 | 0.0000 | 5171.0 | 33.13% | 17133.9 | 23.39% | motif file (matrix) | svg |
| 77 | A T G C C A T G A G C T C A G T C A T G T C G A A G T C G A C T C G A T C G A T C A G T C A G T | WRKY26(WRKY)/colamp-WRKY26-DAP-Seq(GSE60143)/Homer | 1e-135 | -3.111e+02 | 0.0000 | 6239.0 | 39.98% | 21725.9 | 29.66% | motif file (matrix) | svg |
| 78 | A G C T G A T C C T G A A G T C A G T C A C T G C G T A A G T C C T G A G T C A G C A T C G A T G C T A G C A T C G T A | At2g44940(AP2EREBP)/colamp-At2g44940-DAP-Seq(GSE60143)/Homer | 1e-135 | -3.110e+02 | 0.0000 | 6195.0 | 39.69% | 21535.8 | 29.40% | motif file (matrix) | svg |
| 79 | G T C A T G C A T G C A G C T A C G T A G C T A G C T A G C T A | REM19(REM)/colamp-REM19-DAP-Seq(GSE60143)/Homer | 1e-133 | -3.085e+02 | 0.0000 | 2667.0 | 17.09% | 7241.7 | 9.89% | motif file (matrix) | svg |
| 80 | C G A T T C G A G A T C C G A T G C A T T C A G G A C T C G A T G C A T G C T A C T G A A G T C C G T A G T C A C T A G | ANAC005(NAC)/col-ANAC005-DAP-Seq(GSE60143)/Homer | 1e-133 | -3.069e+02 | 0.0000 | 4269.0 | 27.35% | 13478.3 | 18.40% | motif file (matrix) | svg |
| 81 | A G C T A G C T C A T G C T G A G T A C A G T C A G C T A G C T C A G T C T A G | RARa(NR)/K562-RARa-ChIP-Seq(Encode)/Homer | 1e-132 | -3.057e+02 | 0.0000 | 13444.0 | 86.14% | 56889.4 | 77.68% | motif file (matrix) | svg |
| 82 | C G A T T C G A G A T C A C G T A C G T T C A G G C T A C G A T C G T A C G T A C G T A A T G C C G T A T G C A C T A G | ANAC028(NAC)/col-ANAC028-DAP-Seq(GSE60143)/Homer | 1e-131 | -3.036e+02 | 0.0000 | 7358.0 | 47.15% | 26772.0 | 36.55% | motif file (matrix) | svg |
| 83 | T C G A G T A C C A T G A G C T A C G T C A T G G T C A G T A C A G C T G C T A C G A T C A G T | WRKY31(WRKY)/colamp-WRKY31-DAP-Seq(GSE60143)/Homer | 1e-131 | -3.028e+02 | 0.0000 | 7529.0 | 48.24% | 27554.5 | 37.62% | motif file (matrix) | svg |
| 84 | G A T C A G C T C T A G G A T C T G A C C T A G C G T A G T A C C G T A G C A T G T C A C T G A | CBF3(AP2EREBP)/colamp-CBF3-DAP-Seq(GSE60143)/Homer | 1e-130 | -3.016e+02 | 0.0000 | 9774.0 | 62.63% | 38057.7 | 51.96% | motif file (matrix) | svg |
| 85 | C G T A G A C T C A T G C T A G A G T C A C T G A C T G G T A C C A T G T A C G | ERF3(AP2EREBP)/colamp-ERF3-DAP-Seq(GSE60143)/Homer | 1e-130 | -3.012e+02 | 0.0000 | 8652.0 | 55.44% | 32739.2 | 44.70% | motif file (matrix) | svg |
| 86 | C T G A T C A G G T A C G C T A A C T G T G A C G C A T C A T G | SCL(bHLH)/HPC7-Scl-ChIP-Seq(GSE13511)/Homer | 1e-130 | -3.002e+02 | 0.0000 | 14267.0 | 91.41% | 61710.3 | 84.26% | motif file (matrix) | svg |
| 87 | G C A T A G C T A G C T A G C T A C T G A C G T A G T C A C T G A C G T G A C T C G A T G C A T | JKD(C2H2)/col-JKD-DAP-Seq(GSE60143)/Homer | 1e-128 | -2.967e+02 | 0.0000 | 2335.0 | 14.96% | 6116.4 | 8.35% | motif file (matrix) | svg |
| 88 | A G C T G A C T A C G T A C T G A C G T A G T C A C T G A C G T G C A T C G A T | AtIDD11(C2H2)/colamp-AtIDD11-DAP-Seq(GSE60143)/Homer | 1e-128 | -2.965e+02 | 0.0000 | 4123.0 | 26.42% | 12990.4 | 17.74% | motif file (matrix) | svg |
| 89 | A T G C C A T G G C A T G A C T C T A G T C G A G T A C A G C T C G T A G C T A | WRKY75(WRKY)/col-WRKY75-DAP-Seq(GSE60143)/Homer | 1e-128 | -2.950e+02 | 0.0000 | 9163.0 | 58.71% | 35226.7 | 48.10% | motif file (matrix) | svg |
| 90 | G A C T A C T G C G A T A G T C A C T G C T A G A G T C C G T A | Rap210(AP2EREBP)/col-Rap210-DAP-Seq(GSE60143)/Homer | 1e-126 | -2.921e+02 | 0.0000 | 11038.0 | 70.72% | 44394.1 | 60.62% | motif file (matrix) | svg |
| 91 | G C A T T G A C C A T G G A C T C A G T C A T G T C G A G T A C G A C T G C T A C G A T C G A T | WRKY6(WRKY)/colamp-WRKY6-DAP-Seq(GSE60143)/Homer | 1e-126 | -2.919e+02 | 0.0000 | 8621.0 | 55.24% | 32717.9 | 44.67% | motif file (matrix) | svg |
| 92 | A G T C A G T C C T G A A G T C A G T C A C T G C G T A A G T C C T G A T C G A G C A T G A T C C G A T C G A T A C T G | AT3G60490(AP2EREBP)/colamp-AT3G60490-DAP-Seq(GSE60143)/Homer | 1e-126 | -2.904e+02 | 0.0000 | 7254.0 | 46.48% | 26472.1 | 36.14% | motif file (matrix) | svg |
| 93 | C T A G G C A T A C T G C T A G A G T C A C T G A C T G A G T C A C T G T C A G | ERF10(AP2EREBP)/col-ERF10-DAP-Seq(GSE60143)/Homer | 1e-124 | -2.868e+02 | 0.0000 | 8240.0 | 52.80% | 31016.9 | 42.35% | motif file (matrix) | svg |
| 94 | A G C T A G C T A G C T A C T G A C G T A G T C A C T G A C G T G C A T G C A T G C A T A C G T | At5g66730(C2H2)/colamp-At5g66730-DAP-Seq(GSE60143)/Homer | 1e-124 | -2.862e+02 | 0.0000 | 3106.0 | 19.90% | 9073.9 | 12.39% | motif file (matrix) | svg |
| 95 | C A T G A G T C G T A C A C T G A T G C A G T C C A T G G A T C G A T C C T G A | ERF5(AP2EREBP)/colamp-ERF5-DAP-Seq(GSE60143)/Homer | 1e-123 | -2.853e+02 | 0.0000 | 5267.0 | 33.75% | 17849.4 | 24.37% | motif file (matrix) | svg |
| 96 | G A C T G T A C C T G A G A T C A G T C C T A G G C T A G T A C C T G A G C T A G C A T C G A T G C A T G A C T C G T A | AT3G16280(AP2EREBP)/colamp-AT3G16280-DAP-Seq(GSE60143)/Homer | 1e-123 | -2.836e+02 | 0.0000 | 8909.0 | 57.08% | 34180.5 | 46.67% | motif file (matrix) | svg |
| 97 | C G A T T G A C C A T G G A C T A C G T C A T G C G T A G A T C G A C T G C A T G C A T C G A T | WRKY14(WRKY)/colamp-WRKY14-DAP-Seq(GSE60143)/Homer | 1e-123 | -2.833e+02 | 0.0000 | 5780.0 | 37.03% | 20065.3 | 27.40% | motif file (matrix) | svg |
| 98 | C G A T A C G T A C G T A G C T A G C T G A T C G A T C G C T A A G C T A C G T A T C G T A C G | NFATC2(RHD)/Islets-NFATC2-ChIP-Seq(GSE158496)/Homer | 1e-123 | -2.833e+02 | 0.0000 | 10081.0 | 64.59% | 39791.2 | 54.33% | motif file (matrix) | svg |
| 99 | A T G C A G T C A G C T A G C T A C G T A T C G C G T A C G A T T A G C G A C T | LEF1(HMG)/H1-LEF1-ChIP-Seq(GSE64758)/Homer | 1e-122 | -2.823e+02 | 0.0000 | 5551.0 | 35.57% | 19093.1 | 26.07% | motif file (matrix) | svg |
| 100 | T A C G C G T A G A C T T C A G A G C T A G T C A C T G T C A G A G T C C T G A | DDF2(AP2EREBP)/col-DDF2-DAP-Seq(GSE60143)/Homer | 1e-121 | -2.792e+02 | 0.0000 | 2165.0 | 13.87% | 5627.7 | 7.68% | motif file (matrix) | svg |
| 101 | C T G A A T C G G T A C C T G A A G T C A G T C A C T G C G T A A G T C C T G A | TINY(AP2EREBP)/col-TINY-DAP-Seq(GSE60143)/Homer | 1e-119 | -2.744e+02 | 0.0000 | 6431.0 | 41.21% | 23013.4 | 31.42% | motif file (matrix) | svg |
| 102 | A T G C C A G T A G C T A G C T T C A G G T C A T A G C G A C T C G T A C G A T | WRKY20(WRKY)/col-WRKY20-DAP-Seq(GSE60143)/Homer | 1e-118 | -2.739e+02 | 0.0000 | 7768.0 | 49.77% | 29020.0 | 39.62% | motif file (matrix) | svg |
| 103 | C T A G C T A G A T G C G T A C T C A G A T G C A G T C G C A T G A T C G A T C | ZNF91(Zf)/HEK-ZNF91.HA-ChIP-Seq(GSE162571)/Homer | 1e-118 | -2.738e+02 | 0.0000 | 5970.0 | 38.25% | 21004.0 | 28.68% | motif file (matrix) | svg |
| 104 | G C A T C G A T C G T A G A T C C A T G A C G T A C G T A C T G C G T A A G T C A G C T G C A T G C A T C G T A G C T A | WRKY45(WRKY)/col-WRKY45-DAP-Seq(GSE60143)/Homer | 1e-118 | -2.738e+02 | 0.0000 | 4459.0 | 28.57% | 14597.7 | 19.93% | motif file (matrix) | svg |
| 105 | C G A T T C G A G A T C A C G T A C G T T C A G G C A T C T G A C G T A G C T A C G T A A G T C C G T A T G C A C A T G | ANAC050(NAC)/colamp-ANAC050-DAP-Seq(GSE60143)/Homer | 1e-118 | -2.732e+02 | 0.0000 | 6489.0 | 41.58% | 23284.6 | 31.79% | motif file (matrix) | svg |
| 106 | G A C T G C A T C T A G C G A T G A T C T C G A C A T G G A T C | Tgif1(Homeobox)/mES-Tgif1-ChIP-Seq(GSE55404)/Homer | 1e-117 | -2.705e+02 | 0.0000 | 14054.0 | 90.05% | 60765.5 | 82.97% | motif file (matrix) | svg |
| 107 | T G A C T A G C T C A G T C G A T C G A C G T A A G T C C G T A C G T A C G A T C T A G T A C G | Sox7(HMG)/ESC-Sox7-ChIP-Seq(GSE133899)/Homer | 1e-116 | -2.687e+02 | 0.0000 | 3671.0 | 23.52% | 11449.7 | 15.63% | motif file (matrix) | svg |
| 108 | G A C T A G T C C T G A A G T C A G T C A C T G C T G A A G T C G C T A G C A T G T A C C G A T G C A T G A C T C G A T | CBF2(AP2EREBP)/colamp-CBF2-DAP-Seq(GSE60143)/Homer | 1e-116 | -2.681e+02 | 0.0000 | 9846.0 | 63.09% | 38863.7 | 53.06% | motif file (matrix) | svg |
| 109 | A G T C G T A C C T G A A G T C G T A C C T A G G C T A T G A C T G C A G C T A C G T A C G T A | At1g22810(AP2EREBP)/colamp-At1g22810-DAP-Seq(GSE60143)/Homer | 1e-116 | -2.672e+02 | 0.0000 | 8213.0 | 52.62% | 31158.8 | 42.54% | motif file (matrix) | svg |
| 110 | A G T C C T A G A C G T A C G T A C T G C G T A A G T C A G C T G C T A G C A T | WRKY24(WRKY)/colamp-WRKY24-DAP-Seq(GSE60143)/Homer | 1e-115 | -2.657e+02 | 0.0000 | 8721.0 | 55.88% | 33544.0 | 45.80% | motif file (matrix) | svg |
| 111 | T C G A A G T C C G T A A T C G A T G C C G A T A C T G A G T C A G C T A C T G | Tcf12(bHLH)/GM12878-Tcf12-ChIP-Seq(GSE32465)/Homer | 1e-114 | -2.629e+02 | 0.0000 | 5096.0 | 32.65% | 17384.1 | 23.74% | motif file (matrix) | svg |
| 112 | C T A G C T A G T C G A C T A G C G T A A T C G T C G A A C T G C T G A T C G A C T G A T A C G | FRS9(ND)/col-FRS9-DAP-Seq(GSE60143)/Homer | 1e-113 | -2.618e+02 | 0.0000 | 1140.0 | 7.30% | 2280.9 | 3.11% | motif file (matrix) | svg |
| 113 | G C T A G C T A C G T A C G T A C T G A C T A G A C G T A G T C C G T A C T G A G T A C A C T G | WRKY65(WRKY)/colamp-WRKY65-DAP-Seq(GSE60143)/Homer | 1e-113 | -2.606e+02 | 0.0000 | 5929.0 | 37.99% | 20990.7 | 28.66% | motif file (matrix) | svg |
| 114 | G A T C C A G T T A G C A G T C A C T G A G T C A G T C C T A G G A C T G T A C | LEP(AP2EREBP)/col-LEP-DAP-Seq(GSE60143)/Homer | 1e-112 | -2.590e+02 | 0.0000 | 3877.0 | 24.84% | 12375.1 | 16.90% | motif file (matrix) | svg |
| 115 | C G A T G A T C G A T C C T G A G A T C G A T C C A T G T G C A G T A C T C G A G T C A G C A T C G A T C G A T G C A T | At4g32800(AP2EREBP)/colamp-At4g32800-DAP-Seq(GSE60143)/Homer | 1e-112 | -2.588e+02 | 0.0000 | 4005.0 | 25.66% | 12896.9 | 17.61% | motif file (matrix) | svg |
| 116 | G A C T A C T G C G A T A G T C A C T G C T A G A G T C C T G A | AT1G12630(AP2EREBP)/colamp-AT1G12630-DAP-Seq(GSE60143)/Homer | 1e-112 | -2.588e+02 | 0.0000 | 9366.0 | 60.01% | 36692.7 | 50.10% | motif file (matrix) | svg |
| 117 | C A G T T C A G T C G A A G T C C G T A A C T G T G A C C G A T A C T G A C T G A C G T A T C G | Atoh7(bHLH)/Retina-Atoh7-CutnRun(GSE156756)/Homer | 1e-112 | -2.579e+02 | 0.0000 | 4771.0 | 30.57% | 16072.1 | 21.94% | motif file (matrix) | svg |
| 118 | G A C T C T G A A G T C A G T C A C T G C G T A A G T C C T G A | bHLH10(bHLH)/colamp-bHLH10-DAP-Seq(GSE60143)/Homer | 1e-111 | -2.575e+02 | 0.0000 | 6956.0 | 44.57% | 25567.2 | 34.91% | motif file (matrix) | svg |
| 119 | G A T C G C T A G T A C A G T C G C T A T G C A G T A C G A T C C G T A G A C T | MYB83(MYB)/colamp-MYB83-DAP-Seq(GSE60143)/Homer | 1e-111 | -2.569e+02 | 0.0000 | 12131.0 | 77.73% | 50441.0 | 68.87% | motif file (matrix) | svg |
| 120 | G C T A C G T A C G A T C A G T A C T G C G A T G T A C A C T G A T C G G A C T C A T G C T A G G C A T C A G T C A T G | DEAR5(AP2EREBP)/col-DEAR5-DAP-Seq(GSE60143)/Homer | 1e-111 | -2.559e+02 | 0.0000 | 4252.0 | 27.24% | 13938.3 | 19.03% | motif file (matrix) | svg |
| 121 | G A T C C T G A A G T C A G T C A C T G C G T A A G T C C T G A | ERF38(AP2EREBP)/col-ERF38-DAP-Seq(GSE60143)/Homer | 1e-110 | -2.533e+02 | 0.0000 | 9072.0 | 58.13% | 35374.6 | 48.30% | motif file (matrix) | svg |
| 122 | C G A T A G C T T G C A A C T G A G T C T G A C C T A G G T A C A G T C C G T A G C A T G C A T | ERF13(AP2EREBP)/colamp-ERF13-DAP-Seq(GSE60143)/Homer | 1e-109 | -2.516e+02 | 0.0000 | 10639.0 | 68.17% | 42971.3 | 58.67% | motif file (matrix) | svg |
| 123 | A T G C G A T C C G T A A G C T C A G T A T C G G C A T A G C T G A C T A C T G | Sox17(HMG)/Endoderm-Sox17-ChIP-Seq(GSE61475)/Homer | 1e-108 | -2.490e+02 | 0.0000 | 6843.0 | 43.85% | 25177.4 | 34.38% | motif file (matrix) | svg |
| 124 | G A C T T C A G G C A T A G T C G C T A G A T C C T G A A C G T A G T C G T C A | Replumless(BLH)/Arabidopsis-RPL.GFP-ChIP-Seq(GSE78727)/Homer | 1e-107 | -2.482e+02 | 0.0000 | 10935.0 | 70.06% | 44488.7 | 60.74% | motif file (matrix) | svg |
| 125 | A T G C C A T G A C G T A C G T A C T G C G T A A G T C G A C T G C A T C G A T | WRKY71(WRKY)/col-WRKY71-DAP-Seq(GSE60143)/Homer | 1e-107 | -2.474e+02 | 0.0000 | 7644.0 | 48.98% | 28820.1 | 39.35% | motif file (matrix) | svg |
| 126 | C A G T C G T A C G T A G C A T G A C T G C A T G T A C A G C T A C T G G A C T A C G T C A T G | RAV1(RAV)/colamp-RAV1-DAP-Seq(GSE60143)/Homer | 1e-107 | -2.472e+02 | 0.0000 | 5107.0 | 32.72% | 17619.6 | 24.06% | motif file (matrix) | svg |
| 127 | A G T C T G A C C T G A A G T C A G T C A C T G C G T A A G T C G T C A G C T A G C A T C G T A G C A T G C T A C G T A | DEAR3(AP2EREBP)/colamp-DEAR3-DAP-Seq(GSE60143)/Homer | 1e-106 | -2.462e+02 | 0.0000 | 7514.0 | 48.15% | 28244.8 | 38.57% | motif file (matrix) | svg |
| 128 | T C A G T G A C C A T G G C A T C A G T A C T G C G T A T G A C G A C T C G A T C G A T C G T A | WRKY3(WRKY)/col-WRKY3-DAP-Seq(GSE60143)/Homer | 1e-106 | -2.459e+02 | 0.0000 | 6954.0 | 44.56% | 25717.0 | 35.11% | motif file (matrix) | svg |
| 129 | G T C A T C G A T C G A C G T A G C T A C G T A T C G A T G A C A C T G C G T A A G T C C G T A C G T A T C G A G C T A | IDD2(C2H2)/colamp-IDD2-DAP-Seq(GSE60143)/Homer | 1e-105 | -2.422e+02 | 0.0000 | 1424.0 | 9.12% | 3293.8 | 4.50% | motif file (matrix) | svg |
| 130 | G A T C C T G A A G T C G T A C A C T G G C T A G A T C C T G A G C T A G C T A | At4g31060(AP2EREBP)/colamp-At4g31060-DAP-Seq(GSE60143)/Homer | 1e-104 | -2.405e+02 | 0.0000 | 8539.0 | 54.71% | 33055.5 | 45.13% | motif file (matrix) | svg |
| 131 | G T A C T C G A T A G C C G T A C G T A C T G A T G C A T G A C A C T G C G T A A G T C C T G A C T G A T C G A C G T A | NUC(C2H2)/col-NUC-DAP-Seq(GSE60143)/Homer | 1e-103 | -2.384e+02 | 0.0000 | 1228.0 | 7.87% | 2671.3 | 3.65% | motif file (matrix) | svg |
| 132 | G A T C A T G C A G T C C G T A A G T C A G T C A C T G G C T A A G T C C G T A | AT1G44830(AP2EREBP)/col-AT1G44830-DAP-Seq(GSE60143)/Homer | 1e-103 | -2.373e+02 | 0.0000 | 5043.0 | 32.31% | 17469.8 | 23.85% | motif file (matrix) | svg |
| 133 | C G A T C G T A A G T C A C G T A C G T T C G A T G C A G C A T G C T A C G T A A C G T A G C T C G T A C G T A A C T G | ANAC062(NAC)/colamp-ANAC062-DAP-Seq(GSE60143)/Homer | 1e-102 | -2.366e+02 | 0.0000 | 3587.0 | 22.98% | 11438.4 | 15.62% | motif file (matrix) | svg |
| 134 | A T C G T C G A G A C T A T C G T G A C A C G T C T A G A C T G C G T A A C T G A G T C G T A C | ZNF415(Zf)/HEK293-ZNF415.GFP-ChIP-Seq(GSE58341)/Homer | 1e-102 | -2.360e+02 | 0.0000 | 4763.0 | 30.52% | 16298.2 | 22.25% | motif file (matrix) | svg |
| 135 | T A C G C T G A C A T G G A T C G T A C G C A T T C A G T A C G A G C T G T C A G A T C G C A T T A C G C G T A C T A G G A T C G A T C C G A T A C T G T C A G | ZNF322(Zf)/HEK293-ZNF322.GFP-ChIP-Seq(GSE58341)/Homer | 1e-102 | -2.354e+02 | 0.0000 | 1427.0 | 9.14% | 3342.4 | 4.56% | motif file (matrix) | svg |
| 136 | A T G C G T A C A G T C A G T C A C G T A C G T C G A T A C G T | AT5G02460(C2C2dof)/col-AT5G02460-DAP-Seq(GSE60143)/Homer | 1e-101 | -2.334e+02 | 0.0000 | 11203.0 | 71.78% | 46043.1 | 62.87% | motif file (matrix) | svg |
| 137 | T A G C C A T G G A C T G A C T T C A G G T C A G A T C G A C T G C A T G C T A | WRKY15(WRKY)/col-WRKY15-DAP-Seq(GSE60143)/Homer | 1e-100 | -2.319e+02 | 0.0000 | 9303.0 | 59.61% | 36788.8 | 50.23% | motif file (matrix) | svg |
| 138 | G C A T A C T G C T A G A G T C A C T G A C T G A G T C A C G T | ERF105(AP2EREBP)/colamp-ERF105-DAP-Seq(GSE60143)/Homer | 1e-100 | -2.307e+02 | 0.0000 | 11029.0 | 70.67% | 45212.6 | 61.73% | motif file (matrix) | svg |
| 139 | C G A T C T A G A C G T A C G T A C G T C G T A A G C T C G A T A G C T C G T A C T A G T A G C | FoxD3(forkhead)/ZebrafishEmbryo-Foxd3.biotin-ChIP-seq(GSE106676)/Homer | 1e-99 | -2.291e+02 | 0.0000 | 5123.0 | 32.83% | 17914.1 | 24.46% | motif file (matrix) | svg |
| 140 | A T G C G T A C C T G A A G T C A G T C A C T G G T C A A G T C G T C A G C A T G C A T G A C T | At5g65130(AP2EREBP)/colamp-At5g65130-DAP-Seq(GSE60143)/Homer | 1e-98 | -2.279e+02 | 0.0000 | 4185.0 | 26.81% | 13978.4 | 19.09% | motif file (matrix) | svg |
| 141 | C A G T A C T G T C A G T G C A G C T A A T G C T C G A A T C G G T C A T G C A | ZNF189(Zf)/HEK293-ZNF189.GFP-ChIP-Seq(GSE58341)/Homer | 1e-98 | -2.272e+02 | 0.0000 | 5656.0 | 36.24% | 20238.5 | 27.63% | motif file (matrix) | svg |
| 142 | G A C T C T A G A T G C A G T C G T C A T A C G A T G C A T C G | HIC1(Zf)/Treg-ZBTB29-ChIP-Seq(GSE99889)/Homer | 1e-97 | -2.236e+02 | 0.0000 | 11874.0 | 76.08% | 49586.5 | 67.70% | motif file (matrix) | svg |
| 143 | G T A C A C G T A C G T T C A G G A C T G C A T T C A G C G T A C T G A A G T C C G T A G T C A A C T G A C G T G C T A | NTM2(NAC)/col-NTM2-DAP-Seq(GSE60143)/Homer | 1e-96 | -2.227e+02 | 0.0000 | 5812.0 | 37.24% | 20979.3 | 28.64% | motif file (matrix) | svg |
| 144 | C A T G T G A C C A T G G A C T C A G T C T A G G C T A G T A C G A C T G C A T G C A T C G A T | WRKY21(WRKY)/colamp-WRKY21-DAP-Seq(GSE60143)/Homer | 1e-96 | -2.216e+02 | 0.0000 | 1553.0 | 9.95% | 3857.9 | 5.27% | motif file (matrix) | svg |
| 145 | A T C G T G C A G A T C C T A G A C G T A T C G C G T A A G T C T C A G A C T G T C A G G C T A | Knotted(Homeobox)/Corn-KN1-ChIP-Seq(GSE39161)/Homer | 1e-95 | -2.202e+02 | 0.0000 | 12552.0 | 80.43% | 53147.3 | 72.57% | motif file (matrix) | svg |
| 146 | T C A G A T C G G A C T A C T G G A C T C A G T C T A G C G T A G T A C C G T A C T A G A T C G | Tbx20(T-box)/Heart-Tbx20-ChIP-Seq(GSE29636)/Homer | 1e-95 | -2.202e+02 | 0.0000 | 2469.0 | 15.82% | 7212.2 | 9.85% | motif file (matrix) | svg |
| 147 | G C T A T C G A C G T A C T G A A C T G A C G T A G T C C G T A C G T A A G T C C T A G T G C A | WRKY42(WRKY)/colamp-WRKY42-DAP-Seq(GSE60143)/Homer | 1e-95 | -2.191e+02 | 0.0000 | 5518.0 | 35.36% | 19747.9 | 26.96% | motif file (matrix) | svg |
| 148 | G C A T G C A T G C A T A T G C A G C T T C G A T A C G G C T A C G T A C A T G G T A C G C A T G C A T A G T C A G C T | HSFA6B(HSF)/colamp-HSFA6B-DAP-Seq(GSE60143)/Homer | 1e-94 | -2.172e+02 | 0.0000 | 4152.0 | 26.60% | 13967.0 | 19.07% | motif file (matrix) | svg |
| 149 | G A C T A C T G C A G T A G T C A C T G A C T G A G C T A C T G C T A G G T C A | At1g77640(AP2EREBP)/col-At1g77640-DAP-Seq(GSE60143)/Homer | 1e-93 | -2.159e+02 | 0.0000 | 3405.0 | 21.82% | 10926.3 | 14.92% | motif file (matrix) | svg |
| 150 | A G T C G T A C C T G A A G T C A G T C C A T G G C T A A G T C T G C A G C T A G C A T G C A T | RAP21(AP2EREBP)/colamp-RAP21-DAP-Seq(GSE60143)/Homer | 1e-93 | -2.143e+02 | 0.0000 | 4955.0 | 31.75% | 17388.6 | 23.74% | motif file (matrix) | svg |
| 151 | G C A T C G A T G A C T T G C A A C T G A G T C T G A C A C T G G A T C A G T C C G T A G A C T | ERF15(AP2EREBP)/colamp-ERF15-DAP-Seq(GSE60143)/Homer | 1e-89 | -2.069e+02 | 0.0000 | 12241.0 | 78.43% | 51722.0 | 70.62% | motif file (matrix) | svg |
| 152 | A T G C G C A T T A G C C G A T T A G C G C A T T A G C G C A T A T G C G A C T | GAGA-repeat/Arabidopsis-Promoters/Homer | 1e-89 | -2.060e+02 | 0.0000 | 5689.0 | 36.45% | 20674.6 | 28.23% | motif file (matrix) | svg |
| 153 | T A C G T A G C G C T A C G A T C T A G A C G T C A G T C A G T G C T A A G T C G T C A G C A T | FOXK2(Forkhead)/U2OS-FOXK2-ChIP-Seq(E-MTAB-2204)/Homer | 1e-89 | -2.060e+02 | 0.0000 | 5758.0 | 36.89% | 20976.6 | 28.64% | motif file (matrix) | svg |
| 154 | C T A G A C T G T G C A G T C A A T G C C G T A A T C G A T G C A G T C C T A G | ZNF341(Zf)/EBV-ZNF341-ChIP-Seq(GSE113194)/Homer | 1e-89 | -2.057e+02 | 0.0000 | 5420.0 | 34.73% | 19506.5 | 26.63% | motif file (matrix) | svg |
| 155 | A G T C A C G T A C G T T C A G G C T A G C T A A T G C C G T A C G A T A G T C C G T A G T C A A C T G G A T C G C A T | SND3(NAC)/col-SND3-DAP-Seq(GSE60143)/Homer | 1e-88 | -2.045e+02 | 0.0000 | 7663.0 | 49.10% | 29546.6 | 40.34% | motif file (matrix) | svg |
| 156 | C G A T T C G A G T A C A C G T A C G T T C A G G C A T G C A T G C T A C G T A C G T A A G T C C G T A T G C A C A T G | ANAC020(NAC)/col-ANAC020-DAP-Seq(GSE60143)/Homer | 1e-88 | -2.037e+02 | 0.0000 | 7086.0 | 45.40% | 26928.6 | 36.77% | motif file (matrix) | svg |
| 157 | G T A C A C T G A T G C A G T C C T A G G A T C G T A C C T G A G A C T G C A T C G A T G A C T | RAP212(AP2EREBP)/col-RAP212-DAP-Seq(GSE60143)/Homer | 1e-88 | -2.031e+02 | 0.0000 | 9587.0 | 61.43% | 38596.1 | 52.70% | motif file (matrix) | svg |
| 158 | G T A C G C T A T C A G C T G A C T A G C A T G A G C T G A T C T G C A T C G A C T G A A C T G C A G T A G T C G A T C G C T A | HNF4a(NR),DR1/HepG2-HNF4a-ChIP-Seq(GSE25021)/Homer | 1e-88 | -2.028e+02 | 0.0000 | 2954.0 | 18.93% | 9263.8 | 12.65% | motif file (matrix) | svg |
| 159 | T C A G A C G T T C G A T A G C A G T C C G T A A C T G G T A C A C G T A C T G A T C G A G T C | Atoh1(bHLH)/Cerebellum-Atoh1-ChIP-Seq(GSE22111)/Homer | 1e-87 | -2.025e+02 | 0.0000 | 6744.0 | 43.21% | 25403.8 | 34.69% | motif file (matrix) | svg |
| 160 | C G T A C G A T C T A G C G T A A G C T C A G T T A C G C G T A A C G T C A T G C T A G A T G C | HOXA3(Homeobox)/mEmbryo-Hoxa3-ChIP-Seq(E-MTAB-8607)/Homer | 1e-87 | -2.022e+02 | 0.0000 | 2187.0 | 14.01% | 6303.5 | 8.61% | motif file (matrix) | svg |
| 161 | G C A T C G T A C T A G A G T C G T C A C G T A A T G C A C G T A C G T A C T G G A T C G C A T C G T A G C T A G C T A | bHLH122(bHLH)/col100-bHLH122-DAP-Seq(GSE60143)/Homer | 1e-87 | -2.016e+02 | 0.0000 | 7604.0 | 48.72% | 29321.3 | 40.03% | motif file (matrix) | svg |
| 162 | T C G A A G T C A C G T A C G T T C A G C A G T C T G A C T A G T C G A C G T A A T C G C G T A C G T A A C T G A G C T | NTM1(NAC)/col-NTM1-DAP-Seq(GSE60143)/Homer | 1e-87 | -2.009e+02 | 0.0000 | 4231.0 | 27.11% | 14497.9 | 19.80% | motif file (matrix) | svg |
| 163 | T C A G A G C T G T C A C G T A A C G T A T G C C G T A A C G T A C G T C T G A | PHV(HB)/col-PHV-DAP-Seq(GSE60143)/Homer | 1e-87 | -2.005e+02 | 0.0000 | 4461.0 | 28.58% | 15469.2 | 21.12% | motif file (matrix) | svg |
| 164 | C T A G G C T A A G T C A C T G A C G T G A C T G A C T A T G C T C G A C A G T G A T C C G A T G A C T G A T C G A T C | RKD2(RWPRK)/colamp-RKD2-DAP-Seq(GSE60143)/Homer | 1e-86 | -1.987e+02 | 0.0000 | 6752.0 | 43.26% | 25499.0 | 34.82% | motif file (matrix) | svg |
| 165 | G T A C C A T G A G C T A G C T T C A G T G C A T G A C A G C T C G T A C G T A | WRKY33(WRKY)/col-WRKY33-DAP-Seq(GSE60143)/Homer | 1e-85 | -1.975e+02 | 0.0000 | 8510.0 | 54.53% | 33584.9 | 45.86% | motif file (matrix) | svg |
| 166 | C G A T C T A G A C T G A G C T C T G A A C T G A C G T A C G T C T A G C T A G | MYB96(MYB)/colamp-MYB96-DAP-Seq(GSE60143)/Homer | 1e-85 | -1.972e+02 | 0.0000 | 8232.0 | 52.75% | 32293.4 | 44.09% | motif file (matrix) | svg |
| 167 | C T G A A T G C G C T A C G A T A T G C C G T A C G T A C G T A C T A G T A C G | Tcf3(HMG)/mES-Tcf3-ChIP-Seq(GSE11724)/Homer | 1e-85 | -1.960e+02 | 0.0000 | 2531.0 | 16.22% | 7674.5 | 10.48% | motif file (matrix) | svg |
| 168 | C G A T T C G A A T G C C G A T G C A T T C G A A G C T G C A T G C A T C G A T T C G A A G C T T C G A G T C A C T A G | ANAC004(NAC)/colamp-ANAC004-DAP-Seq(GSE60143)/Homer | 1e-84 | -1.957e+02 | 0.0000 | 3237.0 | 20.74% | 10471.4 | 14.30% | motif file (matrix) | svg |
| 169 | A C T G C G T A A C T G A T G C T G A C G A T C A T C G T G C A A C T G A G T C | ZNF519(Zf)/HEK293-ZNF519.GFP-ChIP-Seq(GSE58341)/Homer | 1e-83 | -1.933e+02 | 0.0000 | 1776.0 | 11.38% | 4852.6 | 6.63% | motif file (matrix) | svg |
| 170 | C G A T C G T A G C T A G C A T G C A T C T G A A C T G A C G T A G T C C G T A C G T A G T A C T C A G G C T A C G A T | WRKY25(WRKY)/colamp-WRKY25-DAP-Seq(GSE60143)/Homer | 1e-82 | -1.910e+02 | 0.0000 | 10082.0 | 64.60% | 41182.5 | 56.23% | motif file (matrix) | svg |
| 171 | T C A G T C A G G C T A C G T A T A C G G A C T T C A G T C G A C T G A C G T A T A C G G A C T | IRF8(IRF)/BMDM-IRF8-ChIP-Seq(GSE77884)/Homer | 1e-82 | -1.907e+02 | 0.0000 | 2253.0 | 14.44% | 6655.3 | 9.09% | motif file (matrix) | svg |
| 172 | C G T A G A T C C T A G A C G T G T A C C T G A A G C T G A T C G C T A G A C T | TGA2(bZIP)/colamp-TGA2-DAP-Seq(GSE60143)/Homer | 1e-82 | -1.907e+02 | 0.0000 | 9940.0 | 63.69% | 40500.0 | 55.30% | motif file (matrix) | svg |
| 173 | A G T C G A C T G A T C C G T A G T A C A G T C G C T A C G T A G T A C A G T C G T A C G T A C | MYB63(MYB)/col-MYB63-DAP-Seq(GSE60143)/Homer | 1e-82 | -1.904e+02 | 0.0000 | 5462.0 | 35.00% | 19908.9 | 27.18% | motif file (matrix) | svg |
| 174 | C G T A G A C T C A T G C T A G A G T C A C T G C T A G A G T C C A T G C T A G | ERF7(AP2EREBP)/col-ERF7-DAP-Seq(GSE60143)/Homer | 1e-82 | -1.903e+02 | 0.0000 | 12339.0 | 79.06% | 52478.6 | 71.65% | motif file (matrix) | svg |
| 175 | C A T G G A T C C T G A G T A C C T A G C T G A G C T A G C A T G A T C G A T C A G T C C T A G C G T A C A T G C T A G | AIL7(AP2EREBP)/colamp-AIL7-DAP-Seq(GSE60143)/Homer | 1e-82 | -1.899e+02 | 0.0000 | 6571.0 | 42.10% | 24822.9 | 33.89% | motif file (matrix) | svg |
| 176 | C T A G C T G A C T A G C T G A C T A G C T G A C T A G C T G A C T A G C T G A | SeqBias: GA-repeat | 1e-82 | -1.889e+02 | 0.0000 | 14977.0 | 95.96% | 67232.5 | 91.80% | motif file (matrix) | svg |
| 177 | A G C T C T A G G A T C A G T C C T A G C T G A A G T C G C T A G C A T T G C A | CBF1(AP2EREBP)/colamp-CBF1-DAP-Seq(GSE60143)/Homer | 1e-82 | -1.888e+02 | 0.0000 | 11083.0 | 71.01% | 46135.9 | 62.99% | motif file (matrix) | svg |
| 178 | G C A T C G A T G C A T C G T A C T A G A G T C G T C A C G T A A T C G A C G T A C G T A C T G G T A C G C A T C G A T | bHLH80(bHLH)/col-bHLH80-DAP-Seq(GSE60143)/Homer | 1e-80 | -1.844e+02 | 0.0000 | 7968.0 | 51.05% | 31278.6 | 42.71% | motif file (matrix) | svg |
| 179 | T C A G T G A C G T A C C G T A A C G T T G A C A C G T T C A G A G C T G A C T | NeuroD1(bHLH)/Islet-NeuroD1-ChIP-Seq(GSE30298)/Homer | 1e-79 | -1.838e+02 | 0.0000 | 5177.0 | 33.17% | 18764.5 | 25.62% | motif file (matrix) | svg |
| 180 | C G A T C G T A A G T C A C G T A C G T T C A G G C A T C G T A G C T A G C T A C G T A A G T C C G T A G T C A A C T G | ANAC058(NAC)/col-ANAC058-DAP-Seq(GSE60143)/Homer | 1e-79 | -1.838e+02 | 0.0000 | 6506.0 | 41.69% | 24627.1 | 33.63% | motif file (matrix) | svg |
| 181 | A G T C A C G T A C G T T A C G G C T A G C T A C G T A C G A T C G A T A T G C C G T A G T C A A C T G G A C T G C A T | SND2(NAC)/colamp-SND2-DAP-Seq(GSE60143)/Homer | 1e-78 | -1.796e+02 | 0.0000 | 6727.0 | 43.10% | 25687.9 | 35.07% | motif file (matrix) | svg |
| 182 | C T G A C T A G A C T G G C A T A T G C C G T A C T G A C T A G A C T G A C G T A G T C C T G A | RARg(NR)/ES-RARg-ChIP-Seq(GSE30538)/Homer | 1e-77 | -1.789e+02 | 0.0000 | 427.0 | 2.74% | 567.7 | 0.78% | motif file (matrix) | svg |
| 183 | C G T A C G A T C G T A C G A T C A T G A C T G C G A T A G T C A T C G T C A G G A C T A C T G | At1g36060(AP2EREBP)/colamp-At1g36060-DAP-Seq(GSE60143)/Homer | 1e-76 | -1.772e+02 | 0.0000 | 9995.0 | 64.04% | 40994.2 | 55.97% | motif file (matrix) | svg |
| 184 | A C G T T G C A A G C T G A T C C T A G C T G A A G C T G T C A T C G A C G T A | CUX1(Homeobox)/K562-CUX1-ChIP-Seq(GSE92882)/Homer | 1e-76 | -1.770e+02 | 0.0000 | 10269.0 | 65.80% | 42329.6 | 57.80% | motif file (matrix) | svg |
| 185 | G C T A G C T A C T G A A C T G A C G T A G T C C G T A C G T A G T A C A C T G A T G C G C A T | WRKY47(WRKY)/colamp-WRKY47-DAP-Seq(GSE60143)/Homer | 1e-76 | -1.768e+02 | 0.0000 | 4136.0 | 26.50% | 14412.5 | 19.68% | motif file (matrix) | svg |
| 186 | G C T A C G T A C G T A G A C T C A T G C T A G G A T C A C T G T C A G G A T C A C T G T A C G | ERF9(AP2EREBP)/colamp-ERF9-DAP-Seq(GSE60143)/Homer | 1e-76 | -1.764e+02 | 0.0000 | 3967.0 | 25.42% | 13708.6 | 18.72% | motif file (matrix) | svg |
| 187 | G C A T T G A C C T G A A G T C A G T C A C T G G T C A A G T C G C T A G A C T G C T A C T G A | DREB2(AP2EREBP)/col-DREB2-DAP-Seq(GSE60143)/Homer | 1e-76 | -1.757e+02 | 0.0000 | 8151.0 | 52.23% | 32275.2 | 44.07% | motif file (matrix) | svg |
| 188 | A G C T C A T G G C A T G A T C T G C A C T A G G A T C A C G T | Tgif2(Homeobox)/mES-Tgif2-ChIP-Seq(GSE55404)/Homer | 1e-75 | -1.733e+02 | 0.0000 | 14167.0 | 90.77% | 62565.4 | 85.43% | motif file (matrix) | svg |
| 189 | G A T C G A C T G A C T A C G T A G T C A C G T A G T C A C G T A G T C A C G T A G T C A C G T G T A C C G A T G T C A | BPC6(BBRBPC)/col-BPC6-DAP-Seq(GSE60143)/Homer | 1e-74 | -1.710e+02 | 0.0000 | 230.0 | 1.47% | 160.9 | 0.22% | motif file (matrix) | svg |
| 190 | C G A T T G C A T G C A G A T C C G T A A C T G T G A C G A C T C A T G A C T G | Tcf21(bHLH)/ArterySmoothMuscle-Tcf21-ChIP-Seq(GSE61369)/Homer | 1e-73 | -1.696e+02 | 0.0000 | 5186.0 | 33.23% | 19015.6 | 25.96% | motif file (matrix) | svg |
| 191 | A C T G C T A G A G T C A C T G A C T G A G T C A C G T C T A G | AT5G23930(mTERF)/col-AT5G23930-DAP-Seq(GSE60143)/Homer | 1e-73 | -1.686e+02 | 0.0000 | 10454.0 | 66.98% | 43377.6 | 59.23% | motif file (matrix) | svg |
| 192 | T A G C C A T G A G C T A C G T A C T G C G T A A G T C G A C T G C A T C T G A | AT3G42860(zfGRF)/col-AT3G42860-DAP-Seq(GSE60143)/Homer | 1e-73 | -1.683e+02 | 0.0000 | 4141.0 | 26.53% | 14549.9 | 19.87% | motif file (matrix) | svg |
| 193 | G C T A C T G A T C G A A G T C A G T C C T G A A G T C G T C A C T G A T G C A | RUNX1(Runt)/Jurkat-RUNX1-ChIP-Seq(GSE29180)/Homer | 1e-72 | -1.672e+02 | 0.0000 | 7529.0 | 48.24% | 29546.6 | 40.34% | motif file (matrix) | svg |
| 194 | C G A T T C G A A G T C A C G T A C G T T A C G C G T A G C T A G C T A C G A T G C A T A T G C C G T A G T C A A C T G | ANAC071(NAC)/col-ANAC071-DAP-Seq(GSE60143)/Homer | 1e-71 | -1.644e+02 | 0.0000 | 8999.0 | 57.66% | 36454.3 | 49.77% | motif file (matrix) | svg |
| 195 | G C A T A C G T A C T G A C G T A G T C A C T G A T C G G T C A C G A T C G T A | ARF2(ARF)/col-ARF2-DAP-Seq(GSE60143)/Homer | 1e-71 | -1.642e+02 | 0.0000 | 14219.0 | 91.11% | 62979.5 | 85.99% | motif file (matrix) | svg |
| 196 | T G A C C A T G A C G T A C G T A C T G C G T A A G T C A G C T G C A T T C G A | WRKY30(WRKY)/colamp-WRKY30-DAP-Seq(GSE60143)/Homer | 1e-71 | -1.636e+02 | 0.0000 | 4499.0 | 28.83% | 16139.8 | 22.04% | motif file (matrix) | svg |
| 197 | A T G C G T A C A C T G A G T C A G T C A C T G A G T C G T A C | ERF73(AP2EREBP)/col-ERF73-DAP-Seq(GSE60143)/Homer | 1e-70 | -1.629e+02 | 0.0000 | 6202.0 | 39.74% | 23606.4 | 32.23% | motif file (matrix) | svg |
| 198 | C G A T C T A G C T G A A T G C C T G A T C G A C G T A C T G A T C G A T A G C A G T C C G T A A C T G T C G A A T G C | Hand2(bHLH)/Mesoderm-Hand2-ChIP-Seq(GSE61475)/Homer | 1e-70 | -1.623e+02 | 0.0000 | 2882.0 | 18.47% | 9421.5 | 12.86% | motif file (matrix) | svg |
| 199 | T G A C C G A T C T G A C T A G C T A G A C G T A T G C T G C A T C G A C T G A C T A G C A T G A C G T A G T C C G T A | PPARa(NR),DR1/Liver-Ppara-ChIP-Seq(GSE47954)/Homer | 1e-70 | -1.623e+02 | 0.0000 | 5897.0 | 37.78% | 22258.4 | 30.39% | motif file (matrix) | svg |
| 200 | C T A G A G T C T A C G T A C G T G A C C G T A A C T G T A G C G C A T C A T G A T G C A G C T | Ascl1(bHLH)/NeuralTubes-Ascl1-ChIP-Seq(GSE55840)/Homer | 1e-69 | -1.600e+02 | 0.0000 | 8168.0 | 52.34% | 32631.7 | 44.55% | motif file (matrix) | svg |
| 201 | A G C T T G A C C G A T C G A T C T A G A C G T C A G T C A G T G C T A A G T C | FOXK1(Forkhead)/HEK293-FOXK1-ChIP-Seq(GSE51673)/Homer | 1e-69 | -1.598e+02 | 0.0000 | 8005.0 | 51.29% | 31877.3 | 43.52% | motif file (matrix) | svg |
| 202 | C G T A G C A T C G T A C G T A G C A T A C T G C G A T A G T C A C T G A C T G G A C T C T A G | AT1G71450(AP2EREBP)/col-AT1G71450-DAP-Seq(GSE60143)/Homer | 1e-69 | -1.596e+02 | 0.0000 | 13926.0 | 89.23% | 61399.5 | 83.83% | motif file (matrix) | svg |
| 203 | C G A T A G T C C A T G G A C T A C G T C T A G C G T A G A T C G A C T C G A T G C A T G A C T | WRKY43(WRKY)/colamp-WRKY43-DAP-Seq(GSE60143)/Homer | 1e-69 | -1.589e+02 | 0.0000 | 4136.0 | 26.50% | 14660.4 | 20.02% | motif file (matrix) | svg |
| 204 | C T G A T A C G G C A T A G C T A G C T A G T C T C G A A C T G C A G T A G C T A G C T G A T C | IRF3(IRF)/BMDM-Irf3-ChIP-Seq(GSE67343)/Homer | 1e-68 | -1.586e+02 | 0.0000 | 1794.0 | 11.49% | 5209.8 | 7.11% | motif file (matrix) | svg |
| 205 | A C T G C T A G A G T C A C T G A C T G A T G C A C T G T A C G | ESE1(AP2EREBP)/col-ESE1-DAP-Seq(GSE60143)/Homer | 1e-68 | -1.586e+02 | 0.0000 | 6929.0 | 44.40% | 26956.3 | 36.81% | motif file (matrix) | svg |
| 206 | C G A T C G A T G C A T G A C T A C G T C G T A C G T A A C T G T A G C C G T A C G T A C G T A | AT5G60130(ABI3VP1)/col-AT5G60130-DAP-Seq(GSE60143)/Homer | 1e-68 | -1.577e+02 | 0.0000 | 8334.0 | 53.40% | 33446.4 | 45.67% | motif file (matrix) | svg |
| 207 | C G T A C G T A C G A T A C T G C A G T A G T C A C T G A C T G A G C T A C T G | DREB19(AP2EREBP)/colamp-DREB19-DAP-Seq(GSE60143)/Homer | 1e-67 | -1.563e+02 | 0.0000 | 9039.0 | 57.92% | 36791.9 | 50.24% | motif file (matrix) | svg |
| 208 | C T A G T C G A C G A T C T A G G C A T C A G T C T A G G A T C C G T A G T C A | CEBP:AP1(bZIP)/ThioMac-CEBPb-ChIP-Seq(GSE21512)/Homer | 1e-67 | -1.561e+02 | 0.0000 | 7004.0 | 44.88% | 27340.1 | 37.33% | motif file (matrix) | svg |
| 209 | C A T G A G T C G T C A C G T A A T G C A C G T A C G T A C T G | bHLH130(bHLH)/col-bHLH130-DAP-Seq(GSE60143)/Homer | 1e-67 | -1.544e+02 | 0.0000 | 6694.0 | 42.89% | 25965.7 | 35.45% | motif file (matrix) | svg |
| 210 | G A C T C G A T T C A G G A T C G A C T A G C T A G C T A G T C G A T C C G T A C T A G C T A G T C G A T C G A C T G A | Bcl6(Zf)/Liver-Bcl6-ChIP-Seq(GSE31578)/Homer | 1e-66 | -1.542e+02 | 0.0000 | 6464.0 | 41.42% | 24929.6 | 34.04% | motif file (matrix) | svg |
| 211 | C G T A C T A G T C A G T C A G A G T C A T G C A G T C G C A T A G C T A C G T A T C G C G A T | Sox9(HMG)/Limb-SOX9-ChIP-Seq(GSE73225)/Homer | 1e-66 | -1.538e+02 | 0.0000 | 6255.0 | 40.08% | 23998.3 | 32.77% | motif file (matrix) | svg |
| 212 | G A T C G T A C C T G A A G T C A G T C A C T G G C T A G T A C G T C A G C A T G C A T C G A T | DEAR2(AP2EREBP)/colamp-DEAR2-DAP-Seq(GSE60143)/Homer | 1e-65 | -1.512e+02 | 0.0000 | 12613.0 | 80.82% | 54523.9 | 74.45% | motif file (matrix) | svg |
| 213 | G T C A T G C A G C T A A G T C C G T A A C T G T G A C G C A T T C A G C A G T | Ap4(bHLH)/AML-Tfap4-ChIP-Seq(GSE45738)/Homer | 1e-65 | -1.508e+02 | 0.0000 | 6126.0 | 39.25% | 23472.6 | 32.05% | motif file (matrix) | svg |
| 214 | C T G A T G A C T G A C C G T A A C G T T G A C A G C T C T A G A C G T G A C T | Olig2(bHLH)/Neuron-Olig2-ChIP-Seq(GSE30882)/Homer | 1e-65 | -1.507e+02 | 0.0000 | 10795.0 | 69.17% | 45371.6 | 61.95% | motif file (matrix) | svg |
| 215 | G T C A T C G A C T A G C T A G A G T C G T C A C G A T C T A G G A C T G A T C G A T C T C A G C T A G C T G A A G T C G C T A C A G T T C A G G A T C G A T C | p63(p53)/Keratinocyte-p63-ChIP-Seq(GSE17611)/Homer | 1e-65 | -1.499e+02 | 0.0000 | 3525.0 | 22.59% | 12217.5 | 16.68% | motif file (matrix) | svg |
| 216 | T C A G T A C G T A G C A C G T A C T G C G A T A G T C C G T A T A C G A G T C | Meis1(Homeobox)/MastCells-Meis1-ChIP-Seq(GSE48085)/Homer | 1e-64 | -1.492e+02 | 0.0000 | 11044.0 | 70.76% | 46627.5 | 63.66% | motif file (matrix) | svg |
| 217 | C T G A C T G A C T G A A T G C G A T C C A T G A C T G G A C T G A C T G C A T C G T A C G T A A G T C G T A C C T G A A T C G G C A T G A C T G A C T A G C T | GRHL2(CP2)/HBE-GRHL2-ChIP-Seq(GSE46194)/Homer | 1e-64 | -1.490e+02 | 0.0000 | 3655.0 | 23.42% | 12773.8 | 17.44% | motif file (matrix) | svg |
| 218 | A G C T G A T C G A T C C G T A G T A C A G T C C G A T C T G A G T A C G A T C C G T A G A C T | ATY19(MYB)/col-ATY19-DAP-Seq(GSE60143)/Homer | 1e-64 | -1.488e+02 | 0.0000 | 7312.0 | 46.85% | 28879.9 | 39.43% | motif file (matrix) | svg |
| 219 | C G T A A C G T A C G T A C G T A C G T A G T C A G T C C T G A A G C T A G C T | NFAT(RHD)/Jurkat-NFATC1-ChIP-Seq(Jolma\_et\_al.)/Homer | 1e-64 | -1.488e+02 | 0.0000 | 6141.0 | 39.35% | 23574.7 | 32.19% | motif file (matrix) | svg |
| 220 | A G T C C G T A A C G T A G T C A C G T A C T G | Tal1 | 1e-63 | -1.467e+02 | 0.0000 | 8871.0 | 56.84% | 36176.0 | 49.39% | motif file (matrix) | svg |
| 221 | C A T G A T G C T A G C C T G A A G T C A G T C A C T G G C T A A G T C G T A C G C T A G C A T | At4g28140(AP2EREBP)/colamp-At4g28140-DAP-Seq(GSE60143)/Homer | 1e-63 | -1.458e+02 | 0.0000 | 5761.0 | 36.91% | 21932.9 | 29.95% | motif file (matrix) | svg |
| 222 | A G T C A G T C C T G A A G T C A G T C A C T G C G T A A G T C T C G A G A T C C G A T C G T A | AT1G01250(AP2EREBP)/col-AT1G01250-DAP-Seq(GSE60143)/Homer | 1e-62 | -1.430e+02 | 0.0000 | 2321.0 | 14.87% | 7402.3 | 10.11% | motif file (matrix) | svg |
| 223 | T C G A T G A C G T A C C G T A C A G T T G A C A C G T A C T G A G C T A G C T | NeuroG2(bHLH)/Fibroblast-NeuroG2-ChIP-Seq(GSE75910)/Homer | 1e-61 | -1.417e+02 | 0.0000 | 8854.0 | 56.73% | 36192.3 | 49.42% | motif file (matrix) | svg |
| 224 | A G T C A C G T A C G T T C A G G C T A C G T A G C T A G C A T C G A T A G T C C G T A G T C A A C T G G A C T G C T A | SMB(NAC)/colamp-SMB-DAP-Seq(GSE60143)/Homer | 1e-61 | -1.413e+02 | 0.0000 | 9154.0 | 58.65% | 37621.2 | 51.37% | motif file (matrix) | svg |
| 225 | A T G C A G T C A C T G A T G C A G T C A C T G A G T C G T A C | SHN3(AP2EREBP)/col-SHN3-DAP-Seq(GSE60143)/Homer | 1e-61 | -1.412e+02 | 0.0000 | 2784.0 | 17.84% | 9281.3 | 12.67% | motif file (matrix) | svg |
| 226 | C G T A A C T G C G T A A C G T A T C G C A G T T A G C C G T A T C G A G T A C C T G A T A G C C G T A A C T G C G T A A C G T C G T A C T G A A T C G G C T A | GATA3(Zf),DR8/iTreg-Gata3-ChIP-Seq(GSE20898)/Homer | 1e-60 | -1.388e+02 | 0.0000 | 962.0 | 6.16% | 2362.4 | 3.23% | motif file (matrix) | svg |
| 227 | T C G A C G T A A G T C C G T A C T A G A G T C C G A T A C T G G A C T A G C T A C T G G A C T | HLH-1(bHLH)/cElegans-Embryo-HLH1-ChIP-Seq(modEncode)/Homer | 1e-59 | -1.380e+02 | 0.0000 | 5708.0 | 36.57% | 21832.6 | 29.81% | motif file (matrix) | svg |
| 228 | C G A T C G T A A G T C A C G T A C G T T C A G C G T A C G T A G C A T G C A T G C A T A G T C C G T A G T C A A C T G | VND2(NAC)/col-VND2-DAP-Seq(GSE60143)/Homer | 1e-58 | -1.349e+02 | 0.0000 | 9155.0 | 58.66% | 37752.7 | 51.55% | motif file (matrix) | svg |
| 229 | A C T G A C T G A G T C A C T G A C T G A G T C A C G T C T A G | ERF1(AP2EREBP)/colamp-ERF1-DAP-Seq(GSE60143)/Homer | 1e-58 | -1.342e+02 | 0.0000 | 5615.0 | 35.98% | 21487.0 | 29.34% | motif file (matrix) | svg |
| 230 | T C G A T G A C G C A T A G C T C A G T G A T C G C T A G A T C G A C T A C G T G C A T A G T C | PRDM1(Zf)/Hela-PRDM1-ChIP-Seq(GSE31477)/Homer | 1e-58 | -1.337e+02 | 0.0000 | 3154.0 | 20.21% | 10898.5 | 14.88% | motif file (matrix) | svg |
| 231 | T C G A T C G A A G T C C G T A C T A G T A G C A C G T A C T G | MyoG(bHLH)/C2C12-MyoG-ChIP-Seq(GSE36024)/Homer | 1e-57 | -1.335e+02 | 0.0000 | 5622.0 | 36.02% | 21531.5 | 29.40% | motif file (matrix) | svg |
| 232 | C G A T C T A G A G T C A C G T A C G T T C A G G C T A C G T A G C A T G C A T C G A T A G T C C G T A G T C A A C T G | VND3(NAC)/colamp-VND3-DAP-Seq(GSE60143)/Homer | 1e-57 | -1.329e+02 | 0.0000 | 6827.0 | 43.74% | 26963.6 | 36.82% | motif file (matrix) | svg |
| 233 | C A T G A C T G C T A G T C G A T C G A T C G A T C G A T C A G T C A G T C A G T G A C T G A C C G T A A C T G T G C A C G A T A C T G | RBPJ:Ebox(?,bHLH)/Panc1-Rbpj1-ChIP-Seq(GSE47459)/Homer | 1e-57 | -1.319e+02 | 0.0000 | 1509.0 | 9.67% | 4382.8 | 5.98% | motif file (matrix) | svg |
| 234 | G A C T G T A C T G C A A C G T G A T C G C T A T C G A A C G T A G T C C G T A | Pdx1(Homeobox)/Islet-Pdx1-ChIP-Seq(SRA008281)/Homer | 1e-57 | -1.317e+02 | 0.0000 | 8058.0 | 51.63% | 32655.9 | 44.59% | motif file (matrix) | svg |
| 235 | C A G T T C A G A G C T G A C T A C G T A G T C G A T C G A C T C T G A A C T G G A T C C G T A C T G A A G T C G T A C | Rfx6(HTH)/Min6b1-Rfx6.HA-ChIP-Seq(GSE62844)/Homer | 1e-56 | -1.311e+02 | 0.0000 | 7503.0 | 48.07% | 30095.8 | 41.09% | motif file (matrix) | svg |
| 236 | A C T G A C G T C G A T C A G T C A T G C A T G C A G T G C A T C A G T C A T G | HuR(?)/HEK293-HuR-CLIP-Seq(GSE87887)/Homer | 1e-56 | -1.308e+02 | 0.0000 | 13446.0 | 86.15% | 59230.9 | 80.87% | motif file (matrix) | svg |
| 237 | C G A T G A T C T A C G C T G A G C T A C G T A G C A T A G T C C T A G C G T A G C A T C G A T | AT2G15740(C2H2)/col-AT2G15740-DAP-Seq(GSE60143)/Homer | 1e-56 | -1.305e+02 | 0.0000 | 13934.0 | 89.28% | 61865.5 | 84.47% | motif file (matrix) | svg |
| 238 | G C T A A G T C T A C G T G C A A T C G T C A G G C T A T C G A T C A G A G C T | ELF5(ETS)/T47D-ELF5-ChIP-Seq(GSE30407)/Homer | 1e-56 | -1.302e+02 | 0.0000 | 5507.0 | 35.29% | 21080.7 | 28.78% | motif file (matrix) | svg |
| 239 | G A T C C T G A G A T C G A T C C T A G G C T A A G T C C T G A G C T A C G T A | At4g16750(AP2EREBP)/col-At4g16750-DAP-Seq(GSE60143)/Homer | 1e-55 | -1.271e+02 | 0.0000 | 11713.0 | 75.05% | 50375.8 | 68.78% | motif file (matrix) | svg |
| 240 | T C A G T C A G A C G T G T A C G C T A T C A G C T G A A C T G A C T G A G C T A G T C C G T A | EAR2(NR)/K562-NR2F6-ChIP-Seq(Encode)/Homer | 1e-54 | -1.244e+02 | 0.0000 | 8805.0 | 56.42% | 36307.3 | 49.57% | motif file (matrix) | svg |
| 241 | A C G T T C G A T C G A A G T C G T C A T A C G A T G C A C G T A C T G A G C T | Myf5(bHLH)/GM-Myf5-ChIP-Seq(GSE24852)/Homer | 1e-53 | -1.240e+02 | 0.0000 | 3882.0 | 24.87% | 14107.9 | 19.26% | motif file (matrix) | svg |
| 242 | C G T A C G A T C T A G G T C A G A C T C G A T C T A G C G T A A C G T C A T G | LIN-39(Homeobox)/cElegans.L3-LIN39-ChIP-Seq(modEncode)/Homer | 1e-53 | -1.226e+02 | 0.0000 | 8398.0 | 53.81% | 34430.3 | 47.01% | motif file (matrix) | svg |
| 243 | C T A G A C T G A G T C A C T G A C T G A G C T A C T G T C A G | AT3G57600(AP2EREBP)/col-AT3G57600-DAP-Seq(GSE60143)/Homer | 1e-53 | -1.225e+02 | 0.0000 | 6605.0 | 42.32% | 26155.4 | 35.71% | motif file (matrix) | svg |
| 244 | G C A T C G T A G C A T C G T A T C G A C G T A C T G A A C T G C G T A C G T A C G T A A C G T A C T G G T C A G C A T | AT2G31460(REMB3)/col-AT2G31460-DAP-Seq(GSE60143)/Homer | 1e-53 | -1.225e+02 | 0.0000 | 2521.0 | 16.15% | 8451.8 | 11.54% | motif file (matrix) | svg |
| 245 | G C A T A C G T A C G T A T C G C G T A C G T A C G T A C G T A | At2g41835(C2H2)/col-At2g41835-DAP-Seq(GSE60143)/Homer | 1e-52 | -1.218e+02 | 0.0000 | 3416.0 | 21.89% | 12165.4 | 16.61% | motif file (matrix) | svg |
| 246 | C A G T G C T A G C A T T A C G C T G A C A G T T A G C C T G A | GATA15(C2C2gata)/col-GATA15-DAP-Seq(GSE60143)/Homer | 1e-52 | -1.209e+02 | 0.0000 | 12653.0 | 81.07% | 55260.4 | 75.45% | motif file (matrix) | svg |
| 247 | C G A T T C A G G T A C A C G T A C G T T C A G C G A T C G T A G T C A G C T A C G T A A G T C C G T A G T C A C A T G | ANAC057(NAC)/colamp-ANAC057-DAP-Seq(GSE60143)/Homer | 1e-52 | -1.208e+02 | 0.0000 | 7884.0 | 50.52% | 32068.8 | 43.79% | motif file (matrix) | svg |
| 248 | C G T A C G A T C A G T C A T G C G A T G T A C C A T G A C T G G A C T C A T G | CEJ1(AP2EREBP)/col-CEJ1-DAP-Seq(GSE60143)/Homer | 1e-52 | -1.203e+02 | 0.0000 | 12331.0 | 79.01% | 53625.5 | 73.22% | motif file (matrix) | svg |
| 249 | G A C T C T A G C T A G G T A C A G T C G A T C G A C T G A C T T A G C T C A G | NLP7(RWPRK)/col-NLP7-DAP-Seq(GSE60143)/Homer | 1e-52 | -1.201e+02 | 0.0000 | 11579.0 | 74.19% | 49842.3 | 68.05% | motif file (matrix) | svg |
| 250 | A G C T C T G A C T A G C T A G A C T G T A G C T G C A T C G A C T G A C T A G C A T G A C G T A T G C T C G A | RXR(NR),DR1/3T3L1-RXR-ChIP-Seq(GSE13511)/Homer | 1e-51 | -1.194e+02 | 0.0000 | 5516.0 | 35.34% | 21319.2 | 29.11% | motif file (matrix) | svg |
| 251 | A C T G T C A G A G C T G A C T C A T G A G T C A G T C G C T A C G A T C T A G T C A G G T A C C T G A T C G A | Rfx1(HTH)/NPC-H3K4me1-ChIP-Seq(GSE16256)/Homer | 1e-50 | -1.174e+02 | 0.0000 | 1803.0 | 11.55% | 5651.2 | 7.72% | motif file (matrix) | svg |
| 252 | C G T A G C T A C G A T C T A G A C G T G T C A C G T A C G T A A G T C C G T A T G C A T A C G | FoxL2(Forkhead)/Ovary-FoxL2-ChIP-Seq(GSE60858)/Homer | 1e-50 | -1.174e+02 | 0.0000 | 5935.0 | 38.03% | 23232.5 | 31.72% | motif file (matrix) | svg |
| 253 | C G T A C T A G C A G T A C G T C G T A A C T G C A T G G C A T T C A G C T G A | MYB49(MYB)/col-MYB49-DAP-Seq(GSE60143)/Homer | 1e-50 | -1.161e+02 | 0.0000 | 9364.0 | 60.00% | 39134.0 | 53.43% | motif file (matrix) | svg |
| 254 | A G T C C G A T A C T G A T C G T G A C G C T A C A T G A T C G T G A C C G A T A C T G T A G C G T A C G T C A | Tlx?(NR)/NPC-H3K4me1-ChIP-Seq(GSE16256)/Homer | 1e-50 | -1.159e+02 | 0.0000 | 2109.0 | 13.51% | 6875.7 | 9.39% | motif file (matrix) | svg |
| 255 | A C T G A C T G A G T C A C T G A C T G A G T C A C G T T C A G | ERF2(AP2EREBP)/colamp-ERF2-DAP-Seq(GSE60143)/Homer | 1e-50 | -1.154e+02 | 0.0000 | 5977.0 | 38.30% | 23459.6 | 32.03% | motif file (matrix) | svg |
| 256 | A G T C A C G T A C T G A G C T A C G T A C G T G T C A A G T C | Foxo1(Forkhead)/RAW-Foxo1-ChIP-Seq(Fan\_et\_al.)/Homer | 1e-50 | -1.153e+02 | 0.0000 | 10028.0 | 64.25% | 42337.3 | 57.81% | motif file (matrix) | svg |
| 257 | C A T G T G C A G A C T C A T G C G T A A G T C T C A G G C A T T G A C C G T A | bZIP50(bZIP)/colamp-bZIP50-DAP-Seq(GSE60143)/Homer | 1e-49 | -1.149e+02 | 0.0000 | 11757.0 | 75.33% | 50834.3 | 69.41% | motif file (matrix) | svg |
| 258 | G T A C G C T A C G A T C A G T A G T C G C T A C G A T C G A T A G T C G C T A | WUS1(Homeobox)/colamp-WUS1-DAP-Seq(GSE60143)/Homer | 1e-49 | -1.139e+02 | 0.0000 | 3838.0 | 24.59% | 14084.6 | 19.23% | motif file (matrix) | svg |
| 259 | C T A G C T A G C G T A C G T A T A C G C G A T C T A G C T G A C T G A C G T A T A C G G A C T | PU.1:IRF8(ETS:IRF)/pDC-Irf8-ChIP-Seq(GSE66899)/Homer | 1e-49 | -1.136e+02 | 0.0000 | 1115.0 | 7.14% | 3087.7 | 4.22% | motif file (matrix) | svg |
| 260 | C T G A A G T C G A T C C A T G G C T A G A T C C T G A G C T A G C T A C G A T | AT1G77200(AP2EREBP)/colamp-AT1G77200-DAP-Seq(GSE60143)/Homer | 1e-49 | -1.135e+02 | 0.0000 | 11889.0 | 76.18% | 51525.9 | 70.35% | motif file (matrix) | svg |
| 261 | G T C A C G T A A C G T A T C G C G T A A C G T A C G T C T A G | ATHB7(Homeobox)/col-ATHB7-DAP-Seq(GSE60143)/Homer | 1e-48 | -1.119e+02 | 0.0000 | 7443.0 | 47.69% | 30213.3 | 41.25% | motif file (matrix) | svg |
| 262 | C T A G T C G A T G A C A G T C C G T A A C T G G T A C A C G T A C T G A C T G | BHLHA15(bHLH)/NIH3T3-BHLHB8.HA-ChIP-Seq(GSE119782)/Homer | 1e-47 | -1.103e+02 | 0.0000 | 7723.0 | 49.48% | 31543.7 | 43.07% | motif file (matrix) | svg |
| 263 | G A C T G C T A T G C A A G T C A C G T A C G T A C G T C G A T A C G T T A C G | At3g45610(C2C2dof)/col-At3g45610-DAP-Seq(GSE60143)/Homer | 1e-47 | -1.096e+02 | 0.0000 | 9013.0 | 57.75% | 37608.0 | 51.35% | motif file (matrix) | svg |
| 264 | G T A C A C G T A C G T T A C G A T G C C A T G T A C G G T A C T C A G A T G C C G T A G T C A A C T G A G C T G C T A | AT1G19040(NAC)/col-AT1G19040-DAP-Seq(GSE60143)/Homer | 1e-47 | -1.096e+02 | 0.0000 | 1530.0 | 9.80% | 4678.2 | 6.39% | motif file (matrix) | svg |
| 265 | T A C G T C G A G A C T A C T G C T G A A G T C T C A G G A C T T G A C C T G A | Atf1(bZIP)/K562-ATF1-ChIP-Seq(GSE31477)/Homer | 1e-47 | -1.095e+02 | 0.0000 | 7895.0 | 50.59% | 32360.9 | 44.19% | motif file (matrix) | svg |
| 266 | C T G A G A C T C A T G C T A G A G T C A C T G A C T G A G T C A C T G T C A G | ERF11(AP2EREBP)/col-ERF11-DAP-Seq(GSE60143)/Homer | 1e-47 | -1.091e+02 | 0.0000 | 10205.0 | 65.39% | 43326.8 | 59.16% | motif file (matrix) | svg |
| 267 | C A G T A T C G C T G A A G T C T C A G C A G T T A G C C T G A A T G C T A C G | FEA4(bZIP)/Corn-FEA4-ChIP-Seq(GSE61954)/Homer | 1e-47 | -1.090e+02 | 0.0000 | 10172.0 | 65.18% | 43168.7 | 58.94% | motif file (matrix) | svg |
| 268 | T C A G C T G A C G T A C G T A T A C G G C A T C T A G C T G A C G T A C G T A T A C G G A C T | IRF1(IRF)/PBMC-IRF1-ChIP-Seq(GSE43036)/Homer | 1e-47 | -1.083e+02 | 0.0000 | 816.0 | 5.23% | 2065.5 | 2.82% | motif file (matrix) | svg |
| 269 | G C T A C G T A C G T A G C A T C A T G C T A G G A T C A C T G T C A G G A T C A C T G T C A G | ERF4(AP2EREBP)/colamp-ERF4-DAP-Seq(GSE60143)/Homer | 1e-46 | -1.082e+02 | 0.0000 | 11541.0 | 73.95% | 49896.4 | 68.13% | motif file (matrix) | svg |
| 270 | G C T A A C T G T C G A C T G A C T G A A C G T T A G C C T G A C G T A C G A T | Cux2(Homeobox)/Liver-Cux2-ChIP-Seq(GSE35985)/Homer | 1e-46 | -1.081e+02 | 0.0000 | 7909.0 | 50.68% | 32455.5 | 44.31% | motif file (matrix) | svg |
| 271 | C G A T C G T A G C T A G A C T T C G A A G C T A G T C A C T G T C G A A G C T C T G A C G A T | ZBTB38(Zf)/Hela-ZBTB38-ChIP-seq(GSE108618)/Homer | 1e-46 | -1.066e+02 | 0.0000 | 14970.0 | 95.92% | 68079.9 | 92.96% | motif file (matrix) | svg |
| 272 | C T A G C A G T C G T A A C G T A G T C A C T G C G T A A G C T A G T C G A T C | HNF6(Homeobox)/Liver-Hnf6-ChIP-Seq(ERP000394)/Homer | 1e-46 | -1.062e+02 | 0.0000 | 9544.0 | 61.15% | 40212.6 | 54.91% | motif file (matrix) | svg |
| 273 | G A T C C G T A G A C T C T A G G A T C C T G A G A C T C T G A G A C T C T A G G A T C C T G A G A C T C T G A G A C T | OCT:OCT(POU,Homeobox)/NPC-OCT6-ChIP-Seq(GSE43916)/Homer | 1e-43 | -1.013e+02 | 0.0000 | 524.0 | 3.36% | 1135.7 | 1.55% | motif file (matrix) | svg |
| 274 | C G T A T A C G T C G A A C T G A C T G C G T A C G T A T A C G A G C T T A C G | PU.1(ETS)/ThioMac-PU.1-ChIP-Seq(GSE21512)/Homer | 1e-43 | -9.975e+01 | 0.0000 | 2872.0 | 18.40% | 10222.4 | 13.96% | motif file (matrix) | svg |
| 275 | G T A C A C G T A C G T T C A G G C T A C G T A C G A T G C A T G C A T A G T C C G T A G T C A C A T G G A C T G C T A | ANAC070(NAC)/colamp-ANAC070-DAP-Seq(GSE60143)/Homer | 1e-43 | -9.971e+01 | 0.0000 | 9848.0 | 63.10% | 41816.2 | 57.10% | motif file (matrix) | svg |
| 276 | G A C T A G T C C G A T A C T G C T G A T G A C G T A C C G T A A T C G G C A T C T G A C T A G | Bcl11a(Zf)/HSPC-BCL11A-ChIP-Seq(GSE104676)/Homer | 1e-42 | -9.861e+01 | 0.0000 | 4651.0 | 29.80% | 17895.8 | 24.43% | motif file (matrix) | svg |
| 277 | G T A C A C G T A C G T T C A G C G T A C G T A C G A T G C A T G C A T A G T C C G T A G T C A C A T G G A C T G C T A | VND1(NAC)/col-VND1-DAP-Seq(GSE60143)/Homer | 1e-42 | -9.748e+01 | 0.0000 | 7842.0 | 50.25% | 32384.1 | 44.22% | motif file (matrix) | svg |
| 278 | G C A T C T A G G T A C A G T C C G A T A C T G C T A G C T A G G T A C G C T A | ZNF416(Zf)/HEK293-ZNF416.GFP-ChIP-Seq(GSE58341)/Homer | 1e-42 | -9.708e+01 | 0.0000 | 6072.0 | 38.91% | 24270.3 | 33.14% | motif file (matrix) | svg |
| 279 | G C A T C G A T A T G C A G C T T C G A T A C G G C T A C G T A C A T G T G A C G C A T C G A T A G T C A G C T C G T A | AT3G09735(S1Falike)/col-AT3G09735-DAP-Seq(GSE60143)/Homer | 1e-41 | -9.610e+01 | 0.0000 | 3217.0 | 20.61% | 11736.4 | 16.02% | motif file (matrix) | svg |
| 280 | G A C T A C T G C G T A A G T C T C A G G C A T G T A C C G T A A C G T G A T C | TGA1(bZIP)/colamp-TGA1-DAP-Seq(GSE60143)/Homer | 1e-41 | -9.610e+01 | 0.0000 | 5249.0 | 33.63% | 20595.8 | 28.12% | motif file (matrix) | svg |
| 281 | C G A T C A G T C T A G G C T A A G T C C G T A T C A G A G T C A C G T A C T G A C G T G T A C G C T A G C T A G C T A | bZIP52(bZIP)/colamp-bZIP52-DAP-Seq(GSE60143)/Homer | 1e-40 | -9.421e+01 | 0.0000 | 8454.0 | 54.17% | 35323.6 | 48.23% | motif file (matrix) | svg |
| 282 | C T G A C T G A C T A G T C G A C G T A A T G C C G T A A C T G C G T A A C G T C T G A C G A T A G C T C G T A A C G T A G T C C G A T T A C G G T C A G C A T | GATA(Zf),IR3/iTreg-Gata3-ChIP-Seq(GSE20898)/Homer | 1e-40 | -9.345e+01 | 0.0000 | 1757.0 | 11.26% | 5749.6 | 7.85% | motif file (matrix) | svg |
| 283 | C A G T T C G A A G T C A C G T A C G T T C A G C G A T G C T A G C T A C G T A G C A T A G T C C G T A T G C A A C T G | ANAC045(NAC)/col-ANAC045-DAP-Seq(GSE60143)/Homer | 1e-40 | -9.292e+01 | 0.0000 | 13036.0 | 83.53% | 57768.0 | 78.88% | motif file (matrix) | svg |
| 284 | T G A C A G T C C G T A A C T G G T A C A C G T A C T G A C G T G A C T G A T C | Twist2(bHLH)/Myoblast-Twist2.Ty1-ChIP-Seq(GSE127998)/Homer | 1e-38 | -8.799e+01 | 0.0000 | 9210.0 | 59.01% | 39047.6 | 53.31% | motif file (matrix) | svg |
| 285 | T A C G T A G C C A T G C A G T A C G T C T A G C G T A A G T C G A C T G C A T G C A T C A G T | WRKY11(WRKY)/col-WRKY11-DAP-Seq(GSE60143)/Homer | 1e-38 | -8.782e+01 | 0.0000 | 1817.0 | 11.64% | 6061.6 | 8.28% | motif file (matrix) | svg |
| 286 | A G T C G A T C A G T C C G T A A T C G C A G T A G T C G T A C C T G A A C T G T C A G A G C T A G C T A G C T A G C T | PRDM15(Zf)/ESC-Prdm15-ChIP-Seq(GSE73694)/Homer | 1e-38 | -8.760e+01 | 0.0000 | 6862.0 | 43.97% | 28088.1 | 38.35% | motif file (matrix) | svg |
| 287 | G A T C G A T C A G T C G T A C C G A T G T A C G T A C A G T C A G T C A G T C G C T A G A T C | ZNF148(Zf)/MDAMB231-ZNF148-ChIP-Seq(GSE147020)/Homer | 1e-37 | -8.584e+01 | 0.0000 | 1701.0 | 10.90% | 5622.8 | 7.68% | motif file (matrix) | svg |
| 288 | C T G A A G T C C G A T A G C T A T G C G T A C A C G T A T C G C A G T G C A T | Elf4(ETS)/BMDM-Elf4-ChIP-Seq(GSE88699)/Homer | 1e-35 | -8.285e+01 | 0.0000 | 6871.0 | 44.03% | 28245.4 | 38.57% | motif file (matrix) | svg |
| 289 | T G A C G C T A T C G A T G C A A G T C A G T C C G T A A G T C C G T A C T G A G C T A G T A C | RUNX2(Runt)/PCa-RUNX2-ChIP-Seq(GSE33889)/Homer | 1e-35 | -8.219e+01 | 0.0000 | 5882.0 | 37.69% | 23750.8 | 32.43% | motif file (matrix) | svg |
| 290 | C G A T G A C T C G A T T C A G G A C T A C G T C A G T C T G A G A C T G A C T A G C T C G A T A C T G A T C G G T A C G C T A | NF1:FOXA1(CTF,Forkhead)/LNCAP-FOXA1-ChIP-Seq(GSE27824)/Homer | 1e-35 | -8.167e+01 | 0.0000 | 441.0 | 2.83% | 975.3 | 1.33% | motif file (matrix) | svg |
| 291 | T G A C C T G A C T A G T C G A C T G A A T G C C G T A A C T G G C A T G T A C G C A T A T C G G C A T A G C T G A T C | PR(NR)/T47D-PR-ChIP-Seq(GSE31130)/Homer | 1e-35 | -8.073e+01 | 0.0000 | 11010.0 | 70.55% | 47903.6 | 65.41% | motif file (matrix) | svg |
| 292 | C G T A G C A T C A T G C T A G A G T C A C T G A T C G G T A C A C T G T C A G | At2g33710(AP2EREBP)/colamp-At2g33710-DAP-Seq(GSE60143)/Homer | 1e-34 | -8.051e+01 | 0.0000 | 13594.0 | 87.10% | 60908.7 | 83.16% | motif file (matrix) | svg |
| 293 | T C G A G C A T A C G T C T A G G T A C T C G A G C A T T G A C T C G A A C G T | Chop(bZIP)/MEF-Chop-ChIP-Seq(GSE35681)/Homer | 1e-34 | -7.977e+01 | 0.0000 | 2626.0 | 16.83% | 9516.8 | 12.99% | motif file (matrix) | svg |
| 294 | C T A G A C T G A C G T C G T A A C T G A C T G A G C T C T A G T C A G C T A G | MYB93(MYB)/colamp-MYB93-DAP-Seq(GSE60143)/Homer | 1e-34 | -7.902e+01 | 0.0000 | 9543.0 | 61.15% | 40861.4 | 55.79% | motif file (matrix) | svg |
| 295 | G C T A T C G A C G T A C T A G A G C T G T C A G T C A C G T A A G T C C G T A | FOXA1(Forkhead)/LNCAP-FOXA1-ChIP-Seq(GSE27824)/Homer | 1e-33 | -7.814e+01 | 0.0000 | 6562.0 | 42.05% | 26945.9 | 36.79% | motif file (matrix) | svg |
| 296 | C G A T C A G T C A G T C A T G G T C A G A T C C G T A T C A G A G T C A C G T C T A G A C G T G T A C G T C A G C T A | VIP1(bZIP)/col-VIP1-DAP-Seq(GSE60143)/Homer | 1e-33 | -7.789e+01 | 0.0000 | 1425.0 | 9.13% | 4626.4 | 6.32% | motif file (matrix) | svg |
| 297 | A T C G A G T C A C T G A G T C A G T C A C T G G A C T G A C T | PUCHI(AP2EREBP)/colamp-PUCHI-DAP-Seq(GSE60143)/Homer | 1e-33 | -7.774e+01 | 0.0000 | 8756.0 | 56.10% | 37158.5 | 50.74% | motif file (matrix) | svg |
| 298 | G T A C G T C A G T A C G T C A G T A C G T C A G T A C G T C A G T A C G T C A | SeqBias: CA-repeat | 1e-33 | -7.763e+01 | 0.0000 | 14619.0 | 93.67% | 66458.0 | 90.74% | motif file (matrix) | svg |
| 299 | C G T A C G T A C G T A C G T A C G T A A C T G A C T G A G T C | dof42(C2C2dof)/col-dof42-DAP-Seq(GSE60143)/Homer | 1e-33 | -7.755e+01 | 0.0000 | 5431.0 | 34.80% | 21825.0 | 29.80% | motif file (matrix) | svg |
| 300 | G C A T G A T C T C A G G C T A G A C T A G T C C T A G C G T A C A T G G T C A | GATA20(C2C2gata)/colamp-GATA20-DAP-Seq(GSE60143)/Homer | 1e-33 | -7.705e+01 | 0.0000 | 14502.0 | 92.92% | 65821.3 | 89.87% | motif file (matrix) | svg |
| 301 | G A T C G A T C G A T C C G T A G T A C A G T C G C A T C G T A G T A C G A T C | MYB58(MYB)/colamp-MYB58-DAP-Seq(GSE60143)/Homer | 1e-33 | -7.695e+01 | 0.0000 | 9053.0 | 58.01% | 38584.6 | 52.68% | motif file (matrix) | svg |
| 302 | T A C G T C G A C G T A C G T A C G T A C T G A A C T G A C G T C G T A T C G A | AT2G28810(C2C2dof)/colamp-AT2G28810-DAP-Seq(GSE60143)/Homer | 1e-33 | -7.692e+01 | 0.0000 | 11767.0 | 75.40% | 51722.3 | 70.62% | motif file (matrix) | svg |
| 303 | T C A G G A C T G T C A C G T A A C G T A T C G C G T A A C G T A C G T C T G A | ATHB15(HB)/col-ATHB15-DAP-Seq(GSE60143)/Homer | 1e-33 | -7.682e+01 | 0.0000 | 3685.0 | 23.61% | 14108.5 | 19.26% | motif file (matrix) | svg |
| 304 | C G A T C T G A A G T C A C G T A C G T T C A G C G T A C G T A C G T A G C A T C G A T A G T C C G T A G T C A A C T G | VND4(NAC)/colamp-VND4-DAP-Seq(GSE60143)/Homer | 1e-33 | -7.632e+01 | 0.0000 | 7690.0 | 49.27% | 32199.1 | 43.96% | motif file (matrix) | svg |
| 305 | A C T G G A T C G A C T A C T G A C G T C A T G A C T G A C G T A G C T C G A T | RUNX-AML(Runt)/CD4+-PolII-ChIP-Seq(Barski\_et\_al.)/Homer | 1e-32 | -7.545e+01 | 0.0000 | 4782.0 | 30.64% | 18969.3 | 25.90% | motif file (matrix) | svg |
| 306 | G A C T C T A G G A T C C A G T A C T G C T G A A T G C G C A T A T G C C T G A | MafA(bZIP)/Islet-MafA-ChIP-Seq(GSE30298)/Homer | 1e-32 | -7.469e+01 | 0.0000 | 5933.0 | 38.01% | 24164.2 | 32.99% | motif file (matrix) | svg |
| 307 | T A C G T G C A A G T C C G T A A C G T T G A C A C G T A C T G A C T G G C A T | TCF4(bHLH)/SHSY5Y-TCF4-ChIP-Seq(GSE96915)/Homer | 1e-32 | -7.422e+01 | 0.0000 | 8433.0 | 54.03% | 35730.2 | 48.79% | motif file (matrix) | svg |
| 308 | C T A G T C G A C T G A C G T A T A C G G A C T T C A G T C G A G T C A T G C A T A C G A G C T | IRF2(IRF)/Erythroblas-IRF2-ChIP-Seq(GSE36985)/Homer | 1e-32 | -7.378e+01 | 0.0000 | 840.0 | 5.38% | 2425.9 | 3.31% | motif file (matrix) | svg |
| 309 | C T G A T C A G C T G A C T A G C A T G A C G T A T G C C G T A A T G C G C A T T C A G C T G A A C T G A C G T C A G T A G T C C G T A C A G T C T A G C A T G | VDR(NR),DR3/GM10855-VDR+vitD-ChIP-Seq(GSE22484)/Homer | 1e-32 | -7.373e+01 | 0.0000 | 1673.0 | 10.72% | 5672.1 | 7.74% | motif file (matrix) | svg |
| 310 | A C G T C T A G A G C T A C G T A C G T C T G A A G T C G A C T A G C T C G T A | FOXM1(Forkhead)/MCF7-FOXM1-ChIP-Seq(GSE72977)/Homer | 1e-31 | -7.323e+01 | 0.0000 | 5957.0 | 38.17% | 24310.4 | 33.19% | motif file (matrix) | svg |
| 311 | A C G T T G A C A G T C A G C T A G T C A G C T A C T G G A C T A G C T G A C T | REF6(Zf)/Arabidopsis-REF6-ChIP-Seq(GSE106942)/Homer | 1e-30 | -7.126e+01 | 0.0000 | 3352.0 | 21.48% | 12777.2 | 17.45% | motif file (matrix) | svg |
| 312 | A G T C T A G C G A C T A C G T C T A G A C G T A C G T A C G T C T G A A G T C G C T A G A C T C G T A C T A G A C T G | Foxa3(Forkhead)/Liver-Foxa3-ChIP-Seq(GSE77670)/Homer | 1e-30 | -7.081e+01 | 0.0000 | 2566.0 | 16.44% | 9420.5 | 12.86% | motif file (matrix) | svg |
| 313 | G T A C A C T G A T G C T G A C C T A G G A C T G T A C C G T A G C A T G C A T | ERF8(AP2EREBP)/colamp-ERF8-DAP-Seq(GSE60143)/Homer | 1e-30 | -7.053e+01 | 0.0000 | 11937.0 | 76.48% | 52723.2 | 71.99% | motif file (matrix) | svg |
| 314 | G C T A C G T A C G T A G C A T C A T G C T A G A G T C A C T G T A C G A G T C C A T G T A C G | RAP26(AP2EREBP)/colamp-RAP26-DAP-Seq(GSE60143)/Homer | 1e-30 | -6.980e+01 | 0.0000 | 12216.0 | 78.27% | 54129.6 | 73.91% | motif file (matrix) | svg |
| 315 | C T A G G T A C A C G T A C G T A T C G G C A T A G C T A G C T A G C T G C A T G A C T C G T A G T C A A C T G G A C T | VND6(NAC)/col-VND6-DAP-Seq(GSE60143)/Homer | 1e-30 | -6.977e+01 | 0.0000 | 10259.0 | 65.73% | 44543.5 | 60.82% | motif file (matrix) | svg |
| 316 | G C A T G C A T G A T C G C A T T C G A A C T G C G T A C G T A A T C G T A G C C G A T A C G T A G T C A G C T C G T A | HSF6(HSF)/col-HSF6-DAP-Seq(GSE60143)/Homer | 1e-30 | -6.922e+01 | 0.0000 | 1698.0 | 10.88% | 5838.1 | 7.97% | motif file (matrix) | svg |
| 317 | G A C T G A C T G A T C C G T A G T A C A G T C G C A T C G T A G T A C G A T C G C A T G C T A | MYB74(MYB)/colamp-MYB74-DAP-Seq(GSE60143)/Homer | 1e-29 | -6.829e+01 | 0.0000 | 6150.0 | 39.41% | 25317.8 | 34.57% | motif file (matrix) | svg |
| 318 | C T A G C A T G G A C T C G T A C T A G A C T G A C G T C T A G C T A G T C A G | MYB17(MYB)/colamp-MYB17-DAP-Seq(GSE60143)/Homer | 1e-29 | -6.810e+01 | 0.0000 | 5712.0 | 36.60% | 23330.0 | 31.85% | motif file (matrix) | svg |
| 319 | C G A T C T G A A G T C A C G T A C G T T C A G G C A T C G A T G C T A G C T A C G T A A G T C C G T A G T C A A C T G | CUC1(NAC)/col-CUC1-DAP-Seq(GSE60143)/Homer | 1e-29 | -6.793e+01 | 0.0000 | 5353.0 | 34.30% | 21712.0 | 29.65% | motif file (matrix) | svg |
| 320 | C G A T C T G A G T A C A C G T A C G T T C A G C G T A C G T A G C T A G C A T G C A T A G T C C G T A G T C A C A T G | NST1(NAC)/colamp-NST1-DAP-Seq(GSE60143)/Homer | 1e-29 | -6.730e+01 | 0.0000 | 7868.0 | 50.41% | 33274.5 | 45.43% | motif file (matrix) | svg |
| 321 | A C T G C T A G A G T C A C T G A C T G A G T C A C T G T A C G | ERF104(AP2EREBP)/col-ERF104-DAP-Seq(GSE60143)/Homer | 1e-29 | -6.728e+01 | 0.0000 | 9017.0 | 57.78% | 38678.8 | 52.81% | motif file (matrix) | svg |
| 322 | A T G C G A C T A C G T C T A G A C G T A C G T A C G T C T G A G A T C G C T A A G C T C G T A | Foxa2(Forkhead)/Liver-Foxa2-ChIP-Seq(GSE25694)/Homer | 1e-29 | -6.703e+01 | 0.0000 | 5818.0 | 37.28% | 23839.9 | 32.55% | motif file (matrix) | svg |
| 323 | A G T C C T G A A T C G A G C T A G C T G A C T A G T C G C T A A C G T C G A T G C A T C G A T A T C G C G T A T A G C G C A T A T G C C G T A | bZIP:IRF(bZIP,IRF)/Th17-BatF-ChIP-Seq(GSE39756)/Homer | 1e-28 | -6.641e+01 | 0.0000 | 2091.0 | 13.40% | 7502.1 | 10.24% | motif file (matrix) | svg |
| 324 | C G A T T A C G T G C A G T A C G A T C G A C T A G C T A C G T A T C G G T A C G A T C G T A C G A T C G T C A | PPARE(NR),DR1/3T3L1-Pparg-ChIP-Seq(GSE13511)/Homer | 1e-28 | -6.611e+01 | 0.0000 | 4722.0 | 30.26% | 18925.8 | 25.84% | motif file (matrix) | svg |
| 325 | T G C A C T G A A T G C G T C A A C G T A T G C A C G T A C T G A C T G T G C A | ZBTB18(Zf)/HEK293-ZBTB18.GFP-ChIP-Seq(GSE58341)/Homer | 1e-27 | -6.425e+01 | 0.0000 | 3045.0 | 19.51% | 11595.5 | 15.83% | motif file (matrix) | svg |
| 326 | T A C G A C T G A G C T G T A C C G T A T C G A C T G A A C T G C A T G A C G T A G T C C G T A | COUP-TFII(NR)/K562-NR2F1-ChIP-Seq(Encode)/Homer | 1e-27 | -6.244e+01 | 0.0000 | 9144.0 | 58.59% | 39420.6 | 53.82% | motif file (matrix) | svg |
| 327 | A T G C T C A G T C G A G C A T A C T G C G T A A G T C T C A G G A C T T G A C C G T A A G C T | Atf2(bZIP)/3T3L1-Atf2-ChIP-Seq(GSE56872)/Homer | 1e-27 | -6.231e+01 | 0.0000 | 3043.0 | 19.50% | 11627.2 | 15.88% | motif file (matrix) | svg |
| 328 | T A G C C T A G T C G A G A C T A C T G C T G A A G T C T C A G G C A T T G A C C T G A A G C T | Atf7(bZIP)/3T3L1-Atf7-ChIP-Seq(GSE56872)/Homer | 1e-26 | -6.156e+01 | 0.0000 | 4785.0 | 30.66% | 19322.9 | 26.38% | motif file (matrix) | svg |
| 329 | C T A G A G T C A G C T A C T G C G T A C A G T C G T A C T G A T A G C T G A C | Unknown5/Drosophila-Promoters/Homer | 1e-26 | -6.094e+01 | 0.0000 | 7868.0 | 50.41% | 33458.9 | 45.68% | motif file (matrix) | svg |
| 330 | C A T G C T A G A G T C A C T G A C T G G T A C C A T G T A C G | AT1G28160(AP2EREBP)/colamp-AT1G28160-DAP-Seq(GSE60143)/Homer | 1e-26 | -6.093e+01 | 0.0000 | 12594.0 | 80.69% | 56242.2 | 76.79% | motif file (matrix) | svg |
| 331 | C G A T T C G A A C T G G T C A C G T A C G A T G T A C G A C T | At3g04030(G2like)/col-At3g04030-DAP-Seq(GSE60143)/Homer | 1e-26 | -6.037e+01 | 0.0000 | 8148.0 | 52.21% | 34785.3 | 47.50% | motif file (matrix) | svg |
| 332 | G A C T C A G T G A T C G A T C A C G T G A T C C T G A T A C G C G T A G T C A | STAT6(Stat)/Macrophage-Stat6-ChIP-Seq(GSE38377)/Homer | 1e-26 | -5.997e+01 | 0.0000 | 3476.0 | 22.27% | 13562.4 | 18.52% | motif file (matrix) | svg |
| 333 | G A T C G C A T G C A T A G T C A G C T T C G A T A C G G C T A C G T A C T A G T G A C G C A T C G A T G A T C A G C T | HSFC1(HSF)/col-HSFC1-DAP-Seq(GSE60143)/Homer | 1e-26 | -5.993e+01 | 0.0000 | 1341.0 | 8.59% | 4527.3 | 6.18% | motif file (matrix) | svg |
| 334 | A G T C C G T A T G A C A T G C G C A T C T G A G T A C G A T C | MYB55(MYB)/colamp-MYB55-DAP-Seq(GSE60143)/Homer | 1e-26 | -5.989e+01 | 0.0000 | 9869.0 | 63.23% | 42952.0 | 58.65% | motif file (matrix) | svg |
| 335 | T C G A T G A C G T A C C G T A A C G T G A C T A C G T A C T G A C T G A G C T | Mesp1(bHLH)/ESC-Mesp1-ChIP-Seq(GSE165102)/Homer | 1e-25 | -5.938e+01 | 0.0000 | 4341.0 | 27.81% | 17395.9 | 23.75% | motif file (matrix) | svg |
| 336 | T C A G A G C T A C G T A C G T G T A C G A T C C G T A C T A G C A T G G T C A C G T A T C G A | STAT4(Stat)/CD4-Stat4-ChIP-Seq(GSE22104)/Homer | 1e-25 | -5.936e+01 | 0.0000 | 5484.0 | 35.14% | 22531.1 | 30.76% | motif file (matrix) | svg |
| 337 | G C A T G C A T G T A C G A C T T C G A A C T G G C T A C G T A A T C G T G A C G C A T G A C T A G T C A G C T C T G A | AGL95(ND)/col-AGL95-DAP-Seq(GSE60143)/Homer | 1e-25 | -5.905e+01 | 0.0000 | 782.0 | 5.01% | 2351.7 | 3.21% | motif file (matrix) | svg |
| 338 | T C G A T A G C G T C A A C T G A C T G C G T A C G T A C T A G A G C T T C A G | ERG(ETS)/VCaP-ERG-ChIP-Seq(GSE14097)/Homer | 1e-25 | -5.905e+01 | 0.0000 | 7564.0 | 48.47% | 32097.6 | 43.83% | motif file (matrix) | svg |
| 339 | T A G C G T A C A G T C G T A C C G A T A G T C A G T C A G T C A G T C A G T C C G T A G A T C | Zfp281(Zf)/ES-Zfp281-ChIP-Seq(GSE81042)/Homer | 1e-25 | -5.846e+01 | 0.0000 | 358.0 | 2.29% | 838.8 | 1.15% | motif file (matrix) | svg |
| 340 | G C A T C G T A C G A T G A C T A C T G C T G A G A C T G A T C | Hnf6b(Homeobox)/LNCaP-Hnf6b-ChIP-Seq(GSE106305)/Homer | 1e-24 | -5.733e+01 | 0.0000 | 10012.0 | 64.15% | 43713.4 | 59.69% | motif file (matrix) | svg |
| 341 | T C G A A C T G A C T G C G T A C G T A T C G A A G T C C T G A A T C G G T A C G C A T C A T G | ETS:E-box(ETS,bHLH)/HPC7-Scl-ChIP-Seq(GSE22178)/Homer | 1e-24 | -5.680e+01 | 0.0000 | 510.0 | 3.27% | 1373.0 | 1.87% | motif file (matrix) | svg |
| 342 | G C T A C G T A C G T A G C A T C A T G C T A G A G T C A C T G A C T G A G T C A C T G T C A G | ABR1(AP2EREBP)/colamp-ABR1-DAP-Seq(GSE60143)/Homer | 1e-24 | -5.643e+01 | 0.0000 | 11125.0 | 71.28% | 49117.5 | 67.06% | motif file (matrix) | svg |
| 343 | C T A G T A G C A T G C C T A G A G T C A G T C C T A G G A C T G A C T G C T A | CRF10(AP2EREBP)/col100-CRF10-DAP-Seq(GSE60143)/Homer | 1e-23 | -5.453e+01 | 0.0000 | 12270.0 | 78.62% | 54797.6 | 74.82% | motif file (matrix) | svg |
| 344 | G C A T G C A T G A T C G A C T T C G A T C A G G C T A C G T A A C T G G T A C G C A T G C A T A G T C A G C T C G T A | HSF7(HSF)/colamp-HSF7-DAP-Seq(GSE60143)/Homer | 1e-23 | -5.405e+01 | 0.0000 | 1299.0 | 8.32% | 4442.7 | 6.07% | motif file (matrix) | svg |
| 345 | C G A T G T C A A G T C A C G T A C G T A C T G G A C T C G A T A T C G G C T A G T C A A G T C C G T A G T C A A C T G | ANAC017(NAC)/colamp-ANAC017-DAP-Seq(GSE60143)/Homer | 1e-23 | -5.354e+01 | 0.0000 | 1572.0 | 10.07% | 5565.2 | 7.60% | motif file (matrix) | svg |
| 346 | C G A T G C T A G C T A G C A T G C T A C G T A A G T C A C G T A C G T A C G T C G A T A G C T | At5g62940(C2C2dof)/col-At5g62940-DAP-Seq(GSE60143)/Homer | 1e-22 | -5.276e+01 | 0.0000 | 13742.0 | 88.05% | 62270.0 | 85.02% | motif file (matrix) | svg |
| 347 | A C G T C T A G C G T A A G T C G T A C A C G T A C G T A C G T G T C A G T A C T G A C G A C T | Nur77(NR)/K562-NR4A1-ChIP-Seq(GSE31363)/Homer | 1e-22 | -5.269e+01 | 0.0000 | 1563.0 | 10.01% | 5542.5 | 7.57% | motif file (matrix) | svg |
| 348 | A T C G T G A C A T G C C T G A T C A G G A C T A G T C C G A T T C A G T C G A C A T G C T A G C T A G C G T A C T A G C T A G C T G A C T A G C T A G A T G C | ZSCAN22(Zf)/HEK293-ZSCAN22.GFP-ChIP-Seq(GSE58341)/Homer | 1e-22 | -5.237e+01 | 0.0000 | 308.0 | 1.97% | 710.7 | 0.97% | motif file (matrix) | svg |
| 349 | C G T A C T G A C G T A C T A G T C G A C T A G A C T G C G T A C G T A T A C G A G C T A T C G | SpiB(ETS)/OCILY3-SPIB-ChIP-Seq(GSE56857)/Homer | 1e-22 | -5.213e+01 | 0.0000 | 1466.0 | 9.39% | 5152.2 | 7.03% | motif file (matrix) | svg |
| 350 | A T G C G A C T A G C T C T A G C G T A C T A G C G A T C T A G A T C G G A T C | Nkx2.2(Homeobox)/NPC-Nkx2.2-ChIP-Seq(GSE61673)/Homer | 1e-22 | -5.209e+01 | 0.0000 | 11793.0 | 75.56% | 52512.4 | 71.70% | motif file (matrix) | svg |
| 351 | G A T C C T G A A G T C A G C T A C G T A C G T A C G T A C G T | At1g64620(C2C2dof)/colamp-At1g64620-DAP-Seq(GSE60143)/Homer | 1e-22 | -5.183e+01 | 0.0000 | 8884.0 | 56.92% | 38516.8 | 52.59% | motif file (matrix) | svg |
| 352 | C A T G C T A G A G C T G A C T C A T G A G T C G A T C G C T A C G A T C T A G T C A G G T A C C T G A T C G A | X-box(HTH)/NPC-H3K4me1-ChIP-Seq(GSE16256)/Homer | 1e-22 | -5.145e+01 | 0.0000 | 783.0 | 5.02% | 2436.6 | 3.33% | motif file (matrix) | svg |
| 353 | C G T A C G T A G C A T A C T G C G T A A G C T C T G A C G T A T A C G C T G A | ELT-3(Gata)/cElegans-L1-ELT3-ChIP-Seq(modEncode)/Homer | 1e-21 | -5.055e+01 | 0.0000 | 4348.0 | 27.86% | 17661.2 | 24.11% | motif file (matrix) | svg |
| 354 | T C G A A G C T A C G T A C G T A G T C A G T C A C G T A T C G G A C T A T C G | EWS:ERG-fusion(ETS)/CADO\_ES1-EWS:ERG-ChIP-Seq(SRA014231)/Homer | 1e-21 | -5.024e+01 | 0.0000 | 3740.0 | 23.96% | 14963.2 | 20.43% | motif file (matrix) | svg |
| 355 | A G T C G A T C A G C T C G T A G T A C A G T C G C A T C T G A G T A C G A T C | AT4G26030(C2H2)/col-AT4G26030-DAP-Seq(GSE60143)/Homer | 1e-21 | -4.979e+01 | 0.0000 | 8931.0 | 57.22% | 38805.4 | 52.98% | motif file (matrix) | svg |
| 356 | G C T A C G T A C G A T G A C T G C A T T G C A A G T C A G C T A C G T A C G T C G A T G A C T | DAG2(C2C2dof)/col-DAG2-DAP-Seq(GSE60143)/Homer | 1e-21 | -4.965e+01 | 0.0000 | 8813.0 | 56.47% | 38252.9 | 52.23% | motif file (matrix) | svg |
| 357 | A T G C T C G A A G T C A G C T A C G T G T A C A G T C G C T A C T A G C A T G G T C A C T G A T C A G A G T C | Stat3+il21(Stat)/CD4-Stat3-ChIP-Seq(GSE19198)/Homer | 1e-21 | -4.947e+01 | 0.0000 | 4373.0 | 28.02% | 17803.0 | 24.31% | motif file (matrix) | svg |
| 358 | C T G A A C T G C G T A A C G T G T C A A G C T A G C T G A C T G A C T C A G T | CCA(Myb)/Arabidopsis-CCA.GFP-ChIP-Seq(GSE70533)/Homer | 1e-21 | -4.916e+01 | 0.0000 | 7734.0 | 49.55% | 33201.8 | 45.33% | motif file (matrix) | svg |
| 359 | C A G T C A T G T G C A G T A C C G T A T C A G G T A C G A C T T C A G C T G A | bZIP18(bZIP)/colamp-bZIP18-DAP-Seq(GSE60143)/Homer | 1e-21 | -4.868e+01 | 0.0000 | 14893.0 | 95.43% | 68453.3 | 93.47% | motif file (matrix) | svg |
| 360 | T G C A C G T A A C T G T C A G C A G T C A T G T C A G G A T C T A C G A G T C T G C A A C T G A C T G T G A C G T C A | ZNF165(Zf)/WHIM12-ZNF165-ChIP-Seq(GSE65937)/Homer | 1e-20 | -4.789e+01 | 0.0000 | 653.0 | 4.18% | 1979.4 | 2.70% | motif file (matrix) | svg |
| 361 | C G T A C G T A C T G A A C T G A C G T A G T C C G T A C G T A A G T C A C T G A T G C G A T C | WRKY46(WRKY)/colamp-WRKY46-DAP-Seq(GSE60143)/Homer | 1e-20 | -4.671e+01 | 0.0000 | 1436.0 | 9.20% | 5118.8 | 6.99% | motif file (matrix) | svg |
| 362 | C T A G C A T G C A T G T A C G A G T C G C A T A G C T C T A G A C G T A G T C G A C T A C T G A C T G A C T G T C G A | Zfp809(Zf)/ES-Zfp809-ChIP-Seq(GSE70799)/Homer | 1e-20 | -4.653e+01 | 0.0000 | 1052.0 | 6.74% | 3556.0 | 4.86% | motif file (matrix) | svg |
| 363 | T C G A G A C T T C A G T G C A G T A C G T A C A G C T G T A C C A T G T C G A C A T G C A T G A C G T A G T C C T G A | FXR(NR),ER2/Liver-FXR-ChIP-Seq(GSE133700)/Homer | 1e-19 | -4.585e+01 | 0.0000 | 3894.0 | 24.95% | 15766.3 | 21.53% | motif file (matrix) | svg |
| 364 | T G C A A G C T A C G T C T A G G A T C C T A G G A T C G T C A C T G A A G T C | CEBP(bZIP)/ThioMac-CEBPb-ChIP-Seq(GSE21512)/Homer | 1e-19 | -4.520e+01 | 0.0000 | 7262.0 | 46.53% | 31136.2 | 42.51% | motif file (matrix) | svg |
| 365 | G C T A T C G A C G T A C T A G A G C T G T C A G T C A C G T A A G T C C G T A | FOXA1(Forkhead)/MCF7-FOXA1-ChIP-Seq(GSE26831)/Homer | 1e-19 | -4.499e+01 | 0.0000 | 5103.0 | 32.70% | 21227.7 | 28.98% | motif file (matrix) | svg |
| 366 | C G T A A C T G C G T A A C G T C A G T A G T C A G C T G C A T G C T A C G A T | At2g01060(G2like)/colamp-At2g01060-DAP-Seq(GSE60143)/Homer | 1e-19 | -4.473e+01 | 0.0000 | 14445.0 | 92.55% | 66114.5 | 90.27% | motif file (matrix) | svg |
| 367 | A C T G A G C T A G T C G T C A A G C T T C A G A T G C G A T C G C A T A T C G T C G A T A G C C G A T C A T G T A G C | Pax8(Paired,Homeobox)/Thyroid-Pax8-ChIP-Seq(GSE26938)/Homer | 1e-19 | -4.471e+01 | 0.0000 | 2060.0 | 13.20% | 7779.6 | 10.62% | motif file (matrix) | svg |
| 368 | A G T C G A C T C A G T A C T G C T A G T G A C G C T A A T G C G C A T A T C G C G A T A C T G G A T C G T A C G T C A C T G A | NF1(CTF)/LNCAP-NF1-ChIP-Seq(Unpublished)/Homer | 1e-19 | -4.466e+01 | 0.0000 | 2101.0 | 13.46% | 7956.0 | 10.86% | motif file (matrix) | svg |
| 369 | G A C T G A T C C T G A A G T C A G T C A C T G C G T A A G T C G T A C G C T A G C A T C G A T | At1g19210(AP2EREBP)/colamp-At1g19210-DAP-Seq(GSE60143)/Homer | 1e-19 | -4.433e+01 | 0.0000 | 13236.0 | 84.81% | 59908.3 | 81.80% | motif file (matrix) | svg |
| 370 | C T G A C G A T C T A G C G T A A G C T C G A T C A G T C T G A G A C T C T A G C T A G A T G C | PBX2(Homeobox)/K562-PBX2-ChIP-Seq(Encode)/Homer | 1e-19 | -4.429e+01 | 0.0000 | 7734.0 | 49.55% | 33367.3 | 45.56% | motif file (matrix) | svg |
| 371 | C T G A A T C G A G C T A G C T A C G T T A G C C T G A T A C G C G A T A C G T G A C T A G T C | ISRE(IRF)/ThioMac-LPS-Expression(GSE23622)/Homer | 1e-18 | -4.365e+01 | 0.0000 | 365.0 | 2.34% | 962.8 | 1.31% | motif file (matrix) | svg |
| 372 | C T A G T C G A A C G T A C G T C A T G A G T C C T G A C G A T A G T C C G T A | AARE(HLH)/mES-cMyc-ChIP-Seq/Homer | 1e-18 | -4.364e+01 | 0.0000 | 1263.0 | 8.09% | 4457.0 | 6.09% | motif file (matrix) | svg |
| 373 | T C G A C G T A C G T A T C G A A C T G G T A C C G T A A G C T G T C A G C A T | At3g24120(G2like)/col-At3g24120-DAP-Seq(GSE60143)/Homer | 1e-18 | -4.333e+01 | 0.0000 | 14470.0 | 92.71% | 66276.4 | 90.49% | motif file (matrix) | svg |
| 374 | C G T A G A C T C G T A A C G T C A G T A G T C A G C T G A C T | KAN2(G2like)/colamp-KAN2-DAP-Seq(GSE60143)/Homer | 1e-18 | -4.215e+01 | 0.0000 | 9288.0 | 59.51% | 40760.8 | 55.65% | motif file (matrix) | svg |
| 375 | A T G C A G T C G T A C A G C T T C G A C T A G G A T C C T G A G T C A A G T C G C T A T C A G | Rfx5(HTH)/GM12878-Rfx5-ChIP-Seq(GSE31477)/Homer | 1e-17 | -4.124e+01 | 0.0000 | 2782.0 | 17.83% | 10982.0 | 14.99% | motif file (matrix) | svg |
| 376 | T C G A G T A C T C G A T C G A C A T G A T G C A C G T A C T G A C T G A G T C C G T A C T A G A G T C A T C G A G T C | Unknown3/Drosophila-Promoters/Homer | 1e-17 | -4.110e+01 | 0.0000 | 858.0 | 5.50% | 2857.5 | 3.90% | motif file (matrix) | svg |
| 377 | C G A T T C G A G T A C A C G T A C G T T C A G G C A T G C T A T G C A G C A T C G T A A G T C C G T A T G A C C A T G | ANAC092(NAC)/colamp-ANAC092-DAP-Seq(GSE60143)/Homer | 1e-17 | -4.104e+01 | 0.0000 | 5280.0 | 33.83% | 22159.2 | 30.26% | motif file (matrix) | svg |
| 378 | A T C G A G T C A G T C C G T A A C T G G C A T | hINR(CPE) | 1e-17 | -4.097e+01 | 0.0000 | 8032.0 | 51.46% | 34880.5 | 47.63% | motif file (matrix) | svg |
| 379 | G A T C A G T C G A C T G C T A G T A C A G T C G C A T G C T A G T A C G A T C | MYB61(MYB)/colamp-MYB61-DAP-Seq(GSE60143)/Homer | 1e-17 | -4.059e+01 | 0.0000 | 10925.0 | 70.00% | 48654.0 | 66.43% | motif file (matrix) | svg |
| 380 | T A G C T A G C G A C T C T A G A G C T A G T C G T C A T G C A A C G T A T G C G C T A T G C A | Pbx3(Homeobox)/GM12878-PBX3-ChIP-Seq(GSE32465)/Homer | 1e-17 | -4.035e+01 | 0.0000 | 1824.0 | 11.69% | 6866.2 | 9.38% | motif file (matrix) | svg |
| 381 | G C A T T C A G C T G A A T C G A C T G C G A T G A T C C T G A | THRb(NR)/Liver-NR1A2-ChIP-Seq(GSE52613)/Homer | 1e-17 | -3.998e+01 | 0.0000 | 13964.0 | 89.47% | 63723.4 | 87.01% | motif file (matrix) | svg |
| 382 | A T G C A G C T T C A G T G A C T C A G A T G C T G C A A C G T A T C G G A T C A C T G A G T C | NRF1(NRF)/MCF7-NRF1-ChIP-Seq(Unpublished)/Homer | 1e-17 | -3.979e+01 | 0.0000 | 818.0 | 5.24% | 2716.9 | 3.71% | motif file (matrix) | svg |
| 383 | C G T A T C G A G A T C G C A T C G T A A C G T G T A C T C A G G T C A G A C T C G T A C T A G | DREF/Drosophila-Promoters/Homer | 1e-17 | -3.940e+01 | 0.0000 | 1022.0 | 6.55% | 3541.4 | 4.84% | motif file (matrix) | svg |
| 384 | C A T G G T C A A G T C C G T A C T A G G A T C C G A T A C T G A C G T G T A C C G T A C G T A | bZIP69(bZIP)/col-bZIP69-DAP-Seq(GSE60143)/Homer | 1e-16 | -3.887e+01 | 0.0000 | 805.0 | 5.16% | 2677.6 | 3.66% | motif file (matrix) | svg |
| 385 | A T G C A G T C C T G A A G T C C G A T A C G T A G T C A G T C A C G T A T C G G A C T A C G T | Etv2(ETS)/ES-ER71-ChIP-Seq(GSE59402)/Homer | 1e-16 | -3.874e+01 | 0.0000 | 4924.0 | 31.55% | 20619.0 | 28.15% | motif file (matrix) | svg |
| 386 | C G T A A C T G G T C A A C G T A T C G C A G T C T A G T C A G C G T A A C T G C G T A A C G T C G T A C T G A T A C G | GATA3(Zf),DR4/iTreg-Gata3-ChIP-Seq(GSE20898)/Homer | 1e-16 | -3.863e+01 | 0.0000 | 906.0 | 5.81% | 3084.6 | 4.21% | motif file (matrix) | svg |
| 387 | G C A T G A C T A T G C A G C T T C G A A C T G C G T A C G T A A T C G A T G C G C A T G A C T G A T C A G C T T C G A | HSFA1E(HSF)/col-HSFA1E-DAP-Seq(GSE60143)/Homer | 1e-16 | -3.860e+01 | 0.0000 | 519.0 | 3.33% | 1570.6 | 2.14% | motif file (matrix) | svg |
| 388 | A C T G A G T C G T C A C G T A A G T C C G T A C T A G C T A G G A C T C A T G | SCRT1(Zf)/HEK293-SCRT1.eGFP-ChIP-Seq(Encode)/Homer | 1e-16 | -3.826e+01 | 0.0000 | 3224.0 | 20.66% | 13007.9 | 17.76% | motif file (matrix) | svg |
| 389 | C G T A G A C T C G A T A T C G G T A C G C A T C A T G C G T A T A C G G C A T G T A C C G T A C A T G A T G C G C T A C T A G G C A T G C A T G C A T G A C T | MafB(bZIP)/BMM-Mafb-ChIP-Seq(GSE75722)/Homer | 1e-16 | -3.812e+01 | 0.0000 | 2400.0 | 15.38% | 9395.8 | 12.83% | motif file (matrix) | svg |
| 390 | A G C T G T C A T G C A A G T C A C G T A C G T A C G T C G A T G A C T T A C G | AT3G12130(C3H)/colamp-AT3G12130-DAP-Seq(GSE60143)/Homer | 1e-16 | -3.696e+01 | 0.0000 | 12062.0 | 77.29% | 54310.4 | 74.15% | motif file (matrix) | svg |
| 391 | T C G A G A C T A T C G C G T A A G T C C T A G G C A T G T A C C T G A A C G T G A T C G C T A | TGA4(bZIP)/colamp-TGA4-DAP-Seq(GSE60143)/Homer | 1e-16 | -3.687e+01 | 0.0000 | 3860.0 | 24.73% | 15881.1 | 21.68% | motif file (matrix) | svg |
| 392 | T G A C G C T A T G A C C G T A T C A G G A T C C G T A C A T G C A T G C T A G C T A G C T A G | Unknown-ESC-element(?)/mES-Nanog-ChIP-Seq(GSE11724)/Homer | 1e-15 | -3.682e+01 | 0.0000 | 2523.0 | 16.17% | 9964.0 | 13.60% | motif file (matrix) | svg |
| 393 | T C A G A C T G A C G T C G T A A C T G A C T G A C G T C T A G | MYB51(MYB)/col-MYB51-DAP-Seq(GSE60143)/Homer | 1e-15 | -3.641e+01 | 0.0000 | 8134.0 | 52.12% | 35530.3 | 48.51% | motif file (matrix) | svg |
| 394 | T A C G A T G C G A C T A C T G A G C T A G T C G T C A T G C A A C G T A G T C G C T A T G C A | Pknox1(Homeobox)/ES-Prep1-ChIP-Seq(GSE63282)/Homer | 1e-15 | -3.640e+01 | 0.0000 | 1991.0 | 12.76% | 7668.5 | 10.47% | motif file (matrix) | svg |
| 395 | C G T A C T G A T C A G A C T G G T C A C G T A A C G T G T A C C G A T G C A T | AT5G45580(G2like)/colamp-AT5G45580-DAP-Seq(GSE60143)/Homer | 1e-15 | -3.630e+01 | 0.0000 | 11331.0 | 72.60% | 50772.0 | 69.32% | motif file (matrix) | svg |
| 396 | T C G A T C A G T C G A A C T G C A T G A C G T A G T C C T G A | COUP-TFII(NR)/Artia-Nr2f2-ChIP-Seq(GSE46497)/Homer | 1e-15 | -3.519e+01 | 0.0000 | 10400.0 | 66.64% | 46325.0 | 63.25% | motif file (matrix) | svg |
| 397 | C T A G C T G A A G T C G C T A C G A T A C T G G A C T G A T C G A T C C T G A C T A G C T G A T G A C G C T A C G A T T C A G G A C T G A T C G A T C T G A C | p53(p53)/Saos-p53-ChIP-Seq(GSE15780)/Homer | 1e-15 | -3.501e+01 | 0.0000 | 927.0 | 5.94% | 3224.4 | 4.40% | motif file (matrix) | svg |
| 398 | C T A G C T G A A G T C G C T A C G A T A C T G G A C T G A T C G A T C C T G A C T A G C T G A T G A C G C T A C G A T T C A G G A C T G A T C G A T C T G A C | p53(p53)/Saos-p53-ChIP-Seq/Homer | 1e-15 | -3.501e+01 | 0.0000 | 927.0 | 5.94% | 3224.4 | 4.40% | motif file (matrix) | svg |
| 399 | C G T A T G A C T C G A A G T C C G T A A T C G A T G C A C G T A C T G A G T C | E2A(bHLH)/proBcell-E2A-ChIP-Seq(GSE21978)/Homer | 1e-15 | -3.485e+01 | 0.0000 | 6929.0 | 44.40% | 29968.5 | 40.92% | motif file (matrix) | svg |
| 400 | C A T G G A T C C T G A A G T C C T A G C G T A G C T A G C A T G A T C G A T C A G T C C T A G C G T A C A T G C T A G | PLT1(AP2EREBP)/colamp-PLT1-DAP-Seq(GSE60143)/Homer | 1e-15 | -3.460e+01 | 0.0000 | 1515.0 | 9.71% | 5676.3 | 7.75% | motif file (matrix) | svg |
| 401 | A T G C T A G C A G C T A G C T T G A C G A C T T C A G T A C G G T C A C T G A A T C G T A G C G A C T C A G T A G T C A G C T T C G A A T C G T G C A T G C A | HRE(HSF)/HepG2-HSF1-ChIP-Seq(GSE31477)/Homer | 1e-14 | -3.391e+01 | 0.0000 | 980.0 | 6.28% | 3458.6 | 4.72% | motif file (matrix) | svg |
| 402 | G T A C G T A C G T C A G C T A C G T A C G T A C G T A C T A G C T A G C T A G | SEP3(MADS)/Arabidoposis-Flower-Sep3-ChIP-Seq/Homer | 1e-14 | -3.250e+01 | 0.0000 | 7196.0 | 46.11% | 31303.5 | 42.74% | motif file (matrix) | svg |
| 403 | C T A G A C T G A C G T C G T A A C T G A C T G A C G T T C A G C T G A T C G A | MYB107(MYB)/col-MYB107-DAP-Seq(GSE60143)/Homer | 1e-13 | -3.217e+01 | 0.0000 | 11340.0 | 72.66% | 50969.7 | 69.59% | motif file (matrix) | svg |
| 404 | T C G A G C A T A C T G C T G A A G T C T C A G G A C T G T A C C G T A A G C T A G T C G A T C | c-Jun-CRE(bZIP)/K562-cJun-ChIP-Seq(GSE31477)/Homer | 1e-13 | -3.153e+01 | 0.0000 | 2277.0 | 14.59% | 9031.2 | 12.33% | motif file (matrix) | svg |
| 405 | T C A G C T G A C T A G C A T G A C G T A T G C C T G A C T G A C T G A C T A G C A T G A C G T A T G C C T G A | TR4(NR),DR1/Hela-TR4-ChIP-Seq(GSE24685)/Homer | 1e-13 | -3.128e+01 | 0.0000 | 545.0 | 3.49% | 1753.1 | 2.39% | motif file (matrix) | svg |
| 406 | A G T C G A C T A C T G G A T C G T A C C G T A T G A C A G T C C G A T A G C T A C G T A C G T C T A G G A C T C T G A | ZNF7(Zf)/HepG2-ZNF7.Flag-ChIP-Seq(Encode)/Homer | 1e-13 | -3.078e+01 | 0.0000 | 3989.0 | 25.56% | 16664.1 | 22.75% | motif file (matrix) | svg |
| 407 | G A C T G A T C A G T C C G T A T G A C A G T C G C A T C T G A G T A C G A T C G C A T G A C T | MYB10(MYB)/col-MYB10-DAP-Seq(GSE60143)/Homer | 1e-13 | -3.069e+01 | 0.0000 | 4308.0 | 27.60% | 18108.5 | 24.73% | motif file (matrix) | svg |
| 408 | T G C A A G C T C A T G C G T A A G C T A C T G G A T C G T C A C G T A A G C T | Atf4(bZIP)/MEF-Atf4-ChIP-Seq(GSE35681)/Homer | 1e-13 | -3.058e+01 | 0.0000 | 3266.0 | 20.93% | 13428.6 | 18.34% | motif file (matrix) | svg |
| 409 | T G C A C T G A C A T G C T A G C A G T A G T C C G T A A T G C A T G C T A C G G C A T T C A G G T C A G A T C G T A C | ERE(NR),IR3/MCF7-ERa-ChIP-Seq(Unpublished)/Homer | 1e-13 | -3.053e+01 | 0.0000 | 1840.0 | 11.79% | 7159.7 | 9.78% | motif file (matrix) | svg |
| 410 | C G T A C T A G G A C T G T C A G T C A C G T A A G T C C G T A T C G A T C G A T C G A C G T A C T G A C T A G G C T A C G T A T A G C C G T A C G A T C G T A | FOXA1:AR(Forkhead,NR)/LNCAP-AR-ChIP-Seq(GSE27824)/Homer | 1e-13 | -3.042e+01 | 0.0000 | 224.0 | 1.44% | 569.9 | 0.78% | motif file (matrix) | svg |
| 411 | G C T A C G T A A C T G C G T A C G A T A C G T A G T C A G C T | At3g12730(G2like)/colamp-At3g12730-DAP-Seq(GSE60143)/Homer | 1e-13 | -3.040e+01 | 0.0000 | 11093.0 | 71.08% | 49845.4 | 68.06% | motif file (matrix) | svg |
| 412 | A G T C C G T A C G A T A G T C G T C A A G T C A C G T C T G A | Unknown2/Drosophila-Promoters/Homer | 1e-13 | -3.015e+01 | 0.0000 | 7513.0 | 48.14% | 32879.6 | 44.89% | motif file (matrix) | svg |
| 413 | C G A T C T A G T C A G C A G T C G T A A G T C G C T A A C G T G A C T A T G C A G T C G C T A | PRDM10(Zf)/HEK293-PRDM10.eGFP-ChIP-Seq(Encode)/Homer | 1e-12 | -2.917e+01 | 0.0000 | 4071.0 | 26.08% | 17092.4 | 23.34% | motif file (matrix) | svg |
| 414 | C T G A A G C T A C G T A C G T A G T C G A C T G A C T C T G A C T G A C T A G C G T A C G T A | STAT6(Stat)/CD4-Stat6-ChIP-Seq(GSE22104)/Homer | 1e-12 | -2.865e+01 | 0.0000 | 3119.0 | 19.98% | 12836.9 | 17.53% | motif file (matrix) | svg |
| 415 | C T G A T A C G G C A T C T A G A T G C G A T C C G A T A C T G C T A G G A T C C T G A A T G C | MYRF(MYRF)/CFPAC1-MYRF-ChIP-Seq(GSE145627)/Homer | 1e-12 | -2.836e+01 | 0.0000 | 2418.0 | 15.49% | 9739.5 | 13.30% | motif file (matrix) | svg |
| 416 | T C G A T A G C T G C A A C T G A C T G C G T A C G T A C T A G G A C T T A C G | ETS1(ETS)/Jurkat-ETS1-ChIP-Seq(GSE17954)/Homer | 1e-12 | -2.836e+01 | 0.0000 | 6457.0 | 41.37% | 28046.3 | 38.29% | motif file (matrix) | svg |
| 417 | G T A C C G T A C G T A T A C G G C A T G T A C C G T A C A T G A G T C C G T A C G T A C G A T G C A T G C A T G A C T | MafF(bZIP)/HepG2-MafF-ChIP-Seq(GSE31477)/Homer | 1e-12 | -2.770e+01 | 0.0000 | 1994.0 | 12.78% | 7901.0 | 10.79% | motif file (matrix) | svg |
| 418 | C T A G C T G A C G T A C G T A C G T A C G T A A C T G A C G T C T A G G T C A | COG1(C2C2dof)/col-COG1-DAP-Seq(GSE60143)/Homer | 1e-12 | -2.769e+01 | 0.0000 | 8349.0 | 53.50% | 36904.5 | 50.39% | motif file (matrix) | svg |
| 419 | T G C A C G T A G T C A A G C T A G T C G C T A T A G C C G A T C T A G G A T C | Gfi1b(Zf)/HPC7-Gfi1b-ChIP-Seq(GSE22178)/Homer | 1e-11 | -2.755e+01 | 0.0000 | 4393.0 | 28.15% | 18612.0 | 25.41% | motif file (matrix) | svg |
| 420 | C G A T C T A G A C G T G T C A C G T A C G T A A G T C C G T A | Foxo3(Forkhead)/U2OS-Foxo3-ChIP-Seq(E-MTAB-2701)/Homer | 1e-11 | -2.753e+01 | 0.0000 | 5401.0 | 34.61% | 23214.4 | 31.70% | motif file (matrix) | svg |
| 421 | A G T C C T A G A T C G C A G T C G A T A G C T G T A C A C T G C A T G C A T G | ZBED2(Zf)/SUIT2-ZBED2.HA-ChIP-Seq(GSE141606)/Homer | 1e-11 | -2.748e+01 | 0.0000 | 8384.0 | 53.72% | 37078.4 | 50.63% | motif file (matrix) | svg |
| 422 | A G C T G C T A T G C A A G T C A C G T A C G T A C G T C G A T A G C T T C A G | dof24(C2C2dof)/col-dof24-DAP-Seq(GSE60143)/Homer | 1e-11 | -2.661e+01 | 0.0000 | 10979.0 | 70.35% | 49454.9 | 67.52% | motif file (matrix) | svg |
| 423 | C T A G T A C G G A T C G T A C G C T A A G C T A G C T G T C A T C G A T A G C | Nanog(Homeobox)/mES-Nanog-ChIP-Seq(GSE11724)/Homer | 1e-11 | -2.659e+01 | 0.0000 | 14899.0 | 95.46% | 68912.3 | 94.09% | motif file (matrix) | svg |
| 424 | T C G A C T G A C G T A C G T A C G T A C T G A A C T G A C G T C T G A C T G A | AT5G63260(C3H)/col-AT5G63260-DAP-Seq(GSE60143)/Homer | 1e-11 | -2.615e+01 | 0.0000 | 11539.0 | 73.93% | 52174.7 | 71.24% | motif file (matrix) | svg |
| 425 | G C A T A G T C G A C T T C G A T A C G G T C A T C G A A C T G T A G C G C A T G C A T A T G C | AT2G01818(PLATZ)/col-AT2G01818-DAP-Seq(GSE60143)/Homer | 1e-11 | -2.596e+01 | 0.0000 | 1256.0 | 8.05% | 4765.5 | 6.51% | motif file (matrix) | svg |
| 426 | T A C G T C A G T G C A A G C T T G A C A G C T A G T C A C T G G A T C A C T G T C G A A C T G C T G A C T G A A T G C | ZBTB33(Zf)/GM12878-ZBTB33-ChIP-Seq(GSE32465)/Homer | 1e-11 | -2.569e+01 | 0.0000 | 1248.0 | 8.00% | 4737.2 | 6.47% | motif file (matrix) | svg |
| 427 | A C T G T G A C A C T G A C G T A C G T A C T G C G T A A G T C A G C T C G A T G C A T A C G T | WRKY17(WRKY)/colamp-WRKY17-DAP-Seq(GSE60143)/Homer | 1e-10 | -2.474e+01 | 0.0000 | 187.0 | 1.20% | 482.3 | 0.66% | motif file (matrix) | svg |
| 428 | T C G A C G T A A G T C A G C T C G T A A G T C T C G A G C T A G A C T C G A T A G T C A G T C A G T C C T G A T C A G T G C A T C G A C A G T A T C G A G T C | GFY-Staf(?,Zf)/Promoter/Homer | 1e-10 | -2.473e+01 | 0.0000 | 297.0 | 1.90% | 879.4 | 1.20% | motif file (matrix) | svg |
| 429 | C A T G C T G A A G T C A C T G A C T G A G C T A C T G A T C G | ESE3(AP2EREBP)/col-ESE3-DAP-Seq(GSE60143)/Homer | 1e-10 | -2.465e+01 | 0.0000 | 11500.0 | 73.68% | 52050.1 | 71.07% | motif file (matrix) | svg |
| 430 | C G T A C T G A C T A G C T G A A G T C G C T A C G A T A T C G G A C T G A T C A G T C C T G A C T A G C T A G A G T C G C T A C G A T C T A G G A T C G A T C | p73(p53)/Trachea-p73-ChIP-Seq(PRJNA310161)/Homer | 1e-10 | -2.322e+01 | 0.0000 | 429.0 | 2.75% | 1402.2 | 1.91% | motif file (matrix) | svg |
| 431 | G C A T G C T A G C T A C G T A G C A T G C T A C T A G C G T A C G T A A C T G C G T A C G A T A C G T A G T C G A C T | At1g68670(G2like)/colamp-At1g68670-DAP-Seq(GSE60143)/Homer | 1e-9 | -2.297e+01 | 0.0000 | 3737.0 | 23.94% | 15821.4 | 21.60% | motif file (matrix) | svg |
| 432 | G A C T A G C T G T A C G A C T C T G A A C T G G T C A C T G A A T G C T A C G G A C T A C G T A G T C G A C T C T G A | HRE(HSF)/Striatum-HSF1-ChIP-Seq(GSE38000)/Homer | 1e-9 | -2.268e+01 | 0.0000 | 1119.0 | 7.17% | 4258.0 | 5.81% | motif file (matrix) | svg |
| 433 | C G T A A T G C C G A T A C G T A G T C C G T A C G T A C G T A C T A G A T C G | TCFL2(HMG)/K562-TCF7L2-ChIP-Seq(GSE29196)/Homer | 1e-9 | -2.201e+01 | 0.0000 | 666.0 | 4.27% | 2374.1 | 3.24% | motif file (matrix) | svg |
| 434 | C G A T C G A T G C A T G C A T G T C A A G T C A G C T A C G T A C G T C G A T G A C T A C G T | OBP4(C2C2dof)/col-OBP4-DAP-Seq(GSE60143)/Homer | 1e-9 | -2.178e+01 | 0.0000 | 8239.0 | 52.79% | 36670.2 | 50.07% | motif file (matrix) | svg |
| 435 | G C T A G C A T G A C T G C A T T C A G G T A C G C T A G C A T C T G A G C T A T A G C G C T A C T G A C G A T C T A G | OCT4-SOX2-TCF-NANOG(POU,Homeobox,HMG)/mES-Oct4-ChIP-Seq(GSE11431)/Homer | 1e-9 | -2.165e+01 | 0.0000 | 834.0 | 5.34% | 3077.9 | 4.20% | motif file (matrix) | svg |
| 436 | G C A T C T A G A C T G A C G T C G T A A C T G A C T G C G A T C T A G T C G A T C G A G C T A | MYB40(MYB)/col-MYB40-DAP-Seq(GSE60143)/Homer | 1e-9 | -2.154e+01 | 0.0000 | 3469.0 | 22.23% | 14668.8 | 20.03% | motif file (matrix) | svg |
| 437 | T G C A G C A T C G A T C G T A C A G T A C T G G T A C C G T A C T G A A G C T G T C A A C T G C T A G G T C A C G A T A C T G G T A C T G C A C G T A A G C T | CEBP:CEBP(bZIP)/MEF-Chop-ChIP-Seq(GSE35681)/Homer | 1e-9 | -2.154e+01 | 0.0000 | 1030.0 | 6.60% | 3905.2 | 5.33% | motif file (matrix) | svg |
| 438 | G C A T G A C T T G A C A G C T T C G A A C T G C G T A C T G A T C A G T A G C C G A T C G A T G T A C A G C T T C G A | AT1G23810(Orphan)/col-AT1G23810-DAP-Seq(GSE60143)/Homer | 1e-9 | -2.108e+01 | 0.0000 | 247.0 | 1.58% | 729.1 | 1.00% | motif file (matrix) | svg |
| 439 | T C G A C A T G C T G A C G T A A T C G G T A C G C A T C G A T A G T C A G C T T C G A T A C G C G T A C G T A C A T G | HSFA6A(HSF)/col-HSFA6A-DAP-Seq(GSE60143)/Homer | 1e-9 | -2.082e+01 | 0.0000 | 190.0 | 1.22% | 523.9 | 0.72% | motif file (matrix) | svg |
| 440 | T C A G G C A T A C T G C G T A A G T C C T A G G C A T T G A C | TGA9(bZIP)/colamp-TGA9-DAP-Seq(GSE60143)/Homer | 1e-9 | -2.081e+01 | 0.0000 | 11192.0 | 71.71% | 50743.3 | 69.28% | motif file (matrix) | svg |
| 441 | G A C T A T C G C T G A A G T C T C A G G A C T G T A C C T G A A G C T G T A C | TGA6(bZIP)/colamp-TGA6-DAP-Seq(GSE60143)/Homer | 1e-9 | -2.079e+01 | 0.0000 | 7952.0 | 50.95% | 35374.5 | 48.30% | motif file (matrix) | svg |
| 442 | T C A G A C T G C A G T A G T C A G T C G T C A C G T A C G T A A C T G C A G T A G T C A G T C C T G A T G C A A G C T | dHNF4(NR)/Fly-HNF4-ChIP-Seq(GSE73675)/Homer | 1e-8 | -2.050e+01 | 0.0000 | 429.0 | 2.75% | 1439.3 | 1.97% | motif file (matrix) | svg |
| 443 | C T G A G A T C G C A T A C T G C G T A A C G T C G T A C G T A T A C G T C G A | PQM-1(?)/cElegans-L3-ChIP-Seq(modEncode)/Homer | 1e-8 | -2.037e+01 | 0.0000 | 4025.0 | 25.79% | 17238.7 | 23.54% | motif file (matrix) | svg |
| 444 | T A C G C G T A T C A G G A C T C T A G A C T G C A G T T A G C T C G A A C G T G T A C C T A G A G T C A G T C G A T C | ZNF669(Zf)/HEK293-ZNF669.GFP-ChIP-Seq(GSE58341)/Homer | 1e-8 | -2.004e+01 | 0.0000 | 1281.0 | 8.21% | 5016.7 | 6.85% | motif file (matrix) | svg |
| 445 | G C T A C G T A C G T A C G T A A C T G A C G T A G T C C G T A C G T A A G T C C A T G T A G C G T A C C G T A C G T A | WRKY7(WRKY)/colamp-WRKY7-DAP-Seq(GSE60143)/Homer | 1e-8 | -1.999e+01 | 0.0000 | 63.0 | 0.40% | 109.7 | 0.15% | motif file (matrix) | svg |
| 446 | C T A G T A C G G A T C G T C A T G C A A C G T T G C A G C T A T C G A T G C A | Hoxa9(Homeobox)/ChickenMSG-Hoxa9.Flag-ChIP-Seq(GSE86088)/Homer | 1e-8 | -1.960e+01 | 0.0000 | 13107.0 | 83.98% | 60088.8 | 82.04% | motif file (matrix) | svg |
| 447 | G A C T C A G T A G C T C G A T A G T C G A T C A G T C C G T A A T G C T C A G | Rbpj1(?)/Panc1-Rbpj1-ChIP-Seq(GSE47459)/Homer | 1e-8 | -1.948e+01 | 0.0000 | 7169.0 | 45.93% | 31781.6 | 43.39% | motif file (matrix) | svg |
| 448 | C T G A A C G T A C G T A C G T A G T C G A C T C G A T C T G A A C T G C G T A C G T A T C G A | STAT5(Stat)/mCD4+-Stat5-ChIP-Seq(GSE12346)/Homer | 1e-8 | -1.938e+01 | 0.0000 | 1744.0 | 11.17% | 7052.4 | 9.63% | motif file (matrix) | svg |
| 449 | G C T A C G T A A C G T A T C G C G T A A C G T A C G T C T A G | ATHB6(Homeobox)/col-ATHB6-DAP-Seq(GSE60143)/Homer | 1e-8 | -1.937e+01 | 0.0000 | 9605.0 | 61.54% | 43245.0 | 59.05% | motif file (matrix) | svg |
| 450 | G T A C G A T C C A G T A G T C A G T C A G T C T G C A G A T C C T G A A T G C G T C A A C G T | WT1(Zf)/Kidney-WT1-ChIP-Seq(GSE90016)/Homer | 1e-8 | -1.930e+01 | 0.0000 | 2985.0 | 19.13% | 12578.8 | 17.17% | motif file (matrix) | svg |
| 451 | C G A T C T A G G A T C G C T A A G C T C T A G G A T C C G T A | RBFox2(?)/Heart-RBFox2-CLIP-Seq(GSE57926)/Homer | 1e-8 | -1.875e+01 | 0.0000 | 12048.0 | 77.20% | 54969.9 | 75.06% | motif file (matrix) | svg |
| 452 | G A C T G A C T T C G A C G T A C G A T A G C T C T G A A C T G T G A C G A C T T C G A C G T A A C G T A G C T C T G A C T G A G T C A G C T A G C T A C G T A | Pax7(Paired,Homeobox),longest/Myoblast-Pax7-ChIP-Seq(GSE25064)/Homer | 1e-8 | -1.867e+01 | 0.0000 | 133.0 | 0.85% | 339.1 | 0.46% | motif file (matrix) | svg |
| 453 | G A C T A G T C G A T C C G T A G T A C A G T C G C A T C G T A G T C A G A T C | MYB67(MYB)/col-MYB67-DAP-Seq(GSE60143)/Homer | 1e-7 | -1.836e+01 | 0.0000 | 8520.0 | 54.59% | 38174.8 | 52.12% | motif file (matrix) | svg |
| 454 | C G T A A G T C T G A C A G C T A C G T C G T A A C G T A G T C | At5g05790(MYBrelated)/col-At5g05790-DAP-Seq(GSE60143)/Homer | 1e-7 | -1.835e+01 | 0.0000 | 9531.0 | 61.07% | 42949.7 | 58.64% | motif file (matrix) | svg |
| 455 | T C G A T A G C G T C A A C T G C T A G C G T A C G A T A C T G A C G T A C T G A C T G A C G T | ETS:RUNX(ETS,Runt)/Jurkat-RUNX1-ChIP-Seq(GSE17954)/Homer | 1e-7 | -1.832e+01 | 0.0000 | 511.0 | 3.27% | 1803.4 | 2.46% | motif file (matrix) | svg |
| 456 | C A G T A G C T C G T A G C A T A G T C G A C T C T A G C T A G C A G T C T A G T C G A T G C A C T A G C A T G G A C T | STOP1(C2H2)/colamp-STOP1-DAP-Seq(GSE60143)/Homer | 1e-7 | -1.827e+01 | 0.0000 | 3370.0 | 21.59% | 14359.1 | 19.61% | motif file (matrix) | svg |
| 457 | A G C T G C A T G T C A C G A T T A G C C G T A A C G T G C T A | CRC(C2C2YABBY)/col-CRC-DAP-Seq(GSE60143)/Homer | 1e-7 | -1.752e+01 | 0.0000 | 10121.0 | 64.85% | 45796.8 | 62.53% | motif file (matrix) | svg |
| 458 | T C G A A C G T A C T G C G T A A G T C C T A G A G C T T G A C | TGA10(bZIP)/colamp-TGA10-DAP-Seq(GSE60143)/Homer | 1e-7 | -1.719e+01 | 0.0000 | 7536.0 | 48.29% | 33623.2 | 45.91% | motif file (matrix) | svg |
| 459 | G C A T G A C T A T G C A G C T T C G A C T A G G C T A C G T A C A T G T G A C G C A T G A C T A G T C A G C T C T G A | HSFB4(HSF)/col-HSFB4-DAP-Seq(GSE60143)/Homer | 1e-7 | -1.700e+01 | 0.0000 | 191.0 | 1.22% | 561.5 | 0.77% | motif file (matrix) | svg |
| 460 | G A C T T C G A C G T A C G T A C G T A C G T A C G T A C T A G A G C T C G T A | dof45(C2C2dof)/col-dof45-DAP-Seq(GSE60143)/Homer | 1e-7 | -1.614e+01 | 0.0000 | 11984.0 | 76.79% | 54791.8 | 74.81% | motif file (matrix) | svg |
| 461 | C G A T C T A G T C G A A G C T C G A T C T G A C G T A A G C T A C T G C T A G A T G C G A T C | Hoxb4(Homeobox)/ES-Hoxb4-ChIP-Seq(GSE34014)/Homer | 1e-6 | -1.609e+01 | 0.0000 | 2145.0 | 13.74% | 8943.3 | 12.21% | motif file (matrix) | svg |
| 462 | T A G C G C T A T C G A C T G A A G T C A G T C C T G A A G T C C G T A C T A G | RUNX(Runt)/HPC7-Runx1-ChIP-Seq(GSE22178)/Homer | 1e-6 | -1.606e+01 | 0.0000 | 5551.0 | 35.57% | 24457.3 | 33.39% | motif file (matrix) | svg |
| 463 | T A C G T A C G C T A G T C A G A G T C C G T A A T C G A T G C A C G T A C T G A G T C G A C T | Ascl2(bHLH)/ESC-Ascl2-ChIP-Seq(GSE97712)/Homer | 1e-6 | -1.592e+01 | 0.0000 | 5756.0 | 36.88% | 25413.8 | 34.70% | motif file (matrix) | svg |
| 464 | C T G A T C A G G C T A A G C T A G T C G A C T C T G A C T A G T G C A C T G A A G T C G T A C G A T C A C T G T C G A | ZBTB12(Zf)/HEK293-ZBTB12.GFP-ChIP-Seq(GSE58341)/Homer | 1e-6 | -1.560e+01 | 0.0000 | 3404.0 | 21.81% | 14637.3 | 19.99% | motif file (matrix) | svg |
| 465 | A G C T G C A T A C T G A C G T A G T C A C G T C T A G T A C G | Smad3(MAD)/NPC-Smad3-ChIP-Seq(GSE36673)/Homer | 1e-6 | -1.545e+01 | 0.0000 | 12571.0 | 80.55% | 57665.3 | 78.74% | motif file (matrix) | svg |
| 466 | A C T G C G T A A C G T C G T A C T G A A C T G T C A G G C A T | At3g11280(MYBrelated)/col-At3g11280-DAP-Seq(GSE60143)/Homer | 1e-6 | -1.532e+01 | 0.0000 | 9246.0 | 59.24% | 41776.6 | 57.04% | motif file (matrix) | svg |
| 467 | C T A G T A C G G A C T T G C A T G C A C G A T T A C G C T G A T C G A C T G A | Hoxa10(Homeobox)/ChickenMSG-Hoxa10.Flag-ChIP-Seq(GSE86088)/Homer | 1e-6 | -1.517e+01 | 0.0000 | 4212.0 | 26.99% | 18345.7 | 25.05% | motif file (matrix) | svg |
| 468 | A T G C G A C T A C T G C A G T G A T C A C G T T A C G T A C G | Smad2(MAD)/ES-SMAD2-ChIP-Seq(GSE29422)/Homer | 1e-6 | -1.516e+01 | 0.0000 | 10417.0 | 66.75% | 47340.1 | 64.64% | motif file (matrix) | svg |
| 469 | A T G C C T G A G A C T A C G T A C G T G T A C G A T C C G A T C T A G C A T G C G T A C G T A C T G A G A C T | STAT1(Stat)/HelaS3-STAT1-ChIP-Seq(GSE12782)/Homer | 1e-6 | -1.506e+01 | 0.0000 | 1635.0 | 10.48% | 6716.3 | 9.17% | motif file (matrix) | svg |
| 470 | A G T C G A T C G C T A C G A T A C G T T A C G G C A T C T G A G A C T A C T G A G T C G C T A C T G A T C G A C A G T | Oct4:Sox17(POU,Homeobox,HMG)/F9-Sox17-ChIP-Seq(GSE44553)/Homer | 1e-6 | -1.505e+01 | 0.0000 | 923.0 | 5.91% | 3606.1 | 4.92% | motif file (matrix) | svg |
| 471 | G T C A G C A T G C T A C A G T C T A G G A T C C G T A C T G A C G T A C G A T | Oct2(POU,Homeobox)/Bcell-Oct2-ChIP-Seq(GSE21512)/Homer | 1e-6 | -1.498e+01 | 0.0000 | 1821.0 | 11.67% | 7543.5 | 10.30% | motif file (matrix) | svg |
| 472 | C G T A C G T A C G T A C T G A C T A G A C G T C T A G G T C A | CDF3(C2C2dof)/colamp-CDF3-DAP-Seq(GSE60143)/Homer | 1e-6 | -1.455e+01 | 0.0000 | 9744.0 | 62.43% | 44181.7 | 60.33% | motif file (matrix) | svg |
| 473 | C G A T C T G A G T A C A C T G A C G T T C A G G C A T C G T A C G T A G C A T C G T A A G T C C G T A G T A C C A T G | CUC3(NAC)/col-CUC3-DAP-Seq(GSE60143)/Homer | 1e-6 | -1.444e+01 | 0.0000 | 4443.0 | 28.47% | 19444.5 | 26.55% | motif file (matrix) | svg |
| 474 | T A G C G A T C A G C T T G A C G C T A A G C T C A T G A C T G A C G T T C A G A G T C G A T C G A C T A G C T G C T A A G T C A G C T A G T C G A T C A T G C A G C T G A C T C A T G A C G T A T C G | ZNF41(Zf)/HEK293-ZNF41.GFP-ChIP-Seq(GSE58341)/Homer | 1e-6 | -1.406e+01 | 0.0000 | 142.0 | 0.91% | 408.1 | 0.56% | motif file (matrix) | svg |
| 475 | C T G A C T A G T C G A C G T A A T G C C G T A A T C G C G A T T A G C G C A T A T C G G C A T A G C T G A T C G A C T A G C T | ARE(NR)/LNCAP-AR-ChIP-Seq(GSE27824)/Homer | 1e-5 | -1.373e+01 | 0.0000 | 1406.0 | 9.01% | 5752.9 | 7.85% | motif file (matrix) | svg |
| 476 | C G T A C G T A C T A G A C G T A C G T C G T A A C T G A C T G A C G T C T G A T C G A T C G A | MYB4(MYB)/col200-MYB4-DAP-Seq(GSE60143)/Homer | 1e-5 | -1.324e+01 | 0.0000 | 5783.0 | 37.05% | 25700.6 | 35.09% | motif file (matrix) | svg |
| 477 | T C G A G C T A T G A C G C T A C T A G G A T C C G A T A C T G C G A T A G C T G A C T C T A G | E-box/Drosophila-Promoters/Homer | 1e-5 | -1.271e+01 | 0.0000 | 1396.0 | 8.94% | 5744.5 | 7.84% | motif file (matrix) | svg |
| 478 | C G T A C G T A G C T A C G T A G A T C C T G A A C G T A C G T A G T C A G C T G C A T G C A T | AT2G40260(G2like)/colamp-AT2G40260-DAP-Seq(GSE60143)/Homer | 1e-5 | -1.200e+01 | 0.0000 | 9645.0 | 61.80% | 43879.7 | 59.91% | motif file (matrix) | svg |
| 479 | C G A T C T G A G T A C C A T G G C A T T C A G G C A T C G T A C G T A G C T A C G T A A G T C G C T A G T A C C A T G | CUC2(NAC)/colamp-CUC2-DAP-Seq(GSE60143)/Homer | 1e-5 | -1.195e+01 | 0.0000 | 4333.0 | 27.76% | 19085.9 | 26.06% | motif file (matrix) | svg |
| 480 | T A C G C T G A T C G A C G A T C T A G C T A G T C G A C T G A T C G A T C G A C G T A T C G A G C A T C A T G C G T A T A C G G C A T T G A C C G T A A G C T | NFAT:AP1(RHD,bZIP)/Jurkat-NFATC1-ChIP-Seq(Jolma\_et\_al.)/Homer | 1e-5 | -1.168e+01 | 0.0000 | 781.0 | 5.00% | 3086.0 | 4.21% | motif file (matrix) | svg |
| 481 | T G C A A T G C A C G T A C G T A C G T A T G C C T A G A C G T A C G T A G C T G A T C A G C T | T1ISRE(IRF)/ThioMac-Ifnb-Expression/Homer | 1e-5 | -1.153e+01 | 0.0000 | 92.0 | 0.59% | 249.2 | 0.34% | motif file (matrix) | svg |
| 482 | C G T A G C A T C A G T C T A G A G T C A C T G A C T G G T A C A C T G A T C G | ERF115(AP2EREBP)/colamp-ERF115-DAP-Seq(GSE60143)/Homer | 1e-4 | -1.091e+01 | 0.0000 | 12858.0 | 82.39% | 59301.7 | 80.97% | motif file (matrix) | svg |
| 483 | G T A C A C T G A C G T T C A G G C A T C G T A C G A T G C A T C G T A A G T C C G T A T G A C C A T G G A C T G C T A | ANAC083(NAC)/col-ANAC083-DAP-Seq(GSE60143)/Homer | 1e-4 | -1.085e+01 | 0.0000 | 7837.0 | 50.21% | 35446.9 | 48.40% | motif file (matrix) | svg |
| 484 | C T A G T G A C G A C T A T C G T C G A A G T C C T A G C A G T C T A G A T C G G T A C T C G A | O2(bZIP)/Corn-O2-ChIP-Seq(GSE63991)/Homer | 1e-4 | -1.075e+01 | 0.0000 | 1649.0 | 10.57% | 6947.7 | 9.49% | motif file (matrix) | svg |
| 485 | G C A T G C T A C G T A A G C T G C T A T G C A A G T C A C G T A C G T A C G T G C A T G C A T | At4g38000(C2C2dof)/col-At4g38000-DAP-Seq(GSE60143)/Homer | 1e-4 | -1.063e+01 | 0.0001 | 6354.0 | 40.71% | 28531.6 | 38.96% | motif file (matrix) | svg |
| 486 | A T G C C G T A C G T A C G T A C G T A C G T A A C T G A C G T C G A T C T G A | dof43(C2C2dof)/colamp-dof43-DAP-Seq(GSE60143)/Homer | 1e-4 | -1.055e+01 | 0.0001 | 8788.0 | 56.31% | 39938.4 | 54.53% | motif file (matrix) | svg |
| 487 | G C T A T A G C A G C T A T C G G T C A C G T A G C T A A T G C G A T C C T G A | IRF4(IRF)/GM12878-IRF4-ChIP-Seq(GSE32465)/Homer | 1e-4 | -1.038e+01 | 0.0001 | 3852.0 | 24.68% | 16973.2 | 23.17% | motif file (matrix) | svg |
| 488 | G A C T G T A C G A C T A G T C T C A G C T G A A G T C A G T C C T A G C G A T A G C T A T G C C T G A C A G T A G C T | AT4G27900(C2C2COlike)/col-AT4G27900-DAP-Seq(GSE60143)/Homer | 1e-4 | -1.032e+01 | 0.0001 | 202.0 | 1.29% | 681.5 | 0.93% | motif file (matrix) | svg |
| 489 | T G C A G C A T A G C T G C A T A G T C A G T C A G T C C T G A A C T G T C G A T C G A C A G T A T C G A G T C G A T C | ZNF143\|STAF(Zf)/CUTLL-ZNF143-ChIP-Seq(GSE29600)/Homer | 1e-4 | -1.025e+01 | 0.0001 | 1358.0 | 8.70% | 5670.7 | 7.74% | motif file (matrix) | svg |
| 490 | C T G A G T A C G A C T A G T C C A G T T G C A C T G A A C G T A G C T G A T C C T A G C G A T A C T G A T G C G A C T C T G A G A T C G A C T A G C T G A T C | Mouse\_Recombination\_Hotspot(Zf)/Testis-DMC1-ChIP-Seq(GSE24438)/Homer | 1e-4 | -1.023e+01 | 0.0001 | 475.0 | 3.04% | 1812.6 | 2.47% | motif file (matrix) | svg |
| 491 | G A T C G C T A A G T C A C G T A C T G C G T A A G T C C G T A G C T A C G A T C A G T G C A T G C T A C G T A G C A T | GRF9(GRF)/colamp-GRF9-DAP-Seq(GSE60143)/Homer | 1e-4 | -1.009e+01 | 0.0001 | 6557.0 | 42.01% | 29518.6 | 40.30% | motif file (matrix) | svg |
| 492 | T G C A T A G C G A C T T G C A T G A C T G C A C G T A A G C T A G C T A G T C A G T C G T A C | GFY(?)/Promoter/Homer | 1e-4 | -9.842e+00 | 0.0001 | 505.0 | 3.24% | 1949.9 | 2.66% | motif file (matrix) | svg |
| 493 | T C G A G A C T A G C T T G A C A G C T G T A C T C A G G A T C A T C G T G C A A C T G C T G A | GFX(?)/Promoter/Homer | 1e-4 | -9.782e+00 | 0.0001 | 374.0 | 2.40% | 1396.3 | 1.91% | motif file (matrix) | svg |
| 494 | C A T G A G T C C T G A A T G C C T A G T C G A G C T A G C A T G A T C G A C T A G T C C T A G C G T A C A T G C T A G | PLT3(AP2EREBP)/col-PLT3-DAP-Seq(GSE60143)/Homer | 1e-4 | -9.643e+00 | 0.0001 | 1202.0 | 7.70% | 5001.2 | 6.83% | motif file (matrix) | svg |
| 495 | A G T C A T C G C T A G A G C T G A C T C T A G A G T C A G T C G C T A C A G T T C A G T C A G G A T C C T G A T C G A G A T C | RFX(HTH)/K562-RFX3-ChIP-Seq(SRA012198)/Homer | 1e-4 | -9.601e+00 | 0.0001 | 523.0 | 3.35% | 2032.2 | 2.77% | motif file (matrix) | svg |
| 496 | A T G C C G T A A C T G C G T A A C G T G C T A T C G A A G C T C G A T C G T A A C G T A G T C C G A T A C T G G A T C | GATA(Zf),IR4/iTreg-Gata3-ChIP-Seq(GSE20898)/Homer | 1e-3 | -9.085e+00 | 0.0002 | 852.0 | 5.46% | 3476.5 | 4.75% | motif file (matrix) | svg |
| 497 | A T G C G A C T A G C T A G C T A G T C G C T A C A G T C G A T G C T A A C G T A C T G G C T A T A G C G C A T T G A C | IRF:BATF(IRF:bZIP)/pDC-Irf8-ChIP-Seq(GSE66899)/Homer | 1e-3 | -8.836e+00 | 0.0003 | 577.0 | 3.70% | 2285.1 | 3.12% | motif file (matrix) | svg |
| 498 | C T A G T A C G A G T C C G T A A G T C A C G T A G T C T C G A C G T A T A C G | Nkx2.1(Homeobox)/LungAC-Nkx2.1-ChIP-Seq(GSE43252)/Homer | 1e-3 | -8.787e+00 | 0.0003 | 13352.0 | 85.55% | 61818.5 | 84.41% | motif file (matrix) | svg |
| 499 | C G T A A C G T A G C T C G A T C T A G G T A C C G T A A G C T C G T A G C T A | Oct4(POU,Homeobox)/mES-Oct4-ChIP-Seq(GSE11431)/Homer | 1e-3 | -8.660e+00 | 0.0004 | 2977.0 | 19.07% | 13074.8 | 17.85% | motif file (matrix) | svg |
| 500 | G A C T C T A G C T A G C T A G A C T G T C G A C T G A C T A G C T A G C T A G G T A C G T C A | ZNF467(Zf)/HEK293-ZNF467.GFP-ChIP-Seq(GSE58341)/Homer | 1e-3 | -8.545e+00 | 0.0004 | 3307.0 | 21.19% | 14593.0 | 19.93% | motif file (matrix) | svg |
| 501 | T A G C T C A G C A T G G C A T A G C T C G A T A T G C C G T A C G T A G T C A | CHR(?)/Hela-CellCycle-Expression/Homer | 1e-3 | -8.529e+00 | 0.0004 | 3591.0 | 23.01% | 15898.5 | 21.71% | motif file (matrix) | svg |
| 502 | C G T A C T G A C T A G C G T A C G T A A G T C C G T A C A G T G C A T G T C A C G A T A C T G A C G T G C A T G A T C | PGR(NR)/EndoStromal-PGR-ChIP-Seq(GSE69539)/Homer | 1e-3 | -8.405e+00 | 0.0005 | 1583.0 | 10.14% | 6759.3 | 9.23% | motif file (matrix) | svg |
| 503 | G C T A C G T A A C G T C A T G C G T A A C G T A C G T C T A G | ATHB5(HB)/colamp-ATHB5-DAP-Seq(GSE60143)/Homer | 1e-3 | -8.224e+00 | 0.0006 | 7129.0 | 45.68% | 32340.7 | 44.16% | motif file (matrix) | svg |
| 504 | G C T A G A C T A G T C T C G A T C A G T C G A A C G T A G T C G A C T T C A G | GATA14(C2C2gata)/col-GATA14-DAP-Seq(GSE60143)/Homer | 1e-3 | -8.036e+00 | 0.0007 | 6595.0 | 42.26% | 29862.0 | 40.77% | motif file (matrix) | svg |
| 505 | A C T G G A C T A G T C C T G A G A T C T C A G A T G C G A C T A G T C A T G C T A G C A G C T A T C G T G C A | PAX5(Paired,Homeobox),condensed/GM12878-PAX5-ChIP-Seq(GSE32465)/Homer | 1e-3 | -7.964e+00 | 0.0007 | 1108.0 | 7.10% | 4652.2 | 6.35% | motif file (matrix) | svg |
| 506 | A G C T G A T C A G C T G A C T G A C T T C G A A G T C C G T A A C T G T C A G | SpliceAcceptor/Homer | 1e-3 | -7.945e+00 | 0.0007 | 14591.0 | 93.49% | 67911.1 | 92.72% | motif file (matrix) | svg |
| 507 | A T C G A G C T A C T G A G T C A C T G A G T C C G T A A C G T A C T G A G T C A C T G A G T C | NRF(NRF)/Promoter/Homer | 1e-3 | -7.698e+00 | 0.0009 | 1735.0 | 11.12% | 7481.6 | 10.22% | motif file (matrix) | svg |
| 508 | C T A G A G C T G A C T C A T G A G T C A G T C G T C A C A G T C T A G T C A G G T A C C T G A T C G A G A T C T G A C | Rfx2(HTH)/LoVo-RFX2-ChIP-Seq(GSE49402)/Homer | 1e-3 | -7.607e+00 | 0.0010 | 589.0 | 3.77% | 2374.5 | 3.24% | motif file (matrix) | svg |
| 509 | T C G A C T G A C G T A C G T A C G T A C G T A A C T G A C G T C G A T C T G A | BBX31(Orphan)/col-BBX31-DAP-Seq(GSE60143)/Homer | 1e-3 | -7.597e+00 | 0.0010 | 9168.0 | 58.74% | 41971.2 | 57.31% | motif file (matrix) | svg |
| 510 | A G C T T G A C G A C T A G T C C T A G G A T C C T A G C T G A A C T G T C G A A G T C A G C T | BANP(?)/ESC-Banp-ChIP-Seq(GSE155603)/Homer | 1e-3 | -7.400e+00 | 0.0013 | 2921.0 | 18.72% | 12903.0 | 17.62% | motif file (matrix) | svg |
| 511 | T G C A T G C A A G T C A G T C G A C T C A G T A T G C G A T C C T G A A C G T C T A G C T A G A G T C A C G T A G T C A G T C A G T C G A C T C G T A A C G T A G C T C T A G G A T C G A T C G A T C | ZNF16(Zf)/HEK293-ZNF16.GFP-ChIP-Seq(GSE58341)/Homer | 1e-3 | -7.294e+00 | 0.0014 | 28.0 | 0.18% | 59.4 | 0.08% | motif file (matrix) | svg |
| 512 | C T A G C T A G T C G A C G T A A T G C C G T A A T C G T C G A T A C G G C A T A C T G C A G T T A G C G A T C G A C T | MRE(NR)/Neuro2A-NR3C2-ChIPnexus(GSE115417)/Homer | 1e-3 | -7.116e+00 | 0.0017 | 7829.0 | 50.16% | 35719.5 | 48.77% | motif file (matrix) | svg |
| 513 | G A T C G C A T T C G A A G T C A C G T A C G T A C G T C G A T A C G T A T C G | AT1G47655(C2C2dof)/colamp-AT1G47655-DAP-Seq(GSE60143)/Homer | 1e-3 | -7.084e+00 | 0.0017 | 13640.0 | 87.40% | 63319.9 | 86.46% | motif file (matrix) | svg |
| 514 | T G C A C T G A A T G C G T C A A C T G A C T G C G T A C G T A C T A G A G C T | Ets1-distal(ETS)/CD4+-PolII-ChIP-Seq(Barski\_et\_al.)/Homer | 1e-3 | -7.062e+00 | 0.0017 | 1416.0 | 9.07% | 6075.7 | 8.30% | motif file (matrix) | svg |
| 515 | C T A G A C T G A C G T C G T A A C T G C A T G G C A T T C A G | MYB92(MYB)/colamp-MYB92-DAP-Seq(GSE60143)/Homer | 1e-3 | -7.059e+00 | 0.0018 | 8081.0 | 51.78% | 36907.3 | 50.39% | motif file (matrix) | svg |
| 516 | C A T G T A C G T A G C G A T C G A T C A T G C G T A C G A C T T C A G A T G C C G A T A T C G C A G T A C T G G T A C | Zic3(Zf)/mES-Zic3-ChIP-Seq(GSE37889)/Homer | 1e-2 | -6.538e+00 | 0.0029 | 2527.0 | 16.19% | 11158.1 | 15.24% | motif file (matrix) | svg |
| 517 | G C T A A G C T G T A C G C A T A G C T T C G A C T G A A G T C A G T C T A C G A C G T G A C T T A C G C T A G C G T A | ZML1(C2C2gata)/colamp-ZML1-DAP-Seq(GSE60143)/Homer | 1e-2 | -6.455e+00 | 0.0032 | 539.0 | 3.45% | 2193.5 | 2.99% | motif file (matrix) | svg |
| 518 | G C A T T C G A C T G A G A T C A G T C G A T C G T C A G T C A A C G T A G T C C G T A C T G A | Duxbl(Homeobox)/NIH3T3-Duxbl.HA-ChIP-Seq(GSE119782)/Homer | 1e-2 | -6.446e+00 | 0.0032 | 714.0 | 4.57% | 2964.0 | 4.05% | motif file (matrix) | svg |
| 519 | C G T A C G T A T C G A C G T A C G A T C G T A A C G T A G T C G C A T G C A T | At3g09600(MYBrelated)/colamp-At3g09600-DAP-Seq(GSE60143)/Homer | 1e-2 | -6.192e+00 | 0.0041 | 3090.0 | 19.80% | 13768.5 | 18.80% | motif file (matrix) | svg |
| 520 | G A T C G T A C C G T A A G C T G A C T G C T A C T G A A C G T G A T C G C T A | Hoxc6(Homeobox)/EB-Hoxc6.iFlag-ChIP-Seq(GSE142377)/Homer | 1e-2 | -6.187e+00 | 0.0041 | 13321.0 | 85.35% | 61843.7 | 84.44% | motif file (matrix) | svg |
| 521 | G T A C C T G A A G T C A G T C A C T G G T C A G A T C G C A T | At1g75490(AP2EREBP)/colamp-At1g75490-DAP-Seq(GSE60143)/Homer | 1e-2 | -6.146e+00 | 0.0043 | 14050.0 | 90.02% | 65365.5 | 89.25% | motif file (matrix) | svg |
| 522 | T G A C C T G A A G T C A G T C A C T G G A T C G A C T G C A T | At5g18450(AP2EREBP)/col-At5g18450-DAP-Seq(GSE60143)/Homer | 1e-2 | -6.034e+00 | 0.0048 | 11886.0 | 76.16% | 54991.3 | 75.08% | motif file (matrix) | svg |
| 523 | C A G T A G C T G A C T T G C A A G T C A G C T A C G T A C G T C G A T G A C T | AT3G52440(C2C2dof)/colamp-AT3G52440-DAP-Seq(GSE60143)/Homer | 1e-2 | -5.858e+00 | 0.0057 | 11383.0 | 72.94% | 52616.0 | 71.84% | motif file (matrix) | svg |
| 524 | C G A T T A C G G A T C G C T A T C A G G A C T C G A T G T A C G A T C G T C A T C G A T G A C C G T A C T A G G A C T C T A G C T A G G T A C A G T C C G T A | CTCF-SatelliteElement(Zf?)/CD4+-CTCF-ChIP-Seq(Barski\_et\_al.)/Homer | 1e-2 | -5.780e+00 | 0.0062 | 91.0 | 0.58% | 303.9 | 0.41% | motif file (matrix) | svg |
| 525 | G C A T A C G T A G T C A T G C A G T C C T A G T A G C G T A C C T G A G C T A | DEL1(E2FDP)/colamp-DEL1-DAP-Seq(GSE60143)/Homer | 1e-2 | -5.637e+00 | 0.0071 | 67.0 | 0.43% | 211.6 | 0.29% | motif file (matrix) | svg |
| 526 | A T C G A G T C A G T C G A C T A T G C C T G A C T A G A C T G T A C G G T A C C T G A C G A T | AP-2gamma(AP2)/MCF7-TFAP2C-ChIP-Seq(GSE21234)/Homer | 1e-2 | -5.435e+00 | 0.0087 | 5319.0 | 34.08% | 24160.0 | 32.99% | motif file (matrix) | svg |
| 527 | T G A C G T A C C G T A A C T G T G A C C G A T A C T G A T C G A G C T T A C G T C G A T A G C G T A C C G T A A T C G T G A C G C A T A C T G A C T G A T G C | Twist(bHLH)/HMLE-TWIST1-ChIP-Seq(Chang\_et\_al)/Homer | 1e-2 | -5.398e+00 | 0.0090 | 526.0 | 3.37% | 2173.2 | 2.97% | motif file (matrix) | svg |
| 528 | C T A G A T G C A T G C C G A T A C T G G A C T A T G C G C T A T G A C A G C T T A G C G C T A | PBX1(Homeobox)/MCF7-PBX1-ChIP-Seq(GSE28007)/Homer | 1e-2 | -5.151e+00 | 0.0115 | 406.0 | 2.60% | 1654.8 | 2.26% | motif file (matrix) | svg |
| 529 | C T A G G A C T G A T C A C G T A T C G A G C T C T G A A T C G C G A T C T A G G A T C G A C T C A T G A T C G G T A C G A C T A G T C G C A T A G C T C G A T | ZNF382(Zf)/HEK293-ZNF382.GFP-ChIP-Seq(GSE58341)/Homer | 1e-2 | -5.141e+00 | 0.0116 | 178.0 | 1.14% | 671.6 | 0.92% | motif file (matrix) | svg |
| 530 | T A G C G T A C C T A G C A G T T C G A C G T A C G T A G C A T G A C T T G A C A G T C A C T G A T C G A G T C C T A G | AS2(LOBAS2)/col-AS2-DAP-Seq(GSE60143)/Homer | 1e-2 | -5.131e+00 | 0.0117 | 1435.0 | 9.19% | 6271.6 | 8.56% | motif file (matrix) | svg |
| 531 | C A G T G A C T G C A T T C G A A G T C A C G T A C G T A C G T C G A T G A C T | OBP3(C2C2dof)/col-OBP3-DAP-Seq(GSE60143)/Homer | 1e-2 | -5.053e+00 | 0.0126 | 12827.0 | 82.19% | 59568.2 | 81.33% | motif file (matrix) | svg |
| 532 | G C T A G C T A T G C A A G T C C T A G C T G A G A T C C T A G G A C T G A T C C T A G A C G T C G A T C G A T G A C T | Unknown2/Arabidopsis-Promoters/Homer | 1e-2 | -4.746e+00 | 0.0171 | 273.0 | 1.75% | 1087.5 | 1.48% | motif file (matrix) | svg |
