## Supplemental dataset for "Hybrid CNN and Multi-Head Attention Model for Analyzing Epigenetic Mechanisms and Gene Expression Across Fungal Phylogenetic Distances": NcrassaModel_FgramTest_K4me3locs_homerResults.html

/projects/wg-feeds/SHAP/NcrassaModel\_FgramTest\_K4me3locs\_SHAP\_noDup\_HOMER// - Homer de novo Motif Results


### Homer *de novo* Motif Results (/projects/wg-feeds/SHAP/NcrassaModel\_FgramTest\_K4me3locs\_SHAP\_noDup\_HOMER//)

Non-redundant Motif File of Results  
Known Motif Enrichment Results  
Gene Ontology Enrichment Results  
If Homer is having trouble matching a motif to a known motif, try copy/pasting the matrix file into
STAMP  
More information on motif finding results: HOMER
| Description of Results
| Tips
  
Total target sequences = 14965  
Total background sequences = 69620  
\* - possible false positive  

|  |  |  |  |  |  |  |  |  |
| --- | --- | --- | --- | --- | --- | --- | --- | --- |
| Rank | Motif | P-value | log P-pvalue | % of Targets | % of Background | STD(Bg STD) | Best Match/Details | Motif File |
| 1 | A G T C G C A T A G C T A T G C G C A T A G C T A T G C G C A T A G C T A T G C G C A T A G C T | 1e-2226 | -5.127e+03 | 59.35% | 17.37% | 689.2bp (733.3bp) | Unknown4/Arabidopsis-Promoters/Homer(0.848) More Information \| Similar Motifs Found | motif file (matrix) |
| 2 | A T C G C G A T A G C T A T C G C G T A A G C T A T G C C G A T A G C T A C T G | 1e-2129 | -4.904e+03 | 63.81% | 21.47% | 666.4bp (717.4bp) | CG4360/MA2204.1/Jaspar(0.790) More Information \| Similar Motifs Found | motif file (matrix) |
| 3 | C G A T A G T C T A C G C G A T A G T C T A C G C G A T A G T C T A C G C G A T A G T C T C A G | 1e-1905 | -4.388e+03 | 62.31% | 22.20% | 660.5bp (723.7bp) | SRSF7(RRM,Znf)/Homo\_sapiens-RNCMPT00073-PBM/HughesRNA(0.637) More Information \| Similar Motifs Found | motif file (matrix) |
| 4 | T A G C A C G T A G C T T C A G C T G A T A G C T C G A C T G A A T C G T G A C C G A T A G T C | 1e-1732 | -3.990e+03 | 59.16% | 21.27% | 669.3bp (726.8bp) | RIM101/MA0368.1/Jaspar(0.644) More Information \| Similar Motifs Found | motif file (matrix) |
| 5 | A T G C T C G A C G T A A T C G T C G A C G T A A T C G T G C A G C A T A T G C | 1e-1615 | -3.720e+03 | 69.42% | 31.26% | 677.1bp (731.2bp) | REF2(RRM)/Drosophila\_melanogaster-RNCMPT00059-PBM/HughesRNA(0.780) More Information \| Similar Motifs Found | motif file (matrix) |
| 6 | A G C T A G T C A C G T A G C T C A T G A G T C A G T C A G T C A C G T A G T C | 1e-1368 | -3.151e+03 | 74.42% | 39.21% | 677.1bp (730.0bp) | SeqBias: GA-repeat(0.777) More Information \| Similar Motifs Found | motif file (matrix) |
| 7 | C T A G C T A G G A T C T A G C C G T A C G A T A T C G C G T A G C A T A C T G T C A G G T A C | 1e-1266 | -2.915e+03 | 62.80% | 29.23% | 656.2bp (720.5bp) | Tv\_0259(RRM)/Trichomonas\_vaginalis-RNCMPT00259-PBM/HughesRNA(0.716) More Information \| Similar Motifs Found | motif file (matrix) |
| 8 | A G C T A G C T A G C T A G C T A G C T A G C T A G C T A G C T A G C T A G C T A G C T A G C T | 1e-1187 | -2.735e+03 | 38.89% | 11.85% | 693.9bp (713.9bp) | SeqBias: polyA-repeat(0.891) More Information \| Similar Motifs Found | motif file (matrix) |
| 9 | C G T A G A T C G T A C C G T A G A T C T G A C C G T A G T A C T G A C C T G A G T A C G T A C | 1e-1161 | -2.675e+03 | 61.69% | 29.53% | 673.7bp (704.4bp) | DREB2F/MA1242.1/Jaspar(0.769) More Information \| Similar Motifs Found | motif file (matrix) |
| 10 | T C A G G C A T A T G C C T G A C G T A A T C G T A G C G C A T A G C T A T C G | 1e-1114 | -2.565e+03 | 60.09% | 28.72% | 670.3bp (728.3bp) | Nr2e3/MA0164.2/Jaspar(0.780) More Information \| Similar Motifs Found | motif file (matrix) |
| 11 | T C A G T G C A C A G T C T A G A C G T A G T C G T A C G C A T A G C T T C A G | 1e-913 | -2.103e+03 | 74.31% | 45.67% | 667.6bp (722.2bp) | ttk/dmmpmm(Pollard)/fly(0.668) More Information \| Similar Motifs Found | motif file (matrix) |
| 12 | A T G C C T G A T C A G T A G C T C G A T C A G T A G C C T G A T A C G T A G C C T G A C T A G | 1e-888 | -2.045e+03 | 67.61% | 39.06% | 668.1bp (718.1bp) | GRF4/MA1815.2/Jaspar(0.780) More Information \| Similar Motifs Found | motif file (matrix) |
| 13 | T C A G G T C A T G A C T C G A C T G A T A C G G T C A T G A C T G C A C G T A T A C G T G C A | 1e-875 | -2.017e+03 | 65.90% | 37.55% | 654.8bp (713.3bp) | ARF16/MA1688.2/Jaspar(0.877) More Information \| Similar Motifs Found | motif file (matrix) |
| 14 | G T A C C T A G T G A C T A G C C G T A A C G T A C G T A C T G G A C T A T G C T C G A G C T A | 1e-874 | -2.015e+03 | 73.90% | 45.85% | 676.5bp (723.7bp) | pho/dmmpmm(Bergman)/fly(0.755) More Information \| Similar Motifs Found | motif file (matrix) |
| 15 | A T C G G T C A C G A T T C A G A G C T A G T C C T A G C G T A | 1e-864 | -1.990e+03 | 79.59% | 52.49% | 672.1bp (721.4bp) | DDF2(AP2EREBP)/col-DDF2-DAP-Seq(GSE60143)/Homer(0.697) More Information \| Similar Motifs Found | motif file (matrix) |
| 16 | G A T C A C G T A G T C A G C T A C G T A G T C T G A C G T C A | 1e-791 | -1.823e+03 | 70.93% | 44.07% | 664.9bp (729.9bp) | ZNF189(Zf)/HEK293-ZNF189.GFP-ChIP-Seq(GSE58341)/Homer(0.783) More Information \| Similar Motifs Found | motif file (matrix) |
| 17 | A G T C G T C A G C A T A T G C A C G T A G T C T A G C G T C A | 1e-701 | -1.614e+03 | 83.03% | 59.62% | 669.2bp (717.3bp) | Pp\_0237(RRM)/Physcomitrella\_patens-RNCMPT00237-PBM/HughesRNA(0.705) More Information \| Similar Motifs Found | motif file (matrix) |
| 18 | G C A T A C G T C T A G C G A T A G C T T C G A A C T G G A T C T C G A G C T A | 1e-675 | -1.556e+03 | 41.56% | 19.28% | 669.2bp (716.6bp) | Aef1/dmmpmm(Bergman)/fly(0.714) More Information \| Similar Motifs Found | motif file (matrix) |
| 19 | G A C T A C G T A T C G A C G T A C G T C A G T A T C G G A C T G C A T A C T G | 1e-667 | -1.536e+03 | 58.61% | 34.03% | 671.4bp (718.4bp) | PB0093.1\_Zfp105\_1/Jaspar(0.748) More Information \| Similar Motifs Found | motif file (matrix) |
| 20 | A G T C C T A G C T A G G A T C C T A G C A T G A G T C C T A G | 1e-339 | -7.828e+02 | 48.01% | 30.86% | 678.8bp (697.0bp) | Os05g0497200/MA1034.1/Jaspar(0.917) More Information \| Similar Motifs Found | motif file (matrix) |
| 21 | A C G T C A G T A G C T A G C T A T G C G A C T C T A G G T C A A G C T T C G A A G T C G A T C | 1e-90 | -2.073e+02 | 1.51% | 0.15% | 702.4bp (741.9bp) | Six4/dmmpmm(Noyes\_hd)/fly(0.734) More Information \| Similar Motifs Found | motif file (matrix) |
| 22 | C A T G T C A G G C A T C G T A A T G C T A C G C G A T G C T A A G T C G A T C C A G T A C G T | 1e-85 | -1.971e+02 | 2.40% | 0.52% | 760.4bp (785.6bp) | Zm00001d018571/MA1822.2/Jaspar(0.887) More Information \| Similar Motifs Found | motif file (matrix) |
