## Supplemental dataset for "Hybrid CNN and Multi-Head Attention Model for Analyzing Epigenetic Mechanisms and Gene Expression Across Fungal Phylogenetic Distances": NcrassaModel_FgramTest_K4me3locs_knownResults.html

/projects/wg-feeds/SHAP/NcrassaModel\_FgramTest\_K4me3locs\_SHAP\_noDup\_HOMER/ - Homer Known Motif Enrichment Results


### Homer Known Motif Enrichment Results (/projects/wg-feeds/SHAP/NcrassaModel\_FgramTest\_K4me3locs\_SHAP\_noDup\_HOMER/)

Homer *de novo* Motif Results  
Gene Ontology Enrichment Results  
Known Motif Enrichment Results (txt file)  
Total Target Sequences = 14967, Total Background Sequences = 69613

|  |  |  |  |  |  |  |  |  |  |  |  |
| --- | --- | --- | --- | --- | --- | --- | --- | --- | --- | --- | --- |
| Rank | Motif | Name | P-value | log P-pvalue | q-value (Benjamini) | # Target Sequences with Motif | % of Targets Sequences with Motif | # Background Sequences with Motif | % of Background Sequences with Motif | Motif File | SVG |
| 1 | T A G C G T C A G A C T T A G C G T C A G A C T A G T C G C T A G A C T G A T C | ZML2(C2C2gata)/col-ZML2-DAP-Seq(GSE60143)/Homer | 1e-1256 | -2.894e+03 | 0.0000 | 5216.0 | 34.85% | 6195.6 | 8.90% | motif file (matrix) | svg |
| 2 | C T A G T C A G C T G A C T G A C T A G C T G A C A T G C A T G C T G A C T A G C T A G C G T A C T A G C G T A G T C A | TF3A(C2H2)/col-TF3A-DAP-Seq(GSE60143)/Homer | 1e-1082 | -2.492e+03 | 0.0000 | 8794.0 | 58.76% | 19471.0 | 27.96% | motif file (matrix) | svg |
| 3 | A T G C C A G T A C G T A G T C C A T G A C G T A G T C A C G T A G C T G A T C | Unknown4/Arabidopsis-Promoters/Homer | 1e-1076 | -2.478e+03 | 0.0000 | 10590.0 | 70.77% | 27439.9 | 39.41% | motif file (matrix) | svg |
| 4 | T G A C C G T A C T G A A C T G A C T G G A C T G A T C T G C A G T A C T A C G | SF1(NR)/H295R-Nr5a1-ChIP-Seq(GSE44220)/Homer | 1e-737 | -1.698e+03 | 0.0000 | 5458.0 | 36.47% | 10222.6 | 14.68% | motif file (matrix) | svg |
| 5 | A C G T G A C T A T G C G C T A C T G A C T A G A C T G G A C T A G T C C G T A | Nr5a2(NR)/mES-Nr5a2-ChIP-Seq(GSE19019)/Homer | 1e-647 | -1.491e+03 | 0.0000 | 6127.0 | 40.94% | 13359.3 | 19.19% | motif file (matrix) | svg |
| 6 | A C G T G A C T T A G C C G T A C T G A C A T G C T A G G A C T G A T C C G T A | Nr5a2(NR)/Pancreas-LRH1-ChIP-Seq(GSE34295)/Homer | 1e-630 | -1.453e+03 | 0.0000 | 7341.0 | 49.05% | 18162.5 | 26.08% | motif file (matrix) | svg |
| 7 | G C A T G A T C C T A G C G T A G C A T C G T A G C A T A G T C C T A G C G T A G C A T C G A T | AT5G22990(C2H2)/col-AT5G22990-DAP-Seq(GSE60143)/Homer | 1e-513 | -1.183e+03 | 0.0000 | 8570.0 | 57.27% | 24820.7 | 35.64% | motif file (matrix) | svg |
| 8 | A C G T A T G C A C G T A C G T A G C T A G T C A G C T A G C T A G C T A G C T A G C T | hTCT(CPE) | 1e-443 | -1.022e+03 | 0.0000 | 10792.0 | 72.11% | 36420.3 | 52.30% | motif file (matrix) | svg |
| 9 | G C A T A G C T A C G T A C G T A C T G A C G T G A T C A C G T A C G T A G C T C G A T G C A T A G T C G A C T C A G T | IDD5(C2H2)/colamp-IDD5-DAP-Seq(GSE60143)/Homer | 1e-429 | -9.900e+02 | 0.0000 | 4525.0 | 30.24% | 9929.6 | 14.26% | motif file (matrix) | svg |
| 10 | C A T G A G C T T A C G G T C A G T A C T A G C A G C T G A C T A T C G T C G A | Esrrb(NR)/mES-Esrrb-ChIP-Seq(GSE11431)/Homer | 1e-377 | -8.696e+02 | 0.0000 | 6762.0 | 45.19% | 19121.7 | 27.46% | motif file (matrix) | svg |
| 11 | A C G T A C T G C G T A A C G T A C T G A C T G C G T A C G T A | HAP3(CCAATHAP3)/col-HAP3-DAP-Seq(GSE60143)/Homer | 1e-356 | -8.204e+02 | 0.0000 | 4745.0 | 31.71% | 11573.7 | 16.62% | motif file (matrix) | svg |
| 12 | G C T A G C A T A C G T A C G T A G C T G A T C G A C T G A C T A G C T A C G T A C G T A G C T | RLR1?/SacCer-Promoters/Homer | 1e-326 | -7.507e+02 | 0.0000 | 2122.0 | 14.18% | 3307.1 | 4.75% | motif file (matrix) | svg |
| 13 | G A T C G C A T A G T C A G C T G A T C A G C T G A T C G A C T G A T C G A C T A G T C A C G T G A T C A G C T G A T C | GAGA-repeat/SacCer-Promoters/Homer | 1e-323 | -7.460e+02 | 0.0000 | 12942.0 | 86.48% | 50263.0 | 72.18% | motif file (matrix) | svg |
| 14 | G A C T A G C T A G C T C T A G A C G T G A T C A C G T A C G T G A C T C G A T G C A T A G T C | IDD4(C2H2)/col-IDD4-DAP-Seq(GSE60143)/Homer | 1e-322 | -7.422e+02 | 0.0000 | 6033.0 | 40.31% | 17010.4 | 24.43% | motif file (matrix) | svg |
| 15 | T G A C C T A G T C A G G T C A C G T A T C A G C G A T T C A G T C G A T G C A C T G A T A G C | PU.1-IRF(ETS:IRF)/Bcell-PU.1-ChIP-Seq(GSE21512)/Homer | 1e-321 | -7.406e+02 | 0.0000 | 6942.0 | 46.39% | 20764.2 | 29.82% | motif file (matrix) | svg |
| 16 | T C G A T G C A C A G T T C G A G A T C A G T C C G T A C G T A A C T G A G T C C G T A C G T A T C A G C G A T A G T C | AT5G25475(ABI3VP1)/col-AT5G25475-DAP-Seq(GSE60143)/Homer | 1e-307 | -7.072e+02 | 0.0000 | 9865.0 | 65.92% | 34252.6 | 49.19% | motif file (matrix) | svg |
| 17 | C T G A C G A T C T A G T C A G G A T C C T G A T C A G G A T C C T G A A C T G A G T C G C T A A C G T A G T C G C A T | PRDM9(Zf)/Testis-DMC1-ChIP-Seq(GSE35498)/Homer | 1e-306 | -7.057e+02 | 0.0000 | 3706.0 | 24.76% | 8483.6 | 12.18% | motif file (matrix) | svg |
| 18 | C A T G G A C T T A C G G T C A G T A C G A T C G A C T A G C T A T C G T C G A T A C G T A G C | ERRg(NR)/Kidney-ESRRG-ChIP-Seq(GSE104905)/Homer | 1e-295 | -6.797e+02 | 0.0000 | 7539.0 | 50.38% | 23791.9 | 34.17% | motif file (matrix) | svg |
| 19 | C T G A C G A T C A T G A T C G G C A T C A T G G C T A A G T C | ASHR1(ND)/col-ASHR1-DAP-Seq(GSE60143)/Homer | 1e-285 | -6.577e+02 | 0.0000 | 9704.0 | 64.84% | 33899.8 | 48.68% | motif file (matrix) | svg |
| 20 | A G T C A G T C C G A T A C G T A C G T A C T G A C G T A G C T A G T C A G T C | Sox4(HMG)/proB-Sox4-ChIP-Seq(GSE50066)/Homer | 1e-270 | -6.222e+02 | 0.0000 | 6756.0 | 45.15% | 20885.6 | 29.99% | motif file (matrix) | svg |
| 21 | C A T G G C T A C T A G T A C G C G T A T C A G C G T A A C T G C G T A C A T G C T G A C G T A | BPC1(BBRBPC)/colamp-BPC1-DAP-Seq(GSE60143)/Homer | 1e-253 | -5.842e+02 | 0.0000 | 3615.0 | 24.16% | 8828.7 | 12.68% | motif file (matrix) | svg |
| 22 | G A C T G C A T A C G T A C G T A C T G C G T A A G T C A G C T C G A T A T C G G C A T A C T G C G A T C T A G C G T A | WRKY50(WRKY)/col-WRKY50-DAP-Seq(GSE60143)/Homer | 1e-241 | -5.559e+02 | 0.0000 | 9419.0 | 62.94% | 33460.2 | 48.05% | motif file (matrix) | svg |
| 23 | T C G A A C T G C A T G A G C T A G T C C G T A C T G A C T A G A C T G C G A T A T G C C T G A | RAR:RXR(NR),DR0/ES-RAR-ChIP-Seq(GSE56893)/Homer | 1e-233 | -5.380e+02 | 0.0000 | 1682.0 | 11.24% | 2777.8 | 3.99% | motif file (matrix) | svg |
| 24 | A T G C A G T C G C A T A G C T A C G T T C A G C G A T A G C T G A T C A T C G | Sox10(HMG)/SciaticNerve-Sox3-ChIP-Seq(GSE35132)/Homer | 1e-232 | -5.362e+02 | 0.0000 | 10508.0 | 70.22% | 38963.8 | 55.96% | motif file (matrix) | svg |
| 25 | A G C T A C G T A C T G A T G C A G T C C G T A C T G A T A C G | NF1-halfsite(CTF)/LNCaP-NF1-ChIP-Seq(Unpublished)/Homer | 1e-229 | -5.277e+02 | 0.0000 | 10416.0 | 69.60% | 38587.6 | 55.42% | motif file (matrix) | svg |
| 26 | A T G C G A T C C G A T A C G T A C G T A C T G C A G T A G C T | Sox3(HMG)/NPC-Sox3-ChIP-Seq(GSE33059)/Homer | 1e-224 | -5.172e+02 | 0.0000 | 10919.0 | 72.96% | 41222.3 | 59.20% | motif file (matrix) | svg |
| 27 | C T A G A G T C A G T C A C T G C G T A A G T C C T G A G A C T | DDF1(AP2EREBP)/col-DDF1-DAP-Seq(GSE60143)/Homer | 1e-222 | -5.114e+02 | 0.0000 | 9498.0 | 63.47% | 34269.6 | 49.21% | motif file (matrix) | svg |
| 28 | G T C A G C T A G C T A T C G A A T C G A C G T A G T C T C G A T C G A T G A C | WRKY40(WRKY)/colamp-WRKY40-DAP-Seq(GSE60143)/Homer | 1e-214 | -4.935e+02 | 0.0000 | 6680.0 | 44.64% | 21660.4 | 31.11% | motif file (matrix) | svg |
| 29 | G A C T A C G T A C G T A C T G A C G T A G T C G C A T A G C T G C A T G C A T G A C T A G C T | SGR5(C2H2)/colamp-SGR5-DAP-Seq(GSE60143)/Homer | 1e-208 | -4.793e+02 | 0.0000 | 5302.0 | 35.43% | 16027.2 | 23.02% | motif file (matrix) | svg |
| 30 | C G A T C T A G C G T A G A C T C A G T C T A G C G T A A G C T C A T G C T A G | HOXA1(Homeobox)/mES-Hoxa1-ChIP-Seq(SRP084292)/Homer | 1e-204 | -4.704e+02 | 0.0000 | 3767.0 | 25.17% | 10104.2 | 14.51% | motif file (matrix) | svg |
| 31 | A C G T A C G T A C G T A C G T A C G T A C G T A C G T A C G T A C G T A C G T | VRN1(ABI3VP1)/col-VRN1-DAP-Seq(GSE60143)/Homer | 1e-197 | -4.544e+02 | 0.0000 | 471.0 | 3.15% | 200.6 | 0.29% | motif file (matrix) | svg |
| 32 | A T G C T C G A T A C G A C G T A T G C A G T C A C G T A G T C A G T C G A T C | Znf263(Zf)/K562-Znf263-ChIP-Seq(GSE31477)/Homer | 1e-194 | -4.470e+02 | 0.0000 | 9123.0 | 60.96% | 33144.3 | 47.60% | motif file (matrix) | svg |
| 33 | C A T G G A C T C T A G C A T G C A G T C G A T C T A G C A T G C G A T C G T A C T A G C A G T C G A T C T A G C A T G | AT1G24250(Orphan)/col-AT1G24250-DAP-Seq(GSE60143)/Homer | 1e-192 | -4.427e+02 | 0.0000 | 3569.0 | 23.85% | 9558.5 | 13.73% | motif file (matrix) | svg |
| 34 | A G T C G A T C G C T A C G A T C A G T T A C G C G A T A G C T A G T C A T C G | SOX1(HMG)/NPC-SOX1-ChIP-Seq(GSE138215)/Homer | 1e-185 | -4.272e+02 | 0.0000 | 11502.0 | 76.86% | 45144.0 | 64.83% | motif file (matrix) | svg |
| 35 | A G T C T G C A T C G A C T G A A C T G C A T G A C G T A T G C G T C A T A C G | Erra(NR)/HepG2-Erra-ChIP-Seq(GSE31477)/Homer | 1e-184 | -4.257e+02 | 0.0000 | 10535.0 | 70.40% | 40228.8 | 57.77% | motif file (matrix) | svg |
| 36 | A G C T A G C T A G C T A C T G A C G T A G T C A C T G A C G T G A C T C G A T G C A T A T C G | IDD7(C2H2)/col-IDD7-DAP-Seq(GSE60143)/Homer | 1e-182 | -4.212e+02 | 0.0000 | 4010.0 | 26.80% | 11385.9 | 16.35% | motif file (matrix) | svg |
| 37 | C A T G G T A C A C T G G T C A A G C T T A C G T G C A A T C G T G A C C A G T | TOD6?/SacCer-Promoters/Homer | 1e-179 | -4.126e+02 | 0.0000 | 3172.0 | 21.20% | 8303.4 | 11.92% | motif file (matrix) | svg |
| 38 | G T A C A C G T A C G T A T C G C A G T C G A T A T C G G C T A T G C A T A G C C G T A G T C A C A T G A G C T G C T A | ANAC013(NAC)/col-ANAC013-DAP-Seq(GSE60143)/Homer | 1e-178 | -4.117e+02 | 0.0000 | 4887.0 | 32.66% | 14924.1 | 21.43% | motif file (matrix) | svg |
| 39 | C T G A T C A G A G T C C G T A A T C G A T G C C G A T A C T G A G T C G A C T A T C G A G T C | MyoD(bHLH)/Myotube-MyoD-ChIP-Seq(GSE21614)/Homer | 1e-176 | -4.075e+02 | 0.0000 | 4342.0 | 29.01% | 12786.6 | 18.36% | motif file (matrix) | svg |
| 40 | C G T A T G A C T A G C T G C A A C T G A C T G C G T A C G T A T C A G G A C T | ELF3(ETS)/PDAC-ELF3-ChIP-Seq(GSE64557)/Homer | 1e-168 | -3.878e+02 | 0.0000 | 4669.0 | 31.20% | 14256.0 | 20.47% | motif file (matrix) | svg |
| 41 | C G T A T A G C T A G C T G C A A C T G C T A G C G T A C G T A T C A G G A C T | EHF(ETS)/LoVo-EHF-ChIP-Seq(GSE49402)/Homer | 1e-167 | -3.867e+02 | 0.0000 | 7806.0 | 52.16% | 27706.6 | 39.79% | motif file (matrix) | svg |
| 42 | A G T C G A T C C T G A A G T C A G T C C A T G G T C A G A T C C G T A G A T C | DREB26(AP2EREBP)/col-DREB26-DAP-Seq(GSE60143)/Homer | 1e-166 | -3.833e+02 | 0.0000 | 3995.0 | 26.70% | 11628.1 | 16.70% | motif file (matrix) | svg |
| 43 | T A C G G A C T T G A C C G T A A C G T G A T C G T C A C G T A A C G T A T G C C G T A G A C T | HOXA2(Homeobox)/mES-Hoxa2-ChIP-Seq(Donaldson\_et\_al.)/Homer | 1e-165 | -3.817e+02 | 0.0000 | 1812.0 | 12.11% | 3801.9 | 5.46% | motif file (matrix) | svg |
| 44 | C T G A T C G A C G T A A T G C C G T A C G T A C G A T C T A G T C A G G A T C | Sox15(HMG)/CPA-Sox15-ChIP-Seq(GSE62909)/Homer | 1e-162 | -3.736e+02 | 0.0000 | 7512.0 | 50.20% | 26525.7 | 38.09% | motif file (matrix) | svg |
| 45 | T A G C C G T A C T G A T A C G C G T A A C G T A C T G A C T G A G T C T A C G C T A G G T A C | YY1(Zf)/Promoter/Homer | 1e-162 | -3.732e+02 | 0.0000 | 1012.0 | 6.76% | 1487.1 | 2.14% | motif file (matrix) | svg |
| 46 | A G T C C A T G A C G T A C G T A C T G C G T A A G T C G A C T G C A T G C T A | WRKY28(WRKY)/col-WRKY28-DAP-Seq(GSE60143)/Homer | 1e-160 | -3.696e+02 | 0.0000 | 10247.0 | 68.47% | 39430.5 | 56.63% | motif file (matrix) | svg |
| 47 | C T A G G T A C C A T G G A C T C G A T C A T G G T C A G T A C G A C T C G A T C G A T C G A T | WRKY27(WRKY)/colamp-WRKY27-DAP-Seq(GSE60143)/Homer | 1e-160 | -3.689e+02 | 0.0000 | 8199.0 | 54.79% | 29699.2 | 42.65% | motif file (matrix) | svg |
| 48 | C G T A C T A G C A T G A G C T C T G A C A T G C A G T C G A T C T A G C T A G | MYB30(MYB)/colamp-MYB30-DAP-Seq(GSE60143)/Homer | 1e-157 | -3.619e+02 | 0.0000 | 8980.0 | 60.01% | 33415.2 | 47.99% | motif file (matrix) | svg |
| 49 | C T A G T C A G C T G A T C A G T G C A A C T G T C G A T C A G | Trl(Zf)/S2-GAGAfactor-ChIP-Seq(GSE40646)/Homer | 1e-156 | -3.600e+02 | 0.0000 | 11844.0 | 79.14% | 47669.5 | 68.46% | motif file (matrix) | svg |
| 50 | A G C T C T A G A G T C A G T C A C T G C G T A A G T C C T G A G C A T G C T A C T G A G C A T G C A T C G A T G C A T | CBF4(AP2EREBP)/colamp-CBF4-DAP-Seq(GSE60143)/Homer | 1e-156 | -3.592e+02 | 0.0000 | 11435.0 | 76.41% | 45542.9 | 65.40% | motif file (matrix) | svg |
| 51 | G T A C C T G A T A G C C G T A G C T A T C G A T G C A T G A C C T A G G T C A A G T C C G T A C T G A C T G A C G T A | At1g14580(C2H2)/colamp-At1g14580-DAP-Seq(GSE60143)/Homer | 1e-154 | -3.554e+02 | 0.0000 | 1584.0 | 10.58% | 3206.5 | 4.60% | motif file (matrix) | svg |
| 52 | C G A T C T G A A G T C A C G T A C G T A T C G G A C T C T A G C G A T G A C T C G T A A T G C C G T A G T C A A C T G | ANAC011(NAC)/col-ANAC011-DAP-Seq(GSE60143)/Homer | 1e-152 | -3.519e+02 | 0.0000 | 3439.0 | 22.98% | 9742.3 | 13.99% | motif file (matrix) | svg |
| 53 | C G T A C T A G C A T G G A C T C T G A A C T G A C G T A C G T C T A G C T A G C A T G T C G A | MYB94(MYB)/col-MYB94-DAP-Seq(GSE60143)/Homer | 1e-150 | -3.471e+02 | 0.0000 | 4352.0 | 29.08% | 13350.2 | 19.17% | motif file (matrix) | svg |
| 54 | C T G A A T G C G C T A G C A T A T G C C G T A T C G A C T G A C T A G T C A G T A C G G T C A | Tcf4(HMG)/Hct116-Tcf4-ChIP-Seq(SRA012054)/Homer | 1e-149 | -3.442e+02 | 0.0000 | 3988.0 | 26.65% | 11931.2 | 17.13% | motif file (matrix) | svg |
| 55 | C T G A T C G A G T A C A C G T A C G T A T C G A C G T C G A T A T C G G C T A G T A C A T G C C G T A T G C A C A T G | ANAC103(NAC)/col-ANAC103-DAP-Seq(GSE60143)/Homer | 1e-149 | -3.438e+02 | 0.0000 | 4407.0 | 29.45% | 13601.9 | 19.53% | motif file (matrix) | svg |
| 56 | G A C T G A T C G A T C G C T A G T A C A G T C G C T A C T G A G T A C G A T C G C T A G A C T | MYB13(MYB)/col-MYB13-DAP-Seq(GSE60143)/Homer | 1e-147 | -3.390e+02 | 0.0000 | 6527.0 | 43.62% | 22551.3 | 32.39% | motif file (matrix) | svg |
| 57 | C G A T C G T A G T A C A C G T A C G T T C A G G C A T C A G T T A C G G T C A C G T A A G T C C G T A T G C A C A T G | NAC2(NAC)/colamp-NAC2-DAP-Seq(GSE60143)/Homer | 1e-145 | -3.340e+02 | 0.0000 | 6583.0 | 43.99% | 22852.8 | 32.82% | motif file (matrix) | svg |
| 58 | A G T C A C G T A C G T T A C G G C A T G C A T A T G C G C T A C G T A A T G C C G T A G T C A A C T G G A T C G C A T | ANAC075(NAC)/col-ANAC075-DAP-Seq(GSE60143)/Homer | 1e-145 | -3.339e+02 | 0.0000 | 4750.0 | 31.74% | 15087.0 | 21.67% | motif file (matrix) | svg |
| 59 | C G A T C G T A G T A C A C G T A C G T T C A G G C A T C A G T T A C G G T C A C G T A A G T C C G T A T G C A C A T G | ANAC053(NAC)/colamp-ANAC053-DAP-Seq(GSE60143)/Homer | 1e-144 | -3.332e+02 | 0.0000 | 5879.0 | 39.28% | 19817.5 | 28.46% | motif file (matrix) | svg |
| 60 | A C G T A G T C A G T C C G A T A C G T A C G T A C T G A C G T A T G C G A C T A C T G T A C G | Sox21(HMG)/ESC-SOX21-ChIP-Seq(GSE110505)/Homer | 1e-144 | -3.317e+02 | 0.0000 | 10278.0 | 68.68% | 40034.3 | 57.49% | motif file (matrix) | svg |
| 61 | G T A C C A T G A G C T A C G T A C T G C G T A A G T C G A C T G C A T C G A T | WRKY29(WRKY)/colamp-WRKY29-DAP-Seq(GSE60143)/Homer | 1e-143 | -3.305e+02 | 0.0000 | 9196.0 | 61.45% | 34815.2 | 50.00% | motif file (matrix) | svg |
| 62 | A T G C A G T C G A T C C G T A A C G T A C G T A C T G A C G T A G C T G A T C | Sox2(HMG)/mES-Sox2-ChIP-Seq(GSE11431)/Homer | 1e-142 | -3.286e+02 | 0.0000 | 6885.0 | 46.01% | 24243.9 | 34.82% | motif file (matrix) | svg |
| 63 | G A C T G C A T G C A T A G T C A G C T T C G A T A C G G C T A C G T A A C T G G T A C G C A T C G A T A G T C G A C T | HSF3(HSF)/colamp-HSF3-DAP-Seq(GSE60143)/Homer | 1e-142 | -3.272e+02 | 0.0000 | 6704.0 | 44.80% | 23461.7 | 33.69% | motif file (matrix) | svg |
| 64 | G A C T C A G T G C A T C G A T T G A C A C G T A T G C G T A C C T G A A C T G A C T G A G C T | WIP5(C2H2)/colamp-WIP5-DAP-Seq(GSE60143)/Homer | 1e-141 | -3.264e+02 | 0.0000 | 8814.0 | 58.90% | 33061.5 | 47.48% | motif file (matrix) | svg |
| 65 | G A T C A G C T C T A G G A T C T G A C C T A G C G T A G T A C C G T A G C A T G T C A C T G A | CBF3(AP2EREBP)/colamp-CBF3-DAP-Seq(GSE60143)/Homer | 1e-139 | -3.206e+02 | 0.0000 | 9231.0 | 61.68% | 35105.9 | 50.42% | motif file (matrix) | svg |
| 66 | G A C T A C T G C G A T A G T C A C T G C T A G A G T C C G T A | Rap210(AP2EREBP)/col-Rap210-DAP-Seq(GSE60143)/Homer | 1e-136 | -3.149e+02 | 0.0000 | 10448.0 | 69.82% | 41083.0 | 59.00% | motif file (matrix) | svg |
| 67 | C T G A G C A T A C T G C T A G A G T C A C T G A C T G A G T C A C T G T C A G | AT4G18450(AP2EREBP)/col-AT4G18450-DAP-Seq(GSE60143)/Homer | 1e-136 | -3.144e+02 | 0.0000 | 4939.0 | 33.00% | 16060.6 | 23.06% | motif file (matrix) | svg |
| 68 | A T G C G T A C C G T A A G C T G C A T T A C G A G C T A G C T A G T C A G C T | Sox6(HMG)/Myotubes-Sox6-ChIP-Seq(GSE32627)/Homer | 1e-136 | -3.134e+02 | 0.0000 | 10558.0 | 70.55% | 41647.4 | 59.81% | motif file (matrix) | svg |
| 69 | G C T A C G T A G C T A C G T A C T G A A C T G A C G T G T A C C G T A C T G A G T A C A C T G | WRKY22(WRKY)/colamp-WRKY22-DAP-Seq(GSE60143)/Homer | 1e-136 | -3.134e+02 | 0.0000 | 6134.0 | 40.99% | 21133.8 | 30.35% | motif file (matrix) | svg |
| 70 | C G A T A C G T A C G T A G C T A G C T G A T C G A T C G C T A A G C T A C G T A T C G T A C G | NFATC2(RHD)/Islets-NFATC2-ChIP-Seq(GSE158496)/Homer | 1e-135 | -3.109e+02 | 0.0000 | 9535.0 | 63.72% | 36680.1 | 52.68% | motif file (matrix) | svg |
| 71 | T C G A T C G A C T G A C G T A A C T G A T G C A C G T A G T C | Lola-I(Zf)/Embryo-LolaI-ChIP-Seq(GSE200870)/Homer | 1e-134 | -3.107e+02 | 0.0000 | 4793.0 | 32.03% | 15496.0 | 22.25% | motif file (matrix) | svg |
| 72 | A T G C C A T G A G C T C A G T C A T G T C G A A G T C G A C T C G A T C G A T C A G T C A G T | WRKY26(WRKY)/colamp-WRKY26-DAP-Seq(GSE60143)/Homer | 1e-134 | -3.100e+02 | 0.0000 | 5900.0 | 39.43% | 20163.6 | 28.96% | motif file (matrix) | svg |
| 73 | G T C A T G C A T G C A G C T A C G T A G C T A G C T A G C T A | REM19(REM)/colamp-REM19-DAP-Seq(GSE60143)/Homer | 1e-134 | -3.097e+02 | 0.0000 | 2525.0 | 16.87% | 6664.1 | 9.57% | motif file (matrix) | svg |
| 74 | A G C T G A T C C T G A A G T C A G T C A C T G C G T A A G T C C T G A G T C A G C A T C G A T G C T A G C A T C G T A | At2g44940(AP2EREBP)/colamp-At2g44940-DAP-Seq(GSE60143)/Homer | 1e-134 | -3.090e+02 | 0.0000 | 5828.0 | 38.94% | 19865.5 | 28.53% | motif file (matrix) | svg |
| 75 | G C A T A G C T A G C T A G C T A C T G A C G T A G T C A C T G A C G T G A C T C G A T G C A T | JKD(C2H2)/col-JKD-DAP-Seq(GSE60143)/Homer | 1e-133 | -3.082e+02 | 0.0000 | 2195.0 | 14.67% | 5498.2 | 7.90% | motif file (matrix) | svg |
| 76 | G A C T A C G T A G C T G A C T A C T G C A G T A G T C A T C G A C G T G C A T G C A T G C A T | MGP(C2H2)/colamp-MGP-DAP-Seq(GSE60143)/Homer | 1e-133 | -3.079e+02 | 0.0000 | 3492.0 | 23.33% | 10311.0 | 14.81% | motif file (matrix) | svg |
| 77 | C G A T T G C A A G T C A C G T A C G T T A C G C G A T C G A T T A C G G C T A G C T A A T G C C G T A G T C A C A T G | ANAC016(NAC)/col-ANAC016-DAP-Seq(GSE60143)/Homer | 1e-132 | -3.050e+02 | 0.0000 | 8338.0 | 55.72% | 31109.4 | 44.68% | motif file (matrix) | svg |
| 78 | C G A T C T G A A G T C A C G T A C G T T A C G G C T A C A T G C T A G G C A T C G A T A G T C C G T A G T C A A C T G | ANAC096(NAC)/colamp-ANAC096-DAP-Seq(GSE60143)/Homer | 1e-131 | -3.031e+02 | 0.0000 | 6358.0 | 42.49% | 22226.1 | 31.92% | motif file (matrix) | svg |
| 79 | C A T G A G T C G T A C A C T G A T G C A G T C C A T G G A T C G A T C C T G A | ERF5(AP2EREBP)/colamp-ERF5-DAP-Seq(GSE60143)/Homer | 1e-130 | -2.995e+02 | 0.0000 | 5063.0 | 33.83% | 16733.6 | 24.03% | motif file (matrix) | svg |
| 80 | G T A C A C T G A G T C A G T C C T A G G A T C G T A C C T G A | CRF4(AP2EREBP)/colamp-CRF4-DAP-Seq(GSE60143)/Homer | 1e-129 | -2.985e+02 | 0.0000 | 6679.0 | 44.63% | 23689.0 | 34.02% | motif file (matrix) | svg |
| 81 | A G T C G A C T A G C T C G A T A T C G G C T A C G A T A T C G C G A T A C T G T A C G A C G T | Tcf7(HMG)/GM12878-TCF7-ChIP-Seq(Encode)/Homer | 1e-129 | -2.978e+02 | 0.0000 | 3014.0 | 20.14% | 8565.8 | 12.30% | motif file (matrix) | svg |
| 82 | G A C T A C T G C G A T A G T C A C T G C T A G A G T C C T G A | AT1G12630(AP2EREBP)/colamp-AT1G12630-DAP-Seq(GSE60143)/Homer | 1e-128 | -2.965e+02 | 0.0000 | 8910.0 | 59.54% | 33892.9 | 48.67% | motif file (matrix) | svg |
| 83 | C G T A G A C T C A T G C T A G A G T C A C T G A C T G G T A C C A T G T A C G | ERF3(AP2EREBP)/colamp-ERF3-DAP-Seq(GSE60143)/Homer | 1e-128 | -2.962e+02 | 0.0000 | 8182.0 | 54.67% | 30500.4 | 43.80% | motif file (matrix) | svg |
| 84 | G A T C C A T G A C G T A C G T A C T G C G T A A G T C A G C T C G A T G A C T | WRKY8(WRKY)/colamp-WRKY8-DAP-Seq(GSE60143)/Homer | 1e-127 | -2.943e+02 | 0.0000 | 1553.0 | 10.38% | 3406.3 | 4.89% | motif file (matrix) | svg |
| 85 | A G C T A G C T A G C T A C T G A C G T A G T C A C T G A C G T G C A T G C A T G C A T A C G T | At5g66730(C2H2)/colamp-At5g66730-DAP-Seq(GSE60143)/Homer | 1e-127 | -2.938e+02 | 0.0000 | 2910.0 | 19.45% | 8205.2 | 11.78% | motif file (matrix) | svg |
| 86 | C G A T T G A C C A T G G A C T A C G T C A T G C G T A G A T C G A C T G C A T G C A T C G A T | WRKY14(WRKY)/colamp-WRKY14-DAP-Seq(GSE60143)/Homer | 1e-127 | -2.938e+02 | 0.0000 | 5454.0 | 36.45% | 18442.5 | 26.49% | motif file (matrix) | svg |
| 87 | G A C T A G T C C T G A A G T C A G T C A C T G C T G A A G T C G C T A G C A T G T A C C G A T G C A T G A C T C G A T | CBF2(AP2EREBP)/colamp-CBF2-DAP-Seq(GSE60143)/Homer | 1e-127 | -2.931e+02 | 0.0000 | 9322.0 | 62.29% | 35890.9 | 51.54% | motif file (matrix) | svg |
| 88 | C G T A G A T C A G C T A C G T A C G T A C T G C G T A G T A C A G C T G C T A C G A T C G A T C G A T G C A T G C T A | WRKY18(WRKY)/col-WRKY18-DAP-Seq(GSE60143)/Homer | 1e-127 | -2.930e+02 | 0.0000 | 11723.0 | 78.34% | 47824.2 | 68.68% | motif file (matrix) | svg |
| 89 | A G T C G T A C C T G A A G T C G T A C C T A G G C T A T G A C T G C A G C T A C G T A C G T A | At1g22810(AP2EREBP)/colamp-At1g22810-DAP-Seq(GSE60143)/Homer | 1e-127 | -2.925e+02 | 0.0000 | 7820.0 | 52.26% | 28886.7 | 41.48% | motif file (matrix) | svg |
| 90 | T C G A G T A C C A T G A G C T A C G T C A T G G T C A G T A C A G C T G C T A C G A T C A G T | WRKY31(WRKY)/colamp-WRKY31-DAP-Seq(GSE60143)/Homer | 1e-126 | -2.917e+02 | 0.0000 | 7086.0 | 47.35% | 25580.6 | 36.74% | motif file (matrix) | svg |
| 91 | C G T A G A T C C A T G G C A T G A C T C T A G T C G A T A G C A G C T G C A T | WRKY55(WRKY)/col-WRKY55-DAP-Seq(GSE60143)/Homer | 1e-126 | -2.911e+02 | 0.0000 | 9570.0 | 63.95% | 37104.2 | 53.29% | motif file (matrix) | svg |
| 92 | G A C T G T A C C T G A G A T C A G T C C T A G G C T A G T A C C T G A G C T A G C A T C G A T G C A T G A C T C G T A | AT3G16280(AP2EREBP)/colamp-AT3G16280-DAP-Seq(GSE60143)/Homer | 1e-126 | -2.904e+02 | 0.0000 | 8439.0 | 56.39% | 31766.1 | 45.62% | motif file (matrix) | svg |
| 93 | A G T C A G T C C T G A A G T C A G T C A C T G C G T A A G T C C T G A T C G A G C A T G A T C C G A T C G A T A C T G | AT3G60490(AP2EREBP)/colamp-AT3G60490-DAP-Seq(GSE60143)/Homer | 1e-126 | -2.904e+02 | 0.0000 | 6847.0 | 45.75% | 24533.0 | 35.23% | motif file (matrix) | svg |
| 94 | C G A T T C G A G A T C A C G T A C G T T C A G G C T A C G A T C G T A C G T A C G T A A T G C C G T A T G C A C T A G | ANAC028(NAC)/col-ANAC028-DAP-Seq(GSE60143)/Homer | 1e-125 | -2.890e+02 | 0.0000 | 6883.0 | 45.99% | 24709.1 | 35.48% | motif file (matrix) | svg |
| 95 | G A T C C A G T T A G C A G T C A C T G A G T C A G T C C T A G G A C T G T A C | LEP(AP2EREBP)/col-LEP-DAP-Seq(GSE60143)/Homer | 1e-125 | -2.880e+02 | 0.0000 | 3729.0 | 24.92% | 11420.0 | 16.40% | motif file (matrix) | svg |
| 96 | G C T A G C T A C G T A C G T A C T G A C T A G A C G T A G T C C G T A C T G A G T A C A C T G | WRKY65(WRKY)/colamp-WRKY65-DAP-Seq(GSE60143)/Homer | 1e-123 | -2.848e+02 | 0.0000 | 5612.0 | 37.50% | 19218.6 | 27.60% | motif file (matrix) | svg |
| 97 | G C T A C G T A C G A T C A G T A C T G C G A T G T A C A C T G A T C G G A C T C A T G C T A G G C A T C A G T C A T G | DEAR5(AP2EREBP)/col-DEAR5-DAP-Seq(GSE60143)/Homer | 1e-123 | -2.837e+02 | 0.0000 | 4096.0 | 27.37% | 12927.9 | 18.57% | motif file (matrix) | svg |
| 98 | A G T C T G A C C T G A A G T C A G T C A C T G C G T A A G T C G T C A G C T A G C A T C G T A G C A T G C T A C G T A | DEAR3(AP2EREBP)/colamp-DEAR3-DAP-Seq(GSE60143)/Homer | 1e-122 | -2.817e+02 | 0.0000 | 7163.0 | 47.87% | 26051.8 | 37.41% | motif file (matrix) | svg |
| 99 | C T A G G C A T A C T G C T A G A G T C A C T G A C T G A G T C A C T G T C A G | ERF10(AP2EREBP)/col-ERF10-DAP-Seq(GSE60143)/Homer | 1e-122 | -2.812e+02 | 0.0000 | 7813.0 | 52.21% | 29000.3 | 41.65% | motif file (matrix) | svg |
| 100 | C G A T G A T C G A T C C T G A G A T C G A T C C A T G T G C A G T A C T C G A G T C A G C A T C G A T C G A T G C A T | At4g32800(AP2EREBP)/colamp-At4g32800-DAP-Seq(GSE60143)/Homer | 1e-120 | -2.782e+02 | 0.0000 | 3812.0 | 25.47% | 11842.9 | 17.01% | motif file (matrix) | svg |
| 101 | C T G A A T C G G T A C C T G A A G T C A G T C A C T G C G T A A G T C C T G A | TINY(AP2EREBP)/col-TINY-DAP-Seq(GSE60143)/Homer | 1e-119 | -2.746e+02 | 0.0000 | 6060.0 | 40.49% | 21270.7 | 30.55% | motif file (matrix) | svg |
| 102 | A G C T A G C T C A T G C T G A G T A C A G T C A G C T A G C T C A G T C T A G | RARa(NR)/K562-RARa-ChIP-Seq(Encode)/Homer | 1e-118 | -2.730e+02 | 0.0000 | 12656.0 | 84.57% | 53027.4 | 76.15% | motif file (matrix) | svg |
| 103 | C T A G C T A G A T G C G T A C T C A G A T G C A G T C G C A T G A T C G A T C | ZNF91(Zf)/HEK-ZNF91.HA-ChIP-Seq(GSE162571)/Homer | 1e-118 | -2.720e+02 | 0.0000 | 5641.0 | 37.69% | 19494.0 | 28.00% | motif file (matrix) | svg |
| 104 | G C A T T G A C C A T G G A C T C A G T C A T G T C G A G T A C G A C T G C T A C G A T C G A T | WRKY6(WRKY)/colamp-WRKY6-DAP-Seq(GSE60143)/Homer | 1e-118 | -2.717e+02 | 0.0000 | 8093.0 | 54.08% | 30411.2 | 43.67% | motif file (matrix) | svg |
| 105 | A G C T G A C T A C G T A C T G A C G T A G T C A C T G A C G T G C A T C G A T | AtIDD11(C2H2)/colamp-AtIDD11-DAP-Seq(GSE60143)/Homer | 1e-117 | -2.701e+02 | 0.0000 | 3857.0 | 25.77% | 12102.8 | 17.38% | motif file (matrix) | svg |
| 106 | T A C G C G T A G A C T T C A G A G C T A G T C A C T G T C A G A G T C C T G A | DDF2(AP2EREBP)/col-DDF2-DAP-Seq(GSE60143)/Homer | 1e-116 | -2.689e+02 | 0.0000 | 2055.0 | 13.73% | 5265.9 | 7.56% | motif file (matrix) | svg |
| 107 | C G A T A G C T T G C A A C T G A G T C T G A C C T A G G T A C A G T C C G T A G C A T G C A T | ERF13(AP2EREBP)/colamp-ERF13-DAP-Seq(GSE60143)/Homer | 1e-115 | -2.669e+02 | 0.0000 | 10037.0 | 67.07% | 39691.6 | 57.00% | motif file (matrix) | svg |
| 108 | A T G C C A T G G C A T G A C T C T A G T C G A G T A C A G C T C G T A G C T A | WRKY75(WRKY)/col-WRKY75-DAP-Seq(GSE60143)/Homer | 1e-114 | -2.645e+02 | 0.0000 | 8586.0 | 57.37% | 32798.1 | 47.10% | motif file (matrix) | svg |
| 109 | G C A T C G A T C G T A G A T C C A T G A C G T A C G T A C T G C G T A A G T C A G C T G C A T G C A T C G T A G C T A | WRKY45(WRKY)/col-WRKY45-DAP-Seq(GSE60143)/Homer | 1e-113 | -2.602e+02 | 0.0000 | 4198.0 | 28.05% | 13583.1 | 19.51% | motif file (matrix) | svg |
| 110 | C T G A T C A G G T A C G C T A A C T G T G A C G C A T C A T G | SCL(bHLH)/HPC7-Scl-ChIP-Seq(GSE13511)/Homer | 1e-112 | -2.591e+02 | 0.0000 | 13432.0 | 89.76% | 57521.3 | 82.61% | motif file (matrix) | svg |
| 111 | C G A T C T A G A C G T A C G T A C G T C G T A A G C T C G A T A G C T C G T A C T A G T A G C | FoxD3(forkhead)/ZebrafishEmbryo-Foxd3.biotin-ChIP-seq(GSE106676)/Homer | 1e-112 | -2.586e+02 | 0.0000 | 4825.0 | 32.24% | 16192.8 | 23.25% | motif file (matrix) | svg |
| 112 | G A T C A T G C A G T C C G T A A G T C A G T C A C T G G C T A A G T C C G T A | AT1G44830(AP2EREBP)/col-AT1G44830-DAP-Seq(GSE60143)/Homer | 1e-111 | -2.569e+02 | 0.0000 | 4807.0 | 32.12% | 16138.0 | 23.18% | motif file (matrix) | svg |
| 113 | A T G C G T A C A G T C A G T C A C G T A C G T C G A T A C G T | AT5G02460(C2C2dof)/col-AT5G02460-DAP-Seq(GSE60143)/Homer | 1e-111 | -2.563e+02 | 0.0000 | 10592.0 | 70.78% | 42567.0 | 61.13% | motif file (matrix) | svg |
| 114 | T A C G C T G A C A T G G A T C G T A C G C A T T C A G T A C G A G C T G T C A G A T C G C A T T A C G C G T A C T A G G A T C G A T C C G A T A C T G T C A G | ZNF322(Zf)/HEK293-ZNF322.GFP-ChIP-Seq(GSE58341)/Homer | 1e-111 | -2.557e+02 | 0.0000 | 1403.0 | 9.38% | 3123.7 | 4.49% | motif file (matrix) | svg |
| 115 | G A T C C T G A A G T C G T A C A C T G G C T A G A T C C T G A G C T A G C T A | At4g31060(AP2EREBP)/colamp-At4g31060-DAP-Seq(GSE60143)/Homer | 1e-110 | -2.555e+02 | 0.0000 | 8053.0 | 53.81% | 30448.2 | 43.73% | motif file (matrix) | svg |
| 116 | C G A T T C G A G A T C A C G T A C G T T C A G G C A T C T G A C G T A G C T A C G T A A G T C C G T A T G C A C A T G | ANAC050(NAC)/colamp-ANAC050-DAP-Seq(GSE60143)/Homer | 1e-110 | -2.543e+02 | 0.0000 | 6084.0 | 40.65% | 21628.4 | 31.06% | motif file (matrix) | svg |
| 117 | C A G T T C A G T C G A A G T C C G T A A C T G T G A C C G A T A C T G A C T G A C G T A T C G | Atoh7(bHLH)/Retina-Atoh7-CutnRun(GSE156756)/Homer | 1e-109 | -2.531e+02 | 0.0000 | 4521.0 | 30.21% | 14989.2 | 21.53% | motif file (matrix) | svg |
| 118 | A T G C C A T G A C G T A C G T A C T G C G T A A G T C G A C T G C A T C G A T | WRKY71(WRKY)/col-WRKY71-DAP-Seq(GSE60143)/Homer | 1e-109 | -2.530e+02 | 0.0000 | 7239.0 | 48.37% | 26770.8 | 38.45% | motif file (matrix) | svg |
| 119 | C T A G C T A G T C G A C T A G C G T A A T C G T C G A A C T G C T G A T C G A C T G A T A C G | FRS9(ND)/col-FRS9-DAP-Seq(GSE60143)/Homer | 1e-109 | -2.516e+02 | 0.0000 | 1059.0 | 7.08% | 2060.9 | 2.96% | motif file (matrix) | svg |
| 120 | T C G A A G T C C G T A A T C G A T G C C G A T A C T G A G T C A G C T A C T G | Tcf12(bHLH)/GM12878-Tcf12-ChIP-Seq(GSE32465)/Homer | 1e-108 | -2.506e+02 | 0.0000 | 4833.0 | 32.30% | 16318.2 | 23.43% | motif file (matrix) | svg |
| 121 | A G T C C T A G A C G T A C G T A C T G C G T A A G T C A G C T G C T A G C A T | WRKY24(WRKY)/colamp-WRKY24-DAP-Seq(GSE60143)/Homer | 1e-108 | -2.506e+02 | 0.0000 | 8191.0 | 54.73% | 31153.8 | 44.74% | motif file (matrix) | svg |
| 122 | G A C T C T G A A G T C A G T C A C T G C G T A A G T C C T G A | bHLH10(bHLH)/colamp-bHLH10-DAP-Seq(GSE60143)/Homer | 1e-108 | -2.491e+02 | 0.0000 | 6553.0 | 43.79% | 23757.1 | 34.12% | motif file (matrix) | svg |
| 123 | A T G C C A G T A G C T A G C T T C A G G T C A T A G C G A C T C G T A C G A T | WRKY20(WRKY)/col-WRKY20-DAP-Seq(GSE60143)/Homer | 1e-107 | -2.471e+02 | 0.0000 | 7275.0 | 48.61% | 27014.1 | 38.79% | motif file (matrix) | svg |
| 124 | G A T C G C T A G T A C A G T C G C T A T G C A G T A C G A T C C G T A G A C T | MYB83(MYB)/colamp-MYB83-DAP-Seq(GSE60143)/Homer | 1e-107 | -2.468e+02 | 0.0000 | 11411.0 | 76.25% | 46816.8 | 67.23% | motif file (matrix) | svg |
| 125 | T G A C T A G C T C A G T C G A T C G A C G T A A G T C C G T A C G T A C G A T C T A G T A C G | Sox7(HMG)/ESC-Sox7-ChIP-Seq(GSE133899)/Homer | 1e-107 | -2.466e+02 | 0.0000 | 3473.0 | 23.21% | 10805.2 | 15.52% | motif file (matrix) | svg |
| 126 | G A T C C T G A A G T C A G T C A C T G C G T A A G T C C T G A | ERF38(AP2EREBP)/col-ERF38-DAP-Seq(GSE60143)/Homer | 1e-105 | -2.434e+02 | 0.0000 | 8566.0 | 57.24% | 32999.8 | 47.39% | motif file (matrix) | svg |
| 127 | G T A C T C G A T A G C C G T A C G T A C T G A T G C A T G A C A C T G C G T A A G T C C T G A C T G A T C G A C G T A | NUC(C2H2)/col-NUC-DAP-Seq(GSE60143)/Homer | 1e-105 | -2.428e+02 | 0.0000 | 1169.0 | 7.81% | 2440.6 | 3.50% | motif file (matrix) | svg |
| 128 | C A G T A C T G T C A G T G C A G C T A A T G C T C G A A T C G G T C A T G C A | ZNF189(Zf)/HEK293-ZNF189.GFP-ChIP-Seq(GSE58341)/Homer | 1e-105 | -2.424e+02 | 0.0000 | 5342.0 | 35.70% | 18573.7 | 26.67% | motif file (matrix) | svg |
| 129 | A T G C A G T C A G C T A G C T A C G T A T C G C G T A C G A T T A G C G A C T | LEF1(HMG)/H1-LEF1-ChIP-Seq(GSE64758)/Homer | 1e-105 | -2.424e+02 | 0.0000 | 5128.0 | 34.27% | 17662.7 | 25.37% | motif file (matrix) | svg |
| 130 | A G T C G T A C C T G A A G T C A G T C C A T G G C T A A G T C T G C A G C T A G C A T G C A T | RAP21(AP2EREBP)/colamp-RAP21-DAP-Seq(GSE60143)/Homer | 1e-105 | -2.422e+02 | 0.0000 | 4741.0 | 31.68% | 16032.3 | 23.02% | motif file (matrix) | svg |
| 131 | A T G C G T A C C T G A A G T C A G T C A C T G G T C A A G T C G T C A G C A T G C A T G A C T | At5g65130(AP2EREBP)/colamp-At5g65130-DAP-Seq(GSE60143)/Homer | 1e-104 | -2.414e+02 | 0.0000 | 4011.0 | 26.80% | 13023.3 | 18.70% | motif file (matrix) | svg |
| 132 | G A C T C T A G A T G C A G T C G T C A T A C G A T G C A T C G | HIC1(Zf)/Treg-ZBTB29-ChIP-Seq(GSE99889)/Homer | 1e-104 | -2.410e+02 | 0.0000 | 11240.0 | 75.11% | 46027.0 | 66.10% | motif file (matrix) | svg |
| 133 | C G A T T C G A G A T C C G A T G C A T T C A G G A C T C G A T G C A T G C T A C T G A A G T C C G T A G T C A C T A G | ANAC005(NAC)/col-ANAC005-DAP-Seq(GSE60143)/Homer | 1e-104 | -2.401e+02 | 0.0000 | 3941.0 | 26.33% | 12752.3 | 18.31% | motif file (matrix) | svg |
| 134 | G T C A T C G A T C G A C G T A G C T A C G T A T C G A T G A C A C T G C G T A A G T C C G T A C G T A T C G A G C T A | IDD2(C2H2)/colamp-IDD2-DAP-Seq(GSE60143)/Homer | 1e-103 | -2.391e+02 | 0.0000 | 1336.0 | 8.93% | 2994.1 | 4.30% | motif file (matrix) | svg |
| 135 | C A G T C G T A C G T A G C A T G A C T G C A T G T A C A G C T A C T G G A C T A C G T C A T G | RAV1(RAV)/colamp-RAV1-DAP-Seq(GSE60143)/Homer | 1e-103 | -2.388e+02 | 0.0000 | 4781.0 | 31.95% | 16240.3 | 23.32% | motif file (matrix) | svg |
| 136 | G A C T T C A G G C A T A G T C G C T A G A T C C T G A A C G T A G T C G T C A | Replumless(BLH)/Arabidopsis-RPL.GFP-ChIP-Seq(GSE78727)/Homer | 1e-103 | -2.385e+02 | 0.0000 | 10233.0 | 68.38% | 41046.9 | 58.95% | motif file (matrix) | svg |
| 137 | G C A T A C T G C T A G A G T C A C T G A C T G A G T C A C G T | ERF105(AP2EREBP)/colamp-ERF105-DAP-Seq(GSE60143)/Homer | 1e-103 | -2.381e+02 | 0.0000 | 10417.0 | 69.61% | 41957.0 | 60.25% | motif file (matrix) | svg |
| 138 | A T G C G A T C C G T A A G C T C A G T A T C G G C A T A G C T G A C T A C T G | Sox17(HMG)/Endoderm-Sox17-ChIP-Seq(GSE61475)/Homer | 1e-100 | -2.305e+02 | 0.0000 | 6409.0 | 42.83% | 23371.4 | 33.56% | motif file (matrix) | svg |
| 139 | G A C T A C T G C A G T A G T C A C T G A C T G A G C T A C T G C T A G G T C A | At1g77640(AP2EREBP)/col-At1g77640-DAP-Seq(GSE60143)/Homer | 1e-99 | -2.285e+02 | 0.0000 | 3242.0 | 21.66% | 10072.7 | 14.47% | motif file (matrix) | svg |
| 140 | A G C T C T A G G A T C A G T C C T A G C T G A A G T C G C T A G C A T T G C A | CBF1(AP2EREBP)/colamp-CBF1-DAP-Seq(GSE60143)/Homer | 1e-99 | -2.282e+02 | 0.0000 | 10531.0 | 70.37% | 42662.6 | 61.27% | motif file (matrix) | svg |
| 141 | G A C T G C A T C T A G C G A T G A T C T C G A C A T G G A T C | Tgif1(Homeobox)/mES-Tgif1-ChIP-Seq(GSE55404)/Homer | 1e-98 | -2.273e+02 | 0.0000 | 13195.0 | 88.17% | 56544.5 | 81.20% | motif file (matrix) | svg |
| 142 | T C A G T G A C C A T G G C A T C A G T A C T G C G T A T G A C G A C T C G A T C G A T C G T A | WRKY3(WRKY)/col-WRKY3-DAP-Seq(GSE60143)/Homer | 1e-97 | -2.250e+02 | 0.0000 | 6519.0 | 43.56% | 23935.9 | 34.37% | motif file (matrix) | svg |
| 143 | A T C G T C G A G A C T A T C G T G A C A C G T C T A G A C T G C G T A A C T G A G T C G T A C | ZNF415(Zf)/HEK293-ZNF415.GFP-ChIP-Seq(GSE58341)/Homer | 1e-97 | -2.246e+02 | 0.0000 | 4466.0 | 29.84% | 15091.9 | 21.67% | motif file (matrix) | svg |
| 144 | C G T A G A C T C A T G C T A G A G T C A C T G C T A G A G T C C A T G C T A G | ERF7(AP2EREBP)/col-ERF7-DAP-Seq(GSE60143)/Homer | 1e-94 | -2.183e+02 | 0.0000 | 11689.0 | 78.11% | 48635.8 | 69.85% | motif file (matrix) | svg |
| 145 | T C A G A T C G G A C T A C T G G A C T C A G T C T A G C G T A G T A C C G T A C T A G A T C G | Tbx20(T-box)/Heart-Tbx20-ChIP-Seq(GSE29636)/Homer | 1e-93 | -2.163e+02 | 0.0000 | 2313.0 | 15.46% | 6606.7 | 9.49% | motif file (matrix) | svg |
| 146 | G C A T C G A T G A C T T G C A A C T G A G T C T G A C A C T G G A T C A G T C C G T A G A C T | ERF15(AP2EREBP)/colamp-ERF15-DAP-Seq(GSE60143)/Homer | 1e-93 | -2.162e+02 | 0.0000 | 11549.0 | 77.17% | 47948.3 | 68.86% | motif file (matrix) | svg |
| 147 | C T A G A C T G T G C A G T C A A T G C C G T A A T C G A T G C A G T C C T A G | ZNF341(Zf)/EBV-ZNF341-ChIP-Seq(GSE113194)/Homer | 1e-91 | -2.110e+02 | 0.0000 | 5121.0 | 34.22% | 18031.8 | 25.90% | motif file (matrix) | svg |
| 148 | G C A T T G A C C T G A A G T C A G T C A C T G G T C A A G T C G C T A G A C T G C T A C T G A | DREB2(AP2EREBP)/col-DREB2-DAP-Seq(GSE60143)/Homer | 1e-91 | -2.109e+02 | 0.0000 | 7793.0 | 52.07% | 29896.2 | 42.93% | motif file (matrix) | svg |
| 149 | A G T C G A C T G A T C C G T A G T A C A G T C G C T A C G T A G T A C A G T C G T A C G T A C | MYB63(MYB)/col-MYB63-DAP-Seq(GSE60143)/Homer | 1e-91 | -2.102e+02 | 0.0000 | 5215.0 | 34.85% | 18445.0 | 26.49% | motif file (matrix) | svg |
| 150 | T C A G A G C T G T C A C G T A A C G T A T G C C G T A A C G T A C G T C T G A | PHV(HB)/col-PHV-DAP-Seq(GSE60143)/Homer | 1e-91 | -2.102e+02 | 0.0000 | 4203.0 | 28.09% | 14171.2 | 20.35% | motif file (matrix) | svg |
| 151 | C G A T C T A G A C T G A G C T C T G A A C T G A C G T A C G T C T A G C T A G | MYB96(MYB)/colamp-MYB96-DAP-Seq(GSE60143)/Homer | 1e-90 | -2.089e+02 | 0.0000 | 7783.0 | 52.01% | 29881.6 | 42.91% | motif file (matrix) | svg |
| 152 | G C T A T C G A C G T A C T G A A C T G A C G T A G T C C G T A C G T A A G T C C T A G T G C A | WRKY42(WRKY)/colamp-WRKY42-DAP-Seq(GSE60143)/Homer | 1e-89 | -2.053e+02 | 0.0000 | 5200.0 | 34.75% | 18446.2 | 26.49% | motif file (matrix) | svg |
| 153 | G C A T C G T A C T A G A G T C G T C A C G T A A T G C A C G T A C G T A C T G G A T C G C A T C G T A G C T A G C T A | bHLH122(bHLH)/col100-bHLH122-DAP-Seq(GSE60143)/Homer | 1e-88 | -2.048e+02 | 0.0000 | 7149.0 | 47.77% | 27053.4 | 38.85% | motif file (matrix) | svg |
| 154 | A T G C G C A T T A G C C G A T T A G C G C A T T A G C G C A T A T G C G A C T | GAGA-repeat/Arabidopsis-Promoters/Homer | 1e-88 | -2.044e+02 | 0.0000 | 5376.0 | 35.92% | 19216.4 | 27.60% | motif file (matrix) | svg |
| 155 | G T A C A C T G A T G C A G T C C T A G G A T C G T A C C T G A G A C T G C A T C G A T G A C T | RAP212(AP2EREBP)/col-RAP212-DAP-Seq(GSE60143)/Homer | 1e-88 | -2.035e+02 | 0.0000 | 9074.0 | 60.63% | 35990.5 | 51.69% | motif file (matrix) | svg |
| 156 | T C A G A C G T T C G A T A G C A G T C C G T A A C T G G T A C A C G T A C T G A T C G A G T C | Atoh1(bHLH)/Cerebellum-Atoh1-ChIP-Seq(GSE22111)/Homer | 1e-88 | -2.033e+02 | 0.0000 | 6353.0 | 42.45% | 23509.2 | 33.76% | motif file (matrix) | svg |
| 157 | T A G C C A T G G A C T G A C T T C A G G T C A G A T C G A C T G C A T G C T A | WRKY15(WRKY)/col-WRKY15-DAP-Seq(GSE60143)/Homer | 1e-87 | -2.025e+02 | 0.0000 | 8701.0 | 58.14% | 34246.4 | 49.18% | motif file (matrix) | svg |
| 158 | C T A G G C T A A G T C A C T G A C G T G A C T G A C T A T G C T C G A C A G T G A T C C G A T G A C T G A T C G A T C | RKD2(RWPRK)/colamp-RKD2-DAP-Seq(GSE60143)/Homer | 1e-87 | -2.019e+02 | 0.0000 | 6361.0 | 42.51% | 23565.6 | 33.84% | motif file (matrix) | svg |
| 159 | C G A T T C G A G T A C A C G T A C G T T C A G G C A T G C A T G C T A C G T A C G T A A G T C C G T A T G C A C A T G | ANAC020(NAC)/col-ANAC020-DAP-Seq(GSE60143)/Homer | 1e-86 | -2.000e+02 | 0.0000 | 6660.0 | 44.50% | 24926.3 | 35.80% | motif file (matrix) | svg |
| 160 | A T C G T G C A G A T C C T A G A C G T A T C G C G T A A G T C T C A G A C T G T C A G G C T A | Knotted(Homeobox)/Corn-KN1-ChIP-Seq(GSE39161)/Homer | 1e-86 | -1.991e+02 | 0.0000 | 11793.0 | 78.80% | 49447.7 | 71.01% | motif file (matrix) | svg |
| 161 | C A T G T G A C C A T G G A C T C A G T C T A G G C T A G T A C G A C T G C A T G C A T C G A T | WRKY21(WRKY)/colamp-WRKY21-DAP-Seq(GSE60143)/Homer | 1e-85 | -1.957e+02 | 0.0000 | 1444.0 | 9.65% | 3620.5 | 5.20% | motif file (matrix) | svg |
| 162 | G T A C A C G T A C G T T C A G G A C T G C A T T C A G C G T A C T G A A G T C C G T A G T C A A C T G A C G T G C T A | NTM2(NAC)/col-NTM2-DAP-Seq(GSE60143)/Homer | 1e-84 | -1.950e+02 | 0.0000 | 5454.0 | 36.45% | 19686.5 | 28.27% | motif file (matrix) | svg |
| 163 | C G A T C G T A A G T C A C G T A C G T T C G A T G C A G C A T G C T A C G T A A C G T A G C T C G T A C G T A A C T G | ANAC062(NAC)/colamp-ANAC062-DAP-Seq(GSE60143)/Homer | 1e-84 | -1.948e+02 | 0.0000 | 3273.0 | 21.87% | 10552.2 | 15.15% | motif file (matrix) | svg |
| 164 | A C T G C T A G A G T C A C T G A C T G A G T C A C G T C T A G | AT5G23930(mTERF)/col-AT5G23930-DAP-Seq(GSE60143)/Homer | 1e-84 | -1.945e+02 | 0.0000 | 9942.0 | 66.44% | 40294.7 | 57.87% | motif file (matrix) | svg |
| 165 | C G T A C G A T C G T A C G A T C A T G A C T G C G A T A G T C A T C G T C A G G A C T A C T G | At1g36060(AP2EREBP)/colamp-At1g36060-DAP-Seq(GSE60143)/Homer | 1e-84 | -1.945e+02 | 0.0000 | 9475.0 | 63.31% | 38046.0 | 54.64% | motif file (matrix) | svg |
| 166 | A T G C G T A C A C T G A G T C A G T C A C T G A G T C G T A C | ERF73(AP2EREBP)/col-ERF73-DAP-Seq(GSE60143)/Homer | 1e-84 | -1.937e+02 | 0.0000 | 5984.0 | 39.99% | 22020.4 | 31.62% | motif file (matrix) | svg |
| 167 | A G T C A C G T A C G T T C A G G C T A G C T A A T G C C G T A C G A T A G T C C G T A G T C A A C T G G A T C G C A T | SND3(NAC)/col-SND3-DAP-Seq(GSE60143)/Homer | 1e-84 | -1.936e+02 | 0.0000 | 7179.0 | 47.97% | 27361.0 | 39.29% | motif file (matrix) | svg |
| 168 | G C T A C G T A C G T A G A C T C A T G C T A G G A T C A C T G T C A G G A T C A C T G T A C G | ERF9(AP2EREBP)/colamp-ERF9-DAP-Seq(GSE60143)/Homer | 1e-83 | -1.927e+02 | 0.0000 | 3804.0 | 25.42% | 12735.4 | 18.29% | motif file (matrix) | svg |
| 169 | A C G T T G C A A G C T G A T C C T A G C T G A A G C T G T C A T C G A C G T A | CUX1(Homeobox)/K562-CUX1-ChIP-Seq(GSE92882)/Homer | 1e-83 | -1.923e+02 | 0.0000 | 9697.0 | 64.80% | 39146.9 | 56.22% | motif file (matrix) | svg |
| 170 | T C A G T C A G G C T A C G T A T A C G G A C T T C A G T C G A C T G A C G T A T A C G G A C T | IRF8(IRF)/BMDM-IRF8-ChIP-Seq(GSE77884)/Homer | 1e-83 | -1.921e+02 | 0.0000 | 2120.0 | 14.17% | 6092.9 | 8.75% | motif file (matrix) | svg |
| 171 | C G T A G A T C C T A G A C G T G T A C C T G A A G C T G A T C G C T A G A C T | TGA2(bZIP)/colamp-TGA2-DAP-Seq(GSE60143)/Homer | 1e-82 | -1.909e+02 | 0.0000 | 9436.0 | 63.05% | 37917.0 | 54.45% | motif file (matrix) | svg |
| 172 | G C A T C G A T G C A T C G T A C T A G A G T C G T C A C G T A A T C G A C G T A C G T A C T G G T A C G C A T C G A T | bHLH80(bHLH)/col-bHLH80-DAP-Seq(GSE60143)/Homer | 1e-81 | -1.887e+02 | 0.0000 | 7489.0 | 50.04% | 28851.0 | 41.43% | motif file (matrix) | svg |
| 173 | G C A T A C G T A C T G A C G T A G T C A C T G A T C G G T C A C G A T C G T A | ARF2(ARF)/col-ARF2-DAP-Seq(GSE60143)/Homer | 1e-81 | -1.886e+02 | 0.0000 | 13452.0 | 89.89% | 58458.1 | 83.95% | motif file (matrix) | svg |
| 174 | C T A G C T G A C T A G C T G A C T A G C T G A C T A G C T G A C T A G C T G A | SeqBias: GA-repeat | 1e-80 | -1.865e+02 | 0.0000 | 14202.0 | 94.90% | 62871.7 | 90.29% | motif file (matrix) | svg |
| 175 | G T A C G C T A T C A G C T G A C T A G C A T G A G C T G A T C T G C A T C G A C T G A A C T G C A G T A G T C G A T C G C T A | HNF4a(NR),DR1/HepG2-HNF4a-ChIP-Seq(GSE25021)/Homer | 1e-80 | -1.864e+02 | 0.0000 | 2783.0 | 18.60% | 8695.5 | 12.49% | motif file (matrix) | svg |
| 176 | G C A T G C A T G C A T A T G C A G C T T C G A T A C G G C T A C G T A C A T G G T A C G C A T G C A T A G T C A G C T | HSFA6B(HSF)/colamp-HSFA6B-DAP-Seq(GSE60143)/Homer | 1e-80 | -1.845e+02 | 0.0000 | 3871.0 | 25.87% | 13113.7 | 18.83% | motif file (matrix) | svg |
| 177 | T A C G T A G C G C T A C G A T C T A G A C G T C A G T C A G T G C T A A G T C G T C A G C A T | FOXK2(Forkhead)/U2OS-FOXK2-ChIP-Seq(E-MTAB-2204)/Homer | 1e-80 | -1.844e+02 | 0.0000 | 5375.0 | 35.92% | 19495.0 | 28.00% | motif file (matrix) | svg |
| 178 | C G T A C G A T C T A G C G T A A G C T C A G T T A C G C G T A A C G T C A T G C T A G A T G C | HOXA3(Homeobox)/mEmbryo-Hoxa3-ChIP-Seq(E-MTAB-8607)/Homer | 1e-80 | -1.843e+02 | 0.0000 | 2091.0 | 13.97% | 6052.1 | 8.69% | motif file (matrix) | svg |
| 179 | G T A C C A T G A G C T A G C T T C A G T G C A T G A C A G C T C G T A C G T A | WRKY33(WRKY)/col-WRKY33-DAP-Seq(GSE60143)/Homer | 1e-79 | -1.836e+02 | 0.0000 | 7969.0 | 53.25% | 31139.5 | 44.72% | motif file (matrix) | svg |
| 180 | T C A G T G A C G T A C C G T A A C G T T G A C A C G T T C A G A G C T G A C T | NeuroD1(bHLH)/Islet-NeuroD1-ChIP-Seq(GSE30298)/Homer | 1e-79 | -1.828e+02 | 0.0000 | 4881.0 | 32.62% | 17388.9 | 24.97% | motif file (matrix) | svg |
| 181 | C G A T A G T C C A T G G A C T A C G T C T A G C G T A G A T C G A C T C G A T G C A T G A C T | WRKY43(WRKY)/colamp-WRKY43-DAP-Seq(GSE60143)/Homer | 1e-78 | -1.810e+02 | 0.0000 | 3914.0 | 26.15% | 13335.9 | 19.15% | motif file (matrix) | svg |
| 182 | A C T G C T A G A G T C A C T G A C T G A T G C A C T G T A C G | ESE1(AP2EREBP)/col-ESE1-DAP-Seq(GSE60143)/Homer | 1e-78 | -1.804e+02 | 0.0000 | 6664.0 | 44.53% | 25247.6 | 36.26% | motif file (matrix) | svg |
| 183 | G A T C G T A C C T G A A G T C A G T C A C T G G C T A G T A C G T C A G C A T G C A T C G A T | DEAR2(AP2EREBP)/colamp-DEAR2-DAP-Seq(GSE60143)/Homer | 1e-78 | -1.801e+02 | 0.0000 | 11987.0 | 80.10% | 50729.0 | 72.85% | motif file (matrix) | svg |
| 184 | C G T A C T A G T C A G T C A G A G T C A T G C A G T C G C A T A G C T A C G T A T C G C G A T | Sox9(HMG)/Limb-SOX9-ChIP-Seq(GSE73225)/Homer | 1e-77 | -1.783e+02 | 0.0000 | 5880.0 | 39.29% | 21794.4 | 31.30% | motif file (matrix) | svg |
| 185 | C T G A A T G C G C T A C G A T A T G C C G T A C G T A C G T A C T A G T A C G | Tcf3(HMG)/mES-Tcf3-ChIP-Seq(GSE11724)/Homer | 1e-76 | -1.770e+02 | 0.0000 | 2340.0 | 15.64% | 7066.9 | 10.15% | motif file (matrix) | svg |
| 186 | G C T A C T G A T C G A A G T C A G T C C T G A A G T C G T C A C T G A T G C A | RUNX1(Runt)/Jurkat-RUNX1-ChIP-Seq(GSE29180)/Homer | 1e-76 | -1.764e+02 | 0.0000 | 7163.0 | 47.87% | 27564.3 | 39.59% | motif file (matrix) | svg |
| 187 | C A T G A T G C T A G C C T G A A G T C A G T C A C T G G C T A A G T C G T A C G C T A G C A T | At4g28140(AP2EREBP)/colamp-At4g28140-DAP-Seq(GSE60143)/Homer | 1e-76 | -1.757e+02 | 0.0000 | 5542.0 | 37.03% | 20352.8 | 29.23% | motif file (matrix) | svg |
| 188 | C A T G G A T C C T G A G T A C C T A G C T G A G C T A G C A T G A T C G A T C A G T C C T A G C G T A C A T G C T A G | AIL7(AP2EREBP)/colamp-AIL7-DAP-Seq(GSE60143)/Homer | 1e-75 | -1.745e+02 | 0.0000 | 6158.0 | 41.15% | 23082.8 | 33.15% | motif file (matrix) | svg |
| 189 | C G A T C T A G C T G A A T G C C T G A T C G A C G T A C T G A T C G A T A G C A G T C C G T A A C T G T C G A A T G C | Hand2(bHLH)/Mesoderm-Hand2-ChIP-Seq(GSE61475)/Homer | 1e-75 | -1.743e+02 | 0.0000 | 2718.0 | 18.16% | 8567.9 | 12.30% | motif file (matrix) | svg |
| 190 | C G T A C G T A C G A T A C T G C A G T A G T C A C T G A C T G A G C T A C T G | DREB19(AP2EREBP)/colamp-DREB19-DAP-Seq(GSE60143)/Homer | 1e-75 | -1.734e+02 | 0.0000 | 8560.0 | 57.20% | 34059.4 | 48.91% | motif file (matrix) | svg |
| 191 | C G A T C G T A A G T C A C G T A C G T T C A G G C A T C G T A G C T A G C T A C G T A A G T C C G T A G T C A A C T G | ANAC058(NAC)/col-ANAC058-DAP-Seq(GSE60143)/Homer | 1e-75 | -1.732e+02 | 0.0000 | 6132.0 | 40.98% | 22987.5 | 33.01% | motif file (matrix) | svg |
| 192 | C T G A C T A G A C T G G C A T A T G C C G T A C T G A C T A G A C T G A C G T A G T C C T G A | RARg(NR)/ES-RARg-ChIP-Seq(GSE30538)/Homer | 1e-74 | -1.719e+02 | 0.0000 | 417.0 | 2.79% | 556.2 | 0.80% | motif file (matrix) | svg |
| 193 | T C G A A G T C A C G T A C G T T C A G C A G T C T G A C T A G T C G A C G T A A T C G C G T A C G T A A C T G A G C T | NTM1(NAC)/col-NTM1-DAP-Seq(GSE60143)/Homer | 1e-73 | -1.689e+02 | 0.0000 | 3922.0 | 26.21% | 13526.9 | 19.43% | motif file (matrix) | svg |
| 194 | G A T C G A C T G A C T A C G T A G T C A C G T A G T C A C G T A G T C A C G T A G T C A C G T G T A C C G A T G T C A | BPC6(BBRBPC)/col-BPC6-DAP-Seq(GSE60143)/Homer | 1e-72 | -1.676e+02 | 0.0000 | 210.0 | 1.40% | 129.8 | 0.19% | motif file (matrix) | svg |
| 195 | G C T A G C T A C T G A A C T G A C G T A G T C C G T A C G T A G T A C A C T G A T G C G C A T | WRKY47(WRKY)/colamp-WRKY47-DAP-Seq(GSE60143)/Homer | 1e-72 | -1.672e+02 | 0.0000 | 3892.0 | 26.01% | 13424.5 | 19.28% | motif file (matrix) | svg |
| 196 | C G T A A C G T A C G T A C G T A C G T A G T C A G T C C T G A A G C T A G C T | NFAT(RHD)/Jurkat-NFATC1-ChIP-Seq(Jolma\_et\_al.)/Homer | 1e-71 | -1.655e+02 | 0.0000 | 5804.0 | 38.78% | 21659.4 | 31.11% | motif file (matrix) | svg |
| 197 | C G T A G C A T C G T A C G T A G C A T A C T G C G A T A G T C A C T G A C T G G A C T C T A G | AT1G71450(AP2EREBP)/col-AT1G71450-DAP-Seq(GSE60143)/Homer | 1e-71 | -1.642e+02 | 0.0000 | 13168.0 | 87.99% | 57208.8 | 82.16% | motif file (matrix) | svg |
| 198 | C G A T C G T A G C T A G C A T G C A T C T G A A C T G A C G T A G T C C G T A C G T A G T A C T C A G G C T A C G A T | WRKY25(WRKY)/colamp-WRKY25-DAP-Seq(GSE60143)/Homer | 1e-71 | -1.641e+02 | 0.0000 | 9433.0 | 63.03% | 38353.6 | 55.08% | motif file (matrix) | svg |
| 199 | T A G C C A T G A G C T A C G T A C T G C G T A A G T C G A C T G C A T C T G A | AT3G42860(zfGRF)/col-AT3G42860-DAP-Seq(GSE60143)/Homer | 1e-70 | -1.632e+02 | 0.0000 | 3915.0 | 26.16% | 13574.5 | 19.49% | motif file (matrix) | svg |
| 200 | C G A T T C G A A G T C A C G T A C G T T A C G C G T A G C T A G C T A C G A T G C A T A T G C C G T A G T C A A C T G | ANAC071(NAC)/col-ANAC071-DAP-Seq(GSE60143)/Homer | 1e-70 | -1.624e+02 | 0.0000 | 8428.0 | 56.32% | 33631.4 | 48.30% | motif file (matrix) | svg |
| 201 | A C T G C G T A A C T G A T G C T G A C G A T C A T C G T G C A A C T G A G T C | ZNF519(Zf)/HEK293-ZNF519.GFP-ChIP-Seq(GSE58341)/Homer | 1e-70 | -1.621e+02 | 0.0000 | 1696.0 | 11.33% | 4775.5 | 6.86% | motif file (matrix) | svg |
| 202 | C T G A T A C G G C A T A G C T A G C T A G T C T C G A A C T G C A G T A G C T A G C T G A T C | IRF3(IRF)/BMDM-Irf3-ChIP-Seq(GSE67343)/Homer | 1e-69 | -1.612e+02 | 0.0000 | 1699.0 | 11.35% | 4794.6 | 6.89% | motif file (matrix) | svg |
| 203 | G A T C C T G A G A T C G A T C C T A G G C T A A G T C C T G A G C T A C G T A | At4g16750(AP2EREBP)/col-At4g16750-DAP-Seq(GSE60143)/Homer | 1e-69 | -1.608e+02 | 0.0000 | 11130.0 | 74.37% | 46693.4 | 67.06% | motif file (matrix) | svg |
| 204 | C G A T T C G A A T G C C G A T G C A T T C G A A G C T G C A T G C A T C G A T T C G A A G C T T C G A G T C A C T A G | ANAC004(NAC)/colamp-ANAC004-DAP-Seq(GSE60143)/Homer | 1e-69 | -1.605e+02 | 0.0000 | 2992.0 | 19.99% | 9816.3 | 14.10% | motif file (matrix) | svg |
| 205 | T G A C C G A T C T G A C T A G C T A G A C G T A T G C T G C A T C G A C T G A C T A G C A T G A C G T A G T C C G T A | PPARa(NR),DR1/Liver-Ppara-ChIP-Seq(GSE47954)/Homer | 1e-68 | -1.585e+02 | 0.0000 | 5518.0 | 36.87% | 20514.7 | 29.46% | motif file (matrix) | svg |
| 206 | A G T C A C G T A C G T T A C G G C T A G C T A C G T A C G A T C G A T A T G C C G T A G T C A A C T G G A C T G C A T | SND2(NAC)/colamp-SND2-DAP-Seq(GSE60143)/Homer | 1e-68 | -1.578e+02 | 0.0000 | 6314.0 | 42.19% | 24048.1 | 34.54% | motif file (matrix) | svg |
| 207 | C G A T T G C A T G C A G A T C C G T A A C T G T G A C G A C T C A T G A C T G | Tcf21(bHLH)/ArterySmoothMuscle-Tcf21-ChIP-Seq(GSE61369)/Homer | 1e-68 | -1.578e+02 | 0.0000 | 4898.0 | 32.73% | 17829.8 | 25.61% | motif file (matrix) | svg |
| 208 | T G A C C A T G A C G T A C G T A C T G C G T A A G T C A G C T G C A T T C G A | WRKY30(WRKY)/colamp-WRKY30-DAP-Seq(GSE60143)/Homer | 1e-68 | -1.573e+02 | 0.0000 | 4248.0 | 28.39% | 15057.1 | 21.62% | motif file (matrix) | svg |
| 209 | A G C T C A T G G C A T G A T C T G C A C T A G G A T C A C G T | Tgif2(Homeobox)/mES-Tgif2-ChIP-Seq(GSE55404)/Homer | 1e-68 | -1.567e+02 | 0.0000 | 13337.0 | 89.12% | 58235.4 | 83.63% | motif file (matrix) | svg |
| 210 | A G C T T G A C C G A T C G A T C T A G A C G T C A G T C A G T G C T A A G T C | FOXK1(Forkhead)/HEK293-FOXK1-ChIP-Seq(GSE51673)/Homer | 1e-66 | -1.536e+02 | 0.0000 | 7525.0 | 50.28% | 29606.1 | 42.52% | motif file (matrix) | svg |
| 211 | A T G C A G T C A C T G A T G C A G T C A C T G A G T C G T A C | SHN3(AP2EREBP)/col-SHN3-DAP-Seq(GSE60143)/Homer | 1e-66 | -1.536e+02 | 0.0000 | 2699.0 | 18.04% | 8724.0 | 12.53% | motif file (matrix) | svg |
| 212 | G A C T C G A T T C A G G A T C G A C T A G C T A G C T A G T C G A T C C G T A C T A G C T A G T C G A T C G A C T G A | Bcl6(Zf)/Liver-Bcl6-ChIP-Seq(GSE31578)/Homer | 1e-66 | -1.534e+02 | 0.0000 | 6033.0 | 40.31% | 22871.1 | 32.85% | motif file (matrix) | svg |
| 213 | C A T G A G T C G T C A C G T A A T G C A C G T A C G T A C T G | bHLH130(bHLH)/col-bHLH130-DAP-Seq(GSE60143)/Homer | 1e-66 | -1.528e+02 | 0.0000 | 6271.0 | 41.90% | 23941.0 | 34.38% | motif file (matrix) | svg |
| 214 | A C T G A C T G A G T C A C T G A C T G A G T C A C G T C T A G | ERF1(AP2EREBP)/colamp-ERF1-DAP-Seq(GSE60143)/Homer | 1e-66 | -1.525e+02 | 0.0000 | 5398.0 | 36.07% | 20085.9 | 28.85% | motif file (matrix) | svg |
| 215 | A C T G A C G T C G A T C A G T C A T G C A T G C A G T G C A T C A G T C A T G | HuR(?)/HEK293-HuR-CLIP-Seq(GSE87887)/Homer | 1e-66 | -1.521e+02 | 0.0000 | 12723.0 | 85.02% | 54999.0 | 78.98% | motif file (matrix) | svg |
| 216 | C T A G A G T C T A C G T A C G T G A C C G T A A C T G T A G C G C A T C A T G A T G C A G C T | Ascl1(bHLH)/NeuralTubes-Ascl1-ChIP-Seq(GSE55840)/Homer | 1e-64 | -1.492e+02 | 0.0000 | 7673.0 | 51.27% | 30365.8 | 43.61% | motif file (matrix) | svg |
| 217 | A G C T G A T C G A T C C G T A G T A C A G T C C G A T C T G A G T A C G A T C C G T A G A C T | ATY19(MYB)/col-ATY19-DAP-Seq(GSE60143)/Homer | 1e-64 | -1.489e+02 | 0.0000 | 6923.0 | 46.26% | 26943.8 | 38.69% | motif file (matrix) | svg |
| 218 | A G T C A G T C C T G A A G T C A G T C A C T G C G T A A G T C T C G A G A T C C G A T C G T A | AT1G01250(AP2EREBP)/col-AT1G01250-DAP-Seq(GSE60143)/Homer | 1e-64 | -1.479e+02 | 0.0000 | 2204.0 | 14.73% | 6836.5 | 9.82% | motif file (matrix) | svg |
| 219 | C T A G T C G A C G A T C T A G G C A T C A G T C T A G G A T C C G T A G T C A | CEBP:AP1(bZIP)/ThioMac-CEBPb-ChIP-Seq(GSE21512)/Homer | 1e-63 | -1.459e+02 | 0.0000 | 6556.0 | 43.81% | 25339.5 | 36.39% | motif file (matrix) | svg |
| 220 | T C G A T G A C G T A C C G T A C A G T T G A C A C G T A C T G A G C T A G C T | NeuroG2(bHLH)/Fibroblast-NeuroG2-ChIP-Seq(GSE75910)/Homer | 1e-62 | -1.448e+02 | 0.0000 | 8343.0 | 55.75% | 33553.0 | 48.19% | motif file (matrix) | svg |
| 221 | C G T A C G A T C A G T C A T G C G A T G T A C C A T G A C T G G A C T C A T G | CEJ1(AP2EREBP)/col-CEJ1-DAP-Seq(GSE60143)/Homer | 1e-62 | -1.436e+02 | 0.0000 | 11702.0 | 78.20% | 49860.9 | 71.61% | motif file (matrix) | svg |
| 222 | G C T A A C T G T C G A C T G A C T G A A C G T T A G C C T G A C G T A C G A T | Cux2(Homeobox)/Liver-Cux2-ChIP-Seq(GSE35985)/Homer | 1e-62 | -1.428e+02 | 0.0000 | 7540.0 | 50.38% | 29871.7 | 42.90% | motif file (matrix) | svg |
| 223 | A G T C C G T A A C G T A G T C A C G T A C T G | Tal1 | 1e-61 | -1.414e+02 | 0.0000 | 8334.0 | 55.69% | 33575.3 | 48.22% | motif file (matrix) | svg |
| 224 | G T C A T C G A C T A G C T A G A G T C G T C A C G A T C T A G G A C T G A T C G A T C T C A G C T A G C T G A A G T C G C T A C A G T T C A G G A T C G A T C | p63(p53)/Keratinocyte-p63-ChIP-Seq(GSE17611)/Homer | 1e-61 | -1.409e+02 | 0.0000 | 3302.0 | 22.06% | 11331.0 | 16.27% | motif file (matrix) | svg |
| 225 | C G A T C G A T G C A T G A C T A C G T C G T A C G T A A C T G T A G C C G T A C G T A C G T A | AT5G60130(ABI3VP1)/col-AT5G60130-DAP-Seq(GSE60143)/Homer | 1e-61 | -1.408e+02 | 0.0000 | 7815.0 | 52.22% | 31176.7 | 44.77% | motif file (matrix) | svg |
| 226 | C T G A C T G A C T G A A T G C G A T C C A T G A C T G G A C T G A C T G C A T C G T A C G T A A G T C G T A C C T G A A T C G G C A T G A C T G A C T A G C T | GRHL2(CP2)/HBE-GRHL2-ChIP-Seq(GSE46194)/Homer | 1e-61 | -1.407e+02 | 0.0000 | 3441.0 | 22.99% | 11908.5 | 17.10% | motif file (matrix) | svg |
| 227 | G T C A T G C A G C T A A G T C C G T A A C T G T G A C G C A T T C A G C A G T | Ap4(bHLH)/AML-Tfap4-ChIP-Seq(GSE45738)/Homer | 1e-60 | -1.399e+02 | 0.0000 | 5793.0 | 38.71% | 22037.5 | 31.65% | motif file (matrix) | svg |
| 228 | A G T C A C G T A C G T T C A G G C T A C G T A G C T A G C A T C G A T A G T C C G T A G T C A A C T G G A C T G C T A | SMB(NAC)/colamp-SMB-DAP-Seq(GSE60143)/Homer | 1e-60 | -1.393e+02 | 0.0000 | 8608.0 | 57.52% | 34897.1 | 50.12% | motif file (matrix) | svg |
| 229 | T C A G T A C G T A G C A C G T A C T G C G A T A G T C C G T A T A C G A G T C | Meis1(Homeobox)/MastCells-Meis1-ChIP-Seq(GSE48085)/Homer | 1e-60 | -1.386e+02 | 0.0000 | 10374.0 | 69.32% | 43357.2 | 62.27% | motif file (matrix) | svg |
| 230 | C G A T C G T A A G T C A C G T A C G T T C A G C G T A C G T A G C A T G C A T G C A T A G T C C G T A G T C A A C T G | VND2(NAC)/col-VND2-DAP-Seq(GSE60143)/Homer | 1e-59 | -1.380e+02 | 0.0000 | 8605.0 | 57.50% | 34907.4 | 50.13% | motif file (matrix) | svg |
| 231 | C G T A C G A T C T A G G T C A G A C T C G A T C T A G C G T A A C G T C A T G | LIN-39(Homeobox)/cElegans.L3-LIN39-ChIP-Seq(modEncode)/Homer | 1e-59 | -1.361e+02 | 0.0000 | 7899.0 | 52.78% | 31652.9 | 45.46% | motif file (matrix) | svg |
| 232 | C G A T C T A G A G T C A C G T A C G T T C A G G C T A C G T A G C A T G C A T C G A T A G T C C G T A G T C A A C T G | VND3(NAC)/colamp-VND3-DAP-Seq(GSE60143)/Homer | 1e-58 | -1.350e+02 | 0.0000 | 6419.0 | 42.89% | 24920.0 | 35.79% | motif file (matrix) | svg |
| 233 | A C T G A C T G A G T C A C T G A C T G A G T C A C G T T C A G | ERF2(AP2EREBP)/colamp-ERF2-DAP-Seq(GSE60143)/Homer | 1e-58 | -1.343e+02 | 0.0000 | 5752.0 | 38.44% | 21954.2 | 31.53% | motif file (matrix) | svg |
| 234 | C T G A T G A C T G A C C G T A A C G T T G A C A G C T C T A G A C G T G A C T | Olig2(bHLH)/Neuron-Olig2-ChIP-Seq(GSE30882)/Homer | 1e-57 | -1.333e+02 | 0.0000 | 10085.0 | 67.39% | 42054.3 | 60.39% | motif file (matrix) | svg |
| 235 | C T G A A G T C G A T C C A T G G C T A G A T C C T G A G C T A G C T A C G A T | AT1G77200(AP2EREBP)/colamp-AT1G77200-DAP-Seq(GSE60143)/Homer | 1e-57 | -1.322e+02 | 0.0000 | 11285.0 | 75.41% | 47968.0 | 68.89% | motif file (matrix) | svg |
| 236 | C G A T G A T C T A C G C T G A G C T A C G T A G C A T A G T C C T A G C G T A G C A T C G A T | AT2G15740(C2H2)/col-AT2G15740-DAP-Seq(GSE60143)/Homer | 1e-57 | -1.319e+02 | 0.0000 | 13165.0 | 87.97% | 57659.8 | 82.81% | motif file (matrix) | svg |
| 237 | C A T G A C T G C T A G T C G A T C G A T C G A T C G A T C A G T C A G T C A G T G A C T G A C C G T A A C T G T G C A C G A T A C T G | RBPJ:Ebox(?,bHLH)/Panc1-Rbpj1-ChIP-Seq(GSE47459)/Homer | 1e-57 | -1.319e+02 | 0.0000 | 1456.0 | 9.73% | 4151.1 | 5.96% | motif file (matrix) | svg |
| 238 | C T A G C A G T C G T A A C G T A G T C A C T G C G T A A G C T A G T C G A T C | HNF6(Homeobox)/Liver-Hnf6-ChIP-Seq(ERP000394)/Homer | 1e-56 | -1.301e+02 | 0.0000 | 9061.0 | 60.55% | 37211.2 | 53.44% | motif file (matrix) | svg |
| 239 | C G T A C T A G C A G T A C G T C G T A A C T G C A T G G C A T T C A G C T G A | MYB49(MYB)/col-MYB49-DAP-Seq(GSE60143)/Homer | 1e-55 | -1.283e+02 | 0.0000 | 8842.0 | 59.08% | 36213.8 | 52.01% | motif file (matrix) | svg |
| 240 | T C G A C G T A A G T C C G T A C T A G A G T C C G A T A C T G G A C T A G C T A C T G G A C T | HLH-1(bHLH)/cElegans-Embryo-HLH1-ChIP-Seq(modEncode)/Homer | 1e-55 | -1.276e+02 | 0.0000 | 5378.0 | 35.94% | 20421.1 | 29.33% | motif file (matrix) | svg |
| 241 | G C T A C G T A C G T A G C A T C A T G C T A G G A T C A C T G T C A G G A T C A C T G T C A G | ERF4(AP2EREBP)/colamp-ERF4-DAP-Seq(GSE60143)/Homer | 1e-55 | -1.269e+02 | 0.0000 | 10955.0 | 73.20% | 46430.8 | 66.68% | motif file (matrix) | svg |
| 242 | C A G T G C T A G C A T T A C G C T G A C A G T T A G C C T G A | GATA15(C2C2gata)/col-GATA15-DAP-Seq(GSE60143)/Homer | 1e-54 | -1.266e+02 | 0.0000 | 11901.0 | 79.53% | 51168.4 | 73.48% | motif file (matrix) | svg |
| 243 | A G T C C G A T A C T G A T C G T G A C G C T A C A T G A T C G T G A C C G A T A C T G T A G C G T A C G T C A | Tlx?(NR)/NPC-H3K4me1-ChIP-Seq(GSE16256)/Homer | 1e-54 | -1.244e+02 | 0.0000 | 2031.0 | 13.57% | 6417.6 | 9.22% | motif file (matrix) | svg |
| 244 | C T A G A C T G A G T C A C T G A C T G A G C T A C T G T C A G | AT3G57600(AP2EREBP)/col-AT3G57600-DAP-Seq(GSE60143)/Homer | 1e-53 | -1.242e+02 | 0.0000 | 6294.0 | 42.06% | 24559.7 | 35.27% | motif file (matrix) | svg |
| 245 | G A C T C T A G C T A G G T A C A G T C G A T C G A C T G A C T T A G C T C A G | NLP7(RWPRK)/col-NLP7-DAP-Seq(GSE60143)/Homer | 1e-53 | -1.226e+02 | 0.0000 | 10864.0 | 72.60% | 46063.2 | 66.15% | motif file (matrix) | svg |
| 246 | T C G A T G A C G C A T A G C T C A G T G A T C G C T A G A T C G A C T A C G T G C A T A G T C | PRDM1(Zf)/Hela-PRDM1-ChIP-Seq(GSE31477)/Homer | 1e-53 | -1.224e+02 | 0.0000 | 2996.0 | 20.02% | 10325.7 | 14.83% | motif file (matrix) | svg |
| 247 | G T C A C G T A A C G T A T C G C G T A A C G T A C G T C T A G | ATHB7(Homeobox)/col-ATHB7-DAP-Seq(GSE60143)/Homer | 1e-52 | -1.213e+02 | 0.0000 | 7040.0 | 47.04% | 27995.7 | 40.20% | motif file (matrix) | svg |
| 248 | C T G A G A C T C A T G C T A G A G T C A C T G A C T G A G T C A C T G T C A G | ERF11(AP2EREBP)/col-ERF11-DAP-Seq(GSE60143)/Homer | 1e-52 | -1.198e+02 | 0.0000 | 9687.0 | 64.73% | 40403.2 | 58.02% | motif file (matrix) | svg |
| 249 | A G C T C T G A C T A G C T A G A C T G T A G C T G C A T C G A C T G A C T A G C A T G A C G T A T G C T C G A | RXR(NR),DR1/3T3L1-RXR-ChIP-Seq(GSE13511)/Homer | 1e-51 | -1.189e+02 | 0.0000 | 5216.0 | 34.85% | 19865.3 | 28.53% | motif file (matrix) | svg |
| 250 | C A G T T C A G A G C T G A C T A C G T A G T C G A T C G A C T C T G A A C T G G A T C C G T A C T G A A G T C G T A C | Rfx6(HTH)/Min6b1-Rfx6.HA-ChIP-Seq(GSE62844)/Homer | 1e-50 | -1.167e+02 | 0.0000 | 7058.0 | 47.16% | 28169.0 | 40.45% | motif file (matrix) | svg |
| 251 | C G A T C G T A G C T A G A C T T C G A A G C T A G T C A C T G T C G A A G C T C T G A C G A T | ZBTB38(Zf)/Hela-ZBTB38-ChIP-seq(GSE108618)/Homer | 1e-50 | -1.163e+02 | 0.0000 | 14181.0 | 94.76% | 63532.2 | 91.24% | motif file (matrix) | svg |
| 252 | C G A T T C A G G T A C A C G T A C G T T C A G C G A T C G T A G T C A G C T A C G T A A G T C C G T A G T C A C A T G | ANAC057(NAC)/colamp-ANAC057-DAP-Seq(GSE60143)/Homer | 1e-49 | -1.151e+02 | 0.0000 | 7431.0 | 49.66% | 29912.0 | 42.96% | motif file (matrix) | svg |
| 253 | C G T A A C T G C G T A A C G T A T C G C A G T T A G C C G T A T C G A G T A C C T G A T A G C C G T A A C T G C G T A A C G T C G T A C T G A A T C G G C T A | GATA3(Zf),DR8/iTreg-Gata3-ChIP-Seq(GSE20898)/Homer | 1e-49 | -1.145e+02 | 0.0000 | 903.0 | 6.03% | 2304.6 | 3.31% | motif file (matrix) | svg |
| 254 | G A C T G C T A T G C A A G T C A C G T A C G T A C G T C G A T A C G T T A C G | At3g45610(C2C2dof)/col-At3g45610-DAP-Seq(GSE60143)/Homer | 1e-49 | -1.139e+02 | 0.0000 | 8508.0 | 56.85% | 34936.9 | 50.17% | motif file (matrix) | svg |
| 255 | T C G A T C G A A G T C C G T A C T A G T A G C A C G T A C T G | MyoG(bHLH)/C2C12-MyoG-ChIP-Seq(GSE36024)/Homer | 1e-49 | -1.135e+02 | 0.0000 | 5301.0 | 35.42% | 20341.1 | 29.21% | motif file (matrix) | svg |
| 256 | G C A T C G T A G C A T C G T A T C G A C G T A C T G A A C T G C G T A C G T A C G T A A C G T A C T G G T C A G C A T | AT2G31460(REMB3)/col-AT2G31460-DAP-Seq(GSE60143)/Homer | 1e-49 | -1.132e+02 | 0.0000 | 2378.0 | 15.89% | 7930.4 | 11.39% | motif file (matrix) | svg |
| 257 | T C A G T C A G A C G T G T A C G C T A T C A G C T G A A C T G A C T G A G C T A G T C C G T A | EAR2(NR)/K562-NR2F6-ChIP-Seq(Encode)/Homer | 1e-49 | -1.132e+02 | 0.0000 | 8292.0 | 55.41% | 33939.9 | 48.74% | motif file (matrix) | svg |
| 258 | G C T A A G T C T A C G T G C A A T C G T C A G G C T A T C G A T C A G A G C T | ELF5(ETS)/T47D-ELF5-ChIP-Seq(GSE30407)/Homer | 1e-49 | -1.130e+02 | 0.0000 | 5151.0 | 34.42% | 19688.2 | 28.27% | motif file (matrix) | svg |
| 259 | G C A T C T A G G T A C A G T C C G A T A C T G C T A G C T A G G T A C G C T A | ZNF416(Zf)/HEK293-ZNF416.GFP-ChIP-Seq(GSE58341)/Homer | 1e-48 | -1.122e+02 | 0.0000 | 5776.0 | 38.60% | 22474.4 | 32.28% | motif file (matrix) | svg |
| 260 | C T A G C T A G C G T A C G T A T A C G C G A T C T A G C T G A C T G A C G T A T A C G G A C T | PU.1:IRF8(ETS:IRF)/pDC-Irf8-ChIP-Seq(GSE66899)/Homer | 1e-48 | -1.118e+02 | 0.0000 | 1068.0 | 7.14% | 2907.6 | 4.18% | motif file (matrix) | svg |
| 261 | A C T G T C A G A G C T G A C T C A T G A G T C A G T C G C T A C G A T C T A G T C A G G T A C C T G A T C G A | Rfx1(HTH)/NPC-H3K4me1-ChIP-Seq(GSE16256)/Homer | 1e-47 | -1.105e+02 | 0.0000 | 1654.0 | 11.05% | 5106.6 | 7.33% | motif file (matrix) | svg |
| 262 | G C A T A C G T A C G T A T C G C G T A C G T A C G T A C G T A | At2g41835(C2H2)/col-At2g41835-DAP-Seq(GSE60143)/Homer | 1e-47 | -1.102e+02 | 0.0000 | 3225.0 | 21.55% | 11455.3 | 16.45% | motif file (matrix) | svg |
| 263 | T C A G C T G A C G T A C G T A T A C G G C A T C T A G C T G A C G T A C G T A T A C G G A C T | IRF1(IRF)/PBMC-IRF1-ChIP-Seq(GSE43036)/Homer | 1e-47 | -1.092e+02 | 0.0000 | 759.0 | 5.07% | 1847.7 | 2.65% | motif file (matrix) | svg |
| 264 | C T A G T C G A T G A C A G T C C G T A A C T G G T A C A C G T A C T G A C T G | BHLHA15(bHLH)/NIH3T3-BHLHB8.HA-ChIP-Seq(GSE119782)/Homer | 1e-46 | -1.066e+02 | 0.0000 | 7233.0 | 48.33% | 29179.5 | 41.90% | motif file (matrix) | svg |
| 265 | G T A C G C T A C G A T C A G T A G T C G C T A C G A T C G A T A G T C G C T A | WUS1(Homeobox)/colamp-WUS1-DAP-Seq(GSE60143)/Homer | 1e-46 | -1.062e+02 | 0.0000 | 3575.0 | 23.89% | 12995.5 | 18.66% | motif file (matrix) | svg |
| 266 | T G A C G C T A T C G A T G C A A G T C A G T C C G T A A G T C C G T A C T G A G C T A G T A C | RUNX2(Runt)/PCa-RUNX2-ChIP-Seq(GSE33889)/Homer | 1e-45 | -1.052e+02 | 0.0000 | 5636.0 | 37.66% | 21988.6 | 31.58% | motif file (matrix) | svg |
| 267 | A C G T T C G A T C G A A G T C G T C A T A C G A T G C A C G T A C T G A G C T | Myf5(bHLH)/GM-Myf5-ChIP-Seq(GSE24852)/Homer | 1e-45 | -1.051e+02 | 0.0000 | 3652.0 | 24.40% | 13339.2 | 19.16% | motif file (matrix) | svg |
| 268 | G A C T A C T G C G T A A G T C T C A G G C A T G T A C C G T A A C G T G A T C | TGA1(bZIP)/colamp-TGA1-DAP-Seq(GSE60143)/Homer | 1e-45 | -1.041e+02 | 0.0000 | 4999.0 | 33.40% | 19189.6 | 27.56% | motif file (matrix) | svg |
| 269 | G A C T G T A C T G C A A C G T G A T C G C T A T C G A A C G T A G T C C G T A | Pdx1(Homeobox)/Islet-Pdx1-ChIP-Seq(SRA008281)/Homer | 1e-45 | -1.039e+02 | 0.0000 | 7480.0 | 49.98% | 30374.6 | 43.62% | motif file (matrix) | svg |
| 270 | C A G T T C G A A G T C A C G T A C G T T C A G C G A T G C T A G C T A C G T A G C A T A G T C C G T A T G C A A C T G | ANAC045(NAC)/col-ANAC045-DAP-Seq(GSE60143)/Homer | 1e-44 | -1.029e+02 | 0.0000 | 12293.0 | 82.15% | 53599.8 | 76.97% | motif file (matrix) | svg |
| 271 | A C T G G A T C G A C T A C T G A C G T C A T G A C T G A C G T A G C T C G A T | RUNX-AML(Runt)/CD4+-PolII-ChIP-Seq(Barski\_et\_al.)/Homer | 1e-43 | -1.010e+02 | 0.0000 | 4568.0 | 30.52% | 17361.4 | 24.93% | motif file (matrix) | svg |
| 272 | T A C G T C G A G A C T A C T G C T G A A G T C T C A G G A C T T G A C C T G A | Atf1(bZIP)/K562-ATF1-ChIP-Seq(GSE31477)/Homer | 1e-43 | -1.004e+02 | 0.0000 | 7448.0 | 49.77% | 30303.3 | 43.52% | motif file (matrix) | svg |
| 273 | C A G T A T C G C T G A A G T C T C A G C A G T T A G C C T G A A T G C T A C G | FEA4(bZIP)/Corn-FEA4-ChIP-Seq(GSE61954)/Homer | 1e-43 | -9.962e+01 | 0.0000 | 9527.0 | 63.66% | 40067.0 | 57.54% | motif file (matrix) | svg |
| 274 | G T A C A C G T A C G T T C A G C G T A C G T A C G A T G C A T G C A T A G T C C G T A G T C A C A T G G A C T G C T A | VND1(NAC)/col-VND1-DAP-Seq(GSE60143)/Homer | 1e-43 | -9.910e+01 | 0.0000 | 7380.0 | 49.32% | 30019.6 | 43.11% | motif file (matrix) | svg |
| 275 | C G T A G C T A C G A T C T A G A C G T G T C A C G T A C G T A A G T C C G T A T G C A T A C G | FoxL2(Forkhead)/Ovary-FoxL2-ChIP-Seq(GSE60858)/Homer | 1e-42 | -9.757e+01 | 0.0000 | 5513.0 | 36.84% | 21597.9 | 31.02% | motif file (matrix) | svg |
| 276 | C G T A G C A T C A T G C T A G A G T C A C T G A T C G G T A C A C T G T C A G | At2g33710(AP2EREBP)/colamp-At2g33710-DAP-Seq(GSE60143)/Homer | 1e-42 | -9.699e+01 | 0.0000 | 12875.0 | 86.03% | 56706.3 | 81.44% | motif file (matrix) | svg |
| 277 | G A C T A G T C C G A T A C T G C T G A T G A C G T A C C G T A A T C G G C A T C T G A C T A G | Bcl11a(Zf)/HSPC-BCL11A-ChIP-Seq(GSE104676)/Homer | 1e-42 | -9.671e+01 | 0.0000 | 4424.0 | 29.56% | 16816.3 | 24.15% | motif file (matrix) | svg |
| 278 | A G T C A C G T A C T G A G C T A C G T A C G T G T C A A G T C | Foxo1(Forkhead)/RAW-Foxo1-ChIP-Seq(Fan\_et\_al.)/Homer | 1e-41 | -9.568e+01 | 0.0000 | 9416.0 | 62.92% | 39627.5 | 56.91% | motif file (matrix) | svg |
| 279 | G T A C A C G T A C G T T C A G G C T A C G T A C G A T G C A T G C A T A G T C C G T A G T C A C A T G G A C T G C T A | ANAC070(NAC)/colamp-ANAC070-DAP-Seq(GSE60143)/Homer | 1e-41 | -9.565e+01 | 0.0000 | 9255.0 | 61.84% | 38862.5 | 55.81% | motif file (matrix) | svg |
| 280 | T A C G T C G A C G T A C G T A C G T A C T G A A C T G A C G T C G T A T C G A | AT2G28810(C2C2dof)/colamp-AT2G28810-DAP-Seq(GSE60143)/Homer | 1e-41 | -9.534e+01 | 0.0000 | 11110.0 | 74.24% | 47832.9 | 68.69% | motif file (matrix) | svg |
| 281 | A T C G A G T C A C T G A G T C A G T C A C T G G A C T G A C T | PUCHI(AP2EREBP)/colamp-PUCHI-DAP-Seq(GSE60143)/Homer | 1e-41 | -9.523e+01 | 0.0000 | 8344.0 | 55.76% | 34579.2 | 49.66% | motif file (matrix) | svg |
| 282 | A C T G C T A G A G T C A C T G A C T G A G T C A C T G T A C G | ERF104(AP2EREBP)/col-ERF104-DAP-Seq(GSE60143)/Homer | 1e-40 | -9.354e+01 | 0.0000 | 8590.0 | 57.40% | 35771.3 | 51.37% | motif file (matrix) | svg |
| 283 | C G A T C A G T C T A G G C T A A G T C C G T A T C A G A G T C A C G T A C T G A C G T G T A C G C T A G C T A G C T A | bZIP52(bZIP)/colamp-bZIP52-DAP-Seq(GSE60143)/Homer | 1e-39 | -9.076e+01 | 0.0000 | 7946.0 | 53.10% | 32828.8 | 47.15% | motif file (matrix) | svg |
| 284 | G C T A C G T A C G T A G C A T C A T G C T A G A G T C A C T G T A C G A G T C C A T G T A C G | RAP26(AP2EREBP)/colamp-RAP26-DAP-Seq(GSE60143)/Homer | 1e-38 | -8.896e+01 | 0.0000 | 11588.0 | 77.43% | 50335.0 | 72.29% | motif file (matrix) | svg |
| 285 | G C T A T C G A C G T A C T A G A G C T G T C A G T C A C G T A A G T C C G T A | FOXA1(Forkhead)/LNCAP-FOXA1-ChIP-Seq(GSE27824)/Homer | 1e-38 | -8.854e+01 | 0.0000 | 6119.0 | 40.89% | 24511.3 | 35.20% | motif file (matrix) | svg |
| 286 | C G A T G A C T C G A T T C A G G A C T A C G T C A G T C T G A G A C T G A C T A G C T C G A T A C T G A T C G G T A C G C T A | NF1:FOXA1(CTF,Forkhead)/LNCAP-FOXA1-ChIP-Seq(GSE27824)/Homer | 1e-38 | -8.764e+01 | 0.0000 | 424.0 | 2.83% | 882.4 | 1.27% | motif file (matrix) | svg |
| 287 | C A T G T G C A G A C T C A T G C G T A A G T C T C A G G C A T T G A C C G T A | bZIP50(bZIP)/colamp-bZIP50-DAP-Seq(GSE60143)/Homer | 1e-37 | -8.747e+01 | 0.0000 | 11062.0 | 73.92% | 47770.5 | 68.60% | motif file (matrix) | svg |
| 288 | G T A C A C T G A T G C T G A C C T A G G A C T G T A C C G T A G C A T G C A T | ERF8(AP2EREBP)/colamp-ERF8-DAP-Seq(GSE60143)/Homer | 1e-37 | -8.693e+01 | 0.0000 | 11276.0 | 75.35% | 48835.8 | 70.13% | motif file (matrix) | svg |
| 289 | G A T C C G T A G A C T C T A G G A T C C T G A G A C T C T G A G A C T C T A G G A T C C T G A G A C T C T G A G A C T | OCT:OCT(POU,Homeobox)/NPC-OCT6-ChIP-Seq(GSE43916)/Homer | 1e-37 | -8.562e+01 | 0.0000 | 469.0 | 3.13% | 1034.7 | 1.49% | motif file (matrix) | svg |
| 290 | A G T C G A T C A G T C C G T A A T C G C A G T A G T C G T A C C T G A A C T G T C A G A G C T A G C T A G C T A G C T | PRDM15(Zf)/ESC-Prdm15-ChIP-Seq(GSE73694)/Homer | 1e-37 | -8.558e+01 | 0.0000 | 6443.0 | 43.05% | 26046.9 | 37.41% | motif file (matrix) | svg |
| 291 | T A C G T A G C C A T G C A G T A C G T C T A G C G T A A G T C G A C T G C A T G C A T C A G T | WRKY11(WRKY)/col-WRKY11-DAP-Seq(GSE60143)/Homer | 1e-36 | -8.515e+01 | 0.0000 | 1704.0 | 11.39% | 5598.4 | 8.04% | motif file (matrix) | svg |
| 292 | G T A C A C G T A C G T T A C G A T G C C A T G T A C G G T A C T C A G A T G C C G T A G T C A A C T G A G C T G C T A | AT1G19040(NAC)/col-AT1G19040-DAP-Seq(GSE60143)/Homer | 1e-36 | -8.486e+01 | 0.0000 | 1405.0 | 9.39% | 4428.2 | 6.36% | motif file (matrix) | svg |
| 293 | G T A C G T C A G T A C G T C A G T A C G T C A G T A C G T C A G T A C G T C A | SeqBias: CA-repeat | 1e-36 | -8.433e+01 | 0.0000 | 13816.0 | 92.32% | 61924.5 | 88.93% | motif file (matrix) | svg |
| 294 | A C G T T G A C A G T C A G C T A G T C A G C T A C T G G A C T A G C T G A C T | REF6(Zf)/Arabidopsis-REF6-ChIP-Seq(GSE106942)/Homer | 1e-36 | -8.373e+01 | 0.0000 | 3202.0 | 21.40% | 11801.2 | 16.95% | motif file (matrix) | svg |
| 295 | G A T C G A T C A G T C G T A C C G A T G T A C G T A C A G T C A G T C A G T C G C T A G A T C | ZNF148(Zf)/MDAMB231-ZNF148-ChIP-Seq(GSE147020)/Homer | 1e-36 | -8.296e+01 | 0.0000 | 1648.0 | 11.01% | 5404.2 | 7.76% | motif file (matrix) | svg |
| 296 | G A C T G A C T G A T C C G T A G T A C A G T C G C A T C G T A G T A C G A T C G C A T G C T A | MYB74(MYB)/colamp-MYB74-DAP-Seq(GSE60143)/Homer | 1e-35 | -8.101e+01 | 0.0000 | 5885.0 | 39.33% | 23630.8 | 33.94% | motif file (matrix) | svg |
| 297 | C T A G A C T G A C G T C G T A A C T G A C T G A G C T C T A G T C A G C T A G | MYB93(MYB)/colamp-MYB93-DAP-Seq(GSE60143)/Homer | 1e-35 | -8.094e+01 | 0.0000 | 9019.0 | 60.27% | 38096.9 | 54.71% | motif file (matrix) | svg |
| 298 | G C A T C G A T A T G C A G C T T C G A T A C G G C T A C G T A C A T G T G A C G C A T C G A T A G T C A G C T C G T A | AT3G09735(S1Falike)/col-AT3G09735-DAP-Seq(GSE60143)/Homer | 1e-35 | -8.066e+01 | 0.0000 | 2982.0 | 19.93% | 10924.7 | 15.69% | motif file (matrix) | svg |
| 299 | T G A C A G T C C G T A A C T G G T A C A C G T A C T G A C G T G A C T G A T C | Twist2(bHLH)/Myoblast-Twist2.Ty1-ChIP-Seq(GSE127998)/Homer | 1e-34 | -8.027e+01 | 0.0000 | 8646.0 | 57.77% | 36355.3 | 52.21% | motif file (matrix) | svg |
| 300 | G A T C C T G A A G T C A G C T A C G T A C G T A C G T A C G T | At1g64620(C2C2dof)/colamp-At1g64620-DAP-Seq(GSE60143)/Homer | 1e-34 | -7.985e+01 | 0.0000 | 8453.0 | 56.49% | 35460.6 | 50.92% | motif file (matrix) | svg |
| 301 | C T A G G T A C A C G T A C G T A T C G G C A T A G C T A G C T A G C T G C A T G A C T C G T A G T C A A C T G G A C T | VND6(NAC)/col-VND6-DAP-Seq(GSE60143)/Homer | 1e-34 | -7.937e+01 | 0.0000 | 9662.0 | 64.56% | 41190.7 | 59.15% | motif file (matrix) | svg |
| 302 | C G T A T A C G T C G A A C T G A C T G C G T A C G T A T A C G A G C T T A C G | PU.1(ETS)/ThioMac-PU.1-ChIP-Seq(GSE21512)/Homer | 1e-34 | -7.922e+01 | 0.0000 | 2674.0 | 17.87% | 9653.1 | 13.86% | motif file (matrix) | svg |
| 303 | C T A G T C G A C T G A C G T A T A C G G A C T T C A G T C G A G T C A T G C A T A C G A G C T | IRF2(IRF)/Erythroblas-IRF2-ChIP-Seq(GSE36985)/Homer | 1e-34 | -7.853e+01 | 0.0000 | 793.0 | 5.30% | 2192.7 | 3.15% | motif file (matrix) | svg |
| 304 | T G A C C T G A C T A G T C G A C T G A A T G C C G T A A C T G G C A T G T A C G C A T A T C G G C A T A G C T G A T C | PR(NR)/T47D-PR-ChIP-Seq(GSE31130)/Homer | 1e-33 | -7.787e+01 | 0.0000 | 10280.0 | 68.69% | 44193.9 | 63.47% | motif file (matrix) | svg |
| 305 | A C G T C T A G A G C T A C G T A C G T C T G A A G T C G A C T A G C T C G T A | FOXM1(Forkhead)/MCF7-FOXM1-ChIP-Seq(GSE72977)/Homer | 1e-33 | -7.778e+01 | 0.0000 | 5579.0 | 37.28% | 22331.1 | 32.07% | motif file (matrix) | svg |
| 306 | G C T A C G T A C G T A G C A T C A T G C T A G A G T C A C T G A C T G A G T C A C T G T C A G | ABR1(AP2EREBP)/colamp-ABR1-DAP-Seq(GSE60143)/Homer | 1e-33 | -7.766e+01 | 0.0000 | 10563.0 | 70.58% | 45567.7 | 65.44% | motif file (matrix) | svg |
| 307 | C G A T C T G A A G T C A C G T A C G T T C A G C G T A C G T A C G T A G C A T C G A T A G T C C G T A G T C A A C T G | VND4(NAC)/colamp-VND4-DAP-Seq(GSE60143)/Homer | 1e-33 | -7.702e+01 | 0.0000 | 7240.0 | 48.38% | 29899.4 | 42.94% | motif file (matrix) | svg |
| 308 | T C G A T A G C G T C A A C T G A C T G C G T A C G T A C T A G A G C T T C A G | ERG(ETS)/VCaP-ERG-ChIP-Seq(GSE14097)/Homer | 1e-33 | -7.691e+01 | 0.0000 | 7147.0 | 47.76% | 29474.2 | 42.33% | motif file (matrix) | svg |
| 309 | C G A T G C T A G C T A G C A T G C T A C G T A A G T C A C G T A C G T A C G T C G A T A G C T | At5g62940(C2C2dof)/col-At5g62940-DAP-Seq(GSE60143)/Homer | 1e-33 | -7.654e+01 | 0.0000 | 12999.0 | 86.86% | 57737.8 | 82.92% | motif file (matrix) | svg |
| 310 | C T G A C T G A C T A G T C G A C G T A A T G C C G T A A C T G C G T A A C G T C T G A C G A T A G C T C G T A A C G T A G T C C G A T T A C G G T C A G C A T | GATA(Zf),IR3/iTreg-Gata3-ChIP-Seq(GSE20898)/Homer | 1e-32 | -7.547e+01 | 0.0000 | 1604.0 | 10.72% | 5327.1 | 7.65% | motif file (matrix) | svg |
| 311 | C A T G C T A G A G T C A C T G A C T G G T A C C A T G T A C G | AT1G28160(AP2EREBP)/colamp-AT1G28160-DAP-Seq(GSE60143)/Homer | 1e-32 | -7.529e+01 | 0.0000 | 11961.0 | 79.93% | 52497.7 | 75.39% | motif file (matrix) | svg |
| 312 | C G T A C G T A C G T A C G T A C G T A A C T G A C T G A G T C | dof42(C2C2dof)/col-dof42-DAP-Seq(GSE60143)/Homer | 1e-32 | -7.488e+01 | 0.0000 | 5083.0 | 33.97% | 20183.2 | 28.99% | motif file (matrix) | svg |
| 313 | A T G C T C A G T C G A G C A T A C T G C G T A A G T C T C A G G A C T T G A C C G T A A G C T | Atf2(bZIP)/3T3L1-Atf2-ChIP-Seq(GSE56872)/Homer | 1e-31 | -7.353e+01 | 0.0000 | 2913.0 | 19.47% | 10762.1 | 15.46% | motif file (matrix) | svg |
| 314 | C G A T T A C G T G C A G T A C G A T C G A C T A G C T A C G T A T C G G T A C G A T C G T A C G A T C G T C A | PPARE(NR),DR1/3T3L1-Pparg-ChIP-Seq(GSE13511)/Homer | 1e-31 | -7.326e+01 | 0.0000 | 4460.0 | 29.80% | 17462.8 | 25.08% | motif file (matrix) | svg |
| 315 | G A T C G A T C G A T C C G T A G T A C A G T C G C A T C G T A G T A C G A T C | MYB58(MYB)/colamp-MYB58-DAP-Seq(GSE60143)/Homer | 1e-31 | -7.311e+01 | 0.0000 | 8511.0 | 56.87% | 35906.3 | 51.56% | motif file (matrix) | svg |
| 316 | T C A G G A C T G T C A C G T A A C G T A T C G C G T A A C G T A C G T C T G A | ATHB15(HB)/col-ATHB15-DAP-Seq(GSE60143)/Homer | 1e-31 | -7.303e+01 | 0.0000 | 3457.0 | 23.10% | 13097.2 | 18.81% | motif file (matrix) | svg |
| 317 | C T A G C A T G G A C T C G T A C T A G A C T G A C G T C T A G C T A G T C A G | MYB17(MYB)/colamp-MYB17-DAP-Seq(GSE60143)/Homer | 1e-31 | -7.236e+01 | 0.0000 | 5479.0 | 36.61% | 22013.1 | 31.61% | motif file (matrix) | svg |
| 318 | C G A T C A G T C A G T C A T G G T C A G A T C C G T A T C A G A G T C A C G T C T A G A C G T G T A C G T C A G C T A | VIP1(bZIP)/col-VIP1-DAP-Seq(GSE60143)/Homer | 1e-31 | -7.155e+01 | 0.0000 | 1321.0 | 8.83% | 4259.3 | 6.12% | motif file (matrix) | svg |
| 319 | T A C G T G C A A G T C C G T A A C G T T G A C A C G T A C T G A C T G G C A T | TCF4(bHLH)/SHSY5Y-TCF4-ChIP-Seq(GSE96915)/Homer | 1e-31 | -7.155e+01 | 0.0000 | 7899.0 | 52.78% | 33090.7 | 47.52% | motif file (matrix) | svg |
| 320 | C G A T C T G A A G T C A C G T A C G T T C A G G C A T C G A T G C T A G C T A C G T A A G T C C G T A G T C A A C T G | CUC1(NAC)/col-CUC1-DAP-Seq(GSE60143)/Homer | 1e-30 | -7.086e+01 | 0.0000 | 5066.0 | 33.85% | 20203.5 | 29.01% | motif file (matrix) | svg |
| 321 | C G A T C T G A G T A C A C G T A C G T T C A G C G T A C G T A G C T A G C A T G C A T A G T C C G T A G T C A C A T G | NST1(NAC)/colamp-NST1-DAP-Seq(GSE60143)/Homer | 1e-30 | -7.080e+01 | 0.0000 | 7398.0 | 49.44% | 30789.1 | 44.22% | motif file (matrix) | svg |
| 322 | G C T A C G T A C G A T G A C T G C A T T G C A A G T C A G C T A C G T A C G T C G A T G A C T | DAG2(C2C2dof)/col-DAG2-DAP-Seq(GSE60143)/Homer | 1e-30 | -7.045e+01 | 0.0000 | 8368.0 | 55.92% | 35307.4 | 50.70% | motif file (matrix) | svg |
| 323 | A G T C C T G A A T C G A G C T A G C T G A C T A G T C G C T A A C G T C G A T G C A T C G A T A T C G C G T A T A G C G C A T A T G C C G T A | bZIP:IRF(bZIP,IRF)/Th17-BatF-ChIP-Seq(GSE39756)/Homer | 1e-30 | -7.014e+01 | 0.0000 | 1997.0 | 13.34% | 6995.9 | 10.05% | motif file (matrix) | svg |
| 324 | C T A G A G T C A G C T A C T G C G T A C A G T C G T A C T G A T A G C T G A C | Unknown5/Drosophila-Promoters/Homer | 1e-30 | -6.969e+01 | 0.0000 | 7431.0 | 49.66% | 30971.1 | 44.48% | motif file (matrix) | svg |
| 325 | G C A T C G T A C G A T G A C T A C T G C T G A G A C T G A T C | Hnf6b(Homeobox)/LNCaP-Hnf6b-ChIP-Seq(GSE106305)/Homer | 1e-30 | -6.942e+01 | 0.0000 | 9441.0 | 63.09% | 40395.5 | 58.01% | motif file (matrix) | svg |
| 326 | C T G A A G T C C G A T A G C T A T G C G T A C A C G T A T C G C A G T G C A T | Elf4(ETS)/BMDM-Elf4-ChIP-Seq(GSE88699)/Homer | 1e-30 | -6.916e+01 | 0.0000 | 6415.0 | 42.87% | 26325.2 | 37.81% | motif file (matrix) | svg |
| 327 | C G A T T C G A A C T G G T C A C G T A C G A T G T A C G A C T | At3g04030(G2like)/col-At3g04030-DAP-Seq(GSE60143)/Homer | 1e-29 | -6.890e+01 | 0.0000 | 7643.0 | 51.07% | 31973.9 | 45.92% | motif file (matrix) | svg |
| 328 | A T G C G A C T A C G T C T A G A C G T A C G T A C G T C T G A G A T C G C T A A G C T C G T A | Foxa2(Forkhead)/Liver-Foxa2-ChIP-Seq(GSE25694)/Homer | 1e-29 | -6.878e+01 | 0.0000 | 5408.0 | 36.14% | 21783.5 | 31.28% | motif file (matrix) | svg |
| 329 | C T G A T C A G C T G A C T A G C A T G A C G T A T G C C G T A A T G C G C A T T C A G C T G A A C T G A C G T C A G T A G T C C G T A C A G T C T A G C A T G | VDR(NR),DR3/GM10855-VDR+vitD-ChIP-Seq(GSE22484)/Homer | 1e-29 | -6.813e+01 | 0.0000 | 1595.0 | 10.66% | 5392.4 | 7.74% | motif file (matrix) | svg |
| 330 | T C G A T G A C G T A C C G T A A C G T G A C T A C G T A C T G A C T G A G C T | Mesp1(bHLH)/ESC-Mesp1-ChIP-Seq(GSE165102)/Homer | 1e-29 | -6.793e+01 | 0.0000 | 4117.0 | 27.51% | 16076.6 | 23.09% | motif file (matrix) | svg |
| 331 | G C A T G C A T G A T C G C A T T C G A A C T G C G T A C G T A A T C G T A G C C G A T A C G T A G T C A G C T C G T A | HSF6(HSF)/col-HSF6-DAP-Seq(GSE60143)/Homer | 1e-29 | -6.680e+01 | 0.0000 | 1632.0 | 10.91% | 5560.2 | 7.99% | motif file (matrix) | svg |
| 332 | A T C G T G A C A T G C C T G A T C A G G A C T A G T C C G A T T C A G T C G A C A T G C T A G C T A G C G T A C T A G C T A G C T G A C T A G C T A G A T G C | ZSCAN22(Zf)/HEK293-ZSCAN22.GFP-ChIP-Seq(GSE58341)/Homer | 1e-28 | -6.641e+01 | 0.0000 | 319.0 | 2.13% | 663.1 | 0.95% | motif file (matrix) | svg |
| 333 | C A G T C A T G T G C A G T A C C G T A T C A G G T A C G A C T T C A G C T G A | bZIP18(bZIP)/colamp-bZIP18-DAP-Seq(GSE60143)/Homer | 1e-28 | -6.628e+01 | 0.0000 | 14121.0 | 94.36% | 63885.4 | 91.75% | motif file (matrix) | svg |
| 334 | C T G A A C T G C G T A A C G T G T C A A G C T A G C T G A C T G A C T C A G T | CCA(Myb)/Arabidopsis-CCA.GFP-ChIP-Seq(GSE70533)/Homer | 1e-28 | -6.595e+01 | 0.0000 | 7284.0 | 48.67% | 30394.1 | 43.65% | motif file (matrix) | svg |
| 335 | A G T C T A G C G A C T A C G T C T A G A C G T A C G T A C G T C T G A A G T C G C T A G A C T C G T A C T A G A C T G | Foxa3(Forkhead)/Liver-Foxa3-ChIP-Seq(GSE77670)/Homer | 1e-28 | -6.551e+01 | 0.0000 | 2353.0 | 15.72% | 8549.2 | 12.28% | motif file (matrix) | svg |
| 336 | G C A T G A T C T C A G G C T A G A C T A G T C C T A G C G T A C A T G G T C A | GATA20(C2C2gata)/colamp-GATA20-DAP-Seq(GSE60143)/Homer | 1e-27 | -6.404e+01 | 0.0000 | 13663.0 | 91.30% | 61451.4 | 88.25% | motif file (matrix) | svg |
| 337 | A T G C G A C T A G C T C T A G C G T A C T A G C G A T C T A G A T C G G A T C | Nkx2.2(Homeobox)/NPC-Nkx2.2-ChIP-Seq(GSE61673)/Homer | 1e-27 | -6.394e+01 | 0.0000 | 11160.0 | 74.57% | 48820.5 | 70.11% | motif file (matrix) | svg |
| 338 | T C A G A G C T A C G T A C G T G T A C G A T C C G T A C T A G C A T G G T C A C G T A T C G A | STAT4(Stat)/CD4-Stat4-ChIP-Seq(GSE22104)/Homer | 1e-27 | -6.350e+01 | 0.0000 | 5181.0 | 34.62% | 20900.9 | 30.02% | motif file (matrix) | svg |
| 339 | A G T C G A C T C A G T A C T G C T A G T G A C G C T A A T G C G C A T A T C G C G A T A C T G G A T C G T A C G T C A C T G A | NF1(CTF)/LNCAP-NF1-ChIP-Seq(Unpublished)/Homer | 1e-27 | -6.349e+01 | 0.0000 | 2055.0 | 13.73% | 7343.5 | 10.55% | motif file (matrix) | svg |
| 340 | T C G A G C A T A C G T C T A G G T A C T C G A G C A T T G A C T C G A A C G T | Chop(bZIP)/MEF-Chop-ChIP-Seq(GSE35681)/Homer | 1e-27 | -6.267e+01 | 0.0000 | 2462.0 | 16.45% | 9059.9 | 13.01% | motif file (matrix) | svg |
| 341 | T A C G A C T G A G C T G T A C C G T A T C G A C T G A A C T G C A T G A C G T A G T C C G T A | COUP-TFII(NR)/K562-NR2F1-ChIP-Seq(Encode)/Homer | 1e-26 | -6.200e+01 | 0.0000 | 8598.0 | 57.45% | 36620.0 | 52.59% | motif file (matrix) | svg |
| 342 | C T A G T A G C A T G C C T A G A G T C A G T C C T A G G A C T G A C T G C T A | CRF10(AP2EREBP)/col100-CRF10-DAP-Seq(GSE60143)/Homer | 1e-26 | -6.173e+01 | 0.0000 | 11629.0 | 77.71% | 51178.9 | 73.50% | motif file (matrix) | svg |
| 343 | T C G A A C T G A C T G C G T A C G T A T C G A A G T C C T G A A T C G G T A C G C A T C A T G | ETS:E-box(ETS,bHLH)/HPC7-Scl-ChIP-Seq(GSE22178)/Homer | 1e-26 | -6.073e+01 | 0.0000 | 480.0 | 3.21% | 1226.3 | 1.76% | motif file (matrix) | svg |
| 344 | G A C T C T A G G A T C C A G T A C T G C T G A A T G C G C A T A T G C C T G A | MafA(bZIP)/Islet-MafA-ChIP-Seq(GSE30298)/Homer | 1e-26 | -6.012e+01 | 0.0000 | 5571.0 | 37.23% | 22742.9 | 32.66% | motif file (matrix) | svg |
| 345 | T A G C C T A G T C G A G A C T A C T G C T G A A G T C T C A G G C A T T G A C C T G A A G C T | Atf7(bZIP)/3T3L1-Atf7-ChIP-Seq(GSE56872)/Homer | 1e-25 | -5.920e+01 | 0.0000 | 4487.0 | 29.98% | 17918.8 | 25.73% | motif file (matrix) | svg |
| 346 | T C G A C G T A C G T A T C G A A C T G G T A C C G T A A G C T G T C A G C A T | At3g24120(G2like)/col-At3g24120-DAP-Seq(GSE60143)/Homer | 1e-25 | -5.894e+01 | 0.0000 | 13698.0 | 91.53% | 61733.1 | 88.65% | motif file (matrix) | svg |
| 347 | C G T A G A C T C G T A A C G T C A G T A G T C A G C T G A C T | KAN2(G2like)/colamp-KAN2-DAP-Seq(GSE60143)/Homer | 1e-25 | -5.830e+01 | 0.0000 | 8767.0 | 58.58% | 37522.4 | 53.89% | motif file (matrix) | svg |
| 348 | G A T C G C A T G C A T A G T C A G C T T C G A T A C G G C T A C G T A C T A G T G A C G C A T C G A T G A T C A G C T | HSFC1(HSF)/col-HSFC1-DAP-Seq(GSE60143)/Homer | 1e-25 | -5.791e+01 | 0.0000 | 1278.0 | 8.54% | 4267.9 | 6.13% | motif file (matrix) | svg |
| 349 | T G C A C T G A A T G C G T C A A C G T A T G C A C G T A C T G A C T G T G C A | ZBTB18(Zf)/HEK293-ZBTB18.GFP-ChIP-Seq(GSE58341)/Homer | 1e-25 | -5.766e+01 | 0.0000 | 2855.0 | 19.08% | 10830.7 | 15.55% | motif file (matrix) | svg |
| 350 | G C A T G C A T G T A C G A C T T C G A A C T G G C T A C G T A A T C G T G A C G C A T G A C T A G T C A G C T C T G A | AGL95(ND)/col-AGL95-DAP-Seq(GSE60143)/Homer | 1e-24 | -5.749e+01 | 0.0000 | 736.0 | 4.92% | 2175.8 | 3.12% | motif file (matrix) | svg |
| 351 | A T G C T C G A A G T C A G C T A C G T G T A C A G T C G C T A C T A G C A T G G T C A C T G A T C A G A G T C | Stat3+il21(Stat)/CD4-Stat3-ChIP-Seq(GSE19198)/Homer | 1e-24 | -5.732e+01 | 0.0000 | 4143.0 | 27.68% | 16444.1 | 23.62% | motif file (matrix) | svg |
| 352 | T C G A G A C T T C A G T G C A G T A C G T A C A G C T G T A C C A T G T C G A C A T G C A T G A C G T A G T C C T G A | FXR(NR),ER2/Liver-FXR-ChIP-Seq(GSE133700)/Homer | 1e-24 | -5.618e+01 | 0.0000 | 3786.0 | 25.30% | 14902.4 | 21.40% | motif file (matrix) | svg |
| 353 | T C G A A G C T A C G T A C G T A G T C A G T C A C G T A T C G G A C T A T C G | EWS:ERG-fusion(ETS)/CADO\_ES1-EWS:ERG-ChIP-Seq(SRA014231)/Homer | 1e-24 | -5.591e+01 | 0.0000 | 3483.0 | 23.27% | 13584.7 | 19.51% | motif file (matrix) | svg |
| 354 | A G T C C G T A T G A C A T G C G C A T C T G A G T A C G A T C | MYB55(MYB)/colamp-MYB55-DAP-Seq(GSE60143)/Homer | 1e-24 | -5.553e+01 | 0.0000 | 9272.0 | 61.96% | 39988.6 | 57.43% | motif file (matrix) | svg |
| 355 | C G T A A C T G C G T A A C G T C A G T A G T C A G C T G C A T G C T A C G A T | At2g01060(G2like)/colamp-At2g01060-DAP-Seq(GSE60143)/Homer | 1e-23 | -5.509e+01 | 0.0000 | 13659.0 | 91.27% | 61603.9 | 88.47% | motif file (matrix) | svg |
| 356 | A G T C G A T C A G C T C G T A G T A C A G T C G C A T C T G A G T A C G A T C | AT4G26030(C2H2)/col-AT4G26030-DAP-Seq(GSE60143)/Homer | 1e-23 | -5.402e+01 | 0.0000 | 8442.0 | 56.41% | 36126.8 | 51.88% | motif file (matrix) | svg |
| 357 | A T G C A G T C C T G A A G T C C G A T A C G T A G T C A G T C A C G T A T C G G A C T A C G T | Etv2(ETS)/ES-ER71-ChIP-Seq(GSE59402)/Homer | 1e-23 | -5.396e+01 | 0.0000 | 4665.0 | 31.17% | 18847.9 | 27.07% | motif file (matrix) | svg |
| 358 | G A C T C A G T G A T C G A T C A C G T G A T C C T G A T A C G C G T A G T C A | STAT6(Stat)/Macrophage-Stat6-ChIP-Seq(GSE38377)/Homer | 1e-23 | -5.328e+01 | 0.0000 | 3246.0 | 21.69% | 12615.0 | 18.12% | motif file (matrix) | svg |
| 359 | A C G T C T A G C G T A A G T C G T A C A C G T A C G T A C G T G T C A G T A C T G A C G A C T | Nur77(NR)/K562-NR4A1-ChIP-Seq(GSE31363)/Homer | 1e-22 | -5.230e+01 | 0.0000 | 1475.0 | 9.86% | 5144.4 | 7.39% | motif file (matrix) | svg |
| 360 | C G T A C T G A C G T A C T A G T C G A C T A G A C T G C G T A C G T A T A C G A G C T A T C G | SpiB(ETS)/OCILY3-SPIB-ChIP-Seq(GSE56857)/Homer | 1e-22 | -5.190e+01 | 0.0000 | 1372.0 | 9.17% | 4732.6 | 6.80% | motif file (matrix) | svg |
| 361 | C G T A C G T A C T G A A C T G A C G T A G T C C G T A C G T A A G T C A C T G A T G C G A T C | WRKY46(WRKY)/colamp-WRKY46-DAP-Seq(GSE60143)/Homer | 1e-22 | -5.163e+01 | 0.0000 | 1345.0 | 8.99% | 4627.1 | 6.64% | motif file (matrix) | svg |
| 362 | T A G C G T A C A G T C G T A C C G A T A G T C A G T C A G T C A G T C A G T C C G T A G A T C | Zfp281(Zf)/ES-Zfp281-ChIP-Seq(GSE81042)/Homer | 1e-22 | -5.156e+01 | 0.0000 | 347.0 | 2.32% | 837.8 | 1.20% | motif file (matrix) | svg |
| 363 | G A C T G A T C C T G A A G T C A G T C A C T G C G T A A G T C G T A C G C T A G C A T C G A T | At1g19210(AP2EREBP)/colamp-At1g19210-DAP-Seq(GSE60143)/Homer | 1e-22 | -5.155e+01 | 0.0000 | 12512.0 | 83.61% | 55823.5 | 80.17% | motif file (matrix) | svg |
| 364 | A G C T G T C A T G C A A G T C A C G T A C G T A C G T C G A T G A C T T A C G | AT3G12130(C3H)/colamp-AT3G12130-DAP-Seq(GSE60143)/Homer | 1e-22 | -5.125e+01 | 0.0000 | 11376.0 | 76.02% | 50222.2 | 72.12% | motif file (matrix) | svg |
| 365 | C A T G C T A G A G C T G A C T C A T G A G T C G A T C G C T A C G A T C T A G T C A G G T A C C T G A T C G A | X-box(HTH)/NPC-H3K4me1-ChIP-Seq(GSE16256)/Homer | 1e-22 | -5.095e+01 | 0.0000 | 725.0 | 4.84% | 2201.6 | 3.16% | motif file (matrix) | svg |
| 366 | C T G A C G A T C T A G C G T A A G C T C G A T C A G T C T G A G A C T C T A G C T A G A T G C | PBX2(Homeobox)/K562-PBX2-ChIP-Seq(Encode)/Homer | 1e-21 | -5.060e+01 | 0.0000 | 7237.0 | 48.36% | 30629.8 | 43.99% | motif file (matrix) | svg |
| 367 | C G A T G T C A A G T C A C G T A C G T A C T G G A C T C G A T A T C G G C T A G T C A A G T C C G T A G T C A A C T G | ANAC017(NAC)/colamp-ANAC017-DAP-Seq(GSE60143)/Homer | 1e-21 | -5.009e+01 | 0.0000 | 1466.0 | 9.80% | 5143.0 | 7.39% | motif file (matrix) | svg |
| 368 | A G C T G C T A T G C A A G T C A C G T A C G T A C G T C G A T A G C T T C A G | dof24(C2C2dof)/col-dof24-DAP-Seq(GSE60143)/Homer | 1e-21 | -5.003e+01 | 0.0000 | 10392.0 | 69.44% | 45496.7 | 65.34% | motif file (matrix) | svg |
| 369 | G C A T G C A T G A T C G A C T T C G A T C A G G C T A C G T A A C T G G T A C G C A T G C A T A G T C A G C T C G T A | HSF7(HSF)/colamp-HSF7-DAP-Seq(GSE60143)/Homer | 1e-21 | -4.960e+01 | 0.0000 | 1231.0 | 8.23% | 4198.7 | 6.03% | motif file (matrix) | svg |
| 370 | G C T A T C G A C G T A C T A G A G C T G T C A G T C A C G T A A G T C C G T A | FOXA1(Forkhead)/MCF7-FOXA1-ChIP-Seq(GSE26831)/Homer | 1e-21 | -4.951e+01 | 0.0000 | 4731.0 | 31.61% | 19266.2 | 27.67% | motif file (matrix) | svg |
| 371 | C T G A A T C G A G C T A G C T A C G T T A G C C T G A T A C G C G A T A C G T G A C T A G T C | ISRE(IRF)/ThioMac-LPS-Expression(GSE23622)/Homer | 1e-21 | -4.867e+01 | 0.0000 | 336.0 | 2.25% | 819.8 | 1.18% | motif file (matrix) | svg |
| 372 | A C T G A G T C G T C A C G T A A G T C C G T A C T A G C T A G G A C T C A T G | SCRT1(Zf)/HEK293-SCRT1.eGFP-ChIP-Seq(Encode)/Homer | 1e-20 | -4.688e+01 | 0.0000 | 3062.0 | 20.46% | 11970.2 | 17.19% | motif file (matrix) | svg |
| 373 | C G T A C G T A G C A T A C T G C G T A A G C T C T G A C G T A T A C G C T G A | ELT-3(Gata)/cElegans-L1-ELT3-ChIP-Seq(modEncode)/Homer | 1e-20 | -4.658e+01 | 0.0000 | 4035.0 | 26.96% | 16250.4 | 23.34% | motif file (matrix) | svg |
| 374 | G A C T G A T C A G T C C G T A T G A C A G T C G C A T C T G A G T A C G A T C G C A T G A C T | MYB10(MYB)/col-MYB10-DAP-Seq(GSE60143)/Homer | 1e-19 | -4.588e+01 | 0.0000 | 4143.0 | 27.68% | 16749.9 | 24.05% | motif file (matrix) | svg |
| 375 | A T G C T A G C A G C T A G C T T G A C G A C T T C A G T A C G G T C A C T G A A T C G T A G C G A C T C A G T A G T C A G C T T C G A A T C G T G C A T G C A | HRE(HSF)/HepG2-HSF1-ChIP-Seq(GSE31477)/Homer | 1e-19 | -4.551e+01 | 0.0000 | 956.0 | 6.39% | 3160.8 | 4.54% | motif file (matrix) | svg |
| 376 | G C T A C G T A A C T G C G T A C G A T A C G T A G T C A G C T | At3g12730(G2like)/colamp-At3g12730-DAP-Seq(GSE60143)/Homer | 1e-19 | -4.507e+01 | 0.0000 | 10464.0 | 69.92% | 45998.1 | 66.06% | motif file (matrix) | svg |
| 377 | T C G A G C A T A C T G C T G A A G T C T C A G G A C T G T A C C G T A A G C T A G T C G A T C | c-Jun-CRE(bZIP)/K562-cJun-ChIP-Seq(GSE31477)/Homer | 1e-19 | -4.442e+01 | 0.0000 | 2171.0 | 14.51% | 8194.9 | 11.77% | motif file (matrix) | svg |
| 378 | T G C A C G T A A C T G T C A G C A G T C A T G T C A G G A T C T A C G A G T C T G C A A C T G A C T G T G A C G T C A | ZNF165(Zf)/WHIM12-ZNF165-ChIP-Seq(GSE65937)/Homer | 1e-19 | -4.415e+01 | 0.0000 | 597.0 | 3.99% | 1791.1 | 2.57% | motif file (matrix) | svg |
| 379 | C A T G G T C A A G T C C G T A C T A G G A T C C G A T A C T G A C G T G T A C C G T A C G T A | bZIP69(bZIP)/col-bZIP69-DAP-Seq(GSE60143)/Homer | 1e-18 | -4.326e+01 | 0.0000 | 779.0 | 5.21% | 2496.1 | 3.58% | motif file (matrix) | svg |
| 380 | G C A T T C A G C T G A A T C G A C T G C G A T G A T C C T G A | THRb(NR)/Liver-NR1A2-ChIP-Seq(GSE52613)/Homer | 1e-18 | -4.316e+01 | 0.0000 | 13185.0 | 88.11% | 59426.8 | 85.34% | motif file (matrix) | svg |
| 381 | C G T A C T G A T C A G A C T G G T C A C G T A A C G T G T A C C G A T G C A T | AT5G45580(G2like)/colamp-AT5G45580-DAP-Seq(GSE60143)/Homer | 1e-18 | -4.255e+01 | 0.0000 | 10644.0 | 71.13% | 46945.4 | 67.42% | motif file (matrix) | svg |
| 382 | T G A C G C T A T G A C C G T A T C A G G A T C C G T A C A T G C A T G C T A G C T A G C T A G | Unknown-ESC-element(?)/mES-Nanog-ChIP-Seq(GSE11724)/Homer | 1e-18 | -4.186e+01 | 0.0000 | 2420.0 | 16.17% | 9316.5 | 13.38% | motif file (matrix) | svg |
| 383 | T G C A C G T A G T C A A G C T A G T C G C T A T A G C C G A T C T A G G A T C | Gfi1b(Zf)/HPC7-Gfi1b-ChIP-Seq(GSE22178)/Homer | 1e-18 | -4.151e+01 | 0.0000 | 4182.0 | 27.95% | 17050.9 | 24.49% | motif file (matrix) | svg |
| 384 | C T A G A C T G A C G T C G T A A C T G A C T G A C G T T C A G C T G A T C G A | MYB107(MYB)/col-MYB107-DAP-Seq(GSE60143)/Homer | 1e-17 | -4.093e+01 | 0.0000 | 10705.0 | 71.53% | 47292.2 | 67.92% | motif file (matrix) | svg |
| 385 | C T A G C A T G C A T G T A C G A G T C G C A T A G C T C T A G A C G T A G T C G A C T A C T G A C T G A C T G T C G A | Zfp809(Zf)/ES-Zfp809-ChIP-Seq(GSE70799)/Homer | 1e-17 | -4.093e+01 | 0.0000 | 982.0 | 6.56% | 3328.4 | 4.78% | motif file (matrix) | svg |
| 386 | T G C A A G C T A C G T C T A G G A T C C T A G G A T C G T C A C T G A A G T C | CEBP(bZIP)/ThioMac-CEBPb-ChIP-Seq(GSE21512)/Homer | 1e-17 | -4.085e+01 | 0.0000 | 6769.0 | 45.23% | 28796.2 | 41.35% | motif file (matrix) | svg |
| 387 | C T A G T C G A A C G T A C G T C A T G A G T C C T G A C G A T A G T C C G T A | AARE(HLH)/mES-cMyc-ChIP-Seq/Homer | 1e-17 | -4.043e+01 | 0.0000 | 1194.0 | 7.98% | 4192.0 | 6.02% | motif file (matrix) | svg |
| 388 | C G T A T C G A G A T C G C A T C G T A A C G T G T A C T C A G G T C A G A C T C G T A C T A G | DREF/Drosophila-Promoters/Homer | 1e-17 | -4.026e+01 | 0.0000 | 953.0 | 6.37% | 3222.3 | 4.63% | motif file (matrix) | svg |
| 389 | A T G C A G T C G T A C A G C T T C G A C T A G G A T C C T G A G T C A A G T C G C T A T C A G | Rfx5(HTH)/GM12878-Rfx5-ChIP-Seq(GSE31477)/Homer | 1e-17 | -3.928e+01 | 0.0000 | 2592.0 | 17.32% | 10121.8 | 14.54% | motif file (matrix) | svg |
| 390 | G A T C A G T C G A C T G C T A G T A C A G T C G C A T G C T A G T A C G A T C | MYB61(MYB)/colamp-MYB61-DAP-Seq(GSE60143)/Homer | 1e-16 | -3.899e+01 | 0.0000 | 10284.0 | 68.72% | 45340.8 | 65.11% | motif file (matrix) | svg |
| 391 | C G A T C T A G T C A G C A G T C G T A A G T C G C T A A C G T G A C T A T G C A G T C G C T A | PRDM10(Zf)/HEK293-PRDM10.eGFP-ChIP-Seq(Encode)/Homer | 1e-16 | -3.869e+01 | 0.0000 | 3878.0 | 25.91% | 15783.3 | 22.67% | motif file (matrix) | svg |
| 392 | C T A G C T G A C G T A C G T A C G T A C G T A A C T G A C G T C T A G G T C A | COG1(C2C2dof)/col-COG1-DAP-Seq(GSE60143)/Homer | 1e-16 | -3.804e+01 | 0.0000 | 7923.0 | 52.94% | 34240.3 | 49.17% | motif file (matrix) | svg |
| 393 | T C G A T C A G T C G A A C T G C A T G A C G T A G T C C T G A | COUP-TFII(NR)/Artia-Nr2f2-ChIP-Seq(GSE46497)/Homer | 1e-16 | -3.755e+01 | 0.0000 | 9837.0 | 65.73% | 43261.3 | 62.13% | motif file (matrix) | svg |
| 394 | T C G A C T G A C G T A C G T A C G T A C T G A A C T G A C G T C T G A C T G A | AT5G63260(C3H)/col-AT5G63260-DAP-Seq(GSE60143)/Homer | 1e-16 | -3.719e+01 | 0.0000 | 10867.0 | 72.62% | 48199.2 | 69.22% | motif file (matrix) | svg |
| 395 | A C T G A G C T A G T C G T C A A G C T T C A G A T G C G A T C G C A T A T C G T C G A T A G C C G A T C A T G T A G C | Pax8(Paired,Homeobox)/Thyroid-Pax8-ChIP-Seq(GSE26938)/Homer | 1e-16 | -3.714e+01 | 0.0000 | 1932.0 | 12.91% | 7338.5 | 10.54% | motif file (matrix) | svg |
| 396 | A G T C C T A G A T C G C A G T C G A T A G C T G T A C A C T G C A T G C A T G | ZBED2(Zf)/SUIT2-ZBED2.HA-ChIP-Seq(GSE141606)/Homer | 1e-16 | -3.692e+01 | 0.0000 | 7993.0 | 53.41% | 34608.6 | 49.70% | motif file (matrix) | svg |
| 397 | A T G C A G C T T C A G T G A C T C A G A T G C T G C A A C G T A T C G G A T C A C T G A G T C | NRF1(NRF)/MCF7-NRF1-ChIP-Seq(Unpublished)/Homer | 1e-15 | -3.667e+01 | 0.0000 | 779.0 | 5.21% | 2582.9 | 3.71% | motif file (matrix) | svg |
| 398 | G T A C G T A C G T C A G C T A C G T A C G T A C G T A C T A G C T A G C T A G | SEP3(MADS)/Arabidoposis-Flower-Sep3-ChIP-Seq/Homer | 1e-15 | -3.631e+01 | 0.0000 | 6782.0 | 45.32% | 29020.7 | 41.68% | motif file (matrix) | svg |
| 399 | C G A T T C G A G T A C A C G T A C G T T C A G G C A T G C T A T G C A G C A T C G T A A G T C C G T A T G A C C A T G | ANAC092(NAC)/colamp-ANAC092-DAP-Seq(GSE60143)/Homer | 1e-15 | -3.608e+01 | 0.0000 | 4979.0 | 33.27% | 20802.8 | 29.87% | motif file (matrix) | svg |
| 400 | T C G A G T A C T C G A T C G A C A T G A T G C A C G T A C T G A C T G A G T C C G T A C T A G A G T C A T C G A G T C | Unknown3/Drosophila-Promoters/Homer | 1e-15 | -3.598e+01 | 0.0000 | 825.0 | 5.51% | 2774.9 | 3.99% | motif file (matrix) | svg |
| 401 | C G T A T G A C T C G A A G T C C G T A A T C G A T G C A C G T A C T G A G T C | E2A(bHLH)/proBcell-E2A-ChIP-Seq(GSE21978)/Homer | 1e-15 | -3.568e+01 | 0.0000 | 6553.0 | 43.79% | 27990.7 | 40.20% | motif file (matrix) | svg |
| 402 | T C A G A C T G A C G T C G T A A C T G A C T G A C G T C T A G | MYB51(MYB)/col-MYB51-DAP-Seq(GSE60143)/Homer | 1e-15 | -3.556e+01 | 0.0000 | 7722.0 | 51.60% | 33399.0 | 47.96% | motif file (matrix) | svg |
| 403 | G C A T G A C T A T G C A G C T T C G A A C T G C G T A C G T A A T C G A T G C G C A T G A C T G A T C A G C T T C G A | HSFA1E(HSF)/col-HSFA1E-DAP-Seq(GSE60143)/Homer | 1e-15 | -3.541e+01 | 0.0000 | 491.0 | 3.28% | 1485.5 | 2.13% | motif file (matrix) | svg |
| 404 | C G T A A C T G G T C A A C G T A T C G C A G T C T A G T C A G C G T A A C T G C G T A A C G T C G T A C T G A T A C G | GATA3(Zf),DR4/iTreg-Gata3-ChIP-Seq(GSE20898)/Homer | 1e-15 | -3.527e+01 | 0.0000 | 848.0 | 5.67% | 2876.4 | 4.13% | motif file (matrix) | svg |
| 405 | C A T G G A T C C T G A A G T C C T A G C G T A G C T A G C A T G A T C G A T C A G T C C T A G C G T A C A T G C T A G | PLT1(AP2EREBP)/colamp-PLT1-DAP-Seq(GSE60143)/Homer | 1e-15 | -3.515e+01 | 0.0000 | 1427.0 | 9.54% | 5250.6 | 7.54% | motif file (matrix) | svg |
| 406 | C G T A G A C T C G A T A T C G G T A C G C A T C A T G C G T A T A C G G C A T G T A C C G T A C A T G A T G C G C T A C T A G G C A T G C A T G C A T G A C T | MafB(bZIP)/BMM-Mafb-ChIP-Seq(GSE75722)/Homer | 1e-15 | -3.475e+01 | 0.0000 | 2286.0 | 15.28% | 8913.1 | 12.80% | motif file (matrix) | svg |
| 407 | C T A G T A C G G A T C G T A C G C T A A G C T A G C T G T C A T C G A T A G C | Nanog(Homeobox)/mES-Nanog-ChIP-Seq(GSE11724)/Homer | 1e-15 | -3.464e+01 | 0.0000 | 14118.0 | 94.34% | 64438.2 | 92.54% | motif file (matrix) | svg |
| 408 | C G A T C G A T G C A T G C A T G T C A A G T C A G C T A C G T A C G T C G A T G A C T A C G T | OBP4(C2C2dof)/col-OBP4-DAP-Seq(GSE60143)/Homer | 1e-14 | -3.373e+01 | 0.0000 | 7788.0 | 52.04% | 33776.2 | 48.51% | motif file (matrix) | svg |
| 409 | T C G A T A G C T G C A A C T G A C T G C G T A C G T A C T A G G A C T T A C G | ETS1(ETS)/Jurkat-ETS1-ChIP-Seq(GSE17954)/Homer | 1e-14 | -3.346e+01 | 0.0000 | 6105.0 | 40.80% | 26019.7 | 37.37% | motif file (matrix) | svg |
| 410 | T C G A G A C T A T C G C G T A A G T C C T A G G C A T G T A C C T G A A C G T G A T C G C T A | TGA4(bZIP)/colamp-TGA4-DAP-Seq(GSE60143)/Homer | 1e-14 | -3.339e+01 | 0.0000 | 3640.0 | 24.32% | 14891.3 | 21.39% | motif file (matrix) | svg |
| 411 | C A T G C T G A A G T C A C T G A C T G A G C T A C T G A T C G | ESE3(AP2EREBP)/col-ESE3-DAP-Seq(GSE60143)/Homer | 1e-14 | -3.320e+01 | 0.0000 | 10892.0 | 72.78% | 48462.0 | 69.60% | motif file (matrix) | svg |
| 412 | A T C G A G T C A G T C C G T A A C T G G C A T | hINR(CPE) | 1e-13 | -3.181e+01 | 0.0000 | 7566.0 | 50.56% | 32821.8 | 47.14% | motif file (matrix) | svg |
| 413 | A G T C C G T A C G A T A G T C G T C A A G T C A C G T C T G A | Unknown2/Drosophila-Promoters/Homer | 1e-13 | -3.114e+01 | 0.0000 | 7049.0 | 47.10% | 30453.2 | 43.73% | motif file (matrix) | svg |
| 414 | T A G C T A G C G A C T C T A G A G C T A G T C G T C A T G C A A C G T A T G C G C T A T G C A | Pbx3(Homeobox)/GM12878-PBX3-ChIP-Seq(GSE32465)/Homer | 1e-13 | -3.076e+01 | 0.0000 | 1710.0 | 11.43% | 6540.0 | 9.39% | motif file (matrix) | svg |
| 415 | T A C G A T G C G A C T A C T G A G C T A G T C G T C A T G C A A C G T A G T C G C T A T G C A | Pknox1(Homeobox)/ES-Prep1-ChIP-Seq(GSE63282)/Homer | 1e-13 | -3.042e+01 | 0.0000 | 1876.0 | 12.54% | 7258.2 | 10.42% | motif file (matrix) | svg |
| 416 | G A C T T C G A C G T A C G T A C G T A C G T A C G T A C T A G A G C T C G T A | dof45(C2C2dof)/col-dof45-DAP-Seq(GSE60143)/Homer | 1e-12 | -2.963e+01 | 0.0000 | 11308.0 | 75.56% | 50598.5 | 72.66% | motif file (matrix) | svg |
| 417 | G A C T C A G T A G C T C G A T A G T C G A T C A G T C C G T A A T G C T C A G | Rbpj1(?)/Panc1-Rbpj1-ChIP-Seq(GSE47459)/Homer | 1e-12 | -2.920e+01 | 0.0000 | 6824.0 | 45.60% | 29494.7 | 42.36% | motif file (matrix) | svg |
| 418 | T C A G C T G A C T A G C A T G A C G T A T G C C T G A C T G A C T G A C T A G C A T G A C G T A T G C C T G A | TR4(NR),DR1/Hela-TR4-ChIP-Seq(GSE24685)/Homer | 1e-12 | -2.843e+01 | 0.0000 | 518.0 | 3.46% | 1669.8 | 2.40% | motif file (matrix) | svg |
| 419 | C T A G T A C G G A T C G T C A T G C A A C G T T G C A G C T A T C G A T G C A | Hoxa9(Homeobox)/ChickenMSG-Hoxa9.Flag-ChIP-Seq(GSE86088)/Homer | 1e-12 | -2.821e+01 | 0.0000 | 12344.0 | 82.49% | 55682.7 | 79.97% | motif file (matrix) | svg |
| 420 | T A C G T C A G T G C A A G C T T G A C A G C T A G T C A C T G G A T C A C T G T C G A A C T G C T G A C T G A A T G C | ZBTB33(Zf)/GM12878-ZBTB33-ChIP-Seq(GSE32465)/Homer | 1e-12 | -2.788e+01 | 0.0000 | 1200.0 | 8.02% | 4449.1 | 6.39% | motif file (matrix) | svg |
| 421 | T G C A C T G A C A T G C T A G C A G T A G T C C G T A A T G C A T G C T A C G G C A T T C A G G T C A G A T C G T A C | ERE(NR),IR3/MCF7-ERa-ChIP-Seq(Unpublished)/Homer | 1e-11 | -2.712e+01 | 0.0000 | 1743.0 | 11.65% | 6771.3 | 9.72% | motif file (matrix) | svg |
| 422 | C G T A C T A G G A C T G T C A G T C A C G T A A G T C C G T A T C G A T C G A T C G A C G T A C T G A C T A G G C T A C G T A T A G C C G T A C G A T C G T A | FOXA1:AR(Forkhead,NR)/LNCAP-AR-ChIP-Seq(GSE27824)/Homer | 1e-11 | -2.702e+01 | 0.0000 | 216.0 | 1.44% | 561.9 | 0.81% | motif file (matrix) | svg |
| 423 | G C A T G C T A G C T A C G T A G C A T G C T A C T A G C G T A C G T A A C T G C G T A C G A T A C G T A G T C G A C T | At1g68670(G2like)/colamp-At1g68670-DAP-Seq(GSE60143)/Homer | 1e-11 | -2.681e+01 | 0.0000 | 3505.0 | 23.42% | 14513.3 | 20.84% | motif file (matrix) | svg |
| 424 | C A G T A G C T C G T A G C A T A G T C G A C T C T A G C T A G C A G T C T A G T C G A T G C A C T A G C A T G G A C T | STOP1(C2H2)/colamp-STOP1-DAP-Seq(GSE60143)/Homer | 1e-11 | -2.574e+01 | 0.0000 | 3211.0 | 21.46% | 13242.8 | 19.02% | motif file (matrix) | svg |
| 425 | T C A G G C A T A C T G C G T A A G T C C T A G G C A T T G A C | TGA9(bZIP)/colamp-TGA9-DAP-Seq(GSE60143)/Homer | 1e-11 | -2.570e+01 | 0.0000 | 10579.0 | 70.69% | 47260.6 | 67.87% | motif file (matrix) | svg |
| 426 | C T G A A G C T A C G T A C G T A G T C G A C T G A C T C T G A C T G A C T A G C G T A C G T A | STAT6(Stat)/CD4-Stat6-ChIP-Seq(GSE22104)/Homer | 1e-11 | -2.562e+01 | 0.0000 | 2931.0 | 19.59% | 12004.9 | 17.24% | motif file (matrix) | svg |
| 427 | C T G A T A C G G C A T C T A G A T G C G A T C C G A T A C T G C T A G G A T C C T G A A T G C | MYRF(MYRF)/CFPAC1-MYRF-ChIP-Seq(GSE145627)/Homer | 1e-10 | -2.525e+01 | 0.0000 | 2271.0 | 15.18% | 9112.8 | 13.09% | motif file (matrix) | svg |
| 428 | T G C A A G C T C A T G C G T A A G C T A C T G G A T C G T C A C G T A A G C T | Atf4(bZIP)/MEF-Atf4-ChIP-Seq(GSE35681)/Homer | 1e-10 | -2.516e+01 | 0.0000 | 3061.0 | 20.45% | 12596.8 | 18.09% | motif file (matrix) | svg |
| 429 | C G T A G C A T C A G T C T A G A G T C A C T G A C T G G T A C A C T G A T C G | ERF115(AP2EREBP)/colamp-ERF115-DAP-Seq(GSE60143)/Homer | 1e-10 | -2.476e+01 | 0.0000 | 12182.0 | 81.40% | 55017.7 | 79.01% | motif file (matrix) | svg |
| 430 | A C T G T G A C A C T G A C G T A C G T A C T G C G T A A G T C A G C T C G A T G C A T A C G T | WRKY17(WRKY)/colamp-WRKY17-DAP-Seq(GSE60143)/Homer | 1e-10 | -2.463e+01 | 0.0000 | 186.0 | 1.24% | 475.5 | 0.68% | motif file (matrix) | svg |
| 431 | C T G A T C A G G C T A A G C T A G T C G A C T C T G A C T A G T G C A C T G A A G T C G T A C G A T C A C T G T C G A | ZBTB12(Zf)/HEK293-ZBTB12.GFP-ChIP-Seq(GSE58341)/Homer | 1e-10 | -2.457e+01 | 0.0000 | 3267.0 | 21.83% | 13533.4 | 19.44% | motif file (matrix) | svg |
| 432 | G T A C C G T A C G T A T A C G G C A T G T A C C G T A C A T G A G T C C G T A C G T A C G A T G C A T G C A T G A C T | MafF(bZIP)/HepG2-MafF-ChIP-Seq(GSE31477)/Homer | 1e-10 | -2.412e+01 | 0.0000 | 1861.0 | 12.44% | 7361.8 | 10.57% | motif file (matrix) | svg |
| 433 | G A C T A T C G C T G A A G T C T C A G G A C T G T A C C T G A A G C T G T A C | TGA6(bZIP)/colamp-TGA6-DAP-Seq(GSE60143)/Homer | 1e-10 | -2.373e+01 | 0.0000 | 7565.0 | 50.55% | 33169.4 | 47.63% | motif file (matrix) | svg |
| 434 | T A G C G C T A T C G A C T G A A G T C A G T C C T G A A G T C C G T A C T A G | RUNX(Runt)/HPC7-Runx1-ChIP-Seq(GSE22178)/Homer | 1e-9 | -2.299e+01 | 0.0000 | 5327.0 | 35.60% | 22896.1 | 32.88% | motif file (matrix) | svg |
| 435 | G C A T G A C T T G A C A G C T T C G A A C T G C G T A C T G A T C A G T A G C C G A T C G A T G T A C A G C T T C G A | AT1G23810(Orphan)/col-AT1G23810-DAP-Seq(GSE60143)/Homer | 1e-9 | -2.226e+01 | 0.0000 | 245.0 | 1.64% | 704.0 | 1.01% | motif file (matrix) | svg |
| 436 | T C G A A C G T A C T G C G T A A G T C C T A G A G C T T G A C | TGA10(bZIP)/colamp-TGA10-DAP-Seq(GSE60143)/Homer | 1e-9 | -2.222e+01 | 0.0000 | 7173.0 | 47.93% | 31423.9 | 45.13% | motif file (matrix) | svg |
| 437 | A G C T G C A T A C T G A C G T A G T C A C G T C T A G T A C G | Smad3(MAD)/NPC-Smad3-ChIP-Seq(GSE36673)/Homer | 1e-9 | -2.219e+01 | 0.0000 | 11832.0 | 79.06% | 53424.0 | 76.72% | motif file (matrix) | svg |
| 438 | C G T A A T G C C G A T A C G T A G T C C G T A C G T A C G T A C T A G A T C G | TCFL2(HMG)/K562-TCF7L2-ChIP-Seq(GSE29196)/Homer | 1e-9 | -2.150e+01 | 0.0000 | 607.0 | 4.06% | 2123.2 | 3.05% | motif file (matrix) | svg |
| 439 | A G T C G A C T A C T G G A T C G T A C C G T A T G A C A G T C C G A T A G C T A C G T A C G T C T A G G A C T C T G A | ZNF7(Zf)/HepG2-ZNF7.Flag-ChIP-Seq(Encode)/Homer | 1e-9 | -2.097e+01 | 0.0000 | 3663.0 | 24.48% | 15446.7 | 22.18% | motif file (matrix) | svg |
| 440 | G A C T A G C T G T A C G A C T C T G A A C T G G T C A C T G A A T G C T A C G G A C T A C G T A G T C G A C T C T G A | HRE(HSF)/Striatum-HSF1-ChIP-Seq(GSE38000)/Homer | 1e-9 | -2.095e+01 | 0.0000 | 1049.0 | 7.01% | 3966.7 | 5.70% | motif file (matrix) | svg |
| 441 | C G T A C G T A G C T A C G T A G A T C C T G A A C G T A C G T A G T C A G C T G C A T G C A T | AT2G40260(G2like)/colamp-AT2G40260-DAP-Seq(GSE60143)/Homer | 1e-8 | -2.040e+01 | 0.0000 | 9054.0 | 60.50% | 40291.3 | 57.86% | motif file (matrix) | svg |
| 442 | T G C A G C A T C G A T C G T A C A G T A C T G G T A C C G T A C T G A A G C T G T C A A C T G C T A G G T C A C G A T A C T G G T A C T G C A C G T A A G C T | CEBP:CEBP(bZIP)/MEF-Chop-ChIP-Seq(GSE35681)/Homer | 1e-8 | -2.031e+01 | 0.0000 | 967.0 | 6.46% | 3635.1 | 5.22% | motif file (matrix) | svg |
| 443 | C G A T C T A G A C G T G T C A C G T A C G T A A G T C C G T A | Foxo3(Forkhead)/U2OS-Foxo3-ChIP-Seq(E-MTAB-2701)/Homer | 1e-8 | -2.031e+01 | 0.0000 | 5027.0 | 33.59% | 21652.9 | 31.10% | motif file (matrix) | svg |
| 444 | C T A G T A C G G A C T T G C A T G C A C G A T T A C G C T G A T C G A C T G A | Hoxa10(Homeobox)/ChickenMSG-Hoxa10.Flag-ChIP-Seq(GSE86088)/Homer | 1e-8 | -1.998e+01 | 0.0000 | 4028.0 | 26.92% | 17133.1 | 24.60% | motif file (matrix) | svg |
| 445 | C G T A C G T A C G T A C T G A C T A G A C G T C T A G G T C A | CDF3(C2C2dof)/colamp-CDF3-DAP-Seq(GSE60143)/Homer | 1e-8 | -1.983e+01 | 0.0000 | 9174.0 | 61.30% | 40884.3 | 58.71% | motif file (matrix) | svg |
| 446 | G A T C G C A T T C G A A G T C A C G T A C G T A C G T C G A T A C G T A T C G | AT1G47655(C2C2dof)/colamp-AT1G47655-DAP-Seq(GSE60143)/Homer | 1e-8 | -1.983e+01 | 0.0000 | 12909.0 | 86.26% | 58757.5 | 84.38% | motif file (matrix) | svg |
| 447 | G A C T A G T C G A T C C G T A G T A C A G T C G C A T C G T A G T C A G A T C | MYB67(MYB)/col-MYB67-DAP-Seq(GSE60143)/Homer | 1e-8 | -1.948e+01 | 0.0000 | 8006.0 | 53.50% | 35434.2 | 50.89% | motif file (matrix) | svg |
| 448 | C G T A C G T A C T A G A C G T A C G T C G T A A C T G A C T G A C G T C T G A T C G A T C G A | MYB4(MYB)/col200-MYB4-DAP-Seq(GSE60143)/Homer | 1e-8 | -1.921e+01 | 0.0000 | 5468.0 | 36.54% | 23720.4 | 34.06% | motif file (matrix) | svg |
| 449 | A T G C C G T A C G T A C G T A C G T A C G T A A C T G A C G T C G A T C T G A | dof43(C2C2dof)/colamp-dof43-DAP-Seq(GSE60143)/Homer | 1e-8 | -1.910e+01 | 0.0000 | 8294.0 | 55.42% | 36798.8 | 52.85% | motif file (matrix) | svg |
| 450 | A G C T G C A T G T C A C G A T T A G C C G T A A C G T G C T A | CRC(C2C2YABBY)/col-CRC-DAP-Seq(GSE60143)/Homer | 1e-8 | -1.900e+01 | 0.0000 | 9565.0 | 63.92% | 42768.5 | 61.42% | motif file (matrix) | svg |
| 451 | G C A T G A C T A T G C A G C T T C G A C T A G G C T A C G T A C A T G T G A C G C A T G A C T A G T C A G C T C T G A | HSFB4(HSF)/col-HSFB4-DAP-Seq(GSE60143)/Homer | 1e-8 | -1.895e+01 | 0.0000 | 199.0 | 1.33% | 567.8 | 0.82% | motif file (matrix) | svg |
| 452 | C T A G C T G A A G T C G C T A C G A T A C T G G A C T G A T C G A T C C T G A C T A G C T G A T G A C G C T A C G A T T C A G G A C T G A T C G A T C T G A C | p53(p53)/Saos-p53-ChIP-Seq(GSE15780)/Homer | 1e-8 | -1.880e+01 | 0.0000 | 821.0 | 5.49% | 3056.6 | 4.39% | motif file (matrix) | svg |
| 453 | C T A G C T G A A G T C G C T A C G A T A C T G G A C T G A T C G A T C C T G A C T A G C T G A T G A C G C T A C G A T T C A G G A C T G A T C G A T C T G A C | p53(p53)/Saos-p53-ChIP-Seq/Homer | 1e-8 | -1.880e+01 | 0.0000 | 821.0 | 5.49% | 3056.6 | 4.39% | motif file (matrix) | svg |
| 454 | T C G A C A T G C T G A C G T A A T C G G T A C G C A T C G A T A G T C A G C T T C G A T A C G C G T A C G T A C A T G | HSFA6A(HSF)/col-HSFA6A-DAP-Seq(GSE60143)/Homer | 1e-8 | -1.879e+01 | 0.0000 | 187.0 | 1.25% | 525.1 | 0.75% | motif file (matrix) | svg |
| 455 | G T A C G A T C C A G T A G T C A G T C A G T C T G C A G A T C C T G A A T G C G T C A A C G T | WT1(Zf)/Kidney-WT1-ChIP-Seq(GSE90016)/Homer | 1e-8 | -1.859e+01 | 0.0000 | 2805.0 | 18.74% | 11703.1 | 16.81% | motif file (matrix) | svg |
| 456 | C T G A C T A G T C G A C G T A A T G C C G T A A T C G C G A T T A G C G C A T A T C G G C A T A G C T G A T C G A C T A G C T | ARE(NR)/LNCAP-AR-ChIP-Seq(GSE27824)/Homer | 1e-8 | -1.844e+01 | 0.0000 | 1335.0 | 8.92% | 5250.3 | 7.54% | motif file (matrix) | svg |
| 457 | C G T A A G T C T G A C A G C T A C G T C G T A A C G T A G T C | At5g05790(MYBrelated)/col-At5g05790-DAP-Seq(GSE60143)/Homer | 1e-8 | -1.842e+01 | 0.0000 | 8897.0 | 59.45% | 39657.2 | 56.95% | motif file (matrix) | svg |
| 458 | T A C G C G T A T C A G G A C T C T A G A C T G C A G T T A G C T C G A A C G T G T A C C T A G A G T C A G T C G A T C | ZNF669(Zf)/HEK293-ZNF669.GFP-ChIP-Seq(GSE58341)/Homer | 1e-7 | -1.827e+01 | 0.0000 | 1229.0 | 8.21% | 4799.1 | 6.89% | motif file (matrix) | svg |
| 459 | G C A T C T A G A C T G A C G T C G T A A C T G A C T G C G A T C T A G T C G A T C G A G C T A | MYB40(MYB)/col-MYB40-DAP-Seq(GSE60143)/Homer | 1e-7 | -1.803e+01 | 0.0000 | 3290.0 | 21.98% | 13896.1 | 19.96% | motif file (matrix) | svg |
| 460 | C T A G T A C G A G T C C G T A A G T C A C G T A G T C T C G A C G T A T A C G | Nkx2.1(Homeobox)/LungAC-Nkx2.1-ChIP-Seq(GSE43252)/Homer | 1e-7 | -1.763e+01 | 0.0000 | 12613.0 | 84.28% | 57401.7 | 82.43% | motif file (matrix) | svg |
| 461 | T C G A C T G A C G T A C G T A C G T A C G T A A C T G A C G T C G A T C T G A | BBX31(Orphan)/col-BBX31-DAP-Seq(GSE60143)/Homer | 1e-7 | -1.756e+01 | 0.0000 | 8671.0 | 57.94% | 38645.6 | 55.50% | motif file (matrix) | svg |
| 462 | T A C G T A C G C T A G T C A G A G T C C G T A A T C G A T G C A C G T A C T G A G T C G A C T | Ascl2(bHLH)/ESC-Ascl2-ChIP-Seq(GSE97712)/Homer | 1e-7 | -1.751e+01 | 0.0000 | 5448.0 | 36.40% | 23716.2 | 34.06% | motif file (matrix) | svg |
| 463 | G C T A C G T A A C G T A T C G C G T A A C G T A C G T C T A G | ATHB6(Homeobox)/col-ATHB6-DAP-Seq(GSE60143)/Homer | 1e-7 | -1.681e+01 | 0.0000 | 9017.0 | 60.25% | 40309.9 | 57.89% | motif file (matrix) | svg |
| 464 | C T G A G A T C G C A T A C T G C G T A A C G T C G T A C G T A T A C G T C G A | PQM-1(?)/cElegans-L3-ChIP-Seq(modEncode)/Homer | 1e-7 | -1.678e+01 | 0.0000 | 3745.0 | 25.03% | 16002.3 | 22.98% | motif file (matrix) | svg |
| 465 | T A C G C T G A T C G A C G A T C T A G C T A G T C G A C T G A T C G A T C G A C G T A T C G A G C A T C A T G C G T A T A C G G C A T T G A C C G T A A G C T | NFAT:AP1(RHD,bZIP)/Jurkat-NFATC1-ChIP-Seq(Jolma\_et\_al.)/Homer | 1e-7 | -1.673e+01 | 0.0000 | 772.0 | 5.16% | 2896.7 | 4.16% | motif file (matrix) | svg |
| 466 | T C A G A C T G C A G T A G T C A G T C G T C A C G T A C G T A A C T G C A G T A G T C A G T C C T G A T G C A A G C T | dHNF4(NR)/Fly-HNF4-ChIP-Seq(GSE73675)/Homer | 1e-7 | -1.666e+01 | 0.0000 | 394.0 | 2.63% | 1345.0 | 1.93% | motif file (matrix) | svg |
| 467 | A G C T G A T C A G C T G A C T G A C T T C G A A G T C C G T A A C T G T C A G | SpliceAcceptor/Homer | 1e-7 | -1.639e+01 | 0.0000 | 13816.0 | 92.32% | 63366.7 | 91.00% | motif file (matrix) | svg |
| 468 | G C A T A G T C G A C T T C G A T A C G G T C A T C G A A C T G T A G C G C A T G C A T A T G C | AT2G01818(PLATZ)/col-AT2G01818-DAP-Seq(GSE60143)/Homer | 1e-7 | -1.617e+01 | 0.0000 | 1154.0 | 7.71% | 4535.8 | 6.51% | motif file (matrix) | svg |
| 469 | A C T G C G T A A C G T C G T A C T G A A C T G T C A G G C A T | At3g11280(MYBrelated)/col-At3g11280-DAP-Seq(GSE60143)/Homer | 1e-6 | -1.571e+01 | 0.0000 | 8635.0 | 57.70% | 38582.5 | 55.41% | motif file (matrix) | svg |
| 470 | T A G C T C A G C A T G G C A T A G C T C G A T A T G C C G T A C G T A G T C A | CHR(?)/Hela-CellCycle-Expression/Homer | 1e-6 | -1.566e+01 | 0.0000 | 3413.0 | 22.81% | 14557.4 | 20.91% | motif file (matrix) | svg |
| 471 | C T A G T G A C G A C T A T C G T C G A A G T C C T A G C A G T C T A G A T C G G T A C T C G A | O2(bZIP)/Corn-O2-ChIP-Seq(GSE63991)/Homer | 1e-6 | -1.564e+01 | 0.0000 | 1572.0 | 10.50% | 6365.6 | 9.14% | motif file (matrix) | svg |
| 472 | G C T A C G T A C G T A C G T A A C T G A C G T A G T C C G T A C G T A A G T C C A T G T A G C G T A C C G T A C G T A | WRKY7(WRKY)/colamp-WRKY7-DAP-Seq(GSE60143)/Homer | 1e-6 | -1.545e+01 | 0.0000 | 65.0 | 0.43% | 133.8 | 0.19% | motif file (matrix) | svg |
| 473 | G C A T G C T A C G T A A G C T G C T A T G C A A G T C A C G T A C G T A C G T G C A T G C A T | At4g38000(C2C2dof)/col-At4g38000-DAP-Seq(GSE60143)/Homer | 1e-6 | -1.527e+01 | 0.0000 | 5976.0 | 39.93% | 26265.5 | 37.72% | motif file (matrix) | svg |
| 474 | C T G A G T A C G A C T A G T C C A G T T G C A C T G A A C G T A G C T G A T C C T A G C G A T A C T G A T G C G A C T C T G A G A T C G A C T A G C T G A T C | Mouse\_Recombination\_Hotspot(Zf)/Testis-DMC1-ChIP-Seq(GSE24438)/Homer | 1e-6 | -1.525e+01 | 0.0000 | 467.0 | 3.12% | 1662.7 | 2.39% | motif file (matrix) | svg |
| 475 | C T G A A C G T A C G T A C G T A G T C G A C T C G A T C T G A A C T G C G T A C G T A T C G A | STAT5(Stat)/mCD4+-Stat5-ChIP-Seq(GSE12346)/Homer | 1e-6 | -1.514e+01 | 0.0000 | 1612.0 | 10.77% | 6557.2 | 9.42% | motif file (matrix) | svg |
| 476 | A G T C G A T C G C T A C G A T A C G T T A C G G C A T C T G A G A C T A C T G A G T C G C T A C T G A T C G A C A G T | Oct4:Sox17(POU,Homeobox,HMG)/F9-Sox17-ChIP-Seq(GSE44553)/Homer | 1e-6 | -1.510e+01 | 0.0000 | 838.0 | 5.60% | 3213.2 | 4.61% | motif file (matrix) | svg |
| 477 | T G A C C T G A A G T C A G T C A C T G G A T C G A C T G C A T | At5g18450(AP2EREBP)/col-At5g18450-DAP-Seq(GSE60143)/Homer | 1e-6 | -1.447e+01 | 0.0000 | 11313.0 | 75.60% | 51300.7 | 73.67% | motif file (matrix) | svg |
| 478 | G T C A G C A T G C T A C A G T C T A G G A T C C G T A C T G A C G T A C G A T | Oct2(POU,Homeobox)/Bcell-Oct2-ChIP-Seq(GSE21512)/Homer | 1e-6 | -1.442e+01 | 0.0000 | 1689.0 | 11.29% | 6921.0 | 9.94% | motif file (matrix) | svg |
| 479 | T C G A T A G C G T C A A C T G C T A G C G T A C G A T A C T G A C G T A C T G A C T G A C G T | ETS:RUNX(ETS,Runt)/Jurkat-RUNX1-ChIP-Seq(GSE17954)/Homer | 1e-6 | -1.440e+01 | 0.0000 | 500.0 | 3.34% | 1813.6 | 2.60% | motif file (matrix) | svg |
| 480 | C A G T G A C T G C A T T C G A A G T C A C G T A C G T A C G T C G A T G A C T | OBP3(C2C2dof)/col-OBP3-DAP-Seq(GSE60143)/Homer | 1e-6 | -1.430e+01 | 0.0000 | 12110.0 | 80.92% | 55125.2 | 79.17% | motif file (matrix) | svg |
| 481 | T C G A C G T A A G T C A G C T C G T A A G T C T C G A G C T A G A C T C G A T A G T C A G T C A G T C C T G A T C A G T G C A T C G A C A G T A T C G A G T C | GFY-Staf(?,Zf)/Promoter/Homer | 1e-6 | -1.428e+01 | 0.0000 | 267.0 | 1.78% | 876.4 | 1.26% | motif file (matrix) | svg |
| 482 | C G A T C T A G G A T C G C T A A G C T C T A G G A T C C G T A | RBFox2(?)/Heart-RBFox2-CLIP-Seq(GSE57926)/Homer | 1e-6 | -1.425e+01 | 0.0000 | 11279.0 | 75.37% | 51150.7 | 73.46% | motif file (matrix) | svg |
| 483 | G C T A G C A T G A C T G C A T T C A G G T A C G C T A G C A T C T G A G C T A T A G C G C T A C T G A C G A T C T A G | OCT4-SOX2-TCF-NANOG(POU,Homeobox,HMG)/mES-Oct4-ChIP-Seq(GSE11431)/Homer | 1e-5 | -1.350e+01 | 0.0000 | 723.0 | 4.83% | 2765.4 | 3.97% | motif file (matrix) | svg |
| 484 | G A C T G A C T T C G A C G T A C G A T A G C T C T G A A C T G T G A C G A C T T C G A C G T A A C G T A G C T C T G A C T G A G T C A G C T A G C T A C G T A | Pax7(Paired,Homeobox),longest/Myoblast-Pax7-ChIP-Seq(GSE25064)/Homer | 1e-5 | -1.330e+01 | 0.0000 | 132.0 | 0.88% | 375.5 | 0.54% | motif file (matrix) | svg |
| 485 | T G C A G C A T A G C T G C A T A G T C A G T C A G T C C T G A A C T G T C G A T C G A C A G T A T C G A G T C G A T C | ZNF143\|STAF(Zf)/CUTLL-ZNF143-ChIP-Seq(GSE29600)/Homer | 1e-5 | -1.306e+01 | 0.0000 | 1316.0 | 8.79% | 5334.9 | 7.66% | motif file (matrix) | svg |
| 486 | A G C T T G A C G A C T A G T C C T A G G A T C C T A G C T G A A C T G T C G A A G T C A G C T | BANP(?)/ESC-Banp-ChIP-Seq(GSE155603)/Homer | 1e-5 | -1.293e+01 | 0.0000 | 2815.0 | 18.81% | 11999.0 | 17.23% | motif file (matrix) | svg |
| 487 | G C A T T C G A C T G A G A T C A G T C G A T C G T C A G T C A A C G T A G T C C G T A C T G A | Duxbl(Homeobox)/NIH3T3-Duxbl.HA-ChIP-Seq(GSE119782)/Homer | 1e-5 | -1.270e+01 | 0.0000 | 685.0 | 4.58% | 2624.1 | 3.77% | motif file (matrix) | svg |
| 488 | C G T A C T G A C T A G C G T A C G T A A G T C C G T A C A G T G C A T G T C A C G A T A C T G A C G T G C A T G A T C | PGR(NR)/EndoStromal-PGR-ChIP-Seq(GSE69539)/Homer | 1e-5 | -1.266e+01 | 0.0000 | 1498.0 | 10.01% | 6146.9 | 8.83% | motif file (matrix) | svg |
| 489 | G C T A G A C T A G T C T C G A T C A G T C G A A C G T A G T C G A C T T C A G | GATA14(C2C2gata)/col-GATA14-DAP-Seq(GSE60143)/Homer | 1e-5 | -1.258e+01 | 0.0000 | 6243.0 | 41.72% | 27662.0 | 39.73% | motif file (matrix) | svg |
| 490 | A T G C C T G A G A C T A C G T A C G T G T A C G A T C C G A T C T A G C A T G C G T A C G T A C T G A G A C T | STAT1(Stat)/HelaS3-STAT1-ChIP-Seq(GSE12782)/Homer | 1e-5 | -1.250e+01 | 0.0000 | 1503.0 | 10.04% | 6174.9 | 8.87% | motif file (matrix) | svg |
| 491 | C G A T C T A G T C G A A G C T C G A T C T G A C G T A A G C T A C T G C T A G A T G C G A T C | Hoxb4(Homeobox)/ES-Hoxb4-ChIP-Seq(GSE34014)/Homer | 1e-5 | -1.236e+01 | 0.0000 | 2003.0 | 13.38% | 8392.0 | 12.05% | motif file (matrix) | svg |
| 492 | A G T C A T C G G C A T C A T G A C T G A T C G C G A T C T A G A C T G A G C T T G A C G A C T | Gli2(Zf)/GM2-Gli2-ChIP-Chip(GSE112702)/Homer | 1e-5 | -1.226e+01 | 0.0000 | 1535.0 | 10.26% | 6324.1 | 9.08% | motif file (matrix) | svg |
| 493 | G C T A T A G C A G C T A T C G G T C A C G T A G C T A A T G C G A T C C T G A | IRF4(IRF)/GM12878-IRF4-ChIP-Seq(GSE32465)/Homer | 1e-5 | -1.204e+01 | 0.0000 | 3656.0 | 24.43% | 15845.3 | 22.76% | motif file (matrix) | svg |
| 494 | C A G T A G C T G A C T T G C A A G T C A G C T A C G T A C G T C G A T G A C T | AT3G52440(C2C2dof)/colamp-AT3G52440-DAP-Seq(GSE60143)/Homer | 1e-5 | -1.188e+01 | 0.0000 | 10731.0 | 71.71% | 48687.4 | 69.92% | motif file (matrix) | svg |
| 495 | T A G C G A T C A G C T T G A C G C T A A G C T C A T G A C T G A C G T T C A G A G T C G A T C G A C T A G C T G C T A A G T C A G C T A G T C G A T C A T G C A G C T G A C T C A T G A C G T A T C G | ZNF41(Zf)/HEK293-ZNF41.GFP-ChIP-Seq(GSE58341)/Homer | 1e-5 | -1.167e+01 | 0.0000 | 137.0 | 0.92% | 409.8 | 0.59% | motif file (matrix) | svg |
| 496 | G T A C C T G A A G T C A G T C A C T G G T C A G A T C G C A T | At1g75490(AP2EREBP)/colamp-At1g75490-DAP-Seq(GSE60143)/Homer | 1e-5 | -1.162e+01 | 0.0000 | 13299.0 | 88.87% | 61007.6 | 87.61% | motif file (matrix) | svg |
| 497 | C G T A C T G A C T A G C T G A A G T C G C T A C G A T A T C G G A C T G A T C A G T C C T G A C T A G C T A G A G T C G C T A C G A T C T A G G A T C G A T C | p73(p53)/Trachea-p73-ChIP-Seq(PRJNA310161)/Homer | 1e-4 | -1.134e+01 | 0.0000 | 382.0 | 2.55% | 1386.4 | 1.99% | motif file (matrix) | svg |
| 498 | C T A G A G C T G A C T C A T G A G T C A G T C G T C A C A G T C T A G T C A G G T A C C T G A T C G A G A T C T G A C | Rfx2(HTH)/LoVo-RFX2-ChIP-Seq(GSE49402)/Homer | 1e-4 | -1.125e+01 | 0.0000 | 552.0 | 3.69% | 2097.3 | 3.01% | motif file (matrix) | svg |
| 499 | A T G C G A C T A G C T A G C T A G T C G C T A C A G T C G A T G C T A A C G T A C T G G C T A T A G C G C A T T G A C | IRF:BATF(IRF:bZIP)/pDC-Irf8-ChIP-Seq(GSE66899)/Homer | 1e-4 | -1.102e+01 | 0.0000 | 563.0 | 3.76% | 2149.2 | 3.09% | motif file (matrix) | svg |
| 500 | G T A C A C T G A C G T T C A G G C A T C G T A C G A T G C A T C G T A A G T C C G T A T G A C C A T G G A C T G C T A | ANAC083(NAC)/col-ANAC083-DAP-Seq(GSE60143)/Homer | 1e-4 | -1.101e+01 | 0.0000 | 7398.0 | 49.44% | 33119.5 | 47.56% | motif file (matrix) | svg |
| 501 | T G C A T A G C G A C T T G C A T G A C T G C A C G T A A G C T A G C T A G T C A G T C G T A C | GFY(?)/Promoter/Homer | 1e-4 | -1.083e+01 | 0.0000 | 485.0 | 3.24% | 1825.1 | 2.62% | motif file (matrix) | svg |
| 502 | G A C T C T A G C T A G C T A G A C T G T C G A C T G A C T A G C T A G C T A G G T A C G T C A | ZNF467(Zf)/HEK293-ZNF467.GFP-ChIP-Seq(GSE58341)/Homer | 1e-4 | -1.081e+01 | 0.0000 | 3154.0 | 21.08% | 13640.9 | 19.59% | motif file (matrix) | svg |
| 503 | C G A T C T G A G T A C A C T G A C G T T C A G G C A T C G T A C G T A G C A T C G T A A G T C C G T A G T A C C A T G | CUC3(NAC)/col-CUC3-DAP-Seq(GSE60143)/Homer | 1e-4 | -1.065e+01 | 0.0000 | 4157.0 | 27.78% | 18210.8 | 26.15% | motif file (matrix) | svg |
| 504 | C A T G A G T C C T G A A T G C C T A G T C G A G C T A G C A T G A T C G A C T A G T C C T A G C G T A C A T G C T A G | PLT3(AP2EREBP)/col-PLT3-DAP-Seq(GSE60143)/Homer | 1e-4 | -1.013e+01 | 0.0001 | 1124.0 | 7.51% | 4598.7 | 6.60% | motif file (matrix) | svg |
| 505 | A T G C G A C T A C T G C A G T G A T C A C G T T A C G T A C G | Smad2(MAD)/ES-SMAD2-ChIP-Seq(GSE29422)/Homer | 1e-4 | -1.003e+01 | 0.0001 | 9737.0 | 65.07% | 44123.3 | 63.37% | motif file (matrix) | svg |
| 506 | G A C T G T A C G A C T A G T C T C A G C T G A A G T C A G T C C T A G C G A T A G C T A T G C C T G A C A G T A G C T | AT4G27900(C2C2COlike)/col-AT4G27900-DAP-Seq(GSE60143)/Homer | 1e-4 | -9.945e+00 | 0.0001 | 191.0 | 1.28% | 637.5 | 0.92% | motif file (matrix) | svg |
| 507 | T G A C G T A C C G T A A C T G T G A C C G A T A C T G A T C G A G C T T A C G T C G A T A G C G T A C C G T A A T C G T G A C G C A T A C T G A C T G A T G C | Twist(bHLH)/HMLE-TWIST1-ChIP-Seq(Chang\_et\_al)/Homer | 1e-4 | -9.937e+00 | 0.0001 | 500.0 | 3.34% | 1909.2 | 2.74% | motif file (matrix) | svg |
| 508 | T C G A G A C T A G C T T G A C A G C T G T A C T C A G G A T C A T C G T G C A A C T G C T G A | GFX(?)/Promoter/Homer | 1e-4 | -9.464e+00 | 0.0002 | 363.0 | 2.43% | 1346.0 | 1.93% | motif file (matrix) | svg |
| 509 | C G T A C G T A T C G A C G T A C G A T C G T A A C G T A G T C G C A T G C A T | At3g09600(MYBrelated)/colamp-At3g09600-DAP-Seq(GSE60143)/Homer | 1e-4 | -9.458e+00 | 0.0002 | 2881.0 | 19.25% | 12483.3 | 17.93% | motif file (matrix) | svg |
| 510 | T G C A T G C A A G T C A G T C G A C T C A G T A T G C G A T C C T G A A C G T C T A G C T A G A G T C A C G T A G T C A G T C A G T C G A C T C G T A A C G T A G C T C T A G G A T C G A T C G A T C | ZNF16(Zf)/HEK293-ZNF16.GFP-ChIP-Seq(GSE58341)/Homer | 1e-3 | -9.192e+00 | 0.0002 | 30.0 | 0.20% | 56.0 | 0.08% | motif file (matrix) | svg |
| 511 | T A G C G T A C C G T A C T A G A C T G T G C A C G T A A T G C C G T A A T C G | AR-halfsite(NR)/LNCaP-AR-ChIP-Seq(GSE27824)/Homer | 1e-3 | -9.160e+00 | 0.0002 | 12513.0 | 83.62% | 57344.3 | 82.35% | motif file (matrix) | svg |
| 512 | T G C A A T G C A C G T A C G T A C G T A T G C C T A G A C G T A C G T A G C T G A T C A G C T | T1ISRE(IRF)/ThioMac-Ifnb-Expression/Homer | 1e-3 | -8.976e+00 | 0.0003 | 80.0 | 0.53% | 225.5 | 0.32% | motif file (matrix) | svg |
| 513 | G C T A C G T A A C G T C A T G C G T A A C G T A C G T C T A G | ATHB5(HB)/colamp-ATHB5-DAP-Seq(GSE60143)/Homer | 1e-3 | -8.840e+00 | 0.0003 | 6707.0 | 44.82% | 30077.3 | 43.19% | motif file (matrix) | svg |
| 514 | A T C G A G C T A C T G A G T C A C T G A G T C C G T A A C G T A C T G A G T C A C T G A G T C | NRF(NRF)/Promoter/Homer | 1e-3 | -8.139e+00 | 0.0006 | 1645.0 | 10.99% | 6993.4 | 10.04% | motif file (matrix) | svg |
| 515 | A G T C A T C G C T A G A G C T G A C T C T A G A G T C A G T C G C T A C A G T T C A G T C A G G A T C C T G A T C G A G A T C | RFX(HTH)/K562-RFX3-ChIP-Seq(SRA012198)/Homer | 1e-3 | -8.054e+00 | 0.0006 | 475.0 | 3.17% | 1851.4 | 2.66% | motif file (matrix) | svg |
| 516 | C G A T C T G A G T A C C A T G G C A T T C A G G C A T C G T A C G T A G C T A C G T A A G T C G C T A G T A C C A T G | CUC2(NAC)/colamp-CUC2-DAP-Seq(GSE60143)/Homer | 1e-3 | -8.020e+00 | 0.0007 | 4062.0 | 27.14% | 17957.3 | 25.79% | motif file (matrix) | svg |
| 517 | G A T C G C T A A G T C A C G T A C T G C G T A A G T C C G T A G C T A C G A T C A G T G C A T G C T A C G T A G C A T | GRF9(GRF)/colamp-GRF9-DAP-Seq(GSE60143)/Homer | 1e-3 | -7.897e+00 | 0.0008 | 6121.0 | 40.90% | 27442.4 | 39.41% | motif file (matrix) | svg |
| 518 | T C G A G C T A T G A C G C T A C T A G G A T C C G A T A C T G C G A T A G C T G A C T C T A G | E-box/Drosophila-Promoters/Homer | 1e-3 | -7.698e+00 | 0.0009 | 1293.0 | 8.64% | 5445.9 | 7.82% | motif file (matrix) | svg |
| 519 | A T G C G T A C A C T G A G T C A G T C A C T G G A T C G T C A C G T A C G A T G C A T C G A T | RRTF1(AP2EREBP)/colamp-RRTF1-DAP-Seq(GSE60143)/Homer | 1e-3 | -7.679e+00 | 0.0009 | 4902.0 | 32.76% | 21838.9 | 31.36% | motif file (matrix) | svg |
| 520 | G C T A A G C T G T A C G C A T A G C T T C G A C T G A A G T C A G T C T A C G A C G T G A C T T A C G C T A G C G T A | ZML1(C2C2gata)/colamp-ZML1-DAP-Seq(GSE60143)/Homer | 1e-3 | -7.505e+00 | 0.0011 | 502.0 | 3.35% | 1982.9 | 2.85% | motif file (matrix) | svg |
| 521 | A C T G C G T A C G T A A C G T G T A C G A C T C G T A C G A T C T G A C T G A | AT1G49560(G2like)/colamp-AT1G49560-DAP-Seq(GSE60143)/Homer | 1e-3 | -7.370e+00 | 0.0013 | 9636.0 | 64.39% | 43860.8 | 62.99% | motif file (matrix) | svg |
| 522 | G C T A G C T A T G C A A G T C C T A G C T G A G A T C C T A G G A C T G A T C C T A G A C G T C G A T C G A T G A C T | Unknown2/Arabidopsis-Promoters/Homer | 1e-3 | -7.266e+00 | 0.0014 | 263.0 | 1.76% | 975.3 | 1.40% | motif file (matrix) | svg |
| 523 | C A T G T A C G T A G C G A T C G A T C A T G C G T A C G A C T T C A G A T G C C G A T A T C G C A G T A C T G G T A C | Zic3(Zf)/mES-Zic3-ChIP-Seq(GSE37889)/Homer | 1e-3 | -7.204e+00 | 0.0015 | 2405.0 | 16.07% | 10468.1 | 15.03% | motif file (matrix) | svg |
| 524 | G A T C G A T C G C T A G T C A G A C T A T G C T C G A C G A T C G A T C T A G | HAT2(Homeobox)/colamp-HAT2-DAP-Seq(GSE60143)/Homer | 1e-2 | -6.892e+00 | 0.0020 | 6780.0 | 45.31% | 30583.5 | 43.92% | motif file (matrix) | svg |
| 525 | G C A T C G T A G C T A G A C T C G A T G A C T A G T C C A G T A G T C A G T C A C T G C T A G G T A C C T A G C T G A | AT5G05550(Trihelix)/col-AT5G05550-DAP-Seq(GSE60143)/Homer | 1e-2 | -6.843e+00 | 0.0021 | 11877.0 | 79.37% | 54472.2 | 78.23% | motif file (matrix) | svg |
| 526 | T A G C G T A C C T A G C A G T T C G A C G T A C G T A G C A T G A C T T G A C A G T C A C T G A T C G A G T C C T A G | AS2(LOBAS2)/col-AS2-DAP-Seq(GSE60143)/Homer | 1e-2 | -6.813e+00 | 0.0022 | 1373.0 | 9.17% | 5845.4 | 8.39% | motif file (matrix) | svg |
| 527 | T C G A C T G A C G A T C G T A C G T A C G T A C T A G A G C T C T G A T C A G | Adof1(C2C2dof)/col-Adof1-DAP-Seq(GSE60143)/Homer | 1e-2 | -6.496e+00 | 0.0030 | 11612.0 | 77.59% | 53244.4 | 76.46% | motif file (matrix) | svg |
| 528 | G C A T A C G T A G T C A T G C A G T C C T A G T A G C G T A C C T G A G C T A | DEL1(E2FDP)/colamp-DEL1-DAP-Seq(GSE60143)/Homer | 1e-2 | -6.171e+00 | 0.0041 | 69.0 | 0.46% | 211.4 | 0.30% | motif file (matrix) | svg |
| 529 | C T A G G A C T G A T C A C G T A T C G A G C T C T G A A T C G C G A T C T A G G A T C G A C T C A T G A T C G G T A C G A C T A G T C G C A T A G C T C G A T | ZNF382(Zf)/HEK293-ZNF382.GFP-ChIP-Seq(GSE58341)/Homer | 1e-2 | -6.149e+00 | 0.0042 | 152.0 | 1.02% | 539.4 | 0.77% | motif file (matrix) | svg |
| 530 | A T C G A G T C A G T C G A C T A T G C C T G A C T A G A C T G T A C G G T A C C T G A C G A T | AP-2gamma(AP2)/MCF7-TFAP2C-ChIP-Seq(GSE21234)/Homer | 1e-2 | -6.086e+00 | 0.0045 | 5030.0 | 33.61% | 22566.1 | 32.41% | motif file (matrix) | svg |
| 531 | C T A G A T G C A T G C C G A T A C T G G A C T A T G C G C T A T G A C A G C T T A G C G C T A | PBX1(Homeobox)/MCF7-PBX1-ChIP-Seq(GSE28007)/Homer | 1e-2 | -6.071e+00 | 0.0046 | 392.0 | 2.62% | 1551.2 | 2.23% | motif file (matrix) | svg |
| 532 | C T A G C T A G T C A G G T C A C T A G T C A G G C T A A G T C A T C G A G C T C T A G | DPR(core promoter) | 1e-2 | -5.979e+00 | 0.0050 | 14367.0 | 96.00% | 66491.0 | 95.49% | motif file (matrix) | svg |
| 533 | G C T A T G A C G A T C C G A T G A C T A T G C C T G A A T C G G C A T A C G T | JGL(C2H2)/col-JGL-DAP-Seq(GSE60143)/Homer | 1e-2 | -5.970e+00 | 0.0050 | 9123.0 | 60.96% | 41587.4 | 59.72% | motif file (matrix) | svg |
| 534 | C T A G A C T G A G C T C G T A A C T G A C T G A C G T C T A G | MYB99(MYB)/colamp-MYB99-DAP-Seq(GSE60143)/Homer | 1e-2 | -5.656e+00 | 0.0069 | 8014.0 | 53.55% | 36442.4 | 52.33% | motif file (matrix) | svg |
| 535 | G A T C G T A C C G T A A G C T G A C T G C T A C T G A A C G T G A T C G C T A | Hoxc6(Homeobox)/EB-Hoxc6.iFlag-ChIP-Seq(GSE142377)/Homer | 1e-2 | -5.486e+00 | 0.0081 | 12579.0 | 84.06% | 57913.4 | 83.17% | motif file (matrix) | svg |
| 536 | G A T C G C A T A C T G C T A G C T G A A G C T G C T A G C T A C T G A T C A G G C A T T G C A A C G T A C G T G A T C G A C T G C A T C T A G T A C G G A C T C T A G C A T G C T A G G T A C T C G A | ZNF136(Zf)/HEK293-ZNF136.GFP-ChIP-Seq(GSE58341)/Homer | 1e-2 | -5.460e+00 | 0.0083 | 549.0 | 3.67% | 2253.5 | 3.24% | motif file (matrix) | svg |
| 537 | T C G A C A T G C A T G A C G T A T G C T C G A C T G A A G C T T A C G T G C A G T A C G A T C A G C T A G T C | FXR(NR),IR1/Liver-FXR-ChIP-Seq(Chong\_et\_al.)/Homer | 1e-2 | -5.290e+00 | 0.0099 | 2733.0 | 18.26% | 12098.7 | 17.37% | motif file (matrix) | svg |
| 538 | C T A G C T A G T C G A C G T A A T G C C G T A A T C G T C G A T A C G G C A T A C T G C A G T T A G C G A T C G A C T | MRE(NR)/Neuro2A-NR3C2-ChIPnexus(GSE115417)/Homer | 1e-2 | -5.274e+00 | 0.0100 | 7305.0 | 48.81% | 33183.1 | 47.65% | motif file (matrix) | svg |
| 539 | T A G C A G T C T G A C A G T C C T A G A T C G A G T C C A T G T G A C A G T C G T A C A G T C A G T C G C A T C T A G A T C G G C A T A C T G A T C G G A T C | BORIS(Zf)/K562-CTCFL-ChIP-Seq(GSE32465)/Homer | 1e-2 | -4.865e+00 | 0.0150 | 615.0 | 4.11% | 2567.0 | 3.69% | motif file (matrix) | svg |
| 540 | C G T A A C G T A G C T C G A T C T A G G T A C C G T A A G C T C G T A G C T A | Oct4(POU,Homeobox)/mES-Oct4-ChIP-Seq(GSE11431)/Homer | 1e-2 | -4.749e+00 | 0.0168 | 2758.0 | 18.43% | 12258.1 | 17.60% | motif file (matrix) | svg |
| 541 | G T C A A C G T C T A G G T C A C G T A G C A T C G T A C G T A G C A T C A G T A G T C C G T A C A G T C T A G C T G A | OCT:OCT(POU,Homeobox,IR1)/NPC-Brn2-ChIP-Seq(GSE35496)/Homer | 1e-2 | -4.732e+00 | 0.0171 | 87.0 | 0.58% | 299.4 | 0.43% | motif file (matrix) | svg |
| 542 | G C A T C A G T C G A T A G T C G A T C G C T A C G A T C G A T C G A T G C T A C G A T C T A G A C T G G C T A G C T A | AGL25(MADS)/colamp-AGL25-DAP-Seq(GSE60143)/Homer | 1e-2 | -4.658e+00 | 0.0184 | 218.0 | 1.46% | 845.8 | 1.21% | motif file (matrix) | svg |
| 543 | G A C T C T A G C T A G A G T C T G C A A C T G A C G T A C G T C T A G T C A G | AMYB(HTH)/Testes-AMYB-ChIP-Seq(GSE44588)/Homer | 1e-2 | -4.657e+00 | 0.0184 | 11606.0 | 77.55% | 53381.3 | 76.66% | motif file (matrix) | svg |
| 544 | A T G C G C A T C G A T G A T C A G C T C T G A A C T G C G T A C G T A T C A G T G A C C G A T G C A T G A T C C G A T | HSF21(HSF)/col-HSF21-DAP-Seq(GSE60143)/Homer | 1e-2 | -4.644e+00 | 0.0185 | 492.0 | 3.29% | 2034.3 | 2.92% | motif file (matrix) | svg |
