## Supplemental dataset for "Hybrid CNN and Multi-Head Attention Model for Analyzing Epigenetic Mechanisms and Gene Expression Across Fungal Phylogenetic Distances": NcrassaModel_FgramTest_K27me3locs_homerResults.html

/projects/wg-feeds/SHAP/NcrassaModel\_FgramTest\_K27me3locs\_SHAP\_noDup\_HOMER// - Homer de novo Motif Results


### Homer *de novo* Motif Results (/projects/wg-feeds/SHAP/NcrassaModel\_FgramTest\_K27me3locs\_SHAP\_noDup\_HOMER//)

Non-redundant Motif File of Results  
Known Motif Enrichment Results  
Gene Ontology Enrichment Results  
If Homer is having trouble matching a motif to a known motif, try copy/pasting the matrix file into
STAMP  
More information on motif finding results: HOMER
| Description of Results
| Tips
  
Total target sequences = 1617  
Total background sequences = 7373  
\* - possible false positive  

|  |  |  |  |  |  |  |  |  |
| --- | --- | --- | --- | --- | --- | --- | --- | --- |
| Rank | Motif | P-value | log P-pvalue | % of Targets | % of Background | STD(Bg STD) | Best Match/Details | Motif File |
| 1 | G A C T A G T C T G C A G C A T A G T C G C T A C G A T A T G C C G T A G C A T A T G C T G C A | 1e-233 | -5.372e+02 | 57.58% | 16.64% | 1087.4bp (1120.9bp) | ZML2(C2C2gata)/col-ZML2-DAP-Seq(GSE60143)/Homer(0.868) More Information \| Similar Motifs Found | motif file (matrix) |
| 2 | G C A T A T G C T C A G G C T A A T G C C T G A C G T A A T C G C T G A C G T A A T C G T C G A | 1e-221 | -5.094e+02 | 57.27% | 17.26% | 1025.4bp (1069.8bp) | TRA2(RRM)/Drosophila\_melanogaster-RNCMPT00078-PBM/HughesRNA(0.703) More Information \| Similar Motifs Found | motif file (matrix) |
| 3 | A G T C T A G C C G A T A G C T A C T G C T G A T A C G T A G C G C A T A G C T A T C G C A G T | 1e-214 | -4.934e+02 | 48.48% | 11.96% | 968.5bp (1093.5bp) | NR6A1/MA1541.2/Jaspar(0.842) More Information \| Similar Motifs Found | motif file (matrix) |
| 4 | T C G A T C G A T C G A C T G A T C G A T C G A T C G A T C G A T C G A C T G A T C G A C T G A | 1e-181 | -4.169e+02 | 50.22% | 15.30% | 1147.2bp (1081.6bp) | SeqBias: polyA-repeat(0.879) More Information \| Similar Motifs Found | motif file (matrix) |
| 5 | T C A G T G A C C G T A T C G A T G A C C T G A C T A G T G A C C T G A T C A G T G A C T C G A | 1e-167 | -3.867e+02 | 56.15% | 20.66% | 1100.1bp (1103.0bp) | CG4360/MA2204.1/Jaspar(0.775) More Information \| Similar Motifs Found | motif file (matrix) |
| 6 | C T A G C G T A C T A G C T A G G T A C G A C T T A C G G A C T G T A C C T G A C G T A T A C G | 1e-142 | -3.285e+02 | 55.91% | 22.84% | 1092.3bp (1091.5bp) | RIM101/MA0368.1/Jaspar(0.647) More Information \| Similar Motifs Found | motif file (matrix) |
| 7 | G A C T A T G C T C G A C G T A A C T G T C G A C G A T A G T C T A G C G C A T A G C T T A C G | 1e-137 | -3.168e+02 | 39.89% | 11.83% | 1027.0bp (1117.5bp) | ftz-f1/MA2311.1/Jaspar(0.734) More Information \| Similar Motifs Found | motif file (matrix) |
| 8 | C A T G G T A C T A G C C G T A C G A T A T C G G C T A A G C T A T C G T C A G | 1e-123 | -2.854e+02 | 60.05% | 28.36% | 1176.4bp (1113.6bp) | ZML2(C2C2gata)/col-ZML2-DAP-Seq(GSE60143)/Homer(0.702) More Information \| Similar Motifs Found | motif file (matrix) |
| 9 | T A G C C T G A C A T G G A T C G T A C C G A T A G C T A C T G C T A G G T A C | 1e-118 | -2.722e+02 | 68.03% | 36.51% | 1060.3bp (1100.4bp) | RIM101/MA0368.1/Jaspar(0.733) More Information \| Similar Motifs Found | motif file (matrix) |
| 10 | T A C G G A C T G A C T T A G C C G T A C G A T A T G C C A G T A T G C C G T A C A T G T G A C | 1e-111 | -2.568e+02 | 53.74% | 24.42% | 1033.1bp (1129.7bp) | pros/dmmpmm(Bergman)/fly(0.655) More Information \| Similar Motifs Found | motif file (matrix) |
| 11 | G C T A T G A C T G C A C G T A C T A G G C T A T G A C T G C A C T G A G T C A T G A C C G T A | 1e-105 | -2.430e+02 | 53.99% | 25.35% | 1115.7bp (1180.7bp) | NF1:FOXA1(CTF,Forkhead)/LNCAP-FOXA1-ChIP-Seq(GSE27824)/Homer(0.730) More Information \| Similar Motifs Found | motif file (matrix) |
| 12 | T C G A C T G A T C A G T C G A C T A G G C T A A C G T C T A G C T G A T C A G T G C A A C T G | 1e-95 | -2.210e+02 | 43.17% | 17.87% | 1109.5bp (1067.2bp) | TOD6?/SacCer-Promoters/Homer(0.727) More Information \| Similar Motifs Found | motif file (matrix) |
| 13 | A G C T A T G C C G A T A G T C A C G T T G A C A G C T A G T C A C G T A T G C G A C T A G C T | 1e-93 | -2.163e+02 | 57.51% | 29.91% | 1002.9bp (1121.8bp) | SeqBias: GA-repeat(0.933) More Information \| Similar Motifs Found | motif file (matrix) |
| 14 | G C A T A C T G A T C G T G C A C T G A A T C G T G C A T C G A G C T A A T C G T C G A C T G A | 1e-73 | -1.694e+02 | 41.25% | 19.09% | 1112.1bp (1165.8bp) | NTL8/MA1678.3/Jaspar(0.645) More Information \| Similar Motifs Found | motif file (matrix) |
| 15 | A T G C G T C A C G T A G C A T A C T G T C A G A G T C G C T A | 1e-70 | -1.631e+02 | 50.22% | 26.82% | 1118.2bp (1154.2bp) | pho/dmmpmm(Bergman)/fly(0.866) More Information \| Similar Motifs Found | motif file (matrix) |
| 16 | C A T G A T G C G A T C T C A G C G T A A C G T C T G A G A C T A T G C C T A G | 1e-70 | -1.619e+02 | 42.61% | 20.59% | 986.0bp (1064.3bp) | PB0127.1\_Gata6\_2/Jaspar(0.818) More Information \| Similar Motifs Found | motif file (matrix) |
| 17 | T C A G C T A G G T A C C T A G A C T G G T A C C T A G C T G A G T A C C T A G | 1e-70 | -1.616e+02 | 51.89% | 28.38% | 1010.0bp (1066.6bp) | LEP/MA1246.2/Jaspar(0.825) More Information \| Similar Motifs Found | motif file (matrix) |
| 18 | A C G T A G C T A C T G C T A G T G A C C G T A A C T G G T A C | 1e-66 | -1.523e+02 | 53.25% | 30.24% | 1119.9bp (1088.6bp) | NFIA/MA0670.2/Jaspar(0.824) More Information \| Similar Motifs Found | motif file (matrix) |
| 19 | T G C A T G C A A G T C G C T A C G T A G T A C A G T C C A G T G A T C A G C T | 1e-46 | -1.075e+02 | 17.81% | 5.92% | 996.1bp (1177.7bp) | YBX2(CSD)/Homo\_sapiens-RNCMPT00084-PBM/HughesRNA(0.712) More Information \| Similar Motifs Found | motif file (matrix) |
| 20 | T A G C A G C T C T G A A G T C T G A C G A C T C G T A T G A C G T A C A G C T T G C A A T G C | 1e-41 | -9.556e+01 | 12.06% | 3.12% | 1206.5bp (1031.8bp) | PK06182.1/MA2354.1/Jaspar(0.831) More Information \| Similar Motifs Found | motif file (matrix) |
| 21 | C A T G A C G T C A T G G A C T T C A G G A C T C A T G G A C T C A T G C G A T C A T G G A C T | 1e-20 | -4.762e+01 | 8.84% | 3.12% | 880.8bp (1027.0bp) | cg/MA2107.1/Jaspar(0.930) More Information \| Similar Motifs Found | motif file (matrix) |
