## Supplemental dataset for "Hybrid CNN and Multi-Head Attention Model for Analyzing Epigenetic Mechanisms and Gene Expression Across Fungal Phylogenetic Distances": NcrassaModel_FgramTest_K27me3locs_knownResults.html

/projects/wg-feeds/SHAP/NcrassaModel\_FgramTest\_K27me3locs\_SHAP\_noDup\_HOMER/ - Homer Known Motif Enrichment Results


### Homer Known Motif Enrichment Results (/projects/wg-feeds/SHAP/NcrassaModel\_FgramTest\_K27me3locs\_SHAP\_noDup\_HOMER/)

Homer *de novo* Motif Results  
Gene Ontology Enrichment Results  
Known Motif Enrichment Results (txt file)  
Total Target Sequences = 1617, Total Background Sequences = 7369

|  |  |  |  |  |  |  |  |  |  |  |  |
| --- | --- | --- | --- | --- | --- | --- | --- | --- | --- | --- | --- |
| Rank | Motif | Name | P-value | log P-pvalue | q-value (Benjamini) | # Target Sequences with Motif | % of Targets Sequences with Motif | # Background Sequences with Motif | % of Background Sequences with Motif | Motif File | SVG |
| 1 | T A G C G T C A G A C T T A G C G T C A G A C T A G T C G C T A G A C T G A T C | ZML2(C2C2gata)/col-ZML2-DAP-Seq(GSE60143)/Homer | 1e-137 | -3.172e+02 | 0.0000 | 600.0 | 37.11% | 745.7 | 10.12% | motif file (matrix) | svg |
| 2 | A T G C C A G T A C G T A G T C C A T G A C G T A G T C A C G T A G C T G A T C | Unknown4/Arabidopsis-Promoters/Homer | 1e-102 | -2.361e+02 | 0.0000 | 1116.0 | 69.02% | 2923.4 | 39.66% | motif file (matrix) | svg |
| 3 | C T A G T C A G C T G A C T G A C T A G C T G A C A T G C A T G C T G A C T A G C T A G C G T A C T A G C G T A G T C A | TF3A(C2H2)/col-TF3A-DAP-Seq(GSE60143)/Homer | 1e-100 | -2.316e+02 | 0.0000 | 943.0 | 58.32% | 2192.0 | 29.74% | motif file (matrix) | svg |
| 4 | G C T A G C A T A C G T A C G T A G C T G A T C G A C T G A C T A G C T A C G T A C G T A G C T | RLR1?/SacCer-Promoters/Homer | 1e-71 | -1.649e+02 | 0.0000 | 354.0 | 21.89% | 455.6 | 6.18% | motif file (matrix) | svg |
| 5 | G C A T A G C T A C G T A C G T A C T G A C G T G A T C A C G T A C G T A G C T C G A T G C A T A G T C G A C T C A G T | IDD5(C2H2)/colamp-IDD5-DAP-Seq(GSE60143)/Homer | 1e-61 | -1.428e+02 | 0.0000 | 596.0 | 36.86% | 1270.9 | 17.24% | motif file (matrix) | svg |
| 6 | A C G T G A C T A T G C G C T A C T G A C T A G A C T G G A C T A G T C C G T A | Nr5a2(NR)/mES-Nr5a2-ChIP-Seq(GSE19019)/Homer | 1e-61 | -1.420e+02 | 0.0000 | 677.0 | 41.87% | 1565.0 | 21.23% | motif file (matrix) | svg |
| 7 | A C G T G A C T T A G C C G T A C T G A C A T G C T A G G A C T G A T C C G T A | Nr5a2(NR)/Pancreas-LRH1-ChIP-Seq(GSE34295)/Homer | 1e-57 | -1.331e+02 | 0.0000 | 814.0 | 50.34% | 2142.0 | 29.06% | motif file (matrix) | svg |
| 8 | A C G T A T G C A C G T A C G T A G C T A G T C A G C T A G C T A G C T A G C T A G C T | hTCT(CPE) | 1e-54 | -1.252e+02 | 0.0000 | 1195.0 | 73.90% | 3916.7 | 53.14% | motif file (matrix) | svg |
| 9 | C A T G G C T A C T A G T A C G C G T A T C A G C G T A A C T G C G T A C A T G C T G A C G T A | BPC1(BBRBPC)/colamp-BPC1-DAP-Seq(GSE60143)/Homer | 1e-50 | -1.170e+02 | 0.0000 | 506.0 | 31.29% | 1074.8 | 14.58% | motif file (matrix) | svg |
| 10 | T G A C C G T A C T G A A C T G A C T G G A C T G A T C T G C A G T A C T A C G | SF1(NR)/H295R-Nr5a1-ChIP-Seq(GSE44220)/Homer | 1e-48 | -1.122e+02 | 0.0000 | 554.0 | 34.26% | 1263.4 | 17.14% | motif file (matrix) | svg |
| 11 | A C G T A C T G C G T A A C G T A C T G A C T G C G T A C G T A | HAP3(CCAATHAP3)/col-HAP3-DAP-Seq(GSE60143)/Homer | 1e-48 | -1.114e+02 | 0.0000 | 597.0 | 36.92% | 1422.0 | 19.29% | motif file (matrix) | svg |
| 12 | G A C T A G C T A G C T C T A G A C G T G A T C A C G T A C G T G A C T C G A T G C A T A G T C | IDD4(C2H2)/col-IDD4-DAP-Seq(GSE60143)/Homer | 1e-44 | -1.032e+02 | 0.0000 | 747.0 | 46.20% | 2046.5 | 27.76% | motif file (matrix) | svg |
| 13 | T C G A A C T G C A T G A G C T A G T C C G T A C T G A C T A G A C T G C G A T A T G C C T G A | RAR:RXR(NR),DR0/ES-RAR-ChIP-Seq(GSE56893)/Homer | 1e-44 | -1.017e+02 | 0.0000 | 241.0 | 14.90% | 328.9 | 4.46% | motif file (matrix) | svg |
| 14 | A G C T A G C T A G C T A C T G A C G T A G T C A C T G A C G T G A C T C G A T G C A T A T C G | IDD7(C2H2)/col-IDD7-DAP-Seq(GSE60143)/Homer | 1e-37 | -8.606e+01 | 0.0000 | 556.0 | 34.38% | 1411.6 | 19.15% | motif file (matrix) | svg |
| 15 | G A C T A C G T A C G T A C T G A C G T A G T C G C A T A G C T G C A T G C A T G A C T A G C T | SGR5(C2H2)/colamp-SGR5-DAP-Seq(GSE60143)/Homer | 1e-37 | -8.536e+01 | 0.0000 | 694.0 | 42.92% | 1949.9 | 26.45% | motif file (matrix) | svg |
| 16 | G T A C C T G A T A G C C G T A G C T A T C G A T G C A T G A C C T A G G T C A A G T C C G T A C T G A C T G A C G T A | At1g14580(C2H2)/colamp-At1g14580-DAP-Seq(GSE60143)/Homer | 1e-35 | -8.096e+01 | 0.0000 | 244.0 | 15.09% | 398.0 | 5.40% | motif file (matrix) | svg |
| 17 | C A T G A G C T T A C G G T C A G T A C T A G C A G C T G A C T A T C G T C G A | Esrrb(NR)/mES-Esrrb-ChIP-Seq(GSE11431)/Homer | 1e-34 | -8.021e+01 | 0.0000 | 755.0 | 46.69% | 2234.9 | 30.32% | motif file (matrix) | svg |
| 18 | A G T C G A C T A G C T C G A T A T C G G C T A C G A T A T C G C G A T A C T G T A C G A C G T | Tcf7(HMG)/GM12878-TCF7-ChIP-Seq(Encode)/Homer | 1e-34 | -7.968e+01 | 0.0000 | 453.0 | 28.01% | 1074.4 | 14.58% | motif file (matrix) | svg |
| 19 | C A T G G A C T C T A G C A T G C A G T C G A T C T A G C A T G C G A T C G T A C T A G C A G T C G A T C T A G C A T G | AT1G24250(Orphan)/col-AT1G24250-DAP-Seq(GSE60143)/Homer | 1e-34 | -7.966e+01 | 0.0000 | 483.0 | 29.87% | 1181.7 | 16.03% | motif file (matrix) | svg |
| 20 | C T G A A T G C G C T A G C A T A T G C C G T A T C G A C T G A C T A G T C A G T A C G G T C A | Tcf4(HMG)/Hct116-Tcf4-ChIP-Seq(SRA012054)/Homer | 1e-33 | -7.799e+01 | 0.0000 | 572.0 | 35.37% | 1521.3 | 20.64% | motif file (matrix) | svg |
| 21 | C T G A C G A T C T A G T C A G G A T C C T G A T C A G G A T C C T G A A C T G A G T C G C T A A C G T A G T C G C A T | PRDM9(Zf)/Testis-DMC1-ChIP-Seq(GSE35498)/Homer | 1e-33 | -7.754e+01 | 0.0000 | 457.0 | 28.26% | 1100.0 | 14.92% | motif file (matrix) | svg |
| 22 | T G A C C T A G T C A G G T C A C G T A T C A G C G A T T C A G T C G A T G C A C T G A T A G C | PU.1-IRF(ETS:IRF)/Bcell-PU.1-ChIP-Seq(GSE21512)/Homer | 1e-33 | -7.651e+01 | 0.0000 | 784.0 | 48.48% | 2383.4 | 32.33% | motif file (matrix) | svg |
| 23 | A G T C A C G T A C G T T A C G G C A T G C A T A T G C G C T A C G T A A T G C C G T A G T C A A C T G G A T C G C A T | ANAC075(NAC)/col-ANAC075-DAP-Seq(GSE60143)/Homer | 1e-33 | -7.636e+01 | 0.0000 | 636.0 | 39.33% | 1780.9 | 24.16% | motif file (matrix) | svg |
| 24 | A C G T A C G T A C G T A C G T A C G T A C G T A C G T A C G T A C G T A C G T | VRN1(ABI3VP1)/col-VRN1-DAP-Seq(GSE60143)/Homer | 1e-32 | -7.524e+01 | 0.0000 | 82.0 | 5.07% | 40.4 | 0.55% | motif file (matrix) | svg |
| 25 | A G T C A G T C C G A T A C G T A C G T A C T G A C G T A G C T A G T C A G T C | Sox4(HMG)/proB-Sox4-ChIP-Seq(GSE50066)/Homer | 1e-31 | -7.324e+01 | 0.0000 | 799.0 | 49.41% | 2472.8 | 33.55% | motif file (matrix) | svg |
| 26 | A T G C A G T C A G C T A G C T A C G T A T C G C G T A C G A T T A G C G A C T | LEF1(HMG)/H1-LEF1-ChIP-Seq(GSE64758)/Homer | 1e-31 | -7.264e+01 | 0.0000 | 729.0 | 45.08% | 2183.9 | 29.63% | motif file (matrix) | svg |
| 27 | C G A T C T A G C G T A G A C T C A G T C T A G C G T A A G C T C A T G C T A G | HOXA1(Homeobox)/mES-Hoxa1-ChIP-Seq(SRP084292)/Homer | 1e-31 | -7.256e+01 | 0.0000 | 501.0 | 30.98% | 1289.2 | 17.49% | motif file (matrix) | svg |
| 28 | G C A T G A T C C T A G C G T A G C A T C G T A G C A T A G T C C T A G C G T A G C A T C G A T | AT5G22990(C2H2)/col-AT5G22990-DAP-Seq(GSE60143)/Homer | 1e-31 | -7.250e+01 | 0.0000 | 881.0 | 54.48% | 2833.7 | 38.44% | motif file (matrix) | svg |
| 29 | A G C T A C G T A C T G A T G C A G T C C G T A C T G A T A C G | NF1-halfsite(CTF)/LNCaP-NF1-ChIP-Seq(Unpublished)/Homer | 1e-30 | -7.091e+01 | 0.0000 | 1141.0 | 70.56% | 4055.8 | 55.02% | motif file (matrix) | svg |
| 30 | G T C A T G C A T G C A G C T A C G T A G C T A G C T A G C T A | REM19(REM)/colamp-REM19-DAP-Seq(GSE60143)/Homer | 1e-30 | -6.947e+01 | 0.0000 | 399.0 | 24.68% | 941.8 | 12.78% | motif file (matrix) | svg |
| 31 | C G A T C T G A A G T C A C G T A C G T A T C G G A C T C T A G C G A T G A C T C G T A A T G C C G T A G T C A A C T G | ANAC011(NAC)/col-ANAC011-DAP-Seq(GSE60143)/Homer | 1e-30 | -6.935e+01 | 0.0000 | 470.0 | 29.07% | 1195.8 | 16.22% | motif file (matrix) | svg |
| 32 | G T A C T C G A T A G C C G T A C G T A C T G A T G C A T G A C A C T G C G T A A G T C C T G A C T G A T C G A C G T A | NUC(C2H2)/col-NUC-DAP-Seq(GSE60143)/Homer | 1e-29 | -6.771e+01 | 0.0000 | 191.0 | 11.81% | 295.1 | 4.00% | motif file (matrix) | svg |
| 33 | T C G A T C G A C T G A C G T A A C T G A T G C A C G T A G T C | Lola-I(Zf)/Embryo-LolaI-ChIP-Seq(GSE200870)/Homer | 1e-28 | -6.575e+01 | 0.0000 | 641.0 | 39.64% | 1877.1 | 25.47% | motif file (matrix) | svg |
| 34 | T A C G G A C T T G A C C G T A A C G T G A T C G T C A C G T A A C G T A T G C C G T A G A C T | HOXA2(Homeobox)/mES-Hoxa2-ChIP-Seq(Donaldson\_et\_al.)/Homer | 1e-28 | -6.517e+01 | 0.0000 | 251.0 | 15.52% | 477.6 | 6.48% | motif file (matrix) | svg |
| 35 | A G C T A G C T A G C T A C T G A C G T A G T C A C T G A C G T G C A T G C A T G C A T A C G T | At5g66730(C2H2)/colamp-At5g66730-DAP-Seq(GSE60143)/Homer | 1e-28 | -6.509e+01 | 0.0000 | 416.0 | 25.73% | 1026.5 | 13.93% | motif file (matrix) | svg |
| 36 | C G A T C T A G A C G T A C G T A C G T C G T A A G C T C G A T A G C T C G T A C T A G T A G C | FoxD3(forkhead)/ZebrafishEmbryo-Foxd3.biotin-ChIP-seq(GSE106676)/Homer | 1e-28 | -6.483e+01 | 0.0000 | 677.0 | 41.87% | 2030.7 | 27.55% | motif file (matrix) | svg |
| 37 | C T G A C T A G A C T G G C A T A T G C C G T A C T G A C T A G A C T G A C G T A G T C C T G A | RARg(NR)/ES-RARg-ChIP-Seq(GSE30538)/Homer | 1e-26 | -6.216e+01 | 0.0000 | 83.0 | 5.13% | 58.8 | 0.80% | motif file (matrix) | svg |
| 38 | C T G A A T G C G C T A C G A T A T G C C G T A C G T A C G T A C T A G T A C G | Tcf3(HMG)/mES-Tcf3-ChIP-Seq(GSE11724)/Homer | 1e-26 | -6.209e+01 | 0.0000 | 379.0 | 23.44% | 913.0 | 12.39% | motif file (matrix) | svg |
| 39 | C G A T A C G T A C G T A G C T A G C T G A T C G A T C G C T A A G C T A C G T A T C G T A C G | NFATC2(RHD)/Islets-NFATC2-ChIP-Seq(GSE158496)/Homer | 1e-26 | -6.080e+01 | 0.0000 | 1108.0 | 68.52% | 3986.7 | 54.08% | motif file (matrix) | svg |
| 40 | A T C G T C G A G A C T A T C G T G A C A C G T C T A G A C T G C G T A A C T G A G T C G T A C | ZNF415(Zf)/HEK293-ZNF415.GFP-ChIP-Seq(GSE58341)/Homer | 1e-25 | -5.942e+01 | 0.0000 | 597.0 | 36.92% | 1750.7 | 23.75% | motif file (matrix) | svg |
| 41 | T C G A T G C A C A G T T C G A G A T C A G T C C G T A C G T A A C T G A G T C C G T A C G T A T C A G C G A T A G T C | AT5G25475(ABI3VP1)/col-AT5G25475-DAP-Seq(GSE60143)/Homer | 1e-25 | -5.930e+01 | 0.0000 | 1053.0 | 65.12% | 3737.7 | 50.71% | motif file (matrix) | svg |
| 42 | G T A C A C G T A C G T A T C G C A G T C G A T A T C G G C T A T G C A T A G C C G T A G T C A C A T G A G C T G C T A | ANAC013(NAC)/col-ANAC013-DAP-Seq(GSE60143)/Homer | 1e-25 | -5.917e+01 | 0.0000 | 596.0 | 36.86% | 1748.4 | 23.72% | motif file (matrix) | svg |
| 43 | G A C T G C A T A C G T A C G T A C T G C G T A A G T C A G C T C G A T A T C G G C A T A C T G C G A T C T A G C G T A | WRKY50(WRKY)/col-WRKY50-DAP-Seq(GSE60143)/Homer | 1e-25 | -5.897e+01 | 0.0000 | 1050.0 | 64.94% | 3726.4 | 50.55% | motif file (matrix) | svg |
| 44 | G A T C G C A T A G T C A G C T G A T C A G C T G A T C G A C T G A T C G A C T A G T C A C G T G A T C A G C T G A T C | GAGA-repeat/SacCer-Promoters/Homer | 1e-25 | -5.866e+01 | 0.0000 | 1337.0 | 82.68% | 5175.7 | 70.22% | motif file (matrix) | svg |
| 45 | C A T G G A C T T A C G G T C A G T A C G A T C G A C T A G C T A T C G T C G A T A C G T A G C | ERRg(NR)/Kidney-ESRRG-ChIP-Seq(GSE104905)/Homer | 1e-25 | -5.813e+01 | 0.0000 | 829.0 | 51.27% | 2730.8 | 37.05% | motif file (matrix) | svg |
| 46 | A T G C T C G A T A C G A C G T A T G C A G T C A C G T A G T C A G T C G A T C | Znf263(Zf)/K562-Znf263-ChIP-Seq(GSE31477)/Homer | 1e-25 | -5.779e+01 | 0.0000 | 984.0 | 60.85% | 3429.4 | 46.52% | motif file (matrix) | svg |
| 47 | C T A G C T A G T C G A C T A G C G T A A T C G T C G A A C T G C T G A T C G A C T G A T A C G | FRS9(ND)/col-FRS9-DAP-Seq(GSE60143)/Homer | 1e-24 | -5.604e+01 | 0.0000 | 169.0 | 10.45% | 273.5 | 3.71% | motif file (matrix) | svg |
| 48 | C T A G C T A G A T G C G T A C T C A G A T G C A G T C G C A T G A T C G A T C | ZNF91(Zf)/HEK-ZNF91.HA-ChIP-Seq(GSE162571)/Homer | 1e-24 | -5.600e+01 | 0.0000 | 677.0 | 41.87% | 2103.6 | 28.54% | motif file (matrix) | svg |
| 49 | A T G C G A T C C G A T A C G T A C G T A C T G C A G T A G C T | Sox3(HMG)/NPC-Sox3-ChIP-Seq(GSE33059)/Homer | 1e-23 | -5.445e+01 | 0.0000 | 1191.0 | 73.65% | 4456.4 | 60.46% | motif file (matrix) | svg |
| 50 | A C G T A G T C A G T C C G A T A C G T A C G T A C T G A C G T A T G C G A C T A C T G T A C G | Sox21(HMG)/ESC-SOX21-ChIP-Seq(GSE110505)/Homer | 1e-23 | -5.405e+01 | 0.0000 | 1156.0 | 71.49% | 4286.8 | 58.16% | motif file (matrix) | svg |
| 51 | G T C A T C G A T C G A C G T A G C T A C G T A T C G A T G A C A C T G C G T A A G T C C G T A C G T A T C G A G C T A | IDD2(C2H2)/colamp-IDD2-DAP-Seq(GSE60143)/Homer | 1e-23 | -5.403e+01 | 0.0000 | 206.0 | 12.74% | 387.5 | 5.26% | motif file (matrix) | svg |
| 52 | C A G T A C T G T C A G T G C A G C T A A T G C T C G A A T C G G T C A T G C A | ZNF189(Zf)/HEK293-ZNF189.GFP-ChIP-Seq(GSE58341)/Homer | 1e-23 | -5.333e+01 | 0.0000 | 660.0 | 40.82% | 2056.8 | 27.90% | motif file (matrix) | svg |
| 53 | C G T A C T A G C A T G A G C T C T G A C A T G C A G T C G A T C T A G C T A G | MYB30(MYB)/colamp-MYB30-DAP-Seq(GSE60143)/Homer | 1e-22 | -5.286e+01 | 0.0000 | 1023.0 | 63.27% | 3659.7 | 49.65% | motif file (matrix) | svg |
| 54 | C T G A T C G A G T A C A C G T A C G T A T C G A C G T C G A T A T C G G C T A G T A C A T G C C G T A T G C A C A T G | ANAC103(NAC)/col-ANAC103-DAP-Seq(GSE60143)/Homer | 1e-22 | -5.243e+01 | 0.0000 | 541.0 | 33.46% | 1584.8 | 21.50% | motif file (matrix) | svg |
| 55 | A T G C G T A C A G T C A G T C A C G T A C G T C G A T A C G T | AT5G02460(C2C2dof)/col-AT5G02460-DAP-Seq(GSE60143)/Homer | 1e-22 | -5.195e+01 | 0.0000 | 1197.0 | 74.03% | 4511.5 | 61.20% | motif file (matrix) | svg |
| 56 | C G A T T C G A G A T C A C G T A C G T T C A G G C T A C G A T C G T A C G T A C G T A A T G C C G T A T G C A C T A G | ANAC028(NAC)/col-ANAC028-DAP-Seq(GSE60143)/Homer | 1e-21 | -4.991e+01 | 0.0000 | 835.0 | 51.64% | 2835.3 | 38.46% | motif file (matrix) | svg |
| 57 | G T C A G C T A G C T A T C G A A T C G A C G T A G T C T C G A T C G A T G A C | WRKY40(WRKY)/colamp-WRKY40-DAP-Seq(GSE60143)/Homer | 1e-21 | -4.954e+01 | 0.0000 | 775.0 | 47.93% | 2578.0 | 34.97% | motif file (matrix) | svg |
| 58 | A T G C A G T C G A T C C G T A A C G T A C G T A C T G A C G T A G C T G A T C | Sox2(HMG)/mES-Sox2-ChIP-Seq(GSE11431)/Homer | 1e-21 | -4.904e+01 | 0.0000 | 828.0 | 51.21% | 2813.3 | 38.17% | motif file (matrix) | svg |
| 59 | C G T A C G A T C T A G C G T A A G C T C A G T T A C G C G T A A C G T C A T G C T A G A T G C | HOXA3(Homeobox)/mEmbryo-Hoxa3-ChIP-Seq(E-MTAB-8607)/Homer | 1e-20 | -4.800e+01 | 0.0000 | 312.0 | 19.29% | 763.9 | 10.36% | motif file (matrix) | svg |
| 60 | C T G A C G A T C A T G A T C G G C A T C A T G G C T A A G T C | ASHR1(ND)/col-ASHR1-DAP-Seq(GSE60143)/Homer | 1e-20 | -4.750e+01 | 0.0000 | 1019.0 | 63.02% | 3696.0 | 50.14% | motif file (matrix) | svg |
| 61 | A T G C G C A T T A G C C G A T T A G C G C A T T A G C G C A T A T G C G A C T | GAGA-repeat/Arabidopsis-Promoters/Homer | 1e-20 | -4.688e+01 | 0.0000 | 682.0 | 42.18% | 2207.7 | 29.95% | motif file (matrix) | svg |
| 62 | A T C G T G C A G A T C C T A G A C G T A T C G C G T A A G T C T C A G A C T G T C A G G C T A | Knotted(Homeobox)/Corn-KN1-ChIP-Seq(GSE39161)/Homer | 1e-20 | -4.665e+01 | 0.0000 | 1271.0 | 78.60% | 4942.7 | 67.05% | motif file (matrix) | svg |
| 63 | C A T G G T A C A C T G G T C A A G C T T A C G T G C A A T C G T G A C C A G T | TOD6?/SacCer-Promoters/Homer | 1e-20 | -4.661e+01 | 0.0000 | 367.0 | 22.70% | 968.8 | 13.14% | motif file (matrix) | svg |
| 64 | C T G A T C A G A G T C C G T A A T C G A T G C C G A T A C T G A G T C G A C T A T C G A G T C | MyoD(bHLH)/Myotube-MyoD-ChIP-Seq(GSE21614)/Homer | 1e-20 | -4.627e+01 | 0.0000 | 509.0 | 31.48% | 1509.9 | 20.48% | motif file (matrix) | svg |
| 65 | T A G C C G T A C T G A T A C G C G T A A C G T A C T G A C T G A G T C T A C G C T A G G T A C | YY1(Zf)/Promoter/Homer | 1e-19 | -4.575e+01 | 0.0000 | 117.0 | 7.24% | 167.3 | 2.27% | motif file (matrix) | svg |
| 66 | G A C T C A G T G C A T C G A T T G A C A C G T A T G C G T A C C T G A A C T G A C T G A G C T | WIP5(C2H2)/colamp-WIP5-DAP-Seq(GSE60143)/Homer | 1e-19 | -4.509e+01 | 0.0000 | 1004.0 | 62.09% | 3652.2 | 49.55% | motif file (matrix) | svg |
| 67 | T C G A G T A C C A T G A G C T A C G T C A T G G T C A G T A C A G C T G C T A C G A T C A G T | WRKY31(WRKY)/colamp-WRKY31-DAP-Seq(GSE60143)/Homer | 1e-19 | -4.507e+01 | 0.0000 | 856.0 | 52.94% | 2978.6 | 40.41% | motif file (matrix) | svg |
| 68 | G A C T T C A G G C A T A G T C G C T A G A T C C T G A A C G T A G T C G T C A | Replumless(BLH)/Arabidopsis-RPL.GFP-ChIP-Seq(GSE78727)/Homer | 1e-19 | -4.433e+01 | 0.0000 | 1153.0 | 71.30% | 4373.8 | 59.34% | motif file (matrix) | svg |
| 69 | A T G C A G T C G C A T A G C T A C G T T C A G C G A T A G C T G A T C A T C G | Sox10(HMG)/SciaticNerve-Sox3-ChIP-Seq(GSE35132)/Homer | 1e-19 | -4.403e+01 | 0.0000 | 1129.0 | 69.82% | 4259.6 | 57.79% | motif file (matrix) | svg |
| 70 | C G A T C G T A G T A C A C G T A C G T T C A G G C A T C A G T T A C G G T C A C G T A A G T C C G T A T G C A C A T G | ANAC053(NAC)/colamp-ANAC053-DAP-Seq(GSE60143)/Homer | 1e-19 | -4.379e+01 | 0.0000 | 698.0 | 43.17% | 2305.6 | 31.28% | motif file (matrix) | svg |
| 71 | A G T C T G C A T C G A C T G A A C T G C A T G A C G T A T G C G T C A T A C G | Erra(NR)/HepG2-Erra-ChIP-Seq(GSE31477)/Homer | 1e-18 | -4.280e+01 | 0.0000 | 1127.0 | 69.70% | 4263.3 | 57.84% | motif file (matrix) | svg |
| 72 | C G T A G A T C A G C T A C G T A C G T A C T G C G T A G T A C A G C T G C T A C G A T C G A T C G A T G C A T G C T A | WRKY18(WRKY)/col-WRKY18-DAP-Seq(GSE60143)/Homer | 1e-18 | -4.220e+01 | 0.0000 | 1262.0 | 78.05% | 4943.9 | 67.07% | motif file (matrix) | svg |
| 73 | C G T A A C G T A C G T A C G T A C G T A G T C A G T C C T G A A G C T A G C T | NFAT(RHD)/Jurkat-NFATC1-ChIP-Seq(Jolma\_et\_al.)/Homer | 1e-18 | -4.192e+01 | 0.0000 | 747.0 | 46.20% | 2534.5 | 34.38% | motif file (matrix) | svg |
| 74 | A G C T G A C T A C G T A C T G A C G T A G T C A C T G A C G T G C A T C G A T | AtIDD11(C2H2)/colamp-AtIDD11-DAP-Seq(GSE60143)/Homer | 1e-18 | -4.170e+01 | 0.0000 | 503.0 | 31.11% | 1525.9 | 20.70% | motif file (matrix) | svg |
| 75 | G C A T A G C T A G C T A G C T A C T G A C G T A G T C A C T G A C G T G A C T C G A T G C A T | JKD(C2H2)/col-JKD-DAP-Seq(GSE60143)/Homer | 1e-18 | -4.150e+01 | 0.0000 | 288.0 | 17.81% | 720.3 | 9.77% | motif file (matrix) | svg |
| 76 | T C A G A C G T T C G A T A G C A G T C C G T A A C T G G T A C A C G T A C T G A T C G A G T C | Atoh1(bHLH)/Cerebellum-Atoh1-ChIP-Seq(GSE22111)/Homer | 1e-17 | -4.137e+01 | 0.0000 | 777.0 | 48.05% | 2670.2 | 36.23% | motif file (matrix) | svg |
| 77 | A G T C C G T A A C G T A G T C A C G T A C T G | Tal1 | 1e-17 | -4.129e+01 | 0.0000 | 1006.0 | 62.21% | 3704.6 | 50.26% | motif file (matrix) | svg |
| 78 | C G A T C G T A G T A C A C G T A C G T T C A G G C A T C A G T T A C G G T C A C G T A A G T C C G T A T G C A C A T G | NAC2(NAC)/colamp-NAC2-DAP-Seq(GSE60143)/Homer | 1e-17 | -4.112e+01 | 0.0000 | 772.0 | 47.74% | 2651.8 | 35.98% | motif file (matrix) | svg |
| 79 | C G A T T G A C C A T G G A C T A C G T C A T G C G T A G A T C G A C T G C A T G C A T C G A T | WRKY14(WRKY)/colamp-WRKY14-DAP-Seq(GSE60143)/Homer | 1e-17 | -4.083e+01 | 0.0000 | 670.0 | 41.43% | 2217.3 | 30.08% | motif file (matrix) | svg |
| 80 | C G T A C T A G C A T G G A C T C T G A A C T G A C G T A C G T C T A G C T A G C A T G T C G A | MYB94(MYB)/col-MYB94-DAP-Seq(GSE60143)/Homer | 1e-17 | -4.079e+01 | 0.0000 | 513.0 | 31.73% | 1573.5 | 21.35% | motif file (matrix) | svg |
| 81 | C T G A T C G A C G T A A T G C C G T A C G T A C G A T C T A G T C A G G A T C | Sox15(HMG)/CPA-Sox15-ChIP-Seq(GSE62909)/Homer | 1e-17 | -4.062e+01 | 0.0000 | 877.0 | 54.24% | 3121.1 | 42.34% | motif file (matrix) | svg |
| 82 | T A C G T A G C G C T A C G A T C T A G A C G T C A G T C A G T G C T A A G T C G T C A G C A T | FOXK2(Forkhead)/U2OS-FOXK2-ChIP-Seq(E-MTAB-2204)/Homer | 1e-17 | -4.000e+01 | 0.0000 | 715.0 | 44.22% | 2417.1 | 32.79% | motif file (matrix) | svg |
| 83 | A G T C A C G T A C G T T A C G G C T A G C T A C G T A C G A T C G A T A T G C C G T A G T C A A C T G G A C T G C A T | SND2(NAC)/colamp-SND2-DAP-Seq(GSE60143)/Homer | 1e-17 | -3.999e+01 | 0.0000 | 795.0 | 49.17% | 2764.9 | 37.51% | motif file (matrix) | svg |
| 84 | G C T A G C T A C G T A C G T A C T G A C T A G A C G T A G T C C G T A C T G A G T A C A C T G | WRKY65(WRKY)/colamp-WRKY65-DAP-Seq(GSE60143)/Homer | 1e-17 | -3.952e+01 | 0.0000 | 688.0 | 42.55% | 2307.8 | 31.31% | motif file (matrix) | svg |
| 85 | T C G A A G T C C G T A A T C G A T G C C G A T A C T G A G T C A G C T A C T G | Tcf12(bHLH)/GM12878-Tcf12-ChIP-Seq(GSE32465)/Homer | 1e-17 | -3.943e+01 | 0.0000 | 590.0 | 36.49% | 1898.4 | 25.75% | motif file (matrix) | svg |
| 86 | G C A T C G A T C G T A G A T C C A T G A C G T A C G T A C T G C G T A A G T C A G C T G C A T G C A T C G T A G C T A | WRKY45(WRKY)/col-WRKY45-DAP-Seq(GSE60143)/Homer | 1e-16 | -3.911e+01 | 0.0000 | 554.0 | 34.26% | 1754.6 | 23.80% | motif file (matrix) | svg |
| 87 | C G A T T C G A G A T C C G A T G C A T T C A G G A C T C G A T G C A T G C T A C T G A A G T C C G T A G T C A C T A G | ANAC005(NAC)/col-ANAC005-DAP-Seq(GSE60143)/Homer | 1e-16 | -3.907e+01 | 0.0000 | 518.0 | 32.03% | 1609.3 | 21.83% | motif file (matrix) | svg |
| 88 | G A C T C T A G A T G C A G T C G T C A T A C G A T G C A T C G | HIC1(Zf)/Treg-ZBTB29-ChIP-Seq(GSE99889)/Homer | 1e-16 | -3.879e+01 | 0.0000 | 1204.0 | 74.46% | 4688.9 | 63.61% | motif file (matrix) | svg |
| 89 | C G T A T G A C T A G C T G C A A C T G A C T G C G T A C G T A T C A G G A C T | ELF3(ETS)/PDAC-ELF3-ChIP-Seq(GSE64557)/Homer | 1e-16 | -3.876e+01 | 0.0000 | 548.0 | 33.89% | 1733.6 | 23.52% | motif file (matrix) | svg |
| 90 | T C G A A G T C A C G T A C G T T C A G C A G T C T G A C T A G T C G A C G T A A T C G C G T A C G T A A C T G A G C T | NTM1(NAC)/col-NTM1-DAP-Seq(GSE60143)/Homer | 1e-16 | -3.848e+01 | 0.0000 | 533.0 | 32.96% | 1675.7 | 22.73% | motif file (matrix) | svg |
| 91 | C T A G A C T G T G C A G T C A A T G C C G T A A T C G A T G C A G T C C T A G | ZNF341(Zf)/EBV-ZNF341-ChIP-Seq(GSE113194)/Homer | 1e-16 | -3.847e+01 | 0.0000 | 626.0 | 38.71% | 2057.1 | 27.91% | motif file (matrix) | svg |
| 92 | G C T A C G T A G C T A C G T A C T G A A C T G A C G T G T A C C G T A C T G A G T A C A C T G | WRKY22(WRKY)/colamp-WRKY22-DAP-Seq(GSE60143)/Homer | 1e-16 | -3.830e+01 | 0.0000 | 733.0 | 45.33% | 2513.8 | 34.10% | motif file (matrix) | svg |
| 93 | A G C T C T G A C T A G C T A G A C T G T A G C T G C A T C G A C T G A C T A G C A T G A C G T A T G C T C G A | RXR(NR),DR1/3T3L1-RXR-ChIP-Seq(GSE13511)/Homer | 1e-16 | -3.761e+01 | 0.0000 | 669.0 | 41.37% | 2247.3 | 30.49% | motif file (matrix) | svg |
| 94 | G A C T A C G T A G C T G A C T A C T G C A G T A G T C A T C G A C G T G C A T G C A T G C A T | MGP(C2H2)/colamp-MGP-DAP-Seq(GSE60143)/Homer | 1e-16 | -3.741e+01 | 0.0000 | 431.0 | 26.65% | 1281.4 | 17.38% | motif file (matrix) | svg |
| 95 | A G T C C A T G A C G T A C G T A C T G C G T A A G T C G A C T G C A T G C T A | WRKY28(WRKY)/col-WRKY28-DAP-Seq(GSE60143)/Homer | 1e-16 | -3.688e+01 | 0.0000 | 1121.0 | 69.33% | 4303.4 | 58.38% | motif file (matrix) | svg |
| 96 | T C A G A T C G G A C T A C T G G A C T C A G T C T A G C G T A G T A C C G T A C T A G A T C G | Tbx20(T-box)/Heart-Tbx20-ChIP-Seq(GSE29636)/Homer | 1e-15 | -3.684e+01 | 0.0000 | 314.0 | 19.42% | 845.6 | 11.47% | motif file (matrix) | svg |
| 97 | C G A T T C G A G A T C A C G T A C G T T C A G G C A T C T G A C G T A G C T A C G T A A G T C C G T A T G C A C A T G | ANAC050(NAC)/colamp-ANAC050-DAP-Seq(GSE60143)/Homer | 1e-15 | -3.638e+01 | 0.0000 | 721.0 | 44.59% | 2483.8 | 33.70% | motif file (matrix) | svg |
| 98 | A G T C A C G T A C G T T C A G G C T A G C T A A T G C C G T A C G A T A G T C C G T A G T C A A C T G G A T C G C A T | SND3(NAC)/col-SND3-DAP-Seq(GSE60143)/Homer | 1e-15 | -3.630e+01 | 0.0000 | 868.0 | 53.68% | 3131.7 | 42.49% | motif file (matrix) | svg |
| 99 | G A T C G A C T G A C T A C G T A G T C A C G T A G T C A C G T A G T C A C G T A G T C A C G T G T A C C G A T G T C A | BPC6(BBRBPC)/col-BPC6-DAP-Seq(GSE60143)/Homer | 1e-15 | -3.595e+01 | 0.0000 | 38.0 | 2.35% | 18.6 | 0.25% | motif file (matrix) | svg |
| 100 | C G A T C T A G C T G A A T G C C T G A T C G A C G T A C T G A T C G A T A G C A G T C C G T A A C T G T C G A A T G C | Hand2(bHLH)/Mesoderm-Hand2-ChIP-Seq(GSE61475)/Homer | 1e-15 | -3.564e+01 | 0.0000 | 363.0 | 22.45% | 1036.0 | 14.05% | motif file (matrix) | svg |
| 101 | A G T C G A T C C T G A A G T C A G T C C A T G G T C A G A T C C G T A G A T C | DREB26(AP2EREBP)/col-DREB26-DAP-Seq(GSE60143)/Homer | 1e-15 | -3.560e+01 | 0.0000 | 424.0 | 26.22% | 1270.9 | 17.24% | motif file (matrix) | svg |
| 102 | G T A C G C T A T C A G C T G A C T A G C A T G A G C T G A T C T G C A T C G A C T G A A C T G C A G T A G T C G A T C G C T A | HNF4a(NR),DR1/HepG2-HNF4a-ChIP-Seq(GSE25021)/Homer | 1e-15 | -3.550e+01 | 0.0000 | 373.0 | 23.07% | 1075.5 | 14.59% | motif file (matrix) | svg |
| 103 | C A G T T C A G T C G A A G T C C G T A A C T G T G A C C G A T A C T G A C T G A C G T A T C G | Atoh7(bHLH)/Retina-Atoh7-CutnRun(GSE156756)/Homer | 1e-15 | -3.549e+01 | 0.0000 | 537.0 | 33.21% | 1721.9 | 23.36% | motif file (matrix) | svg |
| 104 | G A C T A C T G C A G T A G T C A C T G A C T G A G C T A C T G C T A G G T C A | At1g77640(AP2EREBP)/col-At1g77640-DAP-Seq(GSE60143)/Homer | 1e-15 | -3.549e+01 | 0.0000 | 378.0 | 23.38% | 1094.9 | 14.85% | motif file (matrix) | svg |
| 105 | C A G T C G T A C G T A G C A T G A C T G C A T G T A C A G C T A C T G G A C T A C G T C A T G | RAV1(RAV)/colamp-RAV1-DAP-Seq(GSE60143)/Homer | 1e-15 | -3.519e+01 | 0.0000 | 625.0 | 38.65% | 2088.4 | 28.33% | motif file (matrix) | svg |
| 106 | G A T C C A G T T A G C A G T C A C T G A G T C A G T C C T A G G A C T G T A C | LEP(AP2EREBP)/col-LEP-DAP-Seq(GSE60143)/Homer | 1e-15 | -3.511e+01 | 0.0000 | 419.0 | 25.91% | 1255.6 | 17.03% | motif file (matrix) | svg |
| 107 | C T A G G T A C C A T G G A C T C G A T C A T G G T C A G T A C G A C T C G A T C G A T C G A T | WRKY27(WRKY)/colamp-WRKY27-DAP-Seq(GSE60143)/Homer | 1e-15 | -3.462e+01 | 0.0000 | 932.0 | 57.64% | 3442.4 | 46.70% | motif file (matrix) | svg |
| 108 | A T G C A G T C G T A C A G C T T C G A C T A G G A T C C T G A G T C A A G T C G C T A T C A G | Rfx5(HTH)/GM12878-Rfx5-ChIP-Seq(GSE31477)/Homer | 1e-15 | -3.462e+01 | 0.0000 | 423.0 | 26.16% | 1275.2 | 17.30% | motif file (matrix) | svg |
| 109 | G A C T G C A T C T A G C G A T G A T C T C G A C A T G G A T C | Tgif1(Homeobox)/mES-Tgif1-ChIP-Seq(GSE55404)/Homer | 1e-15 | -3.456e+01 | 0.0000 | 1398.0 | 86.46% | 5745.2 | 77.94% | motif file (matrix) | svg |
| 110 | C G A T C G T A A G T C A C G T A C G T T C G A T G C A G C A T G C T A C G T A A C G T A G C T C G T A C G T A A C T G | ANAC062(NAC)/colamp-ANAC062-DAP-Seq(GSE60143)/Homer | 1e-14 | -3.441e+01 | 0.0000 | 433.0 | 26.78% | 1316.9 | 17.87% | motif file (matrix) | svg |
| 111 | C T G A C T G A C T A G T C G A C G T A A T G C C G T A A C T G C G T A A C G T C T G A C G A T A G C T C G T A A C G T A G T C C G A T T A C G G T C A G C A T | GATA(Zf),IR3/iTreg-Gata3-ChIP-Seq(GSE20898)/Homer | 1e-14 | -3.379e+01 | 0.0000 | 271.0 | 16.76% | 712.1 | 9.66% | motif file (matrix) | svg |
| 112 | C A T G C T A G A G C T G A C T C A T G A G T C G A T C G C T A C G A T C T A G T C A G G T A C C T G A T C G A | X-box(HTH)/NPC-H3K4me1-ChIP-Seq(GSE16256)/Homer | 1e-14 | -3.376e+01 | 0.0000 | 144.0 | 8.91% | 287.0 | 3.89% | motif file (matrix) | svg |
| 113 | A T G C G T A C C T G A A G T C A G T C A C T G G T C A A G T C G T C A G C A T G C A T G A C T | At5g65130(AP2EREBP)/colamp-At5g65130-DAP-Seq(GSE60143)/Homer | 1e-14 | -3.346e+01 | 0.0000 | 449.0 | 27.77% | 1388.9 | 18.84% | motif file (matrix) | svg |
| 114 | C G A T T C G A G T A C A C G T A C G T T C A G G C A T G C A T G C T A C G T A C G T A A G T C C G T A T G C A C A T G | ANAC020(NAC)/col-ANAC020-DAP-Seq(GSE60143)/Homer | 1e-14 | -3.287e+01 | 0.0000 | 793.0 | 49.04% | 2839.7 | 38.52% | motif file (matrix) | svg |
| 115 | G A C T G C A T G C A T A G T C A G C T T C G A T A C G G C T A C G T A A C T G G T A C G C A T C G A T A G T C G A C T | HSF3(HSF)/colamp-HSF3-DAP-Seq(GSE60143)/Homer | 1e-14 | -3.232e+01 | 0.0000 | 781.0 | 48.30% | 2793.4 | 37.90% | motif file (matrix) | svg |
| 116 | A C T G T C A G A G C T G A C T C A T G A G T C A G T C G C T A C G A T C T A G T C A G G T A C C T G A T C G A | Rfx1(HTH)/NPC-H3K4me1-ChIP-Seq(GSE16256)/Homer | 1e-13 | -3.214e+01 | 0.0000 | 250.0 | 15.46% | 649.3 | 8.81% | motif file (matrix) | svg |
| 117 | T A C G C T G A C A T G G A T C G T A C G C A T T C A G T A C G A G C T G T C A G A T C G C A T T A C G C G T A C T A G G A T C G A T C C G A T A C T G T C A G | ZNF322(Zf)/HEK293-ZNF322.GFP-ChIP-Seq(GSE58341)/Homer | 1e-13 | -3.210e+01 | 0.0000 | 160.0 | 9.89% | 345.7 | 4.69% | motif file (matrix) | svg |
| 118 | T C A G T A C G T A G C A C G T A C T G C G A T A G T C C G T A T A C G A G T C | Meis1(Homeobox)/MastCells-Meis1-ChIP-Seq(GSE48085)/Homer | 1e-13 | -3.206e+01 | 0.0000 | 1159.0 | 71.68% | 4548.3 | 61.70% | motif file (matrix) | svg |
| 119 | G C A T C G T A C T A G A G T C G T C A C G T A A T G C A C G T A C G T A C T G G A T C G C A T C G T A G C T A G C T A | bHLH122(bHLH)/col100-bHLH122-DAP-Seq(GSE60143)/Homer | 1e-13 | -3.195e+01 | 0.0000 | 847.0 | 52.38% | 3091.2 | 41.94% | motif file (matrix) | svg |
| 120 | A T G C C A T G A C G T A C G T A C T G C G T A A G T C G A C T G C A T C G A T | WRKY71(WRKY)/col-WRKY71-DAP-Seq(GSE60143)/Homer | 1e-13 | -3.179e+01 | 0.0000 | 849.0 | 52.50% | 3102.0 | 42.08% | motif file (matrix) | svg |
| 121 | A G C T C A T G G C A T G A T C T G C A C T A G G A T C A C G T | Tgif2(Homeobox)/mES-Tgif2-ChIP-Seq(GSE55404)/Homer | 1e-13 | -3.174e+01 | 0.0000 | 1411.0 | 87.26% | 5847.3 | 79.33% | motif file (matrix) | svg |
| 122 | T A C G A T G C G A C T A C T G A G C T A G T C G T C A T G C A A C G T A G T C G C T A T G C A | Pknox1(Homeobox)/ES-Prep1-ChIP-Seq(GSE63282)/Homer | 1e-13 | -3.152e+01 | 0.0000 | 312.0 | 19.29% | 879.1 | 11.93% | motif file (matrix) | svg |
| 123 | C G T A G A T C C A T G G C A T G A C T C T A G T C G A T A G C A G C T G C A T | WRKY55(WRKY)/col-WRKY55-DAP-Seq(GSE60143)/Homer | 1e-13 | -3.145e+01 | 0.0000 | 1054.0 | 65.18% | 4051.0 | 54.96% | motif file (matrix) | svg |
| 124 | T C G A T G A C G T A C C G T A C A G T T G A C A C G T A C T G A G C T A G C T | NeuroG2(bHLH)/Fibroblast-NeuroG2-ChIP-Seq(GSE75910)/Homer | 1e-13 | -3.133e+01 | 0.0000 | 979.0 | 60.54% | 3701.6 | 50.22% | motif file (matrix) | svg |
| 125 | T C A G T G A C C A T G G C A T C A G T A C T G C G T A T G A C G A C T C G A T C G A T C G T A | WRKY3(WRKY)/col-WRKY3-DAP-Seq(GSE60143)/Homer | 1e-13 | -3.112e+01 | 0.0000 | 778.0 | 48.11% | 2795.6 | 37.93% | motif file (matrix) | svg |
| 126 | C T G A G C A T A C T G C T A G A G T C A C T G A C T G A G T C A C T G T C A G | AT4G18450(AP2EREBP)/col-AT4G18450-DAP-Seq(GSE60143)/Homer | 1e-13 | -3.111e+01 | 0.0000 | 531.0 | 32.84% | 1743.3 | 23.65% | motif file (matrix) | svg |
| 127 | G T A C A C T G A G T C A G T C C T A G G A T C G T A C C T G A | CRF4(AP2EREBP)/colamp-CRF4-DAP-Seq(GSE60143)/Homer | 1e-13 | -3.094e+01 | 0.0000 | 703.0 | 43.48% | 2470.8 | 33.52% | motif file (matrix) | svg |
| 128 | C G A T A G C T T G C A A C T G A G T C T G A C C T A G G T A C A G T C C G T A G C A T G C A T | ERF13(AP2EREBP)/colamp-ERF13-DAP-Seq(GSE60143)/Homer | 1e-13 | -3.086e+01 | 0.0000 | 1065.0 | 65.86% | 4111.2 | 55.77% | motif file (matrix) | svg |
| 129 | C A G T T C A G A G C T G A C T A C G T A G T C G A T C G A C T C T G A A C T G G A T C C G T A C T G A A G T C G T A C | Rfx6(HTH)/Min6b1-Rfx6.HA-ChIP-Seq(GSE62844)/Homer | 1e-13 | -3.049e+01 | 0.0000 | 849.0 | 52.50% | 3119.9 | 42.33% | motif file (matrix) | svg |
| 130 | C G A T C T G A A G T C A C G T A C G T T A C G G C T A C A T G C T A G G C A T C G A T A G T C C G T A G T C A A C T G | ANAC096(NAC)/colamp-ANAC096-DAP-Seq(GSE60143)/Homer | 1e-13 | -3.028e+01 | 0.0000 | 740.0 | 45.76% | 2639.5 | 35.81% | motif file (matrix) | svg |
| 131 | C G A T C G T A A G T C A C G T A C G T T C A G G C A T C G T A G C T A G C T A C G T A A G T C C G T A G T C A A C T G | ANAC058(NAC)/col-ANAC058-DAP-Seq(GSE60143)/Homer | 1e-13 | -3.024e+01 | 0.0000 | 744.0 | 46.01% | 2657.0 | 36.05% | motif file (matrix) | svg |
| 132 | G A C T G A T C G A T C G C T A G T A C A G T C G C T A C T G A G T A C G A T C G C T A G A C T | MYB13(MYB)/col-MYB13-DAP-Seq(GSE60143)/Homer | 1e-13 | -3.018e+01 | 0.0000 | 731.0 | 45.21% | 2601.6 | 35.29% | motif file (matrix) | svg |
| 133 | T C G A T G A C G C A T A G C T C A G T G A T C G C T A G A T C G A C T A C G T G C A T A G T C | PRDM1(Zf)/Hela-PRDM1-ChIP-Seq(GSE31477)/Homer | 1e-13 | -3.011e+01 | 0.0000 | 411.0 | 25.42% | 1271.7 | 17.25% | motif file (matrix) | svg |
| 134 | G C A T A C G T A C G T A T C G C G T A C G T A C G T A C G T A | At2g41835(C2H2)/col-At2g41835-DAP-Seq(GSE60143)/Homer | 1e-13 | -3.009e+01 | 0.0000 | 469.0 | 29.00% | 1502.8 | 20.39% | motif file (matrix) | svg |
| 135 | C G T A T A C G T C G A A C T G A C T G C G T A C G T A T A C G A G C T T A C G | PU.1(ETS)/ThioMac-PU.1-ChIP-Seq(GSE21512)/Homer | 1e-13 | -3.009e+01 | 0.0000 | 381.0 | 23.56% | 1154.1 | 15.66% | motif file (matrix) | svg |
| 136 | G T A C C A T G A G C T A C G T A C T G C G T A A G T C G A C T G C A T C G A T | WRKY29(WRKY)/colamp-WRKY29-DAP-Seq(GSE60143)/Homer | 1e-13 | -3.003e+01 | 0.0000 | 1016.0 | 62.83% | 3891.5 | 52.79% | motif file (matrix) | svg |
| 137 | G A T C A T G C A G T C C G T A A G T C A G T C A C T G G C T A A G T C C G T A | AT1G44830(AP2EREBP)/col-AT1G44830-DAP-Seq(GSE60143)/Homer | 1e-13 | -2.995e+01 | 0.0000 | 540.0 | 33.40% | 1793.9 | 24.34% | motif file (matrix) | svg |
| 138 | T G A C T A G C T C A G T C G A T C G A C G T A A G T C C G T A C G T A C G A T C T A G T A C G | Sox7(HMG)/ESC-Sox7-ChIP-Seq(GSE133899)/Homer | 1e-12 | -2.991e+01 | 0.0000 | 430.0 | 26.59% | 1348.6 | 18.29% | motif file (matrix) | svg |
| 139 | G C A T T G A C C A T G G A C T C A G T C A T G T C G A G T A C G A C T G C T A C G A T C G A T | WRKY6(WRKY)/colamp-WRKY6-DAP-Seq(GSE60143)/Homer | 1e-12 | -2.984e+01 | 0.0000 | 923.0 | 57.08% | 3463.6 | 46.99% | motif file (matrix) | svg |
| 140 | C T A G T C A G C T G A T C A G T G C A A C T G T C G A T C A G | Trl(Zf)/S2-GAGAfactor-ChIP-Seq(GSE40646)/Homer | 1e-12 | -2.982e+01 | 0.0000 | 1224.0 | 75.70% | 4898.4 | 66.45% | motif file (matrix) | svg |
| 141 | G C A T A C T G C T A G A G T C A C T G A C T G A G T C A C G T | ERF105(AP2EREBP)/colamp-ERF105-DAP-Seq(GSE60143)/Homer | 1e-12 | -2.972e+01 | 0.0000 | 1092.0 | 67.53% | 4255.4 | 57.73% | motif file (matrix) | svg |
| 142 | C G T A A T G C C G A T A C G T A G T C C G T A C G T A C G T A C T A G A T C G | TCFL2(HMG)/K562-TCF7L2-ChIP-Seq(GSE29196)/Homer | 1e-12 | -2.971e+01 | 0.0000 | 141.0 | 8.72% | 297.0 | 4.03% | motif file (matrix) | svg |
| 143 | A G T C C T A G A C G T A C G T A C T G C G T A A G T C A G C T G C T A G C A T | WRKY24(WRKY)/colamp-WRKY24-DAP-Seq(GSE60143)/Homer | 1e-12 | -2.952e+01 | 0.0000 | 931.0 | 57.58% | 3504.5 | 47.54% | motif file (matrix) | svg |
| 144 | A G T C G A T C G C T A C G A T C A G T T A C G C G A T A G C T A G T C A T C G | SOX1(HMG)/NPC-SOX1-ChIP-Seq(GSE138215)/Homer | 1e-12 | -2.939e+01 | 0.0000 | 1210.0 | 74.83% | 4834.1 | 65.58% | motif file (matrix) | svg |
| 145 | A T G C C A G T A G C T A G C T T C A G G T C A T A G C G A C T C G T A C G A T | WRKY20(WRKY)/col-WRKY20-DAP-Seq(GSE60143)/Homer | 1e-12 | -2.918e+01 | 0.0000 | 846.0 | 52.32% | 3123.7 | 42.38% | motif file (matrix) | svg |
| 146 | C T G A T G A C T G A C C G T A A C G T T G A C A G C T C T A G A C G T G A C T | Olig2(bHLH)/Neuron-Olig2-ChIP-Seq(GSE30882)/Homer | 1e-12 | -2.907e+01 | 0.0000 | 1140.0 | 70.50% | 4495.9 | 60.99% | motif file (matrix) | svg |
| 147 | C T G A T C A G G T A C G C T A A C T G T G A C G C A T C A T G | SCL(bHLH)/HPC7-Scl-ChIP-Seq(GSE13511)/Homer | 1e-12 | -2.906e+01 | 0.0000 | 1388.0 | 85.84% | 5753.1 | 78.05% | motif file (matrix) | svg |
| 148 | C T A G T C G A T G A C A G T C C G T A A C T G G T A C A C G T A C T G A C T G | BHLHA15(bHLH)/NIH3T3-BHLHB8.HA-ChIP-Seq(GSE119782)/Homer | 1e-12 | -2.879e+01 | 0.0000 | 877.0 | 54.24% | 3268.4 | 44.34% | motif file (matrix) | svg |
| 149 | T C A G A G C T G T C A C G T A A C G T A T G C C G T A A C G T A C G T C T G A | PHV(HB)/col-PHV-DAP-Seq(GSE60143)/Homer | 1e-12 | -2.870e+01 | 0.0000 | 557.0 | 34.45% | 1878.0 | 25.48% | motif file (matrix) | svg |
| 150 | A G T C A C G T A C T G A G C T A C G T A C G T G T C A A G T C | Foxo1(Forkhead)/RAW-Foxo1-ChIP-Seq(Fan\_et\_al.)/Homer | 1e-12 | -2.868e+01 | 0.0000 | 1098.0 | 67.90% | 4298.6 | 58.32% | motif file (matrix) | svg |
| 151 | C G A T T G C A T G C A G A T C C G T A A C T G T G A C G A C T C A T G A C T G | Tcf21(bHLH)/ArterySmoothMuscle-Tcf21-ChIP-Seq(GSE61369)/Homer | 1e-12 | -2.863e+01 | 0.0000 | 605.0 | 37.41% | 2080.3 | 28.22% | motif file (matrix) | svg |
| 152 | T C G A T C G A A G T C C G T A C T A G T A G C A C G T A C T G | MyoG(bHLH)/C2C12-MyoG-ChIP-Seq(GSE36024)/Homer | 1e-12 | -2.847e+01 | 0.0000 | 660.0 | 40.82% | 2316.1 | 31.42% | motif file (matrix) | svg |
| 153 | C G A T T C G A A G T C A C G T A C G T T A C G C G T A G C T A G C T A C G A T G C A T A T G C C G T A G T C A A C T G | ANAC071(NAC)/col-ANAC071-DAP-Seq(GSE60143)/Homer | 1e-12 | -2.843e+01 | 0.0000 | 972.0 | 60.11% | 3708.8 | 50.32% | motif file (matrix) | svg |
| 154 | G C A T C G A T G A C T T G C A A C T G A G T C T G A C A C T G G A T C A G T C C G T A G A C T | ERF15(AP2EREBP)/colamp-ERF15-DAP-Seq(GSE60143)/Homer | 1e-12 | -2.812e+01 | 0.0000 | 1208.0 | 74.71% | 4841.5 | 65.68% | motif file (matrix) | svg |
| 155 | A T G C C A T G A G C T C A G T C A T G T C G A A G T C G A C T C G A T C G A T C A G T C A G T | WRKY26(WRKY)/colamp-WRKY26-DAP-Seq(GSE60143)/Homer | 1e-12 | -2.812e+01 | 0.0000 | 672.0 | 41.56% | 2372.2 | 32.18% | motif file (matrix) | svg |
| 156 | G C A T C G A T A T G C A G C T T C G A T A C G G C T A C G T A C A T G T G A C G C A T C G A T A G T C A G C T C G T A | AT3G09735(S1Falike)/col-AT3G09735-DAP-Seq(GSE60143)/Homer | 1e-12 | -2.808e+01 | 0.0000 | 438.0 | 27.09% | 1399.1 | 18.98% | motif file (matrix) | svg |
| 157 | G T A C A C G T A C G T T C A G G A C T G C A T T C A G C G T A C T G A A G T C C G T A G T C A A C T G A C G T G C T A | NTM2(NAC)/col-NTM2-DAP-Seq(GSE60143)/Homer | 1e-12 | -2.803e+01 | 0.0000 | 668.0 | 41.31% | 2356.6 | 31.97% | motif file (matrix) | svg |
| 158 | A T G C G T A C C G T A A G C T G C A T T A C G A G C T A G C T A G T C A G C T | Sox6(HMG)/Myotubes-Sox6-ChIP-Seq(GSE32627)/Homer | 1e-12 | -2.788e+01 | 0.0000 | 1136.0 | 70.25% | 4492.2 | 60.94% | motif file (matrix) | svg |
| 159 | C G T A A C T G C G T A A C G T A T C G C A G T T A G C C G T A T C G A G T A C C T G A T A G C C G T A A C T G C G T A A C G T C G T A C T G A A T C G G C T A | GATA3(Zf),DR8/iTreg-Gata3-ChIP-Seq(GSE20898)/Homer | 1e-12 | -2.787e+01 | 0.0000 | 139.0 | 8.60% | 300.8 | 4.08% | motif file (matrix) | svg |
| 160 | C G A T T C G A A T G C C G A T G C A T T C G A A G C T G C A T G C A T C G A T T C G A A G C T T C G A G T C A C T A G | ANAC004(NAC)/colamp-ANAC004-DAP-Seq(GSE60143)/Homer | 1e-12 | -2.787e+01 | 0.0000 | 388.0 | 24.00% | 1203.3 | 16.32% | motif file (matrix) | svg |
| 161 | G C A T C G A T G C A T C G T A C T A G A G T C G T C A C G T A A T C G A C G T A C G T A C T G G T A C G C A T C G A T | bHLH80(bHLH)/col-bHLH80-DAP-Seq(GSE60143)/Homer | 1e-12 | -2.774e+01 | 0.0000 | 874.0 | 54.05% | 3269.0 | 44.35% | motif file (matrix) | svg |
| 162 | C A T G A C T G C T A G T C G A T C G A T C G A T C G A T C A G T C A G T C A G T G A C T G A C C G T A A C T G T G C A C G A T A C T G | RBPJ:Ebox(?,bHLH)/Panc1-Rbpj1-ChIP-Seq(GSE47459)/Homer | 1e-12 | -2.770e+01 | 0.0000 | 188.0 | 11.63% | 463.2 | 6.28% | motif file (matrix) | svg |
| 163 | A C G T C T A G A G C T A C G T A C G T C T G A A G T C G A C T A G C T C G T A | FOXM1(Forkhead)/MCF7-FOXM1-ChIP-Seq(GSE72977)/Homer | 1e-12 | -2.769e+01 | 0.0000 | 753.0 | 46.57% | 2730.3 | 37.04% | motif file (matrix) | svg |
| 164 | C T A G T C G A C G A T C T A G G C A T C A G T C T A G G A T C C G T A G T C A | CEBP:AP1(bZIP)/ThioMac-CEBPb-ChIP-Seq(GSE21512)/Homer | 1e-12 | -2.766e+01 | 0.0000 | 813.0 | 50.28% | 2996.0 | 40.65% | motif file (matrix) | svg |
| 165 | A C G T T C G A T C G A A G T C G T C A T A C G A T G C A C G T A C T G A G C T | Myf5(bHLH)/GM-Myf5-ChIP-Seq(GSE24852)/Homer | 1e-11 | -2.759e+01 | 0.0000 | 481.0 | 29.75% | 1578.3 | 21.41% | motif file (matrix) | svg |
| 166 | C G A T C T A G A G T C A C G T A C G T T C A G G C T A C G T A G C A T G C A T C G A T A G T C C G T A G T C A A C T G | VND3(NAC)/colamp-VND3-DAP-Seq(GSE60143)/Homer | 1e-11 | -2.753e+01 | 0.0000 | 798.0 | 49.35% | 2931.4 | 39.77% | motif file (matrix) | svg |
| 167 | C G T A T A G C T A G C T G C A A C T G C T A G C G T A C G T A T C A G G A C T | EHF(ETS)/LoVo-EHF-ChIP-Seq(GSE49402)/Homer | 1e-11 | -2.744e+01 | 0.0000 | 832.0 | 51.45% | 3084.5 | 41.85% | motif file (matrix) | svg |
| 168 | G A T C C G T A G A C T C T A G G A T C C T G A G A C T C T G A G A C T C T A G G A T C C T G A G A C T C T G A G A C T | OCT:OCT(POU,Homeobox)/NPC-OCT6-ChIP-Seq(GSE43916)/Homer | 1e-11 | -2.740e+01 | 0.0000 | 94.0 | 5.81% | 166.2 | 2.26% | motif file (matrix) | svg |
| 169 | C G T A C T G A C G T A C T A G T C G A C T A G A C T G C G T A C G T A T A C G A G C T A T C G | SpiB(ETS)/OCILY3-SPIB-ChIP-Seq(GSE56857)/Homer | 1e-11 | -2.732e+01 | 0.0000 | 227.0 | 14.04% | 602.2 | 8.17% | motif file (matrix) | svg |
| 170 | A G T C C T G A A T C G A G C T A G C T G A C T A G T C G C T A A C G T C G A T G C A T C G A T A T C G C G T A T A G C G C A T A T G C C G T A | bZIP:IRF(bZIP,IRF)/Th17-BatF-ChIP-Seq(GSE39756)/Homer | 1e-11 | -2.675e+01 | 0.0000 | 318.0 | 19.67% | 944.0 | 12.81% | motif file (matrix) | svg |
| 171 | T G C A C T G A A T G C G T C A A C G T A T G C A C G T A C T G A C T G T G C A | ZBTB18(Zf)/HEK293-ZBTB18.GFP-ChIP-Seq(GSE58341)/Homer | 1e-11 | -2.651e+01 | 0.0000 | 412.0 | 25.48% | 1312.9 | 17.81% | motif file (matrix) | svg |
| 172 | G T C A T C G A C T A G C T A G A G T C G T C A C G A T C T A G G A C T G A T C G A T C T C A G C T A G C T G A A G T C G C T A C A G T T C A G G A T C G A T C | p63(p53)/Keratinocyte-p63-ChIP-Seq(GSE17611)/Homer | 1e-11 | -2.642e+01 | 0.0000 | 410.0 | 25.36% | 1305.5 | 17.71% | motif file (matrix) | svg |
| 173 | T C A G T G A C G T A C C G T A A C G T T G A C A C G T T C A G A G C T G A C T | NeuroD1(bHLH)/Islet-NeuroD1-ChIP-Seq(GSE30298)/Homer | 1e-11 | -2.641e+01 | 0.0000 | 588.0 | 36.36% | 2036.0 | 27.62% | motif file (matrix) | svg |
| 174 | G C A T G C A T G A T C G C A T T C G A A C T G C G T A C G T A A T C G T A G C C G A T A C G T A G T C A G C T C G T A | HSF6(HSF)/col-HSF6-DAP-Seq(GSE60143)/Homer | 1e-11 | -2.625e+01 | 0.0000 | 257.0 | 15.89% | 719.3 | 9.76% | motif file (matrix) | svg |
| 175 | G A T C G C T A G T A C A G T C G C T A T G C A G T A C G A T C C G T A G A C T | MYB83(MYB)/colamp-MYB83-DAP-Seq(GSE60143)/Homer | 1e-11 | -2.607e+01 | 0.0000 | 1214.0 | 75.08% | 4899.2 | 66.46% | motif file (matrix) | svg |
| 176 | C G T A G C T A C G A T C T A G A C G T G T C A C G T A C G T A A G T C C G T A T G C A T A C G | FoxL2(Forkhead)/Ovary-FoxL2-ChIP-Seq(GSE60858)/Homer | 1e-11 | -2.593e+01 | 0.0000 | 755.0 | 46.69% | 2763.1 | 37.48% | motif file (matrix) | svg |
| 177 | C T G A C T G A C T G A A T G C G A T C C A T G A C T G G A C T G A C T G C A T C G T A C G T A A G T C G T A C C T G A A T C G G C A T G A C T G A C T A G C T | GRHL2(CP2)/HBE-GRHL2-ChIP-Seq(GSE46194)/Homer | 1e-11 | -2.581e+01 | 0.0000 | 462.0 | 28.57% | 1521.3 | 20.64% | motif file (matrix) | svg |
| 178 | C G A T T A C G T G C A G T A C G A T C G A C T A G C T A C G T A T C G G T A C G A T C G T A C G A T C G T C A | PPARE(NR),DR1/3T3L1-Pparg-ChIP-Seq(GSE13511)/Homer | 1e-11 | -2.580e+01 | 0.0000 | 580.0 | 35.87% | 2010.7 | 27.28% | motif file (matrix) | svg |
| 179 | G C T A C G T A C G A T C A G T A C T G C G A T G T A C A C T G A T C G G A C T C A T G C T A G G C A T C A G T C A T G | DEAR5(AP2EREBP)/col-DEAR5-DAP-Seq(GSE60143)/Homer | 1e-11 | -2.574e+01 | 0.0000 | 446.0 | 27.58% | 1457.9 | 19.78% | motif file (matrix) | svg |
| 180 | G C T A T C G A C G T A C T A G A G C T G T C A G T C A C G T A A G T C C G T A | FOXA1(Forkhead)/LNCAP-FOXA1-ChIP-Seq(GSE27824)/Homer | 1e-10 | -2.515e+01 | 0.0000 | 818.0 | 50.59% | 3054.8 | 41.44% | motif file (matrix) | svg |
| 181 | A G C T A G C T C A T G C T G A G T A C A G T C A G C T A G C T C A G T C T A G | RARa(NR)/K562-RARa-ChIP-Seq(Encode)/Homer | 1e-10 | -2.486e+01 | 0.0000 | 1308.0 | 80.89% | 5388.2 | 73.10% | motif file (matrix) | svg |
| 182 | A T G C G A T C C G T A A G C T C A G T A T C G G C A T A G C T G A C T A C T G | Sox17(HMG)/Endoderm-Sox17-ChIP-Seq(GSE61475)/Homer | 1e-10 | -2.484e+01 | 0.0000 | 744.0 | 46.01% | 2730.3 | 37.04% | motif file (matrix) | svg |
| 183 | G C T A T C G A C G T A C T G A A C T G A C G T A G T C C G T A C G T A A G T C C T A G T G C A | WRKY42(WRKY)/colamp-WRKY42-DAP-Seq(GSE60143)/Homer | 1e-10 | -2.466e+01 | 0.0000 | 613.0 | 37.91% | 2165.7 | 29.38% | motif file (matrix) | svg |
| 184 | A G T C G A C T G A T C C G T A G T A C A G T C G C T A C G T A G T A C A G T C G T A C G T A C | MYB63(MYB)/col-MYB63-DAP-Seq(GSE60143)/Homer | 1e-10 | -2.454e+01 | 0.0000 | 622.0 | 38.47% | 2205.9 | 29.93% | motif file (matrix) | svg |
| 185 | T C G A G A C T T C A G T G C A G T A C G T A C A G C T G T A C C A T G T C G A C A T G C A T G A C G T A G T C C T G A | FXR(NR),ER2/Liver-FXR-ChIP-Seq(GSE133700)/Homer | 1e-10 | -2.434e+01 | 0.0000 | 488.0 | 30.18% | 1646.0 | 22.33% | motif file (matrix) | svg |
| 186 | G C T A T C G A C G T A C T A G A G C T G T C A G T C A C G T A A G T C C G T A | FOXA1(Forkhead)/MCF7-FOXA1-ChIP-Seq(GSE26831)/Homer | 1e-10 | -2.433e+01 | 0.0000 | 684.0 | 42.30% | 2475.1 | 33.58% | motif file (matrix) | svg |
| 187 | C G A T T G C A A G T C A C G T A C G T T A C G C G A T C G A T T A C G G C T A G C T A A T G C C G T A G T C A C A T G | ANAC016(NAC)/col-ANAC016-DAP-Seq(GSE60143)/Homer | 1e-10 | -2.428e+01 | 0.0000 | 896.0 | 55.41% | 3419.8 | 46.39% | motif file (matrix) | svg |
| 188 | A C G T T G C A A G C T G A T C C T A G C T G A A G C T G T C A T C G A C G T A | CUX1(Homeobox)/K562-CUX1-ChIP-Seq(GSE92882)/Homer | 1e-10 | -2.418e+01 | 0.0000 | 1101.0 | 68.09% | 4377.0 | 59.38% | motif file (matrix) | svg |
| 189 | C G T A G A T C C T A G A C G T G T A C C T G A A G C T G A T C G C T A G A C T | TGA2(bZIP)/colamp-TGA2-DAP-Seq(GSE60143)/Homer | 1e-10 | -2.417e+01 | 0.0000 | 1019.0 | 63.02% | 3989.1 | 54.12% | motif file (matrix) | svg |
| 190 | A C T G C G T A A C T G A T G C T G A C G A T C A T C G T G C A A C T G A G T C | ZNF519(Zf)/HEK293-ZNF519.GFP-ChIP-Seq(GSE58341)/Homer | 1e-10 | -2.412e+01 | 0.0000 | 197.0 | 12.18% | 519.3 | 7.05% | motif file (matrix) | svg |
| 191 | C G A T T C G A G T A C A C G T A C G T T C A G G C A T G C T A T G C A G C A T C G T A A G T C C G T A T G A C C A T G | ANAC092(NAC)/colamp-ANAC092-DAP-Seq(GSE60143)/Homer | 1e-10 | -2.412e+01 | 0.0000 | 672.0 | 41.56% | 2426.8 | 32.92% | motif file (matrix) | svg |
| 192 | C A T G A G T C G T A C A C T G A T G C A G T C C A T G G A T C G A T C C T G A | ERF5(AP2EREBP)/colamp-ERF5-DAP-Seq(GSE60143)/Homer | 1e-10 | -2.394e+01 | 0.0000 | 523.0 | 32.34% | 1795.4 | 24.36% | motif file (matrix) | svg |
| 193 | G C T A C T G A T C G A A G T C A G T C C T G A A G T C G T C A C T G A T G C A | RUNX1(Runt)/Jurkat-RUNX1-ChIP-Seq(GSE29180)/Homer | 1e-10 | -2.382e+01 | 0.0000 | 827.0 | 51.14% | 3114.9 | 42.26% | motif file (matrix) | svg |
| 194 | T A G C T A G C G A C T C T A G A G C T A G T C G T C A T G C A A C G T A T G C G C T A T G C A | Pbx3(Homeobox)/GM12878-PBX3-ChIP-Seq(GSE32465)/Homer | 1e-10 | -2.358e+01 | 0.0000 | 264.0 | 16.33% | 769.0 | 10.43% | motif file (matrix) | svg |
| 195 | T C A G C T G A C G T A C G T A T A C G G C A T C T A G C T G A C G T A C G T A T A C G G A C T | IRF1(IRF)/PBMC-IRF1-ChIP-Seq(GSE43036)/Homer | 1e-10 | -2.352e+01 | 0.0000 | 118.0 | 7.30% | 256.6 | 3.48% | motif file (matrix) | svg |
| 196 | G T A C A C T G A T G C A G T C C T A G G A T C G T A C C T G A G A C T G C A T C G A T G A C T | RAP212(AP2EREBP)/col-RAP212-DAP-Seq(GSE60143)/Homer | 1e-10 | -2.349e+01 | 0.0000 | 973.0 | 60.17% | 3785.6 | 51.36% | motif file (matrix) | svg |
| 197 | A C G T T G A C A G T C A G C T A G T C A G C T A C T G G A C T A G C T G A C T | REF6(Zf)/Arabidopsis-REF6-ChIP-Seq(GSE106942)/Homer | 1e-10 | -2.330e+01 | 0.0000 | 438.0 | 27.09% | 1453.5 | 19.72% | motif file (matrix) | svg |
| 198 | A G C T T G A C C G A T C G A T C T A G A C G T C A G T C A G T G C T A A G T C | FOXK1(Forkhead)/HEK293-FOXK1-ChIP-Seq(GSE51673)/Homer | 1e-10 | -2.308e+01 | 0.0000 | 918.0 | 56.77% | 3538.6 | 48.01% | motif file (matrix) | svg |
| 199 | C A T G T G C A G A C T C A T G C G T A A G T C T C A G G C A T T G A C C G T A | bZIP50(bZIP)/colamp-bZIP50-DAP-Seq(GSE60143)/Homer | 1e-9 | -2.302e+01 | 0.0000 | 1200.0 | 74.21% | 4874.8 | 66.13% | motif file (matrix) | svg |
| 200 | C G T A C G T A C G T A C G T A C G T A A C T G A C T G A G T C | dof42(C2C2dof)/col-dof42-DAP-Seq(GSE60143)/Homer | 1e-9 | -2.284e+01 | 0.0000 | 659.0 | 40.75% | 2388.6 | 32.40% | motif file (matrix) | svg |
| 201 | T A C G T G C A A G T C C G T A A C G T T G A C A C G T A C T G A C T G G C A T | TCF4(bHLH)/SHSY5Y-TCF4-ChIP-Seq(GSE96915)/Homer | 1e-9 | -2.282e+01 | 0.0000 | 945.0 | 58.44% | 3666.6 | 49.74% | motif file (matrix) | svg |
| 202 | T A G C C A T G G A C T G A C T T C A G G T C A G A T C G A C T G C A T G C T A | WRKY15(WRKY)/col-WRKY15-DAP-Seq(GSE60143)/Homer | 1e-9 | -2.278e+01 | 0.0000 | 976.0 | 60.36% | 3810.3 | 51.69% | motif file (matrix) | svg |
| 203 | T G A C A G T C C G T A A C T G G T A C A C G T A C T G A C G T G A C T G A T C | Twist2(bHLH)/Myoblast-Twist2.Ty1-ChIP-Seq(GSE127998)/Homer | 1e-9 | -2.274e+01 | 0.0000 | 1004.0 | 62.09% | 3941.4 | 53.47% | motif file (matrix) | svg |
| 204 | C G A T C G A T G C A T G A C T A C G T C G T A C G T A A C T G T A G C C G T A C G T A C G T A | AT5G60130(ABI3VP1)/col-AT5G60130-DAP-Seq(GSE60143)/Homer | 1e-9 | -2.268e+01 | 0.0000 | 948.0 | 58.63% | 3682.3 | 49.96% | motif file (matrix) | svg |
| 205 | A G T C A C G T A C G T T C A G G C T A C G T A G C T A G C A T C G A T A G T C C G T A G T C A A C T G G A C T G C T A | SMB(NAC)/colamp-SMB-DAP-Seq(GSE60143)/Homer | 1e-9 | -2.258e+01 | 0.0000 | 982.0 | 60.73% | 3841.2 | 52.11% | motif file (matrix) | svg |
| 206 | G A C T G C T A T G C A A G T C A C G T A C G T A C G T C G A T A C G T T A C G | At3g45610(C2C2dof)/col-At3g45610-DAP-Seq(GSE60143)/Homer | 1e-9 | -2.245e+01 | 0.0000 | 985.0 | 60.92% | 3857.6 | 52.33% | motif file (matrix) | svg |
| 207 | A T G C C A T G G C A T G A C T C T A G T C G A G T A C A G C T C G T A G C T A | WRKY75(WRKY)/col-WRKY75-DAP-Seq(GSE60143)/Homer | 1e-9 | -2.204e+01 | 0.0000 | 946.0 | 58.50% | 3683.3 | 49.97% | motif file (matrix) | svg |
| 208 | A G C T G A T C C T G A A G T C A G T C A C T G C G T A A G T C C T G A G T C A G C A T C G A T G C T A G C A T C G T A | At2g44940(AP2EREBP)/colamp-At2g44940-DAP-Seq(GSE60143)/Homer | 1e-9 | -2.188e+01 | 0.0000 | 621.0 | 38.40% | 2238.7 | 30.37% | motif file (matrix) | svg |
| 209 | A G T C G A C T A C T G G A T C G T A C C G T A T G A C A G T C C G A T A G C T A C G T A C G T C T A G G A C T C T G A | ZNF7(Zf)/HepG2-ZNF7.Flag-ChIP-Seq(Encode)/Homer | 1e-9 | -2.170e+01 | 0.0000 | 520.0 | 32.16% | 1812.1 | 24.58% | motif file (matrix) | svg |
| 210 | T G A C G C T A T G A C C G T A T C A G G A T C C G T A C A T G C A T G C T A G C T A G C T A G | Unknown-ESC-element(?)/mES-Nanog-ChIP-Seq(GSE11724)/Homer | 1e-9 | -2.151e+01 | 0.0000 | 346.0 | 21.40% | 1105.2 | 14.99% | motif file (matrix) | svg |
| 211 | C A G T G C T A G C A T T A C G C T G A C A G T T A G C C T G A | GATA15(C2C2gata)/col-GATA15-DAP-Seq(GSE60143)/Homer | 1e-9 | -2.144e+01 | 0.0000 | 1290.0 | 79.78% | 5345.3 | 72.52% | motif file (matrix) | svg |
| 212 | C G T A G A C T C A T G C T A G A G T C A C T G A C T G G T A C C A T G T A C G | ERF3(AP2EREBP)/colamp-ERF3-DAP-Seq(GSE60143)/Homer | 1e-9 | -2.143e+01 | 0.0000 | 847.0 | 52.38% | 3241.3 | 43.97% | motif file (matrix) | svg |
| 213 | C A T G A G T C G T C A C G T A A T G C A C G T A C G T A C T G | bHLH130(bHLH)/col-bHLH130-DAP-Seq(GSE60143)/Homer | 1e-9 | -2.142e+01 | 0.0000 | 751.0 | 46.44% | 2812.6 | 38.16% | motif file (matrix) | svg |
| 214 | C A T G G A T C C T G A G T A C C T A G C T G A G C T A G C A T G A T C G A T C A G T C C T A G C G T A C A T G C T A G | AIL7(AP2EREBP)/colamp-AIL7-DAP-Seq(GSE60143)/Homer | 1e-9 | -2.138e+01 | 0.0000 | 705.0 | 43.60% | 2611.0 | 35.42% | motif file (matrix) | svg |
| 215 | C T A G C T G A C T A G C T G A C T A G C T G A C T A G C T G A C T A G C T G A | SeqBias: GA-repeat | 1e-9 | -2.130e+01 | 0.0000 | 1462.0 | 90.41% | 6248.6 | 84.77% | motif file (matrix) | svg |
| 216 | A G C T G C T A T G C A A G T C A C G T A C G T A C G T C G A T A G C T T C A G | dof24(C2C2dof)/col-dof24-DAP-Seq(GSE60143)/Homer | 1e-9 | -2.120e+01 | 0.0000 | 1174.0 | 72.60% | 4775.4 | 64.78% | motif file (matrix) | svg |
| 217 | A C T G C T A G A G T C A C T G A C T G A G T C A C G T C T A G | AT5G23930(mTERF)/col-AT5G23930-DAP-Seq(GSE60143)/Homer | 1e-9 | -2.100e+01 | 0.0000 | 1032.0 | 63.82% | 4100.6 | 55.63% | motif file (matrix) | svg |
| 218 | C T A G T C G A A C G T A C G T C A T G A G T C C T G A C G A T A G T C C G T A | AARE(HLH)/mES-cMyc-ChIP-Seq/Homer | 1e-9 | -2.083e+01 | 0.0000 | 196.0 | 12.12% | 541.7 | 7.35% | motif file (matrix) | svg |
| 219 | A G C T G A T C G A T C C G T A G T A C A G T C C G A T C T G A G T A C G A T C C G T A G A C T | ATY19(MYB)/col-ATY19-DAP-Seq(GSE60143)/Homer | 1e-9 | -2.078e+01 | 0.0000 | 812.0 | 50.22% | 3094.8 | 41.98% | motif file (matrix) | svg |
| 220 | C T A G G C T A A G T C A C T G A C G T G A C T G A C T A T G C T C G A C A G T G A T C C G A T G A C T G A T C G A T C | RKD2(RWPRK)/colamp-RKD2-DAP-Seq(GSE60143)/Homer | 1e-8 | -2.064e+01 | 0.0000 | 719.0 | 44.47% | 2683.9 | 36.41% | motif file (matrix) | svg |
| 221 | T C G A C G T A A G T C C G T A C T A G A G T C C G A T A C T G G A C T A G C T A C T G G A C T | HLH-1(bHLH)/cElegans-Embryo-HLH1-ChIP-Seq(modEncode)/Homer | 1e-8 | -2.058e+01 | 0.0000 | 660.0 | 40.82% | 2426.6 | 32.92% | motif file (matrix) | svg |
| 222 | C T A G C A T G C A T G T A C G A G T C G C A T A G C T C T A G A C G T A G T C G A C T A C T G A C T G A C T G T C G A | Zfp809(Zf)/ES-Zfp809-ChIP-Seq(GSE70799)/Homer | 1e-8 | -2.052e+01 | 0.0000 | 149.0 | 9.21% | 377.3 | 5.12% | motif file (matrix) | svg |
| 223 | C G A T C T G A A G T C A C G T A C G T T C A G C G T A C G T A C G T A G C A T C G A T A G T C C G T A G T C A A C T G | VND4(NAC)/colamp-VND4-DAP-Seq(GSE60143)/Homer | 1e-8 | -2.039e+01 | 0.0000 | 868.0 | 53.68% | 3353.3 | 45.49% | motif file (matrix) | svg |
| 224 | G C T A C G T A C G T A G C A T C A T G C T A G G A T C A C T G T C A G G A T C A C T G T C A G | ERF4(AP2EREBP)/colamp-ERF4-DAP-Seq(GSE60143)/Homer | 1e-8 | -2.027e+01 | 0.0000 | 1153.0 | 71.30% | 4688.2 | 63.60% | motif file (matrix) | svg |
| 225 | G T A C A C G T A C G T T C A G C G T A C G T A C G A T G C A T G C A T A G T C C G T A G T C A C A T G G A C T G C T A | VND1(NAC)/col-VND1-DAP-Seq(GSE60143)/Homer | 1e-8 | -2.020e+01 | 0.0000 | 886.0 | 54.79% | 3438.2 | 46.64% | motif file (matrix) | svg |
| 226 | C G A T C T G A A G T C A C G T A C G T T C A G G C A T C G A T G C T A G C T A C G T A A G T C C G T A G T C A A C T G | CUC1(NAC)/col-CUC1-DAP-Seq(GSE60143)/Homer | 1e-8 | -2.016e+01 | 0.0000 | 635.0 | 39.27% | 2324.0 | 31.53% | motif file (matrix) | svg |
| 227 | C T A G A G T C A G T C A C T G C G T A A G T C C T G A G A C T | DDF1(AP2EREBP)/col-DDF1-DAP-Seq(GSE60143)/Homer | 1e-8 | -2.007e+01 | 0.0000 | 934.0 | 57.76% | 3660.9 | 49.67% | motif file (matrix) | svg |
| 228 | C A G T A T C G C T G A A G T C T C A G C A G T T A G C C T G A A T G C T A C G | FEA4(bZIP)/Corn-FEA4-ChIP-Seq(GSE61954)/Homer | 1e-8 | -2.002e+01 | 0.0000 | 1064.0 | 65.80% | 4267.5 | 57.89% | motif file (matrix) | svg |
| 229 | C T A G A G T C T A C G T A C G T G A C C G T A A C T G T A G C G C A T C A T G A T G C A G C T | Ascl1(bHLH)/NeuralTubes-Ascl1-ChIP-Seq(GSE55840)/Homer | 1e-8 | -1.998e+01 | 0.0000 | 859.0 | 53.12% | 3319.0 | 45.03% | motif file (matrix) | svg |
| 230 | C G A T C G T A A G T C A C G T A C G T T C A G C G T A C G T A G C A T G C A T G C A T A G T C C G T A G T C A A C T G | VND2(NAC)/col-VND2-DAP-Seq(GSE60143)/Homer | 1e-8 | -1.998e+01 | 0.0000 | 969.0 | 59.93% | 3823.6 | 51.87% | motif file (matrix) | svg |
| 231 | A C T G A C T G A G T C A C T G A C T G A G T C A C G T C T A G | ERF1(AP2EREBP)/colamp-ERF1-DAP-Seq(GSE60143)/Homer | 1e-8 | -1.997e+01 | 0.0000 | 589.0 | 36.43% | 2129.3 | 28.89% | motif file (matrix) | svg |
| 232 | G T C A T G C A G C T A A G T C C G T A A C T G T G A C G C A T T C A G C A G T | Ap4(bHLH)/AML-Tfap4-ChIP-Seq(GSE45738)/Homer | 1e-8 | -1.982e+01 | 0.0000 | 691.0 | 42.73% | 2573.1 | 34.91% | motif file (matrix) | svg |
| 233 | G C A T G C A T G C A T A T G C A G C T T C G A T A C G G C T A C G T A C A T G G T A C G C A T G C A T A G T C A G C T | HSFA6B(HSF)/colamp-HSFA6B-DAP-Seq(GSE60143)/Homer | 1e-8 | -1.974e+01 | 0.0000 | 475.0 | 29.38% | 1651.4 | 22.40% | motif file (matrix) | svg |
| 234 | C G A T C T A G A C T G A G C T C T G A A C T G A C G T A C G T C T A G C T A G | MYB96(MYB)/colamp-MYB96-DAP-Seq(GSE60143)/Homer | 1e-8 | -1.968e+01 | 0.0000 | 859.0 | 53.12% | 3324.5 | 45.10% | motif file (matrix) | svg |
| 235 | G A T C G C T A A G T C A C G T A C T G C G T A A G T C C G T A G C T A C G A T C A G T G C A T G C T A C G T A G C A T | GRF9(GRF)/colamp-GRF9-DAP-Seq(GSE60143)/Homer | 1e-8 | -1.968e+01 | 0.0000 | 824.0 | 50.96% | 3166.5 | 42.96% | motif file (matrix) | svg |
| 236 | G T A C C A T G A G C T A G C T T C A G T G C A T G A C A G C T C G T A C G T A | WRKY33(WRKY)/col-WRKY33-DAP-Seq(GSE60143)/Homer | 1e-8 | -1.965e+01 | 0.0000 | 905.0 | 55.97% | 3534.2 | 47.95% | motif file (matrix) | svg |
| 237 | G A C T A G T C C G A T A C T G C T G A T G A C G T A C C G T A A T C G G C A T C T G A C T A G | Bcl11a(Zf)/HSPC-BCL11A-ChIP-Seq(GSE104676)/Homer | 1e-8 | -1.946e+01 | 0.0000 | 549.0 | 33.95% | 1966.4 | 26.68% | motif file (matrix) | svg |
| 238 | A T G C G T A C A C T G A G T C A G T C A C T G A G T C G T A C | ERF73(AP2EREBP)/col-ERF73-DAP-Seq(GSE60143)/Homer | 1e-8 | -1.945e+01 | 0.0000 | 629.0 | 38.90% | 2309.8 | 31.34% | motif file (matrix) | svg |
| 239 | C T A G G C A T A C T G C T A G A G T C A C T G A C T G A G T C A C T G T C A G | ERF10(AP2EREBP)/col-ERF10-DAP-Seq(GSE60143)/Homer | 1e-8 | -1.938e+01 | 0.0000 | 810.0 | 50.09% | 3108.4 | 42.17% | motif file (matrix) | svg |
| 240 | A G T C A G T C C T G A A G T C A G T C A C T G C G T A A G T C C T G A T C G A G C A T G A T C C G A T C G A T A C T G | AT3G60490(AP2EREBP)/colamp-AT3G60490-DAP-Seq(GSE60143)/Homer | 1e-8 | -1.935e+01 | 0.0000 | 710.0 | 43.91% | 2664.8 | 36.15% | motif file (matrix) | svg |
| 241 | G C T A C G T A C G T A G A C T C A T G C T A G G A T C A C T G T C A G G A T C A C T G T A C G | ERF9(AP2EREBP)/colamp-ERF9-DAP-Seq(GSE60143)/Homer | 1e-8 | -1.930e+01 | 0.0000 | 410.0 | 25.36% | 1389.1 | 18.85% | motif file (matrix) | svg |
| 242 | T G A C C G A T C T G A C T A G C T A G A C G T A T G C T G C A T C G A C T G A C T A G C A T G A C G T A G T C C G T A | PPARa(NR),DR1/Liver-Ppara-ChIP-Seq(GSE47954)/Homer | 1e-8 | -1.929e+01 | 0.0000 | 657.0 | 40.63% | 2433.9 | 33.02% | motif file (matrix) | svg |
| 243 | G A C T A G T C C T G A A G T C A G T C A C T G C T G A A G T C G C T A G C A T G T A C C G A T G C A T G A C T C G A T | CBF2(AP2EREBP)/colamp-CBF2-DAP-Seq(GSE60143)/Homer | 1e-8 | -1.923e+01 | 0.0000 | 955.0 | 59.06% | 3771.7 | 51.17% | motif file (matrix) | svg |
| 244 | C G A T C T G A G T A C A C T G A C G T T C A G G C A T C G T A C G T A G C A T C G T A A G T C C G T A G T A C C A T G | CUC3(NAC)/col-CUC3-DAP-Seq(GSE60143)/Homer | 1e-8 | -1.918e+01 | 0.0000 | 589.0 | 36.43% | 2141.1 | 29.05% | motif file (matrix) | svg |
| 245 | T A G C C A T G A G C T A C G T A C T G C G T A A G T C G A C T G C A T C T G A | AT3G42860(zfGRF)/col-AT3G42860-DAP-Seq(GSE60143)/Homer | 1e-8 | -1.911e+01 | 0.0000 | 470.0 | 29.07% | 1639.0 | 22.24% | motif file (matrix) | svg |
| 246 | G C A T G C A T G T A C G A C T T C G A A C T G G C T A C G T A A T C G T G A C G C A T G A C T A G T C A G C T C T G A | AGL95(ND)/col-AGL95-DAP-Seq(GSE60143)/Homer | 1e-8 | -1.898e+01 | 0.0000 | 124.0 | 7.67% | 302.9 | 4.11% | motif file (matrix) | svg |
| 247 | G A T C C T G A A G T C A G C T A C G T A C G T A C G T A C G T | At1g64620(C2C2dof)/colamp-At1g64620-DAP-Seq(GSE60143)/Homer | 1e-8 | -1.868e+01 | 0.0000 | 992.0 | 61.35% | 3952.7 | 53.62% | motif file (matrix) | svg |
| 248 | C G A T A G T C C A T G G A C T A C G T C T A G C G T A G A T C G A C T C G A T G C A T G A C T | WRKY43(WRKY)/colamp-WRKY43-DAP-Seq(GSE60143)/Homer | 1e-8 | -1.865e+01 | 0.0000 | 479.0 | 29.62% | 1683.3 | 22.84% | motif file (matrix) | svg |
| 249 | T C G A G C A T A C G T C T A G G T A C T C G A G C A T T G A C T C G A A C G T | Chop(bZIP)/MEF-Chop-ChIP-Seq(GSE35681)/Homer | 1e-8 | -1.860e+01 | 0.0000 | 344.0 | 21.27% | 1131.9 | 15.36% | motif file (matrix) | svg |
| 250 | G A C T C G A T T C A G G A T C G A C T A G C T A G C T A G T C G A T C C G T A C T A G C T A G T C G A T C G A C T G A | Bcl6(Zf)/Liver-Bcl6-ChIP-Seq(GSE31578)/Homer | 1e-8 | -1.859e+01 | 0.0000 | 721.0 | 44.59% | 2725.3 | 36.97% | motif file (matrix) | svg |
| 251 | G T C A C G T A A C G T A T C G C G T A A C G T A C G T C T A G | ATHB7(Homeobox)/col-ATHB7-DAP-Seq(GSE60143)/Homer | 1e-8 | -1.854e+01 | 0.0000 | 858.0 | 53.06% | 3339.8 | 45.31% | motif file (matrix) | svg |
| 252 | C G T A G A C T C A T G C T A G A G T C A C T G C T A G A G T C C A T G C T A G | ERF7(AP2EREBP)/col-ERF7-DAP-Seq(GSE60143)/Homer | 1e-8 | -1.847e+01 | 0.0000 | 1198.0 | 74.09% | 4936.9 | 66.98% | motif file (matrix) | svg |
| 253 | C T A G A C T G A G T C A C T G A C T G A G C T A C T G T C A G | AT3G57600(AP2EREBP)/col-AT3G57600-DAP-Seq(GSE60143)/Homer | 1e-7 | -1.830e+01 | 0.0000 | 694.0 | 42.92% | 2611.5 | 35.43% | motif file (matrix) | svg |
| 254 | A C T G A C T G A G T C A C T G A C T G A G T C A C G T T C A G | ERF2(AP2EREBP)/colamp-ERF2-DAP-Seq(GSE60143)/Homer | 1e-7 | -1.826e+01 | 0.0000 | 626.0 | 38.71% | 2315.9 | 31.42% | motif file (matrix) | svg |
| 255 | G A C T C A G T G A T C G A T C A C G T G A T C C T G A T A C G C G T A G T C A | STAT6(Stat)/Macrophage-Stat6-ChIP-Seq(GSE38377)/Homer | 1e-7 | -1.822e+01 | 0.0000 | 454.0 | 28.08% | 1585.6 | 21.51% | motif file (matrix) | svg |
| 256 | G C A T A C G T A C T G A C G T A G T C A C T G A T C G G T C A C G A T C G T A | ARF2(ARF)/col-ARF2-DAP-Seq(GSE60143)/Homer | 1e-7 | -1.819e+01 | 0.0000 | 1371.0 | 84.79% | 5806.3 | 78.77% | motif file (matrix) | svg |
| 257 | G C T A C G T A C G T A G C A T C A T G C T A G A G T C A C T G A C T G A G T C A C T G T C A G | ABR1(AP2EREBP)/colamp-ABR1-DAP-Seq(GSE60143)/Homer | 1e-7 | -1.803e+01 | 0.0000 | 1134.0 | 70.13% | 4634.7 | 62.88% | motif file (matrix) | svg |
| 258 | T C G A A G C T A C G T A C G T A G T C A G T C A C G T A T C G G A C T A T C G | EWS:ERG-fusion(ETS)/CADO\_ES1-EWS:ERG-ChIP-Seq(SRA014231)/Homer | 1e-7 | -1.799e+01 | 0.0000 | 476.0 | 29.44% | 1680.8 | 22.80% | motif file (matrix) | svg |
| 259 | G A C T C T A G G A T C C A G T A C T G C T G A A T G C G C A T A T G C C T G A | MafA(bZIP)/Islet-MafA-ChIP-Seq(GSE30298)/Homer | 1e-7 | -1.794e+01 | 0.0000 | 668.0 | 41.31% | 2503.4 | 33.96% | motif file (matrix) | svg |
| 260 | A T G C A G T C A C T G A T G C A G T C A C T G A G T C G T A C | SHN3(AP2EREBP)/col-SHN3-DAP-Seq(GSE60143)/Homer | 1e-7 | -1.792e+01 | 0.0000 | 291.0 | 18.00% | 930.0 | 12.62% | motif file (matrix) | svg |
| 261 | C G A T G C T A G C T A G C A T G C T A C G T A A G T C A C G T A C G T A C G T C G A T A G C T | At5g62940(C2C2dof)/col-At5g62940-DAP-Seq(GSE60143)/Homer | 1e-7 | -1.789e+01 | 0.0000 | 1380.0 | 85.34% | 5857.9 | 79.47% | motif file (matrix) | svg |
| 262 | G C A T C G T A G C A T C G T A T C G A C G T A C T G A A C T G C G T A C G T A C G T A A C G T A C T G G T C A G C A T | AT2G31460(REMB3)/col-AT2G31460-DAP-Seq(GSE60143)/Homer | 1e-7 | -1.788e+01 | 0.0000 | 322.0 | 19.91% | 1053.0 | 14.28% | motif file (matrix) | svg |
| 263 | A G T C T A G C G A C T A C G T C T A G A C G T A C G T A C G T C T G A A G T C G C T A G A C T C G T A C T A G A C T G | Foxa3(Forkhead)/Liver-Foxa3-ChIP-Seq(GSE77670)/Homer | 1e-7 | -1.773e+01 | 0.0000 | 353.0 | 21.83% | 1178.4 | 15.99% | motif file (matrix) | svg |
| 264 | C G T A C T A G T C A G T C A G A G T C A T G C A G T C G C A T A G C T A C G T A T C G C G A T | Sox9(HMG)/Limb-SOX9-ChIP-Seq(GSE73225)/Homer | 1e-7 | -1.760e+01 | 0.0000 | 692.0 | 42.80% | 2614.8 | 35.47% | motif file (matrix) | svg |
| 265 | G A C T C T G A A G T C A G T C A C T G C G T A A G T C C T G A | bHLH10(bHLH)/colamp-bHLH10-DAP-Seq(GSE60143)/Homer | 1e-7 | -1.754e+01 | 0.0000 | 687.0 | 42.49% | 2593.1 | 35.18% | motif file (matrix) | svg |
| 266 | T A C G T C G A C G T A C G T A C G T A C T G A A C T G A C G T C G T A T C G A | AT2G28810(C2C2dof)/colamp-AT2G28810-DAP-Seq(GSE60143)/Homer | 1e-7 | -1.754e+01 | 0.0000 | 1212.0 | 74.95% | 5020.4 | 68.11% | motif file (matrix) | svg |
| 267 | C G T A C G T A G C A T A C T G C G T A A G C T C T G A C G T A T A C G C T G A | ELT-3(Gata)/cElegans-L1-ELT3-ChIP-Seq(modEncode)/Homer | 1e-7 | -1.745e+01 | 0.0000 | 571.0 | 35.31% | 2091.1 | 28.37% | motif file (matrix) | svg |
| 268 | T C G A G T A C T C G A T C G A C A T G A T G C A C G T A C T G A C T G A G T C C G T A C T A G A G T C A T C G A G T C | Unknown3/Drosophila-Promoters/Homer | 1e-7 | -1.738e+01 | 0.0000 | 120.0 | 7.42% | 299.9 | 4.07% | motif file (matrix) | svg |
| 269 | G A C T G T A C C T G A G A T C A G T C C T A G G C T A G T A C C T G A G C T A G C A T C G A T G C A T G A C T C G T A | AT3G16280(AP2EREBP)/colamp-AT3G16280-DAP-Seq(GSE60143)/Homer | 1e-7 | -1.727e+01 | 0.0000 | 861.0 | 53.25% | 3375.1 | 45.79% | motif file (matrix) | svg |
| 270 | C T G A C G A T C T A G C G T A A G C T C G A T C A G T C T G A G A C T C T A G C T A G A T G C | PBX2(Homeobox)/K562-PBX2-ChIP-Seq(Encode)/Homer | 1e-7 | -1.716e+01 | 0.0000 | 904.0 | 55.91% | 3573.6 | 48.48% | motif file (matrix) | svg |
| 271 | T G C A G C A T C G A T C G T A C A G T A C T G G T A C C G T A C T G A A G C T G T C A A C T G C T A G G T C A C G A T A C T G G T A C T G C A C G T A A G C T | CEBP:CEBP(bZIP)/MEF-Chop-ChIP-Seq(GSE35681)/Homer | 1e-7 | -1.711e+01 | 0.0000 | 184.0 | 11.38% | 529.6 | 7.18% | motif file (matrix) | svg |
| 272 | G C A T G A T C T C A G G C T A G A C T A G T C C T A G C G T A C A T G G T C A | GATA20(C2C2gata)/colamp-GATA20-DAP-Seq(GSE60143)/Homer | 1e-7 | -1.696e+01 | 0.0000 | 1430.0 | 88.44% | 6132.7 | 83.20% | motif file (matrix) | svg |
| 273 | T A C G C G T A G A C T T C A G A G C T A G T C A C T G T C A G A G T C C T G A | DDF2(AP2EREBP)/col-DDF2-DAP-Seq(GSE60143)/Homer | 1e-7 | -1.695e+01 | 0.0000 | 212.0 | 13.11% | 635.1 | 8.62% | motif file (matrix) | svg |
| 274 | G A C T T C G A C G T A C G T A C G T A C G T A C G T A C T A G A G C T C G T A | dof45(C2C2dof)/col-dof45-DAP-Seq(GSE60143)/Homer | 1e-7 | -1.689e+01 | 0.0000 | 1257.0 | 77.74% | 5252.0 | 71.25% | motif file (matrix) | svg |
| 275 | G T A C A C G T A C G T T A C G A T G C C A T G T A C G G T A C T C A G A T G C C G T A G T C A A C T G A G C T G C T A | AT1G19040(NAC)/col-AT1G19040-DAP-Seq(GSE60143)/Homer | 1e-7 | -1.688e+01 | 0.0000 | 181.0 | 11.19% | 520.6 | 7.06% | motif file (matrix) | svg |
| 276 | C G A T T C A G G T A C A C G T A C G T T C A G C G A T C G T A G T C A G C T A C G T A A G T C C G T A G T C A C A T G | ANAC057(NAC)/colamp-ANAC057-DAP-Seq(GSE60143)/Homer | 1e-7 | -1.671e+01 | 0.0000 | 853.0 | 52.75% | 3349.2 | 45.44% | motif file (matrix) | svg |
| 277 | T C A G T C A G A C G T G T A C G C T A T C A G C T G A A C T G A C T G A G C T A G T C C G T A | EAR2(NR)/K562-NR2F6-ChIP-Seq(Encode)/Homer | 1e-7 | -1.659e+01 | 0.0000 | 927.0 | 57.33% | 3689.2 | 50.05% | motif file (matrix) | svg |
| 278 | T C A G G A C T G T C A C G T A A C G T A T C G C G T A A C G T A C G T C T G A | ATHB15(HB)/col-ATHB15-DAP-Seq(GSE60143)/Homer | 1e-7 | -1.633e+01 | 0.0000 | 485.0 | 29.99% | 1743.6 | 23.65% | motif file (matrix) | svg |
| 279 | C T G A A G C T A C G T A C G T A G T C G A C T G A C T C T G A C T G A C T A G C G T A C G T A | STAT6(Stat)/CD4-Stat6-ChIP-Seq(GSE22104)/Homer | 1e-7 | -1.629e+01 | 0.0000 | 441.0 | 27.27% | 1559.7 | 21.16% | motif file (matrix) | svg |
| 280 | A G T C G T A C C T G A A G T C A G T C C A T G G C T A A G T C T G C A G C T A G C A T G C A T | RAP21(AP2EREBP)/colamp-RAP21-DAP-Seq(GSE60143)/Homer | 1e-7 | -1.626e+01 | 0.0000 | 507.0 | 31.35% | 1837.8 | 24.93% | motif file (matrix) | svg |
| 281 | C T A G G T A C A C G T A C G T A T C G G C A T A G C T A G C T A G C T G C A T G A C T C G T A G T C A A C T G G A C T | VND6(NAC)/col-VND6-DAP-Seq(GSE60143)/Homer | 1e-7 | -1.617e+01 | 0.0000 | 1070.0 | 66.17% | 4363.8 | 59.20% | motif file (matrix) | svg |
| 282 | C T G A A T C G G T A C C T G A A G T C A G T C A C T G C G T A A G T C C T G A | TINY(AP2EREBP)/col-TINY-DAP-Seq(GSE60143)/Homer | 1e-6 | -1.597e+01 | 0.0000 | 625.0 | 38.65% | 2349.9 | 31.88% | motif file (matrix) | svg |
| 283 | A G T C C G A T A C T G A T C G T G A C G C T A C A T G A T C G T G A C C G A T A C T G T A G C G T A C G T C A | Tlx?(NR)/NPC-H3K4me1-ChIP-Seq(GSE16256)/Homer | 1e-6 | -1.589e+01 | 0.0000 | 249.0 | 15.40% | 788.4 | 10.70% | motif file (matrix) | svg |
| 284 | A G T C G A T C A G T C C G T A A T C G C A G T A G T C G T A C C T G A A C T G T C A G A G C T A G C T A G C T A G C T | PRDM15(Zf)/ESC-Prdm15-ChIP-Seq(GSE73694)/Homer | 1e-6 | -1.588e+01 | 0.0000 | 743.0 | 45.95% | 2870.0 | 38.94% | motif file (matrix) | svg |
| 285 | G C T A C G T A C G T A G C A T C A T G C T A G A G T C A C T G T A C G A G T C C A T G T A C G | RAP26(AP2EREBP)/colamp-RAP26-DAP-Seq(GSE60143)/Homer | 1e-6 | -1.588e+01 | 0.0000 | 1204.0 | 74.46% | 5010.5 | 67.97% | motif file (matrix) | svg |
| 286 | C G A T C G T A G C T A G C A T G C A T C T G A A C T G A C G T A G T C C G T A C G T A G T A C T C A G G C T A C G A T | WRKY25(WRKY)/colamp-WRKY25-DAP-Seq(GSE60143)/Homer | 1e-6 | -1.583e+01 | 0.0000 | 1037.0 | 64.13% | 4214.5 | 57.18% | motif file (matrix) | svg |
| 287 | A T G C G A C T A C G T C T A G A C G T A C G T A C G T C T G A G A T C G C T A A G C T C G T A | Foxa2(Forkhead)/Liver-Foxa2-ChIP-Seq(GSE25694)/Homer | 1e-6 | -1.573e+01 | 0.0000 | 714.0 | 44.16% | 2744.4 | 37.23% | motif file (matrix) | svg |
| 288 | G A T C G C A T G C A T A G T C A G C T T C G A T A C G G C T A C G T A C T A G T G A C G C A T C G A T G A T C A G C T | HSFC1(HSF)/col-HSFC1-DAP-Seq(GSE60143)/Homer | 1e-6 | -1.568e+01 | 0.0000 | 197.0 | 12.18% | 591.2 | 8.02% | motif file (matrix) | svg |
| 289 | G C A T G C A T G A T C G A C T T C G A T C A G G C T A C G T A A C T G G T A C G C A T G C A T A G T C A G C T C G T A | HSF7(HSF)/colamp-HSF7-DAP-Seq(GSE60143)/Homer | 1e-6 | -1.560e+01 | 0.0000 | 192.0 | 11.87% | 573.9 | 7.79% | motif file (matrix) | svg |
| 290 | G A T C C T G A A G T C A G T C A C T G C G T A A G T C C T G A | ERF38(AP2EREBP)/col-ERF38-DAP-Seq(GSE60143)/Homer | 1e-6 | -1.556e+01 | 0.0000 | 876.0 | 54.17% | 3475.1 | 47.14% | motif file (matrix) | svg |
| 291 | T G C A C G T A A C T G T C A G C A G T C A T G T C A G G A T C T A C G A G T C T G C A A C T G A C T G T G A C G T C A | ZNF165(Zf)/WHIM12-ZNF165-ChIP-Seq(GSE65937)/Homer | 1e-6 | -1.549e+01 | 0.0000 | 84.0 | 5.19% | 191.6 | 2.60% | motif file (matrix) | svg |
| 292 | A C T G C T A G A G T C A C T G A C T G A T G C A C T G T A C G | ESE1(AP2EREBP)/col-ESE1-DAP-Seq(GSE60143)/Homer | 1e-6 | -1.544e+01 | 0.0000 | 682.0 | 42.18% | 2608.4 | 35.39% | motif file (matrix) | svg |
| 293 | T G A C C A T G A C G T A C G T A C T G C G T A A G T C A G C T G C A T T C G A | WRKY30(WRKY)/colamp-WRKY30-DAP-Seq(GSE60143)/Homer | 1e-6 | -1.533e+01 | 0.0000 | 498.0 | 30.80% | 1814.4 | 24.61% | motif file (matrix) | svg |
| 294 | G A T C A G C T C T A G G A T C T G A C C T A G C G T A G T A C C G T A G C A T G T C A C T G A | CBF3(AP2EREBP)/colamp-CBF3-DAP-Seq(GSE60143)/Homer | 1e-6 | -1.529e+01 | 0.0000 | 919.0 | 56.83% | 3677.4 | 49.89% | motif file (matrix) | svg |
| 295 | G C A T G A C T A T G C A G C T T C G A A C T G C G T A C G T A A T C G A T G C G C A T G A C T G A T C A G C T T C G A | HSFA1E(HSF)/col-HSFA1E-DAP-Seq(GSE60143)/Homer | 1e-6 | -1.520e+01 | 0.0000 | 89.0 | 5.50% | 209.9 | 2.85% | motif file (matrix) | svg |
| 296 | C T A G T G A C G A C T A T C G T C G A A G T C C T A G C A G T C T A G A T C G G T A C T C G A | O2(bZIP)/Corn-O2-ChIP-Seq(GSE63991)/Homer | 1e-6 | -1.508e+01 | 0.0000 | 234.0 | 14.47% | 739.7 | 10.03% | motif file (matrix) | svg |
| 297 | G A C T A C T G C G A T A G T C A C T G C T A G A G T C C G T A | Rap210(AP2EREBP)/col-Rap210-DAP-Seq(GSE60143)/Homer | 1e-6 | -1.497e+01 | 0.0000 | 1042.0 | 64.44% | 4254.3 | 57.72% | motif file (matrix) | svg |
| 298 | G C A T G A C T A T G C A G C T T C G A C T A G G C T A C G T A C A T G T G A C G C A T G A C T A G T C A G C T C T G A | HSFB4(HSF)/col-HSFB4-DAP-Seq(GSE60143)/Homer | 1e-6 | -1.482e+01 | 0.0000 | 49.0 | 3.03% | 87.2 | 1.18% | motif file (matrix) | svg |
| 299 | G A T C C A T G A C G T A C G T A C T G C G T A A G T C A G C T C G A T G A C T | WRKY8(WRKY)/colamp-WRKY8-DAP-Seq(GSE60143)/Homer | 1e-6 | -1.463e+01 | 0.0000 | 157.0 | 9.71% | 452.6 | 6.14% | motif file (matrix) | svg |
| 300 | G A T C C T G A A G T C G T A C A C T G G C T A G A T C C T G A G C T A G C T A | At4g31060(AP2EREBP)/colamp-At4g31060-DAP-Seq(GSE60143)/Homer | 1e-6 | -1.450e+01 | 0.0000 | 836.0 | 51.70% | 3314.8 | 44.97% | motif file (matrix) | svg |
| 301 | A G C T G C A T A C T G A C G T A G T C A C G T C T A G T A C G | Smad3(MAD)/NPC-Smad3-ChIP-Seq(GSE36673)/Homer | 1e-6 | -1.441e+01 | 0.0000 | 1278.0 | 79.04% | 5399.3 | 73.25% | motif file (matrix) | svg |
| 302 | T G C A C G T A G T C A A G C T A G T C G C T A T A G C C G A T C T A G G A T C | Gfi1b(Zf)/HPC7-Gfi1b-ChIP-Seq(GSE22178)/Homer | 1e-6 | -1.436e+01 | 0.0000 | 551.0 | 34.08% | 2057.8 | 27.92% | motif file (matrix) | svg |
| 303 | G C T A A C T G T C G A C T G A C T G A A C G T T A G C C T G A C G T A C G A T | Cux2(Homeobox)/Liver-Cux2-ChIP-Seq(GSE35985)/Homer | 1e-6 | -1.435e+01 | 0.0000 | 887.0 | 54.85% | 3549.1 | 48.15% | motif file (matrix) | svg |
| 304 | C G A T G A T C G A T C C T G A G A T C G A T C C A T G T G C A G T A C T C G A G T C A G C A T C G A T C G A T G C A T | At4g32800(AP2EREBP)/colamp-At4g32800-DAP-Seq(GSE60143)/Homer | 1e-6 | -1.431e+01 | 0.0000 | 402.0 | 24.86% | 1428.0 | 19.37% | motif file (matrix) | svg |
| 305 | G A C T A G C T G T A C G A C T C T G A A C T G G T C A C T G A A T G C T A C G G A C T A C G T A G T C G A C T C T G A | HRE(HSF)/Striatum-HSF1-ChIP-Seq(GSE38000)/Homer | 1e-6 | -1.425e+01 | 0.0000 | 177.0 | 10.95% | 530.1 | 7.19% | motif file (matrix) | svg |
| 306 | G T A C G T A C G T C A G C T A C G T A C G T A C G T A C T A G C T A G C T A G | SEP3(MADS)/Arabidoposis-Flower-Sep3-ChIP-Seq/Homer | 1e-6 | -1.422e+01 | 0.0000 | 810.0 | 50.09% | 3202.7 | 43.45% | motif file (matrix) | svg |
| 307 | C G T A G C A T C G T A C G T A G C A T A C T G C G A T A G T C A C T G A C T G G A C T C T A G | AT1G71450(AP2EREBP)/col-AT1G71450-DAP-Seq(GSE60143)/Homer | 1e-6 | -1.411e+01 | 0.0000 | 1319.0 | 81.57% | 5608.2 | 76.08% | motif file (matrix) | svg |
| 308 | A G T C G A T C A G C T C G T A G T A C A G T C G C A T C T G A G T A C G A T C | AT4G26030(C2H2)/col-AT4G26030-DAP-Seq(GSE60143)/Homer | 1e-6 | -1.407e+01 | 0.0000 | 979.0 | 60.54% | 3978.0 | 53.97% | motif file (matrix) | svg |
| 309 | A G T C G A C T C A G T A C T G C T A G T G A C G C T A A T G C G C A T A T C G C G A T A C T G G A T C G T A C G T C A C T G A | NF1(CTF)/LNCAP-NF1-ChIP-Seq(Unpublished)/Homer | 1e-6 | -1.399e+01 | 0.0000 | 259.0 | 16.02% | 850.5 | 11.54% | motif file (matrix) | svg |
| 310 | G A C T A C T G C G A T A G T C A C T G C T A G A G T C C T G A | AT1G12630(AP2EREBP)/colamp-AT1G12630-DAP-Seq(GSE60143)/Homer | 1e-6 | -1.399e+01 | 0.0000 | 904.0 | 55.91% | 3634.9 | 49.31% | motif file (matrix) | svg |
| 311 | A T C G A G T C A G T C C G T A A C T G G C A T | hINR(CPE) | 1e-6 | -1.392e+01 | 0.0000 | 877.0 | 54.24% | 3512.8 | 47.66% | motif file (matrix) | svg |
| 312 | C T A G C A T G G A C T C G T A C T A G A C T G A C G T C T A G C T A G T C A G | MYB17(MYB)/colamp-MYB17-DAP-Seq(GSE60143)/Homer | 1e-6 | -1.389e+01 | 0.0000 | 659.0 | 40.75% | 2536.7 | 34.41% | motif file (matrix) | svg |
| 313 | C T A G C T A G T C G A C G T A A T G C C G T A A T C G T C G A T A C G G C A T A C T G C A G T T A G C G A T C G A C T | MRE(NR)/Neuro2A-NR3C2-ChIPnexus(GSE115417)/Homer | 1e-6 | -1.384e+01 | 0.0000 | 904.0 | 55.91% | 3637.6 | 49.35% | motif file (matrix) | svg |
| 314 | A C T G A G T C G T C A C G T A A G T C C G T A C T A G C T A G G A C T C A T G | SCRT1(Zf)/HEK293-SCRT1.eGFP-ChIP-Seq(Encode)/Homer | 1e-6 | -1.383e+01 | 0.0000 | 411.0 | 25.42% | 1472.2 | 19.97% | motif file (matrix) | svg |
| 315 | G A C T C T A G C T A G A G T C T G C A A C T G A C G T A C G T C T A G T C A G | AMYB(HTH)/Testes-AMYB-ChIP-Seq(GSE44588)/Homer | 1e-5 | -1.374e+01 | 0.0000 | 1256.0 | 77.67% | 5303.9 | 71.95% | motif file (matrix) | svg |
| 316 | C T A G C T G A C G T A C G T A C G T A C G T A A C T G A C G T C T A G G T C A | COG1(C2C2dof)/col-COG1-DAP-Seq(GSE60143)/Homer | 1e-5 | -1.374e+01 | 0.0000 | 954.0 | 59.00% | 3869.4 | 52.49% | motif file (matrix) | svg |
| 317 | C G T A G A C T C G T A A C G T C A G T A G T C A G C T G A C T | KAN2(G2like)/colamp-KAN2-DAP-Seq(GSE60143)/Homer | 1e-5 | -1.367e+01 | 0.0000 | 1029.0 | 63.64% | 4219.3 | 57.24% | motif file (matrix) | svg |
| 318 | T A C G C T G A T C G A C G A T C T A G C T A G T C G A C T G A T C G A T C G A C G T A T C G A G C A T C A T G C G T A T A C G G C A T T G A C C G T A A G C T | NFAT:AP1(RHD,bZIP)/Jurkat-NFATC1-ChIP-Seq(Jolma\_et\_al.)/Homer | 1e-5 | -1.366e+01 | 0.0000 | 133.0 | 8.23% | 373.4 | 5.07% | motif file (matrix) | svg |
| 319 | C T G A G A C T C A T G C T A G A G T C A C T G A C T G A G T C A C T G T C A G | ERF11(AP2EREBP)/col-ERF11-DAP-Seq(GSE60143)/Homer | 1e-5 | -1.362e+01 | 0.0000 | 1020.0 | 63.08% | 4178.8 | 56.69% | motif file (matrix) | svg |
| 320 | C T G A T C A G G C T A A G C T A G T C G A C T C T G A C T A G T G C A C T G A A G T C G T A C G A T C A C T G T C G A | ZBTB12(Zf)/HEK293-ZBTB12.GFP-ChIP-Seq(GSE58341)/Homer | 1e-5 | -1.360e+01 | 0.0000 | 452.0 | 27.95% | 1648.2 | 22.36% | motif file (matrix) | svg |
| 321 | C G A T C G T A G C T A G A C T T C G A A G C T A G T C A C T G T C G A A G C T C T G A C G A T | ZBTB38(Zf)/Hela-ZBTB38-ChIP-seq(GSE108618)/Homer | 1e-5 | -1.344e+01 | 0.0000 | 1453.0 | 89.86% | 6305.8 | 85.55% | motif file (matrix) | svg |
| 322 | G T A C G C T A C G A T C A G T A G T C G C T A C G A T C G A T A G T C G C T A | WUS1(Homeobox)/colamp-WUS1-DAP-Seq(GSE60143)/Homer | 1e-5 | -1.342e+01 | 0.0000 | 456.0 | 28.20% | 1668.2 | 22.63% | motif file (matrix) | svg |
| 323 | G C T A C G T A C G A T G A C T G C A T T G C A A G T C A G C T A C G T A C G T C G A T G A C T | DAG2(C2C2dof)/col-DAG2-DAP-Seq(GSE60143)/Homer | 1e-5 | -1.333e+01 | 0.0000 | 985.0 | 60.92% | 4021.8 | 54.56% | motif file (matrix) | svg |
| 324 | T C G A T A G C G T C A A C T G A C T G C G T A C G T A C T A G A G C T T C A G | ERG(ETS)/VCaP-ERG-ChIP-Seq(GSE14097)/Homer | 1e-5 | -1.318e+01 | 0.0000 | 827.0 | 51.14% | 3300.6 | 44.78% | motif file (matrix) | svg |
| 325 | C G A T C T G A G T A C C A T G G C A T T C A G G C A T C G T A C G T A G C T A C G T A A G T C G C T A G T A C C A T G | CUC2(NAC)/colamp-CUC2-DAP-Seq(GSE60143)/Homer | 1e-5 | -1.314e+01 | 0.0000 | 556.0 | 34.38% | 2101.7 | 28.51% | motif file (matrix) | svg |
| 326 | T G C A A G C T C A T G C G T A A G C T A C T G G A T C G T C A C G T A A G C T | Atf4(bZIP)/MEF-Atf4-ChIP-Seq(GSE35681)/Homer | 1e-5 | -1.303e+01 | 0.0000 | 426.0 | 26.35% | 1549.0 | 21.01% | motif file (matrix) | svg |
| 327 | T A G C G T A C A G T C G T A C C G A T A G T C A G T C A G T C A G T C A G T C C G T A G A T C | Zfp281(Zf)/ES-Zfp281-ChIP-Seq(GSE81042)/Homer | 1e-5 | -1.299e+01 | 0.0000 | 35.0 | 2.16% | 55.6 | 0.75% | motif file (matrix) | svg |
| 328 | C T A G A C T G A C G T C G T A A C T G A C T G A G C T C T A G T C A G C T A G | MYB93(MYB)/colamp-MYB93-DAP-Seq(GSE60143)/Homer | 1e-5 | -1.297e+01 | 0.0000 | 998.0 | 61.72% | 4090.0 | 55.49% | motif file (matrix) | svg |
| 329 | C A G T T C G A A G T C A C G T A C G T T C A G C G A T G C T A G C T A C G T A G C A T A G T C C G T A T G C A A C T G | ANAC045(NAC)/col-ANAC045-DAP-Seq(GSE60143)/Homer | 1e-5 | -1.278e+01 | 0.0000 | 1260.0 | 77.92% | 5341.0 | 72.46% | motif file (matrix) | svg |
| 330 | A G C T C T A G A G T C A G T C A C T G C G T A A G T C C T G A G C A T G C T A C T G A G C A T G C A T C G A T G C A T | CBF4(AP2EREBP)/colamp-CBF4-DAP-Seq(GSE60143)/Homer | 1e-5 | -1.269e+01 | 0.0000 | 1109.0 | 68.58% | 4616.5 | 62.63% | motif file (matrix) | svg |
| 331 | G A C T C T A G C T A G G T A C A G T C G A T C G A C T G A C T T A G C T C A G | NLP7(RWPRK)/col-NLP7-DAP-Seq(GSE60143)/Homer | 1e-5 | -1.267e+01 | 0.0000 | 1148.0 | 71.00% | 4802.6 | 65.15% | motif file (matrix) | svg |
| 332 | C A G T C A T G T G C A G T A C C G T A T C A G G T A C G A C T T C A G C T G A | bZIP18(bZIP)/colamp-bZIP18-DAP-Seq(GSE60143)/Homer | 1e-5 | -1.263e+01 | 0.0000 | 1459.0 | 90.23% | 6349.1 | 86.13% | motif file (matrix) | svg |
| 333 | C G T A C G T A C G T A C T G A C T A G A C G T C T A G G T C A | CDF3(C2C2dof)/colamp-CDF3-DAP-Seq(GSE60143)/Homer | 1e-5 | -1.262e+01 | 0.0000 | 1071.0 | 66.23% | 4439.0 | 60.22% | motif file (matrix) | svg |
| 334 | A T C G A G T C A C T G A G T C A G T C A C T G G A C T G A C T | PUCHI(AP2EREBP)/colamp-PUCHI-DAP-Seq(GSE60143)/Homer | 1e-5 | -1.257e+01 | 0.0000 | 880.0 | 54.42% | 3554.9 | 48.23% | motif file (matrix) | svg |
| 335 | T G A C C T G A C T A G T C G A C T G A A T G C C G T A A C T G G C A T G T A C G C A T A T C G G C A T A G C T G A T C | PR(NR)/T47D-PR-ChIP-Seq(GSE31130)/Homer | 1e-5 | -1.253e+01 | 0.0000 | 1122.0 | 69.39% | 4681.9 | 63.52% | motif file (matrix) | svg |
| 336 | T C G A C G T A C G T A T C G A A C T G G T A C C G T A A G C T G T C A G C A T | At3g24120(G2like)/col-At3g24120-DAP-Seq(GSE60143)/Homer | 1e-5 | -1.252e+01 | 0.0000 | 1429.0 | 88.37% | 6195.0 | 84.04% | motif file (matrix) | svg |
| 337 | C G A T C A G T C T A G G C T A A G T C C G T A T C A G A G T C A C G T A C T G A C G T G T A C G C T A G C T A G C T A | bZIP52(bZIP)/colamp-bZIP52-DAP-Seq(GSE60143)/Homer | 1e-5 | -1.250e+01 | 0.0000 | 902.0 | 55.78% | 3656.9 | 49.61% | motif file (matrix) | svg |
| 338 | C G T A C G A T C G T A C G A T C A T G A C T G C G A T A G T C A T C G T C A G G A C T A C T G | At1g36060(AP2EREBP)/colamp-At1g36060-DAP-Seq(GSE60143)/Homer | 1e-5 | -1.247e+01 | 0.0000 | 972.0 | 60.11% | 3979.3 | 53.98% | motif file (matrix) | svg |
| 339 | C T A G C T A G C G T A C G T A T A C G C G A T C T A G C T G A C T G A C G T A T A C G G A C T | PU.1:IRF8(ETS:IRF)/pDC-Irf8-ChIP-Seq(GSE66899)/Homer | 1e-5 | -1.239e+01 | 0.0000 | 131.0 | 8.10% | 377.0 | 5.12% | motif file (matrix) | svg |
| 340 | C G A T G T C A A G T C A C G T A C G T A C T G G A C T C G A T A T C G G C T A G T C A A G T C C G T A G T C A A C T G | ANAC017(NAC)/colamp-ANAC017-DAP-Seq(GSE60143)/Homer | 1e-5 | -1.233e+01 | 0.0000 | 184.0 | 11.38% | 577.1 | 7.83% | motif file (matrix) | svg |
| 341 | G C T A A G T C T A C G T G C A A T C G T C A G G C T A T C G A T C A G A G C T | ELF5(ETS)/T47D-ELF5-ChIP-Seq(GSE30407)/Homer | 1e-5 | -1.226e+01 | 0.0000 | 588.0 | 36.36% | 2257.8 | 30.63% | motif file (matrix) | svg |
| 342 | G A C T G A C T G A T C C G T A G T A C A G T C G C A T C G T A G T A C G A T C G C A T G C T A | MYB74(MYB)/colamp-MYB74-DAP-Seq(GSE60143)/Homer | 1e-5 | -1.224e+01 | 0.0000 | 715.0 | 44.22% | 2817.3 | 38.22% | motif file (matrix) | svg |
| 343 | A T C G T G A C A T G C C T G A T C A G G A C T A G T C C G A T T C A G T C G A C A T G C T A G C T A G C G T A C T A G C T A G C T G A C T A G C T A G A T G C | ZSCAN22(Zf)/HEK293-ZSCAN22.GFP-ChIP-Seq(GSE58341)/Homer | 1e-5 | -1.212e+01 | 0.0000 | 40.0 | 2.47% | 73.0 | 0.99% | motif file (matrix) | svg |
| 344 | T C A G G T C A G C T A A G T C C G T A A T C G T C G A T C A G C G T A A C T G A C T G A C T G | ZNF768(Zf)/Rajj-ZNF768-ChIP-Seq(GSE111879)/Homer | 1e-5 | -1.212e+01 | 0.0000 | 35.0 | 2.16% | 58.5 | 0.79% | motif file (matrix) | svg |
| 345 | G A T C G A T C G A T C C G T A G T A C A G T C G C A T C G T A G T A C G A T C | MYB58(MYB)/colamp-MYB58-DAP-Seq(GSE60143)/Homer | 1e-5 | -1.208e+01 | 0.0000 | 960.0 | 59.37% | 3932.1 | 53.34% | motif file (matrix) | svg |
| 346 | G A C T G T A C T G C A A C G T G A T C G C T A T C G A A C G T A G T C C G T A | Pdx1(Homeobox)/Islet-Pdx1-ChIP-Seq(SRA008281)/Homer | 1e-5 | -1.204e+01 | 0.0000 | 874.0 | 54.05% | 3538.0 | 48.00% | motif file (matrix) | svg |
| 347 | C G T A G C A T C A T G C T A G A G T C A C T G A T C G G T A C A C T G T C A G | At2g33710(AP2EREBP)/colamp-At2g33710-DAP-Seq(GSE60143)/Homer | 1e-5 | -1.203e+01 | 0.0000 | 1301.0 | 80.46% | 5557.5 | 75.39% | motif file (matrix) | svg |
| 348 | C G A T T C G A G T A C A C T G A C G T T C A G G C A T T G C A G C T A G C A T C G T A A G T C C G T A G T A C C A T G | ANAC087(NAC)/col-ANAC087-DAP-Seq(GSE60143)/Homer | 1e-5 | -1.198e+01 | 0.0000 | 543.0 | 33.58% | 2067.9 | 28.05% | motif file (matrix) | svg |
| 349 | C T A G C A G T C G T A A C G T A G T C A C T G C G T A A G C T A G T C G A T C | HNF6(Homeobox)/Liver-Hnf6-ChIP-Seq(ERP000394)/Homer | 1e-5 | -1.196e+01 | 0.0000 | 1012.0 | 62.59% | 4176.5 | 56.66% | motif file (matrix) | svg |
| 350 | A C T G C T A G A G T C A C T G A C T G A G T C A C T G T A C G | ERF104(AP2EREBP)/col-ERF104-DAP-Seq(GSE60143)/Homer | 1e-5 | -1.186e+01 | 0.0000 | 914.0 | 56.52% | 3725.1 | 50.54% | motif file (matrix) | svg |
| 351 | C G T A C T A G C A G T A C G T C G T A A C T G C A T G G C A T T C A G C T G A | MYB49(MYB)/col-MYB49-DAP-Seq(GSE60143)/Homer | 1e-5 | -1.183e+01 | 0.0000 | 986.0 | 60.98% | 4058.5 | 55.06% | motif file (matrix) | svg |
| 352 | A G T C C G T A T G A C A T G C G C A T C T G A G T A C G A T C | MYB55(MYB)/colamp-MYB55-DAP-Seq(GSE60143)/Homer | 1e-5 | -1.181e+01 | 0.0000 | 1040.0 | 64.32% | 4310.3 | 58.47% | motif file (matrix) | svg |
| 353 | A C T G A C G T C G A T C A G T C A T G C A T G C A G T G C A T C A G T C A T G | HuR(?)/HEK293-HuR-CLIP-Seq(GSE87887)/Homer | 1e-5 | -1.180e+01 | 0.0000 | 1331.0 | 82.31% | 5710.6 | 77.47% | motif file (matrix) | svg |
| 354 | T C A G T C A G G C T A C G T A T A C G G A C T T C A G T C G A C T G A C G T A T A C G G A C T | IRF8(IRF)/BMDM-IRF8-ChIP-Seq(GSE77884)/Homer | 1e-5 | -1.173e+01 | 0.0000 | 246.0 | 15.21% | 828.7 | 11.24% | motif file (matrix) | svg |
| 355 | C G A T C T G A G T A C A C G T A C G T T C A G C G T A C G T A G C T A G C A T G C A T A G T C C G T A G T C A C A T G | NST1(NAC)/colamp-NST1-DAP-Seq(GSE60143)/Homer | 1e-5 | -1.164e+01 | 0.0000 | 867.0 | 53.62% | 3515.5 | 47.69% | motif file (matrix) | svg |
| 356 | A G T C A G T C C T G A A G T C A G T C A C T G C G T A A G T C T C G A G A T C C G A T C G T A | AT1G01250(AP2EREBP)/col-AT1G01250-DAP-Seq(GSE60143)/Homer | 1e-5 | -1.161e+01 | 0.0000 | 227.0 | 14.04% | 754.1 | 10.23% | motif file (matrix) | svg |
| 357 | C G T A C G T A C T A G A C G T A C G T C G T A A C T G A C T G A C G T C T G A T C G A T C G A | MYB4(MYB)/col200-MYB4-DAP-Seq(GSE60143)/Homer | 1e-5 | -1.155e+01 | 0.0000 | 713.0 | 44.09% | 2823.4 | 38.30% | motif file (matrix) | svg |
| 358 | A G C T G T C A T G C A A G T C A C G T A C G T A C G T C G A T G A C T T A C G | AT3G12130(C3H)/colamp-AT3G12130-DAP-Seq(GSE60143)/Homer | 1e-4 | -1.151e+01 | 0.0000 | 1226.0 | 75.82% | 5201.0 | 70.56% | motif file (matrix) | svg |
| 359 | C A T G G A T C C T G A A G T C C T A G C G T A G C T A G C A T G A T C G A T C A G T C C T A G C G T A C A T G C T A G | PLT1(AP2EREBP)/colamp-PLT1-DAP-Seq(GSE60143)/Homer | 1e-4 | -1.146e+01 | 0.0000 | 205.0 | 12.68% | 669.0 | 9.08% | motif file (matrix) | svg |
| 360 | C T A G T A G C A T G C C T A G A G T C A G T C C T A G G A C T G A C T G C T A | CRF10(AP2EREBP)/col100-CRF10-DAP-Seq(GSE60143)/Homer | 1e-4 | -1.145e+01 | 0.0000 | 1183.0 | 73.16% | 4995.3 | 67.77% | motif file (matrix) | svg |
| 361 | G C A T C T A G G T A C A G T C C G A T A C T G C T A G C T A G G T A C G C T A | ZNF416(Zf)/HEK293-ZNF416.GFP-ChIP-Seq(GSE58341)/Homer | 1e-4 | -1.132e+01 | 0.0000 | 651.0 | 40.26% | 2553.4 | 34.64% | motif file (matrix) | svg |
| 362 | G T A C A C G T A C G T T C A G G C T A C G T A C G A T G C A T G C A T A G T C C G T A G T C A C A T G G A C T G C T A | ANAC070(NAC)/colamp-ANAC070-DAP-Seq(GSE60143)/Homer | 1e-4 | -1.129e+01 | 0.0000 | 1018.0 | 62.96% | 4219.5 | 57.24% | motif file (matrix) | svg |
| 363 | C A T G T G A C C A T G G A C T C A G T C T A G G C T A G T A C G A C T G C A T G C A T C G A T | WRKY21(WRKY)/colamp-WRKY21-DAP-Seq(GSE60143)/Homer | 1e-4 | -1.126e+01 | 0.0000 | 150.0 | 9.28% | 459.7 | 6.24% | motif file (matrix) | svg |
| 364 | C G T A T G A C T C G A A G T C C G T A A T C G A T G C A C G T A C T G A G T C | E2A(bHLH)/proBcell-E2A-ChIP-Seq(GSE21978)/Homer | 1e-4 | -1.124e+01 | 0.0000 | 764.0 | 47.25% | 3058.6 | 41.49% | motif file (matrix) | svg |
| 365 | G C A T T G A C C T G A A G T C A G T C A C T G G T C A A G T C G C T A G A C T G C T A C T G A | DREB2(AP2EREBP)/col-DREB2-DAP-Seq(GSE60143)/Homer | 1e-4 | -1.123e+01 | 0.0000 | 807.0 | 49.91% | 3252.1 | 44.12% | motif file (matrix) | svg |
| 366 | C G T A C G A T C A G T C A T G C G A T G T A C C A T G A C T G G A C T C A T G | CEJ1(AP2EREBP)/col-CEJ1-DAP-Seq(GSE60143)/Homer | 1e-4 | -1.122e+01 | 0.0000 | 1174.0 | 72.60% | 4957.3 | 67.25% | motif file (matrix) | svg |
| 367 | G A C T A T C G C T G A A G T C T C A G G A C T G T A C C T G A A G C T G T A C | TGA6(bZIP)/colamp-TGA6-DAP-Seq(GSE60143)/Homer | 1e-4 | -1.116e+01 | 0.0000 | 878.0 | 54.30% | 3576.4 | 48.52% | motif file (matrix) | svg |
| 368 | T C G A G A C T A T C G C G T A A G T C C T A G G C A T G T A C C T G A A C G T G A T C G C T A | TGA4(bZIP)/colamp-TGA4-DAP-Seq(GSE60143)/Homer | 1e-4 | -1.114e+01 | 0.0000 | 480.0 | 29.68% | 1812.3 | 24.59% | motif file (matrix) | svg |
| 369 | T G C A A C T G T A C G A T G C A G T C G A C T T C G A A T C G | ZNF711(Zf)/SHSY5Y-ZNF711-ChIP-Seq(GSE20673)/Homer | 1e-4 | -1.104e+01 | 0.0000 | 1041.0 | 64.38% | 4332.7 | 58.78% | motif file (matrix) | svg |
| 370 | T A G C G T A C C G T A C T A G A C T G T G C A C G T A A T G C C G T A A T C G | AR-halfsite(NR)/LNCaP-AR-ChIP-Seq(GSE27824)/Homer | 1e-4 | -1.102e+01 | 0.0000 | 1312.0 | 81.14% | 5631.3 | 76.40% | motif file (matrix) | svg |
| 371 | T C A G G C A T A C T G C G T A A G T C C T A G G C A T T G A C | TGA9(bZIP)/colamp-TGA9-DAP-Seq(GSE60143)/Homer | 1e-4 | -1.098e+01 | 0.0000 | 1157.0 | 71.55% | 4881.5 | 66.22% | motif file (matrix) | svg |
| 372 | G A T C G C A T T C G A A G T C A C G T A C G T A C G T C G A T A C G T A T C G | AT1G47655(C2C2dof)/colamp-AT1G47655-DAP-Seq(GSE60143)/Homer | 1e-4 | -1.089e+01 | 0.0001 | 1366.0 | 84.48% | 5902.1 | 80.07% | motif file (matrix) | svg |
| 373 | C T G A T C A G C T G A C T A G C A T G A C G T A T G C C G T A A T G C G C A T T C A G C T G A A C T G A C G T C A G T A G T C C G T A C A G T C T A G C A T G | VDR(NR),DR3/GM10855-VDR+vitD-ChIP-Seq(GSE22484)/Homer | 1e-4 | -1.069e+01 | 0.0001 | 205.0 | 12.68% | 679.9 | 9.22% | motif file (matrix) | svg |
| 374 | G A T C A G T C G A C T G C T A G T A C A G T C G C A T G C T A G T A C G A T C | MYB61(MYB)/colamp-MYB61-DAP-Seq(GSE60143)/Homer | 1e-4 | -1.056e+01 | 0.0001 | 1134.0 | 70.13% | 4781.7 | 64.87% | motif file (matrix) | svg |
| 375 | T G A C G T C A A G T C G C T A C T A G A G T C C T G A C A T G A C T G C T A G T A C G T C A G | Zic2(Zf)/ESC-Zic2-ChIP-Seq(SRP197560)/Homer | 1e-4 | -1.053e+01 | 0.0001 | 297.0 | 18.37% | 1053.1 | 14.29% | motif file (matrix) | svg |
| 376 | C G A T C T A G G A T C G C T A A G C T C T A G G A T C C G T A | RBFox2(?)/Heart-RBFox2-CLIP-Seq(GSE57926)/Homer | 1e-4 | -1.034e+01 | 0.0001 | 1221.0 | 75.51% | 5202.3 | 70.58% | motif file (matrix) | svg |
| 377 | C T G A T A C G G C A T A G C T A G C T A G T C T C G A A C T G C A G T A G C T A G C T G A T C | IRF3(IRF)/BMDM-Irf3-ChIP-Seq(GSE67343)/Homer | 1e-4 | -1.032e+01 | 0.0001 | 186.0 | 11.50% | 609.7 | 8.27% | motif file (matrix) | svg |
| 378 | T C G A C T G A C G T A C G T A C G T A C G T A A C T G A C G T C G A T C T G A | BBX31(Orphan)/col-BBX31-DAP-Seq(GSE60143)/Homer | 1e-4 | -1.029e+01 | 0.0001 | 1013.0 | 62.65% | 4219.7 | 57.25% | motif file (matrix) | svg |
| 379 | A T G C C T G A A T C G T A C G A G T C C G A T T C A G C G A T C T A G A G C T G T C A G T C A C G T A A G T C C G T A T A C G C T G A | Fox:Ebox(Forkhead,bHLH)/Panc1-Foxa2-ChIP-Seq(GSE47459)/Homer | 1e-4 | -1.028e+01 | 0.0001 | 656.0 | 40.57% | 2598.3 | 35.25% | motif file (matrix) | svg |
| 380 | T C G A C T G A C G T A C G T A C G T A C T G A A C T G A C G T C T G A C T G A | AT5G63260(C3H)/col-AT5G63260-DAP-Seq(GSE60143)/Homer | 1e-4 | -1.024e+01 | 0.0001 | 1196.0 | 73.96% | 5084.2 | 68.97% | motif file (matrix) | svg |
| 381 | C A G T G A C T G C A T T C G A A G T C A C G T A C G T A C G T C G A T G A C T | OBP3(C2C2dof)/col-OBP3-DAP-Seq(GSE60143)/Homer | 1e-4 | -1.021e+01 | 0.0001 | 1306.0 | 80.77% | 5618.6 | 76.22% | motif file (matrix) | svg |
| 382 | C G A T C T A G A C G T G T C A C G T A C G T A A G T C C G T A | Foxo3(Forkhead)/U2OS-Foxo3-ChIP-Seq(E-MTAB-2701)/Homer | 1e-4 | -1.021e+01 | 0.0001 | 684.0 | 42.30% | 2724.4 | 36.96% | motif file (matrix) | svg |
| 383 | G T A C A C T G A T G C T G A C C T A G G A C T G T A C C G T A G C A T G C A T | ERF8(AP2EREBP)/colamp-ERF8-DAP-Seq(GSE60143)/Homer | 1e-4 | -1.011e+01 | 0.0001 | 1168.0 | 72.23% | 4953.4 | 67.20% | motif file (matrix) | svg |
| 384 | A T G C T A G C A G C T A G C T T G A C G A C T T C A G T A C G G T C A C T G A A T C G T A G C G A C T C A G T A G T C A G C T T C G A A T C G T G C A T G C A | HRE(HSF)/HepG2-HSF1-ChIP-Seq(GSE31477)/Homer | 1e-4 | -1.008e+01 | 0.0001 | 130.0 | 8.04% | 396.3 | 5.38% | motif file (matrix) | svg |
| 385 | C A T G T A C G T A G C G A T C G A T C A T G C G T A C G A C T T C A G A T G C C G A T A T C G C A G T A C T G G T A C | Zic3(Zf)/mES-Zic3-ChIP-Seq(GSE37889)/Homer | 1e-4 | -9.989e+00 | 0.0001 | 319.0 | 19.73% | 1153.8 | 15.65% | motif file (matrix) | svg |
| 386 | C A T G A G C T A C T G A C G T A G T C C G T A A C T G C G T A C T G A C G T A G C T A C G T A C G A T G C A T C G T A | AtGRF6(GRF)/col-AtGRF6-DAP-Seq(GSE60143)/Homer | 1e-4 | -9.982e+00 | 0.0001 | 1053.0 | 65.12% | 4413.3 | 59.87% | motif file (matrix) | svg |
| 387 | A G C T C T A G G A T C A G T C C T A G C T G A A G T C G C T A G C A T T G C A | CBF1(AP2EREBP)/colamp-CBF1-DAP-Seq(GSE60143)/Homer | 1e-4 | -9.935e+00 | 0.0001 | 1038.0 | 64.19% | 4344.9 | 58.94% | motif file (matrix) | svg |
| 388 | G T A C G A T C C A G T A G T C A G T C A G T C T G C A G A T C C T G A A T G C G T C A A C G T | WT1(Zf)/Kidney-WT1-ChIP-Seq(GSE90016)/Homer | 1e-4 | -9.907e+00 | 0.0001 | 340.0 | 21.03% | 1242.3 | 16.85% | motif file (matrix) | svg |
| 389 | G C A T C G T A C G A T G A C T A C T G C T G A G A C T G A T C | Hnf6b(Homeobox)/LNCaP-Hnf6b-ChIP-Seq(GSE106305)/Homer | 1e-4 | -9.887e+00 | 0.0001 | 1083.0 | 66.98% | 4556.5 | 61.81% | motif file (matrix) | svg |
| 390 | G A C T A C T G C G T A A G T C T C A G G C A T G T A C C G T A A C G T G A T C | TGA1(bZIP)/colamp-TGA1-DAP-Seq(GSE60143)/Homer | 1e-4 | -9.880e+00 | 0.0001 | 574.0 | 35.50% | 2246.6 | 30.48% | motif file (matrix) | svg |
| 391 | C G A T C G A T G C A T G C A T G T C A A G T C A G C T A C G T A C G T C G A T G A C T A C G T | OBP4(C2C2dof)/col-OBP4-DAP-Seq(GSE60143)/Homer | 1e-4 | -9.876e+00 | 0.0001 | 925.0 | 57.20% | 3823.0 | 51.86% | motif file (matrix) | svg |
| 392 | C T G A G A T C G C A T A C T G C G T A A C G T C G T A C G T A T A C G T C G A | PQM-1(?)/cElegans-L3-ChIP-Seq(modEncode)/Homer | 1e-4 | -9.852e+00 | 0.0001 | 534.0 | 33.02% | 2072.7 | 28.12% | motif file (matrix) | svg |
| 393 | C T G A T C A G C A G T C T A G A C T G C T A G G A T C A T C G A C T G C T G A T C A G G A T C | Sp5(Zf)/mES-Sp5.Flag-ChIP-Seq(GSE72989)/Homer | 1e-4 | -9.841e+00 | 0.0001 | 510.0 | 31.54% | 1968.2 | 26.70% | motif file (matrix) | svg |
| 394 | C A T G A C G T A G T C G A T C G A T C G A T C G C T A C T A G C T A G C T A G T C A G T C G A | EBF1(EBF)/Near-E2A-ChIP-Seq(GSE21512)/Homer | 1e-4 | -9.642e+00 | 0.0002 | 512.0 | 31.66% | 1981.0 | 26.88% | motif file (matrix) | svg |
| 395 | G A T C G A T C G C T A G T C A G A C T A T G C T C G A C G A T C G A T C T A G | HAT2(Homeobox)/colamp-HAT2-DAP-Seq(GSE60143)/Homer | 1e-4 | -9.574e+00 | 0.0002 | 876.0 | 54.17% | 3605.5 | 48.91% | motif file (matrix) | svg |
| 396 | A G T C G T A C C T G A A G T C G T A C C T A G G C T A T G A C T G C A G C T A C G T A C G T A | At1g22810(AP2EREBP)/colamp-At1g22810-DAP-Seq(GSE60143)/Homer | 1e-4 | -9.561e+00 | 0.0002 | 767.0 | 47.43% | 3111.4 | 42.21% | motif file (matrix) | svg |
| 397 | C A T G A G T C C T G A A T G C C T A G T C G A G C T A G C A T G A T C G A C T A G T C C T A G C G T A C A T G C T A G | PLT3(AP2EREBP)/col-PLT3-DAP-Seq(GSE60143)/Homer | 1e-4 | -9.551e+00 | 0.0002 | 171.0 | 10.58% | 560.5 | 7.60% | motif file (matrix) | svg |
| 398 | G C T A C G T A A C T G C G T A C G A T A C G T A G T C A G C T | At3g12730(G2like)/colamp-At3g12730-DAP-Seq(GSE60143)/Homer | 1e-4 | -9.502e+00 | 0.0002 | 1145.0 | 70.81% | 4858.4 | 65.91% | motif file (matrix) | svg |
| 399 | T G A C G C T A T C G A T G C A A G T C A G T C C G T A A G T C C G T A C T G A G C T A G T A C | RUNX2(Runt)/PCa-RUNX2-ChIP-Seq(GSE33889)/Homer | 1e-4 | -9.475e+00 | 0.0002 | 640.0 | 39.58% | 2546.7 | 34.55% | motif file (matrix) | svg |
| 400 | C G T A G C A T C A G T C T A G A G T C A C T G A C T G G T A C A C T G A T C G | ERF115(AP2EREBP)/colamp-ERF115-DAP-Seq(GSE60143)/Homer | 1e-4 | -9.462e+00 | 0.0002 | 1260.0 | 77.92% | 5410.8 | 73.40% | motif file (matrix) | svg |
| 401 | C T G A A C T G C G T A A C G T G T C A A G C T A G C T G A C T G A C T C A G T | CCA(Myb)/Arabidopsis-CCA.GFP-ChIP-Seq(GSE70533)/Homer | 1e-4 | -9.419e+00 | 0.0002 | 862.0 | 53.31% | 3545.1 | 48.09% | motif file (matrix) | svg |
| 402 | C G T A A C T G G T C A A C G T A T C G C A G T C T A G T C A G C G T A A C T G C G T A A C G T C G T A C T G A T A C G | GATA3(Zf),DR4/iTreg-Gata3-ChIP-Seq(GSE20898)/Homer | 1e-4 | -9.360e+00 | 0.0002 | 140.0 | 8.66% | 442.7 | 6.01% | motif file (matrix) | svg |
| 403 | G C A T G C T A G C T A C G T A G C A T G C T A C T A G C G T A C G T A A C T G C G T A C G A T A C G T A G T C G A C T | At1g68670(G2like)/colamp-At1g68670-DAP-Seq(GSE60143)/Homer | 1e-4 | -9.353e+00 | 0.0002 | 487.0 | 30.12% | 1879.8 | 25.50% | motif file (matrix) | svg |
| 404 | C G T A C G A T C T A G G T C A G A C T C G A T C T A G C G T A A C G T C A T G | LIN-39(Homeobox)/cElegans.L3-LIN39-ChIP-Seq(modEncode)/Homer | 1e-4 | -9.340e+00 | 0.0002 | 904.0 | 55.91% | 3739.8 | 50.74% | motif file (matrix) | svg |
| 405 | T C G A C T A G A G T C A G T C C G T A C G T A A C G T T A G C T C A G T A C G | NFY(CCAAT)/Promoter/Homer | 1e-4 | -9.256e+00 | 0.0002 | 986.0 | 60.98% | 4119.8 | 55.89% | motif file (matrix) | svg |
| 406 | G C T A C G T A A C G T C A T G C G T A A C G T A C G T C T A G | ATHB5(HB)/colamp-ATHB5-DAP-Seq(GSE60143)/Homer | 1e-4 | -9.244e+00 | 0.0002 | 863.0 | 53.37% | 3554.3 | 48.22% | motif file (matrix) | svg |
| 407 | G C T A C G T A A C G T A T C G C G T A A C G T A C G T C T A G | ATHB6(Homeobox)/col-ATHB6-DAP-Seq(GSE60143)/Homer | 1e-3 | -9.200e+00 | 0.0003 | 1060.0 | 65.55% | 4465.8 | 60.58% | motif file (matrix) | svg |
| 408 | C T A G T A C G G A C T T G C A T G C A C G A T T A C G C T G A T C G A C T G A | Hoxa10(Homeobox)/ChickenMSG-Hoxa10.Flag-ChIP-Seq(GSE86088)/Homer | 1e-3 | -9.123e+00 | 0.0003 | 544.0 | 33.64% | 2132.3 | 28.93% | motif file (matrix) | svg |
| 409 | C T A G T A C G G A T C G T A C G C T A A G C T A G C T G T C A T C G A T A G C | Nanog(Homeobox)/mES-Nanog-ChIP-Seq(GSE11724)/Homer | 1e-3 | -9.112e+00 | 0.0003 | 1472.0 | 91.03% | 6474.6 | 87.84% | motif file (matrix) | svg |
| 410 | C G T A C G T A C G A T A C T G C A G T A G T C A C T G A C T G A G C T A C T G | DREB19(AP2EREBP)/colamp-DREB19-DAP-Seq(GSE60143)/Homer | 1e-3 | -8.973e+00 | 0.0003 | 879.0 | 54.36% | 3634.2 | 49.30% | motif file (matrix) | svg |
| 411 | T A C G A C T G A G C T G T A C C G T A T C G A C T G A A C T G C A T G A C G T A G T C C G T A | COUP-TFII(NR)/K562-NR2F1-ChIP-Seq(Encode)/Homer | 1e-3 | -8.965e+00 | 0.0003 | 935.0 | 57.82% | 3891.6 | 52.80% | motif file (matrix) | svg |
| 412 | C G T A C T A G T C G A A C G T C G A T G T A C C T G A T C G A G C A T A C T G G C A T C G T A G A C T G A T C G C A T | EIN3(EIL)/col-EIN3-DAP-Seq(GSE60143)/Homer | 1e-3 | -8.915e+00 | 0.0003 | 180.0 | 11.13% | 604.5 | 8.20% | motif file (matrix) | svg |
| 413 | A T G C C G T A C G T A C G T A C G T A C G T A A C T G A C G T C G A T C T G A | dof43(C2C2dof)/colamp-dof43-DAP-Seq(GSE60143)/Homer | 1e-3 | -8.885e+00 | 0.0004 | 974.0 | 60.24% | 4073.9 | 55.27% | motif file (matrix) | svg |
| 414 | C A T G C T A G A G T C A C T G A C T G G T A C C A T G T A C G | AT1G28160(AP2EREBP)/colamp-AT1G28160-DAP-Seq(GSE60143)/Homer | 1e-3 | -8.883e+00 | 0.0004 | 1204.0 | 74.46% | 5154.5 | 69.93% | motif file (matrix) | svg |
| 415 | T C G A A C T G A C T G C G T A C G T A T C G A A G T C C T G A A T C G G T A C G C A T C A T G | ETS:E-box(ETS,bHLH)/HPC7-Scl-ChIP-Seq(GSE22178)/Homer | 1e-3 | -8.826e+00 | 0.0004 | 62.0 | 3.83% | 160.7 | 2.18% | motif file (matrix) | svg |
| 416 | G A T C G T A C C G T A A G C T G A C T G C T A C T G A A C G T G A T C G C T A | Hoxc6(Homeobox)/EB-Hoxc6.iFlag-ChIP-Seq(GSE142377)/Homer | 1e-3 | -8.774e+00 | 0.0004 | 1352.0 | 83.61% | 5875.5 | 79.71% | motif file (matrix) | svg |
| 417 | G A C T G A T C C T G A A G T C A G T C A C T G C G T A A G T C G T A C G C T A G C A T C G A T | At1g19210(AP2EREBP)/colamp-At1g19210-DAP-Seq(GSE60143)/Homer | 1e-3 | -8.583e+00 | 0.0005 | 1268.0 | 78.42% | 5470.0 | 74.21% | motif file (matrix) | svg |
| 418 | G T A C G T C A G T A C G T C A G T A C G T C A G T A C G T C A G T A C G T C A | SeqBias: CA-repeat | 1e-3 | -8.521e+00 | 0.0005 | 1410.0 | 87.20% | 6169.7 | 83.70% | motif file (matrix) | svg |
| 419 | A G T C T G A C C T G A A G T C A G T C A C T G C G T A A G T C G T C A G C T A G C A T C G T A G C A T G C T A C G T A | DEAR3(AP2EREBP)/colamp-DEAR3-DAP-Seq(GSE60143)/Homer | 1e-3 | -8.403e+00 | 0.0006 | 709.0 | 43.85% | 2880.0 | 39.07% | motif file (matrix) | svg |
| 420 | C T A G A C T G A C G T C G T A A C T G A C T G A C G T T C A G C T G A T C G A | MYB107(MYB)/col-MYB107-DAP-Seq(GSE60143)/Homer | 1e-3 | -8.366e+00 | 0.0006 | 1165.0 | 72.05% | 4981.2 | 67.58% | motif file (matrix) | svg |
| 421 | C A G T A G C T C G T A G C A T A G T C G A C T C T A G C T A G C A G T C T A G T C G A T G C A C T A G C A T G G A C T | STOP1(C2H2)/colamp-STOP1-DAP-Seq(GSE60143)/Homer | 1e-3 | -8.338e+00 | 0.0006 | 404.0 | 24.98% | 1544.6 | 20.95% | motif file (matrix) | svg |
| 422 | G T C A C T G A T C A G G A T C T G C A T G C A A C G T T C A G C G T A C G T A C G T A G C T A | Hoxd12(Homeobox)/ChickenMSG-Hoxd12.Flag-ChIP-Seq(GSE86088)/Homer | 1e-3 | -8.268e+00 | 0.0006 | 1092.0 | 67.53% | 4639.6 | 62.94% | motif file (matrix) | svg |
| 423 | C T A G A C T G A C G T C G T A A C T G C A T G G C A T T C A G | MYB92(MYB)/colamp-MYB92-DAP-Seq(GSE60143)/Homer | 1e-3 | -8.210e+00 | 0.0007 | 913.0 | 56.46% | 3810.2 | 51.69% | motif file (matrix) | svg |
| 424 | C T A G A G T C A G C T A C T G C G T A C A G T C G T A C T G A T A G C T G A C | Unknown5/Drosophila-Promoters/Homer | 1e-3 | -8.194e+00 | 0.0007 | 842.0 | 52.07% | 3486.4 | 47.30% | motif file (matrix) | svg |
| 425 | G C T A G C T A C T G A A C T G A C G T A G T C C G T A C G T A G T A C A C T G A T G C G C A T | WRKY47(WRKY)/colamp-WRKY47-DAP-Seq(GSE60143)/Homer | 1e-3 | -8.193e+00 | 0.0007 | 424.0 | 26.22% | 1633.7 | 22.16% | motif file (matrix) | svg |
| 426 | A T G C A G T C C T G A A G T C C G A T A C G T A G T C A G T C A C G T A T C G G A C T A C G T | Etv2(ETS)/ES-ER71-ChIP-Seq(GSE59402)/Homer | 1e-3 | -8.170e+00 | 0.0007 | 555.0 | 34.32% | 2203.6 | 29.90% | motif file (matrix) | svg |
| 427 | C G A T C A G T G T C A G C T A C A G T A G C T A C G T A C T G A G T C C G T A A C G T A C T G A C G T C T G A T C G A | FUS3(ABI3VP1)/col-FUS3-DAP-Seq(GSE60143)/Homer | 1e-3 | -8.122e+00 | 0.0007 | 820.0 | 50.71% | 3388.8 | 45.97% | motif file (matrix) | svg |
| 428 | C G A T C T A G T C A G C A G T C G T A A G T C G C T A A C G T G A C T A T G C A G T C G C T A | PRDM10(Zf)/HEK293-PRDM10.eGFP-ChIP-Seq(Encode)/Homer | 1e-3 | -8.106e+00 | 0.0007 | 477.0 | 29.50% | 1865.0 | 25.30% | motif file (matrix) | svg |
| 429 | C G T A A C T G C G T A A C G T C A G T A G T C A G C T G C A T G C T A C G A T | At2g01060(G2like)/colamp-At2g01060-DAP-Seq(GSE60143)/Homer | 1e-3 | -8.086e+00 | 0.0008 | 1419.0 | 87.76% | 6223.7 | 84.43% | motif file (matrix) | svg |
| 430 | G A C T A G T C G A T C C G T A G T A C A G T C G C A T C G T A G T C A G A T C | MYB67(MYB)/col-MYB67-DAP-Seq(GSE60143)/Homer | 1e-3 | -8.031e+00 | 0.0008 | 935.0 | 57.82% | 3916.5 | 53.13% | motif file (matrix) | svg |
| 431 | C A T G A T G C T A G C C T G A A G T C A G T C A C T G G C T A A G T C G T A C G C T A G C A T | At4g28140(AP2EREBP)/colamp-At4g28140-DAP-Seq(GSE60143)/Homer | 1e-3 | -8.005e+00 | 0.0008 | 547.0 | 33.83% | 2172.8 | 29.48% | motif file (matrix) | svg |
| 432 | A T G C G A C T A C T G C A G T G A T C A C G T T A C G T A C G | Smad2(MAD)/ES-SMAD2-ChIP-Seq(GSE29422)/Homer | 1e-3 | -8.002e+00 | 0.0008 | 1067.0 | 65.99% | 4529.7 | 61.45% | motif file (matrix) | svg |
| 433 | A C T G G A T C C T G A A T C G A G T C T A G C C T G A C G T A T A C G A G T C C T A G C A G T T C A G T C G A T G A C G A T C | PAX5(Paired,Homeobox)/GM12878-PAX5-ChIP-Seq(GSE32465)/Homer | 1e-3 | -7.980e+00 | 0.0008 | 418.0 | 25.85% | 1612.3 | 21.87% | motif file (matrix) | svg |
| 434 | T C G A C G T A A G T C A G C T C G T A A G T C T C G A G C T A G A C T C G A T A G T C A G T C A G T C C T G A T C A G T G C A T C G A C A G T A T C G A G T C | GFY-Staf(?,Zf)/Promoter/Homer | 1e-3 | -7.945e+00 | 0.0009 | 47.0 | 2.91% | 115.4 | 1.57% | motif file (matrix) | svg |
| 435 | G A T C G T A C C T G A A G T C A G T C A C T G G C T A G T A C G T C A G C A T G C A T C G A T | DEAR2(AP2EREBP)/colamp-DEAR2-DAP-Seq(GSE60143)/Homer | 1e-3 | -7.938e+00 | 0.0009 | 1177.0 | 72.79% | 5049.5 | 68.50% | motif file (matrix) | svg |
| 436 | C T G A A T C G A G C T A G C T A C G T T A G C C T G A T A C G C G A T A C G T G A C T A G T C | ISRE(IRF)/ThioMac-LPS-Expression(GSE23622)/Homer | 1e-3 | -7.891e+00 | 0.0009 | 46.0 | 2.84% | 112.6 | 1.53% | motif file (matrix) | svg |
| 437 | T G A C A T G C C G T A A T C G A T G C C A G T C A T G A C T G A G T C G T A C | HEB(bHLH)/mES-Heb-ChIP-Seq(GSE53233)/Homer | 1e-3 | -7.780e+00 | 0.0010 | 940.0 | 58.13% | 3946.1 | 53.53% | motif file (matrix) | svg |
| 438 | A T G C C G T A A C T G C G T A A C G T G C T A T C G A A G C T C G A T C G T A A C G T A G T C C G A T A C T G G A T C | GATA(Zf),IR4/iTreg-Gata3-ChIP-Seq(GSE20898)/Homer | 1e-3 | -7.779e+00 | 0.0010 | 135.0 | 8.35% | 442.8 | 6.01% | motif file (matrix) | svg |
| 439 | C T G A G A C T G A T C C T G A A G T C G C A T A C G T G A C T G C T A G C A T | OBP1(C2C2dof)/col-OBP1-DAP-Seq(GSE60143)/Homer | 1e-3 | -7.734e+00 | 0.0010 | 1289.0 | 79.72% | 5591.7 | 75.86% | motif file (matrix) | svg |
| 440 | C G T A C G T A T C G A C G T A C G A T C G T A A C G T A G T C G C A T G C A T | At3g09600(MYBrelated)/colamp-At3g09600-DAP-Seq(GSE60143)/Homer | 1e-3 | -7.717e+00 | 0.0011 | 434.0 | 26.84% | 1687.7 | 22.90% | motif file (matrix) | svg |
| 441 | C T A G T C G A C T G A C G T A T A C G G A C T T C A G T C G A G T C A T G C A T A C G A G C T | IRF2(IRF)/Erythroblas-IRF2-ChIP-Seq(GSE36985)/Homer | 1e-3 | -7.701e+00 | 0.0011 | 101.0 | 6.25% | 312.0 | 4.23% | motif file (matrix) | svg |
| 442 | C T G A A G T C C G A T A G C T A T G C G T A C A C G T A T C G C A G T G C A T | Elf4(ETS)/BMDM-Elf4-ChIP-Seq(GSE88699)/Homer | 1e-3 | -7.529e+00 | 0.0013 | 716.0 | 44.28% | 2935.2 | 39.82% | motif file (matrix) | svg |
| 443 | A G T C C G T A C G A T A G T C G T C A A G T C A C G T C T G A | Unknown2/Drosophila-Promoters/Homer | 1e-3 | -7.456e+00 | 0.0014 | 817.0 | 50.53% | 3393.6 | 46.04% | motif file (matrix) | svg |
| 444 | A G T C G T A C A G C T C T A G A G T C C G A T A C T G C G T A A C T G G T C A | Zic(Zf)/Cerebellum-ZIC1.2-ChIP-Seq(GSE60731)/Homer | 1e-3 | -7.330e+00 | 0.0015 | 597.0 | 36.92% | 2410.5 | 32.70% | motif file (matrix) | svg |
| 445 | A G C T G A T C A G C T G A C T G A C T T C G A A G T C C G T A A C T G T C A G | SpliceAcceptor/Homer | 1e-3 | -7.321e+00 | 0.0016 | 1427.0 | 88.25% | 6279.6 | 85.19% | motif file (matrix) | svg |
| 446 | A C T G T G A C A C T G A C G T A C G T A C T G C G T A A G T C A G C T C G A T G C A T A C G T | WRKY17(WRKY)/colamp-WRKY17-DAP-Seq(GSE60143)/Homer | 1e-3 | -7.234e+00 | 0.0017 | 31.0 | 1.92% | 67.4 | 0.91% | motif file (matrix) | svg |
| 447 | T A G C C T A G T C G A G A C T A C T G C T G A A G T C T C A G G C A T T G A C C T G A A G C T | Atf7(bZIP)/3T3L1-Atf7-ChIP-Seq(GSE56872)/Homer | 1e-3 | -7.229e+00 | 0.0017 | 547.0 | 33.83% | 2192.3 | 29.74% | motif file (matrix) | svg |
| 448 | G C T A T G A C G A T C C G A T G A C T A T G C C T G A A T C G G C A T A C G T | JGL(C2H2)/col-JGL-DAP-Seq(GSE60143)/Homer | 1e-3 | -7.188e+00 | 0.0018 | 1066.0 | 65.92% | 4547.2 | 61.69% | motif file (matrix) | svg |
| 449 | A T G C G A C T A G C T C T A G C G T A C T A G C G A T C T A G A T C G G A T C | Nkx2.2(Homeobox)/NPC-Nkx2.2-ChIP-Seq(GSE61673)/Homer | 1e-3 | -7.115e+00 | 0.0019 | 1166.0 | 72.11% | 5019.2 | 68.09% | motif file (matrix) | svg |
| 450 | C T A G T A C G A G T C C G T A A G T C A C G T A G T C T C G A C G T A T A C G | Nkx2.1(Homeobox)/LungAC-Nkx2.1-ChIP-Seq(GSE43252)/Homer | 1e-3 | -7.070e+00 | 0.0020 | 1311.0 | 81.08% | 5714.5 | 77.52% | motif file (matrix) | svg |
| 451 | T C G A T A G C T G C A A C T G A C T G C G T A C G T A C T A G G A C T T A C G | ETS1(ETS)/Jurkat-ETS1-ChIP-Seq(GSE17954)/Homer | 1e-3 | -7.059e+00 | 0.0020 | 705.0 | 43.60% | 2899.1 | 39.33% | motif file (matrix) | svg |
| 452 | G A T C G A T C A G T C G T A C C G A T G T A C G T A C A G T C A G T C A G T C G C T A G A T C | ZNF148(Zf)/MDAMB231-ZNF148-ChIP-Seq(GSE147020)/Homer | 1e-3 | -7.000e+00 | 0.0021 | 153.0 | 9.46% | 524.9 | 7.12% | motif file (matrix) | svg |
| 453 | C T G A C T A G T C G A C G T A A T G C C G T A A T C G C G A T T A G C G C A T A T C G G C A T A G C T G A T C G A C T A G C T | ARE(NR)/LNCAP-AR-ChIP-Seq(GSE27824)/Homer | 1e-3 | -6.932e+00 | 0.0023 | 182.0 | 11.26% | 642.1 | 8.71% | motif file (matrix) | svg |
| 454 | G T A C A C T G A C G T T C A G G C A T C G T A C G A T G C A T C G T A A G T C C G T A T G A C C A T G G A C T G C T A | ANAC083(NAC)/col-ANAC083-DAP-Seq(GSE60143)/Homer | 1e-3 | -6.922e+00 | 0.0023 | 870.0 | 53.80% | 3650.3 | 49.52% | motif file (matrix) | svg |
| 455 | C G T A C T A G G A C T G T C A G T C A C G T A A G T C C G T A T C G A T C G A T C G A C G T A C T G A C T A G G C T A C G T A T A G C C G T A C G A T C G T A | FOXA1:AR(Forkhead,NR)/LNCAP-AR-ChIP-Seq(GSE27824)/Homer | 1e-2 | -6.857e+00 | 0.0024 | 31.0 | 1.92% | 69.0 | 0.94% | motif file (matrix) | svg |
| 456 | T A C G T A C G C T A G T C A G A G T C C G T A A T C G A T G C A C G T A C T G A G T C G A C T | Ascl2(bHLH)/ESC-Ascl2-ChIP-Seq(GSE97712)/Homer | 1e-2 | -6.839e+00 | 0.0025 | 645.0 | 39.89% | 2637.4 | 35.78% | motif file (matrix) | svg |
| 457 | C T G A C T A G A T C G A G C T A C T G G A C T A G T C C T G A | Tbx5(T-box)/HL1-Tbx5.biotin-ChIP-Seq(GSE21529)/Homer | 1e-2 | -6.813e+00 | 0.0025 | 1302.0 | 80.52% | 5677.2 | 77.02% | motif file (matrix) | svg |
| 458 | T A C G T C G A G A C T A C T G C T G A A G T C T C A G G A C T T G A C C T G A | Atf1(bZIP)/K562-ATF1-ChIP-Seq(GSE31477)/Homer | 1e-2 | -6.792e+00 | 0.0026 | 816.0 | 50.46% | 3408.4 | 46.24% | motif file (matrix) | svg |
| 459 | G C A T A G T C G A C T T C G A T A C G G T C A T C G A A C T G T A G C G C A T G C A T A T G C | AT2G01818(PLATZ)/col-AT2G01818-DAP-Seq(GSE60143)/Homer | 1e-2 | -6.724e+00 | 0.0027 | 175.0 | 10.82% | 617.2 | 8.37% | motif file (matrix) | svg |
| 460 | T G A C C T G A A G T C A G T C A C T G G A T C G A C T G C A T | At5g18450(AP2EREBP)/col-At5g18450-DAP-Seq(GSE60143)/Homer | 1e-2 | -6.687e+00 | 0.0028 | 1181.0 | 73.04% | 5102.8 | 69.23% | motif file (matrix) | svg |
| 461 | C A T G G T C A A G T C C G T A C T A G G A T C C G A T A C T G A C G T G T A C C G T A C G T A | bZIP69(bZIP)/col-bZIP69-DAP-Seq(GSE60143)/Homer | 1e-2 | -6.641e+00 | 0.0030 | 96.0 | 5.94% | 305.9 | 4.15% | motif file (matrix) | svg |
| 462 | C G T A C T G A T C A G A C T G G T C A C G T A A C G T G T A C C G A T G C A T | AT5G45580(G2like)/colamp-AT5G45580-DAP-Seq(GSE60143)/Homer | 1e-2 | -6.574e+00 | 0.0032 | 1153.0 | 71.30% | 4973.1 | 67.47% | motif file (matrix) | svg |
| 463 | G A T C C T G A G A T C G A T C C T A G G C T A A G T C C T G A G C T A C G T A | At4g16750(AP2EREBP)/col-At4g16750-DAP-Seq(GSE60143)/Homer | 1e-2 | -6.398e+00 | 0.0038 | 1104.0 | 68.27% | 4748.1 | 64.41% | motif file (matrix) | svg |
| 464 | G C T A A C G T C T A G G T A C G C T A G A C T C T G A G C A T C A T G G A T C | Pit1(Homeobox)/GCrat-Pit1-ChIP-Seq(GSE58009)/Homer | 1e-2 | -6.347e+00 | 0.0040 | 851.0 | 52.63% | 3582.0 | 48.59% | motif file (matrix) | svg |
| 465 | C G A T T C G A A C T G G T C A C G T A C G A T G T A C G A C T | At3g04030(G2like)/col-At3g04030-DAP-Seq(GSE60143)/Homer | 1e-2 | -6.295e+00 | 0.0042 | 857.0 | 53.00% | 3610.1 | 48.98% | motif file (matrix) | svg |
| 466 | G C A T G A C T T G A C A G C T T C G A A C T G C G T A C T G A T C A G T A G C C G A T C G A T G T A C A G C T T C G A | AT1G23810(Orphan)/col-AT1G23810-DAP-Seq(GSE60143)/Homer | 1e-2 | -6.289e+00 | 0.0042 | 39.0 | 2.41% | 99.8 | 1.35% | motif file (matrix) | svg |
| 467 | T C G A C T G A C G A T C G T A C G T A C G T A C T A G A G C T C T G A T C A G | Adof1(C2C2dof)/col-Adof1-DAP-Seq(GSE60143)/Homer | 1e-2 | -6.287e+00 | 0.0042 | 1261.0 | 77.98% | 5494.4 | 74.54% | motif file (matrix) | svg |
| 468 | T C G A A G T C C G T A A T C G T A G C A C G T A C T G A G C T A C G T A G T C | Ptf1a(bHLH)/Panc1-Ptf1a-ChIP-Seq(GSE47459)/Homer | 1e-2 | -6.098e+00 | 0.0050 | 1114.0 | 68.89% | 4804.6 | 65.18% | motif file (matrix) | svg |
| 469 | T C A G A G C T A C G T A C G T G T A C G A T C C G T A C T A G C A T G G T C A C G T A T C G A | STAT4(Stat)/CD4-Stat4-ChIP-Seq(GSE22104)/Homer | 1e-2 | -6.081e+00 | 0.0051 | 611.0 | 37.79% | 2508.0 | 34.02% | motif file (matrix) | svg |
| 470 | C G A T G A T C T A C G C T G A G C T A C G T A G C A T A G T C C T A G C G T A G C A T C G A T | AT2G15740(C2H2)/col-AT2G15740-DAP-Seq(GSE60143)/Homer | 1e-2 | -6.077e+00 | 0.0051 | 1328.0 | 82.13% | 5823.0 | 79.00% | motif file (matrix) | svg |
| 471 | G C A T G C A T C T G A A C G T C T G A A C G T C G T A C G T A C G T A A G T C G T C A G T C A | Foxf1(Forkhead)/Lung-Foxf1-ChIP-Seq(GSE77951)/Homer | 1e-2 | -6.018e+00 | 0.0054 | 704.0 | 43.54% | 2927.0 | 39.71% | motif file (matrix) | svg |
| 472 | C T G A A G T C G A T C C A T G G C T A G A T C C T G A G C T A G C T A C G A T | AT1G77200(AP2EREBP)/colamp-AT1G77200-DAP-Seq(GSE60143)/Homer | 1e-2 | -5.929e+00 | 0.0059 | 1118.0 | 69.14% | 4828.6 | 65.51% | motif file (matrix) | svg |
| 473 | A T G C G C A T C G A T G A T C A G C T C T G A A C T G C G T A C G T A T C A G T G A C C G A T G C A T G A T C C G A T | HSF21(HSF)/col-HSF21-DAP-Seq(GSE60143)/Homer | 1e-2 | -5.909e+00 | 0.0060 | 88.0 | 5.44% | 284.0 | 3.85% | motif file (matrix) | svg |
| 474 | C G T A C T G A C T A G C T G A A G T C G C T A C G A T A T C G G A C T G A T C A G T C C T G A C T A G C T A G A G T C G C T A C G A T C T A G G A T C G A T C | p73(p53)/Trachea-p73-ChIP-Seq(PRJNA310161)/Homer | 1e-2 | -5.879e+00 | 0.0062 | 58.0 | 3.59% | 170.4 | 2.31% | motif file (matrix) | svg |
| 475 | T C G A T G A C G T A C C G T A A C G T G A C T A C G T A C T G A C T G A G C T | Mesp1(bHLH)/ESC-Mesp1-ChIP-Seq(GSE165102)/Homer | 1e-2 | -5.863e+00 | 0.0063 | 479.0 | 29.62% | 1931.2 | 26.20% | motif file (matrix) | svg |
| 476 | G C A T C T A G C T A G C G T A A G C T C G T A C T G A C A T G C T A G G C A T | AT5G56840(MYBrelated)/colamp-AT5G56840-DAP-Seq(GSE60143)/Homer | 1e-2 | -5.785e+00 | 0.0068 | 1071.0 | 66.23% | 4613.3 | 62.59% | motif file (matrix) | svg |
| 477 | G A C T G A T C A G T C C G T A T G A C A G T C G C A T C T G A G T A C G A T C G C A T G A C T | MYB10(MYB)/col-MYB10-DAP-Seq(GSE60143)/Homer | 1e-2 | -5.656e+00 | 0.0077 | 504.0 | 31.17% | 2047.8 | 27.78% | motif file (matrix) | svg |
| 478 | C T G A G A C T A C G T A G C T C T G A C G T A C G T A G A C T G T C A G A T C | AT2G20110(CPP)/colamp-AT2G20110-DAP-Seq(GSE60143)/Homer | 1e-2 | -5.631e+00 | 0.0079 | 920.0 | 56.90% | 3920.8 | 53.19% | motif file (matrix) | svg |
| 479 | T C A G A C T G A C G T C G T A A C T G A C T G A C G T C T A G | MYB51(MYB)/col-MYB51-DAP-Seq(GSE60143)/Homer | 1e-2 | -5.620e+00 | 0.0079 | 878.0 | 54.30% | 3728.1 | 50.58% | motif file (matrix) | svg |
| 480 | A T G C G T A C A C T G A G T C A G T C A C T G G A T C G T C A C G T A C G A T G C A T C G A T | RRTF1(AP2EREBP)/colamp-RRTF1-DAP-Seq(GSE60143)/Homer | 1e-2 | -5.605e+00 | 0.0080 | 587.0 | 36.30% | 2416.9 | 32.79% | motif file (matrix) | svg |
| 481 | T C A G C T G A C T A G C A T G A C G T A T G C C T G A C T G A C T G A C T A G C A T G A C G T A T G C C T G A | TR4(NR),DR1/Hela-TR4-ChIP-Seq(GSE24685)/Homer | 1e-2 | -5.562e+00 | 0.0084 | 66.0 | 4.08% | 203.4 | 2.76% | motif file (matrix) | svg |
| 482 | C T G A T A C G G C A T C T A G A T G C G A T C C G A T A C T G C T A G G A T C C T G A A T G C | MYRF(MYRF)/CFPAC1-MYRF-ChIP-Seq(GSE145627)/Homer | 1e-2 | -5.537e+00 | 0.0086 | 270.0 | 16.70% | 1036.5 | 14.06% | motif file (matrix) | svg |
| 483 | C G T A C G T A G C T A C G T A G A T C C T G A A C G T A C G T A G T C A G C T G C A T G C A T | AT2G40260(G2like)/colamp-AT2G40260-DAP-Seq(GSE60143)/Homer | 1e-2 | -5.523e+00 | 0.0087 | 1033.0 | 63.88% | 4445.9 | 60.31% | motif file (matrix) | svg |
| 484 | C A T G C T G A A G T C A C T G A C T G A G C T A C T G A T C G | ESE3(AP2EREBP)/col-ESE3-DAP-Seq(GSE60143)/Homer | 1e-2 | -5.488e+00 | 0.0090 | 1103.0 | 68.21% | 4773.0 | 64.75% | motif file (matrix) | svg |
| 485 | C G A T C T A G T C G A A G C T C G A T C T G A C G T A A G C T A C T G C T A G A T G C G A T C | Hoxb4(Homeobox)/ES-Hoxb4-ChIP-Seq(GSE34014)/Homer | 1e-2 | -5.466e+00 | 0.0091 | 283.0 | 17.50% | 1093.8 | 14.84% | motif file (matrix) | svg |
| 486 | A C T G G A T C G A C T A C T G A C G T C A T G A C T G A C G T A G C T C G A T | RUNX-AML(Runt)/CD4+-PolII-ChIP-Seq(Barski\_et\_al.)/Homer | 1e-2 | -5.460e+00 | 0.0092 | 509.0 | 31.48% | 2075.0 | 28.15% | motif file (matrix) | svg |
| 487 | T C G A T C A G T C G A A C T G C A T G A C G T A G T C C T G A | COUP-TFII(NR)/Artia-Nr2f2-ChIP-Seq(GSE46497)/Homer | 1e-2 | -5.458e+00 | 0.0092 | 1039.0 | 64.25% | 4475.4 | 60.71% | motif file (matrix) | svg |
| 488 | C T A G T A C G G A T C G T C A T G C A A C G T T G C A G C T A T C G A T G C A | Hoxa9(Homeobox)/ChickenMSG-Hoxa9.Flag-ChIP-Seq(GSE86088)/Homer | 1e-2 | -5.458e+00 | 0.0092 | 1302.0 | 80.52% | 5714.8 | 77.53% | motif file (matrix) | svg |
| 489 | T A G C A G T C T G A C A G T C C T A G A T C G A G T C C A T G T G A C A G T C G T A C A G T C A G T C G C A T C T A G A T C G G C A T A C T G A T C G G A T C | BORIS(Zf)/K562-CTCFL-ChIP-Seq(GSE32465)/Homer | 1e-2 | -5.287e+00 | 0.0109 | 87.0 | 5.38% | 288.0 | 3.91% | motif file (matrix) | svg |
| 490 | G C A T C T A G A C T G A C G T C G T A A C T G A C T G C G A T C T A G T C G A T C G A G C T A | MYB40(MYB)/col-MYB40-DAP-Seq(GSE60143)/Homer | 1e-2 | -5.262e+00 | 0.0111 | 433.0 | 26.78% | 1747.8 | 23.71% | motif file (matrix) | svg |
| 491 | A C G T C T A G C G T A A G T C G T A C A C G T A C G T A C G T G T C A G T A C T G A C G A C T | Nur77(NR)/K562-NR4A1-ChIP-Seq(GSE31363)/Homer | 1e-2 | -5.225e+00 | 0.0115 | 189.0 | 11.69% | 702.6 | 9.53% | motif file (matrix) | svg |
| 492 | T G C A A G C T A C G T C T A G G A T C C T A G G A T C G T C A C T G A A G T C | CEBP(bZIP)/ThioMac-CEBPb-ChIP-Seq(GSE21512)/Homer | 1e-2 | -5.184e+00 | 0.0120 | 806.0 | 49.85% | 3415.9 | 46.34% | motif file (matrix) | svg |
| 493 | T C A G A C T G C A G T A G T C A G T C G T C A C G T A C G T A A C T G C A G T A G T C A G T C C T G A T G C A A G C T | dHNF4(NR)/Fly-HNF4-ChIP-Seq(GSE73675)/Homer | 1e-2 | -5.153e+00 | 0.0123 | 55.0 | 3.40% | 166.1 | 2.25% | motif file (matrix) | svg |
| 494 | G A T C G C T A A G C T G C A T A G T C C T G A T A G C G A C T | STZ(C2H2)/colamp-STZ-DAP-Seq(GSE60143)/Homer | 1e-2 | -5.100e+00 | 0.0129 | 1486.0 | 91.90% | 6623.2 | 89.85% | motif file (matrix) | svg |
| 495 | G C A T G C T A C G T A A G C T G C T A T G C A A G T C A C G T A C G T A C G T G C A T G C A T | At4g38000(C2C2dof)/col-At4g38000-DAP-Seq(GSE60143)/Homer | 1e-2 | -5.099e+00 | 0.0129 | 763.0 | 47.19% | 3223.3 | 43.73% | motif file (matrix) | svg |
| 496 | C A G T A G C T G A C T T G C A A G T C A G C T A C G T A C G T C G A T G A C T | AT3G52440(C2C2dof)/colamp-AT3G52440-DAP-Seq(GSE60143)/Homer | 1e-2 | -5.084e+00 | 0.0131 | 1178.0 | 72.85% | 5137.7 | 69.70% | motif file (matrix) | svg |
| 497 | C T A G G A C T G A T C A C G T A T C G A G C T C T G A A T C G C G A T C T A G G A T C G A C T C A T G A T C G G T A C G A C T A G T C G C A T A G C T C G A T | ZNF382(Zf)/HEK293-ZNF382.GFP-ChIP-Seq(GSE58341)/Homer | 1e-2 | -5.076e+00 | 0.0132 | 31.0 | 1.92% | 80.9 | 1.10% | motif file (matrix) | svg |
| 498 | G T A C C A T G T A G C A G T C C T A G C A T G C T G A C G T A G C A T G C A T A C G T G C A T G T A C A C T G A T C G | LOB(LOBAS2)/col-LOB-DAP-Seq(GSE60143)/Homer | 1e-2 | -5.059e+00 | 0.0134 | 476.0 | 29.44% | 1943.0 | 26.36% | motif file (matrix) | svg |
| 499 | C T G A C T G A G C A T G A C T A G T C T C G A C T A G C G T A A C G T G A T C A G C T T C A G | GATA4(C2C2gata)/col-GATA4-DAP-Seq(GSE60143)/Homer | 1e-2 | -5.017e+00 | 0.0139 | 846.0 | 52.32% | 3603.3 | 48.88% | motif file (matrix) | svg |
| 500 | G C T A G A C T A G T C T C G A T C A G T C G A A C G T A G T C G A C T T C A G | GATA14(C2C2gata)/col-GATA14-DAP-Seq(GSE60143)/Homer | 1e-2 | -5.004e+00 | 0.0141 | 747.0 | 46.20% | 3154.3 | 42.79% | motif file (matrix) | svg |
| 501 | C G T A G A C T C G A T A T C G G T A C G C A T C A T G C G T A T A C G G C A T G T A C C G T A C A T G A T G C G C T A C T A G G C A T G C A T G C A T G A C T | MafB(bZIP)/BMM-Mafb-ChIP-Seq(GSE75722)/Homer | 1e-2 | -4.987e+00 | 0.0143 | 287.0 | 17.75% | 1122.2 | 15.22% | motif file (matrix) | svg |
| 502 | C G T A C T G A C G T A G C A T G A C T A G C T T C G A C T A G C G T A A C G T A G T C A G C T C T G A C T G A C G T A | GATA1(C2C2gata)/colamp-GATA1-DAP-Seq(GSE60143)/Homer | 1e-2 | -4.925e+00 | 0.0152 | 468.0 | 28.94% | 1911.0 | 25.93% | motif file (matrix) | svg |
| 503 | T C G A C A T G C A T G A C G T A T G C T C G A C T G A A G C T T A C G T G C A G T A C G A T C A G C T A G T C | FXR(NR),IR1/Liver-FXR-ChIP-Seq(Chong\_et\_al.)/Homer | 1e-2 | -4.900e+00 | 0.0155 | 356.0 | 22.02% | 1422.0 | 19.29% | motif file (matrix) | svg |
| 504 | T C A G T G A C G T A C T G C A G T A C C T A G G T A C A T G C A G T C G T C A A G T C G A C T | Klf9(Zf)/GBM-Klf9-ChIP-Seq(GSE62211)/Homer | 1e-2 | -4.885e+00 | 0.0157 | 157.0 | 9.71% | 576.9 | 7.83% | motif file (matrix) | svg |
| 505 | C T G A C T A G A T C G G C A T A C T G G T A C A T G C C G T A A C T G G C T A A G T C C G T A | Tbox:Smad(T-box,MAD)/ESCd5-Smad2\_3-ChIP-Seq(GSE29422)/Homer | 1e-2 | -4.840e+00 | 0.0164 | 158.0 | 9.77% | 582.0 | 7.90% | motif file (matrix) | svg |
| 506 | G C A T C G T A G C T A G A C T C G A T G A C T A G T C C A G T A G T C A G T C A C T G C T A G G T A C C T A G C T G A | AT5G05550(Trihelix)/col-AT5G05550-DAP-Seq(GSE60143)/Homer | 1e-2 | -4.809e+00 | 0.0169 | 1224.0 | 75.70% | 5363.7 | 72.77% | motif file (matrix) | svg |
| 507 | T C G A G C A T A T G C C T G A A T G C T A G C A G T C G T A C T C G A A G C T | Srebp1a(bHLH)/HepG2-Srebp1a-ChIP-Seq(GSE31477)/Homer | 1e-2 | -4.763e+00 | 0.0177 | 236.0 | 14.59% | 910.5 | 12.35% | motif file (matrix) | svg |
| 508 | A G C T G C A T G T C A C G A T T A G C C G T A A C G T G C T A | CRC(C2C2YABBY)/col-CRC-DAP-Seq(GSE60143)/Homer | 1e-2 | -4.750e+00 | 0.0179 | 1072.0 | 66.30% | 4653.8 | 63.14% | motif file (matrix) | svg |
| 509 | T G C A C T G A A T G C G T C A A C T G A C T G C G T A C G T A C T A G A G C T | Ets1-distal(ETS)/CD4+-PolII-ChIP-Seq(Barski\_et\_al.)/Homer | 1e-2 | -4.705e+00 | 0.0187 | 184.0 | 11.38% | 692.5 | 9.39% | motif file (matrix) | svg |
| 510 | G A C T C T A G C T A G C T A G A C T G T C G A C T G A C T A G C T A G C T A G G T A C G T C A | ZNF467(Zf)/HEK293-ZNF467.GFP-ChIP-Seq(GSE58341)/Homer | 1e-2 | -4.681e+00 | 0.0191 | 355.0 | 21.95% | 1424.1 | 19.32% | motif file (matrix) | svg |
| 511 | C G T A C T G A C T A G C G T A C G T A A G T C C G T A C A G T G C A T G T C A C G A T A C T G A C G T G C A T G A T C | PGR(NR)/EndoStromal-PGR-ChIP-Seq(GSE69539)/Homer | 1e-2 | -4.651e+00 | 0.0196 | 212.0 | 13.11% | 811.5 | 11.01% | motif file (matrix) | svg |
| 512 | T A C G T A G C C A T G C A G T A C G T C T A G C G T A A G T C G A C T G C A T G C A T C A G T | WRKY11(WRKY)/col-WRKY11-DAP-Seq(GSE60143)/Homer | 1e-2 | -4.621e+00 | 0.0202 | 185.0 | 11.44% | 698.7 | 9.48% | motif file (matrix) | svg |
