## Supplemental dataset for "Hybrid CNN and Multi-Head Attention Model for Analyzing Epigenetic Mechanisms and Gene Expression Across Fungal Phylogenetic Distances": NcrassaModel_FgramTest_K36me3locs_homerResults.html

/projects/wg-feeds/SHAP/NcrassaModel\_FgramTest\_K36me3locs\_SHAP\_noDup\_HOMER// - Homer de novo Motif Results


### Homer *de novo* Motif Results (/projects/wg-feeds/SHAP/NcrassaModel\_FgramTest\_K36me3locs\_SHAP\_noDup\_HOMER//)

Non-redundant Motif File of Results  
Known Motif Enrichment Results  
Gene Ontology Enrichment Results  
If Homer is having trouble matching a motif to a known motif, try copy/pasting the matrix file into
STAMP  
More information on motif finding results: HOMER
| Description of Results
| Tips
  
Total target sequences = 28516  
Total background sequences = 131374  
\* - possible false positive  

|  |  |  |  |  |  |  |  |  |
| --- | --- | --- | --- | --- | --- | --- | --- | --- |
| Rank | Motif | P-value | log P-pvalue | % of Targets | % of Background | STD(Bg STD) | Best Match/Details | Motif File |
| 1 | G A T C C T G A G C A T T A G C C T G A G C A T T A G C C T G A G C A T T A G C T C G A G C T A | 1e-2026 | -4.666e+03 | 48.73% | 19.93% | 707.2bp (680.1bp) | ZML2(C2C2gata)/col-ZML2-DAP-Seq(GSE60143)/Homer(0.861) More Information \| Similar Motifs Found | motif file (matrix) |
| 2 | T C A G G A T C T G A C T C G A G C T A C A T G T C A G G A T C G T A C T C G A C G T A T A C G | 1e-1710 | -3.939e+03 | 55.60% | 27.65% | 699.2bp (682.9bp) | SF1(NR)/H295R-Nr5a1-ChIP-Seq(GSE44220)/Homer(0.815) More Information \| Similar Motifs Found | motif file (matrix) |
| 3 | T C A G G C T A A T C G T C A G G C T A A T C G T C A G G C T A A T C G T C A G G C T A A T C G | 1e-1448 | -3.336e+03 | 55.45% | 29.56% | 682.2bp (660.3bp) | TF3A(C2H2)/col-TF3A-DAP-Seq(GSE60143)/Homer(0.668) More Information \| Similar Motifs Found | motif file (matrix) |
| 4 | C A G T A T G C C T G A C T A G G A T C T G A C C G T A G C A T A T C G C G T A G A C T T A C G | 1e-1218 | -2.806e+03 | 59.99% | 35.79% | 724.3bp (698.3bp) | Tv\_0259(RRM)/Trichomonas\_vaginalis-RNCMPT00259-PBM/HughesRNA(0.719) More Information \| Similar Motifs Found | motif file (matrix) |
| 5 | G C A T A G C T A C G T A C G T A G T C C A T G G C A T A T G C C A G T A G C T A C G T C A T G | 1e-1191 | -2.744e+03 | 54.16% | 30.63% | 727.1bp (683.4bp) | Unknown4/Arabidopsis-Promoters/Homer(0.777) More Information \| Similar Motifs Found | motif file (matrix) |
| 6 | G T A C A G C T A G T C T C A G C T G A T A C G T C G A C G T A A T C G G T A C C A G T A G T C | 1e-1132 | -2.607e+03 | 42.01% | 20.79% | 716.0bp (698.5bp) | XBP1/MA0414.2/Jaspar(0.720) More Information \| Similar Motifs Found | motif file (matrix) |
| 7 | T A G C G T C A C A G T T G A C A C G T A G C T A T C G G T C A C T A G T C A G | 1e-1127 | -2.596e+03 | 56.50% | 33.39% | 750.0bp (698.7bp) | MATR3(RRM)/Homo\_sapiens-RNCMPT00037-PBM/HughesRNA(0.745) More Information \| Similar Motifs Found | motif file (matrix) |
| 8 | A T G C T G A C C G T A C A G T A T C G A C G T A G T C G C T A C A G T A T G C T G C A G T C A | 1e-965 | -2.223e+03 | 52.18% | 31.04% | 751.9bp (690.5bp) | pros/dmmpmm(Bergman)/fly(0.670) More Information \| Similar Motifs Found | motif file (matrix) |
| 9 | C A T G G T A C G T A C G C A T A C G T A C G T A C T G G A C T A T G C T C G A C G T A C A T G | 1e-915 | -2.109e+03 | 70.42% | 49.59% | 747.0bp (697.6bp) | Sox3(HMG)/NPC-Sox3-ChIP-Seq(GSE33059)/Homer(0.672) More Information \| Similar Motifs Found | motif file (matrix) |
| 10 | A G T C G A T C T C A G C G T A A C T G T A G C G A T C A C G T A T C G C A T G C A T G T A C G | 1e-849 | -1.955e+03 | 55.59% | 35.45% | 754.7bp (686.6bp) | FEZF2/MA2341.1/Jaspar(0.729) More Information \| Similar Motifs Found | motif file (matrix) |
| 11 | G T A C T G A C G T C A C T G A T A G C G T A C C G T A T C G A T A G C G T C A G T C A G T C A | 1e-748 | -1.724e+03 | 50.60% | 31.97% | 702.5bp (676.6bp) | MYB30/MA1768.2/Jaspar(0.702) More Information \| Similar Motifs Found | motif file (matrix) |
| 12 | T G A C C T G A C G T A A T G C C G T A A G C T A G C T A T C G | 1e-610 | -1.406e+03 | 53.31% | 36.24% | 740.8bp (693.9bp) | ZIPIC/MA2322.1/Jaspar(0.651) More Information \| Similar Motifs Found | motif file (matrix) |
| 13 | G A T C G A C T A G C T A T C G C G T A G A T C C G A T A G C T A C T G G C T A A G T C G C A T | 1e-489 | -1.128e+03 | 42.55% | 27.96% | 757.9bp (694.1bp) | WRKY40/MA1085.3/Jaspar(0.821) More Information \| Similar Motifs Found | motif file (matrix) |
| 14 | A T G C A G C T A G C T A C G T A G T C A G T C C G T A C T A G | 1e-431 | -9.942e+02 | 49.75% | 35.48% | 767.4bp (679.6bp) | Nfatc1/MA0624.3/Jaspar(0.900) More Information \| Similar Motifs Found | motif file (matrix) |
| 15 | C G T A C G T A C T G A C G T A C G T A G C T A C T G A G C A T T C G A G C T A C G T A G C T A | 1e-383 | -8.835e+02 | 27.49% | 16.43% | 749.1bp (677.8bp) | br-Z1/dmmpmm(Down)/fly(0.739) More Information \| Similar Motifs Found | motif file (matrix) |
| 16 | A G T C G T C A C T A G G T C A C G T A T A C G A G T C A T G C | 1e-261 | -6.022e+02 | 18.42% | 10.70% | 725.5bp (669.6bp) | OSR2/MA1646.2/Jaspar(0.889) More Information \| Similar Motifs Found | motif file (matrix) |
| 17 | A C G T A C T G C G A T C T G A A C G T A T C G C G A T C T G A A C G T A C T G | 1e-166 | -3.839e+02 | 26.71% | 19.25% | 776.9bp (684.6bp) | SeqBias: CA-repeat(0.886) More Information \| Similar Motifs Found | motif file (matrix) |
| 18 | T A G C A G C T C G T A T A G C T A G C A C G T C T G A A G T C G A T C G A C T C G T A A G C T | 1e-166 | -3.830e+02 | 2.86% | 0.72% | 663.8bp (695.2bp) | PK06182.1/MA2354.1/Jaspar(0.874) More Information \| Similar Motifs Found | motif file (matrix) |
| 19 | G A C T A C G T A C G T T G A C A G C T T C A G G T C A A G C T C T G A G A T C A G T C A G T C | 1e-145 | -3.353e+02 | 0.97% | 0.05% | 590.5bp (607.4bp) | PB0059.1\_Six6\_1/Jaspar(0.738) More Information \| Similar Motifs Found | motif file (matrix) |
| 20 | C G T A A C T G C G T A A C T G C G T A A C T G C G T A A C T G C G T A A C T G | 1e-127 | -2.930e+02 | 7.93% | 4.31% | 774.5bp (663.2bp) | Trl/dmmpmm(Down)/fly(0.905) More Information \| Similar Motifs Found | motif file (matrix) |
| 21 | A T G C C G A T C T A G C G A T C T G A A G T C A G C T T G A C A C G T A C T G A C G T T C G A | 1e-85 | -1.967e+02 | 1.32% | 0.30% | 740.1bp (672.0bp) | Dmrt1/MA1603.2/Jaspar(0.761) More Information \| Similar Motifs Found | motif file (matrix) |
| 22 | A C T G A C T G A G T C A C T G A C T G A G T C C T G A A C T G A G T C C T G A | 1e-77 | -1.777e+02 | 1.42% | 0.38% | 602.5bp (714.6bp) | IG1/MA2418.1/Jaspar(0.853) More Information \| Similar Motifs Found | motif file (matrix) |
| 23 | A G T C A G T C A G C T C G T A C G T A A C T G G C T A A G T C A C G T A C G T C T G A A C T G | 1e-64 | -1.495e+02 | 0.53% | 0.05% | 511.3bp (557.3bp) | CG2931(RRM)/Drosophila\_melanogaster-RNCMPT00147-PBM/HughesRNA(0.585) More Information \| Similar Motifs Found | motif file (matrix) |
| 24 | A C G T A G T C C G A T A C G T C G T A A C G T A G T C C G A T A C G T G T C A A C G T A G T C | 1e-62 | -1.446e+02 | 0.52% | 0.05% | 442.7bp (637.6bp) | Mecom/MA0029.2/Jaspar(0.856) More Information \| Similar Motifs Found | motif file (matrix) |
| 25 | A G T C A C G T A C G T C G T A A C G T A G T C A C T G C G T A A C G T C G T A | 1e-40 | -9.424e+01 | 0.56% | 0.12% | 347.2bp (700.8bp) | Dref/dmmpmm(Bergman)/fly(0.909) More Information \| Similar Motifs Found | motif file (matrix) |
| 26 | C G T A A C T G A G T C A C T G C G T A C G T A A C G T A G T C A G T C A G T C A G T C A C T G | 1e-31 | -7.299e+01 | 0.19% | 0.01% | 561.9bp (173.9bp) | dif/Rel/dmmpmm(Bergman)/fly(0.752) More Information \| Similar Motifs Found | motif file (matrix) |
