## Supplemental dataset for "Hybrid CNN and Multi-Head Attention Model for Analyzing Epigenetic Mechanisms and Gene Expression Across Fungal Phylogenetic Distances": NcrassaModel_FgramTest_K36me3locs_knownResults.html

/projects/wg-feeds/SHAP/NcrassaModel\_FgramTest\_K36me3locs\_SHAP\_noDup\_HOMER/ - Homer Known Motif Enrichment Results


### Homer Known Motif Enrichment Results (/projects/wg-feeds/SHAP/NcrassaModel\_FgramTest\_K36me3locs\_SHAP\_noDup\_HOMER/)

Homer *de novo* Motif Results  
Gene Ontology Enrichment Results  
Known Motif Enrichment Results (txt file)  
Total Target Sequences = 28519, Total Background Sequences = 131407

|  |  |  |  |  |  |  |  |  |  |  |  |
| --- | --- | --- | --- | --- | --- | --- | --- | --- | --- | --- | --- |
| Rank | Motif | Name | P-value | log P-pvalue | q-value (Benjamini) | # Target Sequences with Motif | % of Targets Sequences with Motif | # Background Sequences with Motif | % of Background Sequences with Motif | Motif File | SVG |
| 1 | A T G C C A G T A C G T A G T C C A T G A C G T A G T C A C G T A G C T G A T C | Unknown4/Arabidopsis-Promoters/Homer | 1e-883 | -2.035e+03 | 0.0000 | 11402.0 | 39.98% | 27989.6 | 21.30% | motif file (matrix) | svg |
| 2 | T A G C G T C A G A C T T A G C G T C A G A C T A G T C G C T A G A C T G A T C | ZML2(C2C2gata)/col-ZML2-DAP-Seq(GSE60143)/Homer | 1e-861 | -1.983e+03 | 0.0000 | 4585.0 | 16.08% | 6086.6 | 4.63% | motif file (matrix) | svg |
| 3 | C T A G T C A G C T G A C T G A C T A G C T G A C A T G C A T G C T G A C T A G C T A G C G T A C T A G C G T A G T C A | TF3A(C2H2)/col-TF3A-DAP-Seq(GSE60143)/Homer | 1e-596 | -1.374e+03 | 0.0000 | 8198.0 | 28.75% | 19803.7 | 15.07% | motif file (matrix) | svg |
| 4 | A C G T G A C T T A G C C G T A C T G A C A T G C T A G G A C T G A T C C G T A | Nr5a2(NR)/Pancreas-LRH1-ChIP-Seq(GSE34295)/Homer | 1e-503 | -1.160e+03 | 0.0000 | 6994.0 | 24.53% | 16730.5 | 12.73% | motif file (matrix) | svg |
| 5 | A C G T A T G C A C G T A C G T A G C T A G T C A G C T A G C T A G C T A G C T A G C T | hTCT(CPE) | 1e-493 | -1.136e+03 | 0.0000 | 13234.0 | 46.41% | 41280.3 | 31.42% | motif file (matrix) | svg |
| 6 | A C G T G A C T A T G C G C T A C T G A C T A G A C T G G A C T A G T C C G T A | Nr5a2(NR)/mES-Nr5a2-ChIP-Seq(GSE19019)/Homer | 1e-463 | -1.066e+03 | 0.0000 | 5405.0 | 18.95% | 11801.5 | 8.98% | motif file (matrix) | svg |
| 7 | T G A C C G T A C T G A A C T G A C T G G A C T G A T C T G C A G T A C T A C G | SF1(NR)/H295R-Nr5a1-ChIP-Seq(GSE44220)/Homer | 1e-454 | -1.046e+03 | 0.0000 | 4572.0 | 16.03% | 9206.3 | 7.01% | motif file (matrix) | svg |
| 8 | G A T C G C A T A G T C A G C T G A T C A G C T G A T C G A C T G A T C G A C T A G T C A C G T G A T C A G C T G A T C | GAGA-repeat/SacCer-Promoters/Homer | 1e-387 | -8.912e+02 | 0.0000 | 17936.0 | 62.90% | 64673.6 | 49.22% | motif file (matrix) | svg |
| 9 | G C T A G C A T A C G T A C G T A G C T G A T C G A C T G A C T A G C T A C G T A C G T A G C T | RLR1?/SacCer-Promoters/Homer | 1e-383 | -8.827e+02 | 0.0000 | 2720.0 | 9.54% | 4359.5 | 3.32% | motif file (matrix) | svg |
| 10 | G C A T A G C T A C G T A C G T A C T G A C G T G A T C A C G T A C G T A G C T C G A T G C A T A G T C G A C T C A G T | IDD5(C2H2)/colamp-IDD5-DAP-Seq(GSE60143)/Homer | 1e-362 | -8.338e+02 | 0.0000 | 5003.0 | 17.54% | 11685.6 | 8.89% | motif file (matrix) | svg |
| 11 | A G T C A G T C C G A T A C G T A C G T A C T G A C G T A G C T A G T C A G T C | Sox4(HMG)/proB-Sox4-ChIP-Seq(GSE50066)/Homer | 1e-341 | -7.873e+02 | 0.0000 | 7594.0 | 26.63% | 21381.3 | 16.27% | motif file (matrix) | svg |
| 12 | A C G T A C T G C G T A A C G T A C T G A C T G C G T A C G T A | HAP3(CCAATHAP3)/col-HAP3-DAP-Seq(GSE60143)/Homer | 1e-339 | -7.823e+02 | 0.0000 | 4874.0 | 17.09% | 11541.0 | 8.78% | motif file (matrix) | svg |
| 13 | G A C T A G C T A G C T C T A G A C G T G A T C A C G T A C G T G A C T C G A T G C A T A G T C | IDD4(C2H2)/col-IDD4-DAP-Seq(GSE60143)/Homer | 1e-320 | -7.377e+02 | 0.0000 | 6935.0 | 24.32% | 19265.1 | 14.66% | motif file (matrix) | svg |
| 14 | A T G C A G T C G C A T A G C T A C G T T C A G C G A T A G C T G A T C A T C G | Sox10(HMG)/SciaticNerve-Sox3-ChIP-Seq(GSE35132)/Homer | 1e-306 | -7.068e+02 | 0.0000 | 12935.0 | 45.36% | 43994.2 | 33.48% | motif file (matrix) | svg |
| 15 | A T G C G A T C C G A T A C G T A C G T A C T G C A G T A G C T | Sox3(HMG)/NPC-Sox3-ChIP-Seq(GSE33059)/Homer | 1e-304 | -7.005e+02 | 0.0000 | 13834.0 | 48.51% | 47996.0 | 36.53% | motif file (matrix) | svg |
| 16 | G A C T G C A T A C G T A C G T A C T G C G T A A G T C A G C T C G A T A T C G G C A T A C T G C G A T C T A G C G T A | WRKY50(WRKY)/col-WRKY50-DAP-Seq(GSE60143)/Homer | 1e-303 | -6.981e+02 | 0.0000 | 11113.0 | 38.97% | 36308.6 | 27.63% | motif file (matrix) | svg |
| 17 | C T G A T C A G G T A C G C T A A C T G T G A C G C A T C A T G | SCL(bHLH)/HPC7-Scl-ChIP-Seq(GSE13511)/Homer | 1e-294 | -6.777e+02 | 0.0000 | 19684.0 | 69.03% | 75424.6 | 57.41% | motif file (matrix) | svg |
| 18 | T G A C C T A G T C A G G T C A C G T A T C A G C G A T T C A G T C G A T G C A C T G A T A G C | PU.1-IRF(ETS:IRF)/Bcell-PU.1-ChIP-Seq(GSE21512)/Homer | 1e-293 | -6.756e+02 | 0.0000 | 7445.0 | 26.11% | 21706.8 | 16.52% | motif file (matrix) | svg |
| 19 | T C G A T G C A C A G T T C G A G A T C A G T C C G T A C G T A A C T G A G T C C G T A C G T A T C A G C G A T A G T C | AT5G25475(ABI3VP1)/col-AT5G25475-DAP-Seq(GSE60143)/Homer | 1e-292 | -6.742e+02 | 0.0000 | 10968.0 | 38.46% | 35943.2 | 27.36% | motif file (matrix) | svg |
| 20 | G A C T A C G T A C G T A C T G A C G T A G T C G C A T A G C T G C A T G C A T G A C T A G C T | SGR5(C2H2)/colamp-SGR5-DAP-Seq(GSE60143)/Homer | 1e-285 | -6.579e+02 | 0.0000 | 6324.0 | 22.18% | 17584.7 | 13.38% | motif file (matrix) | svg |
| 21 | C A T G A G C T T A C G G T C A G T A C T A G C A G C T G A C T A T C G T C G A | Esrrb(NR)/mES-Esrrb-ChIP-Seq(GSE11431)/Homer | 1e-272 | -6.265e+02 | 0.0000 | 6390.0 | 22.41% | 18081.0 | 13.76% | motif file (matrix) | svg |
| 22 | C G T A G A T C A G C T A C G T A C G T A C T G C G T A G T A C A G C T G C T A C G A T C G A T C G A T G C A T G C T A | WRKY18(WRKY)/col-WRKY18-DAP-Seq(GSE60143)/Homer | 1e-272 | -6.264e+02 | 0.0000 | 15500.0 | 54.35% | 56315.1 | 42.86% | motif file (matrix) | svg |
| 23 | C T G A C G A T C A T G A T C G G C A T C A T G G C T A A G T C | ASHR1(ND)/col-ASHR1-DAP-Seq(GSE60143)/Homer | 1e-268 | -6.173e+02 | 0.0000 | 10780.0 | 37.80% | 35749.6 | 27.21% | motif file (matrix) | svg |
| 24 | G A C T G C A T C T A G C G A T G A T C T C G A C A T G G A T C | Tgif1(Homeobox)/mES-Tgif1-ChIP-Seq(GSE55404)/Homer | 1e-254 | -5.868e+02 | 0.0000 | 19531.0 | 68.49% | 75780.4 | 57.68% | motif file (matrix) | svg |
| 25 | C A T G G C T A C T A G T A C G C G T A T C A G C G T A A C T G C G T A C A T G C T G A C G T A | BPC1(BBRBPC)/colamp-BPC1-DAP-Seq(GSE60143)/Homer | 1e-245 | -5.653e+02 | 0.0000 | 3852.0 | 13.51% | 9344.6 | 7.11% | motif file (matrix) | svg |
| 26 | A G T C T G C A T C G A C T G A A C T G C A T G A C G T A T G C G T C A T A C G | Erra(NR)/HepG2-Erra-ChIP-Seq(GSE31477)/Homer | 1e-245 | -5.651e+02 | 0.0000 | 12233.0 | 42.90% | 42540.2 | 32.38% | motif file (matrix) | svg |
| 27 | A G T C G A T C G C T A C G A T C A G T T A C G C G A T A G C T A G T C A T C G | SOX1(HMG)/NPC-SOX1-ChIP-Seq(GSE138215)/Homer | 1e-244 | -5.631e+02 | 0.0000 | 14860.0 | 52.11% | 54192.0 | 41.25% | motif file (matrix) | svg |
| 28 | A G C T A C G T A C T G A T G C A G T C C G T A C T G A T A C G | NF1-halfsite(CTF)/LNCaP-NF1-ChIP-Seq(Unpublished)/Homer | 1e-244 | -5.619e+02 | 0.0000 | 11714.0 | 41.08% | 40339.5 | 30.70% | motif file (matrix) | svg |
| 29 | A G T C C A T G A C G T A C G T A C T G C G T A A G T C G A C T G C A T G C T A | WRKY28(WRKY)/col-WRKY28-DAP-Seq(GSE60143)/Homer | 1e-235 | -5.427e+02 | 0.0000 | 12605.0 | 44.20% | 44428.7 | 33.82% | motif file (matrix) | svg |
| 30 | C T G A T C A G A G T C C G T A A T C G A T G C C G A T A C T G A G T C G A C T A T C G A G T C | MyoD(bHLH)/Myotube-MyoD-ChIP-Seq(GSE21614)/Homer | 1e-229 | -5.281e+02 | 0.0000 | 4221.0 | 14.80% | 10873.8 | 8.28% | motif file (matrix) | svg |
| 31 | C A T G G A C T T A C G G T C A G T A C G A T C G A C T A G C T A T C G T C G A T A C G T A G C | ERRg(NR)/Kidney-ESRRG-ChIP-Seq(GSE104905)/Homer | 1e-228 | -5.264e+02 | 0.0000 | 7356.0 | 25.80% | 22703.6 | 17.28% | motif file (matrix) | svg |
| 32 | C A T G G A C T C T A G C A T G C A G T C G A T C T A G C A T G C G A T C G T A C T A G C A G T C G A T C T A G C A T G | AT1G24250(Orphan)/col-AT1G24250-DAP-Seq(GSE60143)/Homer | 1e-227 | -5.232e+02 | 0.0000 | 3870.0 | 13.57% | 9675.0 | 7.36% | motif file (matrix) | svg |
| 33 | C T A G C T G A C T A G C T G A C T A G C T G A C T A G C T G A C T A G C T G A | SeqBias: GA-repeat | 1e-226 | -5.219e+02 | 0.0000 | 23110.0 | 81.04% | 94638.1 | 72.03% | motif file (matrix) | svg |
| 34 | C G A T C T A G C G T A G A C T C A G T C T A G C G T A A G C T C A T G C T A G | HOXA1(Homeobox)/mES-Hoxa1-ChIP-Seq(SRP084292)/Homer | 1e-224 | -5.171e+02 | 0.0000 | 4221.0 | 14.80% | 10950.2 | 8.33% | motif file (matrix) | svg |
| 35 | G C A T G A T C C T A G C G T A G C A T C G T A G C A T A G T C C T A G C G T A G C A T C G A T | AT5G22990(C2H2)/col-AT5G22990-DAP-Seq(GSE60143)/Homer | 1e-220 | -5.081e+02 | 0.0000 | 7496.0 | 26.29% | 23440.9 | 17.84% | motif file (matrix) | svg |
| 36 | A C G T A C G T A C G T A C G T A C G T A C G T A C G T A C G T A C G T A C G T | VRN1(ABI3VP1)/col-VRN1-DAP-Seq(GSE60143)/Homer | 1e-219 | -5.054e+02 | 0.0000 | 602.0 | 2.11% | 329.9 | 0.25% | motif file (matrix) | svg |
| 37 | C T G A T C G A C G T A A T G C C G T A C G T A C G A T C T A G T C A G G A T C | Sox15(HMG)/CPA-Sox15-ChIP-Seq(GSE62909)/Homer | 1e-219 | -5.043e+02 | 0.0000 | 8800.0 | 30.86% | 28752.4 | 21.88% | motif file (matrix) | svg |
| 38 | A T G C G T A C C G T A A G C T G C A T T A C G A G C T A G C T A G T C A G C T | Sox6(HMG)/Myotubes-Sox6-ChIP-Seq(GSE32627)/Homer | 1e-213 | -4.913e+02 | 0.0000 | 13483.0 | 47.28% | 48949.5 | 37.26% | motif file (matrix) | svg |
| 39 | A G C T A G C T C A T G C T G A G T A C A G T C A G C T A G C T C A G T C T A G | RARa(NR)/K562-RARa-ChIP-Seq(Encode)/Homer | 1e-211 | -4.874e+02 | 0.0000 | 17219.0 | 60.38% | 66052.3 | 50.27% | motif file (matrix) | svg |
| 40 | G T A C C A T G A G C T A C G T A C T G C G T A A G T C G A C T G C A T C G A T | WRKY29(WRKY)/colamp-WRKY29-DAP-Seq(GSE60143)/Homer | 1e-210 | -4.839e+02 | 0.0000 | 11034.0 | 38.69% | 38360.6 | 29.20% | motif file (matrix) | svg |
| 41 | C T G A C G A T C T A G T C A G G A T C C T G A T C A G G A T C C T G A A C T G A G T C G C T A A C G T A G T C G C A T | PRDM9(Zf)/Testis-DMC1-ChIP-Seq(GSE35498)/Homer | 1e-209 | -4.821e+02 | 0.0000 | 3264.0 | 11.45% | 7856.0 | 5.98% | motif file (matrix) | svg |
| 42 | C G T A G A T C C A T G G C A T G A C T C T A G T C G A T A G C A G C T G C A T | WRKY55(WRKY)/col-WRKY55-DAP-Seq(GSE60143)/Homer | 1e-208 | -4.810e+02 | 0.0000 | 11612.0 | 40.72% | 40883.7 | 31.12% | motif file (matrix) | svg |
| 43 | G A C T T C A G G C A T A G T C G C T A G A T C C T G A A C G T A G T C G T C A | Replumless(BLH)/Arabidopsis-RPL.GFP-ChIP-Seq(GSE78727)/Homer | 1e-207 | -4.773e+02 | 0.0000 | 12862.0 | 45.10% | 46386.9 | 35.31% | motif file (matrix) | svg |
| 44 | A G C T A G C T A G C T A C T G A C G T A G T C A C T G A C G T G A C T C G A T G C A T A T C G | IDD7(C2H2)/col-IDD7-DAP-Seq(GSE60143)/Homer | 1e-202 | -4.660e+02 | 0.0000 | 4532.0 | 15.89% | 12452.7 | 9.48% | motif file (matrix) | svg |
| 45 | G T C A G C T A G C T A T C G A A T C G A C G T A G T C T C G A T C G A T G A C | WRKY40(WRKY)/colamp-WRKY40-DAP-Seq(GSE60143)/Homer | 1e-201 | -4.635e+02 | 0.0000 | 7120.0 | 24.97% | 22397.9 | 17.05% | motif file (matrix) | svg |
| 46 | C T A G G T A C C A T G G A C T C G A T C A T G G T C A G T A C G A C T C G A T C G A T C G A T | WRKY27(WRKY)/colamp-WRKY27-DAP-Seq(GSE60143)/Homer | 1e-198 | -4.571e+02 | 0.0000 | 9558.0 | 33.52% | 32437.0 | 24.69% | motif file (matrix) | svg |
| 47 | A T G C A G T C G A T C C G T A A C G T A C G T A C T G A C G T A G C T G A T C | Sox2(HMG)/mES-Sox2-ChIP-Seq(GSE11431)/Homer | 1e-196 | -4.524e+02 | 0.0000 | 7914.0 | 27.75% | 25708.0 | 19.57% | motif file (matrix) | svg |
| 48 | C T G A A T G C G C T A G C A T A T G C C G T A T C G A C T G A C T A G T C A G T A C G G T C A | Tcf4(HMG)/Hct116-Tcf4-ChIP-Seq(SRA012054)/Homer | 1e-195 | -4.505e+02 | 0.0000 | 4544.0 | 15.93% | 12620.1 | 9.61% | motif file (matrix) | svg |
| 49 | A C G T A G T C A G T C C G A T A C G T A C G T A C T G A C G T A T G C G A C T A C T G T A C G | Sox21(HMG)/ESC-SOX21-ChIP-Seq(GSE110505)/Homer | 1e-194 | -4.487e+02 | 0.0000 | 12722.0 | 44.61% | 46152.9 | 35.13% | motif file (matrix) | svg |
| 50 | A G C T C A T G G C A T G A T C T G C A C T A G G A T C A C G T | Tgif2(Homeobox)/mES-Tgif2-ChIP-Seq(GSE55404)/Homer | 1e-188 | -4.336e+02 | 0.0000 | 19928.0 | 69.88% | 79777.7 | 60.72% | motif file (matrix) | svg |
| 51 | G C A T C G A T G A C T T G C A A C T G A G T C T G A C A C T G G A T C A G T C C G T A G A C T | ERF15(AP2EREBP)/colamp-ERF15-DAP-Seq(GSE60143)/Homer | 1e-186 | -4.299e+02 | 0.0000 | 13674.0 | 47.95% | 50639.9 | 38.54% | motif file (matrix) | svg |
| 52 | G C A T T G A C C A T G G A C T C A G T C A T G T C G A G T A C G A C T G C T A C G A T C G A T | WRKY6(WRKY)/colamp-WRKY6-DAP-Seq(GSE60143)/Homer | 1e-184 | -4.260e+02 | 0.0000 | 9414.0 | 33.01% | 32215.2 | 24.52% | motif file (matrix) | svg |
| 53 | C G A T A C G T A C G T A G C T A G C T G A T C G A T C G C T A A G C T A C G T A T C G T A C G | NFATC2(RHD)/Islets-NFATC2-ChIP-Seq(GSE158496)/Homer | 1e-180 | -4.153e+02 | 0.0000 | 11525.0 | 40.41% | 41372.9 | 31.49% | motif file (matrix) | svg |
| 54 | T C G A A G T C C G T A A T C G A T G C C G A T A C T G A G T C A G C T A C T G | Tcf12(bHLH)/GM12878-Tcf12-ChIP-Seq(GSE32465)/Homer | 1e-178 | -4.116e+02 | 0.0000 | 4822.0 | 16.91% | 13985.3 | 10.64% | motif file (matrix) | svg |
| 55 | T A C G G A C T T G A C C G T A A C G T G A T C G T C A C G T A A C G T A T G C C G T A G A C T | HOXA2(Homeobox)/mES-Hoxa2-ChIP-Seq(Donaldson\_et\_al.)/Homer | 1e-178 | -4.108e+02 | 0.0000 | 1977.0 | 6.93% | 4058.6 | 3.09% | motif file (matrix) | svg |
| 56 | T C G A G T A C C A T G A G C T A C G T C A T G G T C A G T A C A G C T G C T A C G A T C A G T | WRKY31(WRKY)/colamp-WRKY31-DAP-Seq(GSE60143)/Homer | 1e-175 | -4.051e+02 | 0.0000 | 7963.0 | 27.92% | 26449.6 | 20.13% | motif file (matrix) | svg |
| 57 | A T C G T G C A G A T C C T A G A C G T A T C G C G T A A G T C T C A G A C T G T C A G G C T A | Knotted(Homeobox)/Corn-KN1-ChIP-Seq(GSE39161)/Homer | 1e-175 | -4.031e+02 | 0.0000 | 15015.0 | 52.65% | 57086.4 | 43.45% | motif file (matrix) | svg |
| 58 | G C A T A C G T A C T G A C G T A G T C A C T G A T C G G T C A C G A T C G T A | ARF2(ARF)/col-ARF2-DAP-Seq(GSE60143)/Homer | 1e-174 | -4.014e+02 | 0.0000 | 19419.0 | 68.10% | 77779.2 | 59.20% | motif file (matrix) | svg |
| 59 | A T G C C A T G A C G T A C G T A C T G C G T A A G T C G A C T G C A T C G A T | WRKY71(WRKY)/col-WRKY71-DAP-Seq(GSE60143)/Homer | 1e-173 | -3.996e+02 | 0.0000 | 8364.0 | 29.33% | 28167.7 | 21.44% | motif file (matrix) | svg |
| 60 | G C T A C G T A G C T A C G T A C T G A A C T G A C G T G T A C C G T A C T G A G T A C A C T G | WRKY22(WRKY)/colamp-WRKY22-DAP-Seq(GSE60143)/Homer | 1e-170 | -3.923e+02 | 0.0000 | 6866.0 | 24.08% | 22146.1 | 16.86% | motif file (matrix) | svg |
| 61 | A T G C C A T G A G C T C A G T C A T G T C G A A G T C G A C T C G A T C G A T C A G T C A G T | WRKY26(WRKY)/colamp-WRKY26-DAP-Seq(GSE60143)/Homer | 1e-170 | -3.923e+02 | 0.0000 | 6317.0 | 22.15% | 19957.8 | 15.19% | motif file (matrix) | svg |
| 62 | A T G C C A T G G C A T G A C T C T A G T C G A G T A C A G C T C G T A G C T A | WRKY75(WRKY)/col-WRKY75-DAP-Seq(GSE60143)/Homer | 1e-169 | -3.895e+02 | 0.0000 | 10064.0 | 35.29% | 35437.3 | 26.97% | motif file (matrix) | svg |
| 63 | A T G C C A G T A G C T A G C T T C A G G T C A T A G C G A C T C G T A C G A T | WRKY20(WRKY)/col-WRKY20-DAP-Seq(GSE60143)/Homer | 1e-168 | -3.891e+02 | 0.0000 | 8301.0 | 29.11% | 28035.5 | 21.34% | motif file (matrix) | svg |
| 64 | A G T C C T A G A C G T A C G T A C T G C G T A A G T C A G C T G C T A G C A T | WRKY24(WRKY)/colamp-WRKY24-DAP-Seq(GSE60143)/Homer | 1e-167 | -3.862e+02 | 0.0000 | 9509.0 | 33.35% | 33125.8 | 25.21% | motif file (matrix) | svg |
| 65 | C T A G A G T C A G T C A C T G C G T A A G T C C T G A G A C T | DDF1(AP2EREBP)/col-DDF1-DAP-Seq(GSE60143)/Homer | 1e-162 | -3.750e+02 | 0.0000 | 9543.0 | 33.46% | 33418.0 | 25.43% | motif file (matrix) | svg |
| 66 | C A T G G T A C A C T G G T C A A G C T T A C G T G C A A T C G T G A C C A G T | TOD6?/SacCer-Promoters/Homer | 1e-161 | -3.712e+02 | 0.0000 | 2958.0 | 10.37% | 7530.7 | 5.73% | motif file (matrix) | svg |
| 67 | T C A G T G A C C A T G G C A T C A G T A C T G C G T A T G A C G A C T C G A T C G A T C G T A | WRKY3(WRKY)/col-WRKY3-DAP-Seq(GSE60143)/Homer | 1e-156 | -3.607e+02 | 0.0000 | 7376.0 | 25.87% | 24582.4 | 18.71% | motif file (matrix) | svg |
| 68 | C G T A C T A G C A T G A G C T C T G A C A T G C A G T C G A T C T A G C T A G | MYB30(MYB)/colamp-MYB30-DAP-Seq(GSE60143)/Homer | 1e-154 | -3.555e+02 | 0.0000 | 9959.0 | 34.92% | 35458.4 | 26.99% | motif file (matrix) | svg |
| 69 | C G A T A G C T T G C A A C T G A G T C T G A C C T A G G T A C A G T C C G T A G C A T G C A T | ERF13(AP2EREBP)/colamp-ERF13-DAP-Seq(GSE60143)/Homer | 1e-153 | -3.531e+02 | 0.0000 | 10552.0 | 37.00% | 38038.9 | 28.95% | motif file (matrix) | svg |
| 70 | T C G A A C T G C A T G A G C T A G T C C G T A C T G A C T A G A C T G C G A T A T G C C T G A | RAR:RXR(NR),DR0/ES-RAR-ChIP-Seq(GSE56893)/Homer | 1e-151 | -3.491e+02 | 0.0000 | 1331.0 | 4.67% | 2399.6 | 1.83% | motif file (matrix) | svg |
| 71 | C G T A C T A G C A T G G A C T C T G A A C T G A C G T A C G T C T A G C T A G C A T G T C G A | MYB94(MYB)/col-MYB94-DAP-Seq(GSE60143)/Homer | 1e-150 | -3.471e+02 | 0.0000 | 4327.0 | 15.17% | 12705.6 | 9.67% | motif file (matrix) | svg |
| 72 | G C T A G C T A C G T A C G T A C T G A C T A G A C G T A G T C C G T A C T G A G T A C A C T G | WRKY65(WRKY)/colamp-WRKY65-DAP-Seq(GSE60143)/Homer | 1e-150 | -3.460e+02 | 0.0000 | 6213.0 | 21.79% | 20055.5 | 15.26% | motif file (matrix) | svg |
| 73 | G A T C G C T A G T A C A G T C G C T A T G C A G T A C G A T C C G T A G A C T | MYB83(MYB)/colamp-MYB83-DAP-Seq(GSE60143)/Homer | 1e-148 | -3.423e+02 | 0.0000 | 14425.0 | 50.58% | 55351.0 | 42.13% | motif file (matrix) | svg |
| 74 | C G A T C G T A G C T A G C A T G C A T C T G A A C T G A C G T A G T C C G T A C G T A G T A C T C A G G C T A C G A T | WRKY25(WRKY)/colamp-WRKY25-DAP-Seq(GSE60143)/Homer | 1e-148 | -3.414e+02 | 0.0000 | 11546.0 | 40.49% | 42532.7 | 32.37% | motif file (matrix) | svg |
| 75 | G C A T C G T A C T A G A G T C G T C A C G T A A T G C A C G T A C G T A C T G G A T C G C A T C G T A G C T A G C T A | bHLH122(bHLH)/col100-bHLH122-DAP-Seq(GSE60143)/Homer | 1e-147 | -3.403e+02 | 0.0000 | 7781.0 | 27.29% | 26503.8 | 20.17% | motif file (matrix) | svg |
| 76 | G T C A T G C A G C T A A G T C C G T A A C T G T G A C G C A T T C A G C A G T | Ap4(bHLH)/AML-Tfap4-ChIP-Seq(GSE45738)/Homer | 1e-147 | -3.396e+02 | 0.0000 | 6298.0 | 22.09% | 20470.6 | 15.58% | motif file (matrix) | svg |
| 77 | C G A T T G A C C A T G G A C T A C G T C A T G C G T A G A T C G A C T G C A T G C A T C G A T | WRKY14(WRKY)/colamp-WRKY14-DAP-Seq(GSE60143)/Homer | 1e-145 | -3.343e+02 | 0.0000 | 5965.0 | 20.92% | 19200.3 | 14.61% | motif file (matrix) | svg |
| 78 | G A C T A C T G C G A T A G T C A C T G C T A G A G T C C G T A | Rap210(AP2EREBP)/col-Rap210-DAP-Seq(GSE60143)/Homer | 1e-143 | -3.309e+02 | 0.0000 | 11230.0 | 39.38% | 41313.0 | 31.44% | motif file (matrix) | svg |
| 79 | A G C T C T A G A G T C A G T C A C T G C G T A A G T C C T G A G C A T G C T A C T G A G C A T G C A T C G A T G C A T | CBF4(AP2EREBP)/colamp-CBF4-DAP-Seq(GSE60143)/Homer | 1e-142 | -3.289e+02 | 0.0000 | 12698.0 | 44.53% | 47812.2 | 36.39% | motif file (matrix) | svg |
| 80 | A G C T A G C T A G C T A C T G A C G T A G T C A C T G A C G T G C A T G C A T G C A T A C G T | At5g66730(C2H2)/colamp-At5g66730-DAP-Seq(GSE60143)/Homer | 1e-142 | -3.274e+02 | 0.0000 | 3284.0 | 11.52% | 9017.6 | 6.86% | motif file (matrix) | svg |
| 81 | G A C T G A T C G A T C G C T A G T A C A G T C G C T A C T G A G T A C G A T C G C T A G A C T | MYB13(MYB)/col-MYB13-DAP-Seq(GSE60143)/Homer | 1e-141 | -3.255e+02 | 0.0000 | 6792.0 | 23.82% | 22635.8 | 17.23% | motif file (matrix) | svg |
| 82 | C G A T C G T A G C T A G A C T T C G A A G C T A G T C A C T G T C G A A G C T C T G A C G A T | ZBTB38(Zf)/Hela-ZBTB38-ChIP-seq(GSE108618)/Homer | 1e-141 | -3.248e+02 | 0.0000 | 22908.0 | 80.33% | 96279.2 | 73.28% | motif file (matrix) | svg |
| 83 | G A C T C A G T G C A T C G A T T G A C A C G T A T G C G T A C C T G A A C T G A C T G A G C T | WIP5(C2H2)/colamp-WIP5-DAP-Seq(GSE60143)/Homer | 1e-140 | -3.225e+02 | 0.0000 | 9418.0 | 33.03% | 33623.5 | 25.59% | motif file (matrix) | svg |
| 84 | A T G C A G T C A G C T A G C T A C G T A T C G C G T A C G A T T A G C G A C T | LEF1(HMG)/H1-LEF1-ChIP-Seq(GSE64758)/Homer | 1e-139 | -3.208e+02 | 0.0000 | 5862.0 | 20.56% | 18945.1 | 14.42% | motif file (matrix) | svg |
| 85 | T A G C C A T G G A C T G A C T T C A G G T C A G A T C G A C T G C A T G C T A | WRKY15(WRKY)/col-WRKY15-DAP-Seq(GSE60143)/Homer | 1e-137 | -3.159e+02 | 0.0000 | 10208.0 | 35.80% | 37108.2 | 28.24% | motif file (matrix) | svg |
| 86 | A G T C G A C T A G C T C G A T A T C G G C T A C G A T A T C G C G A T A C T G T A C G A C G T | Tcf7(HMG)/GM12878-TCF7-ChIP-Seq(Encode)/Homer | 1e-136 | -3.149e+02 | 0.0000 | 3229.0 | 11.32% | 8920.4 | 6.79% | motif file (matrix) | svg |
| 87 | G T A C A C G T A C G T A T C G C A G T C G A T A T C G G C T A T G C A T A G C C G T A G T C A C A T G A G C T G C T A | ANAC013(NAC)/col-ANAC013-DAP-Seq(GSE60143)/Homer | 1e-134 | -3.088e+02 | 0.0000 | 4625.0 | 16.22% | 14219.9 | 10.82% | motif file (matrix) | svg |
| 88 | G C A T C G A T G C A T C G T A C T A G A G T C G T C A C G T A A T C G A C G T A C G T A C T G G T A C G C A T C G A T | bHLH80(bHLH)/col-bHLH80-DAP-Seq(GSE60143)/Homer | 1e-132 | -3.049e+02 | 0.0000 | 8120.0 | 28.47% | 28392.7 | 21.61% | motif file (matrix) | svg |
| 89 | G C A T A G C T A G C T A G C T A C T G A C G T A G T C A C T G A C G T G A C T C G A T G C A T | JKD(C2H2)/col-JKD-DAP-Seq(GSE60143)/Homer | 1e-130 | -3.013e+02 | 0.0000 | 2430.0 | 8.52% | 6186.1 | 4.71% | motif file (matrix) | svg |
| 90 | T C A G T A C G T A G C A C G T A C T G C G A T A G T C C G T A T A C G A G T C | Meis1(Homeobox)/MastCells-Meis1-ChIP-Seq(GSE48085)/Homer | 1e-129 | -2.986e+02 | 0.0000 | 12588.0 | 44.14% | 47820.3 | 36.40% | motif file (matrix) | svg |
| 91 | C T A G C T A G T C G A C T A G C G T A A T C G T C G A A C T G C T G A T C G A C T G A T A C G | FRS9(ND)/col-FRS9-DAP-Seq(GSE60143)/Homer | 1e-129 | -2.985e+02 | 0.0000 | 1100.0 | 3.86% | 1940.2 | 1.48% | motif file (matrix) | svg |
| 92 | C G T A G A C T C A T G C T A G A G T C A C T G C T A G A G T C C A T G C T A G | ERF7(AP2EREBP)/col-ERF7-DAP-Seq(GSE60143)/Homer | 1e-129 | -2.976e+02 | 0.0000 | 13470.0 | 47.23% | 51785.0 | 39.41% | motif file (matrix) | svg |
| 93 | G T A C C A T G A G C T A G C T T C A G T G C A T G A C A G C T C G T A C G T A | WRKY33(WRKY)/col-WRKY33-DAP-Seq(GSE60143)/Homer | 1e-128 | -2.955e+02 | 0.0000 | 9077.0 | 31.83% | 32570.9 | 24.79% | motif file (matrix) | svg |
| 94 | T G A C T A G C T C A G T C G A T C G A C G T A A G T C C G T A C G T A C G A T C T A G T A C G | Sox7(HMG)/ESC-Sox7-ChIP-Seq(GSE133899)/Homer | 1e-128 | -2.948e+02 | 0.0000 | 3631.0 | 12.73% | 10577.5 | 8.05% | motif file (matrix) | svg |
| 95 | C A T G A G T C G T C A C G T A A T G C A C G T A C G T A C T G | bHLH130(bHLH)/col-bHLH130-DAP-Seq(GSE60143)/Homer | 1e-127 | -2.931e+02 | 0.0000 | 6835.0 | 23.97% | 23226.6 | 17.68% | motif file (matrix) | svg |
| 96 | T C G A T C G A C T G A C G T A A C T G A T G C A C G T A G T C | Lola-I(Zf)/Embryo-LolaI-ChIP-Seq(GSE200870)/Homer | 1e-126 | -2.922e+02 | 0.0000 | 4932.0 | 17.29% | 15594.0 | 11.87% | motif file (matrix) | svg |
| 97 | C G T A G C A T C G T A C G T A G C A T A C T G C G A T A G T C A C T G A C T G G A C T C T A G | AT1G71450(AP2EREBP)/col-AT1G71450-DAP-Seq(GSE60143)/Homer | 1e-126 | -2.907e+02 | 0.0000 | 17707.0 | 62.09% | 71423.3 | 54.36% | motif file (matrix) | svg |
| 98 | A G C T G A C T A C G T A C T G A C G T A G T C A C T G A C G T G C A T C G A T | AtIDD11(C2H2)/colamp-AtIDD11-DAP-Seq(GSE60143)/Homer | 1e-124 | -2.860e+02 | 0.0000 | 4243.0 | 14.88% | 12984.3 | 9.88% | motif file (matrix) | svg |
| 99 | G A C T A C G T A G C T G A C T A C T G C A G T A G T C A T C G A C G T G C A T G C A T G C A T | MGP(C2H2)/colamp-MGP-DAP-Seq(GSE60143)/Homer | 1e-123 | -2.838e+02 | 0.0000 | 3697.0 | 12.96% | 10929.6 | 8.32% | motif file (matrix) | svg |
| 100 | A T G C G A T C C G T A A G C T C A G T A T C G G C A T A G C T G A C T A C T G | Sox17(HMG)/Endoderm-Sox17-ChIP-Seq(GSE61475)/Homer | 1e-122 | -2.810e+02 | 0.0000 | 7185.0 | 25.20% | 24834.3 | 18.90% | motif file (matrix) | svg |
| 101 | G C A T C G A T C G T A G A T C C A T G A C G T A C G T A C T G C G T A A G T C A G C T G C A T G C A T C G T A G C T A | WRKY45(WRKY)/col-WRKY45-DAP-Seq(GSE60143)/Homer | 1e-121 | -2.787e+02 | 0.0000 | 4660.0 | 16.34% | 14679.2 | 11.17% | motif file (matrix) | svg |
| 102 | G T A C A C T G A G T C A G T C C T A G G A T C G T A C C T G A | CRF4(AP2EREBP)/colamp-CRF4-DAP-Seq(GSE60143)/Homer | 1e-120 | -2.771e+02 | 0.0000 | 5712.0 | 20.03% | 18872.0 | 14.36% | motif file (matrix) | svg |
| 103 | C G A T C T A G A C G T A C G T A C G T C G T A A G C T C G A T A G C T C G T A C T A G T A G C | FoxD3(forkhead)/ZebrafishEmbryo-Foxd3.biotin-ChIP-seq(GSE106676)/Homer | 1e-119 | -2.762e+02 | 0.0000 | 5598.0 | 19.63% | 18424.9 | 14.02% | motif file (matrix) | svg |
| 104 | G A T C A G C T C T A G G A T C T G A C C T A G C G T A G T A C C G T A G C A T G T C A C T G A | CBF3(AP2EREBP)/colamp-CBF3-DAP-Seq(GSE60143)/Homer | 1e-119 | -2.751e+02 | 0.0000 | 9216.0 | 32.32% | 33479.3 | 25.48% | motif file (matrix) | svg |
| 105 | C T A G T C A G C T G A T C A G T G C A A C T G T C G A T C A G | Trl(Zf)/S2-GAGAfactor-ChIP-Seq(GSE40646)/Homer | 1e-117 | -2.711e+02 | 0.0000 | 14174.0 | 49.70% | 55435.6 | 42.19% | motif file (matrix) | svg |
| 106 | G C A T A C T G C T A G A G T C A C T G A C T G A G T C A C G T | ERF105(AP2EREBP)/colamp-ERF105-DAP-Seq(GSE60143)/Homer | 1e-117 | -2.707e+02 | 0.0000 | 10805.0 | 37.89% | 40427.2 | 30.77% | motif file (matrix) | svg |
| 107 | A C G T T G C A A G C T G A T C C T A G C T G A A G C T G T C A T C G A C G T A | CUX1(Homeobox)/K562-CUX1-ChIP-Seq(GSE92882)/Homer | 1e-117 | -2.703e+02 | 0.0000 | 12086.0 | 42.38% | 46076.9 | 35.07% | motif file (matrix) | svg |
| 108 | G T A C G C T A T C A G C T G A C T A G C A T G A G C T G A T C T G C A T C G A C T G A A C T G C A G T A G T C G A T C G C T A | HNF4a(NR),DR1/HepG2-HNF4a-ChIP-Seq(GSE25021)/Homer | 1e-115 | -2.660e+02 | 0.0000 | 2926.0 | 10.26% | 8229.0 | 6.26% | motif file (matrix) | svg |
| 109 | C G T A G A C T C A T G C T A G A G T C A C T G A C T G G T A C C A T G T A C G | ERF3(AP2EREBP)/colamp-ERF3-DAP-Seq(GSE60143)/Homer | 1e-115 | -2.660e+02 | 0.0000 | 7507.0 | 26.32% | 26388.9 | 20.08% | motif file (matrix) | svg |
| 110 | G A C T A G T C C T G A A G T C A G T C A C T G C T G A A G T C G C T A G C A T G T A C C G A T G C A T G A C T C G A T | CBF2(AP2EREBP)/colamp-CBF2-DAP-Seq(GSE60143)/Homer | 1e-115 | -2.658e+02 | 0.0000 | 9442.0 | 33.11% | 34598.4 | 26.33% | motif file (matrix) | svg |
| 111 | C G A T C T A G A C T G A G C T C T G A A C T G A C G T A C G T C T A G C T A G | MYB96(MYB)/colamp-MYB96-DAP-Seq(GSE60143)/Homer | 1e-115 | -2.649e+02 | 0.0000 | 8606.0 | 30.18% | 31037.0 | 23.62% | motif file (matrix) | svg |
| 112 | C T G A G C A T A C T G C T A G A G T C A C T G A C T G A G T C A C T G T C A G | AT4G18450(AP2EREBP)/col-AT4G18450-DAP-Seq(GSE60143)/Homer | 1e-114 | -2.639e+02 | 0.0000 | 3988.0 | 13.98% | 12235.7 | 9.31% | motif file (matrix) | svg |
| 113 | T C A G A T C G G A C T A C T G G A C T C A G T C T A G C G T A G T A C C G T A C T A G A T C G | Tbx20(T-box)/Heart-Tbx20-ChIP-Seq(GSE29636)/Homer | 1e-114 | -2.638e+02 | 0.0000 | 2228.0 | 7.81% | 5752.8 | 4.38% | motif file (matrix) | svg |
| 114 | C T A G T C G A C G A T C T A G G C A T C A G T C T A G G A T C C G T A G T C A | CEBP:AP1(bZIP)/ThioMac-CEBPb-ChIP-Seq(GSE21512)/Homer | 1e-113 | -2.621e+02 | 0.0000 | 7586.0 | 26.60% | 26776.4 | 20.38% | motif file (matrix) | svg |
| 115 | C T G A T C G A G T A C A C G T A C G T A T C G A C G T C G A T A T C G G C T A G T A C A T G C C G T A T G C A C A T G | ANAC103(NAC)/col-ANAC103-DAP-Seq(GSE60143)/Homer | 1e-113 | -2.617e+02 | 0.0000 | 4251.0 | 14.91% | 13275.0 | 10.10% | motif file (matrix) | svg |
| 116 | C G A T T G C A T G C A G A T C C G T A A C T G T G A C G A C T C A T G A C T G | Tcf21(bHLH)/ArterySmoothMuscle-Tcf21-ChIP-Seq(GSE61369)/Homer | 1e-113 | -2.612e+02 | 0.0000 | 4906.0 | 17.20% | 15848.0 | 12.06% | motif file (matrix) | svg |
| 117 | G T A C C T G A T A G C C G T A G C T A T C G A T G C A T G A C C T A G G T C A A G T C C G T A C T G A C T G A C G T A | At1g14580(C2H2)/colamp-At1g14580-DAP-Seq(GSE60143)/Homer | 1e-113 | -2.607e+02 | 0.0000 | 1550.0 | 5.44% | 3490.5 | 2.66% | motif file (matrix) | svg |
| 118 | G T C A T C G A T C G A C G T A G C T A C G T A T C G A T G A C A C T G C G T A A G T C C G T A C G T A T C G A G C T A | IDD2(C2H2)/colamp-IDD2-DAP-Seq(GSE60143)/Homer | 1e-112 | -2.593e+02 | 0.0000 | 1519.0 | 5.33% | 3398.1 | 2.59% | motif file (matrix) | svg |
| 119 | T C G A C G T A A G T C C G T A C T A G A G T C C G A T A C T G G A C T A G C T A C T G G A C T | HLH-1(bHLH)/cElegans-Embryo-HLH1-ChIP-Seq(modEncode)/Homer | 1e-112 | -2.584e+02 | 0.0000 | 5861.0 | 20.55% | 19713.0 | 15.00% | motif file (matrix) | svg |
| 120 | A T G C G C A T T A G C C G A T T A G C G C A T T A G C G C A T A T G C G A C T | GAGA-repeat/Arabidopsis-Promoters/Homer | 1e-111 | -2.570e+02 | 0.0000 | 5653.0 | 19.82% | 18889.9 | 14.38% | motif file (matrix) | svg |
| 121 | C G A T C G T A G T A C A C G T A C G T T C A G G C A T C A G T T A C G G T C A C G T A A G T C C G T A T G C A C A T G | NAC2(NAC)/colamp-NAC2-DAP-Seq(GSE60143)/Homer | 1e-111 | -2.562e+02 | 0.0000 | 6689.0 | 23.46% | 23132.8 | 17.61% | motif file (matrix) | svg |
| 122 | G A C T G T A C C T G A G A T C A G T C C T A G G C T A G T A C C T G A G C T A G C A T C G A T G C A T G A C T C G T A | AT3G16280(AP2EREBP)/colamp-AT3G16280-DAP-Seq(GSE60143)/Homer | 1e-108 | -2.489e+02 | 0.0000 | 8234.0 | 28.87% | 29710.8 | 22.61% | motif file (matrix) | svg |
| 123 | C A G T T C A G T C G A A G T C C G T A A C T G T G A C C G A T A C T G A C T G A C G T A T C G | Atoh7(bHLH)/Retina-Atoh7-CutnRun(GSE156756)/Homer | 1e-107 | -2.480e+02 | 0.0000 | 4232.0 | 14.84% | 13353.4 | 10.16% | motif file (matrix) | svg |
| 124 | T C G A T C G A A G T C C G T A C T A G T A G C A C G T A C T G | MyoG(bHLH)/C2C12-MyoG-ChIP-Seq(GSE36024)/Homer | 1e-107 | -2.469e+02 | 0.0000 | 5449.0 | 19.11% | 18197.6 | 13.85% | motif file (matrix) | svg |
| 125 | C G A T T G C A A G T C A C G T A C G T T A C G C G A T C G A T T A C G G C T A G C T A A T G C C G T A G T C A C A T G | ANAC016(NAC)/col-ANAC016-DAP-Seq(GSE60143)/Homer | 1e-106 | -2.454e+02 | 0.0000 | 8628.0 | 30.26% | 31443.5 | 23.93% | motif file (matrix) | svg |
| 126 | A C T G C T A G A G T C A C T G A C T G A G T C A C G T C T A G | AT5G23930(mTERF)/col-AT5G23930-DAP-Seq(GSE60143)/Homer | 1e-105 | -2.440e+02 | 0.0000 | 10173.0 | 35.67% | 38129.4 | 29.02% | motif file (matrix) | svg |
| 127 | C A T G A G T C G T A C A C T G A T G C A G T C C A T G G A T C G A T C C T G A | ERF5(AP2EREBP)/colamp-ERF5-DAP-Seq(GSE60143)/Homer | 1e-105 | -2.437e+02 | 0.0000 | 4087.0 | 14.33% | 12837.7 | 9.77% | motif file (matrix) | svg |
| 128 | T A G C C G T A C T G A T A C G C G T A A C G T A C T G A C T G A G T C T A C G C T A G G T A C | YY1(Zf)/Promoter/Homer | 1e-105 | -2.423e+02 | 0.0000 | 794.0 | 2.78% | 1299.2 | 0.99% | motif file (matrix) | svg |
| 129 | C T A G G C A T A C T G C T A G A G T C A C T G A C T G A G T C A C T G T C A G | ERF10(AP2EREBP)/col-ERF10-DAP-Seq(GSE60143)/Homer | 1e-105 | -2.418e+02 | 0.0000 | 6971.0 | 24.45% | 24511.3 | 18.66% | motif file (matrix) | svg |
| 130 | A T G C G T A C A G T C A G T C A C G T A C G T C G A T A C G T | AT5G02460(C2C2dof)/col-AT5G02460-DAP-Seq(GSE60143)/Homer | 1e-104 | -2.410e+02 | 0.0000 | 13296.0 | 46.62% | 52027.2 | 39.60% | motif file (matrix) | svg |
| 131 | G A C T G C A T G C A T A G T C A G C T T C G A T A C G G C T A C G T A A C T G G T A C G C A T C G A T A G T C G A C T | HSF3(HSF)/colamp-HSF3-DAP-Seq(GSE60143)/Homer | 1e-103 | -2.384e+02 | 0.0000 | 7080.0 | 24.83% | 25017.5 | 19.04% | motif file (matrix) | svg |
| 132 | G A C T C T A G A T G C A G T C G T C A T A C G A T G C A T C G | HIC1(Zf)/Treg-ZBTB29-ChIP-Seq(GSE99889)/Homer | 1e-102 | -2.360e+02 | 0.0000 | 12636.0 | 44.31% | 49157.7 | 37.41% | motif file (matrix) | svg |
| 133 | C G A T C T G A A G T C A C G T A C G T A T C G G A C T C T A G C G A T G A C T C G T A A T G C C G T A G T C A A C T G | ANAC011(NAC)/col-ANAC011-DAP-Seq(GSE60143)/Homer | 1e-102 | -2.353e+02 | 0.0000 | 3220.0 | 11.29% | 9605.4 | 7.31% | motif file (matrix) | svg |
| 134 | A C G T T C G A T C G A A G T C G T C A T A C G A T G C A C G T A C T G A G C T | Myf5(bHLH)/GM-Myf5-ChIP-Seq(GSE24852)/Homer | 1e-101 | -2.338e+02 | 0.0000 | 3752.0 | 13.16% | 11650.6 | 8.87% | motif file (matrix) | svg |
| 135 | G T A C A C T G A T G C A G T C C T A G G A T C G T A C C T G A G A C T G C A T C G A T G A C T | RAP212(AP2EREBP)/col-RAP212-DAP-Seq(GSE60143)/Homer | 1e-100 | -2.320e+02 | 0.0000 | 8916.0 | 31.27% | 32900.5 | 25.04% | motif file (matrix) | svg |
| 136 | C T A G A G T C T A C G T A C G T G A C C G T A A C T G T A G C G C A T C A T G A T G C A G C T | Ascl1(bHLH)/NeuralTubes-Ascl1-ChIP-Seq(GSE55840)/Homer | 1e-100 | -2.314e+02 | 0.0000 | 7869.0 | 27.59% | 28445.8 | 21.65% | motif file (matrix) | svg |
| 137 | C G T A C G A T C T A G C G T A A G C T C A G T T A C G C G T A A C G T C A T G C T A G A T G C | HOXA3(Homeobox)/mEmbryo-Hoxa3-ChIP-Seq(E-MTAB-8607)/Homer | 1e-99 | -2.297e+02 | 0.0000 | 2339.0 | 8.20% | 6410.0 | 4.88% | motif file (matrix) | svg |
| 138 | C G A T C G T A G T A C A C G T A C G T T C A G G C A T C A G T T A C G G T C A C G T A A G T C C G T A T G C A C A T G | ANAC053(NAC)/colamp-ANAC053-DAP-Seq(GSE60143)/Homer | 1e-98 | -2.277e+02 | 0.0000 | 5701.0 | 19.99% | 19479.6 | 14.83% | motif file (matrix) | svg |
| 139 | A G C T C T A G G A T C A G T C C T A G C T G A A G T C G C T A G C A T T G C A | CBF1(AP2EREBP)/colamp-CBF1-DAP-Seq(GSE60143)/Homer | 1e-98 | -2.274e+02 | 0.0000 | 11271.0 | 39.52% | 43249.6 | 32.92% | motif file (matrix) | svg |
| 140 | G C T A T C G A C G T A C T G A A C T G A C G T A G T C C G T A C G T A A G T C C T A G T G C A | WRKY42(WRKY)/colamp-WRKY42-DAP-Seq(GSE60143)/Homer | 1e-97 | -2.255e+02 | 0.0000 | 5218.0 | 18.30% | 17548.1 | 13.36% | motif file (matrix) | svg |
| 141 | G C T A T C G A C G T A C T A G A G C T G T C A G T C A C G T A A G T C C G T A | FOXA1(Forkhead)/LNCAP-FOXA1-ChIP-Seq(GSE27824)/Homer | 1e-97 | -2.246e+02 | 0.0000 | 7794.0 | 27.33% | 28239.9 | 21.49% | motif file (matrix) | svg |
| 142 | G A T C C A T G A C G T A C G T A C T G C G T A A G T C A G C T C G A T G A C T | WRKY8(WRKY)/colamp-WRKY8-DAP-Seq(GSE60143)/Homer | 1e-97 | -2.240e+02 | 0.0000 | 1503.0 | 5.27% | 3554.8 | 2.71% | motif file (matrix) | svg |
| 143 | T A C G C T G A C A T G G A T C G T A C G C A T T C A G T A C G A G C T G T C A G A T C G C A T T A C G C G T A C T A G G A T C G A T C C G A T A C T G T C A G | ZNF322(Zf)/HEK293-ZNF322.GFP-ChIP-Seq(GSE58341)/Homer | 1e-96 | -2.218e+02 | 0.0000 | 1146.0 | 4.02% | 2418.5 | 1.84% | motif file (matrix) | svg |
| 144 | G A C T A C T G C G A T A G T C A C T G C T A G A G T C C T G A | AT1G12630(AP2EREBP)/colamp-AT1G12630-DAP-Seq(GSE60143)/Homer | 1e-95 | -2.203e+02 | 0.0000 | 8718.0 | 30.57% | 32252.7 | 24.55% | motif file (matrix) | svg |
| 145 | A T G C T C G A T A C G A C G T A T G C A G T C A C G T A G T C A G T C G A T C | Znf263(Zf)/K562-Znf263-ChIP-Seq(GSE31477)/Homer | 1e-95 | -2.197e+02 | 0.0000 | 8716.0 | 30.56% | 32255.9 | 24.55% | motif file (matrix) | svg |
| 146 | T C A G A G C T G T C A C G T A A C G T A T G C C G T A A C G T A C G T C T G A | PHV(HB)/col-PHV-DAP-Seq(GSE60143)/Homer | 1e-94 | -2.183e+02 | 0.0000 | 4725.0 | 16.57% | 15662.5 | 11.92% | motif file (matrix) | svg |
| 147 | A G T C C G T A A C G T A G T C A C G T A C T G | Tal1 | 1e-94 | -2.181e+02 | 0.0000 | 9404.0 | 32.98% | 35251.9 | 26.83% | motif file (matrix) | svg |
| 148 | A G T C G A C T G A T C C G T A G T A C A G T C G C T A C G T A G T A C A G T C G T A C G T A C | MYB63(MYB)/col-MYB63-DAP-Seq(GSE60143)/Homer | 1e-94 | -2.168e+02 | 0.0000 | 5285.0 | 18.53% | 17939.9 | 13.65% | motif file (matrix) | svg |
| 149 | C T G A A T G C G C T A C G A T A T G C C G T A C G T A C G T A C T A G T A C G | Tcf3(HMG)/mES-Tcf3-ChIP-Seq(GSE11724)/Homer | 1e-93 | -2.163e+02 | 0.0000 | 2524.0 | 8.85% | 7196.4 | 5.48% | motif file (matrix) | svg |
| 150 | G T C A C G T A A C G T A T C G C G T A A C G T A C G T C T A G | ATHB7(Homeobox)/col-ATHB7-DAP-Seq(GSE60143)/Homer | 1e-91 | -2.114e+02 | 0.0000 | 8632.0 | 30.27% | 32039.7 | 24.39% | motif file (matrix) | svg |
| 151 | C A G T A C T G T C A G T G C A G C T A A T G C T C G A A T C G G T C A T G C A | ZNF189(Zf)/HEK293-ZNF189.GFP-ChIP-Seq(GSE58341)/Homer | 1e-91 | -2.111e+02 | 0.0000 | 5205.0 | 18.25% | 17694.6 | 13.47% | motif file (matrix) | svg |
| 152 | C T A G C T A G A T G C G T A C T C A G A T G C A G T C G C A T G A T C G A T C | ZNF91(Zf)/HEK-ZNF91.HA-ChIP-Seq(GSE162571)/Homer | 1e-91 | -2.109e+02 | 0.0000 | 4763.0 | 16.70% | 15911.8 | 12.11% | motif file (matrix) | svg |
| 153 | G A C T G T A C T G C A A C G T G A T C G C T A T C G A A C G T A G T C C G T A | Pdx1(Homeobox)/Islet-Pdx1-ChIP-Seq(SRA008281)/Homer | 1e-91 | -2.105e+02 | 0.0000 | 9373.0 | 32.87% | 35255.1 | 26.83% | motif file (matrix) | svg |
| 154 | C G A T A G T C C A T G G A C T A C G T C T A G C G T A G A T C G A C T C G A T G C A T G A C T | WRKY43(WRKY)/colamp-WRKY43-DAP-Seq(GSE60143)/Homer | 1e-90 | -2.093e+02 | 0.0000 | 4228.0 | 14.83% | 13798.4 | 10.50% | motif file (matrix) | svg |
| 155 | A G T C A G T C C T G A A G T C A G T C A C T G C G T A A G T C C T G A T C G A G C A T G A T C C G A T C G A T A C T G | AT3G60490(AP2EREBP)/colamp-AT3G60490-DAP-Seq(GSE60143)/Homer | 1e-90 | -2.086e+02 | 0.0000 | 6173.0 | 21.65% | 21701.8 | 16.52% | motif file (matrix) | svg |
| 156 | C T A G G C T A A G T C A C T G A C G T G A C T G A C T A T G C T C G A C A G T G A T C C G A T G A C T G A T C G A T C | RKD2(RWPRK)/colamp-RKD2-DAP-Seq(GSE60143)/Homer | 1e-89 | -2.070e+02 | 0.0000 | 6611.0 | 23.18% | 23548.7 | 17.92% | motif file (matrix) | svg |
| 157 | A G C T G A T C C T G A A G T C A G T C A C T G C G T A A G T C C T G A G T C A G C A T C G A T G C T A G C A T C G T A | At2g44940(AP2EREBP)/colamp-At2g44940-DAP-Seq(GSE60143)/Homer | 1e-89 | -2.065e+02 | 0.0000 | 5148.0 | 18.05% | 17527.1 | 13.34% | motif file (matrix) | svg |
| 158 | T A G C C A T G A G C T A C G T A C T G C G T A A G T C G A C T G C A T C T G A | AT3G42860(zfGRF)/col-AT3G42860-DAP-Seq(GSE60143)/Homer | 1e-89 | -2.059e+02 | 0.0000 | 3938.0 | 13.81% | 12696.2 | 9.66% | motif file (matrix) | svg |
| 159 | G A T C C T G A A G T C A G T C A C T G C G T A A G T C C T G A | ERF38(AP2EREBP)/col-ERF38-DAP-Seq(GSE60143)/Homer | 1e-87 | -2.023e+02 | 0.0000 | 8319.0 | 29.17% | 30860.2 | 23.49% | motif file (matrix) | svg |
| 160 | T C A G A C G T T C G A T A G C A G T C C G T A A C T G G T A C A C G T A C T G A T C G A G T C | Atoh1(bHLH)/Cerebellum-Atoh1-ChIP-Seq(GSE22111)/Homer | 1e-87 | -2.020e+02 | 0.0000 | 6266.0 | 21.97% | 22189.7 | 16.89% | motif file (matrix) | svg |
| 161 | G C T A C T G A T C G A A G T C A G T C C T G A A G T C G T C A C T G A T G C A | RUNX1(Runt)/Jurkat-RUNX1-ChIP-Seq(GSE29180)/Homer | 1e-87 | -2.015e+02 | 0.0000 | 7660.0 | 26.86% | 28060.1 | 21.36% | motif file (matrix) | svg |
| 162 | C G T A T A G C T A G C T G C A A C T G C T A G C G T A C G T A T C A G G A C T | EHF(ETS)/LoVo-EHF-ChIP-Seq(GSE49402)/Homer | 1e-87 | -2.014e+02 | 0.0000 | 7698.0 | 26.99% | 28224.3 | 21.48% | motif file (matrix) | svg |
| 163 | C G T A C G A T C T A G G T C A G A C T C G A T C T A G C G T A A C G T C A T G | LIN-39(Homeobox)/cElegans.L3-LIN39-ChIP-Seq(modEncode)/Homer | 1e-86 | -1.987e+02 | 0.0000 | 9999.0 | 35.06% | 38206.7 | 29.08% | motif file (matrix) | svg |
| 164 | C A T G T G A C C A T G G A C T C A G T C T A G G C T A G T A C G A C T G C A T G C A T C G A T | WRKY21(WRKY)/colamp-WRKY21-DAP-Seq(GSE60143)/Homer | 1e-85 | -1.961e+02 | 0.0000 | 1468.0 | 5.15% | 3617.9 | 2.75% | motif file (matrix) | svg |
| 165 | C G A T T C G A G A T C A C G T A C G T T C A G G C T A C G A T C G T A C G T A C G T A A T G C C G T A T G C A C T A G | ANAC028(NAC)/col-ANAC028-DAP-Seq(GSE60143)/Homer | 1e-84 | -1.936e+02 | 0.0000 | 6974.0 | 24.46% | 25292.3 | 19.25% | motif file (matrix) | svg |
| 166 | G T A C G C T A C G A T C A G T A G T C G C T A C G A T C G A T A G T C G C T A | WUS1(Homeobox)/colamp-WUS1-DAP-Seq(GSE60143)/Homer | 1e-84 | -1.935e+02 | 0.0000 | 4242.0 | 14.88% | 14057.9 | 10.70% | motif file (matrix) | svg |
| 167 | A G T C A C G T A C T G A G C T A C G T A C G T G T C A A G T C | Foxo1(Forkhead)/RAW-Foxo1-ChIP-Seq(Fan\_et\_al.)/Homer | 1e-84 | -1.935e+02 | 0.0000 | 11322.0 | 39.70% | 44143.3 | 33.60% | motif file (matrix) | svg |
| 168 | G A T C C T G A A G T C G T A C A C T G G C T A G A T C C T G A G C T A G C T A | At4g31060(AP2EREBP)/colamp-At4g31060-DAP-Seq(GSE60143)/Homer | 1e-83 | -1.928e+02 | 0.0000 | 7811.0 | 27.39% | 28857.0 | 21.96% | motif file (matrix) | svg |
| 169 | G T C A T C G A C T A G C T A G A G T C G T C A C G A T C T A G G A C T G A T C G A T C T C A G C T A G C T G A A G T C G C T A C A G T T C A G G A T C G A T C | p63(p53)/Keratinocyte-p63-ChIP-Seq(GSE17611)/Homer | 1e-82 | -1.906e+02 | 0.0000 | 3199.0 | 11.22% | 9999.7 | 7.61% | motif file (matrix) | svg |
| 170 | G C A T G C A T G C A T A T G C A G C T T C G A T A C G G C T A C G T A C A T G G T A C G C A T G C A T A G T C A G C T | HSFA6B(HSF)/colamp-HSFA6B-DAP-Seq(GSE60143)/Homer | 1e-82 | -1.899e+02 | 0.0000 | 4147.0 | 14.54% | 13726.7 | 10.45% | motif file (matrix) | svg |
| 171 | C A G T C G T A C G T A G C A T G A C T G C A T G T A C A G C T A C T G G A C T A C G T C A T G | RAV1(RAV)/colamp-RAV1-DAP-Seq(GSE60143)/Homer | 1e-82 | -1.895e+02 | 0.0000 | 5428.0 | 19.03% | 18920.6 | 14.40% | motif file (matrix) | svg |
| 172 | A G C T T G A C C G A T C G A T C T A G A C G T C A G T C A G T G C T A A G T C | FOXK1(Forkhead)/HEK293-FOXK1-ChIP-Seq(GSE51673)/Homer | 1e-82 | -1.893e+02 | 0.0000 | 8804.0 | 30.87% | 33189.5 | 25.26% | motif file (matrix) | svg |
| 173 | C T G A A T C G G T A C C T G A A G T C A G T C A C T G C G T A A G T C C T G A | TINY(AP2EREBP)/col-TINY-DAP-Seq(GSE60143)/Homer | 1e-81 | -1.883e+02 | 0.0000 | 5401.0 | 18.94% | 18827.1 | 14.33% | motif file (matrix) | svg |
| 174 | C T G A A C T G C G T A A C G T G T C A A G C T A G C T G A C T G A C T C A G T | CCA(Myb)/Arabidopsis-CCA.GFP-ChIP-Seq(GSE70533)/Homer | 1e-81 | -1.870e+02 | 0.0000 | 9096.0 | 31.90% | 34497.9 | 26.26% | motif file (matrix) | svg |
| 175 | A G T C A C G T A C G T T A C G G C A T G C A T A T G C G C T A C G T A A T G C C G T A G T C A A C T G G A T C G C A T | ANAC075(NAC)/col-ANAC075-DAP-Seq(GSE60143)/Homer | 1e-81 | -1.867e+02 | 0.0000 | 4339.0 | 15.22% | 14537.5 | 11.06% | motif file (matrix) | svg |
| 176 | G T A C T C G A T A G C C G T A C G T A C T G A T G C A T G A C A C T G C G T A A G T C C T G A C T G A T C G A C G T A | NUC(C2H2)/col-NUC-DAP-Seq(GSE60143)/Homer | 1e-79 | -1.841e+02 | 0.0000 | 1191.0 | 4.18% | 2771.9 | 2.11% | motif file (matrix) | svg |
| 177 | G C A T C G T A C G A T G A C T A C T G C T G A G A C T G A T C | Hnf6b(Homeobox)/LNCaP-Hnf6b-ChIP-Seq(GSE106305)/Homer | 1e-79 | -1.830e+02 | 0.0000 | 12045.0 | 42.24% | 47588.6 | 36.22% | motif file (matrix) | svg |
| 178 | C G T A G A T C C T A G A C G T G T A C C T G A A G C T G A T C G C T A G A C T | TGA2(bZIP)/colamp-TGA2-DAP-Seq(GSE60143)/Homer | 1e-79 | -1.826e+02 | 0.0000 | 10205.0 | 35.79% | 39431.2 | 30.01% | motif file (matrix) | svg |
| 179 | A G C T G A T C G A T C C G T A G T A C A G T C C G A T C T G A G T A C G A T C C G T A G A C T | ATY19(MYB)/col-ATY19-DAP-Seq(GSE60143)/Homer | 1e-79 | -1.826e+02 | 0.0000 | 7398.0 | 25.94% | 27281.7 | 20.76% | motif file (matrix) | svg |
| 180 | G A C T C T A G C T A G G T A C A G T C G A T C G A C T G A C T T A G C T C A G | NLP7(RWPRK)/col-NLP7-DAP-Seq(GSE60143)/Homer | 1e-78 | -1.818e+02 | 0.0000 | 13126.0 | 46.03% | 52476.5 | 39.94% | motif file (matrix) | svg |
| 181 | A C G T C T A G A G C T A C G T A C G T C T G A A G T C G A C T A G C T C G T A | FOXM1(Forkhead)/MCF7-FOXM1-ChIP-Seq(GSE72977)/Homer | 1e-78 | -1.812e+02 | 0.0000 | 6832.0 | 23.96% | 24908.0 | 18.96% | motif file (matrix) | svg |
| 182 | A G T C G A T C C T G A A G T C A G T C C A T G G T C A G A T C C G T A G A T C | DREB26(AP2EREBP)/col-DREB26-DAP-Seq(GSE60143)/Homer | 1e-78 | -1.807e+02 | 0.0000 | 3096.0 | 10.86% | 9715.7 | 7.39% | motif file (matrix) | svg |
| 183 | T G A C C A T G A C G T A C G T A C T G C G T A A G T C A G C T G C A T T C G A | WRKY30(WRKY)/colamp-WRKY30-DAP-Seq(GSE60143)/Homer | 1e-78 | -1.806e+02 | 0.0000 | 4207.0 | 14.75% | 14091.2 | 10.72% | motif file (matrix) | svg |
| 184 | G A T C C T G A G A T C G A T C C T A G G C T A A G T C C T G A G C T A C G T A | At4g16750(AP2EREBP)/col-At4g16750-DAP-Seq(GSE60143)/Homer | 1e-78 | -1.799e+02 | 0.0000 | 12587.0 | 44.14% | 50086.1 | 38.12% | motif file (matrix) | svg |
| 185 | T A C G C G T A G A C T T C A G A G C T A G T C A C T G T C A G A G T C C T G A | DDF2(AP2EREBP)/col-DDF2-DAP-Seq(GSE60143)/Homer | 1e-77 | -1.790e+02 | 0.0000 | 1717.0 | 6.02% | 4604.2 | 3.50% | motif file (matrix) | svg |
| 186 | C T G A T A C G G C A T A G C T A G C T A G T C T C G A A C T G C A G T A G C T A G C T G A T C | IRF3(IRF)/BMDM-Irf3-ChIP-Seq(GSE67343)/Homer | 1e-77 | -1.788e+02 | 0.0000 | 1876.0 | 6.58% | 5172.9 | 3.94% | motif file (matrix) | svg |
| 187 | G C T A C G T A C G T A G C A T C A T G C T A G G A T C A C T G T C A G G A T C A C T G T C A G | ERF4(AP2EREBP)/colamp-ERF4-DAP-Seq(GSE60143)/Homer | 1e-77 | -1.787e+02 | 0.0000 | 12018.0 | 42.14% | 47560.7 | 36.20% | motif file (matrix) | svg |
| 188 | C G T A A C G T A C G T A C G T A C G T A G T C A G T C C T G A A G C T A G C T | NFAT(RHD)/Jurkat-NFATC1-ChIP-Seq(Jolma\_et\_al.)/Homer | 1e-77 | -1.782e+02 | 0.0000 | 6396.0 | 22.43% | 23123.3 | 17.60% | motif file (matrix) | svg |
| 189 | A G T C G T A C C T G A A G T C G T A C C T A G G C T A T G A C T G C A G C T A C G T A C G T A | At1g22810(AP2EREBP)/colamp-At1g22810-DAP-Seq(GSE60143)/Homer | 1e-76 | -1.769e+02 | 0.0000 | 7194.0 | 25.23% | 26516.4 | 20.18% | motif file (matrix) | svg |
| 190 | G A T C C A G T T A G C A G T C A C T G A G T C A G T C C T A G G A C T G T A C | LEP(AP2EREBP)/col-LEP-DAP-Seq(GSE60143)/Homer | 1e-76 | -1.767e+02 | 0.0000 | 2779.0 | 9.75% | 8543.6 | 6.50% | motif file (matrix) | svg |
| 191 | G C T A T C G A C G T A C T A G A G C T G T C A G T C A C G T A A G T C C G T A | FOXA1(Forkhead)/MCF7-FOXA1-ChIP-Seq(GSE26831)/Homer | 1e-76 | -1.756e+02 | 0.0000 | 6125.0 | 21.48% | 22033.2 | 16.77% | motif file (matrix) | svg |
| 192 | C G T A C T A G T C A G T C A G A G T C A T G C A G T C G C A T A G C T A C G T A T C G C G A T | Sox9(HMG)/Limb-SOX9-ChIP-Seq(GSE73225)/Homer | 1e-76 | -1.752e+02 | 0.0000 | 6218.0 | 21.80% | 22429.0 | 17.07% | motif file (matrix) | svg |
| 193 | C T A G A C T G T G C A G T C A A T G C C G T A A T C G A T G C A G T C C T A G | ZNF341(Zf)/EBV-ZNF341-ChIP-Seq(GSE113194)/Homer | 1e-75 | -1.745e+02 | 0.0000 | 4693.0 | 16.46% | 16140.0 | 12.28% | motif file (matrix) | svg |
| 194 | G C A T C G T A G C A T C G T A T C G A C G T A C T G A A C T G C G T A C G T A C G T A A C G T A C T G G T C A G C A T | AT2G31460(REMB3)/col-AT2G31460-DAP-Seq(GSE60143)/Homer | 1e-75 | -1.739e+02 | 0.0000 | 2759.0 | 9.67% | 8498.1 | 6.47% | motif file (matrix) | svg |
| 195 | C G A T C T G A A G T C A C G T A C G T T A C G G C T A C A T G C T A G G C A T C G A T A G T C C G T A G T C A A C T G | ANAC096(NAC)/colamp-ANAC096-DAP-Seq(GSE60143)/Homer | 1e-74 | -1.724e+02 | 0.0000 | 6296.0 | 22.08% | 22803.2 | 17.36% | motif file (matrix) | svg |
| 196 | T G A C A G T C C G T A A C T G G T A C A C G T A C T G A C G T G A C T G A T C | Twist2(bHLH)/Myoblast-Twist2.Ty1-ChIP-Seq(GSE127998)/Homer | 1e-74 | -1.723e+02 | 0.0000 | 9761.0 | 34.23% | 37695.0 | 28.69% | motif file (matrix) | svg |
| 197 | C G T A G C T A C G A T C T A G A C G T G T C A C G T A C G T A A G T C C G T A T G C A T A C G | FoxL2(Forkhead)/Ovary-FoxL2-ChIP-Seq(GSE60858)/Homer | 1e-74 | -1.716e+02 | 0.0000 | 6649.0 | 23.32% | 24303.7 | 18.50% | motif file (matrix) | svg |
| 198 | G C T A G C T A C T G A A C T G A C G T A G T C C G T A C G T A G T A C A C T G A T G C G C A T | WRKY47(WRKY)/colamp-WRKY47-DAP-Seq(GSE60143)/Homer | 1e-74 | -1.714e+02 | 0.0000 | 3846.0 | 13.49% | 12773.2 | 9.72% | motif file (matrix) | svg |
| 199 | C G A T T C G A G A T C A C G T A C G T T C A G G C A T C T G A C G T A G C T A C G T A A G T C C G T A T G C A C A T G | ANAC050(NAC)/colamp-ANAC050-DAP-Seq(GSE60143)/Homer | 1e-74 | -1.710e+02 | 0.0000 | 5913.0 | 20.74% | 21224.4 | 16.15% | motif file (matrix) | svg |
| 200 | C T G A T G A C T G A C C G T A A C G T T G A C A G C T C T A G A C G T G A C T | Olig2(bHLH)/Neuron-Olig2-ChIP-Seq(GSE30882)/Homer | 1e-74 | -1.710e+02 | 0.0000 | 11832.0 | 41.49% | 46897.9 | 35.69% | motif file (matrix) | svg |
| 201 | C G A T C T A G C T G A A T G C C T G A T C G A C G T A C T G A T C G A T A G C A G T C C G T A A C T G T C G A A T G C | Hand2(bHLH)/Mesoderm-Hand2-ChIP-Seq(GSE61475)/Homer | 1e-74 | -1.709e+02 | 0.0000 | 2648.0 | 9.29% | 8108.5 | 6.17% | motif file (matrix) | svg |
| 202 | T C G A T G A C G T A C C G T A C A G T T G A C A C G T A C T G A G C T A G C T | NeuroG2(bHLH)/Fibroblast-NeuroG2-ChIP-Seq(GSE75910)/Homer | 1e-74 | -1.707e+02 | 0.0000 | 9058.0 | 31.76% | 34657.2 | 26.38% | motif file (matrix) | svg |
| 203 | G A C T C T G A A G T C A G T C A C T G C G T A A G T C C T G A | bHLH10(bHLH)/colamp-bHLH10-DAP-Seq(GSE60143)/Homer | 1e-73 | -1.698e+02 | 0.0000 | 5870.0 | 20.58% | 21065.7 | 16.03% | motif file (matrix) | svg |
| 204 | T G A C C G A T C T G A C T A G C T A G A C G T A T G C T G C A T C G A C T G A C T A G C A T G A C G T A G T C C G T A | PPARa(NR),DR1/Liver-Ppara-ChIP-Seq(GSE47954)/Homer | 1e-72 | -1.679e+02 | 0.0000 | 5640.0 | 19.78% | 20140.5 | 15.33% | motif file (matrix) | svg |
| 205 | G T C A T G C A T G C A G C T A C G T A G C T A G C T A G C T A | REM19(REM)/colamp-REM19-DAP-Seq(GSE60143)/Homer | 1e-72 | -1.673e+02 | 0.0000 | 2767.0 | 9.70% | 8602.0 | 6.55% | motif file (matrix) | svg |
| 206 | T A C G T A G C G C T A C G A T C T A G A C G T C A G T C A G T G C T A A G T C G T C A G C A T | FOXK2(Forkhead)/U2OS-FOXK2-ChIP-Seq(E-MTAB-2204)/Homer | 1e-72 | -1.672e+02 | 0.0000 | 5649.0 | 19.81% | 20188.6 | 15.37% | motif file (matrix) | svg |
| 207 | A C T G A C G T C G A T C A G T C A T G C A T G C A G T G C A T C A G T C A T G | HuR(?)/HEK293-HuR-CLIP-Seq(GSE87887)/Homer | 1e-72 | -1.672e+02 | 0.0000 | 18049.0 | 63.29% | 75549.4 | 57.50% | motif file (matrix) | svg |
| 208 | G A T C G T A C C T G A A G T C A G T C A C T G G C T A G T A C G T C A G C A T G C A T C G A T | DEAR2(AP2EREBP)/colamp-DEAR2-DAP-Seq(GSE60143)/Homer | 1e-72 | -1.669e+02 | 0.0000 | 14039.0 | 49.23% | 56959.4 | 43.35% | motif file (matrix) | svg |
| 209 | C T A G T C G A T G A C A G T C C G T A A C T G G T A C A C G T A C T G A C T G | BHLHA15(bHLH)/NIH3T3-BHLHB8.HA-ChIP-Seq(GSE119782)/Homer | 1e-71 | -1.657e+02 | 0.0000 | 7691.0 | 26.97% | 28848.4 | 21.96% | motif file (matrix) | svg |
| 210 | T C A G T G A C G T A C C G T A A C G T T G A C A C G T T C A G A G C T G A C T | NeuroD1(bHLH)/Islet-NeuroD1-ChIP-Seq(GSE30298)/Homer | 1e-71 | -1.657e+02 | 0.0000 | 4649.0 | 16.30% | 16093.6 | 12.25% | motif file (matrix) | svg |
| 211 | C A G T T C A G A G C T G A C T A C G T A G T C G A T C G A C T C T G A A C T G G A T C C G T A C T G A A G T C G T A C | Rfx6(HTH)/Min6b1-Rfx6.HA-ChIP-Seq(GSE62844)/Homer | 1e-71 | -1.653e+02 | 0.0000 | 7125.0 | 24.99% | 26437.3 | 20.12% | motif file (matrix) | svg |
| 212 | A T G C G T A C A C T G A G T C A G T C A C T G A G T C G T A C | ERF73(AP2EREBP)/col-ERF73-DAP-Seq(GSE60143)/Homer | 1e-71 | -1.653e+02 | 0.0000 | 4943.0 | 17.33% | 17302.4 | 13.17% | motif file (matrix) | svg |
| 213 | G T A C G T C A G T A C G T C A G T A C G T C A G T A C G T C A G T A C G T C A | SeqBias: CA-repeat | 1e-71 | -1.640e+02 | 0.0000 | 21256.0 | 74.54% | 90985.7 | 69.25% | motif file (matrix) | svg |
| 214 | C G A T T C G A G A T C C G A T G C A T T C A G G A C T C G A T G C A T G C T A C T G A A G T C C G T A G T C A C T A G | ANAC005(NAC)/col-ANAC005-DAP-Seq(GSE60143)/Homer | 1e-71 | -1.639e+02 | 0.0000 | 3918.0 | 13.74% | 13164.4 | 10.02% | motif file (matrix) | svg |
| 215 | C A G T C A T G T G C A G T A C C G T A T C A G G T A C G A C T T C A G C T G A | bZIP18(bZIP)/colamp-bZIP18-DAP-Seq(GSE60143)/Homer | 1e-71 | -1.638e+02 | 0.0000 | 22798.0 | 79.95% | 98574.0 | 75.03% | motif file (matrix) | svg |
| 216 | T C A G T C A G A C G T G T A C G C T A T C A G C T G A A C T G A C T G A G C T A G T C C G T A | EAR2(NR)/K562-NR2F6-ChIP-Seq(Encode)/Homer | 1e-69 | -1.595e+02 | 0.0000 | 8812.0 | 30.90% | 33817.9 | 25.74% | motif file (matrix) | svg |
| 217 | C G T A G C A T C A T G C T A G A G T C A C T G A T C G G T A C A C T G T C A G | At2g33710(AP2EREBP)/colamp-At2g33710-DAP-Seq(GSE60143)/Homer | 1e-68 | -1.581e+02 | 0.0000 | 16531.0 | 57.97% | 68655.4 | 52.25% | motif file (matrix) | svg |
| 218 | C T G A G A C T C A T G C T A G A G T C A C T G A C T G A G T C A C T G T C A G | ERF11(AP2EREBP)/col-ERF11-DAP-Seq(GSE60143)/Homer | 1e-68 | -1.579e+02 | 0.0000 | 9784.0 | 34.31% | 38105.3 | 29.00% | motif file (matrix) | svg |
| 219 | C A G T A T C G C T G A A G T C T C A G C A G T T A G C C T G A A T G C T A C G | FEA4(bZIP)/Corn-FEA4-ChIP-Seq(GSE61954)/Homer | 1e-68 | -1.570e+02 | 0.0000 | 10805.0 | 37.89% | 42636.8 | 32.45% | motif file (matrix) | svg |
| 220 | C G T A C G A T C A G T C A T G C G A T G T A C C A T G A C T G G A C T C A T G | CEJ1(AP2EREBP)/col-CEJ1-DAP-Seq(GSE60143)/Homer | 1e-67 | -1.564e+02 | 0.0000 | 13578.0 | 47.61% | 55111.3 | 41.95% | motif file (matrix) | svg |
| 221 | C G T A C G T A G C A T A C T G C G T A A G C T C T G A C G T A T A C G C T G A | ELT-3(Gata)/cElegans-L1-ELT3-ChIP-Seq(modEncode)/Homer | 1e-67 | -1.562e+02 | 0.0000 | 5085.0 | 17.83% | 18033.2 | 13.73% | motif file (matrix) | svg |
| 222 | C A T G T G C A G A C T C A T G C G T A A G T C T C A G G C A T T G A C C G T A | bZIP50(bZIP)/colamp-bZIP50-DAP-Seq(GSE60143)/Homer | 1e-67 | -1.558e+02 | 0.0000 | 13401.0 | 46.99% | 54320.4 | 41.34% | motif file (matrix) | svg |
| 223 | C T A G A C T G A C G T C G T A A C T G A C T G A G C T C T A G T C A G C T A G | MYB93(MYB)/colamp-MYB93-DAP-Seq(GSE60143)/Homer | 1e-67 | -1.550e+02 | 0.0000 | 10500.0 | 36.82% | 41329.9 | 31.46% | motif file (matrix) | svg |
| 224 | C G T A T G A C T A G C T G C A A C T G A C T G C G T A C G T A T C A G G A C T | ELF3(ETS)/PDAC-ELF3-ChIP-Seq(GSE64557)/Homer | 1e-66 | -1.540e+02 | 0.0000 | 4121.0 | 14.45% | 14124.4 | 10.75% | motif file (matrix) | svg |
| 225 | G A C T A C T G C A G T A G T C A C T G A C T G A G C T A C T G C T A G G T C A | At1g77640(AP2EREBP)/col-At1g77640-DAP-Seq(GSE60143)/Homer | 1e-66 | -1.539e+02 | 0.0000 | 2638.0 | 9.25% | 8260.4 | 6.29% | motif file (matrix) | svg |
| 226 | T C A G T C A G G C T A C G T A T A C G G A C T T C A G T C G A C T G A C G T A T A C G G A C T | IRF8(IRF)/BMDM-IRF8-ChIP-Seq(GSE77884)/Homer | 1e-65 | -1.518e+02 | 0.0000 | 2083.0 | 7.30% | 6179.8 | 4.70% | motif file (matrix) | svg |
| 227 | A G T C G A C T C A G T A C T G C T A G T G A C G C T A A T G C G C A T A T C G C G A T A C T G G A T C G T A C G T C A C T G A | NF1(CTF)/LNCAP-NF1-ChIP-Seq(Unpublished)/Homer | 1e-65 | -1.517e+02 | 0.0000 | 2021.0 | 7.09% | 5949.8 | 4.53% | motif file (matrix) | svg |
| 228 | A G T C C G A T A C T G A T C G T G A C G C T A C A T G A T C G T G A C C G A T A C T G T A G C G T A C G T C A | Tlx?(NR)/NPC-H3K4me1-ChIP-Seq(GSE16256)/Homer | 1e-65 | -1.515e+02 | 0.0000 | 1851.0 | 6.49% | 5324.2 | 4.05% | motif file (matrix) | svg |
| 229 | T A C G T G C A A G T C C G T A A C G T T G A C A C G T A C T G A C T G G C A T | TCF4(bHLH)/SHSY5Y-TCF4-ChIP-Seq(GSE96915)/Homer | 1e-65 | -1.514e+02 | 0.0000 | 8723.0 | 30.59% | 33601.7 | 25.57% | motif file (matrix) | svg |
| 230 | A G T C T G A C C T G A A G T C A G T C A C T G C G T A A G T C G T C A G C T A G C A T C G T A G C A T G C T A C G T A | DEAR3(AP2EREBP)/colamp-DEAR3-DAP-Seq(GSE60143)/Homer | 1e-65 | -1.507e+02 | 0.0000 | 6515.0 | 22.85% | 24121.6 | 18.36% | motif file (matrix) | svg |
| 231 | G A T C G A C T G A C T A C G T A G T C A C G T A G T C A C G T A G T C A C G T A G T C A C G T G T A C C G A T G T C A | BPC6(BBRBPC)/col-BPC6-DAP-Seq(GSE60143)/Homer | 1e-65 | -1.497e+02 | 0.0000 | 226.0 | 0.79% | 180.7 | 0.14% | motif file (matrix) | svg |
| 232 | C T G A A G T C G A T C C A T G G C T A G A T C C T G A G C T A G C T A C G A T | AT1G77200(AP2EREBP)/colamp-AT1G77200-DAP-Seq(GSE60143)/Homer | 1e-64 | -1.494e+02 | 0.0000 | 12785.0 | 44.83% | 51685.5 | 39.34% | motif file (matrix) | svg |
| 233 | C G T A C G T A C G A T A C T G C A G T A G T C A C T G A C T G A G C T A C T G | DREB19(AP2EREBP)/colamp-DREB19-DAP-Seq(GSE60143)/Homer | 1e-63 | -1.464e+02 | 0.0000 | 8557.0 | 30.01% | 32986.6 | 25.11% | motif file (matrix) | svg |
| 234 | C T G A C G A T C T A G C G T A A G C T C G A T C A G T C T G A G A C T C T A G C T A G A T G C | PBX2(Homeobox)/K562-PBX2-ChIP-Seq(Encode)/Homer | 1e-62 | -1.448e+02 | 0.0000 | 9041.0 | 31.70% | 35134.9 | 26.74% | motif file (matrix) | svg |
| 235 | A G C T C T G A C T A G C T A G A C T G T A G C T G C A T C G A C T G A C T A G C A T G A C G T A T G C T C G A | RXR(NR),DR1/3T3L1-RXR-ChIP-Seq(GSE13511)/Homer | 1e-62 | -1.437e+02 | 0.0000 | 5212.0 | 18.28% | 18773.5 | 14.29% | motif file (matrix) | svg |
| 236 | G A T C G C T A A G T C A C G T A C T G C G T A A G T C C G T A G C T A C G A T C A G T G C A T G C T A C G T A G C A T | GRF9(GRF)/colamp-GRF9-DAP-Seq(GSE60143)/Homer | 1e-61 | -1.417e+02 | 0.0000 | 7289.0 | 25.56% | 27604.7 | 21.01% | motif file (matrix) | svg |
| 237 | A G T C G A T C A G T C C G T A A T C G C A G T A G T C G T A C C T G A A C T G T C A G A G C T A G C T A G C T A G C T | PRDM15(Zf)/ESC-Prdm15-ChIP-Seq(GSE73694)/Homer | 1e-60 | -1.402e+02 | 0.0000 | 6781.0 | 23.78% | 25457.1 | 19.38% | motif file (matrix) | svg |
| 238 | T C G A A G T C A C G T A C G T T C A G C A G T C T G A C T A G T C G A C G T A A T C G C G T A C G T A A C T G A G C T | NTM1(NAC)/col-NTM1-DAP-Seq(GSE60143)/Homer | 1e-60 | -1.391e+02 | 0.0000 | 3901.0 | 13.68% | 13458.9 | 10.24% | motif file (matrix) | svg |
| 239 | G A C T G C T A T G C A A G T C A C G T A C G T A C G T C G A T A C G T T A C G | At3g45610(C2C2dof)/col-At3g45610-DAP-Seq(GSE60143)/Homer | 1e-60 | -1.385e+02 | 0.0000 | 10457.0 | 36.67% | 41522.6 | 31.60% | motif file (matrix) | svg |
| 240 | G C A T G A T C T C A G G C T A G A C T A G T C C T A G C G T A C A T G G T C A | GATA20(C2C2gata)/colamp-GATA20-DAP-Seq(GSE60143)/Homer | 1e-60 | -1.383e+02 | 0.0000 | 20717.0 | 72.65% | 88975.2 | 67.72% | motif file (matrix) | svg |
| 241 | C G A T G A T C G A T C C T G A G A T C G A T C C A T G T G C A G T A C T C G A G T C A G C A T C G A T C G A T G C A T | At4g32800(AP2EREBP)/colamp-At4g32800-DAP-Seq(GSE60143)/Homer | 1e-59 | -1.381e+02 | 0.0000 | 3097.0 | 10.86% | 10249.4 | 7.80% | motif file (matrix) | svg |
| 242 | C G A T T C G A G T A C A C G T A C G T T C A G G C A T G C A T G C T A C G T A C G T A A G T C C G T A T G C A C A T G | ANAC020(NAC)/col-ANAC020-DAP-Seq(GSE60143)/Homer | 1e-59 | -1.375e+02 | 0.0000 | 6641.0 | 23.29% | 24915.9 | 18.96% | motif file (matrix) | svg |
| 243 | T C G A G C A T A C G T C T A G G T A C T C G A G C A T T G A C T C G A A C G T | Chop(bZIP)/MEF-Chop-ChIP-Seq(GSE35681)/Homer | 1e-59 | -1.371e+02 | 0.0000 | 2789.0 | 9.78% | 9050.8 | 6.89% | motif file (matrix) | svg |
| 244 | G A T C A T G C A G T C C G T A A G T C A G T C A C T G G C T A A G T C C G T A | AT1G44830(AP2EREBP)/col-AT1G44830-DAP-Seq(GSE60143)/Homer | 1e-59 | -1.368e+02 | 0.0000 | 3914.0 | 13.73% | 13546.2 | 10.31% | motif file (matrix) | svg |
| 245 | C T A G C A T G G A C T C G T A C T A G A C T G A C G T C T A G C T A G T C A G | MYB17(MYB)/colamp-MYB17-DAP-Seq(GSE60143)/Homer | 1e-59 | -1.367e+02 | 0.0000 | 5695.0 | 19.97% | 20922.1 | 15.92% | motif file (matrix) | svg |
| 246 | A T C G T C G A G A C T A T C G T G A C A C G T C T A G A C T G C G T A A C T G A G T C G T A C | ZNF415(Zf)/HEK293-ZNF415.GFP-ChIP-Seq(GSE58341)/Homer | 1e-59 | -1.367e+02 | 0.0000 | 3881.0 | 13.61% | 13414.1 | 10.21% | motif file (matrix) | svg |
| 247 | C A G T G C T A G C A T T A C G C T G A C A G T T A G C C T G A | GATA15(C2C2gata)/col-GATA15-DAP-Seq(GSE60143)/Homer | 1e-59 | -1.363e+02 | 0.0000 | 15474.0 | 54.26% | 64305.8 | 48.94% | motif file (matrix) | svg |
| 248 | C T G A C T G A C T G A A T G C G A T C C A T G A C T G G A C T G A C T G C A T C G T A C G T A A G T C G T A C C T G A A T C G G C A T G A C T G A C T A G C T | GRHL2(CP2)/HBE-GRHL2-ChIP-Seq(GSE46194)/Homer | 1e-58 | -1.347e+02 | 0.0000 | 3407.0 | 11.95% | 11532.9 | 8.78% | motif file (matrix) | svg |
| 249 | G C A T C T A G G T A C A G T C C G A T A C T G C T A G C T A G G T A C G C T A | ZNF416(Zf)/HEK293-ZNF416.GFP-ChIP-Seq(GSE58341)/Homer | 1e-57 | -1.335e+02 | 0.0000 | 5539.0 | 19.42% | 20325.5 | 15.47% | motif file (matrix) | svg |
| 250 | T A G C T A G C G A C T C T A G A G C T A G T C G T C A T G C A A C G T A T G C G C T A T G C A | Pbx3(Homeobox)/GM12878-PBX3-ChIP-Seq(GSE32465)/Homer | 1e-57 | -1.320e+02 | 0.0000 | 1982.0 | 6.95% | 6006.3 | 4.57% | motif file (matrix) | svg |
| 251 | C G A T C A G T C T A G G C T A A G T C C G T A T C A G A G T C A C G T A C T G A C G T G T A C G C T A G C T A G C T A | bZIP52(bZIP)/colamp-bZIP52-DAP-Seq(GSE60143)/Homer | 1e-57 | -1.320e+02 | 0.0000 | 8797.0 | 30.85% | 34358.5 | 26.15% | motif file (matrix) | svg |
| 252 | A C T G A C T G A G T C A C T G A C T G A G T C A C G T C T A G | ERF1(AP2EREBP)/colamp-ERF1-DAP-Seq(GSE60143)/Homer | 1e-57 | -1.319e+02 | 0.0000 | 4393.0 | 15.40% | 15584.6 | 11.86% | motif file (matrix) | svg |
| 253 | G A T C G A T C G A T C C G T A G T A C A G T C G C A T C G T A G T A C G A T C | MYB58(MYB)/colamp-MYB58-DAP-Seq(GSE60143)/Homer | 1e-57 | -1.313e+02 | 0.0000 | 9732.0 | 34.13% | 38486.4 | 29.29% | motif file (matrix) | svg |
| 254 | C G T A C G T A C G T A C G T A C G T A A C T G A C T G A G T C | dof42(C2C2dof)/col-dof42-DAP-Seq(GSE60143)/Homer | 1e-56 | -1.307e+02 | 0.0000 | 5644.0 | 19.79% | 20821.6 | 15.85% | motif file (matrix) | svg |
| 255 | C G A T C G A T G C A T G A C T A C G T C G T A C G T A A C T G T A G C C G T A C G T A C G T A | AT5G60130(ABI3VP1)/col-AT5G60130-DAP-Seq(GSE60143)/Homer | 1e-56 | -1.303e+02 | 0.0000 | 9611.0 | 33.70% | 37975.8 | 28.90% | motif file (matrix) | svg |
| 256 | C T G A C T G A C T A G T C G A C G T A A T G C C G T A A C T G C G T A A C G T C T G A C G A T A G C T C G T A A C G T A G T C C G A T T A C G G T C A G C A T | GATA(Zf),IR3/iTreg-Gata3-ChIP-Seq(GSE20898)/Homer | 1e-56 | -1.299e+02 | 0.0000 | 1866.0 | 6.54% | 5594.0 | 4.26% | motif file (matrix) | svg |
| 257 | G A C T A G T C C G A T A C T G C T G A T G A C G T A C C G T A A T C G G C A T C T G A C T A G | Bcl11a(Zf)/HSPC-BCL11A-ChIP-Seq(GSE104676)/Homer | 1e-55 | -1.277e+02 | 0.0000 | 4335.0 | 15.20% | 15416.1 | 11.73% | motif file (matrix) | svg |
| 258 | C G T A C T A G C A G T A C G T C G T A A C T G C A T G G C A T T C A G C T G A | MYB49(MYB)/col-MYB49-DAP-Seq(GSE60143)/Homer | 1e-55 | -1.276e+02 | 0.0000 | 10072.0 | 35.32% | 40082.2 | 30.51% | motif file (matrix) | svg |
| 259 | G C T A A C T G T C G A C T G A C T G A A C G T T A G C C T G A C G T A C G A T | Cux2(Homeobox)/Liver-Cux2-ChIP-Seq(GSE35985)/Homer | 1e-55 | -1.272e+02 | 0.0000 | 8544.0 | 29.96% | 33363.8 | 25.39% | motif file (matrix) | svg |
| 260 | C G A T G C T A G C T A G C A T G C T A C G T A A G T C A C G T A C G T A C G T C G A T A G C T | At5g62940(C2C2dof)/col-At5g62940-DAP-Seq(GSE60143)/Homer | 1e-54 | -1.261e+02 | 0.0000 | 18874.0 | 66.19% | 80479.2 | 61.25% | motif file (matrix) | svg |
| 261 | T A C G A T G C G A C T A C T G A G C T A G T C G T C A T G C A A C G T A G T C G C T A T G C A | Pknox1(Homeobox)/ES-Prep1-ChIP-Seq(GSE63282)/Homer | 1e-54 | -1.259e+02 | 0.0000 | 2146.0 | 7.53% | 6696.3 | 5.10% | motif file (matrix) | svg |
| 262 | C T A G C A G T C G T A A C G T A G T C A C T G C G T A A G C T A G T C G A T C | HNF6(Homeobox)/Liver-Hnf6-ChIP-Seq(ERP000394)/Homer | 1e-54 | -1.259e+02 | 0.0000 | 10504.0 | 36.83% | 42042.4 | 32.00% | motif file (matrix) | svg |
| 263 | C T A G T C G A A C G T A C G T C A T G A G T C C T G A C G A T A G T C C G T A | AARE(HLH)/mES-cMyc-ChIP-Seq/Homer | 1e-54 | -1.252e+02 | 0.0000 | 1426.0 | 5.00% | 4017.0 | 3.06% | motif file (matrix) | svg |
| 264 | G A C T C T A G G A T C C A G T A C T G C T G A A T G C G C A T A T G C C T G A | MafA(bZIP)/Islet-MafA-ChIP-Seq(GSE30298)/Homer | 1e-54 | -1.250e+02 | 0.0000 | 5808.0 | 20.37% | 21625.4 | 16.46% | motif file (matrix) | svg |
| 265 | C G A T C G T A A G T C A C G T A C G T T C G A T G C A G C A T G C T A C G T A A C G T A G C T C G T A C G T A A C T G | ANAC062(NAC)/colamp-ANAC062-DAP-Seq(GSE60143)/Homer | 1e-53 | -1.239e+02 | 0.0000 | 3182.0 | 11.16% | 10788.6 | 8.21% | motif file (matrix) | svg |
| 266 | G C T A C G T A C G T A G C A T C A T G C T A G A G T C A C T G T A C G A G T C C A T G T A C G | RAP26(AP2EREBP)/colamp-RAP26-DAP-Seq(GSE60143)/Homer | 1e-52 | -1.220e+02 | 0.0000 | 13189.0 | 46.25% | 54224.3 | 41.27% | motif file (matrix) | svg |
| 267 | C G T A C G A T C G T A C G A T C A T G A C T G C G A T A G T C A T C G T C A G G A C T A C T G | At1g36060(AP2EREBP)/colamp-At1g36060-DAP-Seq(GSE60143)/Homer | 1e-52 | -1.212e+02 | 0.0000 | 9478.0 | 33.24% | 37613.3 | 28.63% | motif file (matrix) | svg |
| 268 | A C T G C T A G A G T C A C T G A C T G A T G C A C T G T A C G | ESE1(AP2EREBP)/col-ESE1-DAP-Seq(GSE60143)/Homer | 1e-52 | -1.205e+02 | 0.0000 | 5385.0 | 18.88% | 19927.0 | 15.17% | motif file (matrix) | svg |
| 269 | G T A C A C G T A C G T T C A G G A C T G C A T T C A G C G T A C T G A A G T C C G T A G T C A A C T G A C G T G C T A | NTM2(NAC)/col-NTM2-DAP-Seq(GSE60143)/Homer | 1e-52 | -1.202e+02 | 0.0000 | 5324.0 | 18.67% | 19676.1 | 14.98% | motif file (matrix) | svg |
| 270 | C A T G A C T G C T A G T C G A T C G A T C G A T C G A T C A G T C A G T C A G T G A C T G A C C G T A A C T G T G C A C G A T A C T G | RBPJ:Ebox(?,bHLH)/Panc1-Rbpj1-ChIP-Seq(GSE47459)/Homer | 1e-52 | -1.199e+02 | 0.0000 | 1325.0 | 4.65% | 3700.7 | 2.82% | motif file (matrix) | svg |
| 271 | T C A G A C T G A C G T C G T A A C T G A C T G A C G T C T A G | MYB51(MYB)/col-MYB51-DAP-Seq(GSE60143)/Homer | 1e-51 | -1.192e+02 | 0.0000 | 8597.0 | 30.15% | 33785.9 | 25.71% | motif file (matrix) | svg |
| 272 | C G T A A C T G C G T A A C G T A T C G C A G T T A G C C G T A T C G A G T A C C T G A T A G C C G T A A C T G C G T A A C G T C G T A C T G A A T C G G C T A | GATA3(Zf),DR8/iTreg-Gata3-ChIP-Seq(GSE20898)/Homer | 1e-51 | -1.184e+02 | 0.0000 | 943.0 | 3.31% | 2372.6 | 1.81% | motif file (matrix) | svg |
| 273 | C T G A C T A G A C T G G C A T A T G C C G T A C T G A C T A G A C T G A C G T A G T C C T G A | RARg(NR)/ES-RARg-ChIP-Seq(GSE30538)/Homer | 1e-51 | -1.184e+02 | 0.0000 | 330.0 | 1.16% | 484.9 | 0.37% | motif file (matrix) | svg |
| 274 | A T G C G A C T A C G T C T A G A C G T A C G T A C G T C T G A G A T C G C T A A G C T C G T A | Foxa2(Forkhead)/Liver-Foxa2-ChIP-Seq(GSE25694)/Homer | 1e-50 | -1.170e+02 | 0.0000 | 6248.0 | 21.91% | 23662.6 | 18.01% | motif file (matrix) | svg |
| 275 | G A C T G A C T G A T C C G T A G T A C A G T C G C A T C G T A G T A C G A T C G C A T G C T A | MYB74(MYB)/colamp-MYB74-DAP-Seq(GSE60143)/Homer | 1e-50 | -1.165e+02 | 0.0000 | 6436.0 | 22.57% | 24478.1 | 18.63% | motif file (matrix) | svg |
| 276 | T C G A T G A C G C A T A G C T C A G T G A T C G C T A G A T C G A C T A C G T G C A T A G T C | PRDM1(Zf)/Hela-PRDM1-ChIP-Seq(GSE31477)/Homer | 1e-50 | -1.163e+02 | 0.0000 | 3079.0 | 10.80% | 10487.1 | 7.98% | motif file (matrix) | svg |
| 277 | G T A C A C T G A T G C T G A C C T A G G A C T G T A C C G T A G C A T G C A T | ERF8(AP2EREBP)/colamp-ERF8-DAP-Seq(GSE60143)/Homer | 1e-50 | -1.155e+02 | 0.0000 | 12665.0 | 44.41% | 52025.1 | 39.60% | motif file (matrix) | svg |
| 278 | A C T G T C A G A G C T G A C T C A T G A G T C A G T C G C T A C G A T C T A G T C A G G T A C C T G A T C G A | Rfx1(HTH)/NPC-H3K4me1-ChIP-Seq(GSE16256)/Homer | 1e-49 | -1.147e+02 | 0.0000 | 1511.0 | 5.30% | 4427.8 | 3.37% | motif file (matrix) | svg |
| 279 | G C A T T G A C C T G A A G T C A G T C A C T G G T C A A G T C G C T A G A C T G C T A C T G A | DREB2(AP2EREBP)/col-DREB2-DAP-Seq(GSE60143)/Homer | 1e-49 | -1.131e+02 | 0.0000 | 7365.0 | 25.83% | 28561.5 | 21.74% | motif file (matrix) | svg |
| 280 | G A C T C G A T T C A G G A T C G A C T A G C T A G C T A G T C G A T C C G T A C T A G C T A G T C G A T C G A C T G A | Bcl6(Zf)/Liver-Bcl6-ChIP-Seq(GSE31578)/Homer | 1e-48 | -1.128e+02 | 0.0000 | 6108.0 | 21.42% | 23156.0 | 17.62% | motif file (matrix) | svg |
| 281 | A G T C C T G A A T C G A G C T A G C T G A C T A G T C G C T A A C G T C G A T G C A T C G A T A T C G C G T A T A G C G C A T A T G C C G T A | bZIP:IRF(bZIP,IRF)/Th17-BatF-ChIP-Seq(GSE39756)/Homer | 1e-48 | -1.127e+02 | 0.0000 | 2395.0 | 8.40% | 7824.1 | 5.96% | motif file (matrix) | svg |
| 282 | C G A T T C G A A T G C C G A T G C A T T C G A A G C T G C A T G C A T C G A T T C G A A G C T T C G A G T C A C T A G | ANAC004(NAC)/colamp-ANAC004-DAP-Seq(GSE60143)/Homer | 1e-48 | -1.127e+02 | 0.0000 | 2900.0 | 10.17% | 9824.8 | 7.48% | motif file (matrix) | svg |
| 283 | G C T A C G T A C G A T C A G T A C T G C G A T G T A C A C T G A T C G G A C T C A T G C T A G G C A T C A G T C A T G | DEAR5(AP2EREBP)/col-DEAR5-DAP-Seq(GSE60143)/Homer | 1e-48 | -1.118e+02 | 0.0000 | 3222.0 | 11.30% | 11133.1 | 8.47% | motif file (matrix) | svg |
| 284 | A C T G A C T G A G T C A C T G A C T G A G T C A C G T T C A G | ERF2(AP2EREBP)/colamp-ERF2-DAP-Seq(GSE60143)/Homer | 1e-47 | -1.103e+02 | 0.0000 | 4683.0 | 16.42% | 17177.6 | 13.07% | motif file (matrix) | svg |
| 285 | A G T C C G T A C G A T A G T C G T C A A G T C A C G T C T G A | Unknown2/Drosophila-Promoters/Homer | 1e-47 | -1.103e+02 | 0.0000 | 7745.0 | 27.16% | 30280.5 | 23.05% | motif file (matrix) | svg |
| 286 | A G C T G C A T A C T G A C G T A G T C A C G T C T A G T A C G | Smad3(MAD)/NPC-Smad3-ChIP-Seq(GSE36673)/Homer | 1e-47 | -1.098e+02 | 0.0000 | 15739.0 | 55.19% | 66266.8 | 50.44% | motif file (matrix) | svg |
| 287 | T A C G T C G A G A C T A C T G C T G A A G T C T C A G G A C T T G A C C T G A | Atf1(bZIP)/K562-ATF1-ChIP-Seq(GSE31477)/Homer | 1e-47 | -1.089e+02 | 0.0000 | 7871.0 | 27.60% | 30862.8 | 23.49% | motif file (matrix) | svg |
| 288 | A C T G C G T A A C T G A T G C T G A C G A T C A T C G T G C A A C T G A G T C | ZNF519(Zf)/HEK293-ZNF519.GFP-ChIP-Seq(GSE58341)/Homer | 1e-47 | -1.085e+02 | 0.0000 | 1228.0 | 4.31% | 3452.4 | 2.63% | motif file (matrix) | svg |
| 289 | C G A T C G T A A G T C A C G T A C G T T C A G G C A T C G T A G C T A G C T A C G T A A G T C C G T A G T C A A C T G | ANAC058(NAC)/col-ANAC058-DAP-Seq(GSE60143)/Homer | 1e-46 | -1.078e+02 | 0.0000 | 6022.0 | 21.12% | 22898.2 | 17.43% | motif file (matrix) | svg |
| 290 | C G T A T G A C T C G A A G T C C G T A A T C G A T G C A C G T A C T G A G T C | E2A(bHLH)/proBcell-E2A-ChIP-Seq(GSE21978)/Homer | 1e-46 | -1.071e+02 | 0.0000 | 6593.0 | 23.12% | 25363.5 | 19.30% | motif file (matrix) | svg |
| 291 | A G T C C G T A T G A C A T G C G C A T C T G A G T A C G A T C | MYB55(MYB)/colamp-MYB55-DAP-Seq(GSE60143)/Homer | 1e-46 | -1.061e+02 | 0.0000 | 10784.0 | 37.82% | 43818.2 | 33.35% | motif file (matrix) | svg |
| 292 | G C T A C G T A C G T A G C A T C A T G C T A G A G T C A C T G A C T G A G T C A C T G T C A G | ABR1(AP2EREBP)/colamp-ABR1-DAP-Seq(GSE60143)/Homer | 1e-45 | -1.058e+02 | 0.0000 | 11430.0 | 40.08% | 46725.9 | 35.56% | motif file (matrix) | svg |
| 293 | T G C A C T G A A T G C G T C A A C G T A T G C A C G T A C T G A C T G T G C A | ZBTB18(Zf)/HEK293-ZBTB18.GFP-ChIP-Seq(GSE58341)/Homer | 1e-45 | -1.055e+02 | 0.0000 | 2808.0 | 9.85% | 9563.8 | 7.28% | motif file (matrix) | svg |
| 294 | C G A T T A C G T G C A G T A C G A T C G A C T A G C T A C G T A T C G G T A C G A T C G T A C G A T C G T C A | PPARE(NR),DR1/3T3L1-Pparg-ChIP-Seq(GSE13511)/Homer | 1e-45 | -1.052e+02 | 0.0000 | 4512.0 | 15.82% | 16562.3 | 12.61% | motif file (matrix) | svg |
| 295 | C A T G G A T C C T G A G T A C C T A G C T G A G C T A G C A T G A T C G A T C A G T C C T A G C G T A C A T G C T A G | AIL7(AP2EREBP)/colamp-AIL7-DAP-Seq(GSE60143)/Homer | 1e-45 | -1.041e+02 | 0.0000 | 5843.0 | 20.49% | 22215.8 | 16.91% | motif file (matrix) | svg |
| 296 | G A T C A G T C G A C T G C T A G T A C A G T C G C A T G C T A G T A C G A T C | MYB61(MYB)/colamp-MYB61-DAP-Seq(GSE60143)/Homer | 1e-44 | -1.031e+02 | 0.0000 | 12581.0 | 44.12% | 51999.5 | 39.58% | motif file (matrix) | svg |
| 297 | G C T A C G T A C G T A G A C T C A T G C T A G G A T C A C T G T C A G G A T C A C T G T A C G | ERF9(AP2EREBP)/colamp-ERF9-DAP-Seq(GSE60143)/Homer | 1e-44 | -1.030e+02 | 0.0000 | 2850.0 | 9.99% | 9771.0 | 7.44% | motif file (matrix) | svg |
| 298 | T G A C C T G A C T A G T C G A C T G A A T G C C G T A A C T G G C A T G T A C G C A T A T C G G C A T A G C T G A T C | PR(NR)/T47D-PR-ChIP-Seq(GSE31130)/Homer | 1e-44 | -1.020e+02 | 0.0000 | 12273.0 | 43.04% | 50637.1 | 38.54% | motif file (matrix) | svg |
| 299 | A G T C G T A C C T G A A G T C A G T C C A T G G C T A A G T C T G C A G C T A G C A T G C A T | RAP21(AP2EREBP)/colamp-RAP21-DAP-Seq(GSE60143)/Homer | 1e-44 | -1.017e+02 | 0.0000 | 3898.0 | 13.67% | 14072.5 | 10.71% | motif file (matrix) | svg |
| 300 | G C A T C G A T A T G C A G C T T C G A T A C G G C T A C G T A C A T G T G A C G C A T C G A T A G T C A G C T C G T A | AT3G09735(S1Falike)/col-AT3G09735-DAP-Seq(GSE60143)/Homer | 1e-43 | -1.009e+02 | 0.0000 | 3166.0 | 11.10% | 11081.6 | 8.43% | motif file (matrix) | svg |
| 301 | G A T C C T G A A G T C A G C T A C G T A C G T A C G T A C G T | At1g64620(C2C2dof)/colamp-At1g64620-DAP-Seq(GSE60143)/Homer | 1e-43 | -1.008e+02 | 0.0000 | 10210.0 | 35.80% | 41403.3 | 31.51% | motif file (matrix) | svg |
| 302 | T C A G G A C T G T C A C G T A A C G T A T C G C G T A A C G T A C G T C T G A | ATHB15(HB)/col-ATHB15-DAP-Seq(GSE60143)/Homer | 1e-43 | -1.007e+02 | 0.0000 | 3904.0 | 13.69% | 14116.3 | 10.74% | motif file (matrix) | svg |
| 303 | C G A T G A T C T A C G C T G A G C T A C G T A G C A T A G T C C T A G C G T A G C A T C G A T | AT2G15740(C2H2)/col-AT2G15740-DAP-Seq(GSE60143)/Homer | 1e-43 | -1.001e+02 | 0.0000 | 18270.0 | 64.07% | 78367.3 | 59.65% | motif file (matrix) | svg |
| 304 | A T C G A G T C A G T C C G T A A C T G G C A T | hINR(CPE) | 1e-43 | -9.989e+01 | 0.0000 | 7886.0 | 27.65% | 31156.4 | 23.71% | motif file (matrix) | svg |
| 305 | C G A T C T A G T C A G C A G T C G T A A G T C G C T A A C G T G A C T A T G C A G T C G C T A | PRDM10(Zf)/HEK293-PRDM10.eGFP-ChIP-Seq(Encode)/Homer | 1e-42 | -9.785e+01 | 0.0000 | 3951.0 | 13.85% | 14364.3 | 10.93% | motif file (matrix) | svg |
| 306 | T A C G A C T G A G C T G T A C C G T A T C G A C T G A A C T G C A T G A C G T A G T C C G T A | COUP-TFII(NR)/K562-NR2F1-ChIP-Seq(Encode)/Homer | 1e-42 | -9.738e+01 | 0.0000 | 9313.0 | 32.66% | 37510.7 | 28.55% | motif file (matrix) | svg |
| 307 | C G A T G A C T C G A T T C A G G A C T A C G T C A G T C T G A G A C T G A C T A G C T C G A T A C T G A T C G G T A C G C T A | NF1:FOXA1(CTF,Forkhead)/LNCAP-FOXA1-ChIP-Seq(GSE27824)/Homer | 1e-42 | -9.728e+01 | 0.0000 | 458.0 | 1.61% | 925.9 | 0.70% | motif file (matrix) | svg |
| 308 | A G T C A C G T A C G T T C A G G C T A G C T A A T G C C G T A C G A T A G T C C G T A G T C A A C T G G A T C G C A T | SND3(NAC)/col-SND3-DAP-Seq(GSE60143)/Homer | 1e-41 | -9.578e+01 | 0.0000 | 7258.0 | 25.45% | 28520.8 | 21.71% | motif file (matrix) | svg |
| 309 | C G A T T C A G G T A C A C G T A C G T T C A G C G A T C G T A G T C A G C T A C G T A A G T C C G T A G T C A C A T G | ANAC057(NAC)/colamp-ANAC057-DAP-Seq(GSE60143)/Homer | 1e-41 | -9.550e+01 | 0.0000 | 7636.0 | 26.78% | 30177.2 | 22.97% | motif file (matrix) | svg |
| 310 | T G A C G C T A T C G A T G C A A G T C A G T C C G T A A G T C C G T A C T G A G C T A G T A C | RUNX2(Runt)/PCa-RUNX2-ChIP-Seq(GSE33889)/Homer | 1e-41 | -9.459e+01 | 0.0000 | 5776.0 | 20.25% | 22144.7 | 16.85% | motif file (matrix) | svg |
| 311 | T C G A C G T A C G T A T C G A A C T G G T A C C G T A A G C T G T C A G C A T | At3g24120(G2like)/col-At3g24120-DAP-Seq(GSE60143)/Homer | 1e-40 | -9.319e+01 | 0.0000 | 20928.0 | 73.39% | 91203.2 | 69.42% | motif file (matrix) | svg |
| 312 | T C A G C T G A C G T A C G T A T A C G G C A T C T A G C T G A C G T A C G T A T A C G G A C T | IRF1(IRF)/PBMC-IRF1-ChIP-Seq(GSE43036)/Homer | 1e-40 | -9.309e+01 | 0.0000 | 779.0 | 2.73% | 1994.1 | 1.52% | motif file (matrix) | svg |
| 313 | C T A G C T A G C G T A C G T A T A C G C G A T C T A G C T G A C T G A C G T A T A C G G A C T | PU.1:IRF8(ETS:IRF)/pDC-Irf8-ChIP-Seq(GSE66899)/Homer | 1e-40 | -9.276e+01 | 0.0000 | 1031.0 | 3.62% | 2883.8 | 2.19% | motif file (matrix) | svg |
| 314 | A C G T T G A C A G T C A G C T A G T C A G C T A C T G G A C T A G C T G A C T | REF6(Zf)/Arabidopsis-REF6-ChIP-Seq(GSE106942)/Homer | 1e-40 | -9.269e+01 | 0.0000 | 3260.0 | 11.43% | 11607.0 | 8.83% | motif file (matrix) | svg |
| 315 | A C T G G A T C G A C T A C T G A C G T C A T G A C T G A C G T A G C T C G A T | RUNX-AML(Runt)/CD4+-PolII-ChIP-Seq(Barski\_et\_al.)/Homer | 1e-39 | -9.026e+01 | 0.0000 | 4557.0 | 15.98% | 17055.1 | 12.98% | motif file (matrix) | svg |
| 316 | A G T C G A T C A G C T C G T A G T A C A G T C G C A T C T G A G T A C G A T C | AT4G26030(C2H2)/col-AT4G26030-DAP-Seq(GSE60143)/Homer | 1e-38 | -8.824e+01 | 0.0000 | 9515.0 | 33.37% | 38670.9 | 29.43% | motif file (matrix) | svg |
| 317 | C G A T T C G A A G T C A C G T A C G T T A C G C G T A G C T A G C T A C G A T G C A T A T G C C G T A G T C A A C T G | ANAC071(NAC)/col-ANAC071-DAP-Seq(GSE60143)/Homer | 1e-38 | -8.822e+01 | 0.0000 | 8861.0 | 31.07% | 35768.9 | 27.22% | motif file (matrix) | svg |
| 318 | T G C A A G C T C A T G C G T A A G C T A C T G G A T C G T C A C G T A A G C T | Atf4(bZIP)/MEF-Atf4-ChIP-Seq(GSE35681)/Homer | 1e-38 | -8.782e+01 | 0.0000 | 3512.0 | 12.32% | 12734.3 | 9.69% | motif file (matrix) | svg |
| 319 | C A T G A T G C T A G C C T G A A G T C A G T C A C T G G C T A A G T C G T A C G C T A G C A T | At4g28140(AP2EREBP)/colamp-At4g28140-DAP-Seq(GSE60143)/Homer | 1e-37 | -8.743e+01 | 0.0000 | 4653.0 | 16.32% | 17522.4 | 13.34% | motif file (matrix) | svg |
| 320 | C A T G C T A G A G T C A C T G A C T G G T A C C A T G T A C G | AT1G28160(AP2EREBP)/colamp-AT1G28160-DAP-Seq(GSE60143)/Homer | 1e-37 | -8.729e+01 | 0.0000 | 13669.0 | 47.93% | 57444.0 | 43.72% | motif file (matrix) | svg |
| 321 | A T G C G T A C C T G A A G T C A G T C A C T G G T C A A G T C G T C A G C A T G C A T G A C T | At5g65130(AP2EREBP)/colamp-At5g65130-DAP-Seq(GSE60143)/Homer | 1e-37 | -8.647e+01 | 0.0000 | 2977.0 | 10.44% | 10558.0 | 8.04% | motif file (matrix) | svg |
| 322 | A C T G C T A G A G T C A C T G A C T G A G T C A C T G T A C G | ERF104(AP2EREBP)/col-ERF104-DAP-Seq(GSE60143)/Homer | 1e-37 | -8.573e+01 | 0.0000 | 7942.0 | 27.85% | 31784.8 | 24.19% | motif file (matrix) | svg |
| 323 | C G T A G A C T C G T A A C G T C A G T A G T C A G C T G A C T | KAN2(G2like)/colamp-KAN2-DAP-Seq(GSE60143)/Homer | 1e-36 | -8.335e+01 | 0.0000 | 10691.0 | 37.49% | 44077.5 | 33.55% | motif file (matrix) | svg |
| 324 | T A C G T A C G C T A G T C A G A G T C C G T A A T C G A T G C A C G T A C T G A G T C G A C T | Ascl2(bHLH)/ESC-Ascl2-ChIP-Seq(GSE97712)/Homer | 1e-35 | -8.112e+01 | 0.0000 | 5417.0 | 19.00% | 20927.9 | 15.93% | motif file (matrix) | svg |
| 325 | C G A T C T A G G A T C G C T A A G C T C T A G G A T C C G T A | RBFox2(?)/Heart-RBFox2-CLIP-Seq(GSE57926)/Homer | 1e-35 | -8.109e+01 | 0.0000 | 14534.0 | 50.97% | 61615.7 | 46.90% | motif file (matrix) | svg |
| 326 | A C T G A G C T A G T C G T C A A G C T T C A G A T G C G A T C G C A T A T C G T C G A T A G C C G A T C A T G T A G C | Pax8(Paired,Homeobox)/Thyroid-Pax8-ChIP-Seq(GSE26938)/Homer | 1e-34 | -8.045e+01 | 0.0000 | 1839.0 | 6.45% | 6081.5 | 4.63% | motif file (matrix) | svg |
| 327 | C T A G C A T G C A T G T A C G A G T C G C A T A G C T C T A G A C G T A G T C G A C T A C T G A C T G A C T G T C G A | Zfp809(Zf)/ES-Zfp809-ChIP-Seq(GSE70799)/Homer | 1e-34 | -8.006e+01 | 0.0000 | 894.0 | 3.13% | 2504.7 | 1.91% | motif file (matrix) | svg |
| 328 | G A C T A C T G C G T A A G T C T C A G G C A T G T A C C G T A A C G T G A T C | TGA1(bZIP)/colamp-TGA1-DAP-Seq(GSE60143)/Homer | 1e-34 | -7.969e+01 | 0.0000 | 4863.0 | 17.05% | 18591.8 | 14.15% | motif file (matrix) | svg |
| 329 | T A C G T C G A C G T A C G T A C G T A C T G A A C T G A C G T C G T A T C G A | AT2G28810(C2C2dof)/colamp-AT2G28810-DAP-Seq(GSE60143)/Homer | 1e-33 | -7.810e+01 | 0.0000 | 14171.0 | 49.69% | 60051.9 | 45.71% | motif file (matrix) | svg |
| 330 | C T A G A C T G A C G T C G T A A C T G A C T G A C G T T C A G C T G A T C G A | MYB107(MYB)/col-MYB107-DAP-Seq(GSE60143)/Homer | 1e-33 | -7.801e+01 | 0.0000 | 13130.0 | 46.04% | 55293.3 | 42.08% | motif file (matrix) | svg |
| 331 | T G C A A G C T A C G T C T A G G A T C C T A G G A T C G T C A C T G A A G T C | CEBP(bZIP)/ThioMac-CEBPb-ChIP-Seq(GSE21512)/Homer | 1e-33 | -7.673e+01 | 0.0000 | 7845.0 | 27.51% | 31618.2 | 24.07% | motif file (matrix) | svg |
| 332 | T C G A C T A G A G T C A G T C C G T A C G T A A C G T T A G C T C A G T A C G | NFY(CCAAT)/Promoter/Homer | 1e-33 | -7.649e+01 | 0.0000 | 9636.0 | 33.79% | 39572.9 | 30.12% | motif file (matrix) | svg |
| 333 | A T G C G A C T A C T G C A G T G A T C A C G T T A C G T A C G | Smad2(MAD)/ES-SMAD2-ChIP-Seq(GSE29422)/Homer | 1e-33 | -7.609e+01 | 0.0000 | 11269.0 | 39.52% | 46916.8 | 35.71% | motif file (matrix) | svg |
| 334 | C T A G A G T C A G C T A C T G C G T A C A G T C G T A C T G A T A G C T G A C | Unknown5/Drosophila-Promoters/Homer | 1e-33 | -7.607e+01 | 0.0000 | 7853.0 | 27.54% | 31673.3 | 24.11% | motif file (matrix) | svg |
| 335 | G A T C C G T A G A C T C T A G G A T C C T G A G A C T C T G A G A C T C T A G G A T C C T G A G A C T C T G A G A C T | OCT:OCT(POU,Homeobox)/NPC-OCT6-ChIP-Seq(GSE43916)/Homer | 1e-32 | -7.587e+01 | 0.0000 | 526.0 | 1.84% | 1260.6 | 0.96% | motif file (matrix) | svg |
| 336 | C G T A A C T G C G T A A C G T C A G T A G T C A G C T G C A T G C T A C G A T | At2g01060(G2like)/colamp-At2g01060-DAP-Seq(GSE60143)/Homer | 1e-32 | -7.482e+01 | 0.0000 | 20826.0 | 73.03% | 91289.3 | 69.48% | motif file (matrix) | svg |
| 337 | A T G C A G T C A C T G A T G C A G T C A C T G A G T C G T A C | SHN3(AP2EREBP)/col-SHN3-DAP-Seq(GSE60143)/Homer | 1e-32 | -7.429e+01 | 0.0000 | 1845.0 | 6.47% | 6193.4 | 4.71% | motif file (matrix) | svg |
| 338 | A G T C A C G T A C G T T A C G G C T A G C T A C G T A C G A T C G A T A T G C C G T A G T C A A C T G G A C T G C A T | SND2(NAC)/colamp-SND2-DAP-Seq(GSE60143)/Homer | 1e-32 | -7.417e+01 | 0.0000 | 6004.0 | 21.05% | 23640.0 | 17.99% | motif file (matrix) | svg |
| 339 | G C A T T C A G C T G A A T C G A C T G C G A T G A T C C T G A | THRb(NR)/Liver-NR1A2-ChIP-Seq(GSE52613)/Homer | 1e-31 | -7.322e+01 | 0.0000 | 18574.0 | 65.13% | 80674.0 | 61.40% | motif file (matrix) | svg |
| 340 | G T A C A C G T A C G T T A C G A T G C C A T G T A C G G T A C T C A G A T G C C G T A G T C A A C T G A G C T G C T A | AT1G19040(NAC)/col-AT1G19040-DAP-Seq(GSE60143)/Homer | 1e-31 | -7.276e+01 | 0.0000 | 1218.0 | 4.27% | 3778.5 | 2.88% | motif file (matrix) | svg |
| 341 | A G T C A G T C C T G A A G T C A G T C A C T G C G T A A G T C T C G A G A T C C G A T C G T A | AT1G01250(AP2EREBP)/col-AT1G01250-DAP-Seq(GSE60143)/Homer | 1e-31 | -7.198e+01 | 0.0000 | 1668.0 | 5.85% | 5528.2 | 4.21% | motif file (matrix) | svg |
| 342 | C T G A T C A G G C T A A G C T A G T C G A C T C T G A C T A G T G C A C T G A A G T C G T A C G A T C A C T G T C G A | ZBTB12(Zf)/HEK293-ZBTB12.GFP-ChIP-Seq(GSE58341)/Homer | 1e-31 | -7.150e+01 | 0.0000 | 3355.0 | 11.76% | 12406.5 | 9.44% | motif file (matrix) | svg |
| 343 | C G A T C A G T G T C A G C T A C A G T A G C T A C G T A C T G A G T C C G T A A C G T A C T G A C G T C T G A T C G A | FUS3(ABI3VP1)/col-FUS3-DAP-Seq(GSE60143)/Homer | 1e-30 | -7.079e+01 | 0.0000 | 7719.0 | 27.07% | 31243.9 | 23.78% | motif file (matrix) | svg |
| 344 | C T G A T A C G G C A T C T A G A T G C G A T C C G A T A C T G C T A G G A T C C T G A A T G C | MYRF(MYRF)/CFPAC1-MYRF-ChIP-Seq(GSE145627)/Homer | 1e-30 | -7.034e+01 | 0.0000 | 2120.0 | 7.43% | 7353.9 | 5.60% | motif file (matrix) | svg |
| 345 | C T G A G A T C G C A T A C T G C G T A A C G T C G T A C G T A T A C G T C G A | PQM-1(?)/cElegans-L3-ChIP-Seq(modEncode)/Homer | 1e-30 | -6.981e+01 | 0.0000 | 4497.0 | 15.77% | 17268.9 | 13.14% | motif file (matrix) | svg |
| 346 | C A T G G T C A A G T C C G T A C T A G G A T C C G A T A C T G A C G T G T A C C G T A C G T A | bZIP69(bZIP)/col-bZIP69-DAP-Seq(GSE60143)/Homer | 1e-30 | -6.960e+01 | 0.0000 | 733.0 | 2.57% | 2020.6 | 1.54% | motif file (matrix) | svg |
| 347 | T A G C C T A G T C G A G A C T A C T G C T G A A G T C T C A G G C A T T G A C C T G A A G C T | Atf7(bZIP)/3T3L1-Atf7-ChIP-Seq(GSE56872)/Homer | 1e-29 | -6.875e+01 | 0.0000 | 4522.0 | 15.86% | 17401.8 | 13.24% | motif file (matrix) | svg |
| 348 | T C G A T G A C G T A C C G T A A C G T G A C T A C G T A C T G A C T G A G C T | Mesp1(bHLH)/ESC-Mesp1-ChIP-Seq(GSE165102)/Homer | 1e-29 | -6.828e+01 | 0.0000 | 4110.0 | 14.41% | 15658.1 | 11.92% | motif file (matrix) | svg |
| 349 | C G A T C T A G A G T C A C G T A C G T T C A G G C T A C G T A G C A T G C A T C G A T A G T C C G T A G T C A A C T G | VND3(NAC)/colamp-VND3-DAP-Seq(GSE60143)/Homer | 1e-29 | -6.752e+01 | 0.0000 | 6442.0 | 22.59% | 25735.3 | 19.59% | motif file (matrix) | svg |
| 350 | C G A T C A G T C A G T C A T G G T C A G A T C C G T A T C A G A G T C A C G T C T A G A C G T G T A C G T C A G C T A | VIP1(bZIP)/col-VIP1-DAP-Seq(GSE60143)/Homer | 1e-29 | -6.686e+01 | 0.0000 | 1214.0 | 4.26% | 3833.2 | 2.92% | motif file (matrix) | svg |
| 351 | C T A G T A G C A T G C C T A G A G T C A G T C C T A G G A C T G A C T G C T A | CRF10(AP2EREBP)/col100-CRF10-DAP-Seq(GSE60143)/Homer | 1e-28 | -6.611e+01 | 0.0000 | 12851.0 | 45.06% | 54447.6 | 41.44% | motif file (matrix) | svg |
| 352 | C G A T G T C A A G T C A C G T A C G T A C T G G A C T C G A T A T C G G C T A G T C A A G T C C G T A G T C A A C T G | ANAC017(NAC)/colamp-ANAC017-DAP-Seq(GSE60143)/Homer | 1e-28 | -6.547e+01 | 0.0000 | 1341.0 | 4.70% | 4340.4 | 3.30% | motif file (matrix) | svg |
| 353 | G A C T G A T C A G T C C G T A T G A C A G T C G C A T C T G A G T A C G A T C G C A T G A C T | MYB10(MYB)/col-MYB10-DAP-Seq(GSE60143)/Homer | 1e-28 | -6.511e+01 | 0.0000 | 4251.0 | 14.91% | 16335.8 | 12.43% | motif file (matrix) | svg |
| 354 | A G T C T A G C G A C T A C G T C T A G A C G T A C G T A C G T C T G A A G T C G C T A G A C T C G T A C T A G A C T G | Foxa3(Forkhead)/Liver-Foxa3-ChIP-Seq(GSE77670)/Homer | 1e-28 | -6.503e+01 | 0.0000 | 2547.0 | 8.93% | 9186.0 | 6.99% | motif file (matrix) | svg |
| 355 | A T G C T C A G T C G A G C A T A C T G C G T A A G T C T C A G G A C T T G A C C G T A A G C T | Atf2(bZIP)/3T3L1-Atf2-ChIP-Seq(GSE56872)/Homer | 1e-28 | -6.465e+01 | 0.0000 | 2793.0 | 9.79% | 10209.9 | 7.77% | motif file (matrix) | svg |
| 356 | A C T G A G T C G T C A C G T A A G T C C G T A C T A G C T A G G A C T C A T G | SCRT1(Zf)/HEK293-SCRT1.eGFP-ChIP-Seq(Encode)/Homer | 1e-28 | -6.452e+01 | 0.0000 | 2890.0 | 10.13% | 10614.4 | 8.08% | motif file (matrix) | svg |
| 357 | C G A T C G T A A G T C A C G T A C G T T C A G C G T A C G T A G C A T G C A T G C A T A G T C C G T A G T C A A C T G | VND2(NAC)/col-VND2-DAP-Seq(GSE60143)/Homer | 1e-27 | -6.428e+01 | 0.0000 | 9007.0 | 31.58% | 37171.7 | 28.29% | motif file (matrix) | svg |
| 358 | T G A C G C T A T G A C C G T A T C A G G A T C C G T A C A T G C A T G C T A G C T A G C T A G | Unknown-ESC-element(?)/mES-Nanog-ChIP-Seq(GSE11724)/Homer | 1e-27 | -6.425e+01 | 0.0000 | 2230.0 | 7.82% | 7903.4 | 6.02% | motif file (matrix) | svg |
| 359 | C A G T T C G A A G T C A C G T A C G T T C A G C G A T G C T A G C T A C G T A G C A T A G T C C G T A T G C A A C T G | ANAC045(NAC)/col-ANAC045-DAP-Seq(GSE60143)/Homer | 1e-27 | -6.422e+01 | 0.0000 | 15267.0 | 53.54% | 65603.0 | 49.93% | motif file (matrix) | svg |
| 360 | G C T A C G T A A C G T A T C G C G T A A C G T A C G T C T A G | ATHB6(Homeobox)/col-ATHB6-DAP-Seq(GSE60143)/Homer | 1e-27 | -6.267e+01 | 0.0000 | 11533.0 | 40.44% | 48587.9 | 36.98% | motif file (matrix) | svg |
| 361 | C T A G C T G A A G T C G C T A C G A T A C T G G A C T G A T C G A T C C T G A C T A G C T G A T G A C G C T A C G A T T C A G G A C T G A T C G A T C T G A C | p53(p53)/Saos-p53-ChIP-Seq(GSE15780)/Homer | 1e-27 | -6.236e+01 | 0.0000 | 873.0 | 3.06% | 2598.5 | 1.98% | motif file (matrix) | svg |
| 362 | C T A G C T G A A G T C G C T A C G A T A C T G G A C T G A T C G A T C C T G A C T A G C T G A T G A C G C T A C G A T T C A G G A C T G A T C G A T C T G A C | p53(p53)/Saos-p53-ChIP-Seq/Homer | 1e-27 | -6.236e+01 | 0.0000 | 873.0 | 3.06% | 2598.5 | 1.98% | motif file (matrix) | svg |
| 363 | G A T C G C A T G C A T A G T C A G C T T C G A T A C G G C T A C G T A C T A G T G A C G C A T C G A T G A T C A G C T | HSFC1(HSF)/col-HSFC1-DAP-Seq(GSE60143)/Homer | 1e-26 | -6.144e+01 | 0.0000 | 1313.0 | 4.60% | 4285.0 | 3.26% | motif file (matrix) | svg |
| 364 | C G A T C T A G T C G A A G C T C G A T C T G A C G T A A G C T A C T G C T A G A T G C G A T C | Hoxb4(Homeobox)/ES-Hoxb4-ChIP-Seq(GSE34014)/Homer | 1e-26 | -6.058e+01 | 0.0000 | 2418.0 | 8.48% | 8741.2 | 6.65% | motif file (matrix) | svg |
| 365 | C T A G T A C G G A C T T G C A T G C A C G A T T A C G C T G A T C G A C T G A | Hoxa10(Homeobox)/ChickenMSG-Hoxa10.Flag-ChIP-Seq(GSE86088)/Homer | 1e-26 | -6.014e+01 | 0.0000 | 4666.0 | 16.36% | 18242.8 | 13.88% | motif file (matrix) | svg |
| 366 | G T A C G T A C G T C A G C T A C G T A C G T A C G T A C T A G C T A G C T A G | SEP3(MADS)/Arabidoposis-Flower-Sep3-ChIP-Seq/Homer | 1e-25 | -5.958e+01 | 0.0000 | 7802.0 | 27.36% | 31974.8 | 24.34% | motif file (matrix) | svg |
| 367 | C G T A G A C T C G A T A T C G G T A C G C A T C A T G C G T A T A C G G C A T G T A C C G T A C A T G A T G C G C T A C T A G G C A T G C A T G C A T G A C T | MafB(bZIP)/BMM-Mafb-ChIP-Seq(GSE75722)/Homer | 1e-25 | -5.914e+01 | 0.0000 | 2368.0 | 8.30% | 8563.8 | 6.52% | motif file (matrix) | svg |
| 368 | C G T A A C T G G T C A A C G T A T C G C A G T C T A G T C A G C G T A A C T G C G T A A C G T C G T A C T G A T A C G | GATA3(Zf),DR4/iTreg-Gata3-ChIP-Seq(GSE20898)/Homer | 1e-25 | -5.858e+01 | 0.0000 | 1024.0 | 3.59% | 3208.2 | 2.44% | motif file (matrix) | svg |
| 369 | T A C G T A G C C A T G C A G T A C G T C T A G C G T A A G T C G A C T G C A T G C A T C A G T | WRKY11(WRKY)/col-WRKY11-DAP-Seq(GSE60143)/Homer | 1e-25 | -5.814e+01 | 0.0000 | 1593.0 | 5.59% | 5437.7 | 4.14% | motif file (matrix) | svg |
| 370 | A C G T C T A G C G T A A G T C G T A C A C G T A C G T A C G T G T C A G T A C T G A C G A C T | Nur77(NR)/K562-NR4A1-ChIP-Seq(GSE31363)/Homer | 1e-25 | -5.776e+01 | 0.0000 | 1534.0 | 5.38% | 5209.0 | 3.96% | motif file (matrix) | svg |
| 371 | G C A T A C G T A C G T A T C G C G T A C G T A C G T A C G T A | At2g41835(C2H2)/col-At2g41835-DAP-Seq(GSE60143)/Homer | 1e-24 | -5.718e+01 | 0.0000 | 3426.0 | 12.01% | 13025.4 | 9.91% | motif file (matrix) | svg |
| 372 | A G T C A C G T A C G T T C A G G C T A C G T A G C T A G C A T C G A T A G T C C G T A G T C A A C T G G A C T G C T A | SMB(NAC)/colamp-SMB-DAP-Seq(GSE60143)/Homer | 1e-24 | -5.704e+01 | 0.0000 | 9008.0 | 31.59% | 37433.9 | 28.49% | motif file (matrix) | svg |
| 373 | T C G A T C A G T C G A A C T G C A T G A C G T A G T C C T G A | COUP-TFII(NR)/Artia-Nr2f2-ChIP-Seq(GSE46497)/Homer | 1e-24 | -5.644e+01 | 0.0000 | 11046.0 | 38.73% | 46622.8 | 35.49% | motif file (matrix) | svg |
| 374 | C T A G A C T G A G T C A C T G A C T G A G C T A C T G T C A G | AT3G57600(AP2EREBP)/col-AT3G57600-DAP-Seq(GSE60143)/Homer | 1e-24 | -5.538e+01 | 0.0000 | 5199.0 | 18.23% | 20682.4 | 15.74% | motif file (matrix) | svg |
| 375 | C T G A A G C T A C G T A C G T A G T C G A C T G A C T C T G A C T G A C T A G C G T A C G T A | STAT6(Stat)/CD4-Stat6-ChIP-Seq(GSE22104)/Homer | 1e-23 | -5.513e+01 | 0.0000 | 3162.0 | 11.09% | 11959.8 | 9.10% | motif file (matrix) | svg |
| 376 | G C A T G C A T G A T C G A C T T C G A T C A G G C T A C G T A A C T G G T A C G C A T G C A T A G T C A G C T C G T A | HSF7(HSF)/colamp-HSF7-DAP-Seq(GSE60143)/Homer | 1e-23 | -5.511e+01 | 0.0000 | 1281.0 | 4.49% | 4248.6 | 3.23% | motif file (matrix) | svg |
| 377 | G C A T C T A G A C T G A C G T C G T A A C T G A C T G C G A T C T A G T C G A T C G A G C T A | MYB40(MYB)/col-MYB40-DAP-Seq(GSE60143)/Homer | 1e-23 | -5.476e+01 | 0.0000 | 3533.0 | 12.39% | 13538.7 | 10.30% | motif file (matrix) | svg |
| 378 | A G C T G C A T G T C A C G A T T A G C C G T A A C G T G C T A | CRC(C2C2YABBY)/col-CRC-DAP-Seq(GSE60143)/Homer | 1e-23 | -5.476e+01 | 0.0000 | 12209.0 | 42.81% | 51966.8 | 39.55% | motif file (matrix) | svg |
| 379 | C T A G T A C G G A T C G T A C G C T A A G C T A G C T G T C A T C G A T A G C | Nanog(Homeobox)/mES-Nanog-ChIP-Seq(GSE11724)/Homer | 1e-23 | -5.449e+01 | 0.0000 | 22960.0 | 80.51% | 102234.8 | 77.81% | motif file (matrix) | svg |
| 380 | T G C A G C A T C G A T C G T A C A G T A C T G G T A C C G T A C T G A A G C T G T C A A C T G C T A G G T C A C G A T A C T G G T A C T G C A C G T A A G C T | CEBP:CEBP(bZIP)/MEF-Chop-ChIP-Seq(GSE35681)/Homer | 1e-23 | -5.412e+01 | 0.0000 | 1188.0 | 4.17% | 3898.6 | 2.97% | motif file (matrix) | svg |
| 381 | C G T A C G T A C G T A A C G T C G T A A C G T A G T C G C A T | RVE1(MYBrelated)/col-RVE1-DAP-Seq(GSE60143)/Homer | 1e-23 | -5.334e+01 | 0.0000 | 6248.0 | 21.91% | 25328.0 | 19.28% | motif file (matrix) | svg |
| 382 | A T G C G A C T A G C T C T A G C G T A C T A G C G A T C T A G A T C G G A T C | Nkx2.2(Homeobox)/NPC-Nkx2.2-ChIP-Seq(GSE61673)/Homer | 1e-23 | -5.320e+01 | 0.0000 | 13229.0 | 46.39% | 56684.1 | 43.14% | motif file (matrix) | svg |
| 383 | C G A T C T A G A C G T G T C A C G T A C G T A A G T C C G T A | Foxo3(Forkhead)/U2OS-Foxo3-ChIP-Seq(E-MTAB-2701)/Homer | 1e-22 | -5.185e+01 | 0.0000 | 5797.0 | 20.33% | 23400.6 | 17.81% | motif file (matrix) | svg |
| 384 | C A T G C T A G A G C T G A C T C A T G A G T C G A T C G C T A C G A T C T A G T C A G G T A C C T G A T C G A | X-box(HTH)/NPC-H3K4me1-ChIP-Seq(GSE16256)/Homer | 1e-22 | -5.184e+01 | 0.0000 | 694.0 | 2.43% | 2045.6 | 1.56% | motif file (matrix) | svg |
| 385 | T C A G A G C T A C G T A C G T G T A C G A T C C G T A C T A G C A T G G T C A C G T A T C G A | STAT4(Stat)/CD4-Stat4-ChIP-Seq(GSE22104)/Homer | 1e-22 | -5.183e+01 | 0.0000 | 5314.0 | 18.63% | 21291.8 | 16.21% | motif file (matrix) | svg |
| 386 | C T A G C T G A C G T A C G T A C G T A C G T A A C T G A C G T C T A G G T C A | COG1(C2C2dof)/col-COG1-DAP-Seq(GSE60143)/Homer | 1e-22 | -5.177e+01 | 0.0000 | 9556.0 | 33.51% | 40090.6 | 30.51% | motif file (matrix) | svg |
| 387 | G A C T C A G T G A T C G A T C A C G T G A T C C T G A T A C G C G T A G T C A | STAT6(Stat)/Macrophage-Stat6-ChIP-Seq(GSE38377)/Homer | 1e-22 | -5.175e+01 | 0.0000 | 3306.0 | 11.59% | 12651.0 | 9.63% | motif file (matrix) | svg |
| 388 | T C G A G C T A T G A C G C T A C T A G G A T C C G A T A C T G C G A T A G C T G A C T C T A G | E-box/Drosophila-Promoters/Homer | 1e-22 | -5.111e+01 | 0.0000 | 1448.0 | 5.08% | 4970.7 | 3.78% | motif file (matrix) | svg |
| 389 | G A C T A G T C G A T C C G T A G T A C A G T C G C A T C G T A G T C A G A T C | MYB67(MYB)/col-MYB67-DAP-Seq(GSE60143)/Homer | 1e-21 | -5.024e+01 | 0.0000 | 9048.0 | 31.73% | 37871.5 | 28.82% | motif file (matrix) | svg |
| 390 | G C T A C G T A C G A T G A C T G C A T T G C A A G T C A G C T A C G T A C G T C G A T G A C T | DAG2(C2C2dof)/col-DAG2-DAP-Seq(GSE60143)/Homer | 1e-20 | -4.814e+01 | 0.0000 | 10266.0 | 36.00% | 43435.2 | 33.06% | motif file (matrix) | svg |
| 391 | G C A T A G T C G A C T T C G A T A C G G T C A T C G A A C T G T A G C G C A T G C A T A T G C | AT2G01818(PLATZ)/col-AT2G01818-DAP-Seq(GSE60143)/Homer | 1e-20 | -4.808e+01 | 0.0000 | 1241.0 | 4.35% | 4194.5 | 3.19% | motif file (matrix) | svg |
| 392 | C G T A A G T C T G A C A G C T A C G T C G T A A C G T A G T C | At5g05790(MYBrelated)/col-At5g05790-DAP-Seq(GSE60143)/Homer | 1e-20 | -4.777e+01 | 0.0000 | 10684.0 | 37.47% | 45339.7 | 34.51% | motif file (matrix) | svg |
| 393 | C G T A A T G C C G A T A C G T A G T C C G T A C G T A C G T A C T A G A T C G | TCFL2(HMG)/K562-TCF7L2-ChIP-Seq(GSE29196)/Homer | 1e-20 | -4.726e+01 | 0.0000 | 741.0 | 2.60% | 2269.4 | 1.73% | motif file (matrix) | svg |
| 394 | C T A G T A C G G A T C G T C A T G C A A C G T T G C A G C T A T C G A T G C A | Hoxa9(Homeobox)/ChickenMSG-Hoxa9.Flag-ChIP-Seq(GSE86088)/Homer | 1e-20 | -4.647e+01 | 0.0000 | 17040.0 | 59.75% | 74562.6 | 56.75% | motif file (matrix) | svg |
| 395 | A T G C A G T C G T A C A G C T T C G A C T A G G A T C C T G A G T C A A G T C G C T A T C A G | Rfx5(HTH)/GM12878-Rfx5-ChIP-Seq(GSE31477)/Homer | 1e-20 | -4.637e+01 | 0.0000 | 2420.0 | 8.49% | 9046.3 | 6.89% | motif file (matrix) | svg |
| 396 | G A T C G A T C G C T A G T C A G A C T A T G C T C G A C G A T C G A T C T A G | HAT2(Homeobox)/colamp-HAT2-DAP-Seq(GSE60143)/Homer | 1e-20 | -4.634e+01 | 0.0000 | 8352.0 | 29.29% | 34911.2 | 26.57% | motif file (matrix) | svg |
| 397 | T C G A A G T C C G T A A T C G T A G C A C G T A C T G A G C T A C G T A G T C | Ptf1a(bHLH)/Panc1-Ptf1a-ChIP-Seq(GSE47459)/Homer | 1e-20 | -4.612e+01 | 0.0000 | 12020.0 | 42.15% | 51474.3 | 39.18% | motif file (matrix) | svg |
| 398 | T A C G A T C G T A G C G A T C A C T G A C G T A G T C A C G T C T A G A T C G | Smad4(MAD)/ESC-SMAD4-ChIP-Seq(GSE29422)/Homer | 1e-19 | -4.537e+01 | 0.0000 | 11178.0 | 39.20% | 47681.3 | 36.29% | motif file (matrix) | svg |
| 399 | G C A T G C A T G A T C G C A T T C G A A C T G C G T A C G T A A T C G T A G C C G A T A C G T A G T C A G C T C G T A | HSF6(HSF)/col-HSF6-DAP-Seq(GSE60143)/Homer | 1e-19 | -4.504e+01 | 0.0000 | 1531.0 | 5.37% | 5407.0 | 4.12% | motif file (matrix) | svg |
| 400 | C T A G T G A C G A C T A T C G T C G A A G T C C T A G C A G T C T A G A T C G G T A C T C G A | O2(bZIP)/Corn-O2-ChIP-Seq(GSE63991)/Homer | 1e-19 | -4.493e+01 | 0.0000 | 1510.0 | 5.30% | 5324.9 | 4.05% | motif file (matrix) | svg |
| 401 | G T A C C G T A C G T A T A C G G C A T G T A C C G T A C A T G A G T C C G T A C G T A C G A T G C A T G C A T G A C T | MafF(bZIP)/HepG2-MafF-ChIP-Seq(GSE31477)/Homer | 1e-19 | -4.433e+01 | 0.0000 | 2147.0 | 7.53% | 7953.3 | 6.05% | motif file (matrix) | svg |
| 402 | A T G C T C G A A G T C A G C T A C G T G T A C A G T C G C T A C T A G C A T G G T C A C T G A T C A G A G T C | Stat3+il21(Stat)/CD4-Stat3-ChIP-Seq(GSE19198)/Homer | 1e-19 | -4.418e+01 | 0.0000 | 3946.0 | 13.84% | 15595.2 | 11.87% | motif file (matrix) | svg |
| 403 | T G C A C T G A C A T G C T A G C A G T A G T C C G T A A T G C A T G C T A C G G C A T T C A G G T C A G A T C G T A C | ERE(NR),IR3/MCF7-ERa-ChIP-Seq(Unpublished)/Homer | 1e-19 | -4.410e+01 | 0.0000 | 1593.0 | 5.59% | 5676.1 | 4.32% | motif file (matrix) | svg |
| 404 | A T G C C T G A A T C G T A C G A G T C C G A T T C A G C G A T C T A G A G C T G T C A G T C A C G T A A G T C C G T A T A C G C T G A | Fox:Ebox(Forkhead,bHLH)/Panc1-Foxa2-ChIP-Seq(GSE47459)/Homer | 1e-19 | -4.389e+01 | 0.0000 | 5536.0 | 19.41% | 22525.0 | 17.14% | motif file (matrix) | svg |
| 405 | G C T A C G T A A C G T C A T G C G T A A C G T A C G T C T A G | ATHB5(HB)/colamp-ATHB5-DAP-Seq(GSE60143)/Homer | 1e-19 | -4.383e+01 | 0.0000 | 8451.0 | 29.63% | 35456.5 | 26.99% | motif file (matrix) | svg |
| 406 | C T A G A C T G A C G T C G T A A C T G C A T G G C A T T C A G | MYB92(MYB)/colamp-MYB92-DAP-Seq(GSE60143)/Homer | 1e-18 | -4.312e+01 | 0.0000 | 8454.0 | 29.65% | 35499.5 | 27.02% | motif file (matrix) | svg |
| 407 | T G C A C G T A G T C A A G C T A G T C G C T A T A G C C G A T C T A G G A T C | Gfi1b(Zf)/HPC7-Gfi1b-ChIP-Seq(GSE22178)/Homer | 1e-18 | -4.309e+01 | 0.0000 | 4270.0 | 14.97% | 17029.3 | 12.96% | motif file (matrix) | svg |
| 408 | G T A C A C G T A C G T T C A G G C T A C G T A C G A T G C A T G C A T A G T C C G T A G T C A C A T G G A C T G C T A | ANAC070(NAC)/colamp-ANAC070-DAP-Seq(GSE60143)/Homer | 1e-18 | -4.260e+01 | 0.0000 | 10166.0 | 35.65% | 43223.1 | 32.90% | motif file (matrix) | svg |
| 409 | G A T C G A T C A G T C G T A C C G A T G T A C G T A C A G T C A G T C A G T C G C T A G A T C | ZNF148(Zf)/MDAMB231-ZNF148-ChIP-Seq(GSE147020)/Homer | 1e-18 | -4.242e+01 | 0.0000 | 1280.0 | 4.49% | 4440.6 | 3.38% | motif file (matrix) | svg |
| 410 | G A C T T C G A C G T A C G T A C G T A C G T A C G T A C T A G A G C T C G T A | dof45(C2C2dof)/col-dof45-DAP-Seq(GSE60143)/Homer | 1e-18 | -4.201e+01 | 0.0000 | 15132.0 | 53.06% | 65931.2 | 50.18% | motif file (matrix) | svg |
| 411 | T C A G A C T G C A G T A G T C A G T C G T C A C G T A C G T A A C T G C A G T A G T C A G T C C T G A T G C A A G C T | dHNF4(NR)/Fly-HNF4-ChIP-Seq(GSE73675)/Homer | 1e-18 | -4.162e+01 | 0.0000 | 431.0 | 1.51% | 1188.3 | 0.90% | motif file (matrix) | svg |
| 412 | C G T A C G T A T C G A C G T A C G A T C G T A A C G T A G T C G C A T G C A T | At3g09600(MYBrelated)/colamp-At3g09600-DAP-Seq(GSE60143)/Homer | 1e-17 | -4.114e+01 | 0.0000 | 3730.0 | 13.08% | 14755.8 | 11.23% | motif file (matrix) | svg |
| 413 | C T A G A C T G A G C T C G T A A C T G A C T G A C G T C T A G | MYB99(MYB)/colamp-MYB99-DAP-Seq(GSE60143)/Homer | 1e-17 | -4.103e+01 | 0.0000 | 9082.0 | 31.85% | 38407.4 | 29.23% | motif file (matrix) | svg |
| 414 | A G C T G C T A T G C A A G T C A C G T A C G T A C G T C G A T A G C T T C A G | dof24(C2C2dof)/col-dof24-DAP-Seq(GSE60143)/Homer | 1e-17 | -4.084e+01 | 0.0000 | 12795.0 | 44.87% | 55254.4 | 42.05% | motif file (matrix) | svg |
| 415 | G T A C A C G T A C G T T C A G C G T A C G T A C G A T G C A T G C A T A G T C C G T A G T C A C A T G G A C T G C T A | VND1(NAC)/col-VND1-DAP-Seq(GSE60143)/Homer | 1e-17 | -4.077e+01 | 0.0000 | 7588.0 | 26.61% | 31727.5 | 24.15% | motif file (matrix) | svg |
| 416 | C G A T T C G A A C T G G T C A C G T A C G A T G T A C G A C T | At3g04030(G2like)/col-At3g04030-DAP-Seq(GSE60143)/Homer | 1e-17 | -4.070e+01 | 0.0000 | 8441.0 | 29.60% | 35544.1 | 27.05% | motif file (matrix) | svg |
| 417 | C G T A C G A T C G T A T C G A T C G A A C G T C G T A A C G T A G T C G C A T | LHY(Myb)/Seedling-LHY-ChIP-Seq(GSE52175)/Homer | 1e-17 | -4.037e+01 | 0.0000 | 9705.0 | 34.03% | 41242.5 | 31.39% | motif file (matrix) | svg |
| 418 | A G C T A T G C G A C T G C A T C G T A A G C T G T A C C G A T A T C G A G T C | Gata6(Zf)/HUG1N-GATA6-ChIP-Seq(GSE51936)/Homer | 1e-17 | -4.026e+01 | 0.0000 | 9072.0 | 31.81% | 38396.8 | 29.22% | motif file (matrix) | svg |
| 419 | G A C T C A G T A G C T C G A T A G T C G A T C A G T C C G T A A T G C T C A G | Rbpj1(?)/Panc1-Rbpj1-ChIP-Seq(GSE47459)/Homer | 1e-17 | -4.014e+01 | 0.0000 | 7197.0 | 25.24% | 30011.5 | 22.84% | motif file (matrix) | svg |
| 420 | C A T G A G C T A C T G A C G T A G T C C G T A A C T G C G T A C T G A C G T A G C T A C G T A C G A T G C A T C G T A | AtGRF6(GRF)/col-AtGRF6-DAP-Seq(GSE60143)/Homer | 1e-17 | -3.998e+01 | 0.0000 | 10371.0 | 36.37% | 44269.1 | 33.69% | motif file (matrix) | svg |
| 421 | C G T A G C A T C A G T C T A G A G T C A C T G A C T G G T A C A C T G A T C G | ERF115(AP2EREBP)/colamp-ERF115-DAP-Seq(GSE60143)/Homer | 1e-17 | -3.993e+01 | 0.0000 | 14721.0 | 51.62% | 64137.9 | 48.82% | motif file (matrix) | svg |
| 422 | C G T A C T G A C T A G C T G A A G T C G C T A C G A T A T C G G A C T G A T C A G T C C T G A C T A G C T A G A G T C G C T A C G A T C T A G G A T C G A T C | p73(p53)/Trachea-p73-ChIP-Seq(PRJNA310161)/Homer | 1e-17 | -3.932e+01 | 0.0000 | 413.0 | 1.45% | 1144.8 | 0.87% | motif file (matrix) | svg |
| 423 | C G T A C G T A C T G A A C T G A C G T A G T C C G T A C G T A A G T C A C T G A T G C G A T C | WRKY46(WRKY)/colamp-WRKY46-DAP-Seq(GSE60143)/Homer | 1e-16 | -3.898e+01 | 0.0000 | 1197.0 | 4.20% | 4165.6 | 3.17% | motif file (matrix) | svg |
| 424 | C T A G G T A C A C G T A C G T A T C G G C A T A G C T A G C T A G C T G C A T G A C T C G T A G T C A A C T G G A C T | VND6(NAC)/col-VND6-DAP-Seq(GSE60143)/Homer | 1e-16 | -3.897e+01 | 0.0000 | 10480.0 | 36.75% | 44810.6 | 34.11% | motif file (matrix) | svg |
| 425 | T C G A G A C T T C A G T G C A G T A C G T A C A G C T G T A C C A T G T C G A C A T G C A T G A C G T A G T C C T G A | FXR(NR),ER2/Liver-FXR-ChIP-Seq(GSE133700)/Homer | 1e-16 | -3.863e+01 | 0.0000 | 3149.0 | 11.04% | 12332.3 | 9.39% | motif file (matrix) | svg |
| 426 | C G A T C T G A A G T C A C G T A C G T T C A G G C A T C G A T G C T A G C T A C G T A A G T C C G T A G T C A A C T G | CUC1(NAC)/col-CUC1-DAP-Seq(GSE60143)/Homer | 1e-16 | -3.855e+01 | 0.0000 | 4831.0 | 16.94% | 19624.9 | 14.94% | motif file (matrix) | svg |
| 427 | G C A T A C G T C G A T A G T C A G T C G C A T C G T A C G T A C G A T C G A T C G A T C T A G A C T G G C T A G C T A | AGL15(MADS)/col-AGL15-DAP-Seq(GSE60143)/Homer | 1e-16 | -3.848e+01 | 0.0000 | 1071.0 | 3.76% | 3669.0 | 2.79% | motif file (matrix) | svg |
| 428 | T A G C G C T A T C G A C T G A A G T C A G T C C T G A A G T C C G T A C T A G | RUNX(Runt)/HPC7-Runx1-ChIP-Seq(GSE22178)/Homer | 1e-16 | -3.768e+01 | 0.0000 | 5428.0 | 19.03% | 22276.4 | 16.95% | motif file (matrix) | svg |
| 429 | C G T A C T G A T C A G A C T G G T C A C G T A A C G T G T A C C G A T G C A T | AT5G45580(G2like)/colamp-AT5G45580-DAP-Seq(GSE60143)/Homer | 1e-16 | -3.753e+01 | 0.0000 | 13263.0 | 46.51% | 57559.1 | 43.81% | motif file (matrix) | svg |
| 430 | G A C T G A T C C T G A A G T C A G T C A C T G C G T A A G T C G T A C G C T A G C A T C G A T | At1g19210(AP2EREBP)/colamp-At1g19210-DAP-Seq(GSE60143)/Homer | 1e-16 | -3.746e+01 | 0.0000 | 15596.0 | 54.69% | 68299.1 | 51.98% | motif file (matrix) | svg |
| 431 | T G A C A T G C C G T A A T C G A T G C C A G T C A T G A C T G A G T C G T A C | HEB(bHLH)/mES-Heb-ChIP-Seq(GSE53233)/Homer | 1e-15 | -3.651e+01 | 0.0000 | 8655.0 | 30.35% | 36691.4 | 27.93% | motif file (matrix) | svg |
| 432 | C T G A T C A G C T G A C T A G C A T G A C G T A T G C C G T A A T G C G C A T T C A G C T G A A C T G A C G T C A G T A G T C C G T A C A G T C T A G C A T G | VDR(NR),DR3/GM10855-VDR+vitD-ChIP-Seq(GSE22484)/Homer | 1e-15 | -3.646e+01 | 0.0000 | 1330.0 | 4.66% | 4748.7 | 3.61% | motif file (matrix) | svg |
| 433 | G C T A A G T C T A C G T G C A A T C G T C A G G C T A T C G A T C A G A G C T | ELF5(ETS)/T47D-ELF5-ChIP-Seq(GSE30407)/Homer | 1e-15 | -3.640e+01 | 0.0000 | 4735.0 | 16.60% | 19281.2 | 14.68% | motif file (matrix) | svg |
| 434 | A T G C C G T A A C T G C G T A A C G T G C T A T C G A A G C T C G A T C G T A A C G T A G T C C G A T A C T G G A T C | GATA(Zf),IR4/iTreg-Gata3-ChIP-Seq(GSE20898)/Homer | 1e-15 | -3.635e+01 | 0.0000 | 1028.0 | 3.60% | 3532.1 | 2.69% | motif file (matrix) | svg |
| 435 | A T G C A G T C C T G A A G T C C G A T A C G T A G T C A G T C A C G T A T C G G A C T A C G T | Etv2(ETS)/ES-ER71-ChIP-Seq(GSE59402)/Homer | 1e-15 | -3.616e+01 | 0.0000 | 4406.0 | 15.45% | 17851.1 | 13.59% | motif file (matrix) | svg |
| 436 | C G A T C G T A A C T G C G T A A C G T C G T A A C G T A C G T C G A T G C A T C G A T C G A T | AT2G28920(ND)/col-AT2G28920-DAP-Seq(GSE60143)/Homer | 1e-15 | -3.599e+01 | 0.0000 | 2156.0 | 7.56% | 8184.4 | 6.23% | motif file (matrix) | svg |
| 437 | A T C G T G A C A T G C C T G A T C A G G A C T A G T C C G A T T C A G T C G A C A T G C T A G C T A G C G T A C T A G C T A G C T G A C T A G C T A G A T G C | ZSCAN22(Zf)/HEK293-ZSCAN22.GFP-ChIP-Seq(GSE58341)/Homer | 1e-15 | -3.549e+01 | 0.0000 | 239.0 | 0.84% | 573.4 | 0.44% | motif file (matrix) | svg |
| 438 | A T C G A G T C A C T G A G T C A G T C A C T G G A C T G A C T | PUCHI(AP2EREBP)/colamp-PUCHI-DAP-Seq(GSE60143)/Homer | 1e-15 | -3.503e+01 | 0.0000 | 7201.0 | 25.25% | 30246.1 | 23.02% | motif file (matrix) | svg |
| 439 | C T A G A T G C A T G C C G A T A C T G G A C T A T G C G C T A T G A C A G C T T A G C G C T A | PBX1(Homeobox)/MCF7-PBX1-ChIP-Seq(GSE28007)/Homer | 1e-15 | -3.485e+01 | 0.0000 | 450.0 | 1.58% | 1320.9 | 1.01% | motif file (matrix) | svg |
| 440 | G C A T G C A T G T A C G A C T T C G A A C T G G C T A C G T A A T C G T G A C G C A T G A C T A G T C A G C T C T G A | AGL95(ND)/col-AGL95-DAP-Seq(GSE60143)/Homer | 1e-15 | -3.465e+01 | 0.0000 | 672.0 | 2.36% | 2159.1 | 1.64% | motif file (matrix) | svg |
| 441 | C G T A C G T A C G T A A C G T C G T A A C G T A G T C G C A T | EPR1(MYBrelated)/colamp-EPR1-DAP-Seq(GSE60143)/Homer | 1e-14 | -3.397e+01 | 0.0000 | 3637.0 | 12.75% | 14580.3 | 11.10% | motif file (matrix) | svg |
| 442 | C G T A C G T A C G T A A C G T C G T A A C G T A G T C G C A T | LHY1(MYBrelated)/col-LHY1-DAP-Seq(GSE60143)/Homer | 1e-14 | -3.397e+01 | 0.0000 | 3637.0 | 12.75% | 14580.3 | 11.10% | motif file (matrix) | svg |
| 443 | C G T A T A C G T C G A A C T G A C T G C G T A C G T A T A C G A G C T T A C G | PU.1(ETS)/ThioMac-PU.1-ChIP-Seq(GSE21512)/Homer | 1e-14 | -3.390e+01 | 0.0000 | 2332.0 | 8.18% | 8981.2 | 6.84% | motif file (matrix) | svg |
| 444 | A T G C T A G C A G C T A G C T T G A C G A C T T C A G T A C G G T C A C T G A A T C G T A G C G A C T C A G T A G T C A G C T T C G A A T C G T G C A T G C A | HRE(HSF)/HepG2-HSF1-ChIP-Seq(GSE31477)/Homer | 1e-14 | -3.385e+01 | 0.0000 | 815.0 | 2.86% | 2727.3 | 2.08% | motif file (matrix) | svg |
| 445 | C G T A C G T A C T A G A C G T A C G T C G T A A C T G A C T G A C G T C T G A T C G A T C G A | MYB4(MYB)/col200-MYB4-DAP-Seq(GSE60143)/Homer | 1e-14 | -3.327e+01 | 0.0000 | 5943.0 | 20.84% | 24727.7 | 18.82% | motif file (matrix) | svg |
| 446 | A C T G C G T A A C G T C G T A C T G A A C T G T C A G G C A T | At3g11280(MYBrelated)/col-At3g11280-DAP-Seq(GSE60143)/Homer | 1e-14 | -3.324e+01 | 0.0000 | 10188.0 | 35.73% | 43768.2 | 33.31% | motif file (matrix) | svg |
| 447 | C T G A A T C G A G C T A G C T A C G T T A G C C T G A T A C G C G A T A C G T G A C T A G T C | ISRE(IRF)/ThioMac-LPS-Expression(GSE23622)/Homer | 1e-14 | -3.322e+01 | 0.0000 | 330.0 | 1.16% | 902.9 | 0.69% | motif file (matrix) | svg |
| 448 | G C T A C T G A C G T A C G T A C T G A C T G A C G A T G T C A A C G T A G T C G C A T G C A T | At5g52660(MYBrelated)/colamp-At5g52660-DAP-Seq(GSE60143)/Homer | 1e-14 | -3.318e+01 | 0.0000 | 3342.0 | 11.72% | 13329.6 | 10.15% | motif file (matrix) | svg |
| 449 | T C G A A G C T A C G T A C G T A G T C A G T C A C G T A T C G G A C T A T C G | EWS:ERG-fusion(ETS)/CADO\_ES1-EWS:ERG-ChIP-Seq(SRA014231)/Homer | 1e-14 | -3.227e+01 | 0.0000 | 3392.0 | 11.89% | 13575.7 | 10.33% | motif file (matrix) | svg |
| 450 | T C G A T A G C G T C A A C T G A C T G C G T A C G T A C T A G A G C T T C A G | ERG(ETS)/VCaP-ERG-ChIP-Seq(GSE14097)/Homer | 1e-13 | -3.198e+01 | 0.0000 | 7103.0 | 24.91% | 29946.8 | 22.79% | motif file (matrix) | svg |
| 451 | C G T A C T G A C G T A C T A G T C G A C T A G A C T G C G T A C G T A T A C G A G C T A T C G | SpiB(ETS)/OCILY3-SPIB-ChIP-Seq(GSE56857)/Homer | 1e-13 | -3.192e+01 | 0.0000 | 1311.0 | 4.60% | 4759.8 | 3.62% | motif file (matrix) | svg |
| 452 | A G C T A G T C A T G C A G C T A C G T C G T A A C G T A G T C C G A T A T G C | Gata2(Zf)/K562-GATA2-ChIP-Seq(GSE18829)/Homer | 1e-13 | -3.159e+01 | 0.0000 | 6821.0 | 23.92% | 28705.7 | 21.85% | motif file (matrix) | svg |
| 453 | T C G A C T G A C G T A C G T A C G T A C T G A A C T G A C G T C T G A C T G A | AT5G63260(C3H)/col-AT5G63260-DAP-Seq(GSE60143)/Homer | 1e-13 | -3.127e+01 | 0.0000 | 14004.0 | 49.11% | 61292.9 | 46.65% | motif file (matrix) | svg |
| 454 | G A T C G T A C C G T A A G C T G A C T G C T A C T G A A C G T G A T C G C T A | Hoxc6(Homeobox)/EB-Hoxc6.iFlag-ChIP-Seq(GSE142377)/Homer | 1e-13 | -3.113e+01 | 0.0000 | 18237.0 | 63.95% | 80901.1 | 61.57% | motif file (matrix) | svg |
| 455 | G A C T C T A G C T A G A G T C T G C A A C T G A C G T A C G T C T A G T C A G | AMYB(HTH)/Testes-AMYB-ChIP-Seq(GSE44588)/Homer | 1e-13 | -3.033e+01 | 0.0000 | 14674.0 | 51.46% | 64428.9 | 49.04% | motif file (matrix) | svg |
| 456 | A G C T G T C A T G C A A G T C A C G T A C G T A C G T C G A T G A C T T A C G | AT3G12130(C3H)/colamp-AT3G12130-DAP-Seq(GSE60143)/Homer | 1e-13 | -3.017e+01 | 0.0000 | 14616.0 | 51.25% | 64170.3 | 48.84% | motif file (matrix) | svg |
| 457 | C G T A T C G A G A T C G C A T C G T A A C G T G T A C T C A G G T C A G A C T C G T A C T A G | DREF/Drosophila-Promoters/Homer | 1e-13 | -3.012e+01 | 0.0000 | 970.0 | 3.40% | 3405.4 | 2.59% | motif file (matrix) | svg |
| 458 | C G A T C T G A G T A C A C G T A C G T T C A G C G T A C G T A G C T A G C A T G C A T A G T C C G T A G T C A C A T G | NST1(NAC)/colamp-NST1-DAP-Seq(GSE60143)/Homer | 1e-13 | -2.995e+01 | 0.0000 | 7836.0 | 27.48% | 33326.8 | 25.37% | motif file (matrix) | svg |
| 459 | C G A T C T G A A G T C A C G T A C G T T C A G C G T A C G T A C G T A G C A T C G A T A G T C C G T A G T C A A C T G | VND4(NAC)/colamp-VND4-DAP-Seq(GSE60143)/Homer | 1e-12 | -2.976e+01 | 0.0000 | 7282.0 | 25.54% | 30852.5 | 23.48% | motif file (matrix) | svg |
| 460 | T C G A G A C T A T C G C G T A A G T C C T A G G C A T G T A C C T G A A C G T G A T C G C T A | TGA4(bZIP)/colamp-TGA4-DAP-Seq(GSE60143)/Homer | 1e-12 | -2.947e+01 | 0.0000 | 3495.0 | 12.26% | 14118.1 | 10.75% | motif file (matrix) | svg |
| 461 | C G T A C G T A C G T A C G A T C G T A A C G T A G T C G C A T | At4g01280(MYBrelated)/colamp-At4g01280-DAP-Seq(GSE60143)/Homer | 1e-12 | -2.899e+01 | 0.0000 | 4738.0 | 16.61% | 19580.1 | 14.90% | motif file (matrix) | svg |
| 462 | G C T A C G T A A C T G C G T A C G A T A C G T A G T C A G C T | At3g12730(G2like)/colamp-At3g12730-DAP-Seq(GSE60143)/Homer | 1e-12 | -2.886e+01 | 0.0000 | 12766.0 | 44.77% | 55750.2 | 42.43% | motif file (matrix) | svg |
| 463 | G C T A G C A T G A C T G C A T T C A G G T A C G C T A G C A T C T G A G C T A T A G C G C T A C T G A C G A T C T A G | OCT4-SOX2-TCF-NANOG(POU,Homeobox,HMG)/mES-Oct4-ChIP-Seq(GSE11431)/Homer | 1e-12 | -2.884e+01 | 0.0000 | 972.0 | 3.41% | 3436.4 | 2.62% | motif file (matrix) | svg |
| 464 | T C G A G C A T A C T G C T G A A G T C T C A G G A C T G T A C C G T A A G C T A G T C G A T C | c-Jun-CRE(bZIP)/K562-cJun-ChIP-Seq(GSE31477)/Homer | 1e-12 | -2.848e+01 | 0.0000 | 2052.0 | 7.20% | 7940.4 | 6.04% | motif file (matrix) | svg |
| 465 | T C G A A C G T A C T G C G T A A G T C C T A G A G C T T G A C | TGA10(bZIP)/colamp-TGA10-DAP-Seq(GSE60143)/Homer | 1e-12 | -2.822e+01 | 0.0000 | 7510.0 | 26.34% | 31950.9 | 24.32% | motif file (matrix) | svg |
| 466 | T C A G G C A T A C T G C G T A A G T C C T A G G C A T T G A C | TGA9(bZIP)/colamp-TGA9-DAP-Seq(GSE60143)/Homer | 1e-12 | -2.799e+01 | 0.0000 | 12510.0 | 43.87% | 54629.2 | 41.58% | motif file (matrix) | svg |
| 467 | A T G C G A T C C G A T C T A G A C T G G C T A C G T A A G C T A C T G A G C T | TEAD2(TEA)/Py2T-Tead2-ChIP-Seq(GSE55709)/Homer | 1e-12 | -2.779e+01 | 0.0000 | 3353.0 | 11.76% | 13560.7 | 10.32% | motif file (matrix) | svg |
| 468 | C G T A C G T A C G T A C T G A C T A G A C G T C T A G G T C A | CDF3(C2C2dof)/colamp-CDF3-DAP-Seq(GSE60143)/Homer | 1e-12 | -2.764e+01 | 0.0000 | 11153.0 | 39.11% | 48455.3 | 36.88% | motif file (matrix) | svg |
| 469 | T A C G C T G A T C G A C G A T C T A G C T A G T C G A C T G A T C G A T C G A C G T A T C G A G C A T C A T G C G T A T A C G G C A T T G A C C G T A A G C T | NFAT:AP1(RHD,bZIP)/Jurkat-NFATC1-ChIP-Seq(Jolma\_et\_al.)/Homer | 1e-11 | -2.755e+01 | 0.0000 | 850.0 | 2.98% | 2967.1 | 2.26% | motif file (matrix) | svg |
| 470 | G C A T G C A T C T G A A C G T C T G A A C G T C G T A C G T A C G T A A G T C G T C A G T C A | Foxf1(Forkhead)/Lung-Foxf1-ChIP-Seq(GSE77951)/Homer | 1e-11 | -2.719e+01 | 0.0000 | 6146.0 | 21.55% | 25901.3 | 19.71% | motif file (matrix) | svg |
| 471 | G C A T T C G A C T G A G A T C A G T C G A T C G T C A G T C A A C G T A G T C C G T A C T G A | Duxbl(Homeobox)/NIH3T3-Duxbl.HA-ChIP-Seq(GSE119782)/Homer | 1e-11 | -2.662e+01 | 0.0000 | 841.0 | 2.95% | 2947.7 | 2.24% | motif file (matrix) | svg |
| 472 | C T A G T C G A C T G A C G T A T A C G G A C T T C A G T C G A G T C A T G C A T A C G A G C T | IRF2(IRF)/Erythroblas-IRF2-ChIP-Seq(GSE36985)/Homer | 1e-11 | -2.650e+01 | 0.0000 | 680.0 | 2.38% | 2306.5 | 1.76% | motif file (matrix) | svg |
| 473 | G C T A G A C T A G T C T C G A T C A G T C G A A C G T A G T C G A C T T C A G | GATA14(C2C2gata)/col-GATA14-DAP-Seq(GSE60143)/Homer | 1e-11 | -2.587e+01 | 0.0000 | 6696.0 | 23.48% | 28422.7 | 21.63% | motif file (matrix) | svg |
| 474 | G C T A T G A C G A T C C G A T G A C T A T G C C T G A A T C G G C A T A C G T | JGL(C2H2)/col-JGL-DAP-Seq(GSE60143)/Homer | 1e-11 | -2.581e+01 | 0.0000 | 11136.0 | 39.05% | 48485.3 | 36.90% | motif file (matrix) | svg |
| 475 | C G A T C G A T G C A T G C A T G T C A A G T C A G C T A C G T A C G T C G A T G A C T A C G T | OBP4(C2C2dof)/col-OBP4-DAP-Seq(GSE60143)/Homer | 1e-11 | -2.562e+01 | 0.0000 | 9352.0 | 32.79% | 40394.0 | 30.74% | motif file (matrix) | svg |
| 476 | T C G A C A T G C A T G A C G T A T G C T C G A C T G A A G C T T A C G T G C A G T A C G A T C A G C T A G T C | FXR(NR),IR1/Liver-FXR-ChIP-Seq(Chong\_et\_al.)/Homer | 1e-11 | -2.535e+01 | 0.0000 | 2508.0 | 8.79% | 9985.7 | 7.60% | motif file (matrix) | svg |
| 477 | C T G A A G T C C G A T A G C T A T G C G T A C A C G T A T C G C A G T G C A T | Elf4(ETS)/BMDM-Elf4-ChIP-Seq(GSE88699)/Homer | 1e-11 | -2.534e+01 | 0.0000 | 5992.0 | 21.01% | 25304.0 | 19.26% | motif file (matrix) | svg |
| 478 | G C A T C A G T C G A T A G T C G A T C G C T A C G A T C G A T C G A T G C T A C G A T C T A G A C T G G C T A G C T A | AGL25(MADS)/colamp-AGL25-DAP-Seq(GSE60143)/Homer | 1e-10 | -2.466e+01 | 0.0000 | 332.0 | 1.16% | 990.5 | 0.75% | motif file (matrix) | svg |
| 479 | C A T G T A C G T A G C G A T C G A T C A T G C G T A C G A C T T C A G A T G C C G A T A T C G C A G T A C T G G T A C | Zic3(Zf)/mES-Zic3-ChIP-Seq(GSE37889)/Homer | 1e-10 | -2.453e+01 | 0.0000 | 2082.0 | 7.30% | 8182.8 | 6.23% | motif file (matrix) | svg |
| 480 | G C T A C G T A C G T A C G T A C G T A A C G T C G T A A C G T A G T C A G C T G C A T G C T A G C T A G C T A G C T A | LCL1(MYBrelated)/colamp-LCL1-DAP-Seq(GSE60143)/Homer | 1e-10 | -2.420e+01 | 0.0000 | 1080.0 | 3.79% | 3969.4 | 3.02% | motif file (matrix) | svg |
| 481 | G A C T A G C T G T A C G A C T C T G A A C T G G T C A C T G A A T G C T A C G G A C T A C G T A G T C G A C T C T G A | HRE(HSF)/Striatum-HSF1-ChIP-Seq(GSE38000)/Homer | 1e-10 | -2.410e+01 | 0.0000 | 1063.0 | 3.73% | 3901.6 | 2.97% | motif file (matrix) | svg |
| 482 | C G A T T C G A G T A C A C G T A C G T T C A G G C A T G C T A T G C A G C A T C G T A A G T C C G T A T G A C C A T G | ANAC092(NAC)/colamp-ANAC092-DAP-Seq(GSE60143)/Homer | 1e-10 | -2.345e+01 | 0.0000 | 4798.0 | 16.83% | 20087.0 | 15.29% | motif file (matrix) | svg |
| 483 | A T G C G A C T A G C T A G C T A G T C G C T A C A G T C G A T G C T A A C G T A C T G G C T A T A G C G C A T T G A C | IRF:BATF(IRF:bZIP)/pDC-Irf8-ChIP-Seq(GSE66899)/Homer | 1e-9 | -2.294e+01 | 0.0000 | 689.0 | 2.42% | 2400.0 | 1.83% | motif file (matrix) | svg |
| 484 | C T A G A C G T A G T C C G T A A C T G A G T C G C A T A C T G G C A T A G T C G A C T G A T C G C A T A G T C A G C T | ZNF317(Zf)/HEK293-ZNF317.GFP-ChIP-Seq(GSE58341)/Homer | 1e-9 | -2.282e+01 | 0.0000 | 638.0 | 2.24% | 2198.7 | 1.67% | motif file (matrix) | svg |
| 485 | T A G C G T A C C G T A C T A G A C T G T G C A C G T A A T G C C G T A A T C G | AR-halfsite(NR)/LNCaP-AR-ChIP-Seq(GSE27824)/Homer | 1e-9 | -2.277e+01 | 0.0000 | 16720.0 | 58.63% | 74348.4 | 56.59% | motif file (matrix) | svg |
| 486 | G C T A T A G C A G C T A T C G G T C A C G T A G C T A A T G C G A T C C T G A | IRF4(IRF)/GM12878-IRF4-ChIP-Seq(GSE32465)/Homer | 1e-9 | -2.266e+01 | 0.0000 | 3792.0 | 13.30% | 15681.0 | 11.94% | motif file (matrix) | svg |
| 487 | C G T A C T G A C G T A C G T A C G T A A C G T C G T A A C G T A G T C G C A T C G A T C G A T | AT3G10113(MYBrelated)/col-AT3G10113-DAP-Seq(GSE60143)/Homer | 1e-9 | -2.257e+01 | 0.0000 | 4576.0 | 16.05% | 19145.1 | 14.57% | motif file (matrix) | svg |
| 488 | C T A G T A C G A G T C C G T A A G T C A C G T A G T C T C G A C G T A T A C G | Nkx2.1(Homeobox)/LungAC-Nkx2.1-ChIP-Seq(GSE43252)/Homer | 1e-9 | -2.216e+01 | 0.0000 | 16932.0 | 59.38% | 75371.2 | 57.37% | motif file (matrix) | svg |
| 489 | C G T A C T A G G A C T G T C A G T C A C G T A A G T C C G T A T C G A T C G A T C G A C G T A C T G A C T A G G C T A C G T A T A G C C G T A C G A T C G T A | FOXA1:AR(Forkhead,NR)/LNCAP-AR-ChIP-Seq(GSE27824)/Homer | 1e-9 | -2.200e+01 | 0.0000 | 267.0 | 0.94% | 778.8 | 0.59% | motif file (matrix) | svg |
| 490 | A G T C C T A G A T C G C A G T C G A T A G C T G T A C A C T G C A T G C A T G | ZBED2(Zf)/SUIT2-ZBED2.HA-ChIP-Seq(GSE141606)/Homer | 1e-9 | -2.200e+01 | 0.0000 | 7922.0 | 27.78% | 34142.1 | 25.99% | motif file (matrix) | svg |
| 491 | T C A G C T G A C T A G C A T G A C G T A T G C C T G A C T G A C T G A C T A G C A T G A C G T A T G C C T G A | TR4(NR),DR1/Hela-TR4-ChIP-Seq(GSE24685)/Homer | 1e-9 | -2.198e+01 | 0.0000 | 481.0 | 1.69% | 1591.5 | 1.21% | motif file (matrix) | svg |
| 492 | A T G C C G T A C G T A C G T A C G T A C G T A A C T G A C G T C G A T C T G A | dof43(C2C2dof)/colamp-dof43-DAP-Seq(GSE60143)/Homer | 1e-9 | -2.125e+01 | 0.0000 | 9975.0 | 34.98% | 43485.7 | 33.10% | motif file (matrix) | svg |
| 493 | T G C A C G T A A C T G T C A G C A G T C A T G T C A G G A T C T A C G A G T C T G C A A C T G A C T G T G A C G T C A | ZNF165(Zf)/WHIM12-ZNF165-ChIP-Seq(GSE65937)/Homer | 1e-9 | -2.097e+01 | 0.0000 | 426.0 | 1.49% | 1391.7 | 1.06% | motif file (matrix) | svg |
| 494 | G C A T C T A G C T A G C G T A A G C T C G T A C T G A C A T G C T A G G C A T | AT5G56840(MYBrelated)/colamp-AT5G56840-DAP-Seq(GSE60143)/Homer | 1e-9 | -2.078e+01 | 0.0000 | 11290.0 | 39.59% | 49506.7 | 37.68% | motif file (matrix) | svg |
| 495 | A G C T G A T C A G C T G A C T G A C T T C G A A G T C C G T A A C T G T C A G | SpliceAcceptor/Homer | 1e-8 | -2.068e+01 | 0.0000 | 21164.0 | 74.22% | 95227.3 | 72.48% | motif file (matrix) | svg |
| 496 | G C A T G A C T A T G C A G C T T C G A A C T G C G T A C G T A A T C G A T G C G C A T G A C T G A T C A G C T T C G A | HSFA1E(HSF)/col-HSFA1E-DAP-Seq(GSE60143)/Homer | 1e-8 | -2.032e+01 | 0.0000 | 457.0 | 1.60% | 1521.1 | 1.16% | motif file (matrix) | svg |
| 497 | T A G C G T A C A G T C G T A C C G A T A G T C A G T C A G T C A G T C A G T C C G T A G A T C | Zfp281(Zf)/ES-Zfp281-ChIP-Seq(GSE81042)/Homer | 1e-8 | -2.012e+01 | 0.0000 | 234.0 | 0.82% | 676.6 | 0.51% | motif file (matrix) | svg |
| 498 | T A C G A T G C G C T A A C T G C G T A A C G T C G T A C T G A T A C G T C G A | Gata4(Zf)/Heart-Gata4-ChIP-Seq(GSE35151)/Homer | 1e-8 | -1.962e+01 | 0.0000 | 10213.0 | 35.81% | 44675.8 | 34.00% | motif file (matrix) | svg |
| 499 | A G T C T C G A T C A G C G T A G C A T G A T C A G C T T C A G C T G A C G T A | GATA12(C2C2gata)/col-GATA12-DAP-Seq(GSE60143)/Homer | 1e-8 | -1.955e+01 | 0.0000 | 5880.0 | 20.62% | 25107.5 | 19.11% | motif file (matrix) | svg |
| 500 | G C A T C G T A G C T A G A C T C G A T G A C T A G T C C A G T A G T C A G T C A C T G C T A G G T A C C T A G C T G A | AT5G05550(Trihelix)/col-AT5G05550-DAP-Seq(GSE60143)/Homer | 1e-8 | -1.936e+01 | 0.0000 | 14607.0 | 51.22% | 64819.2 | 49.33% | motif file (matrix) | svg |
| 501 | A G T C G T C A G C T A C G A T T G C A C G T A C G A T C A G T A T C G C T A G | ATHB6(Homeobox)/Arabidopsis-HB6-ChIP-Seq(GSE80564)/Homer | 1e-8 | -1.933e+01 | 0.0000 | 7132.0 | 25.01% | 30742.4 | 23.40% | motif file (matrix) | svg |
| 502 | G A T C G C A T T C G A A G T C A C G T A C G T A C G T C G A T A C G T A T C G | AT1G47655(C2C2dof)/colamp-AT1G47655-DAP-Seq(GSE60143)/Homer | 1e-8 | -1.927e+01 | 0.0000 | 18531.0 | 64.98% | 83001.7 | 63.17% | motif file (matrix) | svg |
| 503 | A G T C G A C T A C T G G A T C G T A C C G T A T G A C A G T C C G A T A G C T A C G T A C G T C T A G G A C T C T G A | ZNF7(Zf)/HepG2-ZNF7.Flag-ChIP-Seq(Encode)/Homer | 1e-8 | -1.908e+01 | 0.0000 | 3710.0 | 13.01% | 15479.5 | 11.78% | motif file (matrix) | svg |
| 504 | C T G A C T G A G C A T G A C T A G T C T C G A C T A G C G T A A C G T G A T C A G C T T C A G | GATA4(C2C2gata)/col-GATA4-DAP-Seq(GSE60143)/Homer | 1e-8 | -1.901e+01 | 0.0000 | 7989.0 | 28.01% | 34628.8 | 26.36% | motif file (matrix) | svg |
| 505 | T C G A C T G A C G T A C G T A C G T A C G T A A C T G A C G T C G A T C T G A | BBX31(Orphan)/col-BBX31-DAP-Seq(GSE60143)/Homer | 1e-8 | -1.891e+01 | 0.0000 | 10552.0 | 37.00% | 46268.5 | 35.22% | motif file (matrix) | svg |
| 506 | T C G A G T A C T C G A T C G A C A T G A T G C A C G T A C T G A C T G A G T C C G T A C T A G A G T C A T C G A G T C | Unknown3/Drosophila-Promoters/Homer | 1e-8 | -1.890e+01 | 0.0000 | 599.0 | 2.10% | 2109.8 | 1.61% | motif file (matrix) | svg |
| 507 | G A C T A T C G C T G A A G T C T C A G G A C T G T A C C T G A A G C T G T A C | TGA6(bZIP)/colamp-TGA6-DAP-Seq(GSE60143)/Homer | 1e-8 | -1.868e+01 | 0.0000 | 7759.0 | 27.21% | 33610.0 | 25.58% | motif file (matrix) | svg |
| 508 | G A C T C T A G C T A G C T A G A C T G T C G A C T G A C T A G C T A G C T A G G T A C G T C A | ZNF467(Zf)/HEK293-ZNF467.GFP-ChIP-Seq(GSE58341)/Homer | 1e-7 | -1.837e+01 | 0.0000 | 2792.0 | 9.79% | 11475.7 | 8.73% | motif file (matrix) | svg |
| 509 | A G T C A G T C C G T A C G T A A C G T C G T A C G T A A C G T | ATHB21(HB)/colamp-ATHB21-DAP-Seq(GSE60143)/Homer | 1e-7 | -1.762e+01 | 0.0000 | 5889.0 | 20.65% | 25259.8 | 19.23% | motif file (matrix) | svg |
| 510 | T C G A A C T G A C T G C G T A C G T A T C G A A G T C C T G A A T C G G T A C G C A T C A T G | ETS:E-box(ETS,bHLH)/HPC7-Scl-ChIP-Seq(GSE22178)/Homer | 1e-7 | -1.717e+01 | 0.0000 | 353.0 | 1.24% | 1160.0 | 0.88% | motif file (matrix) | svg |
| 511 | T A G C G C T A A C T G C G T A A C G T C G T A C G T A T A C G T C A G T C G A | Gata1(Zf)/K562-GATA1-ChIP-Seq(GSE18829)/Homer | 1e-7 | -1.697e+01 | 0.0000 | 6368.0 | 22.33% | 27450.3 | 20.89% | motif file (matrix) | svg |
| 512 | C T G A A T G C C G T A A C G T A G T C A G T C A C G T A C T G A T C G G C A T | SPDEF(ETS)/VCaP-SPDEF-ChIP-Seq(SRA014231)/Homer | 1e-7 | -1.695e+01 | 0.0000 | 6580.0 | 23.07% | 28406.0 | 21.62% | motif file (matrix) | svg |
| 513 | A C T G G A C T A G T C C T G A G A T C T C A G A T G C G A C T A G T C A T G C T A G C A G C T A T C G T G C A | PAX5(Paired,Homeobox),condensed/GM12878-PAX5-ChIP-Seq(GSE32465)/Homer | 1e-7 | -1.675e+01 | 0.0000 | 920.0 | 3.23% | 3475.8 | 2.65% | motif file (matrix) | svg |
| 514 | G T C A C T G A T C A G G A T C T G C A T G C A A C G T T C A G C G T A C G T A C G T A G C T A | Hoxd12(Homeobox)/ChickenMSG-Hoxd12.Flag-ChIP-Seq(GSE86088)/Homer | 1e-7 | -1.671e+01 | 0.0000 | 11754.0 | 41.22% | 51918.3 | 39.52% | motif file (matrix) | svg |
| 515 | A C T G T G A C A C T G A C G T A C G T A C T G C G T A A G T C A G C T C G A T G C A T A C G T | WRKY17(WRKY)/colamp-WRKY17-DAP-Seq(GSE60143)/Homer | 1e-7 | -1.633e+01 | 0.0000 | 175.0 | 0.61% | 498.3 | 0.38% | motif file (matrix) | svg |
| 516 | C G T A C G T A G C A T G A C T A G T C C T G A C T A G C G T A A C G T A G T C A G C T T C A G | GATA11(C2C2gata)/col-GATA11-DAP-Seq(GSE60143)/Homer | 1e-7 | -1.631e+01 | 0.0000 | 4083.0 | 14.32% | 17269.3 | 13.14% | motif file (matrix) | svg |
| 517 | G C A T G C A T G C T A G C A T G A T C C G A T A C G T C G T A A C G T A G T C G A C T G C A T G C A T G C A T G C A T | At5g58900(MYBrelated)/colamp-At5g58900-DAP-Seq(GSE60143)/Homer | 1e-6 | -1.575e+01 | 0.0000 | 11386.0 | 39.93% | 50309.7 | 38.29% | motif file (matrix) | svg |
| 518 | C G A T C G A T A G T C A G T C G C T A C G A T C G T A C G A T G C A T C G A T C T A G A C T G G C A T G C T A C G T A | AGL13(MADS)/col-AGL13-DAP-Seq(GSE60143)/Homer | 1e-6 | -1.570e+01 | 0.0000 | 341.0 | 1.20% | 1134.9 | 0.86% | motif file (matrix) | svg |
| 519 | G A C T A C G T C G A T A G C T A G T C C G T A A C T G A C T G C G A T C T A G | NGA4(ABI3VP1)/col-NGA4-DAP-Seq(GSE60143)/Homer | 1e-6 | -1.542e+01 | 0.0000 | 13207.0 | 46.31% | 58680.8 | 44.66% | motif file (matrix) | svg |
| 520 | G A T C G T A C C G A T A C T G A C T G C G T A C G T A A C G T A C T G G A T C | TEAD(TEA)/Fibroblast-PU.1-ChIP-Seq(Unpublished)/Homer | 1e-6 | -1.531e+01 | 0.0000 | 4009.0 | 14.06% | 16994.4 | 12.93% | motif file (matrix) | svg |
| 521 | T G A C G T A C C G T A A C T G T G A C C G A T A C T G A T C G A G C T T A C G T C G A T A G C G T A C C G T A A T C G T G A C G C A T A C T G A C T G A T G C | Twist(bHLH)/HMLE-TWIST1-ChIP-Seq(Chang\_et\_al)/Homer | 1e-6 | -1.521e+01 | 0.0000 | 473.0 | 1.66% | 1667.3 | 1.27% | motif file (matrix) | svg |
| 522 | C T G A C T A G A T C G G C A T A C T G G T A C A T G C C G T A A C T G G C T A A G T C C G T A | Tbox:Smad(T-box,MAD)/ESCd5-Smad2\_3-ChIP-Seq(GSE29422)/Homer | 1e-6 | -1.498e+01 | 0.0000 | 1160.0 | 4.07% | 4536.6 | 3.45% | motif file (matrix) | svg |
| 523 | C G T A C G T A C T A G C A G T G A C T C G T A C A T G C A T G C G A T C T G A C T G A C T G A | MS188(MYB)/colamp-MS188-DAP-Seq(GSE60143)/Homer | 1e-6 | -1.496e+01 | 0.0000 | 7084.0 | 24.84% | 30810.1 | 23.45% | motif file (matrix) | svg |
| 524 | T G A C T C G A C T G A C T G A A T G C G A T C C T A G T A C G G A C T G A C T G A T C T C G A C T G A C T G A A T G C G A T C C T A G A T C G G A C T G A C T | Tcfcp2l1(CP2)/mES-Tcfcp2l1-ChIP-Seq(GSE11431)/Homer | 1e-6 | -1.494e+01 | 0.0000 | 1108.0 | 3.89% | 4316.3 | 3.29% | motif file (matrix) | svg |
| 525 | C G A T C G A T G T A C G A T C G A T C C G T A G C T A C G A T C G A T C T G A C T A G C A T G G C T A G C T A C G T A | AGL16(MADS)/col-AGL16-DAP-Seq(GSE60143)/Homer | 1e-6 | -1.493e+01 | 0.0000 | 678.0 | 2.38% | 2511.9 | 1.91% | motif file (matrix) | svg |
| 526 | A T G C A G C T T C A G T G A C T C A G A T G C T G C A A C G T A T C G G A T C A C T G A G T C | NRF1(NRF)/MCF7-NRF1-ChIP-Seq(Unpublished)/Homer | 1e-6 | -1.483e+01 | 0.0000 | 497.0 | 1.74% | 1771.9 | 1.35% | motif file (matrix) | svg |
| 527 | C A G T A G C T C G T A G C A T A G T C G A C T C T A G C T A G C A G T C T A G T C G A T G C A C T A G C A T G G A C T | STOP1(C2H2)/colamp-STOP1-DAP-Seq(GSE60143)/Homer | 1e-6 | -1.470e+01 | 0.0000 | 2892.0 | 10.14% | 12078.4 | 9.19% | motif file (matrix) | svg |
| 528 | A G T C G A T C G C T A C G A T A C G T T A C G G C A T C T G A G A C T A C T G A G T C G C T A C T G A T C G A C A G T | Oct4:Sox17(POU,Homeobox,HMG)/F9-Sox17-ChIP-Seq(GSE44553)/Homer | 1e-6 | -1.433e+01 | 0.0000 | 994.0 | 3.49% | 3850.2 | 2.93% | motif file (matrix) | svg |
| 529 | G C A T G C T A C G T A A G C T G C T A T G C A A G T C A C G T A C G T A C G T G C A T G C A T | At4g38000(C2C2dof)/col-At4g38000-DAP-Seq(GSE60143)/Homer | 1e-6 | -1.418e+01 | 0.0000 | 7270.0 | 25.49% | 31707.7 | 24.13% | motif file (matrix) | svg |
| 530 | C T A G T A C G A G T C G T A C T C G A A C G T G T C A G C T A G C T A C G A T A G T C G C T A | HOXA9(Homeobox)/HSC-Hoxa9-ChIP-Seq(GSE33509)/Homer | 1e-6 | -1.415e+01 | 0.0000 | 6289.0 | 22.05% | 27280.2 | 20.76% | motif file (matrix) | svg |
| 531 | C T G A A C T G C G T A C G T A A C G T G T A C G A C T G C A T G C A T C G A T | AT4G37180(G2like)/col-AT4G37180-DAP-Seq(GSE60143)/Homer | 1e-5 | -1.379e+01 | 0.0000 | 9507.0 | 33.34% | 41891.0 | 31.88% | motif file (matrix) | svg |
| 532 | G A C T G T A C G A C T A G T C T C A G C T G A A G T C A G T C C T A G C G A T A G C T A T G C C T G A C A G T A G C T | AT4G27900(C2C2COlike)/col-AT4G27900-DAP-Seq(GSE60143)/Homer | 1e-5 | -1.346e+01 | 0.0000 | 184.0 | 0.65% | 560.2 | 0.43% | motif file (matrix) | svg |
| 533 | C T A G C A T G A C G T C G T A C T A G C A T G C G A T C T A G T C A G T C A G | MYB3(MYB)/Arabidopsis-MYB3-ChIP-Seq(GSE80564)/Homer | 1e-5 | -1.342e+01 | 0.0000 | 12464.0 | 43.71% | 55438.8 | 42.20% | motif file (matrix) | svg |
| 534 | G C T A C G T A C G T A C G T A A C T G A C G T A G T C C G T A C G T A A G T C C A T G T A G C G T A C C G T A C G T A | WRKY7(WRKY)/colamp-WRKY7-DAP-Seq(GSE60143)/Homer | 1e-5 | -1.341e+01 | 0.0000 | 58.0 | 0.20% | 120.2 | 0.09% | motif file (matrix) | svg |
| 535 | A G T C G T A C A G C T C T A G A G T C C G A T A C T G C G T A A C T G G T C A | Zic(Zf)/Cerebellum-ZIC1.2-ChIP-Seq(GSE60731)/Homer | 1e-5 | -1.337e+01 | 0.0000 | 4496.0 | 15.77% | 19279.2 | 14.67% | motif file (matrix) | svg |
| 536 | C G A T C G A T C G A T A G T C A G T C G C T A C G T A C G T A C G T A G C T A C G A T C T A G A C T G G C T A C G T A | AGL6(MADS)/col-AGL6-DAP-Seq(GSE60143)/Homer | 1e-5 | -1.329e+01 | 0.0000 | 903.0 | 3.17% | 3494.0 | 2.66% | motif file (matrix) | svg |
| 537 | C G T A C T G A C G T A G C A T G A C T A G C T T C G A C T A G C G T A A C G T A G T C A G C T C T G A C T G A C G T A | GATA1(C2C2gata)/colamp-GATA1-DAP-Seq(GSE60143)/Homer | 1e-5 | -1.314e+01 | 0.0000 | 3674.0 | 12.88% | 15625.6 | 11.89% | motif file (matrix) | svg |
| 538 | C T G A G T A C G A C T A G T C C A G T T G C A C T G A A C G T A G C T G A T C C T A G C G A T A C T G A T G C G A C T C T G A G A T C G A C T A G C T G A T C | Mouse\_Recombination\_Hotspot(Zf)/Testis-DMC1-ChIP-Seq(GSE24438)/Homer | 1e-5 | -1.309e+01 | 0.0000 | 461.0 | 1.62% | 1658.6 | 1.26% | motif file (matrix) | svg |
| 539 | T G C A A T G C A C G T A C G T A C G T A T G C C T A G A C G T A C G T A G C T G A T C A G C T | T1ISRE(IRF)/ThioMac-Ifnb-Expression/Homer | 1e-5 | -1.293e+01 | 0.0000 | 93.0 | 0.33% | 237.6 | 0.18% | motif file (matrix) | svg |
| 540 | T G C A G C A T A G C T G C A T A G T C A G T C A G T C C T G A A C T G T C G A T C G A C A G T A T C G A G T C G A T C | ZNF143\|STAF(Zf)/CUTLL-ZNF143-ChIP-Seq(GSE29600)/Homer | 1e-5 | -1.281e+01 | 0.0000 | 1168.0 | 4.10% | 4638.9 | 3.53% | motif file (matrix) | svg |
| 541 | T C A G T A G C G A C T C A T G C T G A A T C G G C A T G T A C C G T A A C T G T A G C T G C A | MafK(bZIP)/C2C12-MafK-ChIP-Seq(GSE36030)/Homer | 1e-5 | -1.251e+01 | 0.0000 | 1124.0 | 3.94% | 4460.0 | 3.39% | motif file (matrix) | svg |
| 542 | T C G A G C A T A T G C C T G A A T G C T A G C A G T C G T A C T C G A A G C T | Srebp1a(bHLH)/HepG2-Srebp1a-ChIP-Seq(GSE31477)/Homer | 1e-5 | -1.231e+01 | 0.0000 | 1670.0 | 5.86% | 6827.6 | 5.20% | motif file (matrix) | svg |
| 543 | C G T A A C G T A C T G G T A C C G T A A C G T C G T A C G T A A C G T A C T G A G T C C G T A A C G T C T G A G C A T | OCT:OCT-short(POU,Homeobox)/NPC-OCT6-ChIP-Seq(GSE43916)/Homer | 1e-5 | -1.224e+01 | 0.0000 | 5703.0 | 20.00% | 24774.8 | 18.86% | motif file (matrix) | svg |
| 544 | T C G A C G T A A G T C A G C T C G T A A G T C T C G A G C T A G A C T C G A T A G T C A G T C A G T C C T G A T C A G T G C A T C G A C A G T A T C G A G T C | GFY-Staf(?,Zf)/Promoter/Homer | 1e-5 | -1.221e+01 | 0.0000 | 234.0 | 0.82% | 766.6 | 0.58% | motif file (matrix) | svg |
| 545 | C G T A C T G A C T A G C G T A C G T A A G T C C G T A C A G T G C A T G T C A C G A T A C T G A C G T G C A T G A T C | PGR(NR)/EndoStromal-PGR-ChIP-Seq(GSE69539)/Homer | 1e-5 | -1.218e+01 | 0.0000 | 1526.0 | 5.35% | 6206.9 | 4.72% | motif file (matrix) | svg |
| 546 | T A G C G T A C C T A G C A G T T C G A C G T A C G T A G C A T G A C T T G A C A G T C A C T G A T C G A G T C C T A G | AS2(LOBAS2)/col-AS2-DAP-Seq(GSE60143)/Homer | 1e-5 | -1.192e+01 | 0.0000 | 1056.0 | 3.70% | 4187.2 | 3.19% | motif file (matrix) | svg |
| 547 | A T G C G C A T C G A T G A T C A G C T C T G A A C T G C G T A C G T A T C A G T G A C C G A T G C A T G A T C C G A T | HSF21(HSF)/col-HSF21-DAP-Seq(GSE60143)/Homer | 1e-5 | -1.163e+01 | 0.0000 | 568.0 | 1.99% | 2130.4 | 1.62% | motif file (matrix) | svg |
| 548 | A G C T C T A G T G A C C G T A A C G T C G A T A G T C A G T C C T G A C A T G | TEAD3(TEA)/HepG2-TEAD3-ChIP-Seq(Encode)/Homer | 1e-4 | -1.144e+01 | 0.0000 | 8332.0 | 29.22% | 36740.5 | 27.96% | motif file (matrix) | svg |
| 549 | T A G C T A G C G C A T C A T G A C T G G C T A C G T A A C G T A C T G G A T C | TEAD4(TEA)/Tropoblast-Tead4-ChIP-Seq(GSE37350)/Homer | 1e-4 | -1.144e+01 | 0.0000 | 5722.0 | 20.07% | 24919.9 | 18.97% | motif file (matrix) | svg |
| 550 | G C A T G C T A G C T A C G T A G C A T G C T A C T A G C G T A C G T A A C T G C G T A C G A T A C G T A G T C G A C T | At1g68670(G2like)/colamp-At1g68670-DAP-Seq(GSE60143)/Homer | 1e-4 | -1.109e+01 | 0.0000 | 3851.0 | 13.50% | 16539.4 | 12.59% | motif file (matrix) | svg |
| 551 | C G A T C T G A G T A C C A T G G C A T T C A G G C A T C G T A C G T A G C T A C G T A A G T C G C T A G T A C C A T G | CUC2(NAC)/colamp-CUC2-DAP-Seq(GSE60143)/Homer | 1e-4 | -1.076e+01 | 0.0000 | 3957.0 | 13.88% | 17035.0 | 12.97% | motif file (matrix) | svg |
| 552 | C A G T T A G C A G T C C A T G C A G T C A T G C G A T C G A T G A C T C G A T A T C G G T A C A C T G A T C G G T A C | LBD13(LOBAS2)/colamp-LBD13-DAP-Seq(GSE60143)/Homer | 1e-4 | -1.071e+01 | 0.0000 | 9196.0 | 32.25% | 40740.0 | 31.01% | motif file (matrix) | svg |
| 553 | T C G A T A G C T G C A A C T G A C T G C G T A C G T A C T A G G A C T T A C G | ETS1(ETS)/Jurkat-ETS1-ChIP-Seq(GSE17954)/Homer | 1e-4 | -1.054e+01 | 0.0001 | 5769.0 | 20.23% | 25200.3 | 19.18% | motif file (matrix) | svg |
| 554 | G T A C C A T G T A G C A G T C C T A G C A T G C T G A C G T A G C A T G C A T A C G T G C A T G T A C A C T G A T C G | LOB(LOBAS2)/col-LOB-DAP-Seq(GSE60143)/Homer | 1e-4 | -1.049e+01 | 0.0001 | 3466.0 | 12.15% | 14857.7 | 11.31% | motif file (matrix) | svg |
| 555 | G T A C C T A G T C A G A G C T T A G C C G T A A T G C T A C G A G T C G T A C G T C A A G T C | Srebp2(bHLH)/HepG2-Srebp2-ChIP-Seq(GSE31477)/Homer | 1e-4 | -1.046e+01 | 0.0001 | 979.0 | 3.43% | 3905.1 | 2.97% | motif file (matrix) | svg |
| 556 | C A T G G A T C C T G A A G T C C T A G C G T A G C T A G C A T G A T C G A T C A G T C C T A G C G T A C A T G C T A G | PLT1(AP2EREBP)/colamp-PLT1-DAP-Seq(GSE60143)/Homer | 1e-4 | -1.033e+01 | 0.0001 | 1126.0 | 3.95% | 4543.5 | 3.46% | motif file (matrix) | svg |
| 557 | G C T A A G C T T A C G G T C A C T G A C G A T C T G A G C A T C A G T A G T C | Brn2(POU,Homeobox)/NPC-Brn2-ChIP-Seq(GSE35496)/Homer | 1e-4 | -1.026e+01 | 0.0001 | 840.0 | 2.95% | 3317.0 | 2.52% | motif file (matrix) | svg |
| 558 | C G T A C G T A G C T A C G T A G A T C C T G A A C G T A C G T A G T C A G C T G C A T G C A T | AT2G40260(G2like)/colamp-AT2G40260-DAP-Seq(GSE60143)/Homer | 1e-4 | -1.022e+01 | 0.0001 | 10982.0 | 38.51% | 48944.6 | 37.25% | motif file (matrix) | svg |
| 559 | A T G C A T C G T A C G A G C T A T C G C T G A A G T C C T A G A G C T A T G C C T G A A T G C | CRE(bZIP)/Promoter/Homer | 1e-4 | -1.016e+01 | 0.0001 | 2464.0 | 8.64% | 10419.3 | 7.93% | motif file (matrix) | svg |
| 560 | C T A G G A C T G A T C A C G T A T C G A G C T C T G A A T C G C G A T C T A G G A T C G A C T C A T G A T C G G T A C G A C T A G T C G C A T A G C T C G A T | ZNF382(Zf)/HEK293-ZNF382.GFP-ChIP-Seq(GSE58341)/Homer | 1e-4 | -1.014e+01 | 0.0001 | 164.0 | 0.58% | 523.3 | 0.40% | motif file (matrix) | svg |
| 561 | G C A T A C G T A G T C A G C T C T G A A C T G C G T A C G T A T A C G T G A C G C A T G C A T G A T C G C A T C G T A | HSFB3(HSF)/colamp-HSFB3-DAP-Seq(GSE60143)/Homer | 1e-4 | -9.922e+00 | 0.0001 | 347.0 | 1.22% | 1255.4 | 0.96% | motif file (matrix) | svg |
| 562 | T G C A C T A G C T A G C T G A C A T G A C T G T G C A G A T C G T C A T G C A G T C A G T C A A G C T C T A G G C A T | ZNF675(Zf)/HEK293-ZNF675.GFP-ChIP-Seq(GSE58341)/Homer | 1e-4 | -9.633e+00 | 0.0001 | 884.0 | 3.10% | 3525.0 | 2.68% | motif file (matrix) | svg |
| 563 | T G C A A C T G T A C G A T G C A G T C G A C T T C G A A T C G | ZNF711(Zf)/SHSY5Y-ZNF711-ChIP-Seq(GSE20673)/Homer | 1e-3 | -9.206e+00 | 0.0002 | 9354.0 | 32.80% | 41604.5 | 31.67% | motif file (matrix) | svg |
| 564 | T G A C C T A G A C T G T A G C C G A T A C T G A T G C C A T G A T C G A T C G A T C G T A G C C T G A T A G C G C T A A C T G C G T A A G C T C G T A C T G A | GATA:SCL(Zf,bHLH)/Ter119-SCL-ChIP-Seq(GSE18720)/Homer | 1e-3 | -9.073e+00 | 0.0002 | 935.0 | 3.28% | 3764.2 | 2.87% | motif file (matrix) | svg |
| 565 | T G A C G T C A A G T C G C T A C T A G A G T C C T G A C A T G A C T G C T A G T A C G T C A G | Zic2(Zf)/ESC-Zic2-ChIP-Seq(SRP197560)/Homer | 1e-3 | -9.026e+00 | 0.0002 | 1816.0 | 6.37% | 7615.4 | 5.80% | motif file (matrix) | svg |
| 566 | C T A G C T A G T C G A C G T A A T G C C G T A A T C G T C G A T A C G G C A T A C T G C A G T T A G C G A T C G A C T | MRE(NR)/Neuro2A-NR3C2-ChIPnexus(GSE115417)/Homer | 1e-3 | -8.974e+00 | 0.0002 | 7532.0 | 26.41% | 33326.7 | 25.37% | motif file (matrix) | svg |
| 567 | G T A C G A T C C A G T A G T C A G T C A G T C T G C A G A T C C T G A A T G C G T C A A C G T | WT1(Zf)/Kidney-WT1-ChIP-Seq(GSE90016)/Homer | 1e-3 | -8.889e+00 | 0.0003 | 2306.0 | 8.09% | 9790.1 | 7.45% | motif file (matrix) | svg |
| 568 | T G A C C T G A A G T C A G T C A C T G G A T C G A C T G C A T | At5g18450(AP2EREBP)/col-At5g18450-DAP-Seq(GSE60143)/Homer | 1e-3 | -8.539e+00 | 0.0004 | 12751.0 | 44.71% | 57234.4 | 43.56% | motif file (matrix) | svg |
| 569 | G C A T G A C T T G A C A G C T T C G A A C T G C G T A C T G A T C A G T A G C C G A T C G A T G T A C A G C T T C G A | AT1G23810(Orphan)/col-AT1G23810-DAP-Seq(GSE60143)/Homer | 1e-3 | -8.431e+00 | 0.0004 | 196.0 | 0.69% | 672.7 | 0.51% | motif file (matrix) | svg |
| 570 | A G C T A G T C A G T C A C G T C T A G A C G T A C G T A C G T C G T A A G T C G A T C C G T A | FOXP1(Forkhead)/H9-FOXP1-ChIP-Seq(GSE31006)/Homer | 1e-3 | -8.378e+00 | 0.0004 | 3147.0 | 11.04% | 13571.9 | 10.33% | motif file (matrix) | svg |
| 571 | C A G T G A C T G C A T T C G A A G T C A C G T A C G T A C G T C G A T G A C T | OBP3(C2C2dof)/col-OBP3-DAP-Seq(GSE60143)/Homer | 1e-3 | -8.282e+00 | 0.0005 | 16501.0 | 57.86% | 74545.6 | 56.74% | motif file (matrix) | svg |
| 572 | T C G A C T G A C G A T C G T A C G T A C G T A C T A G A G C T C T G A T C A G | Adof1(C2C2dof)/col-Adof1-DAP-Seq(GSE60143)/Homer | 1e-3 | -8.251e+00 | 0.0005 | 15550.0 | 54.53% | 70156.7 | 53.40% | motif file (matrix) | svg |
| 573 | G T C A G C A T G C T A C A G T C T A G G A T C C G T A C T G A C G T A C G A T | Oct2(POU,Homeobox)/Bcell-Oct2-ChIP-Seq(GSE21512)/Homer | 1e-3 | -8.200e+00 | 0.0005 | 1830.0 | 6.42% | 7720.7 | 5.88% | motif file (matrix) | svg |
| 574 | G C T A C G A T C T A G G T A C G C T A C G A T C G T A G C T A C G A T C A G T G A T C G C T A | Pit1+1bp(Homeobox)/GCrat-Pit1-ChIP-Seq(GSE58009)/Homer | 1e-3 | -8.034e+00 | 0.0006 | 2208.0 | 7.74% | 9405.8 | 7.16% | motif file (matrix) | svg |
| 575 | G A C T G A C T T C G A C G T A C G A T A G C T C T G A A C T G T G A C G A C T T C G A C G T A A C G T A G C T C T G A C T G A G T C A G C T A G C T A C G T A | Pax7(Paired,Homeobox),longest/Myoblast-Pax7-ChIP-Seq(GSE25064)/Homer | 1e-3 | -7.828e+00 | 0.0007 | 144.0 | 0.50% | 476.0 | 0.36% | motif file (matrix) | svg |
| 576 | C G A T C T G A C G T A A G C T A G T C T G C A C T G A A C G T A G C T G C T A | Pax7(Paired,Homeobox)/Myoblast-Pax7-ChIP-Seq(GSE25064)/Homer | 1e-3 | -7.805e+00 | 0.0007 | 1005.0 | 3.52% | 4116.2 | 3.13% | motif file (matrix) | svg |
| 577 | T A C G C G T A T C A G G A C T C T A G A C T G C A G T T A G C T C G A A C G T G T A C C T A G A G T C A G T C G A T C | ZNF669(Zf)/HEK293-ZNF669.GFP-ChIP-Seq(GSE58341)/Homer | 1e-3 | -7.762e+00 | 0.0008 | 925.0 | 3.24% | 3770.9 | 2.87% | motif file (matrix) | svg |
| 578 | A G T C A T C G C T A G A G C T G A C T C T A G A G T C A G T C G C T A C A G T T C A G T C A G G A T C C T G A T C G A G A T C | RFX(HTH)/K562-RFX3-ChIP-Seq(SRA012198)/Homer | 1e-3 | -7.759e+00 | 0.0008 | 387.0 | 1.36% | 1468.3 | 1.12% | motif file (matrix) | svg |
| 579 | C A T G C T G A A G T C A C T G A C T G A G C T A C T G A T C G | ESE3(AP2EREBP)/col-ESE3-DAP-Seq(GSE60143)/Homer | 1e-3 | -7.553e+00 | 0.0010 | 11290.0 | 39.59% | 50642.8 | 38.54% | motif file (matrix) | svg |
| 580 | C T G A C T A G T G C A C T G A T C G A A G C T C A T G T C G A A G T C G A C T A C G T A G T C G A T C G A T C G A C T | ZNF528(Zf)/HEK293-ZNF528.GFP-ChIP-Seq(GSE58341)/Homer | 1e-3 | -7.451e+00 | 0.0011 | 27.0 | 0.09% | 54.4 | 0.04% | motif file (matrix) | svg |
| 581 | G C A T A C G T A G T C A T G C A G T C C T A G T A G C G T A C C T G A G C T A | DEL1(E2FDP)/colamp-DEL1-DAP-Seq(GSE60143)/Homer | 1e-3 | -7.449e+00 | 0.0011 | 65.0 | 0.23% | 182.9 | 0.14% | motif file (matrix) | svg |
| 582 | C T A G A G C T G A C T C A T G A G T C A G T C G T C A C A G T C T A G T C A G G T A C C T G A T C G A G A T C T G A C | Rfx2(HTH)/LoVo-RFX2-ChIP-Seq(GSE49402)/Homer | 1e-3 | -7.417e+00 | 0.0011 | 446.0 | 1.56% | 1725.4 | 1.31% | motif file (matrix) | svg |
| 583 | G C T A A G C T G T A C G C A T A G C T T C G A C T G A A G T C A G T C T A C G A C G T G A C T T A C G C T A G C G T A | ZML1(C2C2gata)/colamp-ZML1-DAP-Seq(GSE60143)/Homer | 1e-3 | -7.363e+00 | 0.0011 | 457.0 | 1.60% | 1773.2 | 1.35% | motif file (matrix) | svg |
| 584 | A T C G A G T C A G T C G A C T A T G C C T G A C T A G A C T G T A C G G T A C C T G A C G A T | AP-2gamma(AP2)/MCF7-TFAP2C-ChIP-Seq(GSE21234)/Homer | 1e-3 | -7.342e+00 | 0.0012 | 4225.0 | 14.82% | 18495.4 | 14.08% | motif file (matrix) | svg |
| 585 | C A T G A C G T C T A G A C G T C A G T A C G T C T A G A G T C | PHA-4(Forkhead)/cElegans-Embryos-PHA4-ChIP-Seq(modEncode)/Homer | 1e-3 | -7.145e+00 | 0.0014 | 15874.0 | 55.67% | 71784.8 | 54.64% | motif file (matrix) | svg |
| 586 | C G T A C A G T G T A C A T G C C T A G C G T A A C G T A G T C T C G A T C A G | GATA19(C2C2gata)/colamp-GATA19-DAP-Seq(GSE60143)/Homer | 1e-3 | -7.134e+00 | 0.0014 | 1593.0 | 5.59% | 6730.5 | 5.12% | motif file (matrix) | svg |
| 587 | C T G A C T A G A G T C A G T C C G T A A G T C C A T G C A T G T C G A C T G A | ZBTB9(Zf)/HEK293T-ZBTB9-ChIP-Seq(GSE251687)/Homer | 1e-3 | -7.082e+00 | 0.0015 | 10791.0 | 37.84% | 48412.0 | 36.85% | motif file (matrix) | svg |
| 588 | C G A T C G T A G C A T C T A G A C T G C G T A A C G T G T C A C G T A C T A G C T A G G C A T | At1g19000(MYBrelated)/colamp-At1g19000-DAP-Seq(GSE60143)/Homer | 1e-3 | -7.068e+00 | 0.0015 | 7901.0 | 27.71% | 35202.5 | 26.79% | motif file (matrix) | svg |
| 589 | C G A T T A C G G A T C G C T A T C A G G A C T C G A T G T A C G A T C G T C A T C G A T G A C C G T A C T A G G A C T C T A G C T A G G T A C A G T C C G T A | CTCF-SatelliteElement(Zf?)/CD4+-CTCF-ChIP-Seq(Barski\_et\_al.)/Homer | 1e-2 | -6.521e+00 | 0.0026 | 77.0 | 0.27% | 235.2 | 0.18% | motif file (matrix) | svg |
| 590 | C T G A A C G T A C G T A C G T A G T C G A C T C G A T C T G A A C T G C G T A C G T A T C G A | STAT5(Stat)/mCD4+-Stat5-ChIP-Seq(GSE12346)/Homer | 1e-2 | -6.325e+00 | 0.0032 | 1515.0 | 5.31% | 6430.7 | 4.89% | motif file (matrix) | svg |
| 591 | C G T A A C G T A G C T C G A T C T A G G T A C C G T A A G C T C G T A G C T A | Oct4(POU,Homeobox)/mES-Oct4-ChIP-Seq(GSE11431)/Homer | 1e-2 | -6.283e+00 | 0.0033 | 3056.0 | 10.72% | 13319.5 | 10.14% | motif file (matrix) | svg |
| 592 | A C T G G A T C C T G A A T C G A G T C T A G C C T G A C G T A T A C G A G T C C T A G C A G T T C A G T C G A T G A C G A T C | PAX5(Paired,Homeobox)/GM12878-PAX5-ChIP-Seq(GSE32465)/Homer | 1e-2 | -6.151e+00 | 0.0038 | 2834.0 | 9.94% | 12332.5 | 9.39% | motif file (matrix) | svg |
| 593 | C T G A C G A T A G T C A T C G C T G A C G T A A G T C A C T G A C T G G A C T A G C T T C G A C G T A C A T G C T G A | ZIM(C2C2gata)/col-ZIM-DAP-Seq(GSE60143)/Homer | 1e-2 | -6.109e+00 | 0.0039 | 431.0 | 1.51% | 1699.1 | 1.29% | motif file (matrix) | svg |
| 594 | A G T C G A T C G A T C C G T A G T C A A G T C A G C T C T G A G A C T G A C T | ATY13(MYB)/col-ATY13-DAP-Seq(GSE60143)/Homer | 1e-2 | -6.100e+00 | 0.0040 | 20369.0 | 71.43% | 92735.7 | 70.58% | motif file (matrix) | svg |
| 595 | C A T G G A C T G C A T A C T G A G C T A C T G A C T G C G T A G C A T A G C T A T C G T A C G | Foxh1(Forkhead)/hESC-FOXH1-ChIP-Seq(GSE29422)/Homer | 1e-2 | -5.968e+00 | 0.0045 | 5539.0 | 19.42% | 24575.3 | 18.70% | motif file (matrix) | svg |
| 596 | C G T A C T A G T C G A A C G T C G A T G T A C C T G A T C G A G C A T A C T G G C A T C G T A G A C T G A T C G C A T | EIN3(EIL)/col-EIN3-DAP-Seq(GSE60143)/Homer | 1e-2 | -5.855e+00 | 0.0050 | 1151.0 | 4.04% | 4846.9 | 3.69% | motif file (matrix) | svg |
| 597 | C T G A C T A G A T C G A G C T A C T G G A C T A G T C C T G A | Tbx5(T-box)/HL1-Tbx5.biotin-ChIP-Seq(GSE21529)/Homer | 1e-2 | -5.822e+00 | 0.0052 | 16571.0 | 58.11% | 75177.6 | 57.22% | motif file (matrix) | svg |
| 598 | G C A T A C G T A T G C C T G A C T A G G A C T G A T C A C T G | Initiator/Drosophila-Promoters/Homer | 1e-2 | -5.524e+00 | 0.0070 | 14450.0 | 50.67% | 65434.2 | 49.80% | motif file (matrix) | svg |
| 599 | G A C T C G A T C T G A G T C A G A C T C G A T T C G A C G T A G C T A G C T A T G A C G T A C C G T A A C T G T G C A C G A T A C T G A C G T | Pitx1:Ebox(Homeobox,bHLH)/Hindlimb-Pitx1-ChIP-Seq(GSE41591)/Homer | 1e-2 | -5.512e+00 | 0.0071 | 1032.0 | 3.62% | 4339.3 | 3.30% | motif file (matrix) | svg |
| 600 | G C T A C G T A A C T G C G T A C G A T A C G T A G T C A G C T | At1g25550(G2like)/colamp-At1g25550-DAP-Seq(GSE60143)/Homer | 1e-2 | -5.355e+00 | 0.0083 | 8302.0 | 29.11% | 37240.1 | 28.34% | motif file (matrix) | svg |
| 601 | T C G A C A T G C T G A C G T A A T C G G T A C G C A T C G A T A G T C A G C T T C G A T A C G C G T A C G T A C A T G | HSFA6A(HSF)/col-HSFA6A-DAP-Seq(GSE60143)/Homer | 1e-2 | -5.213e+00 | 0.0095 | 153.0 | 0.54% | 554.8 | 0.42% | motif file (matrix) | svg |
| 602 | A G C T G A T C G A C T A G T C T C G A C T G A A G T C A G T C C T A G A G C T G A C T T A G C T C G A C G A T G A C T | AT5G59990(C2C2COlike)/colamp-AT5G59990-DAP-Seq(GSE60143)/Homer | 1e-2 | -4.906e+00 | 0.0129 | 471.0 | 1.65% | 1911.0 | 1.45% | motif file (matrix) | svg |
| 603 | C T A G C A T G T G C A A C G T A G T C C G T A C A T G T C A G A C G T A C G T G C T A A G T C | Six1(Homeobox)/Myoblast-Six1-ChIP-Chip(GSE20150)/Homer | 1e-2 | -4.888e+00 | 0.0131 | 2213.0 | 7.76% | 9644.3 | 7.34% | motif file (matrix) | svg |
| 604 | A T G C G T A C A C T G A G T C A G T C A C T G G A T C G T C A C G T A C G A T G C A T C G A T | RRTF1(AP2EREBP)/colamp-RRTF1-DAP-Seq(GSE60143)/Homer | 1e-2 | -4.847e+00 | 0.0136 | 4287.0 | 15.03% | 19015.2 | 14.47% | motif file (matrix) | svg |
| 605 | C T A G C T G A C G A T C G A T G T A C C G T A G C A T G C A T A C T G G C T A C G T A A G C T | EIL4(EIL)/Tomato-EIL4-ChIP-Seq(GSE116581)/Homer | 1e-2 | -4.770e+00 | 0.0147 | 1002.0 | 3.51% | 4246.9 | 3.23% | motif file (matrix) | svg |
| 606 | G A T C G C A T A C G T C G T A A C G T A G T C A G T C C G T A | AT5G61620(MYBrelated)/colamp-AT5G61620-DAP-Seq(GSE60143)/Homer | 1e-2 | -4.708e+00 | 0.0156 | 12925.0 | 45.32% | 58537.3 | 44.55% | motif file (matrix) | svg |
