## Supplemental information for "Hybrid CNN and Multi-Head Attention Model for Analyzing Epigenetic Mechanisms and Gene Expression Across Fungal Phylogenetic Distances"

### Supplementary Information for: Hybrid CNN and multi-head attention machine learning model for elucidating complex relationships between epigenetic regulatory mechanisms and gene expression across fungal phylogenetic distances

Weinstock et al.

This supplementary information packet contains Supplementary Figures 1 to 8, Supplementary Table 1, Supplemental Data descriptions and Supplementary Methods. Supplemental Data files are provided separately.

**Supplementary Figure 1.** Intra-species shallow regressor model battery performance using *N. crassa* on mean absolute error and regression AUROC metrics.

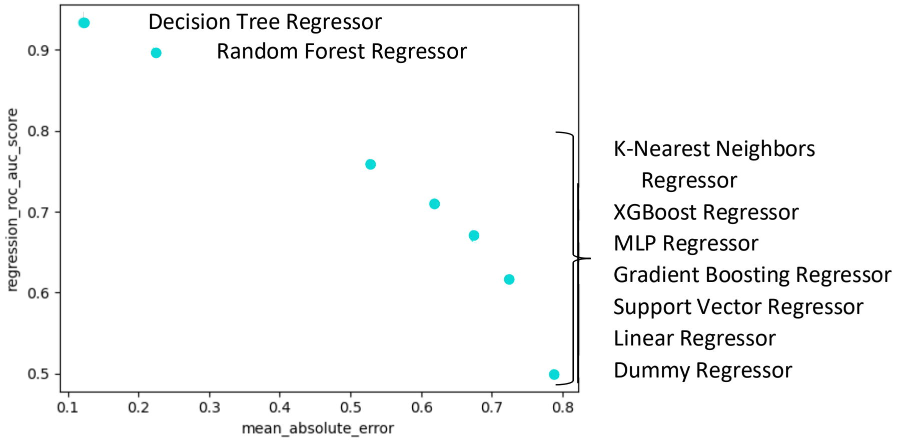

**Supplementary Figure 2.** Confusion matrices and metrics output table show K-nearest neighbors classifier performance across (A) 2, (B) 3, (C) 4, (D) 5 output bins for intra-species predictions using *N. crassa* data. Bin thresholds set to achieve equal # genes/bin. Colorbar indicates number of genes predicted for each cell.

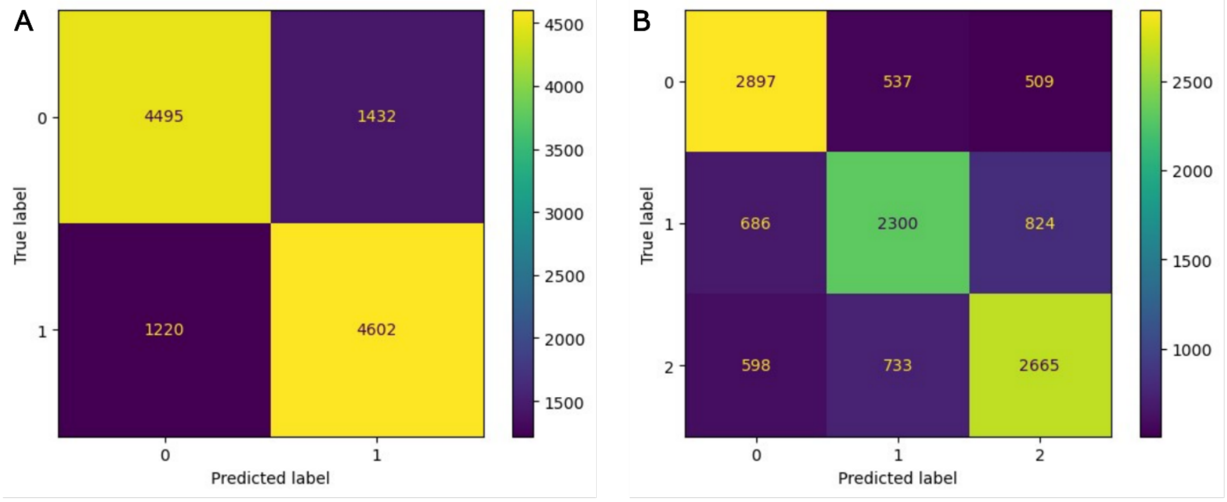


| | precision | recall | f1-score | support |
|---|---|---|---|---|
| 0 | 0.79 | 0.76 | 0.77 | 5927 |
| 1 | 0.76 | 0.79 | 0.78 | 5822 |
| accuracy | | | 0.77 | 11749 |
| macro avg | 0.77 | 0.77 | 0.77 | 11749 |
| weighted avg | 0.77 | 0.77 | 0.77 | 11749 |

| | precision | recall | f1-score | support |
|---|---|---|---|---|
| 0 | 0.69 | 0.73 | 0.71 | 3943 |
| 1 | 0.64 | 0.60 | 0.62 | 3810 |
| 2 | 0.67 | 0.67 | 0.67 | 3996 |
| accuracy | | | 0.67 | 11749 |
| macro avg | 0.67 | 0.67 | 0.67 | 11749 |
| weighted avg | 0.67 | 0.67 | 0.67 | 11749 |

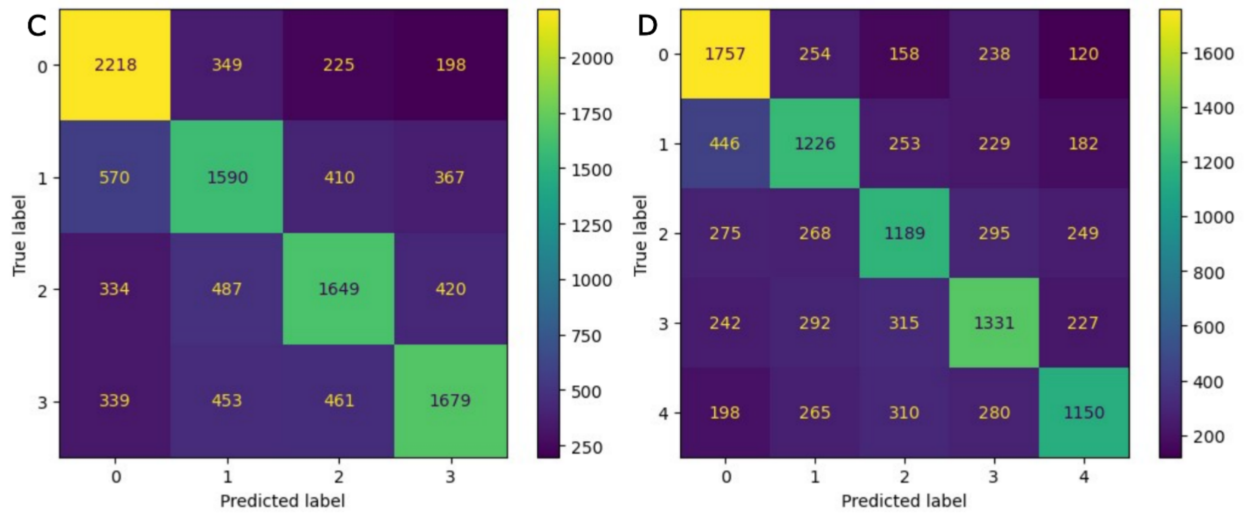


| | precision | recall | f1-score | support |
|---|---|---|---|---|
| 0 | 0.64 | 0.74 | 0.69 | 2990 |
| 1 | 0.55 | 0.54 | 0.55 | 2937 |
| 2 | 0.60 | 0.57 | 0.59 | 2890 |
| 3 | 0.63 | 0.57 | 0.60 | 2932 |
| accuracy | | | 0.61 | 11749 |
| macro avg | 0.61 | 0.61 | 0.60 | 11749 |
| weighted avg | 0.61 | 0.61 | 0.61 | 11749 |

| | precision | recall | f1-score | support |
|---|---|---|---|---|
| 0 | 0.60 | 0.70 | 0.65 | 2527 |
| 1 | 0.53 | 0.52 | 0.53 | 2336 |
| 2 | 0.53 | 0.52 | 0.53 | 2276 |
| 3 | 0.56 | 0.55 | 0.56 | 2407 |
| 4 | 0.60 | 0.52 | 0.56 | 2203 |
| accuracy | | | 0.57 | 11749 |
| macro avg | 0.57 | 0.56 | 0.56 | 11749 |
| weighted avg | 0.57 | 0.57 | 0.56 | 11749 |

**Supplementary Figure 3.** Detailed shallow classifier model prediction performance results for each combination of species, model, TSS window, signal vs averaged features, and class output number for both inter- and intra- species prediction. Colorbar shows performance metric (accuracy, precision, or AUROC) score.

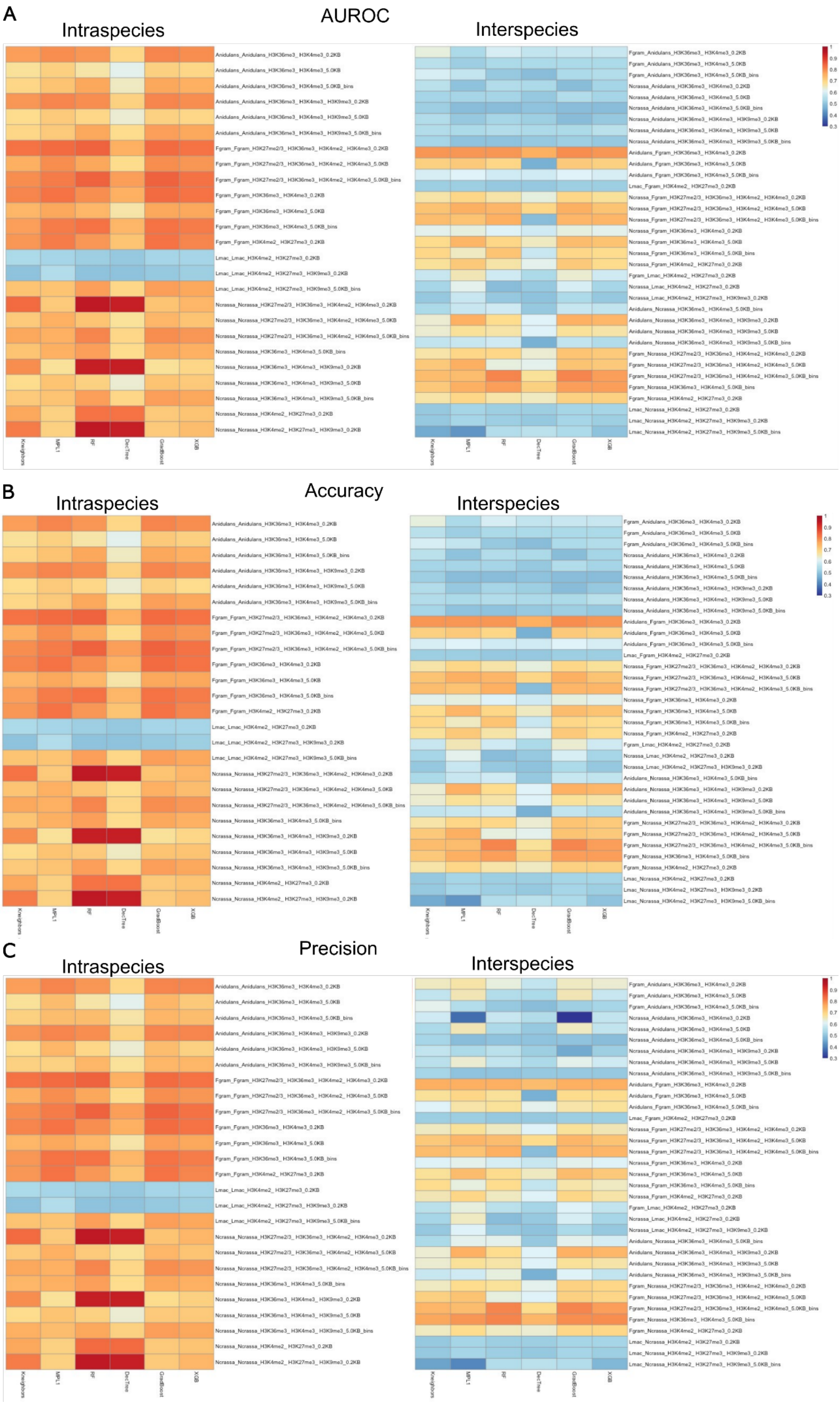
A
AUROC
Intraspecies
Interspecies
B
Accuracy
Intraspecies
Interspecies
C
Precision
Intraspecies
Interspecies

**Supplementary Figure 4**: Average shallow learning classifier performance for each number of overlapping modifications available for use in test for intra- (left) and inter- (right) species predictions. Boxes represent median +/- quartile for minimally n = 36 replicates.

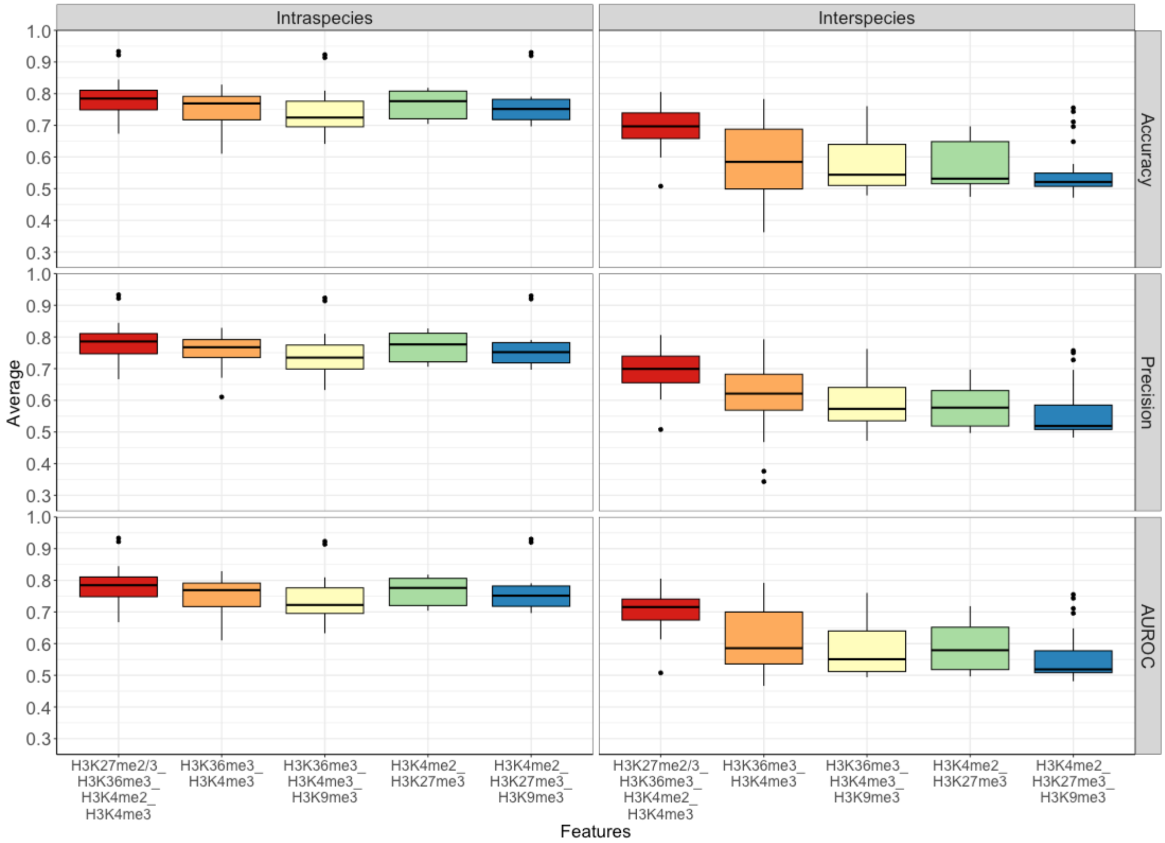

**Supplementary Figure 5**: Feature design impact on prediction performance. Average shallow learning classifier performance for intra- (left) and inter- (right) species predictions using varying feature input approaches: TSS window of 200bp or 5kb and epigenetic ChIPseq signal averaged over full TSS window or in 100bp bins returning a modified signal profile. Boxes represent median +/- quartile for n= 162, 108, 144 replicates for 200 bp average, 5000 bp average, and 5000 bp profile, respectively.

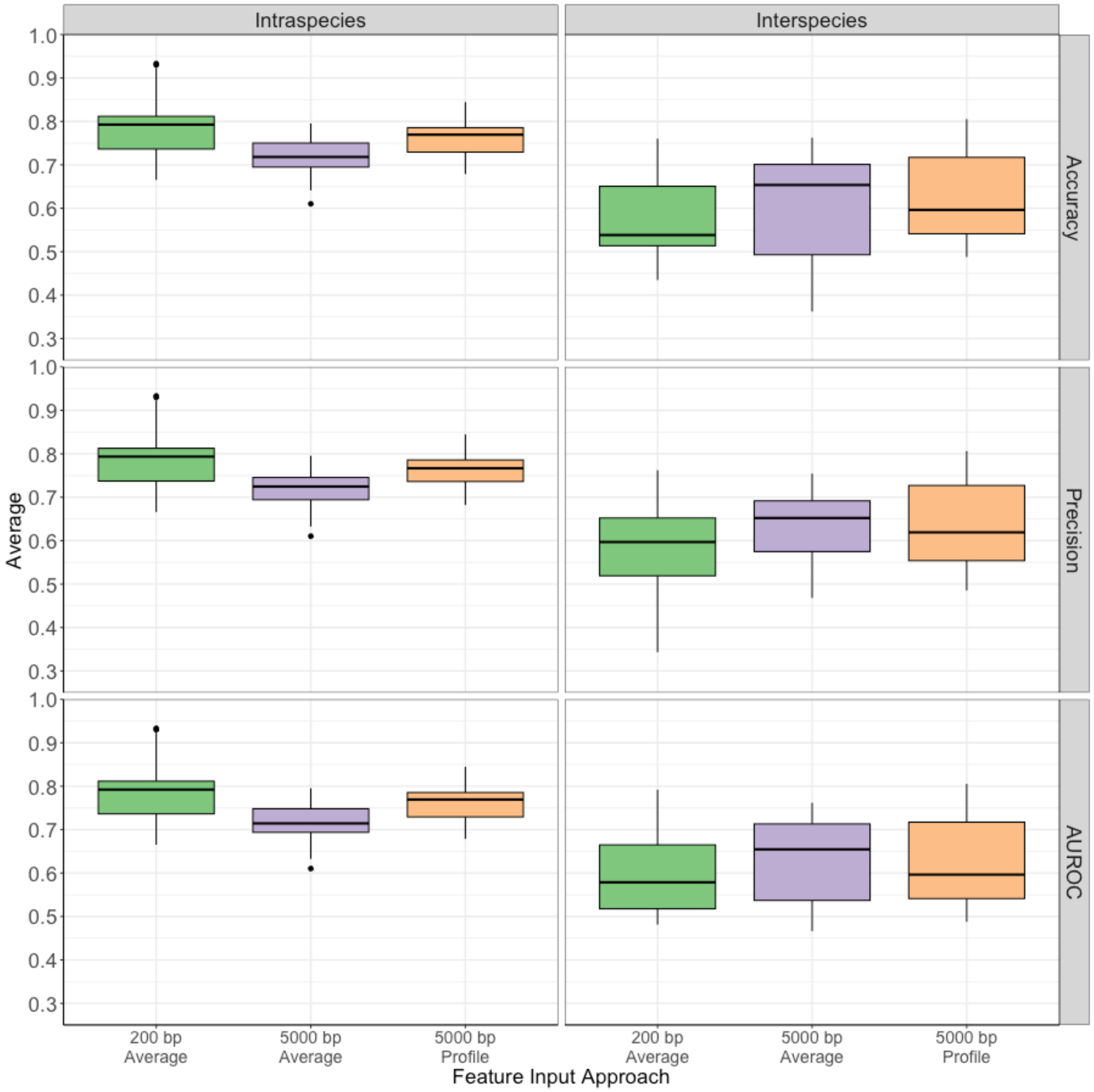

**Supplementary Figure 6**. Data pre-processing pipelines.

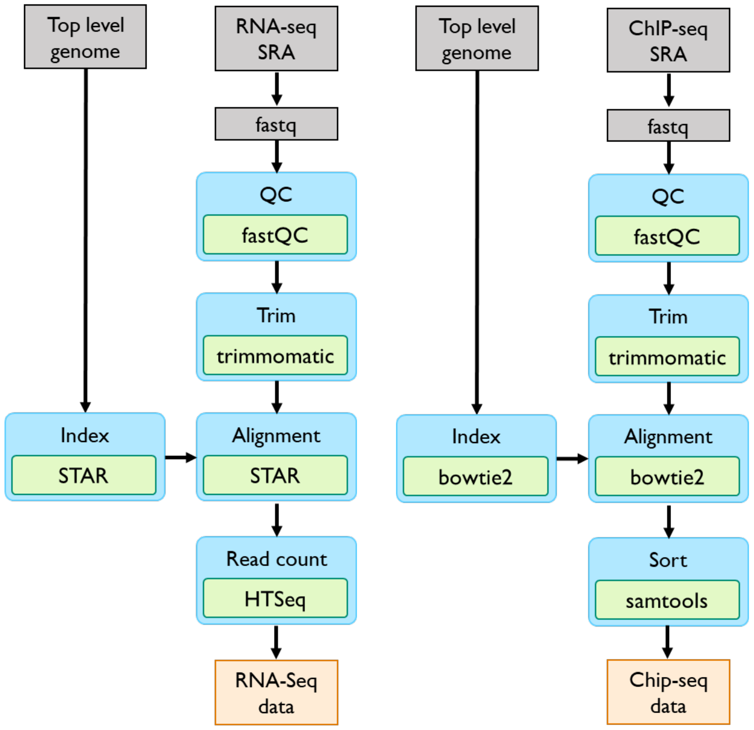

**Supplementary Figure 7**. Feature design strategy.

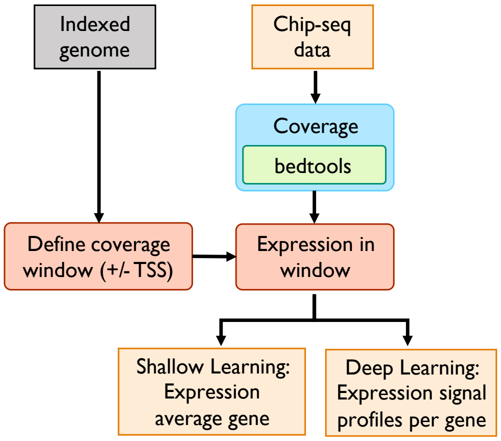

**Supplementary Figure 8**. Machine learning training and evaluation strategy.

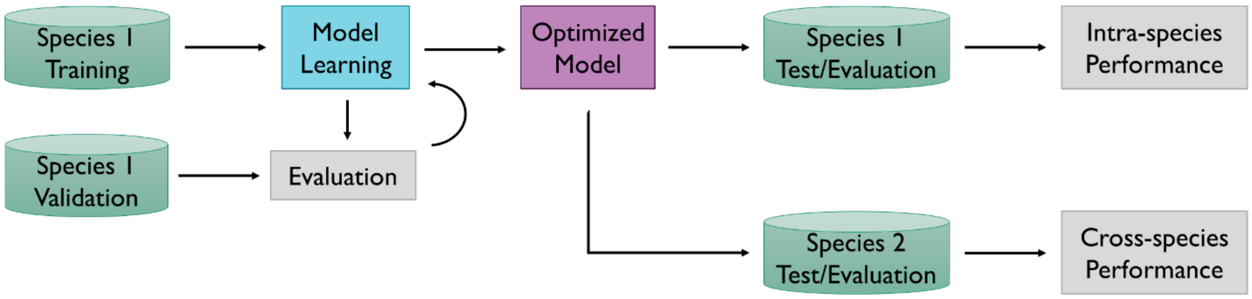

**Supplementary Figure 9**: Intra- and inter-species prediction task performance of the sequence-only 7B parameter Evo2 model using a 5kb TSS window on accuracy (left), precision (center), and area under the receiver operating curve (AUC; right) metrics. Boxes represent median +/- quartile.

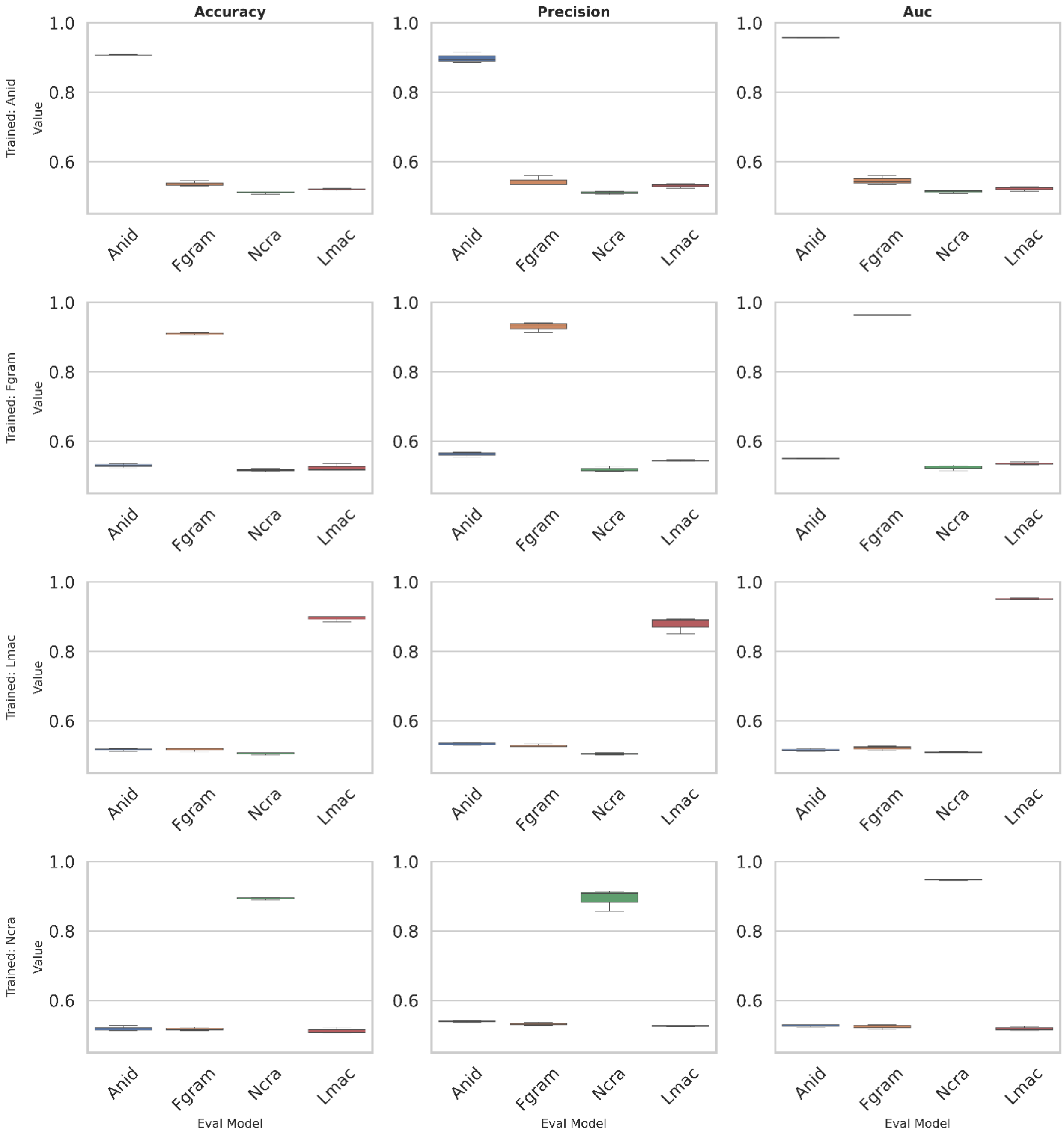

**Supplementary Figure 9**: Intra- and inter-species prediction task performance of the sequence-only 7B parameter Evo2 model using a 200bp TSS window on accuracy (left), precision (center), and area under the receiver operating curve (AUC; right) metrics. Boxes represent median +/- quartile.

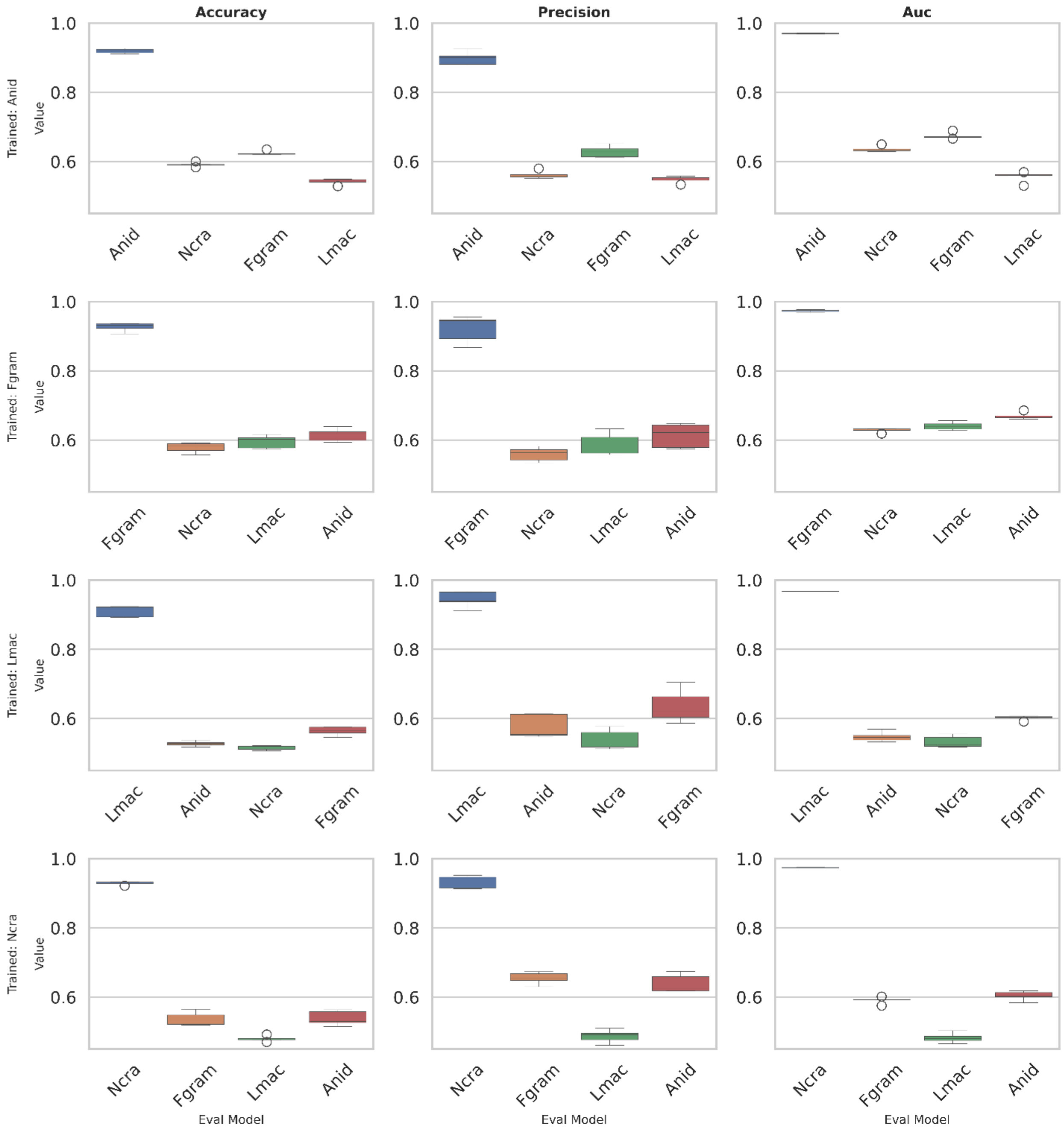

**Supplementary Table 1**: MAPLE vs. benchmark model intra- and inter-species prediction specificity (true negative rate) and sensitivity (true positive rate).

| Model | Trained | Tested | Specificity Mean | Specificity Std | Sensitivity Mean | Sensitivity Std |
|---|---|---|---|---|---|---|
| **Benchmark** | **Anid** | **Anid** | 0.791 | 0.006 | 0.866 | 0.002 |
| | | **Fgram** | 0.479 | 0.019 | 0.899 | 0.006 |
| | | **Lmac** | -- | -- | -- | -- |
| | | **Ncrassa** | 1.000 | 0.000 | 0.001 | 0.001 |
| | **Fgram** | **Anid** | 0.786 | 0.002 | 0.801 | 0.001 |
| | | **Fgram** | 0.866 | 0.005 | 0.762 | 0.003 |
| | | **Lmac** | 0.499 | 0.013 | 0.950 | 0.004 |
| | | **Ncrassa** | 0.975 | 0.030 | 0.178 | 0.182 |
| | **Lmac** | **Anid** | -- | -- | -- | -- |
| | | **Fgram** | 0.961 | 0.000 | 0.492 | 0.005 |
| | | **Lmac** | 0.682 | 0.009 | 0.912 | 0.001 |
| | | **Ncrassa** | 1.000 | 0.000 | 0.000 | 0.000 |
| | **Ncrassa** | **Anid** | 0.002 | 0.000 | 1.000 | 0.000 |
| | | **Fgram** | 0.200 | 0.269 | 0.969 | 0.046 |
| | | **Lmac** | 0.091 | 0.036 | 0.998 | 0.000 |
| | | **Ncrassa** | 0.840 | 0.010 | 0.706 | 0.081 |
| **MAPLE** | **Anid** | **Anid** | 0.691 | 0.042 | 0.723 | 0.072 |
| | | **Fgram** | 0.809 | 0.006 | 0.476 | 0.008 |
| | | **Ncrassa** | 0.091 | 0.004 | 0.938 | 0.004 |
| | **Fgram** | **Anid** | 0.627 | 0.005 | 0.755 | 0.002 |
| | | **Fgram** | 0.824 | 0.012 | 0.746 | 0.017 |
| | | **Lmac** | 0.618 | 0.007 | 0.812 | 0.003 |
| | | **Ncrassa** | 0.727 | 0.006 | 0.793 | 0.006 |
| | **Lmac** | **Fgram** | 0.927 | 0.002 | 0.504 | 0.005 |
| | | **Lmac** | 0.681 | 0.013 | 0.797 | 0.011 |
| | | **Ncrassa** | 0.765 | 0.014 | 0.775 | 0.011 |
| | **Ncrassa** | **Anid** | 0.518 | 0.003 | 0.831 | 0.002 |
| | | **Fgram** | 0.767 | 0.024 | 0.756 | 0.002 |
| | | **Lmac** | 0.555 | 0.008 | 0.825 | 0.007 |
| | | **Ncrassa** | 0.505 | 0.248 | 0.836 | 0.036 |

**Supplementary Table 2**: MAPLE intra-species prediction performance for *A. nidulans* using modification subsets and varying TSS windows.

| TSS window | 40kb | 40kb | 5kb | 5kb | 5kb |
|---|---|---|---|---|---|
| **Modification(s) included** | **H3K36me3** | **H3K36me3, H3K4me3, H3K9me3** | **H3K9me3** | **H3K36me3** | **H3K36me3, H3K4me3, H3K9me3** |
| **AUROC** | -- | -- | 0.528 | 0.734 | 0.820 |
| **Accuracy** | 0.711 | 0.716 | 0.519 | 0.687 | 0.752 |

#### Supplemental Data

**Supplemental Data files consist of outputs from HOMER (for details about the format, please see http://homer.ucsd.edu/homer/).**

**The naming scheme is as follows:**

**[Species1]Model_[Species2]Test_[Modification]locs_[homer/known]Results.html**

**Species1 – the species used to train the model**

**Species2 – the species used as a test set**

**Modification – the specific histone modification evaluated by SHAP**

**homer/known - for details on the difference between these results files, please see http://homer.ucsd.edu/homer/**

### Supplementary Methods

#### MAPLE Hyperparameter Optimization

Hyperparameter optimization for EAGLE was performed using RandomizedSearchCV on the model as a KerasClassifier object with the below ranges defining the parameter search space grid:

Learning rate = [1e-3, 1e-4, 1e-5]

Dense layer dropout = [0.2, 0.3, 0.4, 0.5]

Batch size = [100,500,6000]

Activation = ['softmax', 'softplus', 'softsign', 'relu', 'sigmoid', 'hard_sigmoid', 'linear']

Optimizer = ['SGD', 'Adagrad', 'Adam', 'Adamax', 'Nadam']

Momentum = [0.0, 0.25, 0.5, 0.75, 1.0]

Initialization mode = ['uniform', 'lecun_uniform', 'normal', 'zero', 'glorot_normal', 'glorot_uniform', 'he_normal', 'he_uniform']

Regulation rate=[.1, .01, 0.001]

Dense layer 1 neurons = [100,150,200,250,300]

Dense layer 2 neurons = [25,50,75,100]

CNN layer 1 features = [15,20,25,30]

CNN layer 2 features = [30,40,50,60]

CNN layer 3 features = [50,75,100]

#### Optuna Usage for Benchmark Model Optimization

We instructed Optuna to sample with uniform density in log space, making it more likely to choose smaller values within the search ranges. Twenty optimization trials were completed for each combination of species and number of histone modifications. Each trial ran for fifteen epochs with a batch size of 64 samples. Optuna returned the following parameters that resulted in the highest validation accuracy:

| | **Learning rate** | **L2** | **Gamma** |
|---|---|---|---|
| *F. graminearum* | 3.094e-05 | 2.549e-05 | 0.481 |
| *L. maculans* | 2.440e-04 | 8.367e-05 | 0.134 |
| *N. crassa* | 3.996e-05 | 1.018e-04 | 0.193 |
